# The endosomal TbTpr86/TbUsp7/SkpZ (TUS) complex controls surface protein abundance in trypanosomes

**DOI:** 10.1101/2020.11.10.376269

**Authors:** Kayo Yamada, Farzana K. Yaqub, Martin Zoltner, Mark C. Field

## Abstract

In trypanosomes the orthologs of human USP7 and VDU1 control abundance of a cohort of surface proteins, including invariant surface glycoproteins (ISGs) by functioning as deubiquitinases (DUBs) Silencing TbUsp7 partially inhibits endocytosis and invariant surface glycoprotein turnover. As a component of cullin E3 ubiquitin ligases, S-phase kinase-associated protein 1 (Skp1) has crucial roles in cell cycle progression, transcriptional regulation, signal transduction and other processes in animals and fungi. Unexpectedly, trypanosomes possess multiple Skp1 paralogs, including a divergent paralog designated SkpZ. SkpZ is implicated in suramin-sensitivity and endocytosis and decreases in abundance following TbUsp7 knockdown and physically interacts with TbUsp7 and TbTpr86. The latter is a tetratricopeptide-repeat protein also implicated in suramin sensitivity and located close to the flagellar pocket/endosomes, consistent with a role in endocytosis. Further, silencing SkpZ reduced abundance of TbUsp7 and TbTpr86 and many *trans*-membrane domain surface proteins. Our data indicate that Tb<u>T</u>pr86, Tb<u>U</u>sp7 and <u>S</u>kpZ form the ‘TUS’ complex that regulates abundance of a significant cohort of trypanosome surface proteins.

## Introduction

Trypanosomiasis is a vector-borne parasitic disease caused by infection with trypanosomes and includes multiple African and American species. The trypanosome surface is the interface with the host and possesses adaptations critical for survival in multiple distinct environments; this importance is reflected in the diverse surface protein components exhibited by the major human pathogens *Trypanosoma brucei and T. cruzi.* The *T. brucei* mammalian bloodstream trypomastigote has an efficient endocytic system that enables rapid recycling of surface proteins, antibody clearance and immune evasion mechanisms (Manna et al 2015). The vast majority of the surface is covered by the GPI-anchored variant surface glycoprotein (VSG), although a considerable diversity of additional proteins also populates the surface (Gadelha et al., 2015, Shimogawa et al., 2018). While VSG itself is responsible for antigenic variation and hence immune evasion, several receptors have been characterised recently, but most surface proteins remain functionally uncharacterised. The trypanosome surface is clearly organised as the proteomes of the cell body, flagellum and flagellar pocket are distinct (Oberholzer et al., 2014, Gadelha et al., 2015). Further, this differential composition extends to internal compartments, with many proteins shared between endosomes and surface domains while other proteins have distinct locations (Gadelha et al., 2015). The mechanisms responsible for targeting likely reside within multiple determinants, including *cis*-elements embedded within the proteins themselves, post-translational modification, including phosphorylation, fatty acylation and ubiquitylation, together with so far uncharacterised gating mechanisms at the surface (Allen et al., 2007, Emmer et al, 2011, Gadelha et al, 2009, Mussmann et al, 2004, Graf et al., 2013, Baker et al., 2012).

ISG75 is a member of an extensive superfamily of *T. brucei*-specific type I *transmembrane* proteins with large extracellular domains and comparatively short cytoplasmic domains (Allison et al 2014). ISGs, and their related intracellular glycoproteins (IGPs), are comparatively abundant and the presence of multiple paralogs for each ISG/IGP subfamily suggests a requirement for sequence diversity as well as abundance. Significantly, both ISG75 and its closest relative, ISG65, have a considerable presence within the endosomal system as well as at the surface and are believed to be receptors although physiologically relevant ligands remain unidentified. Further, both are ubiquitylated and while there are clear differences between mechanisms underlying ISG65 and ISG75 targeting and turnover, they share at least one deubiquitylating enzyme in the trypanosome ortholog of Vdu1, while Usp7 only impacts ISG75 turnover (Zoltner et al., 2015). Ubiquitylation is an important regulator of the trypanosome surface but, given the complexity of ubiquitylation systems across the tree of life, it is unclear precisely how these various elements function together.

Protein ubiquitylation proceeds via several steps, depending on the mechanism of action of the ubiquitin ligase responsible for transfer of ubiquitin to the client protein (Hershko and Ciechanover 1998). Cullins are a family of scaffold proteins that support E3 ubiquitin ligases with members present across the eukaryotes, including trypanosomes (Rojas et al., 2017, Lin et al., 2017, Huysman et al., 2014, Desbois et al., 2018, Angers, et al., 2006, Iglesias et al., 2018 and Nagai et al., 2018). Cullins combine with RING proteins to form cullin-RING ubiquitin ligases (CRLs) that are highly diverse and play roles in various cellular processes, most notably protein degradation by ubiquitylation. CRLs, such as the Skp1-Cul1-Fbox (SCF) complex target proteins for ubiquitin-mediated destruction and regulate multiple functions including DNA replication, glucose sensing and limb formation in metazoa. The cullin N-terminus is highly variable and, in the SCF complex, interacts with specific adaptor proteins including Skp1 (S-phase kinase-associated protein 1), to bring substrate proteins close to the E3 ligase Rbx1 (Petroski and Deshaies 2005, Zheng et al., 2002, Goldenberg et al., 2004). Different combinations of adaptor proteins (F-box proteins in the SCF complex) binding to the cullin N-terminus are key to diversifying substrate specificity.

To date functions of Skp1 and its paralogs remain uninvestigated in trypanosomatids, despite the potential for important contributions towards cell cycle progression and differentiation pathways. Here we show that a trypanosomatid-specific Skp1-like protein (SkpZ) forms a novel complex together with TbUsp7 and a trypanosome-specific Tpr protein which we designate as the Tpr86/Ubc7/SkpZ or TUS complex. Further, using unbiased whole cell proteomics we demonstrate that TUS mediates surface *transmembrane* protein turnover, indicating that the complex is a major factor in controlling surface protein expression.

## Results

### Trypanosoma brucei *possesses multiple Skp1 paralogs*

To investigate the complexity and possible roles for Skp1 and paralogs in trypanosomes, we performed a genome search using human Skp1 (P63208) as well as the trypanosome Skp1 paralogs Tb927.11.6130 and Tb927.11.13330 identified previously by cullin affinity isolations (Canavate et al., in preparation) and a Skp1 paralog (Tb927.11610) identified as a suramin sensitivity-associated gene (Alsford et al., 2012) as BLAST queries. We searched across high quality kinetoplastid genomes and retrieved orthologs from the vast majority of these taxa. Phylogenetic reconstruction robustly identified four clades of Skp1 paralogs, which based on associations with cullin ligases we designated as Skp1.1 and Skp1.3, together with a more divergent Skp1-like protein as SkpZ and the fourth paralog with very weak homology that we designated as Skp1-related or Skp1.4 (Figure 1, Tables S1 and S2). Only the genomes of *T. cruzi* (CL Brener and Dm28c), *Angomonas* and *Strigomonas* lack a SkpZ gene, probably the result of incomplete databases.

**Figure 1:**
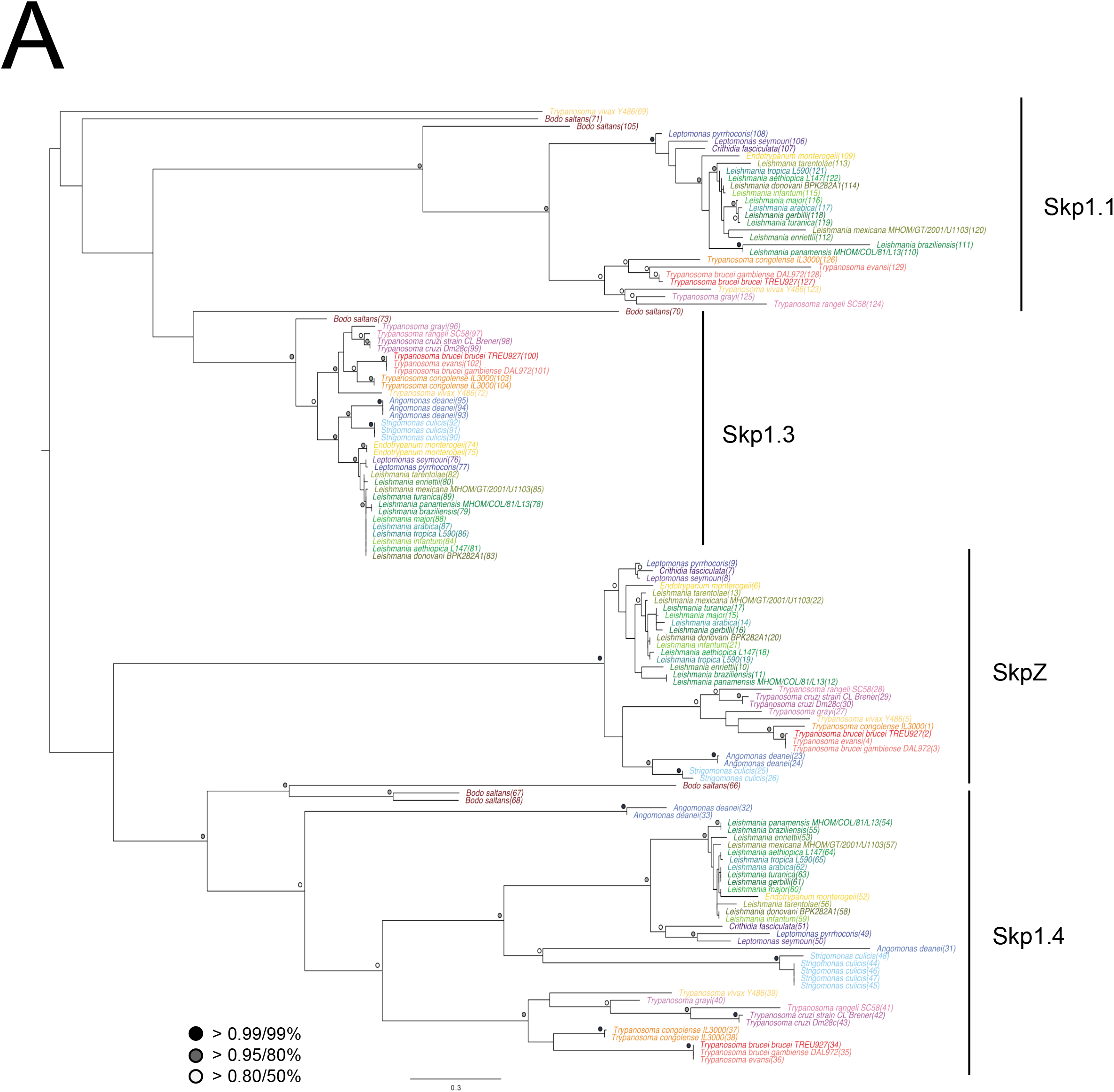
Evolution of selected trypanosomatid Skp1 proteins. Phylogenetic tree of Skp1 paralogs across kinetoplastid genomes. Trees were constructed using MrBayes and PhyML, with the MrBayes topology shown. Statistical support values are shown as symbols at relevant nodes. Species colours are green/blue for Leishmania and related, blue for insect only parasites, purple for American trypanosomes and red/orange for African trypanosomes. The basal kinetoplastid *B. saltans* is in tan. Numbers in parenthesis following the taxon name refer to accessions given in Table S1.

Skp1 possesses a POZ-domain at the N-terminus, consisting of a β/α sandwich double layer with a C-terminal dimerization domain essential for binding of F-box proteins. The interaction stabilises the conformation of Skp1 and increases protein levels by preventing aggregation (Zheng et al., 2002, Yoshida and Tanaka 2011) Trypanosome Skp1.1 and Skp1.3 retain this canonical Skp1 architecture, but SkpZ has an extended N-terminus (Figure S1). Skp1.1 and Skp1.3 associate with cullin ligases in trypanosomes (Canavante et al., in preparation) but SkpZ was never detected in association with a cullin complex, suggesting a divergent function.

### TbSkpZ is an endosomal protein and interacts with TbUsp7 and an 86kDa Tpr-protein

To investigate SkpZ further we established an endogenously tagged SkpZ cell line with three copies of the HA-epitope fused at the C-terminus. While *Homo sapiens* Skp1 localizes to both the nucleus and cytoplasm none of the trypanosome Skp1 orthologs apparently share this localisation and SkpZ localised to the endosomal region, supporting the possibility of a novel function associated with protein trafficking (www.tryptag.org, Dean et al., 2017).

TbSkpZ HA-tagged cells were harvested and cryo-milled (Obado et al., 2016). We identified conditions for isolation of TbSkpZ together with several additional interacting proteins (Figure 2, Table S3), which were identified by LCMSMS and MaxQuant/Perseus (Zoltner et al., 2020). Unexpectedly, TbSkpZ interacted with TbUsp7 rather than a cullin complex, with several additional proteins also identified (Figure 2). Altogether four proteins, GRESAG4, PFC17, Tb927.11.810 and TbUSP7 were significantly enriched. GRESAG4 has multiple paralogs, is highly abundant and frequently identified in proteomics analyses which suggests that this is a contaminant, while PFC17 (paraflagellar component 17) is a small protein, which increases the potential of non-specific binding and/or detection in mass spectrometry and is a flagellar protein which, given no additional evidence for flagellar interaction, we considered unlikely to be a *bona fide* SkpZ partner (Subota et al., 2014).

**Figure 2:**
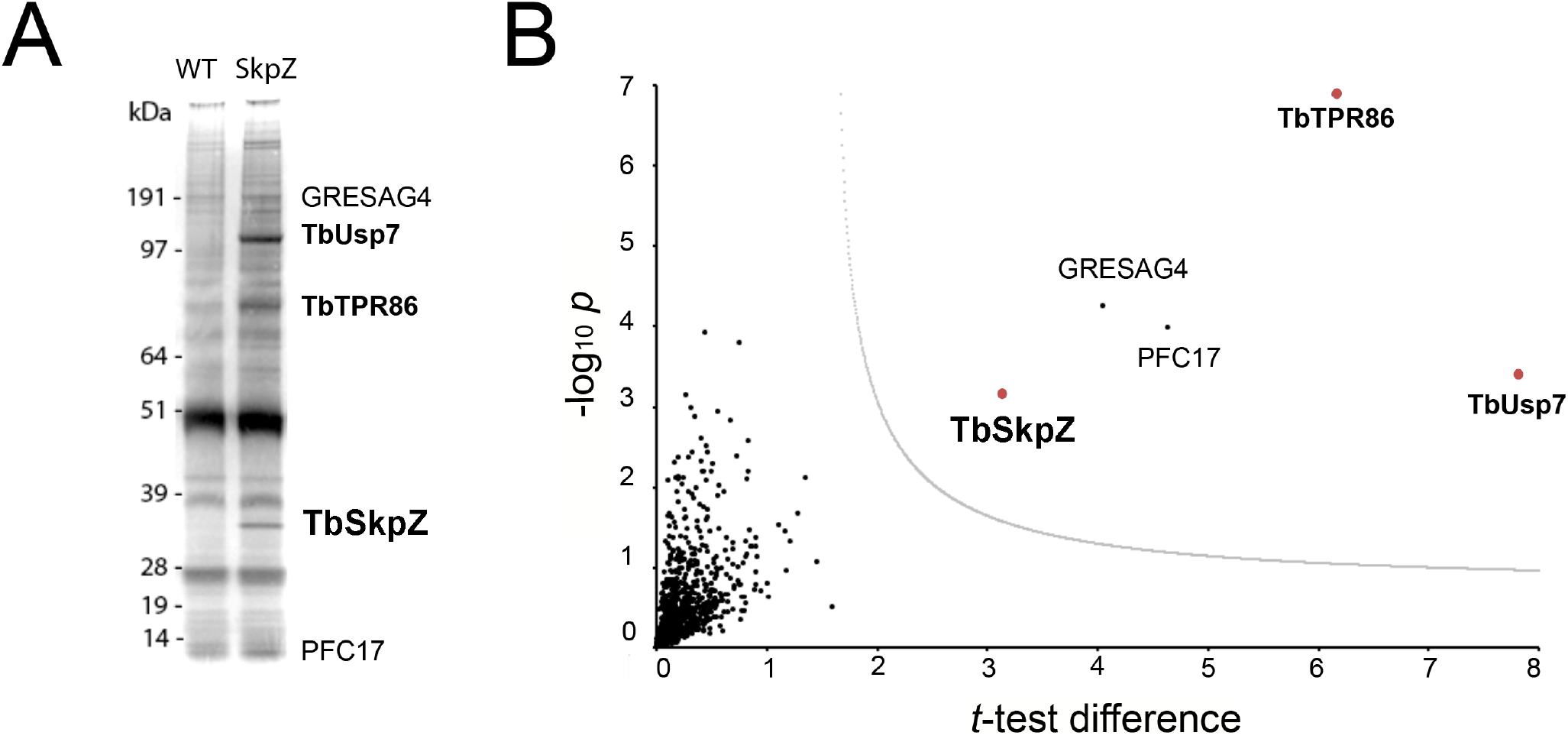
SkpZ interacts with TbUsp7 and TbTpr86. Panel A: SDS-PAGE analysis proteins obtained by affinity isolation of 3xHA-tagged SkpZ and parental cells (WT) as control. The putative TbUsp7 and SkpZ bands, based on estimated molecular weight, were cut from a silver stain gel and individually analyzed by mass spectrometry. Panel B: Volcano plot comparing all proteins coeluted with TbSkpZ with an untagged control elution, and which identifies the TUS complex. The −log10 *t*-test P-value was plotted versus the *t*-test difference (difference between means). The cut-off curve (dotted line) is based on a false discovery rate of 0.05. Members of the Tpr86/Usp7/SkpZ (TUS) complex highlighted in bold and red.

Tb927.11.810 encodes an 86kDa protein with a predicted tetratricopeptide-repeat motif (TPR, IPR011990) located towards the centre of the sequence, between residues 246 and 485, and an overlapping pentatricopeptide repeat region (PPR, IPR002885) at residues 196 to 384, with several experimentally determined phosphorylation sites between 84 and 103; we designated this protein TbTpr86. There is evidence for modest upregulation in the mammalian bloodstage compared to insect forms and decreased expression in stumpy forms at the protein level. Importantly, the protein is located close to the flagellar pocket, similar to SkpZ (www.tryptag.org).

The Tpr motif consists of degenerate 34 amino acid repeats and is present in a wide variety of proteins in other organisms, including anaphase-promoting complex subunits cdc16, cdc23 and cdc27, transcription factors, the major receptor for peroxisomal matrix protein import PEX5 and others (Das et al. 1998). Tandem arrays of three to 16 Tpr motifs form scaffolds to mediate protein–protein interactions including assembly of heteromeric complexes as well as self-assembly (Blatch and Lässle 1999). In *T. brucei* the Tpr motif is retained by the Pex5 ortholog (Gualdrón-López et al., 2013) and predicted for numerous additional proteins.

### TbSkpZ knockdown reduces TbUsp7, ISG75 and Tpr86 levels

Cells harbouring a stemloop RNAi construct specific for TbSkpZ were induced with tetracycline. Quantification revealed that knockdown of TbSkpZ led to enlargement of the flagellar pocket, the ‘BigEye’ phenotype originally observed for knockdown of clathrin (Allen et al., 2003). Specifically, 17% of induced cells possessed the BigEye morphology, constituting a six-fold increase in frequency when compared to uninduced controls. Furthermore, TbSkpZ knockdown resulted in a slight proliferative effect. These impacts effectively mirrored those obtained for TbUsp7 knockdowns (Zoltner et al., 2015).

A whole cell proteome derived from SDS lysates of silenced cells was compared with parental cells using stable isotope-labeling by amino acids in culture (SILAC) and LCMSMS. 3904 protein groups were detected, of which 3116 could be quantified in both replicates (Figure 3; Table S3). Significantly, TbSkpZ knockdown reduced protein levels of TbUsp7 by 38%, ISG75 by ~50%, Tpr86 by ~42% and SkpZ itself by ~70%, which also validates the knockdown as efficient (Figure 3). We were able to reproduce the ISG75 decrease using western blotting providing orthogonal validation of the proteomics data (Figure 3, inset). Significantly, TbUsp7 knockdown was previously shown by us to decrease both ISG75 expression (~40%) and Tpr86 (~58%) (Zoltner et al., 2015). The impact of TbSkpZ was highly biased towards surface/endosomal membrane proteins as well as some of the machinery potentially associated with endocytosis, specifically the VAMP7b SNARE protein (Tb927.5.3560) (Table 1) (Venkatesh et al., 2017). Tb927.11.1750, also present only in kinetoplastida and which lacks an obvious TMD or GPI-anchor signal and a location at the Golgi complex, was also decreased.

**Figure 3:**
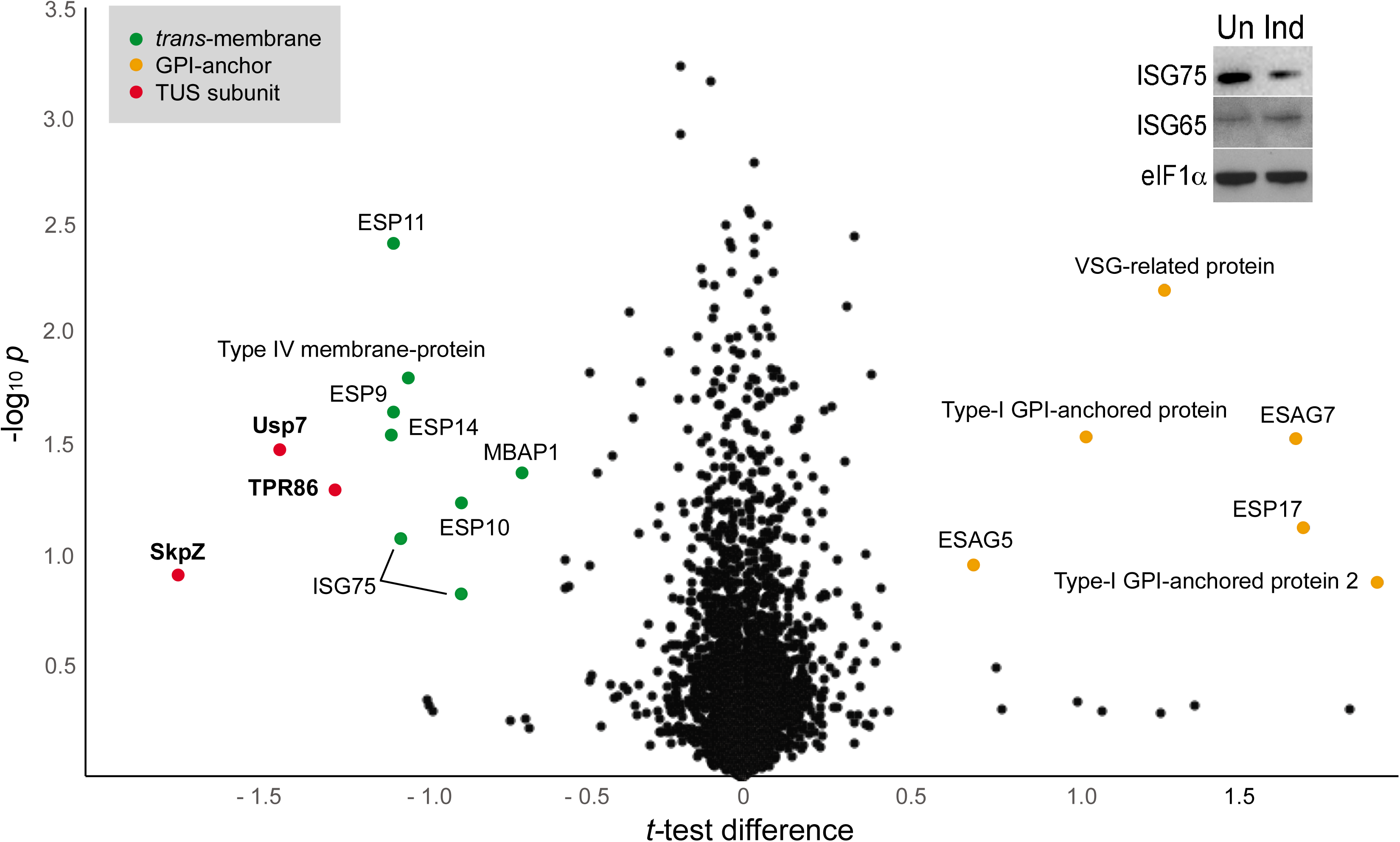
Volcano plot of protein abundance changes following knockdown of SkpZ. t-test difference plotted against −log10 transformed t-test p-value. Data points representing protein groups significantly shifted after 48 hours knockdown induction are labeled (for ratio shifts see Table 1). Points corresponding to TbUsp7, TbSkpZ and TbTpr86 are plotted in red, GPI-anchored proteins in orange and *trans*-membrane proteins in green. Inset: Silencing SkpZ accelerates invariant surface glycoprotein 75 turnover. Cells were silenced by RNA against TbSkpZ using a tetracyclin (Tet) inducible system for 48 hours and protein levels estimated by Western blotting. The data are a representative example of two experiments, which gave similar results.

**Table 1:** Select protein abundance changes in SkpZ knockdown cells. The ratio of abundance of selected protein groups in control and SkpZ-silenced cells is shown and data are ranked based on descending ratio. Note that all membrane anchored proteins exhibit a decrease in abundance possess a *trans*-membrane domain and increased entries possess GP-anchor linkage. The precise mechanism of anchoring ESAG5 is unclear at present. The accession numbers of the SkpZ RNAi target and other TUS components are in bold. Accession numbers of kinetoplastida-specific proteins are italicised.

| Accession | Annotation | Domains, architectural features | Ratio | -Log10 P-value | Description |  |
| --- | --- | --- | --- | --- | --- | --- |
| *Tb927.7.6490* | Hypothetical | Predicted GPI | 3.99 | 0.88 |  | Increased |
| *Tb927.7.3260* | ESAG7 | GPI, VSG-related fold | 3.22 | 1.53 | Transferrin receptor subunit |  |
| *Tb927.7.180* | VSG | Predicted GPI, VSG type B-domain | 2.44 | 2.21 | VSG-related |  |
| *Tb927.7.6600* | Hypothetical | Predicted GPI | 2.07 | 1.54 |  |  |
| *Tb11.v5.0826* | ESAG5 | Tandem bactericidal/permeability-increasing | 1.64 | 0.96 | BES11 ESAG5 |  |
| *Tb927.5.3560* | VAMP7B | v-SNARE coiled-coil domain | 0.72 | 0.96 | R-SNARE, VAMP7B paralog, endosomal | Decreased |
| *Tb927.6.3550* | ATPase | TMD, E1-E2 ATPase | 0.72 | 1.83 | Phospholipid-translocating P-type ATPase (flippase) |  |
| *Tb927.9.9410* | SLS1 | TMD | 0.69 | 0.98 | Inositol phosphorylceramide synthase |  |
| *Tb927.11.13130* | MBAP1 | TMD | 0.63 | 1.38 | Membrane bound acid phosphatase 1 |  |
| *Tb927.2.1700* | ESP10 | TMD, EGF-like domain | 0.55 | 1.24 | Enriched in surface proteome |  |
| *Tb927.5.360* | ISG75 | TMD | 0.56 | 0.83 | 75 kDa invariant surface glycoprotein |  |
| *Tb927.5.390* | ISG75 | TMD | 0.49 | 1.08 | 75 kDa invariant surface glycoprotein |  |
| *Tb927.11.7550* | Hypothetical | Two TMD | 0.49 | 1.81 | Endosomal location |  |
| *Tb927.4.3040* | ESP11 | TMD | 0.48 | 2.42 | Enriched in surface proteome |  |
| *Tb927.9.11480* | ESP9 | TMD | 0.48 | 1.65 | Enriched in surface proteome |  |
| *Tb927.7.470* | ESP14 | TMD | 0.48 | 1.56 | Enriched in surface proteome |  |
| ***Tb927.10.810*** | **TbTpr86** | TPR-like | 0.42 | 1.31 | Tetratricopeptide-repeat protein |  |
| ***Tb927.9.14470*** | **TbUsp7** | Ubiquitin-specific protease | 0.38 | 1.48 | Ubiquitin carboxyl-terminal hydrolase |  |
| ***Tb927.10.11610*** | **SkpZ** | Skp1 dimerisation domain, POZ | 0.31 | 0.92 | SkpZ |  |

Most significantly, TbSkpZ silencing impacted surface protein levels depending on their mechanism of membrane attachment. Specifically, proteins with a predicted TMD were decreased in abundance, while those with a predicted or known GPI-anchor increased. The decrease in abundance is most likely due to the absence of DUB activity via loss of TbUsp7, with a resulting increase in lysosomal delivery of ubiquitylated proteins. Significantly, the observed changes are near identical to those previously described for TbUsp7 (Figure 4), strongly supporting the possibility that both proteins function in the same pathway. Further, many TMD proteins affected are enriched in the *T. brucei* surface-labeled proteome (Gadelha et al., 2014) suggesting that turnover of these surface proteins is controlled through ubiquitylation (Table 1).

**Figure 4:**
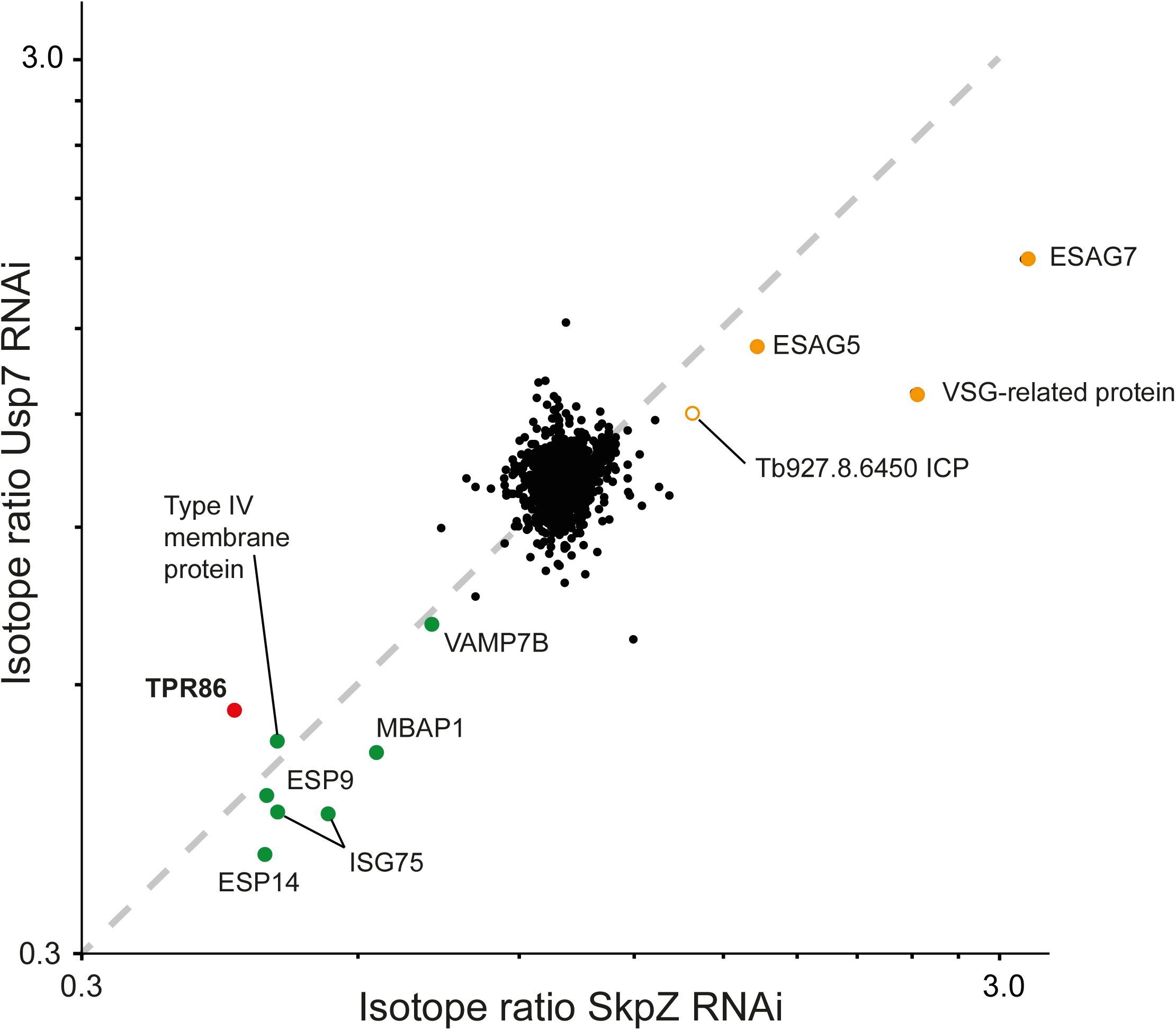
Correlation of proteome changes upon SkpZ and Usp7 silencing. Abundance shifts after 48 hours silencing of SkpZ and Usp7 (Zoltner et al., 2015) were plotted against each other. Selected protein groups in the cohorts are labeled and color coded as green; predicted *trans*-membrane proteins, orange; predicted GPI-anchored, red; TUS complex component. Note that the proteome analysis of the SkpZ RNAi lysate is deeper as extracts were analysed on a more advanced mass spectrometer accounting for the absence of some proteins from the Usp7 data set.

### TbTpr86 is a pan-kinetoplastid protein

TbTpr86 protein abundance is affected by TbSkpZ RNAi and is a TbSkpZ interactor by affinity isolation. Hence, we chose to investigate TbTpr86 in more detail. Initially, we performed comparative genomics screen using the Tpr86 sequence as BLAST query at EuPathDB, NCBI as well as a HMMER search. The TbTpr86 gene is well conserved and syntenic across the kinetoplastida and extends into the bodonids (Figure 5, Table 3). There is, however, no evidence for the protein outside of the lineage indicting an origin post-speciation from the Euglenids. Molecular modelling, using the Phyre2 server, indicates with high confidence that TbTpr86 adopts an a-helical solenoid structure along much of its length (Figure 5).

**Figure 5:**
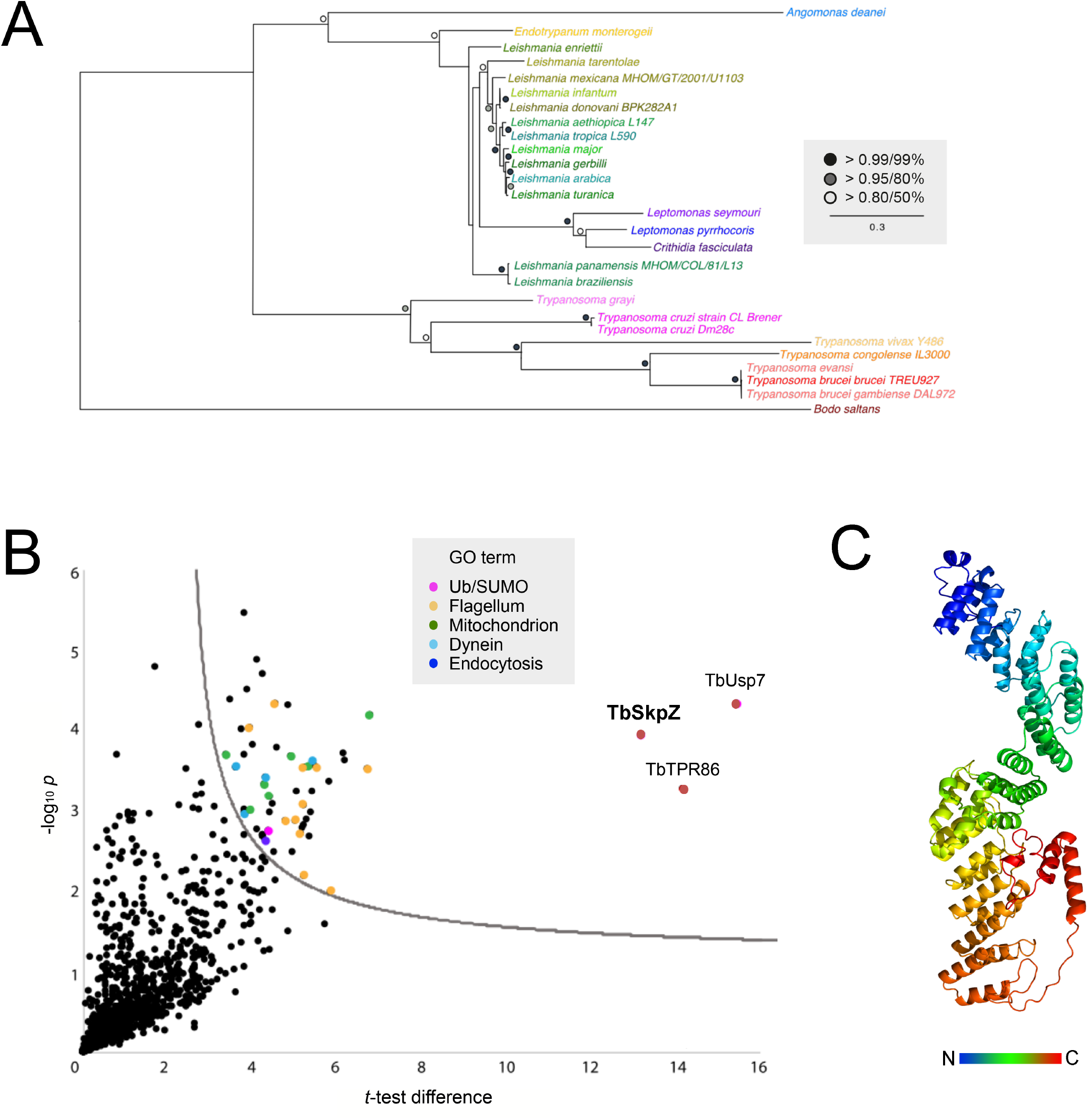
Evolution and interactions of TbTpr86. Panel A: Phylogenetic tree of TbTpr86 paralogs in selected kinetoplastid genomes. Trees were constructed using MrBayes and PhyML, with the MrBayes topology shown. Statistical support values are shown at the relevant nodes. Species colours are arbitrary but consistent throughout all sequence entries and with Figure 1. Accession numbers for entries are given in Table S2. Panel B: Interactions between TbTpr86 and other proteins in trypanosomes validate the TUS complex. Affinity isolations from cryo-milled cells with and without a genomically tagged TbTpr86 were performed in three biological replicates. The −log10 transformed t-test p-values were plotted against *t*-test difference (difference between means). Data points corresponding to TbUsp7, TbSkpZ and TbTpr86 are indicated in red and additional protein groups detected according to the key. Dotted line indicates the confidence limit with a false discovery rate =0.05. Panel C: Model for central region of TbTpr86 generated in Phyre2, demonstrating predicted extensive helical structure of the protein.

We established an endogenously HA-tagged TbTpr86 cell line and cells were harvested and cryo-milled as before. We used essentially the same conditions as developed for the TbSkpZ affinity isolation to identify coenriching proteins with LCMSMS (Table S3). We found strong interactions between TbTpr86, TbUsp7 and TbSkpZ (Figure 5), robustly confirming the presence of a heterotrimeric complex. We designated this complex TUS, based on a composition of Tb<u>T</u>pr86, Tb<u>U</u>sp7 and Tb<u>S</u>kpZ. GRESAG4 and PFC17, enriched in the TbSkpZ affinity isolation, were not detected and confirmed as contaminants. We only detected additional proteins with significantly lower enrichment suggesting that additional interactions were not well retained under the conditions used. *TbSkpZ is required to support vesicle transport.* Transmission electron microscopy revealed a significant accumulation of intracellular membranous structures in TbSkpZ silenced cells, (Figure 6). These structures appear in the region of the cell usually associated with the Golgi complex and endosomes (Figure 6B, C) and also appeared to be associated with the endoplasmic reticulum (Figure 6D) and entirely consistent with localisations for TUS components at TrypTag. We also observed pronounced multi vesicular bodies/autophagosomes (Figure 6D), but no obvious impact to other structures. The contiguous appearance of novel membrane structures suggests a failure to complete budding and hence accumulate vesicular structures.

**Figure 6:**
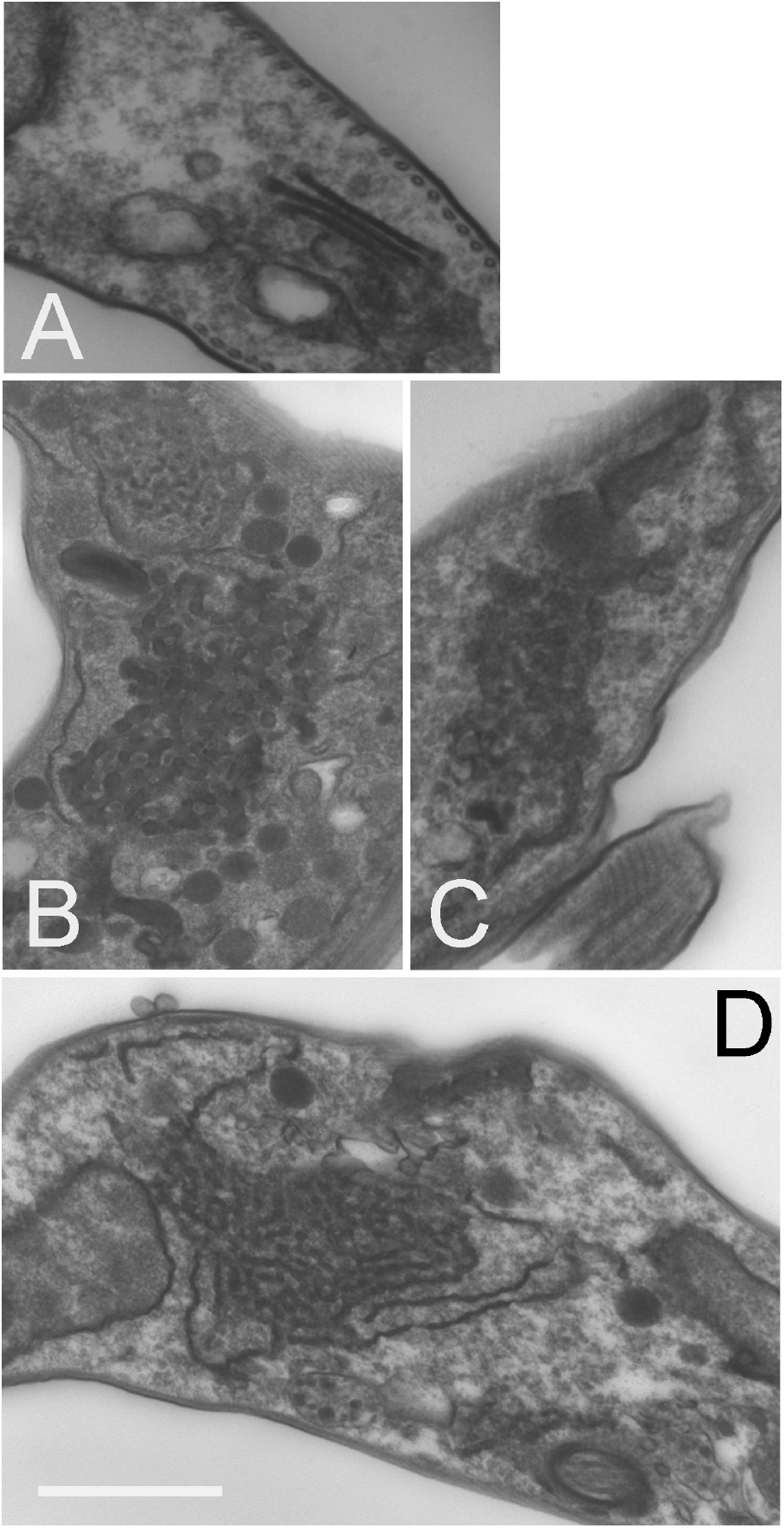
Knockdown of TbSkpZ leads to membrane and vesicle accumulation. Cells were induced for 24 hours, fixed and prepared for electron microscopy. Panel A: Uninduced cell showing the Golgi complex region. Panels B - D: Central regions of indices cells showing accumulation of large clusters of electron dense vesicles. In panel D the formation of these vesicles from tubular structures is suggested by the aligned membrane structure, and which suggests that these are likely ER and/or Golgi-derived structures. These features were observed in ~30% of sections. Other structures, such as the nucleus, flagellum and cytoskeleton appear unaltered. Scale bar is 1.0um in panels A and D and 500nm in panels B and C.

## Discussion

We describe here the TUS complex, which is comprised of the trypanosome ortholog of USP7, a pan-eukaryotic deubiquitinase, together with kinetoplastida-specific proteins SkpZ, and Tpr86. TUS complex interactions are robustly identified by reciprocal isolations using all three subunits as well as *in trans* impacts on abundance following silencing. Significantly, genes encoding all three proteins can be identified from nearly all kinetoplastid genomes, suggesting that the TUS complex is present throughout the lineage and arose early in the evolution of kinetoplastida following speciation from the euglenids. TUS regulates the cell surface proteome, with significant impacts on both TMD and GPI-anchored proteins, but significantly affecting these protein cohorts differentially; TMD proteins are down-regulated when TUS subunits are silenced and GPI-anchored proteins increase in abundance. It is possible that the increase in GPI-anchored protein abundance is a secondary consequence; endocytosis is clearly affected by RNAi against TUS components and in turn GPI-anchored nutrient receptors are upregulated. Further, the complex is associated with the endosomal region of the trypanosome cell, and which suggests that TUS functions to deubiquitylate TMD proteins as they enter the endosomal system, permitting their recycling - silencing prevents this action and increases turnover of TMD proteins (Zoltner et al., 2015). An impact on trafficking machinery proteins, as well as MBAP, is also significant as both participate in endocytic pathways. The increase in GPI-anchored protein abundance may be the result of a simple counting/density sensing mechanism that permits an increase in these proteins due to decreased density of TMD proteins. However, as the GPI-anchored VSG is sorted from other endocytic cargo within sorting endosomes by a clathrin-dependant mechanism, it is also likely that there is a direct impact on the efficiency of this process. However, how GPI-anchored proteins are specifically recognised in trypanosomes is unclear *albeit* that this is likely to be distinct from mechanisms in animals and fungi. Regardless, it is clear that the TUS complex has a major impact on the cell surface proteome and that TUS is an adaptation that occurred early within the kinetoplastida. It is tempting to speculate that this may be associated with the high level of surface GPI-anchored proteins and glycoconjugates in these organisms.

In human cells USP7 is a high abundance DUB with many roles, including control of the cell cycle *via* p53. USP7 is subject to complex regulatory mechanisms and itself ubiquitylated with sophisticated multi-domain architecture involving a self-activation mechanism and several Ubl domains. USP7 can also act as a deSUMOylase (Lecina et al, 2016). Significantly, of many known animal and fungal USP7-interacting proteins, none were identified here and the vast majority are also unlikely to be encoded in any kinetoplastid genome, indicating significant diversity in function. USP7 forms a cellular switch with E3 ligases to facilitate rapid responses to stimuli, coordination of protein level and multiple functions, including endocytosis (Kessler et al, 2007, Hao et al, 2015, Kim and Sixma 2017). This seems unlikely the case in trypanosomes as we failed to identify any E3 ligase, associated subunits or evidence for association of ubiquitylation machinery from isolations of members of the TUS complex. TbTpr86 and TbSkpZ are trypanosome specific, but the known roles of Skp1 proteins as substrate adaptors in association with cullin and Rbx1 containing E3 ligases suggests a possible similar role. While speculative, structure modelling for TbTpr86 confidently predicts formation of an extended α-helical rod, consisting of repeat domains, which is at least conceptually similar in architecture to cullin proteins. Hence, we suggest that the TUS complex may be an ‘anti-cullin’, whereby TbTpr86 provides a scaffold to facilitate TbSkpZ binding substrate and delivering it to TbUsp7 for deubiquitylation, in contrast to the opposite reaction mediated by cullin ligases (Figure 7). Hence, while not intimately associated with an E3 ligase as in metazoan organisms, USP7 may none-the-less function in kinetoplastids as part of a switch mechanism providing highly sensitive and rapid control of surface protein expression.

**Figure 7:**
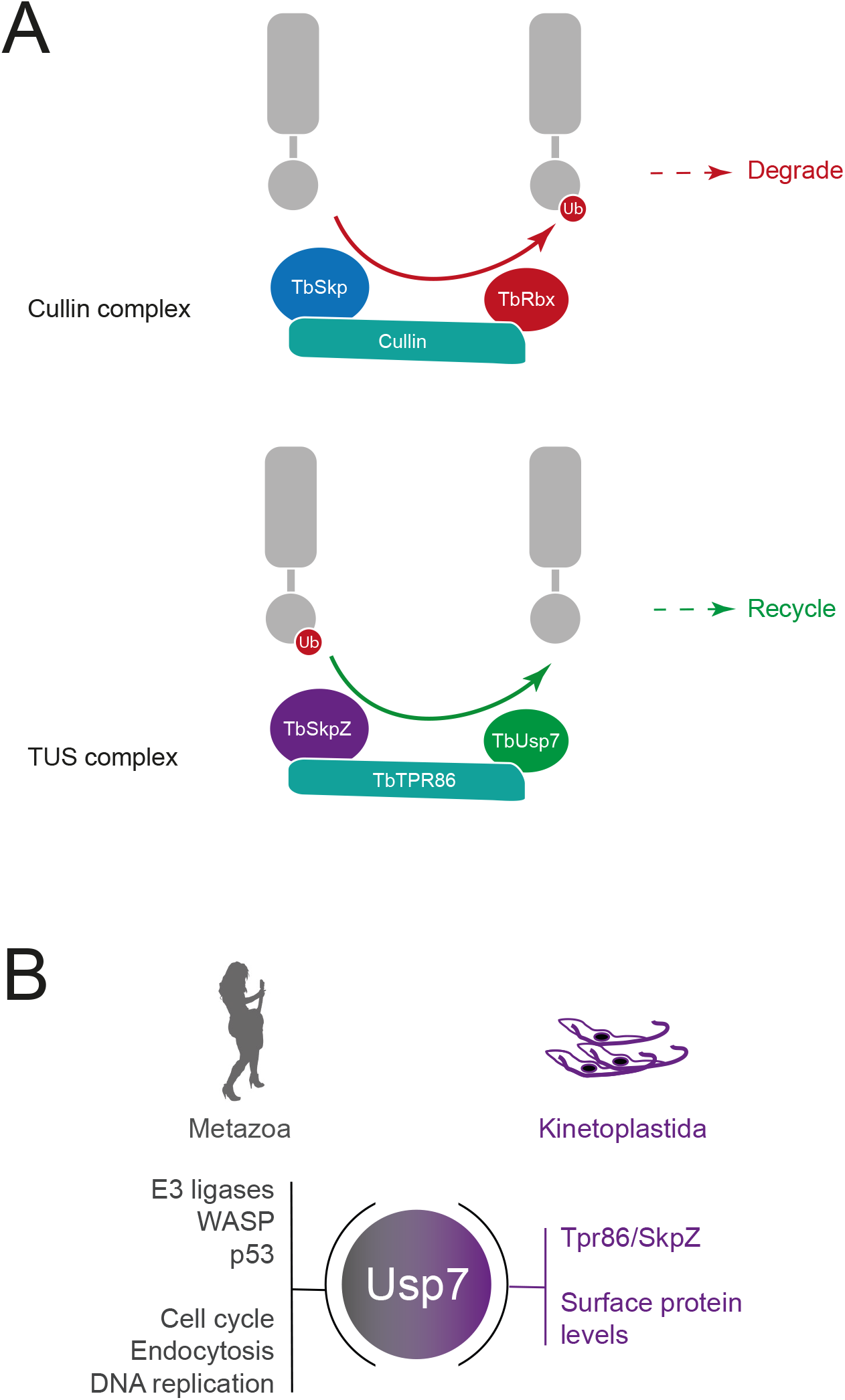
Model for architecture and function of the TUS complex. Top: Cullin 1 contains TbCullin1 (teal), TbRbx1 (red) and TbSkp1.1 (orange) together with numerous F-box proteins (not shown). Substrates are ubiquitylated (red circle) which targets them for degradation. Bottom: TUS contains TbTpr86 (teal), TbSkpZ (orange) and TbUsp7 (green). In contrast to cullin 1, we propose that TUS is responsible for removal of ubiquitin and which is supported by the decrease in abundance of *trans*-membrane domain surface proteins following individual knockdown of all TUS complex subunits.

## Methods and materials

### Trypanosoma brucei brucei culturing and transfection

Bloodstream form (BSF) Molteno Institute Trypanosomal antigen type (MITat) 1.2, derived from Lister strain 427 were cultured in HMI-11 complete medium (HMI-11 supplemented with 10% fetal bovine serum (FBS) non-heat-inactivated, 100 U/ml penicillin, 100 U/ml streptomycin) (Hirumi and Hirumi, 1994) at 37 °C with 5% CO2 in a humid atmosphere, in culture flasks with vented caps. 2T1 cells, a variant of Lister 427 (Alsford and Horn, 2008), were maintained in HMI-11 complete medium in the presence of phleomycin (0.5 μg/ml) and puromycin (1 μg/ ml). Following transfection with stem-loop RNAi plasmids or inducible overexpression plasmids, 2T1 cells were maintained in phleomycin (0.5 μg/ml) and hygromycin (2.5 μg/ml) (Alsford et al., 2005; Alsford and Horn, 2008). Experiments were performed following 48 hours induction with tetracycline (1 μg/ml). Cells were maintained at densities between 1×10^5^ and 2.5×10^6^ cells/ml.

### Recombinant DNA manipulations

Gene-specific RNAi fragments of 400–600 bp were amplified with PCR primers designed using RNAit (Redmond et al., 2004) and cloned into pRPaiSL to generate stem-loop ‘hairpin’ dsRNA to induce RNAi knockdown (Alsford and Horn, 2008). The following primers were used: TbSkp1SL_R (5’-CTAGAGATCTCCTAGGAAGAGGAGACTGTAAGAAGGC-3’), TbSkp1SL_F (5’-GATCGGGCCCGGTACCACTTTGAGCAGCGCCTACAT-3’). All constructs were verified by standard sequencing methods. TbSkpZ or TbTpr86 was C-terminally tagged using the PCR only tagging (pPOT4H) plasmid (Dean et al., 2014). Primer pares for DNA amplification: TbSkpZ Forward:

TCTTGGCGTGGAGAATGACTTCAAGGCTGAAGAAGAGGCTGAACTCAGGAAAGAGTACGGAAGAATGTCAGAGGAGAAGGGTaccGGGcccCCCctcGAG Reverse: ATCCACTAGTTCTAGAGCGGCCGCCAACATGAGGGTGTGAGGCACACTTGTTTTTGCCGATGTGCGCGTATTCGAGAACCAGGTGTGGCGCGTGTTGACGG

TbTpr86 Forward:

TGCGAGGTGCATCTTTACCCTGGGGTTGACGCATTGTTGCTGTCGTTGATGGCTTACT GCCGCTTTAATTGGGAGCAG Reverse: GTAGAACGAAACTAGATAGATAAAGTCACAACACCAGGAGCGCTGCAGTATTAGCTATTATTGTTGTTGCAGCA

### Quantitative real-time polymerase chain reaction (qRT-PCR)

1 x 10^8^ cells were harvested at 800 x g for 10 min at 4°C and washed with ice-cold PBS and quick-frozen in dry ice for 1 min. RNA was purified using the RNeasy mini kit (Qiagen) according to the manufacturer’s instructions. RNA concentration was quantified using an ND-1000 spectrophotometer and Nanodrop software (Nanodrop Technologies). qRT-PCR was performed using iQ-SYBRGreen Supermix on a MiniOpticon Real-Time PCR Detection System (Bio-Rad) and was quantified using Bio-Rad CFX Manager software (Bio-Rad).

### Transfection

3 x 10^7^ bloodstream-form cells were harvested by centrifugation at 800 x g for 10 min at 4°C. Cells were resuspended in 100 ul of Amaxa human T-cell Nucleofector solution (VPA-1002) at 4°C, mixed with 10 ug (in 5 ul) of linearised plasmid DNA and transferred to electrocuvettes. Transfection was achieved using an Amaxa Nucleofector II with Program X-001. Cells were then transferred to tube A containing 30 ml of HMI-9 medium plus any appropriate antibiotic drug for parental cell growth. Serial dilution was performed by transferring 3 ml of cell suspension from tube A into tube B containing 27 ml of HMI-9 medium and repeated again by diluting 3 ml from tube B into tube C. One millilitre aliquots from each dilution were distributed between three 24-well plates and incubated at 37°C. After 6 hours HMI-9 containing antibiotic selection was added to the wells at the desired final concentration. Transformed cells were recovered on days five to six posttransfection. For procyclic cells, 3 x 10^7^ cells per transfection were harvested at 4°C, washed in cytomix and resuspended in 400 μl cytomix. Electroporation was performed with 5–15 μg of linearised DNA using a Bio-Rad Gene Pulser II (1.5 kV and 25 μF). Cells were transferred to 9.5 ml SDM-79 medium and incubated for 16 hours after which selection antibiotics were added. The cells were then diluted into 96-well microtiter plates. Positive transformants were picked into fresh selective medium 10–15 days post transfection.

### Stable isotype-labeling by amino acids in cell culture (SILAC) labeling

HMI11 for SILAC was prepared essentially as described in (Urbaniak et al., 2012): IMDM depleted of L-Arginine, L-Lysine (Thermo) and 10% dialysed (10 kDa molecular weight cutoff) foetal bovine serum (Dundee Cell Products) was supplemented with 4 ug/ml folic acid, 110 μg/ml pyruvic acid, 39 μg/ml thymidine, 2.8 μg/ml bathocuproinedisulfonic acid, 182 μg/ml L-cysteine, 13.6 μg/ml hypoxanthine, 200 μM β-mercaptoethanol, 0.5 μg/ml phleomycin and 2.5 μg/ml hygromycin. Finally, either natural isotopic L-Arginine and L-Lysine (HMI11-R0K0), or L-Arginine ^U-13^C6 and L-Lysine ^4,4,5,5-2^H4 (HMI11-R_6_K_4_) (Cambridge Isotope Laboratories) were added at 120 uM and 240 uM respectively. RNAi was induced by addition of 1 μg/ml tetracycline. Equal numbers of induced and uninduced cells, grown in the presence of HMI11-R0K0 or HMI11-R6K4 respectively, were mixed, harvested by centrifugation, washed twice with PBS containing complete mini protease inhibitor (Roche) and resuspended in Laemmli SDS running buffer containing 1 mM dithiothreitol and stored at −80°C. TbSkpZ and TbTpr86 RNAi samples were generated in duplicate and triplicate, respectively, with each replicate representing a distinct clone. One label swap was performed in each set of replicates. The heavy isotope incorporation at steady state was determined from one gel slice (60–80 kDa) of a control experiment omitting induction. Samples were sonicated and aliquots containing 5 x 10^6^ cells were separated on a NuPAGE bis-tris 4–12% gradient polyacrylamide gel (Invitrogen) under reducing conditions. The sample lane was divided into eight slices that were excised from Coomassie stained gels, destained and then subjected to tryptic digest and reductive alkylation. Liquid chromatography tandem mass spectrometry (LC-MSMS) was performed on an UltiMate 3000 RSLCnano System (Thermo Scientific) coupled to a Q-Exactive Hybrid Quadrupole-Orbitrap (Thermo Scientific) and mass spectra analysed using MaxQuant version 1.6 (Cox et al., 2008) searching the T. brucei brucei 927 annotated protein database (release 24) from TriTrypDB (Aslett et al., 2010). Minimum peptide length was set at six amino acids, isoleucine and leucine were considered indistinguishable and false discovery rates of 0.01 were calculated at the levels of peptides, proteins and modification sites based on the number of hits against the reversed sequence database. SILAC ratios were calculated using only peptides that could be uniquely mapped to a given protein. If the identified peptide sequence set of one protein contained the peptide set of another protein, these two proteins were assigned to the same protein group. Proteomics data have been deposited to the ProteomeXchange Consortium via thePRIDE partner repository (Perez-Riverol et al., 2019) with the dataset identifiers PXD021000 (SkpZ RNAi and immunoprecipitation) and XD021797 (TbTPR86 immunoprecipitation). *Immunoprecipitation:* Three flat culinary smidgen spoons of cryo-milled protein powders, derived from Skp1-HA, Tpr-HA, or wild type *T. brucei* 427 strain PCF, was dissolved into buffer A (20mM HEPES (pH7.4), 250mM NaCl, 0.01mM CaCl2, 1mM MgCl2) with 0.1% Brj58 on ice. The sample was sonicated and centrifuged for 15min, 13000g at 4°C. The supernatant was incubated with anti-HA magnetic beads for two hours at 8°C. After three times washing using buffer A with 0.01% Brij58, the beads were incubated with one times SDS buffer and the supernatant collected for SDS-PAGE as described above for further MS analysis, except samples were run on an OrbiTrap Velos Pro (Thermo Scientific) mass spectrometer. All proteomics manipulations were performed using LoBind tubes (Eppendorf) for efficient protein extraction and removal from each tube.

### Preparation of specimens for conventional (resin-embedded) transmission electron microscopy

Cells were pelleted and fixed in 0.1 M Na cacodylate buffer (pH 7.2) containing 4% paraformaldehyde and 2.5% glutaraldehyde for 60 mins, at room temperature. Cells were stained with 1% OsO4 with 1.5% sodium ferrocyanide in cacodylate buffer for 60 min followed by 1% tannic acid in 0.1M cacodylate buffer for 1hr and 1% uranyl acetate in acetate buffer for 1hr. Fixates were dehydrated with a 50% to 100% ethanol series followed by 100% propylene oxide. Embedding in Durcupan resin followed standard procedures. Sections were cut on an ultramicrotome at 70-100nm and stained with 3% uranyl acetate followed by Reynolds lead citrate. Grids were imaged on a JEOL 1200EX TEM using an SIS camera.

### Protein electrophoresis and immunoblotting

Proteins were separated by electrophoresis on 12.5% SDS-polyacrylamide gels and then transferred to polyvinylidene difluoride (PVDF) membranes (Immobilon; Millipore) using a wet transfer tank (Hoefer Instruments). Non-specific binding was blocked with Tris-buffered saline with 0.2% Tween-20 (TBST) supplemented with 5% freeze-dried milk and proteins were detected by incubation with primary antibody diluted in TBST with 1% milk for 1 hour at room temperature. Antibodies were used at the following dilutions: mouse monoclonal anti-HA (sc-7392, Santa Cruz) at 1:10,000, rabbit monoclonal anti-myc (7E18, Sigma) at 1:5000, mouse monoclonal anti-v5 (37-7500, Invitrogen) at 1:1000. Following three washes of 10 minutes with TBST, the membrane was incubated in secondary antibody diluted in TBST with 1% milk for 1 hour at room temperature. Commercial secondary anti-rabbit peroxidase-conjugated IgG (A0545, Sigma) and anti-mouse peroxidase-conjugated IgG (A9044, Sigma) were used both at 1:10,000. Detection was by chemiluminescence with luminol (Sigma) on BioMaxMR film (Kodak). Densitometry quantification of relative protein level was achieved using ImageJ software (NIH).

### Immunofluorescence (IF)

Samples were prepared as previously described (Leung et al., 2008). Antibodies were used at the following dilutions: mouse and rabbit anti-HA epitope IgG (sc-57594 and sc-7392, Santa Cruz) at 1:1000, mouse 9E10 anti-myc at 1:1000 (Sigma), rabbit anti-ISG75 and anti-ISG65 (from P. Overath, Tubingen) at 1:1000, mouse α-EF1α (Millipore, clone CBP-KK1, 1:20,000). Secondary antibodies were used at the following dilutions: horseradish peroxidase coupled secondary antibodies (α-mouse and α-rabbit, Sigma, 1:20,000), anti-mouse Oregon Green (Molecular Probes) at 1:1000, antirabbit Cy3 (Sigma) at 1:1000. Cells were examined on a Nikon Eclipse E600 epifluorescence microscope fitted with optically-matched filter blocks and a Hamamatsu ORCA CCD camera. Digital Images were captured using Metamorph software (Universal Imaging Corp.), and raw images processed using Adobe Photoshop (Adobe Systems Inc.).

## Acknowledgements

This work was supported by grants from the Wellcome Trust (204697/Z/16/Z to MCF) and the Medical Research Council (MR/P009018/1 to MCF). We thank Alan Prescott for electron microscopy and Samantha Kosto and Kenneth Beattie from the Proteomics Facility at the University of Dundee.

## Supplementary data for

### Supplementary figure legend

**Figure S1: Alignment of trypanosome Skp1 paralogs showing predicted domain architecture.** Panel A: Amino acid sequence alignment of four Skp1-like proteins in *T. brucei.* Protein sequences were retrieved for *Trypanosoma brucei* TRU927 and aligned using Clustal W. indicates a gap introduced in the alignment, “:” indicates conservative substitution and “*” identity. Domains are overlaid with colour: POZ; orange, dimerization; purple. Panel B: Schematic diagram of TbSkp1 protein paralogs. Domain colour codes are as in panel B and amino acid length of the predicted proteins are shown at right. Open symbol in Tb927.10.14310 indicates a divergent POZ domain.

**Tables S1 and S2: Accession numbers for protein sequences included in phylogenetic analysis.**

**Table S3: Proteomics analysis tables for TbSkpZ RNAi SILAC and cryomill affinity isolations of the TUS complex.**

**Data archive 1: Output from Phyre2 prediction of TbTpr86 structure.**

## Notes

### Competing Interest Statement

The authors have declared no competing interest.

