## Supplementary material for "The endosomal TbTpr86/TbUsp7/SkpZ (TUS) complex controls surface protein abundance in trypanosomes": Figure S1

# A

```
Tb927.11.13330      ------------------------------------------------------------ 0
Tb927.10.14310      MVIFDDVENSNRVGGNEGGIRPIFVQLTPNAPLPSEEYLRLVALSKAPPPPDSEGSTLHS 60
Tb927.11.6130       ---------------------------------MAEE----------------------V 5
Tb927.10.11610      ---------------------------------MVNHLVVLIG---------SDGKRATV 18


Tb927.11.13330      ----------MAACDVTLVSPNGET----VSIPAASAWNHIG------------------ 28
Tb927.10.14310      ARNCR----ERSARVNGRRSPTKTNVRPRPPVTATQKWGDNAMSLINSKEEAVQRHEFA- 115
Tb927.11.6130       S---VQDVDFTEYCILESNDPTPVKF--KVRREAAM-MSGLLKDMLEDQNGGDPIIPIPN 59
Tb927.10.11610      SRTAATHASLLLRDLLDGREPNTTVVEGKSGVEHVS-NSDIFEGLVADKEVAVPAIEIPF 77
                                       .*.            .   .

Tb927.11.13330      ------LLQRL--------------------------------------------AELDE 38
Tb927.10.14310      LHVRGARYCRLLETMLDSSDLQLEPLHYDPKDPNMGRRPNGTEGNPANSLPPVILPQATQ 175
Tb927.11.6130       VSART------------------------------------------------------- 64
Tb927.10.11610      THATGSLLQRICAH---------MTYRYDYRPPNGG-----VEG-STVLFEPVAREHRAT 122


Tb927.11.13330      SGCTEMPEIELGFSTEVLRAVADYLEKSSRFVIDARRPLTGSLRDFVPEWNLDFVRGA-- 96
Tb927.10.14310      KGCVAL----FTYLDLITKRVPSML------SKPLR----APLEELVQPWELEFLLYSCM 221
Tb927.11.6130       LKLV------IKYMEHHHKERADPI------EKPLK----SNIEKIISPWDHDFLYTELV 108
Tb927.10.11610      FSCG------GGYGRNVMPEIPRPM------VLPLVDYLDSFDKRFIEDWDEI------- 163
                                :           :               .  . ::  *:

Tb927.11.13330      -----------------------------------------ERREILMPLFECACFLCVV 115
Tb927.10.14310      GDDVRNAITHIDEDGTADKRGRAPTNDTRPSYYSTILEKAPRSVDLLLEVALLSDFLLVE 281
Tb927.11.6130       KDH------------------------------------DEKQHEVLIDVIMAANFLNVR 132
Tb927.10.11610      ---------------------------------------------TTVQMVKLATLLNYE 178
                                                                   : :   : :*

Tb927.11.13330      ELRELLGAYVAERVNEIAKEAPSIMEGAARLRTYLQLENEWTEEETEHLEQEMRYAKQVD 175
Tb927.10.14310      PLRQLTCAFIASLALNA--------SSEEELLQLSGLQRPMTEDELEPLYYQFPFLRPDN 333
Tb927.11.6130       DLLDLTCACVANMIRGK---------SAEQIRELFNIESDFTPEEEEKIREENRWCEEA- 182
Tb927.10.11610      ELLNLASAKLAVYLSEK---------SIEGLRAFLGVENDFKAEEEAELRKEYGRMSEEK 229
                     * :*  * :*               .   :     ::   . :*   :  :

Tb927.11.13330      PFAY 179
Tb927.10.14310      HAKN 337
Tb927.11.6130       ---- 182
Tb927.10.11610      ---- 229
```

# B

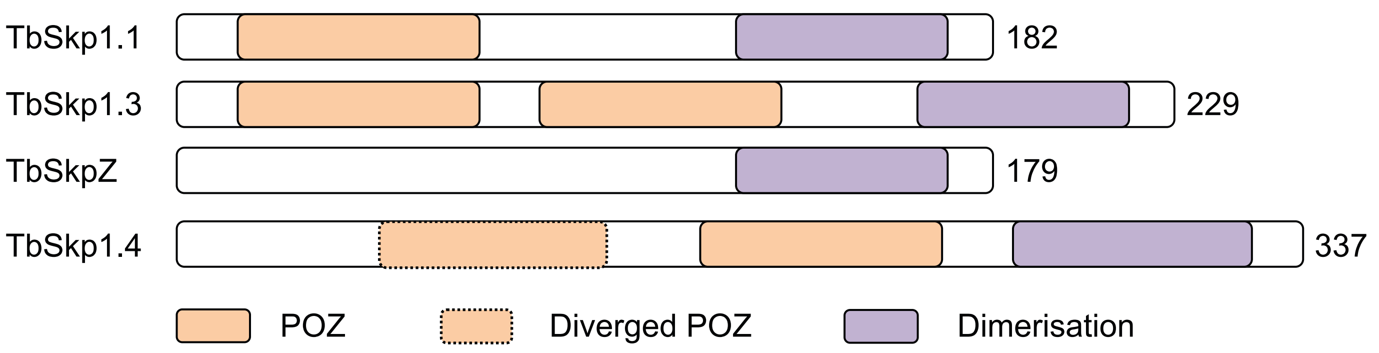
