## Supplementary material for "The endosomal TbTpr86/TbUsp7/SkpZ (TUS) complex controls surface protein abundance in trypanosomes": DA1: dom_report 2.pdf

| | |
|---|---|
| Email | |
| Description | Undefined |
| Date | Tue Dec 31 14:42:05 GMT 2019 |
| Unique Job ID | 55c641d1d7bceb76 |

### Domain analysis

| Rank | Aligned region |
|---|---|
| 1 | c4m57A_ |
| 2 | c3jd5e_ |
| 3 | c5i9fA_ |
| 4 | c5iwwD_ |
| 5 | c4g25A_ |
| 6 | c5dizB_ |
| 7 | c4wslA_ |
| 8 | c6nf8o_ |
| 9 | c3jd5o_ |
| 10 | c6neqo_ |
| 11 | c4n2sA_ |
| 12 | c4leuA_ |
| 13 | c4xgmA_ |
| 14 | c3eiqC_ |
| 15 | c3spaA_ |
| 16 | c4uzyA_ |
| 17 | c5izwA_ |
| 18 | c5mqfO_ |
| 19 | c2xpiA_ |
| 20 | c4ui9C_ |
| 21 | |
| 22 | |
| 23 | |
| 24 | |
| 25 | |
| 26 | |
| 27 | |
| 28 | |
| 29 | |
| 30 | |
| 31 | |
| 32 | |
| 33 | |
| 34 | |
| 35 | |
| 36 | |
| 37 | |
| 38 | |
| 39 | |
| 40 | |
| 41 | |
| 42 | |
| 43 | |
| 44 | |
| 45 | |
| 46 | |
| 47 | |
| 48 | |
| 49 | |
| 50 | |
| 51 | |
| 52 | |
| 53 | |
| 54 | |
| 55 | |
| 56 | |
| 57 | |
| 58 | |

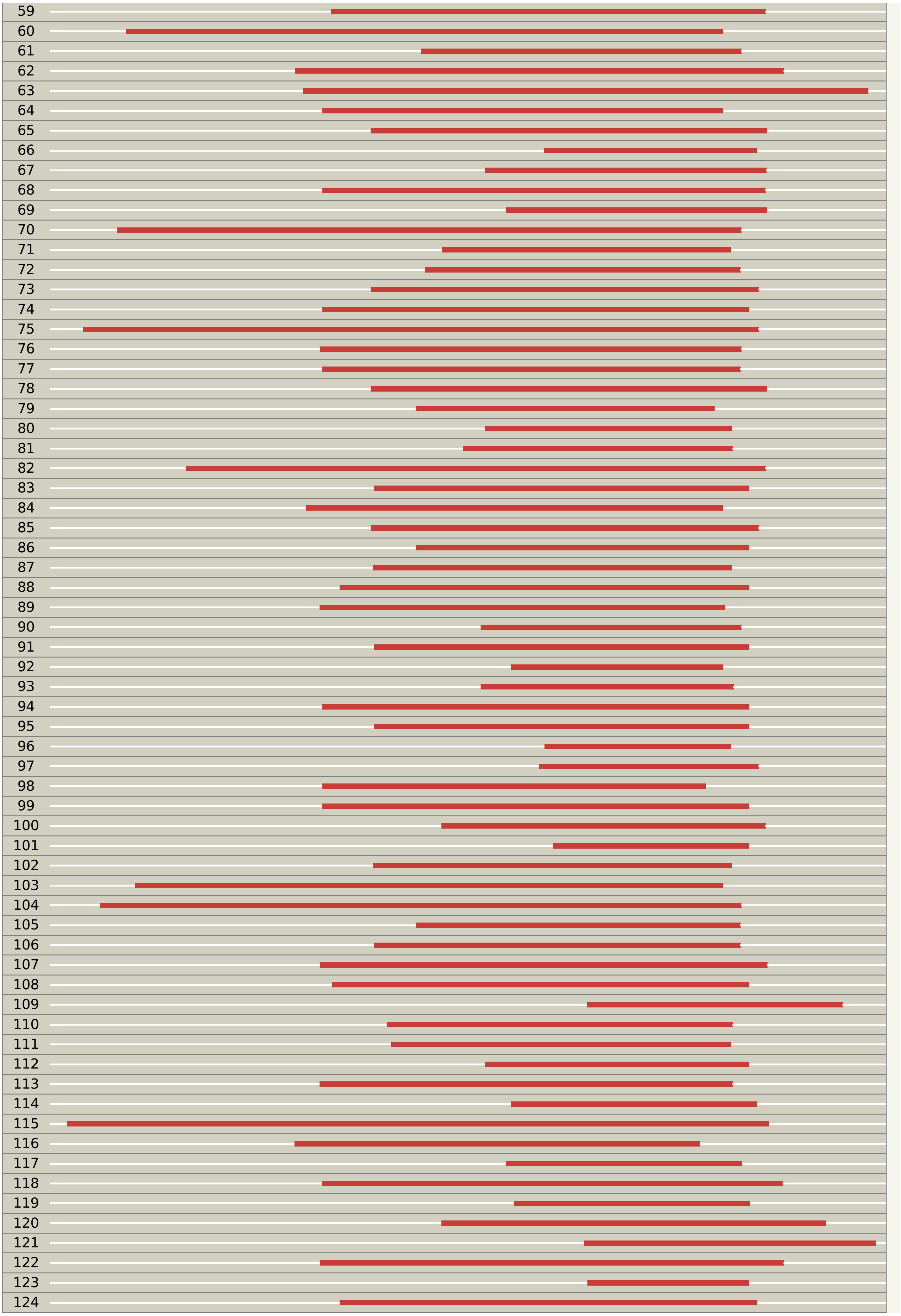

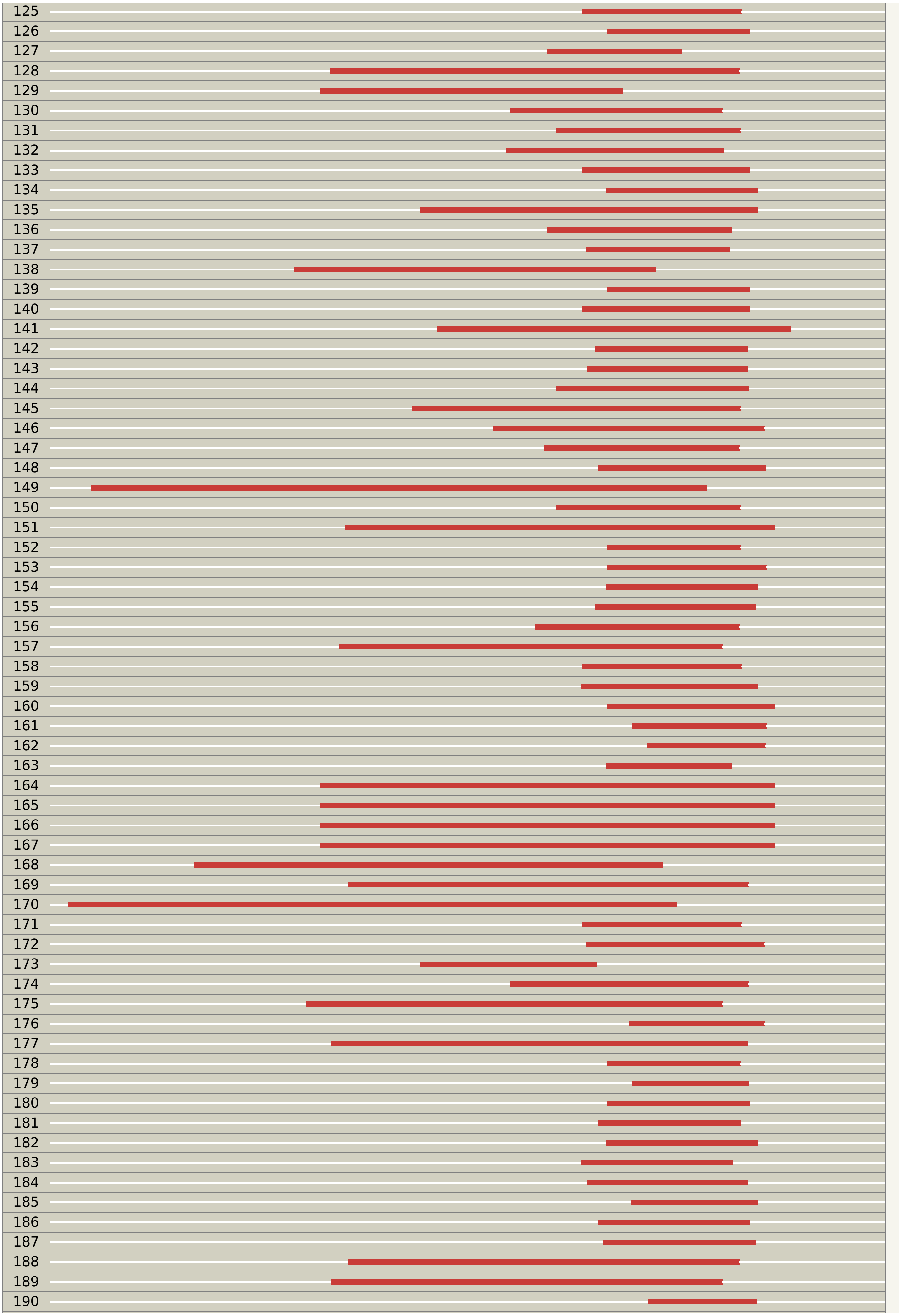

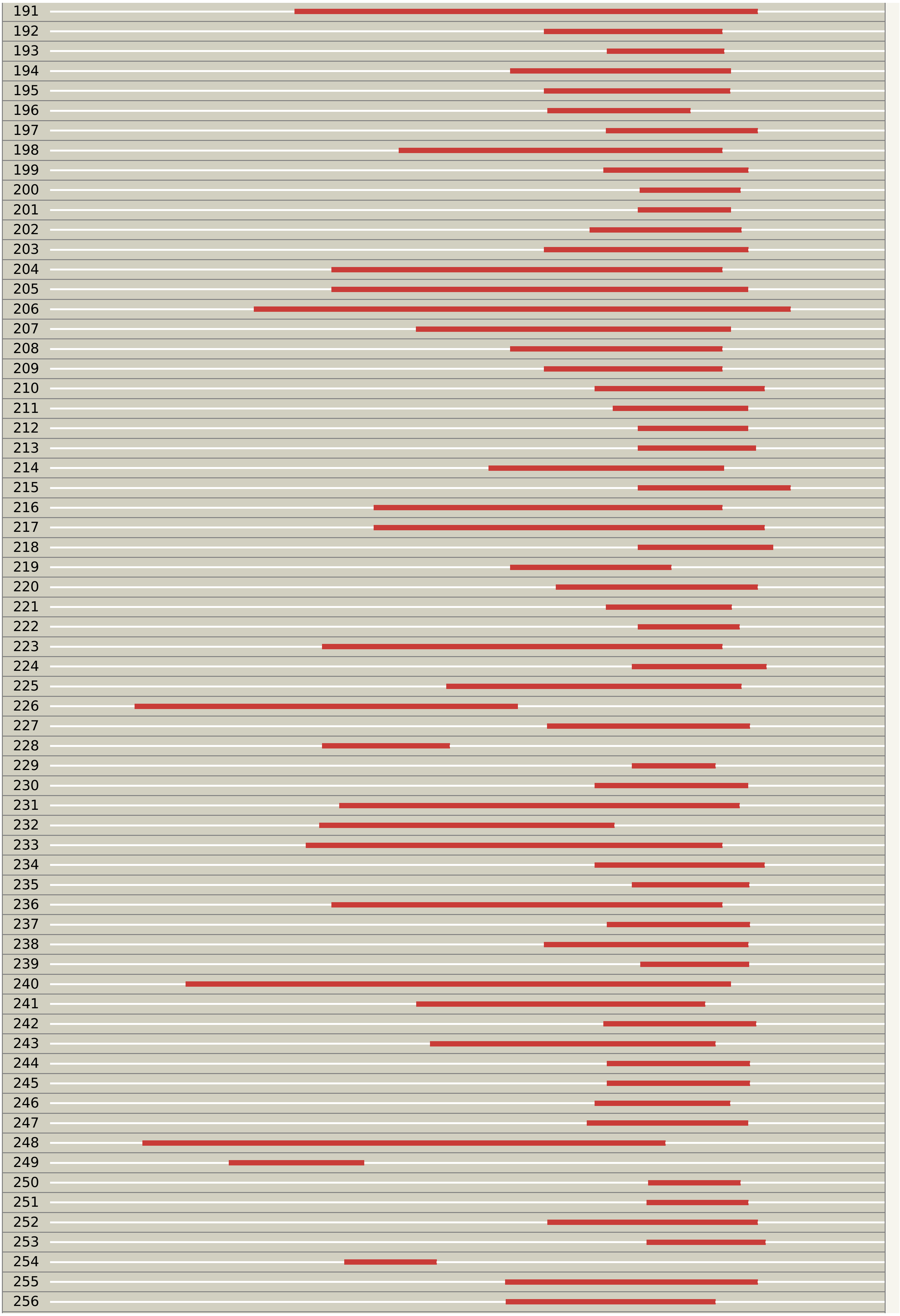

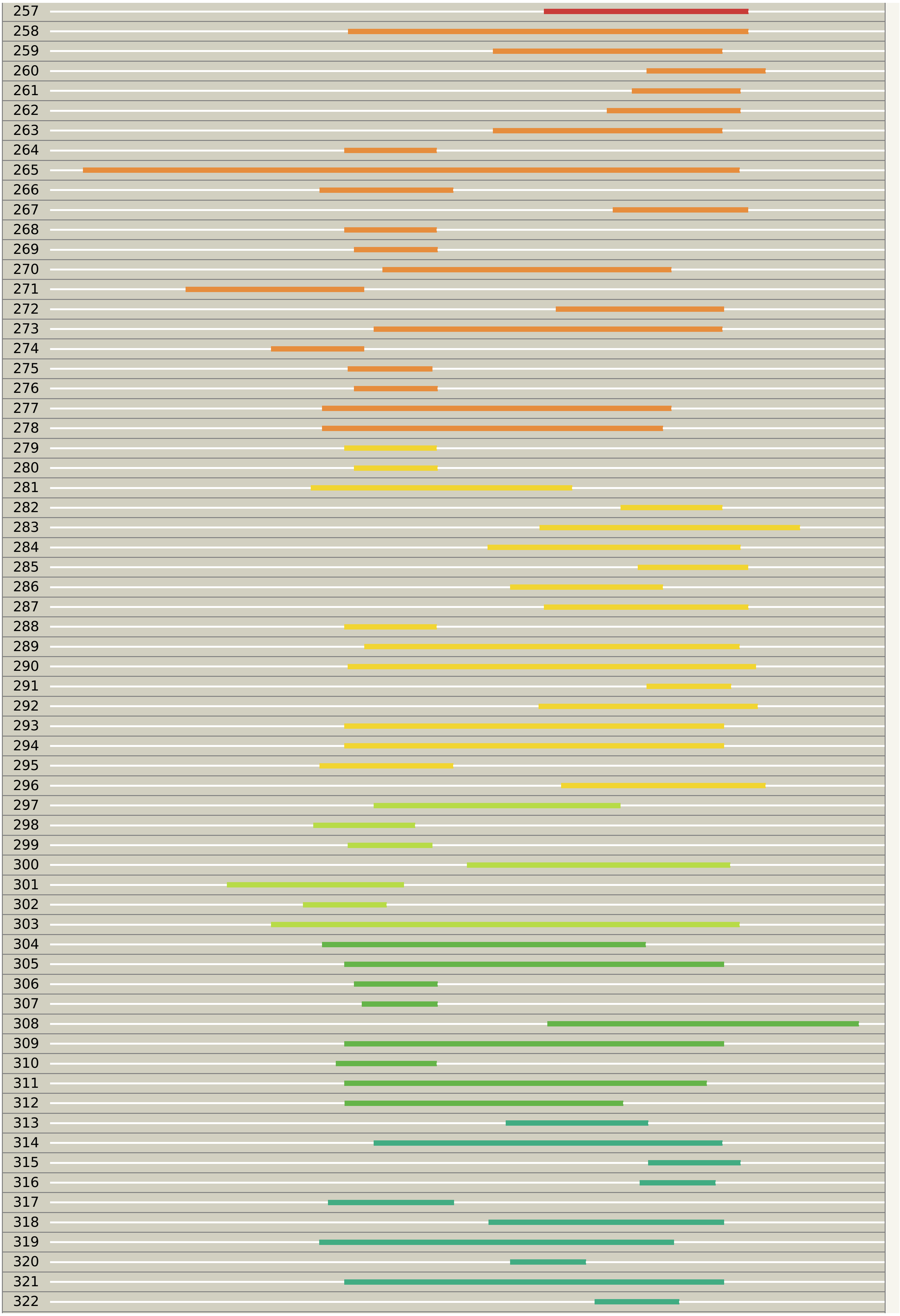

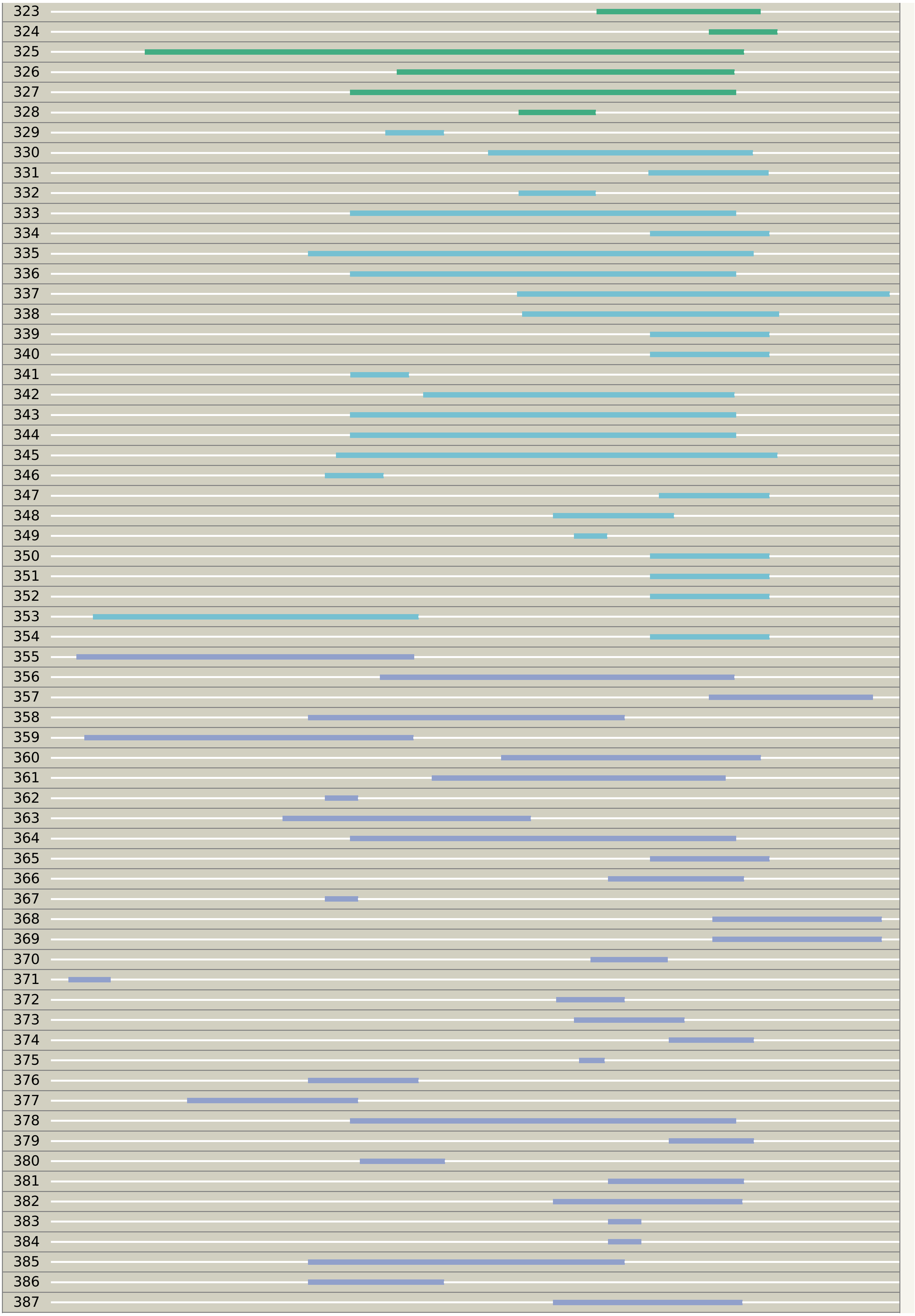
