## Supplementary material for "The endosomal TbTpr86/TbUsp7/SkpZ (TUS) complex controls surface protein abundance in trypanosomes": DA1: dom_summary.html

Phyre 2 Results for Undefined

|  |  |
| --- | --- |
| Email | |
| Description | Undefined |
| Date | Tue Dec 31 14:42:05 GMT 2019 || Unique Job ID | 55c641d1d7bceb76 |

|  |
| --- |
| Domain analysis |

| Rank | Aligned region |
| --- | --- |

|  |  |
| --- | --- |
| 1 | --- \| c4m57A\_ \| --- |
| 2 | --- \| c3jd5e\_ \| --- |
| 3 | --- \| c5i9fA\_ \| --- |
| 4 | --- \| c5iwwD\_ \| --- |
| 5 | --- \| c4g25A\_ \| --- |
| 6 | --- \| c5dizB\_ \| --- |
| 7 | --- \| c4wslA\_ \| --- |
| 8 | --- \| c6nf8o\_ \| --- |
| 9 | --- \| c3jd5o\_ \| --- |
| 10 | --- \| c6neqo\_ \| --- |
| 11 | --- \| c4n2sA\_ \| --- |
| 12 | --- \| c4leuA\_ \| --- |
| 13 | --- \| c4xgmA\_ \| --- |
| 14 | --- \| c3eiqC\_ \| --- |
| 15 | --- \| c3spaA\_ \| --- |
| 16 | --- \| c4uzyA\_ \| --- |
| 17 | --- \| c5izwA\_ \| --- |
| 18 | --- \| c5mqfO\_ \| --- |
| 19 | --- \| c2xpiA\_ \| --- |
| 20 | --- \| c4ui9C\_ \| --- |
| 21 | --- \| --- \| --- |
| 22 | --- \| --- \| --- |
| 23 | --- \| --- \| --- |
| 24 | --- \| --- \| --- |
| 25 | --- \| --- \| --- |
| 26 | --- \| --- \| --- |
| 27 | --- \| --- \| --- |
| 28 | --- \| --- \| --- |
| 29 | --- \| --- \| --- |
| 30 | --- \| --- \| --- |
| 31 | --- \| --- \| --- |
| 32 | --- \| --- \| --- |
| 33 | --- \| --- \| --- |
| 34 | --- \| --- \| --- |
| 35 | --- \| --- \| --- |
| 36 | --- \| --- \| --- |
| 37 | --- \| --- \| --- |
| 38 | --- \| --- \| --- |
| 39 | --- \| --- \| --- |
| 40 | --- \| --- \| --- |
| 41 | --- \| --- \| --- |
| 42 | --- \| --- \| --- |
| 43 | --- \| --- \| --- |
| 44 | --- \| --- \| --- |
| 45 | --- \| --- \| --- |
| 46 | --- \| --- \| --- |
| 47 | --- \| --- \| --- |
| 48 | --- \| --- \| --- |
| 49 | --- \| --- \| --- |
| 50 | --- \| --- \| --- |
| 51 | --- \| --- \| --- |
| 52 | --- \| --- \| --- |
| 53 | --- \| --- \| --- |
| 54 | --- \| --- \| --- |
| 55 | --- \| --- \| --- |
| 56 | --- \| --- \| --- |
| 57 | --- \| --- \| --- |
| 58 | --- \| --- \| --- |
| 59 | --- \| --- \| --- |
| 60 | --- \| --- \| --- |
| 61 | --- \| --- \| --- |
| 62 | --- \| --- \| --- |
| 63 | --- \| --- \| --- |
| 64 | --- \| --- \| --- |
| 65 | --- \| --- \| --- |
| 66 | --- \| --- \| --- |
| 67 | --- \| --- \| --- |
| 68 | --- \| --- \| --- |
| 69 | --- \| --- \| --- |
| 70 | --- \| --- \| --- |
| 71 | --- \| --- \| --- |
| 72 | --- \| --- \| --- |
| 73 | --- \| --- \| --- |
| 74 | --- \| --- \| --- |
| 75 | --- \| --- \| --- |
| 76 | --- \| --- \| --- |
| 77 | --- \| --- \| --- |
| 78 | --- \| --- \| --- |
| 79 | --- \| --- \| --- |
| 80 | --- \| --- \| --- |
| 81 | --- \| --- \| --- |
| 82 | --- \| --- \| --- |
| 83 | --- \| --- \| --- |
| 84 | --- \| --- \| --- |
| 85 | --- \| --- \| --- |
| 86 | --- \| --- \| --- |
| 87 | --- \| --- \| --- |
| 88 | --- \| --- \| --- |
| 89 | --- \| --- \| --- |
| 90 | --- \| --- \| --- |
| 91 | --- \| --- \| --- |
| 92 | --- \| --- \| --- |
| 93 | --- \| --- \| --- |
| 94 | --- \| --- \| --- |
| 95 | --- \| --- \| --- |
| 96 | --- \| --- \| --- |
| 97 | --- \| --- \| --- |
| 98 | --- \| --- \| --- |
| 99 | --- \| --- \| --- |
| 100 | --- \| --- \| --- |
| 101 | --- \| --- \| --- |
| 102 | --- \| --- \| --- |
| 103 | --- \| --- \| --- |
| 104 | --- \| --- \| --- |
| 105 | --- \| --- \| --- |
| 106 | --- \| --- \| --- |
| 107 | --- \| --- \| --- |
| 108 | --- \| --- \| --- |
| 109 | --- \| --- \| --- |
| 110 | --- \| --- \| --- |
| 111 | --- \| --- \| --- |
| 112 | --- \| --- \| --- |
| 113 | --- \| --- \| --- |
| 114 | --- \| --- \| --- |
| 115 | --- \| --- \| --- |
| 116 | --- \| --- \| --- |
| 117 | --- \| --- \| --- |
| 118 | --- \| --- \| --- |
| 119 | --- \| --- \| --- |
| 120 | --- \| --- \| --- |
| 121 | --- \| --- \| --- |
| 122 | --- \| --- \| --- |
| 123 | --- \| --- \| --- |
| 124 | --- \| --- \| --- |
| 125 | --- \| --- \| --- |
| 126 | --- \| --- \| --- |
| 127 | --- \| --- \| --- |
| 128 | --- \| --- \| --- |
| 129 | --- \| --- \| --- |
| 130 | --- \| --- \| --- |
| 131 | --- \| --- \| --- |
| 132 | --- \| --- \| --- |
| 133 | --- \| --- \| --- |
| 134 | --- \| --- \| --- |
| 135 | --- \| --- \| --- |
| 136 | --- \| --- \| --- |
| 137 | --- \| --- \| --- |
| 138 | --- \| --- \| --- |
| 139 | --- \| --- \| --- |
| 140 | --- \| --- \| --- |
| 141 | --- \| --- \| --- |
| 142 | --- \| --- \| --- |
| 143 | --- \| --- \| --- |
| 144 | --- \| --- \| --- |
| 145 | --- \| --- \| --- |
| 146 | --- \| --- \| --- |
| 147 | --- \| --- \| --- |
| 148 | --- \| --- \| --- |
| 149 | --- \| --- \| --- |
| 150 | --- \| --- \| --- |
| 151 | --- \| --- \| --- |
| 152 | --- \| --- \| --- |
| 153 | --- \| --- \| --- |
| 154 | --- \| --- \| --- |
| 155 | --- \| --- \| --- |
| 156 | --- \| --- \| --- |
| 157 | --- \| --- \| --- |
| 158 | --- \| --- \| --- |
| 159 | --- \| --- \| --- |
| 160 | --- \| --- \| --- |
| 161 | --- \| --- \| --- |
| 162 | --- \| --- \| --- |
| 163 | --- \| --- \| --- |
| 164 | --- \| --- \| --- |
| 165 | --- \| --- \| --- |
| 166 | --- \| --- \| --- |
| 167 | --- \| --- \| --- |
| 168 | --- \| --- \| --- |
| 169 | --- \| --- \| --- |
| 170 | --- \| --- \| --- |
| 171 | --- \| --- \| --- |
| 172 | --- \| --- \| --- |
| 173 | --- \| --- \| --- |
| 174 | --- \| --- \| --- |
| 175 | --- \| --- \| --- |
| 176 | --- \| --- \| --- |
| 177 | --- \| --- \| --- |
| 178 | --- \| --- \| --- |
| 179 | --- \| --- \| --- |
| 180 | --- \| --- \| --- |
| 181 | --- \| --- \| --- |
| 182 | --- \| --- \| --- |
| 183 | --- \| --- \| --- |
| 184 | --- \| --- \| --- |
| 185 | --- \| --- \| --- |
| 186 | --- \| --- \| --- |
| 187 | --- \| --- \| --- |
| 188 | --- \| --- \| --- |
| 189 | --- \| --- \| --- |
| 190 | --- \| --- \| --- |
| 191 | --- \| --- \| --- |
| 192 | --- \| --- \| --- |
| 193 | --- \| --- \| --- |
| 194 | --- \| --- \| --- |
| 195 | --- \| --- \| --- |
| 196 | --- \| --- \| --- |
| 197 | --- \| --- \| --- |
| 198 | --- \| --- \| --- |
| 199 | --- \| --- \| --- |
| 200 | --- \| --- \| --- |
| 201 | --- \| --- \| --- |
| 202 | --- \| --- \| --- |
| 203 | --- \| --- \| --- |
| 204 | --- \| --- \| --- |
| 205 | --- \| --- \| --- |
| 206 | --- \| --- \| --- |
| 207 | --- \| --- \| --- |
| 208 | --- \| --- \| --- |
| 209 | --- \| --- \| --- |
| 210 | --- \| --- \| --- |
| 211 | --- \| --- \| --- |
| 212 | --- \| --- \| --- |
| 213 | --- \| --- \| --- |
| 214 | --- \| --- \| --- |
| 215 | --- \| --- \| --- |
| 216 | --- \| --- \| --- |
| 217 | --- \| --- \| --- |
| 218 | --- \| --- \| --- |
| 219 | --- \| --- \| --- |
| 220 | --- \| --- \| --- |
| 221 | --- \| --- \| --- |
| 222 | --- \| --- \| --- |
| 223 | --- \| --- \| --- |
| 224 | --- \| --- \| --- |
| 225 | --- \| --- \| --- |
| 226 | --- \| --- \| --- |
| 227 | --- \| --- \| --- |
| 228 | --- \| --- \| --- |
| 229 | --- \| --- \| --- |
| 230 | --- \| --- \| --- |
| 231 | --- \| --- \| --- |
| 232 | --- \| --- \| --- |
| 233 | --- \| --- \| --- |
| 234 | --- \| --- \| --- |
| 235 | --- \| --- \| --- |
| 236 | --- \| --- \| --- |
| 237 | --- \| --- \| --- |
| 238 | --- \| --- \| --- |
| 239 | --- \| --- \| --- |
| 240 | --- \| --- \| --- |
| 241 | --- \| --- \| --- |
| 242 | --- \| --- \| --- |
| 243 | --- \| --- \| --- |
| 244 | --- \| --- \| --- |
| 245 | --- \| --- \| --- |
| 246 | --- \| --- \| --- |
| 247 | --- \| --- \| --- |
| 248 | --- \| --- \| --- |
| 249 | --- \| --- \| --- |
| 250 | --- \| --- \| --- |
| 251 | --- \| --- \| --- |
| 252 | --- \| --- \| --- |
| 253 | --- \| --- \| --- |
| 254 | --- \| --- \| --- |
| 255 | --- \| --- \| --- |
| 256 | --- \| --- \| --- |
| 257 | --- \| --- \| --- |
| 258 | --- \| --- \| --- |
| 259 | --- \| --- \| --- |
| 260 | --- \| --- \| --- |
| 261 | --- \| --- \| --- |
| 262 | --- \| --- \| --- |
| 263 | --- \| --- \| --- |
| 264 | --- \| --- \| --- |
| 265 | --- \| --- \| --- |
| 266 | --- \| --- \| --- |
| 267 | --- \| --- \| --- |
| 268 | --- \| --- \| --- |
| 269 | --- \| --- \| --- |
| 270 | --- \| --- \| --- |
| 271 | --- \| --- \| --- |
| 272 | --- \| --- \| --- |
| 273 | --- \| --- \| --- |
| 274 | --- \| --- \| --- |
| 275 | --- \| --- \| --- |
| 276 | --- \| --- \| --- |
| 277 | --- \| --- \| --- |
| 278 | --- \| --- \| --- |
| 279 | --- \| --- \| --- |
| 280 | --- \| --- \| --- |
| 281 | --- \| --- \| --- |
| 282 | --- \| --- \| --- |
| 283 | --- \| --- \| --- |
| 284 | --- \| --- \| --- |
| 285 | --- \| --- \| --- |
| 286 | --- \| --- \| --- |
| 287 | --- \| --- \| --- |
| 288 | --- \| --- \| --- |
| 289 | --- \| --- \| --- |
| 290 | --- \| --- \| --- |
| 291 | --- \| --- \| --- |
| 292 | --- \| --- \| --- |
| 293 | --- \| --- \| --- |
| 294 | --- \| --- \| --- |
| 295 | --- \| --- \| --- |
| 296 | --- \| --- \| --- |
| 297 | --- \| --- \| --- |
| 298 | --- \| --- \| --- |
| 299 | --- \| --- \| --- |
| 300 | --- \| --- \| --- |
| 301 | --- \| --- \| --- |
| 302 | --- \| --- \| --- |
| 303 | --- \| --- \| --- |
| 304 | --- \| --- \| --- |
| 305 | --- \| --- \| --- |
| 306 | --- \| --- \| --- |
| 307 | --- \| --- \| --- |
| 308 | --- \| --- \| --- |
| 309 | --- \| --- \| --- |
| 310 | --- \| --- \| --- |
| 311 | --- \| --- \| --- |
| 312 | --- \| --- \| --- |
| 313 | --- \| --- \| --- |
| 314 | --- \| --- \| --- |
| 315 | --- \| --- \| --- |
| 316 | --- \| --- \| --- |
| 317 | --- \| --- \| --- |
| 318 | --- \| --- \| --- |
| 319 | --- \| --- \| --- |
| 320 | --- \| --- \| --- |
| 321 | --- \| --- \| --- |
| 322 | --- \| --- \| --- |
| 323 | --- \| --- \| --- |
| 324 | --- \| --- \| --- |
| 325 | --- \| --- \| --- |
| 326 | --- \| --- \| --- |
| 327 | --- \| --- \| --- |
| 328 | --- \| --- \| --- |
| 329 | --- \| --- \| --- |
| 330 | --- \| --- \| --- |
| 331 | --- \| --- \| --- |
| 332 | --- \| --- \| --- |
| 333 | --- \| --- \| --- |
| 334 | --- \| --- \| --- |
| 335 | --- \| --- \| --- |
| 336 | --- \| --- \| --- |
| 337 | --- \| --- \| --- |
| 338 | --- \| --- \| --- |
| 339 | --- \| --- \| --- |
| 340 | --- \| --- \| --- |
| 341 | --- \| --- \| --- |
| 342 | --- \| --- \| --- |
| 343 | --- \| --- \| --- |
| 344 | --- \| --- \| --- |
| 345 | --- \| --- \| --- |
| 346 | --- \| --- \| --- |
| 347 | --- \| --- \| --- |
| 348 | --- \| --- \| --- |
| 349 | --- \| --- \| --- |
| 350 | --- \| --- \| --- |
| 351 | --- \| --- \| --- |
| 352 | --- \| --- \| --- |
| 353 | --- \| --- \| --- |
| 354 | --- \| --- \| --- |
| 355 | --- \| --- \| --- |
| 356 | --- \| --- \| --- |
| 357 | --- \| --- \| --- |
| 358 | --- \| --- \| --- |
| 359 | --- \| --- \| --- |
| 360 | --- \| --- \| --- |
| 361 | --- \| --- \| --- |
| 362 | --- \| --- \| --- |
| 363 | --- \| --- \| --- |
| 364 | --- \| --- \| --- |
| 365 | --- \| --- \| --- |
| 366 | --- \| --- \| --- |
| 367 | --- \| --- \| --- |
| 368 | --- \| --- \| --- |
| 369 | --- \| --- \| --- |
| 370 | --- \| --- \| --- |
| 371 | --- \| --- \| --- |
| 372 | --- \| --- \| --- |
| 373 | --- \| --- \| --- |
| 374 | --- \| --- \| --- |
| 375 | --- \| --- \| --- |
| 376 | --- \| --- \| --- |
| 377 | --- \| --- \| --- |
| 378 | --- \| --- \| --- |
| 379 | --- \| --- \| --- |
| 380 | --- \| --- \| --- |
| 381 | --- \| --- \| --- |
| 382 | --- \| --- \| --- |
| 383 | --- \| --- \| --- |
| 384 | --- \| --- \| --- |
| 385 | --- \| --- \| --- |
| 386 | --- \| --- \| --- |
| 387 | --- \| --- \| --- |

  
  
  
