## Supplementary material for "The endosomal TbTpr86/TbUsp7/SkpZ (TUS) complex controls surface protein abundance in trypanosomes": DA1: hit_summary 3.html

Phyre 2 Results for Undefined

|  |  |
| --- | --- |
| Email | |
| Description | Undefined |
| Date | Tue Dec 31 14:42:05 GMT 2019 || Unique Job ID | 55c641d1d7bceb76 |

|  |
| --- |
| Detailed template information |

| # | Template | Alignment Coverage | 3D Model | Confidence | % i.d. | Template Information |
| --- | --- | --- | --- | --- | --- | --- |

|  |  |  |  |  |  |  |
| --- | --- | --- | --- | --- | --- | --- |
| 1 | c4m57A\_ | --- \| --- \| --- |  | 100.0 | 17 | **PDB header:**rna binding protein **Chain:** A: **PDB Molecule:**chloroplast pentatricopeptide repeat protein 10;  **PDBTitle:** crystal structure of the pentatricopeptide repeat protein ppr10 from2 maize |
| 2 | c3jd5e\_ | --- \| --- \| --- |  | 100.0 | 12 | **PDB header:**ribosome **Chain:** E: **PDB Molecule:**28s ribosomal protein s5, mitochondrial;  **PDBTitle:** cryo-em structure of the small subunit of the mammalian mitochondrial2 ribosome |
| 3 | c5i9fA\_ | --- \| --- \| --- |  | 100.0 | 18 | **PDB header:**rna binding protein/rna **Chain:** A: **PDB Molecule:**pentatricopeptide repeat protein dppr-u10;  **PDBTitle:** crystal structure of designed pentatricopeptide repeat protein dppr-2 u10 in complex with its target rna u10 |
| 4 | c5iwwD\_ | --- \| --- \| --- |  | 100.0 | 13 | **PDB header:**rna binding protein **Chain:** D: **PDB Molecule:**pls9-ppr;  **PDBTitle:** crystal structure of rna editing factor of designer pls-type ppr/9r2 protein in complex with morf9/rip9 |
| 5 | c4g25A\_ | --- \| --- \| --- |  | 100.0 | 14 | **PDB header:**rna binding protein **Chain:** A: **PDB Molecule:**pentatricopeptide repeat-containing protein at2g32230,  **PDBTitle:** crystal structure of proteinaceous rnase p 1 (prorp1) from a.2 thaliana, semet substituted form with sr |
| 6 | c5dizB\_ | --- \| --- \| --- |  | 100.0 | 13 | **PDB header:**hydrolase **Chain:** B: **PDB Molecule:**proteinaceous rnase p 2;  **PDBTitle:** crystal structure of nuclear proteinaceous rnase p 2 (prorp2) from a.2 thaliana |
| 7 | c4wslA\_ | --- \| --- \| --- |  | 100.0 | 18 | **PDB header:**de novo protein **Chain:** A: **PDB Molecule:**pentatricopeptide repeat protein;  **PDBTitle:** crystal structure of designed cppr-polyc protein |
| 8 | c6nf8o\_ | --- \| --- \| --- |  | 100.0 | 9 | **PDB header:**ribosomal protein **Chain:** O: **PDB Molecule:**28s ribosomal protein s15, mitochondrial;  **PDBTitle:** structure of human mitochondrial translation initiation factor 3 bound2 to the small ribosomal subunit -class i |
| 9 | c3jd5o\_ | --- \| --- \| --- |  | 100.0 | 9 | **PDB header:**ribosome **Chain:** O: **PDB Molecule:**28s ribosomal protein s15, mitochondrial;  **PDBTitle:** cryo-em structure of the small subunit of the mammalian mitochondrial2 ribosome |
| 10 | c6neqo\_ | --- \| --- \| --- |  | 100.0 | 10 | **PDB header:**ribosomal protein **Chain:** O: **PDB Molecule:**28s ribosomal protein s15, mitochondrial;  **PDBTitle:** structure of human mitochondrial translation initiation factor 3 bound2 to the small ribosomal subunit-class-ii |
| 11 | c4n2sA\_ | --- \| --- \| --- |  | 100.0 | 10 | **PDB header:**splicing/rna **Chain:** A: **PDB Molecule:**tha8 rna binding protein;  **PDBTitle:** crystal structure of tha8 in complex with zm1a-6 rna |
| 12 | c4leuA\_ | --- \| --- \| --- |  | 99.9 | 11 | **PDB header:**rna binding protein **Chain:** A: **PDB Molecule:**pentatricopeptide repeat-containing protein at3g46870;  **PDBTitle:** crystal structure of tha8-like protein from arabidopsis thaliana |
| 13 | c4xgmA\_ | --- \| --- \| --- |  | 99.9 | 11 | **PDB header:**hydrolase **Chain:** A: **PDB Molecule:**mitochondrial ribonuclease p protein 3;  **PDBTitle:** structure of the nuclease subunit of human mitochondrial rnase p2 (mrpp3) at 1.98a |
| 14 | c3eiqC\_ | --- \| --- \| --- |  | 99.9 | 10 | **PDB header:**hydrolase/antitumor protein **Chain:** C: **PDB Molecule:**programmed cell death protein 4;  **PDBTitle:** crystal structure of pdcd4-eif4a |
| 15 | c3spaA\_ | --- \| --- \| --- |  | 99.9 | 9 | **PDB header:**transferase **Chain:** A: **PDB Molecule:**dna-directed rna polymerase, mitochondrial;  **PDBTitle:** crystal structure of human mitochondrial rna polymerase |
| 16 | c4uzyA\_ | --- \| --- \| --- |  | 99.9 | 11 | **PDB header:**motor protein **Chain:** A: **PDB Molecule:**flagellar associated protein;  **PDBTitle:** crystal structure of the chlamydomonas ift70 and ift52 complex |
| 17 | c5izwA\_ | --- \| --- \| --- |  | 99.8 | 17 | **PDB header:**rna binding protein **Chain:** A: **PDB Molecule:**pls9-ppr;  **PDBTitle:** crystal structure of rna editing specific factor of designer pls-type2 ppr-9r protein |
| 18 | c5mqfO\_ | --- \| --- \| --- |  | 99.8 | 10 | **PDB header:**splicing **Chain:** O: **PDB Molecule:**crooked neck-like protein 1;  **PDBTitle:** cryo-em structure of a human spliceosome activated for step 2 of2 splicing (c\* complex) |
| 19 | c2xpiA\_ | --- \| --- \| --- |  | 99.8 | 9 | **PDB header:**cell cycle **Chain:** A: **PDB Molecule:**anaphase-promoting complex subunit cut9;  **PDBTitle:** crystal structure of apc/c hetero-tetramer cut9-hcn1 |
| 20 | c4ui9C\_ | --- \| --- \| --- |  | 99.8 | 9 | **PDB header:**cell cycle **Chain:** C: **PDB Molecule:**cell division cycle protein 23 homolog;  **PDBTitle:** atomic structure of the human anaphase-promoting complex |
| 21 | c5mqfM\_ | --- \| --- \| --- | not modelled | 99.8 | 12 | **PDB header:**splicing **Chain:** M: **PDB Molecule:**pre-mrna-splicing factor syf1;  **PDBTitle:** cryo-em structure of a human spliceosome activated for step 2 of2 splicing (c\* complex) |
| 22 | c5o9zG\_ | --- \| --- \| --- | not modelled | 99.8 | 10 | **PDB header:**splicing **Chain:** G: **PDB Molecule:**pre-mrna-processing factor 6;  **PDBTitle:** cryo-em structure of a pre-catalytic human spliceosome primed for2 activation (b complex) |
| 23 | c4ui9K\_ | --- \| --- \| --- | not modelled | 99.7 | 9 | **PDB header:**cell cycle **Chain:** K: **PDB Molecule:**cell division cycle protein 16 homolog;  **PDBTitle:** atomic structure of the human anaphase-promoting complex |
| 24 | c6q6hK\_ | --- \| --- \| --- | not modelled | 99.7 | 9 | **PDB header:**cell cycle **Chain:** K: **PDB Molecule:**cell division cycle protein 16 homolog;  **PDBTitle:** cryo-em structure of the apc/c-cdc20-cdk2-cyclina2-cks2 complex, the2 d2 box class |
| 25 | d2ooea1 | --- \| --- \| --- | not modelled | 99.7 | 8 | **Fold:**alpha-alpha superhelix **Superfamily:**TPR-like **Family:**HAT/Suf repeat |
| 26 | c6ff7M\_ | --- \| --- \| --- | not modelled | 99.6 | 12 | **PDB header:**splicing **Chain:** M: **PDB Molecule:**pre-mrna-splicing factor syf1;  **PDBTitle:** human bact spliceosome core structure |
| 27 | c4hnxA\_ | --- \| --- \| --- | not modelled | 99.6 | 9 | **PDB header:**transferase **Chain:** A: **PDB Molecule:**n-terminal acetyltransferase a complex subunit nat1;  **PDBTitle:** the nata acetyltransferase complex bound to ppgpp |
| 28 | c5ganJ\_ | --- \| --- \| --- | not modelled | 99.6 | 10 | **PDB header:**transcription **Chain:** J: **PDB Molecule:**pre-mrna-splicing factor 6;  **PDBTitle:** the overall structure of the yeast spliceosomal u4/u6.u5 tri-snrnp at2 3.7 angstrom |
| 29 | c4ui9Y\_ | --- \| --- \| --- | not modelled | 99.6 | 9 | **PDB header:**cell cycle **Chain:** Y: **PDB Molecule:**anaphase-promoting complex subunit 7;  **PDBTitle:** atomic structure of the human anaphase-promoting complex |
| 30 | c6erqA\_ | --- \| --- \| --- | not modelled | 99.6 | 10 | **PDB header:**transcription **Chain:** A: **PDB Molecule:**dna-directed rna polymerase, mitochondrial;  **PDBTitle:** structure of the human mitochondrial transcription initiation complex2 at the hsp promoter |
| 31 | c4e85B\_ | --- \| --- \| --- | not modelled | 99.5 | 9 | **PDB header:**structural protein **Chain:** B: **PDB Molecule:**mrna 3'-end-processing protein rna14;  **PDBTitle:** crystal structure of hat domain of rna14 |
| 32 | d1w3ba\_ | --- \| --- \| --- | not modelled | 99.5 | 11 | **Fold:**alpha-alpha superhelix **Superfamily:**TPR-like **Family:**Tetratricopeptide repeat (TPR) |
| 33 | c2uy1A\_ | --- \| --- \| --- | not modelled | 99.5 | 10 | **PDB header:**rna-binding protein **Chain:** A: **PDB Molecule:**cleavage stimulation factor 77;  **PDBTitle:** crystal structure of cstf-77 |
| 34 | c5dseA\_ | --- \| --- \| --- | not modelled | 99.5 | 12 | **PDB header:**protein binding **Chain:** A: **PDB Molecule:**tetratricopeptide repeat protein 7b;  **PDBTitle:** crystal structure of the ttc7b/hyccin complex |
| 35 | c2uy1B\_ | --- \| --- \| --- | not modelled | 99.5 | 10 | **PDB header:**rna-binding protein **Chain:** B: **PDB Molecule:**cleavage stimulation factor 77;  **PDBTitle:** crystal structure of cstf-77 |
| 36 | c4zlhB\_ | --- \| --- \| --- | not modelled | 99.5 | 14 | **PDB header:**metal binding protein **Chain:** B: **PDB Molecule:**lipopolysaccharide assembly protein b;  **PDBTitle:** structure of the lapb cytoplasmic domain at 2 angstroms |
| 37 | c6g70A\_ | --- \| --- \| --- | not modelled | 99.4 | 13 | **PDB header:**splicing **Chain:** A: **PDB Molecule:**pre-mrna-processing factor 39;  **PDBTitle:** structure of murine prpf39 |
| 38 | c5dseC\_ | --- \| --- \| --- | not modelled | 99.4 | 12 | **PDB header:**protein binding **Chain:** C: **PDB Molecule:**tetratricopeptide repeat protein 7b;  **PDBTitle:** crystal structure of the ttc7b/hyccin complex |
| 39 | c5lj3S\_ | --- \| --- \| --- | not modelled | 99.4 | 9 | **PDB header:**splicing **Chain:** S: **PDB Molecule:**clf1;  **PDBTitle:** structure of the core of the yeast spliceosome immediately after2 branching |
| 40 | c4ebaC\_ | --- \| --- \| --- | not modelled | 99.4 | 11 | **PDB header:**structural protein/rna binding protein **Chain:** C: **PDB Molecule:**mrna 3'-end-processing protein rna14;  **PDBTitle:** crystal structure of the rna14-rna15 complex |
| 41 | c5wsgd\_ | --- \| --- \| --- | not modelled | 99.4 | 10 | **PDB header:**rna binding protein/rna **Chain:** D: **PDB Molecule:**u5 snrna;  **PDBTitle:** cryo-em structure of the catalytic step ii spliceosome (c\* complex) at2 4.0 angstrom resolution |
| 42 | c4kvmA\_ | --- \| --- \| --- | not modelled | 99.4 | 10 | **PDB header:**transferase/transferase inhibitor **Chain:** A: **PDB Molecule:**n-terminal acetyltransferase a complex subunit nat1;  **PDBTitle:** the nata (naa10p/naa15p) amino-terminal acetyltransferase complex2 bound to a bisubstrate analog |
| 43 | c5nnrD\_ | --- \| --- \| --- | not modelled | 99.4 | 10 | **PDB header:**transferase **Chain:** D: **PDB Molecule:**n-terminal acetyltransferase-like protein;  **PDBTitle:** structure of naa15/naa10 bound to hypk-thb |
| 44 | c3jb9R\_ | --- \| --- \| --- | not modelled | 99.3 | 12 | **PDB header:**rna binding protein/rna **Chain:** R: **PDB Molecule:**pre-mrna-splicing factor cwf4;  **PDBTitle:** cryo-em structure of the yeast spliceosome at 3.6 angstrom resolution |
| 45 | c6c95A\_ | --- \| --- \| --- | not modelled | 99.3 | 10 | **PDB header:**transferase **Chain:** A: **PDB Molecule:**n-alpha-acetyltransferase 15, nata auxiliary subunit;  **PDBTitle:** the human nata (naa10/naa15) amino-terminal acetyltransferase complex2 bound to hypk |
| 46 | c5c9sB\_ | --- \| --- \| --- | not modelled | 99.3 | 8 | **PDB header:**gene regulation **Chain:** B: **PDB Molecule:**rrna biogenesis protein rrp5;  **PDBTitle:** crystal structure of the c-terminal domain of rrp5 |
| 47 | c5gmkd\_ | --- \| --- \| --- | not modelled | 99.3 | 8 | **PDB header:**rna binding protein/rna **Chain:** D: **PDB Molecule:**u5 snrna;  **PDBTitle:** cryo-em structure of the catalytic step i spliceosome (c complex) at2 3.4 angstrom resolution |
| 48 | c4bujF\_ | --- \| --- \| --- | not modelled | 99.3 | 8 | **PDB header:**hydrolase **Chain:** F: **PDB Molecule:**superkiller protein 3;  **PDBTitle:** crystal structure of the s. cerevisiae ski2-3-8 complex |
| 49 | c5aioA\_ | --- \| --- \| --- | not modelled | 99.2 | 7 | **PDB header:**transcription **Chain:** A: **PDB Molecule:**transcription factor tau 131 kda subunit;  **PDBTitle:** crystal structure of t131 n-terminal tpr array |
| 50 | c3mkrA\_ | --- \| --- \| --- | not modelled | 99.2 | 12 | **PDB header:**transport protein **Chain:** A: **PDB Molecule:**coatomer subunit epsilon;  **PDBTitle:** crystal structure of yeast alpha/epsilon-cop subcomplex of the copi2 vesicular coat |
| 51 | c4g1tB\_ | --- \| --- \| --- | not modelled | 99.1 | 11 | **PDB header:**antiviral protein **Chain:** B: **PDB Molecule:**interferon-induced protein with tetratricopeptide repeats  **PDBTitle:** crystal structure of interferon-stimulated gene 54 |
| 52 | c4rg6B\_ | --- \| --- \| --- | not modelled | 99.1 | 8 | **PDB header:**protein binding **Chain:** B: **PDB Molecule:**cell division cycle protein 27 homolog;  **PDBTitle:** crystal structure of apc3-apc16 complex |
| 53 | c1fchB\_ | --- \| --- \| --- | not modelled | 99.1 | 12 | **PDB header:**signaling protein **Chain:** B: **PDB Molecule:**peroxisomal targeting signal 1 receptor;  **PDBTitle:** crystal structure of the pts1 complexed to the tpr region2 of human pex5 |
| 54 | c6af0A\_ | --- \| --- \| --- | not modelled | 99.0 | 9 | **PDB header:**transcription **Chain:** A: **PDB Molecule:**ctr9 protein;  **PDBTitle:** structure of ctr9, paf1 and cdc73 ternary complex from myceliophthora2 thermophila |
| 55 | c3zpjA\_ | --- \| --- \| --- | not modelled | 99.0 | 14 | **PDB header:**unknown function **Chain:** A: **PDB Molecule:**ton\_1535;  **PDBTitle:** crystal structure of ton1535 from thermococcus onnurineus na1 |
| 56 | c3mv3B\_ | --- \| --- \| --- | not modelled | 99.0 | 12 | **PDB header:**protein transport **Chain:** B: **PDB Molecule:**coatomer subunit epsilon;  **PDBTitle:** crystal structure of a-cop in complex with e-cop |
| 57 | c4r7sA\_ | --- \| --- \| --- | not modelled | 99.0 | 10 | **PDB header:**structural genomics, unknown function **Chain:** A: **PDB Molecule:**tetratricopeptide repeat protein;  **PDBTitle:** crystal structure of a tetratricopeptide repeat protein (parmer\_03812)2 from parabacteroides merdae atcc 43184 at 2.39 a resolution |
| 58 | c3hymB\_ | --- \| --- \| --- | not modelled | 99.0 | 11 | **PDB header:**cell cycle, ligase **Chain:** B: **PDB Molecule:**cell division cycle protein 16 homolog;  **PDBTitle:** insights into anaphase promoting complex tpr subdomain2 assembly from a cdc26-apc6 structure |
| 59 | c1xi4D\_ | --- \| --- \| --- | not modelled | 98.9 | 12 | **PDB header:**endocytosis/exocytosis **Chain:** D: **PDB Molecule:**clathrin heavy chain;  **PDBTitle:** clathrin d6 coat |
| 60 | c5ctqD\_ | --- \| --- \| --- | not modelled | 98.9 | 9 | **PDB header:**immune system, nuclear protein, rna bind **Chain:** D: **PDB Molecule:**squamous cell carcinoma antigen recognized by t-cells 3;  **PDBTitle:** crystal structure of human sart3/tip110 half-a tpr (hat) domain |
| 61 | c2ho1B\_ | --- \| --- \| --- | not modelled | 98.9 | 13 | **PDB header:**protein binding **Chain:** B: **PDB Molecule:**type 4 fimbrial biogenesis protein pilf;  **PDBTitle:** functional characterization of pseudomonas aeruginosa pilf |
| 62 | c4jspA\_ | --- \| --- \| --- | not modelled | 98.9 | 10 | **PDB header:**transferase **Chain:** A: **PDB Molecule:**serine/threonine-protein kinase mtor;  **PDBTitle:** structure of mtordeltan-mlst8-atpgammas-mg complex |
| 63 | c5udjA\_ | --- \| --- \| --- | not modelled | 98.9 | 12 | **PDB header:**rna binding protein **Chain:** A: **PDB Molecule:**interferon-induced protein with tetratricopeptide repeats  **PDBTitle:** ifit1 monomeric mutant (l457e/l464e) with gppp-aaaa |
| 64 | d1hz4a\_ | --- \| --- \| --- | not modelled | 98.8 | 10 | **Fold:**alpha-alpha superhelix **Superfamily:**TPR-like **Family:**Transcription factor MalT domain III |
| 65 | d1fcha\_ | --- \| --- \| --- | not modelled | 98.8 | 11 | **Fold:**alpha-alpha superhelix **Superfamily:**TPR-like **Family:**Tetratricopeptide repeat (TPR) |
| 66 | c3pe3D\_ | --- \| --- \| --- | not modelled | 98.8 | 13 | **PDB header:**transferase **Chain:** D: **PDB Molecule:**udp-n-acetylglucosamine--peptide n-  **PDBTitle:** structure of human o-glcnac transferase and its complex with a peptide2 substrate |
| 67 | c5jqyA\_ | --- \| --- \| --- | not modelled | 98.8 | 10 | **PDB header:**oxidoreductase **Chain:** A: **PDB Molecule:**aspartyl/asparaginyl beta-hydroxylase;  **PDBTitle:** aspartyl/asparaginyl beta-hydroxylase (asph)oxygenase and tpr domains2 in complex with manganese, n-oxalylglycine and factor x substrate3 peptide fragment(39mer-4ser) |
| 68 | c2gw1A\_ | --- \| --- \| --- | not modelled | 98.8 | 9 | **PDB header:**protein transport **Chain:** A: **PDB Molecule:**mitochondrial precursor proteins import receptor;  **PDBTitle:** crystal structure of the yeast tom70 |
| 69 | c4d18J\_ | --- \| --- \| --- | not modelled | 98.8 | 10 | **PDB header:**signaling protein **Chain:** J: **PDB Molecule:**cop9 signalosome complex subunit 2;  **PDBTitle:** crystal structure of the cop9 signalosome |
| 70 | c4hotA\_ | --- \| --- \| --- | not modelled | 98.8 | 9 | **PDB header:**rna binding protein/rna **Chain:** A: **PDB Molecule:**interferon-induced protein with tetratricopeptide repeats  **PDBTitle:** crystal structure of full-length human ifit5 with 5`-triphosphate2 oligoadenine |
| 71 | d2onda1 | --- \| --- \| --- | not modelled | 98.7 | 9 | **Fold:**alpha-alpha superhelix **Superfamily:**TPR-like **Family:**HAT/Suf repeat |
| 72 | c2vq2A\_ | --- \| --- \| --- | not modelled | 98.7 | 18 | **PDB header:**structural protein **Chain:** A: **PDB Molecule:**putative fimbrial biogenesis and twitching motility  **PDBTitle:** crystal structure of pilw, widely conserved type iv pilus biogenesis2 factor |
| 73 | c4ynvA\_ | --- \| --- \| --- | not modelled | 98.7 | 9 | **PDB header:**chaperone **Chain:** A: **PDB Molecule:**acl4;  **PDBTitle:** assembly chaperone of rpl4 (acl4) (residues 28-338) |
| 74 | c3iegB\_ | --- \| --- \| --- | not modelled | 98.7 | 10 | **PDB header:**chaperone **Chain:** B: **PDB Molecule:**dnaj homolog subfamily c member 3;  **PDBTitle:** crystal structure of p58(ipk) tpr domain at 2.5 a |
| 75 | c3fp4A\_ | --- \| --- \| --- | not modelled | 98.6 | 8 | **PDB header:**transport protein **Chain:** A: **PDB Molecule:**tpr repeat-containing protein yhr117w;  **PDBTitle:** crystal structure of tom71 complexed with ssa1 c-terminal2 fragment |
| 76 | d1dcea1 | --- \| --- \| --- | not modelled | 98.6 | 12 | **Fold:**alpha-alpha superhelix **Superfamily:**Protein prenylyltransferase **Family:**Protein prenylyltransferase |
| 77 | c3q75A\_ | --- \| --- \| --- | not modelled | 98.6 | 11 | **PDB header:**transferase **Chain:** A: **PDB Molecule:**farnesyltransferase alpha subunit;  **PDBTitle:** cryptococcus neoformans protein farnesyltransferase in complex with2 fpt-ii and tkcvvm peptide |
| 78 | c3cvpA\_ | --- \| --- \| --- | not modelled | 98.6 | 11 | **PDB header:**transport protein **Chain:** A: **PDB Molecule:**peroxisome targeting signal 1 receptor pex5;  **PDBTitle:** structure of peroxisomal targeting signal 1 (pts1) binding domain of2 trypanosoma brucei peroxin 5 (tbpex5)complexed to pts1 peptide (10-3 skl) |
| 79 | c4houB\_ | --- \| --- \| --- | not modelled | 98.5 | 10 | **PDB header:**rna binding protein **Chain:** B: **PDB Molecule:**interferon-induced protein with tetratricopeptide repeats  **PDBTitle:** crystal structure of n-terminal human ifit1 |
| 80 | c4xi0E\_ | --- \| --- \| --- | not modelled | 98.5 | 11 | **PDB header:**protein binding **Chain:** E: **PDB Molecule:**magnetosome protein mama;  **PDBTitle:** mama 41-end from desulfovibrio magneticus rs-1 |
| 81 | c2q7fA\_ | --- \| --- \| --- | not modelled | 98.5 | 12 | **PDB header:**protein binding **Chain:** A: **PDB Molecule:**yrrb protein;  **PDBTitle:** crystal structure of yrrb: a tpr protein with an unusual peptide-2 binding site |
| 82 | d1qsaa1 | --- \| --- \| --- | not modelled | 98.5 | 10 | **Fold:**alpha-alpha superhelix **Superfamily:**Bacterial muramidases **Family:**Bacterial muramidases |
| 83 | d2h6fa1 | --- \| --- \| --- | not modelled | 98.5 | 8 | **Fold:**alpha-alpha superhelix **Superfamily:**Protein prenylyltransferase **Family:**Protein prenylyltransferase |
| 84 | c5y88I\_ | --- \| --- \| --- | not modelled | 98.4 | 8 | **PDB header:**splicing **Chain:** I: **PDB Molecule:**pre-mrna-splicing factor clf1;  **PDBTitle:** cryo-em structure of the intron-lariat spliceosome ready for2 disassembly from s.cerevisiae at 3.5 angstrom |
| 85 | c4eqfA\_ | --- \| --- \| --- | not modelled | 98.4 | 9 | **PDB header:**protein binding/transport protein **Chain:** A: **PDB Molecule:**pex5-related protein;  **PDBTitle:** trip8b-1a#206-567 interacting with the carboxy-terminal seven residues2 of hcn2 |
| 86 | c5zypA\_ | --- \| --- \| --- | not modelled | 98.4 | 13 | **PDB header:**transcription **Chain:** A: **PDB Molecule:**rna polymerase-associated protein ctr9,rna polymerase ii-  **PDBTitle:** structure of the yeast ctr9/paf1 complex |
| 87 | c3draA\_ | --- \| --- \| --- | not modelled | 98.4 | 7 | **PDB header:**transferase **Chain:** A: **PDB Molecule:**protein  **PDBTitle:** candida albicans protein geranylgeranyltransferase-i2 complexed with ggpp |
| 88 | c5gjqQ\_ | --- \| --- \| --- | not modelled | 98.4 | 9 | **PDB header:**hydrolase **Chain:** Q: **PDB Molecule:**26s proteasome non-atpase regulatory subunit 11;  **PDBTitle:** structure of the human 26s proteasome bound to usp14-ubal |
| 89 | c4n5cH\_ | --- \| --- \| --- | not modelled | 98.4 | 12 | **PDB header:**protein binding **Chain:** H: **PDB Molecule:**cargo-transport protein ypp1;  **PDBTitle:** crystal structure of ypp1 |
| 90 | c3as5A\_ | --- \| --- \| --- | not modelled | 98.4 | 11 | **PDB header:**protein binding **Chain:** A: **PDB Molecule:**mama;  **PDBTitle:** mama amb-1 p212121 |
| 91 | c3uq3A\_ | --- \| --- \| --- | not modelled | 98.3 | 8 | **PDB header:**chaperone **Chain:** A: **PDB Molecule:**heat shock protein sti1;  **PDBTitle:** tpr2ab-domain:phsp90-complex of yeast sti1 |
| 92 | c2r5sB\_ | --- \| --- \| --- | not modelled | 98.3 | 12 | **PDB header:**structural genomics, unknown function **Chain:** B: **PDB Molecule:**uncharacterized protein vp0806;  **PDBTitle:** the crystal structure of a domain of protein vp0806 (unknown function)2 from vibrio parahaemolyticus rimd 2210633 |
| 93 | c3vtxB\_ | --- \| --- \| --- | not modelled | 98.3 | 13 | **PDB header:**protein binding **Chain:** B: **PDB Molecule:**mama;  **PDBTitle:** crystal structure of mama protein |
| 94 | c2y4tA\_ | --- \| --- \| --- | not modelled | 98.3 | 10 | **PDB header:**chaperone **Chain:** A: **PDB Molecule:**dnaj homolog subfamily c member 3;  **PDBTitle:** crystal structure of the human co-chaperone p58(ipk) |
| 95 | c5tqbB\_ | --- \| --- \| --- | not modelled | 98.3 | 10 | **PDB header:**ribosomal protein **Chain:** B: **PDB Molecule:**assembly chaperone of ribosomal protein l4 (acl4);  **PDBTitle:** crystal structure of assembly chaperone of ribosomal protein l4 (acl4)2 in complex with ribosomal protein l4 (rpl4) |
| 96 | c2hyzA\_ | --- \| --- \| --- | not modelled | 98.3 | 13 | **PDB header:**de novo protein **Chain:** A: **PDB Molecule:**synthetic consensus tpr protein;  **PDBTitle:** crystal structure of an 8 repeat consensus tpr superhelix (orthorombic2 crystal form) |
| 97 | c5xi8A\_ | --- \| --- \| --- | not modelled | 98.3 | 13 | **PDB header:**hydrolase **Chain:** A: **PDB Molecule:**beta-barrel assembly-enhancing protease;  **PDBTitle:** structure and function of the tpr domain |
| 98 | d1d8da\_ | --- \| --- \| --- | not modelled | 98.2 | 9 | **Fold:**alpha-alpha superhelix **Superfamily:**Protein prenylyltransferase **Family:**Protein prenylyltransferase |
| 99 | c6gmhQ\_ | --- \| --- \| --- | not modelled | 98.2 | 10 | **PDB header:**transcription **Chain:** Q: **PDB Molecule:**ctr9,rna polymerase-associated protein ctr9 homolog,rna  **PDBTitle:** structure of activated transcription complex pol ii-dsif-paf-spt6 |
| 100 | c5zyqA\_ | --- \| --- \| --- | not modelled | 98.2 | 10 | **PDB header:**transcription **Chain:** A: **PDB Molecule:**rna polymerase-associated protein ctr9 homolog,rna  **PDBTitle:** the structure of human paf1/ctr9 complex |
| 101 | c5m72A\_ | --- \| --- \| --- | not modelled | 98.1 | 9 | **PDB header:**protein transport **Chain:** A: **PDB Molecule:**signal recognition particle subunit srp72;  **PDBTitle:** structure of the human srp68-72 protein-binding domain complex |
| 102 | c6b85J\_ | --- \| --- \| --- | not modelled | 98.1 | 10 | **PDB header:**membrane protein **Chain:** J: **PDB Molecule:**tmhc4\_r;  **PDBTitle:** crystal structure of transmembrane protein tmhc4\_r |
| 103 | c4ui9O\_ | --- \| --- \| --- | not modelled | 98.1 | 12 | **PDB header:**cell cycle **Chain:** O: **PDB Molecule:**anaphase-promoting complex subunit 5;  **PDBTitle:** atomic structure of the human anaphase-promoting complex |
| 104 | c5lj3T\_ | --- \| --- \| --- | not modelled | 98.1 | 10 | **PDB header:**splicing **Chain:** T: **PDB Molecule:**syf1;  **PDBTitle:** structure of the core of the yeast spliceosome immediately after2 branching |
| 105 | c5xw7B\_ | --- \| --- \| --- | not modelled | 98.1 | 14 | **PDB header:**biosynthetic protein **Chain:** B: **PDB Molecule:**cellulose synthase subunit c;  **PDBTitle:** crystal structure of the flexible tandem repeat domain of bacterial2 cellulose synthase subunit c |
| 106 | c1tnoI\_ | --- \| --- \| --- | not modelled | 98.1 | 9 | **PDB header:**transferase **Chain:** I: **PDB Molecule:**geranylgeranyltransferase type i alpha subunit;  **PDBTitle:** rat protein geranylgeranyltransferase type-i complexed with2 a ggpp analog and a kkksktkcvim peptide derived from k-3 ras4b |
| 107 | c4gpkI\_ | --- \| --- \| --- | not modelled | 98.0 | 7 | **PDB header:**transcription, peptide binding protein **Chain:** I: **PDB Molecule:**nprr;  **PDBTitle:** crystal structure of nprr in complex with its cognate peptide nprx |
| 108 | c4cr3Q\_ | --- \| --- \| --- | not modelled | 98.0 | 12 | **PDB header:**hydrolase **Chain:** Q: **PDB Molecule:**26s proteasome regulatory subunit rpn6;  **PDBTitle:** deep classification of a large cryo-em dataset defines the2 conformational landscape of the 26s proteasome |
| 109 | c5djsA\_ | --- \| --- \| --- | not modelled | 98.0 | 15 | **PDB header:**transferase **Chain:** A: **PDB Molecule:**tetratricopeptide tpr\_2 repeat protein;  **PDBTitle:** thermobaculum terrenum o-glcnac transferase mutant - k341m |
| 110 | c3u4tA\_ | --- \| --- \| --- | not modelled | 98.0 | 7 | **PDB header:**structural genomics, unknown function **Chain:** A: **PDB Molecule:**tpr repeat-containing protein;  **PDBTitle:** crystal structure of the c-terminal part of the tpr repeat-containing2 protein q11ti6\_cyth3 from cytophaga hutchinsonii. northeast3 structural genomics consortium target chr11b. |
| 111 | d1xnfa\_ | --- \| --- \| --- | not modelled | 98.0 | 13 | **Fold:**alpha-alpha superhelix **Superfamily:**TPR-like **Family:**Tetratricopeptide repeat (TPR) |
| 112 | c2pl2A\_ | --- \| --- \| --- | not modelled | 98.0 | 19 | **PDB header:**protein binding **Chain:** A: **PDB Molecule:**hypothetical conserved protein ttc0263;  **PDBTitle:** crystal structure of ttc0263: a thermophilic tpr protein in thermus2 thermophilus hb27 |
| 113 | c4lngA\_ | --- \| --- \| --- | not modelled | 97.9 | 10 | **PDB header:**transferase **Chain:** A: **PDB Molecule:**caax farnesyltransferase alpha subunit ram2;  **PDBTitle:** aspergillus fumigatus protein farnesyltransferase complex with2 farnesyldiphosphate and tipifarnib |
| 114 | c6aitD\_ | --- \| --- \| --- | not modelled | 97.9 | 11 | **PDB header:**hydrolase **Chain:** D: **PDB Molecule:**beta-barrel assembly-enhancing protease;  **PDBTitle:** crystal structure of e. coli bepa |
| 115 | c6ah0N\_ | --- \| --- \| --- | not modelled | 97.9 | 10 | **PDB header:**splicing **Chain:** N: **PDB Molecule:**pre-mrna-processing factor 6;  **PDBTitle:** the cryo-em structure of the precusor of human pre-catalytic2 spliceosome (pre-b complex) |
| 116 | c6r7nB\_ | --- \| --- \| --- | not modelled | 97.8 | 12 | **PDB header:**ligase **Chain:** B: **PDB Molecule:**cop9 signalosome complex subunit 2;  **PDBTitle:** structural basis of cullin-2 ring e3 ligase regulation by the cop92 signalosome |
| 117 | c3urzB\_ | --- \| --- \| --- | not modelled | 97.8 | 9 | **PDB header:**protein binding **Chain:** B: **PDB Molecule:**uncharacterized protein;  **PDBTitle:** crystal structure of a putative protein binding protein (bacova\_03105)2 from bacteroides ovatus atcc 8483 at 2.19 a resolution |
| 118 | c4gyoB\_ | --- \| --- \| --- | not modelled | 97.8 | 10 | **PDB header:**hydrolase **Chain:** B: **PDB Molecule:**response regulator aspartate phosphatase j;  **PDBTitle:** crystal structure of rap protein complexed with competence and2 sporulation factor |
| 119 | c4nrhB\_ | --- \| --- \| --- | not modelled | 97.7 | 6 | **PDB header:**chaperone/protein binding **Chain:** B: **PDB Molecule:**chaperone sycd;  **PDBTitle:** copn-scc3 complex |
| 120 | c4abnA\_ | --- \| --- \| --- | not modelled | 97.7 | 11 | **PDB header:**gene regulation **Chain:** A: **PDB Molecule:**tetratricopeptide repeat protein 5;  **PDBTitle:** crystal structure of full length mouse strap (ttc5) |
