## Supplementary material for "The endosomal TbTpr86/TbUsp7/SkpZ (TUS) complex controls surface protein abundance in trypanosomes": DA1: newmodeller.modscript.original 3.py

```
from modeller import * 
from modeller.automodel import * 
log.verbose()    # request verbose output
env = environ()  # create a new MODELLER environment to build this model in
env.io.atom_files_directory = './:../atom_files'
a = automodel(env,
              alnfile  = 'newmodeller.pir',
              knowns   = ('c4m57A_','c4g25A_',),
              sequence = 'query')
a.starting_model= 1
a.ending_model  = 1
a.make()
```
