## Supplementary material for "The endosomal TbTpr86/TbUsp7/SkpZ (TUS) complex controls surface protein abundance in trypanosomes": DA1: psi-blast 3.html

Content-Type: text/html


```
Reference sequence (query): Undefined
Identities normalised by aligned length.
Colored by: property
```

```
HSP processing: ranked
Search cycle: 3
```

```
                                                                                                         1 [        .         .         .         .         :         .         .         .         .         1         .         .         .         .         :         .         .         .         .         2         .         .         .         .         :         .         .         .         .         3         .         .         .         .         :         .         .         .         .         4         .         .         .         .         :         .         .         .         .         5         .         .         .         .         :         .         .         .         .         6  ] 603
     Undefined                                              bits   E-value  N 100.0%   1:603               MGAEKKINRPTNSGEVAENVGAFTEVMVPVWSRLCDKVIAAYKAGEDIVVLQHFEELRATPELDVIAHDSLISAAPDLTVGAVEKKEDDRAAKKERLEYCWGRALLSARHLAGDDDEHINSIVESFEQQFGCVSGYLRALAINAESALEVLQLYQMMEGPVTTRVATAALHRFLTNDASEVSGLLQFLKLNNVPLTREFHDQLEKNFVMRMKTEDAEEVFKDMRNAGVEPVSTHPYNFIFMKTPPFKYMDLLDYYEDMLDRGIVPMNSTFRILSRQTQNGGLVNQHFSALARGTTTGSDRRHETEGHGGTQAFFATRLEELARLKSSTTLDDALRVLRRMDVEGTPLISGSVVLNVLLSVFCRAGHIEDTERVLQLAKERGCTPNGATLSRVLKKSNADRFLQAELILPDALEAGDVRLSDITDVMCHFFNNGMRSQVVRILRAYVVRCYRNPTESIPHTVALWALRLGPFDWALGFLAILLASNDEPLSDRIQRRMHEVVKENRSREILKVLEGHKWDRRSILLAPQLQEPTNSSDGAGEVNGRSSEGGNSGGGHHRRDMGPSWMSAMHLVLKEMGVEKEVLMGYKAKLSALINDAKKKRDN    
   1 UniRef50_A9TFE2        Predicted protein n=1 Tax=P...   543    1e-153  1  14.8%  21:600     2:586     --------------------KGFPSPNVVTYNSLLNALAKAGQCEEAQLLFEELKAAKWT--PDVVSYSCLINSLGRAGKWEAALEVVAEMQAKGCKPNLWTYNTLVDCLGKAGQFDEALRLLAEMRDNGCVPDVRTYNcaGRLSEAFTLFAEMRERGCVPDTFTYNSLIYGLGkgRSQKAMELLEEMERHGCPPDVMTYSSLITGLGKDGETVKAFKLFQEMKRRGRKPDSITFTALMDALGKAGRVDDALELLDEMKERGVKPGVVTYNALIAGFGKVGDLVEAYNLLDEMKR------------NGCKPDVVTYSCLITGLIKASQLDEACQVLKKMEKEGCP--PDTITYNTLINGLGKAGLLNDAGRLFDRMKSKGCNPDVVTYSTLITALGKAARVESACVLFEEMESVGIQPdtYCSIITVLGKAGQVDDADRLFSEMRGKGL-SPDVITYNAFLNSLGRGGRFKEARKIFEDMKESGLLPDVATYDALLLGLSKTKEVDDACGLLKELIEQGCAF-DSLKFDECLEILTSWGNVDEAHELlyNALIDALAKAGRVSEAFNTLEDLKEQGGKPDIV-SYSSLISALGQTGQID---    
   2 UniRef50_B8LQA8        Putative uncharacterized pr...   540    1e-152  1   8.4%   2:582   160:708     -ALASMYTKCGSLENARQVFDRMPKRDVVSWNAIIAGYSQNGQPYEALALFSEMQV--NGIKPNSSTLVSVMPVCAHLLALEQGKQIHCYAIRSGIESDVLVVNGLVNMYAKCGNVNTAHKLFERMprDVASWNAIIGGYSLNshEALAFFNRMQVRGIKPNSITMVSVLPACAHlaLEQGQQIHGYAIRSGFESNDVVGNALVNMYAKCGNVNSAYKLFERMPKK----NVVAWNAIISGYSQHGHPHEALALFIEMQAQGIKPDSFAIVSVLPACAHFLALEQGKQI------------HGYTIRSGFESNVVVGTGLVDIYAKCGNVNTAQKLFERMPEQ------DVVSWTTMILAYGIHGHGEDALALFSKMQETGTKLDHIAFTAILTACSHAGLVDQGLQYFQCMkdYGLApeHYACLVDLLGRAGHLDEANGIIKNMS----LEPDANVWGALLGACRIHCNIELGEQAAKHLFELDPD-NAGYYVLLSNIYAEAQRWEDVAKLRKMMKEKGVKKQP----------GCSVVAVHRDVQTFLVGDRTHPQSEQIYAMLEILYEQMRKAGYVPNT---------------------    
   3 UniRef50_A9T8E9        Predicted protein n=3 Tax=E...   519    1e-146  1  10.8%   2:582   267:815     -ALVKMYAKCGSYKDCRQVFEKLVNRDLIAWNTMIGGLAEGGYWEEASEVYNQMQREGVM--PNKITYVILLNACVNSAALHWGKEIHSRVAKAGFTSDIGVQNALISMYSRCGSIKDARLVFDKMVrdVISWTAMIGGLAKSgaEALTVYQEMQQAGVEPNRVTYTSILNACSsaALEWGRRIHQQVVEAGLATDAHVGNTLVNMYSMCGSVKDARQVFDRMIQR----DIVAYNAMIGGYAAHNLGKEALKLFDRLQEEGLKPDKVTYINMLNACANSGSLEWAREI------------HTLVRKGGFFSDTSVGNALVSTYAKCGSFSDASIVFEKMTKR------NVISWNAIIGGSAQHGRGQDALQLFERMKMEGVKPDIVTFVSLLSACSHAGLLEEGRRYFCSMSQDfpTIEHYGCMVDLLGRAGQLDEAEALIKTMP----FQANTRIWGALLGACRIHGNVPVAERAAESSLKLDLD-NAVVYVALSHMYAAAGMWDSAAKLRKLMEQRGVTKEPGR----------SWIQVGDKLHYFVAEDRSHPQSEKIYAELDRLTHAMKMKGYVPDT---------------------    
   4 UniRef50_A9TNX6        Predicted protein n=9 Tax=E...   512    1e-143  1  10.4%   4:582   354:899     ---LSMYTKCGSMEDALEVFDLVKGRNVVSWTAMIAGFAQHGRIDEAFLFFNKM--IESGIEPNRVTFMSILGACSSPSALKRGQQIQDHIIEAGYGSDDRVRTALLSMYAKCGSLKDAHRVFEKikQNVVAWNAMITAYVQhyDNALATFQALLKEGIKPNSSTFTSILNVCKSSdlELGKWVHFLIMKAGLESDLHVSNALVSMFVNCGDLMSAKNLFNDMPKR----DLVSWNTIIAGFVQHGKNQVAFDYFKMMQESGIKPDKITFTGLLNACASPEALTEGRRL------------HALITEAAFDCDVLVGTGLISMYTKCGSIEDAHQVFHKLPKK------NVYSWTSMITGYAQHGRGKEALELFYQMQQEGVKPDWITFVGALSACAHAGLIEEGLHHFQSMKEFNIEpeHYGCMVDLFGRAGLLNEAVEFIIKMQ----VEPDSRVWGALLGACQVHLNVELAEKAAQK--KLELDPNdgVFVILSNIYAAAGMWKEVAKMRKVMLDRGVVKKPGQ----------SWIEVDGKVHTFYSDDKTHPQTEEIHAELERLHMEMRQLGYVPDT---------------------    
   5 UniRef50_A9RJW2        Predicted protein n=1 Tax=P...   503    1e-140  1  10.7%   4:582    89:636     ---LSMYAKCGSLTDARRVFDSIRDRNIVSWTAMIEAFVAGNKNLEAFKCYETMKLAGC--KPDKVTFVSLLNAFTNPELLQLGQKVHMEIVEAGLELEPRVGTSLVGMYAKCGDISKARVIFDRleKNVVTWTLLIAGYAQQgdVALELLETMQQAEVAPNKITFASILQGCTtaALEHGKKVHRYIIQSGYGRELWVVNSLITMYCKCGGLEEARKLFSDLPHR----DVVTWTAMVTGYAQLGFHDEAINLFRRMQQQGIKPDKMTFTSVLTSCSSPAFLQEGKRI------------HQQLVHAGYNLDVYLQSALVSMYAKCGSMDDASLVFNQMSER------NVVAWTAIITGCcqHGRCREALEYFDQMKKQGIKPDKVTFTSVLSACTHVGLVEEGRKHFRSmlDYGIKpeHYSCFVDLLGRAGHLEEAENVILSMP----FIPGPSVWGALLSACRVHSDVERGERAAENVLKLDPD-DDGAYVALSSIYAAAGRYEDAEKVRQVMEKRDVVKEPGQ----------SWIEVDGKVHVFHVEDKSHPESEQIYVELGKLTEQIKEMGYVPDT---------------------    
   6 UniRef50_A9SDQ1        Predicted protein n=2 Tax=P...   499    1e-139  1  11.6%   6:581   201:780     -----MYSKCGEISVACEVFHKMTERNVVSWTAIIQANAQHRKLNEAFELYEQM--LQAGISPNAVTFVSLLNSCNTPEALNRGRRIHSHISERGLETDMIVANALITMYCKCNSVQEAREIFDRmkRDVISWSAMIAGYAQSgdEVFQLLERMRREGVFPNKVTFMSILRAcaHGALEQGRQIHAELSKVGFELDRSLQTAIFNMYAKCGSIYEAEQVFSKMANK----NVVAWTSFLSMYIKCGDLSSAEKVFSEMPTR----NVVSWNLMIAGYAQNGDIVKVFELLSSMKAEggKLVHAEAVKLGLESDTVVATSLIGMYSKCGQVAEARTVFDKMSNR------DTVAWNAMLAGYGQHGDGLEAVDLFKRMLKERVSPNEITLTAVISACSRAGLVQEGREIFRMMQEdkMTpqHYGCMVDLLGRAGRLQEAEEFIQSMP----CEPDISVWHALLGACKSHNNVQLAERAAHHILE--LEPsaSVYITLSNIYAQAGRWDDSTKVRRVMDDRGLKKDRGE----------SSIEIDGRIHTFVAEDCAHPEIDAIHAELETLTKEMKEAGYTPD----------------------    
   7 UniRef50_B9IFK4        Predicted protein n=2 Tax=M...   493    1e-138  1  12.2%  14:601   257:883     -------------RLLFEMEEKGCSPSLVTYNVVIGGLCRAGEVDEAFELKKLMDKKGLV--ADVFTYSILIDGFGKQKRCTEAKLMLEEMFSKGLKPGHVAYTALIDGFMRQGDSGEAFRVKEEMLARGVKLNLFTYNAlgDMEKADALLNEMIMVGIKPDTQTYNNMIEGYLKenTSRVKDLLSEMKKSNLVPTAYTCGMIINGLCRHGSIEDASRVFEIMVSLGVKPNAVIYTTLIKGHVQEGRFQEAVRILKVMDKKGVQPDVLCYNSVIIGLCKSRKMEEAKDYLVEMIERGLkrYFKEMLGCGIAPNDVVCTALIDGYCKEGSTTEATSIFRCML--GRSVHPDVRTYSALIHGLLRNGKLQGAMELLSEFLEKGLVPDVFTYNSIISGFCKQGGIGKAFQLHEYMCQKGISPntYNALINGLCKAGEIERARELFDGIPGKGLAH-NAVTYATIIDGYCKSGNLSKAFRLFDEMTLKGVPPDSFVYSALIDGCRKEGNTEKALSLFLESVQKGFASTSS--LNALMDGFCKSGKVIEAnhVTYTILIDYHCKTGFLKEAEQFFVDMQKRNLMPNAL-TYTALLSGYNMAGRRSE--    
   8 UniRef50_A9S042        Predicted protein n=1 Tax=P...   493    1e-138  1  10.9%   7:602  297:1002     ------FNRSGRLDPAAemIKSGIDPGVHTFTVLIDALVKSGNIDEACKFFNGMKNLRCS--PNVVTYTTLVNGLAKAGRLEEACEVFVEMKENNCSPDAIAYNTLIDGLGKAGEADMACGLFKEMKDRGLVPNLRTYNIMISVleAWQLFHDLKEQGAVPDVFTYNTLIDVLGKGgmDKVLAIIKEMVEKgiSRdeITYNTLMSAFIHNGHVDEAVKLLEVMKKHECIPTVVTYTTLVDGLGKAGRLDEAVSLLREMEKQGCEPSVVTYSSliNCLCKSDDVDQALDVFGRMKEEgaLQIFNELQESSLVPDTFVYNIMVNGLVKSNRVDEACKLVDSM--KNQNILPDLFTYTSLLDGLGKSGRLEEAFNMFTKMTEEGHEPDVVAYTSLMDVLGKGGKLSHALIIFRAMAKKRCVPdtYSSLIDSLGKEGRVEEAYYFFENSISKGC-TPNVGVYSSLIDSFGKKGMVDRALELFEEMQRRQCPPNIVTYNNLLSGLAKAGRLNVAEKLLEEMEKVGCVPDLVT-YNILIDGVGKMGMVDEAESYFKRMKEKGIVPddKLLEACELFDSMEEEGYNPSVV-TYNVLIDILGRAGKVHEA-    
   9 UniRef50_A9U4W7        Predicted protein n=4 Tax=P...   493    1e-137  1  10.7%   6:582   278:821     -----MYAKCGSIDDARLVFDGMCDRDVISWNAMIGGLAQNGCGHEAFTIFLKMQQEGFV--PDSTTYLSLLNTHVSTGAWEWVKEVHKHAVEVGLVSDLRVGSAFVHMYIRCGSIDDAQLIFDKlvRNVTTWNAMIGGVAQqgREALSLFLQMRREGFFPDATTFVNILSavGEEALEWVKEVHSYAIDAGLV-DLRVGNALVHMYAKCGNTMYAKQVFDDMVER----NVTTWTVMISGLAQHGCGHEAFSLFLQMLREGIVPDATTYVSILSACASTGALEWVKEV------------HSHAVNAGLVSDLRVGNALVHMYAKCGSVDDARRVFDDMLER------DVYSWTVMIGGLAQHGRGLDALDLFVKMKLEGFKPNGYSFVAVLSACSHAGLVDEGrqFLSLTQDYGIEpeHYTCMVDLLGRAGQLEEAKHFILNMPIEPGDAP----WGALLGACVTYGNLEMAEFAAKE--RLKLKpsASTYVLLSNIYAATGNWEQKLLVRSMMQRRGIRKEPGR----------SWIEVDNQIHSFVVGDTSHPESKEIYAKLKDLIKRLKAEGYVPDT---------------------    
  10 UniRef50_Q5W963        PpPPR_77 protein n=14 Tax=E...   492    1e-137  1  10.7%   3:589  469:1026     --ANALMNMYRRCGSLMevFEGTQARDVISWNSMIAGHAQHGSYETAYKLFQEMQ--NEELEPDNITFASVLSGCKNPEALELGKQIHGRITESGLQLDVNLGNALINMYIRCGSLQDARNVFHSlhRDVMSWTAMIGGCADQGekAIELFWQMQNEGFRPVKSTFSSILKVcsSACLDEGKKVIAYILNSGYELDTGVGNALISAYSKSGSMTDAREVFDKMPSR----DIVSWNKIIAGYAQNGLGQTAVEFAYQMQEQDVVPNKFSFVSLLNACSSFSALEEGKRVHAEIVKRK------------LQGDVRVGAALISMYAKCGSQGEAQEVFDNIIEK------NVVTWNAMINAYAQHGLASKALGFFNCMEKEGIKPDGSTFTSILSACNHAGLVLEGYQIFSSMeeYGVLpeHYGCLVGLLGRARRFQEAETLINQMP----FPPDAAVWETLLGACRIHGNIALAEHAANNALKLNAR-NPAVYILLSNVYAAAGRWDDVAKIRRVMEGRGIRKEPGR----------SWIEVDNIIHEFIAADRSHPETAEIYAELKRLSVEMEEAGYFPDTQHVLHDL--------------    
  11 UniRef50_D1IL48        Whole genome shotgun sequen...   490    1e-137  1  11.1%  19:603     1:554     ------------------MGEKGLVPNTYTYTIITAGLCRAKRMNEAKLTFEEMQKTGLKPDYN--ACSALIDGFMREGDIDEVLRIKDVMVSCGIPINLITYNVLIHGLCKFGKMEKAAEILKGMITLGCKPNSRTFCLLinMGRALELLDEMEKRNLVPSAVSYGAMINGLCHcdLSLANKLLEKMTFSGLKPNVVVYSTLIMGYASEGRIEEARRLLDGMSCSGVAPDIFCYNAIISCLSKAGKMEEASTYLLEIQGRGLKPDAVTFGAFILGYSKTGKMTEAAKYFDEML------------DHGLMPNNPLYTVLINGHFKAGNLMEALSIFRHLHALG--VLPDVQTCSAFIHGLLKNGRVQEALKVFSELKEKGLVPDVFTYSSLISGFCKQGEVEKAFELHDEMCLKGIAPniYNALVDGLCKSGDIQRARKLFDGMPEKGL-EPDSVTYSTMIDGYCKSENVAEAFSLFHEMPSKGVQPHSFVYNALVHGCCKEGDMEKAMNLFREMLQKGFATTLS--FNTLIDGYCKSCKI---------------------QEASQLFQEMIAKQIMPDHV-TYTTVIDWHCKAGKMEEAN    
  12 UniRef50_D1I761        Whole genome shotgun sequen...   486    1e-136  1  11.2%  22:601   206:789     ---------------------KGVFPTVKTCTFLLSSLVKANELEKSYWVFETMRQ---GVSPDVYLFSTAINAFCKGGKVEDAIQLFFDMEKLGVSPNVVTYNNLIHGLCKHGNLDEAFRFKEKMVKDGVNATLITYSVLINgnEANSVLKETLEKGFTPNEVVYNTLIDGYCkgNLGDALRIRGDMVSKGINPNSVTLNSIIQGFCKIGQMEQAECILEEMLSRGFSINPGAFTTIIHWLCMNSRFESALRFLREMLLRNMRPNDGLLTTLVGGLCKEGKHSDAVELWFRLLEK------------GFGANLVTTNALIHGLCKTGNMQEAVRLLKKMLERGFVL--DKITYNTLISGCCKEGKVEEGFKLRGEMVKQGIEPDTFTYNLLIHGMCRIGKLDEAVNLWNECKSRDLVPNVYtvMIDGYCKADKIEEGEKLFTELLTQNL-ELNSVVYNTLIRAYCRNGNTVEAFKLHDDMRSKGIPPTTATYSSLIHGMCNIGRMEDAKCLIDEMRKEGL-LPNVVCYTALIGGYCKLGQMDKVvkITYTVMIDGYSKSGDMKTAAKLLHEMVGKGIVPDTV-TYNVLTNGFCKEGKIEE--    
  13 UniRef50_D1I4Z8        Whole genome shotgun sequen...   485    1e-135  1  12.8%   3:602   314:889     --YNALIDGFMRQGDIeeMVACGIEANLIIWNTLLNGVCKAGKMEKALEIMQEM--MEKGVEPDSQTYSLLIEGHCRGQNMARAFELLDEMKKRKLAPTVLTYSVIINGLCRCGNLQGTNAILREMVMNGLKPNAVVYTTlgRVEESRMILERMREQGILPDVFCYNSLIIGFCKarMEEARTYLMEMLERRLRPNAHTYGAFIDGYSKAGEMEIADRYFNEMLSCGVLPNVGIYTALIEGHCKEGNVTEAFSVFRFILSRRVLQDVQTYSVLIHGLSRNGKMHEAFGIFSELQEK------------GLLPNAFTYNSLISGSCKQGNVDKASQLLEEMCIKG--INPDIVTYNILIDGLCKAGEIERAKNLFDDIEGRGLTPNCVTYAAMVDGYCKSKNPTAAFQLLEEMLLRGVPPDayNVILNFCCKEEKFEKALDLFQEMLEKGFA--STVSFNTLIEGYCKSGKLQEANHLLEEMIEKQFIPNHVTYTSLIDHNCKAGMMGEAKRLWLEMQERNVMPTAKT----------------------YTSLLHGYHNIGNMSEVSALFEEMVAKGIEPDKM-TYYVMIDAYCREGNVMEA-    
  14 UniRef50_D1IXS6        Whole genome shotgun sequen...   484    1e-135  1  11.1%   4:582    85:641     ---NSLIskSGFLDEAARLFGSMPEPDQCSWNSMVSGFAQHDRFEESLEYFVKMH--REDFLLNEYSFGSALSACAGLMDLNMGTQVHALVSKSRYSTDVYMGSALIDMYSKCGSVACAEEVFSGmeRNLVTWNSLITCYEQNgsEALEVFVRMMDSGLEPDEVTLASVVSACAscALKEGLQIHARVVKTnfRDDLVLGNALVDMYAKCSKVNEARRVFDRMSIR----NVVSETSMVSGYARAASVKAARFMFSKMTQR----NVVSWNALIAGYTQNGENEEALRLFRLLKRESIWPTH-YTFGNLLKSDIFVGNSLIDMYMKCGSIEDGSRVFEKMKER------DCVSWNAIIVGYAQNGYGAEALQIFRKMLVCGEKPDHVTMIGVLCACSHAGLVEEGRHYFFSMEEHGLIpdHYTCMVDLLGRAGCLNEAKNLIEAMP----VNPDAVVWGSLLAACKVHGNIEMGKHAAEKLLE--IDpnSGPYVLLSNMYAELGRWGDVVRVRKLMRQQGVTKQP----------GCSWIEVESRVHVFLVKDKSHPHRKQIYSVLKMLTEQMKRVGYIPDA---------------------    
  15 UniRef50_A9S537        Predicted protein n=1 Tax=P...   483    1e-135  1   9.4%   5:580   198:744     ----ALVSMygSMDDARQVFDGLHIRDVSTFNVMVGGYAKSGDWEKAFELFYRMQQ--VGLKPNKISFLSILDGCWTPEALAWGKAVHAQCMNAGLVDDIRVATSLIRMYTTCGSIEGARRVFDNMkrDVVSWTVMIEGYagNIEDAFGLFATMQEEGIQPDRITYMHIMNACasANLNHAREIHSQVDIAGFGTDLLVSTALVHMYAKCGAIKDARQVFDAMPRR----DVVSWSAMIGAYVENGYGTEAFETFHLMKRSNIEPDGVTYINLLNACGHLGALDVGMEIY------------TQAIKADLVSHVPLGNALIIMNAKHGSVERARYIFDTMVRR------DVITWNAMIGGYSLHGNAREALYLFDRMLKERFRPNSVTFVGVLSACSRAGFVDEGRRFFTYLLerGIVPTvyGCMVDLLGRAGELDEAELLIKSMPVK----PTSSIWSSLLVACRIHGNLDVAERAAERCLM--IDpdGAVYVQLSHMYAAAGMWENVAKVRKVMESRGIRKEQ----------GCTWIEVAGKVHTFVVEDRSHPLVGEIYAELARLMNAIKREGYIP-----------------------    
  16 UniRef50_D1I3L2        Whole genome shotgun sequen...   482    1e-134  1  11.3%   5:580 1317:1933     ----SLVDMYGKCgdARKVFDSMVEKNVVTWNSMIVGYVQNGLNQEAIDVFYDMRVEG--IEPTRVTVASFLSASANLDALIEGKQGHAIAILNSLDLDNILGSSIINFYSKVGLIEDAELVFSRmeKDVVTWNLLISSYVQHHqkALNMCHLMRSENLRFDSVTLSSILSASatSNIKLGKEGHCYCIRRNLESDVVVANSIIDMYAKCERIDDARKVFdqMQFDSVPPNVISWNSVILGFLRNGQVNEAKDMFSQMQSLGFQPNLITWTTLISGLAQSGFGYEAILFFQKMQEAggRAIHGFITRHEFCLSVPVATSLVDMYAKCGSIDEAKKVFHMMSSKELPI------YNAMISAYALHGQAVEALALFKHLQKEGIEPDSITFTSILSACSHAGLVNEGLNLFADMvkHNMNpeHYGCVVSLLSRCGNLDEALRLILTMP----FQPDAHILGSLLTACREHHEIELGEYLSKHLFK--LEpnSGNYVALSNAYAAAGRWVEVSNMRDLMKVRGLRKNP----------GCSWIQTGGKLNVFVAGDGSHPKTEEIYAMLAMLLSEMRFMGYVP-----------------------    
  17 UniRef50_B9MZK2        Predicted protein n=5 Tax=P...   481    1e-134  1  10.9%   7:602    72:672     ------ISVDDALASFYRMARMNPRPSIVEFGKFLGSIAKMKQYSTVVYLCNQMDLFGVTHT--VYSLNILINCLCRLNHVDFAVSVWGKMFKLGIQPDVITFTTLINGVCNEGKIKVAVELYNEmePDVISYNTLINGLcgNTNMAVHVFKKMEQNGCKPNVVTYNTIIDSLCKDRlnDAMDFLSEMVGRGIPPDAITYNSIVHGLCCLGQLNEATRLFKRMEQNGCKPDVVTYNIIIDSLYKDRLVNDAADFLSEMVDQGIPPDVVTYTTILHGLCYLGQLNEAIRLFKKMEQK------------GCKPDVVAYNTIIDSLCKDRLVNDAMEFLSEMVDRG--IPPNAVTYSTILHGFCNLGQLDEATQLFKEMVGRNVMPNTLTFSILVDGLCQEGMVSEARWVFETMTEKGVEPntYNALMNGYCLRCKMNEARKVFEIMVGKGCA-PDLHSYNILINGYCNSRRMDKAKALLTQMSVKKLTPNTVTYNTIMKGLCYVGRLLDAQELFKKMCSSGMLPTLMT-YSILLNGLCKHGHLDEALKLFKSMKEKKLEPdgKLEVAKGLFSKLSADGIQP-PGRTYNVMIKGLLKEGLSDEA-    
  18 UniRef50_B9HET1        Predicted protein n=2 Tax=M...   478    1e-133  1   9.8%   4:582    63:681     ---NRLIDVYGKCGyaRKVFDRMSERNVFswNSMIAGFAQHDRFEEALDWFVRMH--RDDFVLNDYSFGSGLSACSRLKDLKLGAQIHGLISKSKYSLDVFMGSGLIDFYSKCGLVGCARRVFDGMekNVVSWNCLITCYEQNgiEALEAFGRMTELGFKPDEVTLASVVSACAtaAFKEGVQIHARVVKSdfRNDLILGNALVDMYAKCGRVNEARCVFDRMPVRNaqKDIVSWNALIAGYTQNGENEEALGLFRMLKRESVCPTHYTFGNLLNASANLADLELGRQAHSHVVK------HGFRFQSGEEPDIFVGNSLIDMYMKCGSVEEGLRVFENMVEK------DHVSWNTMIIGYAQNGYGMEALELFQKMLESGEKPDHVTMIGTLCACSHAGLVEEGRRYFFSmkEHGLLpdHYTCMVDLLGRAGCLEEAKDLIESMPK----QPDAVVWSSLLSACKVHRNITLGKYVAEKIFE--IDPTSSgyVLLANMYSELGRWGDAVSVRKLMRRRGVVKQP----------GCSWIDIQSNVHVFMVKDKRHPQKKEIYSILKLLTKHMRQAGYVPDA---------------------    
  19 UniRef50_Q9LTV8        Pentatricopeptide repeat-co...   474    1e-132  1  10.3%  10:589    66:614     ---------FGDITFARQVFDDLPRPQIFPWNAIIRGYSRNNHFQDALLMYSNMQLARVS--PDSFTFPHLLKACSGLSHLQMGRFVHAQVFRLGFDADVFVQNGLIALYAKCRRLGSARTVFEGleRTIVSWTAIVSAYAQNGeeALEIFSQMRKMDVKPDWVALVSVLNAFTCldLKQGRSIHASVVKMGLEIEPDLLISLNTMYAKCGQVATAKILFDKMKS----PNLILWNAMISGYAKNGYAREAIDMFHEMINKDVRPDTISITSAISACAQVGSLEQARSMYE------------YVGRSDYRDDVFISSALIDMFAKCGSVEGARLVFDRTLDR------DVVVWSAMIVGYGLHGRAREAISLYRAMERGGVHPNDVTFLGLLMACNHSGMVREGWWFFNRMADHKINpqHYACVIDLLGRAGHLDQAYEVIKCMP----VQPGVTVWGALLSACKKHRHVELGEYAAQQLFS--IDpnTGHYVQLSNLYAAARLWDRVAEVRVRMKEKGLNK----------DVGCSWVEVRGRLEAFRVGDKSHPRYEEIERQVEWIESRLKEGGFVANKDASLHDL--------------    
  20 UniRef50_B8LLJ0        Putative uncharacterized pr...   473    1e-132  1  10.2%   4:552   104:643     ---NKLVskLGSLVEARRVFDEMPVKNVVSWTAMIAAYARHEHGQEALGFFYEMQD--VGIQPNHFTFASILPACTDLEVLG---EFHDEIVKGGFESNVFVGNGLVDMYAKRGCIEFARELFDKmqRDVVSWNAMIAGYVQNgeDALKLFQEIPKR----DVITWNTMMAGYacGDVENAVELFEKMPEQ----NLVSWNTMIAGYVQNGSVKEAFKLFQIMPER----NVISWNAVISGFAQNGQVEEALKLFKTMPEC----NVVSWNAMIAGYSQNGQAENALKLFGQMqlEQGNEAHEVVIRSGFQSDVLVGNTLVGMYAKCGSIEDARKVFDRMRQQ------DSASLSAMIVGYAINGCSKESLELFEQMQFTGLKPDRVTFVGVLSACCHAGLVDEGRQYFDIMTryHITpeHYGCMIDLLGRAGCFDEANDLINKMPIK----PDADMWGSLLSACRTHNNIDLGEKVAQHLIALNPQ-NPAPYVLLSNIYAAAGRWDDIGSVRNRMKDRKVKK----------KLGCSWIVIKKQVHAFLVG---------------------------------------------------    
  21 UniRef50_UPI00019842B4 PREDICTED: hypothetical pro...   472    1e-131  1  11.3%   6:580   226:838     -----MYGKCGVLEDARKVFDSMVEKNVVTWNSMIVGYVQNGLNQEAIDVFYDMRVEG--IEPTRVTVASFLSASANLDALIEGKQGHAIAILNSLDLDNILGSSIINFYSKVGLIEDAELVFSRmeKDVVTWNLLISSYVQHHqkALNMCHLMRSENLRFDSVTLSSILSASatSNIKLGKEGHCYCIRRNLESDVVVANSIIDMYAKCERIDDARKVFdqMQFDSVPPNVISWNSVILGFLRNGQVNEAKDMFSQMQSLGFQPNLITWTTLISGLAQSGFGYEAILFFQKMQEAggRAIHGFITRHEFCLSVPVATSLVDMYAKCGSIDEAKKVFHMMSSKELPI------YNAMISAYALHGQAVEALALFKHLQKEGIEPDSITFTSILSACSHAGLVNEGLNLFADMvkHNMNpeHYGCVVSLLSRCGNLDEALRLILTMP----FQPDAHILGSLLTACREHHEIELGEYLSKHLFK--LEpnSGNYVALSNAYAAAGRWVEVSNMRDLMKVRGLRKNP----------GCSWIQTGGKLNVFVAGDGSHPKTEEIYAMLAMLLSEMRFMGYVP-----------------------    
  22 UniRef50_Q9LSL9        Pentatricopeptide repeat-co...   470    1e-131  1  11.8%   1:600   193:777     LARFGLVDEMKQV--YMEMLEDKVCPNIYTYNKMVNGYCKLGNVEEANQYVSKIVEAGL--DPDFFTYTSLIMGYCQRKDLDSAFKVFNEMPLKGCRRNEVAYTHLIHGLCVARRIDEAMDLFVKMKDdvRTYTVLIKSLCGSEreALNLVKEMEETGIKPNIHTYTVLIdlCSQCKFEKARELLGQMLEKGLMPNVITYNALINGYCKRGMIEDAVDVVELMESRKLSPNTRTYNELIKGYCKS-NVHKAMGVLNKMLERKVLPDVVTYNSLIDGQCRSGNFDSAYRLLSLM------------NDRGLVPDQWTYTSMIDSLCKSKRVEEACDLFDSLEQKG--VNPNVVMYTALIDGYCKAGKVDEAHLMLEKMLSKNCLPNSLTFNALIHGLCADGKLKEATLLEEKMVKIGLQpsTDTILIHRLLKDGDFDHAYSRFQQMLSSGTK-PDAHTYTTFIQTYCREGRLLDAEDMMAKMRENGVSPDLFTYSSLIKGYGDLGQTNFAFDVLKRMRDTGCEPSQHTFLSLIKHLL----EMKYGKQKGSEPELCAMSNMMEFDTVVELLEKMVEHSVTPNA-KSYEKLILGICEVGNLR---    
  23 UniRef50_Q9SY02        Pentatricopeptide repeat-co...   468    1e-130  1  13.0%   3:579    98:691     --YNGMISGygEFELARKLFDEMPERDLVSWNVMIKGYVRNRNLGKARELFEIM------PERDVCSWNTMLSGYAQNGCVDDARSVFDRMPEKN---DVSWNALLSAYVQNSKMEEaaRQFFDSmvRDVVSWNTIITGYAQSgdEARQLFDESPVQ----DVFTWTAMVSGYIQNRmeEARELFDKMPER----NEVSWNAMLAGYVQGERMEMAKELFDVMPCR----NVSTWNTMITGYAQCGKISEAKNLFDKMPKR----DPVSWAAMIAGYSQSGHSFEALRLFVQMEREGgkQLHGRLVKGGYETGCFVGNALLLMYCKCGSIEEANDLFKEMAGK------DIVSWNTMIAGYSRHGFGEVALRFFESMKREGLKPDDATMVAVLSACSHTGLVDKGRQYFYTmqDYGVMPnhYACMVDLLGRAGLLEDAHNLMKNMP----FEPDAAIWGTLLGASRVHGNTELAETAADKIFAMEPE-NSGMYVLLSNLYASSGRWGDVGKLRVRMRDKGVKKVPGY----------SWIEIQNKTHTFSVGDEFHPEKDEIFAFLEELDLRMKKAGYV------------------------    
  24 UniRef50_A9U2Q7        Predicted protein n=1 Tax=P...   467    1e-130  1  10.1%   5:597    94:670     ----SMYSKCGSIEDANNVFQSMEDKDVVSWNAMISGYALHGRGQEAVDLFYQMQREGLKPNQN--SFISILSACQTPIVLEFGEQIHSHITKAGYESDVNVSTALINMYCKCGSLELARKVFNEmeRNVVSWTAMISGYVQHGdeAFVLFQKLIRSGTQPNKVSFASILGACTnnDLEQGLKLHAYIKQAGLEQEVLVGNALISMYARCGSLANARQVFDNLRS----PNRVSWNAMIAGYG-EGFMEEAFRLFRDMQQKGFQPDRFTYASLLAICADRADLNRGKEL------------HSQIVRTAWEADVTVATALISMYAKCGSLEEARKVFNQMPEK------NAVSWNAFIACCCRHGSEKEAFQVFKQMRRDDVIPDHVTFITLLNSCTSPEDFERGRYIHGKIDQWGMLSnvANALISMYGRCGKLADAREVFYRIRRR-----DLGSWNAMIAAYVQHGANGSAFDLFIKYKSEGGKGDKYTFINVLRAIANLEDLDAGRKIHGLVEKAGLEKDIRI-LTTLIKMYSKCGSLRDAYSVfwNAMLAAYNHSDHGQDALKLFQQMRLEGVNPDS-ATYTSVLNACARLG------    
  25 UniRef50_D1I4P0        Whole genome shotgun sequen...   467    1e-130  1   9.9%   5:582   132:683     ----SLVDMYMKTESVerVFDEMRVKNVVSWTSLLAGYRQNGLNEQALKLFSQMQLEGIKPNP--FTFAAVLGGLAADGAVEKGVQVHTMVIKSGLDSTIFVGNSMVNMYSKSLMVSDAKAVFDSMerNAVSWNSMIAGFVTNglEAFELFYRMRLEGVKLTQTIFATVIKLCANIkmSFAKQLHCQVIKNGSDFDLNIKTALMVAYSKCSEIDDAFKLFCMM--HGVQ-NVVSWTAIISGYVQNGRTDRAMNLFCQmrEEGVEPNEFTFSSVLNACAatASVEQGKQF------------HSCSIKSGFSNALCVSSALVTMYAKRGNIESANEVFKR------QVDRDLVSWNSMISGYAQHGCGKKSLKIFEEMRSKNLELDGITFIGVISACTHAGLVNEGQRYFDLMVkyHIVpeHYSCMVDLYSRAGMLEKAMDLINKMP----FPAGATIWRTLLAACRVHLNVQLGELAAEKLIS--LQpdSAAYVLLSNIYATAGNWQERAKVRKLMDMKKVKK----------EAGYSWIEVKNKTFSFMAGDLSHPQSDRIYLKLEELSIRLKDAGYYPDT---------------------    
  26 UniRef50_D1HSK5        Whole genome shotgun sequen...   466    1e-130  1  11.4%   5:582    19:600     ----KVYSNSGDLQRARHLFDKIPQPDLPTWTILISALTKHGRSLEAIQYYNDFRHKNC-VEPDKLLLLSVAKACASLRDVMNAKRVHEDAIRFGFCSDVLLGNALIDMYGKCRCSEGARLVFEGMprDVISWTSMASCYVNCgrEALGAFRKMGLNGERPNSVTVSSILPACTDldLKSGREVHGFVVRNGMGGNVFVSSALVNMYASCLSIRQAQLVFDSMSRR----DTVSWNVLITAYFLNKECEKGLSVFGRMMSEGVGLNYASWNAVIGGCMQNGRTEKALEVLSRMQnrGGKQIHGYIFRHWFFQDLTTTTALVFMYAKCGDLELSRRVFSMMTKR------DTVSWNTMIIATSMHGNGEEALLLFREMVDSGVRPNSVTFTGVLSGCSHSRLVDEGLLIFDSmrDHSVEpdHHSCMVDVLSRAGRLEEAYEFIKKMP----IEPTAGAWGALLGGCRVYKNVELGRIAANRLFEIESD-NPGNYVLLSNILVSAKLWSEASETRKLMRDRGVTKNP----------GCSWIQVRNRVHTFVVGDKSNDQSDEIYRFLDYMGEKMRIAGYLPNT---------------------    
  27 UniRef50_B9NAU3        Predicted protein n=4 Tax=r...   466    1e-129  1   9.8%   3:581    28:572     --AFCSLQESGSLQYARLVFSQIPNPTSYTCNSIIRGCTDKNLHQEALLFYQEM--MVQGLIPDRYTFPSLFKSCRNSS---EGKQIHCHSTKLGFASDTYAQNTLMNMYSNCGCLVSARKVFDKMekTVVSWATMIGVHAqqPNEAVRLFDrmKSENVKPNEVTLVNVLTACARArlAMVKRIHEYIDEHGFGRHVVLNTVLMDVYCKCGCVQLARDLFDKAQEK----NLFSWNIMINGHVEDSNYEEALLLFREMQTKGIKGDKVTMASLLLACTHLGALELGKWL------------HAYIKKQRIDVDVALGTALVDMYAKCGSIETAIQVFHEMPEK------DVMTWTALILGLAMCGQAENALQYFDEMHIKGVKPDAITFVGVLAACSHAGFVDEGISHFNSMSdyGIQpeHYGGLVDILGRAGRIAEAEELIKSMP----MAPDQFVLGGLLGACRIHGNLEAAERAAKQLLE--IDphSGTYVLLSNIYKSSKKWEEAKRTRELMAERGMRKPP----------GCSQIEVHGVVHEFVKGDSSHRQSSEINEMLEDMISKLKNAGYVPD----------------------    
  28 UniRef50_Q1SMZ4        Tetratricopeptide-like heli...   465    1e-129  1  10.4%   1:597    62:628     LGAIVRMGHYPTAISLfqLQSKGISPSIATFTILINCYFHQSHTAFAFSLLATI--LKSGYQPNLVTFNTIINGFCINGMIFKALDFCQNLLAQGYLFDQFTYGTLINGLSKNGQIKAALHLLQEMEKSSVQPNLVMYSALIDgsDALGLCSQIGERGILLDAVTYNSLIDGCCsgRWQEVTQLLTKMVRENVDPDDYTFNILIDALCKEGRILEAQGVLAMMSKRGEKPDIVTYNALMEGYCSRENVHEARELFNRMVKRGLEPDVLNYNVLIDGYCKTKMVDEAMVLFKELCNKN------------LVPTIASYNSLIDGLCNSGRISHVKKLLDEM--HGSAQPPDVVTYNILIDALCKEGRILEALGVLVMMMKKGVKPNIVTYNAMMDGYCLRNNVNVAKDIFNRMVKSGLEPDiyNVLINGYCKTEMVDEAIVLFKEMRHKNLI-PDIASYNSLIDGLCNLGRIPHVQELLDEMCDSGQSPDVITYNILLDAFCKTQPFDKAISLFRQIVE--GIWPDFYTNHAIVDNLCK---------------------GEKLKMAEDALKHLLMHGCSPN-VQTYTILINALCKDG------    
  29 UniRef50_D1ISZ5        Whole genome shotgun sequen...   464    1e-129  1  10.6%   6:589   115:749     -----LYARSgdLCYARKLFEEMPERNLTAWNTMILAYARVDDYMEAWGIFDRMLKIGVC--PDNFTFASALRVCGALRSRDGGKQVHSKLIACGFKGDTFVGNALIDMYAKCDDEESCLKVFDEmeRNQVTWNSIISAEagHFNDALVLFLRMQEsgIQPDQFTFTTLLTLCANQRNdqGRQIHAHLIRANITKNIIVETELVHMYSECGRLNYAKEIFNRMAER----NAYSWNSMIEGYQQNGETQEALRLFKQMQLNGIKPDCFSLSSMLSScaKCGSMDYAWKVYDQTIkkESFNHFLEMLESDIEYDVLTmtALVDMYSKCGAITKARTVFDNMNGK------NIVSWNAMISGYSKHGCSKEALILYEEMPKKGMYPNEVTFLAILSACSHTGLVEEGLRIFTSMQeyNikAEHYTCMVDLLGRAGRLEDAKEFVEKMP----IEPEVSTWGALLGACRVHKDMDMGRLAAQRLFELDPQ-NPGPYVIMSNIYAAAGRWKEVEDIRQMMKMKGVKKDP----------GVSWIEINSEIQIFHAGSKTHPKTEEIYNNLRHLTLQSKGLGYIPDTSFILQNV--------------    
  30 UniRef50_Q10QC8        Os03g0201300 protein n=4 Ta...   464    1e-129  1  12.1%  11:599   191:786     ----------ASLDVFRSLIELRLHPNHYTFNLLVHTHCSKGTLADALATLSTMQ--GFGLSPDAVTYNTLLNAHCRKGMLGEARALLARMKRDGIAPTQPTYNTLVSAFARLGWIKQATKVVESmePDLRTYNVLAVGLcgKVDEAFRLKDEMERLSTapDVVTYNTLVDACFKWRCsdALRLLEEMRDKGVKPTLVTHNIVVKSLCKEGKLEEALGKLEKIAEEGLAPDVITYNTLIDAYCKAGNVAKAFTLMDEMVGKGLKMDTFTLNTVLYNLCKMKRYEDAEELLHS------------PPQRGFVPDEVSYGTVMAAYFKEYNPEPALRLWDQMIER--KLIPSISTYNTLIKGLCRMERLKEAIDKLNELVEKGLVPDETTYNIIIHAYCKEGDLENAFRFHNKMVENSFKPdtCNTLMNGLCLHGKLDKALKLFESWVEKGKKV-DVITYNTLIQSMCKVGDVDTALHFFDDMEVKGLQPDAFTYNVVLSALSEAGRSEEAHNMLHKLADSGK---LSQSFACPLLKPSSADEADVKEHEgyMERLNGLCTGGQLKEAKAVLDEMMQKGM-PVDSSTYITLMEGLIKRQKR----    
  31 UniRef50_Q0DUV9        Os03g0165100 protein n=3 Ta...   463    1e-129  1  11.7%  12:523    79:583     -----------AADAYRLVFSSNSEVNTYTLNIMVHNYCKALEFDKVDAVISEME--KRCVFPDVVTHNVMVDARFRAGDAEAAMALVDSMVSKGLKPGIVTYNSVLKGLCRSGMWDKAWEVFKEMDDFGVAPDvlIGGFCreIEEALKIYKEMRHRGIKPDLVSFSCLIGLFArgKMDHAMAYLREMRCFGLVPDGVIYTMVIGGFCRAGLMSDALRVRDEMVGCGCLPDVVTYNTLLNGLCKERRLLDAEGLLNEMRERGVPPDLCTFTTLIHGYCIEGKLDKALQLFDTMLNQR------------LRPDIVTYNTLIDGMCRQGDLDKANDLWDDMHSR--EIFPNHVTYSILIDSHCEKGQVEDAFGFLDEMINKGILPNIMTYNSIIKGYCRSGNVSKGQKFLQKMMVNKVSPdtYNTLIHGYIKEDKMHDAFKLLN-MMEKEKVQPDVVTYNMLINGFSVHGNVQEAGWIFEKMCAKGIEPDRYTYMSMINGHVTAGNSKEAFQLHDEMLQRGKEK--------------------------------------------------------------------------------    
  32 UniRef50_O82380        Pentatricopeptide repeat-co...   463    1e-129  1  10.6%   3:580    70:648     --AMAALSSFASLEYARKVFDEIPKPNSFAWNTLIRAYASGPDPVLSIWAFLDMVS-ESQCYPNKYTFPFLIKAAAEVSSLSLGQSLHGMAVKSAVGSDVFVANSLIHCYFSCGDLDSACKVFTTieKDVVSWNSMINGFVQKgdKALELFKKMESEDVKASHVTMVGVLSACAkrNLEFGRQVCSYIEENRVNVNLTLANAMLDMYTKCGSIEDAKRLFDAMEEK----DNVTWTTMLDGYAISEDYEAAREVLNSMPQK----DIVAWNALISAYEQNGKPNEALIVFHELQLQkgRWIHSYIKKHGIRMNFHVTSALIHMYSKCGDLEKSREVFNSVEKR------DVFVWSAMIGGLAMHGCGNEAVDMFYKMQEANVKPNGVTFTNVFCACSHTGLVDEAESLFHQMEsyGIVPehYACIVDVLGRSGYLEKAVKFIEAMP----IPPSTSVWGALLGACKIHANLNLAEMACTRLLE--LEpnDGAHVLLSNIYAKLGKWENVSELRKHMRVTGLKKEP----------GCSSIEIDGMIHEFLSGDNAHPMSEKVYGKLHEVMEKLKSNGYEP-----------------------    
  33 UniRef50_Q9SIT7        Pentatricopeptide repeat-co...   462    1e-128  1  10.2%   7:574    96:669     ------LTKLGFLDEADSLFRSMPERDQCTWNSMVSGFAQHDRCEEALCYFAMMHKEGFV--LNEYSFASVLSACSGLNDMNKGVQVHSLIAKSPFLSDVYIGSALVDMYSKCGNVNDAQRVFDEmdRNVVSWNSLITCFEQNgvEALDVFQMMLESRVEPDEVTLASVISACASLsiKVGQEVHGRVVKNdlRNDIILSNAFVDMYAKCSRIKEARFIFDSMPIRNveRNVVSWNALIAGYTQNGENEEALSLFCLLKRESVCPTHYSFANILKACADLAELHLGMQAHVHVLK------HGFKFQSGEEDDIFVGNSLIDMYVKCGCVEEGYLVFRKMMER------DCVSWNAMIIGFAQNGYGNEALELFREMLESGEKPDHITMIGVLSACGHAGFVEEGRHYFSSMTRdgVApdHYTCMVDLLGRAGFLEEAKSMIEEMP----MQPDSVIWGSLLAACKVHRNITLGKYVAEKLLE--VEpnSGPYVLLSNMYAELGKWEDVMNVRKSMRKEGVTKQP----------GCSWIKIQGHDHVFMVKDKSHPRKKQIHSLLDILIAEMR-----------------------------    
  34 UniRef50_Q9LYZ9        Pentatricopeptide repeat-co...   462    1e-128  1  13.4%   5:602   181:788     ----SMLGKEGRVSSAANMfeDGFSLDVYSYTSLISAFANSGRYREAVNVFKKMEEDGCKPTL--ITYNVILNVFGKMGTPwkITSLVEKMKSDGIAPDAYTYNTLITCCKRGSLHQEAAQVFEEMKAAGFSYDKVtyGKSHRPKEAMKVLNEMVLNGFSPSIVTYNSLISAYARDglDEAMELKNQMAEKGTKPDVFTYTTLLSGFERAGKVESAMSIFEEMRNAGCKPNICTFNAFIKMYGNRGKFTEMMKIFDEINVCGLSPDIVTWNTLLAVFGQNGMDSEVSGVFKEMKR------------AGFVPERETFNTLISAYSRCGSFEQAMTVYRRMLDAG--VTPDLSTYNTVLAALARGGMWEQSEKVLAEMEDGRCKPNELTYCSLLHacSKCDLLPEAERAFSELKERGFSPdtLNSMVSIYGRRQMVAKANGVLDYMKERG-FTPSMATYNSLMYMHSRSADFGKSEEILREILAKGIKPDIISYNTVIYAYCRNTRMRDASRIFSEMRNSGIVPDVIT----------------------YNTFIGSYAADSMFEEAIGVVRYMIKHGCRPNQ-NTYNSIVDGYCKLNRKDEA-    
  35 UniRef50_C5XUZ8        Putative uncharacterized pr...   462    1e-128  1  13.0%  11:602   112:704     ----------GRVADAERVFalGASATVVTYNTMVNGYCRAGRIEDARRLISGM-----PFPPDTFTFNPLIRALCVRGRVPDALAVFDDMLHRGCSPSVVTYSILLDATCKASGYRQAMVLLDEmePDIVTYNVLINAMcgDVDEALNILSDLPSHGCKPDAVTYTPVLklCGSERWKEVEELFAEMASNKCAPDEVTFNTIVTSLCQQGLVDRAIKVVDHMSEHGCIPDIVTYSSILDGLCDVGRVDDAVELLSRLKSYGCKPDTIAYTTVLKGLCSIEQWEHAEELMAEMVCSD------------CPPDEVTFNTVIASLCQKGLVDRAIKVVEQMSENGCN--PDIVTYNSIIDGLCNERCIDDAMELLSNLQSYGCKPDIVTFNTLLKGLCSVDRWEDAEQLMVNMMHSDCPPdtFNTVITSLCQKGLLLQAIETLKIMAENGCI-PNQSTYNIVVDALLKAGKTQEALKLLSGM--TNGTPDLITYNTVISNITKAGKMEEALDLLRVMVSNGLSPDTITYRscREDGTDRAIRMLCRLqtFYNDILLGFCQNWRTDIAIDCFAHMVSSGCMPDE-STYIILLEALAYECLLDEA-    
  36 UniRef50_D1HK23        Whole genome shotgun sequen...   461    1e-128  1  11.5%   7:582   196:737     ------YSKNGNIEEARLVFDQLSEKTAVTWTTIIAGYTKCGRSAVSLELFAQMRE--TNVVPDRYVVSSVLSACSMLEFLEGGKQIHAYVLRRGTEMDVSVVNVLIDFYTKCNRVKAGRKLFDQMvkNIISWTTMISGYMQNswEAMKLFGEMNRLGWKPDGFACTSVLTSCGSRelEQGRQVHAYTIKANLESDEFVKNGLIDMYAKSNLLIDAKKVFDVMAEQ----NVISYNAMIEGYSSQEKLSEALELFHEMRVRLQKPNEFTFAALITAASNLASL------------RHGQQFHNQLVKMGLDFCPFVTNALVDMYAKCGSIEEARKMFN------SSIWRDVVCWNSMISTHAQHGEAEEALGMFREMMKEGIQPNYVTFVAVLSACSHAGRVEDGLNHFNSMPGFGIKpeHYACVVSLLGRSGKLFEAKEFIEKMP----IEPAAIVWRSLLSACRIAGNVELGKYAAEMAISTDPK-DSGSYILLSNIFASKGMWADVKKVRDRM-------DSSEVVKEPGRSW---IEVNNKVNVFIARDTTH-READIGSVLDILIQHIKGAGYVPDA---------------------    
  37 UniRef50_Q9SN39        Pentatricopeptide repeat-co...   461    1e-128  1   9.5%   4:602   234:799     ---NSLVAFYLkvDSARKVFDEMTERDVISWNSIINGYVSNGLAEKGLSVFVQM--LVSGIEIDLATIVSVFAGCADSRLISLGRAVHSIGVKACFSREDRFCNTLLDMYSKCGDLDSAKAVFREmdRSVVSYTSMIAGYAREGleAVKLFEEMEEEGISPDVYTVTAVLNCCARYRldEGKRVHEWIKENDLGFDIFVSNALMDMYAKCGSMQEAELVFSEMRVK----DIISWNTIIGGYSKNCYANEALSLFNLLLekRFSPDERTVACVLPACASLSAFDKGREI------------HGYIMRNGYFSDRHVANSLVDMYAKCGALLLAHMLFDDIASK------DLVSWTVMIAGYGMHGFGKEAIALFNQMRQAGIEADEISFVSLLYACSHSGLVDEGWRFFNIMreCKIEpeHYACIVDMLARTGDLIKAYRFIENMP----IPPDATIWGALLCGCRIHHDVKLAEKVAEKVFE--LEpnTGYYVLMANIYAEAEKWEQVKRLRKRIGQRGLRKNP----------GCSWIEIKGRVNIFVAGDSSNPETENIEAFLRKVRARMIEEGYSP---LTKYALIDA--EEMEKEEA-    
  38 UniRef50_B9HA20        Predicted protein n=3 Tax=r...   461    1e-128  1  12.4%  11:599   242:834     ----------EAEGWFREAKNKGVELDARAYSIVIEAVC---KKPDSVAALGLLREMRDKgvPHEVIFTRVIGVCMKQGKMLEAVKVKGEMLSCGKPMNVVVATTLMKGYCKQGDLDSALELFDKMNENGICPNNVTYacCKNgdKAYEIYNQMKNKDISPTVFNVNSLIRGYLKARseEASKLFDEAVACGIA-NVFTYNSLLSWLCKEGKMSEACSIWEKMVRKGVRPSVVSYNNMILGHCQQGDMDSANGVFVEMLEKGLKPNLITYSVLMDGYFKKGDTEYAFGLYDRMRGEN------------IAPSDFTCNIIINGLCKAGRTSESQDRLKKLVQEG--FIPTCMTYNCIIDGFVKEGSVNSALAVYTEMCKIGVSPNVFTYTNLINGFCKSNNMDLALKVMDEMKNKGIELDvyCALIDGFCRKGDMVNASQLLSELQEVGL-SPNKVVYSSMISGFRKLQNMEAALHLHKRMINEGIPCDLQIYTTLISGLLKEGKLLFASELYAEMLAKGIMPDLIT-YSVLIHGLCNKGQLENAQKILEDMDRkgHFKEGNLQEAFRLHNEMLDKGLVPDD-TTYDILVNGKVKDGNL----    
  39 UniRef50_D1I4W0        Whole genome shotgun sequen...   459    1e-127  1  13.4%   1:546   173:709     LAKFEMVEGARNL--YKQMLNSGIQPSLLTFNTLINILSKKGKVREAELILSQI--FQYDLSPDVFTYTSLILGHCRNRNLDLAFGVFDRMVKEGCDPNSVTYSTLINGLCNEGRVDEALDMLEEMIEKGIEPTVYTYTLPITaeEAIELVARMKKRGCRPNVQTYTALISGLSrgKLEVAIGLYHKMLKEGLVPNTVTYNALINELCVGGRFSTALKIFHWMEGHGSLANTQTYNEIIKGLCLGGDIEKAMVLFEKMLKMGPLPTVVTYNTLINGYLTKGNVNNAARLLDLMKE------------NGCEPDEWTYNELVSGFSKWGKLESASFYFQEMVE--CGLNPNPVSYTALIDGHSKDGKVDIALSLLKRMEEMGCNPNVESYNAVINGLSKENRFSEAEKICDKMVEQGLLPntYTTLIDGLCRNGRTQFAFKIFHDMEKRKCL-PNLYTYSSLIYGLCQEGKADEAEILLKEMERKGLAPDEVTFTSLIDGFVVLGRIDHAFLLLRRMVDMGCKPNYRTYSVLLKGLQKECLLLEEKI---------------------------------------------------------    
  40 UniRef50_Q3E6Q1        Pentatricopeptide repeat-co...   458    1e-127  1  10.1%   6:582   179:723     -----MYAKCRQVNEARKVFDRMPERDLVSWNTIVAGYSQNGMARMALEMVKSMCEENL--KPSFITIVSVLPAVSALRLISVGKEIHGYAMRSGFDSLVNISTALVDMYAKCGSLETARQLFDGmeRNVVSWNSMIDAYVQnpKEAMLIFQKMLDEGVKPTDVSVMGALHACAdgDLERGRFIHKLSVELGLDRNVSVVNSLISMYCKCKEVDTAASMFGKLQSR----TLVSWNAMILGFAQNGRPIDALNYFSQMRSRTVKPDTFTYVSVITAIAELSITHHAKWI------------HGVVMRSCLDKNVFVTTALVDMYAKCGAIMIARLIFDMMSERH------VTTWNAMIDGYGTHGFGKAALELFEEMQKGTIKPNGVTFLSVISACSHSGLVEAGLKCFYMMKEneLSMDHYGAMVDLLGRAGRLNEAWDFIMQMPVK----PAVNVYGAMLGACQIHKNVNFAEKAAERLFELN--PDDGGYHVLlnIYRAASMWEKVGQVRVSMLRQGLRKTP----------GCSMVEIKNEVHSFFSGSTAHPDSKKIYAFLEKLICHIKEAGYVPDT---------------------    
  41 UniRef50_Q5ZDP1        Os01g0355000 protein n=10 T...   457    1e-127  1  10.3%   3:582   240:790     --ANALVDMYMKMGraSVIFEKMPDSDVVSWNALISGCVLNGHDHRAIELLLQMKYSGLV--PNVFTLSSILKACSGAGAFDLGRQIHGFMIKANADSDDYIGVGLVDMYAKNHFLDDARKVFDWmhRDLILCNALISGCsgRHDEALSLFYELRKEGLGVNRTTLAAVLK--STASLEAarQVHALAVKIGFIFDAHVVNGLIDSYWKCSCLSDANRVFEECSS----GDIIACTSMITALSQCDHGEGAIKLFMEMLRKGLEPDPFVLSSLLNACASLSAYEQGKQV------------HAHLIKRQFMSDAFAGNALVYTYAKCGSIEDAELAFSSLPERG------VVSWSAMIGGLAQHGHGKRALELFGRMVDEGINPNHITMTSVLCACNHAGLVDEAKRYFNSMKEmgIDreHYSCMIDLLGRAGKLDDAMELVNSMP----FQANASIWGALLGASRVHKDPELGKLAAEKLFI--LEPesGTHVLLANTYASAGMWNEVAKVRKLMKDSNIKKEPA----------MSWIEVKDKVHTFIVGDKSHPMTKEIYAKLVELGDLMSKAGFVPNV---------------------    
  42 UniRef50_Q9FIX3        Pentatricopeptide repeat-co...   457    1e-127  1  12.5%   3:584   146:747     --RLSLIDKALSIVHLAQAHGFM--PGVLSYNAVLDATIRSKriSFAENVFKEMLE--SQVSPNVFTYNILIRGFCFAGNIDVALTLFDKMETKGCLPNVVTYNTLIDGYCKLRKIDDGFKLLRSmePNLISYNVVINGLCREGreVSFVLTEMNRRGYSLDEVTYNTLIKGYCkgNFHQALVMHAEMLRHGLTPSVITYTSLIHSMCKAGNMNRAMEFLDQMRVRGLCPNERTYTTLVDGFSQKGYMNEAYRVLREMNDNGFSPSVVTYNALINGHCVTGKMEDAIAVLEDMKEK------------GLSPDVVSYSTVLSGFCRSYDVDEALRVKREMVEKG--IKPDTITYSSLIQGFCEQRRTKEACDLYEEMLRVGLPPDEFTYTALINAYCMEGDLEKALQLHNEMVEKGVLPdtYSVLINGLNKQSRTREAKRLLLKLFYEESV-PSDVTYHTLikGFCMKGMMTEADQVFESMLGKNHKPDGTAYNIMIHGHCRAGDIRKAYTLYKEMVKSGFLLHTVteLNSVIVHVLRSCELSEAEQAKVLVEINH--REGNMDVVLDVLAEMAKDGFLPNGIS-------------------    
  43 UniRef50_D1H9V5        Whole genome shotgun sequen...   457    1e-127  1  11.2%  39:601    66:606     --------------------------------------CKSgkRSEAFSVFNHLIDMQ--PTPPISSFNTLLGAVAKIKRYFDVISLYKRMSLIGLAPDFITLNILINCYCNLNKVDFGLAVLGEMLRRGHSPNTVTFTSLVKgsEATGLLRKMVRMGYRPNVVTYGTLLNGLctGNTMLAVKLHEEMLNGniKPNLVCYCTIIDSLCKDGLIDKGKELFLEMKGRGISPDVVAYSSIIHGMCHTGRWEGAKGLFNEMVDEGVHPNVVTFNVLIDALCKAGKMEEANHLLKLMIQR------------GESPDTFTYNTLIDGFCLEGRIDDARDLFVSMESKG--IETDAVSYNVLINGYCKSGRMVEAKKLYREMMCKEIMPTVITYNTLLTGLFREGKVRDAWNLFGEMKVHDLTPetYNILLDGLCKNNHLSEAMELFHYL-ENHDFQPSIQIFNCLIDGLCKARKIEIARELFNRLSHEGLEPNVITYTVMIHGLCKSGQLENAKDLFLGMEEKG----------CAPNLVTFNTLMRGFCQ------------NDEMQKVVELLQEMAEKDFSPDA-STISIVVDLLSKDEKYRE--    
  44 UniRef50_Q9LQ16        Pentatricopeptide repeat-co...   455    1e-126  1  11.6%   9:601    63:625     --------KVDDAVDLfdMVKSRPFPSIVEFNKLLSAVAKMNKFELVISLGEQMQTLGISH--DLYTYSIFINCFCRRSQLSLALAVLAKMMKLGYEPDIVTLSSLLNGYCHSKRISDAVALVDQMVEMGYKPDTFTFTTLIHgsEAVALVDQMVQRGCQPDLVTYGTVVNGLCkgDIDLALSLLKKMEKGKIEADVVIYNTIIDGLCKYKHMDDALNLFTEMDNKGIRPDVFTYSSLISCLCNYGRWSDASRLLSDMIERKINPNVVTFSALIDAFVKEGKLVEAEKLYDEMIKRS------------IDPDIFTYSSLINGFCMHDRLDEAKHMFELMISKDCF--PNVVTYSTLIKGFCKAKRVEEGMELFREMSQRGLVGNTVTYTTLIHGFFQARDCDNAQMVFKQMVSVGVHPntYNILLDGLCKNGKLAKAMVVFEYL-QRSTMEPDIYTYNIMIEGMCKAGKVEDGWELFCNLSLKGVSPNVIAYNTMISGFCRKGSKEEADSLLKKMKEDG----------PLPNSGTYNTLIRARL------------RDGDREASAELIKEMRSCGFAGDASTI--GLVTNMLHDGRLDK--    
  45 UniRef50_UPI0001984A05 PREDICTED: similar to Tetra...   454    1e-126  1   9.8%  27:599   123:668     --------------------------NVFSWNSVIAELARSGDSVEALRAFSSMRKL--SLKPNRSTFPCAIKSCSALLDLHSGRQAHQQALIFGFEPDLFVSSALVDMYSKCGELRDARTLFDEihRNIVSWTSMITGYVQnaHRALLLFKEFLVEevCVDPIAMVSVLSACSRVSEKstEGVHGFLIKRGFEGDLGVENTLMDAYAKCGELGVSRRVFDGMAER----DVISWNSIIAVYAQNGMSTESMEIFHRMVKDgiNYNAVTLSAVLLACAHSGS------------QRLGKCIHDQVIKMGLESNVFVGTSIIDMYCKCGKVEMARKAFDRMREK------NVKSWSAMVAGYGMHGHAKEALEVFYEMNMAGVKPNYITFVSVLAACSHAGLLEEGWHWFKAmhEFDVEpeHYGCMVDLLGRAGYLKEAFDLIKGMKLR----PDFVVWGALLGACRMHKNVDLGEISARKLFELDPK-NCGYYVLLSNIYADAGRWEDVERMRILMKNSGLVKPPGF----------SLVDIKGRVHVFLVGDREHPQHEKIYEYLEKLSMKLQEVGYVPDM----TSVLHDVGHEEKE----    
  46 UniRef50_C9W4B9        PPR-817 n=5 Tax=Zea mays Re...   454    1e-126  1  10.2%   3:582   152:747     --ANHLLEGFCEAKRTDEALdlGCVPDVFSYSILLKSLCDQGKSGQADDLLRMMAEGGAVCSPNVVAYNTVIDGFFKEGDVNKACDLFKEMVQRGIPPDLVTYNSVVHALCKARAMDKAEAFLRQMvpNNWTYNNLIYGYsgQWKEAVRVFKEMRRHSILPDVVTLSMLMGSLCkgKIKEARDVFDTMAMKGQNPDVFSYNIMLNGYATKGCLVDMTDLFDLMLGDGIAPDFYTFNVLIKAYANCGMLDKAMIIFNEMRDHGVKPDVVTYRTVIAALCRIGKMDDAMEKFNQM------------IDQGVAPDKYAYNCLIQGFCTHGSLLKAKELISEIMNNGMHL--DIVFFSSIINNLCKLGRVMDAQNIFDLTVNVGLHPDAVVYNMLMDGYCLVGKMEKALRVFDAMVSAGIEPNvyGTLVNGYCKIGRIDEGLSLFREMLQRGIK-PSTILYSIIIDGLFEAGRTVPAKMKFHEMTESGIAMDICTYNIVLRGLFKNRCFDEAIFLFKELRAMNVKINIIT-LNTMIDGMFQTRRVEEAKDLfySIMITNLIKEGLVEEAEDMFSSMQNAGCEPNS---------------------    
  47 UniRef50_Q9CAA8        Putative pentatricopeptide ...   453    1e-126  1  11.2%  10:587    82:661     ---------YSKAGLISEmfEKLPDRDGVTWNVLIEGYSLSGLVGAAVKAYNTMMR-DFSANLTRVTLMTMLKLSSSNGHVSLGKQIHGQVIKLGFEsnTVMYNSLMGGLLACGMIEDALQLFRGMEKDSVSWAAMIKGLAQNgkEAIECFREMKVQGLKMDQYPFGSVLPACGGLgiNEGKQIHACIIRTNFQDHIYVGSALIDMYCKCKCLHYAKTVFDRMKQK----NVVSWTAMVVGYGQTGRAEEAVKIFLDMQRSGIDPDHYTLGQAISACANVSSLEEG------------SQFHGKAITSGLIHYVTVSNSLVTLYGKCGDIDDSTRLFNEMNVR------DAVSWTAMVSAYAQFGRAVETIQLFDKMVQHGLKPDGVTLTGVISACSRAGLVEKGQRYFKLMteYGIVPshYSCMIDLFSRSGRLEEAMRFINGMP----FPPDAIGWTTLLSACRNKGNLEIGKWAAESLIE--LDPHhaGYTLLSSIYASKGKWDSVAQLRRGMREKNVK-------KEPGQSWIKW---KGKLHSFSADDESSPYLDQIYAKLEELNNKIIDNGYKPDTSFVHH----------------    
  48 UniRef50_Q9M9X9        Pentatricopeptide repeat-co...   453    1e-125  1  13.0%   1:600   312:936     LCEASLFEEAMDF--LNRMRATSCLPNVVTYSTLLCGCLNKKQLGRCKRVLNMMMMEGCYPSPKI--FNSLVHAYCTSGDHSYAYKLLKKMVKCGHMPGYVVYNILIGSIcnCDLLDLAEKAYSEMLAAGVVLnsFTRCLcgKYEKAFSVIREMIGQGFIPDTSTYSKVLNYLCNakMELAFLLFEEMKRGGLVADVYTYTIMVDSFCKAGLIEQARKWFNEMREVGCTPNVVTYTALIHAYLKAKKVSYANELFETMLSEGCLPNIVTYSALIDGHCKAGQVEKACQIFERMCGsdVDMYFKQYDDNSERPNVVTYGALLDGFCKSHRVEEARKLLDAMSMEGC--EPNQIVYDALIDGLCKVGKLDEAQEVKTEMSEHGFPATLYTYSSLIDRYFKVKRQDLASKVLSKMLENSCAPNvyTEMIDGLCKVGKTDEAYKLMQMMEEKGC-QPNVVTYTAMIDGFGMIGKIETCLELLERMGSKGVAPNYVTYRVLIDHCCKNGALDVAHNLLEEMKqaGYRKVIEGFNKEFIESLGLLDEIgaPFLSVYRLLIDNLIKAQRLEMALRLLEEVasATLV-DYSSTYNSLIESLCLANKVE---    
  49 UniRef50_Q6ZLH4        Os07g0590600 protein n=3 Ta...   453    1e-125  1  12.4%   3:599   147:749     --AIRVMARAGRLAEASALLDAAPGPDAGAYTALVSAFSRAGRFRDAVAVFRRMVDSGVQPA--IVTYNVVLHVYSKMAVPweVVELVASMKEHGVAPDRYTYNTLISCCRRRALYKEAAQVFDEMKASGFEPDKVtyGKARRHDEAIEVIQEMERVGCPPSVVTYNSLISSYVKDGleQAVALKQEMEVKGMKPDVVTYTTLISGLDRAGKIDAAIVEYDEMVRNGCKPNLCTYNALIKMHGVRGKFPEMMAVFDEFRSAGFVPDIVTWNTLLAVFGQNGLDSEVSGVFKEMK------------KAGYIPERDTYVSLISSYSRCGLFDLAMQIYKRMMEAG--IYPDVSTYNAVLSALARGGRWEQAEKLFAEMEERDCKPDEYSYSSLLHAYANAKRLdeAEKAFLELRQKRcdINVLNAMVSIYGKNRMVRKVEKILSLMKESAI-NLSAATYNSLMHMYSRLGDCEKCENILTEIKSSGVRPDRYSYNTVIYAYGRKGQMKEASRLFSEMKCSGLKP----------DVVTYNIFVKSYV------------SNSMFEEAIELVRYMVTQGCKPNE-RTYNSIVEGYCRNGKL----    
  50 UniRef50_O81767        Pentatricopeptide repeat-co...   452    1e-125  1  10.4%   6:582   194:736     -----LYSRYKAVGNARILFDEMPVRDMGSWNAMISGYCQSGNAKEALTLSNGLRAM------DSVTVVSLLSACTEAGDFNRGVTIHSYSIKHGLESELFVSNKLIDLYAEFGRLRDCQKVFDRmvRDLISWNSIIKAYELnpLRAISLFQEMRLSRIQPDCLTLISLASILSqgDIRACRSVQGFTLRKgfLEDITIGNAVVVMYAKLGLVDSARAVFNWLPN----TDVISWNTIISGYAQNGFASEAIEMYNIMEEEgiAANQGTWVSVLPACSQAGALRQGMKL------------HGRLLKNGLYLDVFVVTSLADMYGKCGRLEDALSLFYQIPR------VNSVPWNTLIACHGFHGHGEKAVMLFKEMLDEGVKPDHITFVTLLSACSHSGLVDEGQWCFEMMqdYGITPshYGCMVDMYGRAGQLETALKFIKSMS----LQPDASIWGALLSACRVHGNVDLGKIASEHLFE--VEPEHVGyvLLSNMYASAGKWEGVDEIRSIAHGKGLRKTP----------GWSSMEVDNKVEVFYTGNQTHPMYEEMYRELTALQAKLKMIGYVPDH---------------------    
  51 UniRef50_A1YKE2        Auxin efflux carrier n=2 Ta...   452    1e-125  1  10.5%  19:601   234:820     ------------------MEGAGILPDVYTYSTFIEAHCKARDFDAAKKVFEEMR--RRDCAMNEVTYNVMISGLCRSGAVEEAFGFKEEMVDYGLSPDAFTYGALMNGLCKGSRLKEAKALLDEMSCSGLKPNIVVYGTLVDgaEAFDILKEMISAGVQPNKIMYDNLIRGLCkgQLGRASKLLKEMIKVGLRPDTFTYNPLMQGHFQQHDKDGAFELLNEMRNSGILPNVYSYGIMINGLCQNGESKEAGNLLEEMISEGLKPNAFMYAPLIIGHSKEGNISLACEALEKM------------TKANVHPDLFCYNSLIKGLSTVGRMEEAEEYYAQVQKRG--LVPDEFTYSGLIHGYCKTGNLEKADQLLRQMLNSGLKPNADTYTDLLEGYFKSNDYEKVSSILQSMLGSGDKPdiYGIVIRNLSRSENMEVAFMVLTE-VEKNGLVPDLHIYSSLISGLCKIADMEKAVGLLDEMAKEGLEPGIVCYNALIDGFCRSGDISRARNVFDSILAKGLVPNCVT-YTALIDGNCKNGDITDAFDLYKEMlaTGCSDAADLEQALFLTEEMFNRGYA--NVSLFNTLVHGFCKRGKLQE--    
  52 UniRef50_UPI0001983FBF PREDICTED: hypothetical pro...   452    1e-125  1  10.4%   6:589   325:909     -----MYSKCGMVEEAHQVFRSLLERNEVSYNALLAGYVQEGKAEEALELYHDMQS-EDGIQPDQFTFTTLLTLCANQRNDNQGRQIHAHLIRANITKNIIVETELVHMYSECGRLNYAKEIFNRmeRNAYSWNSMIEGYQQNGeeALRLFKQMQLNGIKPDCFSLSSMLSSCVssDSQKGRELHNFIVRNTMEEEGILQVVLVDMYAKCGSMDYAWKVYDQTIKKDVILN----NVMVSAFVNSGRANDAKNLFDQMEQR----NTALWNSILAGYANKGLKKESFNHFLEMLeeHGDQLHSLIIKKGFvcSVVLETALVDMYSKCGAITKARTVFDNMNGK------NIVSWNAMISGYSKHGCSKEALILYEEMPKKGMYPNEVTFLAILSACSHTGLVEEGLRIFTSMQeyNIEaeHYTCMVDLLGRAGRLEDAKEFVEKMP----IEPEVSTWGALLGACRVHKDMDMGRLAAQRLFELDPQ-NPGPYVIMSNIYAAAGRWKEVEDIRQMMKMKGVKKDP----------GVSWIEINSEIQIFHAGSKTHPKTEEIYNNLRHLTLQSKGLGYIPDTSFILQNV--------------    
  53 UniRef50_Q9M907        Pentatricopeptide repeat-co...   451    1e-125  1  10.8%   1:602   175:781     IGAFSAVnsDMMLTLFQQMQELGYEPTVHLFTTLIRGFAKEGRVDSALSLLDEMK--SSSLDADIVLYNVCIDSFGKVGKVDMAWKFFHEIEANGLKPDEVTYTSMIGVLCKANRLDEAVEMFEHLEKntYAYNTMiyGSAGKFDEAYSLLERQRAKGSIPSVIAYNCILTCLRkgKVDEALKVFEEMKK-DAAPNLSTYNILIDMLCRAGKLDTAFELRDSMQKAGLFPNVRTVNIMVDRLCKSQKLDEACAMFEEMDYKVCTPDEITFCSLIDGLGKVGRVDDAYKVYEKmrKEDGHKIYKDMINQNCSPDLQLLNTYMDCMFKAGEPEKGRAMFEEIKAR--RFVPDARSYSILIHGLIKAGFANETYELFYSMKEQGCVLDTRAYNIVIDGFCKCGKVNKAYQLLEEMKTKGFEpvTYGSVIDGLAKIDRLDEAYMLFEEAKSK-RIELNVVIYSSLIDGFGKVGRIDEAYLILEELMQKGLTPNLYTWNSLLDALVKAEEINEALVCFQSMKE----------LKCTPNQVTYGILINGLCKV------------RKFNKAFVFWQEMQKQGMKPSTI-SYTTMISGLAKAGNIAEA-    
  54 UniRef50_B9IK55        Predicted protein n=5 Tax=M...   450    1e-125  1  10.8%  35:582    46:589     ----------------------------------IHILCQQNRLKEALQILHQIDK------PSASVYSTLIQSCIKSRLLQQGKKVHQHIKLSGFVPGLFILNRLLEMYAKCDSLMDSQKLFDEmeRDLCSWNILISGYAKMgqEAKSLFDKMPER----DNFSWTAMISGYVRHDRpeALELFRMMKRsgLDSDEVVWSALSDMYGKCGSIEEARHIFDKMVDR----DIVTWTAMIDRYFQDGRRKEGFDLFADLLRSGIRPNEFTFSGVLNACANQTSEELGKKV------------HGYMTRVGFDPFSFAASALVHMYSKCGNMVSAERVFKE------TPQPDLFSWTSLIAGYAQNGQPDEAIRYFELLVKSGTQPDHITFVGVLSACAHAGLVDKGLDYFHSIKeyGltADHYACIIDLLARSGQFDEAENIISKMSMK----PDKFLWASLLGGCRIHGNLKLAQRAAEALFE--IEpnPATYVTLANIYATAGMWSEVAKIRKTMDDRGVVKKP----------GLSWIAIKRDVHVFLVGDDSHPKSKEINEFLGKLSKRMKEEGFVPDT---------------------    
  55 UniRef50_Q10AR1        Pentatricopeptide, putative...   450    1e-125  1  10.7%   6:575    85:617     -----MYSRCGSLPAAKEVFDRMDSPDVVGYTSLISAFCRNGEFELAAEALIQM--LKQGLKPNEHTMTTILTACPRV----LGQQIHGYLIKklRSQSVYSSTALIDFYSRNGEFKLAKAVFDSLhkNVVSWCSMMQLYIRDgeEALQVFGDMISEGVDPNEFALSIVLGACGSIGL--GRQLHCSAIKHDLITDIRVSNALLSMYGRTGLVEE----LEAMLNKIENPDLVSWTTAISANFQNGFGEKAIALLCQMHSEGFTPNGYAFSSVLSSCADVASLDQGMQF------------HCLALKLGCDSEICTGNALINMYSKCGQMGSARLAFDVM------HTHDVTSWNSLIHGHAQHGDANKALEVFSKMRSNGIKPDDSTFLGVLMGCNHSGMVEEGELFFRLMIdySFtpSHYACMIDMLGRNGRFDEALRMINDMP----FEPDALIWKTLLASCKLHRNLDIGKLAADRLMELSDR-DSASYVLMSNIYAMHGEWEDARKVRRRMDETGVKK----------DAGCSWIEINNEVHTFASRDMSHPNSDSIYQMLGELVAVMQD----------------------------    
  56 UniRef50_A9TMK7        Predicted protein n=1 Tax=P...   449    1e-124  1  11.6%   1:601    92:696     LGRESQLTTVSRL--FKSLREEGYPLDVYAYTSLISALSRNRKFKEALGFFEQMKEAG--PQPSLVTYNVIIDLYGKKGrwENILELFEEMKAQGIQPDEYTYNTAITACASGSLCEEATELFTQMkpDRVTYNALlyGKAGWYNEAANVLKEMESAGCLPNIVTYNELLSAFGRAGlnAAAEMKDSMVSKGIEPDVFTYTSLLSAYSRAGKVEQAMEIYNQMRTSNCTPNSFTFNALIGMHGKNKNFSEMMVIFEDMQACGVEPDIVTWNSLLGAFGKNGMYSEVLKVFRGMK------------KAGFEPDKATFNILIEAYGRCGSSDQALSIYDGMLQAGC--TPDLATFNTLLAALAREGRWEHAELILDELNRSSYKPNDIAYASMLHAYANGglVDEAEDAFLAMRHHGYLSdtFNAMISMYGKKGMMDKATDTF-ALLRSTGLEPDVVTYNCLMGMYGREGMYRKCEATLRECMAAGQTPDLVSYNTVIFSYSKHGQLSSATRIFHEMVSNGIQPDSFTYNTFVGC----------------------YVNGGMFPEALSVVKHMHKTGCKPDEV-TYRTLVDAYCKIGKFEE--    
  57 UniRef50_B9R8H7        Pentatricopeptide repeat-co...   449    1e-124  1  12.0%   6:590   155:708     -----MYCKAGLLFEAREVFDRMPERNEVTWATMISGYAIQRLAGEAFEVFELMR--REEEDVNEFAFTSVLSALAVPEFVDSGKQIHCLAVKTGLLVFLSILNALVTMYAKCGSLDDSLQVFEMsdKNSITWSAMITGYAQSGdkALKLFSRMHFAGINPSEFTLVGVLNACSdcAVEEGKQVHNYLLKLGFESQLYIMTALVDMYAKSGVTEDARKGFNYLQQ----PDLVLWTSMIAGYVQNGENEDALSLYCRMQMEGILPNELTMASVLKACSNLAAFDQGRQI------------HARTIKYGLGLEVTIGSALSTMYAKCGNLEEGNIVFRRMPER------DIISWNAMISGLSQNGYGKEALELFEEMRQQDTKPDDVTFVNVLSACSHMGLVDSGWLYFRMmdEFGLLpeHYACMVDVLSRAGKLYEAKEFIEStiDHGLCL-----WRILLGACRNYRNYELGAYAGEKLMELGSQ-ESSAYVLLSGIYTALGRPEDVERVRSMMKVRGVSKEP----------GCSWIELKSNVHVFVVGDQMHPCIGEIRTEILRLSKQMKDEGYQPASVtsYDFMM-------------    
  58 UniRef50_UPI00019849C5 PREDICTED: hypothetical pro...   449    1e-124  1  10.6%   5:582   170:718     ----SMIskFGKIGEAQRVFDGMPNKDVVCWNSIIGGYVQAGCFDVAFQLF--FEMHGSGIKPSPITMTSLIQACGGIGNLKLGKCMHGYVLGLGLGNDILVLTSFVDMYSKMGDIESARWVFYKMprNLVSWNAMISGCVRNGleSFDLFHRLVRSSGGFDLTTIVSLLQGCSqaSLATGKILHGCAIRS-FESNLILSTAIVDLYSKCGSLKQATFVFNRMKDR----NVITWTAMLVGLAQNGHAEDALRLFAQMQEEGIAANSVTFVSLVHSCAHLGSLKRGRSI------------HGHLFRLGFAFDIVNMTALVDMYAKCGKINLAERIF-----SHGSISKDVVLWNSMITGYGMHGHGYQAVGIYHKMIEEGLKPNQTTFLSLLSACSHSRLVEQGISLFNSMErhNIRPihYACLVDLLSRAGRFEEAQALIEKMP----FQPGTAVLEALLSGCRTHKNINLGIQTSDKLLALD-AMNPGIYIMLSNIYAEARRWDKVDYIRGLMRNRGLKKTPGY----------SLVETGNWVHTFFAGDNSHPNWEEIYHFLESLRSAVETSGYVPDT---------------------    
  59 UniRef50_D1IB08        Whole genome shotgun sequen...   449    1e-124  1  11.2%   1:582   197:774     MALVDMYVKCREIEDARFLFDKMQERDLVTWTVMIGGYAECGKANESLVLFEKMREEGVV--PDKVAMVTVVFACAKLGAMHKARIIDDYIQRKKFQLDVILGTAMIDMYAKCGCVESAREIFDRmeKNVISWSAMiyGYHGQGRKALDLFRMMLSSGMLPDKITLASLLYACINCrlTQVRQVHAQASVHGMLQNLIVANKLVHFYSYYRALDDAYGLFDGM----CVRDSVSWSVMVGGFAkcGNANESLVLFDKMREEGVVPDKVAMVTVVFACAKLGAMHKARTIDDYIQRKK------------FQLDVILGTAMIDMHAKCGCVESAREIFDRMEEK------NVISWSAMIAAYGYHGQGRKALDLFPMMLRSGILPNKITLVSLLYACSHAGLVEEGLRFFSLmeDYSVRAdhYTCVVDLLGRAGRLDEALKLIESM----TVEKDEGLWGAFLGACRTHKDVVLAEKAATSLLE--LQpnPGHYILLSNIYANAGRWEDVAKTRDLMSQRRLKKIPGWT----------WIEVDNKSHQFSVGDTTHPRSKEIYEMLKSLGNKLELVGYVPDT---------------------    
  60 UniRef50_Q8W0G9        Os01g0852900 protein n=3 Ta...   449    1e-124  1   9.7%   1:597    90:699     LSAFSRAIMPDAALDAFRALPsgCNPGIRSHNALLDAFVRARRFSDADAFFASLsaFGRRIAPNLQTYNIVLRSLCARGDLDRAVTLFDSLRRRQVAPDRITYSTLMCGLAKQDRLDHALDLLDEMPrdVVCYNALLGGcaGEFEKVMRVWDKLVKdgARPNLATYNVMLDGLCkgRFKEVGEVWERMVANNLQPDVITYGILIHGLCRSGDVDGAARVYSEIIKTGLVIDAAMYNSLVKGFCQAGRVQEAWKFWDSAGFAGLR-NLRTYNIMIKGLFDSGMVDEAIELWDLLEKDVA-----------CIPDTVTFGTLIHGLCQNGFANKAFTIFEEARVSGKQL--DVFSYSSMINGLCNVGRLVDAVKVYEKMDKDGCKPNSHIYNALISGFCQVYRTSDAVRIYSKMADNGCSPttYNTLIDGLCKAEKYQEASSVAREMVENG-FTPDITTYGSLIRGLFSDKKIDDALSIWKQILYKGLKVDVMMHNILIHGLCSAGKVDEALHVFSDMKEKKNCPPNLVTYNTLMDGLYETGYIDKAATLwyNTRIKGLCSCDRIHEGIQLLDEVLSRGIIP-TVITWNILVRAVIKYG------    
  61 UniRef50_D1H2Y1        Whole genome shotgun sequen...   448    1e-124  1   9.6%   6:589   150:703     -----MYSMCGSPSDSRMVFDKLRRKNLFQWNAIVSAYTRNELFEDAMSIFSELISV-TEHKPDNFTLPCVIKACAGLLDLGLGQIIHGMATKMDLVSDVFVGNALIAMYGKCGLVEEakRVFDLMdkTVSSWNALLCGYAQnpRKALDLYLQMTDSGLDPDWFTIGSLLLACSRMklHYGEEIHGFALRNGLAVDPFIGISLLSLYICCGKPFAAQVLFDGMEHR----SLVSWNVMIAGYSQNGLPDEAINLFRQMLSDGIQPYEIAIMCVCGACSQLSALRLGKEL------------HCFALKAHLTEDIFVSSSIIDMYAKGGCIGLSQRIFDRLREK------DVASWNVIIAGYGIHGRGKEALELFEKMLRLGLKPDDFTFTGILMACSHAGLVEDGLEYFNQMLnhNIEpeHYTCVVDMLGRAGRIDDALRLIEEMPGD----PDSRIWSSLLSSCRIHGNLGLGEKVANKLLE--LEPepENYVLISNLFAGSGKWDDVRRVRGRMKDIGLQK----------DAGCSWIEVGGKVHNFLIGDEMLPELEEVRETWRRLEVKISSIGYTPDTGSVLHDL--------------    
  62 UniRef50_D1I9T3        Whole genome shotgun sequen...   448    1e-124  1  10.7%  68:580    13:497     -------------------------------------------------------------------YDSVLTECISQTAIREGQRVHAHMIKTCYEPPVYLRTRLIVLYNKCRCLGDARRVLDEmeRNVVSWTAMISGYSQRgsEALHLFVEMLMSGTAPNEFTFATVLTScsSSGFQLGRQIHSLVIKTSFESHIFVGSSLLDMYAKAGKICEARRVFDGLPER----DVVSCTAIISGYAQLGLDEEALDLFRRLQREGMRSNYVTYASVLTALSGLAALDHGRQV------------HSHVLRAKLPFYVVLQNSLIDMYSKCGSLTYSRRIFDSMPER------TVISWNAMLVGYSKHGLGREAVELFKLMKEEnvKPDSVTFLAVLSGCSHGGMEDRGLEIFYEMvkDGFEPehYGCVVDLFGRAGRVEEAFEFIKKMP----FEPTAAIWGSLLGACRVHQNVHIGEFVARRLLEIESE-NAGNYVILSNLYASAGRWDDVRTVRELMKEKAVIKEPGR----------SWIELDQTLHTFHASDRSHPRKEEVFAKVRELSIKIKEAGYVP-----------------------    
  63 UniRef50_A5B3U0        Putative uncharacterized pr...   448    1e-124  1  10.3%   7:587   253:805     ------LDMYMKCNNLsqLFNSLPNHNLQSYNAIIVGYARSDKGIEALGMFRLLQKSGLG--LDEVSLSGAXRACAVIKGDLEGLQVHGlmKSLCQSNICVANAILDMYGKCGALVEACLVFEEMVSRDAVSWNAIIAAHEQNgeKTLSLFVWMLQSGMEPDEFTYGSVLKACAgqALNCGMEIHNRIIKSRMGLDSFVGIALIDMYSKCGMMEKAEKLHDRLAEQ----TVVSWNAIISGFSLQKQSEEAQKTFSKMLEMGVDPDNFTYATILDTCANLVTVELGKQIHAQIIKKE------------LQSDAYISSTLVDMYSKCGNMQDFQLIFEKAPNR------DFVTWNAMVCGYAQHGLGEEALKIFEYMQLENVKPNHATFLAVLRACGHMGLVEKGLHYFHSMLsyGLDpeHYSCVVDIMGRSGQVSKALELIEGMP----FEADAVIWRTLLSXCKIHGNVEVAEKAAYSILQ--LEpdSAAYVLLSNIYANAGMWNEVTKLRKMMRFNGLKKEP----------GCSWIEIKSEVHAFLVGDKAHPRSKEIYENLDVLTDEMKWVGYMPDTDFILN----------------    
  64 UniRef50_Q2QNV4        Pentatricopeptide, putative...   448    1e-124  1  10.5%   6:594   187:761     -----LYLRCGSFRLAERVFCDMPHRDTVTFNTLISGHAQCGHGEHALEIFEEMQF--SGLSPDCVTISSLLAACASLGDLQKGTQLHSYLFKAGISSDYIMEGSLLDLYVKCGDVETALVIFNSSDrnVVLWNLMLVAfiNDLAKSFELFCQMQAAGIRPNQFTYPCILRTccTREIDLGEQIHSLSVKTGFESDMYVSGVLIDMYSKYGWLEKARRVLEMLKEK----DVVSWTSMIAGYVQHECCKDALAAFKEMQKCGIWPDNIGLASAISGCAGINAMRQGLQI------------HARIYVSGYSGDVSIWNALVNLYARCGRIREAFSSFEEIEHK------DEITWNGLVSGFAQSGLHEEALKVFMRMDQSGVKHNVFTFVSALSASANLAEIKQGKQIHARVIKTgfETEVGNALISLYGKCGSFEDAKMEFSEMSER-----NEVSWNTIITSCSQHGRGLEALDLFDQMKKEGIKPNDVTFIGVLAACSHVGLVEEGLSYFKSMSDEYGIRPRPDHYACVIDIFGRAGQLDRAKKF----IEEMPIAADAMVWRTLLSACKVHKNiePHDSASYVLLSNAYA---------    
  65 UniRef50_B9T3T5        Pentatricopeptide repeat-co...   447    1e-124  1  10.6%   7:580    65:600     ------FSLCHQMNLAVNVFNQIQDPNVHLYNTLIRAHVQNSQSLKAFATF--FDMQKNGLFADNFTYPFLLKACNGKGWLPTVQMIHCHVEKYGFFGDLFvnSLIDSYSKCGLlvNYAMKLFMEmeKDLVSWNSMIGGLVKAGdrARKLFDEMAER----DAVSWNTILDGYVKAgmSQAFNLFEKMPER----NVVSWSTMVSGYCKTGDMEMARMLFDKMPFK----NLVTWTIIISGFAEKGLAKEATTLYNQMEAAGLKPDDGTLISILAACAESGLLVLGKKV------------HASIKKIRIKCSVNVSNALVDMYAKCGRVDKALSIFNEMSMR------DLVSWNCMLQGLAMHGHGEKAIQLFSKMQQEGFKPDKVTLIAILCACTHAGFVDQGLSYFNSMErhGIVpeHYGCMIDLLGRGGRLEEAFRLVQSMP----MEPNDVIWGTLLGACRVHNAVPLAEKVLDRLITLE-QSDPGNYSMLSNIFAAAGDWNSVANMRLQMKSTGVQKP----------SGASSIELDDEVHEFTVFDKSHPETDKIYQILVKLGQDLKQVAYAP-----------------------    
  66 UniRef50_Q654Y7        Os06g0690900 protein n=4 Ta...   447    1e-124  1  11.6%   1:602   226:844     LGYCRTGELRKACWLFLMMPLMGCQRNEYSYTILIQGLCDAKCVRKALVLFLMMKRDGCSPNVRAFTF--LISGLCKSGRVGDARLLFDAMPQNGVVPSVMTYNAMIVGYSKLGRMNDALKIKELMEKNGctYNTLIYGLcqKTEEAEELLNNAVKEGFTPTVVTFTNLINGYckDRLKEAKELLNEISANGLVPNVITYTSIIDGYCKSGKVDIALEVLKMMERDGCQPNAWTYNSLMYGLVKDKKLHKAMALLTKMQKDGIIPNVITYTTLLQGQCDEHDFDNAFRLFEMMEQ------------NGLKPDEHAYAVLTDALCKAGRAEEAYSF---IVRKGVALTK--VYYTTLIDGFSKAGNTDFAATLIERMIDEGCTPDSYTYSVLLHALCKQKRLNEALPILDQMSLRGIkiFAYTILIDEMLREGKHDHAKRMYNEMTSSGHK-PSATTYTVFINSYCKEGRLEDAEDLILKMEREGVAPDVVTYNILIDGCGHMGYIDRAFSTLKRMVGASCEPNYWTYCLLLKHL------LKGNLAYVRSVDTSGMWNLIELDITWQLLERMVKHGLNP-TVTTYSSLIAGFCKAGRLEEA-    
  67 UniRef50_D1I0N6        Whole genome shotgun sequen...   447    1e-124  1  10.1%   5:528   178:692     ----SMVEQCLSVFD--KMIKSRLSPDVKNCNRILRILRDKDLMSKAVEVYRTMGEFG--IKPTIVTYNTLLDSYCKGGKVQQGLDLLSEMQRRGCAPNDVTYNVLINGLSKKGEFEQAKGLIGEMLKTgyTYNPLIYGYFNKgaEALSLQEEMVLKGASPTVATYNSFIYGLCkgRMSDAMQQLSDMLANNLLPDVVSYNTLIYGYCRLGNLMKAFLLFDELRSIYLFPTIVTYNTLLDGLCRQGELEVAQQLKVEMINEGIAPDIVTYTILVNGSCKMGSLSMAQEFFDEMLHE------------GLELDSYAYATRIVGELKLGDTSRAFSLQEEMLAKG--FPPDLIIYNVVVDGLCKLGNLEEASELLQKMVSDGVIPDYVTYTSIIHAHLENGRLRKGREIFYEMLSKGLTPstYTVLIHGHAGKGRLERAFIYFSEMQEKGIL-PNVITYNSLINGLCKVRRMDQAYNFFAEMVEKGIFPNKYSYTILINENCNMGNWQEALSLYKQMLDRGVQPDSCTH---------------------------------------------------------------------------    
  68 UniRef50_Q9FMF6        Pentatricopeptide repeat-co...   447    1e-124  1  12.7%   1:597   121:683     LGAnkTIDRL-----LIQMKDEGIVFKESLFISIMRDYDKAGFPGQTT---RLMLEMRnsCEPTFKSYNVVLEILVSGNCHKVAANVFYDMLSRKIPPTLFTFGVVMKAFCAVNEIDSALSLLRDMTKHGCVPNSVIysKCNRVNEALQLLEEMFLMGCVPDAETFNDVILGLCkdRINEAAKMVNRMLIRGFAPDDITYGYLMNGLCKIGRVDAAKDLFYRIPK----PEIVIFNTLIHGFVTHGRLDDAKAVLSDMvsYGIVPDVCTYNSLIYGYWKEGLVGLALEVLHDMRNK------------GCKPNVYSYTILVDGFCKLGKIDEAYNVLNEMSADG--LKPNTVGFNCLISAFCKEHRIPEAVEIFREMPRKGCKPDVYTFNSLISGLCEVDEIKHALWLLRDMISEGVVAntYNTLINAFLRRGEIKEARKLVNEMVFQG-SPLDEITYNSLIKGLCRAGEVDKARSLFEKMLRDGHAPSNISCNILINGLCRSGMVEEAVEFQKEMVLRGSTPDIVT-FNSLINGLCRAGRI---------------------EDGLTMFRKLQAEGIPPDTV-TFNTLMSWLCKGG------    
  69 UniRef50_Q6EPV3        Os02g0582300 protein n=4 Ta...   447    1e-124  1  13.8%   7:602   127:710     ------LSRFALAPLALRLFAHmpAPTAPTYNAVIRSLCRRADLARALRYLSLMV--RSGWRPDAYTFNSLIVGYCRTNQVDVARDLFDKMPLRGFAQDVVSYATLIEGLCEAGRIDEAVELFGEmqPDMHMYAALVKGLCnrGEEGLLMLRRMKELGWRPSTRAYAAVVdrCRERKAKEAEEMLQEMFEKGLAPCVVTCTAVINAYCKEGRMSDALRVLELMKLRGCKPNVWTYNALVQGFCNEGKVHKAMTLLNKMRACGVNPDAVTYNLLIRGQCIDGHIESAFRLLRLMEGD------------GLIADQYTYNALINALCKDGRTDQACSLFDSLETRG--IKPNAVTFNSLINGLCKSGKADIAWKFLEKMVSAGCTPDTYTYSSFIEHLCKMKGSQEGLSFIGEMLQKDVKPStyTIVIHKLLKERNYGLVARTWGEMVSSGC-NPDVVTYTTSMRAYCIEGRLNEAENVLMEMSKNGVTVDTMAYNTLMDGHASIGQTDHAVSILKQM----------TSVASVPNQFTYFILLRHLVRMRLVEDVLPLTPAGVWKAIEllFDVMKKNEFLPNS-GTYSSILEGFSEDGRTEEA-    
  70 UniRef50_C5YEK8        Putative uncharacterized pr...   446    1e-123  1   9.2%   2:579   206:786     -CRNSMLVGYVkvDHALELFDSMPERDVVSWNMMVSALSQSGRVREALDMVVDMQ--SKGVRLDSTTYTSSLTACARLSSLRWGKQLHAQVIRNLPCIDPYVASALVELYAKCGCFKEAKGVFNSLhrNNVAWTVLIAGFLQHGceSVELFNQMRAELMTLDQFALATLISGCcrMDLCLGRQLHSLCLKSGQIQAVVVSNSLISMYAKCDNLQSAESIFRFMNEK----DIVSWTSMITAHSQVGNIAKAREFFDGMSTK----NVITWNAMLGAYIQHGAEEDGLRMYNVMLskLGDQIIGRTVKVGLILDTSVANAVITMYSKCGRILEARKVFDFLNVK------DIVSWNAMITGYSQHGMGKQAIEIFDDILKRGAKPDYISYVAVLSGCSHSGLVQEGKSYFDMMKrhNISpeHFSCMVDLLGRAGHLTEAKDLIDEMPMK----PTAEVWGALLSACKIHGNNELAELAAKHVFELD-SPDSGSYMLMAKIYADAGKSDDSAQIRKLMRDKGIKKNPGY----------SWMEVNNKVHVFKADDVSHPQVIAIRKKLDELMEKIARLGYV------------------------    
  71 UniRef50_C5YYT3        Putative uncharacterized pr...   446    1e-123  1   9.6%   5:587   144:796     ----SLVDMYMkvVDGRKVFEAMPKRNVVTWTSLLTGYIQDGALSDVMELFFRMRAEGVW--PNSVTFASVLSVVASQGMVDLGRRVHAQSVKFGCCSTVFVCNSLMNMYAKCGLVEEARVVFCGMerDMVSWNTLMAGLVLNGheALQLFHDSRSSITMLTQSTYATVIKLCANIklGLARQLHSSVLKRGFHSYGNVMTALMDAYSKAGQLGNALDIFLLMSGSQ---NVVSWTAMINGCIQNGDVPLAAALFSRMREDGVAPNDFTYSTILTaySKLCNTEEALSIFKMIDQkgATNIFIKMTMHGLKPNEFTISSVIdmYARKGSIESAQCIFERQTDR------DLVSWNSMLSGYAQHGYSQKALDVFRQMEAEGIEMDGVTFLSVIMGCAHAGLVEEGQRYFDSmrDYGITpeHYACMVDLYSRAGKLDEAMSLIEGMS----FPAGPMVWRTLLGACKVHKNVELGKLAAEKLLS--LEpdSATYVLLSNIYSAAGKWKEKDEVRKLMDTKKVKK----------EAGCSWIQIKNKVHSFIASDKSHPLSEQIYAKLRAMTTKLKQEGYCPDTSFALH----------------    
  72 UniRef50_Q940A6        Pentatricopeptide repeat-co...   446    1e-123  1  10.9%   8:602   253:817     -------NEFQKCCEAFDVVCKGVSPDVYLFTTAINAFCKGGKVEEAVKLFSKMEEAG--VAPNVVTFNTVIDGLGMCGRYDEAFMFKEKMVERGMEPTLITYSILVKGLTRAKRIGDAYFVLKEMTKKGFPPNVIVYNNLIdlNKAIEIKDLMVSKGLSLTSSTYNTLIKGYCKNgaDNAERLLKEMLSIGFNVNQGSFTSVICLLCSHLMFDSALRFVGEMLLRNMSPGGGLLTTLISGLCKHGKHSKALELWFQFLNKGFVVDTRTSNALLHGLCEAGKLDEAFRIQKEILGR------------GCVMDRVSYNTLISGCCGKKKLDEAFMFLDEMVKRG--LKPDNYTYSILICGLFNMNKVEEAIQFWDDCKRNGMLPDVYTYSVMIDGCCKAERTEEGQEFFDEMMSKNVQPNtyNHLIRAYCRSGRLSMALELREDMKHKGI-SPNSATYTSLIKGMSIISRVEEAKLLFEEMRMEGLEPNVFHYTALIDGYGKLGQMVKVECLLREMHSKNVHPNKIT----------------------YTVMIGGYARDGNVTEASRLLNEMREKGIVPDSI-TYKEFIYGYLKQGGVLEA-    
  73 UniRef50_B9GER4        Predicted protein n=3 Tax=r...   446    1e-123  1  11.3%   1:600   203:837     LAKIRRVDMV--LVLFGEILSMGIRPDIYIYVAVIRSFCELKNFAKAKEMIQRME----SSDLNVVVYNVLIHGLCKNKRVWEAVEIKNGLIQKGLTASEVTYCTLVLGLCKVQEFEVGAGVMDEMIELGFVPtsLVEGLRRKGkdAFDLVNRVKKVGAMPSLFVYNALINSLCkgKFDEAELLFKEMGEKGLCANDVTYSILIDSFCRRGKLDTAIHFLGKMIMAGIKITVYPYNSLINGHCKLGNLSAAVSFFDEMIDKGLKPTVVSYTSLISGYCNKGKLHEAFRLYHEMTGK------------GIAPNTYTFTTLISALFRANRMTDAFRLFDEMLEQ--NMMPNEVTYNVMIEGHCKEGNTVKAFELLNQMVQKGLVPDTYTYRPLISSLCSTGRVCEAKKFIDDLHREHFKLncYSALLHGYCKEGRLRDALGVCREMVKRGvlRPDKVIYTSMIDGYSKAGSVKKAFGIWDIMIDEGCTPNIVTYTTLINELCKAGLMDKAELLWKEMLVSNSTPNHVT-YCCFLDHLAREGSMEKAVQLHNDMLkrGFCKLGRVEEATKLLDEMIDNAIFPDCI-TYSTIIYQCCRRGNLD---    
  74 UniRef50_Q9LFL5        Pentatricopeptide repeat-co...   445    1e-123  1   9.5%   2:582   167:763     -ALVAMYSRCRSLSDARKVFDEMSVWDVVSWNSIIESYAKLGKPKVALEMFSRMTN-EFGCRPDNITLVNVLPPCASLGTHSLGKQLHCFAVTSEMIQNMFVGNCLVDMYAKCGMMDEANTVFSNmvKDVVSWNAMVAGyiGRFEDAVRLFEKMQEEKIKMDVVTWSAAISGYAQRGLgeALGVCRQMLSSGIKPNEVTlnQLIDMYAKCKKVDTARAMFDSLSPK--ERDVVTWTVMIGGYSQHGDANKALELLSEMFEEDcrPNAFTISCALVACASLAALRIGKQIHAYALRNQQNAVP-----------LFVSNCLIDMYAKCGSISDARLVFDNMMAK------NEVTWTSLMTGYGMHGYGEEALGIFDEMRRIGFKLDGVTLLVVLYACSHSGMIDQGMEYFNRMKtfGVSpeHYACLVDLLGRAGRLNAALRLIEEMP----MEPPPVVWVAFLSCCRIHGKVELGEYAAEKITELASNHD-GSYTLLSNLYANAGRWKDVTRIRSLMRHKGVKKRP----------GCSWVEGIKGTTTFFVGDKTHPHAKEIYQVLLDHMQRIKDIGYVPET---------------------    
  75 UniRef50_Q0WVK7        Pentatricopeptide repeat-co...   445    1e-123  1  12.6%   7:521   188:697     ------FGLLREARRVFemLNYGLVLSVDSCNVYLTRLSkcYKTATAIIVFREFPE--VGVCWNVASYNIVIHFVCQLGRIKEAHHLLLLMELKGYTPDVISYSTVVNGYCRFGELDKVWKLIEVMKRKGLKPNSYIYGSIikLAEAEEAFSEMIRQGILPDTVVYTTLIDGFCkgDIRAASKFFYEMHSRDITPDVLTYTAIISGFCQIGDMVEAGKLFHEMFCKGLEPDSVTFTELINGYCKAGHMKDAFRVHNHMIQAGCSPNVVTYTTLIDGLCKEGDLDSANELLHEM------------WKIGLQPNIFTYNSIVNGLCKSGNIEEAVKLVGEFEAAG--LNADTVTYTTLMDAYCKSGEMDKAQEILKEMLGKGLQPTIVTFNVLMNGFCLHGMLEDGEKLLNWMLAKGIAPntFNSLVKQYCIRNNLKAATAIYKDMCSRG-VGPDGKTYENLVKGHCKARNMKEAWFLFQEMKGKGFSVSVSTYSVLIKGFLKRKKFLEAREVFDQMRREGL----------------------------------------------------------------------------------    
  76 UniRef50_UPI00019837DE PREDICTED: hypothetical pro...   445    1e-123  1  12.3%  39:601    66:565     --------------------------------------CKSgkRSEAFSVFNHLIDMQ--PTPPISSFNTLLGAVAKIKRYFDVISLYKRMSLIGLAPDFITLNILINCYCN----------------------------LNKVDFGLAVLGEMLRRGHSPNTVTFTSLVKGLCLgrISEATGLLRKMVRMGYRPNVVTYGTLLNGLCKDGLIDKGKELFLEMKGRGISPDVVAYSSIIHGMCHTGRWEGAKGLFNEMVDEGVHPNVVTFNVLIDALCKAGKMEEANHLLKLMIQR------------GESPDTFTYNTLIDGFCLEGRIDDARDLFVSMESKG--IETDAVSYNVLINGYCKSGRMVEAKKLYREMMCKEIMPTVITYNTLLTGLFREGKVRDAWNLFGEMKVHDLTPetYNILLDGLCKNNHLSEAMELFHYL-ENHDFQPSIQIFNCLIDGLCKARKIEIARELFNRLSHEGLEPNVITYTVMIHGLCKSGQLENAKDLFLGMEEKG----------CAPNLVTFNTLMRGFCQ------------NDEMQKVVELLQEMAEKDFSPDA-STISIVVDLLSKDEKYRE--    
  77 UniRef50_C7JA97        Os12g0577900 protein (Fragm...   444    1e-123  1  10.2%  42:580   112:606     -----------------------------------------GAPRDALAAFSAM------PRRDSFTYSFLIKALSAAG-VAPVRAVHSHVVKLGSVEDTFVGNALIDAYSKNGGFSDARKVFDEmeRDVVSWNTAMAAMVREGELarKLFDEMPEK----DTVSWNTILDGYTKAgvEAAFELFQRMPER----NVVSWSTMVSGYCKKGDLEMARVIFDKMPSK----NLVTWTIMVSACAQKGLVDEAGKLFAQMKEASVELDVAAVVSILAACAESGSLSLGKRIHRYVRKRNLGR------------STHVCNALIDMFCKCGCVNRADYVFDT-----ETVQKDSVSWNSIIGGFAMHGHGDKALELFAQMKQQGFNPDAVTMINVLSACTHMGFVDEGRRHFSNMEryGIMpeHYGCMIDLLGRGGLIKEAVDLIKSMP----WEPNEVIWGSLLSACRLHKNVEYAEIAVNEL--SKLQpnAGNYAVLSNIYAEAGQWSDMAKARMQMKGTGSQKT----------AGSSWVELNETFHEFTVGDRKHQDSDQISEMVDRLSSHVKHVGCVP-----------------------    
  78 UniRef50_D1IRU7        Whole genome shotgun sequen...   444    1e-123  1  11.6%   5:581    36:613     ----KLID-LHSIDYARFVLDQTPSPTDFSWNSLIRAYTVHGSPQNSLFLYLKM--LRSSTKPSNFTFPFVLKACSTLGSVLEGEQIHTHVLRLGFGSDLFVCNSLIDMYCKCFRLDSARNFWDDMgrDEVSWNSIISGYvgQVEKARDLFEEMPMRR---NVVCWTAMINGYGkgDFVEMLSLFRQMlvNKIPLNTILVTALIDMYSKCGDVEKAWRIFDGVSCKNLP----SWNAIITGCVQGGLLEEAIDLYRHMKAQSVKPNEITLVNVLSACAGLGALELGREV------------HLYLGRNGLDLNVILATALVDMYAKCGKIDDACLIFVKTSEK------DVALWNAMILGLAYHGDGRDSLAVFSQMVRAGVQPNDVTFIGVLSACNHSGLVEEGRVQFSSmdKHGLSpeHYACMVDLLGRAGHLKEAYELVQNM----LIPPDSIIWGALLSACRIHRNLELADKISETIMASQ-DPNIGFCILLSNIYASSGRWKDVARVRRQVKEKRIKKP----------SGCSWVEVDGVVHRFVVEDTTHLKSGEIYGAYEILVNHLKAEGYVAN----------------------    
  79 UniRef50_C5Y166        Putative uncharacterized pr...   444    1e-123  1   9.3%   2:600   145:765     -GYCRAGGRLADAERMLASLALSGSADVVTYNTLVAGYCREGRLNDARRLVADM-----PFAPNSYTNSTLLKGLCSNKEWDDAEELLSEMIRSGCPPNDLTFGMIIHSLCQNGLADRAMGVLDQMsrGVIVYNEIISCLagRVEEALHLFDQMP---CKPDIFSYNTVMKGLCrgRWEDAGTLIAGMVRKDCPPDEVTFNTVISYLCHRGLVDCAMEVVEQMPKYGCKPDNFTYSALVNALSERGCVDDALELLRTIP---WKPNTVCYRSVLKGLCRADRWDDVGKLVAEMIrdYGAEVLREMTKFGCSPDIIIYNSLINGFSENGSVDDALELFKNM-----SCKRNVVTYNYMLKGLCRAEQWEDAGKLVAEMVKDECLPNEVTFSTLISYLCQKGFVECAIEVFEKMPKYNCMPNvySTLINGLSDQECVDDALKLLNDMP----CKPDTICYSAALKGLCRAERWEDAGELILEMIRKNCLPDEVTFSILINNLCHKGFLESATEVSELMLK----------YECMPNVFIYSSLINGFAEQYRAEDakGLCRAKRWEDARELIAEMFRKQCPLDE-ATFSMLIGSLCQNGLVD---    
  80 UniRef50_Q6YS36        Os07g0300200 protein n=8 Ta...   443    1e-123  1  12.9%   6:601   180:821     -----LVDTYKKSGRVQdmRDRGMAPSIRCCNALLKDLLRADAMALLWKVREFMV--GAGISPDVYTYSTLIEAYCKVREFDTAKKVLVEMRERGCGLNTVTYNVLIAGLCRSGAVEEAFGFKKDMEDYGLVPDGFTYGALINgnEAKALLDEMSCAELKPNVVVYANLIDGFMrgNADEAFKMIKEMVAAGVQPNKITYDNLVRGLCKMGQMDRASLLLKQMVRDSHRPDTITYNLIIEGHFRHHSKKDAFRLLSEMENAGISPNVYTYSIMIHGLCQSGEPEKASDLLEEMTTK------------GLKPNAFVYAPLISGYCREGNVSLACEIFDKMTK--VNVLPDLYCYNSLIFGLSKVGRVEESTKYFAQMQERGLLPNEFTYSGLIHGYLKNGDLESAEQLVQRMLDTGLKPniYGILIHNLSSSGNMEAAFRVLSG-IEKNGSVPDVHVYSSLISGLCKTADREKAFGILDEMSKKGVDPNIVCYNALIDGLCKSGDISYARNVFNSILAKGLVPNCVT-YTSLIDGSCKVGDISNAFYLYNEMLatGCSSAGDLEQAMFLIEEMFLRGHA--SISSFNNLVDGFCKRGKMQE--    
  81 UniRef50_B9I898        Predicted protein n=2 Tax=P...   443    1e-122  1  12.3%   2:528    39:564     -GFCSNYNNLHSVADAvqLLGIRPLPPVVVFNKLLGSLVKKKHYSTVISLCKQMDL--SNIRPNVYTLTILINCLCHSNrvHFAFSALGKMFKLGlpTHVTFGTLLNGLCSKAKIIDAV-KLFDEIGKMGFAPSLITYTTIIKGLcaLQLLKKMEEKGCKPDVVAYNTVIDSLCkrRANEAMYFFSEMVDQGIPPNVVTYSSILHGFCNLGQLNEATSLFKQMIGRNVMPNTVTFTILVDGLCKEGMILEARRVFEMMTENGVEPDAYTYSALMDGYCLQSQMDEAQKLFDIMVGK------------GFAPSVRVYNILINGHCKSRRLNEAKTLLSEMYDR--DLTPDTVTYSTLMQGFCQAGRPQVAQKLFKEMCSYGLLPDSITYSILLDGLCKHGHLDEAFRLLKAMQESKIEPHiyNILIQGMCNFGKLEAARELFSNLFVKGI-QPSVVTYTVMISGLLKEGLSNEACEMFRKMVVNGCLPNSCTYNVAIQGFLRNGDPSNAVRLIEEMVGRGFSADSSTF---------------------------------------------------------------------------    
  82 UniRef50_B9HB63        Predicted protein n=1 Tax=P...   443    1e-122  1  10.8%   1:602  379:1054     LCKAGRINEAFDL--LDTMRKQGVLPNLHTYNTLISGLLRANRLDDALDLFSNMESLGVEpaPNIVACNASLYSLAEMGRLGEAKAMFNELKSSGLAPDSVTYNMMMKCYSKVGQVDEAIKLLSEmePDVIVINSLIDTlaGRVEEAWQMFCRMEEMNLAPTVVTYNILLAGLGkgQIQKAVQLFESMNGHGCSPNTITFNTLLDCLCKNDEVDLALKMFYKMTTMNCRPDVLTFNTIIHGFIKQNQIKNAIWLFHQMKKL-LRPDHVTLCTLLPGVIKSGQIEDAFriLTEAGTEKAILFGERLVCRAICKDDSVLIPIIKVLCKHKKTSVARNVFVKFT-KELGVKPTLKVYNLLIDGFLEVHNVEVAWNLFEEMKSAGCAPDTFTYNSLIDAHGKSGKINELFDLYDEMLTRGCKPNtyNMVISNLVKSNRLDKAMDLYYNLV-SGDFSPTPCTFGPLIDGLLKSGRLDDAHEMFDGMVHYGCRPNSAIYNILVNGYGKLGHVDTACEFFKRMVKEGIRPDLKS-YTILVDILCIAGRVDDALHYfyNLMINGLGRSQRTEEALSLFHEMQNRGIVPD-LYTYNSLILNLGIVGMIEEA-    
  83 UniRef50_Q9LIQ7        Pentatricopeptide repeat-co...   443    1e-122  1  10.9%  55:585    50:549     ------------------------------------------------------DLEGSYIPADRRFYNTLLKKCTVFKLLIQGRIVHAHILQSIFRHDIVMGNTLLNMYAKCGSLEEARKVFEKmqRDFVTWTTLISGYSQhpCDALLFFNQMLRFGYSPNEFTLSSVIKAAAAERRGcgHQLHGFCVKCGFDSNVHVGSALLDLYTRYGLMDDAQLVFDALESR----NDVSWNALIAGHARRSGTEKALELFQGMLRDGFRPSHFSYASLFGACSSTGFLEQGKWVHAYMIKSGEKLV------------AFAGNTLLDMYAKSGSIHDARKIFDRLAKR------DVVSWNSLLTAYAQHGFGKEAVWWFEEMRRVGIRPNEISFLSVLTACSHSGLLDEGWHYYELMKKDGIVPehYVTVVDLLGRAGDLNRALRFIEEMP----IEPTAAIWKALLNACRMHKNTELGAYAAEHVFELDPD-DPGPHVILYNIYASGGRWNDAARVRKKMKESGVKKEPA----------CSWVEIENAIHMFVANDERHPQREEIARKWEEVLAKIKELGYVPDTSHV------------------    
  84 UniRef50_Q7F2F8        P0402A09.8 protein n=5 Tax=...   442    1e-122  1  10.1%   7:589   180:730     ------YCKHGLLAAARRVFLEMHDKDAVTYNAMMMGCSKEGLHTQALQLFAAMR--RAGIPATHFTFSSILTVAAGMAHLLLGHQVHALVLRSTSVLNVFVNNSLLDFYSKCDCLDDMRRLFDEmeRDNVSYNVIIAAYAWNQCAalRLFREMQKLGFDRQVLPYATMLSVAGSldVHIGKQIHAQLVLLGLASEDLLGNALIDMYSKCGMLDAAKSNF----SNRSEKSAISWTALITGYVQNGQHEEALQLFSDMRRAGLRPDRATFSSIIKASSSLAMIGLGRQL------------HSYLIRSGYKSSVFSGSVLVDMYAKCGCLDEALRTFDEMPER------NSISWNAVISAYAHYGEAKNAIKMFEGMLHCGFNPDSVTFLSVLAACSHNGLADECMKYFHLMkqYSISpeHYACVIDTLGRVGCFSQVQKMLVEMP----FKADPIIWTSILHSCRIHGNQELARVAADKLF--GMEPTDAtyVILSNIYARAGQWEDAACVKKIMRDRGVRK----------ESGYSWVEIKQKIYSFASNDLTSPMIDEIKDELDRLYKEMDKQGYKPDITCALHMV--------------    
  85 UniRef50_Q9LFF1        Pentatricopeptide repeat-co...   442    1e-122  1  10.3%   1:602    93:665     LGRSGSFDDMKKI--LEDMKSSRCEMGTSTFLILIESYAQFELQDEILSVVDWMID-EFGLKPDTHFYNRMLNLLVDGNSLKLVEISHAKMSVWGIKPDVSTFNVLIKALCRAHQLRPAILMLEDMPSYGLVPDEKTFTTvgDLDGALRIREQMVEFGCSWSNVSVNVIVHGFCkgRVEDALNFIQEMsqDGFFPDQYTFNTLVNGLCKAGHVKHAIEIMDVMLQEGYDPDVYTYNSVISGLCKLGEVKEAVEVLDQMITRDCSPNTVTYNTLISTLCKENQVEEATELARVLTSK------------GILPDVCTFNSLIQGLCLTRNHRVAMELFEEMRSKGC--EPDEFTYNMLIDSLCSKGKLDEALNMLKQMELSGCARSVITYNTLIDGFCKANKTREAEEIFDEMEVHGVSRntYNTLIDGLCKSRRVEDAAQLMDQMIMEG-QKPDKYTYNSLLTHFCRGGDIKKAADIVQAMTSNGCEPDIVTYGTLISGLCKAGRVEVASKLLRSIQMKGINLTP----------------------HAYNPVIQGLFRKRKTTEAINLFREMLEQNEAPPDAVSYRIVFRGLCNGGGPIRE-    
  86 UniRef50_Q0DEH2        Os06g0152500 protein n=2 Ta...   442    1e-122  1  12.0%  68:602    86:607     -------------------------------------------------------------------FDALIRSYASLPRPSLAAAALAFAASAGYAPSVPAYNAVLLALSDASLPS-ARRFLSSMLRHGVAPNVYTYNIlgRLEEAVGVVGDMRGAGCAPNAVTYNTLVAAFCRAglDGAERVVSLMREEgaKPNLVTFNSMVNGLCKAGRMEGARKVFDEMVREGLAPDVVSYNTLLSGYCKVGCLHESLAVFSEMTQRGLVPDVVTFTSLIHATCKAGNLEQAVALVAQMRER------------GLRMNEVTFTALIDGFCKKGFLDDALLAVEEM--RKCGIQPSVVCYNALINGYCKLGRMDLARELIREMEAKRVKPDVVTYSTIISGYCKVGNLDSAFQLNQKMLKKGVLPdtYSSLIRGLCEEKRLNDACELFENMLQLG-VQPDEFTYTTLIDGHCKEGNVEKALSLHDEMIRKGVLPDVVTYSVLINGLSKSARTKEAHRLLFKLYHEDPVPDNIKYDALMLC--CSKAEFKS-----VVALLKGFCMKGLMKEADKVYQSMLDRNWKLD-GSVYSILIHGHCRGGNVRKA-    
  87 UniRef50_D1H2J7        Whole genome shotgun sequen...   442    1e-122  1  11.0%   1:598    80:651     IGAMRNYGRKGKIQEAVDVFERmcEPSVQSYNAIMNILVEYRYFDQAHKVY--MRMRDKGIVPDVYTFTIRMKSFCRTSRPHAARRLLNNMPSQGCESSAVAYCTVIGGFYEENHRVEAHELFEEMLGLGICPDImiHTLCRKGheSERLLNKVLKRGVSPNLFTVNIFIQGFcrAMLNEAIRLLDG-VGRGLTPDVITYNTLICGLCKNFKVVEAEHYLRKMVNEGYEPDGFTYNSIIDGYCKLGMMQNADQILRDGAFKGFVPDESTYCSLINGLCQDGDIDRAINVFNEAMEK------------GLKPNLVLCNTLVKGLSQQGLILQALKLMNEMSENGCS--PDIWTYNLVINGLCKIGCVSDADNLVIDAIAKGHLPDVFTFNTLIDGYCKKLKLDNAIEIVDRMWNHGVSPdtYNSILNGLCKAGKYEDVMGTFKLMMEKGCV-PNIITYNILTESFCKARKVEEALNLIEEMQNKGLTPDVVNFGTLMKGFCDNGDLDGAYQLFKRVDEQYKFSHTIATYNIMINAF--AGKLN-------------------MNMAEKLFNKMCENGFSPDS-YTYRVMIDGFCKTGN-----    
  88 UniRef50_B9GMS3        Predicted protein n=7 Tax=M...   442    1e-122  1   9.9%   4:576    52:592     ---LKMYCKCGSLADARKVFDEMRERNLVSWNTIISAYAENGVFDKGFCMFSNMLELET--KPNGSTYIGFLRSLLNPSGLEIGKQIHSHAIRSGLGSNASVNTAISNMYVKCGWLEGAELVFEKmeKNAVAWTGIMVGYTQAErdALALFAKMVNEGVELDEYVFSIVLKACAGleLNFGRQIHGHIVKLGLESEVSVGTPLVDFYVKCSNLESATKAFEWISE----PNDVSWSALITGYCQMGEFEEALKTFESLRTRSVDINSFTYTSIFQACSALADFNSGAQAHADAIKSSLVAYQHGE------------SAMITMYSRCGRLDYATRVFESIDD------PDAVAWTAIIAGYAYQGNAPEALKLFRRMQDCGVRPNAVTFIAVLTACSHSGLVIEGRQYLESMSsyGvtIDHYDCMVDIYSRAGFLQEALELIRSMP----FSPDAMSWKCLLGGCWTYRNLEIGELAAENLFQLDPE-DTAGYILMFNLYASFGKWKEAANVRKMMAERNLRK----------ELSCSWITVKGKVHRFIVGDKHHPQTEEIYSKLEALNDSVIKE---------------------------    
  89 UniRef50_Q9LS72        Pentatricopeptide repeat-co...   441    1e-122  1  10.6%   7:571    61:589     ------LSLCRQTNLAVRVFNQVQEPNVHLCNSLIRAHAQNSQPYQAFFVFSEMQRFGLF--ADNFTYPFLLKACSGQSWLPVVKMMHNHIEKLGLSSDIYvnALIDCYSRCGGlvRDAMKLFEKmeRDTVSWNSMLGGLVKAGedARRLFDEMPQR----DLISWNTMLDGYARCrmSKAFELFEKMPER----NTVSWSTMVMGYSKAGDMEMARVMFDKMP--LPAKNVVTWTIIIAGYAEKGLLKEADRLVDQMVASGLKFDAAAVISILAACTESGLLSLGMRI------------HSILKRSNLGSNAYVLNALLDMYAKCGNLKKAFDVFNDIPKK------DLVSWNTMLHGLGVHGHGKEAIELFSRMRREGIRPDKVTFIAVLCSCNHAGLIDEGIDYFYSMEkyDLVpeHYGCLVDLLGRVGRLKEAIKVVQTMP----MEPNVVIWGALLGACRMHNEVDIAKEVLDNLVKLDPC-DPGNYSLLSNIYAAAEDWEGVADIRSKMKSMGVEKP----------SGASSVELEDGIHEFTVFDKSHPKSDQIYQMLGSLIE--------------------------------    
  90 UniRef50_UPI00019842FA PREDICTED: hypothetical pro...   441    1e-122  1  10.7%   3:579   325:872     --ANSmyCKCGRVEDAQALFELMDARDVISWNSLVSGYAQLGNIREVLQLLIRMKTDG--IEPDQQTFGSLVSAAAMQSKLGVGKMVHGHILRAGLEQDSHIETSLIGMYLKCGNVNSAFRIFemMHKDVISWTAMISGLVQnaDMAVTVFRRMLKSRVMPSTATIASVLAACAegSFPLGTSVHGYILRQRIKLDIPSQNSLVTMYAKCGHLEQSCSVFDRMSRR----DIVSWNAIVSGHAQNGHLCKALLLFNEMRKARQRPDSITVVSLLQACASIGALHQGKWI------------HNFVTKSCLGPCILIDTALVDMYSKCGDLGSAQKCFDRMPQQ------DLVSWSSIIAGYGSHGKGETALRMYSDFLHTGIQPNHVIYLSILSACSHNGLVDQGLSFFHSmkDFGIEpeHRACIVDLLSRAGRVEEAYSFYKRM----FPKPSMDVLGILLDACRTTGNVELGDIVAREIVI--LKpnAGNYVQLAHSYASMKRWDGVGEVWTQMKSLHLKKLP----------GWSFIELHGTITTFFTDHSSHPQFEEIMLVLKILGSEMRKVGIH------------------------    
  91 UniRef50_B9RP84        Pentatricopeptide repeat-co...   441    1e-122  1  11.4%  15:602   266:822     --------------FFLEAKSRGVKLDAAAYSIVIQAFCKNLDVELACGLLKDMRDKGWV--PSEGTFTSVIGACVKQGNMVEALRLKDEMVSCGVQMNVVVATTLVKGYCKQDKLVSALEFFDKMNENGPSPNRVTYAVLIEWckAYDLYTQMKNKNICPTVFIVNSLIRGFLKvsREEASKLFDEAVACDIA-NIFTYNSLLSWLCKEGKMSEATTLWQKMLDKGLAPTKVSYNSMILGHCRQGNLDMAASVFSDMLDCGLKPNVITYSILMDGYFKNGDTEYAFYVFDRMVDEN------------IVPSDFTYNIKINGLCKVGRTSEAQDMLKKFVEKG--FVPVCLTYNSIMDGFIKEGSVSSALTAYREMCESGVSPNVITYTTLINGFCKNNNTDLALKMRNEMRNKGLELDiyGALIDGFCKKQDIETASWLFSELLD-GGLSPNSVIYNSLISGYRNLNNMEAALNLQKRMLGEGISCDLQTYTTLIDGLLKEGRLVLALDLYSEMSAKGIIP----------DIIIYTVLINGLCG------------KGQLENAQKILAEMERDSITPN-VPIYNALIAGHFKAGNLQEA-    
  92 UniRef50_P93005        Pentatricopeptide repeat-co...   441    1e-122  1  10.8%   5:579   158:705     ----SLVGMYCKAGLVekVFAYMPERNTYTWSTMVSGYATRGRVEEAIKVFNLFLREKEEGSDSDYVFTAVLSSLAATIYVGLGRQIHCITIKNGLLGFVALSNALVTMYSKCESLNEACKMFDSsdRNSITWSAMVTGYSQNGeeAVKLFSRMFSAGIKPSEYTIVGVLNACSdcYLEEGKQLHSFLLKLGFERHLFATTALVDMYAKAGCLADARKGFDCLQER----DVALWTSLISGYVQNSDNEEALILYRRMKTAGIIPNDPTMASVLKACSSLATLELGKQV------------HGHTIKHGFGLEVPIGSALSTMYSKCGSLEDGNLVFRRTPNK------DVVSWNAMISGLSHNGQGDEALELFEEMLAEGMEPDDVTFVNIISACSHKGFVERGWFYFNMMSDqgLDpdHYACMVDLLSRAGQLKEAKEFIESaiDHGLC-----LWRILLSACKNHGKCELGVYAGEKLMALGSR-ESSTYVQLSGIYTALGRMRDVERVWKHMRANGVSK----------EVGCSWIELKNQYHVFVVGDTMHPMIEETKDLVCLVSRQMIEEGFV------------------------    
  93 UniRef50_D1IWK7        Whole genome shotgun sequen...   441    1e-122  1  11.9%   4:574   175:718     ---NSLLSMYAdmERAEELFDEMCERDVISWSVMIGGYVQTG---EAKMALQLFLEmnASIELDGITMVSVLKACANTGDISMGRSVHGVVICRGLDYDLFVGNSIIDMYSKCDDHESAFKAFNEMprNTVSWNSIISGLVrkHSEALSLFYSMGKAGFRADEVTLVNLLQSCkfVDPFQCKFIHSIVIRWGYELNEFVINSLIDAYSKCDLIELAWKLFDRLKTK----DTVSWSAMIAGFNHCGKPDEAIALFQEMNQAQEKPNGVTILSLLEAFSVSADL------------KRSKWAHGIAIRRGLAAEVAVGTAILDMYAKCGEIGLSRKAFDQIPEK------NIVSWGAMIAACGMNGLARDALALLSEMKLHGLKPNVVTTLSVLSACSHGGLVEEGLSFFENMvdHGVEpeHYSCMVDMLSRAGKLNSAMNLIEKMPER--MRDGAGLWGALLSACRSSGNSRlaFRVLE------LEpsSAGYFLASSMYAASGLWADAARMRWLVKARGVRVVAGYSLVH----------VEDKAWRFVAGDESHPRAGEIWGVVEQLHDCMK-----------------------------    
  94 UniRef50_C5Z4C1        Putative uncharacterized pr...   441    1e-122  1  13.1%  68:603    90:647     -------------------------------------------------------------------FDSLIKSYASlsRASLAAAALAFARSAGYAPSVLAYNAVLLALSDASLPS-ARRFFDSMLSDGVAPNVYTyaLCGRGheALSVLRDMRGAGCDPNAVTYNTLVAAFCRAgvDRAERLVDMMREGGLKPNLVTFNSVVNGICKAGRMEDARKVFDEMVKEGLAPDGVSYNTLVGGYCKVGCSHEALSVFAEMTRKGIMPDVVTFTSLIHVMCKAGNLERAVGLVREMRER------------GLQMNEITFTALIDGFCKKGFLDDALLAVREM--RQCRIQPSVVCYNALINGYCMVGRMDEARELVREMEAKGVKPDVVTYSTILSAYCKNGDTHSAFQLNQQMLENGVLPdtYSSLIRVLCEEKRLGDAHVLFKNMISLGL-QPDEVTYTSLIDGHCKEGNVERALSLHDEMVKAGVLPDVVTYSVLINGLSKSARTKEAQRLLFKllMRCCRNAELKSVLALLKGFCMKGLMNEADKVyySVLIHGHCREGNVMKALSFHKQMLQCGFAPNSTST-ISLIRGLFEKGMVVEAD    
  95 UniRef50_Q6ZC60        Os08g0110200 protein n=4 Ta...   440    1e-122  1  12.7%  15:596    50:604     --------------HAHHLFDELLRRDTTSIFDLNSALSAVARESPAV-ALSLFNRM---PRADLCTYSIVIGCCSRAGHLDLAFAALGRVIRTGWTAQAITFslLKGLCHDKRTSDAMDIALRRMPALGCTPNAFSYNILLKgqQALHLLHTMmrGGCPPDVVSYTTVINGLLrgQLDKAYCLFDEMLDQGMSPNCITYNCLLHGYCSSGKPKEAIGIFRKMCRDGVEPDVVTYNTLMVYLCKNGRSMEARKVFDSMVKKGHKPDSSIYGTLLHGYATEGYLVQMHQLLDVMVR------------NGTQPDHYIFNILIGAYAKHGMVDEAMLAFSKMRQQG--LHPNIVTYGTVMDALCRVGKVDDAMSQFDRLISEGLTPNGVVFRTLIHGLCACDKWDKAEELAVEMIGRGICPNtfNTLLNHLCKEGMVTRAKNIFDLMV-RVDVQCDVITYTTLIDGYCLDGKVDEATKLLEGMVLDGVKPNEVTYNTIINGYCKNGRIEDACSLFRQMASKGVNP----------GIVIYSTILHGLFQT------------RRIAAAKELYLWMIKCGIKL-PIGTYNIILQGLCQN-------    
  96 UniRef50_Q9LQ20        F16P17.1 protein n=2 Tax=Br...   440    1e-122  1  12.0%  19:518     1:504     ------------------MVETGCPANVVTFTTLMNGLCREGRVLQALALVDRMVEEGHQ--PDAVTYGTIVNGMCKLGDTVSALNMLRKMDESQIKANVVIYSAIVDRLCKDGNHIKAQNIFTEMHEKGIFPNVLTYNCMIDgsDAEQLLRDMIERNIDPDVVTFSALINAFVkgKVSGAEELYREMLRRNIFPTTITYSSMIDGFCKHSRLEDAKHMFDLMVSKGCSPDIITLNTLIDGCCRagNVNVAQDLFQEMISNGVSPDIVTCNTLLAGLCENGKLEKALEMFKVFQKSK------------MDLDTATCNIIINGMCKGNKVDEAWDLFNSLPVNG--VETDVVTYNILIGVFVKEGNFLRAEDIYLEMLCKGIIPSTVTYNSMVDGFCKQNRLEEARQMVDSMVSEGCSPdtFSTLIKGYCKAGRVDDGLELFSEMCQRGLVA-DTITYNALIHGFCKVGDLNGAQDIFEEMVSSGVCPDTITFRSMLAGLCTKAELQKGLTMLEDLQK-------------------------------------------------------------------------------------    
  97 UniRef50_Q8LQQ6        Os01g0783100 protein n=5 Ta...   440    1e-122  1  13.0%  32:602    47:581     -------------------------------NARLRRLIARDDLAEAARLVD--RATSRGEAPDVYLCTKLIRNLCRRGRTSDAARVLRAAERSGTAVDVFAYNTLVAGYCRYGQLDAARRLIASMppDAYTYTPIIRGLCDRGreALSLLDDMLHRGCQPSVVTYTVLLEAVCKsgFGQAMEVLDEMRAKGCTPNIVTYNVIINGMCREGRVDDAREFLNRLSSYGFQPDTVSYTTVLKGLCAAKRWEDVEELFAEMMEKNCMPNEVTFDMLVRFFCRGGMVERAIQVLEQM------------SGHGCAANTTLCNIVINTICKQGRVDDAFQFLNNMGSYGCS--PDTISYTTVLKGLCRAERWEDAKELLKEMVRKNCPPNEVTFNTFICILCQKGLIEQATMLIEQMSEHGCEVntYNALVNGFCVQGRVDSALELFYSMP----CKPNTITYTTLLTGLCNAERLDAAAELLAEMLQKDCAPNVVTFNVLVSFFCQKGLMDEAIELVEQMMEHG----------CTPNLITYNTLLDGITKDCNS------------EEALELLHGLVSNGVSPDIV-TYSSIIGVLSREDRVEEA-    
  98 UniRef50_B9GRG5        Predicted protein n=1 Tax=P...   440    1e-122  1   8.5%   7:582    89:710     ------YSAAGDLKLSRKIFSDTPlrDSVFYNAMITAYSHNHDGHAAIELFCDMQ--RDNFRPDNYTFTSVLGALALVAEKehCQQLHCAVVKSGTGFVTSVLNALISSYVKCalMAEARKLFDEmkKLGVAWNAMISGYAHRGleAFEMFRKMIMSKIQLDEFTFTSVISVCANAgfRLGKEMHAYFLkaNPAPDVavNNALITFYWKCGKVDIAQEIFNKMPER----DLVSWNIILSGYVNVRCMDEAKSFFNEMPEK----NILSWIIMISGLAQIGFAEEALKFFNRMKlkHGRQLHAQVVRYGYESSLSAGNALITMYARCGVVDAAHCLFINMP------CVDAISWNAMIAALGQHGQGTQAIELFEEMLKEGILPDRISFLTVISACSHAGLVKEGRKYFDSMHnyGVNPdhYARIIDLLCRAGKFSEAKEVMESMP----FEPGAPIWEALLAGCRIHGNIDLGIEAAERLFELKPQHD-GTYVLLSNMYAVAGQWNDMAKVRKLMRDRGVKKEP----------GCSWIEVENKVHSFLVGDANHPEVRQIYNYLEQLVLEMRKIGYVPDT---------------------    
  99 UniRef50_UPI0001982A13 PREDICTED: hypothetical pro...   440    1e-122  1  10.3%  11:556   328:900     ----------WDVFDLLWATRNVCVPGFGVFDALFSALIELGMLEEASECFLKMRKFRVFPKPR--SCNALLHRLSKVGRGDLSRKFFKDMGAAGIKRSVFTYNIMIDYLCKEGDLEMARSLFTQMKEAGFTPDIVTYNSLIDgdECICIFEQMKDADCDPDVITYNALINCFCkeRMPKAFEFLHEMKANGLKPNVVTYSTFIDAFCKEGMLQEAIKFFVDMRRVALTPNEFTYTSLIDANCKAGNLAEALKLVEEILQAGIKLNVVTYTALLDGLCEEGRMKEAEEVFRAMLN------------AGVAPNQETYTALVHGFIKAKEMEYAKDILKEMKEkeSGINTNAVIYTTLMDAYFKSGQATEALTLLEEMLDLGLIATEVTYCALIDGLCKSGLVQEAMHHFGRMSEIGLQPNvyTALVDGLCKNNCFEVAKKLFDEMLDKGMM-PDKIAYTALIDGNMKHGNLQEALNLRDRMIEIGMELDLHAYTALIWGLSHSGQVQKARNLLDEMIGKGV-LPDEVVYMCLIKKYYALGKVDEALELQNEMAKRG-----------------------------------------------    
 100 UniRef50_Q7Y211        Pentatricopeptide repeat-co...   440    1e-122  1   9.9%   5:589   244:810     ----AMYGKLGKLASSKVLLGSFGGRDLVTWNTVLSSLCQNEQLLEALEYLREMVLEG--VEPDEFTISSVLPACSHLEMLRTGKELHAYALKNglDENSFVGSALVDMYCNCKQVLSGRRVFDGmdRKIGLWNAMIAGYSQNEheALLLFIGMEEsgLLANSTTMAGVVPACVRSgfSRKEAIHGFVVKRGLDRDRFVQNTLMDMYSRLGKIDIAMRIFGKMEDR----DLVTWNTMITGYVFSEHHEDALLLLHKMqrVSLKPNSITLMTILPSCAALSALAKGKEI------------HAYAIKNNLATDVAVGSALVDMYAKCGCLQMSRKVFDQIPQK------NVITWNVIIMAYGMHGNGQEAIDLLRMMMVQGVKPNEVTFISVFAACSHSGMVDEGLRIFYVMkdYGVEpdHYACVVDLLGRAGRIKEAYQLMNMMPRDFNK---AGAWSSLLGASRIHNNLEIGEIAAQNLIQ--LEPNvsHYVLLANIYSSAGLWDKATEVRRNMKEQGVRKEP----------GCSWIEHGDEVHKFVAGDSSHPQSEKLSGYLETLWERMRKEGYVPDTSCVLHNV--------------    
 101 UniRef50_D1J3D5        Whole genome shotgun sequen...   440    1e-122  1  10.5%  10:580    72:613     ---------LKDFNNASLLFSQIPYPNEYAFNIMIRGLTTtqKFNLTIEFYYQMKDFG--IRPNNFTYPFVFIACANLLVLNHGQCAHSGVLKSGLCADGHVRHSLITMYSRCGELGCARRVFDEieKDLVSWNSMISGYSRMggDAVGLFGEMRDAGFEPDEMTLVSILGACGdgDLGLGSWIEGFVVENEMDLNSFVGSALIGMYGKCGDLSSARRVFDRMVKK----DVVTWNAMITGYAQNGVSDEAIILFSGMRESGVNPDKITLVGVLSACASIGALDFGKWL------------DTYASERGLQNDIYVSTALIDMYAKCGSLDDALRVFEDMPQK------NEVSWNAMISALAFHGRPQESLSLFKRMSKEgvRPNDISFIGVLSACVHAGLVDEGRQLFDLMSSsgLVpeHHSCMVDLLARAGHVHEAWDFIEKMPEK----PDEVVLGALLGACQKRRNVDVSERVMHMLLE--MEpnSGNYIISSKIFANMKRWDDSARMRVLMRQRGVTKTP----------GCSWIEIENQVHEFHAGDVLHFISQDMCQVINLLNEEMKVEGYGP-----------------------    
 102 UniRef50_Q7FA49        OSJNBa0013K16.3 protein n=3...   439    1e-121  1   9.3%   2:579   206:786     -CRNSMLAGYAKldHAIEYFEDMAERDVVSWNMMIAALSQSGRVREALGLVVEMH--RKGVRLDSTTYTSSLTACARLFSLGWGKQLHAKVIRSLPQIDPYVASALIELYAKCGSFKEAKRVFNSLqrNSVSWTVLIGGSLqcFSKSVELFNQMRAELMAIDQFALATLISGcnRMDLCLGRQLHSLCLKSGHNRAIVVSNSLISLYAKCGDLQNAEFVFSSMSER----DIVSWTSMITAYSQIGNIIKAREFFDGMATR----NAITWNAMLGAYIQHGAEEDGLKMYSAMLskLGDQIIGHTVKAGLILNVSVANAAITMYSKCGRISEAQKLFDLLNGK------DVVSWNAMITGYSQHGMGKQAAKTFDDMLSKGAKPDYISYVAVLSGCSHSGLVQEGKLYFDMMTrhGISpeHFSCMVDLLGRAGHLTEAKDLIDKMPMK----PTAEVWGALLSACKIHGNDELAELAAKHVFELD-SPDSGSYMLLAKIYSDAGKSDDSAQVRKLMRDKGIKKNPGY----------SWMEVENKVHVFKADDVSHPQVIAIRNKMDELMEKIAHLGYV------------------------    
 103 UniRef50_Q76C99        Protein Rf1, mitochondrial ...   439    1e-121  1  11.9%  14:602    32:670     -------------EDARHVFDELLRRgaSIYG-LNRALADVAR-DSPAAAVSRYNRMARAGapDLCTYGILIGCCCRAGRLDLGFAALGNVIKKGFRVDAIAFtlLKGLCADKRTSDAMDIVLRRmiPNVFSYNILLKGLcnRSQEALELLHMmrGGGSPPDVVSYTTVINGfkEGDSDKAYSTYHEMLDRGILPDVVTYNSIIAALCKAQAMDKAMEVLNTMVKNGVMPDCMTYNSILHGYCSSGQPKEAIGFLKKMRSDGVEPDVVTYSLLMDYLCKNGRCMEARKIFDSMTKRGLKplLDLMVRNGIHPDHYVFSILICAYAKQGKVDQAMLVFSKMRQQG--LNPNAVTYGAVIGILCKSGRVEDAMLYFEQMIDEGLSPGNIVYNSLIHGLCTCNKWERAEELILEMLDRGICLNtfNSIIDSHCKEGRVIESEKLFELMVRIG-VKPNVITYNTLINGYCLAGKMDEAMKLLSGMVSVGLKPNTVTYSTLINGYCKISRMEDALVLFKEMESSGVSPDIITYNIILQGLfaAAKELYVRITetYNIILHGLCKNKLTDDALQMFQNLCLMDLKLEA-RTFNIMIDALLKVGRNDEA-    
 104 UniRef50_Q9FIT7        Pentatricopeptide repeat-co...   438    1e-121  1  10.3%   7:601   256:858     ------LNVDGALKLKESMICKGLVPLKYTYDVLIDGLCKIKRLEDAKSLLVEMDSLGVS--LDNHTYSLLIDGLLKGRNADAAKGLVHEMVSHGINIKPYMYDCCICVMSKEGVMEKAKALFDGMIASGlaYASLIEGYCrnVRQGYELLVEMKKRNIVISPYTYGTVVkmCSSGDLDGAYNIVKEMIASGCRPNVVIYTTLIKTFLQNSRFGDAMRVLKEMKEQGIAPDIFCYNSLIIGLSKAKRMDEARSFLVEMVENGLKPNAFTYGAFISGYIEASEFASADKYVKEMREC------------GVLPNKVLCTGLINEYCKKGKVIEACSAYRSMVDQG--ILGDAKTYTVLMNGLFKNDKVDDAEEIFREMRGKGIAPDVFSYGVLINGFSKLGNMQKASSIFDEMVEEGLTPNvyNMLLGGFCRSGEIEKAKELLDEMSVKGL-HPNAVTYCTIIDGYCKSGDLAEAFRLFDEMKLKGLVPDSFVYTTLVDGCCRLNDVERAITIFGT-NKKGCASSTAP-FNALINWVFKFGkmDGSFDRfyNIMIDYLCKEGNLEAAKELFHQMQNANLMP-TVITYTSLLNGYDKMGRRAE--    
 105 UniRef50_D1HW74        Whole genome shotgun sequen...   438    1e-121  1  11.5%   5:576   207:752     ----SLLNMYAKLGSIEdvFNMMTEHNQVSWNAMISGCTSNGLH---LEAFDLFVRMKNgcTPNMYTLVSVSKAVGKLVDVNMGKEVQNCASELGIEGNVLVGTALIDMYSKCGSLHDARSVFdnFINCGvpWNAMISGYSQSGceALELYVQMCQNGITSDLYTYCSVFNAIAASklQFGRVVHGMVLKCGLDLMVVSVNNAidAYSKCGFLEDVRKVFDRMEER----DIVSWTTLVTAYSQSSLGEEALATFCLMREEGFAPNQFTFSSVLISCASLCFLEYGRQV------------HGLLCKAGLDTEKCIESALIDMYAKCGSITEAGKVFDKISN------PDIVSWTAIISGYAQHGLVEDALQLFRRMELSGIKANAVTLLCVLFACSHGGMVEEGLFYFQQMEdyGVVpeHYACIIDLLGRVGRLDDAMEFIRKMP----MEPNEMVWQTLLGGCRVHGNVELGEIAARKILS--IRPesATYVLLSNTYIETGSYEDGLSLRNVMKDQGVK-------KEPGYSWIS---VKGRVHKFYSGDQQHPQKKEIYVKLEELREKIKAM---------------------------    
 106 UniRef50_Q9SZ52        Pentatricopeptide repeat-co...   438    1e-121  1  11.7%  16:598   176:749     ---------------LRKMREFGFVLNAYSYNGLIHLLLKSRFCTEAMEVYRRMILEGF--RPSLQTYSSLMVGLGKRRDIDSVMGLLKEMETLGLKPNVYTFTICIRVLGRAGKINEAYEILKRMdpDVVTYTVLIDALcrKLDCAKEVFEKMKTGRHKPDRVTYITLLDRFSDNrlDSVKQFWSEMEKDGHVPDVVTFTILVDALCKAGNFGEAFDTLDVMRDQGILPNLHTYNTLICGLLRVHRLDDALELFGNMESLGVKPTAYTYIVFIDYYGKSGDSVSALETFEKMKTK------------GIAPNIVACNASLYSLAKAGRDREAKQIFYGL--KDIGLVPDSVTYNMMMKCYSKVGEIDEAIKLLSEMMENGCEPDVIVVNSLINTLYKADRVDEAWKMFMRMKEMKLKpvTYNTLLAGLGKNGKIQEAIELFEGMVQKGC-PPNTITFNTLFDCLCKNDEVTLALKMLFKMMDMGCVPDVFTYNTIIFGLVKNGQVKEAMCFFHQMKK--LVYPDFVTLCTLLPGVVKASLIEDAYKIITNFLYNCADQPANLFWEDLIGSILAEAGIDNAVSFSERLVANGICRDGD-----    
 107 UniRef50_Q2R497        Os11g0482400 protein n=7 Ta...   438    1e-121  1  11.9%   3:580   116:676     --ATSLVnrCSAARDARRLFDGMPERNVVTWTALVTGYTLNSQPALGLEVFVEMLEMGRY--PSHYTLGATLNACLASCDVDLGKQVHGYAIKYGAEstSMGNSLCSLYAKLGSLDSALRAFWRIPEKNVITWTTMISACAEdvELGMSLFIDMLMDGVMPNEFTLTSVMSLCgrLDLNLGKQVQAFSFKIGCETNLPVKNSTMYLYLRKGETDEAMRLFEQMEDA----SIITWNAMISGYAQIMDSAkaLTIFRDLKRSVMKPDLFTFSSILSVCSAMMALEQGEQI------------HAQTIKSGFLSDVVVNSALVNMYNKCGCIQDANKAFLEMPTRTF------VTWTSMISGYSQHGQPQEAIQLFEEMRLAGVRPNEITFVSLLSACSYAGLVEEAEHYFDMMKkyCIEpdHYGCMIDMFVRLGRVEDAFSFIK----RTGFEPNEAIWSSLVAGCRSHGNMELAFYAADKLLELKPK-GIETYILLLNMYISTERWQDVARVRKLMKQEDVGILRDR----------SWITIKDKVYFFRANDRTHPQATELYQLLENLLEKAKAIGYEP-----------------------    
 108 UniRef50_Q5G1T1        Pentatricopeptide repeat-co...   438    1e-121  1  10.2%   4:582   206:759     ---CSLIDMFVkfENAYKVFDKMSELNVVTWTLMITRCMQMGFPREAIRFFLDMVL--SGFESDKFTLSSVFSACAELENLSLGKQLHSWAIRSGLVDDV-ECSLVDMYAKCsgSVDDCRKVFDRMEDHSvsWTALITGYMKNCneAINLFSEMITQgvEPNHFTFSSAFKACGNLSDPrgKQVLGQAFKRGLASNSSVANSVISMFVKSDRMEDAQRAFESLSEK----NLVSYNTFLDGTCRNLNFEQAFKLLSEITERELGVSAFTFASLLSGVANVGSIRKGEQI------------HSQVVKLGLSCNQPVCNALISMYSKCGSIDTASRVFNFMENR------NVISWTSMITGFAKHGFAIRVLETFNQMIEEGVKPNEVTYVAILSACSHVGLVSEGWRHFNSmeDHKIKpeHYACMVDLLCRAGLLTDAFEFINTMP----FQADVLVWRTFLGACRVHSNTELGKLAARKILE--LDPNeaAYIQLSNIYACAGKWEESTEMRRKMKERNLVK----------EGGCSWIEVGDKIHKFYVGDTAHPNAHQIYDELDRLITEIKRCGYVPDT---------------------    
 109 UniRef50_Q67UW7        Os02g0151000 protein n=5 Ta...   438    1e-121  1   9.7%   7:582   269:835     ------YSRDTSVLdaIKFFESMIERNEYTWSTMIAALSHGGRIDAAIAVYE------RDPVKSIACRTALITGLAQCGRIDDARILFEQIPE----PIVVSWNALITGYMQNGMVNEAKELFDKMprNTISWAGMIAGYAQNgeEALGLLQELHRSGMLPSLSSLTSIFFACSnvALETGTQVHSLAVKVGCQFNSFACNALITMYGKCRNMEYARQVFSRMVTK----DIVSWNSFLAALVQNDLLDEARNTFDNMLSR----DDVSWTTIISAYAHAEQSNEAMGAFKTMFCEhgQQIHTVAIKLGMDSELIVANALISMYFKCGCADS-RRIFDLMEER------DIFTWNTIITGYAQHGLGREAIKMYQHMESAGVLPNEVTFVGLLNACSHAGLVDEGWKFFKSmqDYGLTpeHYACMVDLLGRTGDVQGAEQFIYDMP----IEPDTVIWSALLGACKIHKNAEIGKRAAEKLFR--IEpnAGNYVMLSNIYSSLGMWGEVAEVRKIMKQQGVIKEP----------GCSWTQIKDKMHSFVTGDKQHEQIEEIVATLEELYTLLKATGYVPDT---------------------    
 110 UniRef50_A5ASX4        Putative uncharacterized pr...   438    1e-121  1  12.7%  25:582   101:629     ------------------------EINVFAFNAIISGFITNGFPEEGFEFYQKMRNEGVM--PDKFTFPCAIKAC---LDVLEIKKIHGLLFKFGLELDVFIGSALVNCYLKFGLMEHAQVAFEELprDVVLWNAMVNGYagQFEMVLETFRRMNDESVVPSRFTVTGXLSVFamGDLNNGRIIHGFAMKMGYDSGVAVSNSLIDMYGKCKCIEDALEIFEMMREK----DIFSWNSIVSVHEQCGDHDGTLRLLDRMLGAGIQPDLVTVTTVLPACSHLAALMHGREIHGYMIVSGLGKDGKDID------DVLLKNAVIDMYAKCGSMRDAHLVFERMSNK------DVASWNIMIMGYGMHGYGNEALEMFSRMCEVQLKPDEVTFVGVLSACSHAGFVSQGRNFLVQMkkYDVApeHYTCVIDMLGRAGQLDEAYELALTMP----IEANPVVWRALLAACRLHKHAVLAEVAAQRVFE--LEPEhgSYVLMSNVYGAVGRYEEVLEVRHTMRQQNVRKTP----------GCSWIELKNGVHVFVSADRAHPEAXSIYAGLNSLTARLXEHGYVPDV---------------------    
 111 UniRef50_Q9LFC5        Pentatricopeptide repeat-co...   438    1e-121  1   9.8%   4:526   204:728     ---NALIGSLVRIGWVeeISRSGVGINVYTLNIMVNALCKDGKMEKVGTFLSQVQE--KGVYPDIVTYNTLISAYSSKGLMEEAFELMNAMPGKGFSPGVYTYNTVINGLCKHGKYERAKEVFAEMLRSGLSPDSTTYRSlgDVVETEKVFSDMRSRDVVPDLVCFSSMMSLFTrgNLDKALMYFNSVKEAGLIPDNVIYTILIQGYCRKGMISVAMNLRNEMLQQGCAMDVVTYNTILHGLCKRKMLGEADKLFNEMTERALFPDSYTLTILIDGHCKLGNLQNAMELFQKMKEKR------------IRLDVVTYNTLLDGFGKVGDIDTAKEIWADMVSKEILPTP--ISYSILVNALCSKGHLAEAFRVWDEMISKNIKPTVMICNSMIKGYCRSGNASDGESFLEKMISEGFVPdsYNTLIYGFVREENMSKAFGLVKKMEEegGLVPDVFTYNSILHGFCRQNQMKEAEVVLRKMIERGVNPDRSTYTCMINGFVSQDNLTEAFRIHDEMLQRGFSPDDK-----------------------------------------------------------------------------    
 112 UniRef50_B9P5D9        Predicted protein n=9 Tax=P...   437    1e-121  1  10.3%  12:528   105:615     -----------ALAYFNHMLHRKPRPCIIQFNKLLSAIVKMRHYHDA--VISLSKQMELAgsPDTYTLHMLINCFFQLQRVDLGFSVLAKIIKLGLQLTIVTFNTLINGLCKVGKFGQAVELFDDMVARGYQPDVHTYTTIINGlaAAGLFRKMGEAGCQPDVVTYSTIIDSLCkrRVNEALDIFSYMKAKGISPNIFTYNSLIQGLCNFSRWREASAMLNEMMSLNIMPNIVTFSLLINIFCKEGNVFEARGVLKTMTEMGVEPNVVTYSSLMNGYSLQAEVVEARKLFDVMITK------------GCKPDVFSYNILINGYCKAKRIGEAKQLFNEMIHQG--LTPDIVSYNTLIDGLCQLGRLREAHDLFKNMLTNGNLPDLCTYSILLDGFCKQGYLAKAFRLFRAMQSTYLKPnmYNILIDAMCKSRNLKEARKLFSELFVQGL-QPNVQIYTTIINGLCKEGLLDEALEAFRNMEEDGCPPNEFSYNVIIRGFLQHKDESRAVQLIGEMREKGFVADVATT---------------------------------------------------------------------------    
 113 UniRef50_D1ICI0        Whole genome shotgun sequen...   437    1e-121  1   9.8%   7:580   192:735     ------YSENGCIHDARYLFDRMPSKDGVLWNVMLNGYVKNGDWDNATGVF--MEMRRTETNPNSVTFACVLSVCASEIMINFGSQLHGLVVSSGLEMDSPVAnaLIDIYFKCRDVEMARKIFDQRTpdIVVCTAMISGYVLNgnNALEIFRWLLQERMRANSVTLASVLPACalAALTLGKELHGHILKNGHGGSCYVGSAIMDMYAKCGRLDLAHQTFIGISDK----DAVCWNSMITSCSQNGKPEEAIDLFRQMGMAGTKYDCVSISAALSACANLPALHYGKEI------------HAFMMRGAFRSDLFAESALIDMYSKCGNLDLACRVFDTMEEK------NEVSWNSIIAAYGNHGRLKDSLNLFHGMLGDGIQPDHVTFLAIISACGHAGQVDEGIHYFRCmeELGirMEHYACMVDLFGRAGRLNEAFGMINSMP----FSPDAGVWGTLLGACRLHGNVELAEVASRNLFDLDPQ-NSGYYVLLSNVHANAGQWESVLKIRSLMKERGVQKVP----------GCSWIDVNNTTHMFVAADRSHPQSSQIYLLLKNLFLELRKEGYVP-----------------------    
 114 UniRef50_D1IBS7        Whole genome shotgun sequen...   437    1e-121  1  12.7%   5:593    77:640     ----SLINMYSKCNQMNFALSIFSDpnVFAFNAIISGFITNGFPEEGFEFYQKMRNEGVI--PDKFTFPCAIKAC---LDVLEIKKIHGLLFKFGLELDVFIGSALVNCYLKFGLMEHAQVAFEELprDVVLWNAMVNGYagQFEMVLETFRRMNDESVVPSRFTVTGVLSVFamGDLNNGRIIHGFAMKMGYDSGVAVSNSLIDMYGKCKCIEDALEIFEMMREK----DIFSWNSIVSVHEQCGDHDGTLRLLDRMLGAGIQPDLVTVTTVLPACSHLAALMHGREIHGYMIVSGLGKDGKDID------DVLLKNAVIDMYAKCGSMRDAHLVFERMSNK------DVASWNIMIMGYGMHGYGNEALEMFSRMCEVQLKPDEVTFVGVLSACSHAGFVSQGRNFLVQMkkYDVApeHYTCVIDMLGRAGQLDEAYELALTMP----IEANPVVWRALLAACRLHKHAVLAEVAAQRVFE--LEPEhgSYVLMSNVYGAVGRYEEVLEVRHTMRQQNVRKTP----------GCSWIELKNGVHVFVSADRAHPEAHSIYAGLNSLTARLREHGYVPDVFCFFSPLLLDL----------    
 115 UniRef50_B9ILS5        Predicted protein n=5 Tax=r...   437    1e-121  1  11.4%   4:600    62:621     ---NRLLNLYAKSGeaRKLFDEMTQRDNFSWNAMLSLYAKSGLVEDLRVIFDNM------PSRDSVSYNTVISGFAGNGRGGPALGVFLRMQKEGLKPTEYTHVSVLNAcrRGKQI-HGRIIICNLGGNVFVCNALtyARCGEIDQARRLFDRMVIR----NVVTWNLMISGYLKNRQPecIDLFHEMQVSNLKPDQVTASSVLGAYIQAGYIDEARKVFGEIREK----DEVCWTIMIVGCAQNGKEEDALLLFSEMLLENARPDGYTISSVVSSCAKLASL------------YHGQVVHGKAFLMGVNDDLLVSSALVDMYCKCGVTRDAWTIFSTMQTR------NVVSWNSMIGGYALNGQDLEALSLYENMLEENLKPDSVTFVGVLSACVHAGLVEEGKEYFCSMSdhGLepDHYACMVNLFGRSGHMDKAVDLISSMSQ----EPNSLIWTTVLSVCVMKGDIKHGEMAARCLIELN-PFNAVPYIMLSNMYAARGRWKDVASIRSLMKSKHVKKFSAY----------SWIEIDNEVHKFVADDRTHPDAKIIHVQLNRLIRKLQEAGFSPNT----NLVLHDFGEDEKLE---    
 116 UniRef50_O04659        Pentatricopeptide repeat-co...   436    1e-120  1  10.5%  10:602    52:632     ---------CKDHCSARHVFENfiRSDVYIWNSLMSGYSKNSMFHDTLEVFKRLLNCSICV-PDSFTFPNVIKAYGALGREFLGRMIHTLVVKSGYVCDVVVASSLVGMYAKFNLFENSLQVFDEmeRDVASWNTVISCfsGEAEKALELFGRMESSGFEPNSVSLTVAISACSRlwLERGKEIHRKCVKKGFELDEYVNSALVDMYGKCDCLEVAREVFQKMPRK----SLVAWNSMIKGYVAKGDSKSCVEILNRMIIEGTRPSQTTLTSILMACSRSRNL------------LHGKFIHGYVIRSVVNADIYVNCSLIDLYFKCGEANLAETVFSKTQKDVAE------SWNVMISSYISVGNWFKAVEVYDQMVSVGVKPDVVTFTSVLPACSQLAALEKGKQIHLSISESRLETdlLSALLDMYSKCGNEKEAFRIFNSIPKK-----DVVSWTVMISAYGSHGQPREALYQFDEMQKFGLKPDGVTLLAVLSACGHAGLIDEGLKFFSQMRSKYGIEPIIEHYSCMIDILGRAGRLLEA----YEIIQQTPETSDNAELLSTLfaRLLVENY-PDDASTYMVLFN-LYASGESWDA-    
 117 UniRef50_Q9M2Y7        Pentatricopeptide repeat-co...   436    1e-120  1  11.5%   7:581    84:630     ------YAKDSKIHIARQLFDEIPQPDTVSYNTLISGYADARETFAAMVLFKRMRKLGF--EVDGFTLSGLIAACCD--RVDLIKQLHCFSVSGGFDSYSSVNNAFVTYYSKGGLLREAVSVFYGMdrDEVSWNSMIVAYGQhgAKALALYKEMIFKGFKIDMFTLASVLNALTSLdlIGGRQFHGKLIKAGFHQNSHVGSGLIDFYSKCGGCddSEKVFQEI----LSPDLVVWNTMISGYSMNEEleEAVKSFRQMQRIGHRPDDCSFVCVTSACSNLSSPSQCKQIHGLAIKSHIPSNR-----------ISVNNALISLYYKSGNLQDARWVFDRMPE------LNAVSFNCMIKGYAQHGHGTEALLLYQRMLDSGIAPNKITFVAVLSACAHCGKVDEGQEYFNTMKEtkIEpeHYSCMIDLLGRAGKLEEAERFIDAMPYK----PGSVAWAALLGACRKHKNMALAERAANELMV--MQPLAAtyVMLANMYADARKWEEMASVRKSMRGKRIRKKP----------GCSWIEVKKKKHVFVAEDWSHPMIREVNEYLEEMMKKMKKVGYVMD----------------------    
 118 UniRef50_C5Y5C2        Putative uncharacterized pr...   436    1e-120  1   9.3%  15:603   166:726     --------------LLHRMPHLGCVPDVFSYCIVLKSLCSDRKSGQADELLRMMAEGGAVCLPNAVAYNTVIDGFFKEGDVNKACDLFNEMVQRGISPDLSTYNCVVNALCKARAMDKAEAILRQMVDKGvtYNSLIYGYsgQWKEAVRVSKKMTSQGILPDVVTLNSLmsLCKHGKIKDARDVFDSMAMKGQKTDIFSYKIMLNGYATKGCLVDLTELFNLMLSDGIAPDSHIFNVLIKAYAKCGMLDRATIIFNEMREQGVEPDVVTYSTVIAALCRIGKMDDAVEKFNQM------------IDQGVAPSISTYHFLIQGFCTHGDLLKAKDLVLQMMNKGMR--PDIGCFNFIINNLCKLGRVMDAQNIFDFTISIGLHPNVMVYNTLMDGYCLVGKMENALRVFDVMVSAGIQPnvYGTLVNGYCKVGRIDEGLSLFREILHKGIK-PSTTLYNIILHGLFQAGRTVPAKVKFHEMTESGIAMDRYTYSIVLGGLFKNSCSDEAILLFKELHAMNVKIDITTL---------------------NIMIAGMFQI-RRVEEAKDLFASISRSGLVPSVV-TYSIMMTNLIKEGLVEEAD    
 119 UniRef50_C5XVI6        Putative uncharacterized pr...   436    1e-120  1   9.9%   2:577    81:703     -AAISAACRAGDLAAARDLLGRMPDRNAVSWNTVIAAVARSDSPGEALEMYRGMLQEGL--APTNFTLASVLSACGAVAALDDGRRCHGLAVKVGLDGNQFVENGLLGMYTKCGSVADAVRLFDWMspNEVSFTAMMGGLAQSgdDALRLFARMSRSAIRVDPVAVSSVLGACAQAciRLAQSIHALVVRKGFDSDQHVGNSLIDMYAKGMKMDEAMKVFESMSS----VSIVSWNILVTGYGQLGCYERALEVLDLMQESGFEPNEVTYSNMLASCIKARDVPSAreLFRRMQHQngKQVHSASVKLLLHNDMFVASGLIDMYSKCGQVGIAQIIFNMMTER------DVVCWNSMISGLAIHSLNEEAFDFFKQMRENGMFPTESSYASMINSCARLSSIPQGRQIHAQVLKDGYDQNVysSLIDMYAKCGNMDDARLFFNCMIVK-----NIVAWNEMIHGYAQNGFGEKAVELFEYMLTTKQKPDSVTFIAVLTGCSHSGLVDEAIAYFNSMESNYGIRPLVEHYTCLIDALGRAGRFAEVV----AVIDKMPYKDDAILWEVLLAACVVHHN--------------------------    
 120 UniRef50_Q0DBH2        Os06g0565000 protein n=4 Ta...   435    1e-120  1  11.9%   6:518   166:671     -----RLARDRSGRLVRRLFEQLPAPNVFTFNIVIDFLCKEGELAEARSLFSRMKEMGCL--PDVVTFNSLIDGYGKCGELDEVEQLVEEMRRSGCKADVVTYNALINCFCKFGRMETAYGYFAAMKREGVMANVVTfaFCKEgrEAMKLFAQMRVRGMALNEFTYTCLIDgcKAGRLDDAIVLLDEMVRQGVPLNVVTYTVLVDGLCKERKVAEAEDVLRMMEKAGVRANELLYTTLIHGHFMNKNSEKALGLLSEMKNKGLELDISLYGALIQGLCNVHKLDEAKSLLTKMDES------------GLEPNYIIYTTMMDACFKSGKVPEAIAMLQKILDSG--FQPNVITYCALIDGLCKAGSIDEAISHFNKMRDLGLDPNVQAYTALVDGLCKNGCLNEAVQLFNEMVHKGMSLdvYTALLDGYLKQGNLHDAFALKAKMIDSGLQL-DLFCYTCFISGFCNLNMMPEAREVFSEMIGHGIAPDRAVYNCLISKYQKLGNLEEAISLQDEMER-------------------------------------------------------------------------------------    
 121 UniRef50_Q9LN01        Pentatricopeptide repeat-co...   435    1e-120  1   9.0%  16:582    87:654     ---------------AISVFKTIQEPNLLIWNTMFRGHALSSDPVSALKLYVCMISLGLL--PNSYTFPFVLKSCAKSKAFKEGQQIHGHVLKLGCDLDLYVHTSLISMYVQNGRLEDAHKVFDkpVKDVVSWNAMISGYagNYKEALELFKDMMKTNVRPDESTMVTVVSACAqgSIELGRQVHLWIDDHGFGSNLKIVNALIDLYSKCGELETACGLFERLPYK----DVISWNTLIGGYTHMNLYKEALLLFQEMLRSGETPNDVTMLSILPACAHLGAIDIGRWIHVYIDKRLKGVTNASSLR----------TSLIDMYAKCGDIEAAHQVFNSILHK------SLSSWNAMIFGFAMHGRADASFDLFSRMRKIGIQPDDITFVGLLSACSHSGMLDLGRHIFRTmqDYKMTpeHYGCMIDLLGHSGLF----KEAEEMINMMEMEPDGVIWCSLLKACKMHGNVELGESFAENLIK--IEpnPGSYVLLSNIYASAGRWNEVAKTRALLNDKGMKKVP----------GCSSIEIDSVVHEFIIGDKFHPRNREIYGMLEEMEVLLEKAGFVPDT---------------------    
 122 UniRef50_Q9SUH6        Pentatricopeptide repeat-co...   435    1e-120  1   9.8%   1:582   156:705     LGSnkMYFKFWRVEDARKVFDRMPEKDTILWNTMISGYRKNEMYVESIQVFRDLINESCT-RLDTTTLLDILPAVAELQELRLGMQIHSLATKTGCYSHDYVLTGFISLYSKCGKIKMGSALFREfkPDIVAYNAMIHGYTSNgeLSLSLFKELMLSGARLRSSTLVSLVPV--SGHLMLIYAIHGYCLKSNFLSHASVSTALTTVYSKLNEIESARKLFDESPEKSLP----SWNAMISGYTQNGLTEDAISLFREMQKSEFSPNPVTITCILSACAQLGALSLGKWVHDLVRSTD------------FESSIYVSTALIGMYAKCGSIAEARRLFDLMTKK------NEVTWNTMISGYGLHGQGQEALNIFYEMLNSGITPTPVTFLCVLYACSHAGLVKEGDEIFNSMirYGFEPshYACMVDILGRAGHLQRALQFIEAMS----IEPGSSVWETLLGACRIHKDTNLARTVSEKLFE--LDPDNVGyvLLSNIHSADRNYPQAATVRQTAKKRKLAKAPGYTL----------IEIGETPHVFTSGDQSHPQVKEIYEKLEKLEGKMREAGYQPET---------------------    
 123 UniRef50_D1J475        Whole genome shotgun sequen...   435    1e-120  1  12.8%   6:552   202:757     -----LFRK-GRVDDALHLLDEMLQpnSNTGHIVFSALSKRDKVGRAVdlVSKFAE--HEVFPNSIWLTQLISRLCRSGRTDRAWDVLHGLMKLGGVMEAASCNALLTALGRAREFKRMNTLLAEMKEMDIQPNVVTFGILInvDEALEVFEKmeSNgvEPDVITYNTLIDGLCkgRQEEGLGLVERMRSqrCMPNTVTYNCLIDGYCKASMIEAARELFDQMNKDGVPPNVVTLNTLVDGMCKHGRINGAVEFFNEMQGKGLKGNAVTYTALIRAFCNVNNIEKAMELFDEMLE------------AGCSPDAIVYYTLISGLSQAGKLDRASFVLSKMKEAG--FSPDIVSFNVLINGFCRKNKLDEAYEMLKEMENAGIKPDGVTYNTLISHFSKTGDFSTAHRLMKKMVKEGLVpvTYGALIHAYCLNGNLDEAMKIFRDMSSTSKVPPNTVIYNILINSLCRKNQVDLALSLMDDMKVKGVKPNTNTFNAMFKGLQEKNWLSKAFELMDRMTEHACNPDYITM--EILTEWLSAVGETAKLKSFVQG---------------------------------------------------    
 124 UniRef50_D1J572        Whole genome shotgun sequen...   435    1e-120  1   9.3%   7:566   190:748     ------YSKYGSVKKALELFAKMPERDTVSWNTMISILSQHGFGAETLNTFLEMWNQGF--RPNSMTYASVLSACTSIYDLEWGAHLHARIVRMEPCLDVYAGCGLIDMYAKCGRLESARQVFDGleHNAVSWTSLIGGVAQAgeEALVLFNQMREVPVASDQFTLATVLGVcsQKDISIGEQLHAHTITRGLDSSVPVANALVTMYAKCGDVWKANHAFELMPIR----DIISWTAMITAFSQAGDVEKAREYFDKMPER----NVISWNSMLATYMQRGYWEEGLKVYIQMLREGVkqILAQAEKLGFSSNVSVANSVVTMYSRCGQIEEAQKMFSSIVMK------NLVSWNAMMAGYAQNGQGRKVIEIFEKMLNIGNVPDQISYVSVLSGCSHSGFVSEGQYYFLSmkDHGISpeHFVCMVDLLGRAGQLEQAKNLINQMP----FKPNAAIWGALLAACRIHGNTKLAELAVKNLLELDAE-GPGSYCLLANIYSESGKIQGVTNVRKLMRDKGVRKNP----------GCSWIEVDNRVHVFTVDDTNHPQIKDVHRML-------------------------------------    
 125 UniRef50_C5Y455        Putative uncharacterized pr...   435    1e-120  1  12.9%  15:602   134:787     --------------DALKLFDLLLPhsVTAFNHLLTAVSRAseSQLGISLFDRmrECSDKVAPDRCTYSILIGCFCRMGRLEHGFAVFGLILKTgrVNHIVFNQLLKGLCDAKRLDEATNILLWRMppNVVSYNTLVKGFCnrAEEALELLHVmqGLSCPPDVVSYNTVINGfrEGQVDKAYNLFLQMIDRGIPPDVVTYNTVIDGLCKAQVVDRAKAVFQQMLDKGVKPSNGTYNCLIHGYLSTGKWKEVVRLLEEMSTHDLEPDCFIYALLLDYLCKNGRCTEARNIFDSVIRKGIkcIFDKMRQQGLSPNVVNYGALIDALCKLGRVDDAILKFNQMINedQGIRLDVVVFNTLMCDLCREGRVMEAQRLIELMLRVGVRPDVISYNTLVDGHCLTGRTDEAAKLLDVMVSIGLKPntYNTLLHGYCKARRIDDAYSLLREMLMNG-PTPDVVTYNTILHGLFQTGRFSEAKELYLNMINSRTQWDMYTYNIILNGLCKNNCVDEAFKMFQSLCSKGLQLHIIT-FNIMIGALLKGGKKEDAMDLfyCLIAENLIKEGSLEELGVLFSAMEENGTAPNS-RMLNALVRRFLHRGDISRA-    
 126 UniRef50_Q9FJE6        Putative pentatricopeptide ...   435    1e-120  1  12.3%   9:602   204:841     --------KFRHFGLAMemVSVGIRPDVYIYTGVIRSLCELKDLSRAKEMIAHMEATGCDVnpDVVTYCTLVYGLCKVQEFEIGLEMMDEMLCLRFSPSEAAVSSLVEGLRKRGKIEEALNLVKRVVDFGVSPNLFVYNAlrKFHEAELLFDRMGKIGLRPNDVTYSILIDMFcrGKLDTALSFLGEMVDTGLKLSVYPYNSLINGHCKFGDISAAEGFMAEMINKKLEPTVVTYTSLMGGYCSKGKINKALRLYHEMTGKGIAPSIYTFTTLLSGLFRAGLIRDAVKLFNEMAE------------WNVKPNRVTYNVMIEGYCEEGDMSKAFEFLKEMTEKG--IVPDTYSYRPLIHGLCLTGQASEAKVFVDGLHKGNCELNEICYTGLLHGFCREGKLEEALSVCQEMVQRGVDLdcYGVLIDGSLKHKDRKLFFGLLKEMHDRGLK-PDDVIYTSMIDAKSKTGDFKEAFGIWDLMINEGCVPNEVTYTAVINGLCKAGFVNEAEVLCSKMQPV-SSVPNQVTYGCFLDILtkAVELHNAILKgyNMLIRGFCRQGRIEEASELITRMIGDGVSPDCI-TYTTMINELCRRNDVKKA-    
 127 UniRef50_Q9SI53        Pentatricopeptide repeat-co...   434    1e-120  1  11.8%  10:603     5:558     ---------MSKIKLFRPVVTLRCSYSSTDQTLLLSEFTRLCYQRDLPRAMKAMDSLQSHGlaDSATYSELIKCCISNRAVHEGNLICRHLYFNGHRPMMFLVNVLINMYVKFNLLNDAHQLFDQmqRNVISWTTMISAYSKchQKALELLVLMLRDNVRPNVYTYSSVLRSC--NGMSDVRMLHCGIIKEGLESDVFVRSALIDVFAKLGEPEDALSVFDEM----VTGDAIVWNSIIGGFAQNSRSDVALELFKRMKRAGFIAEQATLTSVLRACTGLALLELGMQAHVHIVKYDQ--------------DLILNNALVDMYCKCGSLEDALRVFNQMKER------DVITWSTMISGLAQNGYSQEALKLFERMKSSGTKPNYITIVGVLFACSHAGLLEDGWYYFRSMKkyGIDpeHYGCMIDLLGKAGKLDDAVKLLNEME----CEPDAVTWRTLLGACRVQRNMVLAEYAAKKVIALDPE-DAGTYTLLSNIYANSQKWDSVEEIRTRMRDRGIKKEP----------GCSWIEVNKQIHAFIIGDNSHPQIVEVSKKLNQLIHRLTGIGYVPET----NFVLQDL--EGEQMEDS    
 128 UniRef50_B9N472        Predicted protein n=2 Tax=M...   434    1e-120  1  12.7%   6:582   170:706     -----MYAVCGNLSDARKVFDGSSVLDMVSWNSMLAGYVLVGNVEEAKDVYDRM------PERNVIASNSMIVLFGKKGNVEEACKLFNEMKQKDLVSW----SALISCYEQNEMYEEALILFKEMNANGIMVDEVVVLSVLSACSRllVHGLVVKVGIETYVNLQNALIHMYSSceVVTAQKLFSE----SCCLDQISWNSMISGYVKCGEIEKARALFDSMPDK----DNVSWSAMISGYAQQDRFTETLVLFQEMQIEGTKPDETILVSVISACTHLAALDQGKWI------------HAYIRKNGLKINIILGTTLINMYMKLGCVEDALEVFKGLEEKG------VSTWNALILGLAMNGLVDKSLKTFSEMKEHGVTPNEITFVAVLGACRHMGLVDEGHRHFNSMieHKIGPnhYGCMVDLLGRAGMLKEAEELIESMP----MAPDVSTWGALLGACKKYGDNETGERIGRKLVE--LHPDHDGFNVLlnIYASKGNWVDVLEVRGMMRQHGVVKTP----------GCSMIEAHGRVHEFLAGDKTHPQNEHIEHMLDEMAKKLKLEGYAPDT---------------------    
 129 UniRef50_C5XD50        Putative uncharacterized pr...   434    1e-120  1  12.3%   3:602   160:765     --AIRVFARAGCLAEASALLEAAPAPDASAYTALVSEFSRAGRFRDAVAVFRRMVDGGVQPAL--VTYNVVLHVYSKMSVPweVVVLVDSMKEDGIELDRYTYNTLISCCRRRGLYREAAQMFDEmePDKVTFNSLlyGKARRHEEAIGVLKKMENAGCTPSVVTYNSLISAYVKDGleEALELKQEMEFRGMKPDVVTYTTLISGLDRIGKIDAALATYSEMVRNGCSPNLCTYNALIKMHGVRGKFTEMMIVFDDLRSAGYVPDVVTWNTLLAVFGQNGLDSEVSGVFKEMK------------KAGYIPERDTYVSLISSYSRCGLFDQAMEIYKRMIEAG--IYPDISTYNAVLSALARGGRWVQAEKLFAEMEDRDCKPDELSYSSLLHAYANAKKLDKMKAlfQELRRRRcdINVLNAMVSIYGKNKMVKKVEEVLTLMKENSINH-SAATYNSLMHMYSRLGDCEKCEAILTEIKSSGMRPDRYSYNTVIYAYGRKGQMKEASRLFSEMKCSGVKPDIVT----------------------YNIFIKSYVANLMFEEAIDLVRYLVAQGCKPNE-RTYNSILEGYCRHGRMVEA-    
 130 UniRef50_Q0D563        Os07g0578800 protein n=7 Ta...   434    1e-120  1   9.6%  29:581   330:872     ----------------------------PSWAYQIRMAASQGQFLHAISLFLQMRA-SVAPRSSvaSLPAALKSCAGLGLCTLAASLHALAIRSGSFADRFTANALLNLCIkgGLESAarKVFDEmeRDAVSWNTLILGCakRHQEALSMVREMWRDGFMPDTFTLSTVLPIFAEcdIKRGMVVHGYAIKNGFDNDVFVGSSLIDMYANCTQMDYSMKVFDSFSDC----DAVLWNSMLAGYAQNGSVEEALGIFRRMLQAGVRPVPVTFSSLIPAFGNLSLLRLGKQL------------HAYLIRARFNDNIFISSSLIDMYCKCGNVDIARRVFNGIQS------PDIVSWTAMIMGYALHGPTTEAFVLFERMELGNVKPNHITFLAVLTACSHAGLVDNGWKYFNSMsqYGFVpeHCAALADTLGRAGDLDEAYNFISEMKIK----PTSSVWSTLLRACRVHKNTVLAEEVAKKIFE--LEpsMGSHVILSNMYSASGRWNEAAQLRKSMRIKGMKKEPA----------CSWIEVKNKLHVFIAHDKSHPWYDRIIDALNVYSEQMIRQGYVPN----------------------    
 131 UniRef50_B9STP1        Pentatricopeptide repeat-co...   434    1e-120  1  10.2%   4:569    73:645     ---NKLMAFcgNLNYAEKMFKYIRYPCLLIYNLIIKAFAKKGNYKRTLVLFSKLREDGLW--PDNFTYPFVFKAIGYLGEVSKAEKLRGLVTKTGLEFDTYVRNSLIDMYAQLALTDVMKMLFDEMprDVISWNVMISGycRRFEDAINVFCRMQEegLMPDEATVVSTLSAcaLKRLELGKKIHHY-VRDNVKFTPIIGNALLDMYCKCGCLSIARAVFEEMPSKNVirDVVIWTAMINGYVQFNRFDEAVALFREMQIRKVKPDKFIVVSLLTGCAQTGAIEQGKWI------------HEFIDENRIPIDAVVGTALIEMYAKCGFIEKALEIFYGLRVK------DTASWTSIICGLAMNGKTSKALELFSKMKQAGVRPDDITFIGVLSACSHGGLVEEGRKFFNSMreYQIKpeHYGCLVDLLGRAGLLNEAEELIKKIPDENKAI-TVPLYGSLLSACRIYGNVEMGERVAKQLVKFESS-DSSVHTLLANIYAFADRWEDVTKVRRKMKDLGVKKTP----------GCSSIEVDSIIHEFFSGHPSHPEMREIYYMLNIM----------------------------------    
 132 UniRef50_Q9STE1        Pentatricopeptide repeat-co...   434    1e-120  1   9.4%   4:580   281:826     ---LSMYSKCGRFDDASKLFRMMSRADTVTWNCMISGYVQSGLMEESLTFFYEM--ISSGVLPDAITFSSLLPSVSKFENLEYCKQIHCYIMRHSISLDIFLTSALIDAYFKCRGVSMAQNIFSQCNsdVVVFTAMISGYLHNGldSLEMFRWLVKVKISPNEITLVSILPVIglLALKLGRELHGFIIKKGFDNRCNIGCAVIDMYAKCGRMNLAYEIFERLSKR----DIVSWNSMITRCAQSDNPSAAIDIFRQMGVSGICYDCVSISAALSACANLPS------------ESFGKAIHGFMIKHSLASDVYSESTLIDMYAKCGNLKAAMNVFKTMKEK------NIVSWNSIIAACGNHGKLKDSLCLFHEMVEkgIRPDQITFLEIISSCCHVGDVDEGVRFFRSmeDYGIQpeHYACVVDLFGRAGRLTEAYETVKSMP----FPPDAGVWGTLLGACRLHKNVELAEVASSKLMDLDPS-NSGYYVLISNAHANAREWESVTKVRSLMKEREVQKIPGY----------SWIEINKRTHLFVSGDVNHPESSHIYSLLNSLLGELRLEGYIP-----------------------    
 133 UniRef50_D1J3N2        Whole genome shotgun sequen...   434    1e-120  1  10.7%   7:577    49:579     ------LGRLGRVEEARRVFNEMIQRDVVSWNSMINGYSQNGKVDEARLLFDAF------VGKNIRTWTILLTGYAKEGRIEEAREVFESMTERNVVSW---NAMISGYVQNGDLKNARKLFDEMPEKNVASWNSVVTGYChrMSEARELFDQMPER----NSVSWMVMISGYVhsDYWEAWDVFVKMCRTR---NEYSWTTMIAAFAQCGRLDDAIQLYERVPE---QTNSASWAAMIAGFVQNEESREALELLIELHRSGSVPSDSSFTSALSACANIGDVEIGRVI------------HSLAIKTGCQFNSYVMNGLISMYAKCGNVEDGSHVFrrMPKR------DVVSWTAIISAYVQAGHGEVALDLFLDMLARGIKPNQLTVTSLLSACGNLGAIKLGEQFHALIFKLGF--DTfnSLITMYFKCG-YEDGFCVFEEMPEH-----DLITWNAVLVGCAQNGLGKEAIKIFEQMEVEGILPDQMSFLGVLCACSHAGLVDEGWAHFNSMTQKYGIMPLVYHYTCMVDLLGRAGYLSEAE----ALIENMPVKPDSVIWEALLGACRIHRN--------------------------    
 134 UniRef50_Q9ZUW3        Pentatricopeptide repeat-co...   434    1e-120  1  11.6%   3:582   232:780     --SNSLINLYLKCGNVRklFDKTEVKSVVTWNSMISGYAANGLDLEALGMFYSMRL--NYVRLSESSFASVIKLCANLKELRFTEQLHCSVVKYGFLFDQNIRTALMVAYSKCTAMLDALRLFKEivGNVVSWTAMISGFLQnkEEAVDLFSEMKRKGVRPNEFTYSVILTALPVISPSE---VHAQVVKTNYERSSTVGTALLDAYVKLGKVEEAAKVFSGIDDK----DIVAWSAMLAGYAQTGETEAAIKMFGELTKGGIKPNEFTFSSILNVCAATN-----------ASMGQGKQFHGFAIKSRLDSSLCVSSALLTMYAKKGNIESAEEVFKRQREK------DLVSWNSMISGYAQHGQAMKALDVFKEMKKRKVKMDGVTFIGVFAACTHAGLVEEGEKYFDIMVRdkIApeHNSCMVDLYSRAGQLEKAMKVIENMPN----PAGSTIWRTILAACRVHKKTELGRLAAEKIIAMKPE-DSAAYVLLSNMYAESGDWQERAKVRKLMNERNVKKEPGY----------SWIEVKNKTYSFLAGDRSHPLKDQIYMKLEDLSTRLKDLGYEPDT---------------------    
 135 UniRef50_Q9SH60        Pentatricopeptide repeat-co...   434    1e-120  1   9.3%  10:601    84:625     ---------FKSLDDAIDFFDYMVRstAVDCNKVIGVFVRMNRPDVAISLYRKM-EIRRIP-LNIYSFNILIKCFCDCHKLSFSLSTFGKLTKLGFQPDVVTFNTLLHGLCLEDRISEALALFGYMVETGFL-------------EAVALFDQMVEIGLTPVVITFNTLINGLceGRVLEAAALVNKMVGKGLHIDVVTYGTIVNGMCKMGDTKSALNLLSKMEETHIKPDVVIYSAIIDRLCKDGHHSDAQYLFSEMLEKGIAPNVFTYNCMIDGFCSFGRWSDAQRLLRDMIERE------------INPDVLTFNALISASVKEGKLFEAEKLCDEMLHR--CIFPDTVTYNSMIYGFCKHNRFDDAKHMFDLMAS----PDVVTFNTIIDVYCRAKRVDEGMQLLREISRRGLVANtyNTLIHGFCEVDNLNAAQDLFQEMISHGVC-PDTITCNILLYGFCENEKLEEALELFEVIQMSKIDLDTVAYNIIIHGMCKGSKVDEAWDLFCSLPIHGVEPDVQT----------------------YNVMISGFCGKSAISDANVLFHKMKDNGHEPDN-STYNTLIRGCLKAGEIDK--    
 136 UniRef50_O64766        Pentatricopeptide repeat-co...   433    1e-119  1   9.3%   7:576    56:612     ------LCKVGKIAEARKLFDGLPERDVVTWTHVITGYIKLGDMREARELFD-----RVDSRKNVVTWTAMVSGYLRSKQLSIAEMLFQEMPERNVVSW---NTMIDGYAQSGRIDKALELFDEMPERNIVSWNSMVKALVQRgdEAMNLFERMPRR----DVVSWTAMVDGLAkgKVDEARRLFDCMPER----NIISWNAMITGYAQNNRIDEADQLFQVMPERDFA----SWNTMITGFIRNREMNKACGLFDRMPEK----NVISWTTMITGYVENKENEEALNVFSKMLRDggQQIHQLISKSVHQKNEIVTSALLNMYSKSGELIAARKMFD----NGLVCQRDLISWNSMIAVYAHHGHGKEAIEMYNQMRKHGFKPSAVTYLNLLFACSHAGLVEKGMEFFKDLVRDESLpeHYTCLVDLCGRAGRLKDVTNFIN----CDDARLSRSFYGAILSACNVHNEVSIAKEVVKKVLETGSD-DAGTYVLMSNIYAANGKREEAAEMRMKMKEKGLKKQP----------GCSWVKVGKQNHLFVVGDKSHPQFEALDSILSDLRNKMRKN---------------------------    
 137 UniRef50_B9SDG1        Pentatricopeptide repeat-co...   432    1e-119  1  12.8%   2:601   156:815     -GYCKAGEMSLALDLVERMRKEGLLSDIVSYNTLINGFCKRGEYDKAKSLLHEISESrlNLEADLITYTTIISTYCKQHGLEEARALYEEMIINGFLPDVVTYSSIVNGLCKDGRLSEAQELLREMKKMGVDPNHVAYtkAGSAWEAFACQSQLVVRGMTLDLVMCTTLVDGlkSSKPKEAEDMFRALSKLNLIPNSITYTALIDGYCKVGDMERVESLLQEMEEKHINPNVITYSSIINGYTKKGILDEAINVMKKMLDQNIIPNAYVYAILIDGYCKAGKQEIATDLYNEMKLSGLkeLLKDVTSRGLLLDHVNYTSLMDGFFKAGKESAALNMVEEMTEKSIPF--DVVTYNVLINGLLEHGKY-EAKSVYSGMIEMGLAPNQATYNIMIKAYCKQGELDNALELWNEMKSHKIMPstCNTLVVGLSEAGEIEKAMNVLNEMSVMGI-HPNLVIHRVLLNASSKSGKANAVLQMHEQLVDMGLKINQEAYNNLIVVFCRLRMTKKATSVLKYMIRDGFVADTVT-YNALIRGYCESSHVKKALATyyNLLLGGLLGAGLMAERDELFDKMKENGLNPDA-STYDTLISGYGKIGNKKE--    
 138 UniRef50_D1HPW7        Whole genome shotgun sequen...   432    1e-119  1  10.7%   3:574   184:726     --ANSmyCKCGRVEDAQALFELMDARDVISWNSLVSGYAQLGNIREVLQLLIRMKTDG--IEPDQQTFGSLVSAAAMQSKLGVGKMVHGHILRAGLEQDSHIETSLIGMYLKCGNVNSAFRIFemMHKDVISWTAMISGLVQnaDMAVTVFRRMLKSRVMPSTATIASVLAACAegSFPLGTSVHGYILRQRIKLDIPSQNSLVTMYAKCGHLEQSCSVFDRMSRR----DIVSWNAIVSGHAQNGHLCKALLLFNEMRKARQRPDSITVVSLLQACASIGALHQGKWI------------HNFVTKSCLGPCILIDTALVDMYSKCGDLGSAQKCFDRMPQQ------DLVSWSSIIAGYGSHGKGETALRMYSDFLHTGIQPNHVIYLSILSACSHNGLVDQGLSFFHSmkDFGIEpeHRACIVDLLSRAGRVEEAYSFYKRM----FPKPSMDVLGILLDACRTTGNVELGDIVAREIVI--LKpnAGNYVQLAHSYASMKRWDGVGEVWTQMKSLHLKKLP----------GWSFIELHGTITTFFTDHSSHPQFEEIIDRVLLLNKRRR-----------------------------    
 139 UniRef50_Q5JNB4        Vegetative storage protein-...   432    1e-119  1   8.2%   5:589    50:618     ----ALLDMYVKCadAAHIFATMPARDLVAWNAMLAGYAHHGMYHHAVAHLLSMQMQMHRLRPNASTLVALLPLLAQQGALAQGTSVHAYCIRACLHPnvLLGTALLDMYAKCGSLLYARRVFDAMprNEVTWSALIGGFVLCSrqAFLLFKAMLAQgcFLSPTSIASALRACASLdlRMGEQLHALLAKSGVHADLTAGNSLLSMYAKAGLIDQAIALFDEMAVK----DTVSYSALVSGYVQNGRAEEAFLVFKKMQACNVEPDAATMVSLIPACSHLAAL------------QHGRCSHGSVIIRGLASETSICNALIDMYAKCGRIDLSRQVFNMMPSR------DIVSWNTMIAGYGIHGLGKEATALFLEMNNLGFPPDGVTFICLLSACSHSGLVIEGKHWFHVmhGYGLTpeHYICMVDLLSRGGFLDEAYEFIQSMPLR----ADVRVWVALLGACRVYKNIDLGKKVSRMIQELGPE-GTGNFVLLSNIYSAAGRFDEAAEVRIIQKVQGFKKSP----------GCSWIEINGSLHAFVGGDQSHPQSPEIYRELDNILVGIKKLGYQPDTSFVLQDL--------------    
 140 UniRef50_B9R997        Pentatricopeptide repeat-co...   432    1e-119  1  10.3%   3:602    82:663     --AMRNYGRKGKVQEAVDVFERmcEPSIFSYNAIMNILVEYGYFNQAHKVYLRMKHEG--IAPDVCTFTIRIKSFCRTKRPLAALRLLNNMPSQGCELNAVVYCTVISGFYEENYQVEAYELFNKMLRLGitFNKLMHILCKKGheGEKLLNKVLKNGVCPNLFTFNIFIQGLCrsKVVEAENYLHKMVNKGLEPDDFTYNTIIYGYSKVGKIQDASRILKDAKFKGFVPDEFTYCSLIIGVCQDGDIDHALALFEEALGKGLKPSIVLYNTLIKGLSQQGLVLKALQLMNDMSKE------------GMSPDIWTYNLVINGLCKMGCVSDANNLLNAAIAKGY--LPDIFTFNTLIDGYCKRLKMDNAIGILDSMWSHGVTPDVITYNSILNGLCKAAKPEDVMETFKMIMEKGCLPntYNILIESLCKARKVTEALDLLEEIRNRGLI-PDPVSFGTVISGFCNNGDLDEAYQLFRRMEQqrICHTVATYNIMINAFSEKLDMDMAQKLFHEMGDKG----------CDPDSYTYRVMIDGFCKVGNVNSGYDFLLKEI------------EIGFVP-SLTTFGRVINCLCVQHRVHEA-    
 141 UniRef50_Q9SVA5        Pentatricopeptide repeat-co...   432    1e-119  1  10.0%   4:574   288:831     ---NVLIDSYVKCGraHKLFNGMPNKNIISWTTLLSGYKQNALHKEAMELFTSMSKFGL--KPDMYACSSILTSCASLHALGFGTQVHAYTIKANLGNDSYVTNSLIDMYAKCDCLTDARKVFDIFAadVVLFNAMIEGYSreLHEALNIFRDMRFRLIRPSLLTFVSLLRASASlsLGLSKQIHGLMFKYGLNLDIFAGSALIDVYSNCYCLKDSRLVFDEMKVK----DLVIWNSMFAGYVQQSENEEALNLFLELQLSRERPDEFTFANMVTAAGNLASVQLGQEFHCQLLKR------------GLECNPYITNALLDMYAKCGSPEDAHKAFDSAASR------DVVCWNSVISSYANHGEGKKALQMLEKMMSEGIEPNYITFVGVLSACSHAGLVEDGLKQFELMLRFGIEpeHYVCMVSLLGRAGRLNKARELIEKMPTK----PAAIVWRSLLSGCAKAGNVELAEHAAEMAILSDPK-DSGSFTMLSNIYASKGMWTEAKKVRERMKVEGVVKEPGR----------SWIGINKEVHIFLSKDKSHCKANQIYEVLDDLLVQIR-----------------------------    
 142 UniRef50_C5Z8N7        Putative uncharacterized pr...   431    1e-119  1  11.8%   1:602   226:914     LGYCRTGDLRKACWLLMMMPLMGCRRNEYSYTILIQGLCETRCVREALVLLLMMLQDGCS--PNLHTYTLLIRGLCKEGRIHDARVLLDEMPRRGVVPSVWTYNAMIDGYCKSGRLKDALGIKTLMEGNGctYNSLIHGLCgkPDEAEELLNGAIARGFSPTVITFTNIINGYCKarLKEAKDTLNEIFANGLSPNVVIYTSIIDAYCKVGKVGAALEVFKLTEHEGCRPNAWTYSSLIYGLIQDQKLHKAMALITKMQEDGITPGVITYTTLIQGQCKKHEFDNAFRLFEMMEQNgaEEAYSFLVKKGVVLTKVTYTSLVDGFSKAGNTEFAAALIEKMVNEGC--KADSHTYSVLLQALCKQKKLNEALSILDQMTLRGVKGNIVAYTIIISEMIKEGKHDHAKSMFNEMISSGHKPstYTVFISSYCKIGRIEEAGHLIGEM-ERNGVAPDVVTYNVFINGCGHMGYMDCAFSTLKRMIDASCEPNYWTYWLLLKHFLksGMwdTVWQLLERMVKHGLNPTAVT-YSSIIAGFCKATRLEEAceEIYTMLIKCCCDIKLFEKAASFVTNMIECGFQP-HLESYHYLITGLCDEGDYDKA-    
 143 UniRef50_Q6ATD8        Os05g0275000 protein n=5 Ta...   431    1e-119  1  10.7%   7:602   154:760     ------LGREGQHDVVCDLLDEMPLpdVRAYTTVLHALSRAGRYERALQLFAELR--RQGVVPTIVTYNVVLDVYGRMGrwPRIVALLEEMRAAGVEPDDFTASTVIAACGRDGLLDQAVAFFEDLKARGHVPCVVTygKAGNYTEALRVLKEMEDSGCQPDAVTYNELAGTYARAGfeEAAKCLDTMTSKGLLPNTFTYNTVMTAYANVGRVDEALALFDRMKKNGYVPNVNTYNLIFGMLGKKSRFTAMLEMLEEMSRSGCTPNRVTWNTMLAVCGKRGMEDYVTRVLNGMKSC------------GVELSRDTYNTLISAYGRCGSRTYAFKMYDEMISSG--FTPCLTTYNALLNVLSRQGDWSTAQSIVSKMLKNGFKPNDQSYSLLLQCYAKGgrLEGVEKAFQEVKAQGYKPdiFNSMLAMYAKNGLYSKATEMFDSIKQSGL-SPDLITYNSLMDMYAKSNESWEAEKILKQLKSSQVKPDVVSYNTVINGFCKQGLIKEAQRILSEMIADGMAP----------CVVTYHTLVGG------------YASLEMFNEAREVVNYMIHHNLKPMEL-TYRRVVDSYCKAKRYDEA-    
 144 UniRef50_C5XG27        Putative uncharacterized pr...   431    1e-119  1  13.1%  13:602    30:677     ------------LGDALKLFDELLPhsVVAFNHLLAAVSRSseSETVVSLFNRmrDCYIKVAPNRCTYSILIGCFCRMGHLEHGFAAFGLILKTgrMDHIVINQLLNGLCDGKRVGEAMDVLLQRMppDTVSYNILLKGLckRAEEALELLHMMADdsCPPNVVSYSTVINGftEGQVDKPYNLFLEMMDRGIPPDVVTYTTVIDGLCKAQLFDRAEAVFQQMIDNGVKPNIDTYNCLIHGYLSIGKWKEVVRMLEEMSAGGPKPNCCTYGSLLNYLCKNGRCREARFFFDSMIGKGIKplLNLMVANGISPNHHIFNIFFSAYAKCGIIDKAMDIFNKMRQQG--LSPDAVSYGALIDALCKLGRVDDAEVKFNQMINEGVTPDIVVFSSLVYGLCTVDKWEKVEELFFEMLNVGIHPNifNTILCNLCKEGRVMEGQRLVDSIECMG-VRPDVISYNTLIDGHCLAGTIDEASKLLEGMVSVGLKPDSFSYNTLLHGYCKAGRIDSAYSHFRKMLSNGITPGVVT-YNTILHGLFQTKRFSEAKELYLNMINsgLCKSNCVDEAIKMFQNLCSKGLQLNII-TFNIMIGALLKGGRKEDA-    
 145 UniRef50_Q9FRI5        Pentatricopeptide repeat-co...   431    1e-119  1   9.3%   7:587    90:707     ------YCASGDITLARGVFEKAPvrDTVMYNAMITGFSHNNDGYSAINLFCKMKHEGF--KPDNFTFASVLAGLALVADDeqCVQFHAAALKSGAGYITSVSNALVSVYSKcsLLHSARKVFDelEKDERSWTTMMTGYVKNgqEALEMVRRMVSSGIELDEFTYPSVIRACATAGlqLGKQVHAYVLRrsFHFD----NSLVSLYYKCGKFDEARAIFEKMPAK----DLVSWNALLSGYVSSGHIGEAKLIFKEMKEK----NILSWMIMISGLAENGFGEEGLKLFSCMKREggQQYHAQLLKIGFDSSLSAGNALITMYAKCGVVEEARQVFRTMP------CLDSVSWNALIAALGQHGHGAEAVDVYEEMLKKGIRPDRITLLTVLTACSHAGLVDQGRKYFDSMEtyRIPpdHYARLIDLLCRSGKFSDAESVIESLP----FKPTAEIWEALLSGCRVHGNMELGIIAADKLF--GLIPEhgTYMLLSNMHAATGQWEEVARVRKLMRDRGVKK----------EVACSWIEMETQVHTFLVDDTSHPEAEAVYIYLQDLGKEMRRLGYVPDTSFVLH----------------    
 146 UniRef50_C5X9V8        Putative uncharacterized pr...   431    1e-119  1   9.6%   9:579    47:614     --------RAGRVPDAERLFAAMSRRSTSTYNAMLGGYAANGRLTLALSFFR------SIPRPDSFSYNTLLHALGVSSSLADARALFDEMPVKDSVSYNVmnGMLAAYIRNGRIQEARDLFDSRTEWDAISWNALMAGYVqqIEEAQKMFNRMPQR----DVVSWNTMVSGHarGDMAEARRLFDVAPIR----DVFTWTAVVSGYAQNGMLEEAKRVFDAMPEK----NAVSWNAMMAAYVQRRMMEEAKELFDAMPCR----NVASWNTMLTGYAQAGMLDEARAIFDMMPQKDAVSWAAMLAAYssEETLQLFKEMGrmYFKCGNMEEAHSAFEEMEER------DIVSWNTMIAGYARHGFGKEALEVFDTMRKTSTKPDDITLVGVLAACSHSGLVEKGISYFYSMHRdgvkPEHYTCMIDLLGRAGRLDEAVNLMKDMP----FEPDSTMWGALLGASRIHRNSELGRNAAEKIFE--LEpnAGMYVLLSNIYASSGKWRDVDKMRLMMHERGVKKVPGF----------SWIEVQNKVHTFSVGDSVHPEREDIYGFLEDLDIRMKKAGYV------------------------    
 147 UniRef50_Q60EM0        Os05g0455900 protein n=5 Ta...   429    1e-118  1  11.0%   5:587   140:792     ----SLVDMYMKCGSviEVFEGMPKKNVVTWTSLLTGCAHAQMHSEVMALFFRMRAEGIWPNP--FTFASVLSAVASQGALDLGQRVHAQSVKFGCRSSVFVCNSLMNMYAKCGLVEDAKSVFNWMerDMVSWNTLMAGLQLNEceALQLFHESRATMGKMTQSTYATVIKLCANLklALARQLHSCVLKHGFHLTGNVMTALADAYSKCGELADALNIF-SMTT--GSRNVVSWTAIISGCIQNGDIPLAVVLFSRMREDRVMPNEFTYSAMLKaySKFGSTEDALSIFKMIEQkgATYLFNKMAIQGIKPNEFTISSVIdmYSRKGNIDSAQIVFERQTDR------DLVSWNSMISGYAQHGYSMKAIETFRQMEASGIQMDGVTFLAVIMGCTHNGLVVEGQQYFDSMVrhKINpeHYACMVDLYSRAGKLDETMSLIRDMP----FPAGAMVWRTLLGACRVHKNVELGKFSADKLLS--LEpdSSTYVLLSNIYAAAGKWKERDEVRKLMDYRKVKK----------EAGCSWIQIKNKVHSFIAFDKSHPMSDQIYKKLKVIITRLKQDGYSPNTSFVLH----------------    
 148 UniRef50_Q9SV26        Pentatricopeptide repeat-co...   429    1e-118  1  10.1%   7:582    33:681     ------YGRCVSLGFANKLFDEMPKRDDLAWNEIVMVNLRSGNWEKAVELFREMQFSGA--KAYDSTMVKLLQVCSNKEGFAEGRQIHGYVLRLGLESNVSMCNSLIVMYSRNGKLELSRKVFNSMkrNLSSWNSILSSylGYVDDAIGLLDEMEICGLKPDIVTWNSLLSGYAsgHLKLGKAIHGYILRNQLWYDVYVETTLIDMYIKTGYLPYARMVFdrMEKEGIKPDAITWNSLASGYATLGKPEKALDVIGKMKEKGVAPNVVSWTAIFSGCSKNGNFRNALKVFIKMQEEggKEVHGFCLRKNLICDAYVATALVDMYGKSGDLQSAIEIFWGIKNK------SLASWNCMLMGYAMFGRGEEGIAAFSVMLEAGMEPDAITFTSVLSVCKNSGLVQEGWKYFDLMrrYGIIpeHCSCMVDLLGRSGYLDEAWDFIQTMSLK----PDATIWGAFLSSCKIHRDLELAEIAWKRLQV--LEpnSANYMMMINLYSNLNRWEDVERIRNLMRNNRVRVQD----------LWSWIQIDQTVHIFYAEGKTHPDEGDIYFELYKLVSEMKKSGYVPDT---------------------    
 149 UniRef50_B9N484        Predicted protein n=6 Tax=r...   429    1e-118  1  12.0%  10:582    77:578     ---------FSNLNFALKVFEYVPNPNVFVFNIIIKGCLQNNEPCKAICCYYKM--MIAHARPNKFTYPTLFKACTAAEAAEEGVQVHAHVIKQGLSGDVHIRSAGIQMYGSFGEVEGAR--------------------------------RMLGEDGNSDVICFNAMIDGYLKCgvEAAKELFWSMEDK----NVGSWNVMVSGMAKCGMIEEARELFNEMKEK----NEISWSAMIDGYIKGGYYKEALEVFNVMQREEIRPRKFVLSSVLAACANLGALDQGRWI------------HAYVNNNSNSFDAVLGTALVDMYAKCGRLDMAWDVFEKMEKKE------VFTWNAMICGLGMHGRAEDAIELFFKMQKQKFRPNGITLLGVLSACAHSGMVDEGLRIFNSMEeyGIEpeHYGCVVDLLGRAGLLGEAEEVMYSMP----MEPSAAVWGALLGACRKHGDVELGERVGKILLE--LEpnSGRYALLSNIYARAGRWDDVANVRKLMKERGVK----------TSTGISMIDFDGVVHEFKMGDGSHPQMKNIYLMLKNMIKRLKMEGFSPNT---------------------    
 150 UniRef50_Q9LVQ5        Pentatricopeptide repeat-co...   429    1e-118  1  10.0%  15:603   180:739     --------------FLKEMLKRKICPDVATFNILINVLCAEGSFEKSSYLMQKME--KSGYAPTIVTYNTVLHWYCKKGRFKAAIELLDHMKSKGVDADVCTYNMLIHDLCRSNRIAKGYLLLRDMRKRMIHPNEVTYNTLINGFsaSQLLNEMLSFGLSPNHVTFNALIDGhsEGNFKEALKMFYMMEAKGLTPSEVSYGVLLDGLCKNAEFDLARGFYMRMKRNGVCVGRITYTGMIDGLCKNGFLDEAVVLLNEMSKDGIDPDIVTYSALINGFCKVGRFKTAKEIVCR------------IYRVGLSPNGIIYSTLIYNCCRMGCLKEAIRIYEAMILEGH--TRDHFTFNVLVTSLCKAGKVAEAEEFMRCMTSDGILPNTVSFDCLINGYGNSGEGLKAFSVFDEMTKVGHHPttYGSLLKGLCKGGHLREAEKFLKSLHAVPAAV-DTVMYNTLLTAMCKSGNLAKAVSLFGEMVQRSILPDSYTYTSLISGLCRKGKTVIAILFAKEAEARGNVLPNKVMYTCFVDGMFKAGQW----------------KAGIY-----FREQMDNLGHTPDIVTT-NAMIDGYSRMGKIEKTN    
 151 UniRef50_Q8LQY2        Os01g0611900 protein n=3 Ta...   429    1e-118  1  10.2%   4:590   223:778     ---CKLRDANRALLVLRKMQDAGFVPWDFTFNSVVDVLVKGGRMEEALHIKDELLATGK--KMSVVLATTLMHGYCLQREVRKALDIFEETLRDGLVPTDVTYTVLIRGCTEEGMPEKAYELCRQMRDHGLLpnMVIKGLLNdwKDAVSLFKEMADSGI-PDAFTYNILIhlCQRRKIREALNLWEKMNETGVKPYIVTYHSLLLCYCVNGCMDEAVKLYTEMPGKGFTPNVVTYTTLMKGHINKAAFDKAYALLAEMKQNGVSCNDYTYNTLINGLCVVGRVCEVGEMLKRFETE------------GFVPTAMTYNSIINGFIKAGMMGSAFAVYQQMCAKG--IPPNIVTYTSFIDGYCKTSCCDLALKMLNDVRCKGLRPDIAAYNSLIFGFCQEGNMSHALQVLVLMLKDGLLPnvYNSFITGYKNLKMMEEALRLYEKMIKEGIDL-DTATYTTLIDGFSKDGNVTFALKLYSEMVAKGNIPDHITFTALTHGLCRNGDIDDARKLLDEMNR----------LDIRPNVLMYNMLING------------YLRNGKLQEAFRLHDEMLERKIMPDD-TTYDILV-------------    
 152 UniRef50_UPI0001983561 PREDICTED: hypothetical pro...   428    1e-118  1  10.7%   4:574 1039:1660     ---NSRISDCMRNgeAQMLFDEMPQRNTVTYNAMIRGYFQNGHFGEGVSLFDEM------PERDIFSYNTMIAGLMKFGDINGASEIFQKMP---FRDVVSWNSMISGYVSNGLIGEALRVFSGMVLKDVVSWNLVIAGlvGKVDLAEEFFKEMGTR----DIASWTTMISGLAsgRIVEARGLFEDMPVR----DVRAWNTMIAGYIENGCIEIGEVLFQKMPQRDFR----SWNEMINGLVRNQRIQDAMRLFVEMPQkglspdTVSWTNLIVGYFETGEVDTAVSIFELMPAreGLKFFVKMKEGGPFPDEATfnAMVTLYARCGNSNSALLLFSAMRSH------DFISWNSIICGLAHNGNGVEAIEVFEKMRSIDIKPNHITFVGVLSACSHAGLVDQGKYYFDFMKYKCctIEHYTCIVDLLGRFGLIDEAMSFLRQMEANGVEVP-ASVWGAVLGACRIHKNMQVGEIAGERILE--IEpnSGVYIILAEMYLSSGKREDAERVWVRMREKGVKKQPA----------CSWMEVNGSGHVFLSGDSSHPQFSRVCGVLGLLHMEME-----------------------------    
 153 UniRef50_D1I316        Whole genome shotgun sequen...   428    1e-118  1  10.1%   3:589   128:684     --SNAFVTMYMKTQSVeqFFKAMMIENLASRNNLLSGFCDTETCDQGPRIL--IQLLVEGFEPNMYTFISILKTCASKGDLNEGKAIHGQVIKSGINPDSHLWNSLVNVYAKCGSANYACKVfeIPERDVVSWTALITGFVAEggSGLRIFNQMLAEGFNPNMYTFISILRSCSssDVDLGKQVHAQIVKNSLDGNDFVGTALVDMYAKNRFLEDAETIFNRLIKR----DLFAWTVIVAGYAQDGQGEKAVKCFIQMQREGVKPNEFTLASSLSGCSRIATLDSGRQL------------HSMAIKAGQSGDMFVASALVDMYAKCGCVEDAEVVFDGLVSR------DTVSWNTIICGYSQHGQGGKALKAFEAMLDEGTVPDEVTFIGVLSACSHMGLIEEGKKHFNSLSkyGITpeHYACMVDILGRAGKFHEVESFIEEMK----LTSNVLIWETVLGACKMHGNIEFGERAAMKLFELEPEIDS-NYILLSNMFAAKGMWDDVTNVRALMSTRGVKKEP----------GCSWVEVNGQVHVFLSHDGSHPKIREIHLKLQDLHQKLMSVGYTPNTDHVLHNV--------------    
 154 UniRef50_Q9LUJ2        Pentatricopeptide repeat-co...   428    1e-118  1   9.1%   7:591   179:761     ------YAECGELDSARKVFDEMSERNVVSWTSMICGYARRDFAKDAVDLFFRMVR-DEEVTPNSVTMVCVISACAKLEDLETGEKVYAFIRNSGIEVNDLMVSALVDMYMKCNAIDVAKRLFDEyaSNLDLCNAMASNYVRQgrEALGVFNLMMDSGVRPDRISMLSAISSCSQLriLWGKSCHGYVLRNGFESWDNICNALIDMYMKCHRQDTAFRIFDRMSNK----TVVTWNSIVAGYVENGEVDAAWETFETMPEK----NIVSWNTIISGLVQGSLFEEAIEVFCSMQSQEgkWIYYYIEKNGIQLDVRLGTTLVDMFSRCGDPESAMSIFNSLTNR------DVSAWTAAIGAMAMAGNAERAIELFDDMIEQGLKPDGVAFVGALTACSHGGLVQQGKEIFYSMLkhGVSPehYGCMVDLLGRAGLLEEAVQLIEDMP----MEPNDVIWNSLLAACRVQGNVEMAAYAAEKIQVLAPE-RTGSYVLLSNVYASAGRWNDMAKVRLSMKEKGLRKPPGT----------SSIQIRGKTHEFTSGDESHPEMPNIEAMLDEVSQRASHLGHVPDLS---NVLMD------------    
 155 UniRef50_D1HPF6        Whole genome shotgun sequen...   428    1e-118  1  10.1%  21:574     4:526     --------------------SENSKPNLVSWSAVIGGFAQNGYDKEALELLCRMQAAGF--EPNARTLASVLPACARLQNLNLGKEIHGYVTRHGFMSNPFVVNGLVDVYRRCADMGSALKIFSGfvKNVVSYNTMIVGYcgNVEKAKELFDQMELVGK--DTISWNSMISGYADndLKAAQLAFDGVTER----DTATWNVLISGYACCNQLENIQNLIQKMKGDGFEPNVYTWNGIISGHVENGHNELALRLFTEMQTSSLRPDIYTVGIILPACARLATIARGKQV------------HAHSIRQGYELDVHIGAALVDMYAKCGSIKHAMQVYNRISN------PNLVSQNAMLTAYAMHGHGDEGIALFRNMLGNGFRPDHVTFLSVLSSCVHAGAVETGHEFFDLMTYYNVTPshYTCIVDLLSRAGRLDEAYELVKKIPRK----PDSVMWGALLGGCVIWGNVELGEIAAESLIE--LEpnTGNYVLLANLYAYAGRWHDLDRTRQMIKDRGMHKSP----------GCSWIEDREDIHVFLSCDKSHEKTEDIYTTLDNLNTHMR-----------------------------    
 156 UniRef50_B9I5M1        Predicted protein n=5 Tax=r...   427    1e-118  1  11.9%  46:599    89:631     ---------------------------------------------HPRYALNTYTYMRkdIEVDSFIIPSVLKACSQISVARMGKEIHGFSVKNGLVSDVFVVNALMQMYSECGSLVSARLLFDKmeRDVVSWSTMIRAycNDLEEGERLFVRMIEENVFPNDITMLSLIISCGfgAVQLGKRLHAYILRNGFGMSLALATALVDMYGKCGEIRSARAIFDSMKNK----DVMTWTAMISAYAQANCIDYAFQLFVQMRDNGVRPNELTMVSLLSLCAVNGALDMGKWF------------HAYIDKQGVEVDVILKTALIDMYAKCGDISGAQRLFSEAIDR------DICTWNVMMAGYGMHGYGEKALKLFTEMETLGVKPNDITFIGALHACSHAGLVVEGKGLFEKMidFGLVpeHYGCMVDLLGRAGLLDEAYKMIESMP----VTPNIAIWGAMLAACKIHKNSNMGELAARELLA--LEPQNCGyvLMSNIYAAANRWNDVAGMRKAVKDTGIKKEP----------GMSSIEVNGLVHDFKMGDTAHPLIEKISEMLAEMSKKLKEAGYLPDT----SVVLHNIDEEEKE----    
 157 UniRef50_C5Y173        Putative uncharacterized pr...   427    1e-118  1  11.9%  25:585   116:665     ------------------------PPAVITCNILIKKLCARRRLADAERVLE---ALKTSGAADAVSHNTLVAGYCRDGSLWDAERVLEAARASGAANVVTYTALIDGYCRSGRLTDALRLIASMPvpDTYTYNTVLKGLckQWEEAEELMREMIRNNCHPNEVTFATQIRAFCQNglDRAVELLEQMPKYGCMPDVVIYSTLVNGFSEHGRVDDALKLLSTML---CRPNTVCYNAALKGLCIAGRWDEVGELIAEMVRKDCPPNDATFSTLINSLCQNRLAEYAIEVLEQMQ------------KYGYMPDVVSYNTIISCFSEQACADDALKLLNSML-----CKPDTISFNAVLKCLCRAERWYDAAELMAKMLKEDCHTNEMTFNILIDSLCQNGQVKDAIEMFELMPKYRCTPdtYSSLINGFSEQGLDKVAFDLFRSMP----CRADIFSYNATLKGLCMAARWDDAGELIADMVTKDCLPNEVTFNILINSLCQKGLVNRAIEVYEQMPKYGITPDIFT-YNALINGYSEQGCLDDALKFlyNSILKGLCRAERWKDAEKLVTEMLRKNCTPNEVTF------------------    
 158 UniRef50_Q9LSB8        Putative pentatricopeptide ...   427    1e-118  1  10.3%  11:582    82:653     ----------GHVSYAYKLFVKIPEPDVVVWNNMIKGWSKVDCDGEGVRLYLNM--LKEGVTPDSHTFPFLLNGLKrgGALACGKKLHCHVVKFGLGSNLYVQNALVKMYSLCGLMDMARGVFdrCKEDVFSWNLMISGymKEYEESIELLVEMERNLVSPTSVTLLLVLSACSKVKDKdcKRVHEYVSECKTEPSLRLENALVNAYAACGEMDIAVRIFRSMKARDvvRDRISWTIMIDGYLRAGCFNESLEIFREMQSAGMIPDEFTMVSVLTACAHLGSLEIGEWI------------KTYIDKNKIKNDVVVGNALIDMYFKCGCSEKAQKVFHDMDQR------DKFTWTAMVVGLANNGQGQEAIKVFFQMQDMSIQPDDITYLGVLSACNHSGMVDQARKFFAKMrdHRIEPshYGCMVDMLGRAGLVKEAYEILRKMP----MNPNSIVWGALLGASRLHNDEPMAELAAKKILE--LEPDNgvYALLCNIYAGCKRWKDLREVRRKIVDVAIKKTPGF----------SLIEVNGFAHEFVAGDKSHLQSEEIYMKLEELAQESTFAAYLPDT---------------------    
 159 UniRef50_Q94LP5        Pentatricopeptide, putative...   427    1e-118  1   9.2%   7:582   337:880     ------YAKANSLTDARRAFFGLPNHTVETSNAMMVGLVRAGLGIEAMGLFQFM--IRSSIRFDVVSLSGVFSACAETKGYFQGQQVHCLAIKSGFDVDICVNNAVLDLYGKCKALMEAYLIFQGMkkDSVSWNAIIAALEQNgdDTILHFNEMLRFGMKPDDFTYGSVLKACAALrlEYGLMVHDKVIKSGLGSDAFVASTVVDMYCKCGIIDEAQKLHDRIGGQQ----VVSWNAILSGFSLNKESEEAQKFFSEMLDMGLKPDHFTFATVLDTCANLATIELGKQIHGQIIKQEMLD------------DEYISSTLVDMYAKCGDMPDSLLVFEKVEKR------DFVSWNAMICGYALHGLGVEALRMFERMQKENVVPNHATFVAVLRACSHVGLFDDGCRYFHLMthYKLEpeHFACMVDILGRSKGPQEAVKFINSMP----FQADAVIWKTLLSICKIRQDVEIAELAASNVLLLDPD-DSSVYILLSNVYAESGKWADVSRTRRLLKQGRLKKEP----------GCSWIEVQSEMHGFLVGDKAHPRSGELYEMLNDLIGEMKLSGYEPDS---------------------    
 160 UniRef50_UPI0001985476 PREDICTED: hypothetical pro...   427    1e-118  1  10.6%   7:579   128:679     ------YAKCGTMETARKVFDELPRRNVVSWTTLMTGYVHDSKPELAVQVFREMLEAGAY--PTNYTLGTALSASSDLHSKELGKQIHGYSIKYRIEFDASIGNSLCSLYSKCGSLECAVKAFRRIrkNVISWTTVISAWGDNGEAalQFFVEMLSECVEPNEFTLTSALSLCCVmsLDIGTQIHSLTIKLGFESNLPIKNSIMYLYLKCGWIHEAKKLFDEME----TISLVTWNAMIAGHARMMDFaeALSIFLKLNRSGMKPDLFTFSSVLSVCSSLVALEQGEQV------------HAQTIKTGFLSDVVVGTALVNMYNKCGSIERASKAFVEMSIR------TLISWTSMITGYAQNGQPQQALLLFEDMRLAGVRPNKITFVGVLSACSHAGMVDEALDYFQMMkeYKITpdHYACLIDMFVRLGRLDEAFDFIKEM----DLEPNEFIWSILIAGCRSQGKLELGFYAAEQLL--NLKpdTETYNLLLNMYLSAGKWKEVSRVRKMMKEEKLGRLKD----------WSWISIKDKIYSFKRNARSHAQSGEMYELLGNLHEKAKSFGYE------------------------    
 161 UniRef50_Q9LUC2        Pentatricopeptide repeat-co...   427    1e-118  1  12.7%   5:573   101:643     ----SLVNMYAKCGLMRRAVLvgSERDVFGYNALISGFVVNGSPLDAMETYREMRANGIL--PDKYTFPSLLKG-SDAMELSDVKKVHGLAFKLGFDSDCYVGSGLVTSYSKFMSVEDAQKVFDElrDDSVLWNALVNGYSqrFEDALLVFSKMREEGVGVSRHTITSVLSAfvSGDIDNGRSIHGLAVKTGSGSDIVVSNALIDMYGKSKWLEEANSIFEAMDER----DLFTWNSVLCVHDYCGDHDGTLALFERMLCSGIRPDIVTLTTVLPTCGRLASLRQGREIHGYMIVSGLLNRKS--------SNEFIHNSLMDMYVKCGDLRDARMVFDSMRVK------DSASWNIMINGYGVQSCGELALDMFSCMCRAGVKPDEITFVGLLQACSHSGFLNEGRNFLAQMEtyNILpdHYACVIDMLGRADKLEEAYEL----AISKPICDNPVVWRSILSSCRLHGNKDLALVAGKRLHE--LEPEhgGYVLMSNVYVEAGKYEEVLDVRDAMRQQNVKKTP----------GCSWIVLKNGVHTFFTGNQTHPEFKSIHDWLSLVISHM------------------------------    
 162 UniRef50_D1IZB5        Whole genome shotgun sequen...   427    1e-118  1  10.0%   4:581    48:631     ---LSNLSKCGRVDDARKLFDVMPDRDECSWNTMIGAYANSGRLNEAreALELFWEMqeGERPNQFTWGSVLRVCSMYVLLEKGKQIHAHAIKTQFDSNAFVVTGLVDMYAKCKCILEAEYLFELakRNHVLWTAMVTGYSQNGdkAIECFRDMRGEGIECNQFTFPSILTACGssACGFGAQVHGCIVRSGFGANVFVGSALVDMYSKCGDLSNARRMLETMEVD----DPVSWNSMIVGCVRQGLGEEALSLFRIMHLRHMKIDEFTynALVDMYAKRGYFDYAFDVFEKMTDK----------------DVISWTSLVTGCVHNGSYEEALRLFCEM--RIMGIHPDQIVinSLVSMYAKCGCIEDANKVFDSMEIQ----DVITWTALIVGYAQNGR---GRDYFQSMEeyGIkpEHYACMIDLLGRSGKLMEAKELLNQM----AVQPDATVWKALLAACRVHGNVELGERAANNLFE--LEpnAVPYVLLSNLYSAAGKWEEAAKTRRLMKLRGVSKEP----------GCSWIEMSSKVHRFMSEDRSHPRTAEIYSKVDEIMILIKEAGYVPD----------------------    
 163 UniRef50_D1IXX6        Whole genome shotgun sequen...   427    1e-118  1  11.2%   1:578    80:587     LCAQSL---PIDPRYALSLLAQLRTPNLPLYNAIIRGLasNNDSIEGLVVYKQM--LSKGIVPDNYTIPFVLKACAESRAVREGEEVHGQAIKMGLASDVYVSNTLMRMYAVCDVIRSARKVFDTSPQR--------------------------------DLVSWTTMIQGYVKMGfrEGVGLY-IIRNSNVNLDVFVGNALVDMYLKCGDANFARKVFQEMPVK----NVVSWNSMISGLAQKGQFKESLYMFRKMQRLGVKPDDVTLVAVLNSCANLGVLELGKWV------------HAYLDRNQIRADGFIGNALVDMYAKCGSIDQACWVFQAMNRK------DVYSYTAMIVGLAMHGQGGKALDLFSEMPKMGIEPDEVTFVGVLTACSHVGLVEEGRKYFEDMStyNLRpeHYGCMVDLLGRAGLINEAEEFIRNMP----IEPDAFVLGALLGACKIHGKVELGESVMKKIEK--IEPrdGAYVLMSNIYSSANRWRDALKLRKTMKERNLEKTP----------GCSSIELDGVIHEFQKGDKSHPKIKEIYKLLDEIMSHLKNNEC-------------------------    
 164 UniRef50_Q6ZIP5        Pentatricopeptide (PPR) rep...   427    1e-118  1  11.0%   1:589    95:728     VAATSLVAAHAAAGRLRDafDAVprRDTVLHNAMMSAFARASLAAPAVSVFHALLGSGS-LRPDDYSFTALISAVGQMHNLAAPhqLHCSVLKSGAAAVLSVSNALIALYMKCDTPEaaRKVLDEMpkFDVVWNAMISGYVQSgaDAFELFRRMVSEKVPLDEFTFTSVLSACANAGFfhGKSVHGQIIRlnFVpdVVSWNTILSGYIDSGCLDKAVEVFKVMPYK----NDLSWMVMVSGYVHGGLSEDALKLFNQMRAEDVKPCDYTYAGAIAACGELGALKHGRQL------------HAHLVQCGFEASNSAGNALLTMYAKCGAVNDARLVFLVMPN------LDSVSWNAMISALGQHGHGREALELFDQMVAEGIDPDRISFLTILTACNHAGLVDEGFHYFESMKRdgISpdHYARLIDLLGRSGRIGEARDLIKTMP----FEPTPSIWEAILSGCRTNGDMEFGAYAADQLFRMIPQHD-GTYILLSNTYSAAGRWVDAARVRKLMRDRGVKKEP----------GCSWIEVGSKIHVFLVGDTKHPEAQEVYQFLEVIGARMRKLGYVPDTKFVLHDM--------------    
 165 UniRef50_Q9FWA6        Pentatricopeptide repeat-co...   427    1e-118  1   9.1%   6:574   325:881     -----MYAKCDNMQDAQILFDNSENLNRQSYNAMITGYSQEEHGFKALLLFHRL--MSSGLGFDEISLSGVFRACALVKGLSEGLQIYGLAIKSSLSLDVCVANAAIDMYGKCQALAEAFRVFDEMRrdAVSWNAIIAAHEQNGkeTLFLFVSMLRSRIEPDEFTFGSILKACTGGSLGYGMEIHSSIVKSGMASNSSVGCSLIDMYSKCGMIEEAEKilEKMHNKRLQEMCVSWNSIISGYVMKEQSEDAQMLFTRMMEMGITPDKFTYATVLDTCANLASAGLGKQIHAQVIKKE------------LQSDVYICSTLVDMYSKCGDLHDSRLMFEK------SLRRDFVTWNAMICGYAHHGKGEEAIQLFERMILENIKPNHVTFISILRACAHMGLIDKGLEYFYMMKryGLDPqhYSNMVDILGKSGKVKRALELIREMP----FEADDVIWRTLLGVCTIhnNVEVAEEATAALLRLDPQ-DSSAYTLLSNVYADAGMWEKVSDLRRNM--RGFKLKKEP--------GCSWVELKDELHVFLVGDKAHPRWEEIYEELGLIYSEMK-----------------------------    
 166 UniRef50_Q0DWA2        Os02g0824000 protein n=5 Ta...   426    1e-117  1  12.2%   2:599   178:850     -GYCRVGDTPAALSVADRMTAQGLPMDVVGYNTLVAGFCRAGQVDAARGVLDMMKEAG--VDPNVATYTPFIVYYCRTKGVEEAFDLYEGMVRNGVLLDVVTLSALVAGLCRDGRFSEAYALFREMDKVGAAPNHVTyaKAGRGKELLSLLGEMVSRGVVMDLVTYTALMDWLGkgKTDEVKDTLRFALSDNLSPNGVTYTVLIDALCKAHNVDEAEQVLLEMEEKSISPNVVTFSSVINGFVKRGLLDKATEYKRMMKERGINPNVVTYGTLIDGFFKFQGQDAALEVYHDMLCEGVkaLFKDASGSGLSLDHVNYTTLIDGLFKAGDMPTAFKFGQELMDRN--MLPDAVVYNVFINCLCMLGKFKEAKSFLTEMRNMGLKPDQSTYNTMIVSHCRKGETAKALKLLHEMKMSSIKPntYNTLVAGLFGTGAVEKAKYLLNEMVSAGflHADITVYNTLLQVLCYHGMTRKATVVLEEMLGSGIAPDTITFNALILGHCKSSHLDNAFATYAQMLHQNISPNIAT-FNTLLGGLESVGRIGEAGTVLIEMEKSGLEPNNLtvEAMRLYCEMVGKGFVP-KVSTYNALISDFTKAGMM----    
 167 UniRef50_Q2QXL4        Os12g0152600 protein n=2 Ta...   426    1e-117  1  11.4%  48:593   181:697     -----------------------------------------------LAAFREMAHHGVPPFIKE--CNLVLRALRDEARWDDMRSVYAEMLQLGIEPSIVTYNTLLDSFFREGRVDQAAKLLREMepSDVTYNVVINGLARKGekAAQLVDRMRMSK-KASAFTFNPLITGyaRGSVEKAGALQLEMENEGIVPTVVTYNTIIHGMFRSGNVEAARMKFVEMRAMGLLPDLITYNSLINGYCKAGNLKEALWLFGDLKRAGLAPSVLTYNILLDGYCRLGDLEEARRFKQEMVEQ------------GCQPDVSTYTILMNGSRKVRNLAMVREFFDEMLSKG--LQPDCFAYNTRISAELILGSTSEAFQLTEVMISRGISSDTVTYNIFLDGLCKSGNLKDAYVLWMKMVSDGLQPdtYTCLIHAHCERGRLREARDIFDGMLVSGL-PPSAVTYTVFIHAYCRRGNLYSAYGWFQKMLEEGVRPNEVTYNVLIHALCRMGRTNLAYQHFHEMLERGLSP----------NKYTYTLLIDGNCKEGN------------WEEAIRLYSEMHQHGIHPDH-CTHNALFKGF----------    
 168 UniRef50_Q3EDF8        Pentatricopeptide repeat-co...   426    1e-117  1  15.8%  27:526   102:590     --------------------------DVESNNHL-RQMVRTGELEEGFKFLENMVYHGNV--PDIIPCTTLIRGFCRLGKTRKAAKILEILEGSGAVPDVITYNVMISGYCKAGEINNALSVLDRMspDVVTYNTILRSLcgKLKQAMEVLDRMLQRDCYPDVITYTILIEacRDSGVGHAMKLLDEMRDRGCTPDVVTYNVLVNGICKEGRLDEAIKFLNDMPSSGCQPNVITHNIILRSMCSTGRWMDAEKLLADMLRKGFSPSVVTFNILINFLCRKGLLGRAIDILEKMPQ------------HGCQPNSLSYNPLLHGFCKEKKMDRAIEYLERMVSRGC--YPDIVTYNTMLTALCKDGKVEDAVEILNQLSSKGCSPVLITYNTVIDGLAKAGKTGKAIKLLDEMRAKDLKPDtySSLVGGLSREGKVDEAIKFFHEFERMGI-RPNAVTFNSIMLGLCKSRQTDRAIDFLVFMINRGCKPNETSYTILIEGLAYEGMAKEALELLNELCNKGLMKKSS-----------------------------------------------------------------------------    
 169 UniRef50_B9H9E4        Predicted protein n=1 Tax=P...   426    1e-117  1   9.8%   4:574   175:774     ---MGLYSKCGKLKEVCQLFEKMTHRDVISWNTMISCYVLKGMYREALDLFDEM--LVSGVLPDEITMVSLVSTCAKLKDLEMGKRLHLYIVDNKLWIrsLLNCLVDMYSKCGKMDEaaRQLFDKmeRSLVSWTTMMSGYVQGgcESLELFQQMRFENVIPDEVALVTVLSACVHldFDLGRSVHAFIVTYGMLVDGFLGNALLDLYAKCGKLDEALRTFEQLPCK----SAASWNSMLDGFCRSGGVDKARDFFNKIPEK----DIVSWNTMVNAYVKHDLFNESFEIFCKMQSSNVkwVNVYIEKNEIGIDAMLGTALIDMYGKCGCVEMAYEIFTQIIEK------NVFVWTAMMAAYAMEGQALEAIDLYLEMEERGVKPDHVTFIALLAACSHGGLVDEGYKYFNKLRsyNIIPthYGCMVDLLGRVGHLEETVKFIERMP----IEPDVSIWSSLMRACRSHHNVELAEQAFKQLIE--IDpnNGAHVLLSNIYADAGRWDDVSKVRTKLHETGVPKQPGFTM----------IEQNGVVHEFVA---SNLVSADILCMLQDIERRLL-----------------------------    
 170 UniRef50_O23169        Pentatricopeptide repeat-co...   426    1e-117  1   9.8%  21:580    34:602     --------------------KKFFNPNHEDGGVVVERLCRANRFGEAIDVlkLLRelGRAKKPPASTYCNLIQVCSQTRALEEGKKVHEHIRTSGFVPGIVIWNRLLRMYAKCGSLVDARKVFDEMprDLCSWNVMVNGYagLLEEARKLFDEMTEK----DSYSWTAMVTGYVKKDQPegKEIHGHIVRAGLDSDEVLWSSLMDMYGKCGCIDEARNIFDKIVEK----DVVSWTSMIDRYFKSSRWREGFSLFSELVGSCERPNEYTFAGVLNACA------------DLTTEELGKQVHGYMTRVGFDPYSFASSSLVDMYTKCGNIESAKHVVD------GCPKPDLVSWTSLIGGCAQNGQPDEALKYFDLLLKSGTKPDHVTFVNVLSACTHAGLVEKGLEFFYSITEKhhTSDHYTCLVDLLARSGRFEQLKSVISEMPMK----PSKFLWASVLGGCSTYGNIDLAEEAAQELFK--IEpnPVTYVTMANIYAAAGKWEEEGKMRKRMQEIGVTKRPGS----------SWTEIKRKRHVFIAADTSHPMYNQIVEFLRELRKKMKEEGYVP-----------------------    
 171 UniRef50_Q9CA58        Putative pentatricopeptide ...   425    1e-117  1  10.9%   5:599    45:653     ----SVIEKLGYYGKFEAMEEVLVDMRENVGNHMLEgyGRKGKVQEAVNVFERMDFYDC--EPTVFSYNAIMSVLVDSGYFDQAHKVYMRMRDRGITPDVYSFTIRMKSFCKTSRPHAALRLLNNMSSQGCEMNVVAYCTVVGgaEGYELFGKMLASGVSLCLSTFNKLLRVLCkgDVKECEKLLDKVIKRGVLPNLFTYNLFIQGLCQRGELDGAVRMVGCLIEQGPKPDVITYNNLIYGLCKNSKFQEAEVYLGKMVNEGLEPDSYTYNTLIAGYCKGGMVQLAERI------------VGDAVFNGFVPDQFTYRSLIDGLCHEGETNRALALFNEALGKG--IKPNVILYNTLIKGLSNQGMILEAAQLANEMSEKGLIPEVQTFNILVNGLCKMGCVSDADGLVKVMISKGYFPdtFNILIHGYSTQLKMENALEILDVMLDNG-VDPDVYTYNSLLNGLCKTSKFEDVMETYKTMVEKGCAPNLFTFNILLESLCRYRKLDEALGLLEEMKNKSVNPDAVT-FGTLIDGFCKNGDLDGAYTLfyNIIIHAFTEKLNVTMAEKLFQEMVDRCLGPD-GYTYRLMVDGFCKTGNV----    
 172 UniRef50_B9GFI1        Predicted protein n=1 Tax=P...   425    1e-117  1  10.7%  23:581    42:578     ----------------------FDRTDVYSWNSLIAELARGGDSCESLRAFSWMRKL--DIKPNRSTFPCAIKSCSALFDLNSGKQAHQQALVFGFESDLFVSSALIDMYSKCGKLSNARVLFDEirRNIVTWTSLITGYVQnaHEALMVFKEfeEVGTSVDSVAMISVLSACSRVSNKAveGVHGVAIKVGLDKVMGVENTLLDAYAKCGEVSLSRKVFDDMAEK----DVVSWNSMIAVYAQNGLSTDAFEVFHGMLKAGgkYNEVTLSTLLLACAHEGAL------------RVGMCLHDQVIKMGYVNNVIMATSIIDMYCKCGQAEMARNAFDGMKEK------NVRSWTAMIAGYGMHGFAREALDVFYQMIWAGVKPNYITFISVLAACSHAGFLEEGWRWFNAmhEYNVEpeHYGCMVDLLGRAGYIKEAYNLIKSMKVR----RDFVLWGSLLAACRIHKDVELAEISARELFKLDPS-NCGYYVLLANIYADAGRWKDVERMRILVKDRGLVKPPGY----------SLVELKGRVHVFLVGDKEHPQHEKIYKYLEELSVKLQEAGYVPN----------------------    
 173 UniRef50_B9MYZ7        Predicted protein n=5 Tax=r...   425    1e-117  1  11.7%   4:593   188:748     ---NSLLSMyaDMECARELFDEMHEKDVIAWSVMIGGYLQ---WEEPQVGLQMFRKMVLVpePDGVVMVSVLKACASSRDVCTGRLVHGLVIHRGFDCDLFVENSLIDMYSKCKDAGSAFKVFNEiqRNNVSWNSMLSGFVlnYSEAQSLISSMRKERVETDEVTLVNILQICkfVHPFHCKSIHCVMIRRGSEANELVLSALIDAYAKCYLIEIAWEVFARMRRR----DVVSWSTMISGFAHCGKPDEAIAVYQEMDRDLVKPNVITIINLLEACSVTAEL------------KRSKWAHGVAIRQGFASEVTVGTAVVDMYSKCGEILASRRAFDQLALK------NIVTWSAMIAAYGMNGLAHEALALFAEMKRHGLKPNPVTTLSVLAACSHGGLVEEGLSLFKSMveLGLEpeHYSCMVDMLGRAGKLDTAIEVIKAMP--HNLKNGASIWGSLLSACRSYGLTELGKEAISRVLE--LEpnSAGYLVASSMYAADGLWDDAARIRVLAKEKGVKVVAGYSLVH----------IDNKACRFVAGDGSHPRSDEIFSMAQQLHDCIKID--EKKEGNTWLAVIECL----------    
 174 UniRef50_D1HDF8        Whole genome shotgun sequen...   425    1e-117  1  10.3%   7:580   269:831     ------YGKCGNLQHASQLFERISRKDVVSWNAMIAANEQRGEGENALGLFRRMLKVEPPVQPNRVTFLSLLSAVSGLSALRCGREIHAHIFRLSLEVDTSITNSLITFYSKCREVGKAREIFERLlrDIISWNSMLAGYEQNEqrCFDIFKRMMLSGIEPDSHSLTIIFNAASRDsfRRGKEIHGYILRrgVSLSVS--NAILKMYAKFNRIADAEKIFKGMKNR----DSYSWNAMMDGYSRNAKFEDVLMIFLDILKQGFPLDHVSLSILLTSCGRLVSLQLGKQFHA-VVAKLFNGQDCPHQDSLLSIN----NALISMYSKCGSIKDAAQVFLKMERK------DVFSWTAMITGCAHHGLAVEALQLFERMKTDGIKPNQVTFLALLMACAHGGLVQEGSYYFDSmnDYGLSpeHYACMIDLFGRSGQFDRAKSLVEFGITLFKPYHDDIlwKVLLGACHASKQLDLGVEAATKILE--LEpdEATYILLANLYASSGLWEDAIKVRKAMRDKGLRK----------EVGCSWIDTGNRRHVFVAGDVYHPQRKEIYEKLAQLNYSCRRMGYVP-----------------------    
 175 UniRef50_Q0JQL2        Os01g0153200 protein n=5 Ta...   424    1e-117  1  10.2%   5:602   177:811     ----RQFALARDLFD--KMLQSGVHLDEYVYTAGIRAYCESRNLDGARGLVVRMESEG--VKASAVPYNVLMYGLCKNMRVQEAVEVKNVMVNIGVTADEVTYRTLVYGFCRMEELEMALRITHDMIRLGFVPSEANCSFMIdvEEAFSLACKLGDLGMVPNVFAYNALIDKLCKnrFDDADRLFKEMAGRGLEPNEVTYAILIHALCKRGMIEDALCLFDKMRDKGIKVTVYPYNSLINGYCKQGSLDRARGLLSGMVKEGLTPTAASYSPLIAGLCRNGDLSSCMELHREMAER------------GIAWNNYTFTALINGFCKDKKMDEAARLFDKMID--SNVIPNEVTFNVMIEGYCLVGNIRKAFQLYDQMVEMGLKPDNYTYRSLISGLcrEGRFTETYHLWDEMAVRGVKLdsFTIIVYAALKQHDKEKSCVLFREMKEQG-VKPDDIFYTCMIDALSKEENMIQALNCWDQMVVDGYSPNTVTHTVLINNLCKSGYLGSAELLCKEMLA-GNVLPNKFTYNCFLDYFATEGDMEKAKDLHSAMLQGhlCKAGKIQEAIDLMSKITESGFSPDCI-SYSTIIHELCKMGDINKA-    
 176 UniRef50_D1J070        Whole genome shotgun sequen...   424    1e-117  1  11.4%  11:588    52:598     ----------KDLGYTLLLFDRLATPYIFLWNTIVRSFSASSQPQMVLVAYSRLRNHGVI--PDRHTFPLLLKAFSKLRNENP-FQFYAHIVKFGLDFDAFVQNSLVSAFAHCGYVDCSRRLfeTAKKDVVSWTALINGCLRNGreALECFVEMRSSGVEVDEVTVVSVLCAAAMLrvWFGRWVHGFYVESgvIWDVYVGSALVDMYSKCGYCDDAVKVFNEMPTR----NLVSWGALIAGYVQCNRYKEALKVFQEMIIEGIEPNQSTVTSALTACAQLGSLDQGRWL------------HEYVDRSKLGLNSKLGTALVDMYSKCGCVDEALLVFEKLPAK------DVYPWTAMINGLAMRGDALSSLNLFSQMIRSRVQPNGVTFLGVLSACAHGGLVDEGLELFRLMidYRLEpdHYGCMVDLLGRAGRLEEAIKFIESMP----MEPTPGVWGALFSGCMIHKAFELGEHIGNHLIK--LQPHHSgyILLANLYSRCQKWEAAANVRRLMKGKGVDKSPA----------CSWIEVNGVIHEFIAFDKSHIESINVYMMLGSVTAQLKLATYALDslLTFTI---------------    
 177 UniRef50_B9S079        Pentatricopeptide repeat-co...   424    1e-117  1  10.6%  28:564   102:606     ---------------------------VYIFNAIIQSLSTSNnfTHIFSLYRQMLLIGLS--PDTYTLPYLLKACSQSHAFIEALQIHAHSIKTGLSSNLFVKNTLMRFYAVSGFIEAVEKVFDqpHWDLISWTTLiySKMGYPSEAIAAFFRM---NCTADRMTLVVVLSACSqgDFTLGKKILAYMDHhdVHSDVFLGNALLDMYLKCGQPHLARQLFHLMPVK----NLVSWNSMISGLAHQGLFKEALHMFRRMQTMGLKPDSVTLVGVLNSCANLGDLELGKWV------------HSYIDKNHMKADGYVANALVDMYAKCGSIDQAFMVFQAMKCK------DVYSYTAMIVGFAMHGKADRALAIFSEMPRMGVRPDHVTLVGVLSACSHAGLLEEGRRHFQDMSryHLQpeHYGCMVDLLGRAGLISEAEAFTNKMP----IVPDASVWGSLLGACKIHAKVELGETVIQKLIEMEPERD-GAYILMSNIYSSANRWRDALKWRKAMKQNNIKKTP----------GCSSIEVDGMVHEFRKGEKSHPKSREMIS---------------------------------------    
 178 UniRef50_D1IYW7        Whole genome shotgun sequen...   424    1e-117  1  13.9%   6:595    93:728     -----LIDAYCklEDALYMYDEmlVPDVVTYTCIMNGLCKSGKVEEAKSVFREMEE--VGVVPNRFSYATLIDSLFKEGNVAEAFVLQGRMVVRGIGFDVVVYTALMDGLFKAGMANNAEDMFQVLLEESLVPNCVTYSALIDGhkGELLLQEMEEKHIFPNVIVYSSIVDGYTKKGlnEAMDVMRKMVQRNILPNVFVYGTLIDGYFKADQRGIALDLFKEMKSRGLEENNFVIDSFVNNLKRSGRMEEADELFKDMMSRGLLPDRVNYTSMMDGFFKAGKESDAFNIAQEMTEkeSESFHTGMRQLGLAPDSATFNTMINAYCKEGNLGNALKLLNEMKSYG--LKPNSITCNILVQRLCAAGEIEKTMDLLNDMLVMGFHPTPTTHKAVLDASSKSRRADVILHMHDQLVGMGVKLdtYNTLISTFCRLGMIRRATLVFKDMMGKGILA-DIITYNALIHGYCISSHLKKAFAVHSQMLTEGVSPNVETYNILLGGLSAARLIKEAAGLVNQMKERGL-VPNATTYDILVSGHGKIGNMKECVKLyyNVLISCFAKGKKMSQAKELMQEMQVRGIPPNS-STYDILICGWYK--------    
 179 UniRef50_D1HJA6        Whole genome shotgun sequen...   424    1e-117  1  10.2%   5:581    39:582     ----SLIAQYASLGSIShlFSTSHSSNVFLWNHIIRAFSNTPHSRNSLRLYSRMLQLG--IQATNFTFPFLLKACGCLADFELGARAHAHVVVFGYESDVFVANSLMAMYGRFGCFDFSRQVFERmeRNVVSWSSMVGAYAHNgeEGLLLFWRMLNEGIAPNrsIVNAM--ACIH-REHEADDFCRVVIDNGLDSDQSVQNAAMGMYARCGRIDVARRFFYGILDK----DLVAWTSMIEAYVQADLPINALELFKQMKLLGIVPDSVTLLSLIHAVSNLASFQLARFVHGVITRS--------FFKNHIALD----TAVIDLYVKCGNLEYARKCFDRMSAR------NLISWSTMISGYGMHGHGREALCLFDQMKAS-IKPDHIAFVMVLSACSHGGLIAEGWECFKAmrDFGVTpeHYACMVDLLGRAGRLSEAQAFIERMP----ITPDAGVWGALLGACRIHSNLETAETAARHLFNLDAE-NPGRYILLSNIYASSGKRKEADDIRALMKSRGVRKTVGHTI----------IEIKNKVYTFVAGDTSNPQTDLIYSELRKLMDRIQEAGYVPD----------------------    
 180 UniRef50_Q9LPX2        Pentatricopeptide repeat-co...   424    1e-117  1  11.2%   1:544    89:630     IDFNRLFSAIAklALCKQMESKGIAHSIYTLSIMINCFCRCRKLS---YAFSTMGkmKLGYEPDTVIFNTLLNGLCLECRVSEALELVDRMVEMGHKPTLITLNTLVNGLCLNGKVSDAVVLIDRMVETGFQPNEVTYGPVLNvaLAMELLRKMEERNIKLDAVKYSIIIDGLCkgSLDNAFNLFNEMEIKGFKADIITYNTLIGGFCNAGRWDDGAKLLRDMIKRKISPNVVTFSVLIDSFVKEGKLREADQLLKEMMQRGIAPNTITYNSLIDGFCKENRLEEAIQMVDLMISK------------GCDPDIMTFNILINGYCKANRIDDGLELFREMSLRG--VIANTVTYNTLVQGFCQSGKLEVAKKLFQEMVSRRVRPDIVSYKILLDGLCDNGELEKALEIFGKIEKSKMELDiyMIIIHGMCNASKVDDAWDLFCSLPLKGVKL-DARAYNIMISELCRKDSLSKADILFRKMTEEGHAPDELTYNILIRAHLGDDDATTAAELIEEMKSSGF-PADVSTVKMVINML-SSGELDK-----------------------------------------------------------    
 181 UniRef50_B2ZAU5        Putative pentatricopeptide ...   423    1e-117  1   9.8%  14:592   202:725     -------------ESAFELFDKLCDRDVISWNSMISGYVSNGLTERGLGIYKQM--MYLGIDVDLATIISVLVGCANSGTLSLGKAVHSLAIKSSFERRINFSNTLLDMYSKCGDLDGALRVFEKmeRNVVSWTSMIAGYTRDGRsaIKLLQQMEKEGVKLDVVAITSILHACArgSLDNGKDVHDYIKANNMESNLFVCNALMDMYAKCGSMEAANSVFSTMVVK----DIISWNTMIG---------------------ELKPDSRTMACVLPACASLSALERGKEI------------HGYILRNGYSSDRHVANALVDLYVKCGVLGLARLLFDMIPSK------DLVSWTVMIAGYGMHGYGNEAIATFNEMRDAGIEPDEVSFISILYACSHSGLLEQGWRFFYIMkdFNIEpeHYACMVDLLSRTGNLSKAYEFMETLP----IAPDATIWGALLCGCRNYHDIELAEKVAERVFE--LEpnTGYYVLLANIYAEAEKWEEVKRLREKIGKQGLRKNP----------GCSWIEIKGKVNLFVSgnSSHPHSKNIESLLKKMRRKMKEEGHFP---KTKYALINA-----------    
 182 UniRef50_Q0WSH6        Pentatricopeptide repeat-co...   423    1e-116  1  10.0%   3:581    45:596     --ANYLINMYSKLDhaRLVLRLTPARNVVSWTSLISGLAQNGHFSTA--LVEFFEMRREGVVPNDFTFPCAFKAVASLRLPVTGKQIHALAVKCGRILDVFVGCSAFDMYCKTRLRDDARKLFDEieRNLETWNAFISNSvgRPREAIEAFIEFRRIDGHPNSITFCAFLNACSdlHLNLGMQLHGLVLRSGFDTDVSVCNGLIDFYGKCKQIRSSEIIFTEMGTK----NAVSWCSLVAAYVQNHEDEKASVLYLRSRKDIVETSDFMISSVLSACAGMAGLELGRSI------------HAHAVKACVERTIFVGSALVDMYGKCGCIEDSEQAFDEMPEK------NLVTRNSLIGGYAHQGQVDMALALFEEMAPRGctPNYMTFVSLLSACSRAGAVENGMKIFDSMRsyGIEpeHYSCIVDMLGRAGMVERAYEFIKKMP----IQPTISVWGALQNACRMHGKPQLGLLAAENLFKLDPK-DSGNHVLLSNTFAAAGRWAEANTVREELKGVGIKK----------GAGYSWITVKNQVHAFQAKDRSHILNKEIQTTLAKLRNEMEAAGYKPD----------------------    
 183 UniRef50_Q2QY23        Cytochrome P450 family prot...   423    1e-116  1  12.2%   1:598  910:1533     LGAlaAYADAGMLGKATEMCErgSLPEVTHCNRLLKLLVEQRRWDDARKLYDEM--LGKDSGADNYSTCVLVRGLCLERRVEEGLKLIEARWGAGCIPHVVFYNVLIDGYCRRGDMGRGLLLLGEMETNGFLPTLVTYGSLINweKIGSLFLEMRKRGFSPNVQIYNSVIDALCNCraTQAMVILKQMFASGCDPDIITFNTLITGLCHEGHVRKAEHFLREAIRRELNPNQLSYTPLIHGFCMRGELMVASDLLVEMMGRGHTPDVVTFGALIHGLVVAGKVSEALIVREKMTERQVF------------PDVNIYNVLISGLCKKRMLPAAKNILEEMLEKN--VQPDEFVYATLIDGFIRSENLGDARKIFEFMEHKGVCPDIVSCNAMIKGYCQFGMMSEAILCMSNMRKVGCIPdtYTTVISGYAKQGNLNGALRWLCDMIKRKCK-PNVVTYSSLINGYCKTGDTDSAEGLFANMQAEALSPNVVTYTILIGSLFKKDKVLRAGLYFETMLLNHCSPNDVTLHycVINSICCNteVHGkrNSAYNAIIFSLCRHNMLREALDFKNRMAKKGYVPNPI-TFLSLLYGFCSVGK-----    
 184 UniRef50_Q9LFI1        Pentatricopeptide repeat-co...   423    1e-116  1  11.5%   4:576   207:752     ---NALIAMYVrmSDASRVFYGIPMKDLISWSSIIAGFSQLGFEFEALSHLKEMLSFGV-FHPNEYIFGSSLKACSSLLRPDYGSQIHGLCIKSELAGNAIAGCSLCDMYARCGFLNSARRVFDQIErdTASWNVIIAGLANNgdEAVSVFSQMRSSGFIPDAISLRSLLCAQTkmALSQGMQIHSYIIKWGFLADLTVCNSLLTMYTFCSDLYCCFNLFEDFRNN---ADSVSWNTILTACLQHEQPVEMLRLFKLMLVSECEPDHITMGNLLRGCVEISSLKLGSQV------------HCYSLKTGLAPEQFIKNGLIDMYAKCGSLGQARRIFDSMDNR------DVVSWSTLIVGYAQSGFGEEALILFKEMKSAGIEPNHVTFVGVLTACSHVGLVEEGLKLYATMqeHGISpeHCSCVVDLLARAGRLNEAERFIDEMK----LEPDVVVWKTLLSACKTQGNVHLAQKAAENILK--IDpnSTAHVLLCSMHASSGNWENAALLRSSMKKHDVKKIPGQ----------SWIEIEDKIHIFFAEDIFHPERDDIYTVLHNIWSQMLDE---------------------------    
 185 UniRef50_Q0JMP2        Os01g0374200 protein n=2 Ta...   423    1e-116  1  11.7%   5:579    52:598     ----SLLNMYCKLgdARRVFDGMPQRNSFSWSTMVAGYAAEKCSEEAFDLFRLMLE-ECPSEKSEFVATAVLSAVSVPLGLLMGEQMHGLIVKDGLLDFVSVENSLVTMYAKAGCMGAAFHVFESseRNSITWSAMITGYAQNgdSAVSMFSQMHAAGFTPTEFTFVGVLNASSDLglAVGKQAHGLMVKLGFEVQIYVKSALVDMYAKCGCIADAKEGFDQL----YEVDIVLWTAMVSGHVQNGEHEEALTLYARMDKEGIIPSKSTIASGLRACAGIAALEPGKQL------------HTQIVKYGLGLGAPVGSALSTMYSKCGNLEDGMSVFRRIPDR------DVIAWNSIISGFSQNGCGNGALDLFEEMKMEGTIPDNITFINILCACSHMGLVDRGWEYFSLmkDYGLTpdHYACMVDILSRAGMLKEAKDFIESITIDHG----TCLWRIVLGACRSLRDFDVGAYAGERLMELG-TGDSSAYILLSNIYASQRKWNDVERVRHLMRLRGVNKDP----------GCSWVELNSRVHVFVVGEQQHPEAENINAQLRRLAKHMKDEGYH------------------------    
 186 UniRef50_B9S4F5        Pentatricopeptide repeat-co...   422    1e-116  1   9.4%   6:577   285:824     -----MYGKCQMIESAEKVFNEVPDKEIELWNALISAYVGNGYAYDALRIYKQMKL--CTVLSDSFTILNVLTSSSMAGLYDLGRLIHTEIVKRPLQSSITIQSALLTMYSKFGDSNYANSIFSTmeRDVVAWGSVISGFCQnyKEALDFFRAMEADLVKPDSDIMASIISAcgLEKVDLGCTIHGFVIKSGLQLDVFVASSLLDMYSKFGFPERAGNIFSDMPLK----NLVAWNSIISCYCRNNLPDLSINLFSQVLRNDLYPDSVSFTSVLAAISSVAALLKGKSV------------HGYLVRLWIPFDLQVENTLIDMYIKCGLLKYAQHIFERISEK------NLVAWNSMIGGYGSHGECSKAIELFDEMRSSGIKPDDVTFLSLLSSCNHSGLIEEGLHLFEMMkkFGIEpeHYVNIVDLYGRAGCLGDAYSFVKNMP----VEPDRSIWLSLLCSCKIHLNLELGEMVANKLL--NMEPSKGsyVQLLNLYGEAELWDRTANLRASMKEKGLKKTP----------GCSWIEVRNKVDVFYSGDCSSPITTEIYDTLSSLKRNMIKKG--------------------------    
 187 UniRef50_B9R9U2        Pentatricopeptide repeat-co...   422    1e-116  1  13.1%  18:544   198:714     -----------------EMLSKGVIPTVYTFGVVMKALCMVNEVDNACSLLRDMTKHGCV--PNSVVYQTLIHALSKRDRVNEALKLLEEMFLMGCLPDvtFNDVIYGLCRLNRIHEGA-KLVDRMLFRGFTPNDITYgrVGKVDEAQVLLNKVP----TPNDVHFTILINGYVkgRLDEANalYDKMIKNGCRPDVFTFNTLIHGLCKKGLMGSAVDMVNDMSANGCTPNLITYTTLLDGFCKKNQLEEAGYVLNEMSAKGFELNIMGYNVLLRALCKNGKVPKALDMLGEMSDK------------GCKPDIFTFNTLIFGLCKVDRKEDALALYRDMLLDG--VIANTVTYNTLIHAFLRGGAIQEALKLVNDMLFRGCPLDEITYNGLIKAFCKLGATEKALGLFDEMVRKDLVPssCNLLINGLCRVGKVCNALELLRDMIHRGLA-PDVVTYNSLINGLCKMGNIREAFNLFNKLQAEGIQPDAITYNTLICWHCRAGMFDDAYLLLLRGVENAFIPNDVTWYILVSNFIKEIGKENG-----------------------------------------------------------    
 188 UniRef50_D1HZ16        Whole genome shotgun sequen...   422    1e-116  1  11.0%  92:592    69:541     -------------------------------------------------------------------------------------------LKRGFEPDAVTVTTLVKGVWMENGIPDAVQLFDEMTEKGLFGDAKTYGILINGllAIKLHEKMKG-NCKGDVFTYGMIIDSLCKDGMteALDMFSEMIGAGILPDVVVYSSLMDGLCRFGRLKEALEFFKEMEGRGISADVYTYNSLIHGLSRAGLWKEVTWFLNLMVDRGFSPDAFTFTILIDGLCKEGKVGEAQQILELMHHK------------GKEPDILTYNTLMNGLCLVGQLEDATKLFESLADRG--IKLNVFSYNILINGYCKDQKIDEAFRLFEEMRPKGLKPSTVTYNTLIGALCQSGRVRTAQKLFVEMQTCglKLSTYCVLLDGLCKNGHLEEAIDLFQSIK-KTEHKPNIEVFSILLDGMCRAGKLEEAWKQFDEISKNGLEPDTIAYNILINGLCNKGMLSEAVKLLWQMEEKGCLPDSIT---------------------FNVIIQNLLKENEIHE-AIQLLEEMRNRNFSPDEAVTSMLLCLA-----------    
 189 UniRef50_C5YI98        Putative uncharacterized pr...   422    1e-116  1  11.4%   4:580   449:995     ---NALISFYAkiDNAVLVFDRMPHQDTISWNSVISGCTSNGLNSEAIELFVRMWMQGH--ELDSTTLLSVLPACARSHYWFVGRVVHGYSVKTGLIGETSLANALLDMYSNCSDWHSTNQIFRNmqKNVVSWTAMiyTRAGLFDKVAGLLQEMVLDGIKPDVFAVTSVLHGFadESLKQGKSVHGYAIRNGMEKLLPVANALMEMYVNCRNMEEARLVFDHVTNK----DIISWNTLIGGYSRNNFANESFSLFSDMLLQ-FKPNTVTMTCILPAVASISSLERGREI------------HAYALRRGFLEDSYTSNALVDMYVKCGALLVARVLFDRLTKK------NLISWTIMIAGYGMHGCGKDAVALFEQMRGSGVEPDTASFSAILYACCHSGLTAEGWKFFNAmkEYKIEPkhYTCIVDLLSHTGNLKEAFEFIESMP----IEPDSSIWVSLLHGCRIHRDVKLAEKVADRVFK--LEpnTGYYVLLANIYAEAERWEAVKKLKNKIGGRGLRE----------NTGCSWIEVRGKVHVFIADNRNHPEWNRIAEFLDHVARRMREEGHDP-----------------------    
 190 UniRef50_Q9FXH1        Pentatricopeptide repeat-co...   422    1e-116  1  10.2%   4:603   122:756     ---LSMYAKCGCIADARKVFDSMRERNLFTWSAMIGAYSRENRWREVAKLFRLM--MKDGVLPDDFLFPKILQGCANCGDVEAGKVIHSVVIKLGMSSCLRVSNSILAVYAKCGELDFATKFFRRmeRDVIAWNSVLLAYCQNgeEAVELVKEMEKEGISPGLVTWNILIGGYNqgKCDAAMDLMQKMETFGITADVFTWTAMISGLIHNGMRYQALDMFRKMFLAGVVPNAVTinSLVDMYSKCGKLEDARKVFDSVKNK----DVYTWNSMITGYCQAGYCGKAYELFTRMQ------------DANLRPNIITWNTMISGYIKNGDEGEAMDLFQRM-EKDGKVQRNTATWNLIIAGYIQNGKKDEALELFRKMQFSRFMPNSVTILSLLPACANLLGAKMVREIHGCVLRRNLDAIHanALTDTYAKSGDIEYSRTIFLGMETK-----DIITWNSLIGGYVLHGSYGPALALFNQMKTQGITPNRGTLSSIILAHGLMGNVDEGKKVFYSIANDYHIIPALEHCSAMVYLYGRANRLEEAleSFLTGCRIHGDIDMAIHAAENLFS------LEPENTATESIviYALGAknKPRRDN    
 191 UniRef50_Q9FLZ9        Pentatricopeptide repeat-co...   422    1e-116  1  10.7%   7:602    59:645     ------YALCGHITYARKLFEEMPQSSLLSYNIVIRMYVREGLYHDAISVFIRMVSEGVKCVPDGYTYPFVAKAAGELKSMKLGLVVHGRILRSWFGRDKYVQNALLAMYMNFGKVEMARDVFDVMkrDVISWNTMISGYYRNgnDALMMFDWMVNESVDLDHATIVSMLPVCGHldLEMGRNVHKLVEEKRLGDKIEVKNALVNMYLKCGRMDEARFVFDRMERR----DVITWTCMINGYTEDGDVENALELCRLMQFEGVRPNAVTIASLVSVCGDALKVNDGK------------CLHGWAVRQQVYSDIIIETSLISMYAKCKRVDLCFRVFSGASKYHTGP------WSAIIAGCVQNELVSDALGLFKRMRREDVEPNIATLNSLLPAYAALADLRQAMNIHCYLTKTGfsLDAATGLVHVYSKCGTLESAHKIFNGIQEK-HKSKDVVLWGALISGYGMHGDGHNALQVFMEMVRSGVTPNEITFTSALNACSHSGLVEEGLTLFRFMLEHYKTLARSNHYTCIVDLLGRAGRLDEA----YNLITTIPFEPTSTVWGALLAACVTHENVQLGEmtGNYVLLANIYAALGRWKD-    
 192 UniRef50_Q9SS83        Pentatricopeptide repeat-co...   422    1e-116  1   8.6%   4:584  467:1021     ---NALVDMYAKCgdARQIFERMCDRDNVTWNTIIGSYVQDENESEAFDLFKRMNL--CGIVSDGACLASTLKACTHVHGLYQGKQVHCLSVKCGLDRDLHTGSSLIDMYSKCGIIKDARKVFSSLPesVVSMNALIAGYSqnLEEAVVLFQEMLTRGVNPSEITFATIVEACHkeSLTLGTQFHGQITKRGFSSeeYLGISLLGMYMNSRGMTEACALFSELSS---PKSIVLWTGMMSGHSQNGFYEEALKFYKEMRHDGVLPDQATFVTVLRVCSVLSSLREGRAI------------HSLIFHLAHDLDELTSNTLIDMYAKCGDMKGSSQVFDEMRRRS-----NVVSWNSLINGYAKNGYAEDALKIFDSMRQSHIMPDEITFLGVLTACSHAGKVSDGRKIFEMMiqYGIEadHVACMVDLLGRWGYLQEADDFI----EAQNLKPDARLWSSLLGACRIHGDDIRGEISAEKLIE--LEpnSSAYVLLSNIYASQGCWEKANALRKVMRDRGVKKVPGY----------SWIDVEQRTHIFAAGDKSHSEIGKIEMFLEDLYDLMKDDAVvpDIVE-------------------    
 193 UniRef50_C5WU11        Putative uncharacterized pr...   422    1e-116  1   9.8%   1:528    95:610     LAATALANMYFKCrdARRVFDRMPARDRVAWNALVAGYARNGLPALAMEAVVRMQGEEGGEQPDSITLVSVLPACADARVLGACREVHAFALRAGLDELVNVSTAVLDAYCKCGAVEAARAVFDWMprNSVSWNAMIDGYagNASEALALFWRMVQEGVDVTDAAVLAALQACRegCLDEARRVHELLVRVGLKSNVSVMNALITTYSKCKRADLAAHAFNEL---GIKKTRISWNAMILGFTQNGCSEDAERLFARMQLENVKPDSFTLVSVIPAVADISDPLQARWI------------HGYSIRHQLDQDIYVLTALIDMYSKCGRVTIARKLFDSARDRH------VITWNAMIHGYGSHGFGQVAVELFEEMKGTGSLPNETTFLSVLAACSHAGLVDEGRKYFASMKDYGLEpeHYGTLVDLLGRAGKVDEAWSFIQNMP----IEPGISVYGAMLGACKLHKNVELAEESAQKIFELG--PEEGSLSQTMNalARcgE-QKVKCLYDSMSHSNGFLPAQTL---------------------------------------------------------------------------    
 194 UniRef50_Q654C7        Os06g0506100 protein n=5 Ta...   422    1e-116  1   8.3%   4:580   191:740     ---NALVTMYSKNGLVDdlFERIKDKDLISWGSIIAGFAQQGFEMEALQVFREMIVEGSH-HPNEFHFGSAFRACGAVGSWEYGEQIHGLSIKYRLDRDLYVGCSLSDMYARCKNLDSARVAFYRIepDLVSWNSIVNAYSVEgsEALVLFSEMRDSGLRPDGITVRGLLCAcgRDALYHGRLIHSYLVKLGLDGDVSVCNSLLSMYARCSDLSSAMDVFHEIKDQ----DVVTWNSILTACAQHNHPEEVLKLFSLLNKSEPSLDRISLNNVLSASAELGYFEMVKQV------------HAYAFKAGLVDDRMLSNTLIDTYAKCGSLDDAMRLFEIM-----GNNRDVFSWSSLIVGYAQFGYAKEAFDLFSRMRSLGIRPNHVTFIGVLTACSRVGFVNEGCYYYSIMeeYGIVpeHCSCIVDLLARAGKLTEAANFIDQMP----FEPDIIMWKTLLAASKMHNDMEMGKRAAEGIL--NIDPSHSayVLLCNIYAASGNWNEFARLKKAMRTSGVKKSPGK----------SWVKLKGELKVFIVEDRSHPESEEIYAMLELIGMEMIKAGYVP-----------------------    
 195 UniRef50_C5XB35        Putative uncharacterized pr...   422    1e-116  1  10.8%  14:596   126:679     -------------GRALHLLDQMPRRFAVtyNAVLSTLARADCHADALLLYRRM--LRDRVPPTTFTFGVAARALCRLGRARDALALLRGMARHGCVPDAVLytVIHALVAQGGVAEAATLLDEMLlaDVNTFNDVVLGLCghVREAARLVDRMMMHGCTPSVVTYGFLLRGLCRTraDEAYAMLGRVPE----VNVVMLNTVIRGCLAEGKLARATELYEMMGSKGCPPDVHTYNILMHGLCKLGRCGSAVRMLDEMEEKGCAPNIVTYSTLLHSFCRNGMWDDARAMLDQMSAK------------GFSMNSQGYNGIIYALGKDGKLDEAMRLVQEMKSQGC--KPDICTYNTIIYHLCNNDQMDEAEHIFGNLLEEGVVANGITYNTLIHALLHSGRWQEGLRLANEMVLHGCPLDvyNGLIKALCKEGNVDRSMMLLEEMMTKGIK-PNNFSYNMLINELCKAGKVRDALELSKEMLNQGLTPDIVTYNTLINGLCKVGWTHAALNLLEKLPNENVHP----------DIVTYNILISWHCKVRLLDDASM------------LLDKAVSGGIVPNE-RTWGMMVQNFVRQ-------    
 196 UniRef50_A5C1A7        Putative uncharacterized pr...   421    1e-116  1  10.1%   7:602   228:861     ------YSNCGDLGSSRSVFHSMQERDVVSWTALISAYMEEGLXDEAKHIFHLMQLDG--VKPDLISWSALLSGFARNGEIDLALETLEEMPERGLQPTVNSWNGIISGCVQNGYLEDALDMFSRMLWYPEDPNIITIASILPACTGLKALRlaXKHGIVGNVYVEGSVIDMYSKCgyDYAEKVFXKAENK----NTAMWNEMIAAYVNEGKVEDALGLLRSMQKDGWKPDVITYNTILSGHARNGLKTQAXELLSEMVQMGLKPNVVSFNVLISGFQQSGLSYEALKVFRIMQscQGKEIHGYTLRNGFEPNIFVSSALVDMYAKCHDMDSANKVFFRIDGR------NTVSWNALMAGYINNKQPEEALKLFLEMLGEGLQPSSITFMILFPACGDIAAIRFGRGLHGYAAkdELKNAIXSALIDMYAKCGSILDAKSVFDSEVEK-----DVPLWNAMISAFSVHGMARNAFAVFXQMELLGIXPDHITFVSLLSACARDGLVEEGWKYFNSMEISYGVAATLEHYTCMVGILGGAGLLDEAL----DFIRQMPYPPDACMWATLLQACRVHSnlEPDNATNYMLLSNIYVSSGMWDFA-    
 197 UniRef50_Q9LW63        Putative pentatricopeptide ...   421    1e-116  1  10.3%  20:582    62:628     -------------------FKTLKSPPVLAWKSVIRCFTDQSLFSKALASFVEMRASGRCPDHNV--FPSVLKSCTMMMDLRFGESVHGFIVRLGMDCDLYTGNALMNMYAkgDeiDSVRRVFEVMPrdVVSYNTIIAGYAQSgeDALRMVREMGTTDLKPDSFTLSSVLPIFSevDVIKGKEIHGYVIRKGIDSDVYIGSSLVDMYAKSARIEDSERVFSRL----YCRDGISWNSLVAGYVQNGRYNEALRLFRQMVTAKVKPGAVAFSSVIPACAHLATLHLGKQL------------HGYVLRGGFGSNIFIASALVDMYSKCGNIKAARKIFDRM------NVLDEVSWTAIIMGHALHGHGHEAVSLFEEMKRQGVKPNQVAFVAVLTACSHVGLVDEAWGYFNSMTkyGleLEHYAAVADLLGRAGKLEEAYNFISKM----CVEPTGSVWSTLLSSCSVHKNLELAEKVAEKIFTVDSE-NMGAYVLMCNMYASNGRWKEMAKLRLRMRKKGLRKKPA----------CSWIEMKNKTHGFVSGDRSHPSMDKINEFLKAVMEQMEKEGYVADT---------------------    
 198 UniRef50_Q9SR82        Putative pentatricopeptide ...   421    1e-116  1  10.9%  26:586    74:602     -------------------------PNIFLYNSLINGFVNNHLFHETLDLFLSIRKHGLY--LHGFTFPLVLKACTRASSRKLGIDLHSLVVKCGfhDVAAMTSLLSIYSGSGRLNDAHKLFDEIPDRSVVTWTALFSGysGRHREAIDLFKKMVEMGVKPDSYFIVQVLSACVhgDLDSGEWIVKYMEEMEMQKNSFVRTTLVNLYAKCGKMEKARSVFDSMVEK----DIVTWSTMIQGYASNSFPKEGIELFLQMLQENLKPDQFSIVGFLSSCASLGALDLGEW------------GISLIDRHEFLTNLFMANALIDMYAKCGAMARGFEVFKEMKEK------DIVIMNAAISGLAKNGHVKLSFAVFGQTEKLGISPDGSTFLGLLCGCVHAGLIQDGLRFFnyALKR--TVEHYGCMVDLWGRAGMLDDAYRLICDMPMR----PNAIVWGALLSGCRLVKDTQLAETVLKELIA--LEpnAGNYVQLSNIYSVGGRWDEAAEVRDMMNKKGMKKIPGY----------SWIELEGKVHEFLADDKSHPLSDKIYAKLEDLGNEMRLMGFVPTTEFVF-----------------    
 199 UniRef50_B9IIT4        Predicted protein n=20 Tax=...   421    1e-116  1  10.6%  72:591     7:492     -----------------------------------------------------------------------ISTLFKCRNIFQIKQVHAQVTTTGIIHDLIVANKLLYMCAKHKDLVTAHLLFNKMerDPVSWSVMIGGFVKNGdrCFQTFRELIRAGSKPDNFSLPFVIKACRdmGLIMGRLIHSTVLKNGLHLDNFVCSTLVDMYAKCGMIDNAKQLFDRMPKK----DLVTRTVMIAGYAECGKPNESWVLFDQMRRDGFVPDKVAMVTIVNACAKLGAMNKARLVHDYVCARR------------YSLDVELGTAMIDMYAKCGSIDSSREIFDRMEQK------NVISWSAMIGAYGYHGQGREALELFHMMLNSGIIPNRITFISLLYACSHAGLVDDGLQLFSLmvSYGVRPdhYTCMVDLLGRAGRLDQALRLIENME----VEKDEGIWCAFLGACRIHRQVDLAEKAAKLLLSLQTQ-NPGHYILLSNIYANAGRWKDVAKIRNLMAKRRLKKIPGYT----------WIEVDNIIYRFGAGDNSHLRSNEIYEMLKSLSQKLESAGYVPDT----NSVLH------------    
 200 UniRef50_Q6Z277        Putative uncharacterized pr...   421    1e-116  1  13.4%  11:576   143:684     ----------GRHADVRRAFgaGARPDTFAWNKAVQACVAAGDLGEAVGMLRRMGRDGA-PPPNAFSYNVVIAGMWRAGRGGDAVEVFDEMTERAVLPNHITYNTMIDGHIKGGDLEAGFRLRDQmkPNAITYNVLLSGLCRAGRMGeaLLDEMASQKMVPDGFTYSILFDGLSRNGDSKAmsLFGKYLKNGVTIGDYTCSILLNGLCKDGKVSIAEEVLQSLVNAGLVPTRVIYNTLINGYCQTGELEGAFSTFGQMKSRHIKPDHITYNALINGLCKAERITNAQDLLMEMQ------------DNGVNPTVETFNTLIDAYGRTGQLEKCFIVLSEMQENG--LKPNVVSYGSIVNAFCKNGKIPEAVAILDDMFHKDVLPNAQVYNAIIDAYVEHGPNDQAFILVEKMKSNGISPstYNLLIKGLCNQSQISEAEEIINSLSNHRLI-PDAVSYNTLISACCYRGNIDKALDLQQRMHKYGIKSTVRTYHQLISGLGGAGRLNEMEYLYQKMMQNNVVPSNAIH----------NIMVEAYSKYGNEI------------KAEDLRKEMLQK---------------------------    
 201 UniRef50_Q9AY70        Pentatricopeptide, putative...   421    1e-116  1  10.9%   3:578   285:831     --ANTLVSMYAKCKCLdkLFGLMPRDDLVTWNGMISGCVQNGFVDQALLLFCDMQ--KSGIRPDSVTLVSLLPALTDLNGFNQGKELHGYIVRNCVHMDVFLVSALVDIYFKCRAVRMAQSVYDSSKadVVIGSTMISGYVLNgqEAVKMFRYLLEQGIRPNAVAIASVLPACAsaAMKLGQELHSYALKNAYEGRCYVESALMDMYAKCGRLDLSHYIFSKISAK----DEVTWNSMISSFAQNGEPEEALNLFREMCMEGVKYSNVTISSVLSACASLPAIYYGKEI------------HGVVIKGPIRADLFAESALIDMYGKCGNLEWAHRVFESMPEK------NEVSWNSIIASYGAYGLVKESVSLLRHMQEEGFKADHVTFLALVSACAHAGQVQEGLRLFRCmeEYQIApeHFACMVDLYSRAGKLDKAMELIVDMP----FKPDAGIWGALLHACRVHRNVELAEIASQELFKLDPH-NSGYYVLMSNINAVAGRWDGVSKVRRLMKDTKVQKIPGY----------SWVDVNNTSHLFVAADKSHPDSEDIYMSLKSILLELREEGH-------------------------    
 202 UniRef50_Q9SMZ2        Pentatricopeptide repeat-co...   421    1e-116  1  10.7%   6:602   359:918     -----MYCKLRKFGFARTVFDNMSERDLISWNSVIAGIAQNGLEVEAVCLF--MQLLRCGLKPDQYTMTSVLKAASSLPelSLSKQVHVHAIKINNVSDSFVSTALIDAYSRNRCMKEAEILFERHNFDLVAWNAMMAGysHDGHKTLKLFALMHKQGERSDDFTLATVFKTCGFlaINQGKQVHAYAIKSGYDLDLWVSSGILDMYVKCGDMSAAQFAFDSIP----VPDDVAWTTMISGCIENGEEERAFHVFSQMRLMGVLPDEFTIATLAKASSCLTALEQGRQI------------HANALKLNCTNDPFVGTSLVDMYAKCGSIDDAYCLFKRIE------MMNITAWNAMLVGLAQHGEGKETLQLFKQMKSLGIKPDKVTFIGVLSACSHSGLVSEAYKHMRSMhdYGIKpeHYSCLADALGRAGLVKQAENLIESMS----MEASASMYRTLLAACRVQGDTETGKRVATKLLE--LEpdSSAYVLLSNMYAAASKWDEMKLARTMMKGHKVKKDPGF----------SWIEVKNKIHIFVVDDRSNRQTELIYRKVKDMIRDIKQEGYVPETDFT---LVD--VEEEEKERA-    
 203 UniRef50_C5XB24        Putative uncharacterized pr...   420    1e-116  1  11.5%   9:564    74:659     --------RSGDLgdALDLFDELLQrsIYALNQLLTTVARasSVRdaVSMFNRMARAgkKVAPDIATFGILISCCCDAGCLNLGFAALGQIIKTGLRAQAVTFtlLRTLCAEKRTSDAMNIVLRRMppDVFSYTTLLKGLckKCEEAAELIHMMAEDgcPPNVVSYTTVIHGfkEGDVGKAYTLFCKMLDHGIPPNVVTCNSVIDGLCKVQAMDKAEAVLQQMIDEHIMPNCTTYNSLIHGYLSSGQWTEAVRILKEMSRDGQRPNVVTYSMLIDCLCKSGLHAEAREIFNSMIQSGQKpvKDLMVQNGMRPGRHVFNIEIYAYCKCGRLDEASLTFNKMQQQG--FMPDIVAYTTVIDGLCKIGRLDDAMSRFCQMIDDGLSPDIITFNTLIHGFALHGKWEKAEELFYEMMDRGIPpnTFNSMIDKLFKEGKVTEARKLFDLMPRAG-AKPNVVSYNTMIHGYFIAGEVGEVMKLLDDMLLIGLKPTAVTFNTLLDG----------------MVSMGLKPDVVT-CKTLIDSCCEDGRIEDILTLFREMLGKADKTDTITE---------------------------------------    
 204 UniRef50_A5AZT2        Putative uncharacterized pr...   420    1e-116  1   9.5%   6:565   183:716     -----LINLYAKCGCLNQALLLFSikTIVTWTSLITHLSHFNMHLQALSLFNQMR--CSGPYPNQFTFSSILSASAATMMVLHGQQLHSLIHKHGFDANIFVGTALVDMYAKCADMHSAVRVFDQmeRNLVSWNSMIVGFFHnyDRAVGVFKDVLREkvIPNEVSVSSVLSACAngGLNFGRQVHGVVVKYGLVPLTYVMNSLMDMYFKCRFFDEGVKLFQCVGDR----DVVTWNVLVMGFVQNDKFEEACNYFWVMRREGILPDEASFSTVLHSSASLAALHQGTAIHDQ------------IIKLGYVKNMCILGSLITMYAKCGSLVDAYQVFEGIEDH------NVISWTAMISAYQLHGCANQVIELFEHMLSEGIEPSHVTFVCVLSACSHTGRVEEGLAHFNSMKkmNPGPEHYACMVDLLGRAGWLDEAKRFIESMPMK----PTPSVWGALLGACRKYGNLKMGREAAERLFE--MEpnPGNYVLLANMCTRSGRLEEANEVRRLMGVNGVRKEP----------GCSWIDVKNMTFVFTAHDRSHSSSDEIYKM--------------------------------------    
 205 UniRef50_Q56XI1        Pentatricopeptide repeat-co...   420    1e-116  1  10.4%   4:581    52:617     ---NSMVAGYFapRDARKLFDEMPDRNIISWNGLVSGYMKNGEIDEARKVFDLM------PERNVVSWTALVKGYVHNGKVDVAESLFWKMPEKNKVSWTV---MLIGFLQDGRIDDACKLYEMIPDKDNIARTSMIHGLcgRVDEAREIFDEMSER----SVITWTTMVTGYGQnrVDDARKIFDVMPEK----TEVSWTSMLMGYVQNGRIEDAEELFEVMPVKP----VIACNAMISGLGQKGEIAKARRVFDSMKER----NDASWQTVIKIHERNGFELEALDLFILMQKQggKQVHAQLVRCQFDVDVYVASVLMTMYIKCGELVKSKLIFDRFPSK------DIIMWNSIISGYASHGLGEEALKVFCEMPLSgtKPNEVTFVATLSACSYAGMVEEGLKIYESMEsfGVKPihYACMVDMLGRAGRFNEAMEMIDSM----TVEPDAAVWGSLLGACRTHSQLDVAEFCAKKLIE--IEpnSGTYILLSNMYASQGRWADVAELRKLMKTRLVRKSP----------GCSWTEVENKVHAfrGGINSHPEQESILKILDELDGLLREAGYNPD----------------------    
 206 UniRef50_D1HY44        Whole genome shotgun sequen...   420    1e-116  1  11.8%  34:603    16:564     ---------------------------------LVANCVKSRKLREAFDIIQTMRKFKFRPAFS--AYTILIgvLCKANRLDEAVELFEQLEQNRKVPCAYAYNTMIMGYGSAGKFDEAYGLLERQKAKGSIPSVIAYNCirRVEEALRIFEEMKRDAV-PNVPTYNILIDMLCrgKLNAALEIRDDMERAGLFPNVLTVNIMIDRLCKAQKLEEACSIFEGMDDKVCTPNAVTFSSLIDGLGKCGRVDDAYSLYEKMLDCGHVPGAIVYTSLIRSFFKCGRKEDGHKIYKEMV------------HTGCSPDLTLINTYMDCVFKAGETEKGRALFREINAHG--FIPDARSYSILIHGLVKAGLANETYELFYAMKEQGCVLDTHAYNAVIDGFCKSGKVNKAYQLLEEMKVKGHPpvTYGSVIDGLAKIDRLDEAYMLFEEAKSNGIKL-NVVVYSSLIDGFGKVGRIDEAYLIMEELMQKGLTPNVYTWNCLLDALVKAEEINEALICFQSMKD----------LKCPPNQITYSILINGLCRV------------RKFNKAFVFWQEMQKLGLKPNTI-TYTTMISGLAKAGNILEAS    
 207 UniRef50_B9I1Z2        Predicted protein n=1 Tax=P...   420    1e-116  1  11.0%   3:574   198:846     --SNALIDMYGKCgdAKKVLVKMPERDSVTWNSVITACAANGMVYEALEFLEKMKSLDYS-MPNVVSWSAVIgaCARLQRLDLGKQLHGYITRHDFISnvVVNALVDVYRRCGDmvSKAKELFDCmeRGLISWNSIISGYVRnfDEAFSMFQNMleEGIEPDSFTLGSVLTACADtsLRQGKEIHAQAIVKGLQSDTFVGGALVEMYSKCQDLTAAQVAFDEVMEK----DVPTWNALISGYTRSNQIERIQYLLEKMKGDGYHPNIYTWNSILAGLVENRQLDLTMQLFSEMqlERGKQAHAHSIKCGYDTDVHIGAALVDMYAKCGSLKYAQLAYDRISN------PNLVSHNAMLTACAMHGHGEEGISLFQTMLALGFIPDHVTFLSVLSSCVHVGSVETGCEFFDLMGYYNVKPthYTSMVDLLSRSGQLHEAYELIKKMPVEC----DSVLWGALLGGCVTHGNIELGEIAAERLIE--LEpnSGNYVLLANLHAYARRWTDLARVRGMMKDRGMHKSP----------GCSWIEDKNEIHSFLACDRSHKRAEEIYATLDYLALHMK-----------------------------    
 208 UniRef50_Q9S7F4        Putative pentatricopeptide ...   420    1e-116  1  10.8%   7:582   194:738     ------YCEVRRLDLACVLFEEIPEKDSVTFNTLITGYEKDGLYTESIHLFLKMRQSGHQ--PSDFTFSGVLKAVVGLHDFALGQQLHALSVTTGFSRDASVGNQILDFYSKHDRVLETRMLFDEMPedFVSYNVVISSYSqqYEASLHFFREMQCMGFDRRNFPFATMLSIAAnsSLQMGRQLHCQALLATADSILHVGNSLVDMYAKCEMFEEAELIFKSLPQR----TTVSWTALISGYVQKGLHGAGLKLFTKMRGSNLRADQSTFATVLKASASFASLLLGKQL------------HAFIIRSGNLENVFSGSGLVDMYAKCGSIKDAVQVFEEMPDR------NAVSWNALISAHADNGDGEAAIGAFAKMIESGLQPDSVSILGVLTACSHCGFVEQGTEYFQAMsiYGITPkhYACMLDLLGRNGRFAEAEKLMDEMP----FEPDEIMWSSVLNACRIHKNQSLAERAAEKLFSMEKLRDAAAYVSMSNIYAAAGEWEKVRDVKKAMRERGIKKVPAY----------SWVEVNHKIHVFSSNDQTHPNGDEIVRKINELTAEIEREGYKPDT---------------------    
 209 UniRef50_Q9SHZ8        Pentatricopeptide repeat-co...   420    1e-116  1   9.1%   7:589    90:706     ------YSKRGDMDSTCEFFDQLPQRDSVSWTTMIVGYKNIGQYHKAIRVMGDMVKEG--IEPTQFTLTNVLASVAATRCMETGKKVHSFIVKLGLRGNVSVSNSLLNMYAKCGDPMMAKFVFDRmeRDIVTWNSMISGFNQRglRALDIFSKMLRDSlsPDRFTLASVLSACANlkLCIGKQIHSHIVTTGFDISGIVLNALISMYSRCGGVETARRLIEQRGTKDLK--IEGFTALLDGYIKLGDMNQAKNIFVSLKDR----DVVAWTAMIVGYEQHGSYGEAINLFRSMvlSHGKQIHGSAVKSGEIYSVSVSNALITMYAKAGNITSASRAFDLIR-----CERDTVSWTSMIIALAQHGHAEEALELFETMLMEGLRPDHITYVGVFSACTHAGLVNQGRQYFDMMKddKIIpsHYACMVDLFGRAGLLQEAQEFIEKMP----IEPDVVTWGSLLSACRVHKNIDLGKVAAERLLL--LEpnSGAYSALANLYSACGKWEEAAKIRKSMKDGRVKKEQGF----------SWIEVKHKVHVFGVEDGTHPEKNEIYMTMKKIWDEIKKMGYVPDTASVLHDL--------------    
 210 UniRef50_A9STA3        Predicted protein n=2 Tax=P...   420    1e-115  1  11.4%  38:589   169:701     -------------------------------------LRELGNRGEWERALQGYEWMVQQGHLRSelASIMISTLGRLGKVEIALDVFNRAQKAGFGNNVYAYSAMVSAYGRSGRCREALKVFQAMKKAGCKPNLITYNTIIDAckALDIFEEMQKEGVEPDRITFNSLIAVCSRGSleDSQRVFAEMQRRGIEQDIFTYNTLIDAVCKGGQMELAASIMSSMRLKNISPNVVTYSTMIDGYGKLGCFEEAIGLYHDMKESGVRPDRVSYNTLIDIYAKLGRFDDALTACKDMER------------VGLKADVVTYNALIDAYGKQGKYKDAAGLFDKMKAEG--LVPNVLTYSALIDAYSKAGMHQDATSIFVEFKRAGLKPDVVLYSSLIDSCCKCGLVEDAVVLLQEMTQAGIQPntYNSLIDAYGRNGQVDNVEAAKGNMPINVFNKVGDRSTEIICKTLTSQQNANdaVSVFHEMQQFGLKPNVVTFSAILNACSRCSSLQEASVLLEQMR--------------FFDSWVY-----GIAHGLLMGLREH-----VWVEAQRLFDEIARMDYATGA-AFYNAL--------------    
 211 UniRef50_Q9SS60        Pentatricopeptide repeat-co...   420    1e-115  1  10.1%   5:581   250:793     ----AMYLKFRRPTDARRVFDEMDVRDSVSYNTMICGYLKLEMVEESV---RMFLENLDQFKPDLLTVSSVLRACGHLRDLSLAKYIYNYMLKAGFVLESTVRNILIDVYAKCGDMITARDVFNSMekDTVSWNSIISGYIQSGdeAMKLFKMMMIMEEQADHITYLMLISVSTraDLKFGKGLHSNGIKSGICIDLSVSNALIDMYAKCGEVGDSLKIFSSM----GTGDTVTWNTVISACVRFGDFATGLQVTTQMRKSEVVPDMATFLVTLPMCASL------------AAKRLGKEIHCCLLRFGYESELQIGNALIEMYSKCGCLENSSRVFERMSRR------DVVTWTGMIYAYGMYGEGEKALETFADMEKSGIVPDSVVFIAIIYACSHSGLVDEGLACFEKMkhYKIDpeHYACVVDLLSRSQKISKAEEFIQAMPIK----PDASIWASVLRACRTSGDMETAERVSRRIIELN--PDDPGYSIlsNAYAALRKWDKVSLIRKSLKDKHITKNPGY----------SWIEVGKNVHVFSSGDDSAPQSEAIYKSLEILYSLMAKEGYIPD----------------------    
 212 UniRef50_Q7XK59        OSJNBa0091C07.1 protein n=2...   419    1e-115  1  11.7%  47:602   156:722     ----------------------------------------------AIELFKRMDrcPRHSPPTIHTYNILIDCYRRVHRPDLGLAIVGRLLKNGLGPDDFSYSLIYGFVKDGEVDKAHCLFLEMMepKILICNSIIKELCKmmDKAESIVQKMVDSGIAPDLFTYSLIIDGLCKSkmDKAERVLEQMVEAGTRPNSITYNSLIHGYSISGMWNESVRVFKQMSSCGVIPTVDNCNSFIHALFKHGRTNEAKCIFDSMVLKGPKPDIISYSTMLHGYATATdlADVHNIFNLMLTK------------GIAPNKHVFNILINAYARCGMMDKAMLIFEDMQNKG--MIPDTVTFATVISSLCRIGRLDDALHKFNHMVDIGVPPSEAVYGCLIQGCCNHGELVKAKELISEMMNKDIPpkYFSSIINNLCKEGRVAEGKDIMDMMVQTG-QRPNVVTFNSLMEGYCLVGNMEEAFALLDAMASIGIEPNCYIYGTLVDGYCKNGRIDDALTVFRDMLHKGVKPTSV-LYSIILHGLFQARRTTAAKKMFHEMIEsgLCRNNCTDEANMLLEKLFAMNVKFDII-TFNIVISAMFKVGRRQEA-    
 213 UniRef50_B9SJZ3        Pentatricopeptide repeat-co...   419    1e-115  1  11.0%  33:596   176:705     --------------------------------VLIGGYLNY----QCLLVFEKM--MRNRFLPDVKNCNRILKILRDKNLLVKALEVYRMMGEYGIRPTVTTYNTMLHSFCKGGEVQRALDLVPKMQERgvTFNVLINGLSKKGElaKGLIQEMAKAGLRVSPYTYNPLICGYCKKGlvEALALWEEMVTRGVSPTVASHNTIMYGFCKEGKMSDARQQLSDMLKKNLMPDIISYNTLIYGFCRLGNIGEAFILLDELRFRNLSFNIVTYNTLIDGLCRLGDLETALKLKEDMINR------------GIHPDVVTYTVLVNGACKLGNMLMAKEFFDEML--HVGLAPDQFAYTARIVGELKLGDTAKAFKLQEEMLTKGFPPDVITYNVFVNGLCKLGNLEEAGELLQKMIRDGHVPDhyTSFMHAHMENGHLREGREIFYDMLSRG-QTPTVVTYTVLIHAHALNGRLDWAMAYFLEMQEKGVVPNVITYNVLINGFCKVRKMDQACKFFIEMQEKGI----------FPNKYTYTILINENCNMGK------------WQEALRLYAQMLGKRIRPDS-CTHGALLKKLDKD-------    
 214 UniRef50_Q9M9E2        Pentatricopeptide repeat-co...   419    1e-115  1  10.4%   2:578   134:777     -AFLAMFVRFGNLVDAWYVFGKMSERNLFSWNVLVGGYAKQGYFDEAMCLYHRMLWVG-GVKPDVYTFPCVLRTCGGIPDLARGKEVHVHVVRYGYELDIDVVNALITMYVKCGDVKSARLLFDRmrRDIISWNAMISGYFENghEGLELFFAMRGLSVDPDLMTLTSVISAclLGDRRLGRDIHAYVITTGFAVDISVCNSLTQMYLNAGSWREAEKLFSRMERK----DIVSWTTMISGYEYNFLPDKAIDTYRMMDQDSVKPDEITvnNLINMYSKCKCIDKALDIFHNIPRKngKEIHAHVLRTGVGLDDFLPNALLDMYVRCGRMNTAWSQFN-------SQKKDVTSWNILLTGYSERGQGSMVVELFDRMVKSRVRPDEITFISLLCGCSKSQMVRQGLMYFSKMEDYGVTPnhYACVVDLLGRAGELQEAHKFIQKMP----VTPDPAVWGALLNACRIHHKIDLGELSAQHIFELDKK-SVGYYILLCNLYADCGKWREVAKVRRMMKENGLT----------VDAGCSWVEVKGKVHAFLSDDKYHPQTKEINTVLEGFYEKMSEVGL-------------------------    
 215 UniRef50_A5C4M4        Putative uncharacterized pr...   419    1e-115  1   9.2%   3:576    90:663     --AAALVDAYGKCGHaaQVFDEMPevDVVSWTALISAYSSNGCVDEAFXAFGRMRWMreCCGVDVVSLGALVSAccGSNCLRRGSAVHGLVVKYGFGVSTHLGNSMVHMYSACKDVGGAWRVFNGiqRDVVSWNSLISGFXLNGerALRTFEDMVSEgvEPNRVTVIALLKSCAELGCVETSsvHEYISSRhvAKDVVVLTALLDMHARCGNLALAREIFDGVEGK----NVVCWSAMIAGYEQGSCPEEALRLFRQMLMEgvKPNAVTLVSVIAACSRLGA------------SRSASMIHKYAVATGLDQDARIASALIDMCAKCGDIEHGRQVFSEMDES----TRTVVSWSSMIGAEGIHGEGKRALELFSEMRTGGYEPNEITYISVLSACSHAGLVEQGKSCFNSMEkyGMSPthYACLVDLLGRAGHLDEAHNVILNMPIK----ADLALWGSLLAACHLHGNCKLGEIVEKKILSLDSN-SVGHHVLLANMYEDAGRWDDVVRMRVELRRSGLRKIPGQSF----------IEIGNEVYSFMAEDRSHPESEMIYKELDGLDERVRKA---------------------------    
 216 UniRef50_Q0J5U1        Os08g0402600 protein (Fragm...   419    1e-115  1  13.3%   5:576     1:547     ----ALLS-LGRHADVRRAFgaGARPDTFAWNKAVQACVAAGDLGEAVGMLRRMGRDGA-PPPNAFSYNVVIAGMWRAGRGGDAVEVFDEMTERAVLPNHITYNTMIDGHIKGGDLEAGFRLRDQmkPNAITYNVLLSGLCRAGRMGeaLLDEMASQKMVPDGFTYSILFDGLSRNGDSKAmsLFGKYLKNGVTIGDYTCSILLNGLCKDGKVSIAEEVLQSLVNAGLVPTRVIYNTLINGYCQTGELEGAFSTFGQMKSRHIKPDHITYNALINGLCKAERITNAQDLLMEMQ------------DNGVNPTVETFNTLIDAYGRTGQLEKCFIVLSEMQENG--LKPNVVSYGSIVNAFCKNGKIPEAVAILDDMFHKDVLPNAQVYNAIIDAYVEHGPNDQAFILVEKMKSNGISPstYNLLIKGLCNQSQISEAEEIINSLSNHRLI-PDAVSYNTLISACCYRGNIDKALDLQQRMHKYGIKSTVRTYHQLISGLGGAGRLNEMEYLYQKMMQNNVVPSNAIH----------NIMVEAYSKYGNEI------------KAEDLRKEMLQK---------------------------    
 217 UniRef50_A9RQM0        Predicted protein n=1 Tax=P...   419    1e-115  1  12.6%  71:600     1:502     ----------------------------------------------------------------------MLVVLASWGILDPLETLLKEMVAEGRPLRPNTLVKLITAYGRGNKSGDAFDLFNQAESFACSPTVHAFTKLIdfERAELVYKKLVQKGCQLDRFAYNVLIRYFGRSglDSAMEMFREMKIKGSEPDEYTYGFLVNALGKAGRVQEARSFFDAMLERGLTPNIPTYNLLMDAFRKVGQLDMALGLFAEMKRRGFQPSVVTYNILLDALCSAGRVGAARKLFHKMTGD------------GCSPDSYTYSTLVNGLGKSGRVEEAHKVFREMVDRGVAV--DLVNYNSLLATLAKAGNMDRVWKLMKEMSRKGFHPDAFSFNTIMDALGKANKPDAAREVFARMVESGCKPdsYNILIDSYARFGDAAQARQMLEEMVEAGFI-PETKTYNSLIHWLATDGQVDEAFAVLEEMETAGCRPDVVTYNRLMDMLGKRGENQRAARLFQQMKDKGVEP----------DTLSYAVRIDGL------------AFDDRLDEALVLFKDMKAVGC-PVDKAMYRILIRAAHRAGDTE---    
 218 UniRef50_D1HW24        Whole genome shotgun sequen...   419    1e-115  1   9.9%   6:578  693:1233     -----MYAKSGSVEDAKKVFDQVLDKEVELRNAMISAFIGNGR---AYDALGLYNKMKAGETpdSFTISSLLSGCSVVGSYDFGRTVHAEVIKRSMQSNVAIQSALLTMYYKCGSTEDADSVFYTmeRDVVAWGSMIAGFCQnfKDALDLFRAMEKEGVKADSDVMTSVISalGLENVELGHLIHGFAIKRGLESDVFVACSLVDMYSKFGFAESAEMVFSSMPNK----NLVAWNSMISCYSWNGLPEMSINLLPQILQHGFYLDSVSITTVLVAVSSVAALLKGKTL------------HAYQIRLQIPSDLQVENALIDMYVKCGCLKYAQLIFENMPRR------NLVTWNSMIAGYGSHGNCEEAVRLFKEMKRSETAPDEVTFLALITSCSHSGMVEEGLNLFQLMreYGVEpeHYASVVDLLGRAGRLDDAYSFIRGMP----IDADRSVWLCLLFACRAHRNMELGELVADNLLK--MEprGSNYVPLLNLYGEVEMWDRAANLRASMKGRGLKKSP----------GCSWIEVKNRVDVFFSGDSSSTRRIEIYKTLSSLKSNMEGKGC-------------------------    
 219 UniRef50_Q7XKS2        OSJNBa0038P21.9 protein n=3...   419    1e-115  1  10.5%  10:602   163:769     ---------CRDvaLAVELFKRMDrpTIYTYNILINCYRRARRPDLGLPVFGRL--LRTGLGPDVFSYNALIDGFSKEGEVDKAHDLFYKMEEQGIMPNVVTYSSLINGLCKTKEMDKAERVLRQMVGAGVRPNNMTYNCLIHgkESVRVFKEMSSSLLVPDVGNCNSFMTALCkgRIKEARDIFDSMVLKGPKPDVISYGALLHGYATAGCIAGMDNLFNVMVCEGVVPDRHVFNTLINAYARLGMMDKSLLMFEDMTKQGVNPDIITFSTVISAFCRLGRLDDAMEKFNHMIDTGvkELISDMLSKGipPCIKFFTSIINNLCKEGRVAEGKDVVDLIIHTGQR--PNLITFNSLVDGYCLVGNMKEAVGLLDSMESVGVEPDIYTYNTLVDGYCKHGRIDDALTLFRDMLHKRVTlvSYNIILHGLFQARRTIVAKEMFHEMIESG-MAVSIHTYATVLGGLCRNNCTDEANMLLEKLFSMNVKFDILTFNIVIRAMFKVGRRQEAKELF----------AAISTYGLVPTILTYRVMITNLIK------------EESFEDADNLFSSMEKSSCTPDS-RILNEIIRMLLNKGEVAKA-    
 220 UniRef50_Q9ZVX5        Pentatricopeptide repeat-co...   419    1e-115  1  13.2%  33:601    83:678     --------------------------------SVVRSLLSHHKFADAKSLLVSYIrlHLSPPPSKALFDIALSAYLHEGKPHVALQIFQKMIRLKLKPNLLtnTLLIGLVrsSFSISSAREVFDDMVKIGVsfNVLVNGYcgKLEDALGMLERMVSekVNPDNVTYNTILKAMSkgRLSDLKELLLDMKKNGLVPNRVTYNNLVYGYCKLGSLKEAFQIVELMKQTNVLPDLCTYNILINGLCNAGSMREGLELMDAMKSLKLQPDVVTYNTLIDGCFELGLSLEARKLMEQMENDGVkkVKELVDMHGFSPDIVTYHTLIKAYLKVGDLSGALEMMREMGQKG--IKMNTITLNTILDALCKERKLDEAHNLLNSAHKRGFIVDEVTYGTLIMGFFREEKVEKALEMWDEMKKVKITpsTFNSLIGGLCHHGKTELAMEKFDELAESGLL-PDDSTFNSIILGYCKEGRVEKAFEFYNESIKHSFKPDNYTCNILLNGLCKEGMTEKALNFFNTLIEERE-----------VDTVTYNTMI------------SAFCKDKKLKEAYDLLSEMEEKGLEPDR-FTYNSFISLLMEDGKLSE--    
 221 UniRef50_Q0DQQ9        Os03g0569800 protein n=7 Ta...   419    1e-115  1  11.5%  28:602    79:620     ---------------------------VVAYNAMVAGYCRAGQLAAARR-----LAAAVPVPPNAYTFFPVVRGLCTRGRIADALEVLDEMSFKGCAPIPPMYHVILEAACRSGGFRNSVRVLEAMHAKGCtcNLVLNAIcgCVDEAVGLLRKLAFFGCEADIVSYNAVLKGLCMarWGDVEELMDEMVRVDCAPNIVTFNTLIGYLCRNGLFERVHEVLAQMSEHGCTPDIRMYATIIDGICKEGHLEVANEILNRMPSYGLKPNVVCYNTVLKGLCSAERWKEAEELLSEMFQKD------------CPLDDVTFNILVDFFCQNGLVDRVIELLEQMLSHGC--MPDVITYTTVINGFCKEGLIDEAVMLLKSMSSCGCKPNTVSYTIVLKGLCSAGRWVDAEELMSQMIQQGCPPntFNTLINFLCKKGLVEQAIELLKQMLVNGC-SPDLISYSTVIDGLGKAGKTEEALELLNVMVNKGISPNTIIYSSIACALSREGRVNKVIQMFDNIKDTTIR-SDAVLYNAVISSLCKRWE---------------------TDRAIDFFAYMVSNGCMPNE-STYTMLIKGLASEGLAKEA-    
 222 UniRef50_D1HSM7        Whole genome shotgun sequen...   419    1e-115  1  11.6%   5:589  415:1003     ----ALVHFYGSIGlaQKLFEEMPDHNVVSWTSLMVGYSDSGNPGEVLNVYQRMRQEGVSGNQN--TFATVTSSCGLLEDQVLGYQVLGHIIQYGFEDSVSVANSLISMFSSFSSVEEACYVFDHMNedIISWNAMISAYAHHgrESLRCFHWMRHLHNETNSTTLSSLLSVCSSVdlKWGRGIHGLVVKLGLDSNVCICNTLLTLYSEAGRSEDAELVFQAMTER----DLISWNSMMACYVQDGKCLDGLKILAELLQMG-KPDRVTWNALIGGHAENEEPNEAVKAYKLIREKggQQLHGLVIKLGFESDLHVTNAAMDMYGKCGEMHDVLKML------PQPINRSRLSWNILISAFARHGCFQKARETFHEMLKLGPKPDHVTFVSLLSACNHGGLVDEGLAYYDSmrEFGVFpeHCVCIIDLLGRSGRLSHAEGFIKEMP----VPPNDLAWRSLLAACRIHGNLELARKTAEHLLELDPS-DDSAYVLYSNVCATSGKWEDVENLRKEMGSNNIKKQPA----------CSWVKLKDKVHSFGMGEKYHPQASRISAKLGELMKMTKEAGYVPDTSFALHDM--------------    
 223 UniRef50_D1HSX8        Whole genome shotgun sequen...   419    1e-115  1  11.5%  32:596     2:538     -------------------------------NIVLKGLCRNGGVFEAMGLIREMG--RKSVSPDIVSYNTLINGLCKAKKLKEAVGLLLEMEAAGCFPNSVTCTTLMDGLCKDGRMDEAMELLEAMKKKGFDADVVLYGTlgNLDRGKELFDEMLGKGISANVVTYSCLVHGLCrgQWKEANTVLNAMAEHGIHPDVVTYTGLIDGLCKDGRATHAMDLLNLMVEKGEEPSNVTYNVLLSGLCKEGLVIDAFKILRMMIEKGKKADVVTYNTLMKGLCDKGKVDEALKLFNSMFDNE----------NCLEPNVFTFNMLIGGLCKEGRLTKAVKIHRKMVKKGSCG--NLVTYNMLLGGCLKAGKIKEAMELWKQVLDLGFVPNSFTYSILIDGFCKMRMLNIAKGLFCEMRTHGLNPAlyNTLMASLCKEGSLEQAKSLFQEMGNANC-EPDIISFNTMIDGTLKAGDFQFVKELQMKMVEMGLRPDALTFSTLINRLSKLGELDEAKSALERMVASGFTPDA----------------------LVYDSLLKGLSSKGDTTEIINLLHQMAAKGTVLDR-KIVSTILTCLCHS-------    
 224 UniRef50_C5Y4P7        Putative uncharacterized pr...   418    1e-115  1  11.6%   2:528   186:702     -AACSTLC----LAAFRVVACHGVAPDVRDCNCVLRVLRDAARWDDVRAVYREMLQLE--IEPTIVTYNTLLDSFLKEGRKDEASMLLKEMETQgcLLNDVTYNVVISFLAREGHLENAAKLVDSmkASSFTYNPLITALLERgvEALQM--EMENEGIMPTLVTYNAIIHGLLKSEQVEAAQlfAEMRAMGLLPDLITYNSMLNGYCKAGNLKEALWLLGDLRRAGLAPTVLTYNTLIDGYCRLGGLEEARRLKEEMVEQGCFPDVCTYTILMNGSHKVRNLPMAREFFDEMLSK------------GLQPDCFAYNTRICAELILGDTHKAFQLREVMMLKG--IYPDTVTYNVIIDGLCKTGNLKDAKDLKMKMVSDGLQPDCITYTCLIHAHCERGLLSEARKLLNGMVSDGLQPstYTILIHTCCRRGNLYSAYGWFRKMLDVGI-EPNEITYNVLIHALCRTGRTLLAYHHFHEMLERGLAPNKYTYTLLIDGNCREGNWADAIRLYFEMHQNGIPPDYCTH---------------------------------------------------------------------------    
 225 UniRef50_A5BUK5        Putative uncharacterized pr...   417    1e-115  1   9.8%  61:601    20:548     ------------------------------------------------------------PNPNSITLnsLIDQCSETKQL---KQIHAQMLRTGLFFDPFSASRLITAAALSpsLDYAQQVFDQihPNLYTWNTLIRAYAsnPHQSLLIFLRMLHQSPdpDKFTFPFLIKAASELelFTGKAFHGMVIKVLLGSDVFILNSLIHFYAKCGELGLGYRVFVNXPRR----DVVSWNSMITAFVQGGCPEEALELFQEMETQNVKPNGITMVGVLSACAKKSDFEFGRWV------------HSYIERNRIXESLTLSNAMLDMYTKCGSVEDAKRLFDKMPEK------DIVSWTTMLVGYAkhGHGKDAIALFSKMQEDKVKPNAVTFTNILCACSHVGLVEEGRTFFNQMevYGVLPghYACMVDILGRAGLLEEAVELIEKMP----MAPAASVWGALLGACTIHENVVLAEQACSQLIE--LEpnHGAYVLLSNIYAKAGKWDRVSGLRKLMRDVGLKKEP----------GCSSIEVDGIVHEFLVGDNSHPSAKKIYAKLDEIVARLETIGYVPNKSH-----LLQLVEEEDVKE--    
 226 UniRef50_A9SZZ6        Predicted protein n=2 Tax=P...   417    1e-115  1  13.2%   5:581   204:760     ----KRGDKHSARAAFESMRAAGIEPNVHSYTNLIHAYAVAQDLRGAIACVEEMEAEGVS--PNAATYSVIISGYGRLGDVEAAERWFQRALSEnhHNDVIYNNIIHAYCKAGNM-ERAEAIMTAMEEQGllYNMLMDGycRAVDKCLNVFRRLKArgLSPTVVSYGCLINLYSkgKMDKALQISNEMEDQGIKHNRKTYSMIIDGYVQLGDTANAFSVFEDMSNAGIKPDGITYNILMNAFCKNGQMNRALELLARMESGDCPPTLRTYTIIIDGFMKIGDLRMAFETVRDMK------------MAGFRPSAATYNVIMHGLAQAGQMDRAASIIDEMVVAG--VHPNERSYTTLIEGYACIGDMGLAFKYFNRIKEVGLKPDVIAYASLLKACCKAGRMQSTLAITAEMAAAGVPMniYNILLDGWAQRGDMWEASDIMQKMRHEGL-TPDIHSYTSFINACCKAGDMLKATETIEQMKQQGVQPNLQAYTTLIHGWASASYPEKALICYDEMKSAGMIPDKPLYHCIMTSLLSRA----------------AVARETVFDGVLRVTSEMVDQGICVD----------------------    
 227 UniRef50_D1IKL2        Whole genome shotgun sequen...   417    1e-115  1  13.4%   2:600   190:761     -AANSMLENAVDVFL--QAKKTGLELSTRSCNFLLKCLAEANRREFLRSLFEEMK--STGPPPNVFTYTIMMNFYCKGndTRQATEILEEMERNGESPTVVTYSTYIYGLCRVGYVESALDFVRSLIsnVYCYNAIIHGLCKKgdEALKVLEEMKSCGISPDVYTYSILIHGFCkgDVEKGLYLIEEMKYSNMEPSLVSYSSLFHGLCKKRLSDISLDIFRDLGAAGYKYDQTAYSILIKGFCMQGDLDSAHKLMEEMVRNNLAPDPSNFESLVHGFCKMGLWVNALEFFNMMLE------------GGILPSIATCNVIIDAHCREGRVEEALNLMNEMQTQG--IFPNLFTYNAVINRLCKERKSERALELFPLMLKRNVLPSVVVYSTLIDGFAKQSNSQKALMLYARMLKIGVTPDmyTILINILCHRSRMCEAYNLFKKMTENG-MTPDKISYTSVIAGFCRIGDMRKAWALFNEMLQRGHLPTVVTYTSLVDGYCKMNRIDIADMLIDEMKRKGITPDVVT----------------------YNVLIAAHRRRGNLDKALEMLNEMKENGVLPDHMT--YMMLEWLLKAKKLK---    
 228 UniRef50_D1HR87        Whole genome shotgun sequen...   417    1e-115  1   9.8%   6:576    48:588     -----MYGKCSRVDLACSVFDRMLERNVVSWTALMCGYLQEGNAKGSLALLCEMGY--SGVKPNEFTFSTSLKACGALGVVENGMQIHGMCVKSGFEWVSVVGNATIDMYSKCGRIGMAEQVFNKMprNLVSWNAMIAGHtgNGRKSLVLFQRMQGQGEVPDEFTFTSTLKACGALgiRGGTQIHASLITRGFPISiiIASAIVDLYAKCGYLFEAQKVFDRIEQK----NLISWSALIQGFAQEGNLLEAMDLFRQLRESVSNVDGFVLSIMMGVFADLALVEQGKQM------------HCYILKVPSGLDISVANSIIDMYLKCGLTEEAERLFSEMQVR------NVVSWTVMITGYGKHGLGEKAIHLFNRMQLDGIELDEVAYLALLSACSHSGLIRESQEYFSRLCNNhmKPnhYACMVDILGRAGQLKEAKNLIENMKLK----PNEGIWQTLLSACRVHGNLEIGREVGEILFRMDTD-NPVNYVMMSNIYAEAGYWKECERVRKLVKAKGLKK----------EAGQSWVEINKEIHFFYGGDDTHPLTEKIHEMLKEMERRVKEE---------------------------    
 229 UniRef50_Q9FIB2        Putative pentatricopeptide ...   417    1e-115  1  11.0%   6:580   358:905     -----MYAKCGSIADARRVFYFMTDKDSVSWNSMITGLDQNGCFIEAVERYKSMR--RHDILPGSFTLISSLSSCASLKWAKLGQQIHGESLKLGIDLNVSVSNALMTLYAETGYLNECRKIFSSmeHDQVSWNSIIGALARSEreAVVCFLNAQRAGQKLNRITFSSVLSAVSSLSFGegKQIHGLALKNNIADEATTENALIACYGKCGEMDGCEKIFSRMAER---RDNVTWNSMISGYIHNELLAKALDLVWFMLQTGQRLDSFMYATVLSAFASVATLERGMEV------------HACSVRACLESDVVVGSALVDMYSKCGRLDYALRFFNTMPVR------NSYSWNSMISGYARHGQGEEALKLFETMKLDgtPPDHVTFVGVLSACSHAGLLEEGFKHFESMSdyGLApeHFSCMADVLGRAGELDKLEDFIEKMPMK----PNVLIWRTVLGACCRAnkAELGKKAAEMLFQ--LEpnAVNYVLLGNMYAAGGRWEDLVKARKKMKDADVKKEAGYSWVTMKDG----------VHMFVAGDKSHPDADVIYKKLKELNRKMRDAGYVP-----------------------    
 230 UniRef50_Q9LW32        Pentatricopeptide repeat-co...   416    1e-114  1  10.1%  27:582    40:572     --------------------------DVFSWNSVIADLARSGDSAEALLAFSSMRKLSLYPTRS--SFPCAIKACSSLFDIFSGKQTHQQAFVFGYQSDIFVSSALIVMYSTCGKLEDARKVFDEikRNIVSWTSMIRGynGNALDAVSLFKDLldDAMFLDSMGLVSVISACSRVPAklTESIHSFVIKRGFDRGVSVGNTLLDAYAKGGevAVARKIFDQIVDK----DRVSYNSIMSVYAQSGMSNEAFEVFRRLVKNKVvfNAITLSTVLLAVSHSGAL------------RIGKCIHDQVIRMGLEDDVIVGTSIIDMYCKCGRVETARKAFDRMKNK------NVRSWTAMIAGYGMHGHAAKALELFPAMIDSGVRPNYITFVSVLAACSHAGLHVEGWRWFNAMkrFGVEpeHYGCMVDLLGRAGFLQKAYDLIQRMKMK----PDSIIWSSLLAACRIHKNVELAEISVARLFELDSS-NCGYYMLLSHIYADAGRWKDVERVRMIMKNRGLVKPPGF----------SLLELNGEVHVFLIGDEEHPQREKIYEFLAELNRKLLEAGYVSNT---------------------    
 231 UniRef50_B9IFS7        Predicted protein n=2 Tax=M...   416    1e-114  1   9.0%   4:603   141:771     ---NAVIAAYIKLgdAYEVFMGMSTPDVVSYSALISSFSKLNRETEAIQLFFRMRI--SGIEPNEYSFVAILTACIRSLELEMGLQVHALAIKLGYSQLVFVANALIGLYGKCGCLDHAIHLFDEmqRDIASWNTMISSLVksYEKALELFRVlqNKGFKADQFTLSTLLTACARCHArqGREIHAYAIRIGLENNLSVSNAIIGFYTRCGSLNHVAALFERMPVR----DIITWTEMITAYMEFGLVDLAVDMFNKMPEK----NSVSYNALLTGFCKNNEGLKALNLFVRMVQEGarQIHGFIIKFGFRSNACIEAALIDMCSKCGRMDDADRMFQSLSTDGG----NSIIQTSMICGYARNGLPEEAICLFYRCQSEgmVLDEVAFTSILGVCGTLGFHEVGKQIHCQALKTGFHAevGNSIISMYSKCYNIDDAIKAFNTMPGH-----DVVSWNGLIAGQLLHRQGDEALAIWSSMEKAGIKPDAITFVLIVSAYksSNLLDECRSLFLSMKMIHDLEPTSEHYASLVGVLGYWGLLEEAEE----LINKMPFDPEVSVWRALLDGCRLHagMEPRDPSTYVLVSNLYAASGRWhrEN    
 232 UniRef50_B9RTF6        Pentatricopeptide repeat-co...   416    1e-114  1   8.4%   4:573    43:634     ---LRLYSKFGAVSYAHKLFDETPEPNSFLWTALIHGFTENNQYENAFAFFIKMHRENIVPL--NFTIASVLKAVSRLGRIKDGDLVYGLAVRCGYEFDLVVKNVMIELFMRCGEMGSARQMFDEmeRDAVSWNSMITGYGNNgdIARKLFDRMEER----NVISWTSMIQGYVkgDLLEARVLFERMPEKDLahDVGTWNLMISGCCKAGEMDAAKEFFDRMQER----NVASWVMIIDGYIKVGDVDAARSVFDQMPEK----NLVAWSTMIGGYAKTGHPYSSLKLYKTFKEQGIksVICDFVGPSLFPNLQVVTSLIDMYAKCGNIERAVQVFEMVDQK------DLHCYSTVITAFANHGLSEDAISLFSEMQKANIKPDGVAFLGVLTACNHGGLVGEGRRLFRQMieYGIQPshYACMVDILGRAGCLEEAHSLICSMP----VAPNATVWGALLSACRVHLNVQLAEAAATELFQ--IEpnSGNYVLLFNIYADAGQWDDFSKVRAMIRENRVRK----------NRGSSWIELGSVIHEFVMGDKSHFDSERIFFMLELLCKDM------------------------------    
 233 UniRef50_D1ICI9        Whole genome shotgun sequen...   416    1e-114  1   9.8%   3:572    43:670     --SNRLIEFYAKCNAirRLFDQMPKRDIYTWNAILGAYCKASELEDAHVLFAEM------PERNIVSWNTLISALTRNgdVECGRRCHGISIKIGLDNNIYVGNALLGMYAKCRCIGDAIQAfdVPEPNEVSFTAMMGGLADsvNEAFRLFRLMLRNRIHVDSVSLSSVLGVCSRGGCGegQQVHCLTIKHGFESDLHLNNSLLDMYAKNGNMDSAEMIFVNMPE----VSVVSWNVMIAGYGQKSQSSKAIEYLQRMQYHGFEPDEITYVNMLVACIKSGDIeeAVKLFREMQFrgGRQVHAVSQKAVFRTDIYLASGLIGMYSKCGKVEMAKRIFDRIAE------LDIVCWNSMMAGLSLNSLDKEAFTFFKKMREKGMFPSQFSYATVLSCCAKLSSLSQGRQVHSQIAREGYMNDAfsALIDMYSKCGDVDAARWVFDMMLGK-----NTVTWNEMIHGYAQNGCGDEAVLLYEDMIGSGEKPDGITFVAVLTACSHSGLVDTGIKIFNSMQQEHGVEPLVDHYTCIIDSLGRAGRL----HEAEVLIDKMPCKYDPIIWEVLLSSC-------------------------------    
 234 UniRef50_Q337S8        Pentatricopeptide, putative...   416    1e-114  1  11.0%  55:591    58:556     ------------------------------------------------------RELAATPRL----YHSLITACARYRSLDDARAIHAHLAGSQFAGSVFLDNSLIHLYCKCGAVADARRVFDGMprDMCSWTSLIAGYAQnpDEALGLLPGMLRGRFKPNGFTFASLLKAaaSASSGIGEQIHALTVKYDWHDDVYVGSALLDMYARCGRMDMAIAVFDQLESK----NGVSWNALIAGFARKGDGETTLLMFAEMQRNGFEATHFTYSSVFSAIAGIGALEQGKWVHAHMIKS------------GERLSAFVGNTILDMYAKSGSMIDARKVFDRVDKK------DVVTWNSMLTAFAQYGLGREAVTHFEEMRKCGVHLNQITFLSILTACSHGGLVKEGKQYFDMMKEYNLEpdHYVTVVDLLGRAGLLNDALVFIFKMPMK----PTAAVWGALLGSCRMHKNAKIGQFAADHVFELDPD-DTGPPVLLYNIYASTGQWDAAARVRKMMKATGVKKEPA----------CSWVEIENSVHMFVANDDTHPRSEEIYKKWEEISIQIRKAGYVPN---TDYVLLH------------    
 235 UniRef50_Q6Z0F9        Os08g0340900 protein n=3 Ta...   415    1e-114  1  10.6%   6:582   265:809     -----LYCKCSRLSAARKLFDCMEYRNLVSWTTMISGYMQNSFNAEAITMFWNMTQAGWQ--PDGFACTSILNSCGSLAAIWQGRQIHAHVIKADLEADEYVKNALIDMYAKCEHLTEARAVFDAleDDAISYNAMIEGYSKNRdeAVNIFQRMRFFSLRPSLLTFVSLLGvsSQLAIELSKQIHGLIIKSGTSLDLYAASALIDVYSKCSLVNDAKTVFNMLHYK----DMVIWNSMIFGHAQNEQGEEAIKLFNQLLLSGMAPNEFTFVALVT----------VASTLASMF--HGQQFHAWIIKAGVDNDPHVSNALIDMYAKCGFIKEGRMLFE------STCGEDVICWNSMITTYAQHGHAEEALQVFRLMGEAEVEPNYVTFVGVLSACAHAGFVGEGLNHFNSMKsyDIEpeHYASVVNLFGRSGKLHAAKEFIERMPIK----PAAAVWRSLLSACHLFGNAEIGRYAAEMALLADPT-DSGPYVLLSNIYASKGLWADVHNLRQQMDSSG----------TVKETGCSWIEVTKEVHTFIVRGREHPEAELIYSVLDELTSLIKNLGYVPDT---------------------    
 236 UniRef50_D1H6T5        Whole genome shotgun sequen...   415    1e-114  1  10.5%  11:602   122:752     ----------GDIGVAQKLFDAMPERDVVSWNSLISGYLHNGDHRKVIDVFLQMGRMGTVF--DRTTFAVVLKSCSSLEDHGGGIQIHGLAVKMGFDCDVVTGSALLDMYAKCKKLDCSIQFFHSmeKNWVSWSAIIAGCVQnlRGGLELFKEMQKAGVGVSQSTFASVFRSCAGLslRLGSQLHGHALKTDFGTDVVIGTATLDMYMKCNNLSDAQKLFNSLpdKGLGLDEVSlnAILDMYGKCGALVEACLVFEEMVSR----DAVSWNAIIAAHEQNGNEEKTLSLF----------IHNRIIKSRLGLDSFVGIALIDMYSKCGMMEKAEKLHDRLAEQ------TVVSWNAIISGFSLQKQSEEAQKTFSKMLEMGVDPDNFTYATILDTCANLVTVELGKQIHAQIIKKELQSDAysTLVDMYSKCGNMQDFQLIFEKAPNR-----DFVTWNAMVCGYAQHGLGEEALKIFEYMQLENVKPNHATFLAVLRACGHMGLVEKGLHYFHSMLSNYGLDPQLEHYSCVVDIMGRSGQVSKALE----LIEGMPFEADAVIWRTLLSICKIHGNvePEDSAAY-VLLSNIYANAGMWNE-    
 237 UniRef50_Q7XD69        Pentatricopeptide, putative...   415    1e-114  1   9.8%   6:584    65:613     -----MYSHCADLASALRLFAAMPRRNAVSWTTLVSGLSQNLMHADALAAFAAMR--RAGVAPTRFALSSAARAAAALGAPLPGAQLHCVGVRLGFDTELFVASNLADMYSKCGLLSEACRVFDQmqKDAVAWTAMIDGYAKNgeAAVLSFRDMKREgvGADQHVFCSVLSA--SGGLKDGwsIHCCVTKAGFELEVAVRNALIDMYAKSMDVESASRVLK------IDpnVVSGTSMIDGYIETDCVEEALVIYVELRRQGVEPNEFTFSSMIKGCAMQALLEQGAQL------------HAQVIKTDLIRDSFVGSTLVDMYGKCGLISLSMQLFNEIEYR------TDIAWNAVINVFAQHGHGREAIQAFDRMIYSGIRPNHIAFVSLLTACSHAGLVDEGLKYFYSMKehGIEpeHYSCIIDTYGRAGRLDEAYKFISEMPIK----PNAYGWCSLLGACRMRGSKELGEVAAQNLMK--LEpnTGIHVSLSGIYASLGQWEDVKAVRKLMRDSRIKKLPGF----------SWVDSNKKTHVFGSEDWSHPQQKDIYEKLEELTTRIKEEGYIPDTSF-------------------    
 238 UniRef50_Q9SVP7        Pentatricopeptide repeat-co...   415    1e-114  1  10.4%   5:589   429:984     ----ALLNLYAKCADIETALDYFLEtnVVLWNVMLVAY---GLLDDLRNSFRIFRQMQIeiVPNQYTYPSILKTCIRLGDLELGEQIHSQIIKTNFQLNAYVCSVLIDMYAKLGKLDTAWDILIRFakDVVSWTTMIAGYTQYndKALTTFRQMLDRGIRSDEVGLTNAVSACAGLqlKEGQQIHAQACVSGFSSDLPFQNALVTLYSRCGKIEESYLAFEQ-TEAG---DNIAWNALVSGFQQSGNNEEALRVFVRMNREGIDNNNFTFGSAVKAASETANMKQGKQV------------HAVITKTGYDSETEVCNALISMYAKCGSISDAEKQFLEVSTK------NEVSWNAIINAYSKHGFGSEALDSFDQMIHSNVRPNHVTLVGVLSACSHIGLVDKGIAYFESmsEYGlkPEHYVCVVDMLTRAGLLSRAKEFIQEMPIK----PDALVWRTLLSACVVHKNMEIGEFAAHHLLE--LEpdSATYVLLSNLYAVSKKWD-ARDlrQKMKEKGVKKEPGQ----------SWIEVKNSIHSFYVGDQNHPLADEIHEYFQDLTKRASEIGYVQDCFSLLNEL--------------    
 239 UniRef50_P0C7Q7        Putative pentatricopeptide ...   415    1e-114  1  12.3%   7:544    67:595     ------IKKDDAIALFQEMIRSRPLPSLVDFSRFFSAIARTKQFNLVLDFCKQLEL--NGIAHNIYTLNIMINCFCRCCKTCFAYSVLGKVMKLGYEPDTTTFNTLIKGLFLEGKVSEAVVLVDRmqPDVVTYNSIVNGICRSGdlALDLLRKMEERNVKADVFTYSTIIDSLCrgCIDAAISLFKEMETKGIKSSVVTYNSLVRGLCKAGKWNDGALLLKDMVSREIVPNVITFNVLLDVFVKEGKLQEANELYKEMITRGISPNIITYNTLMDGYCMQNRLSEANNMLDLMVR------------NKCSPDIVTFTSLIKGYCMVKRVDDGMKVFRNISKRG--LVANAVTYSILVQGFCQSGKIKLAEELFQEMVSHGVLPDVMTYGILLDGLCDNGKLEKALEIFEDLQKSKMdiVMYTTIIEGMCKGGKVEDAWNLFCSLPCKG-VKPNVMTYTVMISGLCKKGSLSEANILLRKMEEDGNAPNDCTYNTLIRAHLRDGDLTASAKLIEEMKSCGFS-ADASSIKMVIDMLLS-GELDK-----------------------------------------------------------    
 240 UniRef50_Q6Z2B6        Os08g0290000 protein n=3 Ta...   415    1e-114  1  11.8%   8:602   165:766     -------SLCRAADAFRVLSSRGAPPSIKTCNAFLEALVRAGQLDAAREVFDEMRESR-NVALNEYSYTAMIKALCKAGKVDAGFEMLAELWRAGLQPTVVTYNVLMDALCKSGRVEEAFRLKGRMEQGGMTPSVVTFGILINGleVGIVLQEMEQLGVSPNEVIYNELIghCRKGHCSQALRLFDEMVLKKMKPTAVTYNLIAKALCKEGEMERAERILEDMLSIGMTVHCGLFNTVVAWLLQrrRLESVVSITNEMVTRGMRPNDPLMTACMRELCKGGKHQEAVGIW------------FKTLNKGLGVNLATSNALIHGLCEGKYMKEATKVIQTMLNKG--IELDSITYNIMIRGCCKDSKMEEAIKLHGDMTRRGFKPDLFTFNTLLHAYCNLGKMEETFHLLDQMKTEGLQPdsYGTIIDGHCKAKDIRKAKEYLTELMDRGLK-PNVFIYNALIGGYGRNGDISGAIDAVETMKSNGIQPTNVTYGSLMYWMCHAGLVEEAKTIFSQARENNVDL-GVIGYTIMIQGYCKLGKMVEAVAYFEEMRSRgySKSGNSEEASKLFDEMVGSGVIPDNI-TYGTLIARCSEVNSLDKD-    
 241 UniRef50_D1IID6        Whole genome shotgun sequen...   415    1e-114  1  10.0%  23:574     1:504     ----------------------MPHRDTVSWNSIITGYWKNGCFDESKRLFGLM------PTKNVVSWNSMIAGCIEDERIDEAWQYFQAMPQRNTASW---NAMISGLVRYDRVEEASRLFEEMPRRNVISYTAMvyAKIGEIEQARALFNCMPQK----NVVSWTVMISGYVegKFDEAENLFEQMPDK----NIVAMTAMITGYCKEGKTDKAKILFDQIPCR----DLASWNAMITGYAQNGSGEEALKLHSQMLKMGMQPDHSTLISVLTACSSLASLQEGRK------------THVLVLKSGYESRISICNALITMYCKCGSILDSELAFRQID------HPDVVSWNAMIAAFARHGFYDRALASFGEMRSNRVEPDGITFLSLLSACGHAGKVHESLNWFNSMIeyKirPEHFACLVDILSRGGQVEKAYKIIQEMP----FEADCGIWGALLAACHVHLNVKLGELAAKKIVE--LEpnSGAYVVLSNIYAAAGMWGEVTRVRGLMREQGVKKQPAY----------SWMEIDNKVHFFLGDDASHPEIHRIRLELKGMKLQMI-----------------------------    
 242 UniRef50_D1I9X1        Whole genome shotgun sequen...   415    1e-114  1   9.6%   6:589   156:755     -----LYSKNGHVDLAKLVFERLFLKDSVSWVAMISGLSQNGREDEAILLFCQMHKSAVIPTP--YVFSSVLSACTKIELFKLGEQLHGFIVKWGLSSETFVCNALVTLYSRWGNLIAAEQIFSKmrRDRISYNSLISGLAQRgdRALQLFEKMQLDCMKPDCVTVASLLSACASVGAgkGKQLHSYVIKMGMSSDLIIEGSLLDLYVKCFDIETAHEYFLTTETE----NVVLWNVMLVAYGQLGNLSESYWIFLQMQIEGLMPNQYTYPSILRTCTSLGALDLGEQI------------HTQVIKSGFQFNVYVCSVLIDMYAKHGELDTARGILQRLREE------DVVSWTAMIAGYTQHDLFAEALKLFQEMENQGIRSDNIGFSSAISACAGIQALNQGQQIHAQSYISGYSEdaSNVLITLYSKCGSIEDAKREFFEMPEK-----NVVSWNAMITGYSQHGYGSEAVSLFEEMKQLGLMPNHvtYVLLSNMYAVSGKWDYRDRTRQMMKDRGVKKEPGR----------SWIEVKNSIHAFFVGDRLHPLAEQIYEYIDDLNERAGEIGYVQDR---YNLL--------------    
 243 UniRef50_B9RDL5        Pentatricopeptide repeat-co...   415    1e-114  1  10.3%   7:520    32:554     ------LSKSGRIDEARHIFDKLPERDEFSWNTMIAAYANSGRLTDaiEAFELYLQMQFEgrPNHYTLGSILRLCSRTGFLQKGERIHGYAIKTRLDANVFVVTSLVDLYAKCECISEAEYIFEMvsKNHATWTAMLTGYSHNGdrAIKCFRDMRAEGVESNQFTFPSILTAcaVSDHVFGVQVHGCIVRSGFGANIFVQSALVDMYAKCRDLNSANRVLENMEVN----DVVSWNSMIVGCVREGFEAEALSLFNEMHARDIKIDDYTYPSVLNSFASMKDIQSAKS------------AHCLIIKTGFGAYTLVNNALVDMYAKQGNLDYAFTVFNQMPDK------DVVSWTSLVTGYSYNGSYEDAIKLFCDMRIAGVCPDQIVLASVLSACAELTVTEFGQQVHATILKSGlsLSIDNSLVTMYAKCGGIEDANYVFDSMQIR-----DVISWTALIVGYAQNGRGKDSLHFYDQMIATGTKPDFITFIGLLFACSHTGLVEDGRRYFESMDKEY-----------------------------------------------------------------------------------    
 244 UniRef50_Q9LNU6        Pentatricopeptide repeat-co...   415    1e-114  1   9.5%   3:581    53:672     --SAKLIASYSnfNDADLVLQSIPDPTIYSFSSLIYALTKAKLFTQSIGVFSRM--FSHGLIPDSHVLPNLFKVCAELSAFKVGKQIHCVSCVSGLDMDAFVQGSMFHMYMRCGRMGDARKVFDRmdKDVVTCSALLCAYARKGceVVRILSEMESSGIEANIVSWNGILSGFNRSglNMGRLIHGYVIKQGLLKDKCVISAMIDMYGKSGhvDKALEMFELFKEQTMELNVVSWTSIIAGCAQNGKDIEALELFREMQVAGVKPNHVTIPSMLPACGNIAALGHGRS------------THGFAVRVHLLDNVHVGSALIDMYAKCGRINLSQIVFNMMPTK------NLVCWNSLMNGFSMHGKAKEVMSIFESLMRTRLKPDFISFTSLLSACGQVGLTDEGWKYFKMmeEYGIKpeHYSCMVNLLGRAGKLQEAYDLIKEMP----FEPDSCVWGALLNSCRLQNNVDLAEIAAEKLF--HLEpnPGTYVLLSNIYAAKGMWTEVDSIRNKMESLGLKKNP----------GCSWIQVKNRVYTLLAGDKSHPQIDQITEKMDEISKEMRKSGHRPN----------------------    
 245 UniRef50_Q2QTL4        Os12g0289800 protein n=5 Ta...   415    1e-114  1   8.3%   3:603    45:673     --YNAMVSAlgRLAEARRLFDEMPRRNPVSWNTMMVACSQHGRVEDARGLFDAM------PARNEYSWTIMVSCYVRAGELTLARELLDRM--PGEKCAACYNTMISGYAKNGRFEDAIALLQEMPAPDIVSWNSVLGGLIRNEerSVQFFDEMPDK----DLVSWNLMLEGYVrgDLDVASAFFSRIP----SPNVISWVNLVNGYCQAGRMGEARELFDRMPER----NVVAWNVLLSGYVQFSQVEAAYNLFIEMPEK----NSISWTTMVSGFVRSGKLQEAKDVLSKMPSDNVGAKTALmfDGIVVRDAVCWNTMISGYVQCGMLDEAMVLFQQMPNK------DMISWNTMIAGCAQGGQIRKAASIFRKMKRR----NTVSWNSIISGFVQNGLFVEALQHFMLMRRDAKSAdtynALISAYAKCGRMLEARQVFDEMVVQ-----DIVSWNALIDGYASNGNGSEVIAVFREMEANSVRPDEITLVVVLSACSHAGLIDEGLHFFNSMIKLYSLKPVAEHYTCMVDLLGRAGRLREAFELVQGMQIqhKNHEIAWLAAEKLFE------LEPCKASNYVLLSNICVEAGKWDDAD    
 246 UniRef50_C5YWY0        Putative uncharacterized pr...   414    1e-114  1  11.5%  11:567   120:686     ----------GSRAAVHMMADDgcPPNVLSYNMVINGLFKEGEVDKAYTLFHEM--LGQGFPPNIVTYNSVIDGLCKagQLEEAVRLLKKMSGGGLQPDVVTYSLLIDYYCKIGRCTEARNIFDSMVRRGQNPDAYTYRTLLHgkGAldLLALMIQDGIPLEDHVFNILIRAYAKNelDKAMTAFIEMRQKGFSPNVVTYTTVIDILCKAGRVEDAVSHFSQMVSEGLSPDIITFTSLIHGLCTIGEWKKVEKLSFEMINRGIHPNAIFLNTIMDSLCKEGRVLEAHDFFDQ------------IIHVGVKPDVVSYTILIDGYCLDGKMDESIKLLGRMVSIG--LRPDNVTYSALLNGYCKNGRVDDALALYREMFSKDVKPNAITYNIILHGLFHAGRVVAAREFYMKIVDSGIqiNTYNIVLGGLCENSFVDEALRMFQGLRSKEFQLE-VRTFNIMIVGLLKVGRIGDAKSLFSAILPSGLVPDAIIYGLMIQSHIEEGLLEESDELFLSMEKNGCTANSRT-LNAIVRKLLEKGDVRRAGTYLTKIDEKEYSVEASTAVLL------------------------------------    
 247 UniRef50_B9HAZ8        Predicted protein n=3 Tax=M...   414    1e-114  1  11.9%   6:569    47:576     -----LYAKCRKLREARQVFDEMSERNLVSWSAMISGYEQ---IGEPISALGLFSKLNIV--PNEYVYASVISACASLKGLVQGKQIHGQALKFGLDSVSFVSNALITMYMKCGKCSDALLAYNelELNPVAYNALITGFVEnpDKGFEVLRMMYQDGFFPDRFTFVGLLGTcsRDDLKRGELLHCQTIKLKLNSTAFIGNLIITMYSKLNLLEEAEKAFRSIEEK----DLISWNTFISSCSHCNDHEKALEAFKEMleCRVRPDEFTFASALAACSGLASM------------CNGKQIHGHLIRTRLYQDVGAGNALINMYAKCGCIAKAYYIFSKMEHQ------NLVSWNTMIAGFGNHGFGGKAFELFAKMKTMGVKPDSVTFVGLLTASNHAGLVDEGLVYFNSMEeyGISpeHFSCLIDLLGRAGRLNEAKEYMKKFP----FGHDTVVLGSLLSACRLHGDVDTGKCFARQLLK--LQptTSPYVLLSNLYASDEMWDGVAEAWKLLKGSGLKKEPGH----------SLIEVNGTFEKFTVVDFSHSRIEEIMDMLKIL----------------------------------    
 248 UniRef50_Q0IZR0        Os09g0555400 protein n=4 Ta...   414    1e-114  1  10.6%  16:563    63:609     ---------------ARQLFDRIPDPDRFVYNSLIRAYCNSHCPQEALPLLRGM--IRRGILPNEFTLPFLLKACARVQAWEHVMVTHGVVVKLGFVGQVFVGNALLHSYASAGSLGDSRRFFDEmdRNVVSWNSMINGyaGNTREACSLFEGMRRQGLLADEFTLVSLLFAcaEGNLEFGKLVHSHLLVRGCRIDLILANALVDMYGKCGDLLMAHTCFDMMPFKNaeKSIISWNAMISCYVQGGRFHEALDLYNRMKLLGLAPDEFTLAAVLSACGQLGDLASGKMIHD------------CIRDNFHNPGVALFNSLLDMYARCGQVDTAISLFSEMPSK------NVISWNAIIGALAMHGRAQDALMFFRSMVFDAFPPDEITFVALLSACNHGGLLEAGQYYFQAMRhyNVKpeHYACMVDLLGRGGQLAKAVDLIKDMPMR----PDVVVWGALLGACRIHGHIQIGKQVIKQLLELE-GMSGGLFVLISNMLYETHQWEDMKRLRKLMREWGMKK----------NMGVSSIETNSNIHESGAEGIGHESSDDMY----------------------------------------    
 249 UniRef50_P0C8Q2        Pentatricopeptide repeat-co...   414    1e-114  1   9.0%   6:569    96:630     -----MFVKCNSVDYAAKVFERMPERDATTWNAMLSGFCQSGHTDKAFSLFREMRL--NEITPDSVTVMTLIQSASFEKSLKLLEAMHAVGIRLGVDVQVTVANTWISTYGKCGDLDSAKLVFEAidRTVVSWNSMFKAYSVFGedAFGLYCLMLREEFKPDLSTFINLAASCQneTLTQGRLIHSHAIHLGTDQDIEAINTFISMYSKSEDTCSARLLFDIMTSRTC----VSWTVMISGYAEKGDMDEALALFHAMIKSGEKPDLVTLLSLISGCGKFGSLETGKWI------------DARADIYGCKRdvMICNALIDMYSKCGSIHEARDIFDNTPEK------TVVTWTTMIAGYALNGIFLEALKLFSKMIDLDYKPNHITFLAVLQACAHSGSLEKGWEYFHIMKqyNISpdHYSCMVDLLGRKGKLEEALELIRNMSAK----PDAGIWGALLNACKIHRNVKIAEQAAESLF--NLEPQMAayVEMANIYAAAGMWDGFARIRSIMKQRNIKKYPGESV----------IQVNGKNHSFTVGEHGHVENEVIYFTLNGL----------------------------------    
 250 UniRef50_C5YIQ0        Putative uncharacterized pr...   414    1e-114  1  11.8%  62:580    69:561     -------------------------------------------------------------RADPVSLTRLVKLCVRHGTAGDGRLIHRHVEAHGQLShfVSNSLVSMYAKFGLLDDALRLFDRMPERNVVTWTTVVAALagRKEEALRFLVAMWRDGVAPNAYTFSSVLGACGTPGVLAA--LHASTVKVGLDSDVFVRSSLIDAYMKLGDLDGGRRVFDEMVTR----DLVVWNSIIAGFAQSGDGVGAIELFMRMKDAGFSANQGTLTSVLRACTGMVMLEAGRQVHAHVLK--------------YERDLILHNALLDMYCKCGSLEDAEALFHRMPQR------DVISWSTMISGLAQNGKSAEALRVFDLMKSEGVAPNRITMVGVLFACSHAGLVEDGWYYFRSMKKlgIQpeHHNCMVDLLGRAGKLDEAVEFIRDM----NLEPDAVIWRTLLGACRMHKSGNLAAYAAREILK--LEPDdgARVLLSNTYADLRQWTDAEKSWKAMRDRGMKKEPGR----------SWIELEKHVHVFIAGDLSHPCSDTIVQELNRLIGRISALGYVP-----------------------    
 251 UniRef50_Q9FHF9        Pentatricopeptide repeat-co...   414    1e-114  1  11.5%   7:581    76:609     ------YTRSNRLVDALNLFDEMPVRDVVSWNSMISGCVECGDMNTAVKLFDEM------PERSVVSWTAMVNGCFRSGKVDQAERLFYQMP---VKDTAAWNSMVHGYLQFGKVDDALKLFKQMPGKNVISWTTMICGLDQNEreALDLFKNMLRCCIKSTSRPFTCVITACANaaFHMGIQVHGLIIKLGFLYEEYVSASLITFYANCKRIGDSRKVFDE----KVHEQVAVWTALLSGYSLNKKHEDALSIFSGMLRNSILPNQSTFASGLNSCSALGTLDWGKEM------------HGVAVKLGLETDAFVGNSLVVMYSDSGNVNDAVSVFIKIFKK------SIVSWNSIIVGCAQHGRGKWAFVIFGQMIRLNKEPDEITFTGLLSACSHCGFLEKGRKLFYYM-SSGikIQHYTCMVDILGRCGKLKEAEELIERMVVK----PNEMVWLALLSACRMHSDVDRGEKAAAAIFNLDSK-SSAAYVLLSNIYASAGRWSNVSKLRVKMKKNGIMKKPGS----------SWVVIRGKKHEFFSGDQPHCSR--IYEKLEFLREKLKELGYAPD----------------------    
 252 UniRef50_B7F8W3        cDNA, clone: J065183B01, fu...   414    1e-114  1  11.8%   4:588   198:755     ---AGMYAENADVAAATRVLDAMGAGSVVPWNAVVACCARLGLVDDALELAARMS--RSGPEPNVATWNTVLSGCSRHGRDREALGVVASMLKQGLRPDATTVSSLLKSVANTGLLRHGMEIHCFFLRNQLEPDVYtyAKCGRLDCAQKVLDALEHR----NLTTWNSLVAGYAngRFDIALELVELMKKNRLDPDITTWNGLITGYSMNGQSSQAVLLLRQIKAAGVTPNVVSWTSLISGSCHNGEYEDSFYFCHEMQKDGVQPSLVTMSVLLRACAGLALQKKGKELHCFALRRAYDC------------DMVVSTALIDMYSKGGSLVSAKVIFESIQQK------NLVLCNAMLTGLAVHGQGREAIELFHDMWNSGLKPDSITFTALLTACRSMGLVTEGWEYFDSMekYGVKPTtyACMVDLLARCGYLDEAMDFI----ERSPIDPGASHWGALLTGCSIHGNLALAEVAARNLFI--LEpnSANYLLMMNLYEYERMYDEAESLKYAMKARGVDSRP----------GWSWIQIEQGIHVFEVDGKPHPETAEIYEELIRLVFQIKKAGYVPDTScaYNV---------------    
 253 UniRef50_D1IYF0        Whole genome shotgun sequen...   414    1e-114  1  10.0%  71:580    42:525     ----------------------------------------------------------------------IISLLQRSRHINQVLPIHAQLIRNGHSQDPFMVFELLRSCSKCHAIDYASRIFqtHNPNVYLYTALIDGFvgNYLEAIQLYSRMLHESILPDNYLMASILKAcsQLALREGREVHSRALKLGFSSNRLVRLRIMELYGKCGELGDARRVFEEMPEDVVAKDTVCWTAMIDGFVRNEETNRALEAFRGMQGENVRPNEFTIVCVLSACSQLGALEIGRWVHSYMR------------KFEIELNLFVGNALINMYSRCGSIDEAQTVFDEMKDR------DVITYNTMISGLSMNGKSRQAIELFRVMVGRRLRPTNVTFVGVLNACSHGGLVDFGFEIFHSmrDYRVEpeHYGCMVDLLGRVGRLEEAYDLIRTMK----MTPDHIMLGTLLSACKMHKNLELGEQVAKELEDRG-QADSGTYVLLSHVYASSGKWKEAAQVRAKMKEAGMQKEP----------GCSSIEVNNEIHEFLLGDLRHPQKERIYEKLEELNRLLRLEGYHP-----------------------    
 254 UniRef50_Q9SJZ3        Pentatricopeptide repeat-co...   413    1e-114  1   8.9%   3:576    93:668     --AFCALSESRYLDYSVKILKGIENPNIFSWNVTIRGFSESENPKESFLLYKQMLRHGCCerPDHFTYPVLFKVCADLRLSSLGHMILGHVLKLRLELVSHVHNASIHMFASCGDMENARKVFDepVRDLVSWNCLINGyiGEAEKAIYVYKLMESEGVKPDDVTMIGLVSSCslGDLNRGKEFYEYVKENGLRMTIPLVNALMDMFSKCGDIHEARRIFDNLEKR----TIVSWTTMISGYARCGLLDVSRKLFDDMEEK----DVVLWNAMIGGSVQAKRGQDALALFQEMQTSngIWIHRYIEKYSLSLNVALGTSLVDMYAKCGNISEALSVFHGIQTR------NSLTYTAIIGGLALHGDASTAISYFNEMIDAGIAPDEITFIGLLSACCHGGMIQTGRDYFSQMKSrnLNPqhYSIMVDLLGRAGLLEEADRLMESMP----MEADAAVWGALLFGCRMHGNVELGEKAAKKLLELDPS-DSGIYVLLDGMYGEANMWEDAKRARRMMNERGVEKIP----------GCSSIEVNGIVCEFIVRDKSRPESEKIYDRLHCLGRHMRSS---------------------------    
 255 UniRef50_UPI000198324E PREDICTED: hypothetical pro...   413    1e-114  1  12.0%  31:580    33:583     ------------------------------YSSLISKCITTRHLKLGKLLHSHF--IKTALIFTTFLANRLIDFYSKCDSIQSAHKVFHVLPVKNTHSW---NIIISAYSRSGLFNEAHNLLDQMPKPNLVSYNSLISGLGHHgkESLNVFKTMLksNVLFDEFTLVSLVGSCASLGAPElrQVHGAAIIIGLNSNIIIGNALIDAYGKCGEPDISFSIFSRMPERDveKNTVSWTALIAGFAQNGRGDEALHLFEQMREEGIPPSAFTFASVLSACADLALIARGKEIHGHIIRSTCIDYFC---------NIFILNALIDMYCKCGQMRSATTLFKGMHEK------DIVSWNSLITGFAQNGHGEESLAVFERMIEADIRPNHVTFLGLLSACCHTGLVSEGLRILDSMEkyGVCpdHYAIMIDLLGRNNRLEEAMGLIKRAPK---GSDHVGMWGALLGACRIHGNMDLARRAAEVLFQ--LEpnAARYVMVYNIYAAASRWDEARQVRRLMMERGLRK----------EAACSWIEVRNTRHQFVAKERSHCQINEVYEVIHNLVDQMYDSGYLP-----------------------    
 256 UniRef50_D1HEB5        Whole genome shotgun sequen...   413    1e-113  1   8.9%   7:578   105:679     ------YGKCSCLDDARELFEEMPQRDGGSWNAMITAYAQGGCAEKALWLFSRMNRLGIW--ANEITFASVLGSCATVLALFLSKQIHGLIVKYGFCWNVILGSSLVDIYGKCRVMSDARRMFDEIepNAISWNVIVRRYlgNEKEAVVMFFKMIRANIRPLNFTFSNALIACSssALQEGIQIHGVAIRIGYDEDEVVSSSLIDMYAKCGDLESACRIF-ELPSSK---NLISWTSIVSGYAMSGQTREARVLFDEMPER----SVISWNAMLAGYTHFCQWEEALEFVFLMRkeSGKQVHGFIYRHGLYSNLFVGNALLHMYGKCGNLRSTRLWFYQM-----SHWRDRISWNALLTSHARHGLSEEAMTIFGEMQWE-TTPSKFTLGTLLSACANIFALEQGKQIHGFMIRNGYEIDVvgALVDMYSKCRCLEYALKVFKEAPSR-----DLILWNSMILGCCHNGRGRDVLGLFGLMEEEGVKPDHITFQGILLGCICEGLAGLGTEYFNSMSNKYCIIPRLEHYESMIELYGR----HGFMDELEDFIKRMPFEP-TVAMLTRVFNACSEHGH-------------------------    
 257 UniRef50_Q8S1M4        Os01g0715900 protein n=6 Ta...   413    1e-113  1   9.7%   6:581    28:601     -----LLSDASTSRAARHLFDAVPRPTPALCGTLISALSRLCSHQELLDAFSSLHRRGSDVPPGCV--PLVVKSCAILAASRQGKQVHCHAIVRGLLGDIFVQTALVDFYAKNGDMDCAVKVFDEmrRTSASWNSMIACYagEYQEALRLFRRMLSEGARPNAITIATMFSICAkgDLETGKWARSLIAEQDLQ-NMIVHTALMEMYVKCRAIDEARREFDRMQQR----DVVAWSTMIAGYAQNGRPHESLELFERMKATSCKPNEVTLVGVLSACAQLGS------------DELGGQIGSHVESQNLPLTSYLGSALIDMYTKCGHVGRARSVFNRMEHKV------VITWNSMMRGLALNGFAQDAITLYKEMTEEDVQPNEITFVALLTACTHAGLVDQGMSFFKEMKthHVSpeHCACIVDLLCKSGRLREAYKFICDME----VEPNAVIWSTLLSACRAHADVELAKLAASKLLV--LEPdsSIYVLLSNIYADAGLWGDVREIRDLMRSKNVQKLSAY----------SWIELDGEVHKFLVQDTYHPKSAEIFNVVDGMGLHLDDVDSDPD----------------------    
 258 UniRef50_Q9ZQ74        Pentatricopeptide repeat-co...   413    1e-113  1  11.7%   2:523   181:681     -GLLDMYAKCGEIKSAHKVFNDITLRNVVCWTSMIAGYVKNDLCEEGLVLFNRMRENNVLG--NEYTYGTLIMACTKLSALHQGKWFHGCLVKSGilSSCLVTSLLDMYVKCGDISNARRVFNEHSHVDLVMWTAMIVGYTHNgnEALSLFQKMKGVEIKPNCVTIASVLSGCGleNLELGRSVHGLSIKVGI-WDTNVANALVHMYAKCYQNRDAKYVF-EMESEK---DIVAWNSIISGFSQNGSIHEALFLFHRMNSESVTPNGVTVASLFSACASLGSLAVGSSLHAYSVKLGFLAS----------SSVHVGTALLDFYAKCGDPQSARLIFDTIEEK------NTITWSAMIGGYGKQGDTIGSLELFEEMLKKQQKPNESTFTSILSACGHTGMVNEGKKYFSSmkDYNFTPshYTCMVDMLARAGELEQALDIIEKMP----IQPDVRCFGAFLHGCGMHSRFDLGEIVIKKMLD--LHPDDASyvLVSNLYASDGRWNQAKEVRNLMKQRGLSK--------------------------------------------------------------------------------    
 259 UniRef50_Q0WPZ6        Pentatricopeptide repeat-co...   413    1e-113  1  11.6%  17:600   136:750     ----------------KDMVLCGIAPQTYTFNLLIRALCDSSCVDAARELFDEMPEKGC--KPNEFTFGILVRGYCKAGLTDKGLELLNAMESFGVLPNKVIYNTIVSSFCREGRNDDSEKMVEKMrpDIVTFNSRISALCKEGkdASRIFSDMELDEyrPNSITYNLMLKGFCKVglEDAKTLFESIRENDDLASLQSYNIWLQGLVRHGKFIEAETVLKQMTDKGIGPSIYSYNILMDGLCKLGMLSDAKTIVGLMKRNGVCPDAVTYGCLLHGYCSVGKVDAAKSLLQEMMR------------NNCLPNAYTCNILLHSLWKMGRISEAEELLRKMNEKGYGL--DTVTCNIIVDGLCGSGELDKAIEIVKGMreNNCLPDLITYSTLLNGLCKAGRFAEAKNLFAEMMGEKLQPDsyNIFIHHFCKQGKISSAFRVLKDMEKKGCHK-SLETYNSLILGLGIKNQIFEIHGLMDEMKEKGISPNICTYNTAIQYLCEGEKVEDATNLLDEMMQKNIA-PNVFSFKYLIEAFCKVPDFDMAQEVFETAVSICGQKEGLYskATELLEAVLDRGFELGT-FLYKDLVESLCKKDELE---    
 260 UniRef50_Q8S5M5        Os10g0368800 protein n=2 Ta...   413    1e-113  1  10.3%   3:534   158:670     --ALALYRRM--------VHRDRVPPTTFTFGVAARALCRLGRADEALALLRGMARHGCV--PDAVLYQTVIHALCDQGGVTEAATLLNEMLLMGCAADvtFDDVVRGMCGLGRVREAARLVDRMMTK-GCMPGVMTYGFlrQADEARAMLGRVPE----LNVVLFNTVIGGcaEGKLAEATELYETMGLKGCQPDAHTYSILMHGLCKLGRIGSAVRLLREMEKKGFAPNVVTYTIVLHSFCKNGMWDDTRALLEEMSAKGLTLNSQGYNGMIYALCKDGRMDEAMGLIQEMRSQ------------GCNPDICSYNTIIYHLCNNEQMEEAEHMFENLLEEG--VVANGITYNTIIHALLRDGRWQDAVRLAKEMILHGCSLDVVSYNGLIKAMCKDGNVDRSLVLLEEMAEKGIKPnsYNILISELCKERRVRDALELSKQMLNQGLA-PDIVTYNTLINGLCKMGWMHAALNLLEKLHNENVHPDIITYNILISWHCKVRLLDDAAMLLNRAMAAVCPVGDRRIMQILPD---------------------------------------------------------------------    
 261 UniRef50_O64624        Pentatricopeptide repeat-co...   412    1e-113  1  11.5%   1:602   185:790     LGRESQYSVAAKL--LDKIPLQEYLLDVRAYTTILHAYSRTGKYEKAIDLFERMKEMG--PSPTLVTYNVILDVFGKMGrwRKILGVLDEMRSKGLKFDEFTCSTVLSACAREGLLREAKEFFAELKSCGYEPGTVTYNALLQvtEALSVLKEMEENSCPADSVTYNELVAAYVRAGfkEAAGVIEMMTKKGVMPNAITYTTVIDAYGKAGKEDEALKLFYSMKEAGCVPNTCTYNAVLSLLGKKSRSNEMIKMLCDMKSNGCSPNRATWNTMLALCGNKGMDKFVNRVFREMKSC------------GFEPDRDTFNTLISAYGRCGSEVDASKMYGEMTRAGFNAC--VTTYNALLNALARKGDWRSGENVISDMKSKGFKPTETSYSLMLQCYAKGgrIKEgeRAFTLFKKHGYKPDmfNSMLSIFTRNNMYDQAEGILESIREDGL-SPDLVTYNSLMDMYVRRGECWKAEEILKTLEKSQLKPDLVSYNTVIKGFCRRGLMQEAVRMLSEMTERGIRPCIFT-YNTFVSGYTAMGM---------------------FAEIEDVIECMAKNDCRPNEL-TFKMVVDGYCRAGKYSEA-    
 262 UniRef50_Q9LSL8        Pentatricopeptide repeat-co...   412    1e-113  1  10.5%   9:589   111:659     --------KCGDIDYARQVFDGMSERHIVTWNSLIAYLIKHRRSKEAVEMYRLM--ITNNVLPDEYTLSSVFKAFSDLSLEKEAQRSHGLAVILGlvSNVFVGSALVDMYVKFGKTREAKLVLDRVekDVVLITALIVGYSQKGeeAVKAFQSMLVEKVQPNEYTYASVLISCGNldIGNGKLIHGLMVKSGFESALASQTSLLTMYLRCSLVDDSLRVFKCIE----YPNQVSWTSLISGLVQNGREEMALIEFRKMMRDSIKPNSFTLSSALRGCSNLAMFEEGRQI------------HGIVTKYGFDRDKYAGSGLIDLYGKCGCSDMARLVFDTLSE------VDVISLNTMIYSYAQNGFGREALDLFERMINLGLQPNDVTVLSVLLACNNSRLVEEGCELFDSFRKDKIMldHYACMVDLLGRAGRLEEA-----EMLTTEVINPDLVLWRTLLSACKVHRKVEMAERITRKILE--IEpdEGTLILMSNLYASTGKWNRVIEMKSKMKDMKLKKNPA----------MSWVEINKETHTFMAGDlsHPNSEQILENLEELIKKSKDLGYVEDKSCVFQDM--------------    
 263 UniRef50_D1HI57        Whole genome shotgun sequen...   412    1e-113  1  10.6%   5:563    84:621     ----SMYSKHDRIKEARFLFDDMPERDDVSWNSMIAGYSQRGLNEEACGLFCSmsCENWKLLVSDFTLATVLKACGGLGCSRIGKCVHGYAVKIGFDSDLFVSGSTVYMYCKCGILDMAGLAFDQIekDIVAWNTMITGYAQNceEAIELFYQMELEGFKPNDTTFCCVLKAsaMSDSAVGRCFHAKVLKLGCSMDVFVATALVDMYSKFYDIEDVERAFGEMSKR----NLVSFNALITGYSLMGKYEEALRVYSQLQSEGMEPDSFTFVGLFSSCSVSSTVAEGAQV------------HVHSVKFGLDSDVSVGNSIVNFYSKCGFTDSALEAFESINR------PNSVCWAGIISGFAQNGEGEKALMQFCKMRKFIDKTDEFSSSSVIKAVSSWAAVEQGRHLHAHVMKSGLDCTIysAVIDMYSKCGMVEDAQKVFSVMPEK-----NVVSWNSMITGYAQNGFCKEALLLFQEMTSSGILPTAVTFVGILFACSHAGLVEEGRNFYNLMVHNYGIPPSMEHCTCMVDLLGRAGYLEEA--EAFLLSSSFSKEPGIW----------------------------------------    
 264 UniRef50_Q2HW11        Tetratricopeptide-like heli...   412    1e-113  1  11.8%   6:582    76:668     -----LYVKSSNLDHAHKLFDEITHKNTQTWTILISGFARA--AGSSELVFSLFREMQADGapNQYTLSSVLKCCSRENNIQFGKGIHAWILRNGVGGDVVLENSILDLYLKCKEFEYAESFFELmeKDVVSWNIMIGAYLrdVEKSLEMFRnmVAHGTEFSPVTFSIALILVSSLSLVegRQLHGRVLTFGLNSDGYIRSSLVEMYGKCGRMDKASTILKDVpcKEPKARMVSWSSMVSGYVWNGKYEDGMKTFRSMVCELIVVDIRTVATIISACANAGILEFGKQI------------HAYIQKIGLRIDAYVGSSLIDMYSKSGSLDDALMIFEQIKE------PNVVLWTSMISGCALHGQGKEAISLFEGMLNLGIIPNEVTFVGVLNACSHVGLIEEGCRYFRMMKdyHINpeHYTSMVNLYGRAGHLIEAKNFIFE----NSISHFTSVWRSFLSSCRLHKNFNMGKSV-SEMLLQSAPSDPDAYILLSNMCSSNHQWDEAAIVRSLMYQRGVKKQPGQ----------SWVQLKDQIHSFTVGDRSHPQDKEIYSYLDSLIGRLKEIGYSLDA---------------------    
 265 UniRef50_Q0WMY5        Pentatricopeptide repeat-co...   412    1e-113  1  11.0%   7:581   319:874     ------YGRRGDMHRARemRARGITPTSRIYTSLIHAYAVGRDMDEALSCVRKMKEEG--IEMSLVTYSVIVGGFSKAGHAEAADYWFDEAKRIHKTLNASIYGKIIYAHCQTCNMERAEALVREMEEEGiiYHTMMDGyvADEKKGLVVFKRLKECGFTPTVVTYGCLINLYTkgKISKALEVSRVMKEEGVKHNLKTYSMMINGFVKLKDWANAFAVFEDMVKEGMKPDVILYNNIISAFCGMGNMDRAIQTVKEMQKLRHRPTTRTFMPIIHGYAKSGDMRRSLEVFDMMRRC------------GCVPTVHTFNGLINGLVEKRQMEKAVEILDEMTLAG--VSANEHTYTKIMQGYASVGDTGKAFEYFTRLQNEGLDVDIFTYEALLKACCKSGRMQSALAVTKEMSARNIPRnvYNILIDGWARRGDVWEAADLIQQMKKEG-VKPDIHTYTSFISACSKAGDMNRATQTIEEMEALGVKPNIKTYTTLIKGWARASLPEKALSCYEEMKAMGIKPDKAVYHCLLTSLLSRASIAEAY----------------IYSGVMTICKEMVEAGLIVD----------------------    
 266 UniRef50_Q9M4P3        Pentatricopeptide repeat-co...   412    1e-113  1  12.6%  34:580    64:572     ---------------------------------LNKIIARCVRSGDIDGALRVFHGMRAK---NTITWNSLLIGISkpSRMMEAHQLFDEIPE----PDTFSYNIMLSCYVRNVNFEKAQSFFDRMpkDAASWNTMITGYARRgeKARELFYSMMEK----NEVSWNAMISGYicGDLEKASHFFKVAPVRG----VVAWTAMITGYMKAKKVELAEAMFKDMT---VNKNLVTWNAMISGYVENSRPEDGLKLFRAMLEEGIRPNSSGLSSALLGCSELSALQLGRQI------------HQIVSKSTLCNDVTALTSLISMYCKCGELGDAWKLFEVMKKK------DVVAWNAMISGYAQHGNADKALCLFREMIDNKIRPDWITFVAVLLACNHAGLVNIGMAYFESMVRdePQPDHYTCMVDLLGRAGKLEEALKLIRSMP----FRPHAAVFGTLLGACRVHKNVELAEFAAEKLLQLNSQ-NAAGYVQLANIYASKNRWEDVARVRKRMKESNVVKVERVKVPGY-----SWIEIRNKVHHFRSSDRIHPELDSIHKKLKELEKKMKLAGYKP-----------------------    
 267 UniRef50_Q0DFR6        Os05g0574800 protein n=4 Ta...   412    1e-113  1   9.7%   4:580   135:782     ---NAMLSMLVRFGeaWRVFAKMPERDVFSWNVMVGGYGKVGFLEEALDLYYRMLWAGM--RPDVYTFPCVLRTCGGIPDWRMGREVHAHVLRFGFGDEVDVLNALVTMYAKCGDIVAARKVFDGmvTDCISWNAMIAGHFEncEAGLELFLTMLENEVQPNLMTITSVTVASglSEVGFAKEMHGFAVKRGFAIDVAFCNSLIQMYTSLGRMGDAGKIFSRMETK----DAMSWTAMISGYEKNGFPDKALEVYALMELHNVSPDDVTinALLEMYAKSKHIDKAIEVFKFMAEKdgKEIHAYVLRCGIGSEGYVPNALLDLYVKCGQTSYAWAQFSVHSEK------DVVSWNIMLSGFVAHGLGDIALSLFNQMVEMGEHPDEVTFVALLCACSRAGMVIQGWELFHMMTEksIVPnhYACMVDLLSRVGKLTEAYNLINRMPIK----PDAAVWGALLNGCRIHRHVELGELAAKVILE--LEPNDVAYHVLldLYTDAGKWAQVARVRKTMREKGLEQ----------DNGCSWVEVKGVTHAFLTDDESHPQIKEINVVLHGIYERMKACGFAP-----------------------    
 268 UniRef50_Q6ZDB7        Os08g0300700 protein n=5 Ta...   411    1e-113  1  12.9%   1:598    51:659     LARHRMFPDMESLAS------RMPARNLRTYTTLINAYCLAGDIPAAKQ--HLTSLLHAGLAPDSYAYTSFVLGYCRAGMLTHACRVFVLMPLRGCLRTAFTYTALLHGLLGAGMVREAMAVFVGMRADSCAPDTHVYATMVHgcEaeVLLEEAMSNGFEPNIVVYNALIDGYCNAgmEHALKVFEGMDGNRCSPNVRTYTELIHGLCKSGKVERAMVLFSRMVEAGLEPNVVTYTALIQGQCNEGHLQCAFRLLHLMETNGLVPNDWTFSVLIDALCKREKVEEAQLFLGSLVKK------------GVKVNEVVYTSLIDGLCKTGKIDAADELMQKMISEG--FVPDAHSYSSLIDGLCRQKKLSQATLMLEDMMEKGIQASPVTYTIIIDELVREVGSEGPKKIFDKMIATGINPdtYTVFVRSYCEEGRMEDAESMIVQMVDRGVF-PNLVTYNTLIRGYANLGLVSQAFSTFEVMVGKGWKPNEDSYTVLLRLVVKKSSSDNSVDIWkdMKdeRQLPLAADIYSCFIRCLCRVDRLEEAKHFfyTSIIDCCCRLKILTDALTLLDSMTKSGYLP-HLESYRIIISSLCEGGN-----    
 269 UniRef50_B9I897        Predicted protein n=25 Tax=...   411    1e-113  1  13.3%  19:498     1:475     ------------------MDLSNIRPDVYTLTILINCFCHS-NHDHFHFAFSVLGNmkLGLQPNHVTFSTLLNGLSSKAKIIDAVKLFDEMVKMGYEPDVITYSTIINGLCKMGSTTMAIQLLKKMEEKGCKPNVVVYSTIIDSleAMEFLSEMVNRGISPNVVTYSSILHGFCngRSNEATSLFKQMVERNVMPDTVTFNILVDGLSKEGMILEAQCVFETMIEKGVEPNVNTYNALMDGYCSQSQMDEAQKLFNIMVRKGCAPSVRSYNILIKGHCKSGRIDEAKGLLAEMSHKA------------LTPDTVTYSTLMKGFCQDGRPQDAQKLLEEM--RSYGLLPDLMTYSIVLDGLCKQGHLDEAFELLKAMQESKIEPNIFIYTILIQGMCNFGKLEAARELFSNLFVKGIQPdtYTVMISGLLKGGLSNEACELFRDMAVHGCL-PNSCTYNVIIQGFLRNGDTSNAGRLIEEMVGRGFSADSSTFQMLSD---------------------------------------------------------------------------------------------------------    
 270 UniRef50_B9IEE4        Predicted protein n=3 Tax=r...   411    1e-113  1  11.4%  13:602   257:817     ------------LRFHKALVQGGFGLDIITCNKILKGIWMQNDIGVADDYFNMVVRIG--PKPNVVTFSTLIDAYCKEGNLDKAFVLFDVMAGNGVTPDLIVYSILIDGLFKAGRLEDGQRLlkGIKLDVVGFSSAMDAYvgDLGRVIQIYKRMLNEGISPNVVSCSILIKGFCqgRILEACGLFVQILKLGFEPSILTYSALIAGFCKSGNLRDGFYLYEDMIKKRCEPDTIVYSVLINGLCKQGLVGDALRFFFQAVNRGLSPNVFTLNTLLDSFCRLKCIVGAMKVYYLM------------GMLNIKADTVTYTILIKGAAQFGRVDEALMLFFQMLKK--DFKPDVITYCTLIDGLCKLKKSSAGLCIFDFMCKNAVAPDIAIYNVLINMHSREGHLEAALGLFVHVVERGPKPdtFNTMICCYCNFKRLDDAVQLFAKMTSE-QLRPNAITFTILIDAFCREGRMDDAMLMFSKMLEEGPEPNLVTYSCLIHGYFKSqmMESGLKLYNEMLENNIAPNIVS-YSILIDGLCKRGLMKEASCAFRCALDKH---------------------LLPDVIA-YTILIRGYCKVGRLTEA-    
 271 UniRef50_Q7XUF2        Os04g0488200 protein n=5 Ta...   411    1e-113  1  12.2%  31:578    80:598     ------------------------------FNALISSLTHAGDPSAAFRAFALlsGAGAGARPDGYTLPAALKACARLGglREGCQAHAVAEKAGFLGRVPVQNALVTFYGACGQCGDARKVFDEmeRDVVSWTALLSAFTrmFMEALGVLAEM---DVTPNEVTLASVLVACGNLgaRAGKAVHGWYFRREKELNLIVGNALLDMYVKCEKLDLARRVFDMLLAR----DIVSWTVMISGLVQCKRPSEALEVFNAMQISGVKPDKVVLSTVLSACASLGALESGRWVHEYIERK------------GIEWDVHVGTSVVDMYVKCGCLDTAVSIFQEMPLK------NVSSWNALINGFALHGRGREALDCFDRMVASGLHPNEVTFITVLGACCHSGLVQEGRQLFELmkSYKLSpdHYGCMVDLLGRAGLIQEAYDVIKAMPMR----PGVFTWVTLLSACQAHGRMDFSQQILMHIHELESSGN-GIYVLLSNMYAVSDRWADVGKARGFMNEKGMQKEPGS----------SVIEVNGKTCEFLVGQQNHLYMDDVRAMLSILMKQIHLDGL-------------------------    
 272 UniRef50_Q338A4        Pentatricopeptide, putative...   411    1e-113  1  12.8%  11:598   104:735     ----------GRVGDMAEVFDVmvKANVGTFAAIFGGLGVEGGLRSAPVALPVMKEAGIV--LNAYTYNGLVYFLVKSGFDREALEVYRVMMVDGVVPSVRTYSVLMVAFGKRRDVETVLWLLREMEAHGVKPNVYSYTICIRvdEAYRILAKMENEGCKPDVITHTVLIQVLCdgRISDAKDVFWKMKKSDQKPDRVTYITLLDKFGDNGDSQSVMEIWNAMKADGYNDNVVAYTAVIDALCQVGRVFEALEMFDEMKQKGIVPEQYSYNSLISGFLKADRFGDALELFKHmeSIKAIQRYELMKSKGIVPDVVAGNAVLFGLAKSGRLGMAKRVFHELKAMG--VSPDTITYTMMIKCCSKASKFDEAVKIFYDMIENNCVPDVLAVNSLIDTLYKAGRGDEAWRIFYQLKEMNLEPttYNTLLAGLGREGKVKEVMHLLEEMYHSNY-PPNLITYNTILDCLCKNGAVNDALDMLYSMTTKGCIPDLSSYNTVIYGLVKEERYNEAFSIFCQMKK--VLIPDYATLCTILPSFVKIGLMKEALHIIKDYfsCHSLMEGILKKaiEFAEIIASSGITLDD-FFLCPLIKHLCKQKK-----    
 273 UniRef50_Q336W7        Os10g0540100 protein n=5 Ta...   410    1e-113  1  11.3%  41:589    84:601     ----------------------------------------HSRRGSPASALRVFRALPPAARPDSTTFTLALSACARLGDLRGGESVRDRAFDAGYKDDVFVCSSLLHLYArgAMGDAVKVFVRMPRRDRVTWSTMVAGfaGQPLDAIQMYRRMREDGVKGDEVVMIGVIQAcaARNVRMGASVHGHLLRHGMRMDVVTATSLVDMYAKNGLLDVACRVFGLMVHR----NDVSWSAMISGFAQNGQSDEALRLFRNMQASGIQPDSGALVSALLACSNIGFLKLGRSVHGFIVRR-------------FDFNCILGTAAIDMYSKCGSLASAQMLFNMISDR------DLILWNAMIACCGAHGRGQDALTLFQEMNETGMRPDHATFASLLSALSHSGLVEEGKLWFGRMVNhkITPahYVCLVDLLARSGLVEEASDLLTSMKA----EPTVAIWVALLSGCLNNKKLELGESIADNILE--LQPDDvvLALVSNLYAATKKWDKVRQVRKLMKDSGSKKMP----------GCSSIEIRGTRHVFVMEDQSHPQREEIVSKVAKLDLEMRKMGYIPRTEFVYHDL--------------    
 274 UniRef50_Q0D4K4        Os07g0621100 protein n=5 Ta...   410    1e-113  1  11.5%  35:556    37:566     ----------------------------------LAVLHRRGRAEAAACLNRHLRLLPLGEarDAVSYNTVLTALCRRGHHDRAGALLRAmpHPACRPNAVSYTVLMRALCADRLADQAVGLLRSMRSAGVRADVVTYGTLIRgdKAVELMGEMCESGIEPNVVVYSSLLQGYCkgRWEDVGKVFVEMSEKGIEPDVVMYTGLIDSLCKVGKAKKAHGVMDMMVRRGLEPNVVTYNVLINCMCKEGSVKEAIGVLKKMSEKGVAPDVVTYNTLIKGLSDVLEMDEAMWLLEEMVR----------GKNIVKPNVVTFNSVIQGLCDIGRMRQAFQVRAMMEETGCMV--NLVTYNLLIGGLLRVHKVRKAMELMDEMTSLGLEPDSFTYSILIKGFCKMWQVDRAEDLLSTMRDRGIEPehYIPLLVAMCEQGMMERARNLFNEM--DNNFPLDVVAYSTMIHGACKAGDLKTAKELLKSIVDEGLTPDAVTYSIVINMFAKSGDMEAANGVLKQMTASGF-LPDVAVFDSLIQGYSTKGEINKVLELIREMITKN-----------------------------------------------    
 275 UniRef50_Q9STF3        Pentatricopeptide repeat-co...   410    1e-113  1  10.7%  32:580    50:568     -------------------------------NQLIQSLCKEGKLKQAIRVLSQ------ESSPSQQTYELLILCCGHRSSLSDALRVHRHILDNGSDQDPFLATKLIGMYSDLGSVDYARKVFDKTRKRTivWNALFRalAGHGEEVLGLYWKMNRIGVESDRFTYTYVLKACVanHLMKGKEIHAHLTRRGYSSHVYIMTTLVDMYARFGCVDYASYVFGGMPVR----NVVSWSAMIACYAKNGKAFEALRTFREMMrkDSSPNSVTMVSVLQACASLAALEQGKLIHGYILRRGLDSILP------------VISALVTMYGRCGKLEVGQRVFDRMHDR------DVVSWNSLISSYGVHGYGKKAIQIFEEMLANGASPTPVTFVSVLGACSHEGLVEEGKRLFETmrDHGIKpeHYACMVDLLGRANRLDEAAKMVQDMRT----EPGPKVWGSLLGSCRIHGNVELAERASRRLFA--LEpnAGNYVLLADIYAEAQMWDEVKRVKKLLEHRGLQKLPGR----------CWMEVRRKMYSFVSVDEFNPLMEQIHAFLVKLAEDMKEKGYIP-----------------------    
 276 UniRef50_D1IGJ5        Whole genome shotgun sequen...   410    1e-113  1  10.9%   3:600    59:624     --SNSLMNacGLLADAKQIFHHTPYKNVVSWTILISGLAKNDCFVEAIDVFREMIM--GNFKPNAVTISSVLPAFANLGLIRIAKSVHCFWVRGGFEGNVFVETALVDMYSKFGCMGVARQLFESmeRNVVTWNAIVSGYSDHgeEAIDLFNLMRRKGLLVDFYTIMSLIPalSVGCLQVGTGIHGFIIRTGYENDKHIKTALMDIYVSHNCVDDAHRVFSEMSVK----DVAAWTLMLTGFSSGRHWDRAIKHFNKMLgqNLKLDSIALMGILSSCSHSGALQQGRRV------------HALAIKTCFANNIFVGSAVIDMYANCGNLEDAKRFFYGMGEK------DVVCWNAMIAGNGMNGYGTDAIDLFLQMKGSGLDPDESTFVSVLYACSHAGMVYEGLQIFYHMVksHVIPnhYACVIDILGRAGQLDAAYSFINNMP----FQPDFDVYSTLLGACRIHGNIKLGHEISQKIFE--MEpdAGYYVLLSNMYALAGNWEGVKMTRASLRSKRMKKDPGF----------SSIEINQEIYTFMAGEKDHPQYFKIEGILKGLILKIKKAGYVPNT----NVLLQDVSDDMKKD---    
 277 UniRef50_B9RLG0        Pentatricopeptide repeat-co...   410    1e-113  1  12.4%  18:597   236:820     -----------------EIKVSGTPQSEYTSSIVVDGLCRQSRFQDAVLFFQ--DTEGKEFQPSVVSFNTIMSRYCKLGFVDVAKSFFCMMLKHGLLPDAYSYNILIHGLCIAGSMGEALDLKNDMENHGLEPDMVTYnlLGLINGAWNIIQKMLIKGPNPNLVTYTVLICGHCqgNVEEALKLYKEMISHGFQLSIISSTVLLGSLCKSRQVDVAFKLFCEMEANGLRPDLITYSTLIHGLCKQGEVQQAILLYEKMCSNRIIPNSLIHGAILMGLCEKGKISQARMYFDYLITSN------------LSLDIILYNIMIDGYIKRGNTREAVKLYKQLGEKG--ISPTIVTFNSLMYGFCINRKLSQARRLLDTIKLHGLEPNAVTYTTLMNVYCEEGNMQSLLELLSEMKAKAIGPThyTVVIKGLCKQWKLQESCQLLEDMDAVGL-TPDQVSYNTIIQAFCKARDMRKAFQLYDKMLLHNLEPTSVTYNILINGFCVYGDLKDADNLLVSLQNRKVNLNKYA-YTTIIKAHCAKGDVDKAVVYfySAVIGRLCKRCLVTEAKYFFCMMLSDGVCPDQ-DLFEVLLNAFHQCG------    
 278 UniRef50_O64705        Pentatricopeptide repeat-co...   410    1e-113  1  10.5%  25:579    93:615     ------------------------EPNHYSFNYMIRGLTNTWNDHEA--ALSLYRRMKFSgkPDKFTYNFVFIACAKLEEIGVGRSVHSSLFKVGLERDVHINHSLIMMYAKCGQVGYARKLFDEieRDTVSWNSMISGyaGYAKDAMDLFRKMEEEGFEPDERTLVSMLGACSHLGDLRTGRLLEEmiTKKIGLSTFLGSKLISMYGKCGDLDSARRVFNQMIKK----DRVAWTAMITVYSQNGKSSEAFKLFFEMEKTGVSPDAGTLSTVLSACGSVGALELGKQI------------ETHASELSLQHNIYVATGLVDMYGKCGRVEEALRVFEAMPVK------NEATWNAMITAYAHQGHAKEALLLFDRMS---VPPSDITFIGVLSACVHAGLVHQGCRYFHEMSSmgLVpeHYTNIIDLLSRAGMLDEAWEFMERFPGK----PDEIMLAAILGACHKRKDveKAMRMLMEMKEAK---NAGNYVISSNVLADMKMWDESAKMRALMRDRGVVKTP----------GCSWIEIEGELMEFLAgdYLQCGREDSGSLFDLLVEEMKRERYE------------------------    
 279 UniRef50_Q9FMQ1        Pentatricopeptide repeat-co...   410    1e-113  1  11.5%   1:602   224:789     LCKGKRMNDAEQLFD--EMLARRLLPSLITYNTLIDGYCKAGNPEKSFKVRERMKA--DHIEPSLITFNTLLKGLFKAGMVEDAENVLKEMKDLGFVPDAFTFSILFDGYSSNEKAEAALGVYETAVDSGVKMNAYTCSILLNaeKAEEILGREMAKGLVPNEVIYNTMIDGYckGDLVGARMKIEAMEKQGMKPDHLAYNCLIRRFCELGEMENAEKEVNKMKLKGVSPSVETYNILIGGYGRKYEFDKCFDILKEMEDNGTMPNVVSYGTLINCLCKGSKLLEAQIVKRDMEDR------------GVSPKVRIYNMLIDGCCSKGKIEDAFRFSKEMLKKG--IELNLVTYNTLIDGLSMTGKLSEAEDLLLEISRKGLKPDVFTYNSLISGYGFAGNVQRCIALYEEMKRSGIKPttYHLLISLCTKEG-IELTERLFGEMSLK----PDLLVYNGVLHCYAVHGDMEKAFNLQKQMIEKSIGLDKTTYNSLILGQLKVGKLCEVRSLIDEMNAREMEPEADT----------YNIIVKGHCEV------------KDYMSAYVWYREMQEKGFLLD-VCIGNELVSGLKEEWRSKEA-    
 280 UniRef50_O49619        Pentatricopeptide repeat-co...   410    1e-112  1   9.6%   4:574   169:707     ---NSLISLYMKLgdAEKVFEEMPERDIVSWNSMISGYLALGDGFSSLMLFKEM--LKCGFKPDRFSTMSALGACSHVYSPKMGKEIHCHAVRSRIEtdVMVMTSILDMYSKYGEVSYAERIFNGmqRNIVAWNVMIGCYARNGrdAFLCFQKmeQNGLQPDVITSINLLPASA---ILEGRTIHGYAMRRGFLPHMVLETALIDMYGECGQLKSAEVIFDRMAEK----NVISWNSIIAAYVQNGKNYSALELFQELWDSSLVPDSTTIASILPAYAESLSLSEGREI------------HAYIVKSRYWSNTIILNSLVHMYAMCGDLEDARKCFNHILLK------DVVSWNSIIMAYAVHGFGRISVWLFSEMIASRVNPNKSTFASLLAACSISGMVDEGWEYFESMkeYGIDpeHYGCMLDLIGRTGNFSAAKRFLEEMP----FVPTARIWGSLLNASRNHKDITIAEFAAEQIFKMEHD-NTGCYVLLLNMYAEAGRWEDVNRIKLLMESKGISRTSSR----------STVEAKGKSHVFTNGDRSHVATNKIYEVLD-VVSRMV-----------------------------    
 281 UniRef50_Q9LIC3        Putative pentatricopeptide ...   410    1e-112  1  11.3%  35:581    27:540     ----------------------------------ISQLCSNGRLQEALL---EMAMLG--PEMGFHGYDALLNACLDKRALRDGQRVHAHMIKTRYLPATYLRTRLLIFYGKCDCLEDARKVLDEmeKNVVSWTAMISRYSQTGheALTVFAEMMRSDGKPNEFTFATVLTSCIRagLGLGKQIHGLIVKWNYDSHIFVGSSLLDMYAKAGQIKEAREIFECLPER----DVVSCTAIIAGYAQLGLDEEALEMFHRLHSEGMSPNYVTYASLLTALSGLALLDHGKQ------------AHCHVLRRELPFYAVLQNSLIDMYSKCGNLSYARRLFDNMPER------TAISWNAMLVGYSKHGLGREVLELFRLMreKRVKPDAVTLLAVLSGCSHGRMEDTGLNIFDGMveYGTKpeHYGCIVDMLGRAGRIDEAFEFIKRMPSK----PTAGVLGSLLGACRVHLSVDIGESVGRRLIE--IEpnAGNYVILSNLYASAGRWADVNNVRAMMMQKAVTKEPGR----------SWIQHEQTLHYFHANDRTHPRREEVLAKMKEISIKMKQAGYVPD----------------------    
 282 UniRef50_Q6K7A0        Os02g0470000 protein n=4 Ta...   409    1e-112  1  11.6%   7:531   134:639     ------YCRDGRLADAERVLGAARaaNVVTYTALIDGYCRSGRLDDALRLIASM-----PVAPDTYTYNTVLKGLCIAKKWEEAEELMAEMIRNRCPPNEVTFATQIRSFCQNGLLDRAVQLLDQMPRYGCTPDVVIYSTLINgdQALDLLNTML---CKPNTVCYNAALKGLCIarWEDIGELMAEMVRKGCSPNEATFSMLISSLCQNNLVDSAVEVLEQMEKYGCEPDTVNYNIIINSLSERGRVDDALRLLNSMV---CKPDALGFNAVLKGFCRAERWHDASELIAQMFRDDCPLI------------EMTFNILIDMLCQNGLVNYATQVFEQMPR--YRCTPDIVTYSSLLNGFSEQGLVEVAIQLFRSMP---CKPDIFSYNAVLKGLCRAARWEDAGELIAEMVGKDCPPntFNILINSLCQKGLVDRAIEVLEQMPNYG-STPDIFTYNALINGFSEQGRLDDALKLLSTM---SCKPDAISYNSTLKGLCRAERWQDAEELVAEMLRNKCTPNEVTFKYA------------------------------------------------------------------------    
 283 UniRef50_B9IHD3        Predicted protein n=3 Tax=r...   409    1e-112  1  10.0%   3:602     9:637     --YNSMISVYAKNGraRNLFDKMPRRNLVSWNTMVSGYLHNGKFDEAYKLFVIM------PRRDLFSWTLMITCYTRNGEVEKARELFDSLPCSYRKGVACWNAMISGYVKKGRVNEAKRLFDEMPVKNLISWNSMLAGYTQnmRLGLEFFNEMDER----DVVSWNLMVDGFIqgDLDSAWKFFQETQK----PNVVSWVTMLSGFARNGNILESRRLFDQMPSR----NIVSWNAMISAYVQRCEIDEASRLFEEMPER----DSVSWTTMINGYVRIGKLDEARELLNEMPYRNIGAQTAMIsdEIGTWDVVCWNAMIAGYAHHGRINEALCLSKRMVNK------DMVTWNTMISCYAQVGQMDRAVKIFEEMGER----DLVSWNSLIAGFMLNGQNLDALKSFALMGHEGKKPdvNNALITMYAKCGRILEAGLVFN-----GICHADVISWNSLIGGYAINGYGKEALKLFEEMASEGMAPDEVTFIGILSACNHAGMVDHGLKLFKCMSKVYAIEPLAEHYACMVDLLGRVGRLDEAFEIVRGmlLGACRAHGNLELGRLAAHKLSE--FEP-HKTSNYVLLSNIHAEANRWNE-    
 284 UniRef50_Q8LIR0        Os07g0635800 protein n=3 Ta...   409    1e-112  1  10.7%   7:602    98:683     ------YAKLGDVAHALRVFDGMPRRNSFAWNAVIKGLVDAGRFSEALEMFWGMVNDGS-VAVDGFTYPPVIKACAALGAVAQGRKVWEMVesGNARPNVFVQCALVDMFAKCGCLDEARNVFESMqrDLAAWTAMIGGTvgNWLEVVDLFNHMRSEGFGVDSLIAATVISACGRAglQVGTALHGCAVKSGASGDIYVSNALVDMYCKCGCVEMADCLFWSTNSK----DVVSWSSLIVGYSQNGMHNVSVSLFCEMISLGINPNSSTLASILPCLSVLKLIRSGKEI------------HCFSIRHGLERSEFVVSALIDLYSKQGLIRVAETIF------WLTLDKDLAIWNSMVAGYAVNGYSDSAFCALRLLQKVGLKPDHVTVVSVLPLCNQHHMLIQGKELHAYVIKYCINssVNNALLDMYCKCGFLEVAKEVFQLMTER-----NTVTYNILISSFGKHNHEDQALSFFDLMKRDGIAPDKVTFVALLSCCSHAGLIDKGLHLYHSMLHDYNISPEKEHYSCIVDLYSRCGKLDEAWCFMSNmlLAACRVHNRMDIAELVGKRIFEQN--PND-PGYHILLSNIYADAGMWSD-    
 285 UniRef50_B9T4Q6        Pentatricopeptide repeat-co...   409    1e-112  1   9.0%  16:596    36:605     ---------------FSDVIKLGVEVNTNTFNILICGCCIENKLSEAIGLIGKMKHFSCF--PDNVSYNTILDVLCKKGKLNEARDLLLDMKNNGLFPNRNTFNILVSGYCKLGWLKEAAEVIDIMARNNVLPDVRTYTTlgKIDEAFRLRDEMENLKLLPHVVTYNMLINGCFECSSskGFELIDEMEGKGVKPNAVTYNVVVKWYVKEGKMDNAGNELRKMEESGFSPDCVTFNTLSNGYCKAGRLSEAFRMMDEMSRKGLKMNSVTLNTILHTLCGERKLDDAYKLLSSASKR------------GYFVDEVSYGTLIMGYFKVGKSXX----WDEMKEK--EIIPSIITYGTMIGGLCRSGKTDQSIDKCNELLESGLVPDQTTYNTIILGYCREGQVDKAFHFRNKMVKKSFKPdtCNILVRGLCTEGMLDKALKLFKTWISKGKAI-DAVTFNTIISGLCEEGRFEEAFDLLAEMEEKKLGPDCYTHNAILSALADAGRMKEAEEFMSGIVEQGKLQDQTISLNKRKTESSSETSQESdnSVAFSEQINELCTQGKYKDAMHMIQESTQKGIIL-HKSTYISLMEGLIKR-------    
 286 UniRef50_B9GNH7        Predicted protein n=2 Tax=M...   409    1e-112  1  10.6%  72:600    36:548     -----------------------------------------------------------------------ISLLQNCKHNNQIPPIYAKIIRNHHHQDPFVVFELLRVCSNLNSIGYASKIFSHTqpNVYLYTALIDGLVLsyTDGIHLYYQMINSSLVPDSYAVTSVLKAccHLALKEGREVHSQVLKLGLSSNRSIRIKLIELYGKCGAFEDARRVFDEMPERDVVadTVCWTAMIDGLVRNGESNRALEVFRNMQREDVMPNEVTIVCVLSACSELGALQLGRWV------------RSYMDKHRIELNHFVGGALINMYSRCGDIDEAQRVFEQMKEK------NVITYNSMIMGFALHGKSVEAVELFRGLIKQGFTPSSVTFVGVLNACSHGGLAELGFEIFHSmkDYGIEpeHYGCMVDLLGRLGRLEEAYSFIRMMK----VAPDHVMLGALLSACKIHGNLELAERVAKSLVACK-NADSGTYILLSNAYSSSGKWKEAAEVRTNMREEGIEKEP----------GCSSIEVNNEIHEFLLGDLRHPQKEKIYKKLEELNQILRLEGYTPAT----EVVLHDIEKSEKEW---    
 287 UniRef50_Q7XSP5        Os04g0686500 protein n=4 Ta...   409    1e-112  1   9.2%   6:582    48:595     -----MYAKCGKLHMAGEVFDGMPERNVVSWTALMVGFLHHG---EARECLRLFGEMRGSgsPNEFTLSATLKACG--GGTRAGVQIHGVCVRTGFEGHDVVANSLVVMYSKGRWTGDARRVFDVIprNLATWNSMISGyaGQGRDSLLVFREMQRRHDepDEFTFASLLKACSGLgaREGAQVHAAMAVRGVSpnAILAGALLDVYVKCHRLPVAMQVFDGLERR----NAIQWTTVIVGHAQEGQVKEAMCLFRRFWSSGVRADGHVLSSVVAVFADFALVEQGKQV------------HCYTAKTPAGLDVSVANSLVDMYLKCGLTGEAGRRFREMPAR------NVVSWTAMINGVGKHGHGREAIDLFEEMQEEGVEADEVAYLALLSACSHSGLVDECRRYFSRirRMRPKAEHYACMVDLLGRAGELREAKELILSMP----MEPTVGVWQTLLSACRVHKDVAVGREVGDVLLAVDGD-NPVNYVMLSNILAEAGEWRECQGIRGAMRRKGLRK----------QGGCSWTEVDKEVHFFygGDDAHPQAGDIRRALREVEARMRErgYSGDA---------------------    
 288 UniRef50_D1IP62        Whole genome shotgun sequen...   409    1e-112  1   9.9%  10:595    27:628     ---------DEALSTFNRMLHKQPPPSTVDFNRLLTSIAKTKHHSTLLSLSRQMDSFG--IPPDVYTLAIVINSLCHLNRVDFAFSALAKILKLGHQPDATTFTTLIRGLCVEGKIGEALHLFDKMIGEGFQPNVVTYGTLINGleAFNIFSEMITKGISPNIVTYNSLIHGLCkcEWKHVTTLMNEMVDSKIMPDVFTLNTVVDALCKEGMVAEAHDVVDMMIHRGVEPNVVTYNALMDGHCLRNEVDVAVKVFDTMVHKDCVANVISYNTLINGYCKIQSVDKAMYLFEEMSRQE------------LTPNTVTYNTLIHGLCHVGRLQDAISLFHEMVARGQ--IPDLVTYRTLSDYLCKNRHLDKAMALLKAIEGSNWDPDIQIYTTILDGMCRAGELEDARDLFSNLSSKGLQPntYNIMIHGLCKQGLLAEASKLFSEM-NKNGCSPNDCTYNLITRGFLRNNEALRTIELLEEMLARGFSVDVSTTTLLVGMLSDDGLDQSVKQIL--CKPSSSSRPSGTQMRSLPPLFLSSShlDDALSSFNRMLHMHpaKMKHYSTVLSLSTQMDSFGIPPD-VYTLNILINSFCH--------    
 289 UniRef50_Q84NV9        Os07g0513200 protein n=5 Ta...   409    1e-112  1  12.8%   4:518   230:733     ---NAVLCRLP-LDEAVQLFQELPEKNTCSYNILLKALCTAGRIKDAHQLFDEMAS-----PPDVVTYGIMVHGYCTLSELETAIKLLSEMAARGLELNPVAYTSVIALLCDEGQVSDAVRVVEDMVMHGvvFTTVMSGFCRKGdaARNWFDEMQKRGLAADGVTYTALINGLCRAglKEAERVLQEMEDKGLDVDAVTYTVLIDGYCKVGKMTEAFLVHNKMVQKRVTPNVVTYTALSDGLCKQGDVCAANELLHEMCSKGLELNIFTYNSLINGLCKAGNLEQAMRTMIDMDE------------AGLKPDVYTYTTIIGALCQSKELDRAHSLLQEMLDKG--IKPTIVTYNVLMNGFCMSGRVEGGKRLLEWMLEKNIHPNTTTYNSLMKQYCIEKNMKSTTEIYKGMLSQEVVpnTYNILIKGHCKARNMKEALYFHSEMIEKG-FRLTASSYNALIRLLNKKKKFTEARRLFEKMRKERLTAEPDVYNFYIDLSFNEDNLESTLALCDELVE-------------------------------------------------------------------------------------    
 290 UniRef50_B9H4S5        Predicted protein n=3 Tax=r...   408    1e-112  1  11.6%   6:523   175:726     -----MYAKCGVMEDASLMFDEIVVRDVVSWTALVIGYVHNDDSEKGLECLCEMRRIGGDGevNSRTLEGGFQACGNLGAMIAGRCLHGLAVKTGlcSQVVQSSLLSMYSKCGNVEEAHNSFCQVVDKDVFSWTSVIGVCARFgnECLNLFWDMQVDDVYPDGIVVSCILLGFGNSMmrEGKAFHGLIVRRNYVLDDTVNNALLSMYCKFGTLNPAEKLFddMYGKgtQRDVVTWNTLISSYTHSGHYAEAITLFDEMISEKLNPNSATLVIVLSACCHLPSLEKGKMVHQYIKE------------GGFELNVSLGTALVDMYAKCGQLEQSRELFNSMKEK------DVISWNVMISGYGLHGDANSAMEVFQQMEQSNVKPNAITFLSLLSACTHAGYVDEGKQLFDRMQYYSIKPnhFACMADLLGRSGNLQEAEDLVQSMP----ICPDGGVWGTLLSACKIHNEIEIGIRVAKCAIESDPE-NDGYYIMLSNMYGSMGKWDEAERARELMKERGIGK--------------------------------------------------------------------------------    
 291 UniRef50_Q2QVE3        Os12g0233200 protein n=5 Ta...   408    1e-112  1   9.2%  24:576    97:657     -----------------------PGPNAYMLGTMMRGFLRARLPARALGLFRRVVRDRLPADARTFVFAVKAAAAahGGTPSGGEAIHCAALKCGFveSVLVGNALVHFYANHKSLDDAGKVFDEmeRDVVSWTTLVDGYARAgdEAWRLFCRMVVvgMRPNAVTLVAAVSAIGQMGLLagIMLHKYVTEGGVARSVNLDNALVDMFGKCGCVRYAREVFDGMEVK----DVYSWTSMVNAYAKCGDLESAEQLFKDMPRR----NVVSWSCMIAAYSQLNQPEEAVWLFREMIAAGVDPIDayIVSNKIGLTVNLGNALIDMFAKCGDVGEASKLFDEMAER------NVVSWNTMIMAHAVHGQSEEAIRLFEQLKGENIVPDQITFLGLLASCSHSGLVSEGRRYFKEMEmyRIEpeHYACMIDLLGKVGLLEEAFEVARGMP----MEADEAGWGALLNACRMHGNVEIGACVADKLVELDPS-DSGIYVLMSQIYASKNKWDQVKMLRMTMRDRGVKKNP----------GCSSIEVEGKFHDFLVADVSHACSEEIYSALKNIYFHLKQE---------------------------    
 292 UniRef50_Q94I34        Os10g0400250 protein n=4 Ta...   408    1e-112  1   9.5%  16:575    69:638     ---------------LRHALALLSSLPASAYNAAIRALSLSDDGDrvRRCLPLYRALLRSgrPDHLTFPFLLKACARLREWGYGDAALAHVLRLGLDSDVFVVNAATHFLSIRGPMEDARRLFdsPVRDLVSWNTLIGGYVRRGneALELFWRMVAEdvRPDEVTMIAAVSGCGQMrlELGRRLHGFVDSDGVSCTVRLMNALMDMYIKCGSLEMAKSVFERIEHR----TVVSWTTMIVGFAKFGLMDDARKVFDEMPERDVFP----WNALMTGYVQCKQCKEALSLFHEMQeeMGMWVHRYIEKHRLVFSVALGTSLIDMYAKCGNIEKAIHIFKEIPEK------NALTWTAMICGLANHGHANEAIEHFRTMIELGQKPDEITFIGVLSACCHAGLVKEGREFFSLMekYHlkMKHYSCMIDLLGRAGHLDEAEQLVNTMP----MEPDAVVWGAIFFACRMQGNISLGEKAA--MKLVEIDpdSGIYVLLANMYAEANMRKKADKVRAMMRHLGVEKVP----------GCSCIELNGVVHEFIVKDKSHMDSHAIYDCLHEITLQIKH----------------------------    
 293 UniRef50_Q8GSH3        Os07g0670000 protein n=2 Ta...   408    1e-112  1  12.9%   7:574   176:713     ------YSRGGSLGEARSVFDDMPARDLVSWNALICGLAQDgcPAEVIGVFLRMLRDG-DVQPDRISVCSVIPACGGEGKLELGRQVHGFAVKLGVEGYVSIGNVLVAMYYKCGAPGSARRLFDAmeRDVVSWTTAIS---MDGEDALTLFNGMRRDGVPPNEVTFVALMSAlaDCPARGAQMVHAACLKAAVSGEAAASNSLITMYAKARRMDDARMVFDLMPRREI----IAWNAVISGYAQNGRCNDALELFSSM-ARCLTPNETTFASVLSAVTAVETVSMA----------YGQMYHSRALSMGFGDSEYVAGALIDMYAKRGNLEESRKAFHETEQR------SLIAWTAIISANARHGSYGAVMSLFGDMARSGVAPDGVVLLAVLTACRYGGMVDAGRDIFDSMadRGVepEHYSCVVDMLGRAGRLAEAEELMMRMPA----GPSVSALQSLLGACRIHGDAEIGERIARILTEK--EPtsGAYVLLSNIYADVGDWDGVAKVRRKMRDRGVK-------KEIGFSWVDAGAGEAllHKFSSDDTTHPRTEEIYAVADVLGWEMK-----------------------------    
 294 UniRef50_B9I8Q2        Predicted protein n=2 Tax=M...   407    1e-112  1  11.8%   9:552   202:760     --------RSGRVKDALKVIDEmcRPNDATGDILFSFLLKRERNEELleIVNLVLKFGEHGVlsSFWMGRLITRLCRNRKTNRGWDLFTEMIKLGAVLESAACNSLLTGLAREGNFNRMNELMEKMVEMDIQPNVVTFGILInvDDALEVLEKmkESgvEPDVVIYNTLIDGLCkgRQQEGLGLMERMrqKGCAPDTITYNCLIDGFCKAGEIEKGKELFDEMNKEGVAPNVVTVNTLVGGMCRTGRVSSAVNFFVEAQRRGMKGDAVTYTALINAFCNVNNFEKAMELFNEMLKS------------GCSPDAIVYYTLISGFSQAGRMADASFVLAELKKLG--IRPDTVCYNTLIGGFCRTNKFHRVFEMLKEMEEAGLKPDTITYNTLIAYASKNGDLKFAQKVMRKMIKAGVVPttYGAVINAYCLNGNGNEAMEIFKDMKAASKVPPNTVIYNILINSLCKNNKVKSAVSLMEDMKIWGVTPNTTTYNAIFKGLRDEKDLEKVFEFMDRMIEHACNPDYIT-MEILTEWLSAVGEI-ERLKKFVAG---------------------------------------------------    
 295 UniRef50_O23337        Pentatricopeptide repeat-co...   407    1e-112  1  10.4%  16:581    63:628     ---------------ALNVFSSIPSPpsIVFNPFLRDLSRSSEPRATILFYQRIRH--VGGRLDQFSFLPILKAVSKVSALFEGMELHGVAFKIATLCDPFVETGFMDMYASCGRINYARNVFDEmhRDVVTWNTMIERYCRFgdEAFKLFEEMKDSNVMPDEMILCNIVSACGrgNMRYNRAIYEFLIENDVRMDTHLLTALVTMYAGAGCMDMAREFFRKMSVRNlkKDLVCWTTMISAYVESDYPQEALRVFEEMCCSGIKPDVVSMFSVISACANLGILDKAKWV------------HSCIHVNGLESELSINNALINMYAKCGGLDATRDVFEKMPRR------NVVSWSSMINALSMHGEASDALSLFARMKQENVEPNEVTFVGVLYGCSHSGLVEEGKKIFASMteYNITpeHYGCMVDLFGRANLLREALEVIESMP----VASNVVIWGSLMSACRIHGELELGKFAAKRILE--LEPDhgALVLMSNIYAREQRWEDVRNIRRVMEEKNV----------FKEKGLSRIDQNGKSHEFLIGDKRHKQSNEIYAKLDEVVSKLKLAGYVPD----------------------    
 296 UniRef50_D1IXU9        Whole genome shotgun sequen...   407    1e-112  1   9.1%   6:590    79:661     -----LLNMYVklCDASKLFDEMPERNTISFVTLIQGYAESVRFLEAIELFVRLHREGHELNP--FVFTTILKLLVSTDCGELGWGIHACIFKLGHESNAFVGTALIDAYSVCGRVDVAREVFDGIlkDMVSWTGMVTCFAEnfKEALKLFSQMRMVGFKPNNFTFASVFKACLGLefDVGKSVHGCALKSRYELDLYVGVALLDLYTKSGDIDDARRAFEEIPKK----DVIPWSFMIARYAQSDQSKEAVEMFFQMRQALVLPNQFTFASVLQACATMEGLNLGNQI------------HCHVIKIGLHSDVFVSNALMDVYAKCGRMENSMELFAE------SPHRNDVTWNTVIVGHVQLGDGEKALRLFLNMLEYRVQATEVTYSSalSACANAGLLDQGQAYFTSMidHGIEpeHYTCMVWLLGRGGHLDKAVKLIDEIP----FQPSVMVWRALLGACVIHNDIELGRISAQRVLE--MEpdKATHVLLSNMYATAKRWDNVASVRKNMKRKGVKKEP----------GLSWIESQGTVHSFTVGDTSHPEVRVINGMLEWLHMKTKKAGYIPN----YNVVL-------------    
 297 UniRef50_D1IQE2        Whole genome shotgun sequen...   407    1e-112  1  11.9%  17:603   183:760     ----------------RRMVNEGFVLGGIDYRTVVNALCKNGFVQAAEMFCCKVLRLGFG--LDTHVCTSLVLANCRRDDLGEAFRVFEKMSKenCRPNSVTYSILIHGLCEAGRLEEAFQLKQEMVEKGCQPSTRTYTVLIKadKAMKMLDEMATKACVPNVHTYTILIDRLCrgKIEEANGVFRKMLKHGLCPGIITFNALINGYCKEGWVVSAFQLLSVMEKGNCKPNIRTYNELMEGLCRVSKSYKAFLLLRRVVDNGLLPDRVTYNILVDGFCKEGQLNMAFNIFNSMNS------------AGLEPDGFTFTALIDGLCKLGRLEQANGILGSMVKKGISL--DEVTFTALIDGHCKIGKAKDVCFLFENMVENRCLTTAHTFNCFLDALGKDYKLNEANAMLGKMMKYGLVPstHTILIEGHCRAGETALSLKMLERMKQAGC-SPNVYTYTIIINGLCNNGRVEEAETILFSMSSFGVSPNHFTYAVLVKAHVKAGRLDRAFQIVSTMVKNGCQPNSHIYSALLSGFVLSNTAIGaaLSSTGDLDARSLSSEENDNNC--LSNEIKKCGV-PTE-DLYNFLVVGLCKEGRIIEAD    
 298 UniRef50_Q9SS81        Pentatricopeptide repeat-co...   407    1e-112  1  10.7%   5:594    57:650     ----RMVNHVSRIVELIRSQECKCDEDVAL--SVIKTYGKNSMPDQALDVFKRMREI-FGCEPAIRSYNTLLNAFVEAKQWVKVESLFAYFETAGVAPNLQTYNVLIKMSCKKKEFEKARGFLDWmkPDVFSYSTVilAKAGKLDDALELFDEMSERGVAPDVTCYNILIDGFLkkDHKTAMELWDRLLesSVYPNVKTHNIMISGLSKCGRVDDCLKIWERMKQNEREKDLYTYSSLIHGLCDAGNVDKAESVFNELDERKASIDVVTYNTMLGGFCRCGKIKESLELWRIMEHKNSV-------------NIVSYNILIKGLLENGKIDEATMIWRLMPAKGY--AADKTTYGIFIHGLCVNGYVNKALGVMQEVESSGGHLDVYAYASIIDCLCKKKRLEEASNLVKEMSKHGVELnvCNALIGGLIRDSRLGEASFFLREMGKNGC-RPTVVSYNILICGLCKAGKFGEASAFVKEMLENGWKPDLKTYSILLCGLCRDRKIDLALELWHQFLQSGLETDVMMH-NILIHGLCSVGKLDDAMTVMANMEHRNcfKVGDSNRATVIWGYMYKMGLQPDII-SYNTIMKGLC---------    
 299 UniRef50_Q9STS9        Putative pentatricopeptide ...   406    1e-111  1   9.4%   5:568   148:682     ----SLLDMYKRVGKIDkvFSEMPFRNAVTWTAIITGLVHAGRYKEGLTYFSEMS--RSEELSDTYTFAIALKACAGLRQVKYGKAIHTHVIVRGFVTTLCVANSLATMYTECGEMQDGLCLFENmeRDVVSWTSLIVAYKRIGQeaVETFIKMRNSQVPPNEQTFASMFSACAssRLVWGEQLHCNVLSLGLNDSLSVSNSMMKMYSTCGNLVSASVLFQGMRCR----DIISWSTIIGGYCQAGFGEEGFKYFSWMRQSGTKPTDFALASLLSVSGNMAVIEGGRQV------------HALALCFGLEQNSTVRSSLINMYSKCGSIKEASMIF------GETDRDDIVSLTAMINGYAEHGKSKEAIDLFEKSLKVGFRPDSVTFISVLTACTHSGQLDLGFHYFNMMQeyNMRpeHYGCMVDLLCRAGRLSDAEKMINEMSWKK----DDVVWTTLLIACKAKGDIERGRRAAERILE--LDPTCAtlVTLANIYSSTGNLEEAANVRKNMKAKGVIKEP----------GWSSIKIKDCVSAFVSGDRFHPQSEDIYNILEL-----------------------------------    
 300 UniRef50_A9TJC6        Predicted protein n=1 Tax=P...   406    1e-111  1  11.9%  14:601   108:700     -------------GLFYEMKVWRCKPDVETYNALMNVHGRAGQWRWALQIFDDMLQAPKPIAPSRSSYNNLINACGSSGQWFKALEICKRMTDNGVGPDLMTYNIILSAFKSGGQPRHAVAYYDHLIsdRYSHNIILNCllGRFEDAINLFKEMRKmgCEPDVVTFNALLHVYacGQITKAQETFDMMIGEGMAPTIVTFNTLLGAYASQGMYTEALQVVGLLVKAKFEPDVVSYSSLLNAFGKAGYPEKAQEVFDLMKQRSRKPNLVTFNGLMDAYASAGKYERARELLHDMAE------------ARIEPNVVTICSLFAACARARCPEKVRDVFHEAKVR--QIALNVPAFNAAITAYIEAGQFDEAKALLEVMEEQNLRPNGVTFLLLINAAGSLGDYREARTLYDKMVDLGIPLTvcSALINAFAKHGMYNEARTIFDDMRKAGCKA-NVVTYTSLMIAYRMYDDAQLAC---EEMVAAGITPDSIAFGTLFVALNKANRSEKVLQIEKSSRARGILFDDVAYTETLlaLFLYEGMKKRRLRLTNTMVisAVGRSGKLDRLHKVFLELQQqgFVP-TLQMYNILIDSYSR-GLKWR--    
 301 UniRef50_B9F9M1        Putative uncharacterized pr...   406    1e-111  1  11.5%  15:597   217:839     --------------FLKESLDRKFPLDVTTCNIVLNSLCTQGKLSKAESMLQKMKNCRL---PNAVTYNTILNWYVKKGRCKSALRILDDMEKNGIEADLYTYNIMIDKLCKLKRSARAYLLLKRMrpDECSYNTLIHGFFGEGklAIYIFNQMLRQSLKPSVATYTALIDGYCrgRTDEARRVLYEMQITGVRPSELTYSALLNGYCKHSKLGPALDLIKYLRSRSISINRTMYTILIDGFCQLGEVSKAKQILKCMLADGIDPDVITYSALINGMCKMGMIHETKEILSRMQKSGvlKYFVDIYRSGLVANSVIHNALLCSFYREGMIAEAEQFKQYMSR--MKISFDVASFNCIIDSYCQRGNVLEAFSVYDNMVRHGWPPDICTYGSLLRGLCQGGHLVQAKEFMVYLLEKACAIdtLNTLLVGICKHGTLDEALDLCEKMVTRNIL-PDTYTYTILLDGFCKRGKVVPALILLQMMLEKGLVPDTIAYTCLLNGLVNEGQVKAASYMFQEIICKEGLYADCIAYNSMMNGYLKGGQINEIERLMRNMHEngYIKKGQLSRTLYLYRDMVKEGIKPDNV-TYRLLIFGLCEYG------    
 302 UniRef50_D1J3C3        Whole genome shotgun sequen...   406    1e-111  1  10.8%   6:562    47:569     -----MYAKCGHTTFARQVFDEMFEKNLVSWSAMISGYDQAGEPQMAIDLYSQM-----FLVPNEYVFASVISACASLSAVTLGQKIHSRSLKFGYESISFVSNSLISMYMKCNQCSDALSVfnTPEPNCVSYNALITGFVEnlERGLEFFKLMRQQGLIPDRFAFMGVLGIctTENLKRGAELHCQTVKLNLDSTPFIGNVIITMYSELNLIQEAEKAFRLIEEK----DVISWNTLIAACSHCDDHAKGLRVFKHmeETNVRPDDFTFTSALAACAGLASMSHGKQIHAHLMRTRLYQ------------DLGVGNALVNMYAKCGCIGYAYDIFSKM------VHHNLVSWNTIIAGFGNHGLGERAVELFEQMNASGIRPDSVTFIGLLTACNHAGLVDKGQLYFNSMEeyGIAPdhFSCLIDMLGRAGRLNEAEEYMRKFPFWN----DPVVLVSLLSASRLHGDVVIGERLAKWLLK--LQptTSPYVLLSNLYASDGMWDSVAEARKRLKGSGLKKEPGH----------SLIEVNGSVEKFTIGDFTHLRIKEI-----------------------------------------    
 303 UniRef50_D1IND5        Whole genome shotgun sequen...   405    1e-111  1  11.4%   3:574    90:726     --SNCMISAygNLVQARLLFDEMPERNEVSWSALISGLMKYGRVEESMWYFErfRLLESGVRPNDVTFTSVVRACGELGDFGLGMSILGLVVKAGFEHYLSVSNSLITLSLRMGEIDLARRVFDRMerDVVSWTAILDAYvgYAEEALKLFSKMVQEGFKPNISCFACTLSALASLrlSAGINIHGHVTKIGIDKDVFIGSSLIDLYCKCGKPDDGRLVFDLILEKNveKNDVSWGTIIAGYLENEQCEKVLEVFNTLLVSGQTPNKSTFSSVLCACASIASLDKGMNV------------HGKIIKLGIQYDIFVGTALTDMYAKCGDIGSSKQVFERMPEK------NEISWTVMIQGLAESGFAVESLILFEEMErsEVAPNELMLLSVLFACSHCGLVDKGLWYFNSMEkyGIKPkhYTCVVDLLSRSGRLYEAEEFIRTIP----FQPEANAWAALLSGCKKYKDEKIAERTAKKllAEN---NSAGYVLLSNIYASAGRWIDVSNIRKLMREKGLKKS----------GGCSWVEVRNQVHSFYSEDGSHSQSNEIYGTLQLLRSEML-----------------------------    
 304 UniRef50_Q9LYU9        Pentatricopeptide repeat-co...   405    1e-111  1   8.1%   6:574   127:664     -----MYCECRSLEDADKLFDEMSELNAVSRTTMISAYAEQGILDKAVGLFSGMLASGD--KPPSSMYTTLLKSLVNPRALDFGRQIHAHVIRAGLCSNTSIETGIVNMYVKCGWLVGAKRVFDQmvKKPVACTGLMVGyaGRARDALKLFVDLVTEGVEWDSFVFSVVLKACASleLNLGKQIHACVAKLGLESEVSVGTPLVDFYIKCSSFESACRAFQEIRE----PNDVSWSAIISGYCQMSQFEEAVKTFKSLRSKNAslNSFTYTSIFQACSVLADCNIGGQVHADAIKRSLIGSQYGE------------SALITMYSKCGCLDDANEVFESMDN------PDIVAWTAFISGHAYYGNASEALRLFEKMVSCGMKPNSVTFIAVLTACSHAGLVEQGKHCLDTmrKYNVApdHYDCMIDIYARSGLLDEALKFMKNMP----FEPDAMSWKCFLSGCWTHKNLELGEIAGEELRQLDPE-DTAGYVLPFNLYTWAGKWEEAAEMMKLMNERMLKK----------ELSCSWIQEKGKIHRFIVGDKHHPQTQEIYEKLKEFDGFME-----------------------------    
 305 UniRef50_Q7XJN6        Pentatricopeptide repeat-co...   405    1e-111  1  10.6%   4:576   314:856     ---LSMYSKCGMVGEAETVFSCVVDKRLEIWNAMVAAYAENDYGYSALDLFGFMRQ--KSVLPDSFTLSNVISCCSVLGLYNYGKSVHAELFKRPIQSTSTIESALLTLYSKCGCDPDAYLVFKSMekDMVAWGSLISGLCKNGkeALKVFGDMKDDdlKPDSDIMTSVTNACAGLelRFGLQVHGSMIKTGLVLNVFVGSSLIDLYSKCGLPEMALKVFTSMSTE----NMVAWNSMISCYSRNNLPELSIDLFNLMLSQGIFPDSVSITSVLVAISSTASLLKGKSL------------HGYTLRLGIPSDTHLKNALIDMYVKCGFSKYAENIFKKMQHK------SLITWNLMIYGYGSHGDCITALSLFDEMKKAGESPDDVTFLSLISACNHSGFVEEGKNIFEFMkdYGIEPnhYANMVDLLGRAGLLEEAYSFIKAMP----IEADSSIWLCLLSASRTHHNVELGILSAEKLLR--MEPERgtYVQLINLYMEAGLKNEAAKLLGLMKEKGLHKQP----------GCSWIEVSDRTNVFFSGGSSSPMKAEIFNVLNRLKSNMVDE---------------------------    
 306 UniRef50_B9SJK1        Pentatricopeptide repeat-co...   405    1e-111  1  11.5%  91:601     1:482     ------------------------------------------------------------------------------------------MKSGYQPNLVTVSTLVKGMCFEGKVMD-ALELFDKMTRSGFQGDILLYGYLINarRAVELHRRMLSEICSGNFFTYGLVIDSLCKDGLakGLKMFREMINMGICPNVIVYSSLINGLCRVGKLREAVNLFDEMVSQGIKANVITYNSLIHASCRFGLWKEATRTFSQMVGEGILPDVVTFTTLIDHLSKKGKVQEAYKIFELM------------IKQGEAPNIVTYNSLLNGLCLHHQMDHAVRLFEVMVERGIKI--DVISYNTLINGYCTSGKTEEAMTLFRKMQYEELTPSITTYTILLKALYQNGRIRTAKELFNNMQICGQSpdTYTVLLDGLCKNGCIEEAIDVFRSLKSIKYK-PSIRIYSILIGGMFQARRWESAMEIFDEIPTVGLVPNIVTYNIMINGLCKEGKLLEAERLFVQMEESGC----------------------EQDEISFNFIIRGFLQENQVQKAMEFLKRMREKNFSPNDSVTL-MLVNLYAADAQSSE--    
 307 UniRef50_B9S753        Pentatricopeptide repeat-co...   405    1e-111  1  11.4%   7:574    54:606     ------YSKWGEMGNAQKLFDRMSERSVISYNILISGYGGMGFYHKAIGLFSEARM--ACLKLDKFSYAGVLSACGQIKDFALGKVIHGLAIVCGLGQQVFLTNLLIDMYCKCERIDHARLLFESSDedNVSWNSLiyARVGAYEEMLKLLVKMHHTGLRLNAFTLGSALKSclNNMVSYGKTLHGYTVKQGLDLDIVVGTALLDMYAKTGYLGDAIQLFRTSPNQ----NVVMYNAMIAGFIQTEDIdeALKLFSQMQRQGIKPSDFTFSSIIKICNHIEAFEYGKQI------------HAHICKHNIQSDEFIGSTLIELYSLLGSTEDQLKCFN------STPKLDIVSWTTMIAGYAQNGQFESALALFYELLASGKKPDEFIITTMLSACADVAAERSGEQVHGYAVKTGilAIVQNSQISMYAKSGNLDSAKITFEEIKN-----PDVVSWSVMICSNAQHGHAKDAINLFELMKSYGIHPNQITFLGVLTACSHGGLVEEGLRYYESMKKDYDMKINVKHCTCIVDLLSRAGrlDAKNFILNSGFGDHPVMWRTLLSGCRIYKDIV-----------------------------    
 308 UniRef50_Q84Q17        Putative pentatricopeptide ...   405    1e-111  1   9.0%  31:577   141:690     ------------------------------YNVLISSCLRHGLPLQALAAYQEMGKNGVL--PDVFTYPSVLRACAEARELVLGRAVHMHAAGAGMDGNLFFQNALMSMYAKCGDLASARKVFDGmqRDVVSWNSMiyAAVGQWAEAMELFRRMRDEGTEVNSVTWNTilNACSrgWLRLGKEIHGLAVRMCCDQVESVSNALITMYARCKDMECARMLFRMLE----CPGVVTWNTMLSSFALSDCAEEASSIFREMICRGVKPNYVTVVTYLALCARVANLQHGQEL------------HGHIVKHGFKGYRLLWNSLIDMYSKSGRLSVAQNVFDTMDD------CDMISYTSMIAGYGMQGKGTVALRLFEQMIDSGIKPDHIIMVTVLSACSHSGLVLEGEELFNKmiSYGIKpeHYSCMIDLYARAGLLEKA----EEMLDHTPFPPTSTMWAALVGACHDRGNIEIGERAARKLLEMRTE-NAGHYVLIANMYAAAGCWDELATVRKLMRDLGVTKAPGLAWTDLGNGFT----------PFLVGDRSNPLAPEIYVVLDELSEQMRNIN--------------------------    
 309 UniRef50_D1H473        Whole genome shotgun sequen...   405    1e-111  1   9.4%   7:601    79:578     ------LSPFPSLDYAQQVFDQIPHPNLYTWNTLIRAYASSSNPHQSLLIFLRMLH-QSPDFPDKFTFPFLIKAASELEELFTGKAFHGMVIKVLLGSDVFILNSLIHFYAKCGELGLGYRVFVNIPRR--------------------------------DVVSWNSMITAFVQGGceEALELFQEMETQN---SLTLSNAMLDMYTKCGSVEDAKRLFDKMPEK----DIVSWTTMLVGYAKIGEYDAAQGIFDAMPNQ----DIAAWNALISAYEQCGKPKEALELFHELQLSKTAKPDEVTLKQGMKLNCHLTTSLIDMYCKCGDLQKALMVFHSVERK------DVFVWSAMIAGLAMHGHGKDAIALFSKMQEDKVKPNAVTFTNILCACSHVGLVEEGRTFFNQM--------------------------ELVYGKMP----MAPAASVWGALLGACTIHENVVLAEQACSQLIE--LEpnHGAYVLLSNIYAKAGKWDRVSGLRKLMRDVGLKKEP----------GCSSIEVDGIVHEFLVGDNSHPSAKKIYAKLDEIVARLETIGYVPNKSH-----LLQLVEEEDVKE--    
 310 UniRef50_D1IBF5        Whole genome shotgun sequen...   405    1e-111  1   9.3%  31:580    71:608     ------------------------------YNIEICRFCELGNLRRA------MELINQSPKPDlrTYCSVLQLCADLKSIQDGRRIHSIIQSNDVEVDGVLGSKLVFMYVTCGDLREGRRIFDKvnEKVFLWNLLmyAKIGNFRESLSLFKRMRELGIRrdVISWNSMISGYVSNGLSegLDLFEQMLLLGINTDLATmnCLLDMYSKSGNLNSAIQVFETMGER----SVVSWTSMIAGYAREGLSDMSVRLFHEMEKEDLFPNSITMACILPACASLAALERGQEI------------HGHILRNGFSLDRHVANALVDMYLKCGALGLARLLFDMIPEK------DLVSWTVMIAGYGMHGYGSEAIAAFNEMRNSGIEPDEVSFISILYACSHSGLLDEGWGFFNMMRNNCckSEHYACIVDLLARAGNLSKAYKFIKMMP----IEPDATIWGALLCGCRIYHDVKLAEKVAEHVFE--LEpnTGYYVLLANIYAEAEKWEEVKKLRERIGRRGLRKNP----------GCSWIEIKGKVHIFVTGDSSHPLANKIELLLKKTRTRMKEEGHFP-----------------------    
 311 UniRef50_Q65XK6        Os05g0572900 protein n=4 Ta...   405    1e-111  1   9.9%   5:581    58:599     ----ALYVKCGRLGLARQVFDAMPSRNPVSGNLLMSGYASSGRHRDALALLRV-----ADFGLNEYVLSSAVAATAHVRSYDMGRQCHGYAIKAGLAEHPYVCSAVLHMYCQCAHMDEAVKVFDNVSsnVFAFNSMINGFLDRgdGSTSIVRSMVRNVGQWDHVSYVAVLGHCASTkvVLGSQVHTQALKRRLELNVYVGSALVDMYGKCDFPHEANRVFEVLPEK----NIVSWTAIMTAYTQNELFEDALQLFLDMEMEGVRPNEFTYAVALNSCAGLATLKNGNAL---------GACTMKTGHWGLLP---VCNALMNMYSKSGSVEDARRVFLSMPCR------DVVSWNSIIIGYAHHGRAREAMEAFHDMLFAEEVPSYVTFIGVLSACAQLGLVDEGFYYLnmMKEVGVKpeHYTCMVGLLCRVGRLDEAERFIES----NCIGTDVVAWRSLLSSCQVYRNYGLGHRVAEQIFQLKPK-DVGTYVLLSNMYAKANRWDGVVKVRRLMRELGVRKEP----------GVSWIQVGSEVHVFTSEDKKHPYMEQITKKLQELIDKIKVIGYVPN----------------------    
 312 UniRef50_D1I1Z5        Whole genome shotgun sequen...   405    1e-111  1  11.9%  18:599   225:803     -----------------EIKASGVPQNEYTNPILIDGLCRQSRLQDAVTFLR--ETGGEEFGPSVVSFNALMSGFCKMGSVDVAKSFFCMMIKYGLLPDVYSYNILLHGLCVAGSMEEALEFTNDMENHGVEPDIVTYNILANgsGAWKVVQRMLLNGLNPDLVTYTILICGHCqgNIEESFKLKEKMLSQGLKLSIVTYTVLLSSLCKSGRIDEAVILLHEMEVIGLKPDLLTYS--------RGAVEEAIELYEEMCSKRIYPNSFVCSAIISGLFEKGAISEAQMYFDSVTKSDVAEE------------IILYNIMIDGYAKLGNIGEAVRSYKQIIEKG--ISPTIVTFNSLIYGFCKKGKLAEAVKLLDTIKVHGLVPTSVTYTTLMNGYCEEGDMHSMFDMLHEMEAKAIKPttYTVVVKGLCKEGRLHESVQLLKYMYARGLF-PDQITYNTVIQSFCKAHDLQKAFQLHNQMLQHSLQPSPVTYNVLINGLCVYGNLKDADRLLVTLQDQSIRLTKVA-YTTIIKAHCAKGDVQNALVFfySAVINRLCKRNLITDAKFFFCMMLTHGIPPDQDICL-VMLNAFHRSGDP----    
 313 UniRef50_B9HD88        Predicted protein n=2 Tax=r...   404    1e-111  1  10.8%  72:580    14:530     -----------------------------------------------------------------------IRLFQGFNSLKHLKHVHAALLRLGLDEDSYLLNKVLRFSFNFGNTNYSHRIFHQtePNIFLFNTMIHGLVLnfQESIEIYHSMRKEGLSPDSFTFPFLLKACArlDSKLGIKLHGLVVKAGCESDAFVNTSLVSLYGKCGFIDNAFKVFDDIPEKNveKDIVSWSSMIQGYASNGLPKEALDLFFKMLNEGFRPDCYAMVGVLCACARLGALELG------------NWASNLMDRNEFLGNPVLGTALIDMYAKCGRMDSAWEVFRGMRKK------DIVVWNAAISGLAMSGHVKAAFGLFGQMEKSGIEPDGNTFVGLLCACTHAGLVDEGRQYFNSMERVfpEIEHYGCMVDLLGRAGFLDEAHQLVKSMP----MEANAIVWGALLGGCRLHRDTQLVEGVLKQLIA--LEpnSGNYVLLSNIYSASHKWEDAAKIRSIMSERGIKKVPGY----------SWIEVDGVVHEFLVGDTSHPLSEKIYAKLGELVKDLKASGYVP-----------------------    
 314 UniRef50_Q9SA46        F3O9.28 n=1 Tax=Arabidopsis...   404    1e-111  1  10.4%   4:582   393:940     ---LRMYAGAGRSVEANLVFKQMPTKDLISWNSLMASFVNDGRSLDALGLLCSM--ISSGKSVNYVTFTSALAACFTPDFFEKGRILHGLVVVSGLfnQIIGNALVSMYGKIGEMSESRRVLLQMPRRDVVAWNALIGGYAEdpDKALAAFQTMRVEGVSSNYITVVSVLSACLLPglERGKPLHAYIVSAGFESDEHVKNSLITMYAKCGDLSSSQDLFNGLDNR----NIITWNAMLAANAHHGHGEEVLKLVSKMRSFGVSLDQFSFSEGLSAAAKLAVLEEGQQL------------HGLAVKLGFEHDSFIFNAAADMYSKCGEIGEVVKML------PPSVNRSLPSWNILISALGRHGYFEEVCATFHEMLEMGIKPGHVTFVSLLTACSHGGLVDKGLAYYDMirDFGLEpeHCICVIDLLGRSGRLAEAETFISKMPMK----PNDLVWRSLLASCKIHGNLDRGRKAAENL--SKLEpdDSVYVLSSNMFATTGRWEDVENVRKQMGFKNIKKKQA----------CSWVKLKDKVSSFGIGDRTHPQTMEIYAKLEDIKKLIKESGYVADT---------------------    
 315 UniRef50_B9SC35        Pentatricopeptide repeat-co...   404    1e-111  1  10.2%   5:525    35:602     ----KLIQMYADCDhaQRLFDKMPQPNVYAWTAIFGFYLRHGMYDKCVQNYGFMKY--SDVLPDNYVFPKVLRACTQLLWFEGGIWIHKDVIVCGCESNLQVCNSLIDMYVKCGNARSARLVFEEMerDLFSWNSMISGYVSNgdLAVELLNCMRLDGFEPDVVTWNTLMDAYCqgRFDEAWEVLKQIEQ----PNIISWTTLISSYSKIGEHDMSLRVFQDMIIREveTNTVFYsaLLTMYAKCGRIQDAINVFELMDKS----DIVTWNAMILGFVELDLGKQAIECFREMQRMDIkpIHAYVRKSITLSSVVtwNAVIHMYCKCGCVRSAYTIFCSMPNK------DVVSWNTMIGGFGMHGHGQAALKLLKEMILSGIFPNSTTFTSVLSACSHSGLVDEGFRLFRSmeDYSITpeHYSCIVDMLARAGQFADAVTFIHKMP----LEPDKSIWGALLAACRAYQNLDFGRLAAEQLIRMEPKC-AGHYVTLSNIYARAGRWDDAGRVRKEIEGKGLVKPS------------------------------------------------------------------------------    
 316 UniRef50_Q7XHP5        Os07g0239600 protein n=3 Ta...   404    1e-111  1   9.3%   9:578    57:629     --------RRRRFHDARGVFDERPTRTAPVWTLTISGCARRGRYADGMRAFAEMLAEGEA-TPNAFVLAAVVRCCAGMGDVESGKRVHGWMLRNGVHLDVVLCNAVLDMYAKCGQFERARRVFGAmeRDAVSWNIAIGACIQSGddALSHLRRMAQAGVVFNHYTYSTAFVLAGMLLLPdgRQLHGRVLIAALEGDAFVRSSLMDMYCKCGLLEAAASVFDHWSPLTRDMN-FAWSTMVAGYVQNGREEEALDLFRRMLREGVAADRFTLTSVAAACANVGMVEQGRQV------------HGCVEKLWYKLDAPLASAIVDMYAKCGNLEDARSIFDRACTK------NIAVWTSMLCSYASHGQGRIAIELFERMTAEKMTPNEITLVGVLSACSHVGLVSEGELYFKQmeEYGIVpeHYNCIVDLYGRSGLLDKAKNFIEE----NNINHEAIVWKTLLSACRLHQHNEYAKLASEKLVQLE-QCDAGSYVMLSNIYATNNKWHDTFELRVSMQERKVRKQPGR----------SWIHLKNTVHTFVAGDASHPQSAEIYAYLEKLVERLKEIGY-------------------------    
 317 UniRef50_C5WXL7        Putative uncharacterized pr...   404    1e-111  1  12.0%  14:579    18:558     -------------GHLRHLDPQLQVPPLLLANSLIAAFSRAALPRLALPLLRRLLAGAHPLRPDSFTFPPLVRAAPG---PASAAQFHACALRLGlpNVFASGSLVHAYLRFGRVAEAYRVFDEMPERDVPAWNAMLSGLCRNArdAVALFGRMVGEGVAGDAVTLSSVLPMCVlgDRALALVMHVYAVKRGLSGELFVCNALIDVYGKLGMLVEARWVFGGMALR----DLVTWNSIISAYEQGGKVAAAVELFHGMMKSGVSPDVLTLVSLASAVAQCGDERGAKSVHCYVRRRGWDVG-----------DIIAGNAVVDMYAKLSKIDAAQMVFDNLPDR------DVVSWNTLITGYMQNGLSNEAVRIYNDMQneGLKPIQGTFVSILPAYSNLGALQQGMRMHALSIKTGLNLDVytCLIDLYAKCGKLVEAMLLFEHMPRRSTGP-----WNAIIAGLGVHGHGAKALNLFSQMQQEGIKPDNVTFVSLLAACSHAGLVDQGRSFFDLMQTVYGIVPIAKHYACMVDMLGRAGQLDEAFEF----IQGMPIKPDSAVWGALLGACRIHGNVE------------------------    
 318 UniRef50_Q9MA95        Putative pentatricopeptide ...   404    1e-111  1  11.8%   5:521    43:552     ----RLIDFCtnLSYARSVFESIDCPSVYIWNSMIRGYSNSPNPDKALIFYQEM--LRKGYSPDYFTFPYVLKACSGLRDIQFGSCVHGFVVKTGFEVNMYVSTCLLHMYMCCGEVNYGLRVFEDIPqnVVAWGSLISGFVNnfSDAIEAFREMQSNGVKANETIMVDLLVACGRcdIVTGKWFHGFLQGLGFDpnVILATSLIDMYAKCGDLRTARYLFDGMPER----TLVSWNSIITGYSQNGDAEEALCMFLDMLDLGIAPDKVTfaSMIQGCSQLGQ-----------------SIHAYVSKTGFVKDAAIVCALVNMYAKTGDAESAKKAFEDLEKK------DTIAWTVVIIGLASHGHGNEALSIFQRMQEKgaTPDGITYLGVLYACSHIGLVEEGQRYFAEMRdhGLEpeHYGCMVDILSRAGRFEEAERLVKTMPVK----PNVNIWGALLNGCDIHENLELTDRI-RSMVAEPEELGSGIYVLLSNIYAKAGRWADVKLIRESMKSKRV----------------------------------------------------------------------------------    
 319 UniRef50_D1IRT6        Whole genome shotgun sequen...   404    1e-111  1   9.4%   7:593    79:707     ------FGLVSDAQRVFEVSPIECFSNLLLWNSILRANVAHGYCEEALEIYCRMRKLGVS--ADGFTFPLVIRACALMGSRKLCRSVHGHVVEMGFQWNLHVGNELMGMYGKIGRMDDARKVFERmvRSCVSWNTMVSGYALNYdgASEMFRMMGSAGLEPNLVTWTSLLSSHARCgfDEGKVIHGYVVKGGFENYLFVKNSLICLYGKHGNVNAARILFLEIKTK----NIVSWNALISSYADLGWCDEAFAIFLQLEKTdvRPNVVSWSAVIGGFASKGQGEEALELFRRMqlHLGREIHGHVVRSLMDGNILVGNGLINMYTKSGSFKEGNLVFEKIENK------DLISWNTMVAGYGIHGLGENAIRTFDQMIKDGFEPDGVTFVAVLSACSHAGLVAEGRELFDKMIKerVEpeHYACMVDLLGRAGLLQEASKVVKSMP----VEPNACVWGALLNSCRMHKNTEVAEETASQIFNLNSEI-AGSYMLLSNIYAASGRWEDSAKVRISAKTKGLKKTPGQ----------SWIQVKKKVYMFSAGNTQHAELEEVYRILKDLGLQMEVEGYIPDIDEEQRSILLGY----------    
 320 UniRef50_B9RCX4        Pentatricopeptide repeat-co...   404    1e-111  1   9.4%   6:567   244:771     -----MYSRCDDFSSAERIFSNMFNRSIASWTAMISCYNRSRWFKQALQVFVEMLEFK--VAPNAVTIMAVLSSCAGFNLLREGKSVHCYAVKHIDLDDDSLGPALIEYYAQFGKLSYCEKVLHTikRNIISWNMLISVYASQgkEALGIFVQMQRQGQIPDSFSLSSSISACANVGlwLGHQIHGYAIKRHI-LDEFVQNSLIDMYSKCGHVDLAYLIFDRIQSK----SVVAWNSMICGFSQIGNSLEAIRLFDQMYLNCLDMNEVTFLTAIQACSHMGHLEKGKWLHHKLIA------------YGVKKDLFIDTALIDMYAKCGDLRIAHRVFDSMSER------SVVSWSAMIGGCGMHGDIDAAISLFAEMIQREMKPNDITFMNILSACSHSGYVEEGKFYFNSMKNFEVEPnhFACMVDLLSRAGDLDEAYRIINSMP----FPAEASIWGALLNGCRIHQRMDMIRNIERDLLDMRTD-DTGYYTLLSNIYAEEGNWDVSRKVRSAMKGIGLKKVPGY----------STIELDKKVYRFGAGDVSHWQVKEINTFLE------------------------------------    
 321 UniRef50_Q7XK39        OSJNBa0044K18.6 protein n=1...   404    1e-111  1  11.6%  67:593   390:881     ------------------------------------------------------------------TSAAALALFKSVLSADKALSPLAVLPHLDGAPSSLPNLLLTASAAV-RPHATSLRLYSRMKSLSLPIStlLSALPSAP--AFALFADMFRLRLPLCTTTFNIMlhLCSAGKPARALELLRQMPR----PNAVTYNTVIAGFCSRGRVQAALDIMREMRERgiAPNQYTYGTVISGWCKVGRVDEAVKVFDEMLTKgvKPEAVMYNALIGGYCDQGKLDTALLYRDRMVER------------GVAMTVATYNLLVHALFMDGRGTEAYELVEEMGGKG--LAPDVFTYNILINGHCKEGNVKKALEIFENMSRRGVRATVVTYTALIYALSKKGQVQETDKLFDEAVRRGIRPdlYNALINSHSTSGNIDRAFEIMGEM-EKKRIAPDDVTYNTLMRGLCLLGRVDEARKLIDEMTERGIQPDLVTYNTLISGYSMKGDVKDALRIRNEMMNKGFNPT----------LLTYNALIQGLCKNGQGDD------------AENMVKEMVENGITPDD-STYISLIEGL----------    
 322 UniRef50_Q2QYR0        Pentatricopeptide, putative...   404    1e-111  1  11.3%  51:580   103:603     --------------------------------------------------LHALRLHCLLPNPSHLTFPIALKSASRLPhlRAGEQLHARSLKLPSHTNPHVLTSLLTLYARCGLLHRAQRVFDEmhPSTVSWTALITAyaGDLREAVHVARNAFANGMRPDSFTAVRVLTACAraDLATGETVWRAAEQEGIAQSVFVATAAVDLYVKCGEMAKAREVFDKMRDK----DAVAWGAMVGGYASNGHPREALDLFLAMQAEGVRPDCYAVAGALSACTRLGALDLGRQAIR------------MVDWDEFLDNPVLGTALIDMYAKCGSTAEAWVVFQQMRKK------DIIVWNAMILGLGMTGHEKTAFTLIGQMEKSGVKLNDNTFIGLLCSCTHTGLIQDGRRYFHNMTkyHISpeHYGCIVDLLSRAGLLQEAHQLIDDMP----MPANAVILGALLGGCKIHRNAELAEHVLTQLIR--LEpnSGNYVMLSNIYSNRGRWEDAAKLRLDMKEKGVEKVPA----------CSWVEFEGKVHEFRVGDKSHPLSDQIYKKLDELGLEMKTMGYEP-----------------------    
 323 UniRef50_Q67WJ3        Pentatricopeptide (PPR) rep...   404    1e-111  1  11.0%   1:600    43:622     LAALLRAGRYGA---ARRLFDALPARSVVTWNSLLAGLARRPDARAAREFFDAM------PVRDAVSWNTLLAAYSASPHPDhaRRLFDEMPQR---DVVTWNTLLGAYARRGLMDEARRLFDEMPQRNAASWNTMVTGfaGQVVKALDVFDAMPAK----DSASLSTMVSGFTKNglHEAEELlkRLSVTDMDKAVDAYNTLIVAYGQAGRFSDAKRLFDMIPkkGFERNVVSWNSMMICYIKAGDVCSARALFNEMPDK----DLVSWNTMISGYTQASDMKESEKLFWEMPD----------------PDTVSWNLIIQGFMQKGEAEHARGFFDRMPERG------TISWNTMISGYEKNGNYISSVKLFSKMLEVGEIPDRHTFSSVLAACASIPMLGLGAQIH-QLVEKSFVPDTanALITMYSRCGALNDAEAIFKQMHTKK----DLVSWNALIGCYEHHGRATKALQLFKEMRRAKVMPTHITFVSLLSACVNAGLVSEGRMVFDTMVHEYGIVARIEHYAALVNLIGRHGQLDDALEV----INSMPMAPDRSVWGAFLGACTAKKniNPDSSAPY-VLIHNLHAHEGKW---    
 324 UniRef50_Q10IY6        Os03g0441400 protein n=6 Ta...   404    1e-111  1   9.1%   7:600   207:764     ------YSLCSLVSDAEHVFNGIVRKDAVVWTAMVSCYSENDCPENAFRVFSKMRVSGCKPNP--FALTSVLKAAVCLPSVVLGKGIHGCAIKTLNDTEPHVGGALLDMYAKCGDIKDARLAFEMIpdDVILLSFMISRYAQSneQAFELFLRLMRSSVLPNEYSLSSVLQACTNmqLDFGKQIHNHAIKIGHESDLFVGNALMDFYAKCNDMDSSLKIFSSLRDA----NEVSWNTIVVGFSQSGLGEEALSVFCEMQAAQMPCTQVTYSSVLRACASTASIRHAGQIHCSIEKSTFNN------------DTVIGNSLIDTYAKCGYIRDALKVFQHLMER------DIISWNAIISGYALHGQAADALELFDRMNKSNVESNDITFVALLSVCSSTGLVNHGLSLFDSMrdHGIKpeHYTCIVRLLGRAGRLNDALQFIGDIPS----APSAMVWRALLSSCIIHKNVALGRFSAEKILE--IEpdETTYVLLSNMYAAAGSLDQVALLRKSMRNIGVRKVP----------GLSWVEIKGEIHAFSVGSVDHPDMRVINAMLEWLNLKTSREGYIPDI----NVVLHDVDKEQKTR---    
 325 UniRef50_Q9LIE7        Pentatricopeptide repeat-co...   404    1e-111  1  10.5%   3:574   257:801     --AISMYAELGDIESSRRVFDSCVERNIEVWNTMIGVYVQNDCLVESIELF--LEAIGSkiVSDEVTYLLAASAVSALQQVELGRQFHGFVSKNFrlPIVIVNSLMVMYSRCGSVHKSFGVFLSMRERDVVSWNTMISAFVQNgdEGLMLVYEMQKQGFKIDYITVTALLSAASNLRNKegKQTHAFLIRQGIQFemNSY--LIDMYSKSGLIRISQKLFE---GSGyeRDQATWNSMISGYTQNGHTEKTFLVFRKMLEQNIRPNAVTVASILPACSQIGSVDLGKQL------------HGFSIRQYLDQNVFVASALVDMYSKAGAIKYAEDMFSQTKER------NSVTYTTMILGYGQHGMGERAISLFLSMQESGIKPDAITFVAVLSACSYSGLIDEGLKIFEEMReyNIQpeHYCCITDMLGRVGRVNEAYEFVKGLGEEGNI---AELWGSLLGSCKLHGELELAETVSERLAKFDKGKNFSGyvLLSNMYAEEQKWKSVDKVRRGMREKGLKK----------EVGRSGIEIAGYVNCFVSRDQEHPHSSEIYDVIDGLAKDMR-----------------------------    
 326 UniRef50_Q9LR69        Pentatricopeptide repeat-co...   404    1e-111  1   9.2%   4:524   100:600     ---NSLLSLYFKLgeTRRVFDGRFVKDAISWTSMMSGYVTGKEHVKALEVFVEMVSFGL--DANEFTLSSAVKACSELGEVRLGRCFHGVVITHGFEWNHFISSTLAYLYGVNREPVDARRVFDEmePDVICWTAVLSAFSKnyEEALGLFYAMHrkGLVPDGSTFGTVLTACGNlrLKQGKEIHGKLITNGIGSNVVVESSLLDMYGKCGSVREARQVFNGMSKK----NSVSWSALLGGYCQNGEHEKAIEIFREMEEK----DLYCFGTVLKACAGLAAVRLGKEI------------HGQYVRRGCFGNVIVESALIDLYGKSGCIDSASRVYSKMSIR------NMITWNAMLSALAQNGRGEEAVSFFNDMVKKGIKPDYISFIAILTACGHTGMVDEGRNYFVLmkSYGIKpeHYSCMIDLLGRAGLFEEAENLLERAECRN----DASLWGVLLGPCAANADASraERIAKRMME--LEPkhMSYVLLSNMYKAIGRHGDALNIRKLMVRRGVAKT-------------------------------------------------------------------------------    
 327 UniRef50_B9GQK7        Predicted protein n=1 Tax=P...   404    1e-111  1   9.7%   7:584    50:595     ------YARIGGITSARHLFDKLPQRGVDAWNAMIVAYSRRYHLTEVLNLYHQMV--NEGGKPDSSTFTVAIKASSSLKDLEAGERIWRRAVDFGYGCDVFVGSSVLNLYVKCGKIDEAKLVFDKmkRDVVCWATMITGLVQNGneAVDMFRRMRKEGIEGDGVLMLGLVQACAngELKLGLSVHGHAVRRelMDDVILQTSLVDMYAKIGDLELASRVFEQMPRK----NAVSWGALISGFAQNGFAEYALDLLVEMQSLEFKPDTAVLVSALLACSQVGHLKLGKSIHGYIVRRLGFEL-------------VLGTALIDMYAKCGSLSCAHAIFDRVDSR------DVILWNTMIASYGIHGDGKEVLSLFLKMKEANISPDHATFASLLSALSHSGQVDVGQYWFNAMveCKIPPshYACMVDLFSRAGRVEEAYQLIESM----NTEPGLAIWVALLSGCHNYRNLLFGEVAAKKILDLNPD-DLGIYALVSNFFSLARMWDRVSILRKIMKETGMKKVPGY----------SAVEVNGKHEAFLVEDKNHHQYEEILQMLDSLDNEMRVIRHVPDSLH-------------------    
 328 UniRef50_B9I396        Predicted protein n=2 Tax=r...   404    1e-111  1   9.9%   3:596    77:665     --KNSLLGMYWKCgnAVDMFERMEERDSVSWNTMISGFCQSGDYVKSLVMFRRMVKECGGSYHNRVACLAALSSCASIKCLTHGLEIHGFLVKKGVDSDEFLVSALIEMYMKCGDIKNAENVFERIrrNMAVWNVMILGYVSNEClaLELFVEMLELGISPDSSTVVVVLVLCSQldLAVGKQIHGLILGLGLDDDVRVGTALMEMYFKCGDPETSLQIF----KRSQNHNLVMWGSVMLNCAQNGYPNEALEFFSemLDCGF-PDPVILLAALRACSFLS------------LKPRGMAIHGFAIKMGFDSDVFVGGALVDFYGKCGDMEYAQQVFYGLSTR------DLVSWNALISGFAQNKCADEALKAFRDMQSKQIKPNTVTMACILSVCTHLSVMILCKEVHCYLLRHWFETNAlnSLISAYAKCGDIHSSRTVFEKLPVR-----NEVTWNSILLGFGMHGRTDEMFATFEKMKEANIKPDHGTFTSLLSSCSHSGKVDAGWKYFNSMMEDYNLEPRVEQYTCMVDLLGRAGNLNQA----YDLIMSMPCSPDDRIWGSLLASC-KNHgvVANHIfgYRVLLANLYED-------    
 329 UniRef50_C5WQX9        Putative uncharacterized pr...   403    1e-111  1  10.5%  68:589    66:560     -------------------------------------------------------------------YATILRSCVLSRAVRPGRQLHARLLVSGLGLDAVLATRLVDLYASCGLVSVARRLFDEMPnnVFLWNVLIRAYARDgeAAIELYREMLACgmEPDNFTYPPVLKACAAldLGAGREVHDRVMRTSWAADVFVCAGLIDMYAKCGCVDEAWAVFDSTTVR----DAVVWNSMIAACGQNGRPAEALALCRNMAAEGIAPTIVTLVSAISAAADAGALPRGRELHGYGWRRGFGSQDKLK------------TSLLDMYAKSGWVTVARVLFDQL------FHRELISWNAMICGFGMHGHADHACELFRRMreAQVMPDHITFVGVLSACNHGGMVQEAKEVFDLMVtySIKpqHYTCLVDVLGHSGRFKEASDVIKGMLVK----PDSGIWGALLNGCKIHKNVELAELALHKLIE--LEpdAGNYVLLSNIYAQSGKWEEAARVRKLMTNRGLKKIIA----------CSWIELKGKSHGFLVGDASHPRSDEIYEELERLEGLISQTGYVPDTTSVFHNV--------------    
 330 UniRef50_UPI0001983DC2 PREDICTED: hypothetical pro...   402    1e-110  1  11.6%   6:599   122:699     -----LYGKFNKLGDAQSVFDEMLVRNTITWTTLIKGHLQ---VNDVESVFRIAREmwVGEEFNEHTCSVILQACDSLENLVRGEQIHGFVIKRGFDEDVFVGTSLISMYSRCGDLGAAEKVYSNLakDVRCLNFMiyGKAGCGEKAIGVFLHLLGSGLEPNDYTFTNVISAcgDIDVEVLRVLHGMCIKCGCGDEISVGNAIVSVYVKHGMLEEAEKSFCGMGER----NLVSWTALLSGYVKNGNGKKALEGFSQILELGVGFDSCCFATLLDGCSECKNLGLGLQI------------HGFVVKLGYVHDVSVGTALIDLYAKCRKLRSARLVFHSLLDK------NIVSFNAILSGYIGADE-EDAMALFSQLRLADIKPDSVTFARLLSLSADQACLVKGKCLHAYIIKTGFepSVGNAVITMYAKCGSIGDACQLFYSM-----NYLDSISWNAVISAYALHGQGRKALILFEEMKKEEFVPDEITILSVLQACSYSGLLEEGFCLFNDMESKYGIKPEIEHFACMVDLLGRAGYLSEAMSFINRsfSGSPLLWRTLVHVCKLHGDlhLLDLAPEEAGSY-ILVSNLYAGGGM----    
 331 UniRef50_A8MQA3        Pentatricopeptide repeat-co...   402    1e-110  1  11.8%  78:603    27:522     -----------------------------------------------------------------------------VSSITKLRQIHAFSIRHGVssDAELGKHLIFYLVspPPMSYAHKVFSKIEknVFIWNTLIRGYagNSISAFSLYREMRVSgvEPDTHTYPFLIKAVTtaDVRLGETIHSVVIRSGFGSLIYVQNSLLHLYANCGDVASAYKVFDKMPEK----DLVAWNSVINGFAENGKPEEALALYTEMNSKGIKPDGFTIVSLLSACAKIGALTLGKRV------------HVYMIKVGLTRNLHSSNVLLDLYARCGRVEEAKTLFDEMVDK------NSVSWTSLIVGLAVNGFGKEAIELFKYMEseGLLPCEITFVGILYACSHCGMVKEGFEYFRRmeEYKIEpeHFGCMVDLLARAGQVKKAYEYIKSMP----MQPNVVIWRTLLGACTVHGDSDLAE--FARIQILQLEPNHSgyVLLSNMYASEQRWSDVQKIRKQMLRDGVKKVPGH----------SLVEVGNRVHEFLMGDKSHPQSDAIYAKLKEMTGRLRSEGYVPQISNVY-------VDVEEEEKEN    
 332 UniRef50_Q9C507        Putative pentatricopeptide ...   402    1e-110  1  11.2%   5:602   141:724     ----SLLCMYgnLSDAEKVFDGMPVRDLVAWSTLVSSCLENGEVVKALRMFKCMVDDG--VEPDAVTMISVVEGCAELGCLRIARSVHGQirKMFDLDETLCNSLLTMYSKCGDLLSSERIFEKIAKKNAVSWTAMISSYNrfSEKALRSFSEMIKSGIEPNLVTLYSVLSSCGLIGlrEGKSVHGFAVRRELDPNYESLsaLVELYAECGKLSDCETVLRVVSDR----NIVAWNSLISLYAHRGMVIQALGLFRQMVTQRIKPDAFTLASSISACENAGLVPLGKQIHGHVIRTDV-------------SDEFVQNSLIDMYSKSGSVDSASTVFNQIKHR------SVVTWNSMLCGFSQNGNSVEAISLFDYMYHSYLEMNEVTFLAVIQACSSIGSLEKGKWVHHKLIISGLK-DLftALIDMYAKCGDLNAAETVFRAMSSRSIVSW-----SSMINAYGMHGRIGSAISTFNQMVESGTKPNEVVFMNVLSACGHSGSVEEGKYYFNLMKSFGVS-PNSEHFACFIDLLSRSGDLKEA----YRTIKEMPFLADASVwgCRIHQKmiKNDLS-DIvtGYYTLLSNIYAEEGEWEE-    
 333 UniRef50_D1HM82        Whole genome shotgun sequen...   402    1e-110  1  12.4%  72:573    21:502     -----------------------------------------------------------------------ISDLHRCSSLNQVKQIHAQVLKANLHRESFVGQKLIAAFSLCRQMTLAVNVFNQIqpDVLLYNTLIRAHVRNSelAFSVFFEMQDSGVCADNFTYPFLLKACskVWVRVVEMIHAQVEKMGFCLDIFVPNSLIDSYFKLGELGEARRLFDEMPER----DTVSWNTILDGYVKAGEMNAAFELFEKMpeAGLKFDDGTVISILSACAVSGLLGLGKRV------------HASIERTRFKCSTPVSNALIDMYAKCGSLENALSIFHGMVRK------DVVSWNAIIQGLAMHGHGEKALQLFSRMKGEGFVPDKVTFVGVLCACTHAGFVDEGLHYFHAMEryGVPPehYGCMVDLLGRGGRLKEAFRLVHSMP----LEPNAIIWGTLLGACRMHSATGLAEEVFDRLVKSELS-DSGNLSMLSNIYAAAGDWDNFANIRLRMKSTSIQKP----------SGGSSIEVDDEVHEFTVFDRSHPKSDRIYKTIDGLGQHI------------------------------    
 334 UniRef50_Q9LYV3        Putative pentatricopeptide ...   402    1e-110  1   9.7%   7:581   191:733     ------YSVCGSVDSARTVFEGILCKDIVVWAGIVSCYVENGYFEDSLKLLSCMRMAGFM--PNNYTFDTALKASIGLGAFDFAKGVHGQILKTCYVLDPRVGVGLLQLYTQLGDMSDAFKVFNEMPkdVVPWSFMIARFCQNgnEAVDLFIRMREAFVVPNEFTLSSILNGCAIGKCslGEQLHGLVVKVGFDLDIYVSNALIDVYAKCEKMDTAVKLFAELSSK----NEVSWNTVIVGYENLGEGGKAFSMFREALRNQVSVTEVTFSSALGACASLASMDLGVQVHGLAIKTNNAKK------------VAVSNSLIDMYAKCGDIKFAQSVFNEME------TIDVASWNALISGYSTHGLGRQALRILDIMKDRDCKPNGLTFLGVLSGCSNAGLIDQGQECFESMIrhGIEpeHYTCMVRLLGRSGQLDKAMKLIEGIP----YEPSVMIWRAMLSASMNQNNEEFARRSAEEILKINPK-DEATYVLVSNMYAGAKQWANVASIRKSMKEMGVKKEP----------GLSWIEHQGDVHYFSVGLSDHPDMKLINGMLEWLNMKATRAGYVPD----------------------    
 335 UniRef50_Q8RWS8        Pentatricopeptide repeat-co...   402    1e-110  1  10.5%   2:601   141:758     -ARNDIYNMMIRlgLFFEMQKWSCKPDAETYDALINA---HGRAGQWRWAMNLMDDmrAAIAPSRSTYNNLINACGSSGNWREALEVCKKMTDNGVGPDLVTHNIVLSAYKSGRQYSKALSYFELMkpDTTTFNIIIYCllGQSSQALDLFNSMREKrcRPDVVTFTSIMHLYskGEIENCRAVFEAMVAEGLKPNIVSYNALMGAYAVHGMSGTALSVLGDIKQNGIIPDVVSYTCLLNSYGRSRQPGKAKEVFLMMRKERRKPNVVTYNALIDAYGSNGFLAEAVEIFRQMEQD------------GIKPNVVSVCTLLAACSRSKKKVNVDTVLSAAQSRGINL--NTAAYNSAIGSYINAAELEKAIALYQSMRKKKVKADSVTFTILISGSCRMSKYPEAISYLKEMEDLSIPleVYSSVLCAYSKQGQVTEAESIFNQMKMAGC-EPDVIAYTSMLHAYNASEKWGKACELFLEMEANGIEPDSIACSALMRAFNKGGQPSNVFVLMDLMREKEIPFTGAVFFEIflQEWKRAIDLIQMMDPYlnQMLHLFGKSGKVEAMMKLFYKIIASGVGIN-LKTYAILLEHLLAVGNWRK--    
 336 UniRef50_Q6Z7E0        cDNA clone:J013162J12, full...   402    1e-110  1  13.1%  35:522    72:540     ----------------------------------LRSLVRRGELDEALRLV------GSARRPDAGTCAALIKKLSASGRTAEARRVLAA-----CGPDVMAYNAMVAGYCGAGQLDAARRLVAEmePDAYTYNTLIRGLCGRGRTalAVLDEMLRRRCVPDVVTYTILLEacKRSGYKQAMKLLDEMRDKGCTPDIVTYNVVVNGICQEGRVDDAIEFLKNLPSYGCEPNTVSYNIVLKGLCTAERWEDAEELMGEMGQKGCPPNVVTFNMLISFLCRKGLVEPALEVLEQIP------------KYGCTPNSLSYNPLLHAFCKQKKMDKAMAFLDLMVSRGC--YPDIVSYNTLLTALCRSGEVDVAVELLHQLKDKGCAPVLISYNTVIDGLTKAGKTKEALELLNEMVSKGLQPdtYSTIAAGLCREDRIEDAIRAFGKVQDMGI-RPNTVLYNAIILGLCKRRETHSAIDLFAYMIGNGCMPNESTYTILIEGLAYEGLIKEARDLLDELCSRGVV---------------------------------------------------------------------------------    
 337 UniRef50_Q9FXA9        Putative pentatricopeptide ...   402    1e-110  1   9.5%  13:580    61:598     ------------VEEARSLFDEMPDRDVVAWTAMITGYASSNYNARAWECFHEMVKQGTS--PNEFTLSSVLKSCRNMKVLAYGALVHGVVVKLGMEGSLYvnAMMNMYATCSVTMEAACLIFRDIkkNDVTWTTLITGflGDGIGGLKMYKQMLLENAEVTPYCITIAVRASASIdvTTGKQIHASVIKRGFQSNLPVMNSILDLYCRCGYLSEAKHYFHEMEDK----DLITWNTLISELERS-DSSEALLMFQRFESQGFVPNCYTFTSLVAACANIAALNCGQQL------------HGRIFRRGFNKNVELANALIDMYAKCGNIPDSQRVFGEIVDR-----RNLVSWTSMMIGYGSHGYGAEAVELFDKMVSSGIRPDRIVFMAVLSACRHAGLVEKGLKYFNVMeeYGINpdIYNCVVDLLGRAGKIGEAYELVERMP----FKPDESTWGAILGACKAHKHNGLISRLAAR-KVMELKPKMvtYVMLSYIYAAEGKWVDFARVRKMMRMMGNKK----------EAGMSWILVENQVFSFAVSDKMCPNASSVYSVLGLLIEETREAGYVP-----------------------    
 338 UniRef50_C5YWL6        Putative uncharacterized pr...   402    1e-110  1   9.9%   4:588   226:785     ---AGMYAESADVGAARRVLETEDAP-PVAWNVVVACCARLGLVDDALDLAERMARSGreLEPSLATWNAVLSGCARNGRDREAFGVVRIMLDQGLQPDSSSMSSLLKSVASLGLLAhaHGFfnQLEPDVYTGTAfmYAKCGHLEYAQKVFDTLELR----NITTWNSLVAGYANAgfDHALDLVEEMKRNRLDPDVTTWNGLINGYSVNGKSSQAVLLLRQIKAIGLTPNVVSWTSLISGSCNNGDYEDSFYFFNEMQKDHVQPSVVTMSVLLRACAGLALLKKGKELHCFALRRAYDR------------DMVVGTALIDMYSKSGNLASAQRIFERIQEN------NLVSSNAMLTGLAAHGQGHEAIALFHDMCNSGLKPDSITFTALLTACRSMDLVTEGWEYFDTMerYGVTPTvyACMVDLLARCGYLDEAMDFINKSPFKSV----SSLWGALLTGCTLHGNLALAEAAARKLFK--LEpnSANYLQMVSLYEHEQMFDEAESLKYAMKARALDTRP----------GWSWIQIEQSIHVFEVEGKPHPDTAEIYEELIRLVFQIRKSGYVPDTScvYNV---------------    
 339 UniRef50_Q6MWE3        B1358B12.23 protein n=5 Tax...   401    1e-110  1  11.6%   5:523   258:756     ----SMYTKCDSTEDARILFPELPEKDLVSWTSLIGAYCRAGHAEKAVELFLGMEE--SGLQPDEVVISCLLAGLGNDAKVRGGKTFHAAIVRRNFGDSVLIGNALISMYAKCKQVDIAATVFRMLhrDTDSWSSMVVAYCKAGlkCLELYREMqkDEFEYDTNSLISIISSCSrgRLRLGQSAHCYSIKHLAGENSSVANALISMYGRCGNFDVARKIFGMVKTK----DVVTWSALISSYSHLGHSKDALLLYDQMLTEGVKPNSATLVSVISSCANLAALEHGE------------LIHSHVKDVGLECDLSICTALVDMYMKCGQLGIARKMFDSMLER------DVVTWNVMISGYGMHGEAIQALKLFSMMERGNVKPNSLTFLAILSACCHAGLVDKGRELFTRMEEYSLEPnhYACMVDLLGKSGHLQEAEDVVSAMP----IEPDGGIWGTLLGACKMHDNFEMGLRVAKKAFASDPE-NDGYYILMSNSYGSAEKWNEIEKLRDMMKNHGVEK--------------------------------------------------------------------------------    
 340 UniRef50_Q9CAY1        Putative pentatricopeptide ...   401    1e-110  1  11.3%  31:569    21:529     ------------------------------WNVRLRELAYQSLFSESISLYRSM--LRSGSSPDAFSFPFILKSCASLSLPVSGQQLHCHVTKGGCETEPFVLTALISMYCKCGLVADARKVFEesSQLSVCYNALISGYTANSkdAAYMFRRMKETGVSVDSVTMLGLVPLCTVPelWLGRSLHGQCVKGGLDSEVAVLNSFITMYMKCGSVEAGRRLFDEMPVKGL----ITWNAVISGYSQNGLAYDVLELYEQMKSSGVCPDPFTLVSVLSSCAHLGA------------KKIGHEVGKLVESNGFVPNVFVSNASISMYARCGNLAKARAVFDIMPVK------SLVSWTAMIGCYGMHGMGEIGLMLFDDMIKRGIRPDGAVFVMVLSACSHSGLTDKGLELFRAMkeYKLepEHYSCLVDLLGRAGRLDEAMEFIESMP----VEPDGAVWGALLGACKIHKNVDMAELAFAKVIE--FEPNNIgyVLMSNIYSDSKNQEGIWRIRVMMRERAFRKKPGYSY----------VEHKGRVHLFLAGDRSHEQTEEVHRMLDEL----------------------------------    
 341 UniRef50_Q2QUQ5        Pentatricopeptide, putative...   401    1e-110  1   9.4%   7:603   164:744     ------YAACGKACDARRVFDEMPERDVVSWNSLVSAFLVNGMFHDARRALVSM--MRSGFPLNVASLVSVVPACGTEQEEKFGLSIHALAVKVGLNTMVNLANALVDMYGKFGDVEASMQVFDGmeQNEVSWNSAIGCFLNAggDVLRMFRKMSEHNVMPGSITLSSLLPALVegSFDLGREVHGYSIKRAMDLDIFVANSLVDMYAKFGSLEKASTIFEQMKDR----NVVSWNAMIANLVQNGAETEAFRLVTDMQKSGECPNSITLVNVLPACARMASLKMGKQI------------HAWSIRRGLMFDLFISNALIDMYSKCGQLSLARNIFERSEK-------DDVSYNTLILGYSQSPWCFESLLLFKQMRSVGIDYDAVSFMGALSACTNLSVFKHGKEIHCVLVRRLLspFLSNSLLDLYTKGGMLVTASKIFNKITKK-----DVASWNTMILGYGMHGQIDIAFELFELMKGDGLDYDHVSYIAVLAACSHGGLVDKGKKYFSQMVAQNIE-PQQMHYACMVDLLGRAGQLSKCAEI----IRDMPFPANSDVWGALLGACRIHGNIepEHSGYYTLMINMYAETGRWNEAN    
 342 UniRef50_D1IBH3        Whole genome shotgun sequen...   401    1e-110  1  12.7%  71:552    10:496     ----------------------------------------------------------------------ILSFAEMATSISELHQAHAHILKSGLIHSTFAASRLIASVSTNSHAQaaHSIFSRIppNSYMWNTIIRAYANSpeAALTIFHQMLHASVLPDKYTFTFALKSCGssGVEEGRQIHGHVLKTGLGDDLFIQNTLIHLYASCGCIEDARHLLDRMLERDVVsnVVSWNAMITGYSHAGRFSEVLVLFEDMQHAGVKPDNCTLVSVLSACAHVGALSQGEWV------------HAYIDKNGISIDGFVATALVDMYSKCGSIEKALEVFNSCLRK------DISTWNSIISGLSTHGSGQHALQIFSEMLVEGFKPNEVTFVCVLSACSRAGLLDEGREMFNLMVhhGIQpeHYGCMVDLLGRVGLLEEAEELVQKMPQKEA----SVVWESLLGACRNHGNVELAERVAQKLLE--LSpeSSSFVQLSNMYASMGRWKDVMEVRQKMRAQGVRKDP----------GCSMIEVDGTVYEFLAG---------------------------------------------------    
 343 UniRef50_D1HN23        Whole genome shotgun sequen...   401    1e-110  1  10.1%   3:569    38:699     --AISNYAKQSKLDVARQLFDQMPQRTVVSWNTMISSYSKHGRFSEA--LFLVYSMHRSHMKLSESTFSSVLSVCARLRCLRDGKLIHCLVLKSGSESFELVGSALLYFYASCFEIGEARRVFdmPRRDVVAWTTLISGFSKNGdkALEIFRLMMRSgtTPNEFTFDCVVRACGRLglSVGRTVHGLLMKCGLEYDPSIGGALVEFYCEcgRIEDAELVFNGMTEMNPvrTIFSSNTMISVYSRNGEIDKALELFEETKNEK---DPVTWNSMISGYIHSGQPEEALKLYITMHRLSIQqlHAHLIKTPFESNVYVGTSLIDMYSKCGSIMEAQTSFVSI------FSPNVAAWTALINGHAYHGLGSEAISLFDRMIEQGLAPNGATFVGVLSACSRAGLVNEGMKIFHSMErySVTpeHYACVVDLLGRSGHIREAEEFIKKMP----LEADGVVWGALLSACWFWMDLEVGERVAEKMFSFDPKP-ISSYVILSNIYAGLGRWRE-----KMMVRK-----ILRGFKVKKDPGCSWIELNNKIHVFSIEDRSHPYCNMIYATLEHL----------------------------------    
 344 UniRef50_UPI00019843BC PREDICTED: hypothetical pro...   400    1e-110  1   9.3%  71:599    38:575     ----------------------------------------------------------------------ILRHLSSCKTLKDLTQIHAQTITTGIFSDNFVASRILSFAAlgSIPYARFLFYRIRKPDIFIANTLIRAYAFSPndAVVFYSEMTESSVvpDVHTFPLLLKACSEisLRLGEAIHSHVFKLGWSSEVSvwNIMIGGYLKCGVFKSARRMFEAMPDR----DVVSWSVMINGYVQESRFKEGLGLFQDMMGEKIEPNESVLVNALSACAHLGAMEQGQWIERYMERKN------------VRLTVRLGTALIDMYSKCGSVERALEVFHKMKEK------NVLAWSAMINGLAINGQGKDALNLFSQMEMQGVKPNEVTFIGILNACSHSKLVDEGCSFFHSMTsyGLKPnhHCCMVDLYGRAGMLDQAQTVIKSMP----FKPNSAIWGALLNACRIHGDTELGEQVGKRLLE--LDPNHGgyVLLSNIYAACGRWDRVAELRRMMRERQVSKTP----------GCSFIDLGDTIHEFVAGDSSHPQLEMIYAKLGEMSQELKAAGYKPDTGQ---VLLdaLCHHSEK----    
 345 UniRef50_B9RM74        Pentatricopeptide repeat-co...   400    1e-110  1  13.1%   1:490   111:594     LAALVRMKHYDSVVSirKMEFFGVSCSVYTLTILINCLCHLHLVGLGFSVLGKIFKLGF--KPSIITFNTLINGLCIEGRIVEAMEQLDYIMSRGYQPTVYTHTMIVNGLCKIGKTSAAIVWMKKmePEVVSYSIIIDSLCKnvNEAVDLFYHMRSIGISPTVVTYNSLigMCNSGQWKQASILFKEMLEWNMKPDVVTFSILVDALCKEGVVLEALSVFGKMIQIAMEPDIVTYSSLIHGVCKSSLWKESSTLLNEMLSRNIRPDVVTFSIWVDVFCKKGMVSEAQSIINLMIER------------GLRPNVVTYNSLMDGYCLHSQMDEARKVFDIMVNKGC--APDVLSYNILIKGYCKSERIDEAKQLFDEMSHKGLTPNSITHTTLISGLCQAGRPYAAKELFKKMGSHGCPPdtYSTLLSGFCKHGHLDEALALFEALK-KSQLKPNHVICKILLGGMCKAGKLEDAKELFSSLSIEELQPDV-----------------------------------------------------------------------------------------------------------------    
 346 UniRef50_Q2QZN2        Pentatricopeptide, putative...   400    1e-110  1  10.0%   4:582   285:835     ---NALVNMYSAAGKLSdlFWNMSRRDLISWNTMISSYVQNCNSTDALKTLGQL--FHTNESPNHLTFSSALGACSSPGALIDGKMVHAIVLQLSLQRNLLVGNSLITMYGKCNSMEDAEKVFQSMphDIVSYNVLiyAVLEDGTKAMQVFSWMRSAGIKPN---YITMINIhsSNDLhyGRPLHAYIIRTGFLSDEYVANSLITMYAKCGNLESSTNIFNSITNK----NIVSWNAIIAANVQLGHGEEALKLFIDMQHAGNKLDRVCLAECLSSCASLASLEEGMQL------------HGLGMKSGLDSDSYVVNAAMDMYGKCGKMDEMLQV---VPDQAIRPQQ---CWNTLISGYAKYGYFKEAEETFKQMVATGRKPDYVTFVALLSACSHAGLVDKGIDYYNSMASsgVSPghCVCIVDLLGRLGRFAEAEKFIEEMP----VLPNDLIWRSLLSSSRTHKNLEIGRKTAKKLLELD-PFDDSAYVLLSNLYATNARWADVDKLRSHMKTININKRPA----------CSWLKLKNEVSTFGIGDRGHKHAEKIYAKLDEMLLKLREVGYIADT---------------------    
 347 UniRef50_Q0J7Z2        Os08g0153600 protein n=4 Ta...   400    1e-109  1  10.8%   7:581     3:525     ------LCRCPFLASAR---SHTCSLSTVA----VAAAVRRGDLTGAEEAFA------STPRKTTATYNCLLAGYArlGRLADARHLFDRIPT---PDAVSYNTLLSCHFASGDADGARRLFASMPVRDVVSWNTMVSGLSKSgeEAKAVFLAMPVR----NSVSWNAMVSGFACSrmSAAEEWFRNAPEKG---DAVLWTAMVSGYMDIGNVVKAIEYFEAMPVR----NLVSWNAVVAGYVKNSHADDALRLFRTmrEANVQPNASTLSSVLLGCSNLSALGFGKQI------------HQWCMKLPLSRNLTVGTSLVSMYCKCGDLSSACKLFGEMHTR------DVVAWNAMISGYAQHGDGKEAINLFERMKDEGVEPNWITFVAVLTACIHTGLCDFGIRCFEGMQeyGIEpdHYSCMVDLLCRAGKLERAVDLIRSMP----FEPHPSAYGTLLAACRVYKNLEFAELAAGKLIEKDPQ-SAGAYVQLANIYAGANQWDDVSRVRRWMKDNAVVKTPGY----------SWIEIKGVMHEFRSNDRLHPQLYLIHEKLGQLAERMKAMGYVPD----------------------    
 348 UniRef50_Q9C866        Pentatricopeptide repeat-co...   400    1e-109  1  10.6%  26:579     9:566     -------------------------PSLLMYNKMLKSLADGKSFTKVLALFGELR--GQGLYPDNFTLPVVLKSIGRLRKVIEGEKVHGYAVKAGLEFDSYVSNSLMGMYASLGKIEITHKVFDEmqRDVVSWNGLISSYVGNgeDAIGVFKRmqESNLKFDEGTIVSTLSACSALklEIGERIYRFVVTE-FEMSVRIGNALVDMFCKCGCLDKARAVFDSMRDKNVkkDVVLWTAMMNGYVQFNRFDEALELFRCMQTAGIRPDNFVLVSLLTGCAQTGALEQGKWI------------HGYINENRVTVDKVVGTALVDMYAKCGCIETALEVFYEIKER------DTASWTSLIYGLAMNGMSGRALDLYYEMENVGVRLDAITFVAVLTACNHGGFVAEGRKIFHSmeRHNVQpeHCSCLIDLLCRAGLLDEAEELIDKMRGESDETL-VPVYCSLLSAARNYGNVKIAERVAEKLEKVEVS-DSSAHTLLASVYASANRWEDVTNVRRKMKDLGIRKFP----------GCSSIEIDGVGHEFIVGdlSHPKMDEINSMLHQTTNLMLDLEHK------------------------    
 349 UniRef50_Q9MA85        Pentatricopeptide repeat-co...   400    1e-109  1  10.4%   4:581    97:641     ---LSLYAKCGKLVDAIKLFDEMPMRDVISQNIVFYGFLRNRETESGFVLLKRM--LGSGG-FDHATLTIVLSVCDTPEFCLVTKMIHALAILSGYDKEISVGNKLITSYFKCGCSVSGRGVFDGmhRNVITLTAVISGLIEnhEDGLRLFSLMRRGLVHPNSVTYLSALAACssQRIVEGQQIHALLWKYGIESELCIESALMDMYSKCGSIEDAWTIF----ESTTEVDEVSMTVILVGLAQNGSEEEAIQFFIRMLQAGVEIDANVVSAVLGVSFIDNSLGLGKQL------------HSLVIKRKFSGNTFVNNGLINMYSKCGDLTDSQTVFRRMPKRNY------VSWNSMIAAFARHGHGLAALKLYEEMTTLEVKPTDVTFLSLLHACSHVGLIDKGRELLNEMKehGIEpeHYTCIIDMLGRAGLLKEAKSFIDSLPLK----PDCKIWQALLGACSFHGDTEVGEYAAEQLFQT--APDSSSAHILinIYSSRGKWKERAKTIKRMKAMGVTK----------ETGISSIEIEHKTHSFVVEDKLHPQAEAIYDVLSGLFPVMVDEGYRPD----------------------    
 350 UniRef50_Q84MA3        Pentatricopeptide repeat-co...   399    1e-109  1  10.1%   6:577   218:759     -----MYSRSGSFRGARRVFDEMSFKDMISWNSLLSGLSQEGTFgeAVVIFRDM--MREGVELDHVSFTSVITTCCHETDLKLARQIHGLCIKRGYESLLEVGNILMSRYSKCGVLEAVKSVFHQmeRNVVSWTTMISS---NKDDAVSIFLNMRFDGVYPNEVTFVGLINAVKCNeiKEGLKIHGLCIKTGFVSEPSVGNSFITLYAKFEALEDAKKAFEDITFREI----ISWNAMISGFAQNGFSHEALKMFLSA-AAETMPNEYTFGSVLN----------AIAFAEDISVKQGQRCHAHLLKLGLNSCPVVSSALLDMYAKRGNIDESEKVFNEMSQK------NQFVWTSIISAYSSHGDFETVMNLFHKMIKENVAPDLVTFLSVLTACNRKGMVDKGYEIFNMMIeyNLEpeHYSCMVDMLGRAGRLKEAEELMSEVP----GGPGESMLQSMLGSCRLHGNVKMGAKVAELAMEMKPEL-SGSYVQMYNIYAEKEEWDKAAEIRKAMRKKNVSKEAGFSWIDVGDTEGSLTM-----QGFSSGDKSHPKSDEIYRMVEIIGLEMNLEG--------------------------    
 351 UniRef50_Q9XE98        Pentatricopeptide repeat-co...   399    1e-109  1  10.6%   6:600    90:668     -----LYAKFGLLAHARKVFEEMRERDVVHWTAMIGCYSRAGIVGEACSLVNEMRFQGIKPGP-----VTLLEMLSGVLEITQLQCLHDFAVIYGFDCDIAVMNSMLNLYCKCDHVGDAKDLFDQMerDMVSWNTMISGYagNMSEILKLLYRMRGDGLRPDQQTFGASLSVSGtcDLEMGRMLHCQIVKTGFDVDMHLKTALITMYLKCGKEEASYRVLETIPNK----DVVCWTVMISGLMRLGRAEKALIVFSEMLQSGSDLSSEAIASVVASCAQLGSFDLGASV------------HGYVLRHGYTLDTPALNSLITMYAKCGHLDKSLVIFERMNER------DLVSWNAIISGYAQNVDLCKALLLFEEMKFKTVqvDSFTVVSLLQACSSAGALPVGKLIHCIVIRSFIRPCSltALVDMYSKCGYLEAAQRCFDSISWK-----DVVSWGILIAGYGFHGKGDIALEIYSEFLHSGMEPNHVIFLAVLSSCSHNGMVQQGLKIFSSMVRDFGVEPNHEHLACVVDLLCRAKRIEDAFKFYKeiILDACRANGKTEVEDIICEDMIE--LKPGDAGHYVKLGHSFAAMKRWD---    
 352 UniRef50_Q6EUH6        Putative uncharacterized pr...   399    1e-109  1   9.5%   4:541   147:666     ---NALISmcGDVGAAEAVFGAMRNRTVVSWNAVIAGCVKNGYAERALEVFGEMAADGVGI--DRATVVSVLPACAQAKDLNTGRAVHRLVEDKGLGDYVAVKNALIDMYGKCRSLEDARRVFDHCkkDVVSWTAMIGAYVLnaFEAISLGCQMLMSGAapNGVTMVYLLSACASmsGKHAKCTHALCIRLGLKSDIAVETALIDAYARCGKMKLMRLTL----ERGSWR-AETWNAALSGYTVSGREKKAIELFKRMIAESVRPDSATMASILPAYAESADLKEGKNI------------HCFLLTLGFLRSTEIATGLIDVYSKAGDLDAAWALFQWLPEK------DVVAWTTIIAGYGIHGHARTAILLYDRMVESGGKPNTVTIATLLYACSHAGMIDEGIKVFKDMRnhGLMPnhYSCLVDMLGRAGRIEEAHRLIQDMP----FEPSTSVWGALLGACVLHKNVEFGEVAAKRLFQLDPE-NTGSYVLLGNIYAAADRWRDVQDVRRMMVERGLLKEPGSSLVEARSVQCRTAM--------------------------------------------------------------    
 353 UniRef50_C7J5C5        Os07g0203900 protein n=7 Ta...   399    1e-109  1   9.4%   7:572    59:605     ------YCRSRRLPEAQHLFDQMPARTAVTWNTLIHGHARSAAPGLAVAAFARMARAGVSPTASSVS--SVLVACVRLEDAAAGATLHSVGLMHGFCASVVVGTALVDMYAKCHHLGAAQQVFREMekNVATFTALVAGFvrRPHDAMLLVREMERSGVAPNLMTYSSLLSSFAseDIDHGKQVHCAVLKKGLEHDQFVLSALVTMYSKCGILENFVKV--QMSVS--CQDQVSFNSVISGLSCLGRGKEAFQHFLEMRRHGTDMDVFTFASVLKAIGSSSSLLEGRQV------------HTLILKIGYDSVVDVQNSLISMYARHGAIGESNGVFISMEA------PNLVSWNSLMSGCAQHGHGKEVVEMFEQMRRLHVQPDHITFLSVLTACSHVGLVDKGLEYFNLMKDKGyrTEHYACMVDLLGRAGYLNEAEYLINGMPIK----PGASVYRALLSACQIHGNLEIVIRVSKRLIELNPH-DSSVHVQLSNAFAGDGRWGNAAEIREAMSEYMSGGSLYDFLHKQHNVLDLPTLLKFAVDLCRGMCYLHQmiHRDLKSANLLMDK-------------------------------    
 354 UniRef50_B6U1A3        Pentatricopeptide repeat pr...   399    1e-109  1   9.0%   9:573   161:786     --------KAGDIASARRLFYGMPEKSVVSYTTMVDALMKRGSVRDAVELYevFRKMlcSVRPNVVTLICVIKACVGAGEFDLAMGVVGLAVKCNLfsIEVHNSLITLYLRMGDAAAAHRVFDDMEVRDVVSWTALLDVYagDTAEALKLYSQMLADGCRPNISCFSSVLSACATldLRGGTRIHANALKMGSSTNLFVSSSLIDMYCKCKQCTYAQRVFNSLPEK----NTVCWNSLISGYSWNGKMVEAEGLFNKMPAR----NSVSWNTMISGYAENRRFGDALNYFYAMLASggRMVHAEIVKLGIEDNIFMGTALSDMYAKSGDLDSSRRVFYQMPEK------NNITWTAMVQGLAENGFAEESISLFEDMIENGIAPNEHTFLAILFACSHCGLVEQAIHYFETMQAHGIPpkHYTCMVDVLARAGCLPEAEELL----MKVSSELDTSSWSSLLSACSTYRNKEIGERAAKKLHELE-KDNTAGYVLLSNMYASCGKWKDAAETRILMQGASLKK----------DAGCSWLQLRGQYHAFFSWKEKHPLSLEIYEILDLLMLEL------------------------------    
 355 UniRef50_Q9SX45        Pentatricopeptide repeat-co...   399    1e-109  1   9.5%   5:584    40:590     ----RLLRRCcqFRYARRLLCQLQTLSIQLWDSLIGHFSgtLNRRLSFLAYRHMR--RNGVIPSRHTFPPLLKAVFKLRDSNP-FQFHAHIVKFGLDSDPFVRNSLISGYSSSGLFDFASRLFDGAekDVVTWTAMIDGFVRNgsEAMVYFVEMKKTGVAANEMTVVSVLKAAGKvdVRFGRSVHGLYLETgvKCDVFIGSSLVDMYGKCSCYDDAQKVFDEMPSR----NVVTWTALIAGYVQSRCFDKGMLVFEEMLKSDVAPNEKTLSSVLSACAHVGALHRGRRVHCYM------------IKNSIEINTTAGTTLIDLYVKCGCLEEAILVFERLHEK------NVYTWTAMINGFAAHGYARDAFDLFYTMLSSHVSPNEVTFMAVLSACAHGGLVEEGRRLFLSMkrFNMEpdHYACMVDLFGRKGLLEEAKALIERMP----MEPTNVVWGALFGSCLLHKDYELGKYAASRVIK--LQPSHSgyTLLANLYSESQNWDEVARVRKQMKDQQVVKSPGF----------SWIEVKGKLCEFIAFDDKKpeSDDLYKTLDTVGVQMR----LPDELE-------------------    
 356 UniRef50_Q3E9N1        Pentatricopeptide repeat-co...   399    1e-109  1   8.4%   7:602   171:753     ------YSKCGFLQDACLVFDEMPDRDVVAWTAIISGHVQNGESEGGLGYLCKMHSAGSDVdpNPRTLECGFQACSNLGALKEGRCLHGFAVKNGLASSKFVQSSMFSFYSKSGNPSEAYLSFREldEDMFSWTSIIASLARSGdeSFDMFWEMQNKGMHPDGVVISCLINELGKMMlpQGKAFHGFVIRHCFSLDSTVCNSLLSMYCKFELLSVAEKLFCRISEEG---NKEAWNTMLKGYGKMKCHVKCIELFRKIQNLGIEIDSASATSVISSCSHIGAVLLGKSL------------HCYVVKTSLDLTISVVNSLIDLYGKMGDLTVAWRMFCEA-------DTNVITWNAMIASYVHCEQSEKAIALFDRMVSENFKPSSITLVTLLMACVNTGSLERGQMIHRYITETEHEMNLsaALIDMYAKCGHLEKSRELFDA-----GNQKDAVCWNVMISGYGMHGDVESAIALFDQMEESDVKPTGPTFLALLSACTHAGLVEQGKKLFLKMHQ-YDVKPNLKHYSCLVDLLSRSGNLEEAESTVM----SMPFSPDGVIWGTLLSSCMteMGIRMAERAvyYIMLANMYSAAGKWEE-    
 357 UniRef50_D1J120        Whole genome shotgun sequen...   398    1e-109  1  11.7%   4:578    80:632     ---NSLIDMYCKCGraRLVFESADELDSVSWNSLIAGYVRIGSNDEMLRLLVKM--LRHGLNLNSYALGSALKACGssSSIECGKMLHGCAVKLGLDLDVVVGTALLDTYAKIGDLEDATKIFKLMppNVVMYNAMIAGFLQmaNEAMYLFFEMQSRGMKPSEFTFSSILKACSTIEAFegKQIHAQIFKYNLQSDEFIGNALVELYSLSGSIEDGLKCFHSTPK----LDVVSWTSLIVGHVQNGQFEGGLTLFHELLFSGRKPDEFTISIMLSACANLAAVKSGEQI------------HAYAIKTGIGNFTIIQNSQICMYAKCGDIDSANMTFKETKN------PDIVSWSVMISSNAQHGCAKEAVDLFELMKGSGIAPNHITFLGVLVACSHGGLVEEGLRYFEIMKkhGITPnhSACIVDLLGRAGRLAEAESFI----MDSGFEGDPVMWRSLLSACRVHKATDTGKRVAERVIE--LEPeaASYVLLYNIYNDAGIQMPATEIRNLMKDRGVKKEP----------GLSWIEVGNVVHSFVAGDRSHPNSQVIYVQLEEMLEEIKKLDY-------------------------    
 358 UniRef50_Q9LER0        Pentatricopeptide repeat-co...   398    1e-109  1  10.7%   7:595   141:867     ------FCKVGRLSFAISLLRNRvsIDTVTYNTVISGLCEHGLADEAYQFLSEMVKMGIL--PDTVSYNTLIDGFCKVGNFVRAKALVDemVMSGFDPDVVTFSSIINRLCKGGKVLEGGLLLREMEEMSVYPNHVTYTTLVdyRHALALYSQMVVRGIPVDLVVYTVLMDGLFkgDLREAEKTFKMLLEDNQVPNVVTYTALVDGLCKAGDLSSAEFIITQMLEKSVIPNVVTYSSMINGYVKKGMLEEAVSLLRKMEDQNVVPNGFTYGTVIDGLFKAGKEEMAIELSKEMRlkEVKGLVKDMVSKGVTLDQINYTSLIDVFFKGGDEEaaDWAYKGMREKG--IEPDIATFNIMMNSQRKQGDSEGILKLWDKMKSCGIKPSLMSCNIVVGMLCENGKMEEAIHILNQMMLMEIHPnvYNTLIATLCKLGMTKKAAMVMGDMEARGFI-PDTVTFNSLMHGYFVGSHVRKALSTYSVMMEAGISPNVATYNTIIRGLSDAGLIKEVDKWLSEMKSRGMRPDDFT-YNALISGQAKIGNMKGSMTIyyNVLISEFANVGKMLQARELLKEMGKRGVSPNT-STYCTMISGLCK--------    
 359 UniRef50_C5Z5U0        Putative uncharacterized pr...   398    1e-109  1  10.4%   7:601    74:655     ------YAALGDLASARMVFDGTPRPDAYSYGVMLWCLVQAERHAEAVALHQDMRRRRPCPEadDFVLSLALKACIRSADYGYGTRLHCDAVKVGGADGFVMNSLVDMYAKAGD-LECARKVFERIprNVVSWTSMLSGCVQNGFAalLLFNKMRQDNVPPSEYTIATVITACSAlgLHQGRWMHGSVIKQGLMSNSFISAALLDMYVKCGELEDAQCVFDELS----YIDLVLWTTMIVGYTQNGNPLDALRLFLDKKFANIVPNSVTIATVLSASAQLRDLSLGRSI------------HGIAVKLGLVEYTVVVNALVDMYAKCQAVSEANRIFGSISNK------DVVAWNSMLSGYAENNMCNDALMLFKQMSLKGPSPDAISVVHALSASVCLGDLLIGKSFHGYAVKHAFLSNIytALLNLYNKCGDLPSARRVFDEMNDR-----NSVTWCAMIGGYGMQGDSAGSIDLFGEMLKDGVHPNDVAFTSILSTCSHTGMVTAAKRYFDSMAQHFNITPSMKHYACMVDVLARAGNLEEALEFIDNmfLHGCELHSRLQFGEEAIKRMMV--LHPERPDLY-VLISNLYTSNGMWE--    
 360 UniRef50_Q9ZUT5        Pentatricopeptide repeat-co...   398    1e-109  1   9.0%   3:578    60:647     --ASKLISFYTrfRQALHVFDEITVRNAFSYNALLIAYTSREMYFDAFSLF--LSWIGSScrPDSISISCVLKALSGCDDFWlaRQVHGFVIRGGFDSDVFVGNGMITYYTKCDNIESARKVFDEmeRDVVSWNSMISGYSQSgeDCKKMYKAMLAcdFKPNGVTVISVFQACGqsDLIFGLEVHKKMIENHIQMDLSLCNAVIGFYAKCGSLDYARALFDEMSEK----DSVTYGAIISGYMAHGLVKEAMALFSEMESIGLS----TWNAMISGLMQNNHHEEVINSFREMIrkGGKEIHAFAIRNGADNNIYVTTSIIDNYAKLGFLLGAQRVFDNCKDR------SLIAWTAIITAYAVHGDSDSACSLFDQMQCLGTKPDDVTLTAVLSAFAHSGDSDMAQHIFDSMlkYDIEpeHYACMVSVLSRAGKLSDAMEFISKMP----IDPIAKVWGALLNGASVLGDLEIARFACDRLFEMEPE-NTGNYTIMANLYTQAGRWEEAEMVRNKMKRIGLKKIPGT----------SWIETEKGLRSFIAKDSSCERSKEMYEIIEGLVESMSDKEY-------------------------    
 361 UniRef50_Q9SZK1        Pentatricopeptide repeat-co...   397    1e-109  1  11.2%   1:529    48:550     LGKSADFASYSSVI----LHSIRSVLSSFSYNTLLSSYAVCDKPRVTIFAYKTFVSNGFS--PDMFTFPPVFKACGKFSGIREGKQIHGIVTKMGFYDDIYVQNSLVHFYGVCGESRNACKVFGEMprDVVSWTGIITGFTRTgkEALDTFSKM---DVEPNLATYVCVLVSSGrgCLSLGKGIHGLILKRASLISLETGNALIDMYVKCEQLSDAMRVFGELEKK----DKVSWNSMISGLVHCERSKEAIDLFSLMqsSGIKPDGHILTSVLSACASLGAVDHGRWV------------HEYILTAGIKWDTHIGTAIVDMYAKCGYIETALEIFNGIRSK------NVFTWNALLGGLAIHGHGLESLRYFEEMVKLGFKPNLVTFLAALNACCHTGLVDEGRRYFHKMkeYNLFpeHYGCMIDLLCRAGLLDEALELVKAMPVK----PDVRICGAILSACKNRglMELPKEILDSFLDIEFE-DSGVYVLLSNIFAANRRWDDVARIRRLMKVKGISKVPGSSY--------------------------------------------------------------------------    
 362 UniRef50_B9T517        Pentatricopeptide repeat-co...   397    1e-109  1  10.9%   4:601    41:626     ---LSRYTRCSslTFACNLFDEMSHRDTVTWNTMISGYVNSGSLGSAWELYKSMKSFGLM--PDAYTFGSILKGVACACRLDVGQQVHSLIVKMGYEEHVYAGSALLDMYAKCERVRDAFMVFKCirRNSVSWNALIAGFVLEGDhaFWLLRCMEEEGVRLDDGTFSPLLTLLDEKKfkLTMQLHCKIIKHGVQFDNTVCNATITSYSQCGSLEDAERVFDGAVG---SRDLVTWNSMLAAFLAHDRKETAFKLFLDMQQFGFEPDIYTYTTIISACSH---------------KDNGKSLHGLVIKRGLEQLVPICNAVIAMysSSNSMEDALNVFHSMESK------DRVSWNSILTGFSQTGHSENALKLFVHMRFAVVDIDHYAFSAVLRSCSDLATLQLGQQIHVLTVKSGFESNDfsSLIFMYSKCGMIEDARKSFEK-----TAKDSSITWNSIMFAYAQHGQGDVALGLFFQMRDKKVKMDHITFVAALTACSHIGLVEQGRYLLKSMASDYGISPRMEHYACAVDLFGRAGYLDEAKALIESmlLGACRACGDIELAAQVASHLLE--LEPEEHCTYVILSNMYGHLKRWDE--    
 363 UniRef50_P0C898        Putative pentatricopeptide ...   397    1e-109  1   8.7%   6:579    50:596     -----MYCKCREPLMAYKVFDSMPERNVVSWSALMSGHVLNGDLKGSLSLFSEMG--RQGIYPNEFTFSTNLKACGLLNALEKGLQIHGFCLKIGFEMMVEVGNSLVDMYSKCGRINEAEKVFRRidRSLISWNAMIAGfaGYGSKALDTFGMMQEANirPDEFTLTSLLKACSSTGmyAGKQIHGFLVRSGFhpSSATItsLVDLYVKCGYLFSARKAFDQIKEK----TMISWSSLILGYAQEGEFVEAMGLFKRLQELNSQIDSFALSSIIGVFADFALLRQGKQM------------QALAVKLPSGLETSVLNSVVDMYLKCGLVDEAEKCFAEMQLK------DVISWTVVITGYGKHGLGKKSVRIFYEMLRHNIEPDEVCYLAVLSACSHSGMIKEGEELFSKLLehGIKpeHYACVVDLLGRAGRLKEAKHLIDTMPIK----PNVGIWQTLLSLCRVHGDIELGKEVGKILLRIDAK-NPANYVMMSNLYGQAGYWNE----------QGNARELGNIKGLKKEAGMSWVEIEREVHFFRSGEDSHPLTPVIQETLKEAERRLREegYV------------------------    
 364 UniRef50_UPI0000E1269A Os06g0152500 n=1 Tax=Oryza ...   397    1e-109  1  11.9% 158:602   113:554     -------------------------------------------------------------------------------------------------------------------------------------------------------------GYAPSVPAYNAVLLALSDASLPSARRFLSSMLRHGVAPNVYTYNILVRALCARGRLEEAVGVVGDMRGAGCAPNAVTYNTLVAAFCRAGELDGAERVVSLMREEgaKPNLVTFNSMVNGLCKAGRMEGARKVFDEMVREGLAplVAQMRERGLRMNEVTFTALIDGFCKKGFLDDALLAVEEM--RKCGIQPSVVCYNALINGYCKLGRMDLARELIREMEAKRVKPDVVTYSTIISGYCKVGNLDSAFQLNQKMLKKGVLPdtYSSLIRGLCEEKRLNDACELFENMLQLG-VQPDEFTYTTLIDGHCKEGNVEKALSLHDEMIRKGVLPDVVTYSVLINGLSKSARTKEAHRLLFKLYHEDPVPDNIKYDALMLC--CSKAEFKS-----VVALLKGFCMKGLMKEADKVYQSMLDRNWKLD-GSVYSILIHGHCRGGNVRKA-    
 365 UniRef50_Q9M302        Pentatricopeptide repeat-co...   397    1e-109  1  10.5%  24:594    72:646     -----------------------FKHTPLTFEVMIRKLAMDGQVDSVQYLLQQMKLQGFHCSEDLFI--SVISVYRQVGLAERAVEMFYRIKEFGCDPSVKIYNHVLDTLLGENRIQMIYMVYRDMKRDGFEPNVFTyaLCKnvDGAKKLLVEMSNKGCCPDAVSYTTVISSMCEVGlkEGRELAER-----FEPVVSVYNALINGLCKEHDYKGAFELMREMVEKGISPNVISYSTLINVLCNSGQIELAFSFLTQMLKRGCHPNIYTLSSLVKGCFLRGTTFDALDLWNQMIR-----------GFGLQPNVVAYNTLVQGFCSHGNIVKAVSVFSHMEEIGCS--PNIRTYGSLINGFAKRGSLDGAVYIWNKMLTSGCCPNVVVYTNMVEALCRHSKFKEAESLIEIMSKENCAPstFNAFIKGLCDAGRLDWAEKVFRQMEQQHRCPPNIVTYNELLDGLAKANRIEEAYGLTREIFMRGVEWSSSTYNTLLHGSCNAGLPGIALQLVGKMMVDG-KSPDEITMNMIILAYCKQGKAERAAQMlyTNVIWGLCRSNCREDGVILLERMISAGIVP-SIATWSVLINCFI---------    
 366 UniRef50_A5BJS3        Putative uncharacterized pr...   397    1e-109  1   9.7%  27:582    44:582     --------------------------DLYTCNALMNMYSKFWSLEegVQRFCDSKMLGGIPEPRE------IGKCSNSHDLPCELDERVAGIDQNGDLNQMSNIlvNTYKKVFDelGSLRKVFEMmkRDIVSWNTVISGNAQNgeDALMMVREMGNADLRPDSFTLSSVLPIFAevNLLKGKEIHGYAIRNGYDADVFIGSSLIDMYAKCTRVDDSCRVFYMLPQH----DGISWNSIIAGCVQNGMFDEGLKFFQQMLIAKIKPNHVSFSSIMPACAHLTTLHLGKQL------------HGYIIRSRFDGNVFIASALVDMYAKCGNIRTARWIFDKME------LYDMVSWTAMIMGYALHGHAYDAISLFKRMEVEGVKPNYVAFMAVLTACSHAGLVDEAWKYFNSmqDYRIIpeHYAAVADLLGRVGRLEEAYEFISDM----HIEPTGSVWSTLLAACRVHKNIELAEKVSKKLFTVDPQ-NIGAYVLLSNIYSAAGRWKDARKLRIAMRDKGMKKKPA----------CSWIEIKNKVHAFVAGDKSHPYYDRINEALKVLLEQMEREGYVLDT---------------------    
 367 UniRef50_Q6YUT6        Os02g0191200 protein n=6 Ta...   397    1e-109  1  11.6%   4:580    40:655     ---NSLVNCLehLRYALHLFDRMP-PSTFLFDTALRACSRAgdPHRPFLLFRRMR--RAGVRPDGFTfnSLIHMYIRLGLAADARRAFDEIH---VKDAVAWTMLISGLAKMGMLCDTQLLLSQAPVRDVISWTSLIAAYSrrAREAVGCFKTMLSHGIAPDEVTVIAVLSACAKldLELGRSLHLLVEEKGMPTSENLVVALIDMYAKCGDFGHAQQVFDAL---GRGPRPQSWNAIIDGYCKHGHVDVARSLFDEMEVR----DIITFNSMMTGYIHSGQLREALLLFMSMRrqQGRALHACIEQRLVEADIYLGTALLDMYMKCGRVDEATIVFQRMGKR------DVHTWTAMIAGLAFNGMGKAALEHFYQMRCDGFQPNSVSYIAVLTACSHSCLLNEGRLYFDEMrlYNIHpeHYGCMIDLLGRSGLLDEAMDLVKTMP----IQPNAVIWASILSACRVHKHIDLAQCAAEHLLK--LEPDegVYVQLYNIYIDSRQWENASKIRMLMEERQVKKT----------AGYSSITVAGQVHKFVVSDKSHPRILEIIAMLEEISHRLKSLGYSP-----------------------    
 368 UniRef50_Q10BR5        Calreticulin family protein...   396    1e-108  1   9.9%  10:565    95:665     ---------AGRLGDAREVFDGMGRRDLLAWSAMIGAYAIRGMYSDVLALAVTMVEEGVL--PDRFLITRILQACAYAEDLELGRALHSMAIRRGfkDVPVGNSVLVMYAKCGDLGWARKVFDKMVCRDLGTWNSMIFGYCRSaeEARHLLDSMRQEGTQPGVVTWNTLISSYArgDLDVAMELLGQMEESGVAPDVVTWTSLVSGFVHMDRSDEALRCFIRMRLAGVEpnVLSGNSLVDMYAKCGEIVAAWRIFSGIPDK----DIFSWNSMISGYAQAGYCGKAYELFCKMEN------------YGVRRNVITWNTMISGYIRNGDDERAFELFQMMESHG--VKRDTATWNILIAGSVHNGYYDRAIRIFRQMQALLRRPDYITILSIIPAFTNLVASWKVREIHACIFHHNLELdiANALINAYSKSGNLASACAVFD-M----HSSRNIISWNCIILAHVLHGSPNEALDLFCQMKQEGVVPDHTTLVTVIKAYGLTGKVSEGSQTFFNMANDYNITPELDHYAAMVDLLGRSGRLKEA----YELIDEMPLIPNSTVW--------------------------------------    
 369 UniRef50_B9GN13        Predicted protein n=2 Tax=P...   396    1e-108  1  10.6%  18:595    75:654     -----------------KIFKSGPKMGSytFYSLINNYANLGDFKSLEKVLDRMKCEKRVIF--EKCFIVIFKAYGKAHLPEKAVDLFDRmkRTGKSFNSVLNVII-----QEGLFHRALEFYnsISPNVLTFNLVIKAMcgLVDDAIQVFRDMTIRKCEPDVYTYCTLMDGLCKarIDEAVSLLDEMQIDGCFPSPVTFNVLINGLCKKGDLSRAAKLVDNMFLKGCIPNEVTYNTLIHGLCLKGKLEKAISLLDRMVSSKCVPNVVTYGTIINGLVKQGRALDGACVLALMEER------------GYCVNEYVYSTLISGLFKEGKSQEAMHLFKEMTVKGYEL--NTIVYSAVIDGLCRDGKPDDAVEVLSEMTNKGCTPNAYTCSSLMKGFFEAGNSHRAVEVWKDMAKHNFTQncYSVLIHGLCKDGKVKEAMMVWTQMLGKGCK-PDVVAYSSMINGLSIAGLVEDAMQLYNEMLCQGpqPDVVTYNILLNTLCKQSSISRAIDLLNSMLDRGCDPDLVT-CTIFLRMLREKLDPPQDGREFLDELVVRLLKRQRVLGASKIVEVMLQK-LLPPKHSTWARVVENLCK--------    
 370 UniRef50_B9R9I2        Pentatricopeptide repeat-co...   396    1e-108  1   9.3%  25:517   118:620     ------------------------QPTLLIFNYLIRGLSQSENPNEAIVMydLMYNQGILG--DNLTFIYLFKACSRVKDVLHGQVFHVQVLKLGFGSYLFIENSLIRMYGYFGELGYAQKVFDKmdRDLVSWNSLICGycNRFKEVLDLFNLMREANVTADSVTmnSLIDMYGRRGldLARRVFDRMQEK----NIVSWNAMLTGYATAGDLVAAKKLFNEMPIR----NVISWTCMISGCAQANQCSDALKLFQEMMDANVKPDEITVSSVLSACSHLGLLDTGQTVHEYM------------CRHDIKSDVYVGNALIDMYCKCGVVDKALEVFHDMKKK------DSVSWTSMILGLAVNGFVDNVFELFSQMLRDGLQPTHGSFIGILLACTHAGLVDKGLEYFESMEhyGLRPehYGCVVDLLSRSGELDRAYEFIKQMP----VVPDVVLWRILLSACKLHRNLVLAEIATSKLLELDPS-NSGNYVLLSNTYAGSDRWDDASRMRDLMV--------------------------------------------------------------------------------------    
 371 UniRef50_A5B7G0        Putative uncharacterized pr...   396    1e-108  1  10.4%  16:523    32:518     ---------------ARRIADQIDGLDTYTWNTIIRGYLEGNDPEEAILIYNHVRKKGLKV--DTYTLVFVIKACGLRPVILEGEQIHGQIFKLGFEFEVIIQTALLSMYGLFDEDCGLQQIFDEmqRDLVMWNALIAAYAHGNckVREVSYDMVSSNVKPNGVTAVSILSVCSSLrlREGKAVHGYVTKNLIEXDVFVHNALIDVYSKCGSIRDAVQVFQLMPMR----NVVSWTSLINGYSDNNCPNEALGFFKQMEAENIRPDEITVLGVVCMCSKLRSFELGEWISQYVVKTGL-----------VKESPAIANVLMDMHAKCGNIKRACQIFDGMEEK------TIVSWTIMIQGLAMYGHGLSALVRFCQMQREGFKPDSLVFLSLLSACSHAGLVDEGWRCFSSMedYHIXpeHYGCMVDILCRAGLVDEAFKFVQNMPIK----PDMIVWRTLLGACXXQGNISLANQVMNXLCELGPKKSED-YVLLSNLFASNAEWDNVXEVRKEMGDRGVTK--------------------------------------------------------------------------------    
 372 UniRef50_Q9LN69        Putative pentatricopeptide ...   396    1e-108  1  11.3%  18:602   215:844     -----------------QMISFEVSPDVFTCSIVVNAYCRSGNVDKAM-VFAKETESSLGLELNVVTYNSLINGYAMIGDVEGMTRVLRLMSERGVSRNVVTYTSLIKGYCKKGLMEEAEHVFELLKEKKLVADQHMYGVLMDgrDAVRVHDNMIEIGVRTNTTICNSLINGYCKSglVEAEQIFSRMNDWSLKPDHHTYNTLVDGYCRAGYVDEALKLCDQMCQKEVVPTVMTYNILLKGYSRIGAFHDVLSLWKMMLKRGVNADEISCSTLLEALFKLGDFNEAMKLWENVLARgaKEILDNVNIFRCKPAVQTYQALSHGYYKVGNLKEAFAVKEYMERKG--IFPTIEMYNTLISGAFKYRHLNKVADLVIELRARGLTPTVATYGALITGWCNIGMIDKAYATCFEMIEKGITLNvcSKIANSLFRLDKIDEACLLLQKIVdkKLLVPNNIVYNVAIAGLCKAGKLEDARKLFSDLLSSdfIPDEYTYTILIHGCAIAGDINKAFTLRDEMALKGIIP----------NIVTYNALIKGLCKLGNV------------DRAQRLLHKLPQKGITPNAI-TYNTLIDGLVKSGNVAEA-    
 373 UniRef50_Q9SUD8        Pentatricopeptide repeat-co...   396    1e-108  1  11.4%  12:553   126:697     -----------AFGVLALMLKRGFAFNVYNHNILLKGLCRNLECGKAVSLLREMR--RNSLMPDVFSYNTVIRGFCEGKELEKALELANEMKGSGCSWSLVTWGILIDAFCKAGKMDEAMGFLKEmeADLVVYTSLIRGFcgELDRGKALFDEVLERGDSPCAITYNTLIRGFCkgQLKEASEIFEFMIERGVRPNVYTYTGLIDGLCGVGKTKEALQLLNLMIEKDEEPNAVTYNIIINKLCKDGLVADAVEIVELMKKRRTRPDNITYNILLGGLCAKGDLDEASKLLYLMLKDSSYTD----------PDVISYNALIHGLCKENRLHQALDIYDLLVEkdSKIVRNSDTYTAMIDGFCKTGMLNVAKGLLCKMRVSELQPSVFDYNCLLSSLCKEGSLDQAWRLFEEMQRDNNFPDvfNIMIDGSLKAGDIKSAESLLVGMSRAGL-SPDLFTYSKLINRFLKLGYLDEAISFFDKMVDSGFEPDAHICDSVLKYCISQGETDKLTELVKKLVDKDIVLDKELTCTVMDYMCNSSANMDLAKRLLRVTD--------------------------------------------------    
 374 UniRef50_A9TCK1        Predicted protein (Fragment...   396    1e-108  1  14.8% 138:597    64:491     -----------------------------------------------------------------------------------------------------------------------------------------YASAGLLEKSVEALKRMEGHRCALTASAYNSLIDAFVKAGyqKALAVYRVMGQSGLRPDTYTFNVLMNAFKKAKRVDSVWKLFEEMQNQNCSPNVITYSILIDAVCKCGGVEKALKVFLDMKSRGCRPNIFTYTSMIDGLGKSGHVDKAFFLFEEMTSE------------GLVATRVVYNSLIHGLGRSGRADAAAKLFREMLSKG--LQPDHVTFTSLVYGLGVAGRASEARRIFQEARDVGCALDVNLYNVLIDTLCKSKRLDEAWEIFGELEEDGLVPDVynALMDGLCKSGRIHDAFILLGDMKRAGC-TPDVTVYNTLIDGLRKSGRVEEAGQLLLEMQSLGYEPDVVTYNTLIDESCKGGRIEDALRLFEEISAKGFA--NTVTYNTILNGLCMAGRVDEAYKLFNGMKQ---------ETVDGVID--------PDFV-TYTTLLNGARQAG------    
 375 UniRef50_Q0DJA4        Os05g0307200 protein n=6 Ta...   396    1e-108  1   9.5%  25:581   115:640     ------------------------APTAFSYNVAIRFFASS-RPHTSLRLFLHM--LRSAIRPDSYTLPFLLLAAARYPAPSLARAAHALLGKIGLNGHDHTVHSLITMYSYLDDPGAARKVFDGIprDVVSWNAMmyGRVGMNGEVGRMFRDMVKEgvAPNVVTVAVVLAACrqGDLVLGRWVEEWSWSARMEMDSLVGSALLGMYEKCGEIAEARRVFDTIIDK----DIVAWNAMITGYAQNGMSNEAISLFHSMRIAGMRPDKITLAGVLSACSAVGALELGSEL------------DGYASRRGLYSNVYVGTALVDMYAKCGDLDKAIEVFRKMPCK------NVASWNALICGLAFNGRGDEAIQHFQLMreEGLKPDDITFIGVLSACVHAGLVKDGKRWFNSLteFQIIpeHYSCMVDLLARSGHLEEAWDFIEKIPDKV----DAVMLGALLAACRKCKNVEIGERVINRIIQ--LEpnSWNYVVSSKIYASSDRLDDSAKMRGLMRERGVNKTP----------GCSWVEVSGKVLEFYAGDEPQHGADDMYQVLDLLVDEMRLEGYVPN----------------------    
 376 UniRef50_B9MU52        Predicted protein n=1 Tax=P...   396    1e-108  1  12.3%  26:601   168:762     -------------------------PDVIACNALLNALIQQKKVEIARKVYEEMV--KRDGCWDNYSVCIMVRGLCKERKVEEGRKLINDRWGKGCIPNIVFYNTLVDGYWKRGDVERANGLFKELKMKGFLPTTETYGIMINGlkAVdlLVEMKERGVDVNVQVYNSIVDAQIKHGCkeVGKTLRWITENGCEPDITTYNTLISGSCRDGKVHEAEELLEHAIKRGLSPNKLSYTPLIHVYCKQGKCLRAFDLFIGMTEKGHPLDLVAYGALVHGLVAAGEVDVALTVRDKMVER------------GVLPDANVYNVLMNGLCKKGRLSAAKLLLVEML--HQNLSLDAFVSATLVDGFIRHGKLDEAKKLFELTIAKGMDPGVVGYNAMIKGYCKFGMMNDALTCVQRMKDGDHSPdtYSTIIDGYVKQNDLHNALKLFGQMV-KQKCKPNVVTYTSLINGFCRTGDSSRAEKTFEEMRSSGLKPNVVTYTILIGCFCKEGKISKACSFFELMLLNRCIPNDVT-FNYLINGLTNniKASLMMDFFRTMISDglCHHKMVNAALQLRDKMTGKGIFPDPV-SFAALVYGLCLEGRSKE--    
 377 UniRef50_Q9FK93        Pentatricopeptide repeat-co...   396    1e-108  1  11.8%   4:603    73:638     ---NSLInkCRETVRARKLFDLMPERNVVSWCAMMKGYQNSGFDFEVLKLFKSMFFSGES-RPNEFVATVVFKSCSNSGRIEEGKQFHGCFLKYGLISHEFVRNTLVYMYSLcgNGEAIRVLDDLPYCDLSVFSSALSGycGAFKEGLDVLRKTANEDFVWNNLTYLSSLRLFSNLrlNLALQVHSRMVRFGFNAEVEACGALINMYGKCGKVLYAQRVFDDTHAQNIFLN----TTIMDAYFQDKSFEEALNLFSKMDTKEVPPNEYTFAILLNSIAELSLLKQG------------DLLHGLVLKSGYRNHVMVGNALVNMYAKSGSIEDARKAFSGMTFR------DIVTWNTMISGCSHHGLGREALEAFDRMIFTGEIPNRITFIGVLQACSHIGFVEQGLHYFNqmKKFDVQPdhYTCIVGLLSKAGMFKDAEDFMRTAP----IEWDVVAWRTLLNACYVRRNYRLGKKVAEYAIEK-YPNDSGVYVLLSNIHAKSREWEGVAKVRSLMNNRGVKKEP----------GVSWIGIRNQTHVFLAEDNQHPEITLIYAKVKEVMSKIKPLGYSPDVAGAFHDV------DEEQREDN    
 378 UniRef50_Q8L8A4        Fertility restorer n=4 Tax=...   396    1e-108  1  13.0%  12:527    56:565     -----------AFSLFRQMVTTKPLPSAVSFSKLLKALVHMKHYSSVVSIFREIHKLRIPV--DAFALSTVVNSCCLMHRTDLGFSVLAIHFKKGIPYNEVTFTTLIRGLFAENKVKDAVHLFKKLVrnICEPDEVMYGTVMDgqKAFDLLRLMEQGITKPDTCIYNIVIDAFCKDglDGATSLLNEMKQKNIPPDIITYTSLIDGLGKLSQWEKVRTLFLEMIHLNIYPDVCTFNSVIDGLCKEGKVEDAEEIMTYMIEKGVEPNEITYNVVMDGYCLRGQMGRARRIFDSMIDK------------GIEPDIISYTALINGYVEKKKMDKAMQLFREISQNG--LKPSIVTCSVLLRGLFEVGRTECAKIFFDEMQAAGHIPNLYTHCTLLGGYFKNGLVEEAMSHFHKLERRREDTNiyTAVINGLCKNGKLDKAHATFEKLPLIGL-HPDVITYTAMISGYCQEGLLDEAKDMLRKMEDNGCLPDNRTYNVIVRGFFRSSKVSEMKAFLKEIAGKSFSFEAAT----------------------------------------------------------------------------    
 379 UniRef50_B9HIP8        Predicted protein n=1 Tax=P...   396    1e-108  1  10.0%   3:580    34:582     --AISLLQNCKNPKDLIQLHTLLIKTSlyAFGRLLLSFASFDNLGSLNYAQKLFDT--VDIPRNSFMYTTMIKAYANFGNPREAFAFYSRMlrYVYPNDFTFTYVFSACSKFNGVFehAQMIKFPFEFGVHSWNSLlyGKVGEVGIVVrvFDKIE----GPDVVSWNCLINGYVkgDLDEARRLFDEMPER----DVVSWTIMLVGYADAGFLSEASCLFDEMPKR----NLVSWSALIKGYIQIGCYSKALELFKEMQVAKVKMDEVIVTTLLSACARLGALDQGRWL------------HMYIDKHGIKVDAHLSTALIDMYSKCGRIDMAWKVFQETGDKK------VFVWSSMIGGLAMHSFGEKAIELFAKMIECGIEPSEITYINILAACTHSGLVDVGLQIFNRMVEnkPRMQHYGCIVDLLGRAGLLHDAFRVVETMPVK----ADPAIWRALLSACKLHRNVELGEQVGRILIK--MEpnDMNYVLFSNVYAAVNRWDISGKLRREMKVRGMQKNP----------GCSSIELNGAVHEFVSRDHSHPQSQVIYELLHILTNHMVQEDHEP-----------------------    
 380 UniRef50_Q9SJK9        Pentatricopeptide repeat-co...   395    1e-108  1  10.6%   7:574    14:618     ------LAKSGRIASARQVFDGMPELDTVAWNTMLTSYSRLGLHQEAIALFTQLRF--SDAKPDDYSFTAILSTCASLGNVKFGRKIQSLVIRSGFCASLPVNNSLIDMYGKCSdfEAALDVFVEMPKRvfAWNIMihAHCGKLESCLSLFKEMLESEFKPDCYTFSSLMNACssSNVVYGRMVHAVMLKNGWslTQVSWNSIIDACMKIGETEKALEVFHLAPEK----NIVTWTTMITGYGRNGDGEQALRFFVEMMKSGVDSDHFAYGAVLHACSGLALLGHGKMI------------HGCLIHCGFQGYAYVGNALVNLYAKCGDIKEADRAFGDIANK------DLVSWNTMLFAFGVHGLADQALKLYDNMIASGIKPDNVTFIGLLTTCSHSGLVEEGCMIFESMVKDylEVDHVTCMIDMFGRGGHLAEAKDLATTYSSLVTDSSNNSSWETLLGACSTHWHTELGREVSKVLKIA--EPSemSFVLLSNLYCSTGRWKEGEDVRREMVERGMKKTP----------GCSWIEVGNQVSTFVVGDSSHPRLEELSETLNCLQHEMR-----------------------------    
 381 UniRef50_Q9SS97        Putative pentatricopeptide ...   395    1e-108  1  10.2%   6:575   105:645     -----MYIKCGRMIEALRMFDELEKPDIVTWSSMVSGFEKNGSPYQAVEFFRRMV-MASDVTPDRVTLITLVSACTKLSNSRLGRCVHGFVIRRGFSNDLSLVNSLLNCYAKSRAFKEAVNLFKMieKDVISWSTVIACYVQNGaeALLVFNDMMDDGTEPNVATVLCVLQACAAadLEQGRKTHELAIRKGLETEVKVSTALVDMYMKCFSPEEAYAVFSRIPRK----DVVSWVALISGFTLNGMAHRSIEEFSIMleNNTRPDAILMVKVLGSCSELGFLEQAK------------CFHSYVIKYGFDSNPFIGASLVELYSRCGSLGNASKVFNGIALK------DTVVWTSLITGYGIHGKGTKALETFNHmkSSEVKPNEVTFLSILSACSHAGLIHEGLRIFKLMvdYRLAPnhYAVLVDLLGRVGDLDTAIEITKRMP----FSPTPQILGTLLGACRIHQNGEMAETVAKKLFE--LESNHAGYYMLmnVYGVKGEWENVEKLRNSVKQRGIKKGLAE----------SLIEIRRKVHRFVADDELHPEKEPVYGLLKELDLHMKE----------------------------    
 382 UniRef50_B9RKB2        Pentatricopeptide repeat-co...   395    1e-108  1  11.1%   6:602   137:723     -----LYAKCGSLAFARKVLVGMRERDVVSWTALIAGYVSEGCGSDGVKAYCEMRKENIC--PNEFTLATVLKASSMCSDIKFGKLIHLEAIKTGLLLDLFVGSALVDLYAKFGEMELADRVFFGmeKNNVSWNALLNGYAQRGDgvLKLFCRMLECEMNFTNYTLSTVLKGCasGNLREGKALHSLSIRRAYELDEFLGCNLVDMYSKCGMAYEALKVFNMIEE----PDIVAWSAIITGLDQQGHSQEAAELFHLMRQKGVRPNQFSFASVISAATNVGDLYLGQSI------------HCCICKYGYESDNSVGNALITMYMKSGFVQDGIRVFDTMTNR------DLVSWNALLSGFYDFETSDQGLRIFCQMLMEGLVPNLYTFVGVLRSCSSLLNVWFGKQVHAHIIKNSLDGNDftALIDMYAKNRCLEDADVAFNKLTNR-----DLFTWTVIIAGHSQTDKAEKAVKYLGQMLREGIKPNEFTLASCLSGCSRMATLGNGQQLHSLAIKSGHS-GDVFVSSALVDMYGKCGCMEDAEAIfwNTIICGYSQHGQGQKALEAFRMMLDEDIDPDEVTFIGVLAagWVEEGKKHFD-    
 383 UniRef50_Q9LU94        Putative pentatricopeptide ...   395    1e-108  1  10.2%   6:585   145:701     -----MYAKCERVEDAFEAFKEISEPNSVSWNALIAGFVQVRDIKTAFWLLGLM-EMKAAVTMDAGTFAPLLTLLDDPMFCNLLKQVHAKVLKLGLQHEITICNAMISSYADCGSVSDAKRVFDGlsKDLISWNSMIAGFSKhkESAFELFIQMQRHWVETDIYTYTGLLSACSGEEHqfGKSLHGMVIKKGLEQVTSATNALISMYIQfgTMEDALSLFESLKSK----DLISWNSIITGFAQKGLSEDAVKFFSYLRSSEIKVDDYAFSALLRSCSDLATLQLGQQI------------HALATKSGFVSNEFVISSLIVMYSKCGIIESARKCFQQISSKH-----STVAWNAMILGYAQHGLGQVSLDLFSQMCNQNVKLDHVTFTAILTACSHTGLIQEGLELLNLMevYKIQpeHYAAAVDLLGRAGLVNKAKELIESMP----LNPDPMVLKTFLGVCRACGEIEMATQVANHLLE--IEpdHFTYVSLSHMYSDLKKWEEKASVKKMMKERGVKKVP----------GWSWIEIRNQVKAFNAEDRSNPLCQDIYMMIKDLTQEMqdNGVDADSLHA------------------    
 384 UniRef50_D1IPH9        Whole genome shotgun sequen...   395    1e-108  1   9.4%   6:527    79:590     -----MYAKCGHLPHAHKLFDHFPHRNAVSWSVLIAGFVQHNRPSCALVLFSQMHAAG--VNINEFTLVSALHACsySYYLSRLYQVYAFVVRLGFEWNVFLMNVFMTALIRSRKLAEALEVFEACrkDVVSWNAVMAGLVQFCceVPGFWRRMCCEGVKPDNFAFSGVLSGLAagDGGMGVQVHGQLVKCGHGGEVCIGNSLVDMYLKCGSLENGIKAFEEMVER----DVCTWNQMAAGCLNCGKPGRALELIEEMRRSGVRMNKFTLATALTACANLASLEEGKK------------AHGLRIKLGNDVDVCVDNALLDMYAKCGSMGDALKVF------GTVEARSVVSWTTMIMGFAQNGQARKALEIFEQMRAEGKAPNYITFICVLYACSQGGLINEGWEYFLSmrDHGISpdHYACMVDMLGRAGHIKEARELIRRMP----FQASVLVWQTLLGACRVHGDVETGLLAAKQALSLD-KHDSSTYVVLSNMFADGRNWKGVGSLRELMETRDVKKMPGS----------------------------------------------------------------------------    
 385 UniRef50_Q6AUZ9        Pentatricopeptide, putative...   395    1e-108  1  11.2%   8:596   278:924     -------GRcsALRILDDMEKNGIEADLYTYNIMIDKLCKLKRSARAYLLLKRMRE--VNLTPDECSYNTLIHGFFGEGKINLAIYIFNQMLRQSlpSVATYTALIDGYCRNGRTDEARRVLYEmmLADGIDPDVITYSALIniAEAEQFKQYMSRMKISFDVASFNCIIDSYcrGNVLEAFSVYDNMVRHGWPPDICTYGSLLRGlcKHGTLDEALDLCEKMVTRNILPDTYTYTILLDGFCKRGKVVPALILLQMMLEKGLVPDTIAYTCLLNGLVNEGQVKAASYMFQEIICKE-----------GLYADCIAYNSMMNGYLKGGQINEIERLMRNMHEN--EVYPSSASYNILMHGYIKKGQLSRTLYLYRDMVKEGIKPDNVTYRLLIFGLCEYGLIEIAVKFLEKMVLEGVFPdaFDILIKAFSEKSKMSNALQLFSYMKWL-HMSPSSKTYVAMVNGLIRKNWLQQSYEILHDMVESGLQPKHTHYIALINAKCRVGDIDGAFELKEDMKALGV-VPSEVAESSIVRGLCKCGKVEEAIIVFSSIMRagLCKEFKIDDAFHLKQLMESCGLKVDVV-TYNVLITGLCNK-------    
 386 UniRef50_C7SI28        Putative PPR repeat protein...   395    1e-108  1  10.9%  19:522     1:551     ------------------MNRNGFKHTIESYCIVAHILFCARMYYDANRILREMveLKDCDVFDVlvFDALFSVLIEEDMFEEALQCFSKMKRCRVFPKTRSCNGLLHKFARLGKTDGMKRFFKDmkPTVFTYNIMIDCMcgDIEAASGLFEEMKFRGLIPDTVTYNSMIDGYGkgRLDDTVYFFEEMKSMSCEPDVITYNTLINCFCKFGKLPKGLEFFREMKQSGLKPNVVSYSTLVDAFCKEDMMQQALKFYVDMRRLGLVPNEHTYTSLVDAYCKIGNLSDAFRLADEMSQ------------VGVEWNVVTYTALIDGLCDVERIKEAEELFGKMVTAG--VIPNLASYTALIHGFVKAKNMDRALELLDEMKGRGIKPDLLLYGTFIWSLCGLEKIEAAKVVMNEMQEKGIKANTfcVLIDGLCKNKLVSKAIDYFGRISNDFGLQANAAIYTAMIDGLCKEDQVEAATTLFEQMAQKGLVPDRTAYTSLIDGKLKQGNVVQALALRDKMAEIGVE---------------------------------------------------------------------------------    
 387 UniRef50_B9N0R5        Predicted protein n=6 Tax=M...   394    1e-108  1   9.3%  77:592    31:548     ----------------------------------------------------------------------------KCQSMSQLRQYHSQIIRLGLSSHNHLIPPLINFCARASTSDaaLKLFDSiqPDAFLYNTIIKGFLHstNSILLLYSHMLQNSVLPNNFTFPSLLIACRK--IQHGMQIHAHLFKFGFGAHSVCLNSLIHMYVTFQALEEARRVFHTIP----HPDSVSWTSLISGYSKWGLIDEAFTIFQLMPQK----NSASWNAMMAAYVQTNRFHEAFALFDRMKAEngKWIHEYIKRNGIELDSKLTTAIVDMYCKCGCLEKALQVFHSLPLPCRWIS----SWNCMIGGLAMHGNGEAAIQLFKEMERQRVAPDDITFLNLLTACAHSGLVEEGRNYFSYMIryGIEpeHFGCMVDLLGRAGMVPEARKLIDEMP----VSPDVTVLGTLLGACKKHRNIELGEEIGRRVIE--LEPNNSgyVLLANLYANAGKWEDAAKVRKLMDDRGVKKAPGF----------SMIELQGTVHEFIAGERNHPQAKELHAKVYEMLEHLKSVGYVADT----NGVLHG-----------    
 388 UniRef50_Q9CA54        Pentatricopeptide repeat-co...   394    1e-108  1   9.5%  16:603    58:570     ---------------ARRLLLCFPEPDAFMFNTLVRGYSESDEPHNSVAVFVEMMRKGFVF-PDSFSFAFVIKAVENFRSLRTGFQMHCQALKHGLESHLFVGTTLIGM----------------------------YGGCGCVEFARKVFDEMHQ----PNLVAWNAVITACFRgdVAGAREIFDKMLVR----NHTSWNVMLAGYIKAGELESAKRIFSEMPHR----DDVSWSTMIVGIAHNGSFNESFLYFRELQRAGMSPNEVSLTGVLSACSQSGSFEFGK------------ILHGFVEKAGYSWIVSVNNALIDMYSRCGNVPMARLVFEGMQEKRC-----IVSWTSMIAGLAMHGQGEEAVRLFNEMTAYGVTPDGISFISLLHACSHAGLIEEGEDYFSEMKryHIEPehYGCMVDLYGRSGKLQKAYDFICQMP----IPPTAIVWRTLLGACSSHGNIELAEQVKQRL--NELDPNNSglVLLSNAYATAGKWKDVASIRKSMIVQRIKKTTA----------WSLVEVGKTMYKFTAGEKKKGIDIEAHEKLKEIILRLkeAGYTPE-------VASALYDVEEEEKED    
 389 UniRef50_Q1KSA8        Putative uncharacterized pr...   394    1e-108  1  10.1%   3:569   158:696     --ANSLVALYAKLgdAERVFDGMPARDIVSWNTMVDGYVSNGMGALALACFREMNDA-LQVGHDSVGVIAALAACCLESALALGREIHGYAIRHGLEQDvvGTSLVDMYCKCGNVFFAENVFAKMPLRTVVTWNCMIGGYALnpVDAFDCFMQMRVDGFQVEVVTAINLLTACAQTESSlgRSVHAYVVRRHFLPHVVLETALLEMYGKVGKVESSEKIFGQITDK----TLVSWNNMIAAYMYMEMYQEAIALFLELLNQPLYPDYFTMTTVVPAFVLLGSIRQCKQM------------HSYIVKLGYGDSTLIMNAVMHMYARCGNIVASREIFDKMPGK------DVISWNTIIIGYAIHGQGKIALEMFDEMKCSGMEPNESTFVSVLTACSVSGLEAEGWKEFNSmqEYGMIpeHYGCMTDLLGRAGELREVLRFIENMP----IAPTSRIWGSLLTASRNKNDIDIAEYAAERIFQLEHN-NTGCYVVLSSMYADAGRWEDVERIRSLMKEKGLRRTEAR----------SLVELNNKECSFVNGDMSHPQSEKIHEFSDIL----------------------------------    
 390 UniRef50_Q0IV66        Os11g0109600 protein n=3 Ta...   394    1e-108  1  11.2%  36:580  526:1035     -----------------------------------RALTSSALHLHALRVHCLL------PNPSHLTIPIALKSASRLPhlRAGEQLHARSLKLPSHTNPHVLTSLLSLYAKCGLLHRAQRVFDEmhPSTVPWTALITAyaGDLREAVHVARNAFANGMRPDSFTAVRVLTACAraDLATGETVWRAAEQEGVAQSVFVATAAVDLYVKCGEMAKAREVFDKMRHK----DAVAWGAMVGGYASNGHPREALDLFLAMQAEGMKPDCYAVAGALSACTRLGALDLGRQAIR------------MVDWDEFLDNPVLGTALIDMYAKCGSTVEAWVVFQQMRKK------DIIVWNAMILGLGMTGHEKIAFALVGQMEKSGVKLNDNTFIGLLCSCTHTGLIQDGRRYFHNMTkyHISpeHYGCMVDLLSRAGLLQEAHQLVDDMP----MPANAVILGALLGGCKIHRNTELAEHVLKQLIL--LEpnSGNYVMLSNIYSNRGRWEDAAKLRLDMKAKGVEKVPA----------CSWVEFEGKVHEFRVGDKSHPLSDQIYKKLDELGLEMKTMGYEP-----------------------    
 391 UniRef50_Q7XQN7        OSJNBa0084A10.18 protein n=...   394    1e-108  1   9.8%   7:567   106:645     ------YARCGRVAEARELFNRIPDRNVVSWNAMVSGYARNGMVKRARELFDMM------PWRDDVSWLTMISGYIKRKHVREARELFDSMPS---PPTSVCNALLSGYVELG-YMRAAEVLFGQMqrNPVSWNVMITGYARAGSmaQRLFDEMPEK----DVLSRTAIMRGYLQNgvDAAWKVFKDMPHR----DTVAWNTMMDGFVRNDRLDDALKLFSEMPDR----DQISWNAILQGYVQQGDMDSANAWFRRAPNK----DAISWNTLISGYKDEG----ALSLLSEMIRGggKMVHLWAIKTGFEHDALVMSSLISMYSKCGLISEASQVFELILQR------DTVTWNAMIATYAYHGLADEALKVFDMMTKAGFRPDHATFLSILSACAHKGYLYEGCYHFRSMQEdnLVpdHYSCMVDLLGRSGFIHQAYDFTRRIPSDHR----TTAWETLFSVCNSHGEIQLGEIIARNVLKARPS-DGGMYTLLSNIYAAKEMWSSAASVRGFMKERGLKK----------ETGCSWIELKGEVVTFSSNDSNHPLIEQICQEAH------------------------------------    
 392 UniRef50_UPI00019833F6 PREDICTED: hypothetical pro...   394    1e-108  1   9.3%   4:602   175:776     ---NSLISMYIKIGKirRVFCDMPKRDIVSWNSLITAYARNGNWAEAFNLFLSMKSAEDLF-PNRITFLGLLLACGQIGNLDLGKSIHGHLICTGLISDLRLGTAMVNMYAKCDRVECAQTIFEefDRSLVSWNSLIAGYSHNgqKATLLYERMieSNLKPDAVTFANVVPAYAsaDIRRIRSIHTFIVKKGLDMdvVLGTAMVDAYGKCLDIKAAKSLFSGIKK----PNTATWNAMIAGNNLNHHADKSMLLFLEMLQSKVFPDAITMVMLFQSCGEIGSLKEGTMVHCYCLSK------------GFSPHLTVGNAIIDMYMRFGCVKSSQLLFNEMSLK------NVVTWNTMLFGYVKIGDSAMAMRVFRQMQSEnyKPDSVTMISSIQalTSCG----AEIAHGFIAKLGLDSETlnSLIDAYAKTGFIEKARSLFMQM----GNLRDQSSWNIMIAGCGMNGQGREACELVSHMEEDGYRPNSITFTSLLSSCSHSGLIEEGCWYFDMMMRKYKIQPGLEHWTCIIDMHARAGRLEEacDAVWGALLSACRMNMNMELGLLAGEKLLK--LAPDN-CGYRTLLSNLYASGKRWDE-    
 393 UniRef50_D1I3L9        Whole genome shotgun sequen...   394    1e-108  1   7.4%  67:569    66:549     ------------------------------------------------------------------TLNKFMVCcpSIGNLHYAERIFNYIDIPGL----FIYNLVIKAFTKNGSFRKAVLLFRQLREEGLSPDNFTygCLGEVREGEKVYGFVVKSGLEFDTYVCNSLMDMYAegRVQNLRQVFEEMPQR----DVVSWNVLISGYVKCRRYEDAVDVFRQIFNDMPIKTVICWTSMVSGYVNCGQLDEARELFERSPVR----DVVLWTAMINGYVQFNRFDDAVALFREMQIKRVSPDRFTLVA--LLTDAVVGTALIEMYAKCGFIEKSLEIFNGLKEK------DTASWTSIICGLAMNGKTSKALELFAEMVQTGVKPDDITFIGVLSACSHGGLVEEGRKHFRSMTayQIEpeHYGCLIDLLGRAGQLDEAEELIEKSPNVNNEVI-VPLYGALLSACRTHGNVEMGERVAKRliESG---DSSVHTLLANIYASADRWEDVTKVRRKMKDLGVKKVP----------GCSSVEVNGIVHEFLVGDASHPEMREIYSMLDSI----------------------------------    
 394 UniRef50_D1I6R6        Whole genome shotgun sequen...   393    1e-108  1  11.9%  81:603   116:595     --------------------------------------------------------------------------------LAYAKTIFHHLQN---PPPSLYNSLIRALSSSKTPLE-ALPLYHTMLQSGLKPDHMTYPFVIKAcfGLLVHTHVVKSGFECDSYIVSSLIHLYANgdLGAAKQLFNLCSAR----DVVSWNAMIDGYVKHVEMGHARMVFDRMVCR----DVISWNTMINGYAICGKPNEALALFDQMRAVGVKPTEATVVSLLSACAHLGALDKGLHL------------HTYINDNRIEVNSIVGTALVDMYAKCGKISLATQVFNAMESK------DVLAWNTIIAGMAIHGNVKEAQQLFKEMKEAGVEPNDITFVAILSACSHAGMVDEGQKLLDCMSsyGIEpeHYGCVIDLLARAGFLEEAMELIGTMP----MEPNPSALGALLGGCRIHGNFELGEMVGKRLI--NLQphSGRYILLSNIYAAAKKWDDARKVRNLMKVNGISKVP----------GVSVIELKGMVHRFVAGDWSHPESNKIYEKLNEIHTRLKSagYSADTG---NVLLD-------MEEED    
 395 UniRef50_UPI00015C94D5 PREDICTED: hypothetical pro...   393    1e-108  1   8.7%  63:588    89:582     --------------------------------------------------------------PDSYSWTTMIRGFVEAKNPEKALEFYGLMRQRGVELNKFTFLFVLKAYGLRPSYQEGRIVHGKLVKVGFCYDVFTRnkCGSITDAHLLFDEMPNH----NVVTWNTMITGcgCGDTERARRLFGEMPER----NVGSWNAVVGGYSKLGHVDIARSLFDLMPER----DVVSWGSMISAYVQNGRAAEALELFKEMMLAGVSADSIIITSILSACAQIGALDMGRWIHAYMKRSKLRN------------DVFLDTALVDMYAKCGCIDTAFGVFNTMPRK------NLCSWNAMLSGLAIHGHGFAALELFKQMESTGVGPNDITFVAVLSACSHIGSVEEGWKKFNQmkEFNITpeHYGCMVDILCRQGLINEAKEMIRTMP----LEPNVVIWGALLNACKVHGYTNVGEDVVGYIQKL-VSEDGGCYVLLSNIFAAKSQWNEVEKTRKMMKQMGVEKKIPGY---------SSIELDSVVHDFFAEDRLHSKWREVSTVIERLNTHLEVKGYEAnsLVLYNI---------------    
 396 UniRef50_B9RQ16        Pentatricopeptide repeat-co...   393    1e-108  1   8.5%  28:533   104:640     ---------------------------VYWWNALIRRAVRLGLLQHSLSLFRTMRRLNWS--PDHYTFPFVFKACGELPSFLHGSCIHAIVCSTGFDSNVFVCNAVVAMYGRCGASSYARQMFDElvFDLVSWNSMIAVYLQSGdsGIELFRRMWKVgiVPDAVSLVNVLPACAsgDWLCGKQVHGFAIRYGLFEDVFVANSLVDMYAKCgkFEDALGLFEKMREEKIQLDVVSWSAVIAGYAQRGLGYEALNVFRQMQVCGLRPNEVTLVSLLSGCASVGALLHGKETHCYSIKCVLNFDRSDPRDELL-----VVNAIIDMYTKCKDINVGRAIFNSIPPKD----RNVVTWTAMIGGYAQHGEANDALELFSQMLkrSVKPNAFTISCALMACARLAALRFGRQIHAFVLRDQYDCDVlnCLIDMYSKSGDMDAARLVFDNMKHR-----NTVSWTSLMTGYGMHGHGEEAIKVFDEMRREGLVSDGITFLVVLYACSHSGMVDEGIKYFHDMCKEFSEKNENSALEIKP----------------------------------------------------------------------    
 397 UniRef50_B9GTD0        Predicted protein n=2 Tax=r...   393    1e-108  1  10.3%  56:581     1:493     -------------------------------------------------------MLRLGIQPDNFTFPFIIKACSCLRHFEFGIRIHQDVVKFGYQSQVFISNSLITMYGKCDKYELSRQVFDEMpkNAVSWSAIIGACLQdcKEGFSLFRQMLSEGSRPSraILNAM--ACVRSH-EEADDVYRVVVENGLDFDQSVQSAAAGMFARCGRVEVARKLFDGIMSK----DLVTWATTIEAYVKADMPLEALGLLKQMMLQGIFPDAITLLGVIRACSTLASFQLAHIVHGIITTGFFYNQLLAVE-----------TALIDLYVKCGSLTYARKVFDGMQER------NIITWSAMISGYGMHGWGREALNLFDQMKAS-VKPDHITFVSILSACSHSGLVAEGWECFNSmrDFGvrPEHYACMVDILGRAGKLDEACDFIERMPVR----PNAAVWGALLGACRIHLNVDLAEMVARALFDLDPH-NAGRYVILYNIYTLTGKRKEADSIRTLMKNRGVKKI----------AGYSVIEIKNKLYAFVAGDRSHPQTDLIYSELERLMDRIRQEGYTPD----------------------    
 398 UniRef50_Q1PEU4        Pentatricopeptide repeat-co...   393    1e-108  1   7.8%  13:578    26:553     ------------IGYARKLFDQRPQrdSFLSNSMIKAYLETRQYPDSFALYRDLRKETC-FAPDNFTFTTLTKSCSLSMCVYQGLQLHSQIWRFGFCADMYVSTGVVDMYAKFGKMGCARNAFDEmhRSEVSWTALISGycGELDLASKLFDQMPHVK---DVVIYNAMMDGFVkgDMTSARRLFDEMTHK----TVITWTTMIHGYCNIKDIDAARKLFDAMPER----NLVSWNTMIGGYCQNKQPQEGIRLFQEMQatSLDPDDVTILSVLPAISDTGALSLGEWCHCFVQRKKLDKK------------VKVCTAILDMYSKCGEIEKAKRIFDEMPEKQ------VASWNAMIHGYALNGNARAALDLFVTMMIEE-KPDEITMLAVITACNHGGLVEEGRKWFHVMREMGlkIEHYGCMVDLLGRAGSLKEAEDLITNMP----FEPNGIILSSFLSACGQYKDIERAERILKKAVE--LEpnDGNYVLLRNLYAADKRWDDFGMVKNVMRKN----------QAKKEVGCSLIEINYIVSEFISGDTTHPHRRSIHLVLGDLLMHMNEEKY-------------------------    
 399 UniRef50_Q0JI98        Os01g0815900 protein n=9 Ta...   393    1e-107  1  11.4%  12:527    36:560     -----------DLVFAHKVFDQIEAPTTFLWNILIRGLAQSDAPADAIAFYK--KAQGGGMVPDNLTFPFILKACARINALNEGEQMHNHITKLGLLSDIFVSNSLIHLYAACGNLCYARSVFDEmvKDVVSWNSLICGycNRFKDILALFKLMQNEGVKADKVTMIKVVSACTrgDYSMADYMVRYIEDYCIEVDVYLGNTLVDYFGRRGQLQSAEKVFFNMKVRNikKDLISWSSMISGYSQANHFSDALEIFRQMQRAKVKPDAIVIASVVSSCAHLGALDLGKWV------------HEYVRRNNIKADTIMENSLIDMYMKCGSAKEALQVFKEMKEK------DTLSWNSIIIGLANNGFEKESLNLFQAMLTEGFRPNGVTFLGVLIACANAKLVEEGLDHFESMKrySLEPqhYGCVVDLLGRAGQLEKALRFITEMP----IDPDPVVWRILLGSCNTHGDVAIAEIVTKKL--NELEpnSGNYTLLSNAYASAHRWSEAMNVRQCMADTDVRKSPGC----------------------------------------------------------------------------    
 400 UniRef50_Q7XJU7        OSIGBa0124N08.1 protein n=2...   393    1e-107  1   8.8%   1:589   199:762     LVANALVGMYAKCGLLDSALRVfdGRDVASWNSAISGCVQNGMFLEALDLFRRMQSDGFS--MNSYTTVGVLQVCAELAQLNHGRELHAALLKCGTEFNIQCNALLVMYARCGWVDSALRVFREIGDKDYISWNSMLSCYVQnyAEAIDFFGEMVQNGFNPDHACIVSLLSAVGhgRLINGREVHAYAVKQRLDSDLQIANTLMDMYIKCYSVECSARVFDRMRIK----DHVSWTTIIACYAQSSRYSEAIGKFRTAQKEGIKVDPMMMGSILEACSGLKSISLLKQVHSYAIRNGL-------------LDLILKNRIIDIYGECGEVCYALNIFEMLDKK------DIVTWTSMVNCFAENGLLHEAVALFGKMLNAGIQPDSVALVGILGAIAGLSSLTKGKEIHGFLIRGKFPVevVSSLVDMYSGCGSMNYALKVFDEAKCK-----DVVLWTAMINATGMHGHGKQAIYIFKRMLETGVSPDHVSFLALLYACSHSKLVDEGKFYLDMMVSKYKLQPWQEHYACVVDLLGRSG----QTEEAYKFIKSMPLEPKSVVWCALLGACRIHKNHELAMIATDKLL--------------    
 401 UniRef50_Q7XUP0        Os04g0514500 protein n=6 Ta...   393    1e-107  1   9.7%   6:592  470:1023     -----MYSKYGAIGDAKALFSLIPYKDSISWNALTVGLAQNLEEEEAVCMLKRMRLHGIT--PDDVSFSTAINACSNIRATETGKQIHCLAIKYGICSNHAVGSSLIDLYSKHGDVESSRKIFAQVDASSIVpnALIAGFVQNndEAIQLFQQVLKDGLKPSSVTFSSILSGCS-GSLNSakQVHCYTLKSGVLYDDTLLgsLAGIYLKSKMLEDANKLLTEMPDHK---NLFEWTAIISGYAQNGYGDHSLVSFWRMRHCNVRSDEATFASVLKACSDVTAFADGKEI------------HGLITKSGFGSYETATSALIDMYSKCGDVISSFEAFKELKNKQ-----DIMPWNSMIVGFAKNGYADEALLLFQKMEELQIKPDEVTFLGVLIACTHSGLISEGRHFFGSMRkyGLTpdHYACFIDLLGRGGHLQEAQEAIDQLP----FRPDGVVWATYLAACRMHKDEERGKIAARKLVE--LEPqsSTYVLLSSLHAATGNWAEAKVTRESMREKGVAKFP----------GCSWITVGNKTSLFLVQDKYHPDNLRIYEMLGDLTGMMKKD----NDIDEYGLLYSA-----------    
 402 UniRef50_Q9LRV9        Pentatricopeptide repeat-co...   393    1e-107  1   9.8%   6:596    91:674     -----MYCKCRELGFARQLFDRMPERNIISFNSLISGYTQMGFYEQAMELFLEARE--ANLKLDKFTYAGALGFCGERCDLDLGELLHGLVVVNGLSQQVFLINVLIDMYSKCGKLDQAMSLFDRceRDQVSWNSLISGYVraAEEPLNLLAKMHRDGLNLTTYALGSVLKACceGFIEKGMAIHCYTAKLGMEFDIVVRTALLDMYAKNGSLKEAIKLFSLMPSK----NVVTYNAMISGFLQMDEisEAFKLFMDMQRRGLEPSPSTFSVVLKACSAAKTLEYGRQI------------HALICKNNFQSDEFIGSALIELYALMGSTEDGMQCF------ASTSKQDIASWTSMIDCHVQNEQLESAFDLFRQLFSSHIRPEEYTVSLMMSACADFAALSSGEQIQGYAIKSGI--DAFTSviSMYAKSGNMPLANQVFIE-----VQNPDVATYSAMISSLAQHGSANEALNIFESMKTHGIKPNQQAFLGVLIACCHGGLVTQGLKYFQCMKNDYRINPNEKHFTCLVDLLGRTGRLSDAEniLSSGFQDHPVTWRALLSSCRVYKdmE---LEPEASGSY-VLLHNIYND-------    
 403 UniRef50_D1IV17        Whole genome shotgun sequen...   392    1e-107  1  10.3%   4:566    83:656     ---LSKFSKNGFIDEARALFEIMPERNVVTYNAMLSGYVQCGRLSDACRFFEEM------PERNVVSWTSLLCGLANAGRIGEARELFNVMPERNVVSW---NSMLVGLIRSGQLEEARRVFNEMPVKSQVSWNVMIAGYAEhmEEARVLFDGMGDR----NVVTWTSMISGYCrgNVQEGYCLFQKMPERNVVsnTQSCNSMINGYIRIGQLEKAQSLFDTIPVR----DKISWTSMINGYFNVGQIAKACYLFNNMPDR----DAVAWTVMVSGHVQNELFAEATYLFSEMRVKggRQFHCLLMKTQFEFDLILQNSLISMYAKCGEIGDAYSIFSKMISR------DLISWNSMIMGFSHHGLTSEALKVFEAMLTSGTHPNSVTFLGILSACSHAGLLNQGWELFDAMSdfAIQpeHYVCMVNLLGRAGKVEEAEEFISKLP----FEPDLTIWGALLGVCGF-GMINtaRRAAKRLLELD-PLNAPAHVVLCNIHASIGQRAEEGQLRKEMGLKGVRKVP----------GCSWILLKGEPYVFLSGDRIHPQADEMLSLL-------------------------------------    
 404 UniRef50_D1ILX5        Whole genome shotgun sequen...   392    1e-107  1  10.4%   6:573   263:795     -----MYSRCGSKSDAIKMFDEMTEPDVVSWTERIGA------AYDAIEAFELFRllSGNMEVNEYMLINVLSAMREPKLLKSGRQIQGLCQKAGYLLVASVNNALIFMYGKCGEMVAARHIFDEMlgDSVSWNSLIAGYAENgkQALKVFSQMRDYLLQPNKYTLASILEVAANSNFPeaMQIHSYIVKLGFIVDDSMLSCLITAYGKCNMICESKRVYSDISQI----NVLHLNAMAATLVHAGCHADALKLFQTGWRLHQEVDCITLSIVLKACGALTDLEYGRNI------------HSMALKSGMSQDNFVESAVIDVYCKCGTVDEAAKTFMNVSKN------NLVAWNAMVMGYAQHGCYHEVFELFNKMLELGIQPDEITYLGVLNSCCHAGLVNEAHTYLSSMLehGVVpeHYACMIDLFGRVGLLEDAKRTIDQMP----IMPDAQIWQILLSGCNIHGNVDLGEVAAKKLIE--LQpnDSAYVLLSNLYASAGRWNAVGKLRRVMKKKIICKEPGS----------SWIQVRGSVHYFFASDTSHPESKEIYMKLQRLYEEM------------------------------    
 405 UniRef50_A5ASR0        Putative uncharacterized pr...   392    1e-107  1   9.5%   9:523   116:627     --------RESHCGAARELFDRMREanDVTWSAVIAGYAQNSRPKDALFVFKEMreMGSPLSPMSHTITSVFSACGQLRDVILGEQIHGYVMksYVENDAFVGSVVVDMYGKCGFEELArvFYSIVDKCVVVWTALIASYVCnpCPAIEVFREMIYEGMEPNCVTLSSLISACSQidLMLGKEIHGFITRRRAapDVFISTSLIDMYGKCNYMVYGHRIFEKDKTYPRCNTTALWNATIAGYMENNFVSDAWDVFRSihGEGINPNTVTMSIVLPLCARSSQLLSGKEI------------HCYALRNGLDQEIFVSNGLIDMYSKCGKLKLAENQFNRMMN-----EKNRISWTSMIDGHGIHGDGEGAIRVFESMVREEVKPDHITFVALISACSHAGLLEQGLRYFEEmrEYGIVpeNYGTVVDLLARAGRFKEARDLIERMPIK----PGANVWGALLGACRIHGNVKEAELAAHCLQK--LEPKDSGfkLLASIYLEAGRRENAEDLRSQMREMGMEK--------------------------------------------------------------------------------    
 406 UniRef50_Q7XK57        OSJNBa0091C07.7 protein n=4...   392    1e-107  1   8.2%  13:571   194:716     ------------LSDALDLLQRMpsRDLFAWTAIISGYAR-GANEYCCKSLELFVQleDGVMPNEFTYDSVLRACVKMGALEFGRSIHGCLIQSGFESEQLITSALVDLYCRSGAVDDAVMVYNGLqpSLITSNTLIAGfmGRTEDAKLVFSQMTEH----DSGSYNLMIKAYaeGRLEDCRRMFEMMPRR----NMVTLNSMMSVLLQNGKLEEGRKLFEQIKD---ERNTVTWNSMISGYVQNDQSSEALKLFAVMCRLSIECSASTFPALLHACATIGTIEQGKMV------------HALLCKTPFESNGYVGTALVDMYSKCGCVSDARAAFS------CIMSPNVASWTSLINGLAQNGHWMEAIVQFARMLKNNVKPNEITFLGILMASARAGLVNKGMRFFHSMESYGVVpeHYTCAVDLLGRARRVREAEKFISKMP----IPADGVVWGALLTACWYTMDLEMGEKVAEKLFYMGTK-HISAYVAMSNIYAKLGKWEDVVKVRTRLRSINAKKEP----------GCSWIEVKDMVHVFLVEDRNHPEREEIYLMLEDLVS--------------------------------    
 407 UniRef50_D1H782        Whole genome shotgun sequen...   392    1e-107  1   9.2%   6:580    99:638     -----MYAKCGLLDYAHQWFDEMLERNIVSWTALVSRYAQHGWPDE---CFRVFTDMLICHRPTEFAFASVISTSG--GDGDCGRQVHALAVKTSFDSCVYvnVLIMMYCRSCGGTDEAWNVYEAMgrNLVSWNFMITGfcGCGNRALEIFSQMHFGGIRFDRATLVNIFSCLCGmgLECCFQLQCLTTKTGFISEIEVPTGLVKAYSSlgEVNDCYRIFLELDGRQ---DVVSWTGIIAVFA-ERDPEEAFLLFRQFLRECLAPDRHMFSIVLKACAGL------------ATEGHALTVQSHVLKVGFEDDIVLTNALIHTCARCGSVALSKQAFDKIGSR------DTVSWNSMLKAYAMHGQGKEALQLFSQMDA---QPDGATFVALISACSHAGMVEEGAKIFEAMsnHGIVpdHYACMVDILGRAGRIYEAKELIDKMP----MEPDSMVWSALLGGCRKHGETKFAKLAAVKLKE--LDPNNsgYILMSNIFSTNGHFNEARLIRREMERKTVRKEP----------GLSWIQVGNQVHEFASGGQQHPEKEALCARLEELVRQLKDLGYVP-----------------------    
 408 UniRef50_Q53RC3        Tetratricopeptide repeat, p...   392    1e-107  1  10.1%   9:524   160:746     --------KAGDIVSARRLFDGMPERSVVSYTTMVDALMKRGSVRDAVELYrvFHEmsCGVSPNEITLVSVIKACIGAGEFDLAMSIVGLAMKSNLLDKNlvRNSLITLYLRKGDADAARRMFDEMerDVVSWTALLDVYagNAKEAVSLYSQMLADGCRPNISCFSSVLGACASldLRSGRKIHNQTLKMACSNNVFVSSALIDMYCKCKQLPDAQMIFYSLPQK----NIVCWNSLISGYSNNSKMVEAEELFKKMPAR----NVASWNSIISGYAQNRQFIDALKSFHAMLASggKMVHAKIIKLGIKESIFVGTALSDMYAKSGDLDSSKRVFYEMPKR------NDVAWTAMIQGLAENGFAEESILLFEDMISAGITPNEQTFLAILFACSHSGLVEHAMHYFEMMQACGISpkHYTCMVDVLARAGHLAEAEDLLLKIESKSE----ANSWAALLSACNIYRNKEMGERAAKRLQELD-KDNTAGYVLLSNMYASCGKWKDAAEMRILMKGINLKKD-------------------------------------------------------------------------------    
 409 UniRef50_Q9SZT8        Pentatricopeptide repeat-co...   392    1e-107  1   9.6%  77:599    38:558     ----------------------------------------------------------------------------KSQSVDEVLQIHAAILRHNLLLHpvLNLKLHRAYASHGKIRHSLALFHqiDPDLFLFTAAINTASINgdQAFLLYVQLLSSEINPNEFTFSSLLKSCST---KSGKLIHTHVLKFGLGIDPYVATGLVDVYAKGGDVVSAQKVFDRMPER----SLVSSTAMITCYAKQGNVEAARALFDSMCER----DIVSWNVMIDGYAQHGFPNDALMLFQKLLAEggRWIHVFVKSSRIRLNVKVCTGLIDMYSKCGSLEEAVLVFNDTPRK------DIVAWNAMIAGYAMHGYSQDALRLFNEMQgtGLQPTDITFIGTLQACAHAGLVNEGIRIFESmqEYGIKpeHYGCLVSLLGRAGQLKRAYETIKNM----NMDADSVLWSSVLGSCKLHGDFVLGKEIAEYLIGLNIK-NSGIYVLLSNIYASVGDYEGVAKVRNLMKEKGIVKEP----------GISTIEIENKVHEFRAGDREHSKSKEIYTMLRKISERIKSHGYVPNT----NTVLQDLEETEKE----    
 410 UniRef50_Q6ZG82        Os02g0132100 protein n=3 Ta...   392    1e-107  1  12.4%  30:595   117:671     -----------------------------VYNRLLLAALQEARFDLVESLYK--DLLLSGAAPDVFTRNILLQALCAAGRMELARRVFDAMPERNEFS---FGILARGYCRAGRSMDALGVLDSMPTMNLVVCNTVVAGFCREgdEAERLVERMRDEGLAPNVVTFNARISALCkgRVLDAYRIFNDMQerGLppDQVTFDVMLSGFCEAGMVDEARVLVDIMRCGGFLRRVESYNRWLSGLVRNGMVGEAQELLREMAHEGVQPNSYTYNIIVDGLCKEGKAFDVRKV------------EDFVKSGVMTPDVVTYTSLLHAYCSEGNAAAANRILDEMAQKGC--APNSFTYNVLLQSLWKAGRITEAERLLERMSEKGYSLDTASCNIIIDGLCRNNKLDMAMDIVDGMWNEgcLPdtYSTLVSALCKEGRFDEAKKKLLEMIGKDI-SPDSVLYDTFIHGYCKHGKTSLAVKVLRDMEKKGCKPSTRTYNLLIWGFREKHNSDEILKLISEMKGKGICPNVMT-YNSLIKSFCERGMVN---------------------KAMPLLDEMLQNEIVPN-ITSFELLIKAFCK--------    
 411 UniRef50_Q9SCT2        Pentatricopeptide repeat-co...   392    1e-107  1  11.2%   5:602   140:724     ----SMYVRCGSLEQARKVFDKMPHRNVVSYNALYSAYSRN--PDFASYAFPLTTHMAFevKPNSSTFTSLVQVCAVLEDVLMGSSLNSQIIKLGYSDNVVVQTSVLGMYSSCGDLESARRIFDCvnRDAVAWNTMIVGSLKniEDGLMFFRNMLMSGVDPTQFTYSIVLNGCSkgSYSLGKLIHARIIVSDSLADLPLDNALLDMYCSCGDMREAFYVFGRIHN----PNLVSWNSIISGCSENGFGEQAMLMYRRLLRMStrPDEYTFSAAISATAEPERFVHGKLL------------HGQVTKLGYERSVFVGTTLLSMYFKNREAESAQKVFDVMKER------DVVLWTEMIVGHSRLGNSELAVQFFIEMYREKNRSDGFSLSSVIGACSDMAMLRQGEVFHCLAIRTGFdmSVCGALVDMYGKNGKYETAETIF-----SLASNPDLKCWNSMLGAYSQHGMVEKALSFFEQILENGFMPDAVTYLSLLAACSHRGSTLQGKFLWNQMKEQGIKAGFK-HYSCMVNLVSKAGLVDEALEL---IEQSPPGNNQAELWRTLLSACVNTRNLqpEDTAT-HILLSNLYAVNGRWED-    
 412 UniRef50_Q0WNP3        Pentatricopeptide repeat-co...   391    1e-107  1  10.7%   9:511   129:607     --------RLGDLVYARKVFDSMPEKNTVTWTAMIDGYLKYGLEDEAFALFEDYVKHGIRFT-NERMFVCLLNLCSRRAEFELGRQVHGNMVKVGVGNLIVESSLVYFYAQCGELTSALRAFDMMEEKDVISWTAVISACSRKGhkAIGMFIGMLNHWFLPNEFTVCSILKACSEEklRFGRQVHSLVVKRMIKTDVFVGTSLMDMYAKCGEISDCRKVFDGMSNR----NTVTWTSIIAAHAREGFGEEAISLFRIMKRRHLIANNLTVVSILRACGSVGALLLGKEL------------HAQIIKNSIEKNVYIGSTLVWLYCKCGESRDAFNVLQQLPSR------DVVSWTAMISGCSSLGHESEALDFLKEMIQEGVEPNPFTYSSALKACANSESLLIGRSIHSIAKKNHALSNVfsALIHMYAKCGFVSEAFRVFDSMPEK-----NLVSWKAMIMGYARNGFCREALKLMYRMEAEGFEVDDYIFATILSTCGDI-ELDEAVE--------------------------------------------------------------------------------------------    
 413 UniRef50_Q9SKQ4        Pentatricopeptide repeat-co...   391    1e-107  1  10.7%  62:576    79:593     -------------------------------------------------------------RPNTLLSNHLIGMYMKCGKPIDACKVFDQMHLRNLYSW---NNMVSGYVKSGMLVRARVVFDSMPERDVVSWNTMVIGYAQDghEALWFYKEFRRSGIKFNEFSFAGLLTACVKSrlQLNRQAHGQVLVAGFLSNVVLSCSIIDAYAKCGQMESAKRCFDEMTVK----DIHIWTTLISGYAKLGDMEAAEKLFCEMPEK----NPVSWTALIAGYVRQGSGNRALDLFRKMIarHGKEIHGYMIRTNVRPNAIVISSLIDMYSKSGSLEASERVF-----RICDDKHDCVFWNTMISALAQHGLGHKALRMLDDMIKFRVQPNRTTLVVILNACSHSGLVEEGLRWFESmvQHGIVPdhYACLIDLLGRAGCFKELMRKIEEMP----FEPDKHIWNAILGVCRIHGNEELGKKAADELIKLDPE-SSAPYILLSSIYADHGKWELVEKLRGVMKKRRVNKEKAV----------SWIEIEKKVEAFTVSDGshARKEEIYFILHNLAAVIEEE---------------------------    
 414 UniRef50_B9I4A0        Predicted protein n=1 Tax=P...   391    1e-107  1  10.8%  28:529    19:500     ---------------------------IVIWNAMIRGYAFNGPFQECIRMFDEMPQRGL--KPHNFTYPYVINSCCELEWYGRGKRVHCEIVKSGfsSYAVANSLFNMYLksNCKLDYARKIFDDmvRPVELWNQMiyVNIGDVKSARELFDIMPER----DIVSWNSMILGYAkgKVANARGLFEKMPEK----NVISWTSMIGAYADTDDLETARSFFETMPQR----NVVSWNSMISSYAKHGKFVESLNLFVQMQSEGVTPDGYTFVSVLSACSNLGDLEFGKYIH-------------YLSGDLSQSEVMVGTALTEMYAQCGDVDKAFAVFIKIGKR------DVFCWNVIIKALALHGRSEEAIKIFLLMRKTGLKPNDFTFTSALFACSHGGLVEEGHIIFNSMEkyKIIptHYGCLIDMLCRNGQLEEAMLLVEDMP----FQPDVAIWGALLGGCRVTGDLKLAEKVVeeMETNE----SGVYVLLSNIHASAGQWIEAADARKKMDEKKISKKTGSSV--------------------------------------------------------------------------    
 415 UniRef50_D1IEQ3        Whole genome shotgun sequen...   391    1e-107  1  11.3%  86:546    45:475     -------------------------------------------------------------------------------------SIHVHALKLGFESYLFVSNALIHMYAMCGQLGFAQKMFDGmdRDLVSWNTLICGYSqkYKEVLRLFDAMTAANIKADAVTMVKIILACShgDWEFADSMVKYIKENNLEIDVYLGNTLIDMYGRLGNLTAARKLFDNMPKR----DVISWTSMITGYSQASQFSDAVKLFQEMMAAKVKPDKVTVASVLSACAHLGKLDVGWAVHHYIRR------------HGVQADIYVGNSLIDMYCKCGMVEKALEVFHRMKDK------DSVSWTSVISGLAVNGFANSALDLFSQMLREGVQPTHGTFVGILLACAHAGLVNKGLEYFESMEshGLVPahYGCVVDLLSRSGNIDKAYEFIKKMP----IVPDVVVWRILLSACKLHGNVVLAEIATKRLLELDPC-DSGNYVLLSNTYAGVDRWDDAMKMRELMEDSDVQKP----------SGSSSIEVDGMV---------------------------------------------------------    
 416 UniRef50_A9SQ58        Predicted protein n=2 Tax=P...   391    1e-107  1  10.7%  35:539    24:498     ----------------------------------IRDLCLNGQPKNSLHILE-----QRGIEPSSYAYVCLLRRCTHMKDLAEGKYVHAHMAKSDFVPTTFVLNALVNMYMKCGSLVDARQVFDRmeRDMFTYTMMlyAKLGYPEDAYKMYEQMQKERVPVDRITFTTILNVCSTLrlEKGMKVHQDMVRGGIRPDIILGNTLIDMYAKCGNLKQAHRVFKEMDNR----DIVTWNIMVGGAARNGYFDEAFEFFKAMLDEGQKPDKVTYISILNACTSL---------------EQGTLLHSVIMKAGFELDVRVGTALVNMFSKCGSVVDALKVFQKLPQR------NVVSWTSVISAYAQAGEPERALECYAKMLNEGMVADKRAYTTILNVCAMLGDIEKGKAVHGHIVQSGIATDIinGLIDMYVKCGRLKDAYRLFQDMNVR-----DVVSWTTLIEGWVQHRQYQEALDTFNDMQLEGVMPNTVTFLGVLKACAGMGSLVDGKRIHARIVEAGLAEN-AHIRHALADMYAKC----------------------------------------------------------------    
 417 UniRef50_B9HGU1        Predicted protein n=3 Tax=r...   391    1e-107  1  10.4%   7:525    42:539     ------YSKLGDTVSARKVFDRMPERTVVSWSAQISGYSQNGCYQDALLVF--LDMLRAGFKANQFTYGSVLRACTGLRCLQRGMQIQGCLEKSRFASNLIVQSALLDLHSKCGKMEDASYLFGMmeRDVVSWNAIIGAYAVQggDSFRMFRSMMQEGMSPDLFTFGSVLKASgaSDVIRVCQIHQLIIQLGYGSHISLSGSLIDAYAKCESLASAHCLYKSMPMK----DMISCTALMTGYARDSNysKALDLFKEIQQMHMKIDDVILCSMFNICANISSLSVGRQI------------HALALKCKPTYDVAMGNALIDMYAKSGEIKDANRAFNEMEEK------NVISWTSLITGYGSHGHGHEAIALFKKMEYEGLKPNDITFLSLLFACNHCGLTGEGWECFNNMikYNILpeHFSCMVDLFARGGLLEEAYNLINKM----NIKPNASLWGAILGACYIYGNMPLAEEAAIHLFKMDPE-NSVNYVVLADIYAAAGSWDNAWKMRKLMEERNLKKAP------------------------------------------------------------------------------    
 418 UniRef50_Q9CA56        Pentatricopeptide repeat-co...   391    1e-107  1  11.1%   6:601   294:873     -----LYAKCGHMAEAMEVFSRIPNPSVVSWTVMLSGYTKS---NDAFSALEIFKEMRHSgeINNCTVTSVISACGRPSMVCEASQVHAWVFKSGFYLDSSVAAALISMYSKSGDIDLSEQVFEDlrQNIV--NVMITSFSqkPGKAIRLFTRMLQEGLRTDEFSVCSLLSVLDCLNL--GKQVHGYTLKSGLVLDLTVGSSLFTLYSKCGSLEESYKLFQGIPFK----DNACWASMISGFNEYGYLREAIGLFSEMLDDGTSPDESTLAAVLTVCSSHPSLPRGKEI------------HGYTLRAGIDKGMDLGSALVNMYSKCGSLKLARQVYDRLPE------LDPVSCSSLISGYSQHGLIQDGFLLFRDMVMSGFTMDSFAISSILKAAALSDESSLGAQVHAYITKIGlePSVGSSLLTMYSKFGSIDDCCKAFSQ-----INGPDLIAWTALIASYAQHGKANEALQVYNLMKEKGFKPDKVTFVGVLSACSHGGLVEESYFHLNSMVKDYGIEPENRHYVCMVDALGRSGRLREAE----SFINNMHIKPDALVwaCKIHGE-VELglEPSDAGAYISLSNILAEVGEWDE--    
 419 UniRef50_Q9FLX6        Pentatricopeptide repeat-co...   391    1e-107  1  11.3%   6:600   168:748     -----LYSKCGQFKEACELFSSLQNADTISWTMMISSLVGARKWREALQFYSEMVKAG--VPPNEFTFVKLLGASSFLG-LEFGKTIHSNIIVRGIPLNVVLKTSLVDFYSQFSKMEDAVRVLNSseQDVFLWTSVVSGFVRnaKEAVGTFLEMRSLGLQPNNFTYSAILSLCSAVrlDFGKQIHSQTIKVGFEDSTDVGNALVDMYMKCSASeeASRVFGAM----VSPNVVSWTTLILGLVDHGFVQDCFGLLMEMVKREVEPNVVTLSGVLRACSKLRHVRRVLEIHAYLLRRHVDGEM------------VVGNSLVDAYASSRKVDYAWNVIRSMKRR------DNITYTSLVTRFNELGKHEMALSVINYMYGDGIRMDQLSLPGFISASANLGALETGKHLHCYSVKSGFsaSVLNSLVDMYSKCGSLEDAKKVFEE-----IATPDVVSWNGLVSGLASNGFISSALSAFEEMRMKETEPDSVTFLILLSACSNGRLTDLGLEYFQVMKKIYNIEPQVEHYVHLVGILGRAGRLEEATGVvnAMIFKTLLRACRYRGNLSLGEDMANKglAPSDPALYILLADLYDESGKPE---    
 420 UniRef50_Q5SMW8        Os06g0185700 protein n=2 Ta...   390    1e-107  1   9.6%   7:601    65:646     ------YAALGDLASARGVLDGTPRPDAYAYRVMLGWLVDAGSHADAVALHRDMRRRCPAAAQADVVLSLALKACVRSADFRYGRRLHCDVVKAGGADGFVMNSLVDMYAKAGDLENARKVFDRVPERNVVSWTSMLSGSIQNgeEGLVLFNEMRQDNVHPSEYTMVSVLAACalGGLHQGRWIHGSVIKYGLSTNSFISASLLDMYAKCEKVEDARRVFDELE----FVDIVLWTAMIVGYTQNKRPLDALQLFLHKKFVSIVPNSVTIATVISASAQLRHLPLGRSV------------HAIGVKLGTMESDVVRNALVDMYAKCQALPEANSIF------GRILIKDVVAWNSMMAGYSENGMANESLVLFNRMRMQGISPDAISVVNALSACVCLADLHIGKGFHTYAIKYAFMSNIytALLNLYSKCADLPSAQRVFNDMTDR-----NSVTWSAMIGGYGMQGDSAGSIDLFNEMLKENIHPNEVVFTSILSACSHTGMVTAGKEYFDSMARHFNITPSMKHYACMVDVMARAGNLEEALEF----IQNMPIKAGISVWGSFLHGCKLHSRLepD----FYVLMSNLYTsgRWDK--    
 421 UniRef50_Q9SR00        Pentatricopeptide repeat-co...   390    1e-107  1  13.8%  32:518    90:571     -------------------------------TQMLKIFHRSCRSGNYIESLHLLETmrKGYNPDVILCTKLIKGFFTLRNIPKAVRVMEILEKFGQPDVFAYNALINGFCKMNRIDDATRVLDRmfSPDTVTYNIMIGSLCSRgdLALKVLNQLLSDNCQPTVITYTILIEamLEGGVDEALKLMDEMLSRGLKPDMFTYNTIIRGMCKEGMVDRAFEMVRNLELKGCEPDVISYNILLRALLNQGKWEEGEKLMTKMFSEKCDPNVVTYSILITTLCRDGKIEEAMNLLKLMKEK------------GLTPDAYSYDPLIAAFCREGRLDVAIEFLETMISDGC--LPDIVNYNTVLATLCKNGKADQALEIFGKLGEVGCSPNSSSYNTMFSALWSSGDKIRALHMILEMMSNGIDPdtYNSMISCLCREGMVDEAFELLVDM-RSCEFHPSVVTYNIVLLGFCKAHRIEDAINVLESMVGNGCRPNETTYTVLIEGIGFAGYRAEAMELANDLVR-------------------------------------------------------------------------------------    
 422 UniRef50_D1H9S6        Whole genome shotgun sequen...   390    1e-107  1  10.6%  27:574    53:601     --------------------------NVFVWNTMIRKQVENGLITRDPPLLYYMQGLRAGLRPNGHTFMYLLKALVSGQEVKEGEEVHASVVRTGFACSEFVSGALLGFYVACGLVGKGRQVFDEMRQPgvLWTLIIRAYVcfPEKALELFRTMREVGLTPDMVAISTVVSACglGDLGVAKAMHCFIEKSGIEVDAFVSSTLISTYGECGSLDYAYRFFQETPMKNIvrDVVSWNSMIGGFARIGQYQEALTWFHEMEFSGVSPNALTLLSTLSACASHGALDTGAWI------------HAYVDKNDMNRDGSLDSSLIDMYSKCGDIDKAVQIFEE------STRRDLFTWTSIVCGLAMHGRGEKALHYFSKMKEAQVQPDDVTMVGVLSACAHAGLLDQGWWYFQSMEkfGLVpeHYGCMVDLLGRMGCLKEAYDLIMGMP----MEANEIIWGAFLSACRVHNNVELGEVAARRLL--GLDpdPWARVMLSNMYAEEAKWDRSMGLR----------KEIKKKGLKKSPGCSSIEVNGSVHGFLVGDNSHPCYTEINSMVEKIEKLMR-----------------------------    
 423 UniRef50_Q9FM64        Pentatricopeptide repeat-co...   390    1e-107  1  10.6%   6:573   217:825     -----MYGKCGVLDDASKVFDEIPDRNAVAWNALMVGYVQNGKNEEAIRLFSDMRKQG--VEPTRVTVSTCLSASANMGGVEEGKQSHAIAIVNGMELDNILGTSLLNFYCKVGLIEYAEMVFDRmeKDVVTWNLIISGYVQQgeDAIYMCQLMRLEKLKYDCVTLATLMSAAAreNLKLGKEVQCYCIRHSFESDIVLASTVMDMYAKCGSIVDAKKVFDsmQLEGVPPNVITWNLIILSLLRNGQVDEAKDMFLQMQSSGIIPNLISWTTMMNGMVQNGCSEEAILFLRKMQESggRTIHGYIIRNLQHSSLVSitSLVDMYAKCGDINKAEKVF------GSKLYSELPLSNAMISAYALYGNLKEAIALYRSLEGVGLKPDNITITNVLSACNHAGDINQAIEIFTDIvkRSMKpeHYGLMVDLLASAGETEKALRLIEEMP----FKPDARMIQSLVASCNKQRKTELVDYLSRKLLESEPE-NSGNYVTISNAYAVEGSWDEVVKMREMMKAKGLKKKP----------GCSWIQieEGVHVFVANDKTHTRINEIQMMLALLLYDM------------------------------    
 424 UniRef50_D1IZD3        Whole genome shotgun sequen...   390    1e-107  1  11.7%  77:582    20:502     ----------------------------------------------------------------------------HCTSISKTKQAHALLLRTHLLHNPLFSSKLISFLALsgDLNYARKLFTQMqpDPFICNTMIRGYArnPYEAVSLYYFMVERGVPVDNYTYPFVLAACARLgvKLGRRFHCEVLKNGFGSDLFVINALIQFYHNCGSFGCACDVFDE----STVRDVVTWNIMINAHLNKGLSEKAFDLLDEmkLDNLRPDEVTMVSLVPACAQLGNLERGKFLHSY---------SKELDKFEINCDLVLETALVDMYAKCGSIDLALQVFRRMRVR------NVFTWNALIGGLAMHGHGEDAISLFDQMEHDKLMPDDVTFIALLCACSHAGLVDEGLAMFQAMkkFQIEpeHYGCVVDLLCRARKVDDALAFIENMPIK----ANSVLWATLLGACRSGGHFDLAEKIGRRVIE--LEPDsgRYVMLSNLYAGVSQWDHALKLRKQMKNKGIEKTP----------GCSWIELNGMIHQFVAGDRSHLQTEQIYAMIEEMTRRvlDGGHVPGT---------------------    
 425 UniRef50_Q8LK93        Pentatricopeptide repeat-co...   390    1e-107  1   9.3%  76:582    37:515     ---------------------------------------------------------------------------SKCNSLRELMQIQAYAIKSHIEDVSFVAKLINFCTESpsSMSYARHLFEAmePDIVIFNSMARGYSrnPLEVFSLFVEILEDGILPDNYTFPSLLKACAVAklEEGRQLHCLSMKLGLDDNVYVCPTLINMYTECEDVDSARCVFDRI----VEPCVVCYNAMITGYARRNRPNEALSLFREMQGKYLKPNEITLLSVLSSCALLGSLDLGKWIHKYAKKHSFCKY------------VKVNTALIDMFAKCGSLDDAVSIFEKMRYK------DTQAWSAMIVAYANHGKAEKSMLMFERMRSENVQPDEITFLGLLNACSHTGRVEEGRKYFSQMVSkgIVPshYGSMVDLLSRAGNLEDAYEFIDKLP----ISPTPMLWRILLAACSSHNNLDLAEKVSERIFELD---DSHgyVILSNLYARNKKWEYVDSLRKVMKDRKAVKVP----------GCSSIEVNNVVHEFFSGDGVKSATTKLHRALDEMVKELKLSGYVPDT---------------------    
 426 UniRef50_B9HKM1        Predicted protein n=1 Tax=P...   390    1e-107  1  10.5%  19:570   293:844     ------------------AFERIKKPARREFGLMVGYYARRGDMHRARQTFESMRV--RGIYPSSHVYTSLIHAYAVGRDMEEALSCVRKMKEEGVEMSLVTYSIVVGGFAksCNMDRAESLVREMEEEGiiYHTMMNGyiGNEEKCLIVFKRLKECGFAPSVISYGCLINMYTkgKVSKALEVSKMMESAGIKHNMKTYSMLINGFLKLKDWTNAFTVFEDVIKDGLKPDVVLYNNIIKAFCGMGNMDRAIHMVKEMQKKRHRPTSRTFMPIIHGFARAGEMRRALEIFDMMRRS------------GCIPTVHTFNALVLGLVEKRQMEKAVEILDEMALAG--VSPDEHTYTTIMNGYAALGDTGKAFEYFTKLRNEGLELDVFTYEALLKACCKSGRMQSALAVTREMSAQNIPRNtyNILIDGWARRGDVWEAADLMQQMKQEG-VQPDIHTYTSFINACCKAGDMLRATKTIQEMEALGVKPNVKTYTTLIHGWACASLPEKALRCFEEMKLAGLKPDKAVYHCLMTSLLSRATVAEAYIYSGILSirEMIESELTVDMGTAVY---------------------------------    
 427 UniRef50_UPI000198324B PREDICTED: hypothetical pro...   390    1e-107  1   8.6%   7:577   264:802     ------YGKCGNMEWGRKVLNSMEWKDSFVWSSIISGYARNGSGDEAVCLFRDM--CRQGMKPDQHALSSTLKACAEIEDLETGVQVHSQMIKNGYHSDCFVASALITLYSDCNEICEAHKLFRRIdkDIVAWNSMILGYAQMEEecIKIFRELCQTplKPDGATIIAILKScnQAGLITGIQIHTMIMKSSLSCETLICNAVINMYSKCGDVDDAYKAFIDIVHR----DAISWSSIIGSYQQNGFELEALSLCKEMLADGVYVTSFSLPLCIAACSELAAIDVGKQF------------HSSVIKLGFDGDVYVGSSIIDMYAKCGFMEESQKAFDE------QEDPNKATFNALISGFAHHGKALEAIELFKEMEKMSLVPNQISFLAALSACSHAGLVEESLFFFDLmkKYNIKPehYSCLVDVLGRAGRLEEAYQIIQ---NDGKEQ----AWRTLLSACRNYGNAEIAEKSARKLME--LDpdHASYVLLSNLYSGEGKWEQAFKLRQRMVEIGVKKDPGS----------SWLIFREQVHQFSVGDFSHHDMREILGELNILNKHINSAG--------------------------    
 428 UniRef50_D1J1Y6        Whole genome shotgun sequen...   390    1e-106  1  10.6%  65:582    26:553     ----------------------------------------------------------------TITLSILETHLHNCHNLKQFNRILSQMILTGFISDTFAASRLLKFSTDSPFidYSLQIFDRIesNGFMWNTMMRAYIQsaEKALLLYKLMVKNNVGPDNYTYPLVVQACAVRLLEFgkEIHDHVLKVGFDSDVYVQNTLINMYAVCGNMRDARKLFDESP----VLDSVSWNSILAGYVKKGdvMEAWKLFNEMDEK----DMVSWSALISGYEQNGMYEEALVMFIEmiVKTGKMIHGLVIRMGIESYVNLQNALIHMYskCGCVENALEVFNGMEEKG------VSSWNALIIGLAVNGLVERSLDMFSEMKNNGVIPNEITFMGVLGACRHMGLVDEGRCHFASmeKHGIEPnhYGCMVDLLGRAGLLNEAEKLIESMP----MAPDVATWGALLGACKKHGDTEMGERVGRKLIE--LQPDHDGfvLLSNIFASKGDWEDVLEVRGMMKQQGVVKTP----------GCSLIEANGVVHEFLAGDKTHPWINKVEGMLNEMAKRLKMEGYAPDT---------------------    
 429 UniRef50_B9HVV2        Predicted protein (Fragment...   390    1e-106  1  10.1%  70:575     3:478     ---------------------------------------------------------------------SLLQFCAKNREPITGKACHGQVIRLGLETDTLTSNMLMNMYSKCGLIRRACKVFDEieRSMVSWNIMIGTCVQNGEeaIDIFLEMQREGSPCSEFTVSSVVCAcaKGDVFFCRQLHAFAIKAVVDADVFVGTALIDVYAKCGSIEDASCVFEGMPER----NDVTWSSIVAGFVQNELYEEGLVLFARGKEMGLENNQFMISSVIRACAGLAALIEGRQV------------HAIVCRTGFGPNNFVASALVDMYAKCGSIEEAYVLFCNIEKK------NVVLWNAIISGFAKHACSSEVMNLFEEMQQTGMQPDEVTYVAVLSACVHVGLVDKGRSYFDLmrEHNVSPnhYSCMVDILGRGGLVHEAHELMTKMP----FAATASMWGSLLASCRIHGNLELAEIAAKNLFE--MEPDrgNYVLLANTYAASKKWEEAAKARKSLKGSEI----------LKERGKSWIEIKDKVHTFMAGERTHPRITDIYLELNNLLEEMTK----------------------------    
 430 UniRef50_D1IPJ5        Whole genome shotgun sequen...   390    1e-106  1  10.4%  68:579    11:555     -------------------------------------------------------------------YLHLLEACIQSKSLTEAKKIHQHFLKNTSNADSSVLHKLTRLYLSCNQVVLARRLFDEIppSVILWNQIIRAYAWNgdGAIDLYHSMLHLGVRPNKYTYPFVLKACslLAIEDGVEIHSHAKMFGLESDVFVCTALVDFYAKCGILVEAQRLFSSMSHR----DVVAWNAMIAGCSLYGLCDDAVQLIMQMQEEGICPneVSWSAMIGGYVASDCMKEALDIFRMMQLSggFCSHGYLIVRGFATDTLICNALIDMYSKCGKISFAREVFNRMDRH------DIVSWNAMIIGYGIHGLGMEALGLFHDLLALGLKPDDITFICLLSSCSHSGLVMEGRLWFDAmrDFSIVpeHCICMVDILGRAGLIDEAHHFIRNMP----FEPDVRIWSALLSACRIHKNIELGEEVSKKIQSLGPE-STGNFVLLSNIYSAAGRWDDAAHIRITQKDWGLKKIP----------GCSWIEINGIVHAFVGGDQSHLQLSQINRKLEELLVEMKRLGYQ------------------------    
 431 UniRef50_D1IJI0        Whole genome shotgun sequen...   389    1e-106  1   9.4%   4:537    98:614     ---NKLLSLYLKFGhaHQLFDTMPRRDVVSFNTMISASVRNNY--DALDLVGLYSKMKkdVKPNHITFAGLIGACDGLIALRLRGIFHAHTVRCGLSSNEFvsSLVDGYAKQMKLEDAIKAFNEIMELDLVSWNIMIDGCARNneHAVRMFSQMLKGNVRVDGFTLTSIIKTCSkgDLKHGMQFHGSAIKLGLAHETPIYNALITMYSKCEkvASPVKIFGSISE----PNIISWTAMISGFMQNEQNEEAIGLFKEMLRLGVRENDFSFSSILPVYGNLANLEQGKQI------------HARIIKSWFGLDLSVNNALIDMYSKCGSLEDAHLVFMKMGKH------DVVSCTTMIMSYGQHGKGKEALEILAEMKSEGLVPDGVTFLGCLYACSHGGLVEEGVRVFKIMIehNLKpeHFACVVDMLGRAGRLNEAENFIDEM----GIESDVLVWETLLGACRVHGEMVLGEKSAQKIME--LQprHGPYVLLANIYAERGSWEDKVMVREKLVSHGLKKQVGCSWVALETMSC------------------------------------------------------------------    
 432 UniRef50_C5WT77        Putative uncharacterized pr...   389    1e-106  1   9.0%  11:598   275:919     ----------AALCVLEDMERDSIPADIYTYNIMIDKLCRIKRSARAFLLLKRMR--KDDLIPDECTYNTLINGFFGEGKINHAHYVFNHMLRQTFVPSVATYTTMIDGYCRNRRIDKALSVLSEMqeDGIDPDIVTYSALIniTEAEHFRQYMSRMKISFDSVSFNCIIDSYcrGNIVEAFTVYDAMVRYGHSPNVCTynALLLGICKYGTLDEALDLCEKMVKNNCLPDIHTYTILLSGFCRKGKILPALIMLQMMLDKGVVPDTVAYTCLLNGLINEGQVKAASYVFQEIICKE-----------GLYADCIAYNSLMNGYLKGGNINTIKRMMSDMYQ--SEVYPNSASYNILMHGYVKGGQFSKSLYLYKYMVRKGIRPDNVTYRLLILGLSECGLIDIAVKFLEKMVLEGIFPdvFDILITAFSEKSKMHNALQLFNCMKWL-RMSPSSKTYSAMINGLIRKNYLDQSHEVLREMLQVGLQPNHTHYIALVNAKCRVGKIDRAFRLKEEMKALGIVPADVAE-SSIIRGLCKCGKLEEAVIVFSTMMRSglCKESKIADALHLKRLMESCRLKVDVV-SYNVLITGLCKDKR-----    
 433 UniRef50_Q9FLL3        Pentatricopeptide repeat-co...   389    1e-106  1  13.7%  40:522    49:527     ---------------------------------------HSLQFNEALDLFTHMVE--SRPLPSIIDFTKLLNVIAKMKKFDVVINLCDHLQIMGVSHDLYTCNLLMNCFCQSSQPYLASSFLGKMMKLGFEPDIVTFTSLINgeEAMSMVNQMVEMGIKPDVVMYTTIIDSLCKNgvNYALSLFDQMENYGIRPDVVMYTSLVNGLCNSGRWRDADSLLRGMTKRKIKPDVITFNALIDAFVKEGKFLDAEELYNEMIRMSIAPNIFTYTSLINGFCMEGCVDEARQMFYLMETK------------GCFPDVVAYTSLINGFCKCKKVDDAMKIFYEMSQKG--LTGNTITYTTLIQGFGQVGKPNVAQEVFSHMVSRGVPPNIRTYNVLLHCLCYNGKVKKALMIFEDMQKrgVAPntYNVLLHGLCYNGKLEKALMVFEDMRKREMDIGIITY-TIIIQGMCKAGKVKNAVNLFCSLPSKGVKPNVVTYTTMISGLFREGLKHEAHVLFRKMKEDGVS---------------------------------------------------------------------------------    
 434 UniRef50_D1I850        Whole genome shotgun sequen...   389    1e-106  1  12.1%  27:586     2:605     --------------------------NQISWthLLKTCIAERDLSTGKSLHSLY--IKSFIPPSTYFSNHFILLYSKCGRLAWARKAFQDISDPNVFS---FNAIIAAYAKESRPLIAHQLFDQIPEPDLVSYNTLISAYADCGETalGLFSGMREMGLDMDGFTLSAVITACCD-DVGLIGQLHSVAVSSGFDSYVSVNNALLTYYGKNGDLDDAKRVFYGM---GGIRDEVSWNSMIVAYGQHQEGSKALGLFQEMVRRGLNVDMFTLASVLTAFTCLEDLSGGLQFHGQLIKTGFHQNsrKVFEEITEPDLVLWNTMVSGYSQNEelEDALECFRQMQGIGYRpeHNTVSLNSMIAGYAQHGIEMESLHLFQWMLERQIAPTSITFISVLSACAHTGRVEEGWNYFNMMKEknIEPehYSCMIDLLGRAGKLSEAENLIARMP----FNPGSIGWASLLGACRTHGNIELAVKAANQVLQ--LEpnAAPYVVLSNMYASAGRWEEVATVRKFMRDRGVKKKP----------GCSWIEVKKRIHVFVAEDSSHPMIKEIYEFLEEMSGKMKRAGYVPDGEFFF-----------------    
 435 UniRef50_C0PGG8        Putative uncharacterized pr...   389    1e-106  1  11.8%   2:596    60:626     -AAVRRLAAAGDVDGVQLALQEMRLRGVptEGALVAafARAGAPDRALKTfrAVHDLGCARPTEPRLYNHLIDALLRENMVGAVVLVYDNMRKDGVHPNVFTYNLLVRALCQNNRVGAARKMLDEMARKGCPPDDVTYakLDRLDEATEVLAAM-----PPVAASYNAIVLALCRerMQEVFSVVSDMVGRGLQPNVITYTTIVDAFCKAGELRMACAILARMVITGCTPNVVTFTALVRGLFDDGRVHDALDMWRWMVAEGWAPSTVSYNILIRGLCSVGDLKGASSILNSMEQ------------HGCFPNVRTYSTLIDGFSKAGDLGGAISIWNDMSRSGC--KPNVVVYTNMVDVFCKKLMFNQAKSLIDKMLLENCPPNTVTFNTLIRSLCDCRRVGRALGVFHEMRRHGCVPntYNELIHGLFREGNCGDALHMVTEMQSHGI-ELSLVTYNTVVSGLCQTRMSREAMVFVGKMIVQGIQPNAFTFSAIIHAYCKEGEVRMAAWMLGAM----------NVVNCHRNILVYTILMAELCNQDKLVDAMTYLLKMLYE------------GIYPNTV-TWNVLVRGVFRN-------    
 436 UniRef50_UPI00019851B0 PREDICTED: hypothetical pro...   388    1e-106  1  11.5%  60:579     2:490     -----------------------------------------------------------KPMLDQGSLVAAIKRCTRVGDL---KAIQAHMVRANLTQDTFLTSKLisAVTLSGHVAYAHRIFstHHPNLFMWNTIIRGYSIspITAIALYRDMFLCGISPNSYTFGFVLKACCKlrLCEGQELHSQIVKVGLDFETPLVNGLIKLYAACGCMDYACVMFDEMPE----PDSASWSTMVSGYAQNGQAVEALKLFREMQAENVSSDAFTLASVVGVCGDLGALDLGKWVHSYM------------DKEGVKIDVVLGTALVGMYSKCGSLDNALKVFQGMAER------DVTAWSTMIAGYAIHGHGEKALQLFDAMKRSKTIPNCVTFTSVLSACSHSGLVEKGHQIFETmtEYKITPqhYGCMVDLFCRAGMVGHAHKFIQTMP----IEPNVVLWRTLLGACKTHGYKDLGEHISRKILK--LDPSseNYVLVSNVYASLGRWSSVCQVRSLMKDK---APKKQH-------GWSSIEINFMVHKFIMGDESHPEREKIYGMLHQMARKLKQVGHV------------------------    
 437 UniRef50_B9HSV8        Predicted protein n=2 Tax=P...   388    1e-106  1  10.8%   3:574    65:617     --ATKLISAYAIFglSRLVFDSLHFKSVYLWNSLINGFVKNRAYNEAFGWFYQM--CCRGVLLDDYTLATMSKVCHEIGDLNAGKLIHGKSLKTGFVLDVIVANSLMSMYSKCGGFGECLKLFDEmeRNVGSWNVILSGYADSGdeVSGFVKDMQIEGLKPDAFTVSSLLTLCnmGKRDHGRELHGFIVrvGSGTEVHLGCCLIDMYSRSNRVDVGRRVFDRMECR----NVYAWTAMINGHVQNGALEEGLVLFHEMqrDGVEPNKVSLVSVLPACSAVAGL------------TGVKQIHGYAIRKQFNNDVSLCNALIDMYSKCGSLDHAKQVFE-----FGSFRRDPISWSSMISGYGLHGKGEEAVFVYNKMLQLGNKPDMITIVGVLSACSRAGLVDEGLCIYKSAikYRIKPTvcACVVDMLGRSGQLGQALDYIKTMP----MEPSPSVWGAVVSASIIHGNSEmaYRFLVQ-----LEpnPSNYVSLSNLHASSRRWDVVSEVRTMMKDRCLTKTP----------GCSWISINNTTHFFYAADKLHPCSKSIYELLGGLILLMK-----------------------------    
 438 UniRef50_C5X4X6        Putative uncharacterized pr...   388    1e-106  1  11.3%   7:574   193:731     ------YSRGGLMDAARKVFEEMPARDLVSWNALVCGLAQDgcSAEVIRVFLRMLKQG-DVRPDRISVCSVIPACGAEGKLELGRQIHGFAVKLGVEGHVSIANVLVAMYYKCGTPGCARRLFEFmeRDVISWTTVMS---MDGEDAVSLFNGMRRDGVAPNEVTFVAMLSSmgDCPAREGQMIHAVCLKTSLSDKAAAANSLITMYAKLRRMDDARMIFSLMPHSEI----IAWNALISGYAQNEMCQDALEAFLAMMKI-MKPNETTFASILSAVTAVETVSMA----------YGQMYHCQALKLGLGASEYVSGALIDMYAKRGSLEESWKAF------GETVHRSLIAWTAIISANSKHGNYDAVVSLFNDMVGSGVAPDGVVLLSVLTACRYSGFVSLGREIFDSMAAKhlWPEHYACVVDMLGRAGRLEEAEELMLQMPS----GPSVSALQSLLGACRIHGNTDVGERVASVLTET--EPtsGAYVLLSNIYAEKGDWGAVARVRRQMRERGVKKEVGFSWVDAGGAGDSL-----HLYKFSSDDTTHPQSEEIYRVAEGLGWEMK-----------------------------    
 439 UniRef50_Q9FI80        Pentatricopeptide repeat-co...   388    1e-106  1   9.1%  77:603    32:573     ----------------------------------------------------------------------------NCRTIRDLSQIHAVFIKSGQMRDTLAAAEILRFCahHRDLDYAHKIFNQmqRNCFSWNTIIRGFSESDEdaITLFYEMMSDevEPNRFTFPSVLKACAkgKIQEGKQIHGLALKYGFGGDEFvwNVMIDGYMRLGDCKAARMLFDKMRQR----SVVSWNTMISGYSLNGFFKDAVEVFREMKKGDIRPNYVTLVSVLPAISRLGSLELGEWL------------HLYAEDSGIRIDDVLGSALIDMYSKCGIIEKAIHVFERLPRE------NVITWSAMINGFAIHGQAGDAIDCFCKMRQAGVRPSDVAYINLLTACSHGGLVEEGRRYFSQMVsdGLEpeHYGCMVDLLGRSGLLDEAEEFILNMPIK----PDDVIWKALLGACRMQGNVEMGKRVANILMDM-VPHDSGAYVALSNMYASQGNWSEVSEMRLRMKEKDIRKDP----------GCSLIDIDGVLHEFVVEDDSHPKAKEINSMLVEISDKLRLAGYRP--ITT-QVLLNL----EEEDKEN    
 440 UniRef50_O23266        Pentatricopeptide repeat-co...   388    1e-106  1   9.3%  68:582     6:525     -------------------------------------------------------------------YLHQLQLCARNRTLTTAKALHAHIVKLGIVQCCPLANTLVNVYGKCGAASHALQVFDEmhRDHIAWASVLTALNqsGKTLSVFSSVGSSSGLRPDDFVFSALVKACAngSIDHGRQVHCHFIVSEYANDEVVKSSLVDMYAKCGLLNSAKAVFDSIRVK----NTISWTAMVSGYAKSGRKEEALELFRILPVK----NLYSWTALISGFVQSGKGLEAFSVFTEMRRErgRQVHGLVIALGFDSCVFISNALIDMYAKCSDVIAAKDIFSRMRHR------DVVSWTSLIVGMAQHGQAEKALALYDDMVSHGVKPNEVTFVGLIYACSHVGFVEKGRELFQSmkDYGIRPshYTCLLDLLGRSGLLDEAENLIHTMP----FPPDEPTWAALLSACKRQGRGQMGIRIADHLVSSFKLKDPSTYILLSNIYASASLWGKVSEARRKLGEMEVRKDPGH----------SSVEVRKETEVFYAGETSHPLKEDIFRLLKKLEEEMrrNGYVPDT---------------------    
 441 UniRef50_D1I4V9        Whole genome shotgun sequen...   388    1e-106  1   9.9%  68:602    16:557     -------------------------------------------------------------------HNLHLSLFQTCSAPQEVEQLHAFSLKTAIFNHPFVSSRLLALYskINDLGYARSIFDRirRSLIHWNTIIKCYVEnsHDGIVLFHELVHE-YLPDNFTLPCVIKGCARLGvqEGKQIHGLALKIGFGSDVFVQGSLVNMYSKCGEIDCARKVFDGMIDKDVVlnLVSWNAMINGYMKSGDFDSALELFYQMP----IWDLVTWNLMIAGYELNGQFMDAVKMFFMMLKLggRWIHSYMEKNGFELDGILGTSLIEMYAKCGCIESALTVFRAIQKKK------VGHWTAIIVGLGIHGMANHALALFLEMCKTGLKPNAIIFIGVLNACNHAGLVDDGRQYFDMMmeYKIEpeHYGCLVDILCRAGHLEEAKNTIENMP----ISPNKVIWMSLLGGSRNHGKIDIGEYAAQRVIE--VAptIGCYILLSNMYAASGMWEKVSHVREMMYKRGFRKDP----------GCSSVEHKGTLHEFIVGDISHPQTKEIYAKMSEMKEKLKCVGHVPDTTQVL------LCIEGEKEKE-    
 442 UniRef50_D1IEQ9        Whole genome shotgun sequen...   388    1e-106  1   9.6%  71:579     8:495     ----------------------------------------------------------------------LLNSIHRCKTIRSLKQIHAHLIASGLLHDEFVLSKVSEFFGKhgFVDYAFDFLNQTDLHVGtyNTLIAAYASspKAAFLVYGRIVGNGFVPDMYTFPVVLKACTklGVQEGEQVHGVAVKMGFLCDLYVQNSLLHFYSVCGKWGGAGRVFDEMLVR----DVVSWTGLISGYVRTGLFDEAINLFLKM---DVVPNVATFVSVLVACGRMGYLSMGKGV------------HGLVYKRAFGIGLVVGNALVDMYVKCECLCEARKLFDELPDR------DIVSWTSIISGLVQCKQPKDSLELFYDMQISGVEPDRIILTSVLSACASLGALDYGRWVQEYIERQGIEWDIhtALVDMYAKCGCIEMALHIFNGIPNR-----NIFTWNALLGGLAMHGHGHEALKHFELMIGAGIRPNEVTFLAILTACCHSGLVAEGRSYFYQMISQPFNfpRLEHYGCMIDLLCRAGLLDEAYNGVYVLLSNIYATNERWDDVTRVRRLMKDKGIR------------------------    
 443 UniRef50_D1IK42        Whole genome shotgun sequen...   388    1e-106  1  11.7% 138:580    72:473     -----------------------------------------------------------------------------------------------------------------------------------------YASLGRLDYSVALFGRTQ----NPSVFFWTAIIHGHarGDVVSAQQLFDTMPEK----SLVSLTAMLTCYAKHGELDAARVLFDGMEER----DGVCWNVMIDGYTQNGMPNEALVLFRRMLKAKAKPNEVTVLSVLSACGQLGALESGRWV------------HSYIENNGIQFNVHVGTALVDMYSKCGSLEDARLVFDKIDDK------DVVAWNSMIVGYAMHGFSQEALQLFKSMCRMGLHPTNITFIGILSACGHSGWVTEGWDIFNKMkeYGIEpeHYGCMVNLLGRAGHVEQAYELVKNM----NIEPDPVLWGTLLGACRLHGKIALGEKIVELLVDQNLA-NSGTYILLSNIYAAVGNWDGVARLRTMMKDSGVKKEP----------GCSSIEVNNKVHEFLAGGLNHPKRKEIYMMLEEINGWLKSHGYTP-----------------------    
 444 UniRef50_D1H2G3        Whole genome shotgun sequen...   388    1e-106  1   9.7%  47:551    50:533     ----------------------------------------------ANQALELFHSVSRRADlnPQLYSAIIHVLTGAKLYAKARCLMRDLIQCLQNS------------RRSRICCSVFNVLSRLESSKFTPNvlIIAFSEMgeEALWVYYKM---DVLPAMQACNMVLDGLVkgRFDTMWKVYGDMVARGASPNVVTYGTLIDGCCRQGDFLKAFRLFDEMIEKKIFPTVVIYTILIRGLCGESRISEAESMFRTMRNSGMLPNLYTYNTMMDGYCKIAHVKKALELYQEMLGD------------GLLPNVVTFGILIDGLCKTDEMVSARKFLIDMAS--FGVVPNIFVYNCLIDGYCKAGNLSEALSLHSEIEKHEILPDVFTYSILIKGLCGVDRMEEADGLLQEMKKKGFLPntYNTLIDGYCKEGNMEKAIEVCSQMTEKGI-EPNIITFSTLIDGYCKAGKMEAAMGLYTEMVIKGLLPDVVAYTALIDGHFKDGNTKEAFRLHKEMQEAGLHPNVFT-LSCLIDGLCKDGRISDAIKLFLA----------------------------------------------------    
 445 UniRef50_Q0WN60        Pentatricopeptide repeat-co...   388    1e-106  1   8.6%   4:585   331:886     ---NALMDMYSKCgnAQMIFKMNNNKNVVSWNTMVGGFSAEGDTHGTFDVLRQMLAGGEDVKADEVTILNAVPVCFHESFLPSLKELHCYSLKQEFVYNELVANAFVASYAKCGSLSYAQRVFHGIRSKTvsWNALIGGHAqdPRLSLDAHLQMKISGLLPDSFTVCSLLSACSKLklRLGKEVHGFIIRNWLERDLFVYLSVLSLYIHCGELCTVQALFDAMEDK----SLVSWNTVITGYLQNGFPDRALGVFRQMVLYGIQLCGISMMPVFGACSLLPSLRLGRE------------AHAYALKHLLEDDAFIACSLIDMYAKNGSITQSSKVFNGLKEK------STASWNAMIMGYGIHGLAKEAIKLFEEMQRTGHNPDDLTFLGVLTACNHSGLIHEGLRYLDQMKSsgLKPnhYACVIDMLGRAGQLDKALRVveEMSE----EADVGIWKSLLSSCRIHQNLEMGEKVAAKLFE--LEPepENYVLLSNLYAGLGKWEDVRKVRQRMNEMSLRK----------DAGCSWIELNRKVFSFVVGERFLDGFEEIKSLWSILEMKISKMGYRPDTMSV------------------    
 446 UniRef50_D1I9Q0        Whole genome shotgun sequen...   387    1e-106  1   9.9%  77:599    66:568     ----------------------------------------------------------------------------KCRSMRVLRQIHARLLTHAMPISSISFGliIGFCALsgDIDYARKLFSQIQrnIFSWNSMIRGCSQssKEPVILFRKMVRRGYppNTFTMAFVLKACsvSALEEGQQVHANVLKSGFGSSPFVETALVNFYAKCEDIVLASKVFDEITDR----NLVAWSTMISGYARIGLVNEALGLFRDMQKAGVVPDEVTMVSVISACAASGALDTGKWV------------HAYINKQLIETDLELSTALVNMYAKCGCIERAKEVFDAMPVK------DTKAWSSMIVGLAINGLAEDALEEFFRMEEAKVKPNHVTFIGVLSACAHSGLVSEGRRYWSSMLEFGIVPSmyGCMVDLLCRASLVEDACTLVETMP----ISPNPVIWRTLLVGCKKSKNLDKSEVVAQRLLE--LEpnAENYILLSNLYASMSQWEKMSQVRKKMKGMGIKAVP----------GCSSIEVDGLVHEFVMGDWSHPEAMEVREILRDISKRVHAVGHQPGISDVLHNVvnALCEHSER----    
 447 UniRef50_Q9SFV9        Pentatricopeptide repeat-co...   387    1e-106  1  11.4%   6:603   140:759     -----MLKLMYCFDELREVF--GFRLNYPCYSSLLMSLAKLDLGFLAYVTYRRMEADGFVVGM--IDYRTIVNALCKNGYTEAAEMFMSKILKIGFVLDSHIGTSLLLGFCRGLNLRDALKVFDVMspNSVSYSILIHGLcgRLEEAFGLKDQMGEKGCQPSTRTYTVLIKALCDRGldKAFNLFDEMIPRGCKPNVHTYTVLIDGLCRDGKIEEANGVCRKMVKDRIFPSVITYNALINGYCKDGRVVPAFELLTVMEKRACKPNVRTFNELMEGLCRVGKPYKAVHLLKRML------------DNGLSPDIVSYNVLIDGLCREGHMNTAYKLLSSM--NCFDIEPDCLTFTAIINAFCKQGKADVASAFLGLMLRKGISLDEVTGTTLIDGVCKVGKTRDALFILedMLSKGCKvvTYTTLVDGLIRSGDITGSFRILELMKLSGCL-PNVYPYTIIINGLCQFGRVEEAEKLLSAMQDSGVSPNHVTYTVMVKGYVNNGKLDRALETVRAMVERGYELNDRIYSSLLQGFVLSQKGIDNSEESTVSDIALRETDPECINELISVVEQL--GGCI-SGLCIF--LVTRLCKEGRTDESN    
 448 UniRef50_Q0J6X9        Os08g0249600 protein n=5 Ta...   387    1e-106  1   9.2% 114:581   155:628     -----------------------------------------------------------------------------------------------------------------GEDQAALVTSGWLRRSADLHNALIRALAgrPHLALPLYAHLLRAGHHPTPHTLPSLLKSLAlgARGLALAVHAHAVKLGlsVDASTFNTLITAYARAGRVTDARALFDEMPAR----NAVSWSAMVNGYVQAGDGREALGLFARMQADGVRPDDTVLVGVLAACAQLGVLEQGKWV------------HGYLKANNIRITVFLGTALVDMYAKCGEMQLAMEVFKVMKEK------NVLAWTTMIKGLAMHGRGSEALELFSQMERLGVKPDDIAFIGALCACTHTGLVDKGRELFDSmrKYGIKpeHYGCMVDLLARNGLLNEAREMVEKMPMK----PDALIWGALMAGCRFHKNVELAEYVVKHWIE--LEPdsGAYVLLGNIYAASGRHASAREIRHLMREKGVDKTP----------GCSTVEIKGVIHQFIVGDLSHPFIEEILSKWDEIDSRIreEGYVPD----------------------    
 449 UniRef50_A9RIA0        Predicted protein n=3 Tax=E...   387    1e-106  1   9.7%  16:602    26:620     ---------------AKTMEGYRNKLTINDFSLIFREFAQRSDWQKALRLFKYMQRHQ-WCKPNEHVYTIMIGIMGREGMLDKASELFEDMPLNDVEWNVYSFTALINAYGRNGQHEASLHLLARMKREKVTPNLITYNTVINACalLGLFAQMRHEGIQPDIITYNTLLSACSSRGleEAGMVFRTMNEAGVVPDSITYNALVDIYGQADRHEGVGELLREMEQAGNAPDVVAYNILIEAYGRAGKYRAAAKMFKQMQEAGCTPDVVTFSTLLEAYGKHGCYDEVRLLFTDMKER------------GTEPDVNTYNTLIQVFGQGGFFQESINLFWDLLDGG--VEPDMSTYAGLLYSCGKGGLHKAAKKIHRHMLQSYVTPTTDGFTGLITAYGNAALYSEATYAFNSMKESGCKPdtYNALIGAHAGGGLYCEAGSAYLTMIDEGI-SADVSTYNSLIEAFGRGGLFDDAIEFSRDMEEARCSPNRHTYEALMGVYCTAGLFDEAKAQFLDL-QVGGELPSVDSYCLLLSVCARRNRWDDASKVLEEMLE--PNMPSIYGVVGGILkeMDddGLKVDGvrFYNALLEALWCLGQRERA-    
 450 UniRef50_Q3ECK2        Pentatricopeptide repeat-co...   387    1e-106  1  11.0%  43:534    65:541     ------------------------------------------KLNDAIDLFSDMV--KSRPFPSIVDFNRLLSAIVKLKKYDVVISLGKKMEVLGIRNDLYTFNIVINCFCCCFQVSLALSILGKMLKLGYEPDRVTIGSLVNgsDAVSLVDKMVEIGYKPDIVAYNAIIDSLCkkRVNDAFDFFKEIERKGIRPNVVTYTALVNGLCNSSRWSDAARLLSDMIKKKITPNVITYSALLDAFVKNGKVLEAKELFEEMVRMSIDPDIVTYSSLINGLCLHDRIDEANQMFDLMVSK------------GCLADVVSYNTLINGFCKAKRVEDGMKLFREMSQRG--LVSNTVTYNTLIQGFFQAGDVDKAQEFFSQMDFFGISPDIWTYNILLGGLCDNGELEKALVIFEDMQKREMDLdtYTTVIRGMCKTGKVEEAWSLFCSLSLKGLK-PDIVTYTTMMSGLCTKGLLHEVEALYTKMKQEGLMKNDCTL--------SDGDITLSAELIKKMLSCGYAPSLLKDIKSGVC---------------------------------------------------------------------    
 451 UniRef50_O04590        Pentatricopeptide repeat-co...   387    1e-106  1   9.6%   9:603    52:632     --------RSGYIAEARDIFEKLEARNTVTWNTMISGYVKRREMNQARKLFDVM------PKRDVVTWNTMISGYVSCgfLEEARKLFDEMPSRDSFSW---NTMISGYAKNRRIGEALLLFEKMPERNAVSWSAMITGFCQNgdSAVVLFRKMPVKDSSP----LCALVAGLIKnrLSEAAWVLGqsLVSGREDLVYAYNTLIVGYGQRGQVEAARCLFDQIprERFCKNVVSWNSMIKAYLKVGDVVSARLLFDQMKDR----DTISWNTMIDGYVHVSRMEDAFALFSEMPNR----------------DAHSWNMMVSGYASVGNVELARHYFEKTPEKH------TVSWNSIIAAYEKNKDYKEAVDLFIRMNIEGEKPDPHTLTSLLSASTGLVNLRLGMQMHQIVVKT-VIPDvhNALITMYSRCGEIMESRRIFDEMKLKRE----VITWNAMIGGYAFHGNASEALNLFGSMKSNGIYPSHITFVSVLNACAHAGLVDEAKAQFVSMMSVYKIEPQMEHYSSLVNVTSGQGQFEEAMYI----ITSMPFEPDKTVWGALLDACRIYNNvePESSTPYVLLYNMYADMGLWDEAS    
 452 UniRef50_D1IDP0        Whole genome shotgun sequen...   387    1e-106  1   8.8%  61:581    42:561     ------------------------------------------------------------PSLSSSTYTNYLhlLSSCKHLNPLLQIHAQIIVSGFKHHHSITHLINLYSLFHKCDLARSVFDSTPNPSRILWNSMIRAysKQYNEALEMYYCMVEKglERDVFIGAGLVDMYSkgDLKRAREVFDKMPKR----DVVAWNAMIAGLSQSEDPYVARRVFDQMVDQ----DDVSWGTMMAGYAHNGCFVEVLELFDKMKLGN-----VTWNVIIAAYMQNGHAKEAISSFHQMRlrEGMAFHACIIQMGFLSNTLVGNSLIDMYAKCGQLDYSEKLFNEMDHK------DTVSWNAMLSGYAVHGHGDRAIALFSLMQESQVQIDSVSFVSVLSACRHAGLVEEGRKIFHSMskYHIKPdhYACMVDLLGRAGLFDETLGFIKVMP----VEPDAGVWGALLGSCRMHSNVKLGEVALDHLVK--LEpnPAHFVVLSSIYAQSGRWADAGKARSKMNDLGLKKTP----------GCSWVELKNKVHAFRVGDKSHPQLESMHLLWNTLLEKMEKIGYVPD----------------------    
 453 UniRef50_B9T205        Pentatricopeptide repeat-co...   387    1e-106  1  10.6%   6:523    85:624     -----LYAKGGNIDEARLVFDGLLEKSAVTWTTIITACVKRGRSEVSLQLFSQMRE--TNVVPDGYILSSVLGACSQLEFVQGGKQIHGHVLRRGIEIDVSFVNVLIDFYTKSGKVQSARKLFDGmdRNVISWTAMIAGYMQNsrEAVKLFIEMTRLGRRPDGFVCTSILTSCGSLelELGRQVHAYSIKGNVESDIFLQNGLIDMYAKCGSLNDARKVFDDMTIR----NVVSYNALIEGYSTLEQLSEAMNLFREMRHGMLSPSFLTFVSLLGASATLSALELGKQI------------HALITKFGISMEIFAGSALIDFYSKCSCLMDARLVFDKMTEK------DIVVWNAMLFGYTQQLENEEALKLYTELQISEPKPNVVTfnSLIDMYAKCGSLEDARkyFESMPKFSIKpeHYACVVSLLGRSGKLYEAKEFIEKMPT----EPEAVVWRSLLSACRVSGNVELGKYAAEKAISIDST-DSGSYTLLSNIYASKGMWVDVKKVRERMDIAGVVK--------------------------------------------------------------------------------    
 454 UniRef50_Q65XT3        Os05g0132000 protein n=3 Ta...   387    1e-106  1  12.0%   4:593    55:617     ---EATVRRLAAAGdlQEMRLRGVACTEDALVAAIGAFARAGSADRALKTFYRASDLGCR-DPGVRVYNHLLDALLRENMVGAVVPVYDNMRKAGVDPNVYTYNLLIRALCQNDRVDAARKMLDEMSRKGCHPDEVSHgkLGRVEEARGFLAE-----TVPVQASYNAVVHacGEFRMWEVFSVVNEMVQRGLQPNVVTYTTIVDAFCKARELRMACAILARMVSMGCTPNVLTFTALVKGFFEDGKVHDALSMWHWMVDEGWAPSTISYNVLIRGLCCIGDLKGALDFFNSMKR------------NALLPNATTYSTLVDGFSNAGDLDGAMLIWNEMKSSGC--KPNVVVYTNMIDVLCKKMMFDQAESLIDKMLMDNCPPNTVTFNTLIGRLCDCGRVGRALNVFHGMRRNGCHPntYNELLHGLFREGNHKDAFAMVIEMLNNG-FELSLVTYNTVINCLCQMCMRKHAMLLLGRMMVQGIQPDAFTFNAIIHAYCKEGKVSIAACLLGQMNA----------VNCPRNVVAYTILISELCNQGKLSNAMVYLLKMLYE------------GICPNE-ATWNVLVRAI----------    
 455 UniRef50_C5XR38        Putative uncharacterized pr...   387    1e-106  1  10.0%   6:587   127:693     -----RYAQDGFLDEAFemEESGIPIGASTWNAVIAGCVRAGNGELAIELLGKMVSIG-GVVPNVATFNTVLHVISVLRGVDVLRELHAFVLrlGPVDLDRLQESLAAGYMCSSFVQYAGRVFRDVrtTCFLGNLMISGFLDTGQraLGVFREM-AFGCGYEARHLPSVSLTLVLPEVNLalEIHAYAYRHGFECDTSVCNALMAMYAKRDKICLAYKIFQGLDNK----DVVSWNTMISSYAAVHDFDLSFELFREMQRNDTRPDDYTFTSVLNACSFTCNHRTVMALHGQMIRMGL-----------CHSYVDDMNSLMDAYGKCGFIDEAQNIFDETDRK------DTISWNIIISCYGYSGFPQQAIRLFHQMQDQGYKPTRVTFISVLAACSHAGLVDEALHYFEGMRryNITAdhYACIIDCFGRAGQLKQAYEFIRGMP----VVPNACVWGALLSSCRIHGNIGLAEIAAKKLIE--LDPQHSGYWMLldIYAKSMRWNDVSQLRTAMRDKGIK-------KCPGYSWIEVG--DSEVHRFLTADKLHTQSHQIYRVLGGLTQQLMDEGYEpdVDLTYT----------------    
 456 UniRef50_Q8GVW4        Os08g0375800 protein n=3 Ta...   387    1e-105  1  10.1%   2:576   130:682     -AASALLdkCGRVRDARQVFDGMPERNTVSWNALIAGYTESGDMASALELFLEMEREGL--APDEATFASLLTA-VEGPSCFLMHQLHGKIVKYGSALGLTVLNAAITAYSQCGSLKDSRRIFDGiiRDLISWNAMLGAYTHNgdEAMKFfrMMQESGVHPDMYSFTSIISSCSEHGHDdgRVIHGLVIKSALEGVTPVCNALIAMYTRYnmMEDAYKCFNSL----VLKDTVSWNSMLTGYSQHGLSADALKFFRCMCSENVRTDEYAFSAALRSSSELAVLQLGKQI------------HGLVIHSGFASNDFVSSSLIFMYSKSGIIDDARKSFEEADKS------SSVPWNAMIFGYAQHGQAENVDILFNEMLQRKAPLDHITFVGLITSCSHAGLVDEGSEILNTMekYGvrMEHYACGVDLYGRAGQLDKAKKLIDSMP----FEPDAMVWMTLLGACRIHGNVELASDVASHLFVA--EPrhSTYVLLSSMYSGLGMWSDRATVQRVMKKRGLSKVP----------GWSWIEVKNEVHSFNAEDKSHPKMDEIYEMLRVLleQMLSN---------------------------    
 457 UniRef50_Q9LRV2        Pentatricopeptide repeat-co...   387    1e-105  1   8.4%   7:552   106:659     ------YGKCGCVDDARELFEEMPERDGGSWNAVITACAQNGVSDEVFRMFRRMN--RDGVRATETSFAGVLKSCGLILDLRLLRQLHCAVVKYGYSGNVDLETSIVDVYGKCRVMSDARRVFDEIvpSDVSWNVIVRRYLEMgdEAVVMFFKMLELNVRPLNHTVSSVMLACSRsaLEVGKVIHAIAVKLSVVADTVVSTSVFDMYVKCDRLESARRVFDQTRSK----DLKSWTSAMSGYAMSGLTREARELFDLMPER----NIVSWNAMLGGYVHAHEWDEALDFLTLMRQemGKQAHGFIYRHGYDTNVIVANALLDMYGKCGTLQSANIWFRQMSE-----LRDEVSWNALLTGVARVGRSEQALSFFEGMQVE-AKPSKYTLATLLAGCANIPALNLGKAIHGFLIRDGYKIDVvgAMVDMYSKCRCFDYAIEVFKEAATR-----DLILWNSIIRGCCRNGRSKEVFELFMLLENEGVKPDHVTFLGILQACIREGHVELGFQYFSSMSTKYHISPQVEHYDCMIELYCKYGCLHQLEEFLLLM---------------------------------------------------    
 458 UniRef50_B9HPD9        Predicted protein n=2 Tax=M...   387    1e-105  1  10.8%   6:525   113:612     -----MYAKCGEISSARVLFDQMPQRTVVTWNSLIYGYLAVNCPKIAIELFIKM--LKAAINVSAFSVSSCLAGCSQLEAREVGTQVHGLILKTGLGYNVVVGTSLVDMYSKCGDVDDSRLVFDHMvrNVITWTSMVTGYSqkPDEAMALVKEMVLQDLRPNCVTYNSLLSSFSGPdlSYCLQVHCCIIQLGLESNVYIAATLVTVYSKCSslED----FKKVCSVVMTCDNIAWNAVIAGYSKLGRHEEALRCFHEMKQAGIDIDSYTLTSVVGAIGNSSFLEEGKAM------------HALIHRTGYISHLNVQNGLVSMYARCGAIGDSKRVFWFMEEH------DVISWNALLTAFAHHGYGREAVELFEQMRKTEIKPNSSTFLAVLCACSHVGFVDKGIEYFDTMKSDilKVEHYASLVDTFGRAGYLNEAEAFINSMP----IVPAPSVYKALLSASLVHGNREIAARSAKKLLEL-WPNDPATYVLLSSVLTVDGNWDDAADLRKLMCDRGLRKKP------------------------------------------------------------------------------    
 459 UniRef50_Q9M1V3        Pentatricopeptide repeat-co...   386    1e-105  1   9.1%   5:589   328:880     ----AMYTRCGKMPQAERILRQMNNADVVTWNSLIKGYVQNLMYKEALEFFSDMIAAGH--KSDEVSMTSIIAASGRLSNLLAGMELHAYVIKHGWDSNLQVGNTLIDMYSKCNLTCYmrAFLRMHDKDLISWTTVIAGYAQNDceALELFRDVAKKRMEIDEMILGSILRASSVLkmLIVKEIHCHILRKGL-LDTVIQNELVDVYGKCRNMGYATRVFESIKGK----DVVSWTSMISSSALNGNESEAVELFRRMVETGLSADSVALLCILSAAASLSALNKGREIHCYLLRKGFCLEGSIAV------------AVVDMYACCGDLQSAKAVFDRIERKGLLQ------YTSMINAYGMHGCGKAAVELFDKMRHENVSPDHISFLALLYACSHAGLLDEGRGFLKIMeeYELEpeHYVCLVDMLGRANCVVEAFEFVKMMKT----EPTAEVWCALLAACRSHSEKEIGEIAAQRLLE--LEpnPGNLVLVSNVFAEQGRWNDVEKVRAKMKASGMEKHP----------GCSWIEMDGKVHKFTARDKSHPESKEIYEKLSEVTRKlrEVGYVADTKFVLHNV--------------    
 460 UniRef50_D1IWE7        Whole genome shotgun sequen...   386    1e-105  1  11.8%  80:580     5:482     -------------------------------------------------------------------------------SLSQAMQLHAQILkrNLTPLFTFAAL-----SPAGDLTYAHLILNSLsqNSFFHNTMIRAYSQTPdqALHLFLSMLCqsPRPDKFTYPFLLKSCARLKQPrgKQLHGLIYKSGLESDRYVSNGLIHMYSSCGKSGRAYKVFGKMRDR----DVVSWTSMIDGFVDDDRALEAIRLFEEMVEDGVEPNEATVVSVLRACADAGAVGMGRRVQGVIEER----------KIGLEANVRT--ALIDMYAKCGSIGSARKVFDGIVNK------DVFAWTAMISGLANHGLCEEAVTLFDQMESFGLRPDERTMTAVLSACRNAGWFSEGFAYFNSmcKYGIKPthYGCMVDLLARTGHLDEAEEFIRKMP----IEPDVVLWRTLIWASKVHGDIDRSEQL---MKDRGlkMDsgSYVLLGNVYASAGKWHDKAKMRELMNQKGLSKPP----------GCSRIEVDGLVHEFAAGDSGHIEAEKIYAKLDEVEERLKAEGYHP-----------------------    
 461 UniRef50_A2Q3R1        Pentatricopeptide repeat n=...   386    1e-105  1  10.5%   1:523   191:719     LSFNFLINRLVKHDEVDMALCLFVRfnEYTYTIVIKALCKKGDWENVVRVFDEMKEAGVDD--DSYCYATFIEGLCKNNRSDLGYAVLQDYRTRNAHVHKYAYTAVIRGFCNETKLDEAESVFLEMEKQGLVPDVYVYCAlrNFDKALAVYKSMISRGIKTNCVIFSCILHCLDegRALEVVDMFEEFKESGLFIDRKAYNILFDALCKLGKVDDAVGMLDELKSMQLDVDMKHYTTLINGYFLQGKPIEAQSLFKEMEERGFKPDVVAYNVLAAGFFRNRTDFEAMDLLNYMESQ------------GVEPNSTTHKIIIEGLCSAGKVEEAEEFFNWL--KGESVEISVEIYTALVNGYckSHELKEAFILLRTMLEMNMKPSKVMYSKIFTALCCNGNMEGAHTLFNLFIHTGFTPdtYTIMINGYCKTNCLPEAHELFKDMKERGI-TPDAVTYTIMINGYCKMNCLREAHELFKDMKERGIKPDVIAYTVIIKGLLNSGHTEIAFQLYNEMIDMGMTP--------------------------------------------------------------------------------    
 462 UniRef50_B9S6P4        Pentatricopeptide repeat-co...   386    1e-105  1  11.2%   7:523    71:567     ------YAKSSCTESALKVFDTMPLKDPISWNCIINGYFQNGYFTQGFIMFKEMYECGIVPKPELVA--SVVSVCVRLGDMKLGRAIHALVLIDGRieIFVLTSLVDWYFKSGDSLMALRVFDQIEVKNVVSWTALVSGCSANLdmALDCFRAMQINGVRPNRVTLIAILPAFAELGCAkgKEIHGYAFRHGFDLDHHFLSSLIHIYCRSRelQLAKLTF----ERSNVKDVVMWSSIIGSYCRSADSIEALKLFRWMRADGTEPNSVTLLAVITACTTLTSL------------KNGYGTHGYIVKCGLNFDTYIGNALINMYSKCGCVVASHQIFKEMHIK------DSVSWSTLITAYGLHGHGKEALCLFHEMQLQGVEPDAMTFLAVLSACNHSGLVKEGQQIFDNVKKDglTVEHYACLVDLLGKSGKVDDAFDILRAMPMK----PSTTIWSSLVSACKIHGRLEIAERLAQELIKS--EpnAANHTLLSMIYAESGNWFAVEDVRRLMRVQGLSK--------------------------------------------------------------------------------    
 463 UniRef50_O49436        Pentatricopeptide repeat-co...   386    1e-105  1  10.2%  17:595    61:643     ----------------EKMFKSAPKMGSftLSSMIESYANSGDFDSVEKLLSRIRLENRVII--ERSFIVVFRAYGKAHLPDKAVDLFHRMvkRSVKSFNSVLNVIINEGLYHRGLEFYDYVVNsiSPNGLSFNLVIKALcrFVDRAIEVFRGMPERKCLPDGYTYCTLMDGLCkeRIDEAVLLLDEMQSEGCSPSPVIYNVLIDGLCKKGDLTRVTKLVDNMFLKGCVPNEVTYNTLIHGLCLKGKLDKAVSLLERMVSSKCIPNDVTYGTLINGLVKQRRATDAVRLLSSMEER------------GYHLNQHIYSVLISGLFKEGKAEEAMSLWRKMAEKGC--KPNIVVYSVLVDGLCREGKPNEAKEILNRMIASGCLPNAYTYSSLMKGFFKTGLCEEAVQVWKEMDKTGCSRncYSVLIDGLCGVGRVKEAMMVWSKMLTIGIK-PDTVAYSSIIKGLCGIGSMDAALKLYHEMlePKSQPDVVTYNILLDGLCMQKDISRAVDLLNSMLDRGCDPDVIT-CNTFLNTLSEKSNSCDKGRSFLEELVVRLLKRQRVSGACTIVEVMLGKYLAPKT-STWAMIVREICK--------    
 464 UniRef50_A9SGY6        Predicted protein n=1 Tax=P...   385    1e-105  1  13.0%   1:515     5:505     LGAARRYDDVQRL--VESMQKDRTKLDTQLFNTLIHIYGEANMMEKALQTLAAFTKEG--GRPTAYTYSSMIQVFMKGGDVQNGLLMYKQMLKAKFVPDHTTFNILIDSLAKADQVEEACNVFGDMFKLNVSPDVYTygTIGRIDAVMKLFESMTAQGCQPNLFTYHSVMHAFGsgRVDEACDIFQQMVQKGLQPDAVTYNILIDAFGKTGQLERA---FDFVGKSRSFTNEVTYNSLLSSLGRKGDIQGLMELFGQMKAKGLVSNELTYAILIERLGWAGRVEDVWQLYLEM------------VDNDIKYDIVTINNVLDCLSKAGRVEAAFELFQDMKFKG--LNADTVTYNILINGLGRAGKLDAAGALLLEMEENGCAPNIITYNTLISSYGKWSNLSAATRLFLEMKERGVAPNvySSLIEGFGKAGRTDAAISLFREMKAEGC-PPNHVTYNLLIDCLIRAGRFGAAMEYLREMRDAGCQMGKVTKSLLVRiy-----MEEAQSFFQE----------------------------------------------------------------------------------------    
 465 UniRef50_Q8H8U1        Os03g0283500 protein n=2 Ta...   385    1e-105  1   9.3%   5:530    57:598     ----SLYSRADRLRDARRVFDSIPQPSLFAWNAILISL--SLHSPDPSAAVRLF--ASSAVSPDEITLSTLLRSLAASGPALSPleLHAVAFLRGFGSDLFVSNALITAYANAGDMRSARAVFDEmrRDVVSWNSLISACARAgrECLDLFQEFVrdGVGPNGVTVTSVLHACAQLKVVdgIGVHRFAAESGLDMDMAVWNSIIGFYAKCGRLQYARQLLDGMTRK----DSISYSAMITGYMNNGHVEEGMQLFRQASARGIS----MWNSVIAGLVQNGRQSDVLRLLQEMIAsgAKQAHGYAIRNDYDQSIRLVSALIDAYAKAGFLDTARKVFKLTEHR------STIVWTSIISAVAAHGEAVEALSLFNQMITAGAKPDTVTFTTVLSACAHSGKVAEARKVFNSMQafGISPViyACMVSALSRAGMLKEAVKLVNKMP----FEPNAKVWGALLNGAAVVGDVEFGRYAFDRLFV--IEpnTGNYIVMANLYSNAGKWEEAETIRSMLWGVGLEKVPGCTWN-------------------------------------------------------------------------    
 466 UniRef50_A5BDH5        Putative uncharacterized pr...   385    1e-105  1  10.3%   4:574   199:808     ---NSRISDCMRNgeAQMLFDEMPQRNTVTYNAMIRGYFQNGHFGEGVSLFDEM------PERDIFSYNTMIAGLMKFGDINGASEIFQKMP---FRDVVSWNSMISGYVSNGLIGEALRVFSGMVLKDVVSWNLVIAGlvGKVDLAEEFFKEMGTR----DIASWTTMISGLAsgRIVEARGLFEDMPVR----DVRAWNTMIAGYLENGCIEIGEVLFQKMPQRDFR----SWNEMINGLVRNQRIQNAMRLFVEMPQkglspdTVSWTNLIVGYFETGEVDTAVSIFELMPAreGLKFFVKMKEGGPFPDEATfnAMVTLYARCGNSNSALLLFSSMTSH------DVISWNSIICGLAHNGNGVEAIEMFEKMRSTDIKPNRITFVGVLSACSHAGLVDQGKYYFDFMKYKCctIEHYTCIVDLLGRFGLIDEAMSFLRQMEANGVEVP-ASVWGAVLGACRIHKNIQVGEIAGERILE--IEPHNF-----------CGKREDAERVWVRMREKGVKKQPA----------CSWMEVNGSGHVFLSGDSSHPQFSRVCGVLGLLHMEME-----------------------------    
 467 UniRef50_D1I6L4        Whole genome shotgun sequen...   385    1e-105  1  11.5%   1:581   173:805     LGSTGLVEEANLLFDQVKMMRL-CVPNSYSFNCLLEAISKSGSIDLVEMRLKEM--CDSGWEPDKYTLTSVLQAYCNSRKFDKALSVFNEIYGRGWVDGHVLSILVLTFSKCG-EVDKAFELIERMEDLGIRLnvLIHGFVrrVDKALQLFKKMQKSGFAPDVSVYDALIGgcAKKEIEKALHLLSEMKELGIDPDIQiyNSVLNGLVNGKSVDKAYYLLRAmvKEMVRPDTTSFSIVIDGLCNTGKLDLALSLFRDMVRVGCKQNVLLYNNLIDKLSNSNRLEECYLLLKEMKGS------------GFRPTQFTHNSIFGCLCRREDVTGALDMVREMRVHGH--EPWIKHYTLLVKQLCKRKRSAEACNFLAEMVREGFLPDIVAYSAAIDGFVKIKAVDQALEIFRDICARGYCPDvyNTLINGFCKVKRVSEAHDILDEMVAKGLV-PSVVTYNLLIDGWCKNGDIDQAFHCLSRMVGKEREPNVITYTTLIDGLCNAGRPDDAIHLWNEMRGKGCSPNRISFI-ALIHGLCKCGWPDAALLYfyVALITSFISNKNPTLAFEILKEMVAKGKFPD----------------------    
 468 UniRef50_Q2HUC3        Tetratricopeptide-like heli...   385    1e-105  1  10.0%   2:601   130:760     -ALNSVICVYAEYGFVDKAVKMfcFPCVVANNSLLNCLVKNGKVDVACELYDKMLehGLDLVVDNYSIVIVVKGLCDVGKVEEGRKLIDDRWGNGCVPNVVFYNVIIDGYCKKGDLKRATRVFEELKLKGFLPTLETYGALIDgcKadQLLNEMNVMGLNVNVKVFNSIIDayKYGLVDKAAEMMRMMTEMGCEPDITTYNILINFSCSGGRIKEAEEFLERAKERTLLPNKFSYTPLMHAYCKQGDYVMASDMLFKIAETGDKPDLVSYGAFIHGSVAGGEIDVALMVREKMMEK------------GVFPDAQIYNVLMSGLCKKGRFPAAKLLLSEMLD--LNLQPDAYMYATLVDGFIRNNELDKATELFEVVMSKGIDPGVVGYNVMIKGLCKCGKMTDAVSYVNKMKIANHAPDEYttVIDGYVKQHDLDSALKMFGQMMKQKYK-PNVVAYTSLINGFCKIADMSRAEKVFRAMQSFNLEPNVVTYTILIGGFSKTGKPEKAASFFELMLMNNC-LPNDTTFHYLINGLtfATMISeqVIATYNSIIVCLCKHGMVDTAQLLQTKMLRKGFLMDSV-CFSALLHGLCQTGKSKE--    
 469 UniRef50_D1H242        Whole genome shotgun sequen...   385    1e-105  1  11.5%  10:583   100:718     ---------CRNLSFARIVFDQFPspPTIIWNLMIQAYSKTPSSQESLYLFHQMLAHGRPTSADKYTFTFVFTACSRHPTLryGENVHGMVVKDGYESDIFVGNSLVNMYSIFSRMVDAKRVFDEmqRDVITWTSVVKGYAMRGeeALQCFNDMlhDEVKPNEAVLVSILSACAHLglDQGKWIHVYIDKNRILLSSNISTALIDMYAKCGRIDCARRVFDGLHKR----DLLTWTSMISGLSMHGLGAECLWTFSEMLAEGFKPDDITLLGVLNGCSHSGLVEE-------------EIVHGMVVKSGFESNLYVGNSVINMCSVFARMEDARKVFNQMSER------DVFSWTSLLGGYAKHGEMDRalTFFCNMldDRVNPNEAVLVCVLSACAHLGALDQGNWIHLYIDKIGIrsNISTALIDMYAKCGRIDCASRVFN-----GICKRDVLSFTSMISGLSYHGLGKDALriLANMESlgIAPKIEHYGCYIDLLGRAGYLERALEVVKTMPMEPDIVIWRALLSalSNLYASLGrvNGLVHEFRVADQLHPQIVEIRNKLNEILKRLSQIGYSANTM--------------------    
 470 UniRef50_C5Y0Y8        Putative uncharacterized pr...   385    1e-105  1  12.0%   1:573    65:690     LAanSLLHAYLQCgrALRLLDGTPRRDAATYASLISAHCRLGAPLDALRAFLDMLDWGCsvRPNEFTAAAVLQACGLARDERLGRMVHGYLVAGGFcdPFVVGSLVNMYAKAGDVVSARRLVLGLPCRDVVSWTAIISGCVLNgeEGLEVFVMMLEDGVLPNNVTMLSVIQACSLMGASELFsvHALVVLLELEHDASVVNSLIMMYAKNGFVEEAIWLFrhTIKIGAFPSISIENSLMGMYARFEQIDAAHFVFEGMKVK----DIVSWNTIISCLAKSDRVNEAMELFSVLHahQGQMLHGYIMKSGFVYDVSICNALITMYAKLGRIDFAEMIFERMDIK------DLVSWNSMINAYGMHGDGHLALRVFHQLKDAGtvPNAITFVSVISACSHSGLISEGYKCFESmrDHSIEpdHYACVVDLLGRSGRFAEAEEFIRDMP----VHPNSSIWGPLLAACQLHGNVDLAEKAAKELSA--LEPESDIwvSLSNTYASAGRWRDAAKIRTEMRRVGLR-------KETGWSFVDVGGVESF--KFVSADTRHHDAEEIYSVWHSMNRHM------------------------------    
 471 UniRef50_Q8L844        Pentatricopeptide repeat-co...   385    1e-105  1  12.7%  31:581   170:688     ------------------------------YSILIHAL---GRSEKLYEAFLLSQKQTLTP----LTYNALIGACARNNDIEKALNLIAKMRQDGYQSDFVNYSLVIQSLTRSNKIDSvlRLYKEIERdvQLVNDIIMGFAKSGdkALQLLGMAQATGLSAKTATLVSIISALasGRTLEAEALFEELRQSGIKPRTRAYNALLKGYVKTGPLKDAESMVSEMEKRGVSPDEHTYSLLIDAYVNAGRWESARIVLKEMEAGDVQPNSFVFSRLLAGFRDRGEWQKTFQVLKEMKS------------IGVKPDRQFYNVVIDTFGKFNCLDHAMTTFDRMLSEG--IEPDRVTWNTLIDCHCKHGRHIVAEEMFEAMERRGCLPCATTYNIMINSYGDQERWDDMKRLLGKMKSQGILPntHTTLVDVYGKSGRFNDAIECLEEMKSVGLK-PSSTMYNALINAYAQRGLSEQAVNAFRVMTSDGLKPSLLALNSLINAFGEDRRDAEAFAVLQYMKENGVKP----------DVVTYTTLMKALIRV------------DKFQKVPVVYEEMIMSGCKPD----------------------    
 472 UniRef50_C5XC86        Putative uncharacterized pr...   385    1e-105  1  11.6%  28:536   111:626     ---------------------------APALGVLVIALSQMGLLDEALYVFRRLWTLPALP-----ACNAVLDGLVKARRSRCAWELFDEMLRRGMVPSVVTYNTLINACRYQGAVAKAQEMWDQmdPNVITYTSMICVLCEEGcdAERLFDAMKEAGMRPNQYTYNVLMSGHCQGdvNSAFVLYQELLNSGLIPNAVVFTTLIDGFCKAKRFSEAKDMFRDMPRFGVAPTVPVYNSLMDGAFRSGDAQEALSLYQEMTRLGLCPDEFTCSIVVRGLCDGGQIQVAARFLEGVRED------------GVNLNAAAYNALIDEYCRNGNLEEALATCTRMTE--VGVEPNVVSYSSLIDGHSKLGKMQIAMAIYTEMVAKGIEPNVVTYTALIHGHAKNGGIDAAFRFHKEMIENGISPntVSVLVDGLCRENRVQDAVRFV--MEHSGIKypNSAIYMTLIYGLYLDGQHYEAGKLFSYMRKSGMISDSFTYTLLIRGQCMLGYVLNAMMLYADMMKIGVKPMRYKIICPEIWSW-------------------------------------------------------------------    
 473 UniRef50_Q9LNP2        Putative pentatricopeptide ...   385    1e-105  1  11.8%   1:577    91:714     LAAnsVYARLGLLLDARNVFETVSLvdLRLWNSILKANVSHGLYENALELYRGMRQRGLTG--DGYILPLILRACRYLGRFGLCRAFHTQVIQIGLKENlvVNELLTLYPKAGRMGDAYNLFVEMPVRNRMSWNVMIKGFSqdCESAVKIFEWMQREEFKPDEVTWTSVLSCHSQCglSIAEKVHGYVIKGGFEEYLPSRNALIHVYGKQGKVKDAEHLFRQIRNKGIE----SWNSLITSFVDAGKLDEALSLFSELEEMnvKANVVTWTSVIKGCNVQGRGDDSLEYFRQMQFSkgREIHGHVIRTSMSENILVQNALVNMYAKCGLLSEGSLVFEAIRDK------DLISWNSIIKGYGMHGFAEKALSMFDRMISSGFHPDGIALVAVLSACSHAGLVEKGREIFYSmkRFGLEpeHYACIVDLLGRVGFLKEASEIVKNMP----MEPKVCVLGALLNSCRMHKNVDIAEGIASQL--SVLEprTGSYMLLSNIYSAGGRWEESANVR----------ALAKKKDLKKVSGSSWIEVKKKKYKFSSGSIVQSEFETIYPVLEDLVSHMLKKG--------------------------    
 474 UniRef50_Q60DN6        Os03g0644100 protein n=9 Ta...   385    1e-105  1  10.9%   7:500   125:594     ------YARFDEVSDAREVFDEMPERSVVSWTAMMNVYLKLGHYGEVVRLF--FDMVGSGVQGNSLTFVCLLKSCGERCDAKLGQQVHCCIVK-GGWSNVIVDSAIAHFYAQCGDVASASAIFDKMarDVISWTTMITAYVQHGhqALRMFSEMVSEGFRPNEFTVCSVLKACAEEkvRFGKQLHCAVLKKMYKNDIHIGSALVTMYARCGEVFDAQAVFDMMPRR----NTITWTSMISGYAQSGHGEKAILLFRKMKMRRVFVNNLTIVGLLSACGSLQSPYLGKEL------------HAQIIKNSMEENLQIGSTLVWFYCKCGEYTYAARILEAMPDR------DAISWTALISGYNNLGHNVEALKSLDDMLWDGVKPNTYTYSSALKACAKLEALQYGRKIHGFVNKTQDFSNVfsSLIDMYMRCGKVDEARSVFDAMPEH-----NLVTWKVIITGFAQNGLCEEALKYMYLMQQEGHEVDDFVLSTVLTSC-------------------------------------------------------------------------------------------------------    
 475 UniRef50_B9R7C6        Pentatricopeptide repeat-co...   385    1e-105  1   9.9%   5:600  482:1026     ----ALLDLYMKLGraHNAFDYMPIRDVVSWNALICGYSRNGYDFSALELFVQMLKLGFCPR--QTTLVGLLPSCGQLELVFQGKSIHGFGIKSGLHLDPQVKNALTSMYAKCGDLEAAEYLFemMDKSVVSWNTMIGAYGQNgdEAMFVFKRMIGAGVEVSQVTIMSLPSANANP-----ESIHCYTIKVGLADDASVVTSLICMYARYGSTDHAELLYWSLPQK----NLVSLTAIITSYAEAGNLGLVMESFSQMHQLNMKPDSVAMLSILHG------------IADPVHICIGHVFHGYAIKSGLDTFNLVTNGLISMYSKFNNVEALFGLFSGMHEKP------LISWNSVISGCVQAGRASHAIELFCQMKMHGCNPDAITIASLLSGCSQLGYLQFGERLHSYILRNKLEMEDftALIHMYTKCGSIVHAERVFKS-----IGKPCLATWNAMISGYSCYGFEHKALTCYSEMQEQGVEPDKITFLGVLAACTHGGLIHEGRRYFQIMTKVYDMVPTLQHCACMVGLLARVGL---------------------FEEALLFIKNMEK---EPDS-AVWGAFLSACCIHQEVK---    
 476 UniRef50_O80958        Pentatricopeptide repeat-co...   384    1e-105  1  12.6%  20:598   161:745     -------------------KRFGFELTPRAFNYLLNAYIRNKRMDYAVDCFGLMVDRKVVPFVPYV--NNVLSSLVRSNLIDEAKEIYNKMVLIGVAGDNVTTQLLMRASLRERKPEEAVKIFRRVMSRGAEPDGLLFSLAvdLVMALDLLREMrkLGVPASQETYTSVIVAFVkgNMEEAVRVMDEMVGFGIPMSVIAATSLVNGYCKGNELGKALDLFNRMEEEGLAPDKVMFSVMVEWFCKNMEMEKAIEFYMRMKSVRIAPSSVLVHTMIQGCLKAESPEAALEIFNDSFESWI--AHGFMCNKIFLL-----------FCKQGKVDAATSFLKMMEQKG--IEPNVVFYNNMMLAHCRMKNMDLARSIFSEMLEKGLEPNNFTYSILIDGFFKNKDEQNAWDVINQMNASNFEAniYNTIINGLCKVGQTSKAKEMLQNLIKEKRYSMSCTSYNSIIDGFVKVGDTDSAVETYREMSENGKSPNVVTFTSLINGFCKSNRMDLALEMTHEMKSMELKLDL-PAYGALIDGFCKKNDMKTAYTLfyNSLISGFRNLGKMDAAIDLYKKMVNDGISCD-LFTYTTMIDGLLKDGN-----    
 477 UniRef50_D1H4J3        Whole genome shotgun sequen...   384    1e-105  1  10.3%   6:510   157:679     -----MYAKCGEVDSAVRVYDKMTSLDAATCNCLISAYARNGFFVQAFQVFMQIGNMGT--RPNHYTYSTMLAVCGTISAIQEGKQLHAHVVKMQYLSETAVGNALLTLYSKCGMMEEAEIVFENlqRNIISWTASINGFYQHGdkALKQFSMMRESGIEPNEFTFSIVLASCG-CDLGKWMklLRKMIEEgfESDAYIISALIYMYSKCGHVEKACRVFDWI------PNVVSWNTLIAG-------------FSQMLDQGFCPSSVTISSLLPACTNVANLRHGKEI------------HGYAMVIGVEKDVYVRSALVDMYAKCGYISEAKILFYMMPER------NTVTWNSLIFGYANHGYCNEAIELFNQMEESDTKLDHLTFTAVLNACSHAGMVELGESLFRKmeKYRIEpeHYACMVDLLGRAGKLSEAYDLIKAMP----VEPDKFVWGALLGACRNHGNIELAEVAAEHLFE--LEpsPGSSLLLSNLYADAGRWGNAA---------------------------------------------------------------------------------------------    
 478 UniRef50_B8BI71        Putative uncharacterized pr...   384    1e-105  1   8.4%   7:599    93:708     ------YSHAGDISTAVALFDGMPDPDVVSWNALVSGYCQRGMFQESVDLFVEM--ARRGVSPDRTTFAVLLKSCSALEELSLGVQVHALAVKTGLEIDVRTGSALVDMYGKCRSLDDALCFFYGmeRNWVSWGSAIAGCVQnyVRGLELFIEMQRLGLGVSQPSYASAFRSCAAMslNTGRQLHAHAIKNKFSSDRVVGTAIVDVYAKANSLTDARRAFFgmIRSSIRFDVVSLssVLDLYGKCKALMEAYLIFQGMKQK----DSVSWNAIIAALEQNGHYDDTILHFNEM----------------LRFDAFVASTVVDMYCKCGIIDEAQKLHDRIGGQQ------VVSWNAILSGFSLNKESEAAQKFFSEMLDMGLKPDHFTFATVLDTCANLATIELGKQIHGQIIKQEMLDDEysTLVDMYAKCGDMPDSLLVFEKVEKR-----DFVSWNAMICGYALHGLGVEALRMFERMQKENVVPNHATFVAVLRACSHVGLFDDGCRYFHLMTTHYKLEPQLEHFACMVDILGRSKGPQEAVKF----INSMPFQADAVIWKTLLSICKIRQDVepDDSSVYILLSNVYAESGKW----    
 479 UniRef50_Q53L25        Os11g0256100 protein n=4 Ta...   384    1e-105  1   9.1%  13:531    72:572     ------------LAYARRVFDAAPSRDAYMWNTLLRAHAHSrhARDALALYARMRAAG--VAPDHYTYPIVLPACAAARAPRLGRAAHGDAVRFALARDGFVSSALISMYSQEGEVRDAERVFAERdrTVVSWTAMVAGYAQnfGEAVTLFSDMVAAGVLPNEITLISFLPClgQEWLPVGEMVHGFVVKLGFDANVPMVNALITMYGRCGSAPVARVLFNGMSSR----TLVSWNTMVAMYEQHGDGVQAIKFFRRMLTEKVGFDCVTLVSVLSACARSGALGTGKWVHELARS------------HGLDTDARIGNVLVDMYAKCGEIAYAKEVFHSHGLRGH----SVVSWSAMINAYANHGEAEEALKLFSLMRNEGVRPNSFTFTAVLMACCHSGLVDEGLKHFNSitDYQMSpeHYACMVDMLGRAGRLVEAYGIIRGMPIR----PDKCVWGAFLGGCRLHGMVELAEYVAKELFDSGS--NDVtyVLMANMYFEAGMLEDAERIRGVMKEMELKKTAGHSLVC------------------------------------------------------------------------    
 480 UniRef50_Q6EPZ3        Os09g0417500 protein n=3 Ta...   383    1e-105  1  12.6%  33:538   111:610     --------------------------------VLVIALSQMALPDEALSVFGRLRELPALP-----ACNAILDGLVKAHMLARVWELFDEMLGRGMVPSVVTYNTLINACRHQGDVAKAWEVWDQmdPNVVTYTTMICALCEEDcdAEGLFLEMKEAGMRPNLYTYNALMSsfKRDNIKHALVFYYDLLKCGLVPNDVIFTTLIDGLCQANRITEAKNIFLDMPRYEVAPTVPVYNSLIHGAFRSGYAQEALAFFQEIIRKGLRPDEFTCSIVVRGLCDGGQMQVATRFLEVMQQS------------GIALNAAAYNVLIDEYCKSGNLDEALVTCTRMSEVG--VEPNVVTYSSLIDGHSKNGEMEIAMAIYTEMVAKGVEPNVVTYTALIHGHAKNGDMDAAFWLQKEMEEKGIYSntVSVLVDGLCRENRVQDAVRFIMEYseKNPSIANSVTYMTLIYGLYIDGQYNEACHFFSCMRDSGMVPDRFTYTLVIRGLCMLGYVLNAMMLYADMVKVGVKPTKCAMV--CPDIWSR-----------------------------------------------------------------    
 481 UniRef50_Q9FJY7        Pentatricopeptide repeat-co...   383    1e-105  1  10.4%  72:580    18:530     -----------------------------------------------------------------------MSCLQRCSKQEELKQIHARMLKTGLMQDSYAITKFLSFctSSDFLPYAQIVFDGFDrdTFLWNLMIRGFSCspERSLLLYQRMLCSSAPHNAYTFPSLLKACSNLsfEETTQIHAQITKLGYENDVYAVNSLINSYAVTGNFKLAHLLFDRIPE----PDDVSWNSVIKGYVKAGKMDIALTLFRKMAEK----NAISWTTMISGYVQADMNKEALQLFHEMQNSdgKWIHSYLNKTRIRMDSVLGCVLIDMYAKCGEMEEALEVFKNIKKK------SVQAWTALISGYAYHGHGREAISKFMEMQKMGIKPNVITFTAVLTACSYTGLVEEGKLIFYSMEryNLKpeHYGCIVDLLGRAGLLDEAKRFIQEMPLK----PNAVIWGALLKACRIHKNIELGEEIGEILIA--IDPYHGgyVHKANIHAMDKKWDKAAETRRLMKEQGVAKVP----------GCSTISLEGTTHEFLAGDRSHPEIEKIQSKWRIMRRKLEENGYVP-----------------------    
 482 UniRef50_Q2R047        Salt-inducible protein, put...   383    1e-105  1  11.9%  10:528   359:921     ---------DEALQVWAEMTEMGVKPDARGYSSFLIGLCDCGKYDLAYVILQEINREKVPVE--AMAYNMVMDGLCKEMRLDEAEKLLENKARQGSNPDVYGYslIRSYCKMGNLINAVDH-YEAMVSHGIETNCHIVSYlgMTSEAIAYFLKFKDSGLHLDKVIYNIAMDTYCkgNMNEAVKLLNEMKYGGLTPDKIHYTCLISGYCLKGEMQNAQQVFEEMLKANIEPDIVTYNILASGFCKSGLVMEVFDLLDRMADQGLEPNSLTYGIAIVGFCRGGNLSEAEVLFNVVEEKGIDHihAYMlrQGNLVDHFSCSKLINDLCRVGNVQGASNVCKIMLE--HDVVPDVISYSKLISIYCQNGDMDKAHLWFHDMVQRGLSVDVIVYTILMNGYCKAGRLQEACQLFVQMTNLGIKPDvyTVLLDGHLkqGWEGIAKelLSSMKDM-QIEPDVPCYTVLIDGKCKAEYLVEARELFDEMLQKGLTPDAYAYTALINGYCSQGEISKAEDLLQEMIDKGIEPDELTF---------------------------------------------------------------------------    
 483 UniRef50_D1HYZ4        Whole genome shotgun sequen...   383    1e-104  1   9.8%   9:582    16:550     --------KSLKSIHARLLIESSVASSEFVINKLLRLYSRFGATDYAHKVFDEITQ------PNAYLWTSLIHGYVENRQYDEAFSLFIQMRREPivLNFTISSVLKALArkGGQAVYGFAFDEMCEKDIVSWNMMISGYGNnvDIARKFFDRMPER----NVVSWTSMICGYVkgDMAEAQVLFDSMPVK----DLASWNVMVSGYMDIGDynGARCLFDQMPMK----NLVTWSTMIGGYARNGQPLKALELFERFKEQDIKPDETFILGIISACSQLGIIDAAESIIH------------NYVGPSLLSDLRVFTSLIDMYAKCGSIEKALQMFEMAHPK------DLLCYSTMIAALANHGLGRDAIFLFDKMQRANIKPDSVTFLGVLTACNHGGLVDEGRKYFKQmeEFGIQPshYACVVDLLGRVGCLEEAYNLIRNMP----IAPHSVVWGALLAACRVHCNVQLAEVAAAELFK--IEPdsGNYILLSNIYAAAGRWGSVAKVRAKIREHRVRK----------NRGSSWIELSHVVHEFVMGDMSHTDSDSISLILYLLCEDMKLSGYLIDS---------------------    
 484 UniRef50_UPI0001985507 PREDICTED: hypothetical pro...   383    1e-104  1  11.3%  69:571    72:587     --------------------------------------------------------------------NSVLALLERCIHLNHLKQLQAFLITLGHatHFYAFKLLRFCTLALSNLSYARFIFDHVepNVYLYTAMITAYASHSDhaLLLYRNMVRRRRppNHFIYPHVLKSCTQvgPGSARMVHCQVLRSGFEQYPVVQTALLDAYLRfsDVESARLLFDEMTER----NVVSWTAMISGYTRLGQIGNAVLLFEEMPER----DVPSWNALIAGYTQNGLFMEALSLFRRMIarLGKWIHGYVYRNGLGLDSFVSNALVDMYGKCGCLKEARRVFDRTLER------SLTSWNSMINCLALHGQSQNAISVFEEMMtsGVKPDEVTFIGLLNACTHGGLVEKGWLYFELmqNYGIEpeHYGCLVDLLGRAGQFEEAMEVVRGMR----IEPDEVIWGSLLNGCKIHGHTDLAEFSIKKLID--MDPNNGGYGIMlnIYGELGKWDEVRKVRKVLKEQNAHKTP----------GCSWIEIDNQVHQFYSVDKTHPRTEEIYNTLESLIS--------------------------------    
 485 UniRef50_Q6YSW0        Os07g0299800 protein n=5 Ta...   383    1e-104  1  10.5%  86:599   114:599     -------------------------------------------------------------------------------------QLHLLALRSGLFPSDPYSalLHMYHHCSRPMDARRAFDEIPDPNPVIVTAMASGYVRnvYHSLELFRAMIASDSASVVDEAAALVAFSASARVPdtASLHALIAKIGFERNAGVVNTMLDSYAKGgdLEVARKVFDTMER-----DVVSWNSMIALYAQNGMSAEAIGLYSKMLngGIKCNAVALSAVLLACAHAGAIQTGKRIHNQ------------VVRMGLEENVYVGTSIVDMYSKCGRVEMASRAFRKIKEK------NILSWSAMITGYGMHGRGQEALEIFTEMKRSGLRPNYITFISVLAACSHAGLLDEGRYWYNAMkeFGIEaeHYGCMVDLLGRAGCLDEAYSLIKEMKVK----PDAAIWGALLSACRIHKNVELAEMSVKRLFELDAS-NSGYYVLLSNIYAEARMWKDVERIRLLVKTRRIEKPPGY----------SSFELKGKIYLFYVGDKSHPQHIEIYSYLEKLLERMQEAGYVPNT----GSVLHDLDEEEKE----    
 486 UniRef50_Q0DFM9        Os05g0581300 protein n=6 Ta...   382    1e-104  1  11.1%   7:581   118:671     ------YAKAGCLGLALKVFDEMPHKNVVSWTTLVASLTRAGRRHEALRRFSEMRASGVHC--DSYAYAAALTACADAGLLSRGREVHAFCAKLGLDSTPYVANTLATLYARCSDVDRALAAVSRmtRDVAAWTTVISAYvgRAKEAIEAFVRMLREESspNEYTYAAVIAACAdaWVCLGEQLHAQAARKGFACARSVANSLVTLYTRagCLSAADAVFRE----SVVKDVVSWSAIISGYAQEGLAEDAFALFREMRhsGcrPNEFTLASLLSVCATAASLDAGRQL------------HTLAVAAGLEHHAMIRSALIDMYGKSGSMLDADIVFS------HRVKDDVVSWTAMIVGYAEHGHSKKALELFQEMCHVGLKPDHVTFIGVLNACCHAGEVELGLRYLNEMNqyGLYpeHYGCVVDLLGRAGRINEAEELIGKIAAN---ERDGVVWTSLLRACAARGEEETGKKAAERAMEA--EpgAGAHVAMANLYASKGQWHEAAQERHMMKQKGV-------VKGAGWSSITVGGEGRRVGVFVASDRTHPQDSAIYRMLELIYFGTGMARYVPD----------------------    
 487 UniRef50_Q0DC25        Os06g0493800 protein n=3 Ta...   382    1e-104  1   8.1%  11:578    51:595     ----------GHLADARGVFDEMPRRNVVSWSVLIAASSRLGVLGDALSLFAGMLRGGERDRPNSFTVAALVAGCARAKDAVAGEQVHASAVKLGVddESVAGTLVDMYAKCGRVGSSWRAFVLTPQRSVLSWTSMIACLVNHGDsaIVLFKKMLVLKVWPTNATFSCILKVF---DVPegKQVHGCLVKMGTEVDPALGTALLAMYGRCGGMDEITRL-----ACRIRHDAFSRTSLLTAYARNGCNMEAVRVFRDMLMGHMPIDQSAITSLLQVCSSLGQLRVVKEI------------HCYALKNFFRLDTLLLNAIVTVYGKCGDIASSEIVFNTLENK------DTISWTALLTCYVQNDLSQEALFFFREMVRKGLESSIFCITSVLRACSATSSLSCGWQIHSRVVKLGVDDDTsnALVTMYAKCGVVQVALKIFNSMRNRG-----IISWNALITSFSQHGNEVAAIQLFDMMQEEMVCPDDYTFVGLLSSCSRMGLVAEGCEYFKQMKTKYNLEPKMEHYTCMVDLFARAGRFSDAMKF----IDAMPCQPDQLVWEALLASCRVHGNL-------------------------    
 488 UniRef50_Q9FHR3        Putative pentatricopeptide ...   382    1e-104  1   8.3%  19:523    65:541     ------------------VFERVPSPGTYLWNHLIKGYSNKFLFFETVSIL--MRMMRTGlrPDEYTFPLVMKVCSNNGQVRVGSSVHGLVLRIGFDKDVVVGTSFVDFYGKCKDLFSARKVFGEmeRNAVSWTALVVAYVKSgeEAKSMFDLMPERNLG----SWNALVDGLVkgDLVNAKKLFDEMPKR----DIISYTSMIDGYAKGGDMVSARDLFEE--ARGV--DVRAWSALILGYAQNGQPNEAFKVFSEMCAKNVKPDEFIMVGLMSACSQMGCFELCEKVDSYLHQRMNKFSSHYVVP-----------ALIDMNAKCGHMDRAAKLFEEMPQR------DLVSYCSMMEGMAIHGCGSEAIRLFEKMVDEGIVPDEVAFTVILKVCGQSRLVEEGLRYFELmkKYSisPDHYSCIVNLLSRTGKLKEAYELIKSMP----FEAHASAWGSLLGGCSLHGNTEIAEVVARHLFE--LEpsAGSYVLLSNIYAALDRWTDVAHLRDKMNENGITK--------------------------------------------------------------------------------    
 489 UniRef50_Q10R82        Pentatricopeptide, putative...   382    1e-104  1   8.8%  11:523    26:483     ----------GAIAHAYLVFDQIPRPDRFMYNTLIRGAAHTAAPRDAVSLYTRmrGGGGGVRPDKLTFPFVLRACTAMGAGDTGVQVHAHVVKAGCESDAFVKNALIGMHASCGNLGIAAALFD-----------------GRARE---------------DAVAWSAMITGCArgDIGAARDLFDECPVK----DLVSWNVMITAYAKRGDMALARELFDQVPER----DVVSWNVMISGYVRCGSHLHALELFEQMQRMGEKPDIVTMLSLLSACADSGDLDVGQRLH--------SSLSDMFSRNGFP--VVLGNALIDMYAKCGSMKSAHEVFWSMRDK------DVSTWNSIVGGLALHGHVLESIDMFEKMLKGKVRPDEITFVAVLIACSHGGMVDKGREFFNLMqkYRVEPnhYGCMVDMLGRAGLLKEAFEFIDTMK----CEPNSVIWRTLLSACRVHGEIELAKHANRQLLKAR---NDEsyVLLSNIYASVGEWFGSEKMRKLMDDSGVNK--------------------------------------------------------------------------------    
 490 UniRef50_Q9C9H9        Pentatricopeptide repeat-co...   382    1e-104  1  12.3%   3:580    99:650     --ANFLINMYAKCGnaRQVFDTMPERNVVSWTALITGYVQAGNEQEG---FCLFSSMLSHCFPNEFTLSSVLTSC----RYEPGKQVHGLALKLGLHCSIYVANAVISMYGRCHDGAAAYevFEAIkkNLVTWNSMIAacCNLGKKAIGVFMRMHSDGVGFDRAtlNICSSLYKSSDlkCCLQLHSLTVKSGLVTQTEVATALIKVYSelEDYTDCYKLFMEMS---HCRDIVAWNGIITAFAVY-DPERAIHLFGQLRQEKLSPDWYTFSSVLKACAGL------------VTARHALSIHAQVIKGGFLADTVLNNSLIHAYAKCGSLDLCMRVFDDMDSR------DVVSWNSMLKAYSLHGQVDSILPVFQKM---DINPDSATFIALLSACSHAGRVEEGLRIFRSMFEKppQLNHYACVIDMLSRAERFAEAEEVIKQMP----MDPDAVVWIALLGSCRKHGNTRLGKLAADKLKEL-VEpnSMSYIQMSNIYNAEGSFNEANLSIKEMETWRVRKEPD----------LSWTEIGNKVHEFASGGRHRPDKEAVYRELKRLISWLKEMGYVP-----------------------    
 491 UniRef50_Q75L17        Os05g0112900 protein n=3 Ta...   381    1e-104  1   9.7%   9:574   135:651     --------KAGQVDRARRLFDGMAVKNVVAWTCMVSGYCRAGLVDEARRLFDLM------PYRNVFSWTTMVQGYAHNGMLREAREMFDKMPERNVVAWTV---MVKAYVDNGCIQEALELFNRMPQMNSYSWNAMATGlaGKVDDAVQLFDKMPHK----NVVSWTIMVTGLAQNgvSRAREFFDRMPKK----DIPAWNSMITAYTNDGQVNDAQRLFDIMPSK----NLVTWNIIIDGYSMNNLKDEALRLFLLMLRSAVSPDS---TTLISVLVVSESTMEVRQI------------HGLSTKLGYQPETNLGNTLVSMYSRSGDLSSAWLAFRRLNEK------DAITWTSMIQALANHGCAPCALQGFAQMLRCGYKPSSTTFTAVLSACNHVGLVEKGRKIFKSIShlGPTIEHYSCLVDLLGRAGYVREAKEVVDGMQRDMC---DEAILGTLLGSCMTHNEVEVGRAVGEDLVK--IDPssGGYTLLANVFASGGMWNEVASVWKIMKGSKVKKTPGF----------SQIEVNARNHVFYSRDQMHSQRTEIYEMlsRLVPQMK-----------------------------    
 492 UniRef50_Q0E0F3        Os02g0555100 protein n=3 Ta...   381    1e-104  1  11.4%   2:573    30:583     -AVLALLERAISAGDVRRlvHALLVKTALTHHTLLslYALLPTPAASLAAFDDL------PHKNAHSYNSLLAALArrGTLPDALRLLDGMPPASR-NVVSYNTVISSLARHGRESEALRVFAQLARDRGlrFTVVSAASarDARHLRELHGAVVVSGMEVTVIMANAMVDAYSkgRVEDARGVFDQMTIR----DSVSWTSMIAGYCRASMLDDAVQVFDMMPAQ----DAIAWTALISGHEQNGEEEIALELFERMTGEGVVPTPFALVSCLGACAKVGLVARGKEVHGFILRRSIGSD---------PFNVFIHNALIDMYSKCGDMVAAMAVFDRMLER------DIISWNSMVTGFSHNGQGKQSLAVFERMLKDEVQPTYVTFLAVLTACSHAGLVSDGRRILESMQDHGVEpeHYAAFIDALGRNRQLEEASEFIKGLSSKIGLG-TTGSWGALLGACHVHGNIEIAEEVAEALFQ--LEpnSGRYVMLSNIYSAAGQWDDARQVRALMKGKGLRKDQAY----------SWIEVQRAKHMFVADDTSHHEANEIYEMLDKLFHHM------------------------------    
 493 UniRef50_D1HBH2        Whole genome shotgun sequen...   381    1e-104  1  12.2%  24:603   119:794     -----------------------LRPNIATCTALLGALFQLGRVREVSDLVSWMEREEFVF--DVVFYSSWICGYFREGVLVEAIRKHKEMIEKGIAPDTVSYTILIDGFSREGYVEKAIGFLEKMKKDGLKPNLVTygFCKKgdEAYTLFKMVENLGIEVDEFMYVTLIDGFcrGDIDCVFGLLEDMEKRGISPSIVTYNSIINGLCKAGRTSEADEvfrRLEEDGVCIDLVMCNTIIKALLMVGALEDAYAFYKGMSGMDLVADSVTYCTMINGYCRVSRIEEALEIFDEFRKTSISSCYLFFVQEGFFpdLVDYSIMIDVLCKEGHLDKALDLCAFVKKKG--IALNIYAYNSVINGLCRQGCLVQAFRLFDSLEKIDLVPSEITYATLIDSLCKEGCLLDAKQLFEKMVIKGFNPNvyNSLIDGYCKFGNMEEALNLLIDLKARCIK-PDEFTVSALINGYCHKGDMEGALGFFFEFKKKDILPDFLGFMYLVRGLCAKGRMEEARGILREMLQTRSVliETESVESFIISLCEQGSIQEAVTVLNEviGRRCRPQNRAEKEEKIYE--GKGSRVPD-FESYYSLIASLCSRGELLEAN    
 494 UniRef50_C5Z5U1        Putative uncharacterized pr...   381    1e-104  1  10.2%   2:510   154:638     -ALAKLYFKLSKVDDARKVFDAVPSPDTILWNTLLAGLPGSV----ALEAFVRMVEVGR-VRPDSTTLASSLRAAAEASHVAMGRCVHGYGVKCGLAEHEHVVTGLMSLYSKCGDMDCARSLFDRMepDLVAYNALISGYSVNgeSSVELFKELAASDWRPNSSTLVAVIPVYSPFGHelARCLHAFVVKARLDADALVSTALTTLYCRLNDMESARSIFDAMPEKTME----SWNAMISGYAQNGLTEMAVELFQLMQELNVQPNPTTISSTLSACAQLGALSLGTWV------------HRIIAKENLELNVYVMTALIDMYAKCGSIAEARSIFDRMDNK------NVVSWNAMISGYGLHGRGAEALKLYKSMLDACILPTSSTFLSVLYACSHGGLVDEGQKVFRVMteYRISpeHCTCMVDLLGRAGKLNEALDLISEFPQSAIGP---GVWGALLSACMVHKNSDLAKLASQKLFELDSE-NAGYYVLLSNLYTSKKHYSEAA---------------------------------------------------------------------------------------------    
 495 UniRef50_D1H5E8        Whole genome shotgun sequen...   381    1e-104  1   7.4%  26:527   113:552     -------------------------PNTFSWNVAIRGFLDSENPREAVVLYKRVLQCDGT-KPDNYTYPLLFKACARLSLIRMGSEILGHVLHLGFDSDIFVSNAVIHLLVSCGDLDGAR--------------------------------KMFDKSCVRDLVSWNSMINGYC-GNLESARKLFDSMTNK----TMVSWTTMVVGYAQSGLLDMAWKLFDEMPDK----DVVPWNAMIGGYVHANRGKEALALFNEMQAMNINPDEVTMVSCLSACSQLGALDVGIWIHHYIE------------KHELSLNVALGTALIDMYAKCGKITKAIQVFQELPGR------NSLTWTAIISGLALHGNAHGAIAYFSEMIDNSVMPDEVTFLGLLSACCHGGLVEEGRKYFSQMSSknLSPkhYSCMVDLLGRAGLLEEAEELIKSMP----IEADAVVWGALFFACRIHGNVLMGERAASKLLQMDPH-DSGIYVLLANMYGEAEMWKEAGKARKLMRQRGVEKTPDS----------------------------------------------------------------------------    
 496 UniRef50_C5YZH4        Putative uncharacterized pr...   381    1e-104  1  11.4%   2:596    66:632     -AAVRRLAAAGDVdeLQEMRLRGVPCTEGALVAAVGAFARAGAPDRALKTfrAVHDLGCARPTAPRLYNHLIDALLRENMVGAVALVCGNMRKDGVQPNVFTYNLLVRALCQNHRVGAARKMLDEMATKGCPPDDVTYGtlGRVDEATEVLSAA-----PPVAASYNAVILALCRerMQEVFAVVGDMVGRGLQPNVITYTTIVNAFCKAGELRMACAILARMVITGCTPNVATFTALVGGLFNDGRVHDALDMWKCMVAEGWAPSTVSYNVLIRGLCSVGDLKGASSVLNDMEQ------------HGCFPNARTYSTLIDGFSKAGDLDGAISIWNDMTRSGC--KPNVVVYTNMVGVFCKKLMFNQAESLIDKMLVENCPPNTVTFNTLIRSLCNCRRVGRALGVFHEMRRHGCPPntYNELLHGLFREGNCGDALQMVIEMQNHGI-ELSLVTYNTVVSGLCQMRMGREAMFFVGRMIVRGIQPDAFTFTAIIHAYCKEGEVRMAAWILGAM----------NVVNCGRNILVYTILMAELC------------NQDKLEDAMVYLLKMLYEGIYPNTV-TWNVLVRGVFRN-------    
 497 UniRef50_Q9T0D6        Pentatricopeptide repeat-co...   380    1e-104  1  13.4%  68:528    97:551     -------------------------------------------------------------------YEVIINSYVQSQSLNLSISYFNEMVDNGFVPGSNCFNYLLTFVVGSSSFNQWWSFFNENKsdVYSFGILIKGCcgEIEKSFDLLIELTEFGFSPNVVIYTTLIDGCCkgEIEKAKDLFFEMGKLGLVANERTYTVLINGLFKNGVKKQGFEMYEKMQEDGVFPNLYTYNCVMNQLCKDGRTKDAFQVFDEMRERGVSCNIVTYNTLIGGLCREMKLNEANKVVDQMKSD------------GINPNLITYNTLIDGFCGVGKLGKALSLCRDLKSRG--LSPSLVTYNILVSGFCRKGDTSGAAKMVKEMEERGIKPSKVTYTILIDTFARSDNMEKAIQLRLSMEELGLVPdtYSVLIHGFCIKGQMNEASRLFKSMVEKNC-EPNEVIYNTMILGYCKEGSSYRALKLLKEMEEKELAPNVASYRYMIEVLCKERKSKEAERLVEKMIDSGIDPSTSIL---------------------------------------------------------------------------    
 498 UniRef50_Q0WQW5        Pentatricopeptide repeat-co...   380    1e-104  1  11.1%  70:581    53:548     ---------------------------------------------------------------------SLAETCSDMSQLKQLHAFTLRTTYPEEPATLFLYGKILQLSSSFSDVNYAFRVFDSIehSSFMWNTLIRACAHDveEAFMLYRKMLERgsSPDKHTFPFVLKACAyfGFSEGKQVHCQIVKHGFGGDVYVNNGLIHLYGSCGCLDLARKVFDEMPER----SLVSWNSMIDALVRFGEYDSALQLFREMQRS-FEPDGYTMQSVLSACAGLGSLSLGTWAHAFLLRK---------CDVDVAMDVLVKNSLIEMYCKCGSLRMAEQVFQGMQKR------DLASWNAMILGFATHGRAEEAMNFFDRmkRENVRPNSVTFVGLLIACNHRGFVNKGRQYFDMMVryCIEpeHYGCIVDLIARAGYITEAIDMVMSMPMK----PDAVIWRSLLDACCKKgsVELSEEIARNikEDNESSnsGAYVLLSRVYASASRWNDVGIVRKLMSEHGIRKEP----------GCSSIEINGISHEFFAGDTSHPQTKQIYQQLKVIDDRLRSIGYLPD----------------------    
 499 UniRef50_B9MU51        Predicted protein n=2 Tax=M...   380    1e-104  1  10.8%  12:529   165:685     -----------KLEAVRSLFdsGPSPNVYTYTIMINFYCKERhmEQASLILEEMEEKGE--NPTVVTYGVYIHGLCRVGSIEDAWNKIQDLRSSNQPLNIYCYNALIQGFCQKGRPDEALKLLEEMKDEGISPDIYSYSIlgDIESGMNLIQEMEHCNKKPPLVCCTSVLMGLrkGLVNDCLNFFHELSAKGYKHDLISYSTLIHGFLKGHNVKSANNLVHEMRKNGLVPDYIIYISLIREYCRKGCLKEALKNFYTMLQDGLQPDIITCNHIVDQYCSRGQFEEALIYINQMK------------DQNILPNSYTYSVIINWLCKYQAVEKAWEVLPVMFK--DNIFPSVIHYTTIMDGYAKqkNPMKAWKLYRKMPKLGCKPDNVTLTVLVDMFSKRGKMSKALNLFKEMAEEGLSRdaFTAIIDGYCRVGNVKRAWSMYKKMK-RNNVTPNVKTYTCLVDGFCKLKRLDMATMLIDDMKRNSVTPDVKTYTALIAGYQRIENIDRAYEVFDEMKKKGTLPDHIAYL--------------------------------------------------------------------------    
 500 UniRef50_Q9S7Q2        Pentatricopeptide repeat-co...   380    1e-103  1  13.0%  11:602   123:687     ----------RSLRLFKYMQRQicKPNEHIYTIMISLLGREGLLDKCLEVFDEMPSQGVS--RSVFSYTALINAYGRNGRYETSLELLDRMKNEKISPSILTYNTVINACARGgdWEGLLGLFAEMrpDIVTYNTLLSACAIRgdEAEMVFRTMNDGGIVPDLTTYSHLVETFGKLrlEKVCDLLGEMASGGSLPDITSYNVLLEAYAKSGSIKEAMGVFHQMQAAGCTPNANTYSVLLNLFGQSGRYDDVRQLFLEMKSSNTDPDAATYNILIEVFGEGGYFKEVVTLFHDMVEEN------------IEPDMETYEGIIFACGKGGLHEDARKILQYMTAN--DIVPSSKAYTGVIEAFGQAALYEEALVAFNTMHEVGSNPSIETFHSLLYSFARGGLVKESEAILSRLVDSGIprDTFNAQIEAYKQGGKFEEAVKTYVDM-EKSRCDPDERTLEAVLSVYSFARLVDECREQFEEMKASDILPSIMCYCMMLAVYGKTERWDDVNELLEEMLSN---------------------RVSNIHQVIGQMIKGDYDDDSNWQIVEYVLDKLNSEGCGLG-IRFYNALLDALWWLGQKERA-    
 501 UniRef50_D1HZ35        Whole genome shotgun sequen...   380    1e-103  1   7.5%   7:569    73:600     ------YSKCCCFHSARLVFDQMPIKSLVSWNSIISAHCRDFHIDQSFGILKQMQLLGL--ELSSATFTGFLASCS----LPQGLSIHGYITKLGLDLHLPLANSIMSMYIRLNQIDGALSVFYTLhkSIVSWTIILGGyaGDVAKVFAVFNQMRCQCVGPDSIVFVNLISCCksGNLLLAMLVHSLLLKSGFDHKDPIDNLLVAMYAKCKDLVSARRVFDAVHEK----SVFLWTSMISGYAQFGYPNEALHLFNMLLRTASRPNELTLATVLSACAEMGSLRMGEEI------------EQYILLNGLGSDLRVQTSLIHMFCKCGSIKKAQALFERIPNK------DLAVWSAMINGYAVHGMGKEALNLFHKMQNegIKPDAIVYTSVLLACSHSGLIEDGLKYFRSMQKdgIEPshYSCLVDLLGRAGYVELALRTIQEMPVLVQ----ARVWAPFLSACYTHHNLELGEFAAKNLFD--LEpsTGNFVLMTNLYTSMGKWKEAAKARSIINARGLVKEP----------GWSQIEIDGAVHVLAAEGQSHLESIDIHELCPEM----------------------------------    
 502 UniRef50_C5Z7W6        Putative uncharacterized pr...   380    1e-103  1  10.2%   5:526   103:593     ----KLLARFSSPADAASFLRDspAPDVSTFNSLLSALGRAGNLRGMTELFTSMRE--ASVKPDIVTYGILLNRLCKAGLVGDALKVLDRMSSPgcADIVILNTVVDGLCKIGRLQDAIVFVDERMR---------------------------QVHGCAPNTVTYNCLADACCrgDIDMACQIVERMEKEGVTPNVITMNTIVGGLCRVGRVGAALDFFRelEARG---NAVTYSTLVSAFLHCNNVALAMELFHEMADQGYPPDAVMYFTMISGLTQAGRLVDACNIAASMK------------KAGFKLDRKAYNILIGGFCRKKRLHEAHQLFEEM--KGAGLQPDVYTYNTLLSCLCKAGDFSAVDELLGNMIDDGCQPSVVTFGTLVHGYCKAGKTDEALKIFRSMGEARIQPNtyNTLIDFLCKSREVDVAIKLFDEMRENN-VPANVTTYNALLKGLQDKNMPEKAFELMDQMKEERCTPDYVTMDVLMEWLPEIGETERLKCFMEQRDEKGGAKLTM-----------------------------------------------------------------------------    
 503 UniRef50_Q9SR01        Pentatricopeptide repeat-co...   379    1e-103  1   9.9%  11:574    82:647     ----------ENLDLAKLLFLNfpNPNVFVYNTMISAVSSSKN-----ECFGLYSSmrHRVSPDRQTFLYLMKA---SSFLSEVKQIHCHIIVSGCLSlnYLWNSLVKFYMELGNFGVAEKVFARmhPDVSSFNVMIVGYAKQglEALKLYFKMVSDGIEPDEYTVLSLLVCCGhsDIRLGKGVHGWIERRgySSNLILSNALLDMYFKCKESGLAKRAFDAMKKK----DMRSWNTMVVGFVRLGDMEAAQAVFDQMPKR----DLVSWNSLLFGYSKKgdQRTVRELFYEmeLSHGRWVHGLVIRLQLKGDAFLSSALIDMYCKCGIIERAFMVFKTATEK------DVALWTSMITGLAFHGNGQQALQLFGRMQEEGVTPNNVTLLAVLTACSHSGLVEEGLHVFNHMKDkgFDPehYGSLVDLLCRAGRVEEAKDIvkKMPMR----PSQSMWGSILSACRGGEDIETAELALTELLK--LEPeeGGYVLLSNIYATVGRWGYSDKTREAMENRGVKKTA---------GYSSVVGVEGLHRFVAAEKQNHPRWTEIKRILQHLYNEMK-----------------------------    
 504 UniRef50_UPI0001986271 PREDICTED: hypothetical pro...   379    1e-103  1  12.1%  27:521    18:492     --------------------------DIMIWNGMIRGYAYNGPVDRCISMFDEM--LQRGLKPNNFTYPYVLSSCSEMGCFGRGRKVHAQIMKSGFESNFIVasLLNMYIKMSDSFedARMIFNGivKPVELWNRMISKYvgDVESARQLFDNMPER----DVISWNSMISGYArgDVANARYLFEQMPEK----NVVTWTSMIGAYSGSGDLETAACLFEKMPCR----NAVSWNAMISSYTQNGKYEKSLDLFVKMQLEGVDSDGFTFVSVLSACSNLGALEFGKWVHS-------------LIKDWSQLGVIVGTALMEMYAKCGDVNRAFAVFVKLGNK------DVFCWNVMIKSLAIHGRTEDAVKLFFLMQKIGLKPNGYTFSSALFACSHGGLVEVGREIFYGMERdgVCpeHFGCLIDLLSRNGRLEEAEALVREMP----FEPDIAIWGALLGGCRVKSDLKLAEEVVER--AGGLEAnsGVYVLLSNIYASAGQWPEALSAREKMEEKKV----------------------------------------------------------------------------------    
 505 UniRef50_Q9LMY5        Putative pentatricopeptide ...   379    1e-103  1   9.7%  26:581   234:798     -------------------------PSVVSFNSIMSGYCKLGFVDMAKSFFCTV--LKCGLVPSVYSHNILINGLCLVGSIAEALELASDMNKHGVEPDSVTYNILAKGFHLLGMISGAWEVIRDMLDKGLSPDVITYTIlgNIDMGLVLLKDMLSRGFELNSIicSVMLSGLCkgRIDEALSLFNQMKADGLSPDLVAYSIVIHGLCKLGKFDMALWLYDEMCDKRILPNSRTHGALLLGLCQKGMLLEARSLLDSLISSGETLDIVLYNIVIDGYAKSGCIEEALELF------------KVVIETGITPSVATFNSLIYGYCKTQNIAEARKILDVI--KLYGLAPSVVSYTTLMDAYANCGNTKSIDELRREMKAEGIPPTNVTYSVIFKGLCreRIFEKCKQGLRDMESEGIPPdtYNTIIQYLCRVKHLSGAFVFLEIMKSRNL-DASSATYNILIDSLCVYGYIRKADSFIYSLQEQNVSLSKFAYTTLIKAHCVKGDPEMAVKLFHQLLHRGFNV-SIRDYSAVINRLCRRHLVNESKFFFCLMLSQGISPdeVMIKSDELLSWTIKWGLLPD----------------------    
 506 UniRef50_Q2HVJ3        Tetratricopeptide-like heli...   379    1e-103  1  12.1%  31:543    72:539     ------------------------------FNKLLSTLIKLKRYNAVVSLYTILQS-KPTIKPSLITFSILINAFAQLGQMGFAFSLLGNILKMGFQIDviLTTLMKGLCLKGRVLE-----------------------------AVSLLHEYVDRGFRFDEVCYGTIINGLCkgKTRDAIQMFPKMKKIRVYPNLIMYNTVIDGLCKQGLVDEACGLCTEMVENGIELDVYSYNSMIHGFCSVGRFQAAVKLLDEMVVRgvYPDVYTFNILIDGLCKLGRVSEAYNVVAVMIKR------------GWKPDIVSYNALMNGYCLSGSVGEAKQVFDKMVERTA--LPNVISYCTLINGYCKVRMVDEAMVLLTEMHNKNLVPDTVTYNCLLDGLSKSGRSLYEWDLVEAMRASGQPAdtYNVLLDDYFKHEKFDKALALFQHIIEIGI-SPNIRTYNILLDGLCKSGRLKYAKEIFQLLSAKGCQPNIRTYNIMIHGLCKEGFLDEAEALLYKMVNNNYLPNYITF-----DTIVRAILVK------------------------------------------------------------    
 507 UniRef50_Q9SII7        Pentatricopeptide repeat-co...   378    1e-103  1  11.0%   9:552    94:610     --------KCGDLCSGLREFDCMNSRDSVSWNVIVFGLLDYGFEEEGLWWFSKLRVWGF--EPNTSTLVLVIHACRSL--WFDGEKIHGYVIRSGFCGISSVQNSILCMYADSDSLSARKLFDEMSERDVISWSVVIRSYVQSKEPvlKLFKEMveAKTEPDCVTVTSVLKACTVmdIDVGRSVHGFSIRRGFDldVFVCNSLIDMYSKGFDVDSAFRVFDETTCR----NIVSWNSILAGFVHNQRYDEALEMFHLMVQEAVEVDEVTVVSLLRVC----------KFFEQ--PLPCKSIHGVIIRRGYESNEVALSSLIDAYTSCSLVDDAGTVLDSMTYK------DVVSCSTMISGLAHAGRSDEAISIFCHMRD---TPNAITVISLLNACSVSADLRTSKWAHGIAIRRSldISVGTSIVDAYAKCGAIEMARRTFDQITEK-----NIISWTVIISAYAINGLPDKALALFDEMKQKGYTPNAVTYLAALSACNHGGLVKKGLMIFKSMVEEDHKPSLQ-HYSCIVDMLSRAGEIDTAVELIKNL---------------------------------------------------    
 508 UniRef50_UPI0001A7B196 unknown protein n=1 Tax=Ara...   378    1e-103  1  12.9%  30:595    84:632     -----------------------------SFSTLlhVLAGQRRFKELQVILEQLLQEerKWESTGLVWDMLLFLSSRLRMVDDSLYILKKMKDQNlvSTQSYNSVLYHFRETDKMWDVYKEIKD---KNEHTYSTVVDGLCrkLEDAVLFLRTSEWKDIGPSVVSFNSIMSGYCKLgvDMAKSFFCTVLKCGLVPSVYSHNILINGLCLVGSIAEALELASDMNKHGVEPDSVTYNILAKGFHLLGMISGAWEVIRDMLDKGLSPDVITYTILLCGQCQLGNIDMGLVLLKDMLSR------------GFELNSIicSVMLSGLCKTGRIDEALSLFNQMKADG--LSPDLVAYSIVIHGLCKLGKFDMALWLYDEMCDKRILPNSRTHGALLLGLCQKGMLLEARSLLDSLISSglDIVLYNIVIDGYAKSGCIEEALELFKVVIETGI-TPSVATFNSLIYGYCKTQNIAEARKILDVIKLYGLAPSVVSYTTLMDAYANCGNTKSIDELRREMKAEGIPPTNVT----------YSVIFKGLCRGWKHENCNHVLRERIFEKCKQGLRDMESEGIPPDQI-TYNTIIQYLCR--------    
 509 UniRef50_Q9FIF7        Putative pentatricopeptide ...   378    1e-103  1  11.0%  57:574    15:532     --------------------------------------------------------FRRDPDSNTLRLstLISVLRSCKNIAHVPSIHAKIIRTFHDQDAFVVFELIRVCSTLDSVDYAYDVFSYVspNVYLYTAMIDGFVSSGRSavSLYHRMIHNSVLPDNYVITSVLKAC---DLKVCREIHAQVLKLGFGSSRSVGLKMMEIYGKSGELVNAKKMFDEMPDRDHVadTVCWTAMIDGLVRNKEMNKALELFREMQMENVSANEFTAVCVLSACSDLGALELGRWV------------HSFVENQRMELSNFVGNALINMYSRCGDINEARRVFRVMRDK------DVISYNTMISGLAMHGASVEAINEFRDMVNRGFRPNQVTLVALLNACSHGGLLDIGLEVFNSMKrfNVEpeHYGCIVDLLGRVGRLEEAYRFIENIP----IEPDHIMLGTLLSACKIHGNMELGEKIAKRLFESE-NPDSGTYVLLSNLYASSGKWKESTEIRESMRDSGIEKEP----------GCSTIEVDNQIHEFLVGDIAHPHKEAIYQRLQELNRILR-----------------------------    
 510 UniRef50_B9RBA6        Pentatricopeptide repeat-co...   378    1e-103  1  11.3%   6:490   150:643     -----LYSKCQFFKYARKMVDESTEPDLVSWSALISGYSQNGFGKEAISAFYEMHLLgsGIRPNEFSLSCMINACTGLEDSSQGRKIHGYLIKLAYDLDLFSANALVDMYAKVGTLEEAIRVFEEikPDIVSWNAIIAGCALrhCWALELFGKMNRSGMCPNMFTISSALKACAGMGLKegRQLHSSLLKMDIRSDSFLAVGLIDMYSKCDLMTDARLLFNLMPER----DLIVWNAAITGHSQNGEDLEAVSLFPSMHKEGVGFNQITLSTVLKSVASLQ------------VDHICSQIHALSLKSGFQFDNYVANSLIDTYGKCGRIKDATRIFQE------SPFVDLVAFTSMITAYSQDGQGEEALKLYLEMQDRKIRPDSFVCSSLLNACANLSAYEQGKQVHVHVLKFGFISDIfnSLVNMYAKCGSIDDADRAFSEIPERG-----IVSWSAMIGGFAQHGHGKEALQLFNRMLEDGIPPNH-----------------------------------------------------------------------------------------------------------------    
 511 UniRef50_Q6YW98        Os09g0110200 protein n=4 Ta...   378    1e-103  1  11.5%  13:578   122:707     ------------FDALADARAAGLPASSSAFAALVAAHSSAGRHADAVQAFSRMDEFQS--RPTAFVYNTILKALVDSGVILLALALYNRMVAAGCAPNRATYNVLMDGLCKQGMAGDALKMFDEMLDRGIMPNVKIycNAGKIDEAVQLLGSMKDKGCLPDEVTYNAFLSGLCkgRVNEAFQRLVMLQDGGFALGLKGYSCLIDGLFQARRFDEGFGYYKTMLERNISPDVVLYTIMIRGCAEAGRIEDALSFLDVMKKKGFVPDTFCYNTVLKVLCDHGDLERAHTLRSEMLQ------------NNLVLDSTTQTIMICGLCKRGLVDEAMQIFDEMGEHGCD--PTVMTYNALIDGFYREGRLEEARMLFHKM-EMGNNpdSESLRKLVHDMCQSGQVLKAYKLLRSIIDSGVVPdtYNTLINGLCKARNLDGAVRLFKELQLKGI-SPDEITYGTLIDGLLRAHRENDAMMLFQNILQSGSSPSLSIYNSMMRSLCRMKKLSQAINLWLDYLPKKYNFPVESEVLANAHKEIEDGSLDDGVRELIKIDQEylCQVRRTDDALRIFHTLQEFGI-------------------------    
 512 UniRef50_B6SXY4        Pentatricopeptide repeat pr...   378    1e-103  1  11.6%  32:573    79:593     -------------------------------NRLVALYSSLPSPAAAVKAFYDL------PHKNAHSYNTLLAALLRRPdlPDAFRLFDTMPADSR-NLVSYNTLMSSLAHHGRQVEALRVVAraMDRFTVVSVATACAGIGAARPLREMHGAVVVSGMEFTVIMANAMLNAYSKArvEDARHLFDQVSIR----DNVTWTSMISGYCQVKKLHEAVQVFDMMPDK----DRIAWTALISGHEQNGEEDTALELFEQMLAKGVSPTPFVLVSALGACAKLGLVTRGKELHCFILRRNIGS---------VPFNIFIHNALVDMYSKCGDMMAAMAVFDWMPER------DFISWNSMVTGFSHNGLGKQSLAIFEEMLVAGVRPTHVTFLAVLTACSHSGLVSNGRLVLESMEDHGVEpeHYAAFIDALGRNHQLEEATEFIKDLPSR-IGPGTAGSWGALLGACRLHGNIELAEEVAEFLFR--LEpnSGRYVMMSNIYAAAGQWDDARRVRGLMKEKGLKKDQAY----------SWIEVRSAKHVFVADDISHCEADKIYEMLGKLLDHM------------------------------    
 513 UniRef50_Q9LT48        Pentatricopeptide repeat-co...   378    1e-103  1  10.5%   6:520    54:553     -----LIDLygDVKHARKLFDRISKRDVVSWTAMISRFSRCGYHPDALLLFKEMH--REDVKANQFTYGSVLKSCKDLGCLKEGMQIHGSVEKGNCAGNLIVRSALLSLYARCGKMEEARLQFDSmeRDLVSWNAMIDGYTANACasFSLFQLMLTEGKKPDCFTFGSLLRASIVVklEIVSELHGLAIKLGFGRSSALIRSLVNAYVKCGSLANAWKLHEGTKKR----DLLSCTALITGFsqNNCTSDAFDIFKDMIRMKTKMDEVVVSSMLKICTTIASVTIGRQIHGFALKSSQ-----------IRFDVALGNSLIDMYAKSGEIEDAVLAFEEMKEK------DVRSWTSLIAGYGRHGNFEKAIDLYNRMEHERIKPNDVTFLSLLSACSHTGQTELGWKIYDTMikHGIEaeHLSCIIDMLARSGYLEEAYALIRS--KEGIVSLSSSTWGAFLDACRRHGNVQLSKVAATQLLS--MEPrpVNYINLASVYAANGAWDNALNTRKLMKESG-----------------------------------------------------------------------------------    
 514 UniRef50_Q9C9I3        Pentatricopeptide repeat-co...   378    1e-103  1   9.4%   6:558   155:688     -----MYTACGSVKDAQKVFDESTSSNVYSWNALLRGTVISgrYQDVLSTFTEMRELG--VDLNVYSLSNVFKSFAGASALRQGLKTHALAIKNGLFNSVFLKTSLVDMYFKCGKVGLARRVFDEieRDIVVWGAMIAGLAHNKreALGLFRTMieEKIYPNSVILTTILPVLGDVklKLGKEVHAHVLkkNYVEQPFVHSGLIDLYCKCGDMASGRRVFYGSKQR----NAISWTALMSGYAANGRFDQALRSIVWMQQEGFRPDVVTIATVLPVCAELRAIKQGKEI------------HCYALKNLFLPNVSLVTSLMVMYSKCGVPEYPIRLFDRLEQR------NVKAWTAMIDCYVENCDLRAGIEVFRLMLLSKHRPDSVTMGRVLTVCSDLKALKLGKELHGHILKKEFepFVSARIIKMYGKCGDLRSANFSFDAVAVKGSL-----TWTAIIEAYGCNELFRDAINCFEQMVSRGFTPNTFTFTAVLSICSQAGFVDEAYRFFNLMLRMYNLQPSEEHYSLVIELLNRCGRVEEAQRLAVMSSSSSLQ---------------------------------------------    
 515 UniRef50_D1HSV4        Whole genome shotgun sequen...   378    1e-103  1  11.5%  70:575     8:541     ---------------------------------------------------------------------SLLKTCIDSKYLKQGKLIHQKIVSLGLQNNITLCKSLINLYFSCHLFQSAKLVFQTIEndITLWNGLMAACTKnpEKALELFEEMKVSGFKPDSVTLTTVISSCARldLERGKEIHMELVRSGFALDGFVSSALVDMYGKCGCLEMAKEVFEQIQRK----NVVSWNSMIAGYSLKGDSKSCIELFRRMDEEGIRPTLTTLSSILMACSRSVNLQLGKFI------------HGYIIRNRVEADIFVNSSLIDLYFKCGNIGSAENVFQNMPK------TNVVSWNVMISGYVKVGSYLEALVIFTDMRKAGVKPDAITFTSvlSACSHAGLVDEGCYYFNQMieYGFKpeHYSCLIDLLGRVGRLREAYEILQRTPD---IREDVGLLSTLFSACHLHKKLDLGEQIGRLLIEKDPD-DPSTYIILSNMYASVKKWDEVRKVRLKIKELGLKKNP----------GCSWIEVGKRIHPFVVEDKSHPQADMIYECMSILASHVEK----------------------------    
 516 UniRef50_D1IYB9        Whole genome shotgun sequen...   377    1e-103  1  10.7%  34:586    26:543     ---------------------------------LIS-FCAVDDAGDLQYAQRMFDQI---PQPNKFMYNSLIRGYSNSDDPIDAVLLFRRMICSGLSPNEFTlfVLKACGCKSAYWEAVLviKLGIGSLVFVQNALIAVycGLIHCARKLFDDITDKTL-IDIIVRNALVDMYAkgNLHSAQAIFDRTQEK----NVVSWTSMISAYAQHGSIEVARQIFDQMPGK----NVVSWNSMISCYLREGQYREALDLFNKMRNSRVVPDEATLVSILAACSQLGDLVMGKKIHNYILSNKGAYG------------VTLYNSLIDMYAKCGPVVTALDIFLEMPGK------NLVSWNVIIGALALHGCGLEAIKLFEEMQADGTLPDEITLTGLLSACSHSGLVDMGLYYFDRM---GveIEHYACMVDLLGRGGLLGEAIELIGRMPMK----PDVVVWGALLGACRIHGNVEIGKQILKQLLE--LEpsGGLYVLISNIYWEAQRWEDVKKIRKLMIDRGIKKGRA----------ISSIEIDGCIYEFMVDDKRHKISSSIYAMLDQLTDHLRSAGYLCNISSVF-----------------    
 517 UniRef50_Q9SSR4        Pentatricopeptide repeat-co...   377    1e-103  1  11.7%   1:600   109:727     LARYRIFNEIEDV--LGNLRNENVKLTHEALSHVLHAYAESGSLSKAVEIYDYVVELYDSV-PDVIACNSLLSLLVKSRRLGDARKVYDEMCDRGDSVDNYSTCILVKGMCNEGKVEVGRKLIEgcIPNIVFYNTIiyCKLGDIENAYLVFKELKLKGFMPTLETFGTMINGFCkgDFVASDRLLSEVKERGLRVSVWFLNNIIDAKYRHgkVDPAESI-GWIIANDCKPDVATYNILINRLCKEGKKEVAVGFLDEASKKGLIPNNLSYAPLIQAYCKSKEYDIASKLLLQMAER------------GCKPDIVTYGILIHGLVVSGHMDDAVNMKVKLIDRG--VSPDAAIYNMLMSGLCKTGRFLPAKLLFSEMLDRNILPDAYVYATLIDGFIRSGDFDEARKVFSLSVEKGVKVdhHNAMIKGFCRSGMLDEALACMNRMNEEHLV-PDKFTYSTIIDGYVKQQDMATAIKIFRYMEKNKCKPNVVTYTSLINGFCCQGDFKMAEETFKEMQLRDLVPNVVTYtkCVPNEVTFNCLLQGFVksGKVLAEPDGSNHGQSSLFSEFFHRMKSDGWS-DHAAAYNSALVCLCVHGMVK---    
 518 UniRef50_D1HYB6        Whole genome shotgun sequen...   377    1e-103  1   8.6%   7:574    15:594     ------LAKLGRITSARRLFDEMPHKDTVAWNAMLASYSQLGLHQQALCLFHHMRIANS--RPDRFTFTATLSACAGLGELRRGMKIHAQVVVSGCQSSLPVGNSLIDMYGKcgLFDVARVVFDGMPKKviAWNIMISGycGDVELCLGLFKKMREDSLQPDQWTFSALVNALCELQEPsgYMMHGFIIKSGWdlTQVSWNAMIDAHMKIGDTHEAFLVFQLAPEK----NVVSWTSMITGYARNGHGEQALSFFVKMMENHIQPDDFTFGAVLHACSSLATLGHGKMI------------HGSIIHYGFHAYVDVGNGLVNMYAKCGDIQGSNTAFKEILGK------DLVSWNAMLFGLGMHGHATQALELYEEMVASGMKPDKVTFIGLLMTCSHSGLIEKGQALFESMVsyGleTEHVVCMVDLLGRGGYLAQARELVDEYSRTGRAETSLP--EALLGACFAHSEVRMGANLGEYLKV--FEPqeMSYVLLSNLYCVSGQWKEAEMVRKTMTDHGVKKMP----------GCSWIEVRNKVTVFVAGNHSHPYMEELCKILNFLKFEMR-----------------------------    
 519 UniRef50_Q5JNP8        Putative PPR986-12 n=3 Tax=...   377    1e-103  1   8.2%   5:577    12:549     ----SLCSR-GAVCHARALFDEMPERDVVAWTAMLSGYASNGLRREALDVFRRMVAAGA--APNEYTLSSVLTACRGPCAPAMAMPLHAVAVRRGVDrpYVVNALIDSYASLAEGVVDARRLFDAlgRTAASWTSMIAGYARWGQegLRLFKTMLKDGVELSTFACSIALHACTLvdLCLGQQLHLQCIKKALDVNLAVVNSLIDMYCTCARILDARSLFDGTPER----NLITWNTMIAGYSQC-DPLMALQLLLEMND---EPNCFTLTSITSACADLAALRCGKQV------------HGAVLRRSYSDDLQMGNALVDMYSKCGSITNAKNVFDRMGCK------DKFSWTSMIAGYGMNGYGNEAVQLFSSMIHAGVHPDHVVFLSLISSCSHAGLVDEGWNFFRSMieYNLQPnvYGSVVNLLARAGRLREALDLIDTMP----FAPDEYVWGALLGASKMHNNVEMGRLAARKITE--INPDDvnYIMLASIYAAGSKWGEYAFTRRSLRGIGSRK----------EAGISWIEVMDKMYSFTAADSSSPQVCLADEVLHILSQHMDDVG--------------------------    
 520 UniRef50_Q8S9M4        Pentatricopeptide repeat-co...   377    1e-103  1  12.7%  14:582    27:563     -------------GNLREAFQRF-RLNIFTNTSLFTPFIQSCTTRQSLPSGKQLHCLlsGFSSDKFICNHLMSMYSKLGDFPSAVAVYGRMRKKNYMSS---NILINGYVRAGDLVNARKVFDEMPDRKLTTWNAMIAGLIQfnEEGLSLFREMHGLGFSPDEYTLGSVFSGSAGLrvSIGQQIHGYTIKYGLELDLVVNSSLAHMYMRNGKLQDGEIVIRSMPVR----NLVAWNTLIMGNAQNGCPETVLYLYKMMKISGCRPNKITFVTVLSSCS------------DLAIRGQGQQIHAEAIKIGASSVVAVVSSLISMYSKCGCLGDAAKAFSEREDE------DEVMWSSMISAYGFHGQGDEAIELFNTmeQTNMEINEVAFLNLLYACSHSGLKDKGLELFDMmeKYGFKPghYTCVVDLLGRAGCLDQAEAIIRSMPIKT----DIVIWKTLLSACNIHKNAEMAQRVFKEILQ--IDpdSACYVLLANVHASAKRWRDVSEVRKSMRDKNVKK----------EAGISWFEHKGEVHQFKMGDRSQSKSKEIYSYLKELTLEMKLKGYKPDT---------------------    
 521 UniRef50_Q9LZ19        Pentatricopeptide repeat-co...   377    1e-103  1  10.4%  71:580    67:546     ----------------------------------------------------------------------ILQLCARNGAVMEAKACHGKIIRIDLEGDVTLLNVLINAYSKCGFVELARQVFDGmeRSLVSWNTMiyTRNRMESEALDIFLEMRNEGFKFSEFTISSVLSACgnCDALECKKLHCLSVKTCIDLNLYVGTALLDLYAKCGMIKDAVQVFESMQDK----SSVTWSSMVAGYVQNKNYEEALLLYRRAQRMSLEQNQFTLSSVICACSNLAALIEGKQM------------HAVICKSGFGSNVFVASSAVDMYAKCGSLRESYIIFSEVQEKNLEL------WNTIISGFAKHARPKEVMILFEKMQQDGMHPNEVTFSSLLSVCGHTGLVEEGRRFFKLMrtYGLSPnhYSCMVDILGRAGLLSEAYELIKSIP----FDPTASIWGSLLASCRVYKNLELAEVAAEKLFE--LEpnAGNHVLLSNIYAANKQWEEIAKSRKLLRDCDVKKVR----------GKSWIDIKDKVHTFSVGESGHPRIREICSTLDNLVIKFRKFGYKP-----------------------    
 522 UniRef50_Q10PA0        Os03g0241800 protein n=4 Ta...   377    1e-103  1  10.4%  19:587    76:582     ------------------VLDASPDRTTVFFNVLLRGLTAASLHRDALLLFASMRPQGHACFPDHYTYPLALKSCAATDGLVLGRQIHSSTARLGLDGNVFVAHSAISM----------------------------YARCGRPDDAYQMFEEMQYR----DVVSWNAMISGFAHAGLfrAMDVFRELVALQckPDAGTMASILPSMGKARVEDIAlgVFDEMRFKGL----ISWNAMLAVYTNNEMHVEAVELFMRMQKDGIEPDAVTLATVLPSCGEVSALSLGKRIHEVIKRRRMCSSMLLE------------NALMDMYANCGCLKEARDVFDSMGTR------DVVSWTSIISAYGRHGHGREAIDLFEKMCGQGLEPDSIAFVAILAACSHAGLLDMGKHYFYSMTSEFhkLEHYACMVDLLGRAGCIREAYDFIMVMPIK----PNERVWGALLGACRIHSNMDIGLLAADSLLRLAPKQTGY-YVLLSNIYARAGRWADVSMVRSVMESKGIKKLP----------GVSNAELGDRVHTFHIGDTSHPQSKMIYKKLSELLRRIREMGYNPEVEATLH----------------    
 523 UniRef50_A5B564        Putative uncharacterized pr...   377    1e-103  1   9.6%   6:525    84:588     -----LYSSAEkmGCACALFDKIPEKTIVTVNCMISGFVKNKRFHAGVGLFNRVLgfDLRVKPNYVTLVILISGCVEFGRFSIGNSLHSYCCKTSLDLKNEVRNALIHLYAEFEYMDAAAKLFHEtvRDLVSWNTMIAGYAKNNDcaFSLFREMRIGNVECDRVSLVSLISACTNXrlHMGKAVHAFIKVSGMEMMIHFETALINMYSKCGSIELGRKVFDELADE----NIASWNSMIYGYVECGFNIEALSLWNVIQSRKIKPDEVTMLGLISACRSSGDLHQGIQINSYIESSD-----------HLSGSTVLCNALIDMYAKCGSMDRAETVFSKMPRR------DVISWTSIIVGYAINGEGEEALLAFRKMGAEKIEPNSVTFLGVLSACDHAGLVDKGKNLYDIMCKyhIEpeHCGCMVDMHARAGMLEEAYKFVKDMP----VEPNAVVWRMLINACRVHGDFNLGLNLVSGLIDVKTEHGPEDHVISSNIFAEAGRWDDVLQERSLMVAQKAVKLP------------------------------------------------------------------------------    
 524 UniRef50_Q8S8P6        Pentatricopeptide repeat-co...   377    1e-102  1  10.8%  10:535    79:619     ---------CIDFFKLLREFESNLKPDLTAVVTLSHRLYSNRRFNEMRSLLNSVVNDGFYKRPVEELGSAMVDcyVDNGMFEEGLRVFDYMVKKGLSIDERSCIVFLVAAKKRRRIDLCLEIFRRMVDSGvsLTIVVEGLCRRGekSKKLIKEFSVKGIKPEAYTYNTIINAYVkrDFSGVEGVLKVMKKDGVVYNKVTYTLLMELSVKNGKMSDAEKLFDEMRERGIESDVHVYTSLISWNCRKGNMKRAFLLFDELTEKGLSPSSYTYGALIDGVCKVGEMGAAEILMNEMQSK------------GVNITQVVFNTLIDGYCRKGMVDEASMIYDVMEQKG--FQADVFTCNTIASCFNRLKRYDEAKQWLFRMMEGGVKLSTVSYTNLIDVYCKEGNVEEAKRLFVEMSSKGVQPntYNVMIYAYCKQGKIKEARKLRANMEANG-MDPDSYTYTSLIHGECIADNVDEAMRLFSEMGLKGLDQNSVTYTVMISGLSKAGKSDEAFGLYDEMKRKGYTIDNKVYTALIGSM--------------------------------------------------------------------    
 525 UniRef50_Q2R1F4        Os11g0607100 protein n=3 Ta...   376    1e-102  1  11.2%  54:598   101:649     -----------------------------------------------------FVARGSLPMAHEVMR-GMVAAFGEAGRLPEAADMVLEMRSHGLPLCVETANWVLRVGLETGSFVYARKVFDGMTRAGVCPdvVVCCREGKVEEVDALLAAMWRYGFSLDNATCTVVVRSLckGRFKDVSEFFRRMLETGTPPNVVNYTAWIDGLCKRRYVKQAFHVLEEMVGRGLKPNVYTHTTLIDGLCKIGWTERAFRLFLKlkSSSYKPNVHTYTVMIGGYCREGKLARAEMLLVRMVEQ------------GLKPNTNTYTTLIGGHCKGGSFDRAFELMNKMKQEG--FLPNIYTYNAVIDGFCKKGKIQEAYKVLRMATSQGLKFDKITYTILITEHCKQGHITYALDLFDRMVENGCCPDiyTSLISTYCQQRQMEESQKFFDKCLMIGLL-PTKQTYTSMIAGYCKVGRSTLALRVFERMVQNGCFADSITYGALISGLCKESRLEEAKALYEGMLDKRLVPCEVTRV-TLTFEYCRREKTSIAVSVLDRLDkrKLSALGDVDAASLFLKKVLDEDYAVDH-ATYTGFINSCYENNR-----    
 526 UniRef50_Q9SVH0        Pentatricopeptide repeat-co...   376    1e-102  1  12.2%   8:603    83:741     -------CKVGDLGEACEVFDGMPERDVVSWNNMISVLVRKGFEEKALVVYKRMVCDGFL--PSRFTLASVLSACSKVLDGVFGMRCHGVAVKTGLDKNIFvnALLSMYAKCGFIVDYGVRVFESlqPNEVSYTAVIGGLArkVLEAVQMFRLMCEKGVQVDSVCLSNILSISacDSLSegKQIHCLALRLGFGGDLHLNNSLLEIYAKNKDMNGAELIFAEMPE----VNVVSWNIMIVGFGQEYRSDKSVEFLTRMRDSGFQPNEVTCISVLGACFRSGDVETGRRifRQMqlEGGKQIHGVVIRTEISKNSHIVSGLIAVYSECEKMEISECIFDD-----CINELDIACWNSMISGF-RHNMLdkALILFRRMHQTAvcPNETSFATVLSSCSRLCSLLHGRQFHGLVVKSGYVSDSftALTDMYCKCGEIDSARQFFDAVLRK-----NTVIWNEMIHGYGHNGRGDEAVGLYRKMISSGEKPDGITFVSVLTACSHSGLVETGLEILSSMQRIHGIEPELDHYICIVDCLGRAGRLEDAEkvLWEILLSSCRVHGDVSLARRVAEKLMR--LDPQSSAAY-VLLSNTYSSLRQWDDS    
 527 UniRef50_C5YF72        Putative uncharacterized pr...   376    1e-102  1  10.5%   3:552   173:712     --RNSLLNLYASCMrvRRLFDVMPKRNAVSWNTLFGWYVKTGRPQEALELFARMLEDGIKPTP--VSFVNVFPAVAKE-DPSWSFVLYGLLVKHGRedLFVVSSAIAMFSELGDLQSAWRVFEYTAKKNTEVWNTMITGYVQNGKFAEavIRLMGSKEVPLDVVTFLSALTAASQsdGRLGQQLHGYLIktHATLPVILGNALVVMYSRCGNVQTAFKLFDRLPEK----DIVSWNTMVTAFVQNDFDLEGLLLVYQMQKSCFAADSVTLTAVLSAASNTGDLQIGKQAHGYLVRRGIE-------------DEGLESYLIDMYAKSGRIEIAQRVFDD----YGNVKRDEVTWNAMIAGYTQSGQPEQAILTFRAMLQASLEPTSVTLASVLPACdlGGGLCAGKQIHCFALRRCLDTNVftALVDMYSKCGEISTAEHVFAVMIEK-----STVSYTTMISGLGQHGFGERALSLFYYMQEKGLKPDGVTFLAAISACNYSGLVDEGLALYRSMETFGVAVTPQ-HRCCIVDMLAKAGRVEEAYEFVQEL---------------------------------------------------    
 528 UniRef50_Q7XV61        Os04g0436800 protein n=5 Ta...   376    1e-102  1   9.9%   3:577    93:671     --AVESLAACGSLTDARKLFDAMPRRDGGSWNAIISAYSRGEEPTEAISLFSAMN--SCGVRPKDVTLASVLGCCAECLDLHGARQLHGHIAKRDFQSNVILGTALVDVYGKCFALSEARRAFDGikPNDISWNVIIRRyaGMGDMAIDMFFRMVWAGVSPLVYTVSQALLACRDNglEEGRRIHTFVLRHGYEQHVHVRSSVVDMYAKCGAIDLAQRLFDLAPVK----DIVMSTSIVSGLASCGRITDAKRVFEGMEEH----NLVSWNAMLTGYVRSMDLTSALQLFQQMRQekGEEVHTFAFKSGFIGYPFLKNALLRMYSKCGCLRSAERLLleM-----GSERDSYSWNSLISGYERHSMSEAALLALSEMHYE-ATPNQSTFSSALAACANIFLLNHGKQIHAYMIRNGYVIDDisALIDMYSKCRLFDYSMRIFEAGPSQ-----DVILWNSMIFGCAYNGKGEYGLELFFEMSKQGITPDSVTFLGALVSCISEGHVGLGRSYFTQMTDEYNIVPRIEHYECMIELLGKHGYMVELEDF-----IGHMPFEPTTAMWLRIFDCCREYG--------------------------    
 529 UniRef50_B9R998        Pentatricopeptide repeat-co...   376    1e-102  1  10.1%   6:561   295:826     -----MYAKCGDIVDAVKEFSRMPVRNVVSWTAIVSGFIKRDDSISALKFFKEMRKMKE--ETNKFTVTTVISACAKPHFIKEAIQIHCWILKTGYYLDPVVGAALINMYAKLHAISSSEMVFREMekNPGIWTIMISSFAKnsQSAIDLLLKLLQQGLRPDKFCLSSVLSVI--DSLYLGREIHCYILKTGFVLDLSVGSSLFTMYSKCGSIGDSYKVFEQIPVK----DNISWTSMISGFTEHGHAYQAFELLRKMLTERSKPDQTTFSAILSAASSIHSLQKGKEI------------HGYAYRARLGDEALVGGALVNMYSKCGALESARKMFDLLAVK------DQVSCSSLVSGYAQNGWLEEALLLFHEMLISNFTIDSFAVSSVLGAIAGLNRLDFGTQLHAHLVKLGLDSDvgSSLVTVYSKCGSIEDCWKAFNQIDDA-----DLISWTTMIASCAQHGKGVEALKIYEQMRREGIRPDSVTFVGVLSACSHANLVEEGYFHFNSMTKDFGLEPNNLDCAcrSEAHNSTKEIVKGIHILLFWDWSAHCVHG------------------------------------------    
 530 UniRef50_D1HIN9        Whole genome shotgun sequen...   376    1e-102  1   9.3%  72:586    11:500     -----------------------------------------------------------------------LDILQACNTLPKLAQLHTHIIKLGFQNNPLVLTKFTSASSNLDAIPYAMSLVFSVEddAFLFSTIIRAYAESSQsaIFYYNLMLGYGISPNKYAFPFVLKACAGLrlNLGKAVHGSLVKFGFDDDIFVQNTMVHMYCCcgGMEFARKLFDEMPK----LDPVTWTAMIGGYARLGQSAGAVGLFRKMQIAGVCPDDVTMVSVLSACTDLGALELGKWIESYIEKERVLK------------TVELSNALVDMFAKCGDVDKALGLFRNMSKR------TIVSWTSVIVGLAMHGRGLEAVSLFEEMKASGMVPEDIAFIGLLSACSHSGLVERGRQYFSEMTRqgIVpeHYGCMVDLLSRAGLVTEALEFVERMP----IEPNPIIWRTLISACRVHGELKLGESISKQLIRNE-PMHESNYVLLSNIYGKMLDWEKKSKIRVAMGKKGIQKVPGSTM----------IELDNEIHEFIVGDRSHNQYNEIIKMVNEMGREMKRAGYAPTTTEVF-----------------    
 531 UniRef50_P0C7R0        Pentatricopeptide repeat-co...   376    1e-102  1  10.8%   7:563   148:738     ------LCDDGRSEDAVELFDEMPERNVVSWNTLVTGLIRNGDMEKAKQVFDAM------PSRDVVSWNAMIKGYIENDGMEEAKLLFGDMSEKNVVTW---TSMVYGYCRYGDVREAYRLFCEMPERNIVSWTAMISGFAWnyREALMLFLEMKKdaVSPNGETLISLAYACGGLGVelGEQLHAQVISNGWefDLQSCNIIINRYLKNGDLERAETLFERVKS---LHDKVSWTSMIDGYLEAGDVSRAFGLFQKLHDK----DGVTWTVMISGLVQNELFAEAASLLSDMVRCggKHIHCVIAkaCYDPDLILQNSLVSMYAKCGAIEDAYEIFAKMVQK------DTVSWNSMIMGLSHHGLADKALNLFKEMLDSGKKPNSVTFLGVLSACSHSGLITRGLELFKAMKeySIQpdHYISMIDLLGRAGKLKEAEEFISALP----FTPDHTVYGALLGLCGLNWRDKDAEGIAErmRLLELDpnAPGHVALCNVYAGLGRHDMEKEMRKEMGIKGVKKTP----------GCSWVVVNGRANVFLSGDKSASEAAQMV----------------------------------------    
 532 UniRef50_C7J029        Os03g0840400 protein n=2 Ta...   375    1e-102  1   9.7%   4:598    17:604     ---LALYCRCGDMRHARLLFDGMPRRDAVSWNTLIAGYSGLGSCRLALDAFRDARASGDGV--DRFTFAAALASCARVGDWRDGRVVHGLAVVSGLARTAFLTNSVIDMYAKCGMIDEVRLLFDRAEERgaSWNLLLSAYVRMgeVAVNVLVWMHRSGVKLDSFALGGILKACSegDSEDVRrlHSCVVKVGLDLDVFVGSAMVDMYAKNGGLEEAIKVFDCIPNQ----NVVVYNAMIAGFARLGnpeeAVTLFSNMLKKRIKPSKFTFKSVLEVC----NLTNAVRC--------GRQIHAHVILSGLQGDEFIASVLINLYSKARCVNDSLRCFDM------TVKEDVFIWTSMITAFVQNEHFEKALYLFRELLYTRKGTDQFTISSVMSACAALSVPTTCKQIHCHAVKTGLdtVSGNSQIAMYRNIGDLKASKQTFEQ-----ISCLDTFSWSAMILSYAVHGYESKALELFEKMKNLGVMMNESASLAALIACSHQGLGDEGLRYYENTIPDDGFSLDVKLKACMVDLLGRVGKIADAEdiLWHALLRACRVHGDKERCTKIGEKLME--LEPFSASSYVMLYNLYMDAGK-----    
 533 UniRef50_D1INA5        Whole genome shotgun sequen...   375    1e-102  1  12.6%  11:580   286:867     ----------RNLEEGKKIHKIAVWKDVVSWVALLSGYAQNGMAYKSMGVFRNM--LSDGIQPDAVAVVKILAASSELGIFQQALCLHGYVVRSGFNSNVFVGASLIELYSKCGSLGDAVKLFKGmvRDVVIWSSMIAAYGIHGreALEIFDQMIqqGipPLAITSCTlvKayMRAHFrhFWEIfaDMSKsgLQYDSRILTKFAIMYVSFNRIDAASIVFEDIPN----PCSFLWNVMIRGFATDGRFLSSLELYSKMMEKGLKPDnvSILSVLLACGNLGALRKGEWF------------HSYVIQTGFEFDILVATAIMDMYSKCGSLDLARCLFDETAGK------DLVCWSAMIASYGIHGHGRKAIDLFDQMVKAGVRPSHVTFTCVLSACSHSGLLEEGKMYFQLMTEEFvkLSNYACMVDLLGRAGQLSEAVDLIENMP----VEPDASIWGSLLGACRIHNNLDLAEKIADHLF--HLDPVHAGyvLLSNIYAAKSRWNEVEKVRKMMARRGANKIQGFSLVEY----------DNQVHKFGVGDRSHPQWEKLYAKLEELAAPMKHLGYVP-----------------------    
 534 UniRef50_B9I8M2        Predicted protein n=4 Tax=r...   375    1e-102  1   9.9%   9:534    35:544     --------KCKDFNYAHRLFDEMHVRNVVTWNTVICGLVDCRGSDYESSvfCYFRKMLlkVGFDAITLNGLLRACLELNDVEIGRELHCFIVKLGFAVNSFVNSALVDLYGKCGLVKEARRAFDEvcRDLVLWNVMLSCYAMnaEEASGFFKLMQEENFMTDGFTFSSMLNSCGtgSCNLGRQIHGLSIKLSFDLDVLVASGLVDMYAKSENIDDARKAFDGMAAR----NVVSWNTMVVGYGRLGDGGEAMKLLIGMFQEDLRPDEITLASIIRSCSSA------------LTSCEIMQVNAYVLKNGFHAFLSIANALINAYSKGGSIAMALQCFN------TVLEPDLVTWTSLIGAYAFHSLPKNSIDTFEEMLADGVWPDQIVFLEVLSACSHAGLVNEGLHYFSLMKDYHILpeHYTCLIDLLGRAGLLDEAFNILNSMS----IGYSSDTLGAFIGACKIHGDVKLAKWAAEKLFE--MEPnpVNYTLMSSVFASEGHWHDVARIHKLMRDrgHGVPGCSWMEYAGD---------------------------------------------------------------------    
 535 UniRef50_P93011        Pentatricopeptide repeat-co...   375    1e-102  1   9.0%  57:580     1:495     --------------------------------------------------------MTTKVAANSAAYEAIVRAGPRVKQL---QQVHAHLIVTGYGRSRSLLTKLITLACSARAIAYTHLLfsVPLPDDFLFNSVIKskLRLPLHCVAYYRRMLSSNVSPSNYTFTSVIKSCADLslRIGKGVHCHAVVSGFGLDTYVQAALVTFYSKCGDMEGARQVFDRMPEK----SIVAWNSLVSGFEQNGLADEAIQVFYQMRESGFEPDSATFVSLLSACAQTGAV------------SLGSWVHQYIISEGLDLNVKLGTALINLYSRCGDVGKAREVFDKMKE------TNVAAWTAMISAYGTHGYGQQAVELFNKMEDdgPIPNNVTFVAVLSACAHAGLVEEGRSVYKRmkSYRLIpeHHVCMVDMLGRAGFLDEAYKFIHQLDATGKATA-PALWTAMLGACKMHRNYDLGVEIAKRLIA--LEPdpGHHVMLSNIYALSGKTDEVSHIRDGMMRNNLRK----------QVGYSVIEVENKTYMFSMGDESHQETGEIYRYLETLISRCKEIGYAP-----------------------    
 536 UniRef50_B9G6F7        Putative uncharacterized pr...   374    1e-102  1  11.5%  14:577    37:573     -------------EDARHVFDELLRRgaSIYG-LNRALADVARHSPAAAvyNRMARAGAgvTPTVHTYAILIGCCCRAGRLDLGFAALGNVVKKGFRVDAITFTPLLKGLCADKRTSDAMDI---------------------------VLRRMTELGCIPDVFSYNNLLKGLceNRSQEALELLHMMADdgSPPDVVSYNTVLNGFFKEgaMDKAMEVLNTMVKNGVMPDCMTYNSILHGYCSSGQPKEAIGTLKKMRSDGVEPNVVTYSSLMNYLCKNGRSTEARKIFDSMTKR------------GLEPDIATYRTLLQGHCKEGRVIESEKLFDLMVRIG--VKPDIITYNTLIDGCCLAGKMDEATKLLASMVSVGVKPDIVTYGTLINGYCRVSRMDDALALFKEMVSSGVSPntYNIILQGLFHTRRTAAAKELYVSITKSGTQLE-LSTYNIILHGLCKNNLTDEALRMFQNLCLTDLQLETRTFNIMIGALLKCGRMDEAKDLFAAHSANGLVPDVRT-YSLMAENLIEQGSLEELDDLFLSMEENGCSADSR--MLNSIVRKLLQRG--------------------------    
 537 UniRef50_Q7XDD5        Os10g0477200 protein n=1 Ta...   374    1e-102  1   8.6%  19:587     1:620     ------------------MRLRGVKPGVNSWNCIISGCVQNARYDEA---LNIFCEMCESESPDAVTVASILPACTGLMALGIGKQLHSYVIRCGIKLNVYIGSSLIGMYSECGEFGYARSVFAAieKNATVWNELIRSYInrMNEAWEAFRLMKENGLQPDTITYNSFIAAYARAGQKeaYTLLSNMVEIGLKPNVVSMNALISGLHHHGRHADALEAFRYMqrSGLISNIFVSSKLVDLYGKTGDVGTADKVFRRISNK----NVVTWNSILASYRQNRKPEIALKLFHEMIKshGRELHGYIRKNW--PdtALASALIDMYGKCGKIEDARLVFE------CTDEKDIATWNAIMSAYLLHRIPGEVKKLFKYIEQSGIQPDPVTFIILLSACKQEGSMEEARRYFYSMEdyGIQPthYTCMVDIMGMAGLLEESLELIQKMQ----LEPDGCLWSILLKACKLHSNLEIGEKAAKALFE--LEpnTSNYMSLSNIYADNGLWESTEALRVAMTEQGLNVERQ----------CSRLYLGTDVHTFEAGDSSHPAFEKILSTWNDLSDRMEQSGYPPQDIEPYS----------------    
 538 UniRef50_Q9SIC9        Pentatricopeptide repeat-co...   374    1e-102  1  10.6%  87:603   182:695     --------------------------------------------------------------------------------------FHSLMlkLCGSDDCTY---IIRELGNRNECDKAVGFYEFAVKrgKLASAMISTLGRYGkiAKRIFETAFAGGYGNTVYAFSALISAYGRSGleEAISVFNSMKEYGLRPNLVTYNAVIDACGKGGmfKQVAKFFDEMQRNGVQPDRITFNSLLAVCSRGGLWEAARNLFDEMTNRRIEQDVFSYNTLLDAICKGGQMDLAFEILAQMPVKRIM------------PNVVSYSTVIDGFAKAGRFDEALNLFGEM--RYLGIALDRVSYNTLLSIYTKVGRSEEALDILREMASVGIKKDVVTYNALLGGYGKQGKYDEVKKVFTEMKREHVLPntYSTLIDGYSKGGLYKEAMEIFREFKSAGLRA-DVVLYSALIDALCKNGLVGSAVSLIDEMTKEGISPNVVTYNSIIDAFGRSATMDRSADY-----SNGGSLpsSSALSALTEtgNRVIQLFGQLTTESNNRTTKDCEEGMQEliLEVFRKMHQLEIKPNVV-TFSAILNACSRCNSFEDAS    
 539 UniRef50_Q9FFG8        Pentatricopeptide repeat-co...   374    1e-102  1  10.3%  69:581    50:569     --------------------------------------------------------------------SSLISKLDDCINLNQIKQIHGHVLRKGLDQSCYILTKLIRTLTKLgmDPYARRVIEPVqrNPFLWTAVIRGYagKFDEAIAMYGCMRKEEITPVSFTFSALLKACGTmdLNLGRQFHAQTFrrGFCF-VYVGNTMIDMYVKCESIDCARKVFDEMPER----DVISWTELIAAYARVGNMECAAELFESLPTK----DMVAWTAMVTGFAQNAKPQEALEYFDRMEKSGIrrAVQIAQKSGYSPsvVIGSALIDMYSKCGNVEEAVNVFMSMNNK------NVFTYSSMILGLATHGRAQEALHLFHYMvqTEIKPNTVTFVGALMACSHSGLVDQGRQVFDSmqTFGVQpdHYTCMVDLLGRTGRLQEALELIKTMS----VEPHGGVWGALLGACRIHNNPEIAEIAAEHLFE--LEPDIinYILLSNVYASAGDWGGVLRVRKLIKEKGLKKTPA---------VSWVVDKNGQMHKFFPGNLNHPMSNKIQDKLEELVERLTVLGYQPD----------------------    
 540 UniRef50_B9MV37        Predicted protein (Fragment...   374    1e-102  1  10.2%  15:601   136:691     --------------LLRGMIHQGPRPCNRNFNAVILGFCRKGHVKVGESLLCVMQKFKCQ--PDVYAYNILINAYCTRGQTFDALGWMRFMIDNGcpSLVTFGTVINAFCNQGNMME-ARNLFDGMkpNVVCYNTLMNGyaRDIGQANMLYEEMKSKAVAPDCTTFNILVAgyRYGREEDRDRLLRDLSQSGSLSISSLYNICVSGLCWAGWLDEAMKFLEDMLEKGITPTVVAFNSIIAAYSRAGLEEKAYKAYRMMVKFGLFPSSLTCSSLIMGLSKLWRLQEARDLLYEMIVE------------GLPINKAAFTLLLDGYFRMGDVAGAYSLWNEMEGRG--IHPDAVAFSAFINGLSIVGLVDEAYDVFLQMSKKGFMPNNFVYNSLIRGFCNSGRLQEALMLEREMARKGLLPDIfnIIINGLCKEGRMKSASDVFRNMHHTGLI-PDIVTYNTLIDGYCKAFDTVSTDEVVNKMYATGWDPDITTYNIRLHGLCTGRKMSRAVMMLEELISAGVVPDTVT-YNTVMNGVC----------------------TDVLERAMIVTAKLLKMAFVPNVVTA-NLLLSHFCKQGMPEK--    
 541 UniRef50_B9ICN8        Predicted protein n=2 Tax=M...   374    1e-102  1   9.6%  70:599   166:665     ---------------------------------------------------------------------CMVRVFAEIGKFQEAVNMVIEMENHGLVLTVRTLNCVTGVAGEMGLVGYAENVFDEMRVRGVCPDSVSyaYCRMGrdTDRWLKDMVRRGFVVDNATCTLMISTFCEKGfsRVFWYFDKWVELGLKPNLINFTSLINGLCKRGSIKQAFEMLEEMVKKGWKPNVYTHTALIDGLCKKGWTEKAFRLFLKLVRSdyKPNVHTYTSMIHGYCKEDKLNRAEMLLSRMKEQ------------GLVPNTKTYTCLIDGHSKAGNFEKAYELMDLMGKEG--FSANIFTYNAFIDSLCKKGRFLEACKLLKKGFRLGLQADTVTYTILISELCRRADTREALVFFSKMFKAGVQPdtYNTLIAAFSRQRRMEESEKLFAEAVGLGLV-PTKETYTSMICGYCRDRNVSLALKFFNRMSDHGCTPDSLTYGALISGLCKESKLDEACQLYEAMVDKGLSP----------------------CEVTRLTLAYEYCKQDDSATAMVILERLDKK----LWIRTVNTLIRKLCSERKV----    
 542 UniRef50_D2STE6        Pentatricopeptide repeat pr...   374    1e-102  1  10.8%  80:580    31:503     -------------------------------------------------------------------------------RLKPLQQIHARIIITGLgtRSLITKLLSFAYAAASPISyrRLFFSIPKPDTFLFHSLITLTSKfpQESLLCYRRMLLANISSSNYTFSAVIKSSADlaFSIGETIHCHVYICGYGLDAYVQAALVSFYAKSGHVMIARKVFDKMPEK----TVVAWNSMISGYEQNGFGKEAVELFFLMQDLGVKPDSSTFVSLLSACAQVGAIGLGFWV------------HEYIARNCFDLNVVLGTALMNMYSRCGNVSKAREVFDSMEEK------NIVAWTAMISGYGMHGHGSQAIELFNEMSFDGPRPNNVTFVAVLSACAHAGLVDEGRQIFTTMkeYGLVpeHQVCMVDMLGRAGHLNEAYQFIK---NTSPKEPAPAVWTAMLGACKMHKNFDLGVEVAEHLLS--IEpnPGHYVMLSNIYALAGRMDRVEKIRNIMIRNRLKK----------EVGYSTIDVDHKVYLFSMGDKSHPKTNQIYLYLDELMSRCREAGYIP-----------------------    
 543 UniRef50_B8LL58        Putative uncharacterized pr...   374    1e-102  1  13.3%  46:532   195:680     ---------------------------------------------DAQNALRFFQWaqEGYDHNTDAYFTMIDILGRAKMFTELQSLLQKMQTQGREITRSMLHSFVMSYGRSGRFKESLEAFNLMKEMGYEpnSVLVSLVKnlDMAENLFAQMINNGVSCNNLTYTSMIQCflKEKMEDAMKLLDDMIQNNYAPDVVTYTIVISALCKRKMIEQAYGVLQKMRENGCEPNIYTYNALIQGLCAVRRPEEALELVTLMEQGGVPPNIYTYTILTHGLCKLRRLDRAKEMFNEALAR------------GLKPNRVTYNTLLNGYCRGSRLIEAMDILKEMHQN--DCTPDHVTYTTLIQGLVQGNQLPDALRMHDEMENKGYDVNFDTLNILARGLARVGNHKDASIFYRRMKDRGfsASDYYLAIHCLSTAGEMEEAQALLYEMINKGY-SPNLTTYNTMIKGFCRQGRLDDADAMLNFMIENGIGPDTGSYNILIKEFHNQGRTQDADQLYATALERGVVLNPKPVIQEP-----------------------------------------------------------------------    
 544 UniRef50_B9H2A2        Predicted protein n=5 Tax=r...   374    1e-102  1  11.4%   3:600   211:783     --ANGKVD--AALAIYKQLKSLGLNPNDYTYSIIIKAFCRKGSLVEASNVFQEMEL--CGVIPNAYAYTTYIEGLCANQRSDFGYQVLQAWKEGNIPIDVYAYVAVIRGFCNEMKMDRAEVVLGDMEkdARCYSELIRGYCKAGdkALALHNDMESKGIKTNCVIVSTILQYFCEKGmsQVVEEFKRFKDLRIFLDEVSYNIVVDALCKLEKVDQAVALLDEMKGKQMDMDIMHYTTLINGYCHVGKLVDAFRVFEEMEGKGLEPDVVTFNILLAAFSRRGLANEALKLYEYMKSQD------------LKPNAITHNVMIEGLCIGGKVTEAEAFFCNMEDKSIDN------YGAMITGYCEAKHTEKASELFFELSERGLLMDRGYIYKLLEKLCEEGEKDRALWLLKTMLDLNMEpdMYGKVITACYRAGDMRNAEAVFDILRKSGL-TPDIFTYTTMINVCCRQNRLSEARNLFQDMKHRGIKPDLVTFTVLLDGHLkaRKRKeaSNIWKEMQNTEIRP----------DVICYTALIDGHCKV------------DRLEDAIGLYDEMMYRGVEPDR-ATCTALLSGCRNRGDVD---    
 545 UniRef50_A9SGM5        Predicted protein n=2 Tax=P...   374    1e-102  1  13.2%   3:516   159:664     --ASEKLNSDEDVMKI--VNETGLRVDVRSMNVVVWQLGQMQNWYAATKVFRAFRSAG--VEPNAYVCTTLIAALGCGRRLSQALKLFRWMEKAGIERPIFTFNALMVACGRCASGDTAVELFEEMEKLGIVPDRITFtaAGLWDRAQSFIDMMQARGFSIGLHEYIEMQWACARArpREAYGLFQVMLEQGYELKLESYNALLCAYERTAQWEDAMRTFIWIQDKGLTPDVMSWSSLISACANAGQAERALEVLERMKTSDCQPNVVSWCGLLKAYQKTGNWEKAEEIFHAMLDS------------GCPPNEVAWCSLLSAYEKGRQWKKVLYTIEKLEELGMKL--DVVAWSTTISALAKAGQWELAEEKFKQMTKSGCLPNIVTYSSLIKAYGDVGLWEKAESVFKLMLRVGIRPnaCCALLRAYGKGKELEKVIIFFESMEPQYGVEPDKYAYAAIFWACWTCGEWQRAVKYIDRMEMAGCTPDSVIYTTLINMYEANGQIDKAMQVLERM---------------------------------------------------------------------------------------    
 546 UniRef50_D1ISD9        Whole genome shotgun sequen...   374    1e-102  1  11.2%  85:602   108:632     ------------------------------------------------------------------------------------IQFFKWakRRNFEHDSTTYMALIHCLDEAGMLGEMWKTIQEMVRSTCvpADlvLGKAKMVNKALSIFYQIKGRKCKPTSNTYNSMILMLmeGHHEKVHELYNEMCNEgcLPDTVTYSALIAAFGKLGRDDSAISLFDEMKENGLHPTAKIYTTILGIYFKLGRVEKALGLVQEMKEKGCALTVYTYTELIKGVGKAGKVEEAYSIFMNMLKE------------GCKPDVVLINNLINLLGKAGRLADAIKLFEEMES--LQCTPNVVTYNTVIKALFESkrASEAFLWYEKMKENGVVPSSFTYSILIDGFCKTNRVEKALLLLEEMDEKGFAPcaYCSLINALGKAKRYEAANELFQELRENCGYS-SARVYAVMIKHLGKCGRLSEAVDLFNEMKKLGCNPDVYAYNALMSGMVRVGMTDEAHSLLRTMEENGCTPDLNSH-NIILNGFARTGGPKGAIEMFTRMKNSklSRAGMFEEAAKLMKEMNSKGFEYDLI-TYSSILEA---VGKIDED-    
 547 UniRef50_B9RIR4        Pentatricopeptide repeat-co...   373    1e-102  1  11.8%   6:520   264:757     -----MYSRCQRIEDAIKVSKLTPEYDVILWTAIISGLAQNMKFQEAVAAFHKMEI--SGVSASNFTYLSMLSVCISILSLDLGRQIHSRVIRTGLEDDvvGNALVDMYMKCSCIVEHGLRMFRGIkpNVISWTSLIAGFAEHgqDSLNLFMEMRTVGVQPNSFTLSIVLRVCSAIKSpqTLKLHGHIIKTKADYDVVVGNALVDAYAGSGRVDDAWRVVKDMNQR----DSITYTSLATRLNQMGYHELALSVISHMFNADVKIDGFSLTCFFSASASLGRIETGKQL------------HCYSLKSGLSCCLSVANGLIDLYGKYGLVHEARRAFTEITE------PDVVSWNGLISGLASNGHISSALSAFDDMRLRGIQPDSITFLLVLSTCSHGGLVDMGLQYFHSMRehDVEpdHYVCLVDILGRAGRLEEAMNIIETMP----LEPDASIYKTLLAACSIHRNMNLGEDVARRGLELN-PLDPAFHLLLVKLYDDCGRYDLGEKTRRSIKQKG-----------------------------------------------------------------------------------    
 548 UniRef50_Q9LJR6        Putative pentatricopeptide ...   373    1e-102  1  11.0%   8:579    61:601     -------GRIGEISYARKVFDELPQRGVSVYNSMIVVYSRGKNPDEVLRLYDQM--IAEKIQPDSSTFTMTIKACLSGLVLEKGEAVWCKAVDFGYKNDVFVCSSVLNLYMKCGKMDEAEVLFGKmkRDVICWTTMVTGFAQAGkkAVEFYREMQNEGFGRDRVVMLGLLQASGdgDTKMGRSVHGYLYRTGLPMNVVVETSLVDMYAKVGFIEVASRVFSRMMFK----TAVSWGSLISGFAQNGLANKAFEAVVEMQSLGFQPDLVTLVGVLVACSQVGSLKTGRLVHCYILKRHV-------------LDRVTATALMDMYSKCGALSSSREIFEHVGRK------DLVCWNTMISCYGIHGNGQEVVSLFLKMTESNIEPDHATFASLLSALSHSGLVEQGQHWFSVMikYKIQPshYVCLIDLLARAGRVEEALDMINSEKLDNALP----IWVALLSGCINHRNLSVGDIAANKILQLN--PDSIGitLVSNFFATANKWKEVAKVRKLM--RNGAMEKVPGY--------SAIEVNGELRTFLMEDLSHHEHYHMLQVLRNLKTEIRdsGVE------------------------    
 549 UniRef50_C5Y1F8        Putative uncharacterized pr...   373    1e-102  1   8.4%   7:580   290:808     ------YAKADSLVDARRAFFGLPNHTVETCNAMMVGLVRTGLGAEAMQLFQFMTRSGIGFG--VVSLSGVFSACAEVK---------------GFDVDVCVRNAILDLYGKCKALVEAYLVFQEMerDSVSWNTIIAALEQnyEDTIVHLNEMLRSGMEADDFTYGSVLKACAGlsLEYGLVVHGKAIKSGLGLDAFVSSTVVDMYCKCGMITEALKLHDRIGGQEL----VSWNSIIAGFSLNKQSEEAQKFFSEMLDMGVKPDHFTYATVLDSCANLATIELGKQIHGQIIKQEMLG------------DEFISSTLVDMYAKCGNMPDSQLMFEKVQK------LDFVSWNAMICGYALHGQGFEALEMFERTQKANVAPNHATFVAVLRACSHVGLLDDGCRYFHLMtrYKLEpeHFACM-------G-PQEALKFIRSMP----LEADAVIWKTLLSICKIRQDVEVAETAASNVLRLDPD-DSSVYILLSNVYAESGKWVDVSRTRRLMRQGRLKKEP----------GCSWIEVQSEMHGFLVGEKVHPRSREVYEMLNNLICEMKLSGYEP-----------------------    
 550 UniRef50_D1I1I8        Whole genome shotgun sequen...   373    1e-102  1  10.2%   5:525   126:679     ----SLYGSFGHVECARLMFDRIRNPDLYSWKVMIRWYFLNDSYSEIVQFYNT-RLRKCLNEYDNVVFSIVLKACSELRETDEGRKLHCQIVKVGSPDSFVLTGLVDMYAKCREVEDSRRVFDEILDRNVVCWTSMIVGYVQnlKEGLVLFNRMREGLVEGNQYTLGSLVTACTKLglHQGKWVHGYVIKSGFDLNSFLVTPLLDLYFKCGDIRDAFSVFDELS----TIDLVSWTAMIVGYAQRGYPREALKLFTDERWKDLLPNTVTTSSVLSACAQTGSLNMGRSVHCLGIKLGSEddARYVFETVFDKDVIAWNSIISGYTQNGYAYEALELFDQM--RSDSVYPDAITLVSVLSACAshGYAIKalELFGDMLKEKLEPNEVIFTTILSACSHSGMLGEGWRYFNTMCqyNFVPshYACMVDLLARAGRLEEALDFIEKIP----IQPDVSLLGAFLHGCRLHSRFDLGEVAVRRMLE--LHPDKACyvLMSNLYASEGRWSQANQVMELMKQRGLAKLP------------------------------------------------------------------------------    
 551 UniRef50_P0C7Q9        Pentatricopeptide repeat-co...   373    1e-102  1  11.8%  88:603   191:706     ---------------------------------------------------------------------------------------FEKMIRKGFLPSVRNCNIVLKVLRDSRMMNKASAVYETMIEHGIMPTVITFnkAGDLERVDKIWLEMKRRNIEFSEVTYNILINGFSkgKMEEARRFHGDMRRSGFAVTPYSFNPLIEGYCKQGLFDDAWGVTDEMLNAGIYPTTSTYNIYICALCDFGRIDDARELLSSMAA----PDVVSYNTLMHGYIKMGKFVEASLLFDDLRAGD------------IHPSIVTYNTLIDGLCESGNLEGAQRLKEEMTTQ--LIFPDVITYTTLVKGFVKNGNLSMATEVYDEMLRKGIKPDGYAYTTRAVGELRLGDSDKAFRLHEEMVATDhdLTIYNVRIDGLCKVGNLVKAIEFQRKIFRVGLV-PDHVTYTTVIRGYLENGQFKMARNLYDEMLRKRLYPSVITYFVLIYGHAKAGRLEQAFQYSTEMKKRGVRPNVMTH-NALLYGMCKAGNIDEanKYSYTMLISKNCDFEKWEEVVKLYKEMLDKEIEPDGYTH-----RALFKHLEKDHES    
 552 UniRef50_D1J5H0        Whole genome shotgun sequen...   373    1e-101  1  12.2%  58:582    71:632     ---------------------------------------------------------GASFSESLQLYSSLIQQCIGIKSITDITKIQSHALKRGFH-HSLGNKLIDAYLKCGSVVYARKVFDEVPHRHIVAWNSMIASYIRNGreAIDIYQRMVPDGILPDEFTFSSVFKAFSDLGlhEGQRAHGQSVVLGvvSNVFVGSALVDMYAKFGKMRDArqVFRNMTKKGIEAnaVTWTSVIVGLVQNGREEIALLKFRQMLRSSITPNSFTLSSVLRACSSLAMLEQGKQI------------HAIVMKFGLDIDKYVGAALIDFYGKCGSTEIARSVFNGLLE------VDVVSVNSMIYSYAQNGFGHEALQLFSGMKDTGLEPNNVTWLGVLSACNNAGLLEEGCHIFSSARNSglTKDHYACMVDLLGRAGRLKEA-----EMLINQVNISDVVIWRTLLSACRIHGDVEMAKRVMNRVIDLAPE-DGGTHVLLSNLYASTGNWSKVIEMKSAMREMRLKKNPA----------MSWVDVEREIHTFMAGDWSHPNFRDIREKLEELIEKVKELGYVPDT---------------------    
 553 UniRef50_Q10MC4        Os03g0314400 protein n=2 Ta...   373    1e-101  1  10.7%  18:529   128:641     -----------------HAVRHGFGRNLYLANALMSMYSACGCLGDARKVFD------AGPVWDAVSWNTILAAYVQAEDVDQAVGVFARMPERG----AAAVSSMVSLFGRRGMVDEARKVFDVVErdVFTWTAMISCFQRNGkeALALFSDMRGEGWPVDEAVMVCVVAACARlnVVAARRLFD----SGQCLDQFSWNSMIAGYLKNGSVKDAKELFTVMPDK----DNVSWTTMISGCVQNDQSSEALTIFNNMQAQGIKPDEVTLVSVISACTNMSSLEQGKSMHEYIREHQ------------YTITVILGTSLIDMYMKCGCLESALEVFDTMEERGTP------CWNAVIVGLAMNGLVMKSLDMFSEMESSSttPNEITFTGVLSACRHAGLVEEGQHFFKLMqkYHIIPnhYGCMVDLLGRAGYVKEAENLIESMP----MSPDVPAWGALLGSCWKHGDNEVGERVGRKLVNLDPHHDGFH-TMLSNIYASEGMWQHVKDLRGSMKQWHVPKIPGSSV--------------------------------------------------------------------------    
 554 UniRef50_B9H6F7        Predicted protein n=1 Tax=P...   373    1e-101  1   8.2%  16:529     4:548     ---------------AENLFDRISNRNLLLWNVMVSGYVRNCLWDTSLAAFCEMQLGGFS--PDAVSIVSVLSACSYLEAVLFGKCAHAFSIRKGIDSSPNVSNALLAFYSDCRQLTSSFKLFHKMhrNTVSWNTLISGCVHSgeKAVDLGHSMQKEGVALDLVTLISVLPVYCDRdlGHGMTLHGHAIKKGFASDVSLVNALISTYCKCGDLDSGRFLFEVMSER----CVVSWNALITGLRHLNLQNEALVLFSQMTEY-QRPNSVtwNAIISVHIQTKYPEKAVCFFYDLLRMgaHSVMAYVICKGFEKDSAVSNALIDMYARCGDIVTAKKLFEGLIEK------DAVSWSVMINGYCLHGDGKAALEILSQMQLSGVIPNVIVFSTILSACSHAGLVEQAWMVLNSMVENGirIEHYACLVDLLGRKGHLKEAYNVVKKLPGK----PSVTLLESLLGACSVHGNVEIGEEISGLLFEMDAD-NPVPYVILSNIYAAAGRWADANKLRSNIDRRRLRKAAGCSL--------------------------------------------------------------------------    
 555 UniRef50_D1HYG4        Whole genome shotgun sequen...   373    1e-101  1   7.8%   3:527    90:615     --KNSLISFYVKCRlaRSLFDTCSVLDPVSCNIMLAGYVKSGSLDNARHLFEKmkDMRFAGvpNEVTLASVISAYSHVGGILNCRMLHALSFKLGlaLNIVATNLVHMYCVCSRLVDLARDLFERIpkDVVSWGTIIDGYVqrLGEALRMYRSMLRTGVGPNEVMIVDLISACGrmAVSEGQQFHGIIVRTGFDCYDFIQATIIHFYAACGEINLAFLQF----ELGSKDHVSSWNALISGFVRNGMIEQARQLFDEMPER----DVFSWSSMISG----------------------RWAHEYILSNSIPLNDNLNAALIDMYAKCGSITIALQLFYEIQDRVSSVSP----WNAIICGLAMHGHANVSLKLFSQLQRVRIKPNSITFIGVLSACCHAGLVDTGEKYFKGMKnyNIEPnhYGCMIDLLGRAGRLKEAAEMIRKMPMK----ADVVIWGTLLAACRTHGNVEIGERAAENLAKLDISHGAGR-VLLSNIYADAGRWDDAFLVRRAMQSQRMKKSPGC----------------------------------------------------------------------------    
 556 UniRef50_Q9ZVF4        Pentatricopeptide repeat-co...   373    1e-101  1   9.4%  76:587    19:502     ---------------------------------------------------------------------------ASSSKPKQLKKIHAIVLRTGFSenSLLTQLLENLVVIGDMCYARQVFDEMHKPRIFLWNTLFKGYVRnpFESLLLYKKMRDLGVRPDEFTYPFVVKAISqgDFSCGFALHAHVVKYGFGCLGIVATELVMMYMKFGELSSAEFLFESMQVK----DLVAWNAFLAVCVQTGNSAIALEYFNKMCADAVQFDSFTVVSMLSACGQLGSLEIGEEIYDRARKEEIDC------------NIIVENARLDMHLKCGNTEAARVLFEEMKQR------NVVSWSTMIVGYAMNGDSREALTLFTTMQNEGLRPNYVTFLGVLSACSHAGLVNEGKRYFSLMvdKNLEpeHYACMVDLLGRSGLLEEAYEFIKKMP----VEPDTGIWGALLGACAVHRDMILGQKVADVLVET--APDIGSyvLLSNIYAAAGKWDCVDKVRSKMRKLGTKKVAAY----------SSVEFEGKIHFFNRGDKSHPQSKAIYEKLDEILKKIRKMGYVPDTCSVFH----------------    
 557 UniRef50_C5Z2D6        Putative uncharacterized pr...   373    1e-101  1   9.0%  14:568    78:613     -------------AHARRLFDEMSRRNLVSWSVVISGSARHGVLAEAFALFSHMLhqGSWDRPDSFMLGALVAGCSRARHVDAGVQVHACVAKFGVDEDESVAAALVDMYAKCGWVDSSWRAFtaPQRSVLSWTSMIACLVNQgdAAMLLFKKMLALKVWPTNVTFSCILKVFNTPDlsVGMQIHGCLLKIGTEVDTALGSALMTMYGRCGGVDEIARL-----ACRIRHDVFSRTSLLGAYARNGYNAEAIGVFKEMILTNMAIDQSAMTCLLQVCSSVGQLRMVREV------------HCYALKTFFKLDTLLLNAIITVYGRCGDTTSAETVFNLMEEQ------DIISWTALLTCYAQNGLDEEVLLFFREMLRRGLGSPVFCMTSVLRACSRTSNLAIGLQIHSRTVKLGIDDdvENALVTLYANCGSVQVALKIFNSMSNR-----DIVTWNALLTSFSQHGNEVAAIQLFDLMQEEEVCPDDYTFVGLLSSCSRMGLVKEGCEYFNEMKAKYNLEPKMVHYTCMVDLFARAGR---FCDAMDFIDAMPYEPDQILWEALL-----------------------------------    
 558 UniRef50_B7EEI3        cDNA clone:J013134G11, full...   372    1e-101  1  10.2%   7:514    57:543     ------YARFGDVAAARKVFDGMPHRSVVSWTAMVSGYARNGRPREALELFALMRASGA--RPNQFTYGSAASACAGAGCARSGEQVHACAAKGRFAGDMFVQSALMDMHLRCGSVEDARQLFAEmkKDVVSWNALIRGFVERGhdALGLFSSMLKEAMIPDHYTLGSALKACGIVGVanVELIHSCIIKLGYWDEKVVIGSLINSYAKCRSMSSARVIYDSISE----PDLVSSTALISGYTMDRNyeDAMELFCKIHRKGLWIDGVLLSSVLCLCASVASARFGTQIHAYMCKKQPMG------------DIALDNALVDMYAKAGEFSDAKRAFDEMPYR------NVISWTSLITACGRNGSGEDAVTLFNRMVEDGVRPNDVTFLSLLSACGHCGLTNKGMEYfsMMSRYGIDpeHYSSAIDLLARGGQLEDAWKLVQ----KTNLKPNSSMLGAMLGACKLHGNMLLGETAAKNLFS--IDpsSVNYAVLANMYAECSLWEDAQRTRE-----------------------------------------------------------------------------------------    
 559 UniRef50_Q8S1U2        Putative uncharacterized pr...   372    1e-101  1  10.7%   3:600   305:891     --ANSLITFYYELGfaEEVFLSTSNKNLVTWNAMIKGLVENDRVNEAMCMFQ---EMRSKNQPDVATLVTIISACGDHGLLPEGKEVHGYIIKKgiYEECSVGNSLLDLYMKCNDPSTARILFRTMprDLISWNTMISGYSRngEEAKAMFKGLLSEGLSCTLSTVVAVisCFCPQDLNFGKSVHSFILKYGFLTGVSAANSLIHMYICCGDSLAAFSLLESIT---PISDIISWNTAIVGCVQNGLYGDALEAFQFMHslTLNPDSITLVSVLSVCGNLK------------LQSLGKSIHCMALKRLIEFNLRVKNALLTMYFRFGDTESAELIFSSLVGR------NLCSWNCMISGFAQNNEGLRAFQFYKKMED--FEPNEISIVGIICACTQLGDLRQGKNIHGHVVRFGLQTNVfaSLVDMYSKCGRLDISIRVFESSAEK-----SIACWNSMISAFGFHGLGLKSIEIFWKMNNSGVKATRSTFIALLSACSHSGLTDEGLKYYHLMIEHFGIIPTPEHHVCVVDMLGRAGRLQEaaHGVWGALLSACSKKSELKMCESVAKHLL--CLEP-ENSGYYVTMSNLYAYQDMW---    
 560 UniRef50_B9N2S4        Predicted protein n=4 Tax=r...   372    1e-101  1   8.7%  86:600    76:572     -------------------------------------------------------------------------------------QIHTQLTVLGLFQHPFAASriKKLCACLNSVSHCVSLYNHIepDAFMCNTIMRSFVndPFGALRFyeKMIAKWVLPNHYTFPLVAKVCAdgSLREGQKVHALVVKFGFELDLFVRNSFIRFYSVCGRTSDARMVFD----NGFVLDLVSWNSMIDGYVKNGELGLAREIFDEMYER----DIFTWNSMISGYVGVGDMEAARGLFDKMPSR----------------DVVSWNCMIDGFARIKDVSMAAKFFDEMPLR------NVVSWNVMLALYLRCKKYSDCLRFFDMMVGGDFVPDEASLVSVLTACAELKMLDQGKWVHSYMKDNGIKPDMltALLTMYAKCGAMDLAREVFDKMPEK-----SVVSWNSMIIGYGIHGHGDKALEMFREMEKGGPMPNDATFMSVLSACSHSGMVWNGWWYFDLMHRKYRIQPKPEHYGCLVDLLGQAGLKeeVEPTLWGDLLSACRAHCISEPGEILAKQLIK--LFPNHVVPYLLLSNTYVAEGRWD---    
 561 UniRef50_D1HCY3        Whole genome shotgun sequen...   372    1e-101  1   9.7%   7:520   203:739     ------YGKLGCIYYAQHLFEEMPRRDVVSWNALICGYSRNGYDYDALEVFVQMLREGFPprPDifSWNTMIGAYGQNGLFDEAMLVFKQMQKERVEVN--YVTIIMCSYAGCGNIESAGLLYNLmqRNLVSLTAMISGYAEKGNMGleCFTQMLQLKMKPDAVAMVSILHGFTDPTfgSGLGIHAYGLKTGLCADCLVVNGLISMYSKFGDIETVFSLFSEMGEKQL----ISWNSVISACIQVGRTSDAMELFCQMRMYGHSPDAITIASLLAGCSEVGFLQFGERL------------HNYVLRNNLDMEDFLETALVDMYIKCGRLESAERVFKSIKE------PCLATWNTMISGYGLSGHEHRALSCYSEMQEQGLKPDRITFLGVLSACTHGGLVWEGKRYFRSMREdgMIpqHNACMVDLLSRAGFLEEAVIFVKNME----VEPDSAIWGALLTSCCIHQELKLGECLAKRLLLLDYY-SGGLYVLMSNLYASKGRWDDVARVRKMMKDTG-----------------------------------------------------------------------------------    
 562 UniRef50_A5C4Z2        Putative uncharacterized pr...   372    1e-101  1  10.1%   3:576    80:626     --ANSLMDMYAKCgdAAKLFDHMPDKTVVSWTSMMSGHCQRGAFDEVISIFWRMLE---TLQPNEYTLAVILQACAQKRDLKLVQLIHCHIIKTGFVMDAFLQNSLIDGYTKSGTLVAAEKLMKRLirDVVSWTSVISGCVLNgeKALLFFFEMQEDGVSPntETIATLLQGCSHSklKLGEQIHGYQIKHGFFPCTIVENSLIYMYAENERDDAAFQLFRKMSCR----DIVSWNTMISSLVKGSSSYQALMLLSEVHSNgiYPDFVTILASIQACSSLASLQLG------------QVIHGYITRAGLICDIFVQNSLVDMYGKCGRLHLAEKVSEEMPVR------DLGSWNSLIAAYGINGNGISALNVFKQLKNTghRPNAITFTNILSACAHAGLVAEGFEIFKSMkeYSLEpeHFACM--------RPE---AFIQKMP----FEPGPEVWGALLGGCGLFGNLDIAERVAKKLXI--LEPKSRAwvALSNVYASVNKWEDAAKVRAEMRR-------SEELQK--EGGWSSVEVRGZEFRFMVGDTMHPEARMVYAVLKGINEHIRDA---------------------------    
 563 UniRef50_C5YH53        Putative uncharacterized pr...   372    1e-101  1  10.3%   9:601    22:574     --------RCADTRSARLLFDGMPRRDVVTWNTLIAGYATQGSARSALGAFRDARRDGA-VAVDRFTYAAVLAACGGAGDWRSGRAAHGLAVVSGLARTAFVSNSVIDMYAKCRMIDEVRLAFDRAerDEVSWNLLLSAYVRMgeVAANVLVWMHRSGVELDAFALGGILKACSEldSEDVRrlHGCVIKVGLDLDVFVGSTMVDMYAKNGGLEEAIKVFGSIPSQ----NVVIYNTMIAGFARLGnpeeAVRIYSNMFRRRIRPSKFTFKSVLEVC----NLTNAVRCWRQI--------HAHVILFGFEDDEFIGNALINLYSKARLVDDSLRCFHRTPKQE------IFTWTSMITAFVRNEHSDKALNLFKGLRYTGVEPDQFTMSSVMNACADLSMPIACEQIHCYAVKSGFDRFTlnSQIEMYRCTGDLKAAKKTFERIP-----SLDTFSWSQMILSYAVHGHEREALLLFKKMRDCSVIINEFAFLAVLVACSHQGLIDEGFRHYESMVSDYSFVPDVKHIACMVDLLGHVGKVADAEDFINS------------------------SGLENDSVL-WHTLLRACRIHGDKDR--    
 564 UniRef50_Q56X05        Pentatricopeptide repeat-co...   372    1e-101  1  10.4%  22:579    85:570     ---------------------QMQEPNVFVYNALFKGFVTCSHPIRSLELYVRM--LRDSVSPSSYTYSSLVKASSFASRF--GESLQAHIWKFGFGFHVKIQTTLIDFYSAT----------------------------GRIREARKVFDEMPER----DDIAWTTMVSAYRrlDMDSANSLANQMSEK----NEATSNCLINGYMGLGNLEQAESLFNQMPVK----DIISWTTMIKGYSQNKRYREAIAVFYKMMEEGIIPDEVTMSTVISACAHLGVLEIGKEV------------HMYTLQNGFVLDVYIGSALVDMYSKCGSLERALLVFFNLPKK------NLFCWNSIIEGLAAHGFAQEALKMFAKMEMESVKPNAVTFVSVFTACTHAGLVDEGRRIYRSMIdySIVSnhYGGMVHLFSKAGLIYEALELIGNME----FEPNAVIWGALLDGCRIHKNLVIAEIAFNKLMV--LEpnSGYYFLLVSMYAEQNRWRDVAEIRGRMRELGIEK---------ICPGTSSIRIDKRDHLFAAADKSHSASDEVCLLLDEIYDQMGLAGYV------------------------    
 565 UniRef50_A9SNP2        Predicted protein n=1 Tax=P...   371    1e-101  1  11.6%  16:599   235:840     ---------------FKDMRAAGHLPDELLYNTIICALGKLGRYQESEALYLDMKKQGIV--PSKFTYTIMINVWSKAGRFASAAETLAEMQRSGCIADEVVYCSIINMYGKAGLYEEAEKIFKEMDTLGlsYTSmvRAEAGRHAEALKLFDVMAEKGLLTTRMTWNTLLHCFVrgDVEQATKVYNDMVEAG-SANVVTYGNMINLYSKFQMVEDAENLLAEMRESGVKPDEYIYGSFVKLYCNSDMIDKATMVVQEMKDDGLeiDTQILNQLLIKRAEAGELREAELLLDKLVEAGgkSLFNSLQKKDHPPSLYVYNTMIKLCAVCKELEEAIFVFDRMEENGRMF--DAVTVSILVHAYTKEGRFKDAAGLMKRAKKVGVAMDTVAYNTSLKANLKSGNLKGALEVYGEMQEADIEPstYTILISLFSKLGDLGRAVQAF-EVLNSSEVGADEIAYSQMIHCYGCAGRPKEAADLFQEMETKGFKPNEVIYNNLLDAFARAGLFAEARLLLSDMRRKGCPPSSVTYL----------------------LLMSAYGSKGKPADAESLLHLMQDRGLYPDCRH-YNEVIRAYGNVGKL----    
 566 UniRef50_D1HTT1        Whole genome shotgun sequen...   371    1e-101  1  11.3%  15:599   136:771     --------------FVGEMESEGHFIDKMMYTSLIHGYCRGKKMRTAMRVFLRMLKMGC--DPDTYTYNTLIHGFVKLGLFDKGWILHNQMSEWGLQPNVVTYHIMIRRYCEEGKVDCALTLLSSMspSVHSYTVLITALYkrLVEVEELYKKMLDIGVVPDHVLFFTLMQKQPKGhlHLALKILQAIAKNGCNLDlvAFGIFISALCAAGKTDAALLFMDKMVSLGCRPLLSTYNSLIKCLFQERLVEDAKSLIDLMQENGIVPDLATYLIMVHEHCNHGDLASAFGLLDQMNER------------GLKPSVAIYDSIIGCLSRRKRILEAENVFKMMLEAG--VDPDAIIYVTMISGYSKNRRAIEARQLFDKMIEHGFQPSSHSYTAVISGLVKENMIDKGCSYLSDMLKDGFVPNtyTSLINQFLRKGELEFAFRLVDLM-DRNQIECDMITCIALVSghVKSgrVReaLNLMQKIKGSSFMPNLYLYNGIISGFCRANMIQDAYNHFELMQTEGVCPNQVT-FTILINGHTRFGEID---------------------HAIGLFNKMNADGLAPDGI-TYNALIKGLCKAGRL----    
 567 UniRef50_D1J2E2        Whole genome shotgun sequen...   371    1e-101  1  10.5% 136:601   101:526     ---------------------------------------------------------------------------------------------------------------------------------------LVCAFTPSFHYAQQIFKCVEKQ--KPETFVWNSCLKALAEGDSpdAIMLFYRLRQYDVCPDTFTCSSVLRACLNLLDLSNGRILHGVVEKVGFR----SWTSMIAGYVQCGKAKEAIHLFAKMEEAGVKCNEVTVVAVLAACADLGALDLGMRI------------HEYSNRHGFKRNVRISNTLIDMYVKCGCLEEACKVFEEMEER------TVVSWSAMIGGLAMHGRAEEALRLFSDMSQVGIEPNGVTFIGLLHACSHMGLISEGRRFFASmrDYGIIpeHYGCMVDLLSRAGLLHEAHEFILNMPMK----PNGVVWGALLGACRVHKNVEMAEEAIKHLLELD-PLNDGYYVVLSNIYAEAGRWEDTARVRKFMKDRQVKKTP----------GWSSITVDGVVHEFVAGEESHPDTEQIFQRWEELLEEMRLKGYVPN---TSVVLLD--IEEGEKVK--    
 568 UniRef50_B9GUE7        Predicted protein n=3 Tax=r...   371    1e-101  1   9.0%   3:593    41:624     --ACNLVSMYAdlIDAYKLFDEMPVKNIVTWTTMVSAYTSNGKPREAIKLYTRMLDSKSEV-PNGFMYSVVLKACGLVGEIELGRLIHKRFSRENLDYDIVLLNALLDMYVKCGCLSDARKVFdmPDRNVVSWNTIIAGLAENGSsaLQFVCKMHREGIKLDKFTFPCALKTCSYAglVAGKQIHCYVLKSGLESSCFAVSALVDMYSNCNGLDDAIRLFDQYSggSICDSLVLWNSMLSGYVVHEKNRAAVNMIAQIHHSGASVDSYTLSSALKVCINLLNVRLGIQV------------HALIVTSGHELDYVVGSILVDLYAKLGNMKDAFKLFHRLPKK------DIVAWSGLLMGCAKMELNSLALSLFRDMVTFGVEVDQYIVSNVLKVCSSLASIGTGKQVHAFCIKRGYETetITALIDMYSKCGEVEDGLVLFG-----CVADRDVVCWTGIIVGCAQNGRANEALEIFRQMVQSGLKPNEVTYLGVLTACRHAGLVVEAQTIFGTMKCDHRLEPQLEHYYCMVDLLCQAGY---------------------FKEVEKLIAEMP---FKPDK-TIWSSMLGAC----------    
 569 UniRef50_Q8GRU0        Os07g0113500 protein n=5 Ta...   371    1e-101  1  11.7%  34:580    40:545     ---------------------------------LLTGCATLRRTGELHAA-----AVRAGVDGDRAVGFRLQRAYAASDRLDLTVTLL----RLTPDPTTVFYTSAIHAHSSRGLHLAALALLSEMLGRGLIPTSHTLSSSLPACHGLALGRALHAYAFKldSYVATALLGMYARgdADAARALFDEMPD----PHVVPVTAMLTCYAKMGALDDARELFDGMPSK----DFICWNAMIDGYTQHGRPNEALRLFRWMLRSGVDPDEVAIILALSAVAQLGTAESGRWLHSYVKNS-----------RRVQLNARVGTALIDMYCKCGSLEDAVSVFNSIGDK------DIVVWNAMINGYAMHGDSRKALEMFSQLRSQGLWPTDITFIGLLNACSHSGLVDEGHQFFQSMEeyAIVpeHYGCMVDLLGRAGLIEEAFHLVQSM----TIAPDTVMWVSLLAACRLHKNMALGQQIADYLVAGGLA-NSGMYILLSNIYAAVGNWEEVARVRSMMKASGIQKEP----------GCSAIEVGRKVYEFVAGDMSHPRTDEIYAMLEKMNGIVKEQGHVP-----------------------    
 570 UniRef50_B9SKC9        Pentatricopeptide repeat-co...   371    1e-101  1  11.2%  16:508    69:546     ---------------ARHLFDKSPQRpvVIWNMLIRAYAWNGPFTEAINLYYKMLELG--IQPSKYTYPFVLKACSALQAIEEGKEIHIHAKRLDLDSDVYVSTAVVDMYAKCGCLDDAEMVFSGMprDVVAWNSMIAGFSLhyNETIQLLAQMQKDLVRPNSSTIVAVLPAVAQAnlSHGKALHGFCVRRGYIDDVVVATGLLDMYGKCQCIIYARNFFDMMS---IVRNEVTWSAMLGAYVMCELMREALDLFQYmkDAAVkpTEVTLATVLRACAELTDVSRGR------------CIHCYTIKSGFISDLMLGNTLLSTYAKCGTVSDAIRYFHEMELK------DEVSYSAIISGCMQNGNAKEALSMFHRMRLSGMDPELATMVGILPACAHLAALQHGSCCHSYALIKGfeITICNALIDMYAKCGRIDTARNVFDRMHKR-----DIVSWNAMIIAYGNHGLGMEALLLFDNMLADGLEPDDVTFICLLSACSHSGLVKE-----------------------------------------------------------------------------------------------    
 571 UniRef50_D1HTH7        Whole genome shotgun sequen...   370    1e-101  1  10.7%  63:576     5:490     --------------------------------------------------------------RNSYDYTYLLQ---RCKGTKTIKSIHAQIIIGGFEENPFLGAKLVGKYAQcsNIEDARKVFDCLprDVFVWNTIiyANLGPFMEALNIYEYMRCSGVAANRYTFPFVLKACGAMKdkKGQAIHGHVVKHGLDLDLFVGNALVAFYAKCNEIGASRRVFDMISEK----DIVTWNSMISGYAINGCADDALVLFHNMLqqgyAPDSATLVAILPACAQAAAIQEGLWI------------HSYVIKSGIELDAALGSGLIAMYANCGLLNSARDVFDRIDDK------NIVVWNAIIRCYGMHGHADEALKMFSGLIDSGLHPDGVIFLCLLSAFSHAGMVAEGMELFEKMGDYGVesEHYASVVDILGRAGFLHEAVEFVKNMP----VQPGKDVYGALLGACRIHNNIELAEEIAEKLFVLDPD-NAGRYIILVKMYEDAGKWENAARLRKALKEKNIRKP----------LGCSAIELGCVLHTFGVEDKSHPFKDQIFDTLERLDNIMEEE---------------------------    
 572 UniRef50_D1HGG7        Whole genome shotgun sequen...   370    1e-101  1  10.1%  68:580    12:487     -------------------------------------------------------------------FNHLAQQIKKCSMVKELESVYASMIKANANQDCFLMNQFIAACSIFHRIDYAILAFTHmePNVFVYNAMIRALVqhPVQALDCYLDMVQAQVSPTSFTFSSLVKACSlsELGFGEAVHGHIWKYGFDSHVFVQTALVDFYGNAGKIVEARRVFDEMSER----DVFAWTTMISVHARTGDMSSARQLFDEMPVR----NTASWNAMIDGYSRLRNVESAELLFSQMPNR----------------DIISWTTMIACYSQNKHLDKSLVVFFKLRKK------NLFCWNSIIEGLAVHGYAEEALAMFSRMQREKIKPNGVTFISVLGACTHAGLVEEGRKRFLSmrDFSIPPehYGCMVDLLGKAGLLEDALELVRSMR----MEPNSVIWGALLGGCKLHRNLKIAQVAVNE--SKVLEPnsGYYTLLVNMYAEVNRWSEVANIRATMKELGVEKTSP---------GSSWIEMDRKIHQFAASDKSHLASDEIYTLLVELDGQLKLSGYVP-----------------------    
 573 UniRef50_B9T607        Pentatricopeptide repeat-co...   370    1e-101  1   8.8%   4:525   168:671     ---NTMIEVYARCGCvrVMFDEMSPRDLVSWTSMISGYVSEGNV---FSAFELFNKMRLEMEPNSVTLIVMLKGCYAYDNFSEGRQLHCYIIKNGLLIYGSVQNSILRMYSITGSAKEVESLfeIYRRDVISWNTLiyALRGDAEEMVCGFNQMRGE-VALSSETLTLVISVFAkgNLVEGEKLHSFSIKVGLCDDV-LLASLLDFYAKCGELRNSVQLFGEIPCR----SSSTWKLMMSGCIQNGYFDEAIHLFRQMQASGVQLQAQILGSLVDACSHLGSLQLCKEIHGYLTRNFFYILEGD--------NIHLGTSILNMYIRCGSISSAREYFNRMVAK------DNITWTSMIEGYGIHGMAIEALKLFNQMLVERVLPNRVTFLSLLSACSHSGLIRQGCELFLSMKwfGMEPdhYTCMVDLLGRCGKIKEALAMIIRMV----VVADSRIWGALVASCRVHGDKKVGEFAAQRLLEMESD-NVGYYTLLSNIQAMVGKWDEVEQVRKVIHEKDLRKTP------------------------------------------------------------------------------    
 574 UniRef50_O22137        Pentatricopeptide repeat-co...   370    1e-101  1   9.5%  27:551    85:604     --------------------------DPFLWNAVIKSHSHGKDPRQALLLLCLMLENGVSV--DKFSLSLVLKACSRLGFVKGGMQIHGFLKKTGLWSDLFLQNCLIGLYLKCGCLGLSRQMFDRmkRDSVSYNSMIDGYVKCGlsARELFDLMPME--MKNLISWNSMISGYAqdGVDIASKLFADMPEK----DLISWNSMIDGYVKHGRIEDAKGLFDVMPRR----DVVTWATMIDGYAKLGFVHHAKTLFDQMPHR----DVVAYNSMMAGYVQNKYHMEALEIFSDMEKekAIDMHLYIVEKQFYLGGKLGVALIDMYSKCGSIQHAMLVFEGIENK------SIDHWNAMIGGLAIHGLGESAFDMLLQIERLSLKPDDITFVGVLNACSHSGLVKEGLLCFELmrKHKIEpqHYGCMVDILSRSGSIELAKNLIEEMP----VEPNDVIWRTFLTACSHHKEFETGELVAKHLiqAGYNPSS--YVLLSNMYASFGMWKDVRRVRTMMKERKIEKIP----------GCSWIELDGRVHEFFV----------------------------------------------------    
 575 UniRef50_Q7XIR4        Os07g0688100 protein n=3 Ta...   370    1e-101  1  12.3%  14:524   146:649     -------------EFLGRVRSQGVAFHRSLYRVLLAGYARAGKFDSVIETFDEMVTSGCREF--GVDYNRFIGVMIKNCCFDLVEKYYNMALAKGFCLTPFTYSRWITALCQSNRIELVEELLTDMDKFGcaCNIYVHYLCGHNrdALQMVEKMTMKGTGPDVVTYTTVVSCLchRRFSEAVGLWEEMVRRGLKPDVVACGALIFGLCKNQKVDEAFELASRMLTLDIQLNVSIYNALISGFWRAGSIEKAYKTVSFMQRNGCEPDVVTYNILLNHYCSIGMTDKAENLIRKME------------MSGVNPDRYSYNILLKGLCKAHQLDKAFAFvdHMEVGGFC---DIVSCNILIDAFCRAKKVNSALNLFKEMGYKGIQADAVTYGILINGLFGIGYSNLAEELFDQMLNTKIVpnVYNIMLHNLCKVGHFKHAQKIFWQMTQK-EVSPDTVTFNTLIYWLGKSSRAVEALDLFKEMRTKGVEPDNLTFRYIISGLLDEGKATLAYEIWEYMMENGIILD-------------------------------------------------------------------------------    
 576 UniRef50_C5XPI6        Putative uncharacterized pr...   370    1e-101  1  11.4%  32:540   418:899     -------------------------------NGLVNMYAKCGAIDSASKIFQLMEA------TDRISWNTIISALDQNGllSAGQQVHCDAVKWGLDLDTSVSNVLVKMYGECGAMSDCWKVFNSmdHDEVSWNSMMGVMASsiSETVEVFNNMMRGGLIPNKVTFVNLLAALSPLslELGKQFHAAVLKHGVMDDNVVDNALISCYAKSGDMNSCEHLFSNMSGR---RDAVSWNSMISGYIYNGYLQEAMDCVWLMLHSGQIMDCCTFSIILNACASVAALERGMEM------------HAFGIRSHLESDVVVESALVDMYSKCGRVDYASKLFNSMTQR------NEFSWNSMISGYARHGLGRKALETFEEMLRSRESPDHVTFVSVLSACSHAGLVERGLEYFEMMPDHGILpeHYSCVIDLLGRAGKIDKIKEYIQRMPMK----PNALIWRTVLVACRQSKdiELGREALRVLLE--IEpnPVNYVLASNFHAATGMWEDTAKARAGMRQ-------ATVKKEAGRSWTYLG---------------------------------------------------------------    
 577 UniRef50_UPI0001983407 PREDICTED: hypothetical pro...   370    1e-101  1   8.5%  62:553     5:502     -------------------------------------------------------------SHNHLKLTTLLLNYAEHFQTKQLLQIQAQLItlSLHPNLIAVKLIGACAANANLYHAQLIFAQLPNPNVFSWNAIIKAhcGAWEHALNCFNRQLSLPsnPNEYIYTSLIKACAGLeiADGLKVHAVVIKSGLDNNIHLGTSLVDMYFKFREPASAGQVFDEMPLR----DVVSWNTMVSGYCLCGDLESARRVFDQMLER----DVISWNAMIGGYVQNGKYSDAIEVFHEMQKVggRWIDRFVGWRGRGLNLYLGNALIDMYAKCGTMEEARRIFDGMRER------DVISWSTMICGSGTHGDADEAFGYYSKMLECGVKPNEVTFMGLLSACSHAGLVDKGIELFSRMieYRIVpgHYGCVIDLLSRAGRLDEAEDLINSMP----IEPNVIVWGALLGGCRIHKDFRRGERVAQHLLELDSEYT-GSYVYIAGAKASVGRVDDAANCWLRMQHKGIIKDP----------GCSKIEVHNTVHEFKECD--------------------------------------------------    
 578 UniRef50_D1J1V9        Whole genome shotgun sequen...   370    1e-100  1  11.9% 106:592   113:577     ---------------------------------------------------------------------------------------------------------LLHLLANAKNYNKIRALLDSFAKNAHYSNSTIF-------HSLSVLG---SWGCA-NSIIVDMLVWAYVKNgmDLALEGFDRAGDYGFRLSALSCNPMLVSLVKEGRIGVVESVYKEMIRRRIGVNVVTFDVVINGLCKVGKFQKAGDVVEDMKAWGFSPSVITYNTIIDGYCKAGKMFKADALLKEMVAKR------------IHPNEITFNILIDGFCRDENVTAAKKVFEEMQRQG--LQPNVVTYNSLINGLCSNGKLDEALGLQDKMSGMGLKPNVVTYNALINGFCKKKMLKEAREMLDDIGKRGLAPntFNTLIDAYGKAGRMDDAF-LLRSMMLDTGVCPNVSTYNCLIVGFCREGNVKEARKLAKEMEGNGLKADLVTYNILVDALCKKGETRKAVRLLDEmmEKKGRRANIVTYNVLIKGFCNKGKLEEANRLLNEMLEKGLIPNRTTY--DILRDEMMEKGFIPddGHLYNVSISS-----------    
 579 UniRef50_A9RVJ5        Predicted protein n=1 Tax=P...   370    1e-100  1  10.5%   1:526    88:605     LSKESRVEEAMLLFD--QLKQLGVSPGPMAYNAIVGTCAREDRLETALELMNQMR--GSGFQPDNVNYTLVFQACAKKrgVDVISRVCANIEQEGLEMDTKLYNDVINAYCRAGDPDKAFQYMGLMqpDVRSYSSLIETLvrRLDDAEAAYAEMKSKSYKINLRTLNALLSAYTRKSleQVEKLMQDAEDAGLKLSTFSYGLLIDAYSRAGRLDQAKAAFHNMKVENVPANAFIYSRLMVAYRNARQWDGTIRLLKDMYASNIKPNQFIFNILIDTYGKFGRLPQAMRTFAQM------------DKEGFKPDVVTWNSLIEAHCRAGLITEALDLLKQMQEREC--VPSLHTYNIILNALGWHNRWKEMALLLDEMRFKGLDPNVVTYTTLVDSYGTSKRYREASEYLKQMKSQGLQPStyCALANSYAKRGLCEQAMEVFKSMEKDG-VELNLSMLNLLINAFGVAGKYVEAFSVFDYILEVGMSPDVVTYTTLMKALIKAEQLDQVPDVYEQMVRAGCTPDDM-----------------------------------------------------------------------------    
 580 UniRef50_O80647        Pentatricopeptide repeat-co...   369    1e-100  1   8.8%   6:552   243:767     -----MYCNCADLYAAESVFEEVWRKDESSWGTMMAAYAHNGFFEEVLELFDLMR--NYDVRMNKVAAASALQAAAYVGDLVKGIAIHDYAVQQGLIGDVSVATSLMSMYSKCGELEIAEQLFINIerDVVSWSAMIASyaGQHDEAISLFRDMMRIHIKPNAVTLTSVLQGCAgaASRLGKSIHCYAIKADIESELETATAVISMYAKCGRFSPALKAFERLPIK----DAVAFNALAQGYTQIGDANKAFDVYKNMKLHGVCPDSRTMVGMLQTCAFCSDYARG------------SCVYGQIIKHGFDSECHVAHALINMFTKCDALAAAIVLFDK-----CGFEKSTVSWNIMMNGYLLHGQAEEAVATFRQMKVEKFQPNAVTFVNIVRAAAELSALRVGMSVHSSLIQCGFCSQtgNSLVDMYAKCGMIESSEKCFIEISNKY-----IVSWNTMLSAYAAHGLASCAVSLFLSMQENELKPDSVSFLSVLSACRHAGLVEEGKRIFEEMGERHKIEAEVEHYACMVDLLGKAGLFGEAVEMMRRM---------------------------------------------------    
 581 UniRef50_D1IC62        Whole genome shotgun sequen...   369    1e-100  1  12.9% 177:580    10:379     --------------------------------------------------------------------------------------------------------------------------------------------------------------------------------DVREGEKVHSIAIRNGFESLVFVQNTLVHMYAACGHAESAHKLFELMAER----NLVTWNSVINGYALNGRPNEALTLFREMGLRGVEPDGFTMVSLLSACAELGALALGRR------------AHVYMVKVGLDGNLHAGNALLDLYAKCGSIRQAHKVFDEMEEK------SVVSWTSLIVGLAVNGFGKEALELFKELERKGLMPSEITFVGVLYACSHCGMVDEGFDYFKRMKeyGIVpeHYGCMVDLLGRAGLVKQAHEFIQNMP----MQPNAVVWRTLLGACTIHGHLALGEVARAQLLQ--LEPKHSgyVLLSNLYASEQRWSDVHKVRRTMLREGVKKTPGH----------SLVELRNRLHEFVMGDRSHPQTEEIYVKLAEITKLLKLEGYVP-----------------------    
 582 UniRef50_Q9FFN1        Pentatricopeptide repeat-co...   369    1e-100  1  10.0%   4:582   223:808     ---MSLYDKDssCDDVLKLFDEIPQRDVASWNTVVSSLVKEGKSHKAFDLFYEMNRVE-GFGVDSFTLSTLLSSCTDSSVLLRGRELHGRAIRIGLMQELSVNNALIGFYSKFWDMKKVESLYEMmeKNTITYNALMAGFCRNGhkALKLFTDMLQRGVELTDFSLTSAVDACGLVSEKKveQIHGFCIKFGTAFNPCIQTALLDMCTRCERMADAEEMFDQWPS-NLDSSKAT-TSIIGGYARNGLPDKAVSLFHrlCEQKLFLDEVSLTLILAVCGTLG--------FREM----GYQIHCYALKAGYFSDISLGNSLISMYAKCCDSDDAIKIFNTMREH------DVISWNSLISCYILQRNGDEALALWSRMNEKEIKPDIITLTLVISayTESNKLSSCRDLFLSMKtyDIEpeHYTAFVRVLGHWGLLEEAEDTINSMP----VQPEVSVLRALLDSCRIHSNTSVAKRVAKLILSTKPepSE--YILKSNIYSASGFWHRSEMIREEMRERGYRKHPAK----------SWIIHENKIHSFHARDTSHPQEKDIYRGLEILIMECLKVGYEPNT---------------------    
 583 UniRef50_C5Y0A6        Putative uncharacterized pr...   369    1e-100  1  11.6%  14:600    51:622     -------------GLARDLFDEFPSPTPRLANALLRAHIRARQWRAAILL-----GPRLRVRLDGFTFSLLLRACAALPSLAHGRAVHAVAIRSCTASefVATAIVQMYARCGDMVGAINAYGVLEKPDIVLRTSVVTGYEQNgeEALEFFArvVGQGVMLTPVTLVSAMSAAAqgHVRKGQACHAYVVRNSLGYDLALVNTVLSFYVKIGDFQASMRLFEGMTDR----DVITWSCMIKGYVQHGDAHEGLRMYREMVKARVQPNSVTLVSVLQACALVVDAEEGKRV------------HRVAVSIGCELEVGVATALVDMYMKCSCHEEAMCLFHRMPKK------DVVAWAAVIGGLTQNELPGESLHVFKCMLLDDHVPDAITMVKVLAACSEFGGTRLAICLHGYLVRNGFNNNAfaALLDLYSKCGDIDSAVRVFE-----GTTEKDIVVWGSMIAGYGAHGLGQEAVALYQRMIASSIQPNSVTFVSVLSACSHSGLVQEGIQIFDSMTQVFGVVPNAEHQSAMVDLLGRAGELQEAIRF----IRGMDGRAVAHTWCALLAACREHNNTkpDHVGYYNLLTNIYAYDEKWE---    
 584 UniRef50_Q9C9U0        Pentatricopeptide repeat-co...   369    1e-100  1  10.2%  14:601   126:777     -------------ERVLRVFRFfyVPNVIHYNIVLRALGRAGKWDELRLCWIEMAHNGVLPTNN--TYGMLVDVYGKAGLVKEALLWIKHMGQRMHFPDEVTMATVVRVFKNSGEFDRADRFfdDFPKNGSAQseLFKVGARNpeKSLHFASGSDSSPRKPrtSTFNTLIDLYGkgRLNDAANLFSEMLKSGVPIDTVTFNTMIHTCGTHGHLSEAESLLKKMEEKGISPDTKTYNILLSLHADAGDIEAALEYYRKIRKVGLFPDTVTHRAVLHILCQRKMVAEVEAVIAEMDRNSIRIDEHSvfQLDCVLSSTTLAAVIDVYAEKGLWVEAETVFYG-KRNMSGQRNDVLEYNVMIKAYGKAKLHEKALSLFKGMKNQGTWPDECTYNSLFQMLAGVDLVDEAQRILAEMLDSGCKPgtYAAMIASYVRLGLLSDAVDLYEAM-EKTGVKPNEVVYGSLINGFAESGMVEEAIQYFRMMEEHGVQSNHIVLTSLIKAYSKVGCLEEARRVYDKMKDSEGGPDVAAS-NSMLSLCADLGIVSEAESIFNALREKgyKGMGMLDEAIEVAEEMRESGLLSDC-TSFNQVMACYAADGQLSE--    
 585 UniRef50_A2Q4A7        Tetratricopeptide-like heli...   368    1e-100  1   8.3%  11:525    61:548     ----------SNLYYAHKLFDTMPnsNCFIWTSLIRAFLSHhhFCHCISTFARMHQKGIL--PSGFTFSLVLNACGRVPAGFEGKQVHARLVQSGFLGNKIVQTALLDMYAKCGHVCDARDVFDGidRDVVAWTAMICGYAKAGrdARFLFDNMGER----NSFTWTTMVAGYAngDMKAAMELYDVMNGK----DEVTWVAMIAGYGKLGNVSEARRIFDEIT---VPWNPSTCAALLACYAQNGHAREAIEMYEKMRRAKIKVTDVAMVGAISACAQLRDIRMSNSLTYNIEEGFCEKTH------------IVSNALIHMQSKCGNIDLAWREFNIMRNR------DLYTYSAMIAAFAEHGKSQDAIDLFLKMQQEGLKPNQVTFVGVLNACSTSGLIEEGCRFFQIMTeyGIepEHYTCMVDLLGRAGQLEKAYSLIKE----NSTSADATTWGSLLAACRVYGNVELGEIAARHLFE--IDpdSGNYVLLANTYASNDKWECAEEVKKLMSKKGMKKPS------------------------------------------------------------------------------    
 586 UniRef50_Q0IWY4        Os10g0479200 protein n=6 Ta...   368    1e-100  1  10.8%   3:596   158:793     --ANS-YARAQMVHDALYVLSKMSSLnvFTYDSLLHGL---RMTDVALELFEEME--SCGVSPSEYSHSIIINGLCKQDKVGEALSFLQEARKEgfKPLGMTFNILMSALCNWGFVQSAKSFLCLMLKYGLVPDRYTFSTLIHgeEALDLFERVTKEGMELEIVTYNSLINGYRLLGlkEIPKIIQMMRGQGVEPDLVTYTILIAGHCESGDVEEGMKVRKDVLDQGLQLNIVTYSVLLNALFKKGMFCEIDNLLGEIYNIGLDMDVIAYSILIHGYCKLGEIEKALQVCNAMCSseARWYLENVARKYQPTDVVFYNVVIDGYAKLGDIVNAVRLYDQITVAG--MHPTIVTCNSLLYGYCKIGDLQLAESYFRAIQLSGLLPTAVTYTTLMDALSEAGEVNTMLSLFDEMVAKRIKAntYSVIVKGLCKQLRFDEAINVLKDMDSKGI-NADPITYNTLIQGFCESENVQMAFHIHDIMLCRGLVPTPVTYNLLINVLCLKGKVIQAEILLESLRENGIKL-RKFAYTTLIKAQCAKGmiNAVLLVGKLLDAGFEASIEDFSAAirLCKrmLSVGIYPDT-QIYCVLGRALQKN-------    
 587 UniRef50_Q9SJG6        Pentatricopeptide repeat-co...   368    1e-100  1  10.3%  56:569    15:530     -------------------------------------------------------AMPSSGSLSGNTYLRLID--TQCSTMRELKQIHASLIKTGLISDTVTAsvLAFCCASPSDMNYAYLVFTRihKNPFVWNTIIRGFSRspEMAISIFIDMLCSsvKPQRLTYPSVFKAYGrgQARDGRQLHGMVIKEGLefDVVAWNSMIMGFAKCGLIDQAQNLFDEMPQR----NGVSWNSMISGFVRNGRFKDALDMFREMQEKDVKPDGFTMVSLLNACAYLGASEQGRWI------------HEYIVRNRFELNSIVVTALIDMYCKCGCIEEGLNVFECAPKKQLS------CWNSMILGLANNGFEERAMDLFSELERSGLEPDSVSFIGVLTACAHSGEVHRADEFFRLMKEKyiEPshYTLMVNVLGGAGLLEEAEALIKNMP----VEEDTVIWSSLLSACRKIGNVEMAKRAAKCLKK--LDPDETcyVLLSNAYASYGLFEEAVEQRLLMKERQMEK----------EVGCSSIEVDFEVHEFISCGGTHPKSAEIYSLLDIL----------------------------------    
 588 UniRef50_Q69XF8        Pentatricopeptide (PPR) rep...   368    1e-100  1   8.8%   3:525   126:632     --ANALLAFyrHLPHALRAFDDLPRRDVVAHSSILAAFLRAGMPRRALASLRDMLaaDDDVSPNAHALSAAVKACAVLRDRNAGACLHGSVLVRGFGDDGVVLSSLVDMYGHVAAPGDARKVFEEMrpDGICYTSLISAFVRnfEEAVRWFRSMLMmgVRPDGCTFGSMMTALGNSKrsQGRQAHAQVVTRGLCGNVIVESSTLDMYAKCGLMVEARKVFDRMQVR----NEVSRCALLGGYCQNGEYEKVIALFREmkEDG---DWYSLGTVLRACAGLSSVKPGKEIHCRFLRMAGWR------------DVVVESALVDLYAKCGAVDYAYSVFE------ASTVRNTITWNAMIGGFAQNGHGERAINLFNRMVREGPRPDYISFIGVLFACSHTGMVEQGRNYFNSmkDYGIApeHYNCMVDLFSRVELLEEAEDLINKSPFRN----DSSLWADILGASATHSNPDVAERVSKKMME--LEPqhLSYILLENVYRTVGRWEDALEIRRLMESRKVKKEP------------------------------------------------------------------------------    
 589 UniRef50_D1HHF7        Whole genome shotgun sequen...   368    1e-100  1  10.3%  96:580    21:479     -----------------------------------------------------------------------------------------------VDPNLIAVKLIGACADHANVRQAALIFAHLASPNIFAHNATLKALAQNShhAIQFFNHQVSSppNPDEFTFTSVLKACalAHVVNGQKIHAMVTKQGFESNLFVRNSLIDMYFKAGYLLLARHLFDEMFVRD--RNVVCWNAMIAGYAQNEKYSDAIEVFRMMQQFgvVPNDVTLVSVLPACAHLGALDLGKWI------------DGFISRREMALGLFLGNALADMYAKCGCITEARRVFNKMEER------DVISWSIIICGLAMYGHADEAFGCFYEMLDCGEKPNDVVFMGLLTACTHAGLVKKGLNCFNTmkEYGVSpeHYGCVVDLLSRAGELDKAEDMISSMPMK----PNVIIWGALLGGCRIYRDSGRGQRVVQHILELDSD-HSGSYVYLANVYSSMGRLDDAAKCRLRMRENGVLKTP----------GCSWIEVDNTVHEFFMGDLSHPESNKIYSMIRELMWKMKLAGYKP-----------------------    
 590 UniRef50_Q67X39        Putative uncharacterized pr...   368    1e-100  1  10.1%  11:523    47:500     ----------GRMGYARKVFDGIPRPDLFMHNAMVRGYAHAGAPGDAFAVYRRMTEA-SRLRPDAFTFCYLLRACAGLPGSRAGRQVHGAVVKLGFLKDAYVRNALINMFAKCGDLRVASVLLDEaeGDVVAWSAVIAGHA--------------------------------ARGDMAAARKMFDECTHK----DIVCWNVMLGAYAKHGEMENARELFDRAPEK----DVVSWNTIITGYAAQGMLKHALEVFDEMRAAGWTPDEATIVSLLSCCANTGLLDAGRMIHHQLHLERRPWI-----------SIVVGNALVSMYAKCGDLHTAVEGFNTMKD------TDVWTWNSVIGGLAFHGQAEQSVRFFKKMLEKRIHPNEISFLCVLGACSHAGLVEDGQKFFYLMkrYGIEPnhYSCIVDMLGRAGLLDEAYAIVSNMR----CEPSAVVWRTLLGACRTHGNMTLGKLVREKLLNMNEDASAD-YVLLSGIYASSGEWLGVETERRSMDRRGIRK--------------------------------------------------------------------------------    
 591 UniRef50_Q9LND4        Pentatricopeptide repeat-co...   368    1e-100  1   9.4%  25:523    69:548     ------------------------KRNRHSWNTILSGYSKscCYSDVLLLYNRMRRHCDGV--DSFNLVFAIKACVGLGLLENGILIHGLAMKNGLDKDDYVAPSLVEMYAQLGTMESAQKVFDEIprNSVLWGVLMKGYLKYSkeVFRLFCLMRDTGLALDALTLICLVKACGngKV--GKCVHgsIRRSFIDQSDYLQASIIDMYVKCRLLDNARKLFETSVDR----NVVMWTTLISGFAKCERAVEAFDLFRQMLRESILPNQCTLAAILVSCSSLGSLRHGKSV------------HGYMIRNGIEMDAVNFTSFIDMYARCGNIQMARTVFDMMPER------NVISWSSMINAFGINGLFEEALDCFHKMKSQNVVPNSVTFVSLLSACSHSGNVKEGWKQFESmrDYGVVpeHYACMVDLLGRAGEIGEAKSFIDNMPVK----PMASAWGALLSACRIHKEVDLAGEIAEKLLSMEPEKSSV-YVLLSNIYADAGMWEMVNCVRRKMGIKGYRK--------------------------------------------------------------------------------    
 592 UniRef50_B9IQQ4        Predicted protein n=3 Tax=r...   368    1e-100  1   9.0%   2:577   153:728     -ALLGLFGRCGWLDEAFHVFEDMPDKSLVTWNSMISLLGHHGFVEDCVVLFRKLV--RKEGSLSKCSFEGVLSGLVCEEDLEFGGQIHGLVIKSGLDCEVLVSNSLINMYARRSSMSQVEKLFEEVdrDVVTWNTIISAFSknPGKALEVFLKMSEDGIMPNQTTFVSVINSCTSLLVPmgEYVHGKIVKTALETDVYLGSALVDYYAKCGKLDNAHYCFREIHQK----NVVSWNSLILGYANKCSFASV-SLLLEMLKLGFRPNEFSFSAVLKS-------SLVLEL---------KQIHSLTIRLGYENNEYVLTSLITSYGRNGLITDAlqYFETLKFLSQLEEPDTVSWNIVIAACARNGNYNEVFELFKHMRVAQMLPDNYTYTSLLCVSSKVCNLALGSSIHGLLIKTNfdIVVRNVLIDMYGKCGNLESSAKIFDSMTER-----NLITWTALISALGINGCAQEALERFNDMEFLGSRPDKVAFIAVLTACRHGALVREGMQLFGKM-NNYHIEPDMDHYHCLVDLLARNGHLEEAEKV----ISCMPFPPDAQIWRSFLEGCKKRRN--------------------------    
 593 UniRef50_Q6K9G0        Os02g0620800 protein n=4 Ta...   368    1e-100  1  11.9%  61:574     3:524     ------------------------------------------------------------PAPPFSSHGDFVAALHRCATLAHLKQLHAHAVVTGRAAatTTFHLLRFASLRLSCLPYARRLFDA----TPSPNVFLYSAMlhARDSLALFLRMLRRGRppNQFVYPLVLRAACAIGVQLVRSIHCHACKDGFYGHDFIRTSLLDGYSRYGMMGDARKLFDGLTDR----NVVSWTALVSGYARAGKVGDAIVLFERMPQR----DVPAWNAIIAGCTQNGLFVEAVGIFRRMVDEggKVIHGYAWRSCVGFGSSVVNGLIDMYGKCGNLMEAKWIFDAFSDRG------LTTWNSLINCLALHGCSESAIAVFNSMRNEGVQPDEVTFVGLLNACTHGGFVDEGLRYFELMceHGIEpeHYGCVVDLLCRAGRFQDAMNFINDMK----VQPDEVIWGSLLNACRIHRHLELAEHAIRNLLDLN--PSNANYVVMlnLYSEGGFWEEVRKVRKLMKEdgKKLP-----------GCSWIEVDRKTRRFYSGDDGHPESEDIYDTLDKLATTME-----------------------------    
 594 UniRef50_B9SIR2        Pentatricopeptide repeat-co...   368    1e-100  1  11.5%  11:602    82:790     ----------SALSFFNQLKDSGFKHDISTYAAIIRILCYWGLHKQLRSIfsHFLDTLSDGFVDVDsyDALVKAYVSVGMFDDAIDVLFQMGRRRFVPHIFICNFLMNSLIKNSKLDMALAVYKQLKRLGLSPNDYTYAIVIKaeEAMYVIKEMEESGITPTGFAYTAYIEGLceLKFDKAESVLRDMEKEGMVPDMHCYTALICRFCKAGNLLKAYAFLNEMMSKGVKVNCVIVGSILHCLCELGMHSEVVDQFNQFKSLGLFLDGVSYNNVVDALCKLGKLEEAITLLDEMKMKQ------------INMDVMHYTTLINGYCCQGNVVDAFKVFEEMRENGIEI--DVVTYDVLVSGFCRNGLATEALNLLDYMQTQKLKPNSITYNVVVESLCMGGKVKEAEAVFNSIEDKSL--DNYFAMINGYCKANHTAGAAKLFFRLSVKGHVKRS-CCYN-LLKNLCEEGDNDGILMLLETMLNLNVEpdLIAYTIMITSYCRMNCLKEAVDLFHDMKQRGIKPDLVthHKAHIKKVYSAANAKGGNeiFYTVLIDGYCKVDSLHDAIGVFDEMIERGLEPDII-TYTALLSGCCQRGDVDRA-    
 595 UniRef50_D1H3V2        Whole genome shotgun sequen...   368    1e-100  1  11.5%   6:500   121:592     -----MYAECSCFVEAWRVFDGMPHRNSFSWTVMIAGSKKNGLFLDGFRFF--YDMLVDGILPDEFVYSAVIQSCIGLGCIELGEAVHGQIVKRGFWDDVIVGTSLLSMYAKLHNSEASVRVFNAieHNQVSWGAVISGLSSNGleAFHQFLAMITQGFTPNMYTFSSVLKAVGkrDATKGREVHHCVMEYGMESNVVVGTSLIDMYSKCGHLSDARSVFDryKSKVNNP----WNAMISSYTQCGYWQEALDLFIEMSLNDVKPDLYTYGGVFSAIAALKWLYFGRQVHGMVVKSGNGSRVLSLN-----------NAIVDAYFKCQSLEDARKVFDRMQER------DMVSWTTLVSAYVQCYQPSEALSIFSQMREQGFMPNQFTFSSVLVACASLSLLEYGRQVHGLICKAGLDDDNcsSLTNMYAKSGNIIDAVEVFEK-----IVCPDVVSWSAIIYGYAQHGFLDKAVELVQKMEQSGIQPNS---NILLSHL-------------------------------------------------------------------------------------------------------    
 596 UniRef50_C5Y600        Putative uncharacterized pr...   368    1e-100  1  11.5%  35:598    61:649     ----------------------------------IASLTSVADNEGSAAALALFRRLSTRPDLRHLmyATAATLFVARGNLPMAHemLLELRSHGLPLVvtANWVLRVGLRQRHHFAHARQAFDGmrPDARSFRALVLGCcgRFEEADALLAAMWREGFSLDSATCTVVVRAFCrgRFRDVPGLFGRMVEMGNPPNVVNYTAWVDGLCKRGYVKQAFHVLEEMVGKGLKPNVYTHTSLIDGLCKIGWTERAFRLFLKlkSSSYKPNVHTYTVMIGGYCKEGKLARAEMLLGRMVEQ------------GLAPNTNTYTTLIDGHCRGGSFDRAFELMNKMKLEG--FLPNIYTYNAIIGGFCKKGKIQQAYKVLRMATSQGLCPDKVTYTMLITEHCKQGHITYALDLFNQMAENSCHPdtYTTIIAMYCQQRQMEQSQQLFDKCLSIGLV-PTKQTYTSMIAGYCRVGKSTSALKVFERMVQHGCLPDPITYGALISGLCKESRLEEARALFETMLDKHMVPCDVT-CVTLAYEYCRRDklDGLDkhAADALVRKLSAVDNLDAASLFLKNVLDKHYAVDH-ATYTSFINSCYESNR-----    
 597 UniRef50_A5AIJ1        Putative uncharacterized pr...   368    1e-100  1  10.4%   4:579    80:636     ---NTLLSFYGNCgdAGRVFDEMPEKDLVSWNTMIGVFSVNGCWXEVLDLFGEMR-LRSGLRPNVVSVVSVLPVCAGVEDEVTASEIHGYVVKVGLEFQVIVGNALLDVYGKCGNVAALKQVFGEmeKNLVSWNAIifGYKGHYRDALDMFRLMIDEGLKPNSITISSFLPVLVELEFFKarEVHGSSIRMGLESDIFIANSLIDMYAKSGHSTEASNVFYKLDAK----NVVSWNAMIANFAQNRFELVAVGLVRQMQDYGELPNSVTFTNVLPACARMGLVRPGKEI------------HARSIHMGCAFDLFVSNALTDMYAKSGHLKLARNVFD-------TSLRDEVSYNILIVGXSQTSDCSESLSLFSEMQLMGLKQDNVSFMGALSACANLTAIKQGKEIHGFLLRkhIHLFVANSLLDFYTKCGRIGLARNIFDRMTNK-----DVASWNTMILGYGMLGELDTAIDLLTemRKDDVEsdSVSFIAVLSACShaGLMEEAAELIKGLP----IVPDANIWGALLGACRIYGNLELAAWAAEHLFELKPekTGRWDEANRIRELMKSRGVK------------------------    
 598 UniRef50_Q8LQG2        Os01g0914600 protein n=9 Ta...   368    1e-100  1  10.5%  25:569    79:582     ------------------------SRHAFPWNTLVRLHAAAS-PRRSLLYFSRMR--RAAVAPDAYTFPAVLKACGCAPGCRVGLVVHGEAVRTGLDADLFTRNALISFYCR----------------------------IGDCRSGRKVFD----HGVR-DLVSWNSMVAGygCGEVDLAQDLFDEMRQR----DAFSWATMIDGYGeaGGVDRARELFDQMPDR----DLVCWNSMIDGYARHGRMDEARVLFEEMPER----NVISWSIVIDGYVRFGEPNEALEFFQRMLRCggRWLHSYLEKKKVLFDVVVQTALIDMYVKCGRLDLAKLIFESMPKK------SVVTWNVMIVGLGTHGYGLDAIKLFNQMETERAPMDDLSILAVLTSCTHAGLVSEGLGIFYRMEKdgLEpeHYGALIDLLGRAGRVDQARNTIETMP----MEPTPELWGSLLASCRSHRCVELAELSVERLASLGAD-DSGVYVLLSNIYADEGMWDDVFRIRKLMSAEGMKK----------NIGRSVIEVDGQIHEFVNGGSSHPHKEEIYLTLWNL----------------------------------    
 599 UniRef50_B9H9T3        Predicted protein (Fragment...   367     1e-99  1  11.7%  18:582   186:771     -----------------RMMKLNCLPNVATFSILIDGLCKSGNVKDALHLFDEMTQ--RGILPDAFTYCVVISGLCRSKRVDDAYRLFDKMKDSGVGPDFVTCNALLNGFCMLDRVDEAFSLLRLFEKDGYvySCLIRGlaKRYEDVQLLYRKMIEDNVKPDVYLYTIMMKGLAegKVRDALELLNEMTESGVVPDTVCYNVLIKGFCDMGLLSEARSLQLEISRHDCFPNVKTYSILISGMCRNGLTRDAQEIFNEMEKLGCYPSAVTFNSLIDGLCKTGQLEKAHLLFYKMegRNPSLFLRLSQGPSHVLDSASLQKMVEQLCDSGLIHKAYRIL--MQLADSGDAPGIYTYNILVNGFCKLGNFNGAYKLFREMQFKGLSPDTVTYGTLINGLLRFQREEDAYKVFDQMEKNGCTPdvYRTMMTWMCRRMELPRAFSLWLKYLRNIRSQEDEAI--KAIEGYFEKQEVEKAVRGLLEMDFKLNDFDLGPYAIWLIGLCQTRRVGEALKIF-LILEEYKVVITPPCCVKLIYFLLKEGDLDRAIDVFLYTIEKGYLLRRRvdRAIYLLCRMKSVGYDLDA---------------------    
 600 UniRef50_C3SA54        Vegetative storage protein ...   366     1e-99  1  10.8%   3:574    28:570     --AAALVHMYAKC---RGMVGsfSEPDMVLRTSMVTGYEQNRMAAEALEFFSR-HVVGQGFMPSPVTLVSVISAAAQLKDVLNGQACHAFVIRNNFEYDLVLVNAILGFYMRIGAVQAARRLFEGmeKDVVTWSCMVTGYVQSGdeALTAYKKMVEAGIKPNAVTVVSVVQACSLadIEEGRRVHDIAVKIGCELEMTVATALVDMYMKCSCHEEAMQLFCRMSKK----DAVAWAVVISGFTQNGLPDESMRVFKCMLFGGPFPDAVTMVKVLAACSESGVMRQAF------------CLHGYLVITGFCDKIFVAAALVDLYSKCGNLGCAVRVFESAMEK------DVVLWSSMISGYGVHGLGQQAVELFQMMVASSVKPNSLTFVSVLSACSHSGLVQEGKRIFESMTqyGVVPnhHSAMVDLLGRAGELQEAAKLLH-----GNGRADAHTWCALLAACRAHHDTEMSEVVAAKLLK--LDPDHAGYYNLlnIYTFDENWSSAKENRNIIRDRGLNKVP----------GCSAVELNNVMHTFTAGERPHKDWEKISRLLREFSRTLK-----------------------------    
 601 UniRef50_C5XGF1        Putative uncharacterized pr...   366     1e-99  1  12.2%  71:578    14:492     ----------------------------------------------------------------------LLSFLPRLTSPRHLLQAHAFLLPRGGHRHPRLVSALILASLRlqTSDHAAALLRR-----VHPSVYLRAAARLPSHLLrlHSLLLRTgvASDTHVSASLVQLYCTCrvADARSVFHEMAVR----DVVAWNVMIAGYVKAGDLAHARELFDAMPER----NVVSWTTVIGGYAQMRRPEEAVEVFRRMQVEGIEPDGVALLSVLSACGDLGAVDLGEWVHRFVVRRGLRQEIPLM------------NSIIDMYMKCGCIEKAVEVFEGMEEK------SVVTWTTLIAGFALHGLGLQAVEMFRRMERENMAPNDVTFLAILSACSHVGLTDLGRWYFNIMVsyRIKpeHYGCMVDLLGRAGCLKEAQDLVKDMPLK----ANAAIWGALLAASRTHGDADLGEEALVHLIE--LEpnSGNYILLSNIFAEQERWDDVSKLRKAMKERGLRNVP----------GASSIEVDGMVHEFTSRDGSHPFLHKICEVLCEINTNMKSVGF-------------------------    
 602 UniRef50_Q75IC8        Pentatricopeptide, putative...   366     1e-99  1  10.9%  62:585    61:570     -------------------------------------------------------------DPTPRAFSALLKAAsrPSSLGLGAQLHAQAVVRGfdDSTILATAVLSFYASCREPDLARKVFDGmrRNAVTWNALIKGYAQAgeEAILLFRDMKREgvAPDRYTFPALLSGIGREglELGGALHAHVIKAGLERDPFVGASLVSLYAARRTLEDAKVAFDQVGSS----DPIVWSSMISAYVNCEEEEGALLIFFNMLCQDIKPTQFVYSTVFSVCGRMGILEMGKQV------------HAHSLKSNTEKDAAMFNALLTMYSDCGCINDAQKVFS------SNDCVNVISYNSMISALGQHGYPKEALEHFRQMKFAGLMPDEVTLLNLISSFNHAGLVHEGLQMFNSMVdeGIKpqHYACVVDMLARSGEIGKAMKTINEMP----FEAEAPLWRIVLGACSKHRDIETGKRIAEMLFE--MEPYEAtyILLGNIYARLGRWTEAEKVRSLMGERGVYKDDAF----------SWIEMGQRTYRFGVDDRSHPISREIYRNLDRLISTIKVAGYVPDISFA------------------    
 603 UniRef50_Q9SUF9        Pentatricopeptide repeat-co...   366     1e-99  1  10.4%   3:600    43:663     --ANNVISMYVdlSDAHKVFDEMSERNIVTWTTMVSGYTSDGKPNKAIELYRRMLD-SEEEAANEFMYSAVLKACGLVGDIQLGILVYERIGKEnrPSSTSWNTLISGYCKAGLMDEAVTLFHRMPQPNVVSWNCLISGFVDKGsrALEFLVRMQREGLVLDGFALPCGLKACSFGglTMGKQLHCCVVKSGLESSPFAISALIDMYSNCGSLIYAADVFHQ-EKLAVNSSVAVWNSMLSGFLINEENEAALWLLLQIYQSDLCFDSYTLSGALKICINYVNLRLGLQV------------HSLVVVSGYELDYIVGSILVDLHANVGNIQDAHKLFHRLPNK------DIIAFSGLIRGCVKSGFNSLAFYLFRELIKLGLDADQFIVSNILKVCSSLASLGWGKQIHGLCIKKGyePVTATALVDMYVKCGEIDNGVVLFDGMLER-----DVVSWTGIIVGFGQNGRVEEAFRYFHKMINIGIEPNKVTFLGLLSACRHSGLLEEARSTLETMKSEYGLEPYLEHYYCVVDLLGQAGLFQEA----NELINKMPLEPDKTIWTSLLTAchKNAGLvpDDPSVYTSLSNAYATLGMWD---    
 604 UniRef50_D1HEE7        Whole genome shotgun sequen...   365     2e-99  1   9.7%  69:534     4:442     --------------------------------------------------------------------TSLVCMYAKCGNVVDSRKVFDYMPERN---AVTWNAMICGYLGNGDSKSAVLLFEKMSIRTAVTWIEMIDGFARSGDtaRRFFDDVPSE--LRNVVTWTVMVDGYARneMEAAREVFEGMPQRNF----FAWSSMISGYCKKGNVKEARSIFDRIPVR----NLVNWNSLISGYAQNGFSEEALEAFGKMQAEGFEPDEVTIASVLSACSQLGLLDAGKKIHHMMNHK------------GIKLNQFVLNGLVDMYAKCGDLANARLIFEGMAHR------NRACWNSMISGFAIHGQSKEALEFFGRMEDSHEGPDEITFLSVLSACAHGGFVNAGLEIFSRMEKYGLtiKHYGCLIDLLGRAGRIKEAYDLIKRMPVK----PNDVVWGALLGACRVHLDMEMADRVVEEIVKVDSNISSghYVLLSNIYAASDRWEKAEKMRMEMANKGFQKTSGCSSIMPGN---------------------------------------------------------------------    
 605 UniRef50_C5WVQ1        Putative uncharacterized pr...   365     3e-99  1   9.2%   2:602   382:964     -ALVSMYSKLGKLDSSSLLFCCFTEKNNILWNSMISGYLVNNEWNMALDAFCKMQIAG--VAPDATTVINVISGCRYTKDLHVAKSIHAYAVRNRFESYQSVMNALLAMYADCGDISTSYTLFQKMerMLISWNTMISGFagDSETSLTLFCQMFHEEVWFDLVTLIGLISSLssEDAIVGESVHSLAIKSGCISDVSLTNALITMYANCGIVEAGQQLFN----SFCSRNTITYNALMSGYRKNNVSEKILPLFTQMVKNDEKPNLVTLLNLLPVCQS---------------QLQGKCIHSYAVRNFTRLETPLFTSAMGMYSRFNNIEYCRTIFSLVSAR------NLIVWNAFLSACVQCKQADMVVDYFKHMLFLNVRPDEVTMLALISACSQLGNADFAACIMAVILQKGFSMnvLNALIDTHSRCGSISFARELFDSSVEK-----DSVTWGAMINAYSMHGNGEAALDLFSMMIDSGVDPDDITFVSILSACSHNGLVEQGRTLFKSLQADHGITPRMEHYACMVDLLGRTGHLDEAYDIVRSmlLGACRFHGNYKIGESVGKLLIKSEY--GKSRSY-VMLSNIYASAGKWSD-    
 606 UniRef50_UPI0000DF05BD Os02g0191200 n=1 Tax=Oryza ...   365     3e-99  1  10.4%   4:580    40:588     ---NSLVNCLehLRYALHLFDRMP-PSTFLFDTALRACSRAgdPHRPFLLFRRMR--RAGVRPDGFTFHFLFKCSSSSSRPHSLlmLHAACLRTMLPspFVSNSLIHMYIRLGLAADARRAFDEIHVKDAVAWTMLISGLAKMgcDTQLLLSQAPVR----DVISWTSLIAAYSrnRAREAVGCFKTMLSHGIAPDEVTVIAVLSACAKLKDLELGRSLHLLVEEKGMPTSE----NLVVALIDIGQLREALLLFMSMRRHDLRVDNFTVVNLLTACASLGALQQGRALHACIEQRL------------VEADIYLGTALLDMYMKCGRVDEATIVFQRMGKR------DVHTWTAMIAGLAFNGMGKAALEHFYQMRCDGFQPNSVSYIAVLTACSHSCLLNEGRLYFDEMrlYNIHpeHYGCMIDLLGRSGLLDEAMDLVKTMP----IQPNAVIWASILSACRVHKHIDLAQCAAEHLLK--LEPDegVYVQLYNIYIDSRQWENASKIRMLMEERQVKKT----------AGYSSITVAGQVHKFVVSDKSHPRILEIIAMLEEISHRLKSLGYSP-----------------------    
 607 UniRef50_D1ISB2        Whole genome shotgun sequen...   365     4e-99  1  12.1% 159:587    71:468     --------------------------------------------------------------------------------------------------------------------------------------------------------------LTPTYSNYASLLQScaRKAIKPGKQLHAQVCLAGFGFDTVIATKLVNLYCVCDSLSSARLLFDRIPKH----NIFLWNVLIRGYAWNGPYEAAVQLYYQMFDYGLVPDNFTFPFVLKACAALSAIEHGREI------------HEHVVQTGWEKDVFVGAALIDMYAKCGCVGSAREVFDKILVRDAVL---VVSWNAMITGYAMHGHATEALDLFEEMNRV-AKPDHITFVGVLSACSHGGLLEEGWMFFETMIryKIDPthYTCMVDLLGHSGRLDEAYNLIMQMK----VLPDSGVWGALLNSCKIHANVELGEIALERLIE--LEPdaGNYVILSNIYAQAGKWEGVAKLRKLMTDRRLKKSIA----------CSWIEVKNKVHAFLSGDTSHPLSDEIYSELERVGGLMKEAGYSPSTGSVFH----------------    
 608 UniRef50_B9HA38        Predicted protein n=6 Tax=M...   365     4e-99  1   9.6% 154:591     1:406     ---------------------------------------------------------------------------------------------------------------------------------------------------------MLERKKLPNKFTYPFVLKACAggNLNLGKSVHGSVMKFGFGDEVNVQNTLVHMYCCcgGIEFARKVFDEM----YKSDSVSWSAMIGGYVRVGRSSDAINLFREMQIKGVCPDEITMVSVLSACTGLGALELGKWV------------ESYVEKERVQKNVELSNALIDMFAKCGDVDKATNLFRSMRER------NIVSWTSVIGGLAMHGRGVEAVAVFEEMVRSGVTPDDVVFIGLLSACSHSGLVDKGKRYFDSMRKdsIVpeHYGCMVDMLCRAGLVKEALKFVQEMP----IDPNPVVWRTLINACRAHGELKLGEKITRQLIRNE-PMHESNYVLLSNIYAKMSDWEKKTRIREAMDMKGMKKIPGSTM----------IELDNEIYEFVAGDKSHAQSKEIYEMVDEMGKEMKRAGYMP---TTTEVLLD------------    
 609 UniRef50_Q0J946        Os04g0672700 protein n=5 Ta...   365     4e-99  1   8.1%   6:574   165:706     -----MYSQCGAPRDANQVFDEMATRDGVAWNCVMHCNVRYGYLDRALGQFCRMV--RGGLPPTESTLSSVLSGCGRAGDCLRGRVLHGWVVKLelDPDMPLQNALLDMYSSCGDLDTALRVFERIepDLVSWNTLIAGfvGDGCSAVHAFVQLkdERVVPDEYTLAAVVSASATlaMFGGKPLHAEVIKAGLENSVFVGNTLLNMYFTNDDPHSARILFDSITQE----DVIMWTEMVAGHSSLGEGELALRYFVSMLQEGYKVDSFSLSSALNSTTELAGLKQGEML------------HAQVVKSGYEGNICASGSLVDMYAKNGALPGAYLVFRNIQR------PDLKCWNSIIGGYGNHGNSEMAFKLFGEMIRDGLQPDHVTYISILSACSHCGLVEKGKFYWFCMMTDGIVPghYTSMVSLLGRAGLLDEAVDL---MMKSPFAKKCPELWRILLSSCITFRNLSIGVHAAEQALEQDPD-DISTHILLSNLYASLGKWDFVAEIRKRM--RGLMADKEPGL--------SWVEMKNVVHVFSADDECHSHIDDCHSELLRLKGNME-----------------------------    
 610 UniRef50_D2V8U1        Predicted protein n=1 Tax=N...   364     4e-99  1  12.0%   7:582   158:708     ------YTKCGHTNVALQLFEDLKktNAISWNSAIQALVLAKDTQKAIDLLAEMQLLN--VKPDTITFTILIGLCAENQYYKEGVKLHEMIKVLNISSLELSNSLMN-FYTKCGQPDVALQLFEqmKMNVVSWSCAIQALgkDIRGAIGLLEEMQANNIKADTILFTILLTLCADNQleEGVKLHEIIKVLKIS-SLELSNTLMNFYTKCGQTSISLEIFEELKKEN-KTNVVSWNSAIQALRSRKDITRAFELLEEMKKLNIEPDEITYVSLLTLCADSVAIKQGEEVVQQFKTKHIAK------------SVFWIGATISFHAKVGRLERAESVFNEYLSNNPPFDPILTS--VMIDAYAKHGKGKEAVYLYQQIIDRNQLPSAEAVTSVLTACSHAGLVDEARRIYKSIENTqcTPSHKACMVDVLARSGYLEEAEQFI-------GSSRDVVSWVTLLGACRIHTNVEIAERTFDILLEIDNE-QASHYVQMSHIYNQMGQIDKANEVRELMKRNNATKIPGIT----------TIEVDGETHNFVSDDSYHPRMQEINQELEKLNKEMLDAGYAPDT---------------------    
 611 UniRef50_D1ILY2        Whole genome shotgun sequen...   364     5e-99  1   9.9%   3:576    80:701     --ANSLMDMYAKCgdAAKLFDHMPDKTVVSWTSMMSGHCQRGAFDEVISIFWRMLE---TLQPNEYTLAVILQACAQKRDLKLVQLIHCHIIKTGFVMDAFlnSLIDGYTKSggMVEKALLFFFEMQEDGVSPNTvmYSINGYFKEGFQIFCNfeGDGQYLSTETIATLLQGCSHSklKLGEQIHGYQIKHGFFPCTIVENSLIYMYAENERDDAAFQLFRKMSCR----DIVSWNTMISSLVKGSSSYQALMLLSEVHSNgiYPDFVTILASIQACSSLASLQLG------------QVIHGYITRAGLICDIFVQNSLVDMYGKCGRLHLAEKVSEEMPVR------DLGSWNSLIAAYGINGNGISALNVFKQLKNTghRPNAITFTNILSACAHAGLVAEGFEIFKSMkeYSLEpeHFACMVDLLGRAGRLEEAEAFIQKMP----FEPGPEVWGALLGGCGLFGNLDIAERVAKKL--YILEPKSRAwvALSNVYASVNKWEDAAKVRAEMRR-------SEELQK--EGGWSSVEVRGQEFRFMVGDTMHPEARMVYAVLKGINEHIRDA---------------------------    
 612 UniRef50_Q69UA3        Os08g0538800 protein n=13 T...   364     5e-99  1  11.1%  63:576    91:577     --------------------------------------------------------------LDNYSLNIAISAAARVPSLDVGSQFHALSLKLSLASDSFVLNALINMYSSCNYPASARLVLDSAPrdVVSWNTIIAGYIrmPNKALQSFHQMAKEQVRLDEVTLLNVLVACARTGAMKVGSlhALVVLNGFEINCYIGSSLVSMYAKCGMVEEARRVFNRMPER----NVVCWTSMIAGCTQSGRFKEAVDLFRDMQIAGVKADDATIATVVSSCGQMGALDLGRYL------------HAYCDGHGLGKELSVKNSLIDMYSKCGDVNKAYQIFHGLTKR------DVFTWTVMIMGFAMNGLCVEALDLFAQMeeDKVMPNEVIFLGVLTACSHGGLVEQGYHHFHRMSkyNLVpeHYGCMVDLLGRAKLLAEAEQFIKDMP----VAPDVVVWRSLLFACRASGQVGLAEYAAERIEQ--LEPKrgGHVLLSNVYATTLRWVDVNNVRTGMGNSRTSKKP----------GCSFIEVDGCIHEFFAGDESHFETEAICNTLFGINELLVAE---------------------------    
 613 UniRef50_O82363        Pentatricopeptide repeat-co...   364     6e-99  1  10.8%   7:534    81:587     ------YTKIREFDDADKLFDEMPLRNIVTWNILIHGVIQRdnHRAHLGFCYLSRILFTDVSLDHVSFMGLIRLCTDSTNMKAGIQLHCLMVKQGLESSCFPSTSLVHFYGKCGLIVEARRVFEalDRDLVLWNALVSSYVLNgdEAFGLLKLmdKNRFRGDYFTFSSLLSAC---RIEQGKQIHAILFKVSYQFDIPVATALLNMYAKSNHLSDARECFESMVVR----NVVSWNAMIVGFAQNGEGREAMRLFGQMLLENLQPDELTFASVLSSCAKFSAIWEIKQV-QAMVTKKGSADF-----------LSVANSLISSYSRNGNLSEALLCFHSIRE------PDLVSWTSVIGALASHGFAEESLQMFESMLQK-LQPDKITFLEVLSACSHGGLVQEGLRCFKRMTeyKIEaeHYTCLIDLLGRAGFIDEASDVLNSMPT----EPSTHALAAFTGGCNIHEKRESMKWGAKKLLE--IEPtpVNYSILSNAYVSEGHWNQAALLRKR-ERRNCYNPKTPGCSWLGD---------------------------------------------------------------------    
 614 UniRef50_D1IC39        Whole genome shotgun sequen...   364     7e-99  1  10.3%  56:581   223:755     -------------------------------------------------------RYRHDPIRRPEAFGYVMVSYSRAGKLRNAMRVLTMMQKAGIEPDLSICNTAIHVLVMGNRLDKAVRFLERMQIVEIEPNVITYNCLIKgeDAMELIAEMPFKGCSPDKISYYTVMGFLCkkRIKEVRLLMEKMLksNLLPDQVTYNTFVHMLSKHGHGDEALEFLREAEERRFRVDKVGYSAIVHSFCREGRMDKAKEIVNEMFSKGCIPDVVTYTSVINGLCQERKVDQAKKMLRQM------------YKHGCKPNTVSYTALLNGLCKNGNSLEAREMMN-MSEEDWWI-PNAITYSVLMHGFRREGKSSEACDLVREMIKKGFFPTPVEINLLIQSLCQEEKVDEAKRFMEQCLNNGCAVNvfTTVIHGFCQKDDLEAALSLLDDMYLSN-KHPDVVTYTTIIDALGKKGRIEEATKLAMKMLRVGLIPTPVTYRTVIHQYCRMGRVEDLLKLLEKMLSR---QECRTAYNQVIEKLCSFGNLEQAYKLLGkmLIESYLSKGIPLMSYNVACRMFNRNLIPD----------------------    
 615 UniRef50_Q9SAJ5        Pentatricopeptide repeat-co...   364     7e-99  1  14.1%  84:602    76:673     -----------------------------------------------------------------------------------GFRFFIWASRRERLRSRESFGLVIDMLSEDNGCDLYWQTLEELKSGGVSVDSYcyAKMGMAEKAVESFGRMKEFDCRPDVFTYNVILRVMMREEVflAFAVYNEMLKCNCSPNLYTFGILMDGLYKKGRTSDAQKMFDDMTGRGISPNRVTYTILISGLCQRGSADDARKLFYEMQTSGNYPDSVAHNALLDGFCKLGRMVEAFELLRLFEKDGffELYANMLKKNIKPDIILYTILIQGLSKAGKIEDALKLLSSMPSKG--ISPDTYCYNAVIKALCGRGLLEEGRSLQLEMSETESFPDACTHTILICSMCRNGLVREAEEIFTEIEKSGCSPstFNALIDGLCKSGELKEARLlfDTMVESGSisPDIVSYNVLINGFCRAGDIDGALKLLNVLQLKGLSPDSVTYNTLINGLHRVGREEEAFKLFY---AKDDFRHSPAVYRSLMTWSCRKRKVLVAFNLWMKYLKKISCLDDETaeTERALRRLIELDTRKDELTlyTIWLIGLCQSGRFHEA-    
 616 UniRef50_Q9LS88        Pentatricopeptide repeat-co...   364     7e-99  1  10.8%  11:602   168:771     ----------ERAVEIFEWFKsgceLNVIHYNIMLRILGKACKWRYVQSLWDEM--IRKGIKPINSTYGTLIDVYSKGGLKVHALCWLGKMSKIGMQPDEVTTGIVLQMYKKAREFQKAEEFFKKWSCdsYTYNTMiyGKSGQIKEASETFKRMLEEGIVPTTVTFNTMIHIYGNNglGEVTSLMKTM-KLHCAPDTRTYNILISLHTKNNDIERAGAYFKEMKDDGLKPDPVSYRTLLYAFSIRHMVEEAEGLIAEMDDDNVEIDEYTQSALTRMYVEAEMLEKSWSWFKRFHVAGNMSSEG-------------YSANIDAYGERGYLSEAERVF---ICCQEVNKRTVIEYNVMIKAYGISKSCEKACELFESMMSYGVTPDKCTYNTLVQILASADMPHKGRCYLEKMRETGYVSDcyCAVISSFVKLGQLNMAEEVYKEMVE-YNIEPDVVVYGVLINAFADTGNVQQAMSYVEAMKEAGIPGNSVIYNSLIKLYTKVGYLDEAEAIYRKLLqnKTQYPDVYTSNCMINLYSERSMVRKAEAIffAMMLCMYKKNGRFEEATQIAKQMREMKILTDPL-SYNSVLGLFALDGRFKEA-    
 617 UniRef50_Q9ZQF1        Pentatricopeptide repeat-co...   363     8e-99  1  11.6%  62:602    99:593     -------------------------------------------------------------RLDFQTQCLAIAVISKLSSPKPVTQLLKEVVTSRKNSI---RNLFDELVLAHDRLET---------KSTILFDLLVRCCcrMVDEAIECFYLMKEKGFYPKTETCNHILTLLSrnRIENAWVFYADMYRMEIKSNVYTFNIMINVLCKEGKLKKAKGFLGIMEVFGIKPTIVTYNTLVQGFSLRGRIEGARLIISEMKSKGFQPDMQTYNPILSWMCNEGR---ASEVLREMKE------------IGLVPDSVSYNILIRGCSNNGDLEMAFAYRDEMVKQG--MVPTFYTYNTLIHGLFMENKIEAAEILIREIREKGIVLDSVTYNILINGYCQHGDAKKAFALHDEMMTDGIQPttYTSLIYVLCRKNKTREADELFEKVVGKG-MKPDLVMMNTLMDGHCAIGNMDRAFSLLKEMDMMSINPDDVTYNCLMRGLCGEGKFEEARELMGEMKRRGIKPD----------------------HISYNTLISGYSKKGDTKHAFMVRDEMLSLGFNP-TLLTYNALLKGLSkeGELAEE-    
 618 UniRef50_C5Z626        Putative uncharacterized pr...   363     9e-99  1  10.5%  68:581    38:500     -------------------------------------------------------------------YDDAITECVERRALREGRQVHARMVAAGYRPALYLATRLVIMYARCGALEDARNVLDGmeRNVVSWTAMISGYSQnpDQAWELFIMMLRAGI-------------------HQVKQVHAFAVKTNFELHMFVGSSLLDMYAKSENIQEARRVFDMLPAR----DVVSYTAILSGYTQLGLDEEALDLFRLLYNEGMQCNQVTFTALLNALSGLSSMDYGKQVHGLILRRELPFFMALQ------------NSLIDMYSKCGKLLYSRRVFDSMPER------SVVSWNAMLMGYGRHGLAHEVVQLFRSMCDE-VKPDSVTLLAVLSGYSHGGLVDEGLDMFDHIVKEqlNTEHYGCVIDLLGRSGQLQKALNLIEKMP----FQPTRAIWGSLLGACRVHTNVHVGEFVAQKLLD--IEpnAGNYVILSNIYAAAGMWKDVFRVRKLMLKK-------TVIKEPGRSW---MILDKVIHTFHSCERFHPRKEDINAKIKEIYVAIKAAGFVPD----------------------    
 619 UniRef50_A0D323        Chromosome undetermined sca...   363     1e-98  1  12.6%  63:516   283:741     --------------------------------------------------------------PTDIFFNKLIDYAFRKSQVEMAEYLLVFMrqAKIQPTIVTINTMIDQYFKNNQREKAWKTFENLKLTQTKPDNFTYTTLINgkLAFSLFEEYKQYN-QPDQIIYNCLLDACIngDLNRGFQLLNEMKQSqiQLDEITYNTLIKGCGRKKRLNDALNLFEEMKLIGIKPNRISFNSLLDSCVKCNKMNIAWRYFEEMRKqgIFPDNFTYSILVNghSNRDELLRAITLLEQIQETGQ-----------FKPDEILYNSLIDACVKFNEIQKGMELFREMKNK--QIEPSSVTYGILIKAYGKQNDLNGAFKIFEEMKQKKIPINDVTYGCLVDACVRNDRLDQALQFIDQMKSQNLPINtyTTIIKGFCKLNQTEEAIKYFDLMKQSPKTYPNLITYNSLLDGMVKNGLLNQADKLFQELVDSKIKPDLITFSTLLKGHCRKGNMKRLNETVQIM---------------------------------------------------------------------------------------    
 620 UniRef50_D1H2D4        Whole genome shotgun sequen...   363     1e-98  1  11.2%  77:586    52:533     ----------------------------------------------------------------------------SISNHAHMNQMLSQMIMNYIPIDHLNLMKLIDFSVSSHGFAASALLFTQFYgdSDLCNSMIRCYTDSNKHLHSVFiqMWKNGIFPDSSTFPTVLKSVAqcRQELGKAIHCCIIQMGFESNVYVSTALVNMYGTCSSVSDARQVFDEIPDR----NIVSWNALITGYNHNRMFRKVIDVFREMQIAGAKPVEVTMVGVLLACAHLGALNQGRWI------------DDYIDHNRLRLNVFVGTALIDMYAKCGVVDEAEKIFKAMRVK------NVYTWNVLISGYAMNGRGESALQAFSRMIMEKFKPDEVTFLGVLCACCHQGLVNEGRTYFTSMKEegLRpeHYGCMVDLLGRAGFLDEAQQLIQAMS----MQPDPIIWRELLGACRIHGNIQLGEFAIKKLLE--LEPNNgnYVLLANLYARDQRWDKVGEVREMMDCRRVRKVP----------GCSSIEIDNVVYEFVVSNYIKPGFEEVYKLLADMNKKLKLAGYVADtmASY-----------------    
 621 UniRef50_B9RW91        Pentatricopeptide repeat-co...   363     1e-98  1   8.1%   6:601    16:641     -----MYAKCGAFSDSNKLFGenCSHDPIFWNILLSGFAASPIHD--AQTFSFFNRMRvqVKPTSVTAAVILPVCARMRDIYVGKSFHCYMIKTGMETHTLVGNALVSMYAKCGLvdAYAAFQSIYHKDVVSWNAIIAGFSEnvDSALRLFFFMLKTQIKPNHATIATVLPLLASLATDTarEIHCYILRHNESldVSVCNALLSFHLMVGRVKEAELLFRRMELR----DLISWNAIISGFASNGEWSKSLELFQEllESN-EPDSVTLVSILPACAQLQNLKAGREIHSYVLRHPYLYQ-----------DTSVGNALVSFYAKCNNLVAAYHTFLMIPSR------DLISWNSMLDAFAIGGRNAQFFKLLHWMHTEGIRPDSITILTILHFCANVLKVDKVKETHCYSLRCgfEPTTRNAMLDTYAKCSNVEYAFKVFQTLSDN----RNLVTFNSMISGYVNCGLYDDAYMIFEKMPAS----DLTTWNLMVRGCAENDCPDQAFSLFRELQARGMKPDAVTIISLLpdLYAKCGDLGYAFKLFHsaMVGGYAMHGMGEEALRIFSDMLDLGIKPDHIVI-TAVLSACCHAGLVDE--    
 622 UniRef50_D1I8R2        Whole genome shotgun sequen...   363     1e-98  1  13.1%  42:581     3:588     -----------------------------------------GLKSRALHHLSHTHKVLALPSLHHF-YDHLLQCCTSLTTLKL---IHSSLSTRGFLLHtfLARLIILYSKLGDLHS-ARTLFDHRhpNSFLCNTMLRAYANAGreAIDLYIYMQRMGVGVNNFTYPFVLKVCaeLGAVFGEVVHGQVVRTGFGSDLFVEAALVDMYAKCGEIGDAHEVFDRM----LIRDVVCWTAMITLYEQAERPLKALMLFRKMQEEGFLGDEItwNSMLSGYTQNGRPTDALSLFNQMQASegRKLHNFVISSKMDIDTTLRNAIMDMYMKCGDLDTAVEMFN----NCELGERDVSSWNVLISGYGVHGHGKEALELFSRMQVEGVEPNDITFTSILSACSHAGLIDEGRKCFADMTKLSVRPehYACMVDMLGRAGFLNEAFRLIKKIPSR----PSDEVWGALLLACRIHGNTELGEIAANNLFQ--LEPEhgYYVLMSNIYAASNKWKEVEMVRQNMKSRGLKKPAAFSV----------IEFGTEVHGFHTADQSSPYYREVYRKVESLAIEMKMVGYVPD----------------------    
 623 UniRef50_Q69VD4        Os06g0597500 protein n=3 Ta...   363     1e-98  1   9.9%  31:545    17:530     ------------------------------CNVLIRTLARRGSFARVMAVYYDLRARGLV--ADSFTYPFVLRAVGVLKLSVEGRKAHAAAVKTGFRWDAYTGSSLMEMYTMLGRVDIARKVFDEMprALVLWNMMVRCYIRCgsAAVALSEQMERSGVTPDRVTLVTAVTACSRArlSLGRRIHVYM-DNVFGFNLPVANALLDMYTKNDCLEEAVKLFEQMPARNieKDLILWTAMINACVQHGCFEEALTLFRDMQMQRVEPDRFTVVTLLTCCANLGALDQGEWIHQYAEQRK------------MKIDAVLGTALIDMYSKCGHIEKSLEVFWRMQGR------DATAWTAIICGLATNGQAGRALELFQDMQRSKVKPDGVTFIGVLSACCHGGLVDEGRKQFHAMReyQIEpeHYSCLVNLLGRAGLLDEAERLIGDVPINKDAMP---LFGALLTACKAHGNVEMSERLTKRICEQDSQITDVNL-LMSNVYATASRWEDVIRVRGKM----------AHPTVKKNAGCSLIEVKGY----------------------------------------------------------    
 624 UniRef50_Q9M9R6        Pentatricopeptide repeat-co...   363     2e-98  1  11.4%  17:532    60:536     ----------------RLIFDSVTFPNVFVVNSMFKYFSKMDMANDVLRLYE--QRSRCGIMPDAFSFPVVIKSAGRFGILFQAL-----VEKLGFFKDPYVRNVIMDMYVKHESVESARKVFDQiqRKGSDWNVMISGywGNKEEACKLFDMMPEN----DVVSWTVMITGFAkkDLENARKYFDRMPEK----SVVSWNAMLSGYAQNGFTEDALRLFNDMLRLGVRPNETTWVIVISACSFRADPSLTRSLVKLIDEKRVRLNCFVKTALLDMHAKCRDIQSARRIFNEL---------------GTQRNLVTWNAMISGYTRIGDMSSARQLFDTMPKR------NVVSWNSLIAGYAHNGQAALAIEFFEDMIDYgsKPDEVTMISVLSACGHMADLELGDCIVDYIRKNQIKLNdyRSLIFMYARGGNLWEAKRVFDEMKER-----DVVSYNTLFTAFAANGDGVETLNLLSKMKDEGIEPDRVTYTSVLTACNRAGLLKEGQRIFKSIRN-----PLADHYACM-----------------------------------------------------------------------    
 625 UniRef50_Q7XKS5        OSJNBa0038P21.6 protein n=2...   362     2e-98  1  11.0%  88:603    17:531     ---------------------------------------------------------------------------------------HAPMRLLMLPPSTPTTSLIDVLPAECTGPELALAVFGRLLrdVCSYNTLIDGFSKEgdKAYELFYKMIEQSVSPDVVTYNSLIDGLCKTkmVKSERVLEQMVDAGIRPNNKTYNSLIYGYSTAGMWKESVRVFKEMSSSGLIPCVVNCNSFIHALCRHNRIKEAKDIFDSMVLKGPKPNIISYSTLLHGYAAEGCFANMNSLVNLMVSK------------GIVPNHRFFNILINAYARCGMMDKAMLIFEDMQNKG--MIPDTVTFATVISSLCRIGRLDDALHKFNHMVDIGVPPSEAVYRCLIQGCCNHGELVKAKELISEMMNKDIPpkYFSSIINNLFKEGRVAEGKDIMDLMVQTG-QRPNVVTFNSLMEGYCLVGNMEEAlkMFHEMIESGTTVSIQTYGVVLGGLCRNNCTDEANMLLEKLFAMNVKFDIIT----------------------FNIVISAMLKVGRRQEAKELFAAISTYGLVP-TVHTYNLMISNLIKEESYEEAD    
 626 UniRef50_C7J306        Os05g0313900 protein n=4 Ta...   362     3e-98  1  12.2%   1:540   148:769     LGAAGLPDTAVRAFDAARA-SFGCTPNSYTYNCLLDALAKAGRADDAQARLREMVARCGDGSVDKYTLTSLLRCYCNAGRPDDANDVFQRMSELGWVDEHVLTTLMVAFSKWggRVDKAMDMFAKMVSYGFvySVLIEGLCQQKdrAVKLFKEMKSSGVAPDVRLLKKVIEAFCrgEVEAAYQLLRSmiREDAKPNSDSFNIVVCGLCKVKKLDMALALTKDMISLGCKGKILMFNDLIHELCNMDRLEEGYGIFNQMKDLGLTPSEFTYNSLFYGICRRKDPKAALDLLREMQT------------NGHPPWIKNCTEMVQQLCFSGRVTEAVQFLDGMLQIG--FLPDIVTYSAAMNGMCNTGEVDDALHLFRDISCKYYLPDVVAHNILINGFRKSSKLDEAQKIMEEMLEKGLFPstYNLMIDVCCKTGRIEKAISYLDKMVYE-EKQPTVITYTSLIDGFCSAGRPDEAIKLWCEMREKGCAPNNIAYTAFINGLRKCGRIETALTYFEEMVTKGFELDTFSLLY-FINFLISNG---------------------------------------------------------------    
 627 UniRef50_D1I468        Whole genome shotgun sequen...   362     3e-98  1  10.2%  67:555    31:546     ------------------------------------------------------------------SFNYLLNCCSSLPDLS---RIHALVVTNGCGQNLLLStlIITACCLAPTMDYARKMFDQmkRDVFLWNTLIRGYADAgeEALALYSNMHGAGLFPDNYTFPFVVRSCAVLslREGKEVHCNIVKHGFDSDVFVQSSLVAMYSQSGETLGMELVFGEMVVR----NIVSWTAVIAGYVQNRYFKEGLGVFREMVGSGTQPNAVTLVSVLPACA-------GLEFLNL-----GKLIHGYGIKLGVDPDVSLTNALIALYGKCGNVETArkLFRRMQAEKVDfcRSVVSWTSMIGACASHGHGEDALKLFSRMKDEGVKPNSFTFAAVFTACRHSGLVEEGRKHFESmrDYSIMpeHCACMVDLLGRAGSLMEAYEFIDKMP----VEPDVSVWGALLGSCRIHSNLELAELVAEKLFL--LDPQTVtyVLMSNIYAEAGRWEDAARLRKLMEERELKKIPGH----------SLVEVNRRFHTFLSGSRS------------------------------------------------    
 628 UniRef50_D1I3E0        Whole genome shotgun sequen...   362     3e-98  1  12.2%  99:572    70:544     --------------------------------------------------------------------------------------------------TVRSRTKLMNILIEKGKPQEAQLIFNSLTEEGHRPTLITYTTlkRFKSIPSLISKLEKNGLKPDSVFFNAMINAFSegNVKEAMKIFRKMKDRGCKPTTSTFNTLIKGYGNAGMPEECLKLLDLmqEENVKPNDRTFNSLIRAWCNKKRITEAWNVVYKMAASGLQPDVVTYNTLARAYAQNGETSRAEGMILEMQN------------NRVMPNERTCGIIINGYCKEGKMKDALRFLYRM--RNYGVHPNLVIFNSLIKGFLDITDTDGVDEALTLMEEFGVKPDVVTFSTIMNAWSSVGLMDKCQEIFDDMVKAGIEPDifSILAKGYVRAGEPEKAESLLTAMGKSG-VQPNVVIFTTIISGWCSAGKMEYASRVYEKMCEMGICPNLKTFETLIWGYGEAKEPQKAEELLQIMEQKGVAPVKST-IQLVADAWHALGLANEavEEAPKVMISTKEDDVAAESLERIYQK-------------------------------    
 629 UniRef50_P0C8Q8        Pentatricopeptide repeat-co...   362     3e-98  1  11.6%  18:602    66:662     -----------------RVLKSGLDSNGYICNSVLNMYAKCRLLADAESVFR------DHAKLDSASFNIMVDGYVRSRRLWDALKLFDVMPERSCVS----YTTLIKGYAQNNQWSEAMELFREMRNLGIMLNEVTLATVISAcdCRMLQSLAIKLKLEGRVFVSTNLLHMYccLCLKDARKLFDEMPER----NLVTWNVMLNGYSKAGLIEQAEELFDQITEK----DIVSWGTMIDGCLRKNQLDEALVYYTEMLRCGMKPSEVMMVDLLSASARSVGSSKGLQLHGTIVKRGFDciHFYAVSNDIKlhIASRNALIAGFVKNGMVEQAREVFDQTHDK------DIFSWNAMISGYAQSLSPQLALHLFREMISSsvKPDAITMVSVFSAISSLGSLEEGKRAHDYLNFSTIPPnlTAAIIDMYAKCGSIETALNIFHQ--TKNISSSTISPWNAIICGSATHGHAKLALDLYSDLQSLPIKPNSITFVGVLSACCHAGLVELGKTYFESMKSDHGIEPDIKHYGCMVDLLGKAGRLEEA----KEMIKKMPVKADVMIWGMLLSASRTHGNVepSHGGC-KVMLSNVYADAGRWED-    
 630 UniRef50_Q9LUL5        Pentatricopeptide repeat-co...   362     3e-98  1  11.0%  37:582   104:623     ------------------------------------GLSKSTKLDEAVTLIEN-SSSSPSNLSTPEAYTDLLHACISAKSLHHGIKICSLILnpSLRHNPKLLSKLITLFSVCRRLDLARKIFDDVTDSSlvWAAMAIGYSRNgrDALIVYVDMLCSFIEPGNFSISVALKACVDldLRVGRGIHAQIVKRKEKVDQVVYNVLLKLYMESGLFDDARKVFDGMSER----NVVTWNSLISVLSKKVRVHEMFNLFRKMQEEMIGFSWATLTTILPACSRVAALLTGKEIHAQILKSKE------------KPDVPLLNSLMDMYGKCGEVEYSRRVFDVMLTK------DLASWNIMLNCYAINGNIEEVINLFEWMIESGVAPDGITFVALLSGCSDTGLTEYGLSLFERMkeFRVSpeHYACLVDILGRAGKIKEAVKVIETMP----FKPSASIWGSLLNSCRLHGNVSVGEIAAKELFV--LEpnPGNYVMVSNIYADAKMWDNVDKIREMMKQRGVKK----------EAGCSWVQVKDKIQIFVAGGGYEFRNSDEYKKvtELQEAIEKSGYSPNT---------------------    
 631 UniRef50_A2WPS0        Putative uncharacterized pr...   361     4e-98  1   9.4%   7:582    47:554     ------YSKCRRPCCARRVFDEIPDPCHVSWSSLVTAYSNNGLPRSAIQAFHGMRAEGVCC--NEFSLPVVLKC---VPDARLGAQVHAMAMATGFGSDVFVANALVAMYGGFGFMDDARRVFDesERNAVSWNGLMSAYVKncGDAIQVFGEMVWSGIQPTEFGFSCVVNAcgSRNIEAGRQVHGMVVRMGYDKDVFTANALVDMYVKMGRVDIASLIFEKMPDS----DVVSWNALISGCVLNGHDHRAIELLLQMKSSGLVPNVFTLSSILKACAGTGAFDLGRQIHGFMIKVNAD------------SDDYIGVGLVDMYAKNHFLDDARKVFDWM------FHRDLILCNALISGCSHGGRHDEALSLFYELRKEGLGVNRTTLTAVLKSTASLEAASTTRQVHALAEKIgfDAHVVNGLIDSYWKCSCLSDANRVFEE-----CSSGDIIAFTSMITAlcDHGMWNEVAKV-----------------------------------RKLMKDSNIKKEPA----------MSWVEVKDKVHTFIVGDKSHPLTKEIYAKLDELGDLMSKAGYVPNV---------------------    
 632 UniRef50_Q9LUS3        Pentatricopeptide repeat-co...   361     5e-98  1   8.9%   5:530    43:650     ----RLYASCNEVELARHVFDEIPHpnPIAWDLMIRAYASNDFAEKALDLYYKM--LNSGVRPTKYTYPFVLKACAGLRAIDDGKLIHSHVNCSDFATDMYVCTALVDFYAKCGELEMAIKVFDEmkRDMVAWNAMISGFSLHCcdVIGLFLDMRrdGLSPNLSTIVGMFPALGRAglREGKAVHGYCTRMGFSNDLVVKTGILDVYAKSKCIIYARRVFD----LDFKKNEVTWSAMIGGYVENEMIKEAGEVFFQMlkAGFILDLTVQNTIISFYAKYGSLCDAFRQFSEirPEESFRLFHEMRTSGIRPDITtcNALMDMYTKCGKLDVAKRVFDTMHKR------DIVSWNTMLFGFGIHGLGKEALSLFNSMQETGVNPDEVTLLAILSACSHSGLVDEGKQLFNSMSRGDfrIDHYNCMTDLLARAGYLDEAYDFVNKMP----FEPDIRVLGTLLSACWTYKNAELGNEVSKKMQSLGE--TTESLVLLSNTYSAAERWEDAARIR--MIQKKRGLLKTPGYS-------------------------------------------------------------------------    
 633 UniRef50_Q9LW84        Pentatricopeptide repeat-co...   361     5e-98  1  12.9%  58:592   119:628     ---------------------------------------------------------RRNFQHDCSTYMTLIRCLEEARLYGEMYRTIQEVVRNTYVSVsaVLSELVKALGRAKMVSKALSVfrKCKPTSSTYNSVILMLMQEgeKVHEVYTEMCNEgcFPDTITYSALISSYEkgRNDSAIRLFDEMKDNCMQPTEKIYTTLLGIYFKVGKVEKALDLFEEMKRAGCSPTVYTYTELIKGLGKAGRVDEAYGFYKDMLRDGLTPDVVFLNNLMNILGKVGRVEELTNVFSEM------------GMWRCTPTVVSYNTVIKALFESkhVSEVSSWFDKM--KADSVSPSEFTYSILIDGYCKTNRVEKALLLLEEMDEKGFPPCPAAYCSLINALGKAKRYEAANELFKELKEnnVSSRVYAVMIKHFGKCGKLSEAVDLFNEMKNQG-SGPDVYAYNALMSGMVKAGMINEANSLLRKMEENGCRADINSHNIILNGFARTGVPRRAIEMFETIKHSGIKP----------DGVTYNTLLGCFAH------------AGMFEEAARMMREMKDKGFEYDAI-TYSSILDA-----------    
 634 UniRef50_B9SZC0        Pentatricopeptide repeat-co...   361     5e-98  1  12.6%  42:595    62:572     -----------------------------------------RRQNDETAALRLLSWASKQprPNSSIYEEILRKLGKVGSFNSMKDILQEMKGLDCQIDRGVLLIFIDSYAKFELYEEILCIVEVME---------------------------VEFGLALDTLLYNFLLNVLVDgkLKLVENVHSTMVSKGIKPDVSTFNILIKALCKAHQIRPAILVMEEMSSYGLVPDEKTFTTLMQGFIEEGNMDGAFRVKEQMLDAGCPVTNVTVNVLVHGFCKEGRIEEALSFIDEMSNEGKheLYNEMKNKGCQPDEFTYNMLIDSLCYRGKLQEALGLLREMEISGC--ARNVITYNTLIDGFCKNKKIEEAEEIFDQMEIQGLSRNSVTYNTLIDGLCKSRRVQEAAQLMDQMIMEGLKPdtYNSMLTYFCREGDIQKAADIVQTMTSNGC-EPDIVTYGTLIGGLCKAGRVEVASRLLRSIQLKGMVLTPHAYNPVIQALFKRKRTKEAMRLFREMEEKGNTPDAFTYKIVFRGLSNSGGPIGEAVDFVI---------------------EMIEKGFLP-EFSSFYMLAEGLCS--------    
 635 UniRef50_B9R8S4        Pentatricopeptide repeat-co...   361     5e-98  1   9.1%   7:579    67:689     ------LSKAEQIDKAKKVFDEMPERDSVSWNSMMSGYFRNGKYEEVVKVFVLMIR-NFTCIVNLLSFSCAMKACGALGCFKLAIQLHGLFEKFDFGSDKSIETSVMGMYIKCGAFTYADRVfrIPERDSVSWSTIISILSRHGFGvlSMFIEMWTQGFRPNSMTFACVLSACTshDLEWGAHLHARIIRKEIILDDYVGNGLVYMYAKCGHLKFARRVFNSLTEH----NAVSWTSLINGVARCGLKEEALLLFNKMREVLVALDEFTFATVLKVCSHPN------------FNFTGRQLHALTTKTGMGSIVTVGNALITMYSKCGDTQKADCVFKMMPVRNIipERNVITWNSMISMYIQHGFQEWGLKLYVRMQRERIAPDDITFATSISACADLAMLKLGNQIVAQAEKLGfdVSVANSLVTMYSRCGQIEEARKAFDLVSMK-----NLVSWNSMIAGYAQNGHGRVAIEVFKNMLEMEYIPDHISysVLSDLLGRAGLLEQAKNLIDDMPFTPNglLGACKIHCNSKLAEYAveLDVDNSGSYALLAHIYSSRGNLEGVADLRKLIKEKGIQ------------------------    
 636 UniRef50_O04504        Pentatricopeptide repeat-co...   361     5e-98  1  11.7%  33:529   122:599     --------------------------------SFLDGFVRNGSDHQVHSIFHAIS-MCDNVCVNSIIADMLVLAYANNSRFELGFEAFKRSGYYGYKLSALSCKPLMIALLKENRSADVEYVYKEMIRRKIQPNVFTFNVVINanKARDVMEDMKVYGCSPNVVSYNTLIDGYCkgKMYKADAVLKEMVENDVSPNLTTFNILIDGFWKDDNLPGSMKVFKEMLDQDVKPNVISYNSLINGLCNGGKISEAISMRDKMVSAGVQPNLITYNALINGFCKNDMLKEALDMF------------GSVKGQGAVPTTRMYNMLIDAYCKLGKIDDGFALKEEMEREG--IVPDVGTYNCLIAGLCRNGNIEAAKKLFDQLTSKGL-PDLVTFHILMEGYCRKGESRKAAMLLKEMSKMGLKPRhyNIVMKGYCKEGNLKAATNMRTQMEKERRLRMNVASYNVLLQGYSQKGKLEDANMLLNEMLEKGLVPNRITYEI----------------VKEEMVDQGFVPDIEGHL--------------------------------------------------------------------------    
 637 UniRef50_A5C7D6        Putative uncharacterized pr...   361     5e-98  1   9.5%   6:552   128:730     -----LYSKCGSLSFACQVFDTTPERDLVTWNAILGAYASsgNAQEGLHLFRLLRE--SLGSTTRMTLAPVLKLCSNSXCLWAAKGVHGYAIKIGLVWDVFVFGTLMNIYSKCGRMXDARLLFDGmeRDVVLWNMMLKGYVQLgkEAFQLFSEFHRSGLXPDEFSVQLILNGVFEVNXDEGKWHADQVQAYXXKldVFCWNKKLSEYLWAGDNWGAIECFVNMNGLNVXYDXVTltALIDVYSRSGKMEEAELLF----QNKDDLDLACWNAMMFGYIISNDGNKALGLFSLINRSggKQIHAHVIKAGFXSDLYVNSGILDMYIKCGDMVNAGIVFNYISA------PDDVAWTSMISGCVDNGNEDQALRIYHQMRQSGVMPDEYTFATLIKASSYVTALEQGRQLHANVIKLDcdPFVGTSLVDMYAKCGNIEDXYRLFKKMNVR-----NIVLWNAMLVGIAQHGNAEEAVNLFKSMKSHGIEPDRVSFIGILSACSLAGLTSEAYEYFHSMPNDCGIEPEIEHYSCLVDALGXAGLVQEXDKVIETM---------------------------------------------------    
 638 UniRef50_B9MU86        Predicted protein n=3 Tax=M...   361     6e-98  1  11.0%  85:582     5:472     ------------------------------------------------------------------------------------KQVHAQVLKWE-NSFCASNLVATCALSDWGSMDYACSIFRQIDQPgfEFNTMIRGYVnnMENALFLYYEMLERGVESDNFTYPALFKACASLriEEGMQIHGYIFKRGLEGDLFVQNSLINMYGKCGKIELSCSVFEHMDRR----DVASWSAIIAAHASLGMWSECLSVFGEMSREgcRPEESILVSVLSACTHLGALDLGR------------CTHVTLLRNIREMNVIVQTSLIDMYVKCGCIEKGLSLFQRMVKKNQL------SYSVMITGLAMHGRGMEALQVFSDMLEEGLKPDDVVYLGVLSACNHAGLVDEGLQCFNRMkeHGIEPthYGCIVHLMGRAGMLNKALEHIRSMPIK----PNEVVWRGLLSACKFHHNLEIGEIAAKSLGELNSS-NPGDYVVLSNMYARAKRWEDVAKIRTEMARKGFTQTPGF----------SLVQVERKIYKFVSQDMSHPQCKGMYEMIHQMEWQLKFEGYSPDT---------------------    
 639 UniRef50_Q9ZUA2        Pentatricopeptide repeat-co...   360     7e-98  1   9.8%  47:599     2:524     ----------------------------------------------VREALQFLSRLRKSSnpDPFTCNKHIHQLINSNCGILSLKFLAYLVSRGYTPHRSSFNSVVSFVCKLGQVKFAEDIVHSmePDVISYNSLIDGHCRNGdsASLVLESLRAShcKPDIVSFNSLFNGFSKMklDEVFVYMGVMLKC-CSPNVVTYSTWIDTFCKSGELQLALKSFHSMKRDALSPNVVTFTCLIDGYCKAGDLEVAVSLYKEMRRVRMSLNVVTYTALIDGFCKKGEMQRAEEMYSRMVEDR------------VEPNSLVYTTIIDGFFQRGDSDNAMKFLAKMLNQGMRL--DITAYGVIISGLCGNGKLKEATEIVEDMEKSDLVPDMVIFTTMMNAYFKSGRMKAAVNMYHKLIERGFEPDVvsTMIDGIAKNGQLHEAIVYF------CIEKANDVMYTVLIDALCKEGDFIEVERLFSKISEAGLVPDKFMYTSWIAGLCKQGNLVDAFKLKTRMVQEGL----------LLDLLAYTTLIYGL------------ASKGLMVEARQVFDEMLNSGISPDS-AVFDLLIRAYEKEGNM----    
 640 UniRef50_D1IH95        Whole genome shotgun sequen...   360     8e-98  1  11.0% 125:580     1:490     ----------------------------------------------------------------------------------------------------------------------------MPERNVVSWTSVIAGYSQNGQgaLEFYFQMLQSGVMPDQFTFGSIIKACSsgDIGLGRQLHAHVLKSEFGAHIIAQNALISMYTKSNLIIDALDVFSRMATRDlrPDLVAWNAIIAGFA-YGELRDAIFFFEEMR---CNADLVSWNAILTACMRHDQAEEVFRLLKLMCISqgNQVHCYALKTGLNCDTSVTNGLIDLYAKCGSLKTAHKIFDSM------INPDVVSWSSLILGYAQFGYGEEALKLFKTMRRLDVKPNHVTFVGVLTACSHVGLVEEGWKLYGTMEKegIApeHCSCMVDLLARAGCLNEAEGFIHQM----AFDPDIVVWKTLLAACKTHGNVDVGKRAAENILK--IDpnSAAHVLLCNIYASKGNWEDVARLRSLMKQRGVRKVPGQ----------SWIEVKDRIHVFFVEDSLHPERNKIYTMLEELLLQMLDAGYVP-----------------------    
 641 UniRef50_Q9FLD8        Pentatricopeptide repeat-co...   360     1e-97  1   9.9%  34:578   125:672     ---------------------------------MVSLLSRENDWQRSLALLDWVHE-EAKYTPSVFAYNVVLRNVLRAKQFDIAHGLFDEMRQRALAPDRYTYSTLITSFGKEGMFDSALSWLQKMEQDRVSGDLVLYSNLidYSKAISIFSRLKRSGITPDLVAYNSMINVYGKAKlrEARLLIKEMNEAGVLPNTVSYSTLLSVYVENHKFLEALSVFAEMKEVNCALDLTTCNIMIDVYGQLDMVKEADRLFWSLRKMDIEPNVVSYNTILRVYGEAELFGEAIHLFRLMQRKDIEQ------------NVVTYNTMIKIYGKTMEHEKATNLVQEMQSRG--IEPNAITYSTIISIWGKAGKLDRAATLFQKLRSSGVEIDQVLYQTMIVAYERVGLMGHAKRLLHELKldNIPRET---AITILAKAGRTEEATWVFRQAFESGEVK-DISVFGCMINLYSRNQRYVNVIEVFEKMRTAGYFPDSNVIAMVLNAYGKQREFEKADTVYREMQEEGCVFPDEVHFQ-MLSLYSSKKDFEMVESLFQRLeaALYERADKLNDASRVMNRMRERGI-------------------------    
 642 UniRef50_Q9M316        Pentatricopeptide repeat-co...   360     1e-97  1  11.9%  19:589   143:729     ------------------AKEKNIPLTVVATNLLIRWFGRMGMVNQSVLVYERLD----SNMKNSQVRNVVVDVLLRNGLVDDAFKVLDEMLQKefPPNRITADIVLHEVWKGRLleKIIALISRFSSHGVSPNSviSSLCKNARaaWDILSDLMKNKTPLEAPPFNALLSCLGRndISRMNDLVLKMDEVKIRPDVVTLGILINTLCKSRRVDEALEVFEKMRGKriKADSIHFNTLIDGLCKVGRLKEAEELLVRMkeERCAPNAVTYNCLIDGYCRAGKLETAKEVVSRMKEDE------------IKPNVVTVNTIVGGMCRHHGLNMAVVFFMDMEKEG--VKGNVVTYMTLIHACCSVSNVEKAMYWYEKMLEAGCSPDAKIYYALISGLCQVRRDHDAIRVVEKLKEGGFSLdaYNMLIGLFCDKNNTEKVYEMLTDMEKEG-KKPDSITYNTLISFFGKHKDFESVERMMEQMREDGLDPTVTTYGAVIDAYCSVGELDEALKLFKDMGLHSKVNPNTVIYNILINAFSKLGNFGQALSLKEEMKMKMVRPNVEteTLLKLMDEMVEQSCEPNQITMEILM--------------    
 643 UniRef50_B9H9W5        Predicted protein n=2 Tax=M...   360     1e-97  1  11.6%   5:574    53:587     ----SMYSRSGNFDEAEKVFNGVGCKDLRCLNCMILEYGKAGYEKRAIGVFIYL--ISVGLDPNDYTFTNIISTC----NVEEGKQLHGLAVKYGVLLQTSVGNAVITMYGKNGMVEEAARMFSVmkKNLISWTALISGYTRNgeKAVDGFLELRGCGVECDSGLLATILDGCSECklDLGTQIHGLVIKLGYPCDINIGTALIDLYAKCKNFQSARTVFN-----GLSpsTASFNAILVGFIENdnEEDPMVFLSQLRLAGIKPDSVSFSRLLSLSANRASLVKGRGL------------HAYSIKTGFAGHISVSNALITMYAKCGIVEDAYQAFNSMSAN------DCISWNAIISAYSLHGQGEKALLLYQEMEEKGFTPDEITILVILQACTYSGLSEDGLHLFNTMekYGIQpeHYACMVDLLGRAGYLSQAMDIINRSP----FSESTLLWRTLVNVCKLCGDLNLGKLASKHLLD--LSPDegSYVLVSNIYAGEGMIDEASKVRTTMKD----------LKLSKEAGSSWVEIDNMVHYFVASGTDHPESIEIYARLDLLRNEMR-----------------------------    
 644 UniRef50_A3BML6        Putative uncharacterized pr...   360     1e-97  1  10.8% 137:580     1:403     ----------------------------------------------------------------------------------------------------------------------------------------MYAKAGRVDLARDAFDEAPLR----DVFLCNVMLAAysRSEVAEARKVFDGMPMR----DLVSWNTMIHGYAMRGEVGLAREIFDGTEDR----DAFSWSSMISAYAKSRRSKEALELWREMHAASIIPDCITLVSVVSACSDLGALAVGAEV------------HRFVESNRIELDLKLGTALIDMYAKCGDIESAQRVFDRMPEK------DVQTWSSMIIGLANHGLGHESLSLFSKMISEGMKPNGVTFVGVLIACTHVGLVSEGKKYFRSMNehGIEpeHYGCMVDLLGRSGHVEEARQLIRSM----TFEPDTIIWRALLGACRIHKNVEIAEEAMAKLRVLDPLGD-GHYVLLSNIYAQANSWEGVAEMRKTIRRDNIQRIPGR----------SSIEWDEKIHEFVSGDRLHPRSKEIYRVLEEMMDRLKQAGYKP-----------------------    
 645 UniRef50_B9IEF8        Predicted protein n=5 Tax=r...   360     1e-97  1  11.7%  81:524     1:427     --------------------------------------------------------------------------------MKDLKQIHARVIQLGFEQNRFVVGKVIVFCAahGDMNYAVSVFEKidPDAFIFNTMIRGfaNDPRKAFDYYKRMQERGLVSDSFTFSFLLKVCGQLGLvlGRLMHCSTLKRGLNSHVFVRNTLVHMYGTFKDIEASRQLFEEIPN----PELVAWNIIIDCHVSCGKFNEALEMFSRMLKFGIEPDEATFVVILSACSALGALDFGRWVHSCISNIGHGCITE------------VNNSLLDMYAKCGALQEAFEIFNGMNKK------NTVTWNTMILGLASHGYANEALALFSNMLEQKlaPDDITFLVVLSACSHGGMVDKGWRFFDIMKkyHIQPthYGCMVDILGRAGFVEEAYRLISNMPMQC----NAIVWRTLLAACRLHGNVELGKQVRKQLLE--LEPDHSsyVLLANTYASAGRWNEAMRVRKTMHKRGVQKP-------------------------------------------------------------------------------    
 646 UniRef50_Q5ZBJ9        Os01g0357800 protein n=9 Ta...   359     1e-97  1  10.1%   7:529   295:795     ------YTRCGAYQSAATVLQN-SKGTVASWNALLSGLNRAGQSFNAIQYLPVMLH-EHKVTPDSVTFANVLSACAELCYFCFAASIHAYFIRRFIpdVVLTTALIEVYTKCTRVMRSKYLFDQLIIKDVVSYNAMIYGYLQnaNEATSLLNYMMAEGVAPDFATVLSLLAAFaqRDLVRGRWIHGFAIRHGFCSDVDVENQILYMYSACGKIAAARAIFDSLEKK----NLVSWTAMMKGCLSNGHADEVVQLFQVMQKYGEKPDSVSLVTAVQAVSDLGHLN------------GLKQIHCFVYRSLLEKDKITANSLISAYAKCGKLDLSAGLFFSLKYR------NLDTWNAMISAYAMHGFHINVLEMFKQMEEENIQPDELTFSTVLTACSHAGLVKDGWRIFNSMTsySVLpeHYGCMVDLLGRAGHLEDGYKFIktLKDK------STIFCALLSACRTHGNTRLAHAISKELLEHGPQ-NPGIYALISEVYAQEGQWNEVANTKARADLSGLKKHPGSSL--------------------------------------------------------------------------    
 647 UniRef50_B9HGU3        Predicted protein n=3 Tax=r...   359     1e-97  1  10.7%   3:589   169:732     --ANSLISMFGYFGSVEevFSGMDEHDTISWNSMIAAYIRNGLCKESLRCFSWM--FRVHKEINSTTLSTMLAGCGSVDNLKWGRGIHSLVLKFGWNSNVCASNTLITMYSDAGRCEDAELVFQGmeKDMISWNSMMACYAQDGndALKLLATMFYMRRGANYVTFTSALAACSDPEFaeGKILHALVIHVGLHENVIVGNALVTLYAKSGLMIEAKKVFQTMPKR----DGVTWNALIGGHADSEEPDEALKAFKLMREEGVPINYITISNVLGACLAPNDLLE-----------HGMPIHAFIILTGFQSDEYVQNSLITMYAKCGDLNSSNNIFDRLTSKNASAWNAMM------AANAHHGHMEEALKFLLEMRRAGVNVDEFSFSECLAAAAKLAILEEGQQLHGLAVKLGCDSnvASATMDMYGKCGEIDDVLRIIPR-----PINRSRLSWNILTSSFSRHGFFEKAKETFHEMINLGVKPDHVTFVSLLSACSHGGMVEEGLAYYDSMIKEFGIPAKIGHCVCIIDLLGRSGRFAEAE----TFIKEMPVSPTDHVWRSLLAACKTHGNLELGRKAVENLL--------------    
 648 UniRef50_Q1PFA6        Pentatricopeptide repeat-co...   359     1e-97  1   9.0%  59:602    25:590     ----------------------------------------------------------LSHSPNKFTFPPLLKSCAKLGDVVQGRILHAQVVKTGFFVDVFTATALVSMYMKVKQVTDALKVLDEMPERGisVNAAVSGLLENgrDAFRMFGDARVSGSGMNSVTVASVLGGC--GDIEGGMQLHCLAMKSGFEMEVYVGTSLVSMYSRCGEWVLAARMFEKVPHK----SVVTYNAFISGLMENGVMNLVPSVFNLMRksSEEPNDVTFVNAITACASLLNLQYGRQL------------HGLVMKKEFQFETMVGTALIDMYSKCRCWKSAYIVFTELKD-----TRNLISWNSVISGMMINGQHETAVELFEKLDSEGLKPDSATWNSLISGFSQLGKVIEAFKFFERMLSVVMVPsvLTSLIDMYMKCGLSSWARRIFDRFEPKPK---DPVFWNVMISGYGKHGECESAIEIFELLREEKVEPSLATFTAVLSACSHCGNVEKGSQIFRLMQEEYGYKPSTEHIGCMIDLLGRSGRLREAKEVIDQMSEPSSSVYSSlqHLDPVLGemKLAELEPENPAPF-VILSSIYAALERWED-    
 649 UniRef50_UPI00019842FD PREDICTED: hypothetical pro...   359     2e-97  1   9.1%  85:579     5:506     ------------------------------------------------------------------------------------KEAHTQLIINGLTHPPPSLRPIISFSasGDIDYALLLLLrsTPPTVFLFNTTIRGFSrsLLSSVLLFVRMGVLSLAPNNFTFTFLFQGCSNcaFDLGRQFHGMVIKNSFEMDVFVRNSIIRFYSVCGRLNDARWVFDESSE----LDVVSWNSMIDGCIRNGNILEALSLFSKMTER----NDISWNMMISGYAQNGQPKEALALFREMqlDHGCWVHCYIGKKCVRVDSILSAALIDMYAKCGSIDLAMQAFSTSRKR------DVSAYTAAISGLAMNGCSEEALQLFEQMKGEGISPDGVSYIAVLCACSHAGWVEKGFHYFASMSdhGIRpdHYACMVDLLGRAGLLEEAEKFVASMPIK----PDNVIWGALLGACRVYGNAEMGQRVGSLLVESDQNHDGR-YILLSNIYAESMKGEDAEQVRKTMRRRKVDRVP----------GCSLIEVAGFVHEFFSGDRSHEKTEEIYLMWEEIVKEIKKFGYR------------------------    
 650 UniRef50_A5BC56        Putative uncharacterized pr...   359     2e-97  1  11.6%  81:581     1:470     --------------------------------------------------------------------------------MNHIYKLHARLLKTGHHNhaLRRLLLSCAASAPASLSYARSIFDLIapDTFAFNTIIRAHasSPSFSLSLFSKMTMAGVSPDHFTFPFVLKACA--RLQTGLDLHSLLFKLGFDSDVYVQNGLIHFYGCCGFLDFALKAFEEMPER----DLVSWSSMIACFAKNGFGYEALALFQRMQLVgvKPDEVIVLSVVSAISILGDLELGKWI------------RGFISRNGLEFTVSLGTALVDMFSRCGCIEESMRVFDEMGER------NVLTWTALINGLAVHGRSAEALRMFYEMRNHGFQPDHVTFTGVLVACSHGGLVSEGWHVFESIreYGMepEHYGCMVDLLGRAGLLNEACKFVDGMPIR----PNSIIWRTLLGACVNHNYIELAEKVKEKINELDSYHDGD-YVLLSNVYGGVGRWAEKAGVRNSMREKRISKKP----------GCSLINVNHLIHEFVAGDNNHPQFESIREFLVSMIDSLKVVGYTPD----------------------    
 651 UniRef50_Q94DV3        Pentatricopeptide (PPR) rep...   359     2e-97  1  11.5%   5:588    48:609     ----SLYARAAAPALHRHvlDRPSPPPLPCFNA--------GLSLPHPLALAVFRGLRHAYSPDAFSFPPLVSSAPSPPHLLALH---ALALRCGLAHDLFCASAMLRSCLRFGLADHARRLfeLPRRDVVVWNAMVNGFARLGchAVECFRMMREDgvEISSFTVTGILSVcaTADLGHGAAVHGLVVKSAFEQEVSVCNALVDLYGKCHKVDDAAMVFEGM--ADIVKDLFSWNSMLSALHYSADHAGTMKLFSRMRRVAVWPDAVTVAAVLPACAQTAALQVGREVHGYIVTSGLACHGA--------LDSFACNALVDMYAKSGALDEARRIFDRMQQR------DVASWNIMIDGYASHGHGKEALELFCQMTeeRLLPDEITLLGALSACSHSGLVEEGKNFLKRMKEesLEpeHYACVTDMLGRAGRLEEARKVVEDAGDVG-----VGAWRTYLAACRMHGDKERAQEAA-RMLMTTEQSESGGWVLLANTYGWEGNFDELEEVRGEMKRRGVQKAAP---------GCSWVEVGgvMHAFVSGDRGHPEADMIYEMLHALISWMRDCSHLSNTTPLYSV---------------    
 652 UniRef50_Q9FLJ4        Pentatricopeptide repeat-co...   358     2e-97  1  12.4%   9:598    50:599     --------KCRSAEEAFKLFETSSRsdLQSFSAVIHVLTGAHKYTLARCLIklIERLKRHSEPSNMSH-RLFNALEDIQSPKFSIGVF---------------SLLIMEFLEMGLFEEALWVSREMkpDSKACLSILNGLVRrfDSVWVDYQLMISRGLVPDVHIYFVLFQCCFKQGlsKKEKLLDEMTSLGIKPNVYIYTIYILDLCRDNKMEEAEKMFELMKKHGVLPNLYTYSAMIDGYCKTGNVRQAYGLYKEILVAELLPNVVVFGTLVDGFCKARELVTARSLFVHMV------------KFGVDPNLYVYNCLIHGHCKSGNMLEAVGLLSEMES--LNLSPDVFTYTILINGLCIEDQVAEANRLFQKMKNERIFPSSATYNSLIHGYCKEYNMEQALDLCSEMTASGVEPntFSTLIDGYCNVRDIKAAMGLYFEMTIKGIV-PDVVTYTALIDAHFKEANMKEALRLYSDMLEAGIHPNDHTFACLVDGFWKEGRLSVAIDFYQE----------NNQQRSCWNHVGFTCLIEGLCQNGYILR------------ASRFFSDMRSCGITPD-ICSYVSMLKGHLQEKR-----    
 653 UniRef50_Q6ZH58        cDNA clone:J023139G21, full...   358     3e-97  1  12.4% 126:583    97:520     -----------------------------------------------------------------------------------------------------------------------------PVPNDVTLLTVLSACADSprLALSVHARVLK--LFPSHLLltCLVRFYFASRLPhaLQLFDAMPVR----SAVTYNTVISGLMRNGLVAAAFEVFDGMP----APDKVSWTALIDGCVKNGRHDEAIDCFRAMLLDGVEPDYVTLIAVISACAEVGALGLGMWV------------HRLVVRQGLERNVRIANSLIDMYARCGQVELARQVFSGMRKR------TVVSWNSMIVGFAANGRCADAVEHFEAMRREGFKPDAVTFTGVLTACSHGGLTDEGLRYYDLMreYGirMEHYGCVVDLLGRSGQLEEAMRVVTTMPMR----PNEVVLGALLAGCRMHGDVGMAEQLMQHLLELDPGGDA-NYVLLSNIYAAVGKWDGAGKVRSLMKARGLRKRPGY----------SAVEIDGDVHEFVSGDRSHPQAEEISQMLGLLMHEM--AGHDYDHV--------------------    
 654 UniRef50_Q0WN01        Pentatricopeptide repeat-co...   358     3e-97  1  10.9%   4:565   156:717     ---LCLYGRLDLLEMAEQVFEDMPFKSLETWNHMMSLLGHRGFLKECMFFFRELVRMGAS--LTESSFLGVLKGVSCVKDLDISKQLHCSATKKGLdeISVVNSLISAYGKCGNTHMAERMFQDAGSWDIVSWNAIICATAknPLKALKLFVSMPEHGFSPNQGTYVSVLGVSSLVQlsCGRQIHGMLIKNGCETGIVLGNALIDFYAKCGNLEDSRLCFDYIRDK----NIVCWNALLSGYA-NKDGPICLSLFLQMLQMGFRPTEYTFSTALKSCC----------------VTELQQLHSVIVRMGYEDNDYVLSSLMRSYAKNQLMNDALLLLDWAsrRGQYhqPDTVSWNIAIAACSRSDYHEEVIELFKHMLQSNIRPDKYTFVSILSLCSKLCDLTLGSSIHGLITKTDFsaDTfnVLIDMYGKCGSIRSVMKVFEETREK-----NLITWTALISCLGIHGYGQEALEKFKETLSLGFKPDRVSFISILTACRHGGMVKEGMGLFQKMKDYGVE-PEMDHYRCAVDLLARNGYLKEAEH----LIREMPFPADAPVW--------------------------------------    
 655 UniRef50_UPI0001984327 PREDICTED: hypothetical pro...   358     3e-97  1  10.0%   5:544   152:688     ----ALIHKLCSFRrvKEVLDEMPspDESIFVTIVRGLGRARMVRQMIKVLDLITKFGENPSLKI--FNSILDVLVKE-DIDLAREFYRkmMMNGVSGDDYTFGILMKGLCLTNRIGDAFKLLQVMkpNTVIYNTMIHALCKNGkrARSLMNEM----VEPSDVTFNVLISAYceENLVQALVLLEKSFSMGFVPDVVTATKVVGILCKAGRVTEGVEVLERVESMGGVVDVVAYNTLIKGFCMLGKAKVGHRVLKDMEIKGCLPNVDTYNILASGYCDSGMLDSAIDLFNDMKTDGINW------------NFMTYDTLIRGLCSGGRMEDGFKILELMEESRGGAGGRISPYNSIIYGLYKKNQFEEALEFLTKMEKLFPRAVDRSLRIL--GFCNEGSIGDAKRVYDQMIKEGGVPsvYVCLIHGFCQDGNVREAFELINEMVDHGYF-PTAPTFNALISAFCGQGKVGSALKLMEDMVGRGCLPDMGSYSPMVDALCNKGDFQKAVRLFLQMVEKDILPDYSTWNSMLLCLTQETVWLEG-----------------------------------------------------------    
 656 UniRef50_Q8LQE8        Os01g0819800 protein n=5 Ta...   358     3e-97  1   8.8%  31:576    26:551     ------------------------------WSARVRTLTRLGRHREALALLRHGD---PSPPPHALALPAAVISCAKLHLASGVAQIHALAAKRGLLpdAYLLSALLSSYSRLRLLPLARQLLDELPLASTPpnSLISGCALHGVPafSLFRRMraAGVRFDAVTLLALVPVAPLGIVP---QLHALAARSGLAANTSVANCLVSVYARGGgaSLARQVFEEMPRA--SRDLVSWNAVISAHAQNGLAVEAVELYRRMRggGVEPDAVTLVGVLSSCAHTGARRVGLDVERYVRD----------TIPGFRDNLPLCNALINFHARCGSLAQAQELFDEMPER------SVVSWTTLIIGYGMHGQGDIAVDLFETMVSEATAPDSVAMVGLLSACSHAGMYDEGRKYFSTMERDYckLEHYTCMVDLLGRAGRLDEARELIASMP----MAADGAVWGALLGACKIHKNVEMGEEAFERVVS--LEpnVGYYVLMANIYADAGQLDGVAKVRAAMRRRGLRKEP----------GCSYVEHKGKVHLFMADDHSHPQARRIYELVVEL-ERMVKE---------------------------    
 657 UniRef50_P0C8Q5        Pentatricopeptide repeat-co...   358     3e-97  1   9.4%  16:536    57:572     ---------------VKRILKGFNGHDSFSWGCLVRFLSQHRKFKETVDVYIDMHNSGIPPSSHAVT--SVLRACGKMENMVDGKPIHAQALKNGLCGCVYVQTGLVGLYSRLGYIELAKKAFDDieKNTVSWNSLLHGYLESgdEARRVFDKIPEK----DAVSWNLIISSYAkgDMGNACSLFSAMPLK----SPASWNILIGGYVNCREMKLARTYFDAMPQK----NGVSWITMISGYTKLGDVQSAEELFRLMSKK----DKLVYDAMIACYTQNGKPKDALKLFAQMLERngTWVESYITEHGIKIDDLLSTSLIDLYMKGGDFAKAFKMFSNLNKK------DTVSYSAMIMGCGINGMATEANSLFTAMIEKKIPPNVVTFTGLLSAYSHSGLVQEGYKCFNSMKDHNLEpdHYGIMVDMLGRAGRLEEAYELIKSMP----MQPNAGVWGALLLASGLHNNVEFGEIACSHCVKLETDPTGY-LSHLAMIYSSVGRWDDARTVRDSIKEKK-------LCKTLGCSW-------------------------------------------------------------------    
 658 UniRef50_D1IAI7        Whole genome shotgun sequen...   358     4e-97  1  10.0% 133:582     2:420     ------------------------------------------------------------------------------------------------------------------------------------WNTMIRGYSrnPREAIVLYMSMIAKGipPNNFTFPFLLNSCArsSLEPGHEVHSHIIKHGFESDLFVRNALIHLYSVFGNLNLARTLFDE----SLVRDLVSYNTMIKGYAEVNQPESALCLFGEMQNS----DEVTLVAVLSACARLGAFDLGKRLYHQYIENGVFNQ-----------NTILTAAVMDMYAKCGSIDSALEIFRRVGKNMKTG----FVFNSMIAGLAQHGLGETAITVFRELISTGLKPDEVTFVGVLCACGHSGLIEEGKKLFESmnAYGIKpeHYGCMVDLLGRYGCLEEAYDLVQKMP----FEANSVIWRALLSACRTHGNVKIGEIAGQKLLEMEAQHGAR-YVLLSNILADANQWEEARQVRKVMEDHGIRKPPGWSY----------IELGGAIHRFVASDKSHPQGKEIELMLKDMAMRLKSAGYVPNT---------------------    
 659 UniRef50_Q7EZR5        Os08g0131000 protein n=16 T...   358     5e-97  1  10.9%  32:602    42:646     -------------------------------NALLTSYGRRGRLRDAQLLFDQM------PSRDVISWTALLTAYADGGDLASARLVFDDMPRRNAPSW---NALLSVYLRAARPRAAHALFYKMPAKNAVSYGAIISGLAkmLHEAELVYEEMPWQWR--DPVGSNALMAGYLrgELAMALRVFEGMTVR----DVISWSAMVDGLCKHGSVSEARRVFDAMPER----NVVSWTSMIRGYVKRGMCRDGLLLFLNMRREGVQVNTTTLSVALDACAAASLAREGIQIHNLIISMGFELDikRAFDCMQQKDIVSWNSLITGYVQHDMVEEAHVLFKLMHQKDAvpVKDEVAWTAIISSFITNGDYLSAVRWFCRMSQEGCKPNTIAFSCLLSALASLAMLNQGRQAHAYSINMGWVFdvHTSLVSMYAKCGRLAEAYHVFSS-----ISNPSLIAINSMITAFVQHGFVEDALKLFTKMQNAGYKPNHVTFLGILTGCARAGFVQQGYNYFESMRPVYGVEPNPEHYTCMVDLLGRAGLLAEALE----MINSMPQNDHSDAWAALLSASSLHSNLapYDATAYTVLSRMFSSAGMEDEE-    
 660 UniRef50_D1IWZ7        Whole genome shotgun sequen...   357     5e-97  1   9.1%   9:599    38:582     --------KSGDIINAGKLFEEIPKRDVVSWSIMIHGYNRNGFRRKSMELFSQMRI--SSLVPTSFTMVGVLVSIAGLGDPVLGQCVHGLILKYGLDSDfvVTALLNAYAKCGNVVDSYRVFEQLENPGLVSCSAIVSGFVYnfEEAVVLFNQFRKLGMVPNAATVLTLIRACVAlsRRLCESIHGMVVKLSLVLDVAVNNSVLDMYSSMLDLDAATRVFEGMECR----DVISWTTMINLLVCLEYASDALMLFRQMRNTGICNDVVVVMNLISACAILGDLKRGREI------------HTQAIVCGFGSELPLTNSIIAMYSKCGDLDSSRTVFDQTTGK------SLVSWTAMILGCVQNGYPREALKLLIKMreESFYLDSIMLIGVLSASGELALLELCQQLHCYAFESGFPR--ynSLISAYSKCGDVEPAYNVFGQM----GYIRDIVSWNAILNGYGINGHGEIAVALYHEMRKGRENPDAATYLCVLSACSHSGLVDDGLVIFNQMVEERTIRPSQDHCGCIVDLLARAGCFSDAREF--------------------VSRYMEKMG--PNA---WRALLSGCQLHGNV----    
 661 UniRef50_UPI00015CA2E9 PREDICTED: hypothetical pro...   357     6e-97  1  11.0%   4:541    68:619     ---NDLFEESTDAALALYFFrmGSKHTVESVCTMIHILVSGNMNHKAMDLLLHLISYNSGEehNIfvYGMLVNCYVKENMTQVALKLICKMRHLNIFPLIGVCNSLLKALLESEQLNLAWDFLKEMKSQGLGLNafISGYcgNIDTGWKLLMEMKYLGIKPDVVAYTIVIDSLCKMSlkEATSILFKMTQMGVFLDSVSVSSVVDGYCKVGKSEEAMDVLE---VFNLSPNIFVFNSFISKLCTDGNMLKAAKVFQDMCEMGLIPDCFSYTTMMAGYCKVKDISNALKYLGKMLKR------------GIRPSVATYTLLIDSCCKPGNMEMAEYLFQRMITEG--LVPDVVSYNTLMNGYGKKGHLQKAFELLSMMRSAGVSPDLVTYNILIHGLIKRGLVNEAKDILDELTRRGFSPdtFTNIIGGFSNKGNFEEAFLLFFYMSEH-HLEPDVVTCSALLNGYCRTRCMAEANVLFHKMLDAGLKADVILYNSLIHGFCSLGNIDDACHLVSMMIEHGIMPNNITHHALVLGYEKKCVE--------------------------------------------------------------    
 662 UniRef50_D2STE4        Pentatricopeptide repeat pr...   357     6e-97  1   9.3%  72:574    14:517     -----------------------------------------------------------------------LSLLEQCRTMSQIKQMHSHLIVSASRLDPFAAGKIISLFAVSSNAdaYKLFLSLPHRTTFIWNTIIRIFVEKNetALSLYKNMLQTGFLPNNYTFSFVLRACTDNS-PVGLASHAQVIKLGWESYDFVLNGLIHLYANWSSVEAARKLFD----VSTCRDVITWTALINGYVKSGHVEFARELFDQMPER----NEVSWSAMITGYVHMGMFREALELFNDLQldHGRWIHAYVDRNGTELDRVLGTALVDMYAKCGCIEIACSVFEKMPDK------DAFAFTSLISGLANHGQSADAIQLFGRMQSEKVIPNEVTFICVLSACSRMGLVDEGLRIFNCmvVYGIEpqHYGCMVDLLGRAGLLEEAKRLVREMP----MEPDSYVLGALLNSCRVHGDVELGKETVESLVERGLD-HGGVHVLLSNMYASSNQWDWVVKVRKEMGAKKVKKVP----------GCSSIEIDGSVSEFIAGDMSYLRVEDVMLVLLGIDNHLK-----------------------------    
 663 UniRef50_D1HZK9        Whole genome shotgun sequen...   357     6e-97  1  10.8%  10:602   109:663     ---------LKSEEFVADVLDDRKVQMTPTDFCFVVKWVGQSSWQRALEVYE-WLNLRHWYSPNARMLATILSVLGKANQEALAVEIFARAEAASGNTVQVYNAMMGVYARTGRFT-----------------------------KVQELLDLMRSRGCEPDLVSFNTLINARLKSGtnLAIELLNEVRRSGIQPDIITYNTLISACSRESNLEEAVKVYNDMVAHRCQPDLWTYNAMISVYGRCGMSREAGRLFKDLESKGFLPDAVTYNSLLYAFAREGNVDKVKEICEDM------------VKMGFGKDEMTYNTIIHMYGKRGQHDLAFQLYSDMKLSGRS--PDAVTYTVLIDSLGKANMIKEAAEVMSEMLNARVKPTLRTFSALICGYAKAGKRVEAEETFDCMLRSGIKPDhySVMLDILLRFNESGKAMKLYQEMV-LHSFKPDHALYEVMLRVLGKENREEDVHKVVKDMEELCGMNSQVICSILVKG----ECFDHAANMLRLAISQGCELDQYGKARDFG-LFCGTEELFelDRSFYHIMMKMFRNSGNHSKAEKLLGVMKEAGVEP-TIATMHLLMVSYSGSGQPEEA-    
 664 UniRef50_A5AYV4        Putative uncharacterized pr...   357     7e-97  1   9.9%   6:598    98:725     -----MYSKFGDFGETFKVFEGMDERNLFSWTVMIGACSKNGDAEKAIELYGKM--ISSGVKADCFLYPLVLKSCGAVKDLRRGQCVHGEVLITGLLGDVVVMNSLIDMYMKCEMVGDSERVFDEMdrDVITWTTMIVGYMQMGreGLELLKEMLSSQVRPSSATLAGVLPLFSDLglKLAKQIHGLAAVTGFEYEKHVGTALVDVYANCGGLYFGRLVFDRVKEK----DVSCWNTMIKSYVQANLSDEAISLLKFMHLDGIHPVKSTWYCFFPHYSKSkcLEHadTLLDQMXEnqVKELHLFFKRSGGISNSSVGSSLXRMYSKFAVLEPAQEIFSCLGVKELDX------WNSILACYAHNGYANKASELFNKMRKTGIEPNVRSWNTLIAGFVDVGDFEAALETFSEMKWTNQRPN--------YWKGGFLXEASRIFHEMEVAGI-EGNTITWTTLVAGYAQXGLXDESLKHFRELQLKGLKPNSITIAsvLSACSQAGMVNEGWEHFNNMDPKYGLIPTGKHYTCIVDLLGRAGLFEDARNFIFQmlLSACKMHRNVEMAELAASHLLE--LQPENSENYMILSDIYAKARR-----    
 665 UniRef50_B9S8B7        Pentatricopeptide repeat-co...   357     9e-97  1  10.2%   5:464   139:577     ----KLVFMYVNCGDIReiFDKIANEKVFLWNLMLSGYAKIGAFEESVYLFRKMLDLGIQV--NSHTVSCILKCFAALGSVKEGEWVHGYLLKLGFGSYNTVVNSLISFYFKTRKIEAAYEVFDELkrDIVSWNSMISGTVAnpEKGIQVFKEMLYLGVSFDLVTLVNVLAACasGNFPLGRVLHAFAIKAQLDQRMTFVNTLLDMYSKCGDLNNAIRVFQKMGER----SVVSWTSLIAGYAREGLSGEGIRLFHEMEREGVRPDNFTVTAVLHACACSGSLEIGKDVHDYVKE------------NNMQKDRIVCNSLMDMYAKCGSMEDANLVFLEMPNK------DIVSWNTMIGGYSKNGRPNETLHLFVAMVQE-LKPDGRTMACILPACASLAALDRGREIHGYIQRNGcdLHVANALIDMYAKCGALALARLFFDMIPVK-----DLISWTVMIAG-------------------------------------------------------------------------------------------------------------------------------------------    
 666 UniRef50_D1ISS9        Whole genome shotgun sequen...   357     9e-97  1  13.3%  27:576    93:625     --------------------------NGFAYSSLLKLLARSRVFSEMEVVLENMRVEEMSPTREAMS--IVIQAYSDSGLVEKALELYYFVLKTyfPDVIACNSLLNMLVKLGR-IEIARKFTCIMVKGL--------CKEGKLEEGRKLIEDRWGQGCIPNIIFYNTLIDGYCkgDMEMANGLFIELKLKGFLPTVETYGAIINGFCKKGDFKAIDRLLMEMNSRGLTVNVQVYNTIIDARYKHGHIVKAVETIEGMIECGCKPDIVTYNTLISGSCRDGKVSEADQLLEQALGK------------GLMPNKFSYTPLIHAYCKQGGYDRASNWLIEMTERGH--KPDLVTYGALVHGLVVAGEVDVALTIREKMLERGVFPDAGIYNILMSGLCKKFKLPAAKLLLAEMLDQSVLPdvYATLVDGFIRNGNLDEARKLFELTIEKG-MNPGIVGYNAMIKGYCKFGMMKDAMACINRMKKRHLAPDEFTYSTVIDGYVKQHDLDGAQKMFREMVKMKCKPNVMDGLqiIPSLFAFANMECSELHCINYSSILDQYLPQGTSEASVILQTMFEE---------------------------    
 667 UniRef50_D1HF32        Whole genome shotgun sequen...   357     1e-96  1   9.0%   3:522    41:510     --ANNLLAMYVdlKDARRLFDETLERNVATWTTMISAYSSSGRPDTALKFYVQMLESKSE-TPNGFLYSAVLKACGLVGDLESGKLIHGRVFRANlfDTVLMNTLLDMYVKCGSLSSARKVFDDILCPSSTSWNTMISGYGKEgeEAVNLFYQMPE----PDTVSWNSIIAGFG---------------------------ILIDLYANDGKIKDALRLFYRLPEK----DIVVWSSLISWCTKMGLNSLVFSLFRDMVNLDIEVDQFIISSVLKACSSLVGLGSGKQV------------HSFCVKSGYESERITVTSLIDLYAKCGEIEDGLALF------YCTSERDTVCYTGIIMGCGQNGRAMEAVGFFQEMIELGLKPNEITFLGVLSACRHAGLVEEAWTIFKYMkeYKMEpeHYYCIVELLSQAGCFKEAEELIAEMP----FEPDQTIWNSLLGACGTHKKTELVNFIAERLLTT-LPEDPSILVTLSNVYATLEMWDDSRKMREVIKKVGMK---------------------------------------------------------------------------------    
 668 UniRef50_D1I6I7        Whole genome shotgun sequen...   356     1e-96  1   9.8%   4:542   222:732     ---NSLMDMYFKNgyALKVFDRLQDKDIISWNTVFAGLSQGDDAREIGRFFHKL--MLTGLKPNCVTFSILFRFCGEALDLVSGLQFHCLAFRFGISDEASVTSSLINMFSRCGAMRMACLVFDSApkSIHTCNEMISGYNLNCheALNLFCNLNGLGLEADECTFSSALEACFreNQKLGRQMHGTIVKSGFASQGYVCSSLLKCYVGFGLLDDSFEFFNGVER----LDLVSWGAMISALVHKGYSSEAIGLLNRLKEAGGKPDEFIFGSIFNCCAGIAAYRQTKSV------------HSLVVKMGYEAHVFVASAVIDAYAKCGDIENARRVFDQ-----TSRFRDVILFNTMVMAYAHHGLVREAVETFEKMKLATLEPSQATFVSVISACSHLGLVEQGDIFFKSmlDYGMDpdNYGCLVDLFSRNGFLEDAKHIIETMP----FPPWPAIWRSLLNGCRIHGNKELGEWAAKKLLQ--LVpnDAAYVLLSKVYSEEGSWSDAAKVRKGMIERGLWKDP----------GCSWIEI-------------------------------------------------------------    
 669 UniRef50_A5BZ17        Putative uncharacterized pr...   356     1e-96  1  10.2%  81:531     1:480     --------------------------------------------------------------------------------MQDLKQLHAQMIKTAQIRDPLAAAELLRFSAvhRDLDYARKIFRSMHrnCFSYNTLIRAlsNDPCDALLVFIEMVecSVEPNCFTFPSVFKACGRarLREGRQVHGLAVKFGLDSDEFVVSNVVRMYLSCGVMEDAHRLFYRrkKRRVDGDVVLWNVMIDGYVRIGELEVARNLFDEMPQR----SVVSWNVMIAGYAQSGHFKEAVEVFREMqlELGKWVHLYAVRNNIGVDDVLGSALIDMYAKCGSIEKALQVFEGLPKR------NVVTWSTIIAGLAMHGRAKDTLDHFEDMERAGVMPSDVTYIGLLSACSHAGLVNEGRWFFDHMVrsGLEpeHYGCMVDLLGRAGLLEESEELILNMPIK----PDDVIWKALLGACKMHGNVEMGKRVAEHLME--LApdSGSYVALSNIYASLGNWEGVAKVRLMMKEMDVSARWHPEDAC------------------------------------------------------------------------    
 670 UniRef50_B9N886        Predicted protein n=1 Tax=P...   356     1e-96  1  12.5%  60:578    43:603     -----------------------------------------------------------SPTNTIITHPILL-AMESCTSMLQLKQIQAHMTKTALISHTFPVSRVLAFCALSddINHAHLLFSQLqpNTYIWNTMIRGyaKMGQTGFLFFCQMVQKGVEMDCRSFVFALKACEqlGVLEGKSVHCVVWKMGFVYTLLVQNGLVHFYGLRGCLGLARLVFDEISER----DVVSWTSMIDGYSKHKWCDEALKLFDSMLMYGvePNEVTmnAILDMYVKCGCLDSAREIFDTmlDVGQWIHRNYVRMRYNEISVILANALIDMYAKCGVIHEAAKVFNDMPER------NLVSWNSMITAYASHGHAKQALSVFERMISGGFKPDDITLVGVLSACSHGGLVAEGQEYFQNMkkYGIEpeHYACMIDLLGRVGLLEDAYELITKMP----MEPSAAAWGALVHACRMHGNVEVAKIAAPRLLELDPE-DSGIYVLLANICASGRRWGDV-------KMARRMMRERRVKKIPGRSI---VEVEGQFHEFLAGDESHPQSEGIYNALDQLFAMSKLEGC-------------------------    
 671 UniRef50_Q9C8L6        Pentatricopeptide repeat-co...   356     2e-96  1  12.0%   5:589    58:647     ----SKHARNGNLQEAEAIFRQMSNRSIVSWIAMISAYAENGKMSKAWQV---FDEMPVRV---TTSYNAMITAMIKNkdLGKAYELFCDIPEKNAVSYAT---MITGFVRAGRFDEAEFLYAETPvrDSVASNVLLSGYLRAGkeAVRVFQGMAVKEV----VSCSSMVHGYCkgRIVDARSLFDRMTER----NVITWTAMIDGYFKAGFFEDGFGLFLRMRQEglEFDLFLGNSLMSMYSKLGYMGEAKAVFGVMKNK----DSVSWNSLITGLVQRKQISEAYELFEKMPGK----------------DMVSWTDMIKGFSGKGEISKCVELFGMMPEK------DNITWTAMISAFVSNGYYEEALCWFHKMLQKEVCPNSYTFSSVLSATASLADLIEGLQIHGRVVKMNidLSVQNSLVSMYCKCGNTNDAYKIF-----SCISEPNIVSYNTMISGYSYNGFGKKALKLFSMLESSGKEPNGVTFLALLSACVHVGYVDLGWKYFKSMKSSYNIEPGPDHYACMVDLLGRSGLLDDASNLISTmlLSASKTHLRVDLAELAAKKLIE--LEPDSATPYVVL--------------    
 672 UniRef50_C5WYF8        Putative uncharacterized pr...   355     2e-96  1  10.2%  19:589     1:624     ------------------MRQRGVDPGVNSWNCIISGCVQNALYDEA---LEFFLEMCESERPDAVTVASILPACAGLQALGIGKQLHSYVLRCGIKINVYVGASLIALYSECGEFDDARVVFSTieKNVNVWNELVQSYIREgdKAWEAFDLMQEDGLEPDIVTYNSFIAAYAkgQNEQAYELFSRMADVGLKPNVVSMNALICGLHRHGLYTDALEAFRYMqkNGLASNIYISSKLVDLYGKIGDMTSAANVFQKIGNK----NVVTWNSLMAAYKHNRMPEVTLKLLGEMFQSNLHPNLVTvmALGYGRELHSysALINMYGKCGNIEDARLVFK------STVPKDIAVWNAMMSCYLLHRMPMDIIDLFNYLEQSGIQPDHITFILLLSACKQEGLFEEAQSYFYNMEdyGIKPshYTCMVDIMGSAGLLAESLTLIQKMPLK----PDACLWSTVLKACKLHSNLEIGEKAAKALFE--LEpnPSNYMVLSNIYADTGLLDASEAVRDAMTEQGLHVDRQ-------CSWLYNG---TTVHSFEAGNLSHPAIDAILSTWKHLTVRMEQSRYSTEDIGPYYNV--------------    
 673 UniRef50_D1HYS8        Whole genome shotgun sequen...   355     3e-96  1   8.9%  62:591    18:587     -------------------------------------------------------------EADPMTKDSCLnsLRCCKTLNQLKQLHCQITKNGLDQIPSTLTKLVNAGasPESLDYARKAFELFKEDvfMLNSLIRGyaGLGREAILLYVRMLVLGVTPNHYTFPFVLSGCTkaAFCEGIQVHGSVVKMGLEEDVFIQNCLIHFYAECGHMDHGHKVFEGMSER----NVVSWTSLICGYARGDRPKEAVSLFFEMldECVDRNLVLYNTILSNYARQGLAREALAILDEMLQQggYWVHTYIEKNGIPCDMRLNTALVDMFARCGDPQSAMQVFNKMTER------DVSAWTAAIGTMAMEGNGEGATGLFNQMLIQGVKPDVVLFVQVLTACSHGGQVEQGLHIFSLMEDHGISpeHYGCMVDLLGRAGLLREAFDLIKSMP----MEPNDVVWGSLLAACRVHKNVEMATYAAERI--NELAprAGVHVLLSNIYASAGKWTDVARVRLNLREKGVRKVPGS----------SSVQVNGVIHEFTSGDESHPEMTHIALMLQEMNCRFSDAGHIPDLS---NVLLD------------    
 674 UniRef50_O49399        Pentatricopeptide repeat-co...   355     3e-96  1  13.2%   8:521    47:538     -------ERAKSLTEIQqmLKTGLFHDTFSASKLVAFAATNPEPKTVSYAHSILNRIGS---PNGFTHNSVIRAYANSSTPEVALTVFREMLLGPVFPDKYSFTFVLKACAAFCGFEEGRQIHGLFIKSGLVTDVFvyGRSGYFEIARKVLDRMPVR----DAVSWNSLLSAYLEKGldEARALFDEMEER----NVESWNFMISGYAAAGLVKEAKEVFDSMPVR----DVVSWNAMVTAYAHVGCYNEVLEVFNKMLDDStkPDGFTLVSVLSACASLGSLSQGEWV------------HVYIDKHGIEIEGFLATALVDMYSKCGKIDKALEVFRATSKR------DVSTWNSIISDLSVHGLGKDALEIFSEMVYEGFKPNGITFIGVLSACNHVGMLDQARKLFEMMSsyRVEpeHYGCMVDLLGRMGKIEEAEELVNEIPADEA----SILLESLLGACKRFGQLEQAERIANRLLELNLR-DSSGYAQMSNLYASDGRWEKVIDGRRNMRAERV----------------------------------------------------------------------------------    
 675 UniRef50_D1IW91        Whole genome shotgun sequen...   355     3e-96  1   9.9%  67:571     8:483     ------------------------------------------------------------------SLSCILRKCVPHSAISQAKQTHAQILVHGFIPNITLQTDLLLVYSKCGVLQDARKVFDKmeRNMHSWNILIASYAHnfYDALGVFDSFLKMGFRPDHFTLPPVFKACAggDSYLGKMLHSWVIRIGFEEYVVVGSSVLDFYAKCGGLVDAWRCFVNMSWR----DSVVWNLMIVGLGKACFFRDALECFRDMLSEGVKMDSRTVPSILSVCGGEGDLMKGKEIHGQVVKNQIF---------GCE--VAIGNSLIDMYAKCGCLHDSEKVFTTMSE------LNLVTWTSMISCYGVHGKGHEALALFKKMKYCGFQPNCVTITAILASCSHSGLIEQGRKIFYSilDYGFEpeHYACMVDLLGRFGYLEEAFELIKNMKS----AATASVWGALLAGCLMHKNIEIGEIAAHCLFE--LEpnSSNYIALCSMYDSLGIWDGVSRTRAKMRELGLVKTP----------GCSWITVAGRVHKFYQEDLSHPETQMIYETLHGIVN--------------------------------    
 676 UniRef50_Q9FND7        Putative pentatricopeptide ...   355     3e-96  1   9.4%  81:582    19:525     --------------------------------------------------------------------------------FKEVRQIHAKLYVDglKDDHLVGHFVKaaLSDHKYLDYANQILDRSepTLFALNSMIRAHCKSpeKSFDFYRRILSSglKPDNYTVNFLVQACTGLRMRegLQVHGMTIRRGFdpDFVCRTAMVTACARCGDVVFARKLFEGMPER----DPIAWNAMISGYAQVGESREALNVFHLMQLEGVKVNGVAMISVLSACTQLGALDQGRW------------AHSYIERNKIKITVRLATTLVDLYAKCGDMEKAMEVFWGMEEK------NVYTWSSALNGLAMNGFGEKCLELFSLMKQDGVTPNAVTFVSVLRGCSVVGFVDEGQRHFDSMreFGIEpeHYGCLVDLYARAGRLEDAVSIIQQMPMK----PHAAVWSSLLHASRMYKNLELGVLASKKMLELE-TANHGAYVLLSNIYADSNDWDNVSHVRQSMKSKGVRKQP----------GCSVMEVNGEVHEFFVGDKSHPKYTQIDAVWKDISRRLRLAGYKADT---------------------    
 677 UniRef50_D1HRG6        Whole genome shotgun sequen...   355     3e-96  1   8.7%  72:555    30:516     -----------------------------------------------------------------------LHFSVTHKSLKLTRQSHSRILSLGLSQNSLLATKLIFAYAICQHPYHSRLVFDSlhKNVFLWNSLINGYAKnyNEAFQLFNQMCSSDVLPDDFTLSTLSKVSSELglFSGKSIHGKSIRIGFVSDTVVANSIMSMYCKCGNFEESRKVFDEMTIR----NSGSWNVLIAGYAVSGNLVVGRRVFDRMKCR----NVFSWTAMINGYVENGDSDEALSLFRDMQvlSGRQIHGFAVRKELNNEVSLCNALIDMYSKCGSLDSARRVFED-----DSLCKDAISWSSMISGYGLHGKGQEAILLYDKMLQAGIRPDMITTVGILSACSRSGLVNEGLNIYSSVidYGIEpeIFACIVDMLGRAGQLDPALDFIKAMP----VEPGPSVWGALVSCSIIHGDLEMQELAYRFLIQ--LEpnPSNYVSISNLYASSRRWDAVAEVRRMMKDKRLRKVP----------GCSWISINNKTHCFYTLKYS------------------------------------------------    
 678 UniRef50_A5BF28        Putative uncharacterized pr...   355     4e-96  1  13.5% 124:537    92:495     ---------------------------------------------------------------------------------------------------------------------------SSFHPDVICYNLLIDAYGQkyKKAESTYLELLEARCVPTEDTYALLLKAYCTSGleKAEAVFAEMRKYGFPPSAVVYNAYIDGLMKGGDTQKAVEIFERMKRDRCQPSTATYTMLINLYGKASKSYMALKVFHEMRSQKCKPNICTFTALVNAFAREGLCEKAEEIFEQLQE------------AGLEPDVYAYNALMEAYSRAGFPYGAAEIFSLMQHMGC--EPDRASYNIMVDAYGRAGLHEDAQAVFEVMKRLGITPTMKSHMLLLSAYSRAGKVAKCEEIVNQMHKSGIKPDTfnSMLNLYGRLGQFEKMEEVLTAM-EKGPYPADISTYNILINIYGRAGFFARMEELFRSLPARNLIPDVVTWTSRIGAYSRRKQYNRCLEVFEEMIDAGCYPDGGTAKNVLLKVAI------------------------------------------------------------------    
 679 UniRef50_B8BCQ1        Putative uncharacterized pr...   355     4e-96  1  11.5% 132:591    47:481     -----------------------------------------------------------------------------------------------------------------------------------ACASYTARMRLNPQLALRLFDHLLRSGADPDHVAYALALGRCARGrhRAAAQLHSHAAKRGAASHRRVCNGLIHAYAVCGSLLDARKVFD----RGHEGDAVAWNSLLRGYAAAGDVNALREFFVGMQAR----DTVSWNTIIAWCVENGEYEEAIAVFREMLASMECLPDRVTLfkGIEVEERLSSALINMYSKCACIEGAVHVFE-----NLGAQMNVDTWNAMLAGFTANGCSEKALELFARMEITGLVPNKITFNTVLNACSHGGFVEEGMGCFERMTkyGIEPdhYGCMVDLFCRAGLFDKAEKMIQMMPMK----PDAAVWKALVGACKTHRNFELGRKAGHMLIEA--APNDHayVLLSNIYALDGNWTGVHKVRKLMLDRGVQKVPGS----------SSIEIDGVIHEFISGDKSHSSKEDIYEMLSEMCQQLKVAGYVPD---TSHVLLD------------    
 680 UniRef50_B9H0N2        Predicted protein n=8 Tax=r...   355     4e-96  1   9.4%  12:573    69:669     -----------NLAQAKSIFDSASVRDLVTYNSMLSGYVSvgYERNALELFVEMQSKRNEIEIDDLTITSMVNLFSKLCNSCYGRQLHSYMVKTGNDRSGFVVSSLIDMYSKCGCFKEACQVFKGCEREgvSWNTLISGYVQNgvEALKLFVCMGENGVKWNEHTFGSVLSACADLrlKIGKEMHAWILKNGLGSSAFVESGIVDVYCKCGNMKYAESLH---LTRGVR-SSFSITSMIVGYSSQGNMVEACRLFDSLEEK----NSIVWAALFSGYVKLKQCEAFFELLREYIAKegKQIHGYVFRMGIEMDMKTTTAMIDMYSKCGSIPYAEKLFLKVIER------DLVLYNVMLAGYAHHGHEIKAINLFQEMLERGVGPDAVTFVALLSACRHRGLVDLGEKTFYSmeDYHILpdHYACMIDLYGRASQLEKMVLFMQRIPVEHQ---DAAVVGAFFNACRLNNNTELAKEAEEKLL--NIEGDSGayVQLANVYAAEGNWAEMGRISREMRGKEAKKF----------AGCSWVYLDNEVHSFTSGDRTHTKAESIYSKLEFLMAEL------------------------------    
 681 UniRef50_D1I733        Whole genome shotgun sequen...   354     5e-96  1   8.7%   4:602   153:821     ---NALIDMYAKCGELSsvFGGMEYRDIISWNSMMRGCAYNNYPKKSLWYFKKM--AYSSEQADNVSLTCAVSASALLGELSFGQVIHGWGIKLGyiSHNSFENSLISLYSQCRDIQAAEILFKEMkkDIVSWNAMLDGLaqRIWEAFDLLHEMQLLGcqPDSVTVVIIIPLCAELmlREGRAVHGLTLRREMGLDFSVTNSLIDMYSKCKDVkeAQHLFRQLLQsgFANNPLAVNSLMLMYINCGDLVACFSLLQTVSAA---ADIVCWNTVMAGCTQNGHFWEALKAFNLMRQdaGGSLHGLALKTLMESDIRVQNALITMYGRCGEIENARIIF------GFSCNRNLCSWNCMISAFSQNKDGRRALELFCHIE---FEPNEITIVGILSACTQLGVLRHGKQIHGHVIRSRLQGNSfaALEDMYSNCGRLDTAFQIFQSSPER-----SVAAWNSMISAFGFHSNGGKAIELFHEMRECGTRPTKSTFISLLSACSHSGLVNEGLWYYSNMLELFNVEADTEHHVCMVDMLGRAGRLGEAYEFIRQmlLSACSYHGDLKMGREVAELLFE--LEPENVGYYISLSNMYVAAGRWKDA-    
 682 UniRef50_Q0DD68        Os06g0249500 protein n=3 Ta...   354     5e-96  1  12.2%  24:497    95:570     -----------------------PPPSAPLLSAILSKLLARFSSPEPAARFLAASLAAGAPAPDVFAFNSILAALARVRDVPGMARIYALMqaSVRPDVVTYGILVNGLCKAGRVGDALRVLDGmiRPDVVTLNTVVDGLcgRVQEALAFVerMSshGCPPNTVTYNCLIDAFCrgNISMAYELVEKMENEGVPQNIVTLNTIVGGLCRAGRTGAALEFFRE--KRTVWpnAVTYSTLVGALLHTNNVGMAMELFHEKMSEGHSPDAIMYFTMISGLTQAGRLEDACSMASSMKE------------AGFKLDTKAYNILIAGFCRKKRLHEAYELLQEMKEVG--IRPDVCTYNTLLSGSCKAGDFAAVDELLGKMIDDGCQPSVITFGTLVHGYCKVGKIDEALRILRSMDESGIHPniYNTLIDFLCKRGDVDLAIELFDEMKEK-SVPANVTTFNALLKGLRDKNMPEKAFELMDQMREERCFPDYVTVDVLM----------------------------------------------------------------------------------------------------------    
 683 UniRef50_C5Z016        Putative uncharacterized pr...   354     5e-96  1   9.2%   5:557    66:590     ----AMYARCGRRGDAARAFDEMPLRDSVSWNS--LLAASASAADALALLRRMLRAAPGAAACDHATLTTVLSACARADGGGGTGavHGLAVSCGLDAEVSVGNALITAYFECSSPGSAERVFDAmdRNVVTWTAMVSGMArrYRESLSLFRQMRHA-VDANRASYSSSLLACasLAASEGQQIHGLVVKAGFETDLHVESELMDMYSKCGLMEDALRVFRSCQD----PDEVFLTVILVGFAQNGLEEKAFELFAEMVGKGIDIDANMVSAVLGAFGASAPFALGKQI------------HALVIKKCFGGNTYVCNGLINMYSKCGELQESIEVFDGTPSK------NTVTWNSIIAAFARHGHGSEVFGLFESMKADGVKPTNVTFLSLLHGCSHVGSASKGLEILNSMSsyGIHpeHYACVVDMLGRASQLDDAKAFIED----GPFKDSALLWQALMGACSFHKDSEVGKYAAEKLLL--LDPdtAAYVLLSNIYSSEGRWDDRARILKRMREKGLRK----------DTGKSWIELEKEVRSFVMAPRSRP----------------------------------------------    
 684 UniRef50_Q9SIL5        Pentatricopeptide repeat-co...   354     5e-96  1   9.2%  69:568    11:512     --------------------------------------------------------------------NYFIPFLQRVKSRNEWKKINASIIIHGLSQSSFMVTKMVDFCDKIEDMDYATRLFNQVspNVFLYNSIIRAYTHnyCDVIRIYKQLLRKSFepDRFTFPFMFKSCAsgSCYLGKQVHGHLCKFGPRFHVVTENALIDMYMKFDDLVDAHKVFDEMYER----DVISWNSLLSGYARLGQMKKAKGLFHLMLDK----TIVSWTAMISGYTGIGCYVEAMDFFREMQLAggKWIHLYAERRGFLKQTGVCNALIEMYSKCGVISQAIQLFGQMEGK------DVISWSTMISGYAYHGNAHGAIETFNEMQRAKVKPNGITFLGLLSACSHVGMWQEGLRYFDMMrdYQIEpeHYGCLIDVLARAGKLERAVEITKTMPMK----PDSKIWGSLLSSCRTPGNLDVALVAMDHLVE--LEpdMGNYVLLANIYADLGKWEDVSRLRKMIRNENMKKTP----------GGSLIEVNNIVQEFVSGDNSKPFWTEISIVLQL-----------------------------------    
 685 UniRef50_B9S6W0        Pentatricopeptide repeat-co...   354     7e-96  1   9.6%   5:545    77:614     ----ALIQKLCAFHrvYQLLDEMPhpDEEIFLTVIRGLGRARMIPHVIKVLDLISKFGKNPSLKI--FNSILDVLVKV-DIDVAREFYrqMMGSGVQGDDYTFAILMKGLCLTNRIGDGFRLLQVMKSRGVKPNAVVyaLCKNGkrARSLMDEIEE----PNDVTFNVLIAAYCKenLVQALVLLEKSFSLGFVPDVVTMTKVVEILCNAGRVTEAVEMLERVEYKGGLVDVVAYNTLLRGFCRLGKIKVAHRFLKEMERKGCLPNVETYNILISGFCDSGMFDMALDMFNDMKTDGISW------------NFDTYDTLIKGLFFGGRIEEGLKILELMEESKGGSGGRISPYNSVLYGLYKKNMWDEALEFLMKMEKLFPRAVDRSLRIL--GFCEKGAVKNAKMVFDQMINEGGTPnvYDCLVHGFCQEGNLREAFELMNEMVGHGYFLV-ASGFNALIHGFCGQGKDESALKLLDDMVGRGCVPDRGTYSPLIDALCRKGNFQKALSIFNQMIEKGITPDSSTWNSLLIRLSKEIIWLENK----------------------------------------------------------    
 686 UniRef50_Q5Z4A4        Os06g0314100 protein n=9 Ta...   354     7e-96  1   8.9%   4:529   484:987     ---NALLAMYADCGQLskLFHTMEVRTLISWNTIISGFAENG---DSVACLRFFCQMRLAdqFDLVTLIALISSLSAIEDITVGESVHSLAIRSGCNLDVSVANALITMYTNCGIIQAGEKLFDSLSsnTISYNALMTGYRKnfEEILPLFYHMIKNDQKPNIITLLNLLPIC--HSQLQGKTVHSYAIRNFSKLETSLFTSAICMYSRFNNLEYCHNLFCLVGER----NNIVWNAILSACVQCKQAGVAFDYFRQIQFLNVKTDAVTMLALISACSQLGKADLAE------------CVTAIALQKGFDGTIIVLNALIDMHSRCGSISFARKIFD------ISMEKDSVSWSTMINAYSMHGDGGSALDLFLMMVSSGIKPDDITFVSVLSACSRSGFLEQGRTLFRSMldHGITpeHYACMVDLLGRTGHLDEAYDIVTTMP----FRPSKSLLESLLGACRFHGNSKLGESVGKILTESD-HGNPRSYVMLSNIYASAGKWSDYERLRSDMEAKGLIKDVGVSL--------------------------------------------------------------------------    
 687 UniRef50_UPI00000A777A Os01g0197500 n=1 Tax=Oryza ...   354     7e-96  1  10.5%  27:576   230:823     --------------------------SAVPWNRLLRAHLGRSRGDLALALYRRMRALSPAL-PNSYTLPLALRAAA---SPRVASAVHAHALHLGLHAQHDVAGQILAAYSRLGRAADARRVFDAMprTTFHWNALISAYSsdPDAARDAFARMAAAGARPDAVTWTALLSAHArgKHADVLQLFGEMQRSGCEGnlFVTNSLICMYGKLGEMDDAKKAFRDATAK----NTVTWNTLITSYAAARLCDKALDVLAQMEQIgvAPNVVSWSAVIGGFASSGDTDRALELFRRMQqrLGRELHGHAMKAELDRHSLVENGLINMYAKCGKVSGARKVFDGMKTR------DLISWNSMLAGYGMHGLCDEALALFTDMAGATVEPDGVTFVAVLSACGHAGRVTEGRRLFDRMVrhKISpeHYTCMVYLLGRAGLLRDASELVETMPVR----PDLCVWGALLNSCRIHGDAAMAEATIANVLQSE---DQSTGNHMlnLYAMCGMWDESKKVRVMTKEAGLRKNPGQ----------SWIEVDNKVVAFAAGSanLTGAEDVFGMLDDLYAEMEDE---------------------------    
 688 UniRef50_D1ICR6        Whole genome shotgun sequen...   353     8e-96  1  10.1%  70:582    10:557     ---------------------------------------------------------------------TLLQNPSSVKSKSQAKQLHAQILRTSLPSPSLLSTILSIYSNLNLLHDSLL-IFNSLppTTLAWKSIIRCYTSHGlhSLSFFIQMLASGKYPDHNVFPSVLKSCTLmdLRFGESVHGCIIRLGMGFDLYTCNALMNMYSKFWSLEEvrKVFEMMPKR----DIVSWNTVISGNAQNGMHEDALMMVREMGNADLRPDSFTLSSVLPIFAEYVNLLKGKEI------------HGYAIRNGYDADVFIGSSLIDMYAKCTRVDDSCRVFYMLPQH------DGISWNSIIAGCVQNGMFDEGLKFFQQmrMEGVKPNYVAFMAVLTACSHAGLVDEAWKYFNSmqDYRIIpeHYAAVADLLGRVGRLEEAYEFISDM----HIEPTGSVWSTLLAACRVHKNIELAEKVSKKLFTVDPQ-NIGAYVLLSNIYSAAGRWKDARKLRIAMRDKGMKKKPA----------CSWIEIKNKVHAFVAGDKSHPYYDRINEALKVLLEQMEREGYVLDT---------------------    
 689 UniRef50_Q6K8H8        Os02g0468800 protein n=3 Ta...   353     9e-96  1   9.4%   6:525    96:594     -----MYAKCGRLPDARRLFDEMPRPTLVSWNCMVAAYGRSSQVEESVAVFNAMR--RAGVRPSEGTLVGVLSGCVDssASNPGMCVYGFSVKSGLDAGLPVLNSVLTMLVRGSHLDAARLLFDGinKSVVTWTALASGYLLRGdeVFDLFNRMRGVGKNVDSVVLVNLISAAVlgNLSVAKGVHALIIKLGFECEEDLAASLINLYAKCGDLESAREVFDAV----HMANVVVWTSMISGYVEGGHLNEALVMFDSMVCANIEPNEATLSSVLSACAKLGSANLGKKV------------EEQAIATGLHSEPRVATGLIDMYSKFGSINLARKIFEGVTNR------DIAVWSAMINGYACNGEGSEALVLFKEMKNKGFQPDGIAFTHVLTACNYSGLVDEGLECFHSmmEYGIEpeHHMCMVDLLCKAGHFGSALKFFKQMpqNKVLAPIISSYSA--------RCADSSIDFieELLNLETQ-DSDHCVLMSNMLSCLGKWKKATSYRRQLSKQGLMKEP------------------------------------------------------------------------------    
 690 UniRef50_Q9LVD3        Pentatricopeptide repeat-co...   353     9e-96  1  10.8%  12:602   190:849     -----------ALGFFESAVDSGvvPNLVTYTTLVSALCQLGKVDEVRDLVRRLEDEGF--EFDCVFYSNWIHGYFKGGALVDALMQDREMVEKGMNRDVVSYSILIDGLSKEGNVEEALGLLGKMIKEGVEPNLITycKMGKLEEAFVLFNRILSVGIEVDEFLYVTLIDgcRKGNLNRAFSMLGDMEQRGIQPSILTYNTVINGLCMAGRVSEalEIRRRFLEAKIPMDLVMCNILLKAFLLMGAYGEADALYRAMPEMDLTPDTATYATMIKGYCKTGQIEEALEMFNELRKSSVSA-------------AVCYNRIIDALCKKGMLDTATEVLIELWEKGLYL--DIHTSRTLLHslCKRGSFEAAIEVYMIMRRKGLtmDVIDYTIIINGLCKEGFLVKALNLCSFAKSRGVTLNtyNSLINGLCQQGCLVEALRLFDSLENIGLV-PSEVTYGILIDNLCKEGLFLDAEKLLDSMVSKGLVPNIIIYNSIVDGYCKLGQTEDAMRVVSR-KMMGRVTPDAFTVSSMIKGYCKKGDM---------------------EEALSVFTEFKDKNISAD-FFGFLFLIKGFCTKGRMEEA-    
 691 UniRef50_B8AKY4        Putative uncharacterized pr...   353     9e-96  1  10.6%   5:502    80:562     ----RLVGMYVLARRFRDAVarGAAACALPWNWLIRGLTMAGDYRSALLFYLKMWAHPSAPLPDSHTFPYVVKSCAALGAIALGRLVHRTARTLGLDGDMFVGSALIKMYANGGLLWDARQVFDGmeRDCVLWNVMMDGyaGSVSSAVELFGDMRASGCEPNFATLACFLSVSaeSDLFFGVQLHTLAVKYGLESEVAVANTLVSMYAKCKCLDDGWKLFGLMPRD----DLVTWNGMISGCVQNGFVDQALLLFCDMQKSGIRPDSVTLVSLLPALTDLNGFNQGKEL------------HGYIVRNCVHMDVFLVSALVDIYFKCRAVRMAQSVYD------SSKAIDVVIGSTMISGYVLNGMSQEAVKMFRYLLEQGIRPNAVAIASVLPACASMAAMKLGQELHSYALKNAYEGRCysALMDMYAKCGRLDLSHYIFSKISAK-----DEVTWNSMISSFAQNGEPEEALNLFREMCMEGVKYSNVTISSVLSACAS-----------------------------------------------------------------------------------------------------    
 692 UniRef50_D1HJV4        Whole genome shotgun sequen...   353     1e-95  1   8.8% 117:525   414:827     --------------------------------------------------------------------------------------------------------------------SSAQALFDKlqPNVFAWTAILGFYSRNgdECVRTYSEMKLKGVLPDKYVFPKVFRACGQlwLEVGIQVHKDVVICGCEFDLQVCNSLIDMYSKSGDVGSGRRVFDEMVER----DVLSWNSMISGYCK--RIQDALNVFELMDRF----DVVTWNAMILGFVDLEMGHLALECFSKMQRSggKQVHAYITKNSFSSVIPVWNALIHMYSKCGCIGTAYSIFSNMISR------DLVSWNTMIGGFGMHGLGQFALQLLRDMSHSDVCPNSVTFTSALSACSHSGLVDEGMELFHTMTRdgFTpeHFSCVVDLLARADRLEDAVGFIEKMPLK----PSKHIWSALLAACRAQQNVSVAKLAAEQLFQ--LEPEhgNYVTLSNIYARAGRWDDAVAVRKLMEDRGLVKPS------------------------------------------------------------------------------    
 693 UniRef50_Q8RZK3        Os01g0814300 protein n=4 Ta...   353     1e-95  1  11.1% 112:580    73:515     ---------------------------------------------------------------------------------------------------------------CGAAPSYARNLFDQiePTAFCYNSLIRALSSAAgdTVLVYRRMLRAGSppNSFTLAFALKACSVvaLGEGRQLHSQAFRRGLEPSPYVQTGLLNLYAKCEEVALARTVFDGMVGDK---NLAAWSAMIGGYSRVGMVNEALGLFREMQAADVNPDEVTMVSVISACAKAGALDLGRWV------------HAFIDRKGITVDLELSTALIDMYAKCGLIERAKGVFDAMAER------DTKAWSAMIVGLAIHGLVEVALKLFSRMLELKVRPNNVTFIGVLSACAHSGLVNEGRRYWSTMQELGIKPSmyGCMVDLFCRSSLLDEAYSFVTGM----AIPPNSVIWRTLLVASKNSNRFDIVQSASKKLLE--LEpnPENYVLLSNLYASNSQWDRVSYMRKKMKENNVNVV----------AGCSSIEINGYLHKFVVSDDSHPEIKEIRLLLREIADRVVRAGHKP-----------------------    
 694 UniRef50_B9MWY4        Predicted protein n=5 Tax=r...   353     1e-95  1   8.4%  99:588    42:533     --------------------------------------------------------------------------------------------------DPYAAAKIISLHAHSNARsyAERLFLCLqkSTFIWNTMMQAFVEKNerAFSLYKHMLESNYLPNNFTFSFVIRACIDvnLQMGLCFHGQVVKFGWESYDFVQNGLIHLYANCGFMDLARNMFD----MSIKRDVVTWTCLISGYLNSGQVLIARELFDRMPEK----NPVSWGALIAGYVRIGFFKEALEVFYDMQVSggRWIHAYVKRHHMSLDRMLGTALIDMYAKCGCIEMACSVFDEMDDR------DVYAFTCLISGLANHDKSEAAIDLFNRMQDEGVVPNEVTFVCVLNACSRMGMVDEGLRIFESMSNRYvqIQHYGCLVDLLGRAGKIEEAKQVVREMP----LQPDSYTLGALLDACRVHGDVQLGEEMVDSLVQRCLD-HGGVHVLLSNMYASADKWEDASKIRKKMEDKNIRKLP----------GCSSIEVNGTVCEFVTGDRSYALEEDIMFLLFGIDKQLKS--LFLDDDEHYNV---------------    
 695 UniRef50_A5AVZ9        Putative uncharacterized pr...   353     1e-95  1   9.4%   3:525  655:1154     --SNALLAFYSDCGKlfKLFQKMPLRNAISWNTLISGCVHNGDTKKAVALLHKMQQEKM--ELDLVTLISIIPICRVXENLIQGMTLHGYAIKTGFACDVSLVNALISMYFNCGDINAGKFLFEVMprSIVSWNALITGYRFHynEVMASFCQMIXEGQKPNYVTLLNLLPSCXT--LLQGKSIHAFAVRTGVIVETPIITSLISMYARFENINSFIFLF----EMGGKEDIALWNAIMSVYVQTKNAKESVTFFCELLHARVEPDYITFLSLISACVQLSSLNL------------SNSVMAYVIQKGFDKHIVISNALIDLFARCGNISIAKKIFEGLSSK------DAVSWSTMINGYGLHGDSEAALALLSQMRLSGMKPDGITYASVLSACSHGGFIDQGWMIFNSMVEEGvrMEHYACMVDLLGRTGQLNEAYDFVEKLPCKPSVSLLESL----LXACIIHGNVKLGEKICSLLFELDPK-NSGSYVMLYNIYAAAGRWMDANRVRSDMEERQLRKIP------------------------------------------------------------------------------    
 696 UniRef50_D1I9L5        Whole genome shotgun sequen...   353     2e-95  1  13.5%  79:602    75:611     ------------------------------------------------------------------------------RRPELGFRFFIWTTRRRSFRSWVTHNLVIDMLAKDDGFDTYWKILEELKNSNitFSVLIAAYAKSgeKAVESFGKMKDFGCKPDVFTYNSILHVMVQKEVFlaLAVYNQMLKLNYNPNRATFVILLNGLCKNGKTDDALKMFDEMTQKGIPPNTMIYTIILSGLCQAKRTDDVHRLLNTMKVSGCCPDSITCNALLDGFCKLGQIDEAFALLQLFEKE------------GYVLGIKGYSSLIDGLFRAKRYDEVQEWCRKMFKAG--IEPDVVLYTILIRGFCEVGMVDYALNMLNDMTQRGLSPDTYCYNALIKGFCDVGLLDKARSLQLEISKNDCFPttYTILICGMCRNGLLDEARQIFNQMENLGC-SPSIMTFNALIDGLCKAGELEEARHLFYKME---IGKNPSLFLRLsnGFHRVDREEDAFRVLDQMVKNGCT-PSSAVYKCLMTWSCRKGKLSVAFSLWLKYLRSLPSQEDETleLEKAVRCLLEMNFKLneIAPYTIWLIGLCQARRSEEA-    
 697 UniRef50_Q0JPW4        Os01g0197500 protein n=4 Ta...   352     2e-95  1  10.5%  27:576   311:904     --------------------------SAVPWNRLLRAHLGRSRGDLALALYRRMRALSPAL-PNSYTLPLALRAAA---SPRVASAVHAHALHLGLHAQHDVAGQILAAYSRLGRAADARRVFDAMprTTFHWNALISAYSsdPDAARDAFARMAAAGARPDAVTWTALLSAHArgKHADVLQLFGEMQRSGCEGnlFVTNSLICMYGKLGEMDDAKKAFRDATAK----NTVTWNTLITSYAAARLCDKALDVLAQMEQIgvAPNVVSWSAVIGGFASSGDTDRALELFRRMQqrLGRELHGHAMKAELDRHSLVENGLINMYAKCGKVSGARKVFDGMKTR------DLISWNSMLAGYGMHGLCDEALALFTDMAGATVEPDGVTFVAVLSACGHAGRVTEGRRLFDRMVrhKISpeHYTCMVYLLGRAGLLRDASELVETMPVR----PDLCVWGALLNSCRIHGDAAMAEATIANVLQSE---DQSTGNHMlnLYAMCGMWDESKKVRVMTKEAGLRKNPGQ----------SWIEVDNKVVAFAAGSanLTGAEDVFGMLDDLYAEMEDE---------------------------    
 698 UniRef50_Q9FNN7        Pentatricopeptide repeat-co...   352     2e-95  1   9.9%  71:574    22:494     ----------------------------------------------------------------------LLQRLLLIPNLVYARKLFDHHQNSC---TFLYNKLIQAYYVHHQPHESI-VLYNLLSFDGLRPSHHTFNFIFAASAsrLLHSQFFRSGFESDSFCCTTLITAYAKLglCCARRVFDEMSKR----DVPVWNAMITGYQRRGDMKAAMELFDSMPRK----NVTSWTTVISGFSQNGNYSEALKMFLCmkDKSVKPNHITVVSVLPACANLGELEIGRRL------------EGYARENGFFDNIYVCNATIEMYSKCGMIDVAKRLFEEL-----GNQRNLCSWNSMIGSLATHGKHDEALTLFAQMLREGEKPDAVTFVGLLLACVHGGMVVKGQELFKSMEehKISpeHYGCMIDLLGRVGKLQEAYDLIKTMPMK----PDAVVWGTLLGACSFHGNVEIAEIASEALFK--LEpnPGNCVIMSNIYAANEKWDGVLRMRKLMKKETMTKAA---------GYSYFVEVGVDVHKFTVEDKSHPRSYEIYQVLEEIFRRMK-----------------------------    
 699 UniRef50_B9SI11        Pentatricopeptide repeat-co...   352     2e-95  1  10.0%   7:415    75:496     ------LSSFSSLDYARKVFEEISQPNLYTWNTLIRAFASSPEPIHSLLIFIRML-YDSPDFPNKFTFPFVIKAAAGVASLPFSQAIHGMAIKASLGSDLFILNSLIHCYASCGDLDSAYSVFVKIekDVVSWNSMIKGFvgCPDKALELFQLMKAENVRPNDVTMVGVLSACAkmDLEFGRRVCHYIERNGINVNLTVSNAMLDMYVKNGSLEDARRLFDKMEEK----DIFSWTTMIDGYAKRRDFDAARSVFDAMPRQ----DISAWNVLISAYEQDGKPKEALAIFHELQLSkgGWIHVYIKKQDIKLNCHLTTSLIDMYSKCGEVEKALDIFYSVDRR------DVFVWSAMIAGLAMHGRGRAAIDLFFEMQETKVRPNAVTFTNLLCACSHTGLVNEGRMFFNQMESS--------------------------------------------------------------------------------------------------------------------------------------------------------------------------------------------    
 700 UniRef50_C7IXD4        Os01g0959600 protein n=3 Ta...   352     2e-95  1  10.2%   1:511    50:542     LAA---YCRLGGPLPARRLLDEMPRRNAVSFNLLIDAYSREGLAPLSLETLA--RARRAGVDVDRFSYAAALAACSRAGHLRAGRAVHALAILDGLSSGVFVSNSLVSMYSKCGEMGEARRVFdaEERDDVSWNSLVSGYVRAgeEMVRVFAMMRRGGMGLNSFALGSVIKCCsdGTMDIAEAVHGCVIKAGLDSDVFLVSAMIDMYAKKGALVEAAALFRSVQE----PNVVMFNTMIAGFCRtgkaSEALTLYSEVQSRGMQPTEFTFSSVLRACNLAGYLEFGKQI------------HGQVIKYTFQEDDFIGSALIDLYFNSGCMEDGFRCFR------SSPKHDIVTWTAMVSGCVQNELHEKALSLFHESLGAGLKPDLFTISSVMNACASLAVARAGEQIQCFATKSGFDRFTvnSCVHMYARSGDVDAATRRFQEMESH-----DVVSWSAVISCHAQHGCARDALHFFDEMVDAKVVPNEITFLGVLTACSHGGLVDEGLR--------------------------------------------------------------------------------------------    
 701 UniRef50_Q8S9D1        Pentatricopeptide repeat-co...   352     2e-95  1  12.8% 105:540   325:756     --------------------------------------------------------------------------------------------------------LMNGLIERGRPQE-AHSIFNTLIEEGHKPSLITYTTlkHFHSLLSLISKVEKNGLKPDTILFNAIINASSegNLDQAMKIFEKMKESGCKPTASTFNTLIKGYGKIGKLEESSRLLDMmrDEMLQPNDRTCNILVQAWCNQRKIEEAWNIVYKMQSYGVKPDVVTFNTLAKAYARIGSTCTAEDM-----------IIPRMLHNKVKPNVRTCGTIVNGYCEEGKMEEALRFFYRMKELG--VHPNLFVFNSLIKGFLNINDMDGVGEVVDLMEEFGVKPDVVTFSTLMNAWSSVGDMKRCEEIYTDMLEGGIDPDifSILAKGYARAGEPEKAEQILNQM-RKFGVRPNVVIYTQIISGWCSAGEMKKAMQVYKKMCgvGLSPNLTTYETLIWGFGEAKQPWKAEELLKDMEGKNVVPTRKT-MQLIADGWKSIG---------------------------------------------------------------    
 702 UniRef50_C5YQV5        Putative uncharacterized pr...   352     2e-95  1  10.8%   7:526    56:555     ------YARRGAVGRARQLFDEMQTRDVVSWTSLVSAYAGARDAQAVSRLVSEMRADGC--EPSAATLAVLLQACMAERDAAAGGQLHCYAVKSGWsdVVVLNSILTHLSRITGVDVAVRLFEQSPRSDAVSWNIIiySSEGRVSKVVDMYERMRREEVCPSCETLTSVVAAFAKCrlRQGQKLHSFGLKSGL-IDTILVASFVDFYAKCGELPSSVQLFEEFRGK----SNCIWSAMLAAFIHHGQFLDAIHLFGRMIDSSLVPSADVLRALVICYTELGDLRLGKVVHGYIIRNNCAAESHSCALE---------TSIVKLYARCGNIHLAERCFSSILHK------DSISWSSMIETYTIHGDGRKALALFRQMLEEGARPNGVTFLSLLSACGHSGLVSEARELFDCMTRkgIAPehYTCMVDVLGRSGNLEEAVQLINGM----TVKPDGRIWGALLASCKTHSNSKLANLAARKLME--LEPNNVGYHVvsNVQAGGSRWGEVEDIRSSMVDKDMQKSPA-----------------------------------------------------------------------------    
 703 UniRef50_Q6ZI32        Os02g0552100 protein n=3 Ta...   352     2e-95  1  10.8% 156:582   106:498     -----------------------------------------------------------------------------------------------------------------------------------------------------------KCGALAHPVVTNSLLKLYCSLGldRARRVL---YSGGAALDVVSWNTMVSGYGKGGDLGAAREVFARMPER----NLVSWSAMVDACVRAGEFGEALWVFDRMMREEFRPDVVVLVSVLKACAHLGAVERGRWVHRYLETGSFG---------GRRGNLMLETALVDMYCKCGCMEDAWQVFDGVHRR------DVVLWNAMIGGLAMNGYGERALELFRRMLQKGFMPNESTFIAVLCACTHTGRVDEGKRVFKSMQDYGIKpeHYGCLADLLGRAGNVEEAEALLLDMP----MEPHASQWGALMSSCQMHNDINVGERVGKRLIE--LEpdGGRYVVLFNLYAVNGRWEEARTIRQMMEDRGAKK----------ETGLSFIELNGLVHEFISGDTRHPLTRKIYALLEDIERRLQLIGYVKDT---------------------    
 704 UniRef50_A9TXR2        Predicted protein n=1 Tax=P...   352     2e-95  1  12.7%  95:582    19:479     ----------------------------------------------------------------------------------------------GFEPHFHLYVAFAHCLMRVQKWVALEILVDEMIGKNCPPDIKFYALvgRFQTAQHWFSKMKLLGCPPDIVIYNILILEYGkgNFSQAMKYFSQLKDEGLLPDSGTYCAVLSACRKVGNIDKGNETLKEMREAGIKPDQVAYSILIDMFGKAGRHEDAAATFRELQMSGYLPDSVAYNTLIHAFAKVGMVDQATLVFKEAQRSF------------CLDDPVIFYTMIQVYAKARMADQALHVLKLMKEIG--LQPNELGYGSVLNAFMRANQPMGAIKLFEEMQESNCRFGESTYITMLNVYSKAGFHSAAEELLAKMRQLGFRRNvySTLIDMYGKVGKLQEATRLFSTMKQQGCKG-NLIVYNTMLDMYGKAGRINDAERLMAELKDSKLVPDTVTYTTLINAYNRLGRFEECIRVFEQHRADGLKLD----------------------RILVAIMINVYGKARQYSKLAELLEVMPKSGIQPDT---------------------    
 705 UniRef50_Q9SJ73        Pentatricopeptide repeat-co...   352     2e-95  1   7.6%   3:579    38:585     --KTSLLNLYLKKgsAQMLFDEMPERDTVVWNALICGYSRNGYECDAWKLFIVMLQQGFS--PSATTLVNLLPFCGQCGFVSQGRSVHGVAAKSGLELDSQVKNALISFYSKCAELGSAEVLFREMkkSTVSWNTMIGAYSQSgeEAITVFKNMFEKNVEISPVTIINLLSAHVSH-----EPLHCLVVKCGMVNDISVVTSLVCAYSRCGCLVSAERLYASAKQD----SIVGLTSIVSCYAEKGDMDIAVVYFSKTRQLCMKIDAVALVGILHGCKKSSHIDIGMSL------------HGYAIKSGLCTKTLVVNGLITMYSKFDDVETVLFLFEQLQETP------LISWNSVISGCVQSGRASTAFEVFHQMMLTglLPDAITIASLLAGCSQLCCLNLGKELHGYTLRNNFENevCTALIDMYAKCGNEVQAESVFKSIKA-----PCTATWNSMISGYSLSGLQHRALSCYLEMREKGLKPDEITFLGVLSACNHGGFVDEGKICFRAMIKEFGISPTLQHYALMVGLLGRACLFTEALYLIWKMDI----KPDSAVWGALLSACIIHRELE------------------------    
 706 UniRef50_Q9FND6        Pentatricopeptide repeat-co...   352     3e-95  1   9.7%  34:579     1:518     ---------------------------------MIKANVYSCSKFRFLYRRRFLSQSSFVHSLDANV-SSLIAAVKSCVSIELCRLLHCKVVKSVSYRHGFIGDQLVGCYLRLGHDVCAEKLFDEmeRDLVSWNSLISGYSGRggKCFEVLSRMmeVGFRPNEVTFLSMISACVYGGseEGRCIHGLVMKFGVLEEVKVVNAFINWYGKTGDLTSSCKLFEDLSIK----NLVSWNTMIVIHLQNGLAEKGLAYFNMSRRVGHEPDQATFLAVLRSCEDMGVVRLAQGI------------HGLIMFGGFSGNKCITTALLDLYSKLGRLEDSSTVFHEITS------PDSMAWTAMLAAYATHGFGRDAIKHFELMVHYGISPDHVTFTHLLNACSHSGLVEEGKHYFETmkRYRIDpdHYSCMVDLLGRSGLLQDAYGLIKEMP----MEPSSGVWGALLGACRVYKDTQLGTKAAERLFE--LEpdGRNYVMLSNIYSASGLWKDASRIRNLMKQKGLVRA----------SGCSYIEHGNKIHKFVVGDWSHPESEKIQKKLKEIRKKMkeMGYK------------------------    
 707 UniRef50_Q9C6S6        Putative pentatricopeptide ...   352     3e-95  1  11.8%  29:595   100:683     ----------------------------PSFYTIAHVLIRNGMFDVADKVFDEMITNrrDRSLDADVCKFLMECCCRYGMVDKALEIFVYSTQLGVvpQDSVYRMLNSLIGSDRVDLIADH-FDKLCRGGIEPSGVshGFVLDAlkALDFHRLVMERGFRVGIVSCNKVLKGLSVDQIEVASRLLSLVLDCGPAPNVVTFCTLINGFCKRGEMDRAFDLFKVMEQRGIEPDLIAYSTLIDGYFKAGMLGMGHKLFSQALHKGVKLDVVVFSSTIDVYVKSGDLATASVVYKRMLCQ------------GISPNVVTYTILIKGLCQDGRIYEAFGMYGQILKRG--MEPSIVTYSSLIDGFCKCGNLRSGFALYEDMIKMGYPPDVVIYGVLVDGLSKQGLMLHAMRFSVKMLGQSIRLnvFNSLIDGWCRLNRFDEALKVFRLMGIYGIK-PDVATFTTVMRVSIMEGRLEEALFLFFRMFKMGLEPDALAYCTLIDAFCKHMKPTIGLQLFDLMQRNKISADIA-VCNVVIHLLFKCHRIEDASKFFNNLIEggYCSLRRLDEAERIFELLKVTPFGPNTV-TLTILIHVLCK--------    
 708 UniRef50_Q84SD0        Os08g0162200 protein n=4 Ta...   351     4e-95  1  10.8%  57:527    78:529     --------------------------------------------------------LRLGLLAHPYPRAALAHLYARLPDPSRAHSLLDETprAGAHSFLVSRNSLLASLLRSGDLAAARALFDRMprDVVSWNSMVAGLagHLDAAIELFDKMPER----NAASWNAVMCGyaQGDLAQARELFEQMPVR----SNVSWITMISGYANSGDVHAAGELFERMENKK---DLYAWNAMIACYAKNGCAREALAVFNRMLKPHvmPNEKTFSSVISACSQLGDLRFGLW------------AESFMGSVGIELDDHLRTALVDLHTKSGRIDRAFDLFRGLGMR------DVVSYSAIIVGCGMNGKFNEAVSLFKEMSDAKISPNAVTFVGLLSAYSNAGLMEEARACFASmeKYKISpeHYTIMVDLLGRSGKLDEAYQLIMQMPMK----PDASIWGALLLACRLHNNVELGEIVASKCFE--LEPGESGYYILlnIYSEANKWEKVKRLRKIMAERGLNKMPGS----------------------------------------------------------------------------    
 709 UniRef50_Q0J1T8        Os09g0413300 protein n=5 Ta...   351     4e-95  1   9.7%   3:526   309:808     --KEALISTYTNFGFIekVFEegTVSNRSIWSAFISGVSNHSLLRSVQLLRRMFHQ---GLRPNDKCYASV---FSSVNSIEFGGQLHSSAIKEGFIHGILVGSALSTMYSRCDNVQDSYKVFEEmeRDGVSWTAMVAGFATHGheAFLTFRNMILDGFKPDHVSLTAILSACNreCLLKGKEVHGHTLRV-YGETTFINDCFISMYSKCQGVQTARRIFDATPRK----DQVMWSSMISGYATNGCGEEAISLFQLMVAASIRIDSYICSSILSLCA-----DIARPFYC-------KPLHGYAIKAGILSDQSVSSSLVKVYSRSGNMDDSRKVFDEIS------VPDLVAWTTIIDGYAQHGSSQNALAMFDLMVQLGVRPDTVVLVSVLSACSRNGLVEQGFNYFNSMraYGVEPehYCCMVDLLGRSGRLAEAKYFVDSMPMK----PDLMVWSTLLAACRVHDDTVLGRFVENKIREGNY--DSGSFATLSNILANSGDWEEVARIRKTMKGVNKEPGWS-----------------------------------------------------------------------------    
 710 UniRef50_Q9FK33        Pentatricopeptide repeat-co...   351     4e-95  1  11.6%  35:578    94:610     ----------------------------------IEKLVLCNRFREAFELFEIL-EIRCSFKVGVSTYDALVEACIRLKSIRCVKRVYGFMMSNGFEPEQYMMNRILLMHVKCGMIIDARRLFDEieRNLYSYYSIISGFvgNYVEAFELFKMMWEELSDCETHTFAVMLRASalGSIYVGKQLHVCALKLGVVDNTFVSCGLIDMYSKCGDIEDARCAFECMPEK----TTVAWNNVIAGYALHGYSEEALCLLYDMRDSGVSIDQFTLSIMIRISTKLAKLEL------------TKQAHASLIRNGFESEIVANTALVDFYSKWGRVDTARYVFDKLPRK------NIISWNALMGGYANHGRGTDAVKLFEKMIAANVAPNHVTFLAVLSACAYSGLSEQGWEIFLSMSehGIKpmHYACMIELLGRDGLLDEAIAFIRRAPLKT----TVNMWAALLNACRMQENLELGRVVAEKLYGMGPE-KLGNYVVMYNMYNSMGKTAEAAGVLETLESKGLSMMPA----------CTWVEVGDQTHSFLSGDrnETVKRQIYQKVDELMEEISEYGY-------------------------    
 711 UniRef50_Q84TA9        Os03g0821700 protein n=3 Ta...   351     5e-95  1  11.9% 129:591    87:521     --------------------------------------------------------------------------------------------------------------------------------DAFLANTLIRAHATSPIHSLRlfPLMLRGAVVPNKFTFPFLLKACAagSPDVGLQAHAAALKFGFATDQYVSNTLIHMYSCFgfLGDARNVFDRMPKE----SAVTWSAMIGGYVRGGLSSDAVDLFREMQANGVQADEVTVIGVLAAATDLGALELARWV------------RRFVEREGIGKSVTLCNALIDTLAKCGDVDGAVAVFEGMQQR------SVVSWTSVIDALAMEGRGKEAVRVFEEMKVAGVPPDDVAFIGVLTACSHAGMVDEGCGYFDAMkeYGIEpeHYGCMVDMFGRAGMVERAMEFVRTMP----IQPNPVIWRSLVSACRAHGRLELGESITRSLL-HEYPAHEANYIMLSNVFALTQRWKEKSEIRREMSKRGIKKVP----------GCSIVELDGEVHEFIAGDESHPQYKDIYRMVEEMARELRRVGH---IAATSEVLLD------------    
 712 UniRef50_O49287        Putative pentatricopeptide ...   351     5e-95  1  10.4%   7:527   134:690     ------FAKAGELSVARRLFNAMPEKDVVTLNSLLHGYILNGYAEEALRLFKEL-----NFSADAITLTTVLKACAELEALKCGKQIHAQILIGGvpDDHSLSALISGYANCGRVNESRGLFDRKSNRCVILWNSMISGyaNNMkeALVLFNEMRNETRE-DSRTLAAVINACIGLGFLegKQMHCHACKFGLIDDIVVASTLLDMYSKCgrIDDAKRVFERIENK----SLISWNSMTNGFSQNGCTVETLEYFHQMHKLDLPTDEVSLSSVISACASISSLELGEQVFAR------------ATIVGLDSDQVVSSSLIDLYCKCGFVEHGRRVFDTMVKS------DEVPWNSMISGYATNGQGFEAIDLFKKMSVAGIRPTQITFMVVLTACNYCGLVEEGRKLFESMkdHGFVPdhFSCMVDLLARAGYVEEAINLVEEMP----FDVDGSMWSSILRGCVANGYKAMGKKAAEKIIE--LEpnSVAYVQLSAIFATSGDWESSALVRKLMRENNVTKNPGS----------------------------------------------------------------------------    
 713 UniRef50_Q9SN85        Pentatricopeptide repeat-co...   351     5e-95  1  11.0%  71:579    14:500     ----------------------------------------------------------------------LLSLIVSSTGKLHLRQIHALLLRTSlsDVFHHFLSRLALSLIPRDINYSCRVFSqlNPTLSHCNTMIRAFSLspCEGFRLFRSLRRNSSLpnPLSSSFALKCCIkgDLLGGLQIHGKIFSDGFLSDSLLMTTLMDLYSTCENSTDACKVFDEIPKR----DTVSWNVLFSCYLRNKRTRDVLVLFDKMKNdcVKPDGVTCLLALQACANLGALDFGKQVHD------------FIDENGLSGALNLSNTLVSMYSRCGSMDKAYQVFYGMRER------NVVSWTALISGLAMNGFGKEAIEAFNEMLKFGISPEEQTLTGLLSACSHSGLVAEGMMFFDRMreFKIKPnhYGCVVDLLGRARLLDKAYSLIKSMEMK----PDSTIWRTLLGACRVHGDVELGERVISHLIELKAE-EAGDYVLLLNTYSTVGKWEKVTELRSLMKEKRIHTKP----------GCSAIELQGTVHEFIVDDVSHPRKEEIYKMLAEINQQLKIAGYV------------------------    
 714 UniRef50_O65543        Pentatricopeptide repeat-co...   351     5e-95  1   9.0%   3:523    96:600     --SNSLISMYAKFSrvRKVFDEMLHRDTVSYCSIINSCCQDGLLYEAMKLIKEMYFYGFIPKSELVA--SLLALCTRMGSSSKVARMFHalVDERMQESVLLSTALVDMYLKFDDHAAAFHVFDQMekNEVSWTAMISGCvqNYEMGVDLFRAMQRENLRPNRVTLLSVLPACVELNysLVKEIHGFSFRHGCHADERLTAAFMTMYCRCGNVSLSRVLFETSKVR----DVVMWSSMISGYAETGDCSEVMNLLNQMRKEGIEANSVTLLAIVSACTNSTLLSFA------------STVHSQILKCGFMSHILLGNALIDMYAKCGSLSAAREVFYELTEK------DLVSWSSMINAYGLHGHGSEALEIFKGMIKGGHEVDDMAFLAILSACNHAGLVEEAQTIFTQAGKYhvTLEHYACYINLLGRFGKIDDAFEVTINMPMK----PSARIWSSLLSACETHGRLDVakIIANELMKS--EPdpANYVLLSKIHTESGNYHAAEEVRRVMQRRKLNK--------------------------------------------------------------------------------    
 715 UniRef50_A5BV08        Putative uncharacterized pr...   351     6e-95  1   9.5%   9:578    31:630     --------MYKAPVLHASLVKNGLIAHVYQCNILLQAYINSQALSDAHKLLHFM------PQPSVVSYNTILSGYFKFGLVSEAIKLFDGTSKRDCHSW---NIVLSGCVKNHKLGEGLTHFMK-MRCSSVRPDNFTYAIIIPCCdgQQVHADIVKVCSDLDAFIGTNLLRMYAegEIGDARKVFDGMPSRGL----VTWNAMISCYSKYGRGDKSIGLFRQLXREGISADEYTynALVNLYSKCGYVASASRLFEEIPDQ----DVVSWTVIIVGFLQSGHMEEAMWLFYQMQlqKGRHFHGLVLKFGLlaDVVVGSAVVDMYSKCGEMGDALRAFQEMPER------DIASWNGIICGYAQNGAGMKALKLYNEMVlsGIAPNEVTFVGVLCACSHNGLLKEGYSYFKEMVDKhiKpeHYTCMVDLLGRAGLLQEAEALILALPIK----PDNVMWGALLGACKLHGDVQMTRRTAEHLYTNEPR-NSSNYVLLANSYTDIGEWGEAVEIREVMEARGVEKT----------AGCSWVEIGTCMHSFLAGDXLHPQIEVASQVLPRLYLQMRERSC-------------------------    
 716 UniRef50_Q8S090        Os01g0912900 protein n=3 Ta...   351     6e-95  1  11.7%  26:519    42:508     -------------------------PPPVIRSLLNRAIRRLSKPHPRAALRLLLLMPRLPVSPDHFSLPFALNAAASLRLLPLGASLHALALRLALLplPVANALVDLYAKCDDLPAAHTALADIAAPDAVSFNSLLCAHArsVPDAESLFVAMPSR----TQVSWNAMVVVYVngDVSSARRVFDQMPTR----DSTSWSVLIVGYCKCGSMRSAREVFDRMPAK----NLVAWTAMINGYAQSGVPKESLALFREMEAAGIEPDAATMVGVISAASQIGSTELA------------GWVGSYVDKKRIERNDKVLTALVDMHAKCGNVDEALSAFREI------AQPDAYPYTALISGLAAHGHAKLALQVFERMQAQSVWPDPITFVGVLTACSHAGLVDKGLDYWEAMVkyGMeaDHYACVVDMLGRAGRLEEAFEMVQTMP----MGPHPGALGALLSACKTHGNVEIAEIVANKLFE--LEpnTGNYIMLSNIYAEKEQWEEAERIRSVMRTR------------------------------------------------------------------------------------    
 717 UniRef50_Q9LTF4        Putative pentatricopeptide ...   350     6e-95  1   9.2%  71:582    21:501     ----------------------------------------------------------------------LLLSSARTRSTIKGLQLHGYVVKSGllIPLVANNLINFYSKSQLPFDSRRAFEDSPQKSSTTWSSIISCFAQnpWMSLEFLKKMMAGNLRPDDHVLPSATKSCalSRCDIGRSVHCLSMKTGYDADVFVGSSLVDMYAKCGEIVYARKMFDEMPQR----NVVTWSGMMYGYAQMGENEEALWLFKEALFENLAVNDYSFSSVISVCANSTLLELGRQIHGLSIKSSFDSSSFVGSS------------LVSLYSKCGVPEGAYQVFNEVPVKNLGI------WNAMLKAYAQHSHTQKVIELFKRMKLSGMKPNFITFLNVLNACSHAGLVDEGRYYFDQMKESRIEPthYASLVDMLGRAGRLQEALEVITNMP----IDPTESVWGALLTSCTVHKNTELAAFAADKVFELGPV-SSGMHISLSNAYAADGRFEDAAKARKLLRDRGEKK----------ETGLSWVEERNKVHTFAAGERRHEKSKEIYEKLAELGEEMEKAGYIADT---------------------    
 718 UniRef50_Q5NAR1        Pentatricopeptide (PPR) rep...   350     1e-94  1  10.0%  71:578    18:489     ----------------------------------------------------------------------VLSLLPRLASPRHLLQAHAYLLPRGGHRHARVASALLLASLRLPLRDHAAALVRRVHPSVSLRAAARLRGRGGGGLAAQLHSLLVRAGHAADPHASASLVQAYCSCgvASARRVFDETAAS---ADVVSWNVMIDGYVKSGDLARARELFDVMPGR----NVVSWTMVIGAYAQMKQPEEAIEVFRRMQVEGIEPDGVALLSVLSACGDLGVVDLGEWVHRFVLRRGLCQEIPLM------------NAIIDMYVKCGSVKKALEVFEGMEQK------SIVTWTTMIAGFALHGLGLEAVELFRRMEKENVSPNDITFLAVLSVCSHVGLTDLGRWYFKTMVsyKIKpeHYGCMIDLLGRAGCLMEARGLIQDMP----FKANAAIWGALLAAARTHGDTELGEQALLHLIE--LEpnSGNYILLSNIYAEQERWDAVRELRISMRDRGLRNVP----------GASSIDVDGMVHEFTSRDGSHPSLHKIREVLCAINSNIKSVGH-------------------------    
 719 UniRef50_D1IC67        Whole genome shotgun sequen...   350     1e-94  1  10.0%  23:544     6:516     ----------------------GSPPDESIFVTIVRGLGRARMVRQMIKVLDLITKFGENPSLKI--FNSILDVLVKE-DIDLAREFYRkmMMNGVSGDDYTFGILMKGLCLTNRIGDAFKLLQVMkpNTVIYNTMIHALCKNGkrARSLMNEM----VEPSDVTFNVLISAYceENLVQALVLLEKSFSMGFVPDVVTATKVVGILCKAGRVTEGVEVLERVESMGGVVDVVAYNTLIKGFCMLGKAKVGHRVLKDMEIKGCLPNVDTYNILASGYCDSGMLDSAIDLFNDMKTDGINW------------NFMTYDTLIRGLCSGGRMEDGFKILELMEESRGGAGGRISPYNSIIYGLYKKNQFEEALEFLTKMEKLFPRAVDRSLRIL--GFCNEGSIGDAKRVYDQMIKEGGVPsvYVCLIHGFCQDGNVREAFELINEMVDHGYF-PTAPTFNALISAFCGQGKVGSALKLMEDMVGRGCLPDMGSYSPMVDALCNKGDFQKAVRLFLQMVEKDILPDYSTWNSMLLCLTQETVWLEG-----------------------------------------------------------    
 720 UniRef50_Q9ZU27        Pentatricopeptide repeat-co...   350     1e-94  1  11.4%  47:528   115:620     ----------------------------------------------AVEFFKLVPSLCPYSQNDPFLYNRIILILSRSnrFDRVRSILDSMVKSNVHGNISTVNILIGFFGNTEDLQMCLRLVKklKMNSFTYKCLLQAYLrdYSKAFDVYCEIRRGGHKLDIFAYNMLLDALAKD--EKACQVFEDMKKRHCRRDEYTYTIMIRTMGRIGKCDEAVGLFNEMITEGLTLNVVGYNTLMQVLAKGKMVDKAIQVFSRMVETGCRPNEYTYSLLLNLLVAEgrLDGVVEISKRYMTQGihRLFCDMWSFPVKGERDSYMSMLESLCGAGKTIEAIEMLSKIHEKG--VVTDTMMYNTVFSALGKLKQISHIHDLFEKMKKDGPSPDIFTYNILIASFGRVGEVDEAINIFEELERSDCKPdsYNSLINCLGKNGDVDEAHVRFKEMQEKGL-NPDVVTYSTLMECFGKTERVEMAYSLFEEMLVKGCQPNIVTYNILLDCLEKNGRTAEAVDLYSKMKQQGLTPDSITY---------------------------------------------------------------------------    
 721 UniRef50_Q7XTG3        Os04g0414800 protein n=2 Ta...   350     1e-94  1   8.4%  32:529    74:542     -------------------------------NAVLASLARHAPLRDALAQFNLLR--GGASRPDAFSFPPLLCACARASSLPTGASLHAAAIRLGVDADLFVRTALIQFYGRCGAAAAARALFDSlnPSEVSWTAIVTAYVNSGdtARELFDQIPHR----NVVHWNAMVDGYVkgDLEGARKLFDEMPER----TPAAYTSLIGGYLNAGNMGAARTLFDKLEDR----DLFAWSTMISGCAQNGYPGEALRIFNEFQKQEICPDELVIVGLMSACSQLGNITLARWIEGYI-----------MIYPIDMNNVHVMAGLINMNAKCGNMERATLLFESMSVR------DVFSYCSMMQGHCLHGSASKAVELFSQMLLEGITPDNAAFTVVLTACSHAGLVEEGKRYFDMMkeYMIVpdHYACLVSLLGRFGMLRDAYELIKSMP----GEPHPGAWGALLGGCKFHCDIELGKIAAKKLFE--IEpnAGNYVSLSNIYANIDRWGNVSETRAEMTGRGITKIAGCTL--------------------------------------------------------------------------    
 722 UniRef50_B9I9F9        Predicted protein n=1 Tax=P...   350     1e-94  1  10.6%  45:498    44:454     --------------------------------------------DDALASFRLM--LLRKPLPCIVEFSKLFSSIVRMGHYETVVYLSKEIEFLGIEQDIHSLSILINC----------------------------FCHLHRVDFGLSVLSKILKLGLEPSIVTFTTLLNGLceGKMDQVMMLYDDMLVRGLQPNVYTYNVIINSLSKSGKANEALGFLKQMEKVGCVPNVVNYSTLIDGYCLRGQMDEARSVFDLMVSKGCTPNVYTYTSLMNGYCKIERIEEAVQLLDE------------TLRKGLVPDIVTFTTIISGLCRAGRPLAAQQLFRYICAHGH--TPNIMTYGVLLDGLCKHGNLEEAFALFQEMQRSTVKPNLVIYTILIDSLCKCGKIKDGKELFSRLIDEGLKPntYTALVGALCKEGLIIEAHKLFRKMEEDGC-TPDKCAYNVIIQGFLQHKDPSMARQLVEEMVNRGFSADAAT-RALLN---------------------------------------------------------------------------------------------------------    
 723 UniRef50_B9MWE8        Predicted protein n=2 Tax=M...   350     1e-94  1  12.7%  77:589   116:634     ----------------------------------------------------------------------------SLNSPSLALRFFHFclSPNFHHDCFTYSRIILILSKSNLPDraRSIVSEMERNGvtVNILI-GFYENSEDLQKCIGLIKKWGLRMTGYTYKCLVQAYLRSRneKGFAVYLEMKKKGHMLDIFAFNMLLDALVKDSEVDHAYKVFEDMKRKHCEPDEYTYTIMIRMTGKIGKPDESLELFEEMLNKGYSPNVIAYNTMIQALANARMVDKAILLFLKMVEKECRPslLATERKLHKLDEVvyAYLVRTLKKLGHASEAHRLFCNMWNCHERGDRDACV--SMLECLCSAGKTTEAIDLLGKIHEKGVSVDTVMYNTVFSALGKLKQISPLHDLYEKMKQDGPLPdtYNILISSFGRAGKVDEAIKIFEELEDSDYK-PDTCSYNSLINCLGKNGHLDEAHMKFKEMCEKGLNPDVVTYSTLIECFGKTDKVEMACRLFDEMLAEG----------CYPNIVTYNILLDCLERSGRTA------------EAVDLYAKLKQQGLIPDSI-TYSVL--------------    
 724 UniRef50_Q75HW7        Os05g0439300 protein n=6 Ta...   350     1e-94  1   9.7%  70:552    16:507     ---------------------------------------------------------------------AVMARLRACVTFRDLLRVHGHVVRLRISQSSYLATQIVHLCNAHRRVTHAARVFAQVrpNLHLHNAMIKAYAQnhRDAVAVYIRMLRCPTSpdRFTYPFLLKAcgTAALELGKQVHTHVVRSGCDSSAIVQNSLIEMYTRAGDLALAHKVFDEMRER----DVVSWNMLISAHARLGQMRKATALFNSMPDK----TIVTWTAMVSGYTTVGDYPGAVDAFRSMQTEggRWIYAYCKRHGMLTSTHICNALMEMYAKCGCIDQALQLFDGMADK------DVISWSTVIGGLAAHGRAHEAVWLFTEMEKEgvRPNVITFVGLLSACSYAGLVDEGLSHFDRMNdyGVEpeHYGCVVDLLGRSGQIRRALDLVRDMP----VPADAKVWGSLLSACRSHGDVDTAVLAAERLVE--LEPDDvnLVMLANVYAAARRWSDVASTRKAIRSRSMRKTP----------GCSLIEVGNVVREFVAG---------------------------------------------------    
 725 UniRef50_B9G7I7        Putative uncharacterized pr...   350     1e-94  1  12.6%  43:597   203:764     ------------------------------------------RGGEVRAAWKVFEEMarGPRPNLAIFNAMIFGLCHRGLVRVSGGLLGVMGRFHVVPDACSYNILMKGHCLYGQAEDAFNLFDEMRVTGCCPTIVTYNIlgRMVEARMLFDEMLQAGVEVNTITFNVLIDGYAKAgmENANMACAEMKAMGLMPDCCTFNIliCRLCWDGRLDDAWELLCSAIEKGVQVSVSGFNALIAAYSKEGFDEEAFELYKTMNKLGLAPSSSTLNYLIMGLCNRGRLDQARLFLEYMVRMGYCV-------------IASFTIYLDSSFRAGDVLGALKCWDDM--KIVGLQPDFVAFSAYVNGLCRLDFLDEAYNGFIEMIRRGLVPNNFTYNSLISAFCRAGNVSEALKLEQKMRQSGLVPDIfnILIDGFCREGGLNTANNLFFGMYSIGL-TPDVVTYNTMLNAYCRSKDINGAMIFMNKMLADGCEPDIFTYNIWMHSLCSNHLLNRAMMLLDELAA----------TDCAPNSVTYNTLMDGICS-------------DVLDRAMILTGRLIKLAFQPNTVTI-NILFSHFCKNG------    
 726 UniRef50_Q69MX4        Os09g0407800 protein n=4 Ta...   349     2e-94  1  10.1%  81:575   142:597     --------------------------------------------------------------------------------PASARRLLADIPR---PDAVTWNTLLRACLRLGLLPAARRLFDEMPERDVVSYNSMVAGYVAEGDLarNLFDGMARR----DVVTWNSMISGYSrgDMENARKMFDAMPER----DVVSWNSMLDGYAQAGDVEMARLVFDGMPKR----SIVSWNVILALYAKLRDWRECLGLFDVMIAEgtVPNEKTFVSVLTACANLGDLEKGRWVHDLVQERWDR----------LVPDVLLLTTLLTMYAKCGVMETAREIFNSMGEK------SVPSWNSMIIGYGLHGQSEKALELFLEMERDGPRPNETTFICVLSSCAHGGLVLEGWWCFDRMVrySIEpeHFGCMMDLLGRAGLLEQSENLIENLQGKV----SEALWGILMSASQTQNNIKLGEFVGKKLIE--MRPTEVgyILLSNIYAAEGRWDDVEKVRKVMEEKGVEK----------DAGLSLVGSREGGHFINESGASAPRNDVMLCMLGEMSVHMKQ----------------------------    
 727 UniRef50_D1H6S1        Whole genome shotgun sequen...   349     2e-94  1  10.9%  50:574    85:583     -------------------------------------------------AFFKWVESNLNHKHSLQSSWTMIHTLAKHKQFKTAQNLLERI-------------AVRDYLSSPSVLNAVVRIHDDPDSniLSWLVIFYANSRMTQDAIQVFEHMRVHGFKPHLHACTVLLNSLAKARlnMVWKVYKKMVRVGVVPNIHIFNVLIHACCKSGDVEKAEQLLNEMESRFIFPDLFTYNTLISLYCKKGMHYEALGIQDRMERGGVSPDIVTYNSLIYGFCrvNDLEEALRLREVMEVE------------GLHPGVVTYNSILRKLCEEGKIKDANRLLNEMSER--KVEPDNVTCNTLINAYCKIGDMGSAMKVKKKMLEAGLKPDQFTFKALIHGFCKLHEVDSAKEFLFEMLDAGFSPstYSWLVDSYYDQDNKEAVIRLPDELSRKGLFV-DISVYRALIRRLCKIDKIESAERIFTLMQGKGMKGDSVVYTSLAYAYFKAGKAIAASDMLDEMDKRRLMITLKI-YRCFSASYAGDGSILGL--FWDHVIERRLISKNILKYIQQL-DYMR-----------------------------    
 728 UniRef50_Q9C7V5        Putative pentatricopeptide ...   349     2e-94  1   9.9%  12:525    55:547     -----------DLISARKLFDVFPERSVFLWNSIIRAYAKAHQFTTVLSLFSQI--LRSDTRPDNFTYACLARGFSESFDTKGLRCIHGIAIVSGLGFDQICGSAIVKAYSKAGLIVEASKLFCSIppDLALWNVMiyGCCGFWDKGINLFNLMQHRGHQPNCYTMVALTSGLIDPSLLlaWSVHAFCLKINLDSHSYVGCALVNMYSRCMCIASACSVFNSISE----PDLVACSSLITGYSRCGNHKEALHLFAELRMSGKKPDCVLVAIVLGSCAELSDSVSGKEV------------HSYVIRLGLELDIKVCSALIDMYSKCGLLKCAMSLFAGIPEK------NIVSFNSLILGLGLHGFASTAFEKFTEILEMGLIPDEITFSALLCTCCHSGLLNKGQEIFERMkeFGIEpeHYVYMVKLMGMAGKLEEAFEFVMSLQK----PIDSGILGALLSCCEVHENTHLAEVVAENIHKNGEERRSVYKVMLSNVYARYGRWDEVERLRDGISESYGGKLP------------------------------------------------------------------------------    
 729 UniRef50_Q9MA50        Pentatricopeptide repeat-co...   349     2e-94  1   8.5%  98:579    35:489     -------------------------------------------------------------------------------------------------TTVSWTSRINLLTRNGRLAEAAKE-FSDMTLAGVEPNHITFIALLSGceALglHGYACKLGLDRNHVMvtAIIGMYSkgRFKKARLVFDYMEDK----NSVTWNTMIDGYMRSGQVDNAAKMFDKMPER----DLISWTAMINGFVKKGYQEEALLWFREMQISGVKPDYVAIIAALNACTNLGALSFGLWVHRYVLSQD------------FKNNVRVSNSLIDLYCRCGCVEFARQVFYNMEKR------TVVSWNSVIVGFAANGNAHESLVYFRKMQEKGFKPDAVTFTGALTACSHVGLVEEGLRYFQIMkdYRISpeHYGCLVDLYSRAGRLEDALKLVQSMPMK----PNEVVIGSLLAACSNHGniVLAERLMKHLTDLNVK-SHSNYVILSNMYAADGKWEGASKMRRKMKGLGLKKQPGF----------SSIEIDDCMHVFMAGDNAHVETTYIREVLELISSDLRLQGCV------------------------    
 730 UniRef50_P0C8Q3        Pentatricopeptide repeat-co...   349     2e-94  1  11.0%  54:596   120:616     -----------------------------------------------------FEKFRHFMRLYLVTADSLLA----NGNLQKAHEVMRCM--------------LRNFSEIGRLNEAVGMVMD-MQNQGLTPSSITMNCVLEieYAENVFDEMSVRGVVPDSSSylMVIGCFRDGKIQEADRWLTGMIQRGFIPDNATCTLILTALCENGLVNRAIWYFRKMIDLGFKPNLINFTSLIDGLCKKGSIKQAFEMLEEMVRNGWKPNVYTHTALIDGLCKRGWTEKAFRLFLKLVRSDT-----------YKPNVHTYTSMIGGYCKEDKLNRAEMLFSRMKEQG--LFPNVNTYTTLINGHCKAGSFGRAYELMNLMGDEGFMPNIYTYNAAIDSLCKKSRAPEAYELLNKAFSCGLEAdtYTILIQEQCKQNDINQALAFFCRM-NKTGFEADMRLNNILIAAFCRQKKMKESERLFQLVVSLGLIPTKETYTSMISCYCKEGDIDLALKYFHNMKRHG----------CVPDSFTYGSLISGLCK------------KSMVDEACKLYEAMIDRGLSPPEV-TRVTLAYEYCKR-------    
 731 UniRef50_B9GYU2        Predicted protein n=1 Tax=P...   348     3e-94  1  10.0%  11:572    42:567     ----------RQLALTHKVLDEIPLSDTFAWNNLIHTHLSNRDPGGALSIYHHM--MMRGACPDRRTLPRVLTASRICGDLFLGKQLHGQAIKLGFFDEHYVITALIEIYGRLDGIEAGKWLFdsPRRNSVAWTMILKLYlnKPDLAINVFYQMVELNARIDSVVLITAAGACGLLkvEHGRRVHDVARKFRLESDILVSNSLLKMQIDCQRMEDARGFFNQMTTK----DVISWTEIICGYVKKGEFNEALKLFRKMNMDGIKPDSLSVSSVLPACART------------VAHKNGKEIHGYSLRNGMDNNLIVQNATTDMYAKSGLVDYALKVFERMKKR------DVISWTVMILGFSLHGKGELGVELFCRMEkqRVEADQFTYAAVLHCCTAACMVEEGKFYFNCIKEPNIT--HYALMVSLLARACLFDEARAFMEE----HHIERHAEVLRALLDGCWMHHRRNIGKQVFEQLCD--LEpdAENYVLLSNWYSDNGKWDLVDKLRETIMSMGSKPKKAY----------SWIEFQNKVHVFGTGDISHPRSERIYTELQCLMKK-------------------------------    
 732 UniRef50_D1HCJ3        Whole genome shotgun sequen...   348     3e-94  1  15.5% 173:526    93:433     ----------------------------------------------------------------------------------------------------------------------------------------------------------------------------CKESGVGQAMKLLDEMRNKGSKPDVVTYNVLINGICKEGRLDEAIKFLNNMPSYGCQPNVITHNIILRSMCSTGRWMDAEKLLSDMLRKGCSPSVVTFNILINFLCRQGLLGRAIDILEKMP------------MHGCTPNSLSYNPLLHGFCKEKKMDRAIEYLDIMVSRGC--YPDIVTYNTLLTALCKDGKVDVAVEILNQLSSKGCSPVLITYNTVIDGLSKVGKTERAIKLLDEMRRKGLKPdtYSSLVSGLSREGKVDEAIKFFHDLEGLGI-RPNAITYNSIMLGLCKSRQTDRAIDFLAYMISKRCKPTEATYTILIEGIAYEGLAKEALDLLNELCSRGLVKKSS-----------------------------------------------------------------------------    
 733 UniRef50_Q9FG16        Pentatricopeptide repeat-co...   348     3e-94  1   7.8%  72:586    16:539     -----------------------------------------------------------------------LALLQSCSSFSDLKIIHGFLLRTHLISDVFVASRLLALckPTNLLGYAYGIFSQIqpNLFVFNLLIRCFSTGAekAFGFYTQMLKSRIWPDNITFPFLIKASSEMECVlgEQTHSQIVRFGFQNDVYVENSLVHMYANCGFIAAAGRIFGQMGFR----DVVSWTSMVAGYCKCGMVENAREMFDEMPHR----NLFTWSIMINGYAKNNCFEKAIDLFEFMKREggERAYEYVVKSHMTVNLILGTALVDMFWRCGDIEKAIHVFEGLPE------TDSLSWSSIIKGLAVHGHAHKAMHYFSQMISLGFIPRDVTFTAVLSACSHGGLVEKGLEIYENMKkhGIEpeHYGCIVDMLGRAGKLAEAENFILKM----HVKPNAPILGALLGACKIYKNTEVAERVGNMLIK--VKPEHSGyvLLSNIYACAGQWDKIESLRDMMKEKLVKKPP----------GWSLIEIDGKINKFTMgdQKHPEMGKIRRKWEEILGKIRLIGYKGNTGDAF-----------------    
 734 UniRef50_Q6MWD7        B1358B12.18 protein n=6 Tax...   348     3e-94  1  12.7% 109:533   183:594     ------------------------------------------------------------------------------------------------------------YAEEGLLPDACTLVLLALRrePTSCTGLMSRFPTAP-EACAFYLQLLDAGLPPEAKLFNVLMRDLVrgELASAQNVFDEMQSRGVRRTVVSFNTMISGMCRAGDLDGAETLHRRMSEAGVTPDVYTYGALIQGLCRVGRIEDARGVFEKMCGRGMKPNAVVFTILIDAHCKKGDAETMLELHREMRER------------GVRPDAVTYNAIVNGLCRARDLKSASGIVVEMRSAG--LRPDTVTYTTLIDGYCKEEELDMAMEIKQNMVAEGVGLDEVTYTALISGLSKAGRSADAERVLGEMMEAGLEPdtYTMVIDAFCRKGDVKTGLRLLKEMQNKG-RKPGVVTYNVIMNGFCKLGQMKNADMLLNAMINIGVSPDDITYNILLDGHCKHGKVTD----IEELKSAKGTVPDLGVYTSIV----------------------------------------------------------------------    
 735 UniRef50_D1I4M3        Whole genome shotgun sequen...   348     3e-94  1   9.1%  65:581     3:516     ----------------------------------------------------------------AISTNPVVSVLDKCKSLCELRQIHAQMIKTNLLNHQFTVSRLIAFCSLSGvlDYASSVFSRihPNSFIFFALIKGFssNPVESLILYARMLScgVE---FSIPSVLKACGKlaFDEGRQVHGQVLKTHLWFDPFVGNSMVRMYIDFGEIELARRVFDRMPNR----DVVSWNSMIAGYLKAGEIELASELFDEMPER----DLVSCNAMIDGYGKE-MLSLGLriADLGFVEEGKWLHAYVSMNKIELSsfIGSALIDMYSKCGYIENAYHVFRSISHR-----RNIGDWNSMISGLAIHGLAREALDIFVEMERMDIEPNEITFLGLLSTCSHGGLVEEGQFYFESmeKYKIVpqHYGCMIDLFGRAGRLEDALGVIQNMP----FEADLLAWKAILSASMKHGHIEIGKSAALRAIE--LAPdsSSYVLLSNIYAKAGRWDDVAKIRLMMRQRGVKKI----------AGCSSMLVNGKVHEFLLGKedSSYSGQVLAKIAEVVSRLKLQGYEPD----------------------    
 736 UniRef50_Q6ZKI7        Os08g0138600 protein n=4 Ta...   348     3e-94  1  12.3%  35:579    85:600     ----------------------------------IEALAAEGRHDEARDAFRGARAGAPFTPLPASTYDALVASAAALREPGFAAAVLWHVESSGFEPGQRAWNRVLRMQLACGMLAEARQVFDGMprSEATWGTMMGGlaGRPRGALALFRELWEegDAAPRVVVVAVRAVTALGSARAGQQLHCCVAKTGMYEDQYLSCALIDMYNKCGKLDEARRVFDGVPQK----SVVAWNSMLASYSLHGCSEEALDLYHSMCEGGVDIDQFTFSTMLRVFSRLGLLEHAKQ------------AHAGLIQTGLPLDIVGNTALVDLYCKWGLMEDARNVFERMPIR------NLISWNALIAGYGYHGMGQKAIEMFEELIAEGIAPNHVTFLAVLNACRFSGFVEEGKRIFQLmqRTKPRAMHYACIIELFGQQGRLDEAY----SMIRKAPFIPTANMWGALLTASRIHRNLQLARLSAEQLLA--MEPQKInyVELLNLYINSGRQTEVSKVVNTLKRRGL----------CIHAACSWITVRKKDHRFFFKDSLHPQSSEIYRKLDSLLKEIKQLGYV------------------------    
 737 UniRef50_D1ITR6        Whole genome shotgun sequen...   348     3e-94  1  10.6%  68:547     5:542     -------------------------------------------------------------------YNSILSLLEKCKTMAELKRLHGLMITTSVIQDVIPLSRLVDFCadSGNLNYAKSVFNQIDrsLYIWNSMIKGYSIspDEALTMYREMRQKGYAPDHFTFPFVLKACSlnGYNLGQCVHNCIVKTGFEVDVYAATALLQMYAACGDMEAALKVFDDIPK----WNVVAWTSLIAGCISNDCPSEAVRVYKDMELWSVAPNEITMVNVLVACARSRDLneALDLFSDMRIAggQALHAYVSKTNLTDDTAIGTALVDMYAKSGDAERAQQVFSELQKK------DVTAWTSLIIGLAMHGHGEEALTFFKKMQEdaLIPDEITYIGVLSACSHVGKVEDGKNHFISMKnhGIEPthYGCMVDLLSRAGRLGEAERLVEKMP----VEPNTAIWSALLNGCKIYQNIDVADRVRRRVRELEVD-GSGVYVLLSNIYAGACRWEEVKMARELMKERKIQ-------KSLGHSSVEMKMVGGALH--------------------------------------------------------    
 738 UniRef50_Q9LR67        Pentatricopeptide repeat-co...   348     4e-94  1  12.9%  94:598    78:622     ---------------------------------------------------------------------------------------------RNLLPSPWVSQILnsASMESNLDGFCRKFLIKLSPNFVSFVLKSDEIREKPDIAWSFFCWSRkkKYTHNLECYVSLVDVLALadVDRIRFVSSEIKKFEFPMTVSAANALIKSFGKLGMVEELLWVWRKMKENGIEPTLYTYNFLMNGLVSAMFVDSAERVFEVMESGRIKPDIVTYNTMIKGYCKAGQTQKAMEKLRDMETRGHeaLYQEMDEKGIQVPPHAFSLVIGGLCKEGKLNEGYTVFENMIRKGS--KPNVAIYTVLIDGYAKSGSVEDAIRLLHRMIDEGFKPDVVTYSVVVNGLCKNGRVEEALDYFHTCRFDGLAINsySSLIDGLGKAGRVDEAERLFEEMSEKGCTR-DSYCYNALIDAFTKHRKVDEAIALFKRmeEEGCDQTVYTYTILLSGMFKEHRNEEALKLWDMMIDKGITPTAAC-FRALSTGLCLSGKVARACKILDElmINTLCKAGRIKEACKLADGITERGRepGRIRT--VMINALRKVGK-----    
 739 UniRef50_Q7XH82        Putative uncharacterized pr...   348     4e-94  1  12.5%   5:602    59:741     ----RLGDISSALSHFHRLVQSGAAPSPASTAALLRAMCSASMSTEAMDVLVL--SMGNPSPLPVSDFALLIPGLCSEGAVDKARFLFDAMLRSGlpPVRVYRSLAFAYCKARRSLDAslMLIKGMYLDRELSTALIRVFCREGRlaLDVFRRMkdEHVQLDAYAYTTMIWgfEHGRVDHGLQMYHEMIDRGIQPDAATYNVMIRWYCKSKWVGAAMDIYKVMIRTGVAPDLRCYTILMASLCKDGKLGEAENLFDKMLESGLFPDHVMfsSLAGGCSNMSLQKEADHLLDEIVTSNVLPvlGKLVAYGCEPSVLTYNIVIKCLCEQNRMDDARALITIMQSRG--VRPDMSTNSIMVTAYCKIGEIESALHLFGEMAKDGIEPSIAVYDSIIVCLCRMRRLKEAEATLRQMIREGLAPdiYTSLINGYSLTRQTRNVCRIFDEMLKRGL-QPGPHAYGSLINGLVKGNKIRKALGYLERMLEEGIAPQTVIYTMLINQFFRKGDVRLGLDLVVLMMKTNVAPDLIT-YGALVTGICRNisLAKKLKEARYMLFRMlcTEEMIQVAQGIIQDLEENGMVPD-LHIYNGMINGLCRANKMDDA-    
 740 UniRef50_Q683I9        Pentatricopeptide repeat-co...   348     4e-94  1  11.4% 109:582     1:486     ------------------------------------------------------------------------------------------------------------MSKGAAIIAYANPIfrHLKLESFLWNIIIRAIVHNvhSPISVYLRMRNHRVSPDFHTFPFLLPSFHnlHLPLGQRTHAQILLFGLDKDPFVRTSLLNMYSSCGDldDARKLFDEMPER----NVISWSCLINGYVMCGKYKEALDLFREMqeAFVRPNEFTMSTVLSACGRLGALEQGKWV------------HAYIDKYHVEIDIVLGTALIDMYAKCGSLERAKRVFNALGSKK-----DVKAYSAMICCLAMYGLTDECFQLFSEmtSDNINPNSVTFVGILGACVHRGLINEGKSYFKMMIEegITPshYGCMVDLYGRSGLIKEAESFIASMP----MEPDVLIWGSLLSGSRMLGDIKTCEGALKRLIELD-PMNSGAYVLLSNVYAKTGRWMEVKCIRHEMEVKGINKVP----------GCSYVEVEGVVHEFVVGDESQQESERIYAMLDEIMQRLREAGYVTDT---------------------    
 741 UniRef50_B9F234        Putative uncharacterized pr...   348     4e-94  1  12.3%  53:574     9:528     ----------------------------------------------------ELLALRRGGMNRSVCH----HLLTQCKTIRELQRIHAQALTHGLHPnsISCKIFRSYAEFGRPADAGRLFDEIPHPDIISFTSLMSLHLkhHWKAISVFSHAIASGHRPDGFAAVGALSASglGDQRIGSVVHGLIFRCGLDSELVVCNALVDMYCRCGKFEPARTVFDRMLVKDEVtsTVSWTALITGHVQDKQPIQALELFGKMLLEGHRPNHITIVGVLSACADIGALDLGRAI------------HGYGSKSNATTNIIVTNALMDMYAKSGSIASAFSVFEEVQMK------DAFTWTTMISSFTVQGNGRKAVELFWDMLRSGILPNSVTFVSVLSACSHAGLIQEGRELFDKMReyHIDpeHYGCMVDLLGRGGLLEEAEALIDHM----DVEPDIVIWRSLLSACLAHGNDRLAEIAGMEIIKR--EpdDGVYVLLWNMYALSNRWKEALDMRKQMLSRKIYKKP----------GCSWIEVDGVVHEFLVEDKTHDARREIYVTLEIMARHLK-----------------------------    
 742 UniRef50_A5C7H1        Putative uncharacterized pr...   348     4e-94  1  10.2%   5:536    68:519     ----GMYASCGDLQSARLVLERTQNPNVFAFNWMVSALAFHGYHEEAIGYFSLMQELGIV--ANKFTFSIVLKQCVGLMDFNKGKEVHCVISRTGLGNVVSVANSXIDMYCKCGHVGYGRK----------------------------VFDGMIER----DVVSWTSMICGYCngTLEEALVLFERMKVEGLEPNDFTWNXMIAGYARDGDCNGAFVLFSRMVREGLVPDLVTWNAMISGFTQSLKAVEAWRLFQDMXVAGIKPNQVTVTGLLPACGLMGSIHRGKEL------------HGLIYRMGXDMNVFVATALIDXYSKCGTVKDAWDVFDRIPIK------NVASWNAMIGCYGKHGLVDSSIQLFERMQAEGMQANHITLISVLSACSHGGLVEKGLTIF------------------------RMVEAYELFKEMP----IEVTDSIVGAFFNGCKIHGRRDLAKXMAEDILRMELK-RPGGFVTLSNIYAADGEWEEVENIRKVMKEQGVHKKPGFSWVEKRDGF-------------------------------------------------------------------    
 743 UniRef50_B9SJ30        Pentatricopeptide repeat-co...   348     4e-94  1   9.4%  23:530     2:461     ----------------------GSLPDIVTYNSLIDGYCKYGQVEEACLTIKRIRNAEC--RVDLVSFNALFNGFCKRKMKEEVFIYMGLMWKCCLPNVITYGTWIDTLCK-----------------------------VGDLDTGYKFFKEMRKDGIVPNLIAFTCLIDGYSkgNLDFAYQLYKDMCKSMHLPNVYTYAALINGFCKRGMLERAEWFFLKMLEVGIMPNSTVYTSIIDGHFKKGNIDVAMKYFSEMRKESFRLDIVAYGVVISGLVNNGRLDKVLEVMEDMVR------------NGLAPDKVVLTTLMHAHFKAGNTKAAYGVYRELLNRG--FEPDAVTISSLIDGLCKDGRYHDAKGYFCK-----EKANEVSYTALIDGICKEGNLDEVERVVMEMSESGFVPdvYTSWIAELCRQGKIVEAFKVKNKMVEEGIDL-DLLTYSSLIFGLANKGLMIEAKQLFDDMLKRGIIPDSMVFDILIRGYLKQDNPVAISHLHEEMRRRGLLTIDKITIS-------------------------------------------------------------------------    
 744 UniRef50_B9HK21        Predicted protein n=6 Tax=r...   348     5e-94  1   9.3%  84:569    51:545     -----------------------------------------------------------------------------------AFLVFGLVFALGDFLPSGSNS------SAHHLLSAHYVFKDIQSPSTILWNQMIRAHARspAKSIQFFNQMLLTDSRPDAHTYSFLLAAcsSLSLREGQQVHSKVLTNGyeRNVVCWNSMLAGYMRRGNLDGARRIFYEMPER----NVVSWTTMISGYAKNGKCKQALNLFDQMRKAGVELDQVVLLAALTACAELGDLKMGMWIHSYIQDTFVGSNQRVL--------VSLNNALIHMYASCGMIDEAYEVFRWMPER------SAVSWTSLITAFAKQGYAQAVLEIFRSMQRLGtrPDGITFIGVLCACSHAGLVDEGRQLFKDMIQrgIKpeHYGCMVDLLSRAGFLDEAQELIATMPVK----PNNAVWGALLGGCRFYRNAELASLVSQKLVAE-PDPDKAayLSLlqVYASAEKWQDVATVRQKMVAMGVKKP----------AGQSWVQINEVFHDSVAGDRTHENTSSIYEMLGKI----------------------------------    
 745 UniRef50_UPI0001984F80 PREDICTED: hypothetical pro...   348     5e-94  1  10.0%  78:577    16:531     -----------------------------------------------------------------------------CKSMRKLKQIHSQIITSPHLPKAHNHFLITRLLFVcgSLSYATDVFRLIqpSLSVYNIMIRAYACKDNGVDelYKQMLCSGIAPNYLTFPFLVKECTKRDGglGLSVHGQIIKFGFHGDLFVQNSLISFYSACGFLSTARILFDEMSNR----DVVSWNALITGYLRSGEIDSALGLFRKMEER----NVITWNSTITGLVQGGRPKEAVEYFFEMQvdHGKWVHSYLKRSGLECDMVIGTALVDMYGKCGCVESAIEVFRKMPKK------DILAWTAMISAFALHGLGKEAFDLLEEMETLGVRPNHVTFVGLLLACAHSGLVEKGRLCFDLMrvYLIEphHYACMVDLLSRAGLFEEAEGLIRCMP----MQPDVFVWGALLGGCQMHGNVELGKKVAQYLIDLDPS-NHAFYVILCDLYAKARKFDDLKRVRALMKERGIKKAVP---------GCSMIEICGVVYEFSVTGSPEVVMGEVEWVLNGLNNVMKMEG--------------------------    
 746 UniRef50_Q9C501        Pentatricopeptide repeat-co...   347     5e-94  1  10.6%  32:571    24:537     -------------------------------NQFISAVISKSRHLNHLKQVQSF-MIVSGLSHSHFLCFKLLRFCtrLCNLSYARFIFD---RFSFPNTHLYAAVLTAYSSSLPLhsSAFSFFRLMvrSVPRPNHFIYPLVLklSSAFsvHTHLFKSGFHLYVVVQTALLHSYASshITLARQLFDEMSER----NVVSWTAMLSGYARSGDISNAVALFEDMPER----DVPSWNAILAACTQNGLFLEAVSLFRRMINesIRPNEVTVVCVLSACAQTGTLQLAKGI------------HAFAYRRDLSSDVFVSNSLVDLYGKCGNLEEASSVFKMASKK------SLTAWNSMINCFALHGRSEEAIAVFEEMMKLNikPDHITFIGLLNACTHGGLVSKGRGYFDLMtrFGIEpeHYGCLIDLLGRAGRFDEALEVMSTMKMK----ADEAIWGSLLNACKIHGHLDLAEVAVKNLVALN--PNNGGYVAMmnLYGEMGNWEEARRARKMIKHQNAYKPP----------GWSRIEIDNEVHQFYSLDKSHPETEEIYMILDSLIS--------------------------------    
 747 UniRef50_C7J3Y1        Os06g0114366 protein n=19 T...   347     6e-94  1  14.1%  70:552    56:503     ---------------------------------------------------------------------ALIKALA-AEHGEAAPAVHAHAAKLGLDRRRAVRDGLVELYLARGELASARALVDGFprDVVSCTAMVTGHARHgdEAVVLFFAMADDRCVAIDAVAAAAAFSACagDLALGREAHRRVAERKVAMDVVAWNALVDMYAKCGDAAAAHRWFRRMP---VKKNVVSWNTMMSAFARAGELEEALALFQEMQAAAVRPDDATFVAALGACAQLGALDTGRWLHAYM------------GRMGHSADGVVGNALLDMYAKCGAVDQATEVFDGMARR------DVYTYTSMILGLAMHGRGEDALSLFAGMQRAGVTPNEVTLLGVLTACCHAGLVEEGLQQLNAMPE--PRIEHYGCVVDMLGRAGRLDKAEELIAAMP----VHSDALIWSSLLAACRAHGDVERAERV---MRRRVADADAGDYVLMSNTYASNGRHGEAVKVRGQMRRNEIDKVP----------GCSLIEIDGVVHEFKAI---------------------------------------------------    
 748 UniRef50_B9HPK8        Predicted protein n=2 Tax=r...   347     7e-94  1   9.0%   4:525   295:797     ---NALMDMYFKNGGMKsvFKKMHDRDVVTWNTVFGSFSQHEDPKDIASLFHSF--LLTSMRPNHITFSILFRECGKLLNLDLGLQFCCLALHFGLFDEANITSALINMFSRCGKMEMAHLVFKsvSENIIIWNELISGYKLNCCDAEAlfYDLLQLGVEANEYTFSNVLETCSRSENQLmrQIHGVAFKSGFASHGYVCSSLIKGYIKCGLLDDSLKVFN-MLDR---PDMAAWGTMISAFVHQGWDCEAIRSLNLLIEAGEKPDEFILGSILSSCA------------STVAYCQTKSVHSLIIKLGFEGHVFVASAVLDAYAKCGDIQSAKMAFNQ-----SCKSSDVVIYNAMIIAYAHHGRVVEALDTYDKMKLANLQPSQATFVSVIAACGHIGHVEKGCRLFKSMDLYGMepDIYGCLVDMFSRNGYLEDAKQIIESLP----YPAWPAILRSLLSGCRMYGNRELGEWAAKKLLQL-VPHNDAAHALLFKVYSELGNWEDAAKMRREMAERGLRKDP------------------------------------------------------------------------------    
 749 UniRef50_Q9FMA1        Pentatricopeptide repeat-co...   347     9e-94  1  10.5%  80:576    27:529     -------------------------------------------------------------------------------NLKTLKQSHCYMIITGLNRDNLNVAKFIEACSNAGHLRYAYSVfhQPCPNTYLHNTMIRALSlnAHsaITVYRKLWALCAKPDTFTFPFVLKIAVrsDVWFGRQIHGQVVVFGFDSSVHVVTGLIQMYFSCGGLGDARKMFDEMLVKDvvRNEVSWTCVISGYAKSGRASEAIEVFQRMLMENVEPDEVTLLAVLSACADLGSLELGERI------------CSYVDHRGMNRAVSLNNAVIDMYAKSGNITKALDVFE------CVNERNVVTWTTIIAGLATHGHGAEALAMFNRMVKAGVRPNDVTFIAILSACSHVGWVDLGKRLFNSMrkYGIHPnhYGCMIDLLGRAGKLREADEVIKSMP----FKANAAIWGSLLAASNVHHDLELGERALSELIK--LEPnsGNYMLLANLYSNLGRWDESRMMRNMMKGIGVKKMAGE----------SSIEVENRVYKFISGDLTHPQVERIHEILQEMDLQIQSK---------------------------    
 750 UniRef50_Q7XUX2        Os04g0446100 protein n=5 Ta...   347     1e-93  1  11.3%  49:515   125:558     ------------------------------------------------SALAVFTALHCSptPSAHSCQQIIVVLCRSGRQADALQLFDQMthYGYSPDARFLSFLVSSCTCANLLDASAT----------------------------LLSKASEFGCRVEAYAYNKLMSSlgRGRVHDVVALFewIQDRVYSPDVWSFNVVIKGVCRVGQVQKALELVERMNEFGCSPDTVTHNILVDGLCRTNEVSRGHEVLRRLQRDGvmPNVVTFTSVISGYCKAGKLEDAMAVYNDMVAS------------GIMPNTVTYNVLINGYGKVGDLGSAVEVYQQMTR--LRCPPDVVTFSSLIDGYCRCGQLDDALRIWSDMAQHRIQPNVYTFSIIIHSLCKQNRSDEAIGLLNEL---NLRPDiyNPVIDVLCKCGKVDEANLIRKGMEEKGC-RPDKYTYTILIIGYCMKSRISEAIMFFHEMVEAGCSPDSITVNCFISCLLKAGMPNEVDHVMRL----------------------------------------------------------------------------------------    
 751 UniRef50_B9SGL5        Pentatricopeptide repeat-co...   347     1e-93  1  11.1%   9:469   262:728     --------KCATLHLAEKKFEEAACKDIVTWNTMIVALAKSEKPIKALELF--FKMPRDAIRPNQITFASLISSCANLQIPMYAEFIHAKVIMHAFDTDVYVGSALVDYYAKCDKLDDARcfVKIHEKNVVSWNSLILGCANkpYAAISLLVEMLQCGYQPNEFSFSAVL---ISSSILELQQLHCLIIRMGYDNNDYVLSSLITSYGRNGLISDALVFLaeLLSQLEEPDNVSWNIAIAACARNGNYKEVFELFKQMLVAQIHPDNYTYVSLLSSSSQICDLALGSSIHGFLIKNNFSSC-----------DTFVCNVLLDMYGKCGCLRSSVKIFNSMRDR------NLITWTALISALGINSCAHEALERFKDMEHQGLRPDKVAFIAVLTACRHGALVGEGIELFKKMKSYGLEpdHYHCLVDLFSRHGHVKEAEKVISSMP----CPPNALIWRTFLEGCKKYR--------------------------------------------------------------------------------------------------------------------------------------    
 752 UniRef50_Q9LXF2        Pentatricopeptide repeat-co...   347     1e-93  1  10.5%  44:529    59:521     -------------------------------------------PGALKYAHKLFDEI---PKPDVSICNHVLRGSAQSMKPEKTVSLYTEMEKRGVSPDRYTFTFVLKACSKLEWRSNGFAFHGKVVRHGFVLNEYvhANCGDLGIASELFDDSAKAH----KVAWSSMTSGYAkgKIDEAMRLFDEMPYK----DQVAWNVMITGCLKCKEMDSARELFDRFTEK----DVVTWNAMISGYVNCGYPKEALGIFKEMRDAGEHPDVVTILSLLSACAVLGDLETGKRLHIYILETASVSSSIYVGTP-------IWNALIDMYAKCGSIDRAIEVFRGVKDR------DLSTWNTLIVGLALH-HAEGSIEMFEEMQRLKVWPNEVTFIGVILACSHSGRVDEGRKYFSLMRdyNIEPnhYGCMVDMLGRAGQLEEAFMFVESMK----IEPNAIVWRTLLGACKIYGNVELGKYANEKLLSMR-KDESGDYVLLSNIYASTGQWDGVQKVRKMFDDTRVKKPTGVSL--------------------------------------------------------------------------    
 753 UniRef50_C5XHZ2        Putative uncharacterized pr...   347     1e-93  1  10.0%   5:526    57:564     ----SLYAAAGRLADALRAFRAhpSANPRTYAVLVSALAR----PHPGLAFSLFSGSRRQLRPSPHVISAVLAACAGL-QPLCGRQVHACAAKVVPPGDVFVYTGMVDAYAKVGDMESSRKVFDEMPSRGasWNALLVGYARncLETMSVFRALAAQgvSLDQVSVSSVLSAcvAGALDFGRQVHACAAKVGLELSAvvSNALLDMYTRCGCSRESLALLNAVGCR----DVVTWNIIIRGCIHDNRFMEACVLFRSMVRTGTLPDDVAFATVLQASACM----SAWAIGASM--------HASVVKTGFLDSDGVASSLITMYSKCGCLDDAHQAFVGAKDHLH-----VLSWTAMITALQQHGHGMQAIDVFENMLENGIPPDHITFVSVLSSCSHSGLVEQGRKYFDSMTqhRITpeHYACMVDMFGRAGLLTEAKRFINQMR----PKPDASVLGALLSACINCRDLEMAKEVANKLFV--IEpnTGNYVLLANIYASHGRLEEAKEVRRWmkEKGCSLINN-----------------------------------------------------------------------------    
 754 UniRef50_B9FDV7        Putative uncharacterized pr...   346     1e-93  1  11.3%  61:511    65:526     ------------------------------------------------------------PAPGRRALPSLLRAIAFARSLGTAGAAHGLAFRVGAEVDGFVGTALVRAYAACGRVEDARRVFDGMprDLVAWGVMLDCYcrNYEEALLLFHSMKRSRVVPDQVIIATVLSTCAhrNLRFGKAIHSYMLVSDTLIDAQVSCALMNMYASCADMEMAEKLYNRVSEKEIVldVVSWSAMIAGYAESSKPMEALNLFHDMQRSGVKPDEITMLSVISACANVGALEKARCIHSFVENHSMCKILP------------IGNALIDMFSKCGSLTLALDVFNAMPQK------NVVTWTSIITASAMHGDGRSALTLFENMKSEGIQPNGVTFLGLLYACCHAGLVEEGRLLFKIMvqYRIEpeHYGCMVDLLGRAKLLGQAADLIQSM----HLRPNVVIWGSLLAACRMHGDLELGTFAAKKILE--LDPNHGGavLLSNIYAEYGNWNDVKE--------------------------------------------------------------------------------------------    
 755 UniRef50_B9HF30        Predicted protein n=3 Tax=r...   346     1e-93  1   9.2%  19:527    48:534     ------------------LFDELPHRDLYSLNSQLASYSRDGNFLATWDLFSRIH--SAFLDLDAYTFTPVLRACSALPDTKCGRQVHALMIKTGTDLGTITKTAVMDMYSKYGCLGESVKVFEEMerDVVTWNALVSSFLRHgkEALGVFRAMRRESVEITEFTLCSVLKACAFIkfRQGKQVHGLVIVMGRDL-VVLGTALIDFYSNVGYISEAMKVFSCLS---CRKDEVLRNSLIAGCVKHRRYEEAFLVMSTMR-----PNAVALTTALAACSDNSDLWIGMQI------------HCVALRFGFIANTQVCNVLLDMYAKCGRILKSRSIFDGICHK------TVVSWTSMIDAYGRHGHGDEALKLFKEMGQEgvLPNSLTLLAVLSACGHSGLVKEGQELFNSaeKYGLDpeHYSCVIDILGRAGQIEDAWCLFHDMVKKGI-GPTAAVWAALVNACCLNLDVSRGEFAAKHLLE--LEPndGIHVLVSKFYASIDRWDVVESLRNNMRKKGLTKVLGS----------------------------------------------------------------------------    
 756 UniRef50_UPI00015C91EC PREDICTED: hypothetical pro...   346     1e-93  1   9.8%  71:580     1:451     ----------------------------------------------------------------------MIQAYSKTPSSQESLYLFHQMLAHGRPTSADKYTFTFVFTACSRHPT-------------------LRGYGEN------VHGMVVKDGYESDIFVGNSLVNMYsfSRMVDAKRVFDEMPQR----DVITWTSVVKGYAMRGELVRARELFDMMPGR----NDVSWAVMVAGYVGHRFYNEALQCFNDMLCHdvKPNEAVLVSILSACAHLGALDQGKWI------------HVYIDKNRILLSSNISTALIDMYAKCGRIDCARRVFDGLHKR------DLLTWTSMISGLSMHGLGAECLWTFSEMLAEGFKPDDITLLGVLNGCSHSGLVEEGLSIFHDMilWGIVpeHYGCLIDLLGRAGRLESAFEAIKSMP----MEPDVVAWRALLSACRIHGDVDLGERIINHIAELCPGSHGGGYVLLSNLYASMGQWESVTKVRKAMSQRGSE----------GCPGCSWIEIDGVVHEFLAADKLHPRISEINKKLNEVLRRiiEGGYVP-----------------------    
 757 UniRef50_D1J1G9        Whole genome shotgun sequen...   346     1e-93  1  11.7% 140:599   104:541     -------------------------------------------------------------------------------------------------------------------------------------------LIRTPAKALHFFRWVEAKGFTHNEQSYFLMIEILGRSrlNAARNFVFSIEKKsvKLGDRFFNSLIRSYGWAGLFQESIKVFKTMKEIGVSPSVVTFNSLLSIVLKRGRTSMAKQLFDEMLdyGVTPDTYTFNILIRGFCMNSMVDEGFWFFKEM------------SRFKCDPDVVTYNTLVDGLCRAGKVKIAHNVVKGMVKKSPNLSPNVVTYTTLIRGYCMKQDMAEALSLLAEMVSRGLKPNKITYNTLIQGLCEAQKLDKIKEILEGMvdGGFIPDTcnTLIKAHCTMGKLEEAFSVFEKMSEL-RVQPDSATYSVLVRSLCQRGDFRRAEEFFDELAEkgCKPLVAAYNPMFEYLCSNGKTKKAERVFRQLMKRGTQDPPS--YKTLILGHCR---------------------EGTPEAGFDLLVLMLRRDFVPDA-ETYGLMIDGLLKKGDP----    
 758 UniRef50_A3AU91        Putative uncharacterized pr...   346     1e-93  1  14.0%  98:525     5:426     -------------------------------------------------------------------------------------------------PNAYTYNPLIHTLCERGQVRDALSVLDDMLCRGCAPDVVTYNILLEarQAMELIDLMRAEGCTPNNVTYNVLMdmCGEGDVDDALELLRNLPSHGCKPSTVNYNTVLKGLCSAERWGDADELVTEMLRENCPPNEATFNVVIYSLCRKGLLQQAIQLLEKMSKHGCTANIVTYNAIINGLCEQRNVDGAMGLLSKMKS------------YGCKPDIVTYNTLLKGLCSAARWVDAEELMDNMTQNGC--LPDNVTFNTLIGFLCQKGLMVDAIEVFKQMPDKGCTPNSITYSTIISGLAKATKLDQALELFNEMGHKGFNPdiYQLLAECLNDDDTIEEAIQTVRKLQDSGI-SPHTVLYNAILLGLCRNGKTEFAIDIMAYMVSSGCMPDDLTYVILIEGLAYEGYLNEARELLIKLCSRDVLVNS------------------------------------------------------------------------------    
 759 UniRef50_Q9LXE8        Pentatricopeptide repeat-co...   346     1e-93  1   9.0%  70:601    11:547     ---------------------------------------------------------------------LLLRHCAHRSFLRPGKELHAVLTTSGlaPRSYLSNALFQFYASSGEMVTAQKLFDEieKDNVDWTTLLSSFSRYGLLvmKLFVEMRRKRVEIDDVSVVCLFGVCAKldLGFAQQGHGVAVKMGVLTSVKVCNALMDMYGKCGLVSEVKRIFEELEEKSveRNAVAWTVMVAGYLGAGFTREVLELLAEMvrCGHGLNFVTLCSMLSACAQSGNLVVGRWVHVYALKKEMMMGEEASYD-----DVMVGTALVDMYAKCGNIDSSMNVFRLMRKR------NVVTWNALFSGLAMHGKGRMVIDMFPQMIRE-VKPDDLTFTAVLSACSHSGIVDEGWRCFHSLRFYGLEpdHYACMVDLLGRAGLIEEAEILMREMP----VPPNEVVLGSLLGSCSVHGKVEIAERIKRELIQ--MSPGNTEYQILmnMYVAEGRSDIADGLRGSLRKRGIRKIP----------GLSSIYVNDSVHRFSSGDRSHPRTKEIYLKLNEVIERIRSAGYVPDV--------SGLVShgDLEE--    
 760 UniRef50_Q7XSV8        Os04g0110500 protein n=3 Ta...   346     2e-93  1  11.2%  63:569     6:524     --------------------------------------------------------------LSSTALLRLIKSLSParAHLSASSIHCLLLKAGLLHVGAHltALLSAYAALGSPDHARRLFDEMPEQGLVptAMARAHvgQAAQAIAVFGDMVADGVFPDNVALAVALGACHGAGSWTArkIHALIVTSGIEPDVFVSTELIRVYGECGKLAVSRRVFDDMPSR----STITWNAMLHQYARHGKVDTAYELFLAMPRR----DVVSWNTVMAGYCVAGRCREALGLFRQMVseTGIWVHAYIERNRMNDDGYLDRCLIDMYCKCGSIDNALQVFEKAPRK-----RDLFSWTTVICGLAMHGRATDALRMFDMMQDNGICPDDVTLVGVLNACAHGGLVDEGLGYFYSMEAkrITpeHYGCMIDLLGRVGRLQEAYSMIRTMP----MDPNTVIWGAFLSACKVHGNMELGKIAAEELTRLDPD-DPWGRVMLSSMYAKAQDWIGLARERREMNSMQVKKTP----------GCSSIELKGEVHEFVAGGSQHPQYAEICSVLQDV----------------------------------    
 761 UniRef50_Q6AVR3        Pentatricopeptide, putative...   346     2e-93  1   9.9%   4:598    17:604     ---LALYCRCGDMRHARLLFDGMPRRDAVSWNTLIAGYSGLGSCRLALDAFRDARASGDGV--DRFTFAAALASCARVGDWRDGRVVHGLAVVSGLARTAFLTNSVIDMYAKCGMIDEVRLLFDRAEERgaSWNLLLSAYVRMgeVAVNVLVWMHRSGVKLDSFALGGILKACSegDSEDVRrlHSCVVKVGLDLDVFVGSAMVDMYAKNGGLEEAIKVFDCIPNQ----NVVVYNAMIAGFARLGnpeeAVTLFSNMLKKRIKPSKFTFKSVLEVCNLTNAVRCGRQI------------HAHVILSGLQGDEFIASVLINLYSKARCVNDSLRCFDM------TVKEDVFIWTSMITAFVQNEHFEKALYLFRELLYTRKGTDQFTISSVMSACAALSVPTTCKQIHCHAVKTGLdtVSGNSQIAMYRNIGDLKASKQTFEQ-----ISCLDTFSWSAMILSYAVHGYESKALELFEKMKNLGVMMNESASLAALIACSHQGLGDEGLRYYENTIPDDGFSLDVKLKACMVDLLGRVGKIADAEdiLWHALLRACRVHGDKERCTKIGEKLME--LEPFSASSYVMLYNLYMDAGK-----    
 762 UniRef50_D1HYF1        Whole genome shotgun sequen...   346     2e-93  1  11.9%  27:537   118:623     --------------------------DFPSFNFllSHFVEHQKNALISLRFFHWLSSQSGFSPDSSSCNVLFDALVEAGACNAAKSFLD---STNFNPKPASLEAYIRCLCKGGLVEEAISVFGQLKGIGvtWNSVLRgrAGRIDFVWELYGEMVESSVVADVHTVGYLVQAFceNRISDGHNLLRRVLEDGVVPRNAAFNKLISGFCKDKAYGRVSDLLHSMIARNRAPDIFTYQEVVNGLCKGGKGPEGFRVFKDLKDRGYAPDRVMYTTMIHGLCRMKWLGDARKLWFEMIQK------------GFLPNEYTYNAMIHGYFKIGNLEEAWKMYREMCDKGYGEK--TVSYNVMIKGLCSHGKIKEAHDLFEEMSHKGILRNHITYNALVRGFCKEGKIVEGANLLYELLDQGIQPssYAPLIDKLCQEGDMQNAKILWDDMQNRG-MEPAVCTHDFMITGFCKQGCAMEGMEWLTTMLRSKLRPQKKSFESLIQCLSQIDRLDDALLVLDSMLKIGFRL-SISICNSLVTKLC------------------------------------------------------------------    
 763 UniRef50_Q9SB36        Pentatricopeptide repeat-co...   345     2e-93  1   9.5%  68:522    95:526     -------------------------------------------------------------------FASLLETCYSLRAIDHGVRVHHLIPPYLLRNNLGISSKLVRLYASCGYAEVAHEVFDRMSKrpFAWNSLISGYagQYEDAMALYFQMAEDGVKPDRFTFPRVLKACGggSVQIGEAIHRDLVKEGFGYDVYVLNALVVMYAKCGDIVKARNVFDMIPHK----DYVSWNSMLTGYLHHGLLHEALDIFRLMVQNGIEPDKVAISSVLA---------------RVLSFKHGRQLHGWVIRRGMEWELSVANALIVLYSKRGQLGQACFIFDQMLER------DTVSWNAIISAHSKNSNG---LKYFEQMHRANAKPDGITFVSVLSLCANTGMVEDGERLFSLmkEYGIDpeHYACMVNLYGRAGMMEEAYSMiqEM----GLEAGPTVWGALLYACYLHGNTDIGEVAAQRLFE--LEPdeHNFELLIRIYSKAKRAEDVERVRQMMVDRGLE---------------------------------------------------------------------------------    
 764 UniRef50_Q7EXY6        Os08g0511000 protein n=3 Ta...   345     3e-93  1   9.9%  32:538    16:512     -------------------------------NTLIRAAADHRSL---LLAFRAM--LREGVAADHFTFPFALKALAQAHhtTTLGCLHAQMVKSGHGANVYAASALVHAYVSRADAASARAVFDAArrNVVTWTAMihAAAGEAREAVALFREAVARGQEVNGITVAQVMGACAQSrlESGRWVHATLRRWGVEpdVALATAVLHMYASCGGLDAAFEVFDKMPRR----NEVSWNAMAEVCSRHGRQDKVLEVFPGMHSVGMKPDKVTWLSILRACTSKGAAGLGQGVHAYMEKTIGHRH------------VAVCTSLMDMYSKIGNARSALQIFQCLKRK------DLMAWTSMIIGLAKHGHGKDAVQLFNQMQHGGVVPDHVAFVGVLTACSHAGMVDEARKYFDSmnHYGIRPthYGCMIDLFSRAGRLAEAEGMMQMMP----IQPSVTMWGSMMNGCKVHGRADIAERVGRQVAELNPQFGAI-YVIMSNIYAEIGRWHAVEHTRRLMWQTGLKKIVGSS-GTEVQMLCS-----------------------------------------------------------------    
 765 UniRef50_C5Y9W9        Putative uncharacterized pr...   345     3e-93  1  13.8%  97:538   174:603     ------------------------------------------------------------------------------------------------DPAPLVSAVAKAYADEGLLPDGCSLLLLLALRRGVrwSDLM-NRLPSVPEAYAFYLQLLDAGVPPEAKQFNMLmdMIRSGKLASARNMFDEMLRRGVPPTVVTFNTLMSGMCKASDLNSANALRGLMAKAGVAPDVYTYGAFMQGLCKAGRIQDAVEMFEEMRERGVNPNTVVFTTLIDAHCKEGNVAAGLELHREMATR------------GIKTDLVAYNALANGLCRVRDLKAANDIVEEMRNNG--LKPDKVTYTTLIDGFCKEGELDMAMEMKQEMSDEGVALDEVTYTTLISGLSKSGRSVDSERILCEMMEAGLEPdtYTMVIDAFCKNSDVKTGFKLLKEMQNKG-RKPGVVTYNVVMNGFCILGQMKNADMLLNAMLNIGVCPDDITYNILLDGHCKHGKVRDAEEL----KSAKGMVSDFGLYTSLINEFVK-----------------------------------------------------------------    
 766 UniRef50_D1HM64        Whole genome shotgun sequen...   345     3e-93  1  11.1%  34:527    13:471     ---------------------------------LIHALarANLHHHALVLAKIFRFAAVSPN----------------GCLHYADRLFSQIHQ----PNTFFYNTLIRGYSKSSSPSQSVQLFNQMRRNCVDPDGFTFTFLLKGRSRmeIHGAVLKLGFCFHLFVMNALIHLYAARGVPAahQVFNEM----VGADVVSWSGLVVAHVRAGELELARQVFYEMPER----DVVSWTVMVSGYAQAKRSREALELFREMRDVGVRPDEVAMVSVISACTSLGDLETGFEVHRYIDENGFGWM------------VSLCNALIDMYAKCGCMDLAWQVFNNMERK------SLITWNSMISACANHGNAEDAFRVFTLMLYSGIRPDGVTFLALLTAYTHKGWVDDGYGLFESMQryGVEaeHYGCMVDMLGRAGRLEEAYKLITSMS----MPCNDVVWGALLAACRIYGDVEMGERVLKKLIE--LKPDEGGYYILldIYVAAGRRAEAIELRRAMDVNGAKKTLGC----------------------------------------------------------------------------    
 767 UniRef50_B9H463        Predicted protein n=1 Tax=P...   345     3e-93  1  10.1%  65:589     6:497     ----------------------------------------------------------------AYHLSILLRHCANHSLIYQAKQINAQIVLSNYLSNVTLQtlLLAYSKSGFLLEARKVFDRMPERNMHSWNIMMSSYAQNYcdAICIFDEFLKMGFRPDHYTLPPLFKAVAggDCYLGFVLHGWVIRLGFEGYVVVGSSVLDFYLKGGKLVEAKRVFSNMLWRDCG----VWNLMISGFGRAGFYVEALSLVRNMVEEEEKLDALVVPSILNACGGEGDLMKGKEIHGRVVKSTL-----------FNVDVVISNSLIDMYARCGCLNDSEKVFRNMRS------LNVVTWTTMISCYGVHGRAEESLEVFKKMKGFGLKPNPVTLTAVLASCSHSGLIDEGRRIFYSMQsyGFEpeHYACMVDLLGRFGYLEEALGLVQRMK----LEATASVWGALLGGCVMHKNVNIGEIAAHCLFE--LEpnPGNYIALCYIYKSHGISDGITITRAKMRELGLAKTP----------GCSWITIAGTVHKFYQGCHSHPLTKVTCE----ILDRMIKVLMLPDIWHGMTSL--------------    
 768 UniRef50_Q2R1Y4        Os11g0587100 protein n=6 Ta...   345     3e-93  1   8.9%   5:556    57:596     ----SRYFRLGDAGAARNVFDAAATAAAplYNAMLRGYLAGGLPRMAVGIFGEMAAAACL--PDRHTYHLAVAACARASEFEVGWRIGAEAAAKGFASDLLVATALIGMYAEAGDMGAARKVFDGmqRDAVAWNAVIAGYArhLREVVELFMRMRSVDVvpTEATLVTLVSGYAGFGSWEGrmMHTIVIKLGFQLNLFVSNALLDLYVEFGCLREAVMLFRQMAVK----DSVTWSAMIGGLVRNGRPDSALKLFRWMVTNsvLVTRSILLNVIMACAELGEWREGKWVEENYVCCNGFE---------FKRDPSVVTALIYMYAKCGMLDSSVSLLYGVAE----VRDDVFAWNAMIKGCGELGLVEKAVGFVVEMQKIGVDPDAITYLEILPMISSIPSLKTGMEAHAQIVRRGfeRAIANSLVSMYGRCGSLRHSIEVFSRIVVK-----DVISWTSMMQVYAWNGHVNEVVKLYEVMKKTETEPNHYTFLAVLSACKNTGLVEEGMELIKYMQEKCGLKPEIEHISCVVDMLCRAGRLTDAYHLIKYNNSEH-----------------------------------------------    
 769 UniRef50_O04491        Putative pentatricopeptide ...   345     4e-93  1  11.4%  44:541    97:602     -------------------------------------------PQRSIFAFFKFISSQPGFRFTVETYFVLARFLAVHEMFTEAQSLIELVvrKGKNSASSVFISLVEMrmCGFLVDALMITYTDL---GFIPDAiiRGCGNLLDRMMKlyMEILDAGFPLNVYVFNILMNKFCkgNISDAQKVFDEITKRSLQPTVVSFNTLINGYCKVGNLDEGFRLKHQMEKSRTRPDVFTYSALINALCKENKMDGAHGLFDEMCKRGLIPNDVIFTTLIHGHSRNGEIDLMKESYQKMLSK------------GLQPDIVLYNTLVNGFCKNGDLVAARNIVDGMIRRG--LRPDKITYTTLIDGFCRGGDVETALEIRKEMDQNGIELDRVGFSALVCGMCKEGRVIDAERALREMLRAGIKPdtYTMMMDAFCKKGDAQTGFKLLKEMQSDGHV-PSVVTYNVLLNGLCKLGQMKNADMLLDAMLNIGVVPDDITYNTLLEG--HHRHANSSKRYIQK-PEIGIVADLAS-YKSIVNELDRASK--------------------------------------------------------------    
 770 UniRef50_Q6K892        Os02g0290000 protein n=4 Ta...   345     4e-93  1   9.7%   7:574   348:915     ------YSKVCEMRAVESIFRSSKMGDIVTWNTVIAGYVMNRYPSRALKLFqlFAGM----APDSVSLISLLTACAQVGNLRVGIRVHGYIFRHplQETSLMNALVSFYSQCDRFDAAFRSFITIQNKDSVSWNAILSACahHIEQFFRLLGEMWHDVTQWDSVTILNIIRMSTFCGIKMVQESHGYSLRVGYTGDSSVANAILDAYAKCGYLHDAETLFRSLAGR----NIVTGNTMISCYLKNNCVEDAEMTFNHMAEK----DLTTWNLMSRLYAQNDLCDQAFCLFHQLQSEgvKQCHGYMLRASLE-DIHLEGALLDAYSKCGNIANAYNLFQ------VSLHKDLVIFTAMIGAYAMHGMAEKAVELFSKMLTLDIKPDHVVLTALLSACSHAGLVDAGIKIFKSIReyGVEpeHCACMVDLLARSGRLQDAYSFALDMPPHVV---NANAWGSLLGACKVHGEVKIGQLAADRLFSMEAE-DIGNYVIMSNIFAADDKWESVEHVRKLMKSKDMKKP----------AGCSWIEVEKTRHLFVASDVQHQDRFSIYDVLGSLYQQIR-----------------------------    
 771 UniRef50_D1I593        Whole genome shotgun sequen...   344     5e-93  1   8.9%   6:546   155:684     -----LYMQMGDCDTALRLFSNLPRRNLAICNLVLRGFCELSRSEWLFTMYSAMKE--ECVKPNGLTFCYLIRGCGNGRFCDEGKQLHCHAIKVGWvnLFAANALVDFYCVCGNLIDAKKSFEAIRDEDVISWNSIVSVFAASgsDALELFARMQSWGKKPSIRSFVGFLNAagCDNIIFGKQIQSHVVKLGFApsVHVESALINMYGKCGDIKSSVSVFESVPERTLE----ICNSLMSSLLHCGIVEDVLELFGLMVDEGIGVDEVTLSTSLKA-----SLVSAFGSLACCR-----LLHCCAIKSGFEFDTAVSCSLIDAYSKYGHVELSHQIFEQLHS------PNAICFTSIINGYARNGMGREGLEMLEIMAKQGLKPDRVTFLCALTGCSHSGLVEEGRSVFNSMKshGIHPdhYSCMVDLLGRAGLLDEAEGLLKHAPAKS----DSVMWSSLLQSCRIHGDTIVGRRAAKVLME--LEpdPATYLQASSFYSEIGELEISMQIRerMkeMGHSLivNTHGVAENRVYDDIAMIEELE---------------------------------------------------------    
 772 UniRef50_D1IT63        Whole genome shotgun sequen...   344     7e-93  1   9.2%   6:579    51:636     -----KLSHLKAIDQASLLFSTIPNPDLFLYNVLIRAFSLNNSPSSAVSLYTHLRK-STPLEPDNFTYAFVISGASSLGLGLLLH---AHSIVAGFGSDLFVGSAIVACYFKFSRVAAARKVFDGmeRDTVLWNTMVSGLVKnfDEAILIFGDMVKGGIGFDSTTVAAVLPGVAEldLALGMGIQCLAMKVGFHSHAYVITGLACLYSKCGEIETARLLFGQIGQ----PDLVSYNAMISGYTCNNETESSVRLFKELLVSGEKVNSSSIVGLIPVFFPFGHLHL------------TRCIHGFCTKSGVVSNSSVSTALTTVYSRLNEIESARLLFDESSEK------SLASWNAMISGYAQNGLTEKAISLFQEMQKCEVRPngVTFLSVLYACSHAGLVREGDEIFRSMvdHGFepEHYACMVDLLGRAGNLDKALDFIRKMP----VEPGPPVWGALLGACMIHKDANLARLASDKLFELDPQ-NVGYYVLLSNIYSAGQNYPEAASVRGVVKRRKLAKTP----------GCTLIEVANTLHIFTSGDQSHPQATAIYAMLEKLTGKMREAGFQ------------------------    
 773 UniRef50_B9RTG3        Pentatricopeptide repeat-co...   344     7e-93  1  10.9%  11:597   185:831     ----------ERAMEIFEWFKSryELNVIHYNIMIRILGKAKQWRYLECLCNEMSFKRISPINS--TYGTLIDVYSKGGLREKALDWLEKMNKQGMEPDEVTMGIVVQMYKKAGEFQKAEEFFKKWSLrsHTYNTMiyGKAGQIKEASDIFAEMLRKRILPTTVTFNTMIHICgqGQLEEVALLMQKMEELRCPPDTRTYNILIFIHAKHNDINMAASYFKRMKKDQLQPDLVSYRTLLYAFSIRHMVNDAENLVSEMDEKGIEIDEYTQSALTRMYIEAGMLEKSWLWFWrmSSECYSANIDAYGERGhkLTVLEFNVMIKAYGFGKNYEKACDLFDSMESHG--VVPDKCSYSSLVQILASADLPDKAKHYLKKMQEAGLVSDCVQYCAVISSFVKLGKLEMAEEVYKEMVGFDVKPdvYGVLINAFADSGCVKEAISYIDAMKGAGLPG-NTVIYNSLIKLYTKVGYLREAQETYKLLQSSDVGPETYSSNCMIDLYSEQSMVKPAEEIFESMKRKGDA--NEFTYAMMLCMYKRLGWFEQAIQIAKQMRegLYALDGRFKEAVGTFKEMVGAGIQPDD-CTFKSLGIVLVKCG------    
 774 UniRef50_P0C8Q7        Pentatricopeptide repeat-co...   343     8e-93  1  10.1%  40:573    17:518     ---------------------------------------QCKSMSELYKIHTLLITLGLSEEEPFVSQTLSFSALSSSGDVDYAYKFLSKLS----DPPNYGWNFVIRGFSNSRNPEKSISVYIQMLRFGLLPDHMTYPFLMKSSSRLslHCSVVKSGLEWDLFICNTLIHMYGsrDQASARKLFDEMPHK----NLVTWNSILDAYAKSGDVVSARLVFDEMSER----DVVTWSSMIDGYVKRGEYNKALEIFDQMMRMGskANEVTMVSVICACAHLGALNRGKTV------------HRYILDVHLPLTVILQTSLIDMYAKCGSIGDAWSVFYRASVKET----DALMWNAIIGGLASHGFIRESLQLFHKMRESKIDPDEITFLCLLAACSHGGLVKEAWHFFKSLKESGAEpeHYACMVDVLSRAGLVKDAHDFISEMPIK----PTGSMLGALLNGCINHGNLELAETVGKKLIE--LQpnDGRYVGLANVYAINKQFRAARSMREAMEKKGVKKIAGHSI----------LDLDGTRHRFIAHDKTHFHSDKIYAVLQLTGAWM------------------------------    
 775 UniRef50_Q0DF75        Os06g0111300 protein n=1 Ta...   343     8e-93  1  10.6%  17:602   264:875     ----------------EQMLEANVAPNQFTYTVVIGSYAKEGMLEEAMDAFGEMKRRRFVPE--EATYSLLISLCAKHGKGEEALGLYDEMKVKSIVPSNYTCASVLTLYYKNEDYSKALSLFSEMEQNKIVPDEVIYGILVryEDAQRMFEEIDKAGLLSDEQTYVAMAQVHMNvnYDRALQVLDAMRARNVKPSQFSYSALLRCHVAKEDVDAAEDTFRALSNYG-PPDVFCCNDLLRLYMRLGHLDKARALILKMRKEALQFDEDLCVTVLEVCCKTsnLTEVIqtLSMMLKseAKFLYEHLTELGAKPDDTAIATLIVQYGQAQQLEQAQKLFETAS---TSFPVGGSVYNAMVDALCRCGKTEEAYRLFMELIDQGHNGDAVTISILVTHLTKQGKLYSAVSIYDRMISSGIPRstFNIMISVYGQGGKLEKAVEMFSAAQELGLPI-DEKTYTNMLSFYGKAGKHHEASLLFSRMKEDGIRPGKISFNTMINAYATSGLHNEAEIIFQEMQKNNHVPDSHTYL----------------------ALIRAYTEGKCYSKAEEAIQMMLRSNMTP-SCTHFNHLISAFLKEGQIDEA-    
 776 UniRef50_Q8LNU8        Os10g0484300 protein n=7 Ta...   343     8e-93  1  11.4% 135:581   126:562     --------------------------------------------------------------------------------------------------------------------------------------VLALARASQPLRAYDAFLLAGEShhRPSTSSVNALLAGlgAKRVDLAEKAFRSALRRRVSPDIYTFNTVISGLCRIGQLRKAGDVAKDIKAWGLAPSVATYNSLIDGYCkaGNMYHVDMLLKEMVEAGISPTAVTFGVLINGYCKNSNTAAAVRVFEEMKQQ------------GIAASVVTYNSLISGLCSEGKVEEGVKLMEEMEDLG--LSPNEITFGCVLKGFCKKGMMADANDWIDGMTERNVEPDVVIYNILIDVYRRLGKMEDAMAVKEAMAKKGISPntYNCLITGFSRSGDWRSASGLLDEMKEKGI-EADVVTYNVLIGALCCKGEVRKAVKLLDEMSEVGLEPNHLTYNTIIQGFCDKGNIKSAYEIRTRMEKCRKRANVVT-YNVFIKYFCQIGKMDEANDLLNEMLDKCLVPNGITYETIK--EGMMEKGYTPD----------------------    
 777 UniRef50_Q9LMH5        Putative pentatricopeptide ...   343     9e-93  1  10.4%  29:528   292:869     ----------------------------IAYRKVVRGLCYEMRIEDAESV--VLDMEKHGIDPDVYVYSAIIEGHRKNMNIPKAVDVFNKMLKKRKRINCVIVSSILQCYCQMGNFSEAYDLFKEFRETNISLDRVCygKLGKVEEAIELFREMTGKGIAPDVINYTTLIGGCcqGKCSDAFDLMIEMDGTGKTPDIVIYNVLAGGLATNGLAQEAFETLKMMENRGVKPTYVTHNMVIEGLIDAGELDKAEAFYESLEHKSRENDA----SMVKGFCAAGCLDHAFERFIRLeiSKAQDLLDRMWKLGVEPEKSMYGKLIGAWCRVNNVRKAREFFEILVTK--KIVPDLFTYTIMINTYCRLNEPKQAYALFEDMKRRDVKPDVVTYSVLLNsyCHLNDLKKVYALFKDMKRREIVPDvyTVLIDWQCKIGDLGEAKRIFDQMIESG-VDPDAAPYTALIACCCKMGYLKEAKMIFDRMIESGVKPDVVPYTALIAGCCRNGFVLKAVKLVKEMLEKGIKPTKASL---------------------------------------------------------------------------    
 778 UniRef50_UPI000198605B PREDICTED: hypothetical pro...   343     1e-92  1  10.0%  72:573    15:517     -----------------------------------------------------------------------LRLLERCSSMHELKQIYAHVITLGLARFTYITSRVLSFCAVsgDLSYAETVFNKIapTIFDFNSMITGFLNrfEKGLSVYAQMRTLGVEPNARTLTVMIKAF--LRLSSINQVHCQIMKFGHGSDLYAISLLITRYSTCGAMELARQVFNESSNR----NIVCWTGLISGYCSNGFVDEARCVFDAMPER----NDVSYSAMASGYVRNDRFNEAIELFRELKseEGKWIHSYIDENGLDYDLKMGTALIDFYAKCGCIKTAEEIFNKMPLK------DVTAWSSMILGLAINGNNEMGLELFHEMETRGPRPNAITFIGVFTACNHKTLVNEALRLLGRMSkyGMFpeHYGCMVDLLSRTGQVKEAEILINTMP----MEPDGAIWGSLLNGCLMHGHMKLGERVGKLLIQ--LEPQHSgyVLLANMYATMGSWEGVMRLRKMMKERGVLTVPA----------WSFIEINGVVHKFLVDDKCHPQWRYIYSVLNQLGREL------------------------------    
 779 UniRef50_B8AB40        Putative uncharacterized pr...   343     1e-92  1  10.8%   5:580   317:863     ----ALVSAFARHGMLDEAKDIYleRNAVTLNGLIAGLVKQ-QHGEA--AAEIFMGARDSAAVNVDTYVVLLSAIAEFStlRKGREVHAHVLRAgiYRKIAVSNGLVNMYAKCGAIGKACRVFQLMerDRISWNTIITALDQNgeAAMMNYCLMRQNSIGPSNFAAISGLSSCAGLGLLAaqQLHCDAVKWGLYLDTSISNALVKMYGECGRMSECWEIFNSMSAH----DVVSWNSIMGVMASSQAptESVQVFSNMMKSGLVPNKVTFVNFLAALTPLSVLELGKQIHSVML------------KHGVTEDNAVDNALMSCYAKSGDVDSCERLFSRMSGR-----RDAISWNSMISGYIYNGHLQEAMDCVWLMMLSEQMMDHCTFSIVLNACASVAALERGMEMHAFGLRSHLESDVvsALVDMYSKCGRIDYASKVFHSMSQK-----NEFSWNSMISGYARHGLGRKALEIFEEMQESGESPDHVTFF-----HAAIGRWEDTAIARAAMKGAAVKKEAGRSWVTLTDG----------VHTFIAGDRSHPNTKEIYEKLNFLIQKIRNAGYVP-----------------------    
 780 UniRef50_B9SL37        Pentatricopeptide repeat-co...   343     1e-92  1   8.7%  67:516     8:443     ------------------------------------------------------------------TSPSLINALnsFNSLKQLKHVHAALLRLGFDQGSYLWSMIIRSSFDFNDTTYTRLIFNQiePDVVLYNTMIRGLVSSDceSIEFYYSMRKKGFLPNNFTFPFVIKACARldFQLGIMMHTLVVKLGFDCNLFVNTSLVSLYAKCGSIDDALKMLDLIPES----NIVSWTAIISGYIGVGKYREAIDLFRRMLETGLRPDSFTIVQVLSACIQIGDLANGEWI------------DRYITENVMARNVFVTTSLVDLYAKHGNMEKARCLFDGMAER------DIVSWSAMIQGYASNGLPKDALDLFFKMLNAGLKPDHYAMVGFLCACARLGALELGDWGSNLMDRTEFFTNHvtALIDMYAKCGSMAKAWEVF-----KGIKGKDRAVWNAAISGLAMNGHEKAAFGLFGQMERFGIQPDGNTFVGLLCGCTHAGLVDEGRKYFHSM---------------------------------------------------------------------------------------    
 781 UniRef50_B9I4V1        Predicted protein n=1 Tax=P...   343     1e-92  1  11.1% 109:587     1:457     ------------------------------------------------------------------------------------------------------------MCKLGDID---------MRPDVVSLVNVlcASMGAWLHGKAVHGIAVRSGSFEDLFVGNALVDMYAKCGmdEASKVFDRIKEK----DVVSWNAMVNGYSQIGRFEDALGLFEKMREENIELNVVSWSAVIAAFAQRGLGCETLDVFREmkQDgvKPNCFTISCALIACARLAALRLGRQIHAYILRNHFDSAF-----------LYVANCLIDMYAKSGDIDVARFVFDNLKQK------NFVSWTSLMTGYGMHGRGKEALEVFDEMRRVGLQPDGVTLLVVLYACSHSGMIDQGIEFFNSmkEFGVIpeHYACMVDLLGRAGRLNEAMELIEGMQ----MEPSSIVWVALLSGCRIHANVELGEHAAKQLLELNSE-NDGSYTLLSNIYANARRWKDVARVRSLMKNSGIRKRP----------GCSWVQGKKGTTTFYVADKTHPQSKQIYEILRSLTQRIKVLGYVPETSFALH----------------    
 782 UniRef50_B9I431        Predicted protein n=2 Tax=M...   343     2e-92  1  10.9%  68:572    16:532     -------------------------------------------------------------------HSIIIKLIKQCKTISQLHQLHAYTITNTPLSFHSSPSLLTKFlsSTSLLHYAKSIFNSIqpSTFCYNTIihTLLSFPIPALHFFTQMRRLSVPLDSHSFPFTLKACAqgGVFSARCLHCQVLKFGFLSDLYVMNSLIHGYMVSDMSNDAYKVFDESPQR----DVVSYNVLIDGFVKAGDVVKARELFDLMPVR----DSVSWNTIIAGCAKGDYCEEAIELFDFMMdeKGKKIHDYIERNAMKVDTFLSTGLVDFYAKCGCVDIALKIFDSSSDK------NLFTWNAMLVGLAMHGYGELLLEYFSRMIEAGVKPDGISILGVLVGCSHSGLVDEARKLFDEMEsyGvePKHYGCMADLLGRAGLIKKVMEMIKDMPRGG----DMSVWSGLLGGCRIHGDVEIAEKAAKHLME--LKPDDgvYSILANVYANAERWEDVMNIRRSLSSNRVVTKIAGF---------SLIQLDGVAHEFIAGDSLHSESDKIYLVLNGIREH-------------------------------    
 783 UniRef50_Q9M2Y4        Pentatricopeptide repeat-co...   343     2e-92  1  11.8%   4:522    99:733     ---LSLYERLGNLASLKKKFDEIDEPDVYSWTTLLSA---SFKLGDIEYAFEVFDKM---PERDDVawNAMITGccDYGSLDFGKQVHSLVIKAGFFIASSVVNALITMYFNCQVVVDACLVFEETdrDQVTFNVVIDGLAgkRDESLLVFRKMLEASLRPTDLTFVSVMGSCSCAAM--GHQVHGLAIKTGYEKYTLVSNATMTMYSSFEDFGAAHKVFESLEEK----DLVTWNTMISSYNQAKLGKSAMSVYKRMHIIGVKPDEFTfnALISAYSKNGQIEKADLLFERSLRKNLIsyHNGFPFEGLepDAYtgNALINMYSQCGTIQNSLEVFNQMSEK------DVVSWNSLISAYSRHGEGENAVNTYKTMQDEgvIPDAATFSAVLSACSHAGLVEEGLEIFNSMVehGvnVDHFSCLVDLLGRAGHLDEAESLVK--ISEKTIGSRVDVWWALFSACAAHGDLKLGKMVAKLLMEKE-KDDPSVYVQLSNIYAGAGMWKEAEETRraMKQRGCS---------------------------------------------------------------------------------    
 784 UniRef50_Q5JK99        Os01g0545900 protein n=5 Ta...   342     2e-92  1   8.5%  11:516    61:550     ----------GDLATAARLLDEAPPPSPLLHNALIRAHARRLDLRAALALFARMRRSRSSaaPDAHTFACVLRACADCSRPDVARVVNGIVVCAGMSSHPVVGSALVSAYAKLGPVGDARRVFDGlePDLVLWNCMmyGYRGMWNDGLDLFSAMRKAGERPDGYSMVSLVSSfnREALAFGQAIHGMCIKGGYDSGHHVRSALVSMYFRCGCMDSGHSLFGNLLDA----DLVTWSSLITGLLHICKYEESFGLFRQMCNSGRRPDSILVASLLSACASMVNI------------SYSKEIHCYAFRVGAHTDIRVSSSLMDAYSKCGFADLAYGVFFQMPDK------NLVMYNTVISNLGSHGFAMKAIEVLDEMVSDKLKPDSVTFSALLAACCHAGLLEEGWKLFRRMRDEfvKTEHYVYIVRLLATFGQLKEAYGLIQTMPA----PADSGVWGALLWGCCVHRDSSLGRIVADKLFE--LYPDKASyvMLSNLYASQEMWWDAEEVREEL---------------------------------------------------------------------------------------    
 785 UniRef50_Q2QM66        Pentatricopeptide, putative...   342     2e-92  1  14.1% 117:597   124:597     --------------------------------------------------------------------------------------------------------------------RHAPEVFDEML-------SLLSNTRLH-DPARRVVRLMIRRGVRRGTEQFAQLMLSYSrgKLRSAMRVLHLMQKDGCAPDISICNMAVNVLVVAGRVDKALEFAERMRRVGVEPDVYTYNCLIKGLCGARRVVDAMEMIGVMLQNGCPPDKISYYTVMSFLCKEKRVEEVRGLLQRMRN-----------DAGLFPDQVTYNVLIHGLAKHGHADEALEFLRE--SEGKRFRVDEVGYSAIVHSFCLNGRMAEAKEIVGEMISKGCQPDVVTYSTVVDGFCRIGELDQARKMMKHMYKNDCKPntHTALLNGLCKVGKTSEAWELLNK-SEEEWWTPSDITYSVVMHGFRREGKLKESCDVVVQMLQKGFFPTTVEINLLIHALCKDRKPAEAKDFMEQCQSKGCTINVV-NFTTVIHGFSRQGDLESALSLMDDMYLshPdkKGRLKEATGLVEKMLNRGVLPTPV-TYRTVIHRYCEKG------    
 786 UniRef50_Q9LVF9        Pentatricopeptide repeat-co...   342     2e-92  1  10.7%  69:525    49:515     --------------------------------------------------------------------PLILRAccVVPRVVLGKLLHSESIKFGVCSDVMVGSSLISMYGKCGCVVSARKVFDEmeRNVATWNAMIGGYMSNGDAVLalFEEI---SVCRNTVTWIEMIKGYGKreIEKARELFERMP---FElnVKAWSVMLGVYVNNRKMEDARKFFEDIPEKnfARDLVIWNTLIAGYAQNGYSDDAIDAFFNMQGEGYEPDAVTVSSILSACAQSGRLDVGREV------------HSLINHRGIELNQFVSNALIDMYAKCGDLENATSVFESISVR------SVACCNSMISCLAIHGKGKEALEMFSTMESLDLKPDEITFIAVLTACVHGGFLMEGLKIFSEMKTQDVKPnhFGCLIHLLGRSGKLKEAYRLVKEM----HVKPNDTVLGALLGACKVHMDTEMAEQVMKIIETAGSITNSYslASISNLYAHTERWQTAEALRVEMEKRGLEKSP------------------------------------------------------------------------------    
 787 UniRef50_Q4V389        Pentatricopeptide repeat-co...   342     2e-92  1  11.1%  29:552    48:587     ----------------------------VLFNSF-RHCISHGQLYEAFRTFSLLRYQSGSHEFVLYSSASLLSTCVGFNEFVPGQQLHAHCISSGLEFDSVLVPKLVTFYSAFNLLDEAQTITENSelHPLPWNVLIGSYIRnfQESVSVYKRMMSKGIRADEFTYPSVIKACAAldFAYGRVVHGSIEVSSHRCNLYVCNALISMYKRFGKVDVARRLFDRMSER----DAVSWNAIINCYTSEEKLGEAFKLLDRMYLSGVEASIVTWNTIAGGCLEAGNYIGALNCVVGMRNCNVRIGSVAMI-NGLKACSHIgnSLITMYSRCSDLRHAFIVFQQVEAN------SLSTWNSIISGFAYNERSEETSFLLKEMLLSGFHPNHITLASILPLFARVGNLQHGKEFHCYILRRQSYKDcwNSLVDMYAKSGEIIAAKRVFDSMRKR-----DKVTYTSLIDGYGRLGKGEVALAWFKDMDRSGIKPDHVTMVAVLSACSHSNLVREGHWLFTKMEHVFGIRLRLEHYSCMVDLYCRAGYLDKARDIFHTI---------------------------------------------------    
 788 UniRef50_A5AH22        Putative uncharacterized pr...   342     2e-92  1  10.3%   7:541   120:649     ------YGKLGCIYYAQHLFEEMPRRDVVSWNALICGYSRNGYDYDALEVFVQMLREGFPPCQRTL-----------------GRLFHGFGIKSGLDLDCRVKNALTSMYAKCADLQAAEVLFEefEKTEVSWNTMIGAYGQNgdEAMLVFKQMQKERVEVNYVTIISLLSANAHLD-----STHCYVIKTGFATDASVITSLVCSYAGCGNIESAGLLYNLMPQR----NLVSLTAMISGYAEKGNMGLVVECFTQMLQLKMKPDAVAMVSILHGFTDPTFIGSGLGI------------HAYGLKTGLCADCLVVNGLISMYSKFGDIETVFSLFSEMGEKQ------LISWNSVISACIQVGRTSDAMELFCQMRMYGHSPDAITIASLLAGCSEVGFLQFGERLHNYVLRNNLDMehNACMVDLLSRAGFLEEAVIFVKNME----VEPDSAXWGALLTSCCIHQELKLGECLAKRLLLLDYN-SGGLYVLMSNLYASKGRWDDVARVRKMMKdsGISLIEVSSLEKMNNSLCLSDL--------------------------------------------------------------    
 789 UniRef50_Q8VYH0        Pentatricopeptide repeat-co...   341     3e-92  1   9.7%  67:579    31:520     ------------------------------------------------------------------TKAFLVHKLSESTNAAFTNLLHTLTLKLGFASDTFTVNHLVISYVKLKEINTARKLFDEmePNVVSWTSVISGymGKPQNALSMFQKMHEDrvPPNEYTFASVFKAcaLAESRIGKNIHARLEISGLRRNIVVSSSLVDMYGKCNDVETARRVFDSMI--GYGRNVVSWTSMITAYAQNARGHEAIELFRSFNAasDRANQFMLASVISACSSLGRL------------QWGKVAHGLVTRGGYESNTVVATSLLDMYAKCGSLSCAEKIFLRIR------CHSVISYTSMIMAKAKHGLGEAAVKLFDEMVAGRINPNYVTLLGVLHACSHSGLVNEGLEYLSLmeKYGVVPdhYTCVVDMLGRFGRVDEAYELAKTI--EVGAEQGALLWGALLSAGRLHGRVEIVSEASKRLIQSNQQVTS-AYIALSNAYAVSGGWEDSESLRLEMKRSGNVKERA----------CSWIENKDSVYVFHAGDLSCDESGEIERFLKDLEKRMKERGHR------------------------    
 790 UniRef50_UPI0001984B81 PREDICTED: hypothetical pro...   341     3e-92  1  13.5%  70:596   141:654     ---------------------------------------------------------------------STIRSCCVAAHLLAAEELRHVAQ-----DVIWWVIARIGVLRSGDLVEFMWRGHHVYESDFSVLDSLMRAFVnmGFQALEILGRMREVGVRPS---------ASGEGNVVEARKLFDGLKEMGFSPNAIMYNTLMNGYVKMREIDQANMLYEEMRKKGIAPDGITFNIlvSGLCWAGRLDEAMEFLMDMLEKGLSPSIIAFNSVIAAYSQAGLEDKAFEAYKLMV------------HFGLTPSPSTCSSLLMGLSINGRLQEATELIGQMIEKGLSV--NNMAFTVLLDKFFKRGDVVGAQSLWGEMERRGIFPDVVAFSAFIDGLSKQGLVEEAYNVFLEMLRKGLIPnaYNSLICGFCKCGKLNEALKLEKVMRHRGLL-PDIFTTNMIIGGLCKQGRMRSAINVFMDMHQTGLSPDIITYNTLINGYCKAFDMVNADNLVNRMYASGSNPDLTT----------YNIRIHGFC------------SSRRMNRAVLMLDELVSAGIVPNTV-TYNSMLNGVCSD-------    
 791 UniRef50_Q8H473        Os07g0671200 protein n=7 Ta...   341     4e-92  1  11.5% 141:593   120:538     --------------------------------------------------------------------------------------------------------------------------------------------AGHLDTAADLLKQAATS--CPDSVtaSVVVDGYCkgRVAHARQLLDEMPRHGVKVNALCYNSLLDAYTREKDDDRVAEMLKVMENEGIEPTVGTYTILVDGLSAARDITKVEAVFEEMKSKNLSGDVYFYSSVINAYCRAGNVRRASEVFDECVG------------NGIEPNEHTYGALINGFCKIGQMEAAEMLVTDMQVRGVGI--NQIVFNTMIDGYCRKNMVDKALEIKMIMEKMGIELDVYTYNTLACGLRRANRMDEAKNLLRIMIEKGVRPNhyTTLISIHCNEGDMVEARRLFREMAGNG-AEPSLVTYNVMMDGYIKKGSIREAERFKNEMEKKGLVPDIYSYAALVHGHCVNGKVDVALRLFEEMKQRGSKPNL----------------------VAYTALISGLAKEGRSEEAFQLYDNMLGDGLTPDD-ALYSALVGSL----------    
 792 UniRef50_B9F7W0        Putative uncharacterized pr...   341     4e-92  1  10.9%  96:580   112:565     -----------------------------------------------------------------------------------------------RAIFPYTNAILRASLEKGSPEKSLTDYNSMLRFTAFCPDYRTYVLLLKACaiMEIHSHLVKLGLLSNQRIVTHLFKLYihGRVMEACKLFWLMLEWSAEP--FYGNLMLMGFLKCGQIDKAYQIFKRMPVK----DLVSWNSMIAGTARNSCLKDAMNIFSKLVNSGLVPDGFSFSSVLSACAQAGA------------RCYGMWVHQLMAELGLEMNHILSSSLVDMYAKCGRIDVAIGIFMTIKRNHVSV------WNTMIGGLAAHGLGSDAVMFFREMESEGLVPDGVTFVALLTACSHSGMVEEARQYFEAMtkYSITpeHYGAMVDTLSRAGLLDEAYDLVKTM----NVKPDAVIWRALLSACCRYRQTKLGEITVKEIAFQG----SGDYTLLSNIYSSANRWEDSEEVWKERKKKGIRKSK----------GLSWVELGRSTHEFKAGDRSHPDTDGIYKVLHRLSNKAKSEGYIP-----------------------    
 793 UniRef50_B9GMA8        Predicted protein n=2 Tax=r...   341     4e-92  1  10.5%   5:578    65:591     ----KLVGMYASCGDVKsvFKRIRNPNVFALNWMVLASAFEGYYKEAIGYFCSMKD--SVFIYNKYTFSIVLKAFVGLLDLNKGKEVHSMVKQLGFESDVCVANALVDMYSKCGCIGY----------------------------ARTVFDRMAKR----DIVSWTSMISGYCngKIEEALVLFERMKLEGLEPNDFTWNALISGYARRGDSDGAFSLLSKMTREGLVPDLVTWNAMIAGFVQGERAGDAFKLFQDMLVLGVKPNLVTVAGLLPACGMVSSIQRGRAIHGLVYRLEFDI-----------SNAFIASALIDMYSECGSFKEARTVFEKIHNK------NVASWNAMIGCYGKHGMVNTSIQLFERMHGEGIQANDVTLLCVLSACSHSGYVEKGLEIFWSMKERYmkKEHYACVVDMLSRSGRLVDAYELVKEMP----IEVTKSIAGAFFNGCMIHGRRDLAEKMIDDVTRGDLK-KPGSFAMLSAIYATSGERKEVRNTMKKIVKERkkEPACSQVEEKDEFVGVEIEKENNEVSIKAAFQLIQMVVRGAEEADEDSKEVHRDGV-------------------------    
 794 UniRef50_Q9FX24        Pentatricopeptide repeat-co...   341     6e-92  1  10.4%  77:603    12:505     ----------------------------------------------------------------------------KCVSFSQIKQLQSHFLTAGHFQSSFLRSRLLERCAIsgDLSFAVQIFriPKPLTNDWNAIIRGFAghPSLAFSWYRSMLQQscRVDALTCSFTLKACARALCSSAmqLHCQINRRGLSADSLLCTTLLDAYSKNGDLISAYKLFDEMPVR----DVASWNALIAGLVSGNRASEAMELYKRMETEGIRRSEVTVVAALGACSHLGDVKEGENIF-----------HGYSND-----NVIVSNAAIDMYSKCGFVDKAYQVFEQ-----FTGKKSVVTWNTMITGFAVHGEAHRALEIFDKLEDNGIKPDDVSYLAALTACRHAGLVEYGLSVFNNMACKGVERnhYGCVVDLLSRAGRLREAHDIICSMS----MIPDPVLWQSLLGASEIYSDVEMAEIASREIKEMGVN-NDGDFVLLSNVYAAQGRWKDVGRVRDDMESKQVKKIPGLSY----------IEAKGTIHEFYNSDKSHEQWREIYEKIDEIRFKIREDGY----VAQTGLVLHDI---GEEEKEN    
 795 UniRef50_Q6ZK14        Os08g0500600 protein n=4 Ta...   341     6e-92  1  10.5%  75:523     2:426     --------------------------------------------------------------------------YGALRDAASMREVFGSC--AGVADVVSWNTVIGGYVKCGEMESAKRVFDEMPRRNGVSWSTMvyAAAGELDVAREMFDRMPAIGR--NVVTWNSMVTGFARHglPLARKMFDEMPVR----NLVSWNAMLRGYSVNSDMDGARELFDVMPEK----DVVSWTCMISGYAQAGRYQDTLELFRAMqeSNVQPNEVTMVSVLSACANLTALEEGRWVHAFIDKHKMVLNNEY----------NLGAALIDMYAKCGRTDIAVKIFNSLDRK------NVSAWNALITRLAMNGNARDSVDAFEQMKRTGEKPNDITFVGVLTACSHGGLVDEGRRCFQSMAStgVQPehYGCMVDMLGRAGLLEEAEELIRSMP----MAPDVMVFGALLGACRMHKRFDVAERVQSEIHSLNLNAQQSGCHVLIsiYAAAGKWFDVLEARQVVQRSGIRK--------------------------------------------------------------------------------    
 796 UniRef50_C5WRU1        Putative uncharacterized pr...   341     6e-92  1   8.9%   3:595    91:642     --ACSlyDDCEDWSSATRVFAEMPRPDAVSYTAMISALLRAGDCHGALALYPCMLPL---CAPTQHTFSKLLAPCASMRLHRHGTQLHAQLLRWGclGLVLKTALVHMYATCGAMVSARAVLHATPETDVVLWTAIITAYTRRGQlaMLAFRDMERSAVPPNAFTYAALIAAcaDHSLQIGQQLHARLFKSCLEHDTSACNALLDLYSKSsdLLHAFHAVDR-------PNVVTWTSFIAGLMRHGREQEAFAAFAHMRATGVQPNSFTLSTLLKGCTTAHACLHAAKIHAYVLKTSSESL-----------DAAVGNSLVHVYARSERLDDAWEVATTM-----SFVRDRFTYTSLAKGLNQIGLHHRALDLILRMFHEEVSIDGFSLACFLSAAATLASIESGKQLHCCAVKLGlqVSLSNSLINMYSRCKCLEDAKRAFQSIRE-----PSVGSWNAIISGMAFNASYTEALSVFEDMILAGAQPDGVTFTVVLSTCSRGGLVDIGIKHFNSMTNLFDVSPQKSHYTWFLDMLGRAGRFTEVAHTIEAMP------------------------VQPD-ISIYRTLL-AYCK--------    
 797 UniRef50_B9HGU6        Predicted protein n=4 Tax=M...   340     7e-92  1  13.4% 139:527    16:392     ------------------------------------------------------------------------------------------------------------------------------------------GYFGNLDDALASFNRMLDRNPQPPIIEFNRLLSSVVRmkYETVVSLFKEMEFRGIKYNVCTLSILINCFCFLHHVDYGFSVFGKTLKRGFKPDVVIFTTLIDGVCRIGKTELAAGLLKEMGLVGCVPDVVTCNSLMRGYCSQGKIDKVRKIFHLMVSK------------GLKPDVYSYSIFINGYCKVEKIDEAMELFDEMSHRG--VVPNAVTYNTLIKALCQALRPRDAQVLFRKMCACGLSPDILAYSTLLDGFCKQGNLDEMLVLFQEMQRRLVKPdvYTIIINGMCRSRKVKDAKEVLSRLIVEGLK-PDVHTYTALVDGLCREGSIIEALRLFRKMEEDGCMPGSCSYNVLLQGCLQHNDTSTVVQLIHEMADRGFYADAVT----------------------------------------------------------------------------    
 798 UniRef50_A5ADS7        Putative uncharacterized pr...   340     7e-92  1   9.5%  71:587    80:673     ----------------------------------------------------------------------LLDLSVRYDDVELIKAVHASIFKL-AEDIHLANALIVAYLKLGMVXNAXKVFVGLSCPNVVSYTAMISGFAKSNReaMEIFFRMRSSGIELNEFSFVAILTVCIRldLELGCQLHAIVIKMGFLNYTFVSNALMGLYGKCGYLDXVLQLFDEMXHR----DIASWNTVISSVVKEMMYERAFELFRDMRrdGFRIDHFtyNAILSGFCQNGEGSKALAFFCRMVEEGvkQIHGFILKFGFGSNACIEAALLDMCTRCGRMADAQKMF----SQGXFXQSGSIIWTSMICGYARNAQPEEAISLFCQSQLEgmVVDXVASTAVLGVCGTLAFHEMGKQIHCHALKSGFLSdvGNSIITMYSKCSNMXDAIKVFNVMPAH-----DIVSWNGLIAGHLLHRQGDEALSVWSKMEKAGIKPDTVTFVLIISAYRhsNLVDNCRRLFLSMKTIYHIDPTVXHYTSLVGVLGYWGLLEEavHSFYARDKSHPQAKDIHSGLELLIMECLKAGYVPDTSFVLH----------------    
 799 UniRef50_B9RFN5        Pentatricopeptide repeat-co...   340     7e-92  1  10.3%  24:520    13:465     -----------------------VSLSAISSNSQISHFARIGQINRARKIFDEL------PYKTIISWNAIVAGYFQNKQPREAHDLFNKMPDRNTVSW---NGLISGYVKNGMINEAREVFDKMPERNVVSWTAMVRGYIQEgkQAESLFWEMPEK----NVVSWTVMLGGLIegRVNEAIKLYDLMPLK----DVVARTNMIGGLCMEGRLSEAREIFDEMPKR----NVVAWTTMISGYAMNNKVDVARKLFEVMPDK----NEVTWTAMLMGYTRSGRIKEAAELFEAMPMKP----------------VPACNEMIIGFGQSGEVGKAKWTFDQMREK------DDGTWSAMIKVYERKGLELEALDLFRLMQREGVRPNFPSIISILSVCGSLASLDYGRQVHTQLVRSQFDFDVysVLITMYIKCGDLVKAKRVFDRFSMK-----DTVMWNSIITGYAQHGLGEEALEVFFEMLSSGISPDEITFIGVLTACSYSGKHRRkcKLIVRMPKKR-----------------------------------------------------------------------------------    
 800 UniRef50_Q7XE15        Os10g0442700 protein n=4 Ta...   340     8e-92  1  10.6% 144:586    78:492     -----------------------------------------------------------------------------------------------------------------------------------------------PRLARAVHAQPLRRGLLpdLHVSNSLVHMYAACALplARRLFDEIPR----PNHVSWNALLDGYAKCRDLPAARRVFARMPQR----DVVSWSAMIDGCVKCGEHREALAVFEMmrHDGVRANDVTMVSVLGACAHLGDLVRGRKMHRYLEE------------HGFPLNIRLATSLVDMYAKCGAIVEALEVFQAVPVENT----DVLIWNAVIGGLAVHGMSRESLQMFQKMEHAGVVPDEITYLSLLSACVHGGLVDEAWSFFRSLEAQGLRpeHYACLVDVLGRAGRLEEAYGVVKSMPMK----PSVSVLGALLNACHLHGWVELGEAVGRQLV--HLQPDHDgyIGLSNIYAVARRWQEAKKARKVMEERGVKKVPGFSEIDVGRGICR----------FIAQDKTHPGSAEIYALLKLIAMDMKMKdtVPDYTCMY-----------------    
 801 UniRef50_A5B8S4        Putative uncharacterized pr...   340     8e-92  1  10.8%  96:569    44:479     -----------------------------------------------------------------------------------------------RKEDVLMLTKRISDYSLKGEVDYARTIFEEmhPDTVSWNVMIRGYVENHrdARELFDKMPVR----SSVSWNTMIMAYAkgKTHIAMKLFIVMPDK----DVVSWTAIITALSRGSHIEDAWRLFKLMPE----PSSVSWASIISGFQQNGLAAETLCRFKEMLSVGVQPTSHSFTSALTASADLAMLSLSQQLYSQLLKR------------GFESNTXIGNSAISMFIKSGSFRNARRVLEDLPQ------PDIVTWNAMVVGYGQNGYGIEAIMSFHQMQKAKFLPDRVSFLGLLHGCSHCGYVEKGKQYFRSMEsyGISpeHYACMVDLLSRAGFLKEAHKLIKEMP----FEPTCIFWRTLLNGCRIWGDLELGFYAANRILE--LDpnSSACLMVIDIYASAGRWKEVLEMRRQMRKR----------EARKELGCSWIEIKGRIHLFTTRDDTHPEADHIYLTLLLL----------------------------------    
 802 UniRef50_A9S733        Predicted protein n=2 Tax=P...   340     8e-92  1   9.6% 114:593    46:496     -----------------------------------------------------------------------------------------------------------------KSPQDVWDALDALPRCVGSWEDLmfRRQRKWRSAIVIYEWILQGsfKPDVGCFNMLMDAYGRTkwTEAENTFHLMKKFQCLPTETSFNVLMAAYSRGGQLERAERVLHEMKESNCSPGLVTYNTYLEVLNKSGSWQLAEDVFREMQNRGVPPAVNTFTLMINIYGKAHHSAKAEHLFQSMR------------KALCPPSLFTYTALINAHAREGNCVRAEEIFAELQS--VGFVPDIYTYNALLEAYSRGGHPAGAKEVFETMLEAGVKADHVSYNILIDAFGRAGLISDAQAIYDSMKKVGFKPtsHILLLSAFVKAGRVTDAENFVRRLESMG-VEPDTFMFNSLLGAYGNSGRMDKMESLYESMQGSVCKPDIITLNTLINVYAQGGYIERAEEIFNSLESKGFTPDVMS----------------------WTSLMGAYSKRKLYRKCVSVYQKMLIAGCIPDR-ATAKVLLSSC----------    
 803 UniRef50_B9HZ40        Predicted protein n=1 Tax=P...   340     9e-92  1  11.1%  33:589   121:667     --------------------------------SVIHVLVNSRRYDDALSllEVLEALNNscESNHAVFDALVRACTQIGATVGACEVIKKLQIEGCWVTIHAWNNFLSHLIKVNEIHRFWIVYKEMveNVNTFNVVVHALCKdlQEALSVFYRILKSGIWPNVVTFNMMVDGACkgDMDLALKLVRKMeaGSIKPNSVTYNSLIDGFCKIGGITVAEELRNEMMKIDIEPNVRTYATMIEGYSRAGCLEEALRLCDEMVERGLLPNSVVYNSIMHWLYMEGDVDGASLVFTDMSDKQ------------IPLDKFTCSILTRGLCRNGYITTALKFLNQVLENN--LIEDAFSHNILINFLCKSNNFAAARQLLARMYVRGLVPDVVTFGTLIDGHCKEGNIESAVQVYDKMVKGEEKPnvYNSIINGLCKDGLVDVARSLVDVLQRMGLV--DTITYNTLINGYFNCGKFDKAFKLSTLMQNAGILASSATYNTVIKFLCKFGCVQEAKELMTMMVLWGVLPDNITYRTLVIN-------INKNCSA---------------EEVIELHDYMVLKGVVPDKLTYENIV--------------    
 804 UniRef50_B9SHV3        Pentatricopeptide repeat-co...   340     1e-91  1  12.1%  42:528   161:639     -----------------------------------------GEEELCDLLFKLVYDTGFGTKDLETVYSMLVDCYVTESKVSLALNLIHEIKLLNIFPSMGVCNSLLKALLRSHQLDLAWDILEGMQSFGMHLNASiyCAEGNIQSGWKILMEMKNYGIKADVIAYTIVIDALCksCVKVATSLLFKMIHCGISVDSVSVSSVIDGYCKKGRSDKAINLLN---FFNVRPNIFVYSSFMTKLCKDGNMLEASKTFQEMSEFGLYPDCFCYTIIIGGYCKVGDINKAFQYLGEML------------KFGITPSVTTYTLLINACCKSGDMAMAEYLLQRMMLEG--LIPDVVTFNCLIDGFGKKGYLQKAFELLDIMRSAGMSPDTVTYNALIHSLIARGYPNEAKDILHELIRRGLSPdtFTDVIDGFSKKGNFEEAFLVWSYMSER-HVKPDVVTCSALLNGYCRARRMDEANALFLKMLDAGLKPDLVLYNNLIHGFCSAGNMNAACNLVAMMIDDGFLPNITTH---------------------------------------------------------------------------    
 805 UniRef50_C5XA99        Putative uncharacterized pr...   340     1e-91  1  11.9%  57:600    84:562     --------------------------------------------------------FRCASCASAFASNHLLRVLPHPLPL----RLFPRLPRRNPHSFTFLLASLS----------------NHLDADPAVAVSIASRLMGAHVHALAV-----KAGAAGDLYVRNALIHFYgcGDVAAMRKVFDELP---LVRDVVTWNAVLAGYVRAGMVGVAREVFDGMPMR----DEVSWSTVVGGYVKEGELEVALGVFKNMVVQGVKANEAAIVTALSAAAQLGLLEQGKFV------------HEVIKRAGVAMSMNLGAALVDMYSKCGSVAAAKEVFDAMPWR------DVFAWNSMICGLATHGLGHDAVQLFEKFVSEGFCPTNITFVGVLNACSRTGLVDEGRWYFKLmeKYGIemEHYGCMVDLLSRAGLVQEAIELIEGM----HIPPDPVLWGTVLSACKRHGLVDLGITVGNKLIELDPAHDGY-YVLLASIYAKAKKWDEVRKVRKLMSNRGTSKS----------AGWSLMEAHGIVHKFLVGDMDHKDSVQIYNMLCMIDRRLAEAGYVPDV----SSVLHDIGDEEKVH---    
 806 UniRef50_B9GGZ8        Predicted protein n=3 Tax=r...   340     1e-91  1   9.1%  27:518    38:544     --------------------------STPTWNCLIRAYSRSPTApkAILVYNFFIK-TCSTRPDNYTYPCLLKACSRLFTDSNGKQVHTHVIKTGLDSDIYVQNALVHFYGsgNSTDACFLFDRMPDRDVASWNSliYNTNNSFTEVMVLFKKLMCGCVKADKISLVIVLSACAQagLEYGRSVHGYVIKVGfeGDVVSYTILINAYVEMGLIDLARDVFDEIVDK----DRVLWNLMVHAYVKARCPNEALDLFEKMDSAGVIPDENTMVSVLLACASISDLQCARLLHRFINR-----------NSNVRQDVFLKTALITMYSKCGSVEEALVTFYKME------YTDVFTWTAMIEGLANNGYGNEALSMFNRMENQGIRPNESTLVSVLTACIHSGLVKEGCQLFKSMvkMQLKMEHFGCLIDLLSRAGLLHQAEEFIQLLLPEER----LIAYKTLLSACMKYSEFDFGKKVANEMSKLSSN-SHETRILLSNFYALAGQWAQVAETRKFIKE-------------------------------------------------------------------------------------    
 807 UniRef50_O49680        Pentatricopeptide repeat-co...   339     1e-91  1  10.0%   5:602   334:921     ----SMYSKCGDTEAAETVFEELVCRDVISSNAILNGFAANGMFEEAFGILNQMQSV-DKIQPDIATVVSITSICGDLSFSREGRAVHGYTVRMEMQSRAlvINSVIDMYGKCGLTTQAELLFktTHRDLVSWNSMISAFSQNghKAKNLFKEVVSE-YSCSKFSLSTVLAILTSCDSSdgKSVHCWLQKLGFGDNMLSANSVINMYIGCRDLTSAFLRLETMSE---TRDLTSWNSVISGCASSGHHLESLRAFQAMSREgiRHDLITLLGTISASGNLGLVLQGR------------CFHGLAIKSLRELDTQLQNTLITMYGRCKDIESAVKVF------GLISDPNLCSWNCVISALSQNKAGREVFQLFRNLK---LEPNEITFVGLLSASTQLGSTSYGMQAHCHLIRRGFQANPfaALVDMYSSCGMLETGMKVF-----RNSGVNSISAWNSVISAHGFHGMGEKAMELFKELSSneMEPNKSSFISLLSACSHSGFIDEGLSYYKQMEEKFGVKPVTEHRVWIVDMLGRAGKLREA-YEFITGI-GEPQKAGVwkEVAEVLFEM-----EPDNASYYISLANTYVGLGGWEEA-    
 808 UniRef50_D2VN29        Predicted protein n=1 Tax=N...   339     2e-91  1  10.5%  19:581    12:573     ------------------MKEYGVSPDIIHFNCLIKALIENGESEEAMNIYRQ----EKELIPNEVTFILLLKGCAITRSLVNGKKLHeiMKNYELYRSITVQNNLINMFSKCGETQTAIELFMKMLksVFVFNVLLDSFlgKVEHAESIFNQMEFK----DVITWNSFLKGCSqgDTKRLLLYFKIMPKY-TQPNDVTMTVMLTFFANHSMEQEACFLFDNMNtfKIARITAHYGCMIKLWANLKKMPEAISLYEDMINNNLVPDVPTLILLLSVCSEFGNLDFGKIIHRNISELRIR-------------DHNLFNALINMYAKCGDLTSAHEVFLSVPNK------DVVSWTCIINAYAIHGMGLKAIELLNQMSKENIKPNEHTLTCVLHACGHCGMVDEAKEIYNHMekYSIKPshLACLIDALARKGNIKEAEKLLGS-----CISPDVVTVSTLFGVCRKFNNVTLAEKLFNMLIERN-EVESSHFILMANIYASQDMYEDAKRVKELMEFGGFKKLPSMTQVC----------IEGKMHSFVCEEKEHPEIKMIEKELKKMVADLIKAGYQPD----------------------    
 809 UniRef50_D1H1G5        Whole genome shotgun sequen...   339     2e-91  1  10.8% 154:580     1:428     ---------------------------------------------------------------------------------------------------------------------------------------------------------MLRNGTLPDNYSFPFVLKACARLSlhKGREIHSSTLKLGVHLDVFVQNALISAFSSCGAVEAARAVFDMLPA--LVRDVVSWNSMISGYLQSHRYELALKVFDRIPHR----NTVCWTSMIAGYAQSDLFKEAIELFREMQIGggRWIHLYCERNSIEMDLNARNALIGMYSKCGDIQKALEIFHGLTQ------PDIFSWSAVISGLAMNGESDKALHLFSQMEmsDIRPNEITFLGVLCACNHGGFVDKGLYYFNAMTqyNLTpeHYGCMVDLLGRANLLVEAEKFIRTLP----IQPDVVIWRSLLFACRNHGNIELAEFAAKQIEE--LEPRrgARVLLSNVYASASRWGDVKRVRKDMATQRIKKQP----------GCSFVEIDGLVHELFVADRSHPEMGAIYETMISINKALQSKGFDP-----------------------    
 810 UniRef50_Q9ZUE9        Pentatricopeptide repeat-co...   339     2e-91  1  14.2%  93:528    96:524     --------------------------------------------------------------------------------------------KLNIRHSFWTYNLLTRSLCKAGLHDLAGQMFECMKSDGVSPNnlVSSFAEKGKLHFATALLLQSFEVEGCCMVVNSLLNTLVKlrVEDAMKLFDEHLRFQSCNDTKTFNILIRGLCGVGKAEKALELLGVMSGFGCEPDIVTYNTLIQGFCKSNELNKASEMFKDVKSGSvsPDVVTYTSMISGYCKAGKMREASSLLDDMLR------------LGIYPTNVTFNVLVDGYAKAGEMLTAEEIRGKMISFGCF--PDVVTFTSLIDGYCRVGQVSQGFRLWEEMNARGMFPNAFTYSILINALCNENRLLKARELLGQLASKDiqPFMYNPVIDGFCKAGKVNEANVIVEEM-EKKKCKPDKITFTILIIGHCMKGRMFEAVSIFHKMVAIGCSPDKITVSSLLSCLLKAGMAKEAYHL-NQIARKGQSNNVVPL---------------------------------------------------------------------------    
 811 UniRef50_C5YNT9        Putative uncharacterized pr...   339     2e-91  1  10.2%  61:579     2:489     ------------------------------------------------------------PPPTAGRLSVLIRRCAASGALSPGAQVHAQGLVGGLLPNTTLDTDFVLLYSRCGALHRARQVFDGMspSMHAYNVLLAAS--PPGTAVELISRLLASGLRPDRYSIPAVLRACAELrmLLGAVFHGLALRLGLLANVVVSGALLDMYAKTGMLADATRVFDEMPER----DAVVWNCMVTGYARAGRSEETLELFrqIESVDMARDLRAVPNVLNVCANEGQLMKGREIHGRMVRCFA-----------FDSDIAVGNALVDMYAKCGQVDGAQAVFASMKVR------NVMSWSTLISCYGVHGMGKEALRIYEEMVSRGVKPNCITFISILSSCSHSGLVSDGRRIFESMSkhAVEpdHYACMVDLLGRAGAIEEAVGIIRKMP----MEPGASLWGALLSACATHNNVDVGEVAAYKLFELE-EGNASNYVTLCGIYDAIGRSDSVSGIRSRMRELGMVKTP----------GCSWVHVKGRAHAFYQGCIPRYLRRQMLWVLGHLLEDMTASEYE------------------------    
 812 UniRef50_Q67X36        Os06g0231400 protein n=3 Ta...   339     2e-91  1  11.8% 177:582   146:515     --------------------------------------------------------------------------------------------------------------------------------------------------------------------------------RPDLARVLFRT--SGGGALDVVSWTTMVGGLCRLGLVDDAREVFDAMPAR----NLVSWNSMISGYVKADRFLDALEVFDEMRALGVEGNGFVATSALVACTGAGALGRGREIYRWVEQS------------GIEVDAKLATAVVDMYCKCGCVDEAWGVFDSLPARG------LTTWNCMIGGFAVHGRCDDALELFHQMEAAGVAPDDVTLLNVLTACAHAGEVSEGRRYLNHIvrHGIEpeHYGCMVDLFGRAGQLDEAKKVIDEMP----MDPDLAVLGALLGACKIHGDVDLGEAIGWRVIDLDPD-NSGRYVLLANLLAGAGRWDEVGKVRRLMDERNVSK----------EAGRSVIEVDGEACEFRCGNLRHPQAREIYAMAVDMVSRIRAEGYVPDT---------------------    
 813 UniRef50_Q9SAD9        Pentatricopeptide repeat-co...   338     3e-91  1   9.0%   1:502    10:512     LAYRSRIakSGMIDNAVQVFDEMRHSssFDYNRFIGVLVRESRFELAEAIYWDMKPMGFSLIP--FTYSRFISGLCKVKKFDLIDALLSDMETLGFIPDIWAFNVYLDLLCRENKVGFAVQTFFCmePDVVSYTILINGlaGKVTDAVEIWNAMIRSGVSPDNKACAALVVGLCHArvDLAYevAEEIKSARVKLSTVVYNALISGFCKAGRIEKAEALKSYMSKIGCEPDLVTYNVLLNYYYDNNMLKRAEGVMAEMVRSGIQLDAYSYNQLLKRHCRVSHPDK---CYNFMVKE--------MEPRGF-CDVVSYSTLIETFCRASNTRKAYRLFEEMRQKGMVM--NVVTYTSLIKAFLREGNSSVAKKLLDQMTELGLSPDRIFYTTILDHLCKSGNVDKAYGVFNDMIEHEITPdsYNSLISGLCRSGRVTEAIKLFEDMKGKECC-PDELTFKFIIGGLIRGKKLSAAYKVWDQMMDKGFTLDRDVSDTLIKASCS-----------------------------------------------------------------------------------------------------    
 814 UniRef50_Q9LQD1        F28C11.9 n=7 Tax=rosids Rep...   338     3e-91  1   9.9%   4:548    50:578     ---NRRIdkSGNLLSAHEAFDEMSVRDVVTYNLLISGNSRYGCSLRAIELYAEMV--SCGLRESASTFPSVLSVCSDELFCREGIQVHCRVISLGFGCNMFVRSALVGLYACLRLVDVALKLFDEmdRNLAVCNLLLRCFCQTGEslFEVYLRMELEGVAKNGLTYCYMIRGCSHDRLveGKQLHSLVVKSGWnsNIFVANVLVDYYSACGDLSGSMRSFNAVPEK----DVISWNSIVSVCADYGSVLDSLDLFSKMQFWGKRPSIRPFMSFLNFCSRNSDIQSGKQIHCYVLKMGFD-----------VSSLHVQSALIDMYGKCNGIENSVLLYQSLP------CLNLECCNSLMTSLMHCGITKDIIEMFGLMIDEGTGIDEVTLSTVLKALSLseSLHSCTLVHCCAIKSGYAADVacSLIDAYTKSGQNEVSRKVFDEL-----DTPNIFCLTSIINGYARNGMGTDCVKMLREMDRMNLIPDEVTILSVLSGCSHSGLVEEGELIFDSLESKYGISPGRKLYACMVDLLGRAGLVEKAERL-------------------------------------------------------    
 815 UniRef50_B9HGK2        Predicted protein n=2 Tax=M...   338     3e-91  1  10.7% 126:574     1:420     -----------------------------------------------------------------------------------------------------------------------------MLQVGIYPDHLTysRLLRKELGVSIHGHVIKSGFEIDRFVANSLIHMysCGDIVYARKVFDGTPVK----NLVSWNSMVDGYAKCGYLDLARGLFDLMPER----DVRSWSCLIDGYAKSGNYGDAMAVFEKMRTSGPKANEVTMVSVLCACAHLGALDKGRMM------------HQYLVDNGFELNLVLRTSLIDMYAKCGAVEEAFAVFRGVSLR----KSDVLIWNAMIGGLATHGLVKESLDLYTEMQIAGIKPDEITFLCLLSACAHGGLVKQASYVFEGLGKNGMTpeHYACMVDVMARAGQVAEAYQFLCQMP----LEPTASMLGALLSGCMNHGKLDLAELIGKKLIELDPEHDGR-YVGLSNVYAIGRRWDEARIMREAMERRGVKKTPGYSF----------LEMSGAHHRFIAHDKSHPSSEQIYTMLSFIVSQMQ-----------------------------    
 816 UniRef50_A9TCH6        Predicted protein (Fragment...   338     4e-91  1  11.1%  62:530    31:497     -------------------------------------------------------------KFSNVQANIILEALCGDDLTEQVLSFFRWVrhEPCLCDPRSFTILFTFLGRMAMPDQALVFFGMLPKdvHVYNTLITCLTNCNRYDLELYHNLAFYEVFPDSVTFSVLMNSAVKGglKEVWALYQDMVQRGISPSVSVFGTFIKAFCDAGRLKEALLITTEMEKLNVPFNVVIYNILIDAYGRVGQLEEAEGLMTEMRERGIQPNVGTYNALITGYLEarQFVVAEGLIEEMEAS------------GLYPDVVTFTMLLGAYGHEGLTEQAEQVFNRMKARG--VQPNSYSYTALINAYAERRCPEKAARAFEMIRKQGVNPTVETYTALLDAYRRAGDLEMVQAVWKSMKVEGCVavTYMTILDAFQKQGRYKEARDLIEEFKNQGHK-PDLMVYNMLLNSYMRGGRHVKASDILLEMKTAGFLPDSFTYCTLIYGFLRVRDQTKALKYHREMMNRGQLPDPKTYAK-------------------------------------------------------------------------    
 817 UniRef50_UPI0001983752 PREDICTED: hypothetical pro...   338     4e-91  1  10.5% 126:582     1:459     -----------------------------------------------------------------------------------------------------------------------------MLQKNFSWARRITSLANQvrKAITEFHQLQKSGQKITEFLLSSVVKVCGRVelQEGKQAHCLILKLGFNADLILMTTLLDMYSKCTNIQEACRVFEEMPQR----DVVVTNSMISGLCRCHLPSYAIDLFTKMNES--ERDVGTWNSLISGLGQNLEGRTALIIFGRMRVehAQQVHGLVIKSGFEQYLPVGNAIVDMYAKCGCMDDACLCFQNMPFK------NVISWTSLISGYAKHGLGIEALKAFQQMEMEGVVPNKITFLGTLYACSHAGLLQEGWSNFTTMVHKYltMEHYTCMVDLLARAGHLEEAHDFIERMP----INPDAKLLTAFLSSCCHHKNVELARTVGQKLLQ--LQpeAGAYISLSNFYGLVGDLEGVANVRRLMLERGIRKEKA----------CTWIEISRKVHSFESGDRSHPDYLKICKYLEVLIQKMKNIGYVPDT---------------------    
 818 UniRef50_B9N9H8        Predicted protein n=3 Tax=r...   338     5e-91  1   9.9%  70:574     8:512     ---------------------------------------------------------------------LLLQSCA---DLVTLKQIHAQALTQGLleQPLACKLVNRYAKLGNPRDAQKVFGYIQDPDRVTYTSLINLYlqLPIKAFSVFSKLVNEGLRPDSHSVVGALSACGKkdLLNGKLVHGMIFRFQLGANSIVGNALIDMYCRNGEIKIAQLVFKQM---GIK-DVSSWTSLLNGFVMCNGLESARRVFDEMPWR----NDVAWTAMITGYVRGGMPIRGLEMFRQMKAEggQAVHGYISKVNLDKGVTVSNALMDMYSKGGCVESAMKIFDRLVKK------DVFSWTTMISAHSSHGKGNHALEVFYDMLESGVIPNDVTFLLVLSGCSHSGLLVEANKLFNGMIqyGFEpeHYGCMVDLLCRAGLLEEAKELIDNMP----MDPDAVIWRSLLSACMNQRNLGLAEIAGKKIIE--LEpdDGVYVLLSNIYHVANRMKDARKMRKMMGDQKVMKKPA----------CSYIELNGVVHQFHTENATHHASTKIYMLLEMINEHLR-----------------------------    
 819 UniRef50_Q9FKR3        Pentatricopeptide repeat-co...   338     5e-91  1  11.7%  16:552    30:543     ---------------LKHKVDSGLLKSAIT-TQVISELSLFSGYGGPSLSWSFFIWTDSLPSSKHSLQSsmILILTKHKHFKTAHQLLDKLAQRELLSSPLVLRSLVG--GVSEDPEDVSHVF--------SWLMIYYAKAGMINDSIVVFEQIRSCGLKPHLQACTVLLNSLVKQRLTdvWKIFKKMVKLGVVANIHVYNVLVHACSKSGDPEKAEKLLSEMEEKGVFPDIFTYNTLISVYCKKSMHFEALSVQDRMERSGVAPNIVTYNSFIHGFSREGRMREATRLFREIKDD-------------VTANHVTYTTLIDGYCRMNDIDEALRLREVMESRG--FSPGVVTYNSILRKLCEDGRIREANRLLTEMSGKKIEPDNITCNTLINAYCKIEDMVSAVKVKKKMIESGLKLDMYsaLIHGFCKVLELENAKEELFSMIEKG-FSPGYATYSWLVDGFYNQNKQDEITKLLEEFEKRGLCADVALYRGLIRRICKLEQVDYAKVLFESMEKKGL-VGDSVIFTTMAYAYWRTGKVTEASALFDVM---------------------------------------------------    
 820 UniRef50_Q84ZD2        Os07g0548300 protein n=6 Ta...   338     5e-91  1  13.9%  85:574   168:659     ------------------------------------------------------------------------------------LRLLGLLREHSFLPDLASYslLASLLNTRDPPDAALLDrsRLEPDAPLFSDLISAFARapDAALELLASAQAIGLTPRSNAVTALISSLGSArvAEAEALFLEFFLAgiKPRTRAYNALLKGYVKIGSLKNAEQVLDEMSQCGVAPDEATYSLLVDAYTRAGRWESARILLKEMEADGVKPSSYVFSRILAGFRDRGEWQKAFAVLREMHAS------------GVRPDRHFYNVMIDTFGKYNCLGHAMDAFDRMREEG--IEPDVVTWNTLIDAHCKGGRHDRAIELFDEMRESNCPLGTTTYNIMINLLGEEQRWEGVEAMLAEMKEQGLVPntYTTLVDVYGRSGRFKEAVDCIEAMKADGLK-PSPTMYHALVNAYAQRGLADHALNVVKAMRADGLEASTVVLNSLINAFGEDRRIAEAFSVLQFMKENGLRPDVIT-YTTLMKALIRVEQFEKVPVIYEEMITSGCAPDrmLRSALRYMKHMR-----------------------------    
 821 UniRef50_A3PM02        Pentatricopeptide repeat co...   337     6e-91  1  11.2%   4:590   412:991     ---NTLIAKAPDfgVRAEMVAAGIKPNEVTFNTLI---AKAADFARAEGVRAEMVAAG--IKPNEFTFNTLI---AKAPDFTGAEGVRAEMVAAGIKPNEVTFNTLIAKAADFARAEAVRA---EMVAAGIKPNEVTFNTLidFTGAEAVRAEMVAAGIKPNEVTFNTLIA--KAADFARAEAVRAEMVAAGIKPNEVTFNTLI---AKAPDFTGAEAVRAEMVAAGIKPNEVTFNTLI---AKAADFARAEAVRAEMVAAGIKPNEVTFNTLV---AKAPDFTGAEGVRAEMVAAGIKpvRAEMVAAGIKPNEVTFNALV---AKAPDFARAEAVRAEMVAAG--IKPNEVTFNTLI---AKAPDFTGAEGVRAEMVAAGIKPNEVTFNTLV---AKAPDFTGAEAVRAEMVAAGIKPntFNTLV---AKAPDFTGAEGVRAEMVAAGIK-PNEVTFNALI---AKAPDFARAEAVRAEMVAAGIKPNEFTFNTLI---AKAADFARAEAVRAEMVAAGIKPNEFTFSTLIAKAPDFTGAEGVRAEMVAAGIktLIAKAPDFARAEAVRAEMVAAGIKPNE-FTFSTLV-------------    
 822 UniRef50_Q2QXB0        Os12g0163600 protein n=3 Ta...   337     7e-91  1   9.4%  69:579    16:504     --------------------------------------------------------------------PLLIHLLRGASDLASVSATHAKLLKLGAASSVASSNHLLAAYGRCGCTALARHLFDGMrrNVVSWTALMSGyaGRPRAAVALLRAMARSGVPPNAFTFSTAASSCAraDAGLGRQVHARAEVAGYACDSVVSTALVDMYGKAGRVGDARAVFDAMPV--TARNAVSWASMLSVYAQNALGHDAIQLFAELRTNgmAPNHFMLSSVVNACAGVGRL------------GIGKCIHGMVLHHGHEDNDVVAVALVDMYSKCGIYEYSKKVFDRI------NQPSVIPYTSIIVATAKYGLGRCALALFSDMVGRGVQPNSVTLLGVLHACSHSGLVDTGLQLLRSMreYGIDpsHYTCAVDMLGRAGQLEAAFELADEAQVNGNDA--LMLWSSLLSACRNHKRLDLATMAGQKVSeqDV---AGALVVMSNAYASAGQADDAAAVWSNMRRRGIRKDP----------GCSWIEIKDISYVFYAGMVsgSRAGEVMLLLDELEGKMREKGYK------------------------    
 823 UniRef50_Q0DLJ0        Os03g0861900 protein n=3 Ta...   337     7e-91  1   8.0%   3:552    93:632     --ACSLLAFYAAAAsaCNLFDEMAHKDAVSYTAMISALVRAghWRQALALYPCMLQAGA--PPTQHTFAQLLSACASGRLHHQGTQLHAQLLRWGAGLNLVLKTALLHMYCNCGNMGYAHTVLHspQTDVVLWTAIITGYARSGdsALQMFRSMTRAAVLPNAFTYAALISACSSFRALqgRQIHARLFKFGLEHDTSVCNALVDLYSKSsrLLDLLHTFHAVDK----PNVVSWTALIAGLACHGRDEEAFLAFSQMRLSGVLPNSFTVSTLLKGSSSSLAFLHARKIHGYILKTSFESL-----------DAAVGNSLVDVYVRFARMDDAWAVATTM-----AFIRDRFTYTSLARGLNQMGLQQRTLEMFVRMFHEDVDIDGFSIASFLSSAASLASIETGKQLHSCSVKLGLSSdvSNSLIDMYSKCKCMGDARSVFQSISE-----PKVVSWNALMSALVSNEYYNEALSAFEDMTLVGAKPDGITFSLMLFACNRSGLVDIGIKYFNSMGNLFGVLPQRSHYTLFLDMLGRSGRLTEAASTIDVI---------------------------------------------------    
 824 UniRef50_Q5Z8I7        Os06g0694300 protein n=2 Ta...   337     8e-91  1  10.6% 167:580   181:559     ----------------------------------------------------------------------------------------------------------------------------------------------------------------------NSLIGAyaCGDVGAARKVFDEMVEK----DVISWTSIVVAYTRSGDMRSAEEVFGRCPVK----DMVAWTAMVTGYAQNAMPVKALEVFDRMAELGMVIDEVSLTGAISACAQLGALRRAAWV------------QEIAERTGFGNNVVVGSGLVDMYAKCGLIDEASKVFYGMQEK------NVYTYSSMIAGLASHGRASEAIALFKEMvrANVEPNHVTFIGVLTACSHAGMVGEGRYYFAQMkkYGIMpdHYACMVDLLGRAGLVDEALDLVRSMS----VTPHGGVWGALLGACRIHGKSEIAKVVAEHLFK--LEpsIGNYVLLSNILASAGKWEEVSKVRILMRKQRLKKDPAVSLFE--------GR-DGLVHQFFAGDNAHPRTHEIKKALLELVAKLKLEGYVP-----------------------    
 825 UniRef50_C5Z2R1        Putative uncharacterized pr...   337     8e-91  1   9.9%   4:597   181:831     ---LRLYGKVGKIKLAEemLQVGCEPDAVACGTLLCMYARWGRHKDMMLFYSAVR--RRDLVPPVSVYNYMISSLQKQKLHGKVIHVWKQMREAGALPNQFTYTVVISSFVKEDLLEKAMDVFGEMRQCRFVPEEATYSLlgKGEQALRLFEEMKAQGIVPSNYTCASLLALYYKndYSKALSLLSEMENSKVIPDEVIYGILIRIYGKLGLYEEAEQTFEKIEKAGLLSDEQTYVAMAQVHLNAGDYDRALEVLESMMMRNVKPSHFSYSAILRCYVAKEDIVAAEDTFRALSQHGL-------------PDVFCCNDLLRLYMRLGHLEKARALVLKMREDDFRLDEDLSM--TVMKFYCKSNMINDAEKIFKDIQRNNKTVKIPTMLLLIEMYARngSTDEAKFIYDQLTELGFKPDhiATLVVQYGQGKQLEQAQELFESASALFPEGANV--YNAMVDAFCKCGKTEDAYHLFMEMVDQGSNRDAVTVSILVTHLTKHGKLHSAISIYDRMISSGTSQSMQT-FNLMISVFGKGGKLDKAVEMfyTNMLSLYGKAGRHQEASLMFKRMKEDGIRPGKI-SFNSMINAYATSG------    
 826 UniRef50_Q9MAT2        Pentatricopeptide repeat-co...   337     9e-91  1  10.7%  19:580    82:576     ------------------IFRNSEERNPFVLNALIRGLTENARFESSVRHFILM--LRLGVKPDRLTFPFVLKSNSKLGFRWLGRALHAATLKNFVDCDSFVRLSLVDMYAKTGQLKH----------------------------AFQVFEESPDRIKKESILIWNVLINGYCRadMHMATTLFRSMPER----NSGSWSTLIKGYVDSGELNRAKQLFELMPEK----NVVSWTTLINGFSQTGDYETAISTYFEMLEKGLKPNEYTIAAVLSACSKSGALGSGIRI------------HGYILDNGIKLDRAIGTALVDMYAKCGELDCAATVFSNMNHK------DILSWTAMIQGWAVHGRFHQAIQCFRQMMYSGEKPDEVVFLAVLTACLNSSEVDLGLNFFDSMrdYAIEPthYVLVVDLLGRAGKLNEAHELVENMP----INPDLTTWAALYRACKAHKGYRRAESVSQNLLELDPEL-CGSYIFLDKTHASKGNIQDVEK---------RRLSLQKRIKERSLGWSY-IELDGQLNKFSAGDYSHKLTQEIGLKLDEIISLAIQKGYNP-----------------------    
 827 UniRef50_Q5JML3        Os01g0754701 protein n=8 Ta...   336     1e-90  1  10.4%   7:565    91:644     ------FGRNGRFDDALRLFGEMPVRSVVTWNCLISSFAQHGRACDAVIWFRALVRSG-----NSVSDGSLVAVLPAFGTP---EQVHGLVKKIAIDSFSTVANSLLNSYCTHSSTSLAENLFSemFRDVVSWNTMIIAFARrpQRALEFFWMMEGWSVLPNEITFSSVIYACTsnGHEHGKSIHAKVIKHNLNTRPFVNTSLFDFYSKCGCTKDAHKVLEEAPEDSTTR----WNGLISCHSDT-DVPTSFFILREMLRSGVMPNEVSLSSLL----KDPSVFDLLQIHSLVTRLGYdaSHEMVSDamLDPdaVSMNVLAGVYNRVRMYDEAKKLL------LHQGCNDTVSWSILITACARNGDYAEALKFFELMRILGHHFDSYVSVSLLSICTKSNSLVLGRLLHGLIIksGC-LDTynMLLDMYAKCGRIEECLKAFKEMEDR-----NIISWTAIISGLALNGFSRKALAWFKAMEEDGFKPDKVAITAVLSACRHGGLVHEGMNIFRHMKSEYSIEPEMEHYICVVDMLCKCGHLKEAE----VVIRGMPFQPSAVVW--------------------------------------    
 828 UniRef50_B9MV11        Predicted protein n=3 Tax=r...   336     1e-90  1  11.9% 146:599    42:473     -------------------------------------------------------------------------------------------------------------------------------------------------KAFEFFNWASRNGFAHDSRSYFMMLDILGrgNLNIARNFLFSIERRsvKIEDRFCNTLLRSYGNAGLFNEAIKLFSLMKSSGVSPSVITFNSLLLILLKRGRTNMAHSVFDEMctYGVTPDTYTFNILIRGFCKNSMVDEGFRFFKEM------------SRFNCEPDVVTYNTLVDGLCRAGKVRIAHNVVKGMVKKMKDLSPDVVTYTTLVRGYCMKQEIDEALVVFEEMVSRGLKPNDITYNTLIKGLCEVQKFDKIKEILGGAvgRGFVPdtYNTLMNAQCDAGNFDEALKMFKKMKELK-VQPDSATYSVLIRNLCQRGDFERAEQLFDklRDDGCTPLVAAYNPIFDFLCKNGKTHKAERVFRQLMKKGTQDPPS----------------------YKTLIIGHC-KEGTFEAGYKLLLFMLRRDYVPD-FETYVLLINGFLQKGEP----    
 829 UniRef50_A5AME8        Putative uncharacterized pr...   336     1e-90  1  11.1%  81:575     1:498     --------------------------------------------------------------------------------MQEAKRLHAHLITSGLHqeNHLRKLITLYTSSSSSLHHARLLFDAvhPSTYLYNTmvYAASPTPLHALLLHRHMFRHGPPPDTYTFPFLLKAcaLAHLLKGQELHCQALKFGLGGHVFVENSLIHLYGSNSRMDSARRVFDGMGYR----DIASWTTLLACYANSCSVEAARKVFDEMPER----SVVSYSAMLAAYVRGNRFREALELFRELFSVKIEPSDSCVMskGDYVDSRIATALIDMFFKCGSIEHALLVFEGAKEKHVGE------WTAMLSGLAMHGLGEQLIEAFEKMVDSGIKPDEVTFVALLSGCSHSGLVNEGLYYFDRMESdgVEpeHFGCVVDLLGRAGLIDQAMQLISEMP----FEPNAAIWGALLNACRVYKNVEVGELAAQWLIKD--EpnGALYMTLLSLYREAGRWDEVEKVKLKMKEVGCRKGP----------GCSLIEVGGICYEFVVGDKSHPCAVEICLNLSKANMEFKE----------------------------    
 830 UniRef50_B9SZK3        Pentatricopeptide repeat-co...   336     1e-90  1  10.7%   7:493    36:485     ------YAKNGQIAKAKKLFDEMPERTTATYNAMITAYIRNSCM--VDEAFSLFSRMS---ERNAVSYGAMITGFLKAGMFEKAENLYREIPAKWREP-VCSNAMISGYLKVGRLEEAIKVFEGTVEKDVVSWSSMVDGYCKKGreARELFNMMPVR----NVVTWTSMIDGYMNVGFFegFSLFLSMRRV-IEVNPLTLTIMFEACRHFGRYREAMQVHGLVLRIGFEFDIFLGNSIIAMYCEFGCMDEAKRMFQMMNNK----DVVSWNSLISGYIQHDEIEEAYKLFEKIPGK----------------DVVSWTTMITGFSAKGNVQKGIQLFKIMPEK------DDVAWTAVISGFVSNGEYEEAFLWFIEMLKKAVKPNSLTFSSMLTASASLATLNQGLQIHAHVEKMDvdLSIRNSLVSMYSKCGNIAEAYQVFTS-----INAPNIISFNSMITGFSQNGHGEEALDLFSKMRKENQEPNEITF--------------------------------------------------------------------------------------------------------------    
 831 UniRef50_D1HRL6        Whole genome shotgun sequen...   336     1e-90  1  10.2% 141:599   120:541     --------------------------------------------------------------------------------------------------------------------------------------------LSRVCMGRCVHGQGLRVGLEGDFYVGASLIDMYVKCGvgDARKLFDKMIVR----DMASWNALIAGYMKEGEIGVAEDLFERMEHR----NIVSWTAMISGYTQNGFAEQALGLFDEMLQDgmKPNWVTIVSVLPACAQSAALERGRRIHD------------FANGIGLHLNSSVQTALAGMYAKCYSLVEARCCFDMIAQNGK----NLIAWNTMITAYASHGCGVEAVSIFENMLRAGVQPDAVTFMGLLSGCSHSGLIDAGLNHFNDMGthSVEpeHYACVVDLLGRAGRLVEAKELISQMPMQAGPS----VWGALLAACRSHRNLEIAELAARRLFV--LEPdsGNYVLLSNLYAEAGMWEEVKKLRALLKYQGMKKSP----------GCSWIEINGKSHLFMGADKSHPQAKEIYKFLEALPEKIKMAGYIPDTSF----VLHDISEEEKE----    
 832 UniRef50_Q9LHN5        Putative pentatricopeptide ...   336     1e-90  1   9.5%   6:575    32:672     -----LYSKSGLLREARNVFDEMLERNVYSWNAVIAAYVKFNNVKEARELFemFGEMHRKEKDDifTVTTMVKLSAKLTNVFYGEQLHGVLVKTgfVDSVARNAMIAAYCREGDIDKALSVFWRNPenDTISWNTLIAGYAQNgeEALKMAVSMEENGLKWDEHSFGAVLNVLSSLklKIGKEVHARVLKNGSYSNKFVSSGIVDVYCKCGNMKYAESAH---LLYGFG-NLYSASSMIVGYSSQGKMVEAKRLFDSLSEK----NLVVWTAMFLGYLNLRQPDSVLELArqAYMEPGKEIHGHSLRTGILMDKKLVTAFVDMYSKCGNVEYAERIFD------SSFERDTVMYNAMIAGCAHHGHEAKSFQHFEDMTEGGFKPDEITFMALLSACRHRGLVLEGEKYFKSMIeyNISPehYTCMIDLYGKAYRLDKAIEL---MEGIDQVEKDAVILGAFLNACSWNKNTELVKEVEEKLLVIEGS-NGSRYIQIANAYASSGRWDEMQRIRHQMRGKELE----------IFSGCSWANIDKQFHMFTSSDISHYETEAIYAMLHFVTKDLSE----------------------------    
 833 UniRef50_B9ID27        Predicted protein n=1 Tax=P...   336     2e-90  1   8.7%   7:571    16:516     ------IQRCKKLKIFKKIHAQLITSGVVSNDLVVNRAAeaKGPNFADYACDFLSQMEDGFLPYMFTFPAVLKSCAKFMAVDEGRQVHGVIAKMDFVCDIYVENSLVHF----------------------------YSVCGSLDDASRVFDEMLVR----DAVSWTGVIYPGIKGYLILGKGIHGLIIERGFGLGLEVSNALMDMYVKCESLPEEKQVFDELPEK----DIVSWTSIICGMVQCKFPKEVLELFCDVQSSGIEPDGIILTSVLSARASPGALDYGRLVREYIDHKAIKW------------DIQIGTAVVDMYAKCGCIEMAMQIFNVMPHK------NVLTWNAMLNGLAMHGHGQKALQLFEEMVREGMRPNEVTFLVTLTACCHCGFVGEGRRYFHWMKsyNLPpeHYGCMVDLLCRAGLLDEALELTKAMP----MLPDVRIMGALLSTCKANGNVELPREILDRLVEFDSR-DSGVYVLLSNILAINQRWADVTRIRRLMK-MGIEKTPGS----------SVIEVDGKAHEIIVGDLRYPQDKHIRLFLKSLSD--------------------------------    
 834 UniRef50_Q65XP2        Os05g0153400 protein n=5 Ta...   336     2e-90  1   8.7%  78:573    31:501     -----------------------------------------------------------------------------VRSATHVMPLYAVLLKLGLSASAILATSLAHLALRCGLPGYARRVFDEmhRDVVSWTSLITGHAHQgqDSLALLRRMVISGVVPNGYSLSGALLACAggALAAGKEIHARVVKMSLhvDAVVENGVLDMYTRCGKIDYARKLFGVMLVR----DIVAWNSMMAGCLRSGQAEEALGLFSSMVSSGVDADGFSFAISVDACGELALLKQGMQ------------AHARVIRGGFDSDVVVRNSLVDMYAKCGCVDSAGLVFRDALSS------DAVLWTTMISAYGKFGRVHDAICMFDRMSQLGIKRDGVAYLAVLSACSHSGLVKEGWNYFKLmnSVKMQPEHYGCMADLLCRSGYLEEALDFITNMP----FESSIAAWSALLNSCRIHGNAKLGQLAASRLVQLDPE-NHSNWVALSNVHASESDWHETWMIRESMSIECVKKEP----------GCSWVELHDGVHVFLMADQSQPELVDVLQTLDSLKEDI------------------------------    
 835 UniRef50_A2ZMR2        Putative uncharacterized pr...   336     2e-90  1  14.5% 135:597   100:564     --------------------------------------------------------------------------------------------------------------------------------------AVLRAQArdVRRAFEFFRWADRQWRYRHAPEFAQLMLSYSrgKLRSAMRVLHLMQKDGCAPDISICNMAVNVLVVAGRVDKALEFAERMRRVGVEPDVYTYNCLIKGLCGARRVVDAMEMIGVMLQNGCPPDKISYYTVMSFLCKEKRVEEVRGLLQRMRN-----------DAGLFPDQVTYNVLIHGLAKHGHADEALEFLRE--SEGKRFRVDEVGYSAIVHSFCLNGRMAEAKEIVGEMISKGCQPDVVTYSTVVDGFCRIGELDQARKMMKHMYKNDCKPntHTALLNGLCKVGKTSEAWELLNK-SEEEWWTPSDITYSVVMHGFRREGKLKESCDVVVQMLQKGFFPTTVEINLLIHALCKDGKPAEAKDFMEQCQSKGCTINVV-NFTTVIHGFSRQGDLESALSLMDDMYLshPdkKGRLKEATGLVEKMLNRGLLPTPV-TYRTVIHRYCEKG------    
 836 UniRef50_D2UZ94        Predicted protein n=1 Tax=N...   335     2e-90  1  11.4%   7:601   139:725     ------LDMCEEI--LQELESQGLC-NVFSYIYYTKACLQNGNTEKAFSVIGRME--SKQIEPSFGIYIPLLKEATNQNNVEVGLKLYDMIKKSNLASFQLSSCIATFALKVNraIEEYEKIKQSELMNVITWTCLIQAYIqdTEKAIDTLIDMKKHGIYPDSATFTILLSACANSNmeEGVKIHELAKQTLKDFTIGLSNTIINFYTKCGYSNIAIEECEKLKQNGLM-NVVTWTCLIQAYIQQNDIEKAIDMIIDMKKH----DSTTFTILLSACANSNMYEEGVKIHELAKQTLKDFTIELS------------NTIINFYTKCGYSNIAIEEYEKIKQSELM---NVITWTCLIQAYIQQNDIEKAIDTLIDMKKHGIYPDSATFTILLSACANSNMYEEGVKIHELAKqkDFTIELSNTTINFYTKCGYSNIAIEECEKLKQNGLM--NVVTWTCLIQAYIQQNDIEKAIDMIIDMKKHGIYPDSATFTILLSACAKSNMYEEGVKIHELAKQTLKIFssTATINFYTKCGYSNIAIEECEKLKQKGLMnaYIQQNDVEKALNTITNMKNSGVVPNS-TTFTILLSACANSNMYEE--    
 837 UniRef50_Q5G1S8        Pentatricopeptide repeat-co...   335     2e-90  1  12.1%   1:602   232:910     MGVYSRSGKFSKAQELVdmRQRGCVPDLISFNTLINARLKSgtPNLAVELLDMVR--NSGLRPDAITYNTLLSACSRDSNLDGAVKVFEDMEAHRCQPDLWTYNAMISVYGRCGLAAEAERLFMELepDAVTYNSLLYAFArnTEKVKEVYQQMQKMGFGKDEMTYNTIIHMYGkgQLDLALQLYKDMKgsGRNPDAITYTVLIDSLGKANRTVEAAALMSEMLDVGIKPTLQTYSALICGYAKAGKREEAEDTFSCMLRSGTKPDNLAYSVMLDVLLRGNETRKAWGLYRDMISD------------GHTPSYTLYELMILGLMKENRSDDIQKTIRDMEenGYELENDTLL--SILGSYSSSGRHSEAFELLEFLKEhhCKVNNLsyETLLHCCVANEHYAEASQVFSDLRLSGCEAsvCKSMVVVYCKLGFPETAHQVVNQAETKGFHFACSPMYTDIIEAYGKQKLWQKAESVVGNLRQSGRTPDLKTWNSLMSAYAQCGCYERARAIFNTMMRDG-PSPTVESINILLHALCVDGRLEELYVVVEelMLDAFARAGNIFEVKKIYSSMKAAGYLP-TIRLYRMMIELLCK-GKRVRD-    
 838 UniRef50_Q9ZVG8        Pentatricopeptide repeat-co...   335     2e-90  1   9.5%  12:485    74:529     -----------DIRYASSIFEHVSNTNLFMFNTMIRGYSISDEPERAFSVFNQLRAKGLT--LDRFSFITTLKSCSRELCVSIGEGLHGIALRSGFMVFTDLRNALIHFYCVCGKISDARKVFDEMPQSvvTFSTLMNGYLqkKALALDLFRIMRKSEVVVNVSTLLSFLSAISdgDLSGAESAHVLCIKIGLDLDLHLITALIGMYGKTGGISSARRIFDCAIRK----DVVTWNCMIDQYAKTGLLEECVWLLRQMKYEKMKPNSSTFVGLLSSCA----YSEAAFV--------GRTVADLLEEERIALDAILGTALVDMYAKVGLLEKAVEIFNRMKDK------DVKSWTAMISGYGAHGLAREAVTLFNKMEEENCkpNEITFLVVLNACSHGGLVMEGIRCFKRMVeySFTpeHYGCVVDLLGRAGQLEEAYELIRNLP----ITSDSTAWRALLAACRVYGNADLGESVMMRLAEMG----------------------------------------------------------------------------------------------------------------------    
 839 UniRef50_D1IS18        Whole genome shotgun sequen...   335     3e-90  1  11.8%  97:589   150:618     ------------------------------------------------------------------------------------------------HNSFTYNRLLvsALRDSERLDLIRAILDDMERCGvtINILI-GIFGGGADVERCFDLVKKWDLQMNCYTYKCLLQAHLrnNSNKAFEVYVELRRRGYKLDIFAYNMLLDALAKDNKVDQVYMVFKDMKRKHCEPDEYTYTIMIRMTGKIGKADESLTLFQEMTEKGYTPNLIAYNTMIQALANNRMVDKTIFLFSKMVVEVSNKFM----------NKSIYAYLVRTLSKLGHASEAHRLFCNMWSFHDEGDRD--AYMSMLESLCDAGKTTEALDLLSKIHEKRISTDTVMYNTVLSALGKLKKTSDLHDLYEKMKQDGPSPdsYNILISSFGRAGRVEEAVKIFEEL-ENSSCKPDIISFNSLINCLGKNGDIDEAHMRFKEMREEGLSPDVVTYSTLIECFGKTDKVEMACRLFDEMLAEG----------CSPNIVTYNILLDCLERSGRTA------------EAVDLYAKLKQQGLTPDSI-TYAVL--------------    
 840 UniRef50_B9FPN0        Putative uncharacterized pr...   335     3e-90  1  10.9% 104:600    72:533     -------------------------------------------------------------------------------------------------------ALVSAYAKSRLLLDALHLFDETPRRNIYLYFSLLAAVSRSaeLALPVLRRMLSAdlRPDHFVIASVASVSARLrlRLGRQLHEHFVVSPYSGDDVVKSSLVDMYCKCGFPDDARKVFDSMGAK----NSVVWTALVSGYASNGRGEEALQLFRSMPGR----NLFAWTALISGLVKTGESVGAVELFVEMRRDDIRID-----------DAFVLSIVIGGAGQacDILSAREVFEGITFRA------VISWTTMVVGEAQHGRAEEALALYDRMVLAGAKPNKVTFVGLIYACSHAGLVQKGRQLFESMkeYGITpqHYTCYLDLLSRSGHLLEAEELMTTMP----YEPDEATWGALLSACKKYKDAEMCIRIADKLLELRPK-DSSTYILLSNVYAVNGKWDSVAKARK-------CTIGLEIRKEPVYSWIEAGR---EFRLFHAGEVPLDVREEIMGFLEEMVLEMRKRGYVPDT----SSVMHDLEENEKEH---    
 841 UniRef50_A9SWH0        Predicted protein n=1 Tax=P...   335     4e-90  1  12.1%  34:578    92:655     ---------------------------------LIEEMRDLLKQRRTDEAWQVMQLLSRSGkpDRQCTSRLVAQLSHRgcLARAQQVLTNLRKNNQVDLLDSDSLGLLasARSGAARYALNVLNLmyPSVKVWSAVVSRLGRHVDdaLELFDEVCrrSARPDTGAFNAALNACATLGFakGEDLMNSMRRCGLQPDSITFNTLIKLYAKCDQRKLLKSLPDEMVENKVMPDESTVNSLIAGYVGLGDLREAEALLRRLQDkhLRPDVRTYTTLMKGYVQKGRRSDAMRTLLAMQMDKDPRS---------SPNEVSYTTAISSCVRLGLMDEATVVLQEMLKHNIPV--NVITYNILLKGYCSARKLQKAHSVVKDMEEAGVALDVVSYNTLINGCIETGDNAAALDYFKKMRETGISPssYTTLMKAFGRNGQPKQVHLVFEEMRNDPRMKTDAVAWNVLLDSYCRNGRVADAKRMFLQMKEDRVLPTAFTYATLVKGYALGGMLGEVLVLWKEIKER----TDDEVNPLQPDEVLLNCLVDTCVRAGY------------FQKALEVVACMEEKGI-------------------------    
 842 UniRef50_Q9LWW0        Pentatricopeptide (PPR) rep...   335     4e-90  1  10.3% 162:579   121:503     -----------------------------------------------------------------------------------------------------------------------------------------------------------------DPFLSSALVSFYAKSKlvEARKVFDELTCR----DTAVYNALLSAYAKGGLVDSAEKLFEEMPDR----NVVSWTAMVSGYAQNGRHEEAVETFLEmeRAGVQPNELTVSSVLPACAAVGAMELGRKV------------EEYARGKGLLRNVYVANALLEMYSKCGSIRQAWQVFQ-----GIGRQQDLCSWNSMIMAFAVHGLWREALALFYKLRMAGVKPDGITFVGVILACTHGGLVNEGKLFFDSMeeFSLKpeHYGCMVDLLGRAGLLIESYSLIASMP----VEPDAVIWGALLGACSFHGNVELAELAMDKLI--HLEpnTANLVILSNIYASSGKWDGVAQVWKLLKEKDHKKS----------AGYSFIELDGTMHKFLVEDKSHPRFEEVYNTLNSVTMTMKLVGLE------------------------    
 843 UniRef50_Q2QNQ2        Pentatricopeptide, putative...   335     4e-90  1  12.7%  44:545    94:663     -------------------------------------------LGEPLLCVRLysRFGAHFARDGGVRRALVGALWRRGPVVLSGRLVAEVRGCGCEVSEELVCALVESWGRLGLARYAHEVFVQMPRLGLRPSTAVYnrAGAVDTAYLRFQQMPADGCRPDHFTYNSLVHGVCRRgvDEAVRLVRQMEGEGIRPNVFTYTMLVDGFCNAGRVEEAFRMLDKMKEKGVAPSEATYRTLVHglSKKDMAKEAVEFEKRMSARGYLLDSTAFGIVMSCALKCLEVSDLCELLDSFIKNGGNPGfgHMVLKGLLSSVMSYNMVIDCFVKAGAVDKAEEIVKEMQDKG--FLPNLVTFNTLISGYSKLGNVHNAKVVLKMLMEHGFMPDIITFTSLIDGLCNTHQLDDAFVCFEEMAEWGVRPntYNVLMHTLCSAGHVNKAIDLLNKMKIDG-VTPDAYSFNALILSFCRMRKVDKAEDIFNDMVRFGVVPDSYTYNSLIKALCDERRVNKAKEILFARERSGCSTSNNQSYWPIVAALAKMGQFSEA----------------------------------------------------------    
 844 UniRef50_B9T0C2        Pentatricopeptide repeat-co...   334     6e-90  1  10.0%  85:516    25:428     ------------------------------------------------------------------------------------KQLHALSVKTNTFSHPSISSCLLSLYAEpnNLEYVLSIFNKiePALVLYNILIKCYIQnsHDAITLFSQLLHE-FNPDGFTLPCVIKGCARLHakEGKQIHGLVLKLGFGLNKFVSSSLVNMYSKCKDIDSAKKVFLSMDDK----DLVSWNSLIDGYVKCGQVELGMKLFEEMPER----DLFSWTVLIDGFSKCGKVDVARELFDKMPSR----------------NLVSWNAMINGYMKAGDFVLASELFDKMPEK------NLISWNSMVAGYDLNERFKEALDLFLTMLERDFTPNHATLVSTFSAVSGLASLSTGKWMHSYMVKNGFQLdlATSLINMYSKCGNIESALSVFQLITKKKLGH-----WTAIIVGLGMHGMADNALEFFHEMCRTGMRPHAITFIGVLNACSHAGLVEDGRKYFDMM---------------------------------------------------------------------------------------    
 845 UniRef50_Q9FGR2        Pentatricopeptide repeat-co...   334     7e-90  1  11.6%   3:509    54:510     --AAKLVteCGKVLDARKVFDEMPKRDISGCVVMIGACARNGYYQESLDFFREM--YKDGLKLDAFIVPSLLKASRNLLDREFGKMIHCLVLKFSYESDAFIVSSLIDMYSKFGEVGNARKVFSDLGEQ--------------------------------DLVVFNAMISGYANNsaDEALNLVKDMKLLGIKPDVITWNALISGFSHMRNEEKVSEILELMCLDGYKPDVVSWTSIISGLVHNFQNEKAFDAFKQMLTHGLYPNSATIITLLPACT----------TLAYM--KHGKEIHGYSVVTGLEDHGFVRSALLDMYGKCGFISEAMILFRKTPKK------TTVTFNSMIFCYANHGLADKAVELFDQMEATGEKLDHLTFTAILTACSHAGLTDLGQNLFLLMqkYRIVpeHYACMVDLLGRAGKLVEAYEMIKAMR----MEPDLFVWGALLAACRNHGNMELARIAAKHLAE--LEpnSGNGLLLTSLYANAGSWESV----------------------------------------------------------------------------------------------    
 846 UniRef50_O81028        Pentatricopeptide repeat-co...   334     7e-90  1  12.6%  33:597   151:758     --------------------------------ALVKAYVSLGMFDEATDVL--FQSKRLDCVVDIKACNFLMNRMTEFGKIGMLMTLFKQLKQLGLCANEYTYAIVvaLCRKGNLEEAAMLLIEN--ESVFGYKTFINGLcgETEKAVALILELIDRKYLAgrAVLGMVVRGFceMKMKAAESVIIEMEEIGFGLDVYACLAVIDRYCKNMNLPEALGFLDKMLGKGLKVNCVIVSLILQCYCKMDMCLEALEKFKEFRDMNIFLDRVCYNVAFDALSKLGRVEEAFELLQEMKDR------------GIVPDVINYTTLIDGYCLQGKVVDALDLIDEMI--GNGMSPDLITYNVLVSGLARNGHEEEVLEIYERMKAEGPKPNAVTNSVIIEGLCFARKVKEAEDFFSSLEQKC--PENKASFVKGYCEAGllEKAHDVLKKMSA-YRVEPGRSMCGKMIGAFCKLNNVREAQVLFDTMVERGLIPDLFTYTIMIHTYCRLNELQKAESLFEDMKQRGIKPDVVT-YTVLLDRYLKlgEVgeVLREfyTVLIDRQCKMNNLEQAAELFDRMIDSGLEPDMVA-YTTLISSYFRKG------    
 847 UniRef50_B9SJB1        Pentatricopeptide repeat-co...   334     7e-90  1  11.6%  34:508    49:478     ---------------------------------LARSYMHCGkyFDALKVYSEMLRSGYCF-PDNYTYPVVIKACSELGLIELGKMIHGQTVK-----------------------EAAERIFDVmeRSVISWNIMINGYFKNGckALMVFNQMVDLGVEIDCASVVSVLPACGHLklEMGRRVHATVEDKDLGNKIAARNALIDMYAKCGSMAEARLVFDGMDER----DVISWTSMINGYILNGDEKSALTLFRTMQMERIIPSSVTIASVLSACINLKD---------------GRCFHGWTMRQNLDSEIIVETSLIDMYAKCNRVDLSYGVLRR--TSGERTAP----WNAMLSGCIHNDLATEAITLFKQMLLKKVEPDGATFNSLLPAYAILADLLPAKNIHCYLKKTGFllEVATCLIDIYSKCGSLESAHRIFNAIP---IGVRDIFAWSVIISGYGMHGHGETAVSLFRQMVQSGVRPNEVTFTSILHACSHAGLVDE-----------------------------------------------------------------------------------------------    
 848 UniRef50_Q9C6T2        Pentatricopeptide repeat-co...   333     8e-90  1  10.0%  72:601    34:534     -----------------------------------------------------------------------LYLLKRCHNIDEFKQVHARFIKLSLFYSSSFSavLAKCAHSgnSMNYAASIFRGIdpCTFDFNTMIRGYVnsFEEALCFYNEMMQRGNEPDNFTYPCLLKACTRLkiREGKQIHGQVFKLGLEADVFVQNSLINMYGRCGEMELSSAVFEKLESK----TAASWSSMVSARAGMGMWSECLLLFRGMCSenLKAEESGMVSALLACANTGALNLGMSI------------HGFLLRNISELNIIVQTSLVDMYVKCGCLDKALHIFQKMEKR------NNLTYSAMISGLALHGEGESALRMFSKMIKEGLEPDHVVYVSVLNACSHSGLVKEGRRVFAEMLKEgvEpeHYGCLVDLLGRAGLLEEALETIQSIP----IEKNDVIWRTFLSQCRVRQNIELGQIAAQELLKLSSH-NPGDYLLISNLYSQGQMWDDVARTRTEIAIKGLKQTPGFSI----------VELKGKTHRFVSQDRSHPKCKEIYKMLHQMEWQLKFEGYSPD----LTQILLNVDEEEKKER--    
 849 UniRef50_Q9ZUT4        Pentatricopeptide repeat-co...   333     9e-90  1  11.7% 155:574   112:498     ----------------------------------------------------------------------------------------------------------------------------------------------------------KRDGWSFDAYGLSSAVRSCGLNrfRTGSGFHCLALKGGFISDVYLGSSLVVLYRDSGEVENAYKVFEEMPER----NVVSWTAMISGFAQEWRVDICLKLYSKMRKSTSDPNDYTFTALLSACTGSGALGQGRSV------------HCQTLHMGLKSYLHISNSLISMYCKCGDLKDAFRIFDQFSNK------DVVSWNSMIAGYAQHGLAMQAIELFemMPKSGTKPDAITYLGVLSSCRHAGLVKEGRKFFNLMAEHGLKpnHYSCLVDLLGRFGLLQEALELIENMPMK----PNSVIWGSLLFSCRVHGDVWTGIRAAEE--RLMLEPdaATHVQLANLYASVGYWKEAATVRKLMKDKGLKTNP----------GCSWIEINNYVFMFKAEDGSNCRMLEIVHVLHCLIDHME-----------------------------    
 850 UniRef50_C5X339        Putative uncharacterized pr...   333     1e-89  1  11.1% 143:582    58:499     ----------------------------------------------------------------------------------------------------------------------------------------------NPHLALRLFDHLLRSGADPDPAAYELALASCARGKDRaaAQLHAHAAKRGFLAsrRVRCRLVHAYAVCGMLPHARRVFDG----GTDNDMVAWNCLLRGYAQEgdADLLRDFFARMPSR----DSVSWNTVLSLCVVNGEYDEAIAVFREMLASqgLWAHAYVIRKGVEVEEKLSSALINMYSKCGFLEGAVYVFD-----NVRGKRSLDTWNAMLAGFTANGCSGRALALFTRMETTGLMPNKITFNCILNACSHGGLVEEGMRCFQRMSryGIEPdhYGCMVDLFCRAGMFEKAEEIIQIMP----MEPDASMLKALLGACRTHKNLELGKKAGHRLIEA--APNDHayVLLSNIYALDGNWGRVHKVRKLMLDRGVLKTPGS----------SSVELNGVIHEFISGDKSHSRKRDIYKMLDEICQQLKSAGYTPDT---------------------    
 851 UniRef50_Q8H2U1        Os07g0662700 protein n=2 Ta...   333     1e-89  1   9.5% 118:579    32:454     ---------------------------------------------------------------------------------------------------------------------HALLVKLGLQPYARVHNALiyAASGLVDDARRVFDGMSHR----DTVSFNSMIHAHasGDVVSARRLFERVP----SPTPVTWTSMVAGLCRAGDVAAARRLFEEMPVR----DLVSWNAMMSGLAGNRRPVEALCLFRRMMAEGFAPNRGTVLSALAACAGAGALETGKWIHAFVERKRLFRWW----------DEFLGTALLDMYAKCGAVELALDVFTKLRSR------NTCTWNAMINGLAMNGYSAKALDMFRKMedRTVVPDEVTFVGVLLACSHGGFVDVGREHFHMikKYGIRleHYACMVDLLARSGHLQEAHKIIAGMPMK----PDAVVWRALLGGCRLHKDVKMAETAISEM-EATCSGDHV---LLSNLYAAVGRWSGVEDVRRTMRSKGIEKIP----------GCSSIEINDSIHEFVSGDKSHPSYNDIHAKLAEISARMQQQGYV------------------------    
 852 UniRef50_Q2R8W9        Os11g0213500 protein n=3 Ta...   333     1e-89  1   9.8%  71:559    12:469     ----------------------------------------------------------------------LLRRCAAAGAVRPGEQAHARAVVGGWLPDATLETDLVLMYCRCGERRRARRVFDGMrpSMHAYNVLLAAS--PPRDAMEVFSRLLASGLRPDGYSVPAVVRACAEldAVLGGVIHGFAVRLGLMGNVVVAAALLDMYAKAGFLDDAVRVFDEMTER----DAVVWNCMVAGYARAGRAVETFEIFSRAQVEAV--NMvaVPSVLNICAKEGELMKGREIHGRMVRCLA-----------FDSDIAVGNALINMYAKCGRVNVSQAVFSGMQQR------DVVSWSTMIHSYSIHGKGEQALKVYMEMLSEGVKPNWITFTSVLSSCSHSGLVTEGRKIFESMTkhGVHpeHYACMVDLLGRAGAIEEAVGLIRKMP----MEPCASVWGALLSACAMHNNVDVGEIAAFRLFELE-EGSASNYVTLCGIYDAVGQSDGVAGLRSRMRELGMVKTP----------GCSRIDVKGRAHAFYQGSIPRYLR--------------------------------------------    
 853 UniRef50_D1HZG5        Whole genome shotgun sequen...   333     1e-89  1  10.1%  18:601   301:943     -----------------EMKNLGFVPEEVTYSLLISLSSKTGNRDEAIKLYEDMRYRRIV--PSNYTCASLLTLYYKNGDYSRAVSLFSEMEKNKIVADEVIYGLLIRIYGKLGLYEDAEKTFKETEQLGLLTNEKTYIAmgNFEKALTIMELMRSRNIWFSRFSYIVLLQCYvkEDLASAEATFQALSKTGL-PDAGSCNDMLNLYIKLDLLEKAKDFIFQIRKDPVEFDMELCKTVMKVYCKKGMLRDAKQLIQEMGTNGLFKDSetLSLILKmfTREGDISKAQNLNDQLVKLGRGAEDASIAseGCTSGKLIYISMIDAYAKCGKAEEAYHLYEEVTGKG--IELGVVSISKVVHALANYGKHQEAENVIRRSFEDGLELDTVAYNTFINAMLGAGRLHFANSIYDRMVSLGVAPstYNTMISVYGRGRKLDKAVEMFNKARCSGvvSLDEKTYTNLISYYGKAGKSHEASLLFREMQEEGIKPGKVSYNIMINVYATAGLHHEAQELFQAMLRDGCSPDSLTYL-ALIRAYTQSFKFLEAEETIMSmlLSAFAKAGFTEEAERVYHTLLSAGLSPD-VACYRTMLRGYLDYGCVEK--    
 854 UniRef50_Q9FGL1        Putative pentatricopeptide ...   333     1e-89  1   9.8%  23:521    16:558     ----------------------GSTASSNSWSTIVPALARFGSIGVLRAAVEL---INDGEKPDASPLVHLLRVSGNYGYVSLCRQLHGYVTKHGFVSNTRLSNSLMRFYKTSDSLEDAHKVFDEMppDVISWNSLVSGYVQSGreGICLFLELHRSDVFPNEFSFTAALAACARLHlpLGACIHSKLVKLGlkGNVVVGNCLIDMYGKCGFMDDAVLVFQHMEEKDTvpDTVTYNELIDAFVKSGDFNNAFQVLSDMPN----PNSSSWNTILTGYVNSEKSGEATEFFTKMHSSGVrlIHACAHKLGLDSRVVVASALIDMYSKCGMLKHAELMFWTMPRK------NLIVWNEMISGYARNGDSIEAIKLFNQLKQErlKPDRFTFLNLLAVCSHCEvmEVMLGYFEMMieYRIKpeHCCSLIRAMGQRGEVWQAKQVIQE----FGFGYDGVAWRALLGACSARKDLKAAKTVAAKMIELgaDKDEYLYIVMSNLYAYHERWREVGQIRKIMRESGV----------------------------------------------------------------------------------    
 855 UniRef50_B9SI50        Pentatricopeptide repeat-co...   333     1e-89  1   9.8%  24:511    11:472     -----------------------VKRPLYLWNLMIRSSTKAGLFFQALDIYSSMLQSGVHG--NGFTFPLLLKACSNTNSIRDGTKIHSHLIQLGFQHVFVMTTLLDMYSKCYDLASSRKVFDEMPMRSTVSWNSIISAYCRfvDEAISMLQKMRLIGFVPTSTTFLCFLPICL---LQHGLSIQCCAFKFGLlsDIPLTNALLNMYVKHGQVHEARTLFDMMHEK----SLISWTTVIGGYVDFGNVREAFSLFNQMRIS-MRLDFIVFITLISGCAREGNLLLASSVHSLLVKYGCDDKDPID------------NLLVTMYSKCGDLISAQRAFDIAREK------SLYLWTSMIAAYTHLGYPVQALRLFNTLLGTAIKPNEATLATILSACADLGSLSMGEEIEEYILANGLHSsvQTSLIHMFCRCGSLEKAKAVFERLATK-----DLAAWSSMINGYAIHGMAEEAFSLFHKMQteGIKPDAVIYTSILLACSHSGLIEDVFN--------------------------------------------------------------------------------------------    
 856 UniRef50_Q1PFQ9        Pentatricopeptide repeat-co...   333     1e-89  1  13.0%  72:524    41:511     -----------------------------------------------------------------------LQEHINSPAPKAGKKIHADIIKTGFQPDLNISIKLLILHLKCGCLSYARQVFDElkPTLSAYNYMISGYLKHgkELLLLVQRMSYSGEKADGYTLSMVLKASNSRgrSLCRLVHARIIKCDVELDDVLITALVDTYVKSGKLESARTVFETMKDE----NVVCCTSMISGYMNQGFVEDAEEIFnsMQRAGFHPNISTFASVIGACSVLTSHEVGQQV------------HAQIMKSGVYTHIKMGSSLLDMYAKCGGINDARRVFDQMQEK------NVFSWTSMIDGYGKNGNPEEALELFTRMKEFRIEPNYVTFLGALSACSHSGLVDKGYEIFESMQRdkPKMEHYACIVDLMGRAGDLNKAFEFARAMPER----PDSDIWAALLSSCNLHGNVELASIAASELFKLNADKRPGAYLALSNVYASNDKWDNVSKIREVMKRRRISKT-------------------------------------------------------------------------------    
 857 UniRef50_A9T9D8        Predicted protein n=1 Tax=P...   333     2e-89  1  14.1% 143:517    25:401     ----------------------------------------------------------------------------------------------------------------------------------------------DWRCTLEVFSWSKKQsyCPNSRLYTRLIGFLGrgRVHNATLLFQEMLLEKCQADQYTYTALVNAYGKAKMFEEALAVFSHMKESncQPNTVTCNALIDALVKGGLYDQAIQVFFDMreHGCEPNVITYNVLIDALCKEGLLDIGMKVLQYMRDGNTDQS--------VQPNSATYNTLINACGKGGLYEKAEELVDLMVEHG--VQPDHITYTALIDAYGKAGLWENAENTFKGMKGTNVSVDVMAYTAMIDAYGREGLYQKAEEMFKMMQHSGLRPntYLSLMEAYGKAGLPEESRNVFNIMRESGY-EGNVLIYSSLIDAYGKAGNYLEAARMLDMMRRAGCQPNLITYSAILSSCCKSNCWVEAQVLLRRLQ--------------------------------------------------------------------------------------    
 858 UniRef50_Q5N7W0        Os01g0913400 protein n=4 Ta...   332     2e-89  1   9.4%  10:574   103:647     ---------CSARARPRQMFDEMPVRDLVACSAAIYRHAKSGLFGEAVRLFVGM--MRVGVSPNSFTLVGALIAAAGMGNLVLAECIHGWAMKSLLESNPFVRTALLDSYAKCGRPTKAWALFGEMrpGIVTWNALISGLVHnfEEALLVFKRLlsFGPVHNVVTMISIAQASacGDLGLCESAHAYSVKIGLDSDVSVTNSILGMYLSFGSLAIGREIFKKIAVN----DVVSWTMMMGFLLEEAQAIEVIHMFVQMRLSGIVPDRVALVTVAQACAHLGD------------GRIGKLVHNEIVICGFSGELPAVNSLITMYSKCEDLSSARLLFDGTMEK------SLVSWTAMVSAYIENGYALEGMYLFAKMRHEgfMIDSVTLVTLLLACYEVAKFELCIQLHAYCYKSGlkP-VLNTLIAVYGKCGYATLAHKVFDEMISR-----NAVSWNTMILSYGVNGQGEKAVSLFNEMEKSSEDQDSVTYLNTLLACSHSGLVDDGMLVFRRMVNDKGIIPCPEHVGCIVDMLARAGRLEEA--AGVASLTHNKLGANAWKALMglHGDMK-----------------------------    
 859 UniRef50_Q9SNP3        Putative uncharacterized pr...   332     2e-89  1   9.2%   9:572    47:699     --------RAGRLEAAREVFDGMPRRDVVSWNTIMAVQARAGSHGRAVGAFLEMRRQGF--RPDHTSFSTALSACARLEALEMGRCVHGLVFKSCSSGNVFVGASLITMYANCGVVsyARKAFDRMprNVVSWTAMIKGhaHEVDMAFQLFKLMpmNGEEVTNVILVKIVNAFAGMkiRGGRCIHGLAVKSGFAYDLVLEASLVLMYCKSLDITEARLEFDKMEGNHVG----SWNAMLCGYIYSDKIDEARKLFDSMNNR----DKISWNSMINGYINDGRIADATELYSKMTEkgMLDKARDMFYNMPQIDVMSCTTLLFGYVKGGHMDDALDLFHMMQKrnESPTRDSVTWSCLVAGLATNGLIHEALQFYKKMLLSNIRPSESVVSSLISCLSNYSMMVHGQQFHATTIKIGLDSHLlnSLISLYCKCGEMIIAQSIFDLMAKR-----DKVTWNTIIHGYALNNLGQNAVEMFESMTKAQVDPDDITFLGVLSACNHMSLLEEAKYFFNAMTCTYGILPNIMHYACMVDLFCRKCMIKEAEG----LVKSMPFEPDSAIWTSLLSGC-------------------------------    
 860 UniRef50_B9FLT7        Putative uncharacterized pr...   332     3e-89  1  16.6% 139:517    13:380     ------------------------------------------------------------------------------------------------------------------------------------------CAAGDVRSALAMLARGTKSgaALDVTACTALVNGCCKgdVAEARRVFDEMPLLGLAPNEVTYTALMHGYFTHGQREKGFALFEEMRRGGVEPNLYTYNCLIGEWCRTGEFERARSLFDEMPVRGIVRNVVSYNTLIAGLCRHGKLWDAAKLLDMMRTE------------GTRPSIITFNLLVDGYGKAGKMSNALPFFNQMKAAG--FQPSAVTYNMLIAGFCRARDMTRANRALSDMKERGLEPTKVTYTILIDSFARENHMGKAFEILAGMEKAGLEVdtYGVLVRALCMEGNMKDARKLFQSMGEKG-VEPSNVIYDMMIYGYGREGSSYKALKLIMEMRQKGLIPNSASYGLTIRVLCKDDKCQEAEALLDDME--------------------------------------------------------------------------------------    
 861 UniRef50_Q9FG85        Pentatricopeptide repeat-co...   332     3e-89  1  10.4%  72:525    13:454     -----------------------------------------------------------------------LNLISKCKSLQNLKQIHAQIITIGLSHHTYPLSKLLHLSSTVCLSYALSILRQIPNPSVFLYNTLISSIVSNhhLAFSLYDQILSSrvRPNEFTYPSLFKASGFDahRHGRALHAHVLKfpVNHDRFVQAALVGFYANCGKLREARSLFERIRE----PDLATWNTLLAAYANSEEieEVLLLFMRMQ---VRPNELSLVALIKSCANLGEFVRGVW------------AHVYVLKNNLTLNQFVGTSLIDLYSKCGCLSFARKVFDEMSQR------DVSCYNAMIRGLAVHGFGQEGIELYKSLISQGLVPDSATFVVTISACSHSGLVDEGLQIFNSMKayGIEpeHYGCLVDLLGRSGRLEEAEECIKKMPVK----PNATLWRSFLGSSQTHGDFERGEIALKHLLGLEFE-NSGNYVLLSNIYAGVNRWTDVEKTRELMKDHRVNKSP------------------------------------------------------------------------------    
 862 UniRef50_D2VKI6        Predicted protein n=3 Tax=N...   332     3e-89  1  10.3%   5:599  569:1166     ----SIINFYTKCgeCEKLKLKGLMNNVAVWNCLIQAHIQNNEVQKAIEIITEMQQFGL--NPDSTTYTIILKACTDSNLFEEGKFIHEQAIENVKDCSTELLTSIISFYTKCGHPeeCEKLqkGLMNNIAVWNCLIQAHIQnvQKSIKIITEMQDYGIEPNSTTFNILLTACADSKleEGKAIHKLATGTLKDYSAELLASIVNFYTKCGHPEVSIEECEQLKSKGLMNNVAIWNCLIQAHIQng-LNKAMVVFNEMKEH-IEPDKITYLILLAGCGEYVARKMGLDIIDELKLKG-MDVHEEIV-----------SVMIYFYSRIAYQSTAQQLFEKLVEESAQKRLSIITYNNMLSCYAQHGKGTQAMNLFNEMVDKNVEMNEQTFTIMLNTFSHCGMIDEANQLYGEMkkYNHNPSVicVMVDVFARSGRFHEAEKIINE-------RMNYSLWSALLAGCRKFNNVELGEKVFKILleKKAPP----YILLSNIYTYYGLNEKADNIRQLMIENGIKKIPGIVYQ----------EEDGKSYQFVSGSKKYERFKEIEKMREEIYRKIIQAGFEPDtvTNSKLllDSLCRHAEK----    
 863 UniRef50_Q9FJY9        Pentatricopeptide repeat-co...   332     3e-89  1   7.9%  13:529    34:530     ------------ATHADHLFDELPQRDLSSLNSQLSSHLRSGNPNDTLALFLQIH--RASPDLSSHTFTPVLGACSLLSYPETGRQVHALMIKQGAETGTIstALIDMYSKYGHLVDSVRVFESVEEKDLVSWNALLSGFLRNGkeALGVFAAMYRERVEISEFTLSSVVKTCASLklQQGKQVHAMVVVTGRDL-VVLGTAMISFYSSVGLINEAMKVYNSL---NVHTDEVMLNSLISGCIRNRNYKEAFLLMSR-----QRPNVRVLSSSLAGCSDNSDLWIGKQI------------HCVALRNGFVSDSKLCNGLMDMYGKCGQIVQARTIFRAIPSK------SVVSWTSMIDAYAVNGDGVKALEIFREMCEEgvLPNSVTFLVVISACAHAGLVKEGKECFGMMkkYRLVpeHYVCFIDILSKAGETEEIWRLVERMMENDNQSIPCAIWVAVLSACSLNMDLTRGEYVARrmEETGPE-NASIYVLVSNFYAAMGKWDVVEELRGKLKNKGLVKTAGHSL--------------------------------------------------------------------------    
 864 UniRef50_C6JS32        Putative uncharacterized pr...   331     3e-89  1  12.2%  77:525    27:464     ----------------------------------------------------------------------------RCATPSALARVHAAMIVSGRldAFAASRLLTAYAALSPDPAAAALALLscAPNSFMLNTTLRVLASSPdaAFPFFSRLRVTglAPGRHTFPFLLKAAARLPLPltEQLHALAVRHGVLLDAYVANGLVRGYSVAGRLQAARRVFDELPER----NPAVYTTMVSAYAQNGRHEDAMAAFDEMVRVGFEPCGATLASVLSACArsGGLEMGCHVHDLMTA------------WGVPVSTILGTALIDMYVKNGAIKEAFAVLDGLPERHV-----TGAWNALISGLAHHGHRERALNLFRQMQQEGVPPNATTLVGALSACSHPGLLDEARRLFSSMEKdgITpqHYGCMVDLLGRAGLLSEAEDLIRGMS----CEADTVIWGALLTACKNHSGIEIAERAAVEMLKLDPS-NHGVYVVLSNLYAEAGRWQDVDKLRKEMKGARLSKIP------------------------------------------------------------------------------    
 865 UniRef50_B9FHI9        Putative uncharacterized pr...   331     3e-89  1  11.2%  49:484     9:490     ------------------------------------------------SVLNFLRHVSFPPDPR-----LLPSALKSCSALRLARALHAAAAVAGVSRDAFVASSLLHAYLRFGATADARSVLDGmhRTVVGWSALIAAHagDAEGAWGLLERMRSDGVEPNVITWNGLVSGLNrgDVAVGEQLHGYVVKAGCRLDACVATALIDMYGKCGRADEIVRVFDESSHM----DVASCNALVAGLSRNAQVSEALRLFREFVGRGIELNVVSWTSIVACCVQNGRDLEAVDLFREMQSEggRSAHCFSLRKGFHHDIYVGSALVDMYAKCGRVRDARMIFEAMPYR------NVVSWNAMIGGYAMHGEAENAVRLFRSMQSSKEKPDLVTFTCVLGACSQAGWTEEGRSYFNEMqkHGISpeHYSCMVTLLGRAGKLDDAYDIINQMP----FEPDGCIWGSLLGPCRVHGNVVLAEVAAENLFQL-----------------------------------------------------------------------------------------------------------------------    
 866 UniRef50_D1H1K9        Whole genome shotgun sequen...   331     4e-89  1  10.0%  71:528   177:632     ----------------------------------------------------------------------VLGLMCEEDLVMGCLYFFEWmlQEPSLVTARACSLLFPMLGRAGMGDDLMVLLRNLprDVRIYNSAISGlcGRYDDAWKVYDEMETNNIRPDHVTCSIMITVMRKDGhkDAWEFFQRMNRKGVKWSLEVLGALIKSFCDEGLKNEALIIQSEMEKKGISSNAIVYNTLMDAYSKSNRVEEAEGLFGEMKAKGVMPTSATYNILMDAYSR------------RMQPEIIENLLLEMQDMGLEPNVKSYTCLISAYGRQKKmdMAADAFLRMKK--VGIKPTSHSYTALIHAYSVGGWHEKAYTAFENMKREGIKPSIETYTALLDAFRRAGDTQTLMKIWKLMLSDKIerVTFNILLDGFAKQGHYMEARDVIFEFGKIG-FQPTVMTYNMLMNAYARGGQHSRLPQLLKEMTSLNLKPDSITYSTMIYAYVRVRDFKRAFFYHKQMVKSGQVPDPQSY---------------------------------------------------------------------------    
 867 UniRef50_A9RN06        Predicted protein n=1 Tax=P...   331     4e-89  1  11.9%  16:527   176:702     ---------------AATLREVGDSLEIMDLNSVLRRFAKMKQWRDACAVFEYMKENNKLSPP---TYTSYFSVLGKSGQSRKAIRVFNELPLDsrKNVFVCNSILSTMIYNGK-IDKAFRLFEQLKKEGLQPDTVTYSTVIAGckALALIEDIKASGLTQDAVIYGAMMNVCASQGLdeAQATFQEMCDAGFTPNLFHYSSLLNAYASRGKYQQAERVLEAIRAAGLTPNLVVYNSLMNAYANCQMPSKAREVFDRIEAEGLQRDMVSFCSMMDAYAKSDMLKEAQETFDSMIAQGMKPVYMLMQLLDLSGSLCsySILISAYAKLGEMDGVLKLVDQI-EKGQNGECDILLFNTLLKVYCNAGYMDGVTAVLKKMDAEGCGSDRGTFNILIGYFAKEDLGDLALKTLEDMETRGHRPntYNALFQGYVKNGKMEEMFEALEAMKQRGI-TPTNLSLQTIMLGCIEHNRIDVVLAYFPYLKATGIALSYSVMERLVEALVETGQMDVVKDALHLMQTTGQQPSQKT----------------------------------------------------------------------------    
 868 UniRef50_C5YAN5        Putative uncharacterized pr...   331     6e-89  1  11.2%   6:477    46:501     -----LVRRATELGdaDLLFSSFRepGAVayNAMIRGCAYHGPHEGALELFAEmrARGEGLAPDSFTYPHVVDACARLRMWRAAEAVHCRVLKEGLAAVPAVSSSLLAFYVARGSLGDARRVFDGFrkSIGLSNRMVSVYVRAGdsAREVFDAMETR----DVVSWNAMLTAYVraDIVAAKELFAAMPVK----NVISWTTMIRALSDAGDFAGMRGLFDRMPER----NLVSWNCILSSYTRHGRFWQALQMFPRMLLEGLNPDSFTVVSVLSACENLRKLRLGRWI------------HANLVTPALQVHAEVGTALVEMYAMCGDVARAFVVFLKMYSK------DVFSWNVMIRALAVHRRADDALRLFGVMRKQGFGPNHFTFMGVLLACRYGSLVDEGRRMFDMmkEYCIPPshYTCLIDLLSCSGHVDEAVTVLQGMP----CRPDSDVWRALLGGCRIEADLGSAKEA------------------------------------------------------------------------------------------------------------------------------    
 869 UniRef50_D1H5I6        Whole genome shotgun sequen...   331     6e-89  1   7.8%  81:580     1:455     --------------------------------------------------------------------------------MNQIKQTHAHLITTGLILHPITANKLLkiASSFGSLSYAHQLFDQikPDVFIYNTMIKAHAvsSHNSMRIFLSMVrsGFLPNRYTFVFVFKACGNggVLEGEQIRVHAIKIGLESNLFVTNAMIRMYANWGLVDEARRVFDWSLDQ----DLYSWNIMIGGYVGSGEIGRAKEMFDEMSER----DVVSWTTIIAGYVQVGCFKEALDLFHEMLqdQGRWIHVYIDKSEIKMNERLLASLLDMYAKCGEIDFAAKVFHD----EYGLKLKVWPWNAMIGGYAMHGKSKEAIDLFEQMKVEKVSPNKVTFVALLNACSHGKLVEEGR------------------------------EAEETVFNMP----MAPDATIWGALLGACRIHKDIERGQRIGKIIKELDSD-HIGCHVLLANLYSASGQWDEAKAVRQKIEVSGRKKTP----------GCSSIELNGVFHQFLVGDRSHPQTKQLYLFLDEMTTKLKNAGYVP-----------------------    
 870 UniRef50_Q6K957        Pentatricopeptide (PPR) rep...   330     8e-89  1  11.8%   3:576   166:710     --ANSLLSLYATLGDFAslFGEMPDRDVASWTSMIGACLGSGYADQALRLFREMLADGA-LQPDGVVAVVVLRACAMLEDVRAGASVHAVATRRGLQGDLFVDNSLVDMYAKCLDLRSARKVFDLivKNVVSWNTMLSGLVHAGSYPEALHLLALQIGVVGDETTLAVLLQLCKKKRlqAARSVHGAAIRRRL-LSMALLNALLDAYGKCGLVEDVLRLFQGMRER----NVITWSTVIAACAHNARPHAAMACFVAMLVTGERPNSITVLSLVEACGSC----------AEMWAS--RRAHGVAVRSGLGFELAVGNALVHMYGKCGELGASARVFDTMPVK------DVLTWNSMIGALGMNGRARDALALLHRMEAEgvRPNGVTMLAALWACAHGGLVEEGIGCLESMARQSLQpeHVSCVVDMLARAGDLDGAAEIVRR--SSGGGSP--AAWSALLSACRRRGDGGgaARVLE------LEPGKSAGYLMSMGMGLGKGW--AAGMRWAMREKGVKVESGHSVVQHAGGSERFVWWDGA----------HPRRAEVYAMLHLLHRHMKHH---------------------------    
 871 UniRef50_D1IJ43        Whole genome shotgun sequen...   330     1e-88  1   9.7% 136:601     1:437     ---------------------------------------------------------------------------------------------------------------------------------------MITCYTRNGekARNLFNLLP---YKWNPVCCNAMVAGYAKNrfDEARRLFDAMPAK----DLVSWNSMLTGYTRNGEMRLGLQFFEEMAER----DVVSWNLMVDGFVEVGDLNSSWEFFEKIPN----PNTVSWVTMLCGFARFGKIAEARRLFDQMPIR----------------NVVAWNAMIAAYVQNCHVDEAISLFMEMPEK------NSISWTTVINGYVRMGKLDEARQLLNQMPYRN-KPDQSTFACGLSSCAHLAALQVGKQLHQLVMKSGYATDLfnALITMYAKCGSISSAELLFKD-----IDHFDVVSWNSLIAAYALNGNGREALKLFHKMEVEGVAPDEVTFVGILSACSHVGLIDQGLKLFKCMVQAYNIEPLAEHYACMVDLLGRAGRLEEAFQLVRGMKInaCRIHGNLELAKFAAEKLLE--FEP-HKTSNYVLLSNmaEAGRWDE--    
 872 UniRef50_C5XWQ9        Putative uncharacterized pr...   330     1e-88  1  14.1%  88:575    32:509     ---------------------------------------------------------------------------------------FSHLLADRFLPNQLLRSLIPAhlGALRLFPRLRRIFPDFRPNNYTFSFLLKAAADsgAHAisLHALAVVLAWDAHAYVANGLIHAYATHGvpSARRLFEDALASR-AADVCSWTSLLTACAKAGQVEEARALFDGMPRR----NDVAWSAMLSAYVAAGSFDDAVRLFEDMLRSGVRPNRAAVVGVLAACGALGALDQGRWVHALLVS------GTHGGDGATAMDGVVATALVDMYAKCGSLDTARQVF---AAAAPRSQRDVFAYTAMISGLSDHGRCGEAIDLFGQMQAEGVRPNEVTFICVLTACGRAGLVGRAKEVFRSMAahGMEpeHYGCLVDVLGRAGLLAEAMETVRSMTMR----PDAYVLGALLNACAAHGDVEAGEQVVRWLAELGLD-HSGVHVQLSNMYAGWSKWEEVLKVRRTMDQRKVAKVP----------GCSMLEVDGVASEFVAGDRSHPRMQEIMSAIRDLHGQLRQ----------------------------    
 873 UniRef50_A3KPF8        Pentatricopeptide repeat-co...   330     1e-88  1  11.0%  60:522   101:561     -----------------------------------------------------------GHKPNVAHSTQLLYDLCKANRLKKAIRVIELMVSSGIIPDASAYTYLVNQLCKRGNVGYAMQLVEKMesNTVTYNALVRGLcgSLNQSLQFVERLMQKGLAPNAFTYSFLlaAYKERGTDEAVKLLDEIIVKGGEPNLVSYNVLLTGFCKEGRTDDAMALFRELPAKGFKANVVSYNILLRCLCCDGRWEEANSLLAEMDGGDRAPSVVTYNILINSLAFHGRTEQALQVLKEMSK----------GNHQFRVTATSYNPVIARLCKEGKVDLVVKCLDEMIYR--RCKPNEGTYNAIGSLCEHNSKVQEAFYIIQSLSNKQKCCTHDFYKSVITSLCRKGNTFAAFQLLYEMTRCGFDPdtYSALIRGLCLEGMFTGAMEVLSIMEESENCKPTVDNFNAMILGLCKIRRTDLAMEVFEMMVEKKRMPNETTYAILVEGIAHEDELELAKEVLDELRLRKVI---------------------------------------------------------------------------------    
 874 UniRef50_D1I4Q2        Whole genome shotgun sequen...   329     1e-88  1   9.0%   3:602    57:684     --YLRMYVNAGAMQEARKLFDEMPERSLVSWTIVMSGYARHGPASEVLMMFWDMlsGGGLLRPDSFVFAVVLRACGMVECLSYGRGVHGLVVKQSSVVDSFVENALVSMYGSCGALEDAAVVFGGikPDLVGWSSILSGYVKNgeEGLRIFCDMVSGGIEPDAFAFSMVLGACTNLewDFGTQAHCYIIKMGFDSCLYLENSLMDFYAKCGDLEGMRRVFSHMSEK----NLVSWNTFINGYVHNFHYLEALRIFQILMEEVSQCDDFSLLSILKAVSGLGHLDHGKEIHGYILRAGIemYIGCIDHESLYpdEFIMTSLLKWCSLESSLETAKRVFTRVE------QPDTAPWSALISGHSWNGCFAEALKLFRKMQFDGIKANEFTFTSVILACLALENLRKGKELHCKILRSGYESnvVNTLINLYSELWQHKQALKLCSMIPD------SEISWNFLIRACLGAEDYEIIHKLLWRIQVSHGNLDPVSACDIFASCSSPVLLNVGTQAHAYMTKRGLISHP-TISNSLIQMYSACGKFDEAVQAFNLmiLSARVEHGHPSEALNLISQMRWKN-KPADQSTFRSVLNACAQMGLVDEA-    
 875 UniRef50_Q75HY8        Os05g0583900 protein n=5 Ta...   329     2e-88  1  16.6% 139:517    13:380     ------------------------------------------------------------------------------------------------------------------------------------------CAAGDVRSALAMLARGTKSgaALDVTACTALVNGCCKgdVAEARRVFDEMPLLGLAPNEVTYTALMHGYFTHGQREKGFALFEEMRRGGVEPNLYTYNCLIGEWCRTGEFERARSLFDEMPVRGIVRNVVSYNTLIAGLCRHGKLWDAAKLLDMMRTE------------GTRPSIITFNLLVDGYGKAGKMSNALPFFNQMKAAG--FQPSAVTYNMLIAGFCRARDMTRANRALSDMKERGLEPTKVTYTILIDSFARENHMGKAFEILAGMEKAGLEVdtYGVLVRALCMEGNMKDARKLFQSMGEKG-VEPSNVIYDMMIYGYGREGSSYKALKLIMEMRQKGLIPNSASYGLTIRVLCKDDKCQEAEALLDDME--------------------------------------------------------------------------------------    
 876 UniRef50_Q656A1        Os06g0499301 protein n=3 Ta...   329     2e-88  1  10.7%  34:601    95:630     ---------------------------------LVAALSRSRGGGgpLLALRLFLAPLHPAAVTHHSFNSALLRFPLPPHLLPPFFSRSLRRFPGRlpTLLSFNLLLKCVCsgRYLDVALRILHEIIpdKFTYSTVVSALagRVDDAVALVHEMVADGVVA-AEAFNPVLRAMLrgDVKGAAKLFGFMQLKGCVPTTATYNVLVHGLLVCGRAGAAMGVMRRMEREGVVPGVMTYGAVVDGLVRCGRVKDAWKVAEEMERNGLAWNEFVYSTVITGFCKSGEIDCALKVWEAMVASP------------VRPNVVLYSAMIGGLANFGKMTEAELLFREMI--HSKCAPNIITYGSIIQGYFKIGDTSRALSVWEEMIGAGCVPNAVSYSILINGLCNVGRLKDAMMVWKHMLDRGCAPDtyTSMIKGLCVSGMVDGGLRLFYDMLASGHADPDVISYNVLLDGLLLAKDLPRAMDLLNRMLDQGCDPDTVTCNIFLREFGAGER---------------------------------------KGREFLEGLVVRLCDRRRNMAAGEVLMVMLAKYIVPEA-PIWEMVVRDVCRRKRVWR--    
 877 UniRef50_B7FM66        Putative uncharacterized pr...   329     2e-88  1  13.2%  87:517    39:464     --------------------------------------------------------------------------------------HFNFIPCSSSKINLIPYSSTSTTFHSNNDVDDAVSLFNCLLRQNPTpnKILGSLVkhYHTVLYLSKKMEFRGIKPNLVNCNILINCFCQLGlpFAFSVFAKILKMGYNPNTITFTTLIKGLCLKGQIHQALNFHDKLVALGFQFNQISYGTLIDGLCKVGETRAALDLLRRVDGKLVQPNVVMYSTIIDGMCKDKHVNDAFDLYSEMVSK------------GISPNVVTYSALISGFFTVGKLKDAVDLFNKMISEN--IKPDVYTFNILVDVFCKSGKISYALKLVDEMHDRGQPPNIVTYSSILDALCKTHRVDKAVALLTKLKDQGIRPntYTILIDGLCTSGKLEDARNIFEDLLVKGYDI-TVVTYIVMFYGFCKKGLFDEASALLSKMEENGCIPDAKTYELIKLSLFKKGENDMAEKLHRECE--------------------------------------------------------------------------------------    
 878 UniRef50_C5Z8R3        Putative uncharacterized pr...   329     2e-88  1  10.4% 167:580   186:564     ----------------------------------------------------------------------------------------------------------------------------------------------------------------------NSLIGAyaCGDVGAARKVLDEMVVK----DVISWTSIVAAYSRSRDMGSAEEVFALCPVK----DMVAWTAMVTGYAQNAMPVKALEAFEQMAGAGMPIDEVSLTGAISACAQLGAVRRAVWI------------QEIADRSGLGRNVVVGSGLVDMYAKCGLIDEACRVFEGMQEK------NVYTYSSMIVGLASHGRANDAIALFNDmrRADVEPNHVTFIGVLTACSHAGMVKEGRYYFAQMkrYGILpdHYTCMVDLLGRAGLVIEALDLVKSM----TVEPHGGVWGALLGACRIHGNTKVAKVAAQHLFK--LEPeiGNYVLLSNTLASAGEWDEVSKVRKLMRIRGLKKDP---------AVSSFEGRDGLVHQFFAGDNTHPWMHEIKKTLLELRARLKLAGYVP-----------------------    
 879 UniRef50_Q3ECB8        Pentatricopeptide repeat-co...   328     3e-88  1   9.2% 131:531    83:464     ----------------------------------------------------------------------------------------------------------------------------------FLWNNIMRSYIRhpLDAIQVYLGMVRSTVLPDRYSLPIVIKAAVqhDFTLGKELHSVAVRLGFVGDEFCESGFITLYCKAGEFENARKVFDENPERKLG----SWNAIIGGLNHAGRANEAVEMFVDMKRSGLEPDDFTMVSVTASCGGLGDLSLAFQLHKCVLQAKTEE----------KSDIMMLNSLIDMYGKCGRMDLASHIFEEMRQR------NVVSWSSMIVGYAANGNTLEALECFRQMREFGVRPNKITFVGVLSACVHGGLVEEGKTYFAMMkeFELEpsHYGCIVDLLSRDGQLKEAKKVVEEMPMK----PNVMVWGCLMGGCEKFGDVEMAEWVAPYMVE--LEpnDGVYVVLANVYALRGMWKDVERVRKLMKTKKVAKIPAYSYAS------------------------------------------------------------------------    
 880 UniRef50_B9T0L0        Pentatricopeptide repeat-co...   328     3e-88  1  10.8% 129:516    86:436     --------------------------------------------------------------------------------------------------------------------------------DVILSNQVITSFIRSGdsALELFNNMTIK----TTVTWNSILAGYSrgKMKNARELFDKIPE----PDTISYNTMLACYVHNSDMEKAQAFFDLIPNK----DPASWNTLISGFSQNGKMAKAHKLFLQMPYK----NVVTWNAMISGYIACGDLTSAWKLFKTMPVK----------------NVVACTAMITGYMKLGFIKLAEKLFKEMS------TDNVVTWNAMISGFIENSRAEDGVKLFRTMVGFGIRPNPSTLSSLLLGCSELSALQLGRQVHQLVCKSPLASDmgTSLVSMYCKCGDLEDAWKLFLELPRK-----DVVTWNAMISGYALHGAGEKALRLFDEMKKEGITPDWITFVAVLLACNHAGFADLGLKYFHSM---------------------------------------------------------------------------------------    
 881 UniRef50_A2WLV6        Putative uncharacterized pr...   328     3e-88  1  12.0%   6:487   138:607     -----MYVKCGDLENARKVFDEMPQvdVRVWTSLMSGYAKAGEFQDGVLLFRQMH--CSGVRPDAHAISCVLKCIAGLGSIADGEVVHGYLEKLGLGVQCAVGNALIALYSRCGHVDGSLQVFDGmhRDVISWNSVISGCFSNGwkSIELFAKMWSEGLEINPVTMLGVLPACAELGYNlgRVLHGYSVKTGLliDENLGSKLVFMYVKCGELGYARKVFDAMSSKS---NLHAWNLMMGGYAKLGKFQESLLLFEKMHDCGITPDEHTISCLLKCITGLSGVMDGLVV------------HGYLVKYGFGAQCAVCNALISFYAKSNRIEDALMVFDEMPQR------DIISWNSIIGGCASNGLYDKAVELFVRMWLEGQELDSTTLLSVMPACVQSHYSFIGGVVHGYSVRTGLISEtgNALLDMYSNCSDWRSTNKIFRNMEQK-----NVVSWTAMITSYTRAGHFDKVAGLFQEMGLEGWK--------------------------------------------------------------------------------------------------------------------    
 882 UniRef50_Q5VS02        Pentatricopeptide (PPR) rep...   328     3e-88  1   9.8%   1:602   212:910     LCAYARWGKLNdlMFYAAVRRRDIVPSISVFNFMVSSLQKQKLHGKVIHLWEQMLE--ANVAPNQFTYTVVIGSYAKEGMLEEAMDAFGEMKRRRFVPEEATYSLLISLCAKHGKGEEALGLYDEMKVKSIVPSNYTCASVldYSKALSLFSEMEQNKIVPDEVIYGILVRIYGKLGleDAQRMFEEIDKAGLLSDEQTYVAMAQVHMNVQNYDRALQVLDAMRARNVKPSQFSYSALLRCHVAKEDVDAAEDTFRALSNYG-PPDVFCCNDLLRLYMRLGHLDKARALILKMRKeeVIQNEGSSSKVLNPTDSSTLSMMLKsfAREGSTDEAKFLYEHLTELGAKPDDTAIayNAMVDALCRCGKTEEAYRLFMELIDQGHNGDAVTISILVTHLTKQEKFQEAENIIYRCleAELDTVVYNTFIKSMLESGKLYSAVSIYDRMISSGIPR-SMQTFNIMISVYGQGGKLEKAVEMFSAAQELGLPIDEKTYTNMLSFYGKAGKHHEASLLFSRMKEDGIR-PGKISFNTMINAYATSGLHNEAEIIFQEMQKNNHVPdkCYSKAEEAIQMMLRSNMTP-SCTHFNHLISAFLKEGQIDEA-    
 883 UniRef50_Q84MQ1        Pentatricopeptide, putative...   328     3e-88  1  11.3%   7:535    66:611     ------VSFPHHARAAETLFEHfpRPNLYIYNLMLSSAAAAAAAASSSPSRRpmSMLASSIHPDEQTFLSLLKS-VDAERRSVGKQVHAHVVVTGLHSRVYLRNSLIKMYLDAGDVEAAEAMFraPTADAVSCNIMLSGYVKGGckALRFFRGMASRGIGVDQYTAVALLACCGRLkaVLGRSVHGVVVRRIGVAdlILSNALLDMYAKCGEMNTAMRVFDEAGER----DGISWNTMVAGFANAGLLDLASKYFGEAPAR----DIISWNALLAGYARYEEFSATMILFHDMLASSvrSVHGWVVKEHGTQDSFLASALVDMYCKCGSIKLAYAVFEKAVDK------DVTLWTAMISGLAFHGHGDVALDLFWKMQAEGTEPNGVTLVAVLSACSHAGLFDEGCKIFDTLKQrnIEpeHFGCMVDLLARSGRLSDAVDLARRMPMK----PSRSIWGSILSASSACQNTEVAEIASKELL--CLEpeEGGYVLLSNMYAAEGQWNYSNEVRENMERKGVRKSADTETAKYKEL--------------------------------------------------------------------    
 884 UniRef50_B9GKL9        Predicted protein n=2 Tax=P...   328     3e-88  1   8.7%  65:549    22:505     ----------------------------------------------------------------SFSLKSCLSLLEICKFTTEFAQLHARLTKLGFIKNPLALTRLLCYSSISqnINYAQSIFndKNPNTFAYNVMIRGYAQrpENALSLFYSMlnANSGQNKLTFPFVLKACSQVriEEGKQVHGLVFKHGLSEDLFVQNSLISMYSSCGLIGFACQVFNKIDD----PDVVSWNSMISGLVDLGFVEEGKQMFDRMSKR----SLVTWNCLIDGYVKAGLLMEARELFDQMrlEQGEWMQAHIEKNGIDVNSVLGTALVEMFAKCGSIERALSVFKSIEER------DVGAWNSIIHKLAAHGHGQEAFAIFSDMLRSNTLLDGIIFLGLLSVCRHLGLVDEGKRFFQLmeEYGLVpeHYGCMVDLLCHADLLEEARDLIDS-SHASQMKSSVPMWGALLGVSCQLKNIVMGEYAAKHLLQLD-PFDGNCYTVLSNLYSAAGLYEKAIEVRNEMKKQGLEKIPGS----------SSMEIDGLVLDF------------------------------------------------------    
 885 UniRef50_UPI0001983420 PREDICTED: hypothetical pro...   328     3e-88  1   9.0%   7:530   156:634     ------LSPLGSLVYAQAIFEDTAMDNPFVCNTMIRAYAKSVFPIKAIYLYNFMH--RKDIKSDHFTYPFGLKACARVLwaCKGAEIHCRLFRLGLDQDCFIQNSLVYLYSQCGFLDLARC----------------------------VFDEMTEK----TITSWNVMISAYDqnDFDSADSLIGSMPGK----NVVSWNMLIARYIRLGKIEDAKVVFQEMPER----DAVSWNSMIAGYVQIKDYARALALFREMEIANVEATDITLISVLGACAETGALEIGRKIHLSLKQREYKIEG------------YLSNALVDMYAKCGHLNLAWEVFSELKMKH------ISCWNAMIVSLAVHGYCEEALRLFseMSVDGARPNRVTFIGVLIACSHKGLVDEGRTFFNRMveYKIMPdhYGCMVDLLSRCGLLKEAHHMIKTMP----FEANSVLWRTLLGACRVHHHVDLAEESFQQLGKMEPLRDGD-YVLLSNIYAEAQRWNDVERVRSEMIGSGvkKPGSSHIG-------------------------------------------------------------------------    
 886 UniRef50_A9SF53        Predicted protein n=3 Tax=P...   328     3e-88  1   9.7%  18:602    37:729     -----------------KMQEDECEPDQYTFNTLIMSASKANYVDYATITFQLM--LQTGVLPSILTYSMMLLLYQRHRKVAEAESVFSHMLNSG-VQAVVAYSVMIALYNREGLFEKSEQIVTEMRRNNVTPDrnTYGQQGKIEEAERIMDTVEESGMSLGLVGYNSMITAYGKACldKAARLVEKMREEDLVPDSITYSCMIGACGRVGKLKEALSWFAEMKRLEIKPASSNFNTLISLYGKAKDVEGIVRVITEMKKYGCKPDWQtyGTLLHVYLKCNLPKEALRIFLAMRkdGAKSVFKEMQVAGVTPSLESSCTMIYVYGMNGDVKEAEGLFHSLRSSVKRL--DIFAYNVMINVYMRCDMPEGAIKVYKLMEEDHLLPDAYTYHSMLRMCQKCNllEEMHKLFQEMIDVGYVpiTFNVMIDLYGKAGMPERAHKALK--LAQQFGSADKISFSTLVHAYAKKQDFPNMEAALWEMQNAGYGSSLEAYNSILDAYGKAGQMEKVSDVLARMENSGMRMDLAS-YNILINTYGKNYMIAEMETLFRTmiIRTYGFADYPDRAVYWFKAMQDAGISPDRV-TYMILVSTFERAGNIDEA-    
 887 UniRef50_Q6NKW7        Pentatricopeptide repeat-co...   328     3e-88  1  12.2%  34:535   168:663     ---------------------------------MLLCLVksGEERSLCLVMKDLFETRIDRRVLETVFSILIDCCIRERKVNMALKLTYKVDQFGIFPSRGVCISLLKEILRVHGLELAREFVEHMLSRGRHLNafIRKYCSDgdKGWELLMGMKHYGIRPDIVAFTVFIDKLCKAglKEATSVLFKLKLFGISQDSVSVSSVIDGFCKVGKPEEAIKL---IHSFRLRPNIFVYSSFLSNICSTGDMLRASTIFQEIFELGLLPDCVCYTTMIDGYCNLGRTDKAFQYF------------GALLKSGNPPSLTTSTILIGACSRFGSISDAESVFRNMKTEG--LKLDVVTYNNLMHGYGKTHQLNKVFELIDEMRSAGISPDVATYNILIHSMVVRGYIDEANEIISELIRRGFVPStfTDVIGGFSKRGDFQEAFILWFYMADL-RMKPDVVTCSALLHGYCKAQRMEKAIVLFNKLLDAGLKPDVVLYNTLIHGYCSVGDIEKACELIGLMVQRGMLPNESTHHALVLGL--------------------------------------------------------------------    
 888 UniRef50_D1IYP2        Whole genome shotgun sequen...   328     4e-88  1   9.8% 125:581    54:476     ----------------------------------------------------------------------------------------------------------------------------QPPRNSFHLDETLKGLcgRVMEAVGLLC---RTGLQVEPATYALLLQECikKEFKTGRRIHAQMIVVGYYPDEYLKTKLLILHAKTGDLDTSHILFDDLSKK----SLISWNAMIAGYVQKGLEEEGLNLYDEMRQSGLTPDQYTFASVFRACATLATLEKGKQAHCVMIKSQIKE------------NVVVNSALMDMYFKCSSLYDGHRVFNK------SLNRNVITWTALISGYGQHGRVAEVLVFFSKMKTEGFRPNYVTFLAVISACSHGGLVNEGWEYFMSmrDYGIQpqHYAAMVDLLGRAGRLHEAFEFVQNSP----CGEHSVIWGALLGACRVHGNIDMLRFAAKKYFD--LEpnAGKYVVLSNAYATFGLWDNVAEVRGVMRESGMRKDPGY----------SRIEVQREVHFFLMGDKSHKQTEQIYEVIKEVTCILKDAGYVPD----------------------    
 889 UniRef50_Q0DZE7        Os02g0625100 protein n=4 Ta...   328     4e-88  1   9.6%   1:523    13:473     LAAVA--TKPADFRRAHalLTSGHLSSRYSVNCLIRAASAPSASALLLRFL-----LNHRLLPDHLSLSFSLHPCSRLPSHPVASVVHSLAVRLGHARDVYVVNAAASAYFAAADVASAERLFSEASRDVA------------------------------DVVTWTTMVTGHAngDVERARWFFDAMPEK----NVVSWNTMLGAYARAGMLPKARKLFDRMPSR----NAATWGSMITGLLQSDHCEEALRVFSDMVASGVVPNEPALVSTVSACTQLRWMEHGTWVHGYAERELNG-----------AMSAVLATAIVDMYGKCGSIRDAVRVFAAMPVR------NIYSWNSMIAGLAMNGSERQALSLFWKMQLAGVRPNDITFIGLLSACSHSGLVDEGRWLFYKMVNhgIQpeHYGLMVDLLGRSGHVREAVDFVKSMP----VEPHPGLWGALAGACKIHGEVELGEEVAKKLIE--LEPQHGsyILLSNIYATSNRWDDMANVRRILKDRKVPK--------------------------------------------------------------------------------    
 890 UniRef50_Q8LH44        Os01g0390600 protein n=5 Ta...   328     4e-88  1  10.2%   3:522    47:646     --RNSLLAFYSRValAYHLFDEIPprDATAHNILLSALARAGRLERAQCLLAEM------PQRDAVSFTTVISALSRSGHPERALAVFRDmvRRGLDGFVIVATNLVHAYGAVSQVPSARSIFELMprNTVTWNTMLNCyaGMINMAAEVFGVIPErgIRANEVILVGLVKACSRHsvSEGQQLHTVILKNGfkSHVASWNALMASLLRRNLVHEARQLFDDMPER----DTISWSTLISGYVQSGNSNMALQIFCSMLDAGVEPNEITLASALSAVANSGTLGQARWIHDYIISRSIQLTDKLSAG------------LINVYAKCGSIAEAVQLFNHVKDKSISVSP----WNSIICNLAIHGYVNMSLELFSQLQstHIKPNSITYLGVLNACCHAGMVAEGKRQFESMrqYGIQPehYGCMVDLLCRAGYLEEAELLIKTMPMK----ADVVAWGCILAAARTQGNVALGEKAAEEL--SKLDPSHgsKIALSNLYADAGHWSNVSVVRKELQNENLE---------------------------------------------------------------------------------    
 891 UniRef50_B9RA32        Pentatricopeptide repeat-co...   328     4e-88  1  11.7%  61:512    33:467     ------------------------------------------------------------PDSTEIVYDNLVKFC---DGLAVLKQIHSALTTNSLITKSPHLAaiIIKYAKFGYLNCARSLFDGInrSSFLWNTMIRAYANAGleTLELYMLMRRAGVSSNNYTFPFIFKACASNSLlqGKVAHGDAVKADFDSDVYVKAALVDMYAKCGQFCDGRKIFDEMPVK----DLVCWTAMITAYEQGEKPDEALILLRKMQQHGLFPDEVTMVSVASAIGQLWDAKWAQSVHAYAIRRSFLKE------------IFVANSILAMHTKCGNMEKSCLIFDMMDER------NVISWNSMLSGYTQNGQASEALFLFDKMRDSGCEPNSVTALIMVAACAYLGSRHLGGKFHDFILDSKMKIDmrNALMDMYAKCGDLKTAVEMFNDV---HPSERNVSSWNVLISGYGMHGHGKEALRLYSRMQEESVEPNHITFTSILSACSHAGLIDEGRNL-------------------------------------------------------------------------------------------    
 892 UniRef50_Q8L6Y3        Pentatricopeptide repeat-co...   328     4e-88  1  12.9%  68:532   124:588     -------------------------------------------------------------------HSSIMRDLCLQGKLDAALWLRKKMIYSGVIPGLITHNHLLNGLCKAGYIEKADGLVREMrpNCVSYNTLIKGLcnNVDKALYLFNTMNKYGIRPNRVTCNIIVHALCQKGvnNNKKLLEEILDsnAPLDIVICTILMDSCFKNGNVVQALEVWKEMSQKNVPADSVVYNVIIRGLCSSGNMVAAYGFMCDMVKRGVNPDVFTYNTLISALCKEGKFDEACDLHGTMQN------------GGVAPDQISYKVIIQGLCIHGDVNRANEF--LLSMLKSSLLPEVLLWNVVIDGYGRYGDTSSALSVLNLMLSYGVKPNVYTNNALIHGYVKGGRLIDAWWVKNEMRSTKIHPDtyNLLLGAACTLGHLRLAFQLYDEMLRRGC-QPDIITYTELVRGLCWKGRLKKAESLLSRIQATGITIDHVPFLILAKKYTRLQRPGEAYLVYKklATRNRGVSCPSILNHM-----------------------------------------------------------------------    
 893 UniRef50_B8LN20        Putative uncharacterized pr...   328     5e-88  1  12.1% 162:500    90:418     -----------------------------------------------------------------------------------------------------------------------------------------------------------------DVISYSRKINGLCkgQIAEAMEIFREMEEKGSAPDSFTYNILINGLCKDAQLEMGVSLMDEMQARGFSPTVVTYTTLIHGFCTAGRVDAAMDILDEMEHRGITPNTVTYTALIQGLCDCGEVERAILVLKEMKLRG----------NECCPNVVTYTQLIHGLCQKGLFEMGMEFLDEMIISGCF--PNNATYTALIHGLCQKGSFLEAAVLLNEMVAIGCELDVVIYNSLIDGLCKMGELDFAREMQILMVEKGVAPdtYTSLIQGLCDAGLLDEALGLMQEMRAKGIV-PNSVTYTTLINGFTQADRTDHALQLVNEMKSNGLMPAGDIYKILFKSL-------------------------------------------------------------------------------------------------------    
 894 UniRef50_Q9FGR7        Pentatricopeptide repeat-co...   327     6e-88  1  11.2%  47:555   173:690     ----------------------------------------------AREILELAKNLKEN-----QTLGEMLSGFERRVSDTECVEALVMMGESGFVKSCLycSVLFTLLGRERMADYILLLLSNLprDVRLYNAAISGLsqRYDDAWEVYEAMDKINVYPDNVTCAILITTLRKAgaKEVWEIFEKMSEKGVKWSQDVFGGLVKSFCDEGLKEEALVIQTEMEKKGIRSNTIVYNTLMDAYNKSNHIEEVEGLFTEMRDKGLKPSAATYNILMDAYARRMQPDIVETLLREME------------DLGLEPNVKSYTCLISAYGRTKKmdMAADAFLRMKKVG--LKPSSHSYTALIHAYSVSGWHEKAYASFEEMCKEGIKPSVETYTSVLDAFRRSGDTGKLMEIWKLMLREKIKGttYNTLLDGFAKQGLYIEARDVVSEFSKMGL-QPSVMTYNMLMNAYARGGQDAKLPQLLKEMAALNLKPDSITYSTMIYAFVRVRDFKRAFFYHKMMVKSGQVPDPRSYEKLRAILEDKAKTKNRKDKTAILGIIN------------------------------------------------    
 895 UniRef50_A9SSM7        Predicted protein n=1 Tax=P...   327     9e-88  1  12.4% 175:600     7:398     ------------------------------------------------------------------------------------------------------------------------------------------------------------------------------RGARRQALQVFRFLQEheYELKEHNCVTIISILGREGKLGLAREIFEGMSKAGVAPSVHAYTALLSGYAKQGLLKEAWALFEAMKEKGCSPNVLTYNTLINACTKRarLPDLVGLFEEMKQ------------AGVQPNDITYNCMVNACVCLSLFDTASQILKEM--KAVNCLPNVISYTTMINSLGRSGRLDEAVELFEEMKELGRSPNSWTYNSLLKAYAREGRYEKAMCLFVGMEDEGCIPdtYNTVIDMCGRGGLFAEAEGVFLEMQRKGC-TPDRVTYNTMLDAYSKWSRRGRARDLLKTMKRAGCTPDLWTYNILLDAAGKAGSASEAMQIFHELKAAGHSPNLVS----------------------FSALINMYGRLGYFEEAERAWVEMRATGCVPNA-TAYCGLMNSYSHHGMYK---    
 896 UniRef50_C5WX20        Putative uncharacterized pr...   327     9e-88  1  11.2%  69:569    51:530     --------------------------------------------------------------------TALLSAYAALGRPRHARDLFDEMPDPGL----VTRTAMARAHAASGQTAQALAVFGDMLADGVLPdaLGMAAARRPGKM--VHALIVTSGIVPDVFVSTELIRLYGegELLVSRRLFDAMPVK----STVSWNAMVHQYIRHSNVGDAYELFLAMPKR----DVVSWNTIIAGYCLIGRFMEALELFRQMmsCPVHPNGPTISTVLAACAGAGCLETGIWV------------HAYIDRNRMNDDGSLDRSLIDMYAKCGSIEKALQVFEKAPGK-----RDLYSWTTVICGLAMHGKAADALRMFGMMQDNGIRPDDVTLVGVLNACAHGGLVDEGLHHFYSLEKYAITpeHYGCVIDLLGRVGRLQEAYSIIRTMRMK----PNAVIWGAFLNACKVHSNVELGEIAAAELTRLDPD-DPWAKVMLSSLYAKAQDWSSLARERREMNSLQMKKTP----------GCSSIELDGEVHEFVAGSFQHPQHGEICTVLENV----------------------------------    
 897 UniRef50_D1HMV4        Whole genome shotgun sequen...   326     1e-87  1  13.9% 146:576    87:537     -------------------------------------------------------------------------------------------------------------------------------------------------DALELKNQMVEKGIKPDVFTYTTLLSGFEkgKDKAAVQIFEEMRNEGCKPNICTFNALIKMHGNRGKFTEMMKVFEDIKTFQCSPDIVTWNTLLSVFGQNGMDSEVSGVFKEMKRAGFVPERDTFNTLISSYSRCGSFDQAMAVYKRMLE------------AGVNPDLSSYNAVLAALARGGLWKQSEKVLAEM--KDGRCKPNELTYCSLLHAYakCDLLMETERAFLELRQRGFSPDITTLNAMVSIYGRRQMVAKANEILDCMKRGGFTPstYNSLMYMYSRSANFERSEEILREILAKGI-RPDIISYNTVIYAYCRNGRMRDASRVLSEMRESGPAPDIITYNTFIASYAADSMFVEAIDVVCYMIKHGCKPNQST-YNSIVDWYCKLNRRDEASMFVNNLRKLDPHI--SMDEECRLSERMAKK---------------------------    
 898 UniRef50_Q9FX57        T6J4.15 protein n=7 Tax=ros...   326     1e-87  1  11.8% 125:582     1:417     ----------------------------------------------------------------------------------------------------------------------------MVEKNVVLWTSMINGYLLNKdsARRYFDLSPER----DIVLWNTMISGYIegNMLEARSLFDQMPCR----DVMSWNTVLEGYANIGDMEACERVFDDMPER----NVFSWNGLIKGYAQNGRVSEVLGSFKRMVDEgvVPNDATMTLVLSACAKLGAFDFGKWVHKYGETLGYNKV-----------DVNVKNALIDMYGKCGAIEIAMEVFKGIKRR------DLISWNTMINGLAAHGHGTEALNLFHEMKNSGISPDKVTFVGVLCACKHMGLVEDGLAYFNSmtDFSIMPehCGCVVDLLSRAGFLTQAVEFINKMPVK----ADAVIWATLLGASKVYKKVDIGEVALEELIK--LEpnPANFVMLSNIYGDAGRFDDAARLKVAMRDTGFKK----------EAGVSWIETDDGLVKFYSSGEKHPRT----EELQRILRELKSFNILRDE---------------------    
 899 UniRef50_Q0DXS6        Os02g0735800 protein (Fragm...   326     1e-87  1   8.8%  30:541   126:613     -----------------------------SASAAIVALAHAGRHAEVVELFCRMR--RGGVPVSRFVLPSVLAACAGLRDIGMLRAVHALVIKCGLCQHVIVGTALVDGYTDFGLVDDARKAFDEidANIVSWSVLIGGYArrWEETLDAFSAMRRAGVLPNDSVLVMAIQAcaLGRLVHGKQLHGLAVVLGFDRNATVWNCLMDMYGKCGDIDSCKMVFETMIGR----DQVSWNTLISSYARVGLCEEALDMIVQMQESGYIVDRFTLGSGVTACARLADIDSGRAF------------HGYLVRRLLDTDVIQGSALVDMYGKCHNMELAHIVFDRMDERNY------VSWDALLSGYVENEQVDLALEIFRQMGCANIKYNQHNFANLLKLCGSQRYKEYGRQIHGHAIKTINKMNVvtELIDMYAKCGCIEVARLLFLRMNER-----NLISWNALLSGYAADGQPVATINIYRQMELACIRPDKYTLAGLLSLCRYQGLLHYGRQIHAHLIKMG-SEMNVVMQTILVHMYIKCMR--------------------------------------------------------------    
 900 UniRef50_Q0D7D9        Os07g0259400 protein n=6 Ta...   326     1e-87  1  10.3%  86:571    27:491     -------------------------------------------------------------------------------------AVHPLAVKSGSGSDARVATALADAYAKSGLVDRARRVFDepLRDQVLWNVMVSCYSSHgrECWDVFGSMRRSGFPGDGFTFSALLSvhANLLLVLGSSVHGIVIRLGLHLDVVVATALLDMYAKCGQVAEASRVFDAM----VLRNTVSWNAIIVCYGKHDRGKEAFDLFVSMMRHGFCPDELTLASLLSSCADMAAANEATQL------------HAYTVRRGLQDFLQVGNALIMAYGKNGFVQEAKRTF------GMIHNPDLVTWSSMVSSFAYLGLAKSAIDLFDRMLQQGIRADGIAFLGVLSACSHAGLIEDGFKYFLLmrDYKIDPthLACLVDLLGRAGRIRDAYEFLVNMS----CDANVDVIGAFLGACRMRGNIESAKWAASRLFS--LKpdPINYLLISNTYAAAGDWNELAKVRSVMRNmgNKVP-----------GCSWIEIGGIVQTFVSNDMMLHQSREMQRMMELLVS--------------------------------    
 901 UniRef50_Q60DD6        PPR repeat containing prote...   326     2e-87  1  10.9%  23:593     1:552     ----------------------MPPPPVPTISTLSALLAGCASLSTAAALHAHLLKSSRLFRP-VFLANCLAAAYCRLGAAPSAVAVLRHAPEPNIFSR---NILLGAMLKSRDLLSARRLFDEMPDRDAVAYNSMMSGYigRNNEALSLVWTMLEAGVRPSGFTFSIILSAVRVAR--HGVQVHAAAVRHCFAhnSVVGNALINMYRRVGLLEYAVQVFWSMNGH----DIVSWNSVMSVYRDDGQRRQVFECFRMIRSHGLFFDECSLSTVLSACIDAEDSSKGDQLL------------THCVKMGLLRNSLICSAVIGLLCASDRLADAVYLFKGM------ATWDSETCNAMISGYARSGLMEQALGLFTMALQNGILPTGFTFASVLRWSSCFGLVEQGTQIHALIFKLGLEDdiATALVDMYCKLASLKHAKKIFSRVSFK-----DLVLWNTMIIGLSHNGRGKEALQVFRQMLKCNIQPDRITLSGVLSACSFEGLVNEGIKMVSLFEDKYHIVPGVEHYTCVVDMLSRAGMLGEAVDFVESKLQkaSLIKRDFRMAELIAEKMTK--LKPRSSLPYVVLAQSY----------    
 902 UniRef50_C5Z6K1        Putative uncharacterized pr...   326     2e-87  1  10.3%  54:521     1:460     -----------------------------------------------------MRMLSQSHAP-----AALLRAITACTSCREAQALHSLACKLGLASDMVISTALLTRYAKRGLVARAQRVFDEmrRDVVAINAMLAAltGRVTDARALFGRMSYR----TPASWNTMVTCYCrgDLTSARDVFEASLRA-MSSSVVSWNAMIDGYCKAGRMDAARELFDRMGSS--LPDVITWNTMMAGYLRGGDPASAIAMFHlmreQRLRPTNGTMATVVAACAQVGNLALGRQIHHLIQQK------------GIWIDTVLSNALMDMYFKSGSVDCALDVFRAMPR---PRSPNLFCWNTVISGLGMNGHGEDAIAAFHDMVenSVKPDGVTFVALLSACSHSGLVLEARKFFSQMlvYGLPpeHYGCMVDLLCRAGHVDEAARLVQTMPGR----PNAKVLGSLLLDARVSERWQEgeWAARRISRLDLH-DGAAY-GLSNVYASLQRWDRVEEHRRQVRAAGC----------------------------------------------------------------------------------    
 903 UniRef50_A5AJ01        Putative uncharacterized pr...   326     2e-87  1   9.0%   5:581   130:647     ----RMSQKSIDFGLTHQLFDEIPVSNTFAWNNLIQTHLTNGDSGRVVSTYRQM--LLRGVRPDKHTIPRILTAARHTSSFSFGKQVHGHALKLGLSSESYVISALLEMYGRLDGABAAKLVfkSARRNSVSWTLISRLYImkPGLAVDMFKQMVESKSEIDPLALVTAIVACG-------------MLKSLPGG---------EMYIDCGSIKDARAVFDRMPSK----DVISWTEIFRGYVKNGGFNEGLKLFRQMSMEGVKPDSLAISSILPACGRGAAHKQGKEI------------HAYLLRNGIDLNVTVQNAVLDMYVKSGFIESAAKIFAGMKDR------DAISWTVMILGYSLHGQGELGVDLFRKMEksSVEIDQIAYAAALHACTTARLVEQGRFYFNCITA--PKSRHYALMVALLSRVGLFDEARVFMEEHKLEGHVE----VLRALLDGCRIHHNMRTAKRVIEQLCDLQ-TLNADNYVLLSNWYSSFAKWDMVNELRETIRDMGLKPRKAY----------SWIEFRNKIHVFGTGDVSHPRSEKIYWELHSLMKKIEEEGTRLN----------------------    
 904 UniRef50_B9T3M8        Pentatricopeptide repeat-co...   326     2e-87  1  10.5%  83:589     8:494     ----------------------------------------------------------------------------------KALSLFNSSINQGINPTHQSVFFIVHYLLKFNLKSEVQSLLLQILSNRIspFFTVSSLLENLTEPISI------PCPKISSLLYESIISVYIqgSIELAVLYFNQMVDKGLVFEPNIFNTLLDSLVKSKCFEKLWFVFNEVKGK-IELDVYTYGIMIKGCCEAGVFYKGFELLTDLEKMGLSPNVVIYTTLIDGCCKNGDVRRAKLLFNKM------------GEVNLVANQYTYTVLINGFFKKGLRKDGFELFEKMQLDG--VFPNLYTYNCMINEFCNEGKLHKAYELFDEMCKRGVVSNVVTYNTLISGLCRKTRIWEAEKLVdeMEERGISPstYTILTNAHVRLGDMEKAFHFYSSMEKVGLV-PDVHIYGVLLHGLCVKGNMKEASKLFRSMAEMKLEPNEVVYNTMINGYCKAGNSFRALRFLKEMEDKRLVPNVAS----------------------YSSTIGILCKDGKWQEAEILLNKMRGSGLEP-SVSIYNII--------------    
 905 UniRef50_A9SEH7        Predicted protein n=2 Tax=P...   326     2e-87  1  12.4% 101:593   240:730     ----------------------------------------------------------------------------------------------------FYIDLYASLLKAGRIKDCLVLllSLMKVNHSKFYEVCKTK-GTVEDAFTFAKIMR---VYSTLQHYTMLLSVCCHadIDGALRVLALVESRGFKADCVFYTTLISACSKAGKVDLLFQILHEMDVAGVEANVHTFGAMIDGCARAGQLPKAFGAYGIMISKDVKPDRVIFNTLINACSRVGAVQRAFDVLADMKSEA----------TPVKPDHVTYGALISACAKGGEVGRALEVYQSMRENDVKGF--LACYTAAVHACSQKGDLDYALAIYDDLLKDGVKPDEVFFSALVDAAGHSGDVDKAFSILDIMRKSGMKpvVYSSLMGVCSNLGEWEKGLEVYEGIRSSKL-RPTVSTYNALMTALCEAKQFDRALSVLNDLKDAGRTPNQVSYSILLKACEREGKADMALDLYTTARAEGIKPNLVI-CDSIIGLCLQQIQTSVAAPLVSWSisNISSQQQWVTWALTIYRQTIEAGVLP-TVEILSQLLGCL----------    
 906 UniRef50_D1H7S8        Whole genome shotgun sequen...   325     2e-87  1  10.4%  71:600    46:517     ----------------------------------------------------------------------LIHA---SNTLPQLHQIHAQIFLHNLFSNSRVVTQLISSSCSLKSLDYALSIFRCfhPNLFVFNALIRGLAEnfEGSVSHFVLMLRLSIRPDRLTLPFVLKSVAAldVGLGRCLHGGVMKLGLEFDSFVRVSLVDMYVKIGELGFGLQLFDESPQRNKAESILLWN-------------------------GVRPNDLTVVSALLACTKIGALQVGERI------------HNYLSSNGFQLNRGIGTALVDMYAKCGNIKSASRVFVETKGK------DLLTWSVMIWGWAIHGCFDQALQCFVKMKSAGINPDEVIFLAILTACSHSGNVDQGLNFFESMrdYSIEPthYTLIVDLLGRAGRLDEALSFIQSMP----INPDFVIWGALFCACRAHKNIEMAELTAEKLLQ--LEphPGSYVFLSNVYAAVGRWEDVERVRTLMKNRGVEKDPGWSY----------IEVEGQVHSFVAGDHAHVRAEEISLKLEEITASAKQEGYMPETAW----VLHNIEEEEKED---    
 907 UniRef50_Q0DLS5        Os03g0849700 protein n=7 Ta...   325     2e-87  1  10.8%   3:565    42:611     --SNSLLLMYARCsdARRLFDEMPVKNCFSYNSVITSHLNSRDHHAALNIFRSM------PERNTFSWNTIITGIVSTGNLDMARSLLIEMP---VKDPVACNAVLHRYVRRGRADE-AFALLRTVGqdPFVLATIVGACamKYDFGRQAHARMVVSKIEQDLVLSCALVNMYCkgDLDSARYVLNGLTQ----VDEFSLSALIYGYASCGHLHEAIRLFD----RKEEPSIAMWNSLISGCAFACCGNDAFALFARMMRSDVLPDSSTYASILNVCG-----------FSVMVN-PGQQIHGCGLKCGAVNDIIVASALIDFYSKCGLWEDACRAFRELR------FHDTIVLNSMITVYSNCGQIEEARRIFDMITGK----SVISWNSMVVGLSQNGHARDALGLFCEMHRLGLRLdvASSLIDLYCKCGSLANGCRIFEE-----IDKPDEVLWNSMLIGHASNGYGHEALELLELMKTKGIKPSERTFIAVLSACCHSGLVKEGLTWFHRMQADFSVSPSAEHYACVTDLLVRAGRLEESVEF----IENMPFEADAVSW--------------------------------------    
 908 UniRef50_Q10RD7        Os03g0162900 protein n=4 Ta...   325     3e-87  1  10.2%  20:597   133:752     -------------------RRRRHEVNVIHYNVV---LCAVGRARRWDLVARLWHEMHSGgaPDNSTYGTLIDVHCKGGRERMALLWLGDMIKRGLLPDEITMSIVLQVHKKAGEYEKAELFFksLESDvyTYNTLiyGKAGQLEKVSDTFNQMLREGVSPNVVTFNTMIHVWGKHrmEQVASLMRTMEEFQCLPDTRTYNILISLYREINDIDVAEYYFRKMKAENLLPDVVSCRTLLYGYSIKGMVTEAQALLKEMYERNMVIDEYTQSAVTRMYVDAGMLEQAWCWFEKfiDAFGERGHILLAEKAFmlSTCVCNVMIKAYGLVEKLDEACEIADGMERYG--ILPDYLTYSSLIHLMSTAKLPEKALYYLRKMQAAKLLIDCVPYSVVISCFAKNGNLHMVDCLFREMITSGIQADTysILIDAYAEVGDVQKAEAYFGLLKKAGLCE-SATIYNSLIKLYTKVVYLAEAQKTYKLLKSLNADTNLYASNCMIDLYSDHCMVKEAREIFENLKVTG--KANEFSHAMMVCLYKKIARFDEAHRISKKMQAsmYVSGGRMEDALKIFQKMLASNTPPND-ATFKALKIILVKRG------    
 909 UniRef50_O82178        Pentatricopeptide repeat-co...   325     3e-87  1  12.6% 124:558   149:572     ---------------------------------------------------------------------------------------------------------------------------SSFQPDVICFNLLIDAYGQKfkEAESLYVQLLESRYVPTEDTYALLIKAYCMAGleRAEVVLVEMQNHHVSpgVTVYNAYIEGLMKrgNTEEAIDVFQRMKRDRCKPTTETYNLMINLYGKASKSYMSWKLYCEMRSHQCKPNICTYTALVNAFAREGLCEKAEEIFEQLQED------------GLEPDVYVYNALMESYSRAGYPYGAAEIFSLMQHMGC--EPDRASYNIMVDAYGRAGLHSDAEAVFEEMKRLGIAPTMKSHMLLLSAYSKARDVTKCEAIVKEMSENGVEPDTfnSMLNLYGRLGQFTKMEKILAEM-ENGPCTADISTYNILINIYGKAGFLERIEELFVELKEKNFRPDVVTWTSRIGAYSRKKLYVKCLEVFEEMIDSGCAPDGGT-----AKVLLSACSSEEQVEQVTSVLRTMHK---------------------------------------------    
 910 UniRef50_Q9LXY5        Pentatricopeptide repeat-co...   325     3e-87  1   8.6%  71:580     8:491     ----------------------------------------------------------------------IVRMLQGCNSMKKLRKIHSHVIINGLQHHPSIFNHLLRFCAVsgSLSHAQLLFDHFDSDPStwNYLIRGFSNspLNSILFYNRMLLSSvrPDLFTFNFALKSCERIkiPKCLEIHGSVIRSGFLDDAIVATSLVRCYSANGSVEIASKVFDEMPVR----DLVSWNVMICCFSHVGLHNQALSMYKRMGNEGVCGDSYTLVALLSSCAHVSALNMGVML------------HRIACDIRCESCVFVSNALIDMYAKCGSLENAIGVFNGMRKR------DVLTWNSMIIGYGVHGHGVEAISFFRKMVASGVRPNAITFLGLLLGCSHQGLVKEGVEHFEIMSSqhLTPnhYGCMVDLYGRAGQLENSL----EMIYASSCHEDPVLWRTLLGSCKIHRNLELGEVAMKKLVQLE-AFNAGDYVLMTSIYSAAND----AQAFASMR----KLIRSHDLQTVP--GWSWIEIGDQVHKFVVDDKMHPESAVIYSELGEVINRAILAGYKP-----------------------    
 911 UniRef50_D1IVG6        Whole genome shotgun sequen...   325     3e-87  1   9.7%  62:546    76:594     -------------------------------------------------------------KPNAYALLHLVRASTNLGLIPFGQQLHSYILRSGFGSNVFVSTALLNFYIRIESLKDAHNVfeIPEPSVVSWNSLISGyaGQFRAALGLFLKLERSDICADAFSFTAALATCGHLslRLGESIHSKIVKSGLEWSVVVANCLVDMYGKCGYVQEAIRVFDSIIDK----DIISWNTVISASARNRKLEQAFSYLRQMPE----PDTISYNELINGIAQFGNIeeALDFFSKMQSKDIelIHCCTIKCGLDESIVVGSALIDMYSKCGQVKNAEMLFQSLPRK------NLVTWNAMISGFAHNGNFTEVIKLFKQLkeRDLKPDGITFLNVLLACSHNQilQVAIQYFESMIkhGIEpeHCCSMIRLMGQRGEVWRAERMIYELGFSSCGL----VWRSLLGACGACGDLDVAEVAAAQVIELEGH-NEFVYVMMSNMYACYGKWGDVSVVRKLMRERGVRK----------GAGCSWIEVENVI---------------------------------------------------------    
 912 UniRef50_B9G1K2        Putative uncharacterized pr...   325     4e-87  1   9.4%  16:515    57:541     ---------------ALSLFAAIPTPTLFHHTALLRALSASAGGSLAAALSVLASARARLPALDEFAFQPLLALCAKRgaASLGRQVHALVVRYGFSGVVSLGNALCHFygGGEGMADARRVFDEmeRDAVSWNTVIGGYVRAGekAVDMFSEMMCCSVGVSVTALVTMI----RCGW-QAEPVHGFCIKVGFCRDAKVASAMVGMYVREKSVECGRKVFDEATKR----DLVLYNCMVDGYAEAGQVEEAMGLVDRMRLEGVRPSSGTLVGVLSACGASGAMAAGRRL------------HEIALEAGLELDTALGTALMDMYFKCGYPSEAVAVFDAMQER------DVKAWTVMIMGFGVNGQAGEVISLFRSMEEDGVVPNEVTFLAVLNACSHGGLVSEGKKFMESMvqYGIFPnhYGCIIDLLGRAGRLDEAYELIASLSSQS----DATAWRALLAACRVHGNANLGRMvlDNM--DDYHPSDAI--LLSNTYALESRWDEIAHVRDS----------------------------------------------------------------------------------------    
 913 UniRef50_C5XTD9        Putative uncharacterized pr...   325     5e-87  1   9.1%   3:551   155:757     --ANGLVTMYSSCQSLrkVFGSITSPDLVSWTSMLSAYTENGCDAEALMLF--MEMARDGIACDAFTLSVALRAASSLGHVGLGHQLHCCMIKMGLVGKEFLDNCLIGFYGRSGELQLMRNVFDEMnkDLVSWNTIIQCYAENLceASAHFRAMMFEFAECDEFTLGSILHVvaTGAFGHGMEIHGYLIRAGLDSDKHVMSALIDMycASgdLVAGRMLHACILKSNMNPDSFVTSSLVDMYAKCGALEeaVHLFRRMQSEHVQPNEFTYTAILTACMALGDTDSGMEI------------HSNSIRSGYGTNTSVLKSLITFYLRQGRYHQALKL-------CLALSNHDISWDTLVESFSQVDHHVGIVNLFHVIQRCGANLDYHTARLILSSCGKLGLLEEGLQAHAYMTKRGLASTACTnlIDMYSSCGSLRHAFDAFNYMPDK-----DASSWTSIVAANVENGCPETAIRLFSQMQKEKCRPTPEAFLSVLKACARTGLVNEAFRFFASMTEVYKIQPSEEHYSHMIEVLSRAGMFKEAEHFIDS----------------------------------------------------    
 914 UniRef50_Q0D7D7        Os07g0260000 protein n=6 Ta...   324     5e-87  1   8.5%  22:575   105:585     ---------------------QGVPPNGFTFTFLLRACALLGLPRPCGCVHGQIVRCGFG--SDVFVQNALMDVYHRCGGGGgaRQVFDEMVDRDVVSW----NSIVGVYMSSGDATGAMGFFEAmeRNVVSWNTVVAGFARMGDmaRAVFDRMPSR----NAVSWNLMISGYasGDVEAARSVFDRMDQK----DVVSWTAMVSAYAKIGDLDTVNELFDHMPVK----NLVSWNAMITGYNHNSRYDEALRTFQLMMLEgfRPDEATLVSVVSACAQLGSVE------------YCNWISSFIGKSNIHLTVALGNALIDMFAKCGDVGRAESIFYKMETRC------IITWTTMISGFAFNGLCRDALLVYNNMCREGVQLDDTVFIAALAACAHGGLLQEGWSIFNEmeRYNIQpeHYGCMVDLLGRAELIEYVSKKITEL------EPFNSSYQVLVSNCS-------ALE---------------------------GRWDGVIDARTSMRNWGIEKVPGS----------SSIQVGSEVHEFLAKDTRHKRRKEIYETVDGLMALMRH----------------------------    
 915 UniRef50_B9MZJ2        Predicted protein n=7 Tax=P...   324     5e-87  1  12.6%  44:463    67:479     -------------------------------------------IDDAMASFYRMIHMN--PRPSFVELGKFLGSIAKKKQYSTVVSLCNQMDLFGVAHNDYTLNILINSLCRLSHIHFAVSVLSKMFKLGIQPDSVTFTTLINgkEAVELFNEMAGRDAMPNTVTFTILVDVLCKKGmsEARCVFETMTEKGVEPNIYTYNALMNGYCLRLEMNDASKVFEIMVGKGCAPSVHSYSILINGYCKSRRIDEAKALLTQMSEKELIPNTVTYNTLMQGLCHASSLLEAQELFKKMCSS------------GMLPNLRTYSILLDGLCKHGHLEEALKLLTSMQER--KLEPDIVLYNILIQGMFIAGKLEVAKELFSKLFANGIRPSVRTYNIMIKGLLKEGLSDEAYKLFRKMEDDGFLPdsYNVIIQGFLQNQDPSTAIQLIDEMVGR-RFSADSSTFKMLLD--------------------------------------------------------------------------------------------------------------------------------------------    
 916 UniRef50_O81908        Pentatricopeptide repeat-co...   324     5e-87  1  12.5%  51:602    85:655     --------------------------------------------------LRFFDwsNKGFSHKEQSFFLMLEFLGRARNLNVARNFLFSIERRskLQDRYFNSLIRSYGNAGLFQESV-KLFQTMKQMGISpnSLLSILLKRGrmAHDLFDEMRryGVTPDSYTFNTLINGFCKNSmdEAFRIFKDMELYHCNPDVVTYNTIIDGLCRAGKVKIAHNVLSGMLKkdVHPNVVSYTTLVRGYCMKQEIDEAVLVFHDMLSRGLKPNAVTYNTLIKGLSEAHRYDEIKDIL----------IGGNDAFTTFAPDACTFNILIKAHCDAGHLDAAMKVFQEML--NMKLHPDSASYSVLIRTLCMRNEFDRAETLFNELFEKecKPLAAAYNPMFEYLCANGKTKQAEKVFRQLMKRGvdPPSYKTLITGHCREGKFKPAYELLVLMLRREFV-PDLETYELLIDGLLKIGEALLAHDTLQRMLRSSYLPVATTFHSVLAELAKRKFANESFCLVTLMLEKRIRQNIDLSTQVVRLLFSSAQKenGYLVKMEELLGYLCENRKLLDAHTLVLFCLEKSQMV--DIDTCNTVIEGLCKHKRHSEA-    
 917 UniRef50_D1HT27        Whole genome shotgun sequen...   324     6e-87  1   9.0%  31:555    48:535     ------------------------------FNTLLPSLS----PQDALLLYNQMVLHRTSH--NHFTFTHALIASSSLHALHKTLEIHARAIKSGHYSDIFIQNTLLHSYVVENNFVFAKSVFKSIspDVVSWTSIISGLSKCgeEAIGEFLSM---DVKPNTSTLVSVVSACCGLrvRFGKAIHGYSLRSMDGDNIILDNALLDFYVKCGYLVSAKYLFMKM----FRRDVISWTTMVGGLAQGGLCEEAVEVFQAMVKGGevPNEVTLVNVLTACSSLSALNLGRWVHSYI-----------STRYDLVVDGNVGNALINMYAKCSDMYMAVRVFNELTHK------DMISWSTIIGGMAMNGHGMHALQFFSLMLVHGVSPDDVTFIGLLSACSHAGLVEQGLIFFKAMNnyGIAPqhYACMVDMYGRAGLLEEAEAFIRGMPMEAEGP----IWGALLNACKIHGN----EKMFGRIDqkGV--SIGTLALLSNTYASSNRWDDANKVRDMMRDMGLKKM----------SGCSWIEVDAMINRYDGANRS------------------------------------------------    
 918 UniRef50_B9S8R9        Pentatricopeptide repeat-co...   324     6e-87  1   9.4%  85:521     5:453     ------------------------------------------------------------------------------------KQIHAQIILNNLTDQTLTLGKLVSFCAVSdnLDYAHLVFNQIspNKFMYNSLIRGYCNspIKSMFLYRQLIDSGLSPNEFTFPFVLKACakSAHWMSMIVHGHAQKLGFASLICVQNGLINAYIACGFIRYARKMFDDMSER----SLVSWNSMIGGYSKLGWCKEVFLLFKEMREIGTEADDFTLVNLLLVCSRRCDINLGRFV------------HLYIQITGMKIDLVARNALIDMYAKCGALVLAERVFQQMPNKNVvpEKNVVSWNSMISSYVQGGQCREALDLFHEMHSFRVVPNEATLLSVLSACGQIGDLVMGKKIHNYitSSMYSVTLCNSLIDMYAKCGALRIAIDVFNEMPNK-----NLVSWNVIIGALALHGYGVEAVELFRKMQAAGVWPDEITFMGLLSACSHSGLVDSGLYYFDKMSSIYV----------------------------------------------------------------------------------    
 919 UniRef50_O80488        Pentatricopeptide repeat-co...   323     1e-86  1  12.1%  71:525     7:474     ----------------------------------------------------------------------LLRLLHGHNTRTRLPEIHAHLLRHFLHGSNLLLAHFISICGSLSNSDYANRVFSHIqpNVLVFNAMiySLVGPPLESLSFFSSMKSRGIWADEYTYAPLLKSCSssDLRFGKCVHGELIRTGfeRNVVVWNLMIRGFCDSGDVERGLHLFKQMSER----SIVSWNSMISSLSKCGRDREALELFCEMIDQGFDPDEATVVTVLPISASLGVLDTGKWIHSTAESSGLFKDF-----------ITVGNALVDFYCKSGDLEAATAIFRKMQRR------NVVSWNTLISGSAVNGKGEFGIDLFDAMIEEgvAPNEATFLGVLACCSYTGQVERGEELFgmMERFKLEaeHYGAMVDLMSRSGRITEAFKFLKNMP----VNANAAMWGSLLSACRSHGDVKLAEVAA--MELVKIEpnSGNYVLLSNLYAEEGRWQDVEKVRTLMKKNRLRKST------------------------------------------------------------------------------    
 920 UniRef50_Q9SZ20        Pentatricopeptide repeat-co...   323     1e-86  1  13.0% 116:545    66:492     -------------------------------------------------------------------------------------------------------------------FDDAFHLFVLmlPSIVEFNKVLTAIaqMYDVVINLWKRIENAeiSPDLYTCNILVNCFCRcqPSSALSYLGKMMKLGIEPDIVTASSLVNGFCLSNSIKDAVYVAGQMEKMGIKRDVVVDTILIDTLCKNRLVVPALEVLKRMKDRGISPNVVTYSSLITGLCKSGRLADAERRLHEMDSKK------------INPNVITFSALIDAYAKRGKLSKVDSVYKMMIQ--MSIDPNVFTYSSLIYGLCMHNRVDEAIKMLDLMISKGCTPNVVTYSTLANGFFKSSRVDDGIKLLDDMPQRGVAANtcNTLIKGYFQAGKIDLALGVFGYMTSNGLI-PNIRSYNIVLAGLFANGEVEKALSRFEHMQKTRNDLDIITYTIMIHGMCKACMVKEAYDLFYKLKFKRVEPDFKA-YTIMIAELNRAGMRTEA----------------------------------------------------------    
 921 UniRef50_B8LKN6        Putative uncharacterized pr...   323     1e-86  1  11.4%  69:464    15:394     --------------------------------------------------------------------SLLLHLCTKAKTLAEAKQVHAHMLLTGILRIPSVEtlLNLYVKCGSLPDARLAFDNMTKGDVFPWNVMIGGYVKHGeeALELYHQMQksSTNPDNYTYSSVLNACAraSLSEGKLIYDEIISKGCEMDVIVENALINMFMKCGSIEDARRVFDKMCER----NLVSWTAMVSGYAQGGFADEALRMFYEMQGEDVKANYVTVASVLPACAQLSDLQQGKEI------------HGYIIRRGLDLGIVVGNALIDMYAKCGSIGSAQKVFDKMLQR------DVVSWNVAIAGYAQNGRFDECMELFRKMQCAGLKIDVITWNTLITAYAQNGYGDQTLELFQQMQLRGVKPntIASVLSACAAVSALQEGKRI-HDLVNRSECKSDICVGNALIDM-------------------------------------------------------------------------------------------------------------------------------------------    
 922 UniRef50_Q9FLS9        Pentatricopeptide repeat-co...   323     1e-86  1   9.7%  27:532     3:498     --------------------------NYSSCSYLLKLCRTLKHLHQFHAQFITSGRISNDFKQNSVFANVLFAITSISPSASASKEVVSYATSVFRfpSTFCFNTIIRICTLHEPSssSKRFFVEMRRRSVPPDFHTFPFVFKACAAKklHCQALRFGLLSDLFTLNTLIRVYSlaPIDSALQLFDE----NPQRDVVTYNVLIDGLVKAREIVRARELFDSMPLR----DLVSWNSLISGYAQMNHCREAIKLFDEMVALGLKPDNVAIVSTLSACAQSGDWQKGKAIHDYTKRKRLFI------------DSFLATGLVDFYAKCGFIDTAMEIFELCSDK------TLFTWNAMITGLAMHGNGELTVDYFRKMVSSGIKPDGVTFISVLVGCSHSGLVDEARNLFDQMRsyDVNRehYGCMADLLGRAGLIEEAAEMIEQMPKDGGNREKLLAWSGLLGGCRIHGNIEIAEKAANRVKALSPE-DGGVYKVMVEMYANAERWEEVVKVREIIDRDKKVKKNVGFSKVL-----------------------------------------------------------------------    
 923 UniRef50_Q6YWP8        Os02g0730900 protein n=3 Ta...   323     1e-86  1  12.0% 143:584    39:484     ----------------------------------------------------------------------------------------------------------------------------------------------SGDAALRAFRRHHLDGRALHAnaLVPALAACArsAAAEAEQIHALLVKSGDPPSvyASTSLVRVYARLGRLGDARKMFDGMPVK----TVVSWNVLLDGIVRASDLDAAWEVFVEMPER----NVVSWNTVIAGFVRHGWVQEAVDLFAEMTVVYgrCAHGYVIRREFSLDGALGVALINMYTRCGSMGDAFRCFSSVASK------NVEQWTSVIGGFAAHGHPEMALRLFTEMRQLGIEPNGVTFLAVLNACSHGGLVNEGFKYFNLMRSMGIKSthYGCLIDLLGRAGFLEEAFELASSLPED----PGLVIWSSLLAACQSHGNVEMAEVAARKLAHA--EPRHgsYVLLSNTYARAGQWEDLKRTRRKMEEYGVMKQP----------GLSWIELDGRVHSFVSADKLHTESEDIYQMLEDLKVNLVSAGSEPETLA-------------------    
 924 UniRef50_B9RPP2        Pentatricopeptide repeat-co...   323     1e-86  1  10.2%  70:490    46:447     ---------------------------------------------------------------------TVLQTLTSIKSFTKGQQLHAYIITSglQNNTYLSTKLAAFYASCGLMAAAQIIFDGIvkNSFLWNFMIRGYACNgvKALFLYQDMSSFGQKADKFTYPFVIKACGDLrvEFGWRVHCEVVITGFNLDIYVGNSLLAMYSKFGNMKMARMVFDRMPVR----DLTSWNTMISGYLKNGKPREVLAIFNLMKQIGLSVDDMTLIGLLCTCAELFAEKQGKEIHGYVVRNRHS-----------VFNHFLINSLIEMYCKCNSMVDARKLFEHMAWK------DTVSWNSMISGYARNRDAFESLRVFCRMVLEGTKPDKITFITVLGACEQITAMEFGRSVHSYLSKKGFSATIftALIDMYAKCGNLACAHLVFEEMPEK-----NLFCWSAMISGYGIHGMGREAISLFHEMIKNHIIPDE-----------------------------------------------------------------------------------------------------------------    
 925 UniRef50_B9RWW5        Pentatricopeptide repeat-co...   322     2e-86  1   9.6%  19:602   138:735     ------------------LRNYGMLPSSFTFCSVIHSFVLQGNMSGAIQVLELMNDEKINYPFCNFVCSSIVSGFCKMGKPELAMGFFENSLKLG-----------------------------ALKPNLVTYTAVVSSLcgRADEVFDLVCEMEEEGLAFDVVFYSCWICGYFRNGVfeAIRKHKEMVKKGISSDTIGYTILIDGFSKEGSVEKSVGFLHHMLANGSEPNLVTYTAIILGFCRKGKIDEAFAIFKLVENLGIKLDEFIYAILVDGFCLKGDFDRAYQLIEEMEKK------------GITPTIVAYNILINSLCKAGRTFDADEV-------SKALQGDKITYSALLHGYIKEENSIGILEVRQRLEEARIQMDIIMFNIILKALFVVGAFEDVLVLYNGMQEMNLVAntYCTIIGGFCKVGRIDEALEIFDEF--RHGLGSSVACYNCMINGLCKNGMVDMAAEIFVELIEKGLTLDIGICMTLIKAIVKEKSADGVLDLIYrmVARRNKLVLTSkyYLIIKGLIGDGkvKSALYFFNKMKEDntFPEGYLLDAKQLFESMVLKGFKWN-IRIYNSFINGYCKFGQFEEA-    
 926 UniRef50_Q2R484        Os11g0483950 protein n=5 Ta...   322     2e-86  1  13.3%  77:527    27:470     ----------------------------------------------------------------------------RCATPRALAAVHAAMLVSGRldDAFAASRLLAAHAALSPPGAVLRLLASLppNSFMLNITLRALASSPdsALRFFSLLRRGSGGGGghTFTFLLKASARLPLRASEQLHALAVRHGLERDAYVANGLVRAYSLAGLVPLARRVFDGLPER----SAVVCTTMVSGYAQNGMHEDAMRSFEEMVGDGIEPHGAALASVLSSCARSGslEMGRRVHELMESRRVTAPVVG---------AILGTALVDMYAKTGAMEEATAVFDRMPERQ------TATWNALITGLAHHGHGEVALATFHRMRRDGVPPNGATLVGVLSAYGCTGRLDEARRVFASMEKDfpTIQHYGCMVDLLGRSGLLTEAEEMIRGM---TTCDADTVIWGALLNACKNHGDIDVAERAVQEMLK--LDpnHGVYVVLSNMYAEAGRWQDVDRLRKVMKRARLSKIPGS----------------------------------------------------------------------------    
 927 UniRef50_Q5Z851        Os06g0700100 protein n=2 Ta...   321     5e-86  1  11.8%  84:516    24:440     -----------------------------------------------------------------------------------ACQIHARALKLGVLPSSLhcSALVKSYAASGSLAAARKLFDEIPrdVPLWNTLLSACARSgqHALVTASTMARAgsRPDNVSVTILLSACARLrlVHGREIHGYAVRNLAALDLPLLNALVSMYGRCGRLVNARMVFDSI---GSMKSVVSWTCMINACCENGKPAEALQVFEQMRLAGVKVDEVTLLAVISACTML-----------DCRSELGEWVEEYAHENGFLENTRVANALIHMHGKMGRVRRSCEIFDSITVR------TVVSWTAIIQALAVHGHGVAALVQFSQMLRQGFQPDELVFLSVINACSHSGLVNEARQLFKSmeEYHITpeHYGSMVDLLCKSGMLEEAFEFVLAMPVR----PDPVIWRVLTGACRDLGNAILARKVVDHVIE--MEPEygNYVLASNLYAANEDWRRVVDVRMEM---------------------------------------------------------------------------------------    
 928 UniRef50_D1ISY3        Whole genome shotgun sequen...   320     7e-86  1  10.3%  70:603    24:532     ---------------------------------------------------------------------SLIKTC---KSMAQLKQIHSQTICTGLISNPIVPAQIIAFCCKhgDMEYARMVFDTMP--------------------------------GPNHFVWNNMIKGYSRVGCpsAVSMYCEMLERGVMPDEYTYPFLLKRFTRDTAVKCGRELHDHIVKLGFSSNVFVQNALIHLYSLSGEVSVARGVFDR-SSKG---DVVTWNVMISGYNRSKQFDESMKLFDEMERMRVL------------PSSITLVSVLSACSKLKDLNVGKRvfDKMPER------DFVSWTAMIDGYLQVNRFKEVLSLFREMQAANIKPDEFTMVSILTACAHLGALELGEWIKAYIDKNEIKIDSfnALIDMYFNCGNVEKAIRIFNAMPHR-----DKISWTAVIFGLAINGYGEEALDMFSQMLKASITPDEViyVLLCNIYAACNRWEKLHEVRKLMMDRGIKKTP----------GCSLIEMNGSVHEFVAGDQVHPQSKEIYSKLDEMSVDLKFAGYSPDTSEVFLDI-------GEEEKES    
 929 UniRef50_Q2QYQ4        Pentatricopeptide, putative...   320     7e-86  1   7.7%   6:552   137:664     -----MYAKHGRIAAAQRAFAHIRNKDLTCWNAMLEGYVSNGFGHHAISTVLVMHH--SGLAPDRYTYISAVKACSISAQWGLGRQLHCLVIHSMLESNTSVMNSLVDMYFRARQKETAASVFRKIrkDTVSWNTMFSGFAHDEDdvFGYLIDMSRTGFKPNEVTFSVLLRLsaKENASLGLQIFALAYRHGYTDNVLVANAVINMLFRCGLLDRAYGFFCSLTFR----NIVTWNEIIAGYGLFSHSEDAMRLFRSLVCIGERPDEFTYSAVLSAFQEAH------------GARDHEQIHAIILKQGFASCQFVSTSLIKaaAAFGSVQSSLKIIED------SGKMELVSWGAIISAFLKHGLNDEVIFLFNLFRGDStkPDEFILATVLNACANAALIRHCRCIHSLVLKTGHS-NHfsAVVDAYAKCGEITSAESAFTAVSSATN---DAIMYNTMLTAYANHGLIHEALNLYEEMTKAKLNPTPATFVAILSACSHLGLVEQGKLAFSTMLSAYGMHPARANYACLVDLLARKGLLDEAKGVIDAM---------------------------------------------------    
 930 UniRef50_Q6K7Q1        UDP-glucoronosyl/UDP-glucos...   320     9e-86  1  13.6% 115:577   125:577     ------------------------------------------------------------------------------------------------------------------RPHDALRVFDHMRAREVVPDAHAsaKSRMTATARKVFDQMTRAGVAMNTHVYNAMLHVCLkgDAALAESLMTRMDAAGVPLDRFSFNTVIALYCRKGMQYEAMCVRERMENQGVKADVVTWNSLIHGLCKERRVKEASQLLREMAMAGVAPDHVTYTTLVDGYCRAGDLEEAVKLRGEMEA------------MGMLPGVATYNAILRKLCEDGKMKEVNVLLNEMDER--KVQADHVTCNTLINAYCKRGDMTSALKVKRRMMESGLQLDQFTYKALVHGFCKAKELDEAKEALFEMMGAGFSPN-YSVlvDGLCNKNNAVAVLAIPDELMKRG-FPPDKAVYRSLIRRLCKKGFIDLAGNVFNEMQGKGLEADCLVYATLACAYLTAGKPVAALDILNEMAKKQLYITPQI-YNCMCTS--YADEKGSLNMLWVHAIERGLITKSVYKVMHQ--ARMKSSN--------------------------    
 931 UniRef50_B9HQ22        Predicted protein n=1 Tax=P...   320     1e-85  1  13.4%  71:599    96:675     ----------------------------------------------------------------------LLRKLVDLGEPLSAHEFydHVIARgePDSNIVNSMIICLAKLGKLDDAVRLFDRLIGSDDCVLSNAACIMIlrFLEAFDYLVRISDANVKLGMWVYTVLINGLCHQrvGEAIQVFDIMCKrgSPPTLHMFKTLFFGLCKAGWLVEAELVFEEMEVQGFFVDKVMYTSLMNAHGKNKKMKMAMRVYFRMLKKGCDPDICTYNTLIHGFLKMGLFDKAWALWSLMSDSGIQPNEA-IAKNGCGLDrvAGSIYVSALCEDGKTETALACFKNVASAGC--IPLLFTFNSLIKRLFQDGLFEDVKSLMDIMQNEGIVPNLETYLIMVNEYCKQEDLASAFGILDQMKEMGPKPSvyDCIIACLSQQRRISEAETLFHRMRRDGL-DPDEVAYMTMINAYARDGRGDKALNLFEKMIENAIQPSSYSYTSLISG----ELKDLASKLMQKIKGTRF-MPYLYLYNIIIFGFCWANMMQDAHHQFKLmlIGAHGRAGEIDRAIGLFNRMSADGCTPDR-CAYSTLLKSLCESGDP----    
 932 UniRef50_D1H8Z4        Whole genome shotgun sequen...   319     2e-85  1  10.8%  63:582    34:541     --------------------------------------------------------------PKSYWYSS--STFSDSMNYPNSEVLHAKLIKNGCVG--IRGNHLLNLYAKSQNLEQAHKMFEEIPqdVFSWTVLISGFARIGLSalGLFTKMQDQGVCPNQFTLSIVLKSCSSndSRIGKGIHGWILRNGLDLDAVLNNSILDYYVKCRCFGYAEKLFGLMAEK----DTVSWNIMMSSYLQIGDMQKSVDLFRQLPGK----DAASWNTMIDGLMRNGCERVALELLYKMVA------------AGPAFNKLTFSIALVLASSLSVLGLGKQIHTQVLKVG--VLDDGFVRNSLIDMYCKCGEMEKASVIFKHLPQE-----SSMMNsvVSACASAGVLELGRQVHGYIQKIGHGLDVfsSIIDMYVKCGSLNDAWLIFNQAKDR-----NVVLWTSMISGCALHGQGREAVRLFELMINEGITPNEVSFVGVLTACSHAGLLEEGCKYFRLMREVYGIRPGAEHFTCMVDLYGRAGRLNevWRSFLSSYRSHPQDTKIYSYLDELIGRLKEIGYSTDV---------------------    
 933 UniRef50_Q9SAA6        Pentatricopeptide repeat-co...   319     2e-85  1  10.4%  21:538   143:652     --------------------ACGSSPDV--FDSLVRACTQNGDAQGAYEVIEQTRAEGFCV--SVHALNNFMGCLLNVNEIDRFWKVYKEMDSLGYVENVNTFNLVisFCKESKLFEALSVFYRMLkpNVVSFNMMIDGACKTGDmaLQLLGKMGMMsvSPNAVTYNSVINGFCkgRLDLAERIRGDMVKSGVDCNERTYGALVDAYGRAGSSDEALRLCDEMTSKGLVVNTVIYNSIVYWLFMEGDIEGAMSVLRDMNSKNMQIDRFTQAIVVRGLCRNGYVKEAVEFQRQISEKKLVE------------DIVCHNTLMHHFVRDKKLACADQILGSMLVQGLSL--DAISFGTLIDGYLKEGKLERALEIYDGMIKMNKTSNLVIYNSIVNGLSKRGMAGAAEAVVNAMEIKDI--VTYNTLLNESLKTGNVEEADDILSKmkQDGEKSVSLVTFNIMINHLCKFGSYEKAKEVLKFMVERGVVPDSITYGTLITSFSKHRSQEKVVELHDYLILQGVTPHEHIYLSIVRPLLDR-----------------------------------------------------------------    
 934 UniRef50_Q8H3J6        Selenium-binding protein-li...   319     2e-85  1   9.5% 148:600   126:578     ---------------------------------------------------------------------------------------------------------------------------------------------------LTLFSRFRRR----NPHSFTFLLAssASACSFLGSHVHALAVKAGAAGDLFVRNALVHFYGVSGDVGAMRRVFDELPR---VRDVLTWNEVLAGYVRAGMMTVAREVFDEMPVR----DEISWSTLVGGYVKEEELEVALGVFRNMVEQggKFVHNVVQRSGMPVCMNVGAALVDMYAKCGCVAVAREVFDGMRRR------DVFAWNAMICGLAAHGLGRDAVELFERFISEGLSPTNVTFVGVLNGCSRSGLVAEGRRYFKLieKYRIEpeHYGCMVDLLGRAGLVPEAIELIEGM----HIAPDPVLWGTILSSCKTHGLVDLGVSVGNKLIE--LDPTHSGYYVLLSgyAKANKWDEVREVRKLMSSRGTNKS----------AGWSLMEAHGKVHKFLVGDTYHKDSVQIYDTLDMINKRLTEAGYVPDV----SSVLHDIGEEEKVH---    
 935 UniRef50_D1IH11        Whole genome shotgun sequen...   319     2e-85  1  11.4%  46:581   138:643     ---------------------------------------------RAQSIIQRLRNERQLHRLDANSLGLLAVSAAKAGHTLYAASLIKSMLRSGYLPHVKAWSAVVSRLAASGDDgeALKLFDSVTRRIHRFTDAT-----------------LVADSRPDTAAYNAVLNACAngDTKRFLQVFEEMTQLGAEPDVLTYNVMIKLCARVDRKDLLVFVLERILDKGIQLCMTTLHSLVAAYVGFGDLETAEKLVQAMREGryAPDSRIYTTLMKGYMKEGRVTDTVRMLEAMRHQDDSTS---------QPDHVTYTTVVSALVKAGSMDRARQVLAEMTRIG--VPANRVTYNILLKGYCEQLQIDKAKELVREMVdeGIVPDVVSYNTLIDGCILVDDSAGALAYFNEMRARGIAPtsYTTLMKAFALSGQPKLANKVFDEMLRDPRVKVDLVAWNMLVEAHCRLGLVEEAKKTVQRMRENGFYPNVATYGSLANGIALARKPGEALLLWNEVKER---CPDEGLLDTLADICVRAA---------------------FFRKALEIVACMEENGIPPN----------------------    
 936 UniRef50_B9MYV4        Predicted protein n=2 Tax=M...   319     2e-85  1  10.8%   6:599    96:772     -----LIRKLVDLGhaRevVARGIEPDSNIVNSLVICLAKLGKLDDAVKLFD--RHIGSgcLVSNAACSTILKGFYEQDKFVEAFDYFVRISDANVKLGMWAYNVLIDGLCQQGYVGEAIEVLDIMCrtGLPPttLFYGLCKRGweAEWIFEEMEAQGFFVDKVMYTSLMNAYGrkKMKMALRVYFRMLKNGYDPDICTCNTLIYGFSKMGLFDKAWVLYNLMNDLGIQPNEVTYSIMIHNYCKKGKLDCAMSLLNSMAPCNLTPCVHCYTPIMVTLYKLNRCLEVDEWCERMLESGIVPDhqAIAKNGCGLDCSSLtiYVSALCEGGKTESALDCLENMVNAGC--VPLLFTFNSLIKRLFQDGLSEDVKSLIEIMQNWGISPNLETYLIMVNEYCKQEDLALAFGILEQMDEMGLKPSvyDCIIACLSQQRRISEAETLFCRMLENG-VDPDEVAYMTMINAYARNGKGVKALHLFEMMIKNAIQPSSYSYTALV-----AGNRRLSIEAHAENKRTGF-MPNLYLYNVTVSGFCWVNLIEDAYHQLRLmlIGAHGRAGEIDRAIGLFNRMNADGCspDR-CTYNTLLKSLCRSGRE----    
 937 UniRef50_UPI0001985272 PREDICTED: hypothetical pro...   318     3e-85  1   9.7%  71:566     5:432     ----------------------------------------------------------------------MLRILQHCKNIRELKQTHLQIFIHGLQHSDFMLPKLITVSSAFNLLDYATH-----------------------KEALRVYSRMKALMVEANSFTFTFLLKCFETSqlEDGRVIHGEILKLGFGSSVFVQNSLLGFYANCsnLGSAYRVFEEMPER----DVISWNSMISAYMTRGEIQSAIGLLDKMPER----NIVTWNSVVCGLSKAGNMXXXXXLH---------------------LSIPLGNVLIDMFAKCGDVENAKEVFHHMSKRC------IITWTTMVSGLAVNGKCREAINLFDKMCLEGTKPDDVIFIAVLSACTHGGLVEEGKRVFDQMveFGIKpeHYGCMVDLLGRAGKLEEAVRFTARM----HLKPNAVIWATLLFCCKIHGNGDLLKSVTEKIMDQ--EPSNPSYLTLVSNLSAsgRWEDVLSFRVAMRQQRMEKVP----------GCSSIQVGNRVHEFLAKDTRHVQRKEIYRAL-------------------------------------    
 938 UniRef50_Q9LJI9        Pentatricopeptide repeat-co...   318     4e-85  1  10.9%  67:569    10:496     ------------------------------------------------------------------SWKSLILASQRCNTVKQIKSTHSLFIIHGLHRNTYAIslLTAFLHLPNLNKhaSSIFDSIepNSFVYDTMIRICSRspHLGLRYFLLmeEEDITPSYLTFHFLIVACLKACFFsgKQIHCWVVKNgfLSDGHVQTGVLRIYVEDKLLFDARKVFDEIPQ----PDVVKWDVLMNGYVRCGLGSEGLEVFKEMLVRGIEPDEFSVTTALTACAQVGALAQGKWIHEFVKKK-----------RWIESDVFVGTALVDMYAKCGCIETAVEVFEKLTRR------NVFSWAALIGGYAAYGYAKKATTCLDRIERegIKPDSVVLLGVLAACAHGGFLEEGRTMLENMerYGITpeHYSCIVDLMCRAGRLDDALDLIEKMPMK----PLASVWGALLNGCRTHKNVELGELAVQNLLDlnVEEEEAALVQLSNIYFSVQRNPEAFKVRGMIEQRGIRKTP----------GWSLLEVDGIVTKFVSGDVSHPNLLQIHTLIHLL----------------------------------    
 939 UniRef50_Q69P44        Os09g0423300 protein n=5 Ta...   318     4e-85  1  10.9%   7:521   412:957     ------YGKVSDMQKAMEVFDrgLAPDETTYRSMIEGFGRADKYKQAILYYRKLR--NSGFKPNASNFYTMINLLARHDDSEGATEILEDMRAAGCQCSSIVTVLVRAYGSVGRMHKVLqcFYKKILFDATSCSILVTGFVQnvEEAMRVLREKKWKDSDFEDNLYHILICSCKEAGcdDAVRIYNQMPKSATHPNLRIYCSMIDVFSIMERFTDAEALYLELKASSCVLDMIAYSVIVRMYTKAGRPEDACLVLEDmkQKEIVPDKYLFLDMLRTYQKCGLLEKLSDTYYWILKSQVerIFDEMIQQGHLANTVTLNVLLDIYGKAGLFNKAEKVFLMARKQGMA---DIISYNTIIAAHAKNGDFRSMIYFVQRMQEAGFPVSLEAYNCMLDAYGKAGQLEEFAAVLQKMERAGCEFDhyNIMINIYGRKGWIEGVANVLAELKSRGG-EPDLYSYNTLIKAYGIAGMPEDAVKLMQEMRIKGIAADRVTYTNLIAALQRNENFLEAVKWSLWMKQTGV----------------------------------------------------------------------------------    
 940 UniRef50_Q6ES36        Os09g0412900 protein n=6 Ta...   318     4e-85  1  10.0% 143:580    84:492     ----------------------------------------------------------------------------------------------------------------------------------------------DPSTAAALFLRLRRGGLRPDAHTLPHVLKALArgSLPVVGSVHSEAVKDGLA-SAVVynALMAAYSACGQLERAVQVFDEMPRR----TVVSWNTALTACAGNGRHELCVELFAEMLEAGSVPDPTTFVVMLSAAAELGNLALGKW------------AHGQVVSRRLDMTLQLGTAAVNMYSKCATVSYALRLFERMPAR------NVWTWTAMIVGFAQNGLAQEALELFDRMKLSKIIPNYVTFLGLLCACSHAGLVDEGRRFFHEMQhyGIVptHYSAMVDVLGRNGLLQEAYDFVVGMP----VEADPVVWRTLLAACQLHS-SKDCIEIIDKVQGkeLEPrsGNYVMVSNIYCEIGSWDEAAKARRVMREGGMKKVAGE----------SCVEVGGRVHRFIFGDDSCPEFHGACRILHELNLNMRK--CEP-----------------------    
 941 UniRef50_Q5NBA9        Fertility restorer-like n=4...   318     4e-85  1  10.6%   5:601   244:914     ----SLVHGYMSSGLLMkmLDKGIAADVVSYTTVIDGLCRERSVEKVMGFLDEMN--RRDAKPNLITYTSLIGGFCKRNRLDDAFSIVRKLEQTGVVVDEYVYSILIDNLCKKGYLDRAFSLLEEMDKKgvTYNSVINGLcgHTKKAVEIYerLESSGVTIDVVTCNVLIKAlmIKKVDDACSLFLRMPEMRLRPNIVTYHTVIDMMCKLGETDRALQLFDDYKEDILFSSTIVHNCLIIALCNGGKVNMAEQIFNDLIQRNLRPDSCTYKKLIHTHFKEGGEHGVldLFSSLCnqGALDVYKLLRMKSFTVTSKTCYRLLKCLLRNGHEQTIQPLLSQFIKIHGLDEPRMI---NMLSCHlrEGRILDACNFLKEAEQNGYSVDLAMYSILVEGLCKSGYLEKALDLCESMKEEGIQPNihNSVLNGLCQQGCLTEAFRLFDYL-ESSEVLPTLVTYSILIAALCREGFLDDAHELFQKMSTKGIKPTTRVYNLLISGYCNYGLTEKALELISHFEEI-FLFPDAITIGAIINGHCLKGDIESALGFFSEYY----------------HKEM-----VPDFVG-FMSLVKGLYAKGRMEE--    
 942 UniRef50_Q9ZQA1        Pentatricopeptide repeat-co...   318     5e-85  1   9.4%  78:568    23:487     -----------------------------------------------------------------------------CSSIKHLLQIHGQIHLSSLQNDSFIISELVRVSSLSLAKDLAFalLHSSDSTPSTWNMLSRGYSSspVESIWVYSEMKRRGIKPNKLTFPFLLKACAslGLTAGRQIQVEVLKHGFDFDVYVGNNLIHLYGTCKKTSDARKVFDEMTER----NVVSWNSIMTALVENGKLNLVFECFCEMIGKRFCPDETTMVVLLSACG--GNLSLGKLVHSQVMVRE------------LELNCRLGTALVDMYAKSGGLEYARLVFERMVDK------NVWTWSAMIVGLAQYGFAEEALQLFSKmkESSVRPNYVTFLGVLCACSHTGLVDDGYKYFHEMEkhKIKpiHYGAMVDILGRAGRLNEAYDFIKKMP----FEPDAVVWRTLLSACSIHHDEDdgEKVKKRLIE--LEprSGNLVIVANRFAEARMWAEAAEVRRVMKETKMKKIAGE----------SCLELGGSFHRFFSGYDPRSEYVSIYELLDL-----------------------------------    
 943 UniRef50_B6SVA6        Leaf protein n=3 Tax=Androp...   317     6e-85  1  12.8% 141:593    80:497     --------------------------------------------------------------------------------------------------------------------------------------------SKKPYQAHSVFKHLMDEGHKPSLVTYTILLTALTNQRMFESisLLAEVELAGVRPDSIFFNALINAFVEAKRMGEAINTFWKMKHSGCHPTTSTFNTLIKGYGIVGKPEESQRVFDMmvEGSVRPNLTTYNILVKAWCDHRNLEEAWGVVGKMQA------------GGFEPDIVTYNTIASAYANNDETWRAEEL---IVEIQTRVRTSERTWGIIIGGYCREGRLEEALRCVQQMKDAGSVPNVVIFNTLLKGFLDANDMAAVNKILGLMKKFGIKPdtYSHQLNTFSSLGHMAKCMHVFDKMIEAGI-EPDPQVYSILAKGFVRAQQPEKAEDLLLQMSHLGLCPNVVTFTTVISGWCSVADMESAMRVYDKMCKSGVYPNLRT----------------------FETLIWGYSEQKQPWKAEEVLQMMRETGVKP-KQSTYCLIADAW----------    
 944 UniRef50_Q7XV94        OSJNBb0012E08.2 protein n=3...   317     9e-85  1  10.1% 142:519   112:463     ---------------------------------------------------------------------------------------------------------------------------------------------RHLPLALSLHAVAVTRNLLPFPHVSNALISLYARSALPdaRRVFDGMR----APDVVSHNALVHGYVKAGRLGLAVRVFEGMPER----DAVSWGTVVAGCAKAGRLEKAVRLFDRMRREGYRPDDVALAAVLSCCAQLGALDKGQEV------------HEYVRRTRPRPNVYLCTGIVDLYAKCGRVEVAREVFDACPEK------NVFTWNALIVGLAMHGHGTVALDYFDRMLVEGFRPDGTTFLGVLIGCSHAGLVDTARRIFYEMqnHGVPRehYGCMADLLGRAGLIDEAMEMISSMP----MEADTYVWGGILAGCRMHGnvEFAEVAARRLLELNPD-DGGVYSAMAGIYADAGRWEDVARVRRSMDEM------------------------------------------------------------------------------------    
 945 UniRef50_D1HCJ8        Whole genome shotgun sequen...   317     9e-85  1  10.6% 119:579    86:586     ----------------------------------------------------------------------------------------------------------------------ARKVFDElrRNVVSWTTLMTGYVHdpELAVQVFREMLEAGAYPTNYTLGTALSASSDLHSKegKQIHGYSIKYRIEFDASIGNSLCSLYSKCGSLECAVKAFRRIRDK----NVISWTTVISAWGDNGEAATGLQFFVEMLSECVEPNEFTLTSALSLCCVMQSLDIGTQIHSLTIKLGFESNlkKLFDEMETISLVTWNAMIAGHARMMDFaeALSIFLKLNRSG--MKPDLFTFstIKTGfvVNGQPQQALLLFEDMRLAGVRPNKITFVGVLSACSHAGMVDEALDYFQMMkeYKITpdHYACLIDMFVRLGRLDEAFDFIKEM----DLEPNEFIWSILIAGCRSQGKLELGFYAAEQLL--NLKpdTETYNLLLNMYLSAGKWKEVSRVRKMMKEEKLGRLKD----------WSWISIKDKIYSFKRNARSHAQSGEMYELLGNLHEKAKSFGYE------------------------    
 946 UniRef50_Q0DWE5        Os02g0818800 protein (Fragm...   317     9e-85  1  10.2%  36:514    80:589     -----------------------------------RLLCPHLRSGNPSATLRLFLRVLRDRdpDSSSFSAALAACSHHASPSPGFSIHAFLLKSGFASDVFAANSLLHFYGSFGLHSLARNLFDEMprDTVSFNTLISSYVHSCcdAFEVFRIMVDSGLRPDGWTVTALIGACAEldLRAAKAVhrRMLEPRVFHSGEVATSLVDAYVKCRSMELARQVFdeMPEK----DTVAWTALIGGFVRAGRYKEAVVLFEEMEEAGFEADEATIVTVLSACVGYGNIDLAKRLHCLVGRDGL-----------INRNAKLATTFVDMYAKHGCIQTAQEVFSGVDDDFKTLEL----FNAMINGLARCKFGEKAIALFDEMGSLGLHPDKITFVCVLSACRHSGLVSQGFHIFDSmeKYGVKpeHYTCMADLLARDGQLDDAYIFIQNMP----FKANSVVWSSLVRACMLHGNIKI-RKLAEE-QLLWLDPNYKPENllSNLFSEGKRKERTARVRK-----------------------------------------------------------------------------------------    
 947 UniRef50_Q8L4S1        Crp1-like protein n=5 Tax=P...   317     1e-84  1  10.7%   5:601    87:735     ----ALLSRQRHWRRARDLLDRmyAPSAIHYGVVLRHLAGARRWGELRRVWARMAREGA-PPPTNQAYAALADALAKAGHARDSLLLLRHMRARGVAPDEVSMSTFVRILKDVGRYSDAATLFDNWcrFEVGFIDLDYSALDSGGPMQFLLEEMCHGNVdpkaVTYNTMIDLYGkgKLKDAMDMFMGMPDYGVAADTCTFNTLINIFGSCGNVKEAEALFASMIVRGTNPDVKTFNVMMTLFASKGDVEGVLKHYYHIGKMGLSADTVSYGIVLRVLCERKMVHEAEDVIVGIMRSGTCVPEQSLPvrGDEVSSKTFAAIMDAFAERGLWEEAEHVfrGVRRRG------IVEYNVMVKAYGAAKRYDRVSPLLEHMNESGISPDECTFNSLVQMFATGGYPQRAKKLLGKMKYAGFKpeTYAAAITTLSRNYLVSEAIYLYNEMKASG-VEPNVVVYGVLIDTFAETGQLEEALHYNNLMEESGIAPNQIVLTSLIKAYSKANCWKEAQDLYSRMKSMDGGPDIVAS-NSMLNLYANLGMVTKVKEIFDCLRRNNQADDVSYTSMISLYKNmqNSGLLSDC-ASYNAVMACFVAKGKLRE--    
 948 UniRef50_B9S184        Pentatricopeptide repeat-co...   316     1e-84  1   9.6%  67:514   110:524     ------------------------------------------------------------------SYRDLLSDLLKNQKLDQARFLFAKIPS----PEAYLCTLMINGYARSGRLNDALMLFDRmvRDIISWNSMIKGccGDLGMARRLFDEMPER----NVVSWTTMVNGYLkgRVEIAERFFWDMPVR----DVAAWNAMIFGYCGNRRIADGLNLFEKMPCR----NVISWTSMIGGLDQNGKSEEALILFGKMMDSGVKPTSTTYACVLTACANAMDFNMGVQV------------HGRVSKLGYKSDEFISASLITFYSNCKQVKNAHQVFNETLSK------NVVIWTALLTGCELNSKHEDALRVFSDMIKMGVFPNQSTFISAFNSCRGLEALDRGKEIHTVTAKLGLETDVfnSLVVMYNECGNVNDAVAVFKKIKEK-----NDVSWNSIIVGSSQHGHGIWALIFFNQMIRTCVDPDEITLTGLLSACSHSGMLQKGRRFFQ-----------------------------------------------------------------------------------------    
 949 UniRef50_Q84VG6        Pentatricopeptide repeat-co...   316     1e-84  1  10.6%   6:535    85:608     -----KLCVFRRFDTVYQLLDEMppPDDAIFVTIIRGFGRARLIKRVISVVDLVSKFGIKPSLKV--FNSILDVLVKE-DIDIAREFfrKMMASGIHGDVYTYGILMKGLSLTNRIGDGFKLLQIMkpNAVVYNTLLHALCKNGkrARSLMSEMKE----PNDVTFNILISAYceQKLIQSMVLLEKCFSLGFVPDVVTVTKVMEVLCNEGRVSEALEVLERVESKGGKVDVVACNTLVKGYCALGKMRVAQRFFIEMERKGYLPNVETYNLLIAGYCDVGMLDSALDTFNDMKTDAIRW------------NFATFNTLIRGLSIGGRTDDGLKILEMMQDSDTVHGARIDPYNCVIYGFYKENRWEDALEFLLKMEKLFPRAVDRSFK-LISLCEKGGM-DDLKTAYDQMIGEGGVPSIihCLIHRYSQHGKIEESLELINDMVTRGYL-PRSSTFNAVIIGFCKQDKVMNGIKFVEDMAERGCVPDTESYNPLLEELCVKGDIQKAWLLFSRMVEKSIVPDPSMWSSLMFCL--------------------------------------------------------------------    
 950 UniRef50_Q9LSQ2        Putative pentatricopeptide ...   316     1e-84  1  12.7%  16:457   203:637     ---------------FQQMRSDGCKPDRFTYNILIHGVCKKGVVDEAIRLVKQMEQEGN--RPNVFTYTILIDGFLIAGRVDEALKQLEMMRVRKLNPnaTIRTFVHGIFRCLPPCKAFEVLVGFMEKDsvGYDAVLYCLSNnaKETGQFLRKIGERGYIPDSSTFNAAMSCLLKgdLVETCRIFDGFVSRGVKPGFNGYLVLVQALLNAQRFSEGDRYLKQMGVDGLLSSVYSYNAVIDCLCKARRIENAAMFLTEMQDRGISPNLVTFNTFLSGYSVRGDVKKVHGVLEKLL------------VHGFKPDVITFSLIINCLCRAKEIKDAFDCFKEMLEWG--IEPNEITYNILIRSCCSTGDTDRSVKLFAKMKENGLSPDLYAYNATIQSFCKMRKVKKAEELLKTMLRIGLKPdtYSTLIKALSESGRESEAREMFSSIERHGCV-PDSYT--------------------------------------------------------------------------------------------------------------------------------------------------    
 951 UniRef50_UPI00019839A8 PREDICTED: hypothetical pro...   316     1e-84  1   9.3%  86:537     6:440     -------------------------------------------------------------------------------------QIHCLMITTGMSRDAFPASRlsSFSLSPPNIHLAYVVFSQMqpDVYSYNIMIKAFSqhPLDSLHFYHKLLHQGLYPNEYTFCFVLTSCAQaaLQEGKQIQAQLIKHLSLGSVYASTSLVSLYGKCGDIRRALQAFEEMPHR----TEVSWGAVIDAYIDQGFLTNALQLYAKMRSLGHSASNAILVGAFGACAKLQDLALGR------------IIHGQVEAMGLQLNVTLGTSLVDMYSKCGAIEMAMEAFSRMPVK------SVASWNCMINSLAMNDGGSAAIGLFKEMQASGLGPNSATFLGVLHACSHGGLVNEALEYFSCMReyGIETnhYGCMVDLYGRAGKLEKAVDVIRTMPMK----PDIVIWGALLGGCKKHGNAKLGELVGAQIMK--LQPHKVscSSLSDMYAMDGRWEDVISVRRMMLDMDVGREPGFLISSQSCLEC------------------------------------------------------------------    
 952 UniRef50_Q94DX0        Os01g0764950 protein n=1 Ta...   316     1e-84  1  10.1%  61:534    16:466     ------------------------------------------------------------PPPSVTRALHAINTCTTAAALAPIRG--AILGDRALLRSTAVVSAFFLACGRLRHLDPALSLfcHPRPHVFVFNSLLRSLGRAPARslPLFRHFLrrGVRPNRFTFPL--LLTSLSSLADLRAVHCQVVKSGFGPDLHVRNALLARYADCddLAHAEQMFDEMPR----PEVVAWTTMITSYRNRGRTFHALATFRRMLAAHVAPNRVTMVAALGACAAHCAVDTGIWI------------HEYVQKQGWEMDVVLGTALVDMYGKCGKVSDGMHVFSKMAKR------NVYTWNSIIGALALAQDGKTALQWFSRMQNDGVQPDEVTLICVLCACAHAGFVDIGRKIFNLAieYGFQPghFGCMVDLLSRSGHLDDAFRVVETMPS----QPNAVIWGLLLRGCRARGDSWLSEHV--TMRLVELEpnASHYVLLSNLYAETGRWQEAQGILHWMKKKGLRKDAGWSLRMLED---------------------------------------------------------------------    
 953 UniRef50_B9RLV9        Pentatricopeptide repeat-co...   316     1e-84  1  12.7%  40:510    36:492     ---------------------------------------HSKTPEQALDTfsVLKQNPNNPTKKLHLYSAVIHYLTGAKVYPTARCL----------TKDLIQTLLQScrRVNSLVFNALSQLRGSKFNPSVFGVLIIAfvGLVDEALRVY---LKVGAFPAVQACNALLNGlkKSSFDIMWELYNNMVSRRLFPTVVTYNVLVDACCRQGDVLRAKSLISEMVKKGIEPTVVIYTTLIHGLCSESKLMEAESMFRQMKDSGVFPNLYTYNVLMDGYCKTANVKQALHLYQGML------------DDGLQPNVVTFGILIDALCKVRELLAARRFFVQMAK--FGVVPNVVVFNSLIDGYSKAGNCSEATDLLLEMEKFKISPDVFTYSILIKNACRLGTVEEADDILKRMEKEGVPAnvYNSLIDGYCKEGNMEKALEIFkdMRRNGL-RPDALAYAVMLQGHLNAKHMADVMMLHADMIKMGIVPNEVTSQVVRRGYQDNGYLKSAL---------------------------------------------------------------------------------------------    
 954 UniRef50_Q7FB20        Os04g0675600 protein n=4 Ta...   316     2e-84  1  13.6%  27:504    52:519     --------------------------TLHDYNRLLDAFARDGDGDAALRVLRRMRHSSPACAPTAASYTSAMSALAKAGRPADAAALFDDMLANGVAPDRCAFslLHVYSSHLHLPSAAHSVLVWMSRLGLPPtvFSFCRAGRLPDALQLLDEMRALNYPLTLHSYTPILQvcANADMQSADALISSMRSTGCHPDVVFYNIYVNGLCKVGDFDAVQRTIDESGRNGWVPDAVTYSTYIAGLCRFGYVEEALRQLEIMVTMGLQPTVVGLNILLDYVAQDLDMWAGKEVLERCQE------------LGFVVDVVTYNTVMDHFCKKRKWLRVLKLFTDLLKK--PITPNVQTCNIFISCLCRAGKFQFAKFVFS---SKGFMADTVTCNILIHAFYEAGKEDELGFLFADVNAGKIAPdtYNTLVDCLFRSGRRAEAVNLIRHIDDGYPVEPVAR----LAYWLVRSGNVREALRLFDDMLEKGLLLDSRIFANVIKAFCRKG---------------------------------------------------------------------------------------------------    
 955 UniRef50_B9RPH6        Pentatricopeptide repeat-co...   316     2e-84  1  12.3%  68:563    70:595     -------------------------------------------------------------------YASVLDSC---NSQNLGTQVHAHAIKTGFHCHDFVQTKLLQMYAKFGCLECAHLLFDSVprNLHSWLAILNVYFDHgdEAFSLFQELLFEDIELEFFAFPLVFKICSGLGMVegRQLHAmmKRCgsLRNEVSYNTMIVGYCAIGDVSKAKEFFDQMEVSGVKRERISWNSMISGYVDNFMFDEALNMFRNLLKEGIEPDSFTLGSVLTACADTASLRQGKEI------------HSYAIVKSLQSNTFVGGALIEMYSKCQDPMAAQLVFNEVIER------DAPTWNVLISCYARCNQNEEIRNLLQKMQEDGFEPNIYTWNGILAGYVENGHLDLAMQLFSEMHTGDVRPDIFTvlVDMYAKCGSLQYAQLAYSRISNH-----NLVCHNVMLTAYAMHGYGEEGIALFRTIRATGFQPDNVTFLSVLASCVHAGLVEhtWILYHFLRPDKTFNNKRPHYSSPCQETANNSRANACRQNIIFSDCSNNLLEEIL----------------------------------------    
 956 UniRef50_B9GHK0        Predicted protein n=4 Tax=r...   316     2e-84  1  10.8%  72:562    17:488     -----------------------------------------------------------------------IISLRTCHDFQEVKQLHAQFVVSGLLGYHPLCalLEAYVTMSQIYYAFSIFERIPSPDVFVYNTMIRGllKAPFEGKQVHCQIIKAGIVPDTHIHSSLIHMYtsGSIDDAERVLGEFSEE----NTLAKNAMISGYLTEGRVDKARKMFDDMAAK----DAASWSALITGYTKNGMHTEALALFQDMMVSHILPNEAALVSLLSACGQLGTLHQGRWI------------HAYIDKTRVLMSTKLTTALIDMYAKSGSIECGYGLFQKMARR------DVVTWGVMISAFAIHGHASKCFQLFDEMLADGIRPNKVIFVAILSACSHAGCVEEGRQYFSQMEhfGIKpeHYGCMVDLLGRAGLLADAEQLILSMPK----QPNSIIWGSLLGACRTHNDLKRGTWAFENLME--LEPTSGDRHKLAgmFANAGEKEEAAKIRKMIHEKEMVTT----------CGSSFIEVEGAVHEFTVGYTVHNKARDI-----------------------------------------    
 957 UniRef50_Q2QQL5        Pentatricopeptide, putative...   315     2e-84  1  10.5%  85:531    28:523     ------------------------------------------------------------------------------------RQLHAHLLIRGLPLPARAAVALVASSSSPR---HARAVFDSVPanVYLWTATISAYARhaGEAVALFALMLWSGPRPNAFTVTSVIKCCSKVrvHEGLQVHAFLAKAGlcSSVHVGAALLDLYGNIGLVGDARRVFDEMPVKNveRDPISWNTLMSGYLHLGDTATVRELFDEMTVR----NVNSWNTMIAACSEEGSLADTVKVFNRMRATGFeqVHGLLQKSCVEMNCHVQNSLIDMYAKCGCISQAQFLFTE------TYPKDTVSYNVMIIAFAQHGHARDALDLFNAMVDSGLQPDAVTFLGVLSACAHAGLVEYGKNYFELMkaVQQSPDHYACVVDLYGRAGLIEEAHCFVKTMPVK----PHAGVWGALLNACRKHCHVDVGEIAAKELIR--IEpnPGNYVLLRNTLARGQQWDAVEDVQLSMRRK--LLLSDCHIGC------------------------------------------------------------------------    
 958 UniRef50_C5LK59        Pentatricopeptide repeat-co...   315     2e-84  1  10.3%  18:600   156:753     -----------------DMQPSGLSPDPALCSLIVFAAVRAGRHELADMLFR----MHPKKDDDFGMHMAMIRSYGLQGDLGSALELFTELksSGELSPFVYNCLLDA-AVQCDDMDVVGKVFEEMKgdVVSFNTVMKGHlgNMRQARKTMREMQEAGFPPNVITYNELLHSMVQGKDKRgwEVVEEMKANGLPPNKVTCSILLKALTSHSHASDVVRTMELVERMRGEMDEVLFSSVIEACIRIGKLD----IlnKLQQygCrlTAPTYGSMIKAYGHAKDLRRVWSLWSEMRQRA------------VKPTAITLGCMVDALVTNRYPEDALSLIHEMLNtdCADCVNTIVYSTVLKGFALSKQVDRVFDIYQEMRSNKIPLNTVSFNTIMDACARSGSMERVSDIFREMDEQGIEPdtYSTVVKGYCLAGDVDTAFSVLRDMSgrKKFAPDEIMYNSLLDGCAKQHRVDQALQLLDEMRANGVAPSNYTLSILVKLLGRARRLLEAFNMVEDL---------STAYSFRANVHVYTCLIQACVH------------NKQLQRAMKLHDAMIQerVDPDQ-KTYAVLARGCIYSNQLE---    
 959 UniRef50_Q6K9R6        Selenium-binding protein-li...   315     2e-84  1   9.9%  36:569    96:661     -----------------------------------RLLCPHLRSGNPSATLRLFLRVLRDRdpDSSSFSAALAACSHHASPSPGFSIHAFLLKSGFASDVFAANSLLHFYGSFGLHSLARNLFDEMprDTVSFNTLISSYVHSCcdAFEVFRIMVDSGLRPDGWTVTALIGACAEldLRAAKAVhrRMLEPRVFHSGEVATSLVDAYVKCRSMELARQVFdeMPEK----DTVAWTALIGGFVRAGRYKEAVVLFEEMEEAGFEADEATIVTVLSACVGYGNIDLAKRLHCLVGRDGL-----------INRNAKLATTFVDMYAKHGCIQTAQEVFSGVDDDFKTLEL----FNAMINGLARCKFGEKAIALFDEMGSLGLHPDKITFVCVLSACRHSGLVSQGFHIFDSmeKYGVKpeHYTCMADLLARDGQLDDAYIFIQNMP----FKANSVVWSSLVRACMLHGNIKI-RKLAEE-QLLWLDPNYKPENLPLSNLFSEGKRKearkLINLMLV-GQLIFTGRVFKNLLKIFNKDIWLNNRATSMKWKQEACPCMDNGDAFADLV----------------------------------    
 960 UniRef50_Q9S7R4        Pentatricopeptide repeat-co...   315     3e-84  1  10.0% 147:515   110:468     --------------------------------------------------------------------------------------------------------------------------------------------------VWSLIHRMRSLRIGPSPKTFAIVAERYAsgKPDKAVKLFLNMHEHGCFQDLASFNTILDVLCKSKRVEKAYELFRALRGR-FSVDTVTYNVILNGWCLIKRTPKALEVLKEMVERGINPNLTTYNTMLKGFFRAGQIRHAWEFFLEMKKRD------------CEIDVVTYTTVVHGFGVAGEIKRARNVFDEMIREG--VLPSVATYNAMIQVLCKKDNVENAVVMFEEMVRRGYEPNVTTYNVLIRGLFHAGEFSRGEELMQRMENEGCEPntYNMMIRYYSECSEVEKALGLFEKM-GSGDCLPNLDTYNILISGMFVRKRSEdaGKLLLEMVERGFIPRKFTFNRVLNGLLLTGNQAFAKEILRL----------------------------------------------------------------------------------------    
 961 UniRef50_A4PRI6        Tetratricopeptide-like heli...   315     3e-84  1   7.8%  35:536    34:473     ----------------------------------ISTHMRNGHCHLALRVFDSM------PYKNLFSWNLMLTGYVKNRRLVDARNLFDLMPQKDAVSW---NVMLSGYVRSGCVDEAKLVFDNMPYKDSISWNGLLAVYVQNgeEARRLF----ESKVDWELISWNCLMGGYVKRklGDARRLFDHMPVR----NAISWNTMISGYARDGDLLQARRLFEESPVR----DVFTWTAMVFAYVQSGMLDEARRVFDEMPGKR----EMAYNVMIAGYVQYKKMDMARELFEAMPCR----------------NVGSWNTIISGYGQNGDIAQARELFDMMTQR------DCVSWAAIIAGYAQTGHYEKVMHMLVKMKRDGKSLNRSTFCCALSTCAGMAALVLGKQVHGQAVKTGYDNgvGNALLEMYCKCGSIGEAYDVFERMQLK-----DIISWNTMLAGYARHGFGRQALLVFDSMKTAGFKPDEITMLIWSN---------------NRLRKVGNTWVPTSLMNPLGNTW-------------------------------------------------------------------    
 962 UniRef50_Q6H6V8        Os02g0164600 protein n=4 Ta...   315     3e-84  1  10.0% 131:519    91:455     ----------------------------------------------------------------------------------------------------------------------------------FSFNSLIRAS--PPRLALQLFDQMRHRGVTTDRYTLPFLIHACSSGDRPLCESLHGQSLRLGYSSVLFTQTALMNAYFACRFEVAARRVFDEMQAK----DVVAWTGMVSGYVDSGMFLRGVEVFQEMreEAVRPNVATVVSVASACAGLGSLEYAKGL------------HAYVEKVGLEGELIVKNSLIDMYGKCGSIELARGLFGLMPQK------DLHSWTAMISGLASHGHGKEAVALFFSMKEAGVLPDSTTFVVVLSACSHAGLVDEGISIFNSMeeYKVTPdhYGCMVDLFSRAGLIARGYQFIESMP----FEPNLAILGALLSACSINNELDIGELVIKRIES--VRSDKGGavLLSNIYANQNLWHEVDSLRRKIRDD------------------------------------------------------------------------------------    
 963 UniRef50_C5XI41        Putative uncharacterized pr...   315     3e-84  1   8.8% 102:577     6:515     -----------------------------------------------------------------------------------------------------ATTLIKSLCARG-AVRHARALFDEMprDVVAWTAMLSGYagRHREALDLFRSMAAAGVIPNEFTLSSVLTACRAAGGGGAASIHALALRRGVDHMPYVVNALIDAYASRggLVDARCLFDAL---GEGRTAASWTSMIAGYARWGQENTGLQLFQNMIQDDVELSPFTCSIAIHACSLVGNLYFGQQI------------HVLSIRKALGANLAVANSLVDLYCTCASILDARRLFDEMPErnMDIEPniLSWTSMIGGYGMNGYVNEAIKLFNSMVHAGIHPDHVVFLGLISACSHAGLVDEGWNLFRSmlEYNIQPniYGCVINLLGRAGRLREAFILIDTMP----LTPDEYIWGALLGACKMHRNLELGRLAAGKIIE--INPDAvtYILLANICAADSKWGEYAATRVLLRGTGSSK----------EVGMSWIELSDKMYTFSTADSSSPQVSLADEVLQILVQHMDEAG--------------------------    
 964 UniRef50_Q8W0A6        Os01g0228400 protein n=4 Ta...   315     4e-84  1  10.7%  35:511    86:527     ----------------------------------MRAMIDARRPHQAHSAFRHLVDDGH--RPSLVTYTTLLAALTSQRAFDTIPWLLAEVEDAGLRPDSIF------------------------------FNALINAlaRRMGEATNTFLRMGHSGCRPTASTFNTLIKGYgaGRPEESQRVFDMMASGgvRPNLTTYNILVKAWCDAGRLEEAWRVVARMRASGADPDVVTYNTLASAYAKNDETWRAEELVVEmqQAGLRTSERTWGIIVGGYCREGRLGEALRCVRQMKDS------------GVLPNVIVFNTLLKGFLDANDMAAVDDVLGLMEQ--FGIKPDIVTYSHQLNALSSMGHMAKCMKVFDKMIEAGIEPDPQVYSILAKGYVRAQQPEKAEELLRQMGRLGVRPntFTTVISGWCSVADMGNAVRVYAAMRDAG-VRPNLRTFETLIWGYSELKQPWKAEEVLQMMQDAGVRPKQTTYCLVADAWKAVGLVENANR--------------------------------------------------------------------------------------------    
 965 UniRef50_B9RGG4        Pentatricopeptide repeat-co...   315     5e-84  1  12.6%  72:602  534:1068     -----------------------------------------------------------------------IRDAFRKNSNKWGPPVVAELRKLRRVTPDLVSEVL----KVENDPHLASQFFHwyRHNFASYNAYAYCLNRsfRAADQLPELMDSQGKPPTEKQFEILIRMHSDANRgrVYHVYQKMKKFGVKPRAFLYNRIMDALIKTAHLDLALVVYDDFKSDGLVEDSVTYMILIKGLCKFGRIDEMMEVWEEMKRDGVNPDVMAYATVVTGLCKGGRVAEGYELFKEMKE------------NKVLIDRAIYGVLIEAFVKDGKIGSACDLLQGLVDSGYR--ADLGIYNSLIEGLCNVKRVDKARKLFQIMVQEGLELDFKTVNPMLVSYAEMKRMDEFCKLLVQMERLGFsmDDISKLFSFLVRREEitLALEVFEELKVKGYI--SVLIYNTLMEALLKVGEVRKALSLFSEMKDLNCEPDSNTYSIAVICFVEDGNIQEACVCHNKIIEM-SSVPSVAAYCSLTKGLCDIGEIDEAMMLVRDCLGnhVCRSGDAEKVIEVLNEMMHENCPPNEV-ILSAIISGMCKHGTLEEA-    
 966 UniRef50_B9SQY5        Pentatricopeptide repeat-co...   314     5e-84  1  10.2%   1:521   372:955     LNAYSQQGRLEEAEQVleMQEASFSPNIVAFNTLITGYGKLSNMAAAQRLF--LDIQNAGLEPDETTYRSMIEGWGRTGNYKEAEWYYKELKRLGYMPNSSNLYTLINLQAKHDDDEGAIGTLDDMLKIGcsCSILVMTYVKnvDEALKVLGDKKWKDQTFEDNLYHLLICSCKegNLESAVRIYTQMPKSEDKPNLHISCTVIDIYSVLGCFAEAEKLYQQLKCSGIALDMVAFSIVVRMYVKAGSLKDACSVLATmkQENIIPDIYLYRDMLRIYQQCGMMSKLKDLYHKILKSEVDwlFSEMLQRGFSPNTITFNVMLDVYGKAKLFNKAKELFWMARKRGLV---DVISYNTVIAAYGHNKDFKNMASAVRNMQFDGFSVSLEAYNCMLDGYGKEGQMECFRNVLQRMKQSSYTSDHynIMINIYGEQGWIDEVAGVLTELRECGL-RPDLCSYNTLIKAYGVAGMVEDAIDLVKEMRENGIEPDKITYSNLITALQKNDKYLEAVKWSLWMKQLGL----------------------------------------------------------------------------------    
 967 UniRef50_Q7XKC9        OSJNBa0064G10.2 protein n=2...   314     6e-84  1   8.6%  55:564    27:516     ------------------------------------------------------QRHQDPIIPGIYSYASLLRRSAATSDPRLAVSLHAALLKRGFlsHIFLCNHLLIACFKSRLHRHGLRLLDEmrRNAVSWTTAIAGLtgQPREALALFKRLRRAGLPPNEFTLVSALNASSfaGVGRARQLFALAVRLGFDSNVFLTNAFLAAMVRHGQLADAVRLFD----NANAWDIVSWNTLLTAFAHRSSLRLC-TLWRRMAIEGVSADGFSFSTVLSGLSGSANV-----------AATGLQVHAQLVKSGFVDDVCVGNSLLEMYMKNKQLESGIRAFTEMRHR------DVVSWTELAAGLLHCGEPAESLRVVSDMILDGIRPNNYTIVAAANACANLANLNQGRKIHGYVIKLGGDSNVgnALIDMYAKCRSVTGAHKVFQSMQRQ-----SVVSWTAMIMAFAQNGQAREAVKVFDDMLLEGMAPNHAAFYSVLYACSQGGFVDEGWIYFDAMADKFGVEPGEDHFACMVDLLTKAGRIEEAEE----LISRMPFRPGVVA---------------------------------------    
 968 UniRef50_A9TYP4        Predicted protein (Fragment...   314     7e-84  1  12.0% 142:581     4:410     ---------------------------------------------------------------------------------------------------------------------------------------------GDHRQVRSTLKNLRRDGHRPAKETYTLLLSSlsDGRLGEGPLLLQQMEEDNVIPDAIFFNTLLNGYCEAGKMKEAEALVLEMKERGCPLITSSYNTLIKGYGLDGQPEEAMKLLQEmkDVNSKPNDRTFNMLIKAWAHDKNPEKARNVISMMRA------------AGVGPDVVSYNTLAQLFAKLGKPWDAENVITEMQ--NVYLRPNERTYGIVINGYIAAGQIDESLRLIERMKMDGISPTIAVFNHLVKGYSESMKPHKVDKVFKLMENFGVKPdtFGLIMNAWCSTGLIDEAKAVLLRMKE-YDLQPDVMAYSILVKGYSRQGRPEEGEAFLNIMIDENLKPNVITYTTVISGYCSLAQMDDAMRVFQEMRSRSVFPN---------------------MQTFRTLI-WGFKEARWPRRAEEILEYIRKAGFTPD----------------------    
 969 UniRef50_Q7XDD8        Os10g0476900 protein n=5 Ta...   313     9e-84  1  13.4% 106:545   144:576     ---------------------------------------------------------------------------------------------------------LRRYCNSGKLIQACCVIDEMVlpETKCCVRIIRGLvgKANKARHVLEVMVLSGGVPDTISCNMLIAQLCRGglSSALQVLEDMRFSGCSPSGITFNTLIRCMFNQRMYDRAISFWKEQLRLGWPPYEMTSTLLVDLVCKNCGPQKALEVLDELSLEGCQPDVVTYNALISASCKAGRLNDAKFILTRLAAE------------GLEPNGTTYCILLHALCNMRRWDEVCDLLEDM--NQVNRDPDVTTYNIFINYFCKYGHLDQAIDVLERMVINNCSPDIVTCNTLLNAISKKGMVEEALGIARCIRENGYKLvtYNTLIDALAKKAEVKKAMVLFDEMLSDGI-SPDDITYGSLVMGFCKKDMADEALDLLNQMLTLGFEVKTTTFVMVIQALCRDGKVEAAAEITKVMLSKN-SIPGNSLCSSIVTKVAKSGWNKEA----------------------------------------------------------    
 970 UniRef50_B9IEL1        Predicted protein n=1 Tax=P...   313     1e-83  1   9.6% 101:515    69:478     ----------------------------------------------------------------------------------------------------LVNTILKRLWNDGPKALQFFNLLSHHPSYSHHPSSyiSARLRDSPSLRSLVYRMRSARLGPTPKTFAIIAERYAsgKPHRAVKVFLSMHQFGCFQDLQSFNTILDVLCKSKRVEMAYNLFKVFKGK-FRADCVSYNVMVNGWCLIKRTNKALEMLKEMVKRGLTPNLTSYNTMLKGYFRAGQINEAWDFFLEMKKRD------------CEIDVITYTTVIHGFGVAGEIKRARKVFDTMVKKG--VLPSVATYNAFIQVLCKKDNVDNAIVIFEEMVVKGYVPNSITYNLVIRGLCHRGEMERAMEFMGRMKDDGCEPntYNLVIRYFCDEGEIDKALDLFQKM-TSGDCLPNLDTYNILISAMFVRKKSDdaGNLLIEMVDRGFVPRKFTFNRVLNGLLLTGNQGFAKEILRL----------------------------------------------------------------------------------------    
 971 UniRef50_D1HY71        Whole genome shotgun sequen...   313     1e-83  1  10.5%  66:579    28:547     -----------------------------------------------------------------HTKTQLIQRLQDCNDITSVSFSHTNVLKSGFLNDTFTTNHLINSYVRIRRTNDASQLFEEmePNVISFTSLMAGfvGRPKSALLIFCKMLESWVLPNAFTFATVVNACslADLRTGQKIHSHVEMFGVQCNLVVCSSLIDMYGKCNNVDDARRVFDGMDYR----NVVSWTSMITAYGQNALGECALQLFREF--NGlsPNHFMLASVINACASLGRLVSGK------------VAHAAVIRRGHESNDVVASALVDMYAKCGCIGYSDKVFRRIPD------PSVIPYTSMIVGAAKYGLGKFSLDIFKEMLDRRIKPNDVTFVGVLHACSHSGLVDDGLEYLHSmkKHGVVPdhYTCVVDMLGRTGRLDEAHQLAKSIQ----VEPdaLLWGTLLSASRLYGRVDIAVEASKWLIESniRDSAYvyAGDVSLCARG---SEVASLLRemKERGYVGGSRGLVSIDVEEEAkvGLHSERLALAFGLIskNLRMCRDCHEAFKLISEIVERDFV------------------------    
 972 UniRef50_Q2QU89        Pentatricopeptide, putative...   313     2e-83  1  12.8% 160:589   108:504     ---------------------------------------------------------------------------------------------------------------------------------------------------------------RPDALSLSFALKASARcdAHTTVQLHALVLRLGVAADVRLLTTLLDSYAKCGDLASARKVFDEMTVR----DVATWNSLLAGLAQGTEPNLALALFHRLAnsREEPNEVTIVAALSACAQIGLLKDGMYV------------HEFAKRFGLDRNVRVCNSLIDMYSKCGSLSRALDVFHSIK----PEDQTLVSYNAAIQAHSMHGHGGDALRLFDEMPTR-IEPDGVTYLAVLCGCNHSGLVDDGLRVFNSMR---VAPnhYGTIVDLLGRAGRLTEAYDTVISMP----FPADIVLWQTLLGAAKMHGVVELAELAANKLAELGSNVDGD-YVLLSNVYASKARWMDVGRVRDTMRSNDVRKVPGFSY----------TEIDGVMHKFINGDKEHPRWQEIYRALEDIVSRISELGYEP---ETSNVL--------------    
 973 UniRef50_B9SL84        Pentatricopeptide repeat-co...   313     2e-83  1  11.0% 137:518   450:807     ----------------------------------------------------------------------------------------------------------------------------------------LNSLCSQGRLQEALF-EMACQGLEMKFDGYNTLLNECVNNkiREGQRVHAHIIKTYYLPSVYLRTRLIVFYTKCDLLMDARHVFDEMSER----NVVSWTAMISGYSRRGFAFETLHLFVRMLRSDIEPNEFTLATVLTSCTDASGSELGRQI------------HSLIIKCNYDSHIFVGSSLLDMYAKAGRILEARMVFESLPER------DVVSCTAIISGYAQLGLDEEALELFRRLEREGLSSNCVTYASVLTSLSGLAALDHGKQVHSHVLRRELPFYVvnSMIDMYSKCGSLNYARRIFDSMPER-----TVISWNAMLVGYGKHGMGREVVELFKLMreNKVKPDSVTILAVLSGCSHGGFEDKGLEIFDEMLK-------------------------------------------------------------------------------------    
 974 UniRef50_D1HB84        Whole genome shotgun sequen...   311     3e-83  1  11.8%  12:529    84:578     -----------NFFHHRRAFDHMV--DVV-----------ARLTQKSGTIRGVMDELGsgCVVKAQTFLLLLRICWRGKLYGLVFDVYEEMIAFGYVPNAFVCNVIIDVLFKIKRGEVALKFLKETrsNFLTFSIALCNLCrdLVGVGDVLRMMLRKGYCPSVEIFSMVLNCFCkgRFAEAFQVLGLMATLGIPISVTVWSILIDGYRRIGKIGRAGNLLEKMVETGCSPNVVTYTTLIKGFMESGMVSRALDVLRIMGTKGCDYDLVLYNVLIDCLSKVGRYDDALGIFFSLPKRK------------MVPDSYTFSSILSAICLSQRFSLLPKLF----SSGLEVHTDLVVCNSLLSYFCKAGFPSLAVEFYNDMLDRGFMIDQYSFVGLLSGLCGSGRVDEAVNVYSGIlhSDLDAHIHTVIIGGLIKAGKFHRAVRLFKkmVERYTL--DAVSYTVAIYGLLKGGRTGEACILYNQMKEAGLAPNAHTYNIILSCFCKERDMKMVKQMIQEMIAAGVELECDTLI--------------------------------------------------------------------------    
 975 UniRef50_Q9LVA2        Pentatricopeptide repeat-co...   311     4e-83  1  11.3%   2:594   246:907     -CFCKRGCAAEAEALFDHMEVDGYYVDKVMYTCLMKEYCKDNNMTMAMRLYLRMVE--RSFELDPCIFNTLIHGFMKLGMLDKGRVMFSQMIKKGVQSNVFTYHIMIGSYCKEGNVDYALRLfeDISRNVHCYTNLIFGFYKKgdKAVDLLMRMLDNGIVPDHITYFVLLKMLPKChlKYAMVILQSILDNGCGINPPvlAVVTTALCSQRNYIAALSRIEKMVNLGCTPLPFSYNSVIKCLFQENIIEDLASLVNIIQELDFVPDVDTYLIVVNELCKKNDRDAAFAIIDAMEE------------LGLRPTVAIYSSIIGSLGKQGRVVEAEETFAKMLESG--IQPDEIAYMIMINTYARNGRIDEANELVEEVVKHFLRPSSFTYTVLISGFVKMGMMEKGCQYLDKMLEDGLSPNvyTALIGHFLKKGDFKFSFTLFGLMGENDIKH-DHIAYITLLSGlrKKKrvEPgeKLLQRLikKSIIPNLYLHNTIITGYCAAGRLDEAYNHLESMQKEGIVPNLVT-YTILMKSHIEAGDIESAIDLfySTLLKGLCDFKRPLDALALMLEMQKSGINPNK-DSYEKLLQCLC---------    
 976 UniRef50_C5Z8W5        Putative uncharacterized pr...   311     5e-83  1  10.9%  96:516    34:440     -----------------------------------------------------------------------------------------------LPSCLHLCSALLKCYAASGRIAAARQLFDEIprDVPLWNALVSAYARSGhlAAASAMARDVAGCRPNNVSVTSLLSVCTQLRSslGRELHGYAIRNVPVLDLPVLNALVNMYGKCGWLRDARMVFTGM---GvmRSAVSWTSMINACCQNGHPAEALEVFEEMRIARVKVDEVTLLAVISACTKL-----------DCTPGLGDWVEECAVENGFLQNTHVANALIHMHGKMGRVKRSCEIFDSM-----GAVKTVVSWTAMIQALAMNGHGVAALVRFTQMLREGFLPDEVIFLCVINACGHSRLVSEGHRLFKSmeEYHITpeHYGSMVDLLCRAGWLDEAFEFVLAMP----VTPDPVIWRVLAGACRDHGNASLARRVMDHVI--GMEPDhgNYVLASNLCAADEDWRRVVDVRLDM---------------------------------------------------------------------------------------    
 977 UniRef50_B9GYG8        Predicted protein n=5 Tax=r...   311     5e-83  1   9.1% 102:529    20:430     -----------------------------------------------------------------------------------------------------SYNTWVLAIKNASSPHKALQLYTHMHRQSIPFDTFscTHLKNLTIIHHLHSHIIKLGFNTHVYVSTCLLHAYAVMSFDQACDLFDEMPQR----NIVTWNTMITGYSRSGSINKARSLFEAMPVR----DAASWSAMITCYINNGFRDQGLSFFQEMMaeNPKPDQVTVGSVLSGCAHMGSLGL----------LAGKSVHGFVVKNGWELNVDIGTLLVDMYAKCGFFKNAVWVFVLMQER------NVSTWTALICGAAQHGFCQEVLSLFKMMQEAGVRPNEMTFTGILNACARKGLIEEGRKYFKMIKETGLEpqHYGCMVDMFGKAGLLEEAYEVIKEME----FEPNIVIWGSFLSACKMHKQFDIADRVIGQVL-RDIKpdGGIYSLVSDLYVLNKKWDDAERVRNLVLNQNVRKARGSSC--------------------------------------------------------------------------    
 978 UniRef50_Q0IQS7        Os12g0102600 protein (Fragm...   311     6e-83  1  15.8%  38:581    82:650     -------------------------------------LLSLLRDGHTDAAYHLFASNPSLP-LSPVSASRLLAQLssSFSRASALLHRLRARQALHLLDANSLSLASSASARSNNPHLAYSLLLSMLRRGLLPdaYTAALARLpsRALRLFDALlkTNSLPDTAAFNAALSACAngDCIRFRHLFDQMPAWNAPPDALTYNVLIKMCARAGRKDLVARVLHRILSSGLTPCATTFHSLVAAYVGFGDIPTAERIVQAMRERryPPNSRVYTTLMKGYMNAGRVEDVVAMLRAMRREGETSPASR-------PDHVTYTTVISTLVAAGDMERARAVLEEMGQAG--VAASRVTYNVLIKGYCQQLQAGKAKELLadMAEAGIQPDVVTYNTLIDGCVLTDDSAGAVALFNEMRERGIAPssYTTLMKAFAASGQPKLAHKVFDEMEKDPRVAVDRAAWNMLVEAYCRLGLLESAKKVVERMKARGVQPDVATYGSLAKGIAVARRPGEALLLWEEIKEK---------------------EVDGEVVEALADV---CVRAALFRKALEMVARMEEMGVEPN----------------------    
 979 UniRef50_Q6ZA49        Os07g0172600 protein n=3 Ta...   311     7e-83  1   9.3%   9:565    91:674     --------RAGALADARGVFERMPARDAASYSALISGHARLGSPAAavELLGRMRLAGM--APTEYTFVGLLTACARRGNPRLGSQVHALAVKGNSpgSLLVDNALLGMYVKGGRFDDALKVFDGMerDVSSWNTVLSGllGRYDEAFELFGDMRDSGVGADRFSLSALLAAAAEggLHEGAAVHALSLKSGLEMDLSVGNALVGFYAEHGhiEDVVDVFERMPAK----DVISWTGLLNGYMEFGLVDMAMDVFDRMPVRNF----VTYNAVLTGFNHNKEGvrGLGLFKQMLEDgsEQVQAFAIKCGCGSTPWIDAALIDMCIKCGRSGDAHLLFEKWRHEESFH----IAWNSLLAASFRDGEYEKALSTFLKmrSNDVqiDEFILTTVLGACGALGFAEFGKQMHCFAAKSGLLsgVGNAIISMYGKCGALETAVNVFKRMPCR-----DLVSWNALITSHLLHRQGDEILDLWSQMERLPIKPDSVTFLLVISSCSYTSSndKCRELFLSMSSIYGIEPAVEHYAAFVHVLGCWGHFEEAEQ----LIGKMPFKPSALVW--------------------------------------    
 980 UniRef50_Q0J0A3        Os09g0520300 protein n=6 Ta...   310     8e-83  1  11.3%  19:549    84:625     ------------------VFDAMPRRDAVAWNAMLTAYARAARPRAALALFARMRAPDAFSLTAALAanALVSMYAKCARAADAARAFREMPERNALSWC---SLLHAFVVSG-HMELAHELFDEMpkSNVAWNTLlhSRSGNAKQCLALFNQMWMSGLTCDDATLCILVDACAEldPSTGFAIHKVVVQSGwtRTTASWNSLIDAHARFGYIEQAALLFESAPE----TNIISWTAMIGGFARNSLTSEALAHFVKMlqEYIQPDDFTFGAVLHACASAPCLASGRMV------------HSCAFQGGFASYLYVANNLVDMYAKCGDVEGANNVFDAIHQK------DLVSWNTMLFGFAINGLPKEALEVYEIMTYHNVSPDEVTFTGLLTACSHSGLLEQGRAFFESMMshGvkPEHLSCVLDMYARSGNIAKAIEMMEQYPEI-VKSPGSGLSEALLSFCSSEndFWVGRKVGDDVVARAPARDTG-YVMLSNLLCASGRWDEAERVRRAMAEQGIKKSP----------GCSWIEVKGKVKMG------------------------------------------------------    
 981 UniRef50_Q10GC5        Pentatricopeptide, putative...   310     1e-82  1  10.1%  67:551    34:523     ------------------------------------------------------------------SHPALTLLHTQCASMAHLRQLHAALVKSGLARDPIAASRAVAFCAGDGRDAaaRLVRHHPRPNAFMWNTAIRALADGPgaAVALFVDMLGSPTPPERRTFPSLFAAYArgRAGDGAGLHGMVVKLGLGGDAYVRNSVIAMYarAGRVDEARAVFDGMPAR----TVATWSAMVSAYSRDSRCHDAVELFSAMQAEGVEPNANVLVSVLGCCASLGALEQGAWV------------HAYIDKHDVAMNALVVTALVDMYCKCGDIRKAREVFDASRSRGQAK---LSSWNSMMLGHAVHGQWREAAALFSELRPHGLRPDNVTFIAILMAYGHSGMADEAKAVLASmsEHGVVpeHYGCLVDALARAGRLREAEGAIAAMP----VAPDAAVWGALLSGCRLHGDAEAAARAAREAVRCDPR-DSGAYVLAASALARGGEARRGAAVRGRMREEGVGKVP----------GCSMIEVDGVVHEFVS----------------------------------------------------    
 982 UniRef50_A9T7E2        Predicted protein (Fragment...   310     1e-82  1  12.1% 129:500     1:350     --------------------------------------------------------------------------------------------------------------------------------NLISCNAAISRLAQQGreALAFFFRMQDEHVRPNRVTFLSLLKAfgPGDLESGRRVHNFILERQLESDVRVGTALTCMYFRCGSALDAHRVFKRVQNR----DIIVWNAMIEGYCQAGLRDKALEMFKDMKIAGLAPDVASFLSILKSCNGSVHLDDGKQL------------HKAIIESGFELSVRVQTALVNMYCRCESLLDARKMFNKLPRK------DIGLWTTMIAGYAQTDHGEEAFALFQAMRDTGLQPNKMTFLALLKACTRPEFLQKGKWLHNHIRCAGLDSdvGTALINAYSKCRSLVDARKVFD-----NLCTRNVISWTALIAAYSRNDRAEEALDLFEQMKREGVQPNEVTRNTVVKAC-------------------------------------------------------------------------------------------------------    
 983 UniRef50_A9S0K8        Predicted protein n=1 Tax=P...   309     2e-82  1  12.0% 132:524   270:672     -----------------------------------------------------------------------------------------------------------------------------------AFNALINAaaGQHKLAESLFSEMRNLGLRLSIYTFNALLRAVVEGrvSHALRVVRSMEAVGVRPDTHTFTTLLDGYSRSGKLDKAEALLAKMEDsnQRPSIYTYNILIKACASKNDPERALRVFAKMKESGTAPDSHTYCALLSAcaVSGRPDLSDAWSRKEVAQRVIAIEKDMAQSGVHHTAASVSALMSVLGSEGIVDAMLQNLRACQETDLTTrlDTTAYNTAMNAFVQSKKFDEAWDIFTEMRSLDVRPDIYTYNILINACSHKQSVSTALELMDIMRKAKIAPdtYNSLIKVMCHCGELDSGLRVLGEMQDAG-VQPDVTTFNTLLASASYHKRKGLAEYLVEEMRLCNVAPDTHTCSQVISLYMRSGNLAEASEALSVLSSRMRKPM-------------------------------------------------------------------------------    
 984 UniRef50_Q9LP03        Pentatricopeptide repeat-co...   309     3e-82  1  12.0%  13:522    98:583     ------------LNNALDLFDEMPERDVVSWNTMISGLVSCGFHEYGIRVF--FDMQRWEIRPTEFTFSILASL---VTCVRHGEQIHGNAICSGVSrnLVVWNSVMDMYRRLGVFDYALSVFLTMerDVVSWNCLILScsGNKEVALDQFWLMREMEIQPDEYTVSMVVSICSDLrlSKGKQALALCIKMGFLSNSIVLGAGIDMFSKCNRLDDSVKLFRELEK----WDSVLCNSMIGSYSWHCCGEDALRLFILAMTQSVRPDKFTFSSVLSS-------------MNAVMLDHGADVHSLVIKLGFDLDTAVATSLMEMYFKTGSVDLAMGVFAKTDGK------DLIFWNTVIMGLARNSRAVESLAIFNQLlnQSLKPDRVTLMGILVACCYAGFVNEGIQIFSSMEkhGVNpeHYACIIELLCRVGMINEAKDIADKIP----FEPSSHIWEPILCASLDLGDTRLAETVAKTMLESEPK-SSFPYLVLIKIYEMTWRWENSVKLRYAMNEHKLK---------------------------------------------------------------------------------    
 985 UniRef50_D1H9A2        Whole genome shotgun sequen...   309     3e-82  1  11.0%  20:521   144:673     -------------------KQKGYKHNFASYNAFAYCLNRSNQFRAADQVPELMNMQGK--PPSEKQFEILIRMHIDANRGLRVYYVYEKMKKFGIKPRVFLYNRIMDGLVKTGHLDLAMSVYEDFKEDGLVEESVTycKAGRIDEVLELLDRMRGNLCKPDVFAYTAMVKVlaEGNLDGCLRVWEEMRKDKVEPDVMAYTTLVAALCNGNRVGEGFELFKEMKQKKYLIDRAIYGSLIEGFVVNERVGSACDLLKDLMDSGYRADLAIYNSLIEGMCNVKQVDKAYKLFqeMKrdDFCSLLGQMQKLGFPvdDLsfSVMIE---KGERLKLALEVFEHLKAKGYC---SISIYNILMEAIHRTGEVKKALSLFDDIKDSNFKPDSSTYSNAIICFVEVGDVQEACACYNKIIEMCQLPSvyRSLVKGLCKSEEIDAAIMLVRDCLANVTSGPMEFKYttILHAC-KSGNAEKVIDVLNEMMQEGCTPDEVTYSALISGMCKHGTLEEARKVFSNMRERKL----------------------------------------------------------------------------------    
 986 UniRef50_D2W5L8        Predicted protein n=1 Tax=N...   308     3e-82  1  10.5%  18:579   314:902     -----------------EMQDYGVEPNSTTFNILLTACADSKLFEEGKAIHKL--ATGTLKDYSSELLASIVNFYTKCGHPETAIKECEQLKSKGLMNNIVWNCLIQAYIQN-KEVRQAIEVITEMQDYGIEPNSTTFNILLKACADSKLFEemQQFGLNPDSTTYTIILKACTDFNleEGKLIHEQAIENVKDCSTELLTSIISFYTKCGHPEVSIGECEKLQSKGLMNNVAVWNCLIQAHIQTKEVPKSIKIITEMQDCGVEPNSTTFNILLTACADSKLFEEGKAIHKLATGTLKDHSSELLA------------SIVNFYTKCGHPEVSIEECEQLKSKG--LMNDVAIWNCLIQAHIQsg-LNKAMVVFNEMKEH-IEPDKITYLILLAGCGEYFARKMGLDIIDELKLKGMDveIVSVMIYFYSRIAYQSTAQQLFEKLVEEsqKRLSIITYNNMLSCYAQHGKGTQAMNLFNEMVAKNVEMNENTFTIMLNTFSHCGMIDEANQLYGEMKTKYNHNPSVIVDCVMVDVFARSGRFHEAEKilWSALMAGCRKFNNVEKADNIRQLMIENGIK------------------------    
 987 UniRef50_Q8S7V8        Os03g0169300 protein n=3 Ta...   308     3e-82  1  11.0%  53:524    33:496     ----------------------------------------------------LLRRRHLLPTADAVAALAKLLRFadLRHAAALLSLHLPFLSSASSHLAFFYNTLMRGLAASSSPAAAIELFSAMRRAGAAPDAFTFTFALKSCSRcdLHAQAIKHGcsSHVHVHNALLHGYsrAAVGDARRVFDEMPAR----DVISFSGLLTLHLKANDLDAARVVFDQMPHR----DVVSWTAMISAYARARRPQEALALFDAMP---VQPDEVTMLSVVSACTALGDLATGERVCQYVDSNGFGWM------------VSLRNALMDMYAKCGSLTEARNLFDGMTVR------SLASWNTFISAYASHGDVESTVALFHRMLADgvKPDGTTLLAVLTAYAHKGFVEEGRAMFNAILSGnlTIEHYGCMVDLLGRAGQLEEAYKMIEQMP----IPSNSVVWGALLGACRTHGDINMAERAVQKLRSLNPE-EGGYYILLSDMYAASGRTTEAMEIRRIMNQAGARKT-------------------------------------------------------------------------------    
 988 UniRef50_Q0DT34        Os03g0271300 protein (Fragm...   308     3e-82  1  13.7% 162:526    97:476     -----------------------------------------------------------------------------------------------------------------------------------------------------------------DAVAATALLDMYSKCGlaSARKVFDEMASRG---DPVVWNALLACYARHGLPEHALALAVKMRGIGLCPDLVTWNIVVSGFALAGDDLMAGDLVGAMKDDGFQPDVVTWTSRVSGSVLNFQYGRARALFRAMmvKHGKEVHGYSVVAGVEQELTVSSALVDMYAKCGLVLEARRLFDKMPQR------STVTWNSMIFGLANSGHCQEAVGLFDRMLRDGAKPDHLTFTAVLTACSYGGMVEVGKILYRAmvEYGVEpeHYACMVHLLGRAGRLDEAYGFIRAMP----LEPDRFVWGALLGACRSHGNIELAELAASRLLA--VEPdaASCLLLSSALANAGKQDDVFKIKRLVKRRRLKKLDS-----------------------------------------------------------------------------    
 989 UniRef50_Q9FRS4        Pentatricopeptide repeat-co...   308     4e-82  1  10.0%  27:475   103:544     --------------------------DEETNNEILHNLCSNGKLTDACKLVEVMARHNQVPHFP--SCSNLVRGLARIDQLDKAMCILRVMVMSGGVPDTITYNMIIGNLCKKGHIRTALVLLEDmpPDVITYNTVIRCMfgNAEQAIRFWKDQLQNGCPPFMITYTVLVELVCrcGSARAIEVLEDMAVEGCYPDIVTYNSLVNYNCRRGNLEEVASVIQHILSHGLELNTVTYNTLLHSLCSHEYWDEVEEILNIMYQTSYCPTVITYNILINGLCKARLLSRAIDFFYQMLEQK------------CLPDIVTYNTVLGAMSKEGMVDDAIELLGLL--KNTCCPPGLITYNSVIDGLAKKGLMKKALELYHQMLDAGIFPDDITRRSLIYGFCRANLVEEAGQVLKEtrGNGIRGSTYRLVIQGLCKKKEIEMAIEVVEIMLT-GGCKPDETIYTAIVKGVEEMGMGSEAV--------------------------------------------------------------------------------------------------------------------------------    
 990 UniRef50_C5WS24        Putative uncharacterized pr...   308     4e-82  1  10.2%  77:574   129:617     ----------------------------------------------------------------------------RERDLPQAEALFRAAPAAARG--LYLDTVMLGGYAKAGRVDRARELFDGMpkNVVTWTAMVSGyaGRVREARELFDVMPDR----NDYSWTTVVQGYTSNglMEARETFDRMPQR----NVVAWTAMVKAYVDSGQIQDAWELFDMMPER----NSYSWNAMISGFLSIGEVAEAVQLFERMPHR--LRNVVSWTTMVTGLARNGLAYRAREFFDRMPVK----------------DAAAWNAMITAYANNGQLNEAQRLFDSMLAK------DLVSWSTIIEAYSKNERRHEALNMFLLMRRSAVSPnvITfnALLTMYSRSGDLMSAWSAFEKLEEK--DAITWTSIvdLLGRSGYVREAKELVNGMQQ---GMRDEAILATLLGACVMHNEVEVAREVGEDLIR--FDPssGGYRLLANVFASHGMRDETANVWKVMRGSKAKRTPGF----------SQIEVNMRNHVFYSGDQEHPQCPEIYEMLNdiVPQMK-----------------------------    
 991 UniRef50_Q0WLC6        Pentatricopeptide repeat-co...   308     4e-82  1  12.3% 142:580   450:875     ---------------------------------------------------------------------------------------------------------------------------------------------RAVKEAFRF----TKLILNPTMSTFNMLMSVCASsdIEGARGVLRLVQESGMTADCKLYTTLISSCAKSGKVDAMFEVFHQMSNSGVEANLHTFGALIDGCARAGQVAKAFGAYGILRSKNVKPDRVVFNALISACGQSGAVDRAFDVLAEMKAE----------THPIDPDHISIGALMKACCNAGQVERAKEVYQMIHKYGIRGTPEV--YTIAVNSCSKSGDWDFACSIYKDMKEKDVTPDEVFFSALIDVAGHAKMLDEAFGILQDAKSQGIRLGtySSLMGACCNAKDWKKALELYEKIKSIKL-RPTISTMNALITALCEGNQLPKAMEYLDEIKTLGLKPNTITYSMLMLASERKDDFEVSFKLLSQAKGDGVSPNLI--MCRCITSLCKR-RFEKAcePVVSFKSGRPQIENKwsMALMVYRETISGGTVP-----------------------    
 992 UniRef50_B9N1T3        Predicted protein n=2 Tax=P...   307     6e-82  1  11.3% 142:521    63:441     ---------------------------------------------------------------------------------------------------------------------------------------------RNIDDALASFNHMLHRKPLPCIIQFTKLLSAIVKMryDAVISLSRPMELAGLSPNTYTLNILINCFCKLQHVDLGFSVLAKVIKLGLQPDVVPFTTLINGLCKVGKFSQAVELFDETVERG-------YNSLIHGLCNCSQWKEASAILNEMTSLNIM------------PDIVTFNVLVDTFCKEGKVSEAQGVLKTMTE--MSVELDVVTYNSLMYGHSLWMEVVEARKLFGFMITKGCKPNVFSYNILINGYCKAEMIDEAKQLFNEMIHQGLTPnsYNT--HGFCQLGKLREAQELNENMHTNGNL-PDLCTYSILLDGFCKQGnlDEALETFRKMEEDGCPPNEWSYNVIIRGFLRHRDESRAVQLIGEMRDRGR----------------------------------------------------------------------------------    
 993 UniRef50_A2Q1P5        Tetratricopeptide-like heli...   307     7e-82  1  10.2%  35:474    36:460     ----------------------------------IKTLVSMGLYHQTLQFFTQLHFSAHHFNSIPFVLPSVIKACSFTHFHAFGTQLHCLAFITGSYTDPIVSNSIISMYAKFFDIESARQVFDTmhRDTITWNSMINAYLQNGleALQMLKDFYFLGFLPKPELLASMVSMCGrlGWRIGRQIHGLVVVDgiQHSVFLSTAFVDFYFRCGDSLMARSVFDEMEVK----NEVSWTAVISGCANNQDYDVALACYREMQVEGVSPNRVTLIALLAACARPGFVKYGKEIHGYAFRRGFDSCHSFSP------------ALIYLYCECGqlHLAERIFE------GSSLRDVVLWSSIIGSYARRGESDKALKLFNKMRTEETEPNYVTLLAVISACTNLSSFKHGGVIHGYILKFGisIFVCNALINMYAKCGSLDDSRKIFLEMPSR-----DSVTWNSMISAYGLHGYGEQA---------------------------------------------------------------------------------------------------------------------------------    
 994 UniRef50_Q0E1N8        Os02g0321000 protein n=3 Ta...   307     9e-82  1  11.5%  20:504    51:521     -------------------KRRRLERD---YNVAMKALALAGDLDEVVAVFAELKRTAADgpPNVLCYNTLVNALAEAGREGEALKAFDEMLASGVAPNASSQNILIKMHARRSEFDLAWELIHkvEPDVGTYSTLIAGLCRAGkeAWGVLDWMLEKNCRPMVQTYTPIVQAYCrgRIVEAKLLMAEMERLGCLPNVVTYNVLIRALCDDDKFDEVEQVLMESSTKDWKPSTVTYNIYMNGLCKKGKAKEALELLDVMLGEGLEPTAYTWSILLNCLCHSSRLLDAIYLLERSTELKWYAG------------VVAYNTVMSSLCEMGKWRGIMKLLTDMIKKG--IEPNTRTFNILIRSLCVGGKSSLAKSLIH---SLGFAANVVTYNILLHWFYYHGKLTEANRLISVMEEKNIAPdtYTIIIDGLCRERKFDAATACFLKSLTSGL---SMDVLTVLLNRLVYADKIWEINRIFD---GKDFVPDHHVFDLTIRTFCRVG---------------------------------------------------------------------------------------------------    
 995 UniRef50_C5YAH4        Putative uncharacterized pr...   306     1e-81  1  13.3%  25:520    55:545     ------------------------SHTLHDYNRLLAALARDGDGDAALRVLRRMRLSSPPCAPTAASYTSAMSALAKAGRPADAASLFDDMLANGVAPDRAAFSLLLHVYSShhLPAAAHSVLLWmpPTPIDYTDLIFSFCRAGClaLQLLDEMRALNYPLTPHTFAPILKAICDndIQAADALISSMRSSGCLPDVVIYNIYIQGLCKIGDFDAVERVIYDSARNGWEPDAVTYSTYIVGLCRSGYIEEAFQQLETMVAKGLQLTVVGLNILLDHVAQNLDMWVGKEVLERCQE------------LGFVVDVVTYNTVMDHLCKKGKWLRVLKLFTDLLKK--PITPNVQTYNILISCLCRAGKFEFAKYVFS---SKGFVADTVTCNILIHEFYEAGKEGELGFLFADVNAGKIAPdtYNTLVDCLFRSGRRSQAVNFVRHIDGGYPTEPVAhtY-----WLVRSGYIREALRLFDDIQIKGLVLDCRIFANVIKAFCRKgdMSQLCFVLDRMLGSG-----------------------------------------------------------------------------------    
 996 UniRef50_B9RTQ7        Pentatricopeptide repeat-co...   306     2e-81  1  10.4%   3:415    45:439     --SNHLISMYSKLDlaQLVLHLTPTRSVVTWTSLISGSVQNGHFSFA--LYHFFNMRRDNIQPNDFTFPCAFKASASLLLPFVGKQIHAIAVKFGQINDVFVGCSAFDMYSKTGLKQDAQKLFDEleRNVVTWNAYISNAvgRYQNAAVAFVELRRAGCEPDSTTFCVFFNACaqLYVDLGRQLHGFVIRSGFEKSVSVLNGLIDFYGKCKEVRLAEMVFGKMENR----NAVSWCSMVAACEQNGEEEKACLFFVEGRKEGIEPTDYMVSSVISACAGLAGLELGRSF------------HALAVKACLEGDIFVGSALVDMYGKCGGIEDSEQAFHEMSER------NLVTWNALIGGYAHQGHAEMAVRLFKEMTTE-VVPNYVTLVCVLSACGRGGAVELGMEIFGSMRDR--------------------------------------------------------------------------------------------------------------------------------------------------------------------------------------------    
 997 UniRef50_Q9LN22        Pentatricopeptide repeat-co...   306     2e-81  1  10.3% 122:579    72:533     -------------------------------------------------------------------------------------------------------------------------IKDHYRKNPTSPNDAILNPSLTLH-ALSlfSQIETSQVSPSVV--RCVIEKCGSVrlHQSLAFFNWATSRDdhKSPHPYNEMIDLSGKVRQFDLAWHLIDLMKSRNVEISIETFTILIRRYVRAGLASEAVHCFNRMEDYGCVPDKIAFSIVISNLSRKRRASEAQSFFDSLKDR-------------FEPDVIVYTNLVRGWCRAGEISEAEKVFKEMKLAG--IEPNVYTYSIVIDALCRCGQISRAHDVFADMLDSGCAPNAITFNNLMRVHVKAGRTEKVLQVYNQMKKLGCEPDtyNFLIEAHCRDENLENAVKVLNTMIKKKC-EVNASTFNTIFRYIEKKRDVNGAHRMYSKMMEAKCEPNTVTYNILMRMFVGSKSTDMVLKMKKEMDDKEVEPNVNT-YRLLVTMFCGMGHWNNAYKLFKEMVEEKcrRAGQLKKHEELVEKMIQKGLV------------------------    
 998 UniRef50_A9TMJ9        Predicted protein n=2 Tax=P...   306     2e-81  1  12.3%  20:450   239:704     -------------------ESGFFVRNLIDPVSVVSSCAKLRNIDLARRFVELLSR-------NPWHYNLIIREFSRHGNLRSCFKVLDFMKELGVMPDVFTYRVLIDSCGRHKEPSKATIVFEKMLNDGFEPNRFVYNsvGNLSEVRRYFQHMQGAGIAADVTSYNILLKAYasGRADLAGDLYAQIKKAgvTLDVVFYSTLINVFGKAKMWEEALRVKADMLEAGVTPNVVTWTSIIGACATAGLVDQSFREFDEMLNAGCRPNVNCYNMLLQACIMASQFDRAFLLFEelQSNLGEKIDEDVRLACCRPNIITYNCMMKACGSAS--EKVRSLMREMRALG--LVPDVKSWSILLDSYGSKGDVEGASEALEEMKAAGFKPDVVAYTSLIQACVQAGHLEKAFDYFSKMKAAGVRPntYNSLLRRHRNYgeVQRALALYEEMREAGH---------------------------------------------------------------------------------------------------------------------------------------------------------    
 999 UniRef50_Q75IT2        Os05g0365500 protein n=10 T...   305     3e-81  1  10.5%  96:551   114:568     -----------------------------------------------------------------------------------------------LSLHNRVQSLIRSGDLDGASVAARAAVTSRVRPTVFTCNAVAAsrAGRHDDAVSLFDFfrRSNIVPNVVSYNTLILAHCEarVDSALQVYREMlkAPFSPSAVTYRHLTKGLVAAGRIQDALDLLREMLNRGQGADSLVYNNLIAGYIEEDNWDKAFELFEELQGRCLVYDGVVHTSFMEGYWKKGMDKEAMENYQSLLDKK------------FRMTPATCNVLLETLFKHDKDKEANDLWETMIDNHTpiGINSESYNVMVNQCFKKCKFHEAIEVFHRQPRKNVQMDVGCFNNIIGKLCENGMLEEAEKLFEEMETKSVLPdtYTYLVDSCFKVGRVDDTIQYFYKMADGREHGpnIGFFNRMFEGLTEAGRMDDALKVYGRMPDKEIKPNTTTFETLVNALCKEGDLDKARELVSDM-AKGGVTPPPEFRDSVIDIFKNAGRQEEIEKAFEE----------------------------------------------------    
1000 UniRef50_B9R967        Pentatricopeptide repeat-co...   305     3e-81  1  11.0%  69:485    34:425     --------------------------------------------------------------------TVLVRKLLNLSDIDYARQLFDQVPQPGQ----ILYNSLISTYSKLSLHKDALKTFFSMHHSDTRLSCFTGPPVIKACSSlqVHSLIVICGIDCNVYVQTSLMDFYAkgELGSARKIFDGILVK----DPISYNCLITGYSKAGDVIAARRLFDSMTER----TVVSWNAMISCYAHNGDLNEGLKTFERMQAEDISPNEITLVTLLSICAKLGDLEMGLRI------------KKYIEDNNLCVNMIVSTAILEMYVKCGAVDDARKEFDRMGQR------DIVAWSAMIAGYAQNGRSNEALELFECMRREKVKPNDVTLVSVLSACVQLGSVEMGNYIGSYVESQDLASNVysALVGMYSKCGNISKAREVFGKTPQK-----DIVTWNSMIVGLAVNGFAKDAIALYRNMKEAD----------------------------------------------------------------------------------------------------------------------    
     consensus/100%                                                                                        ...........................................................................................................................................................................................................................................................................................................................................................................................................................................................................................................................................................................................................................    
     consensus/90%                                                                                           .......................................................................................................h...............h...........s.hh.s.......s..hh..h.......s.h.hs.hh..h...th..s..hht.h.......p....s.hlthh.p.t.h..u..hht.h.......s.hsas.hltsh.t.t...ts..hh..h.......s.hsh..hl.sh.t.t....s..........................t....ssllphhspts.ht.u..hh..h.........s.hsasshl.shs..s...tuh.hhtth...th.ss.hsh..ll.uhsp.s.ht.u..hht.h.t.th..phhs.hlphhsc.s.httu.thh.th........s...ht.hltuh...t.ht.u..hht.h.......p...h..h.thh...t.ht.h..hh..h.....................................................................................      
     consensus/80%                                                                                             ..................................................................hh..h........h..hht.h...th.....h.s.lhthh.t.t..t.t..hhp.h..p.h..s.hh.shhtp.ptuhthh.th.ttth..s.hshs.hlpshsp.tht.uhtlht.h.p.th..shhhhssllphhsptsthttAhtlhpth.tt....shhsasshlsuhspts..tpAhthhtpM.ttth..s.hshsshltuhsp.s.ht.u..h....................h..s.hhhssllshas+sGphptA.tlapth..t......shhsasshltuhs.pG.hppAhthappM.ttsh.Ps.hoa.sllpuhsp.uhhppuhphht.h.p.th.sptasshlshhs+sG.lppA.phhpph........s.hhassllsuhthttphp.u..hhpphht...t.s.hsa.hl.phhst.t.hpps.phht.h.t..h............................................................................          
     consensus/70%                                                                                              .............................................h............p...hs.hltsh.t.ttht.uhtlhs.hhp.sh..s.hh.shlhthhtptt.hptuh.hhpth.tshhshsshltuhhtphppAhtlatpMhppsh.ss.hohssllpuhsptphp.utplas.hhctsht.shhshssllshYs+sGphppAhplFpph.pp....shloasshlsuasptuphpcAhplappM.ppsh.ss.hohssllsuhsphuthp.uhtl...................th..shhhhsuLlshYsKsGplppAhplFpph.pp.  ...shhoasshIsuhshpGpspcAlplFppMhppuhpPstlTahsllsuhs+sGhlpcuhphappMhchslpPppYsshlchhs+sGplpcAhphhppM.......sshhhassllsuhphpsphchu..hhpphhp.thpsstssashl.shhsptsphpcshplhp.Mtppth.............................................................
```

MView 1.51.1, Copyright © 1997-2009 Nigel P. Brown
