## Supplementary material for "The endosomal TbTpr86/TbUsp7/SkpZ (TUS) complex controls surface protein abundance in trypanosomes": DA1: query.dom.html

### Undefined

|  |  |  |  |  |  |  |  |  |  |  |  |  |  |  |  |  |  |  |  |  |  |  |  |  |  |  |  |  |  |  |  |  |  |  |  |  |  |  |  |  |  |  |  |  |  |  |  |  |  |  |  |  |  |  |  |  |  |  |  |  |  |  |  |  |  |  |  |  |  |  |  |  |  |  |  |  |  |  |  |  |  |  |  |  |  |  |  |  |  |  |  |  |  |  |  |  |  |  |  |  |  |  |  |  |  |  |  |  |  |  |  |  |  |  |  |  |  |  |  |  |  |  |  |  |  |  |  |  |  |  |  |  |  |  |  |  |  |  |  |  |  |  |  |  |  |  |  |  |  |  |  |  |  |  |  |  |  |  |  |  |  |  |  |  |  |  |  |  |  |  |  |  |  |  |  |  |  |  |  |  |  |  |  |  |  |  |  |  |  |  |  |  |  |  |  |  |  |  |  |  |  |  |  |  |  |  |  |  |  |  |  |  |  |  |  |  |  |  |  |  |  |  |  |  |  |  |  |  |  |  |  |  |  |  |  |  |  |  |  |  |  |  |  |  |  |  |  |  |  |  |  |  |  |  |  |  |  |  |  |  |  |  |  |  |  |  |  |  |  |  |  |  |  |  |  |  |  |  |  |  |  |  |  |  |  |  |  |  |  |  |  |  |  |  |  |  |  |  |  |  |  |  |  |  |  |  |  |  |  |  |  |  |  |  |  |  |  |  |  |  |  |  |  |  |  |  |  |  |  |  |  |  |  |  |  |  |  |  |  |  |  |  |  |  |  |  |  |  |  |  |  |  |  |  |  |  |  |  |  |  |  |  |  |  |  |  |  |  |  |  |  |  |  |  |  |  |  |  |  |  |  |  |  |  |  |  |  |  |  |  |  |  |  |  |  |  |  |  |  |  |  |  |  |  |  |  |  |  |  |  |  |  |  |  |  |  |  |  |  |  |  |  |  |  |  |  |  |  |  |  |  |  |  |  |  |  |  |  |  |  |  |  |  |  |  |  |  |  |  |  |  |  |  |  |  |  |  |  |  |  |  |  |  |  |  |  |  |  |  |  |  |  |  |  |  |  |  |  |  |  |  |  |  |  |  |  |  |  |  |  |  |  |  |  |  |  |  |  |  |  |  |  |  |  |  |  |  |  |  |  |  |  |  |  |  |  |  |  |  |  |  |  |  |  |  |  |  |  |  |  |  |  |  |  |  |  |  |  |  |  |  |  |  |  |  |  |  |  |  |  |  |  |  |  |  |  |  |  |  |  |  |  |  |  |  |  |  |  |  |  |  |  |  |  |  |  |  |  |  |  |  |  |  |  |  |  |  |  |  |  |  |  |  |  |  |  |  |  |  |  |  |  |  |  |  |  |  |
| --- | --- | --- | --- | --- | --- | --- | --- | --- | --- | --- | --- | --- | --- | --- | --- | --- | --- | --- | --- | --- | --- | --- | --- | --- | --- | --- | --- | --- | --- | --- | --- | --- | --- | --- | --- | --- | --- | --- | --- | --- | --- | --- | --- | --- | --- | --- | --- | --- | --- | --- | --- | --- | --- | --- | --- | --- | --- | --- | --- | --- | --- | --- | --- | --- | --- | --- | --- | --- | --- | --- | --- | --- | --- | --- | --- | --- | --- | --- | --- | --- | --- | --- | --- | --- | --- | --- | --- | --- | --- | --- | --- | --- | --- | --- | --- | --- | --- | --- | --- | --- | --- | --- | --- | --- | --- | --- | --- | --- | --- | --- | --- | --- | --- | --- | --- | --- | --- | --- | --- | --- | --- | --- | --- | --- | --- | --- | --- | --- | --- | --- | --- | --- | --- | --- | --- | --- | --- | --- | --- | --- | --- | --- | --- | --- | --- | --- | --- | --- | --- | --- | --- | --- | --- | --- | --- | --- | --- | --- | --- | --- | --- | --- | --- | --- | --- | --- | --- | --- | --- | --- | --- | --- | --- | --- | --- | --- | --- | --- | --- | --- | --- | --- | --- | --- | --- | --- | --- | --- | --- | --- | --- | --- | --- | --- | --- | --- | --- | --- | --- | --- | --- | --- | --- | --- | --- | --- | --- | --- | --- | --- | --- | --- | --- | --- | --- | --- | --- | --- | --- | --- | --- | --- | --- | --- | --- | --- | --- | --- | --- | --- | --- | --- | --- | --- | --- | --- | --- | --- | --- | --- | --- | --- | --- | --- | --- | --- | --- | --- | --- | --- | --- | --- | --- | --- | --- | --- | --- | --- | --- | --- | --- | --- | --- | --- | --- | --- | --- | --- | --- | --- | --- | --- | --- | --- | --- | --- | --- | --- | --- | --- | --- | --- | --- | --- | --- | --- | --- | --- | --- | --- | --- | --- | --- | --- | --- | --- | --- | --- | --- | --- | --- | --- | --- | --- | --- | --- | --- | --- | --- | --- | --- | --- | --- | --- | --- | --- | --- | --- | --- | --- | --- | --- | --- | --- | --- | --- | --- | --- | --- | --- | --- | --- | --- | --- | --- | --- | --- | --- | --- | --- | --- | --- | --- | --- | --- | --- | --- | --- | --- | --- | --- | --- | --- | --- | --- | --- | --- | --- | --- | --- | --- | --- | --- | --- | --- | --- | --- | --- | --- | --- | --- | --- | --- | --- | --- | --- | --- | --- | --- | --- | --- | --- | --- | --- | --- | --- | --- | --- | --- | --- | --- | --- | --- | --- | --- | --- | --- | --- | --- | --- | --- | --- | --- | --- | --- | --- | --- | --- | --- | --- | --- | --- | --- | --- | --- | --- | --- | --- | --- | --- | --- | --- | --- | --- | --- | --- | --- | --- | --- | --- | --- | --- | --- | --- | --- | --- | --- | --- | --- | --- | --- | --- | --- | --- | --- | --- | --- | --- | --- | --- | --- | --- | --- | --- | --- | --- | --- | --- | --- | --- | --- | --- | --- | --- | --- | --- | --- | --- | --- | --- | --- | --- | --- | --- | --- | --- | --- | --- | --- | --- | --- | --- | --- | --- | --- | --- | --- | --- | --- | --- | --- | --- | --- | --- | --- | --- | --- | --- | --- | --- | --- | --- | --- | --- | --- | --- | --- | --- | --- | --- | --- | --- | --- | --- | --- | --- | --- | --- | --- | --- | --- | --- | --- | --- | --- | --- | --- | --- | --- | --- | --- | --- | --- | --- | --- | --- | --- | --- | --- | --- | --- | --- | --- | --- | --- | --- | --- | --- | --- | --- | --- | --- | --- | --- | --- | --- | --- | --- | --- | --- | --- | --- | --- | --- | --- | --- | --- | --- | --- | --- | --- | --- | --- | --- | --- | --- | --- | --- | --- | --- | --- | --- | --- | --- | --- | --- | --- | --- | --- | --- | --- | --- | --- | --- | --- | --- | --- | --- | --- | --- | --- | --- | --- | --- | --- | --- | --- |
| Secondary Structure |  |  |  |  || Index |  |  |  |  |
| Disorder |  |  |  |  | D | D | D | D | D | D | D | D | D | D | D | D | D | O | O | O | O | O | D | D | D | D | D | D | O | O | O | D | D | D | O | O | O | O | O | O | O | O | O | O | O | D | D | D | D | D | D | D | O | O | O | O | O | O | O | O | D | D | D | D | D | D | D | O | O | O | O | O | D | D | D | D | D | D | O | O | O | O | O | O | D | O | O | D | D | D | D | D | D | D | D | D | O | O | O | O | O | O | O | O | O | O | O | O | O | O | O | O | O | D | D | O | O | D | D | O | O | O | O | O | O | O | O | O | O | O | O | O | O | O | O | O | O | O | O | O | O | O | O | O | O | O | O | O | O | O | O | O | O | O | O | O | O | O | O | O | D | D | O | D | O | O | O | O | O | O | O | O | O | O | O | O | O | O | O | O | O | O | O | O | O | O | O | O | O | O | O | O | O | O | O | O | O | O | O | O | O | O | O | O | O | O | O | O | O | O | O | O | O | O | O | O | O | O | O | O | O | O | O | O | O | O | O | O | O | O | O | O | O | O | O | O | O | O | O | O | O | O | O | O | O | O | O | O | O | O | O | O | O | O | O | O | O | O | O | O | O | O | O | O | O | O | O | O | O | O | O | O | O | O | O | O | O | O | O | O | O | O | O | O | O | O | O | O | O | O | O | O | O | O | O | O | O | O | O | O | D | O | O | O | O | O | O | O | O | O | O | O | O | O | O | O | O | O | O | O | O | O | O | O | O | O | O | O | O | O | O | O | O | O | O | O | O | O | O | O | O | O | O | O | O | O | O | O | O | O | O | O | O | O | O | O | O | O | O | O | O | O | O | O | O | O | O | O | O | O | O | O | O | O | O | O | O | O | O | O | O | O | O | O | O | O | O | O | O | O | O | O | O | O | O | O | O | O | O | O | O | O | O | D | D | O | O | O | O | O | O | O | O | O | O | O | O | O | O | O | O | O | O | O | O | O | O | O | O | O | O | O | O | O | O | O | O | O | O | O | O | O | O | O | O | O | O | O | O | O | O | O | O | O | O | O | O | O | O | O | O | O | O | O | O | O | O | O | O | O | O | O | O | O | O | O | O | O | O | O | O | O | O | O | O | O | O | O | O | O | O | O | O | O | O | O | O | O | O | O | O | O | O | O | O | O | O | O | O | O | O | O | O | O | O | O | O | O | O | O | O | O | O | D | O | D | O | O | D | O | O | D | D | D | D | O | O | D | D | D | D | D | D | O | O | O | D | O | D | D | D | D | D | D | D | D | D | D | D | D | D | D | O | O | O | D | O | O | O | O | O | O | O | O | O | O | O | O | O | D | O | O | O | O | O | O | O | O | O | O | O | O | O | O | O | O | O | O | O | D | D | D | D | D | D | D | D |
| Template | QueryhitRange | Confidence | Percent\_sequence i.d. | Rawscore |
 **c4m57A\_** | 17-516 | 100.0 | 17% | 484.1 |
 c3jd5e\_ | 178-519 | 100.0 | 12% | 376.9 | c5i9fA\_ | 183-513 | 100.0 | 18% | 285.5 | c5iwwD\_ | 195-515 | 100.0 | 13% | 250.6 | **c4g25A\_** | 313-593 | 100.0 | 14% | 265.6 | c5dizB\_ | 313-506 | 100.0 | 13% | 250.3 | c4wslA\_ | 196-500 | 100.0 | 18% | 237.8 | c6nf8o\_ | 176-481 | 100.0 | 9% | 267.6 | c3jd5o\_ | 179-481 | 100.0 | 9% | 263.5 | c6neqo\_ | 183-483 | 100.0 | 10% | 263.7 | c4n2sA\_ | 192-385 | 100.0 | 10% | 201.4 | c4leuA\_ | 235-423 | 99.9 | 11% | 198.7 | c4xgmA\_ | 231-562 | 99.9 | 11% | 212.2 | c3eiqC\_ | 234-496 | 99.9 | 10% | 206.2 | c3spaA\_ | 193-503 | 99.9 | 9% | 199.2 | c4uzyA\_ | 31-515 | 99.9 | 11% | 141.1 | c5izwA\_ | 351-448 | 99.8 | 17% | 153.5 | c5mqfO\_ | 52-514 | 99.8 | 10% | 118.5 | c2xpiA\_ | 8-497 | 99.8 | 9% | 97.3 | c4ui9C\_ | 94-502 | 99.8 | 9% | 101.7 | c5mqfM\_ | 8-516 | 99.8 | 12% | 96.6 | c5o9zG\_ | 30-498 | 99.8 | 10% | 96.5 | c4ui9K\_ | 8-498 | 99.7 | 9% | 93.6 | c6q6hK\_ | 8-502 | 99.7 | 9% | 92.4 | d2ooea1 | 31-491 | 99.7 | 8% | 100.7 | c6ff7M\_ | 8-514 | 99.6 | 12% | 83.8 | c4hnxA\_ | 32-508 | 99.6 | 9% | 83.5 | c5ganJ\_ | 94-498 | 99.6 | 10% | 85.8 | c4ui9Y\_ | 28-500 | 99.6 | 9% | 82.0 | c6erqA\_ | 250-497 | 99.6 | 10% | 115.3 | c4e85B\_ | 31-498 | 99.5 | 9% | 78.4 | d1w3ba\_ | 102-505 | 99.5 | 11% | 78.0 | c2uy1A\_ | 32-483 | 99.5 | 10% | 77.6 | c5dseA\_ | 99-516 | 99.5 | 12% | 82.0 | c2uy1B\_ | 59-483 | 99.5 | 10% | 94.0 | c4zlhB\_ | 179-483 | 99.5 | 14% | 85.6 | c6g70A\_ | 177-511 | 99.4 | 13% | 76.3 | c5dseC\_ | 43-490 | 99.4 | 12% | 72.7 | c5lj3S\_ | 55-516 | 99.4 | 9% | 72.4 | c4ebaC\_ | 46-500 | 99.4 | 11% | 72.2 | c5wsgd\_ | 249-502 | 99.4 | 10% | 74.5 | c4kvmA\_ | 27-509 | 99.4 | 10% | 70.9 | c5nnrD\_ | 27-516 | 99.4 | 10% | 70.6 | c3jb9R\_ | 248-490 | 99.3 | 12% | 77.2 | c6c95A\_ | 26-515 | 99.3 | 10% | 70.2 | c5c9sB\_ | 206-483 | 99.3 | 8% | 77.0 | c5gmkd\_ | 214-502 | 99.3 | 8% | 70.2 | c4bujF\_ | 100-503 | 99.3 | 8% | 68.5 | c5aioA\_ | 176-483 | 99.2 | 7% | 66.3 | c3mkrA\_ | 204-490 | 99.2 | 12% | 71.8 | c4g1tB\_ | 179-516 | 99.1 | 11% | 66.3 | c4rg6B\_ | 32-516 | 99.1 | 8% | 62.0 | c1fchB\_ | 231-515 | 99.1 | 12% | 61.5 | c6af0A\_ | 29-525 | 99.0 | 9% | 59.9 | c3zpjA\_ | 204-531 | 99.0 | 14% | 76.0 | c3mv3B\_ | 204-465 | 99.0 | 12% | 60.9 | c4r7sA\_ | 234-498 | 99.0 | 10% | 59.1 | c3hymB\_ | 196-503 | 99.0 | 11% | 58.4 | c1xi4D\_ | 201-514 | 98.9 | 12% | 64.8 | c5ctqD\_ | 55-483 | 98.9 | 9% | 57.1 | c2ho1B\_ | 269-499 | 98.9 | 13% | 56.6 | c4jspA\_ | 175-526 | 98.9 | 10% | 61.2 | c5udjA\_ | 182-590 | 98.9 | 12% | 56.1 | d1hz4a\_ | 194-483 | 98.8 | 10% | 55.3 | d1fcha\_ | 231-515 | 98.8 | 11% | 55.1 | c3pe3D\_ | 353-508 | 98.8 | 13% | 61.4 | c5jqyA\_ | 310-515 | 98.8 | 10% | 56.5 | c2gw1A\_ | 198-515 | 98.8 | 9% | 54.7 | c4d18J\_ | 328-515 | 98.8 | 10% | 58.3 | c4hotA\_ | 50-500 | 98.8 | 9% | 54.1 | d2onda1 | 281-491 | 98.7 | 9% | 55.3 | c2vq2A\_ | 269-497 | 98.7 | 18% | 53.3 | c4ynvA\_ | 231-511 | 98.7 | 9% | 54.0 | c3iegB\_ | 196-502 | 98.7 | 10% | 52.7 | c3fp4A\_ | 25-511 | 98.6 | 8% | 51.7 | d1dcea1 | 197-499 | 98.6 | 12% | 59.4 | c3q75A\_ | 197-495 | 98.6 | 11% | 53.1 | c3cvpA\_ | 231-515 | 98.6 | 11% | 50.5 | c4houB\_ | 265-480 | 98.5 | 10% | 51.5 | c4xi0E\_ | 313-492 | 98.5 | 11% | 52.0 | c2q7fA\_ | 298-494 | 98.5 | 12% | 50.1 | d1qsaa1 | 99-515 | 98.5 | 10% | 48.6 | d2h6fa1 | 235-503 | 98.5 | 8% | 48.5 | c5y88I\_ | 182-483 | 98.4 | 8% | 48.1 | c4eqfA\_ | 231-511 | 98.4 | 9% | 48.1 | c5zypA\_ | 264-503 | 98.4 | 13% | 48.0 | c3draA\_ | 235-494 | 98.4 | 7% | 48.0 | c5gjqQ\_ | 209-503 | 98.4 | 9% | 49.0 | c4n5cH\_ | 197-488 | 98.4 | 12% | 49.5 | c3as5A\_ | 310-497 | 98.4 | 11% | 53.6 | c3uq3A\_ | 234-505 | 98.3 | 8% | 47.7 | c2r5sB\_ | 328-483 | 98.3 | 12% | 57.4 | c3vtxB\_ | 313-494 | 98.3 | 13% | 46.9 | c2y4tA\_ | 196-502 | 98.3 | 10% | 45.9 | c5tqbB\_ | 231-501 | 98.3 | 10% | 45.7 | c2hyzA\_ | 353-490 | 98.3 | 13% | 48.7 | c5xi8A\_ | 353-511 | 98.3 | 13% | 48.5 | d1d8da\_ | 197-474 | 98.2 | 9% | 45.0 | c6gmhQ\_ | 194-501 | 98.2 | 10% | 44.4 | c5zyqA\_ | 282-516 | 98.2 | 10% | 44.1 |
