## Supplementary material for "The endosomal TbTpr86/TbUsp7/SkpZ (TUS) complex controls surface protein abundance in trypanosomes": DA1: query.ini

```
EXPDTA    THEORETICAL MODEL, MODELLER 9.10 2019/12/31 19:47:54
REMARK   6 MODELLER BEST TEMPLATE % SEQ ID: 100.000
ATOM      1  N   MET     1      20.328 -25.919   5.484  1.00  0.00           N
ATOM      2  CA  MET     1      18.923 -26.115   5.786 99.00  0.00           C
ATOM      3  CB  MET     1      18.722 -27.272   6.805  1.00  0.00           C
ATOM      4  CG  MET     1      17.260 -27.575   7.205  1.00  0.00           C
ATOM      5  SD  MET     1      17.204 -28.951   8.398  1.00  0.00           S
ATOM      6  CE  MET     1      15.402 -29.038   8.644  1.00  0.00           C
ATOM      7  C   MET     1      17.585 -22.637   7.206  1.00  0.00           C
ATOM      8  O   MET     1      18.339 -21.672   7.310  1.00  0.00           O
ATOM      9  N   GLY     2      16.961 -23.814   7.004  1.00  0.00           N
ATOM     10  CA  GLY     2      16.319 -24.560   8.076 99.00  0.00           C
ATOM     11  C   GLY     2      14.526 -28.347   6.361  1.00  0.00           C
ATOM     12  O   GLY     2      14.625 -28.604   5.163  1.00  0.00           O
ATOM     13  N   ALA     3      14.158 -28.531   7.652  1.00  0.00           N
ATOM     14  CA  ALA     3      13.974 -27.492   8.659 99.00  0.00           C
ATOM     15  CB  ALA     3      12.956 -26.461   8.118  1.00  0.00           C
ATOM     16  C   ALA     3      12.997 -28.699  11.859  1.00  0.00           C
ATOM     17  O   ALA     3      12.829 -29.895  12.092  1.00  0.00           O
ATOM     18  N   GLU     4      13.216 -27.432  11.458  1.00  0.00           N
ATOM     19  CA  GLU     4      12.476 -26.279  11.936 99.00  0.00           C
ATOM     20  CB  GLU     4      12.579 -26.152  13.479  1.00  0.00           C
ATOM     21  CG  GLU     4      11.822 -24.948  14.109  1.00  0.00           C
ATOM     22  CD  GLU     4      11.924 -24.822  15.631  1.00  0.00           C
ATOM     23  OE1 GLU     4      11.306 -23.843  16.126  1.00  0.00           O
ATOM     24  OE2 GLU     4      12.600 -25.649  16.286  1.00  0.00           O
ATOM     25  C   GLU     4      13.959 -23.067  10.087  1.00  0.00           C
ATOM     26  O   GLU     4      14.880 -23.169   9.277  1.00  0.00           O
ATOM     27  N   LYS     5      13.065 -22.324  10.770  1.00  0.00           N
ATOM     28  CA  LYS     5      11.675 -22.701  10.945 99.00  0.00           C
ATOM     29  CB  LYS     5      11.545 -24.089  11.637  1.00  0.00           C
ATOM     30  CG  LYS     5      10.103 -24.589  11.872  1.00  0.00           C
ATOM     31  CD  LYS     5      10.040 -25.965  12.559  1.00  0.00           C
ATOM     32  CE  LYS     5       8.598 -26.440  12.782  1.00  0.00           C
ATOM     33  NZ  LYS     5       8.586 -27.747  13.435  1.00  0.00           N
ATOM     34  C   LYS     5       9.852 -19.706  13.194  1.00  0.00           C
ATOM     35  O   LYS     5      10.491 -18.730  13.577  1.00  0.00           O
ATOM     36  N   LYS     6       9.315 -20.836  12.692  1.00  0.00           N
ATOM     37  CA  LYS     6       8.945 -21.978  13.505 99.00  0.00           C
ATOM     38  CB  LYS     6       7.913 -21.585  14.603  1.00  0.00           C
ATOM     39  CG  LYS     6       7.449 -22.730  15.528  1.00  0.00           C
ATOM     40  CD  LYS     6       6.431 -22.279  16.592  1.00  0.00           C
ATOM     41  CE  LYS     6       5.985 -23.434  17.499  1.00  0.00           C
ATOM     42  NZ  LYS     6       5.022 -22.962  18.490  1.00  0.00           N
ATOM     43  C   LYS     6       7.097 -25.448  10.435  1.00  0.00           C
ATOM     44  O   LYS     6       7.025 -25.243   9.227  1.00  0.00           O
ATOM     45  N   ILE     7       7.110 -25.823  11.729  1.00  0.00           N
ATOM     46  CA  ILE     7       6.699 -24.967  12.831 99.00  0.00           C
ATOM     47  CB  ILE     7       5.243 -24.412  12.659  1.00  0.00           C
ATOM     48  CG2 ILE     7       4.194 -25.495  12.325  1.00  0.00           C
ATOM     49  CG1 ILE     7       4.786 -23.497  13.823  1.00  0.00           C
ATOM     50  CD1 ILE     7       3.359 -22.945  13.665  1.00  0.00           C
ATOM     51  C   ILE     7       7.149 -27.081  16.092  1.00  0.00           C
ATOM     52  O   ILE     7       7.567 -28.237  16.036  1.00  0.00           O
ATOM     53  N   ASN     8       6.681 -25.875  16.470  1.00  0.00           N
ATOM     54  CA  ASN     8       5.297 -25.461  16.333 99.00  0.00           C
ATOM     55  CB  ASN     8       4.344 -26.444  17.086  1.00  0.00           C
ATOM     56  CG  ASN     8       2.859 -26.079  16.992  1.00  0.00           C
ATOM     57  OD1 ASN     8       1.976 -26.750  17.529  1.00  0.00           O
ATOM     58  ND2 ASN     8       2.561 -24.977  16.267  1.00  0.00           N
ATOM     59  C   ASN     8       5.155 -21.405  17.915  1.00  0.00           C
ATOM     60  O   ASN     8       6.174 -20.872  18.348  1.00  0.00           O
ATOM     61  N   ARG     9       4.174 -22.173  17.396  1.00  0.00           N
ATOM     62  CA  ARG     9       3.706 -22.022  16.028 99.00  0.00           C
ATOM     63  CB  ARG     9       4.888 -22.150  15.025  1.00  0.00           C
ATOM     64  CG  ARG     9       4.525 -22.008  13.528  1.00  0.00           C
ATOM     65  CD  ARG     9       5.726 -22.142  12.576  1.00  0.00           C
ATOM     66  NE  ARG     9       5.201 -21.981  11.177  1.00  0.00           N
ATOM     67  CZ  ARG     9       5.985 -22.042  10.091  1.00  0.00           C
ATOM     68  NH1 ARG     9       5.415 -21.885   8.898  1.00  0.00           N
ATOM     69  NH2 ARG     9       7.298 -22.225  10.176  1.00  0.00           N
ATOM     70  C   ARG     9       2.297 -23.304  15.669  1.00  0.00           C
ATOM     71  O   ARG     9       1.915 -24.092  16.528  1.00  0.00           O
ATOM     72  N   PRO    10       1.409 -23.820  14.813  1.00  0.00           N
ATOM     73  CA  PRO    10       1.116 -23.221  13.516 99.00  0.00           C
ATOM     74  CD  PRO    10       0.606 -25.013  15.079  1.00  0.00           C
ATOM     75  CB  PRO    10       0.382 -24.357  12.780  1.00  0.00           C
ATOM     76  CG  PRO    10      -0.353 -25.117  13.889  1.00  0.00           C
ATOM     77  C   PRO    10      -0.404 -21.099  13.790  1.00  0.00           C
ATOM     78  O   PRO    10      -0.815 -20.764  14.901  1.00  0.00           O
ATOM     79  N   THR    11      -0.443 -20.857  12.465  1.00  0.00           N
ATOM     80  CA  THR    11      -1.533 -21.275  11.587 99.00  0.00           C
ATOM     81  CB  THR    11      -2.959 -20.793  12.033  1.00  0.00           C
ATOM     82  OG1 THR    11      -3.978 -21.232  11.139  1.00  0.00           O
ATOM     83  CG2 THR    11      -3.127 -19.282  12.222  1.00  0.00           C
ATOM     84  C   THR    11      -0.513 -19.963   7.776  1.00  0.00           C
ATOM     85  O   THR    11       0.539 -19.354   7.591  1.00  0.00           O
ATOM     86  N   ASN    12      -1.637 -20.465   7.228  1.00  0.00           N
ATOM     87  CA  ASN    12      -2.551 -21.355   7.920 99.00  0.00           C
ATOM     88  CB  ASN    12      -3.079 -20.698   9.236  1.00  0.00           C
ATOM     89  CG  ASN    12      -4.054 -21.573  10.030  1.00  0.00           C
ATOM     90  OD1 ASN    12      -4.563 -21.205  11.091  1.00  0.00           O
ATOM     91  ND2 ASN    12      -4.317 -22.793   9.511  1.00  0.00           N
ATOM     92  C   ASN    12      -5.107 -22.042   5.600  1.00  0.00           C
ATOM     93  O   ASN    12      -5.075 -21.546   4.477  1.00  0.00           O
ATOM     94  N   SER    13      -4.915 -22.453   6.868  1.00  0.00           N
ATOM     95  CA  SER    13      -5.942 -23.068   7.699 99.00  0.00           C
ATOM     96  CB  SER    13      -6.578 -24.353   7.087  1.00  0.00           C
ATOM     97  OG  SER    13      -7.582 -24.926   7.935  1.00  0.00           O
ATOM     98  C   SER    13      -3.801 -24.009  11.985  1.00  0.00           C
ATOM     99  O   SER    13      -2.620 -23.721  12.164  1.00  0.00           O
ATOM    100  N   GLY    14      -5.054 -24.462  12.191  1.00  0.00           N
ATOM    101  CA  GLY    14      -5.976 -24.752  11.104 99.00  0.00           C
ATOM    102  C   GLY    14      -9.723 -26.140  12.773  1.00  0.00           C
ATOM    103  O   GLY    14      -9.874 -26.236  13.989  1.00  0.00           O
ATOM    104  N   GLU    15      -9.574 -26.041  11.438  1.00  0.00           N
ATOM    105  CA  GLU    15      -8.958 -27.056  10.603 99.00  0.00           C
ATOM    106  CB  GLU    15      -9.692 -28.416  10.747  1.00  0.00           C
ATOM    107  CG  GLU    15      -9.118 -29.585   9.896  1.00  0.00           C
ATOM    108  CD  GLU    15      -9.841 -30.927  10.037  1.00  0.00           C
ATOM    109  OE1 GLU    15      -9.366 -31.863   9.342  1.00  0.00           O
ATOM    110  OE2 GLU    15     -10.839 -31.027  10.789  1.00  0.00           O
ATOM    111  C   GLU    15      -9.052 -25.752   6.941  1.00  0.00           C
ATOM    112  O   GLU    15      -9.543 -24.657   6.670  1.00  0.00           O
ATOM    113  N   VAL    16      -8.451 -26.817   6.374  1.00  0.00           N
ATOM    114  CA  VAL    16      -8.299 -28.118   7.012 99.00  0.00           C
ATOM    115  CB  VAL    16      -9.665 -28.760   7.429  1.00  0.00           C
ATOM    116  CG1 VAL    16      -9.528 -30.140   8.108  1.00  0.00           C
ATOM    117  CG2 VAL    16     -10.484 -27.836   8.351  1.00  0.00           C
ATOM    118  C   VAL    16      10.618 -50.620 -19.251  1.00  0.00           C
ATOM    119  O   VAL    16      10.945 -50.167 -20.346  1.00  0.00           O
ATOM    120  N   ALA    17      10.217 -51.029 -18.023  1.00  0.00           N
ATOM    121  CA  ALA    17      -0.166 -41.382  -6.009 50.00  0.00           C
ATOM    122  CB  ALA    17      11.601 -51.281 -20.026  1.00  0.00           C
ATOM    123  C   ALA    17      11.393 -53.146 -18.381  1.00  0.00           C
ATOM    124  O   ALA    17      10.917 -54.280 -18.443  1.00  0.00           O
ATOM    125  N   GLU    18      12.520 -52.852 -17.729  1.00  0.00           N
ATOM    126  CA  GLU    18       0.711 -43.032  -6.727 50.00  0.00           C
ATOM    127  CB  GLU    18      14.517 -53.275 -16.319  1.00  0.00           C
ATOM    128  CG  GLU    18      15.405 -54.321 -15.642  1.00  0.00           C
ATOM    129  CD  GLU    18      16.589 -53.599 -15.016  1.00  0.00           C
ATOM    130  OE1 GLU    18      16.664 -52.350 -15.161  1.00  0.00           O
ATOM    131  OE2 GLU    18      17.435 -54.288 -14.384  1.00  0.00           O
ATOM    132  C   GLU    18      12.443 -54.692 -16.089  1.00  0.00           C
ATOM    133  O   GLU    18      12.588 -55.909 -16.006  1.00  0.00           O
ATOM    134  N   ASN    19      11.530 -54.027 -15.385  1.00  0.00           N
ATOM    135  CA  ASN    19       1.257 -43.665  -5.758 50.00  0.00           C
ATOM    136  CB  ASN    19       9.749 -53.757 -13.735  1.00  0.00           C
ATOM    137  CG  ASN    19       8.956 -54.534 -12.694  1.00  0.00           C
ATOM    138  OD1 ASN    19       9.519 -55.073 -11.742  1.00  0.00           O
ATOM    139  ND2 ASN    19       7.606 -54.633 -12.817  1.00  0.00           N
ATOM    140  C   ASN    19       9.866 -55.767 -15.171  1.00  0.00           C
ATOM    141  O   ASN    19       9.838 -56.930 -14.792  1.00  0.00           O
ATOM    142  N   VAL    20       9.232 -55.331 -16.253  1.00  0.00           N
ATOM    143  CA  VAL    20       0.358 -45.411  -5.505 50.00  0.00           C
ATOM    144  CB  VAL    20       7.862 -55.406 -18.258  1.00  0.00           C
ATOM    145  CG1 VAL    20       7.217 -56.267 -19.346  1.00  0.00           C
ATOM    146  CG2 VAL    20       6.755 -54.461 -17.787  1.00  0.00           C
ATOM    147  C   VAL    20       9.366 -57.328 -17.705  1.00  0.00           C
ATOM    148  O   VAL    20       8.898 -58.421 -17.991  1.00  0.00           O
ATOM    149  N   GLY    21      10.651 -57.038 -17.854  1.00  0.00           N
ATOM    150  CA  GLY    21       0.374 -47.285  -6.639 50.00  0.00           C
ATOM    151  C   GLY    21      11.849 -59.133 -17.374  1.00  0.00           C
ATOM    152  O   GLY    21      12.403 -60.170 -17.732  1.00  0.00           O
ATOM    153  N   ALA    22      11.390 -58.948 -16.141  1.00  0.00           N
ATOM    154  CA  ALA    22       1.318 -48.981  -6.466 50.00  0.00           C
ATOM    155  CB  ALA    22      12.530 -59.742 -14.099  1.00  0.00           C
ATOM    156  C   ALA    22      10.111 -60.193 -14.403  1.00  0.00           C
ATOM    157  O   ALA    22      10.090 -60.415 -13.203  1.00  0.00           O
ATOM    158  N   PHE    23       9.009 -60.108 -15.134  1.00  0.00           N
ATOM    159  CA  PHE    23       1.165 -49.285  -5.369 50.00  0.00           C
ATOM    160  CB  PHE    23       6.590 -60.065 -15.581  1.00  0.00           C
ATOM    161  CG  PHE    23       5.283 -60.321 -14.914  1.00  0.00           C
ATOM    162  CD1 PHE    23       4.673 -59.367 -14.069  1.00  0.00           C
ATOM    163  CD2 PHE    23       4.619 -61.545 -15.119  1.00  0.00           C
ATOM    164  CE1 PHE    23       3.421 -59.619 -13.438  1.00  0.00           C
ATOM    165  CE2 PHE    23       3.363 -61.823 -14.499  1.00  0.00           C
ATOM    166  CZ  PHE    23       2.763 -60.855 -13.653  1.00  0.00           C
ATOM    167  C   PHE    23       7.583 -61.700 -13.967  1.00  0.00           C
ATOM    168  O   PHE    23       7.146 -61.888 -12.823  1.00  0.00           O
ATOM    169  N   THR    24       7.963 -62.683 -14.758  1.00  0.00           N
ATOM    170  CA  THR    24       1.409 -52.991  -5.720 50.00  0.00           C
ATOM    171  CB  THR    24       8.308 -65.006 -15.385  1.00  0.00           C
ATOM    172  OG1 THR    24       7.512 -64.867 -16.552  1.00  0.00           O
ATOM    173  CG2 THR    24       8.198 -66.448 -14.862  1.00  0.00           C
ATOM    174  C   THR    24       8.631 -64.163 -13.043  1.00  0.00           C
ATOM    175  O   THR    24       8.150 -64.645 -12.041  1.00  0.00           O
ATOM    176  N   GLU    25       9.861 -63.664 -13.084  1.00  0.00           N
ATOM    177  CA  GLU    25       2.327 -52.394  -6.296 50.00  0.00           C
ATOM    178  CB  GLU    25      12.156 -63.256 -12.314  1.00  0.00           C
ATOM    179  CG  GLU    25      12.731 -64.009 -13.504  1.00  0.00           C
ATOM    180  CD  GLU    25      13.954 -64.827 -13.140  1.00  0.00           C
ATOM    181  OE1 GLU    25      15.083 -64.359 -13.402  1.00  0.00           O
ATOM    182  OE2 GLU    25      13.788 -65.937 -12.592  1.00  0.00           O
ATOM    183  C   GLU    25      10.235 -62.835 -10.760  1.00  0.00           C
ATOM    184  O   GLU    25      10.556 -63.104  -9.603  1.00  0.00           O
ATOM    185  N   VAL    26       9.449 -61.804 -11.064  1.00  0.00           N
ATOM    186  CA  VAL    26       2.530 -49.439  -5.377 50.00  0.00           C
ATOM    187  CB  VAL    26       8.736 -59.489 -10.559  1.00  0.00           C
ATOM    188  CG1 VAL    26       8.103 -58.543  -9.537  1.00  0.00           C
ATOM    189  CG2 VAL    26      10.022 -58.791 -11.007  1.00  0.00           C
ATOM    190  C   VAL    26       7.715 -61.377  -9.311  1.00  0.00           C
ATOM    191  O   VAL    26       7.639 -61.325  -8.091  1.00  0.00           O
ATOM    192  N   MET    27       6.725 -61.847 -10.054  1.00  0.00           N
ATOM    193  CA  MET    27       2.426 -50.888  -4.444 50.00  0.00           C
ATOM    194  CB  MET    27       4.482 -62.749 -10.469  1.00  0.00           C
ATOM    195  CG  MET    27       4.927 -63.970 -11.276  1.00  0.00           C
ATOM    196  SD  MET    27       3.724 -64.534 -12.516  1.00  0.00           S
ATOM    197  CE  MET    27       2.543 -65.193 -11.304  1.00  0.00           C
ATOM    198  C   MET    27       5.759 -63.468  -8.427  1.00  0.00           C
ATOM    199  O   MET    27       4.868 -63.855  -7.672  1.00  0.00           O
ATOM    200  N   VAL    28       6.984 -63.995  -8.416  1.00  0.00           N
ATOM    201  CA  VAL    28       3.325 -53.387  -4.998 50.00  0.00           C
ATOM    202  CB  VAL    28       8.234 -66.107  -8.094  1.00  0.00           C
ATOM    203  CG1 VAL    28       8.723 -67.162  -7.100  1.00  0.00           C
ATOM    204  CG2 VAL    28       7.542 -66.896  -9.207  1.00  0.00           C
ATOM    205  C   VAL    28       8.004 -64.462  -6.214  1.00  0.00           C
ATOM    206  O   VAL    28       7.940 -65.053  -5.139  1.00  0.00           O
ATOM    207  N   PRO    29       8.634 -63.304  -6.356  1.00  0.00           N
ATOM    208  CA  PRO    29       3.990 -50.789  -5.121 50.00  0.00           C
ATOM    209  CD  PRO    29       9.312 -62.036  -8.011  1.00  0.00           C
ATOM    210  CB  PRO    29       9.976 -61.375  -5.628  1.00  0.00           C
ATOM    211  CG  PRO    29      10.334 -61.333  -7.115  1.00  0.00           C
ATOM    212  C   PRO    29       8.282 -62.369  -4.083  1.00  0.00           C
ATOM    213  O   PRO    29       7.064 -62.339  -4.281  1.00  0.00           O
ATOM    214  N   VAL    30       8.830 -62.203  -2.891  1.00  0.00           N
ATOM    215  CA  VAL    30       4.803 -49.361  -4.410 50.00  0.00           C
ATOM    216  CB  VAL    30       8.775 -61.957  -0.506  1.00  0.00           C
ATOM    217  CG1 VAL    30       8.116 -61.328   0.723  1.00  0.00           C
ATOM    218  CG2 VAL    30       8.713 -63.467  -0.264  1.00  0.00           C
ATOM    219  C   VAL    30       8.390 -60.040  -2.070  1.00  0.00           C
ATOM    220  O   VAL    30       9.414 -59.480  -1.656  1.00  0.00           O
ATOM    221  N   TRP    31       7.463 -59.410  -2.784  1.00  0.00           N
ATOM    222  CA  TRP    31       4.163 -46.044  -4.183 50.00  0.00           C
ATOM    223  CB  TRP    31       6.489 -57.689  -4.225  1.00  0.00           C
ATOM    224  CG  TRP    31       6.732 -56.456  -5.063  1.00  0.00           C
ATOM    225  CD2 TRP    31       7.693 -56.305  -6.118  1.00  0.00           C
ATOM    226  CD1 TRP    31       6.114 -55.240  -5.014  1.00  0.00           C
ATOM    227  NE1 TRP    31       6.570 -54.371  -5.895  1.00  0.00           N
ATOM    228  CE2 TRP    31       7.563 -54.982  -6.616  1.00  0.00           C
ATOM    229  CE3 TRP    31       8.657 -57.158  -6.694  1.00  0.00           C
ATOM    230  CZ2 TRP    31       8.368 -54.484  -7.676  1.00  0.00           C
ATOM    231  CZ3 TRP    31       9.470 -56.666  -7.759  1.00  0.00           C
ATOM    232  CH2 TRP    31       9.311 -55.339  -8.232  1.00  0.00           C
ATOM    233  C   TRP    31       7.828 -57.037  -2.203  1.00  0.00           C
ATOM    234  O   TRP    31       8.399 -55.960  -2.392  1.00  0.00           O
ATOM    235  N   SER    32       7.354 -57.427  -1.026  1.00  0.00           N
ATOM    236  CA  SER    32       5.123 -44.346  -3.640 50.00  0.00           C
ATOM    237  CB  SER    32       6.432 -57.187   1.228  1.00  0.00           C
ATOM    238  OG  SER    32       6.904 -58.454   1.663  1.00  0.00           O
ATOM    239  C   SER    32       8.807 -56.503   0.688  1.00  0.00           C
ATOM    240  O   SER    32       9.284 -55.424   1.034  1.00  0.00           O
ATOM    241  N   ARG    33       9.504 -57.623   0.729  1.00  0.00           N
ATOM    242  CA  ARG    33       5.702 -45.168  -4.604 50.00  0.00           C
ATOM    243  CB  ARG    33      11.470 -58.983   1.112  1.00  0.00           C
ATOM    244  CG  ARG    33      10.956 -59.838   2.272  1.00  0.00           C
ATOM    245  CD  ARG    33      11.472 -61.278   2.247  1.00  0.00           C
ATOM    246  NE  ARG    33      10.869 -61.990   3.409  1.00  0.00           N
ATOM    247  CZ  ARG    33      11.100 -63.323   3.587  1.00  0.00           C
ATOM    248  NH1 ARG    33      11.904 -63.746   2.568  1.00  0.00           N
ATOM    249  NH2 ARG    33      10.436 -63.714   4.714  1.00  0.00           N
ATOM    250  C   ARG    33      11.698 -56.713   0.136  1.00  0.00           C
ATOM    251  O   ARG    33      12.624 -56.030   0.562  1.00  0.00           O
ATOM    252  N   LEU    34      11.337 -56.743  -1.150  1.00  0.00           N
ATOM    253  CA  LEU    34       4.831 -43.849  -5.140 50.00  0.00           C
ATOM    254  CB  LEU    34      11.594 -56.367  -3.588  1.00  0.00           C
ATOM    255  CG  LEU    34      11.793 -57.770  -4.172  1.00  0.00           C
ATOM    256  CD1 LEU    34      11.669 -57.750  -5.690  1.00  0.00           C
ATOM    257  CD2 LEU    34      13.125 -58.350  -3.767  1.00  0.00           C
ATOM    258  C   LEU    34      11.906 -54.489  -1.963  1.00  0.00           C
ATOM    259  O   LEU    34      12.882 -53.734  -1.999  1.00  0.00           O
ATOM    260  N   CYS    35      10.667 -54.062  -1.742  1.00  0.00           N
ATOM    261  CA  CYS    35       4.823 -40.547  -4.293 50.00  0.00           C
ATOM    262  CB  CYS    35       8.924 -52.455  -1.182  1.00  0.00           C
ATOM    263  SG  CYS    35       8.170 -52.567  -2.380  1.00  0.00           S
ATOM    264  C   CYS    35      11.166 -52.272  -0.162  1.00  0.00           C
ATOM    265  O   CYS    35      11.810 -51.225  -0.144  1.00  0.00           O
ATOM    266  N   ASP    36      11.092 -53.095   0.880  1.00  0.00           N
ATOM    267  CA  ASP    36       5.929 -40.336  -4.363 50.00  0.00           C
ATOM    268  CB  ASP    36      11.598 -53.913   3.159  1.00  0.00           C
ATOM    269  CG  ASP    36      10.174 -53.799   3.685  1.00  0.00           C
ATOM    270  OD1 ASP    36       9.522 -52.757   3.405  1.00  0.00           O
ATOM    271  OD2 ASP    36       9.720 -54.751   4.373  1.00  0.00           O
ATOM    272  C   ASP    36      13.302 -52.587   1.881  1.00  0.00           C
ATOM    273  O   ASP    36      13.900 -51.611   2.362  1.00  0.00           O
ATOM    274  N   LYS    37      13.896 -53.494   1.118  1.00  0.00           N
ATOM    275  CA  LYS    37       5.875 -40.969  -5.520 50.00  0.00           C
ATOM    276  CB  LYS    37      15.767 -54.621  -0.014  1.00  0.00           C
ATOM    277  CG  LYS    37      15.810 -55.912   0.805  1.00  0.00           C
ATOM    278  CD  LYS    37      16.267 -57.132   0.003  1.00  0.00           C
ATOM    279  CE  LYS    37      16.287 -58.428   0.816  1.00  0.00           C
ATOM    280  NZ  LYS    37      16.707 -59.558  -0.042  1.00  0.00           N
ATOM    281  C   LYS    37      15.603 -52.083   0.061  1.00  0.00           C
ATOM    282  O   LYS    37      16.575 -51.388   0.383  1.00  0.00           O
ATOM    283  N   VAL    38      14.746 -51.725  -0.895  1.00  0.00           N
ATOM    284  CA  VAL    38       5.067 -38.270  -5.406 50.00  0.00           C
ATOM    285  CB  VAL    38      13.961 -50.336  -2.779  1.00  0.00           C
ATOM    286  CG1 VAL    38      13.964 -48.950  -3.427  1.00  0.00           C
ATOM    287  CG2 VAL    38      14.274 -51.310  -3.917  1.00  0.00           C
ATOM    288  C   VAL    38      14.795 -49.237  -0.681  1.00  0.00           C
ATOM    289  O   VAL    38      15.479 -48.235  -0.898  1.00  0.00           O
ATOM    290  N   ILE    39      13.970 -49.338   0.357  1.00  0.00           N
ATOM    291  CA  ILE    39       5.697 -35.855  -4.648 50.00  0.00           C
ATOM    292  CB  ILE    39      12.627 -48.492   2.231  1.00  0.00           C
ATOM    293  CG2 ILE    39      12.529 -47.496   3.399  1.00  0.00           C
ATOM    294  CG1 ILE    39      11.285 -48.375   1.489  1.00  0.00           C
ATOM    295  CD1 ILE    39      10.091 -48.862   2.309  1.00  0.00           C
ATOM    296  C   ILE    39      15.108 -48.129   2.183  1.00  0.00           C
ATOM    297  O   ILE    39      15.580 -47.023   2.446  1.00  0.00           O
ATOM    298  N   ALA    40      15.648 -49.267   2.608  1.00  0.00           N
ATOM    299  CA  ALA    40       6.613 -36.663  -5.399 50.00  0.00           C
ATOM    300  CB  ALA    40      16.989 -50.676   4.106  1.00  0.00           C
ATOM    301  C   ALA    40      18.116 -48.973   2.632  1.00  0.00           C
ATOM    302  O   ALA    40      19.180 -48.779   3.213  1.00  0.00           O
ATOM    303  N   ALA    41      18.024 -48.915   1.308  1.00  0.00           N
ATOM    304  CA  ALA    41       5.979 -36.075  -6.279 50.00  0.00           C
ATOM    305  CB  ALA    41      19.473 -49.504  -0.592  1.00  0.00           C
ATOM    306  C   ALA    41      19.096 -47.067  -0.010  1.00  0.00           C
ATOM    307  O   ALA    41      20.050 -46.546  -0.592  1.00  0.00           O
ATOM    308  N   TYR    42      17.943 -46.434   0.183  1.00  0.00           N
ATOM    309  CA  TYR    42       5.518 -32.734  -5.637 50.00  0.00           C
ATOM    310  CB  TYR    42      19.133 -44.373  -0.385  1.00  0.00           C
ATOM    311  CG  TYR    42      19.806 -44.278   0.941  1.00  0.00           C
ATOM    312  CD1 TYR    42      20.790 -45.212   1.291  1.00  0.00           C
ATOM    313  CD2 TYR    42      19.482 -43.260   1.864  1.00  0.00           C
ATOM    314  CE1 TYR    42      21.450 -45.154   2.534  1.00  0.00           C
ATOM    315  CE2 TYR    42      20.144 -43.186   3.132  1.00  0.00           C
ATOM    316  CZ  TYR    42      21.127 -44.145   3.448  1.00  0.00           C
ATOM    317  OH  TYR    42      21.787 -44.117   4.658  1.00  0.00           O
ATOM    318  C   TYR    42      16.914 -44.858  -1.390  1.00  0.00           C
ATOM    319  O   TYR    42      16.541 -43.731  -1.747  1.00  0.00           O
ATOM    320  N   LYS    43      16.590 -45.979  -2.032  1.00  0.00           N
ATOM    321  CA  LYS    43       4.491 -33.880  -5.441 50.00  0.00           C
ATOM    322  CB  LYS    43      16.391 -47.093  -4.226  1.00  0.00           C
ATOM    323  CG  LYS    43      17.863 -46.981  -4.628  1.00  0.00           C
ATOM    324  CD  LYS    43      18.329 -48.100  -5.562  1.00  0.00           C
ATOM    325  CE  LYS    43      19.809 -48.004  -5.941  1.00  0.00           C
ATOM    326  NZ  LYS    43      20.186 -49.149  -6.800  1.00  0.00           N
ATOM    327  C   LYS    43      14.398 -45.929  -3.213  1.00  0.00           C
ATOM    328  O   LYS    43      13.721 -46.657  -3.911  1.00  0.00           O
ATOM    329  N   ALA    44      13.879 -45.063  -2.350  1.00  0.00           N
ATOM    330  CA  ALA    44       4.479 -32.821  -4.291 50.00  0.00           C
ATOM    331  CB  ALA    44      12.061 -43.863  -1.123  1.00  0.00           C
ATOM    332  C   ALA    44      11.704 -44.678  -3.505  1.00  0.00           C
ATOM    333  O   ALA    44      10.626 -45.194  -3.714  1.00  0.00           O
ATOM    334  N   GLY    45      12.299 -43.919  -4.409  1.00  0.00           N
ATOM    335  CA  GLY    45       3.393 -31.852  -4.273 50.00  0.00           C
ATOM    336  C   GLY    45      11.477 -44.834  -6.504  1.00  0.00           C
ATOM    337  O   GLY    45      10.489 -45.022  -7.228  1.00  0.00           O
ATOM    338  N   GLU    46      12.480 -45.693  -6.381  1.00  0.00           N
ATOM    339  CA  GLU    46       3.211 -35.362  -4.946 50.00  0.00           C
ATOM    340  CB  GLU    46      13.834 -47.671  -6.896  1.00  0.00           C
ATOM    341  CG  GLU    46      14.895 -47.020  -7.786  1.00  0.00           C
ATOM    342  CD  GLU    46      14.405 -47.083  -9.225  1.00  0.00           C
ATOM    343  OE1 GLU    46      14.042 -48.202  -9.678  1.00  0.00           O
ATOM    344  OE2 GLU    46      14.388 -46.014  -9.891  1.00  0.00           O
ATOM    345  C   GLU    46      11.403 -47.864  -6.310  1.00  0.00           C
ATOM    346  O   GLU    46      10.706 -48.623  -6.973  1.00  0.00           O
ATOM    347  N   ASP    47      11.285 -47.753  -4.989  1.00  0.00           N
ATOM    348  CA  ASP    47       3.858 -36.670  -4.179 50.00  0.00           C
ATOM    349  CB  ASP    47      10.413 -48.284  -2.760  1.00  0.00           C
ATOM    350  CG  ASP    47       9.589 -49.342  -2.040  1.00  0.00           C
ATOM    351  OD1 ASP    47       9.114 -50.288  -2.723  1.00  0.00           O
ATOM    352  OD2 ASP    47       9.423 -49.217  -0.797  1.00  0.00           O
ATOM    353  C   ASP    47       8.910 -48.224  -4.786  1.00  0.00           C
ATOM    354  O   ASP    47       8.170 -49.104  -5.205  1.00  0.00           O
ATOM    355  N   ILE    48       8.576 -46.944  -4.787  1.00  0.00           N
ATOM    356  CA  ILE    48       3.187 -34.721  -3.275 50.00  0.00           C
ATOM    357  CB  ILE    48       7.408 -44.915  -5.426  1.00  0.00           C
ATOM    358  CG2 ILE    48       6.263 -44.276  -6.230  1.00  0.00           C
ATOM    359  CG1 ILE    48       7.320 -44.282  -4.027  1.00  0.00           C
ATOM    360  CD1 ILE    48       7.641 -42.788  -4.013  1.00  0.00           C
ATOM    361  C   ILE    48       7.191 -46.954  -6.844  1.00  0.00           C
ATOM    362  O   ILE    48       6.101 -47.416  -7.219  1.00  0.00           O
ATOM    363  N   VAL    49       8.257 -46.877  -7.643  1.00  0.00           N
ATOM    364  CA  VAL    49       2.296 -35.954  -3.948 50.00  0.00           C
ATOM    365  CB  VAL    49       9.526 -47.134  -9.722  1.00  0.00           C
ATOM    366  CG1 VAL    49       9.617 -47.798 -11.097  1.00  0.00           C
ATOM    367  CG2 VAL    49       9.825 -45.654  -9.971  1.00  0.00           C
ATOM    368  C   VAL    49       7.777 -48.859  -9.119  1.00  0.00           C
ATOM    369  O   VAL    49       6.901 -49.215  -9.920  1.00  0.00           O
ATOM    370  N   VAL    50       8.394 -49.698  -8.284  1.00  0.00           N
ATOM    371  CA  VAL    50       2.601 -39.598  -4.149 50.00  0.00           C
ATOM    372  CB  VAL    50       9.077 -51.809  -7.216  1.00  0.00           C
ATOM    373  CG1 VAL    50       8.724 -53.270  -6.928  1.00  0.00           C
ATOM    374  CG2 VAL    50      10.529 -51.836  -7.699  1.00  0.00           C
ATOM    375  C   VAL    50       6.686 -51.378  -7.757  1.00  0.00           C
ATOM    376  O   VAL    50       5.996 -52.214  -8.328  1.00  0.00           O
ATOM    377  N   LEU    51       6.246 -50.684  -6.715  1.00  0.00           N
ATOM    378  CA  LEU    51       2.867 -39.244  -3.053 50.00  0.00           C
ATOM    379  CB  LEU    51       4.613 -50.009  -5.009  1.00  0.00           C
ATOM    380  CG  LEU    51       3.704 -50.594  -3.922  1.00  0.00           C
ATOM    381  CD1 LEU    51       2.722 -49.551  -3.484  1.00  0.00           C
ATOM    382  CD2 LEU    51       2.977 -51.873  -4.352  1.00  0.00           C
ATOM    383  C   LEU    51       3.895 -50.533  -7.351  1.00  0.00           C
ATOM    384  O   LEU    51       2.996 -51.304  -7.648  1.00  0.00           O
ATOM    385  N   GLN    52       4.088 -49.374  -7.968  1.00  0.00           N
ATOM    386  CA  GLN    52       1.931 -37.581  -2.704 50.00  0.00           C
ATOM    387  CB  GLN    52       3.768 -47.595  -9.601  1.00  0.00           C
ATOM    388  CG  GLN    52       3.567 -46.420  -8.641  1.00  0.00           C
ATOM    389  CD  GLN    52       4.238 -45.197  -9.249  1.00  0.00           C
ATOM    390  OE1 GLN    52       4.890 -45.282 -10.289  1.00  0.00           O
ATOM    391  NE2 GLN    52       4.116 -43.991  -8.632  1.00  0.00           N
ATOM    392  C   GLN    52       3.189 -49.917 -10.208  1.00  0.00           C
ATOM    393  O   GLN    52       2.151 -50.093 -10.830  1.00  0.00           O
ATOM    394  N   HIS    53       4.321 -50.550 -10.504  1.00  0.00           N
ATOM    395  CA  HIS    53       1.365 -40.281  -3.406 50.00  0.00           C
ATOM    396  ND1 HIS    53       7.960 -50.751 -12.666  1.00  0.00           N
ATOM    397  CG  HIS    53       6.586 -50.775 -12.550  1.00  0.00           C
ATOM    398  CB  HIS    53       5.818 -51.890 -11.904  1.00  0.00           C
ATOM    399  NE2 HIS    53       7.280 -48.957 -13.567  1.00  0.00           N
ATOM    400  CD2 HIS    53       6.163 -49.626 -13.127  1.00  0.00           C
ATOM    401  CE1 HIS    53       8.350 -49.670 -13.271  1.00  0.00           C
ATOM    402  C   HIS    53       3.621 -52.757 -11.194  1.00  0.00           C
ATOM    403  O   HIS    53       2.801 -53.264 -11.951  1.00  0.00           O
ATOM    404  N   PHE    54       3.753 -53.110  -9.910  1.00  0.00           N
ATOM    405  CA  PHE    54       1.911 -42.948  -3.117 50.00  0.00           C
ATOM    406  CB  PHE    54       3.200 -54.767  -7.924  1.00  0.00           C
ATOM    407  CG  PHE    54       4.606 -55.192  -7.676  1.00  0.00           C
ATOM    408  CD1 PHE    54       5.513 -55.436  -8.732  1.00  0.00           C
ATOM    409  CD2 PHE    54       5.065 -55.361  -6.357  1.00  0.00           C
ATOM    410  CE1 PHE    54       6.857 -55.838  -8.485  1.00  0.00           C
ATOM    411  CE2 PHE    54       6.407 -55.763  -6.081  1.00  0.00           C
ATOM    412  CZ  PHE    54       7.306 -56.004  -7.152  1.00  0.00           C
ATOM    413  C   PHE    54       1.581 -54.200  -9.517  1.00  0.00           C
ATOM    414  O   PHE    54       0.904 -55.132  -9.960  1.00  0.00           O
ATOM    415  N   GLU    55       0.997 -53.065  -9.141  1.00  0.00           N
ATOM    416  CA  GLU    55       1.634 -41.652  -1.974 50.00  0.00           C
ATOM    417  CB  GLU    55      -1.093 -51.654  -8.622  1.00  0.00           C
ATOM    418  CG  GLU    55      -2.622 -51.665  -8.680  1.00  0.00           C
ATOM    419  CD  GLU    55      -3.125 -50.356  -8.088  1.00  0.00           C
ATOM    420  OE1 GLU    55      -2.272 -49.541  -7.645  1.00  0.00           O
ATOM    421  OE2 GLU    55      -4.368 -50.154  -8.070  1.00  0.00           O
ATOM    422  C   GLU    55      -0.902 -52.859 -10.665  1.00  0.00           C
ATOM    423  O   GLU    55      -1.743 -53.643 -11.115  1.00  0.00           O
ATOM    424  N   GLU    56      -0.345 -51.929 -11.416  1.00  0.00           N
ATOM    425  CA  GLU    56       0.547 -40.687  -2.056 50.00  0.00           C
ATOM    426  CB  GLU    56      -0.725 -50.353 -13.414  1.00  0.00           C
ATOM    427  CG  GLU    56      -1.360 -49.170 -12.680  1.00  0.00           C
ATOM    428  CD  GLU    56      -1.039 -47.904 -13.462  1.00  0.00           C
ATOM    429  OE1 GLU    56      -0.395 -48.021 -14.538  1.00  0.00           O
ATOM    430  OE2 GLU    56      -1.434 -46.804 -12.993  1.00  0.00           O
ATOM    431  C   GLU    56      -0.408 -52.703 -13.754  1.00  0.00           C
ATOM    432  O   GLU    56      -1.032 -52.962 -14.787  1.00  0.00           O
ATOM    433  N   LEU    57       0.772 -53.283 -13.488  1.00  0.00           N
ATOM    434  CA  LEU    57       0.409 -43.275  -3.110 50.00  0.00           C
ATOM    435  CB  LEU    57       2.944 -54.181 -14.654  1.00  0.00           C
ATOM    436  CG  LEU    57       3.528 -52.832 -15.079  1.00  0.00           C
ATOM    437  CD1 LEU    57       5.052 -52.789 -15.174  1.00  0.00           C
ATOM    438  CD2 LEU    57       3.073 -52.334 -16.450  1.00  0.00           C
ATOM    439  C   LEU    57       1.182 -55.613 -14.120  1.00  0.00           C
ATOM    440  O   LEU    57       1.786 -56.507 -14.719  1.00  0.00           O
ATOM    441  N   ARG    58       0.310 -55.921 -13.151  1.00  0.00           N
ATOM    442  CA  ARG    58       0.836 -46.263  -2.803 50.00  0.00           C
ATOM    443  CB  ARG    58       0.209 -58.361 -13.944  1.00  0.00           C
ATOM    444  CG  ARG    58      -0.666 -58.199 -15.189  1.00  0.00           C
ATOM    445  CD  ARG    58      -0.407 -59.258 -16.262  1.00  0.00           C
ATOM    446  NE  ARG    58      -1.293 -58.950 -17.419  1.00  0.00           N
ATOM    447  CZ  ARG    58      -1.197 -59.682 -18.567  1.00  0.00           C
ATOM    448  NH1 ARG    58      -0.222 -60.619 -18.382  1.00  0.00           N
ATOM    449  NH2 ARG    58      -2.114 -59.192 -19.451  1.00  0.00           N
ATOM    450  C   ARG    58       0.607 -58.048 -11.732  1.00  0.00           C
ATOM    451  O   ARG    58       0.326 -59.228 -11.502  1.00  0.00           O
ATOM    452  N   ALA    59       1.492 -57.385 -10.984  1.00  0.00           N
ATOM    453  CA  ALA    59       2.000 -46.719  -3.182 50.00  0.00           C
ATOM    454  CB  ALA    59       3.506 -57.278  -9.550  1.00  0.00           C
ATOM    455  C   ALA    59       1.264 -57.769  -8.632  1.00  0.00           C
ATOM    456  O   ALA    59       1.544 -56.984  -7.722  1.00  0.00           O
ATOM    457  N   THR    60       0.162 -58.502  -8.604  1.00  0.00           N
ATOM    458  CA  THR    60       2.350 -46.828  -2.162 50.00  0.00           C
ATOM    459  CB  THR    60      -2.263 -58.548  -6.930  1.00  0.00           C
ATOM    460  OG1 THR    60      -2.374 -59.909  -6.539  1.00  0.00           O
ATOM    461  CG2 THR    60      -3.380 -58.214  -7.933  1.00  0.00           C
ATOM    462  C   THR    60      -0.582 -58.754  -6.178  1.00  0.00           C
ATOM    463  O   THR    60      -1.043 -58.190  -5.182  1.00  0.00           O
ATOM    464  N   PRO    61       0.226 -59.809  -6.092  1.00  0.00           N
ATOM    465  CA  PRO    61       3.333 -48.536  -2.519 50.00  0.00           C
ATOM    466  CD  PRO    61       2.589 -61.884  -6.242  1.00  0.00           C
ATOM    467  CB  PRO    61       1.639 -61.126  -3.988  1.00  0.00           C
ATOM    468  CG  PRO    61       2.300 -62.252  -4.785  1.00  0.00           C
ATOM    469  C   PRO    61       1.550 -59.233  -4.178  1.00  0.00           C
ATOM    470  O   PRO    61       1.384 -58.863  -3.013  1.00  0.00           O
ATOM    471  N   GLU    62       2.521 -58.724  -4.939  1.00  0.00           N
ATOM    472  CA  GLU    62       3.484 -45.935  -2.952 50.00  0.00           C
ATOM    473  CB  GLU    62       4.605 -57.238  -5.290  1.00  0.00           C
ATOM    474  CG  GLU    62       5.653 -58.316  -5.573  1.00  0.00           C
ATOM    475  CD  GLU    62       5.124 -59.204  -6.691  1.00  0.00           C
ATOM    476  OE1 GLU    62       4.036 -58.881  -7.238  1.00  0.00           O
ATOM    477  OE2 GLU    62       5.801 -60.217  -7.012  1.00  0.00           O
ATOM    478  C   GLU    62       2.693 -56.441  -4.112  1.00  0.00           C
ATOM    479  O   GLU    62       2.871 -55.836  -3.051  1.00  0.00           O
ATOM    480  N   LEU    63       1.838 -55.984  -5.015  1.00  0.00           N
ATOM    481  CA  LEU    63       3.101 -43.076  -2.134 50.00  0.00           C
ATOM    482  CB  LEU    63      -0.065 -54.376  -5.825  1.00  0.00           C
ATOM    483  CG  LEU    63      -0.670 -52.997  -5.554  1.00  0.00           C
ATOM    484  CD1 LEU    63       0.326 -51.839  -5.577  1.00  0.00           C
ATOM    485  CD2 LEU    63      -1.748 -52.559  -6.544  1.00  0.00           C
ATOM    486  C   LEU    63       0.149 -54.900  -3.578  1.00  0.00           C
ATOM    487  O   LEU    63      -0.339 -53.915  -3.018  1.00  0.00           O
ATOM    488  N   ASP    64      -0.004 -56.043  -3.104  1.00  0.00           N
ATOM    489  CA  ASP    64       3.822 -44.492  -1.482 50.00  0.00           C
ATOM    490  CB  ASP    64      -0.607 -57.877  -1.613  1.00  0.00           C
ATOM    491  CG  ASP    64       0.642 -58.040  -0.804  1.00  0.00           C
ATOM    492  OD1 ASP    64       1.726 -58.222  -1.402  1.00  0.00           O
ATOM    493  OD2 ASP    64       0.544 -57.973   0.440  1.00  0.00           O
ATOM    494  C   ASP    64      -0.605 -55.539  -0.780  1.00  0.00           C
ATOM    495  O   ASP    64       0.347 -54.776  -0.759  1.00  0.00           O
ATOM    496  N   VAL    65      -1.389 -55.790   0.265  1.00  0.00           N
ATOM    497  CA  VAL    65       4.681 -42.774  -0.976 50.00  0.00           C
ATOM    498  CB  VAL    65      -1.707 -54.961   2.954  1.00  0.00           C
ATOM    499  CG1 VAL    65      -1.056 -53.838   3.764  1.00  0.00           C
ATOM    500  CG2 VAL    65      -3.213 -54.777   3.153  1.00  0.00           C
ATOM    501  C   VAL    65       0.105 -54.833   2.080  1.00  0.00           C
ATOM    502  O   VAL    65       0.510 -53.741   2.488  1.00  0.00           O
ATOM    503  N   ILE    66       0.864 -55.914   2.160  1.00  0.00           N
ATOM    504  CA  ILE    66       5.347 -43.403  -1.882 50.00  0.00           C
ATOM    505  CB  ILE    66       3.305 -56.870   2.925  1.00  0.00           C
ATOM    506  CG2 ILE    66       4.629 -56.289   3.449  1.00  0.00           C
ATOM    507  CG1 ILE    66       2.940 -57.991   3.912  1.00  0.00           C
ATOM    508  CD1 ILE    66       3.909 -59.172   3.879  1.00  0.00           C
ATOM    509  C   ILE    66       3.145 -54.889   1.987  1.00  0.00           C
ATOM    510  O   ILE    66       3.746 -53.970   2.551  1.00  0.00           O
ATOM    511  N   ALA    67       3.294 -55.123   0.688  1.00  0.00           N
ATOM    512  CA  ALA    67       4.714 -42.110  -2.494 50.00  0.00           C
ATOM    513  CB  ALA    67       4.343 -54.848  -1.536  1.00  0.00           C
ATOM    514  C   ALA    67       3.645 -52.876  -0.219  1.00  0.00           C
ATOM    515  O   ALA    67       4.391 -51.894  -0.174  1.00  0.00           O
ATOM    516  N   HIS    68       2.334 -52.754  -0.356  1.00  0.00           N
ATOM    517  CA  HIS    68       4.259 -39.317  -1.601 50.00  0.00           C
ATOM    518  ND1 HIS    68      -1.587 -49.557  -1.010  1.00  0.00           N
ATOM    519  CG  HIS    68      -0.272 -49.916  -0.797  1.00  0.00           C
ATOM    520  CB  HIS    68       0.178 -51.316  -0.501  1.00  0.00           C
ATOM    521  NE2 HIS    68      -0.470 -47.765  -1.189  1.00  0.00           N
ATOM    522  CD2 HIS    68       0.431 -48.765  -0.914  1.00  0.00           C
ATOM    523  CE1 HIS    68      -1.683 -48.283  -1.239  1.00  0.00           C
ATOM    524  C   HIS    68       2.016 -50.660   0.799  1.00  0.00           C
ATOM    525  O   HIS    68       2.596 -49.571   0.750  1.00  0.00           O
ATOM    526  N   ASP    69       1.631 -51.199   1.948  1.00  0.00           N
ATOM    527  CA  ASP    69       5.285 -38.150  -1.276 50.00  0.00           C
ATOM    528  CB  ASP    69       1.430 -51.310   4.472  1.00  0.00           C
ATOM    529  CG  ASP    69      -0.091 -51.316   4.532  1.00  0.00           C
ATOM    530  OD1 ASP    69      -0.716 -50.535   3.767  1.00  0.00           O
ATOM    531  OD2 ASP    69      -0.647 -52.102   5.345  1.00  0.00           O
ATOM    532  C   ASP    69       3.296 -50.164   3.534  1.00  0.00           C
ATOM    533  O   ASP    69       3.578 -49.144   4.169  1.00  0.00           O
ATOM    534  N   SER    70       4.243 -50.986   3.094  1.00  0.00           N
ATOM    535  CA  SER    70       5.647 -38.218  -2.329 50.00  0.00           C
ATOM    536  CB  SER    70       6.859 -51.466   2.869  1.00  0.00           C
ATOM    537  OG  SER    70       6.951 -51.387   1.454  1.00  0.00           O
ATOM    538  C   SER    70       6.034 -49.373   2.701  1.00  0.00           C
ATOM    539  O   SER    70       6.727 -48.540   3.291  1.00  0.00           O
ATOM    540  N   LEU    71       5.569 -49.195   1.455  1.00  0.00           N
ATOM    541  CA  LEU    71       4.847 -35.735  -2.431 50.00  0.00           C
ATOM    542  CB  LEU    71       5.511 -48.008  -0.727  1.00  0.00           C
ATOM    543  CG  LEU    71       5.881 -46.741  -1.500  1.00  0.00           C
ATOM    544  CD1 LEU    71       7.377 -46.434  -1.559  1.00  0.00           C
ATOM    545  CD2 LEU    71       5.460 -46.734  -2.969  1.00  0.00           C
ATOM    546  C   LEU    71       5.339 -46.709   1.417  1.00  0.00           C
ATOM    547  O   LEU    71       5.978 -45.662   1.436  1.00  0.00           O
ATOM    548  N   ILE    72       4.124 -46.806   1.954  1.00  0.00           N
ATOM    549  CA  ILE    72       5.083 -33.344  -1.396 50.00  0.00           C
ATOM    550  CB  ILE    72       2.044 -45.915   3.000  1.00  0.00           C
ATOM    551  CG2 ILE    72       1.436 -44.814   3.886  1.00  0.00           C
ATOM    552  CG1 ILE    72       1.120 -46.000   1.773  1.00  0.00           C
ATOM    553  CD1 ILE    72      -0.284 -46.507   2.102  1.00  0.00           C
ATOM    554  C   ILE    72       4.339 -45.314   3.880  1.00  0.00           C
ATOM    555  O   ILE    72       4.616 -44.130   4.144  1.00  0.00           O
ATOM    556  N   SER    73       4.741 -46.338   4.626  1.00  0.00           N
ATOM    557  CA  SER    73       6.164 -33.518  -1.725 50.00  0.00           C
ATOM    558  CB  SER    73       5.811 -47.414   6.574  1.00  0.00           C
ATOM    559  OG  SER    73       6.676 -47.175   7.674  1.00  0.00           O
ATOM    560  C   SER    73       6.953 -45.553   5.320  1.00  0.00           C
ATOM    561  O   SER    73       7.510 -44.666   5.980  1.00  0.00           O
ATOM    562  N   ALA    74       7.457 -46.038   4.186  1.00  0.00           N
ATOM    563  CA  ALA    74       5.837 -33.069  -2.760 50.00  0.00           C
ATOM    564  CB  ALA    74       9.213 -46.426   2.486  1.00  0.00           C
ATOM    565  C   ALA    74       8.731 -44.102   3.306  1.00  0.00           C
ATOM    566  O   ALA    74       9.522 -43.328   3.822  1.00  0.00           O
ATOM    567  N   ALA    75       7.838 -43.720   2.401  1.00  0.00           N
ATOM    568  CA  ALA    75       5.149 -30.019  -2.361 50.00  0.00           C
ATOM    569  CB  ALA    75       6.663 -42.177   0.886  1.00  0.00           C
ATOM    570  C   ALA    75       7.377 -41.423   3.155  1.00  0.00           C
ATOM    571  O   ALA    75       7.786 -40.269   3.195  1.00  0.00           O
ATOM    572  N   PRO    76       6.617 -41.956   4.106  1.00  0.00           N
ATOM    573  CA  PRO    76       5.900 -28.629  -1.578 50.00  0.00           C
ATOM    574  CD  PRO    76       5.385 -43.618   3.984  1.00  0.00           C
ATOM    575  CB  PRO    76       5.453 -42.213   6.123  1.00  0.00           C
ATOM    576  CG  PRO    76       4.639 -43.159   5.238  1.00  0.00           C
ATOM    577  C   PRO    76       7.589 -40.887   6.086  1.00  0.00           C
ATOM    578  O   PRO    76       7.832 -39.727   6.384  1.00  0.00           O
ATOM    579  N   ASP    77       8.398 -41.895   6.392  1.00  0.00           N
ATOM    580  CA  ASP    77       6.735 -28.913  -2.401 50.00  0.00           C
ATOM    581  CB  ASP    77      10.261 -43.060   7.508  1.00  0.00           C
ATOM    582  CG  ASP    77      11.420 -42.802   8.460  1.00  0.00           C
ATOM    583  OD1 ASP    77      11.175 -42.208   9.544  1.00  0.00           O
ATOM    584  OD2 ASP    77      12.567 -43.196   8.116  1.00  0.00           O
ATOM    585  C   ASP    77      10.642 -40.848   6.268  1.00  0.00           C
ATOM    586  O   ASP    77      11.462 -40.102   6.802  1.00  0.00           O
ATOM    587  N   LEU    78      10.507 -40.943   4.956  1.00  0.00           N
ATOM    588  CA  LEU    78       5.960 -27.650  -3.053 50.00  0.00           C
ATOM    589  CB  LEU    78      11.409 -40.933   2.708  1.00  0.00           C
ATOM    590  CG  LEU    78      12.063 -42.314   2.795  1.00  0.00           C
ATOM    591  CD1 LEU    78      12.046 -43.119   1.496  1.00  0.00           C
ATOM    592  CD2 LEU    78      13.539 -42.305   3.188  1.00  0.00           C
ATOM    593  C   LEU    78      10.865 -38.753   3.874  1.00  0.00           C
ATOM    594  O   LEU    78      11.533 -37.941   3.226  1.00  0.00           O
ATOM    595  N   THR    79       9.710 -38.435   4.465  1.00  0.00           N
ATOM    596  CA  THR    79       5.742 -24.606  -2.108 50.00  0.00           C
ATOM    597  CB  THR    79      10.172 -36.046   4.766  1.00  0.00           C
ATOM    598  OG1 THR    79      11.280 -36.117   3.881  1.00  0.00           O
ATOM    599  CG2 THR    79      10.652 -36.302   6.205  1.00  0.00           C
ATOM    600  C   THR    79       8.531 -36.724   3.030  1.00  0.00           C
ATOM    601  O   THR    79       8.410 -35.552   2.700  1.00  0.00           O
ATOM    602  N   VAL    80       8.144 -37.725   2.247  1.00  0.00           N
ATOM    603  CA  VAL    80       4.653 -25.311  -2.055 50.00  0.00           C
ATOM    604  CB  VAL    80       8.287 -38.545  -0.069  1.00  0.00           C
ATOM    605  CG1 VAL    80       7.860 -38.369  -1.528  1.00  0.00           C
ATOM    606  CG2 VAL    80       9.816 -38.512  -0.093  1.00  0.00           C
ATOM    607  C   VAL    80       6.139 -37.508   0.760  1.00  0.00           C
ATOM    608  O   VAL    80       5.543 -38.408   0.155  1.00  0.00           O
ATOM    609  N   GLY    81       5.525 -36.466   1.309  1.00  0.00           N
ATOM    610  CA  GLY    81       4.448 -24.223  -0.908 50.00  0.00           C
ATOM    611  C   GLY    81       3.417 -36.238   0.018  1.00  0.00           C
ATOM    612  O   GLY    81       2.379 -36.854  -0.239  1.00  0.00           O
ATOM    613  N   ALA    82       4.012 -35.449  -0.866  1.00  0.00           N
ATOM    614  CA  ALA    82       3.350 -23.426  -0.985 50.00  0.00           C
ATOM    615  CB  ALA    82       4.374 -34.404  -3.062  1.00  0.00           C
ATOM    616  C   ALA    82       3.239 -36.624  -2.867  1.00  0.00           C
ATOM    617  O   ALA    82       2.115 -36.980  -3.268  1.00  0.00           O
ATOM    618  N   VAL    83       4.313 -37.399  -2.942  1.00  0.00           N
ATOM    619  CA  VAL    83       3.145 -26.931  -1.640 50.00  0.00           C
ATOM    620  CB  VAL    83       5.643 -39.247  -3.751  1.00  0.00           C
ATOM    621  CG1 VAL    83       5.660 -40.687  -4.268  1.00  0.00           C
ATOM    622  CG2 VAL    83       6.525 -38.462  -4.724  1.00  0.00           C
ATOM    623  C   VAL    83       3.326 -39.669  -2.911  1.00  0.00           C
ATOM    624  O   VAL    83       2.678 -40.478  -3.583  1.00  0.00           O
ATOM    625  N   GLU    84       3.233 -39.607  -1.583  1.00  0.00           N
ATOM    626  CA  GLU    84       3.806 -28.501  -0.945 50.00  0.00           C
ATOM    627  CB  GLU    84       2.382 -40.352   0.635  1.00  0.00           C
ATOM    628  CG  GLU    84       1.430 -41.289   1.381  1.00  0.00           C
ATOM    629  CD  GLU    84       1.649 -41.091   2.874  1.00  0.00           C
ATOM    630  OE1 GLU    84       2.524 -40.261   3.239  1.00  0.00           O
ATOM    631  OE2 GLU    84       0.945 -41.768   3.670  1.00  0.00           O
ATOM    632  C   GLU    84       0.856 -40.148  -1.266  1.00  0.00           C
ATOM    633  O   GLU    84       0.088 -41.038  -1.647  1.00  0.00           O
ATOM    634  N   LYS    85       0.498 -38.872  -1.154  1.00  0.00           N
ATOM    635  CA  LYS    85       3.269 -26.548   0.051 50.00  0.00           C
ATOM    636  CB  LYS    85      -0.903 -36.884  -1.415  1.00  0.00           C
ATOM    637  CG  LYS    85      -2.276 -36.309  -1.767  1.00  0.00           C
ATOM    638  CD  LYS    85      -2.373 -34.794  -1.577  1.00  0.00           C
ATOM    639  CE  LYS    85      -3.731 -34.211  -1.975  1.00  0.00           C
ATOM    640  NZ  LYS    85      -3.743 -32.750  -1.739  1.00  0.00           N
ATOM    641  C   LYS    85      -1.188 -38.830  -2.961  1.00  0.00           C
ATOM    642  O   LYS    85      -2.292 -39.331  -3.201  1.00  0.00           O
ATOM    643  N   LYS    86      -0.249 -38.645  -3.887  1.00  0.00           N
ATOM    644  CA  LYS    86       2.278 -27.574  -0.489 50.00  0.00           C
ATOM    645  CB  LYS    86       0.763 -38.740  -6.101  1.00  0.00           C
ATOM    646  CG  LYS    86       0.897 -37.238  -6.362  1.00  0.00           C
ATOM    647  CD  LYS    86       2.114 -36.871  -7.213  1.00  0.00           C
ATOM    648  CE  LYS    86       2.262 -35.367  -7.454  1.00  0.00           C
ATOM    649  NZ  LYS    86       3.480 -35.099  -8.251  1.00  0.00           N
ATOM    650  C   LYS    86      -0.710 -40.612  -5.334  1.00  0.00           C
ATOM    651  O   LYS    86      -1.629 -41.058  -6.034  1.00  0.00           O
ATOM    652  N   GLU    87       0.098 -41.384  -4.607  1.00  0.00           N
ATOM    653  CA  GLU    87       2.647 -31.198  -0.849 50.00  0.00           C
ATOM    654  CB  GLU    87       0.973 -43.407  -3.570  1.00  0.00           C
ATOM    655  CG  GLU    87       0.953 -44.935  -3.502  1.00  0.00           C
ATOM    656  CD  GLU    87       2.006 -45.375  -2.494  1.00  0.00           C
ATOM    657  OE1 GLU    87       2.689 -44.482  -1.925  1.00  0.00           O
ATOM    658  OE2 GLU    87       2.141 -46.609  -2.280  1.00  0.00           O
ATOM    659  C   GLU    87      -1.433 -43.197  -4.161  1.00  0.00           C
ATOM    660  O   GLU    87      -2.100 -43.995  -4.823  1.00  0.00           O
ATOM    661  N   ASP    88      -1.885 -42.600  -3.061  1.00  0.00           N
ATOM    662  CA  ASP    88       2.990 -31.233   0.179 50.00  0.00           C
ATOM    663  CB  ASP    88      -3.150 -42.566  -0.960  1.00  0.00           C
ATOM    664  CG  ASP    88      -2.169 -43.442  -0.194  1.00  0.00           C
ATOM    665  OD1 ASP    88      -1.741 -44.484  -0.758  1.00  0.00           O
ATOM    666  OD2 ASP    88      -1.835 -43.081   0.966  1.00  0.00           O
ATOM    667  C   ASP    88      -4.345 -42.476  -3.150  1.00  0.00           C
ATOM    668  O   ASP    88      -5.450 -42.963  -2.960  1.00  0.00           O
ATOM    669  N   ASP    89      -4.158 -41.488  -3.977  1.00  0.00           N
ATOM    670  CA  ASP    89       2.096 -29.547   0.733 50.00  0.00           C
ATOM    671  CB  ASP    89      -4.838 -39.692  -5.384  1.00  0.00           C
ATOM    672  CG  ASP    89      -5.785 -38.637  -4.998  1.00  0.00           C
ATOM    673  OD1 ASP    89      -6.868 -38.601  -5.587  1.00  0.00           O
ATOM    674  OD2 ASP    89      -5.472 -37.857  -4.086  1.00  0.00           O
ATOM    675  C   ASP    89      -5.422 -41.966  -5.951  1.00  0.00           C
ATOM    676  O   ASP    89      -6.444 -41.976  -6.573  1.00  0.00           O
ATOM    677  N   ARG    90      -4.407 -42.781  -6.220  1.00  0.00           N
ATOM    678  CA  ARG    90       1.515 -32.439   0.095 50.00  0.00           C
ATOM    679  CB  ARG    90      -3.092 -43.943  -7.938  1.00  0.00           C
ATOM    680  CG  ARG    90      -2.576 -42.696  -8.659  1.00  0.00           C
ATOM    681  CD  ARG    90      -1.204 -42.888  -9.308  1.00  0.00           C
ATOM    682  NE  ARG    90      -0.840 -41.606  -9.975  1.00  0.00           N
ATOM    683  CZ  ARG    90       0.397 -41.452 -10.529  1.00  0.00           C
ATOM    684  NH1 ARG    90       1.089 -42.612 -10.333  1.00  0.00           N
ATOM    685  NH2 ARG    90       0.460 -40.198 -11.064  1.00  0.00           N
ATOM    686  C   ARG    90      -5.011 -45.092  -6.811  1.00  0.00           C
ATOM    687  O   ARG    90      -5.457 -45.922  -7.603  1.00  0.00           O
ATOM    688  N   ALA    91      -4.697 -45.429  -5.565  1.00  0.00           N
ATOM    689  CA  ALA    91       2.205 -35.162   0.249 50.00  0.00           C
ATOM    690  CB  ALA    91      -4.080 -47.781  -5.079  1.00  0.00           C
ATOM    691  C   ALA    91      -5.500 -46.502  -3.513  1.00  0.00           C
ATOM    692  O   ALA    91      -5.217 -45.469  -2.900  1.00  0.00           O
ATOM    693  N   ALA    92      -6.135 -47.512  -2.930  1.00  0.00           N
ATOM    694  CA  ALA    92       3.103 -35.559   0.924 50.00  0.00           C
ATOM    695  CB  ALA    92      -8.005 -47.666  -1.305  1.00  0.00           C
ATOM    696  C   ALA    92      -5.751 -48.546  -0.763  1.00  0.00           C
ATOM    697  O   ALA    92      -5.692 -49.709  -1.173  1.00  0.00           O
ATOM    698  N   LYS    93      -5.156 -48.168   0.369  1.00  0.00           N
ATOM    699  CA  LYS    93       4.114 -36.999   0.468 50.00  0.00           C
ATOM    700  CB  LYS    93      -2.899 -49.176   1.244  1.00  0.00           C
ATOM    701  CG  LYS    93      -2.237 -47.888   1.738  1.00  0.00           C
ATOM    702  CD  LYS    93      -0.708 -47.937   1.711  1.00  0.00           C
ATOM    703  CE  LYS    93      -0.045 -46.775   2.454  1.00  0.00           C
ATOM    704  NZ  LYS    93      -0.237 -45.516   1.699  1.00  0.00           N
ATOM    705  C   LYS    93      -4.726 -48.719   2.670  1.00  0.00           C
ATOM    706  O   LYS    93      -4.967 -47.551   2.990  1.00  0.00           O
ATOM    707  N   LYS    94      -4.713 -49.699   3.569  1.00  0.00           N
ATOM    708  CA  LYS    94       5.154 -37.079   0.957 50.00  0.00           C
ATOM    709  CB  LYS    94      -5.533 -50.580   5.873  1.00  0.00           C
ATOM    710  CG  LYS    94      -6.899 -51.124   5.451  1.00  0.00           C
ATOM    711  CD  LYS    94      -7.459 -52.181   6.406  1.00  0.00           C
ATOM    712  CE  LYS    94      -8.828 -52.721   5.986  1.00  0.00           C
ATOM    713  NZ  LYS    94      -9.309 -53.707   6.980  1.00  0.00           N
ATOM    714  C   LYS    94      -3.828 -49.008   5.811  1.00  0.00           C
ATOM    715  O   LYS    94      -2.692 -48.910   5.340  1.00  0.00           O
ATOM    716  N   GLU    95      -4.096 -48.734   7.085  1.00  0.00           N
ATOM    717  CA  GLU    95       6.046 -35.547   0.807 50.00  0.00           C
ATOM    718  CB  GLU    95      -2.980 -46.774   8.292  1.00  0.00           C
ATOM    719  CG  GLU    95      -4.260 -46.184   8.887  1.00  0.00           C
ATOM    720  CD  GLU    95      -4.038 -44.692   9.093  1.00  0.00           C
ATOM    721  OE1 GLU    95      -3.842 -43.977   8.075  1.00  0.00           O
ATOM    722  OE2 GLU    95      -4.062 -44.248  10.272  1.00  0.00           O
ATOM    723  C   GLU    95      -3.182 -48.822   9.437  1.00  0.00           C
ATOM    724  O   GLU    95      -4.259 -49.135   9.951  1.00  0.00           O
ATOM    725  N   ARG    96      -2.019 -48.930  10.077  1.00  0.00           N
ATOM    726  CA  ARG    96       7.226 -36.275   0.736 50.00  0.00           C
ATOM    727  CB  ARG    96      -1.968 -50.787  11.929  1.00  0.00           C
ATOM    728  CG  ARG    96      -1.968 -50.951  13.450  1.00  0.00           C
ATOM    729  CD  ARG    96      -1.940 -52.410  13.909  1.00  0.00           C
ATOM    730  NE  ARG    96      -0.603 -52.964  13.557  1.00  0.00           N
ATOM    731  CZ  ARG    96      -0.411 -54.315  13.526  1.00  0.00           C
ATOM    732  NH1 ARG    96      -1.594 -54.911  13.854  1.00  0.00           N
ATOM    733  NH2 ARG    96       0.890 -54.543  13.178  1.00  0.00           N
ATOM    734  C   ARG    96      -0.456 -48.990  11.911  1.00  0.00           C
ATOM    735  O   ARG    96       0.451 -49.826  11.891  1.00  0.00           O
ATOM    736  N   LEU    97      -0.256 -47.753  12.430  1.00  0.00           N
ATOM    737  CA  LEU    97       7.785 -34.073   0.160 50.00  0.00           C
ATOM    738  CB  LEU    97       1.861 -48.281  13.527  1.00  0.00           C
ATOM    739  CG  LEU    97       1.224 -48.642  14.871  1.00  0.00           C
ATOM    740  CD1 LEU    97       1.939 -49.745  15.649  1.00  0.00           C
ATOM    741  CD2 LEU    97       1.144 -47.494  15.876  1.00  0.00           C
ATOM    742  C   LEU    97       1.865 -46.851  11.499  1.00  0.00           C
ATOM    743  O   LEU    97       2.769 -46.027  11.592  1.00  0.00           O
ATOM    744  N   GLU    98       1.520 -47.430  10.351  1.00  0.00           N
ATOM    745  CA  GLU    98       6.800 -34.242  -0.593 50.00  0.00           C
ATOM    746  CB  GLU    98       1.769 -48.170   8.045  1.00  0.00           C
ATOM    747  CG  GLU    98       2.192 -49.606   8.362  1.00  0.00           C
ATOM    748  CD  GLU    98       3.711 -49.639   8.455  1.00  0.00           C
ATOM    749  OE1 GLU    98       4.373 -49.216   7.469  1.00  0.00           O
ATOM    750  OE2 GLU    98       4.230 -50.087   9.512  1.00  0.00           O
ATOM    751  C   GLU    98       1.949 -45.784   8.550  1.00  0.00           C
ATOM    752  O   GLU    98       2.622 -45.358   7.606  1.00  0.00           O
ATOM    753  N   TYR    99       0.973 -45.084   9.129  1.00  0.00           N
ATOM    754  CA  TYR    99       6.400 -31.062   0.164 50.00  0.00           C
ATOM    755  CB  TYR    99      -1.028 -43.956   8.323  1.00  0.00           C
ATOM    756  CG  TYR    99      -1.399 -45.110   7.434  1.00  0.00           C
ATOM    757  CD1 TYR    99      -1.373 -44.985   6.054  1.00  0.00           C
ATOM    758  CD2 TYR    99      -1.755 -46.331   7.974  1.00  0.00           C
ATOM    759  CE1 TYR    99      -1.703 -46.035   5.240  1.00  0.00           C
ATOM    760  CE2 TYR    99      -2.089 -47.405   7.165  1.00  0.00           C
ATOM    761  CZ  TYR    99      -2.061 -47.259   5.794  1.00  0.00           C
ATOM    762  OH  TYR    99      -2.393 -48.342   4.987  1.00  0.00           O
ATOM    763  C   TYR    99       0.862 -42.579   9.278  1.00  0.00           C
ATOM    764  O   TYR    99       0.879 -41.474   8.707  1.00  0.00           O
ATOM    765  N   CYS   100       1.204 -42.766  10.548  1.00  0.00           N
ATOM    766  CA  CYS   100       7.246 -28.658   0.300 50.00  0.00           C
ATOM    767  CB  CYS   100       2.186 -42.265  12.747  1.00  0.00           C
ATOM    768  SG  CYS   100       2.573 -41.251  13.662  1.00  0.00           S
ATOM    769  C   CYS   100       2.546 -40.684  10.822  1.00  0.00           C
ATOM    770  O   CYS   100       2.189 -39.510  10.661  1.00  0.00           O
ATOM    771  N   TRP   101       3.736 -41.145  10.442  1.00  0.00           N
ATOM    772  CA  TRP   101       7.017 -27.367  -0.603 50.00  0.00           C
ATOM    773  CB  TRP   101       6.009 -41.072   9.352  1.00  0.00           C
ATOM    774  CG  TRP   101       6.926 -41.461  10.487  1.00  0.00           C
ATOM    775  CD2 TRP   101       7.779 -40.591  11.244  1.00  0.00           C
ATOM    776  CD1 TRP   101       7.147 -42.689  11.040  1.00  0.00           C
ATOM    777  NE1 TRP   101       8.020 -42.683  12.028  1.00  0.00           N
ATOM    778  CE2 TRP   101       8.452 -41.394  12.202  1.00  0.00           C
ATOM    779  CE3 TRP   101       8.046 -39.206  11.208  1.00  0.00           C
ATOM    780  CZ2 TRP   101       9.385 -40.854  13.129  1.00  0.00           C
ATOM    781  CZ3 TRP   101       8.982 -38.657  12.136  1.00  0.00           C
ATOM    782  CH2 TRP   101       9.634 -39.489  13.080  1.00  0.00           C
ATOM    783  C   TRP   101       4.149 -39.461   8.636  1.00  0.00           C
ATOM    784  O   TRP   101       4.354 -38.241   8.565  1.00  0.00           O
ATOM    785  N   GLY   102       3.387 -40.113   7.755  1.00  0.00           N
ATOM    786  CA  GLY   102       5.928 -26.761  -0.297 50.00  0.00           C
ATOM    787  C   GLY   102       1.885 -38.301   7.159  1.00  0.00           C
ATOM    788  O   GLY   102       2.008 -37.127   6.790  1.00  0.00           O
ATOM    789  N   ARG   103       0.991 -38.692   8.063  1.00  0.00           N
ATOM    790  CA  ARG   103       6.189 -24.978   0.801 50.00  0.00           C
ATOM    791  CB  ARG   103      -0.795 -38.405   9.735  1.00  0.00           C
ATOM    792  CG  ARG   103      -1.782 -37.444  10.400  1.00  0.00           C
ATOM    793  CD  ARG   103      -2.648 -38.102  11.476  1.00  0.00           C
ATOM    794  NE  ARG   103      -1.740 -38.561  12.564  1.00  0.00           N
ATOM    795  CZ  ARG   103      -1.437 -39.886  12.687  1.00  0.00           C
ATOM    796  NH1 ARG   103      -2.093 -40.557  11.696  1.00  0.00           N
ATOM    797  NH2 ARG   103      -0.595 -40.023  13.753  1.00  0.00           N
ATOM    798  C   ARG   103       0.918 -36.603   9.251  1.00  0.00           C
ATOM    799  O   ARG   103       0.654 -35.439   8.967  1.00  0.00           O
ATOM    800  N   ALA   104       1.974 -36.948   9.979  1.00  0.00           N
ATOM    801  CA  ALA   104       6.914 -22.986   0.364 50.00  0.00           C
ATOM    802  CB  ALA   104       3.782 -36.592  11.587  1.00  0.00           C
ATOM    803  C   ALA   104       3.419 -35.036   9.618  1.00  0.00           C
ATOM    804  O   ALA   104       3.424 -33.815   9.751  1.00  0.00           O
ATOM    805  N   LEU   105       3.909 -35.646   8.546  1.00  0.00           N
ATOM    806  CA  LEU   105       6.150 -22.214  -0.370 50.00  0.00           C
ATOM    807  CB  LEU   105       5.183 -35.847   6.479  1.00  0.00           C
ATOM    808  CG  LEU   105       6.340 -36.692   7.015  1.00  0.00           C
ATOM    809  CD1 LEU   105       6.850 -37.769   6.059  1.00  0.00           C
ATOM    810  CD2 LEU   105       7.602 -35.908   7.371  1.00  0.00           C
ATOM    811  C   LEU   105       3.644 -33.944   6.845  1.00  0.00           C
ATOM    812  O   LEU   105       4.052 -32.861   6.413  1.00  0.00           O
ATOM    813  N   LEU   106       2.369 -34.327   6.771  1.00  0.00           N
ATOM    814  CA  LEU   106       5.434 -20.945   0.535 50.00  0.00           C
ATOM    815  CB  LEU   106       0.111 -34.203   5.786  1.00  0.00           C
ATOM    816  CG  LEU   106       0.187 -35.042   4.497  1.00  0.00           C
ATOM    817  CD1 LEU   106      -0.867 -36.173   4.438  1.00  0.00           C
ATOM    818  CD2 LEU   106       0.069 -34.125   3.287  1.00  0.00           C
ATOM    819  C   LEU   106       1.150 -32.285   7.074  1.00  0.00           C
ATOM    820  O   LEU   106       1.119 -31.116   6.647  1.00  0.00           O
ATOM    821  N   SER   107       1.066 -32.600   8.360  1.00  0.00           N
ATOM    822  CA  SER   107       6.229 -18.823   1.167 50.00  0.00           C
ATOM    823  CB  SER   107       0.676 -32.201  10.736  1.00  0.00           C
ATOM    824  OG  SER   107       0.683 -31.191  11.725  1.00  0.00           O
ATOM    825  C   SER   107       1.812 -30.497   9.338  1.00  0.00           C
ATOM    826  O   SER   107       1.488 -29.309   9.432  1.00  0.00           O
ATOM    827  N   ALA   108       3.076 -30.889   9.205  1.00  0.00           N
ATOM    828  CA  ALA   108       6.453 -17.102   0.362 50.00  0.00           C
ATOM    829  CB  ALA   108       5.451 -30.557   9.789  1.00  0.00           C
ATOM    830  C   ALA   108       4.297 -29.125   8.055  1.00  0.00           C
ATOM    831  O   ALA   108       4.880 -28.037   8.065  1.00  0.00           O
ATOM    832  N   ARG   109       3.724 -29.631   6.962  1.00  0.00           N
ATOM    833  CA  ARG   109       5.329 -16.410   0.174 50.00  0.00           C
ATOM    834  CB  ARG   109       3.908 -29.828   4.535  1.00  0.00           C
ATOM    835  CG  ARG   109       5.199 -30.649   4.544  1.00  0.00           C
ATOM    836  CD  ARG   109       5.298 -31.648   3.390  1.00  0.00           C
ATOM    837  NE  ARG   109       5.376 -30.864   2.125  1.00  0.00           N
ATOM    838  CZ  ARG   109       5.460 -31.510   0.925  1.00  0.00           C
ATOM    839  NH1 ARG   109       5.453 -32.853   1.175  1.00  0.00           N
ATOM    840  NH2 ARG   109       5.518 -30.562  -0.055  1.00  0.00           N
ATOM    841  C   ARG   109       2.597 -27.940   5.512  1.00  0.00           C
ATOM    842  O   ARG   109       2.482 -27.268   4.478  1.00  0.00           O
ATOM    843  N   HIS   110       1.714 -27.881   6.506  1.00  0.00           N
ATOM    844  CA  HIS   110       5.194 -14.517   1.272 50.00  0.00           C
ATOM    845  ND1 HIS   110       1.851 -24.601   8.076  1.00  0.00           N
ATOM    846  CG  HIS   110       2.039 -24.970   6.760  1.00  0.00           C
ATOM    847  CB  HIS   110       0.985 -25.620   5.913  1.00  0.00           C
ATOM    848  NE2 HIS   110       3.846 -24.060   7.610  1.00  0.00           N
ATOM    849  CD2 HIS   110       3.315 -24.620   6.474  1.00  0.00           C
ATOM    850  CE1 HIS   110       2.931 -24.067   8.560  1.00  0.00           C
ATOM    851  C   HIS   110      -0.509 -27.580   5.546  1.00  0.00           C
ATOM    852  O   HIS   110      -1.510 -26.923   5.249  1.00  0.00           O
ATOM    853  N   LEU   111      -0.294 -28.826   5.124  1.00  0.00           N
ATOM    854  CA  LEU   111       4.548 -17.248   1.388 50.00  0.00           C
ATOM    855  CB  LEU   111      -0.545 -30.437   3.297  1.00  0.00           C
ATOM    856  CG  LEU   111       0.386 -29.676   2.351  1.00  0.00           C
ATOM    857  CD1 LEU   111       1.141 -30.548   1.348  1.00  0.00           C
ATOM    858  CD2 LEU   111      -0.300 -28.635   1.469  1.00  0.00           C
ATOM    859  C   LEU   111      -2.233 -30.308   5.201  1.00  0.00           C
ATOM    860  O   LEU   111      -2.194 -31.536   5.301  1.00  0.00           O
ATOM    861  N   ALA   112      -3.107 -29.538   5.845  1.00  0.00           N
ATOM    862  CA  ALA   112       5.041 -17.553   2.425 50.00  0.00           C
ATOM    863  CB  ALA   112      -4.539 -28.816   7.657  1.00  0.00           C
ATOM    864  C   ALA   112      -5.266 -30.761   6.229  1.00  0.00           C
ATOM    865  O   ALA   112      -5.748 -31.768   6.770  1.00  0.00           O
ATOM    866  N   GLY   113      -5.770 -30.264   5.112  1.00  0.00           N
ATOM    867  CA  GLY   113       4.171 -18.670   2.949 50.00  0.00           C
ATOM    868  C   GLY   113      -6.576 -32.276   4.047  1.00  0.00           C
ATOM    869  O   GLY   113      -7.369 -33.196   4.193  1.00  0.00           O
ATOM    870  N   ASP   114      -5.369 -32.445   3.502  1.00  0.00           N
ATOM    871  CA  ASP   114       4.040 -21.619   1.981 50.00  0.00           C
ATOM    872  CB  ASP   114      -3.596 -33.759   2.295  1.00  0.00           C
ATOM    873  CG  ASP   114      -3.914 -33.228   0.905  1.00  0.00           C
ATOM    874  OD1 ASP   114      -5.124 -33.152   0.563  1.00  0.00           O
ATOM    875  OD2 ASP   114      -2.950 -32.891   0.166  1.00  0.00           O
ATOM    876  C   ASP   114      -4.698 -34.689   4.309  1.00  0.00           C
ATOM    877  O   ASP   114      -5.041 -35.863   4.254  1.00  0.00           O
ATOM    878  N   ASP   115      -4.126 -34.145   5.385  1.00  0.00           N
ATOM    879  CA  ASP   115       5.165 -22.416   1.974 50.00  0.00           C
ATOM    880  CB  ASP   115      -3.436 -34.006   7.700  1.00  0.00           C
ATOM    881  CG  ASP   115      -3.196 -34.859   8.938  1.00  0.00           C
ATOM    882  OD1 ASP   115      -2.375 -35.810   8.850  1.00  0.00           O
ATOM    883  OD2 ASP   115      -3.831 -34.570   9.987  1.00  0.00           O
ATOM    884  C   ASP   115      -5.302 -35.502   7.046  1.00  0.00           C
ATOM    885  O   ASP   115      -5.407 -36.696   7.333  1.00  0.00           O
ATOM    886  N   ASP   116      -6.333 -34.668   7.042  1.00  0.00           N
ATOM    887  CA  ASP   116       5.077 -22.626   3.097 50.00  0.00           C
ATOM    888  CB  ASP   116      -8.554 -33.925   7.790  1.00  0.00           C
ATOM    889  CG  ASP   116      -8.083 -33.367   9.125  1.00  0.00           C
ATOM    890  OD1 ASP   116      -7.284 -34.062   9.808  1.00  0.00           O
ATOM    891  OD2 ASP   116      -8.515 -32.238   9.481  1.00  0.00           O
ATOM    892  C   ASP   116      -8.358 -36.003   6.408  1.00  0.00           C
ATOM    893  O   ASP   116      -9.244 -36.744   6.778  1.00  0.00           O
ATOM    894  N   GLU   117      -7.991 -35.916   5.132  1.00  0.00           N
ATOM    895  CA  GLU   117       4.126 -24.650   2.860 50.00  0.00           C
ATOM    896  CB  GLU   117      -8.188 -36.399   2.720  1.00  0.00           C
ATOM    897  CG  GLU   117      -8.973 -35.228   2.220  1.00  0.00           C
ATOM    898  CD  GLU   117      -8.257 -34.526   1.077  1.00  0.00           C
ATOM    899  OE1 GLU   117      -7.292 -35.140   0.560  1.00  0.00           O
ATOM    900  OE2 GLU   117      -8.640 -33.378   0.712  1.00  0.00           O
ATOM    901  C   GLU   117      -7.881 -38.241   4.318  1.00  0.00           C
ATOM    902  O   GLU   117      -8.552 -39.265   4.165  1.00  0.00           O
ATOM    903  N   HIS   118      -6.594 -38.279   4.634  1.00  0.00           N
ATOM    904  CA  HIS   118       4.674 -27.212   1.992 50.00  0.00           C
ATOM    905  ND1 HIS   118      -3.688 -40.061   2.769  1.00  0.00           N
ATOM    906  CG  HIS   118      -3.737 -39.149   3.803  1.00  0.00           C
ATOM    907  CB  HIS   118      -4.449 -39.389   5.101  1.00  0.00           C
ATOM    908  NE2 HIS   118      -2.592 -38.377   2.096  1.00  0.00           N
ATOM    909  CD2 HIS   118      -3.034 -38.079   3.362  1.00  0.00           C
ATOM    910  CE1 HIS   118      -3.009 -39.585   1.770  1.00  0.00           C
ATOM    911  C   HIS   118      -6.585 -40.152   6.184  1.00  0.00           C
ATOM    912  O   HIS   118      -6.932 -41.335   6.214  1.00  0.00           O
ATOM    913  N   ILE   119      -6.747 -39.337   7.214  1.00  0.00           N
ATOM    914  CA  ILE   119       5.557 -27.215   2.719 50.00  0.00           C
ATOM    915  CB  ILE   119      -7.356 -38.732   9.481  1.00  0.00           C
ATOM    916  CG2 ILE   119      -8.173 -39.081  10.736  1.00  0.00           C
ATOM    917  CG1 ILE   119      -5.943 -38.414   9.997  1.00  0.00           C
ATOM    918  CD1 ILE   119      -5.877 -37.152  10.856  1.00  0.00           C
ATOM    919  C   ILE   119      -8.822 -40.315   8.170  1.00  0.00           C
ATOM    920  O   ILE   119      -9.223 -41.371   8.675  1.00  0.00           O
ATOM    921  N   ASN   120      -9.589 -39.550   7.392  1.00  0.00           N
ATOM    922  CA  ASN   120       4.839 -27.472   3.604 50.00  0.00           C
ATOM    923  CB  ASN   120     -11.602 -38.853   6.192  1.00  0.00           C
ATOM    924  CG  ASN   120     -13.066 -39.219   5.996  1.00  0.00           C
ATOM    925  OD1 ASN   120     -13.838 -39.276   6.952  1.00  0.00           O
ATOM    926  ND2 ASN   120     -13.528 -39.487   4.745  1.00  0.00           N
ATOM    927  C   ASN   120     -10.987 -41.284   6.323  1.00  0.00           C
ATOM    928  O   ASN   120     -11.865 -42.135   6.562  1.00  0.00           O
ATOM    929  N   SER   121     -10.010 -41.458   5.431  1.00  0.00           N
ATOM    930  CA  SER   121       4.356 -30.366   2.854 50.00  0.00           C
ATOM    931  CB  SER   121      -8.726 -42.458   3.607  1.00  0.00           C
ATOM    932  OG  SER   121      -8.557 -43.632   2.826  1.00  0.00           O
ATOM    933  C   SER   121      -9.583 -43.867   5.499  1.00  0.00           C
ATOM    934  O   SER   121     -10.196 -44.911   5.327  1.00  0.00           O
ATOM    935  N   ILE   122      -8.653 -43.722   6.435  1.00  0.00           N
ATOM    936  CA  ILE   122       5.354 -32.193   2.558 50.00  0.00           C
ATOM    937  CB  ILE   122      -7.349 -44.362   8.410  1.00  0.00           C
ATOM    938  CG2 ILE   122      -7.219 -45.322   9.605  1.00  0.00           C
ATOM    939  CG1 ILE   122      -5.952 -44.328   7.768  1.00  0.00           C
ATOM    940  CD1 ILE   122      -4.888 -43.687   8.657  1.00  0.00           C
ATOM    941  C   ILE   122      -9.747 -45.066   8.185  1.00  0.00           C
ATOM    942  O   ILE   122     -10.068 -46.224   8.371  1.00  0.00           O
ATOM    943  N   VAL   123     -10.495 -44.041   8.592  1.00  0.00           N
ATOM    944  CA  VAL   123       5.564 -31.584   3.672 50.00  0.00           C
ATOM    945  CB  VAL   123     -12.388 -42.899   9.649  1.00  0.00           C
ATOM    946  CG1 VAL   123     -13.803 -43.014  10.219  1.00  0.00           C
ATOM    947  CG2 VAL   123     -11.599 -42.106  10.693  1.00  0.00           C
ATOM    948  C   VAL   123     -12.779 -45.071   8.523  1.00  0.00           C
ATOM    949  O   VAL   123     -13.448 -45.950   9.074  1.00  0.00           O
ATOM    950  N   GLU   124     -12.881 -44.787   7.232  1.00  0.00           N
ATOM    951  CA  GLU   124       4.599 -33.155   3.839 50.00  0.00           C
ATOM    952  CB  GLU   124     -13.517 -44.971   4.925  1.00  0.00           C
ATOM    953  CG  GLU   124     -14.078 -43.562   4.724  1.00  0.00           C
ATOM    954  CD  GLU   124     -13.701 -43.101   3.323  1.00  0.00           C
ATOM    955  OE1 GLU   124     -12.963 -43.852   2.632  1.00  0.00           O
ATOM    956  OE2 GLU   124     -14.147 -41.991   2.926  1.00  0.00           O
ATOM    957  C   GLU   124     -13.495 -47.040   6.303  1.00  0.00           C
ATOM    958  O   GLU   124     -14.308 -47.812   5.797  1.00  0.00           O
ATOM    959  N   SER   125     -12.361 -47.460   6.850  1.00  0.00           N
ATOM    960  CA  SER   125       5.034 -36.398   3.145 50.00  0.00           C
ATOM    961  CB  SER   125     -10.551 -49.009   6.324  1.00  0.00           C
ATOM    962  OG  SER   125     -10.501 -48.593   4.967  1.00  0.00           O
ATOM    963  C   SER   125     -12.010 -49.379   8.280  1.00  0.00           C
ATOM    964  O   SER   125     -11.481 -50.454   8.565  1.00  0.00           O
ATOM    965  N   PHE   126     -10.847 -49.744   8.806  1.00  0.00           N
ATOM    966  CA  PHE   126       6.109 -37.429   2.985 50.00  0.00           C
ATOM    967  CB  PHE   126     -10.795 -51.738  10.520  1.00  0.00           C
ATOM    968  CG  PHE   126     -10.805 -51.853  12.006  1.00  0.00           C
ATOM    969  CD1 PHE   126     -11.977 -51.640  12.766  1.00  0.00           C
ATOM    970  CD2 PHE   126      -9.621 -52.183  12.691  1.00  0.00           C
ATOM    971  CE1 PHE   126     -11.977 -51.756  14.186  1.00  0.00           C
ATOM    972  CE2 PHE   126      -9.594 -52.306  14.113  1.00  0.00           C
ATOM    973  CZ  PHE   126     -10.779 -52.089  14.863  1.00  0.00           C
ATOM    974  C   PHE   126      -9.371 -49.881  10.687  1.00  0.00           C
ATOM    975  O   PHE   126      -8.345 -50.294  10.141  1.00  0.00           O
ATOM    976  N   GLU   127      -9.328 -49.094  11.756  1.00  0.00           N
ATOM    977  CA  GLU   127       6.867 -35.619   2.541 50.00  0.00           C
ATOM    978  CB  GLU   127      -7.606 -47.211  12.041  1.00  0.00           C
ATOM    979  CG  GLU   127      -8.549 -46.156  12.623  1.00  0.00           C
ATOM    980  CD  GLU   127      -8.026 -44.784  12.221  1.00  0.00           C
ATOM    981  OE1 GLU   127      -6.861 -44.466  12.583  1.00  0.00           O
ATOM    982  OE2 GLU   127      -8.783 -44.036  11.547  1.00  0.00           O
ATOM    983  C   GLU   127      -8.076 -48.706  13.829  1.00  0.00           C
ATOM    984  O   GLU   127      -9.128 -48.653  14.471  1.00  0.00           O
ATOM    985  N   GLN   128      -6.895 -48.815  14.428  1.00  0.00           N
ATOM    986  CA  GLN   128       8.027 -35.532   2.495 50.00  0.00           C
ATOM    987  CB  GLN   128      -5.579 -49.477  16.610  1.00  0.00           C
ATOM    988  CG  GLN   128      -5.467 -50.991  16.417  1.00  0.00           C
ATOM    989  CD  GLN   128      -4.198 -51.461  17.113  1.00  0.00           C
ATOM    990  OE1 GLN   128      -3.491 -50.674  17.741  1.00  0.00           O
ATOM    991  NE2 GLN   128      -3.840 -52.771  17.040  1.00  0.00           N
ATOM    992  C   GLN   128      -6.685 -47.443  16.453  1.00  0.00           C
ATOM    993  O   GLN   128      -7.057 -46.457  15.812  1.00  0.00           O
ATOM    994  N   GLN   129      -6.181 -47.336  17.678  1.00  0.00           N
ATOM    995  CA  GLN   129       8.648 -32.475   2.782 50.00  0.00           C
ATOM    996  CB  GLN   129      -6.735 -45.641  19.614  1.00  0.00           C
ATOM    997  CG  GLN   129      -8.255 -45.568  19.452  1.00  0.00           C
ATOM    998  CD  GLN   129      -8.573 -44.399  18.530  1.00  0.00           C
ATOM    999  OE1 GLN   129      -7.878 -43.385  18.529  1.00  0.00           O
ATOM   1000  NE2 GLN   129      -9.644 -44.478  17.695  1.00  0.00           N
ATOM   1001  C   GLN   129      -4.568 -45.905  18.822  1.00  0.00           C
ATOM   1002  O   GLN   129      -4.012 -46.800  19.465  1.00  0.00           O
ATOM   1003  N   PHE   130      -3.948 -44.770  18.520  1.00  0.00           N
ATOM   1004  CA  PHE   130       9.103 -30.955   1.888 50.00  0.00           C
ATOM   1005  CB  PHE   130      -1.389 -45.224  18.282  1.00  0.00           C
ATOM   1006  CG  PHE   130      -1.391 -44.835  16.844  1.00  0.00           C
ATOM   1007  CD1 PHE   130      -0.699 -43.695  16.375  1.00  0.00           C
ATOM   1008  CD2 PHE   130      -2.095 -45.614  15.908  1.00  0.00           C
ATOM   1009  CE1 PHE   130      -0.703 -43.333  14.997  1.00  0.00           C
ATOM   1010  CE2 PHE   130      -2.115 -45.274  14.521  1.00  0.00           C
ATOM   1011  CZ  PHE   130      -1.417 -44.126  14.065  1.00  0.00           C
ATOM   1012  C   PHE   130      -2.196 -43.088  18.710  1.00  0.00           C
ATOM   1013  O   PHE   130      -3.017 -42.248  18.331  1.00  0.00           O
ATOM   1014  N   GLY   131      -0.924 -42.772  18.931  1.00  0.00           N
ATOM   1015  CA  GLY   131       9.116 -27.819   1.627 50.00  0.00           C
ATOM   1016  C   GLY   131      -1.200 -40.698  17.628  1.00  0.00           C
ATOM   1017  O   GLY   131      -1.221 -39.467  17.536  1.00  0.00           O
ATOM   1018  N   CYS   132      -1.790 -41.395  16.765  1.00  0.00           N
ATOM   1019  CA  CYS   132       8.011 -27.622   2.019 50.00  0.00           C
ATOM   1020  CB  CYS   132      -3.559 -42.185  15.199  1.00  0.00           C
ATOM   1021  SG  CYS   132      -4.421 -41.823  14.130  1.00  0.00           S
ATOM   1022  C   CYS   132      -3.581 -39.887  16.131  1.00  0.00           C
ATOM   1023  O   CYS   132      -3.669 -38.841  15.477  1.00  0.00           O
ATOM   1024  N   VAL   133      -4.233 -40.103  17.269  1.00  0.00           N
ATOM   1025  CA  VAL   133       8.424 -25.550   3.039 50.00  0.00           C
ATOM   1026  CB  VAL   133      -5.388 -39.518  19.355  1.00  0.00           C
ATOM   1027  CG1 VAL   133      -5.971 -38.406  20.230  1.00  0.00           C
ATOM   1028  CG2 VAL   133      -6.462 -40.607  19.323  1.00  0.00           C
ATOM   1029  C   VAL   133      -4.165 -37.750  18.108  1.00  0.00           C
ATOM   1030  O   VAL   133      -4.620 -36.694  17.710  1.00  0.00           O
ATOM   1031  N   SER   134      -2.967 -37.853  18.670  1.00  0.00           N
ATOM   1032  CA  SER   134       8.844 -23.122   2.483 50.00  0.00           C
ATOM   1033  CB  SER   134      -0.804 -37.139  19.555  1.00  0.00           C
ATOM   1034  OG  SER   134      -1.060 -37.621  20.866  1.00  0.00           O
ATOM   1035  C   SER   134      -1.826 -35.897  17.633  1.00  0.00           C
ATOM   1036  O   SER   134      -1.913 -34.660  17.632  1.00  0.00           O
ATOM   1037  N   GLY   135      -1.504 -36.595  16.552  1.00  0.00           N
ATOM   1038  CA  GLY   135       7.876 -22.622   1.948 50.00  0.00           C
ATOM   1039  C   GLY   135      -2.460 -35.217  14.760  1.00  0.00           C
ATOM   1040  O   GLY   135      -2.404 -34.072  14.270  1.00  0.00           O
ATOM   1041  N   TYR   136      -3.591 -35.913  14.848  1.00  0.00           N
ATOM   1042  CA  TYR   136       7.268 -22.221   2.923 50.00  0.00           C
ATOM   1043  CB  TYR   136      -5.971 -36.393  14.341  1.00  0.00           C
ATOM   1044  CG  TYR   136      -7.193 -35.722  13.816  1.00  0.00           C
ATOM   1045  CD1 TYR   136      -7.365 -35.569  12.434  1.00  0.00           C
ATOM   1046  CD2 TYR   136      -8.198 -35.233  14.680  1.00  0.00           C
ATOM   1047  CE1 TYR   136      -8.508 -34.943  11.899  1.00  0.00           C
ATOM   1048  CE2 TYR   136      -9.366 -34.594  14.152  1.00  0.00           C
ATOM   1049  CZ  TYR   136      -9.502 -34.458  12.756  1.00  0.00           C
ATOM   1050  OH  TYR   136     -10.606 -33.846  12.203  1.00  0.00           O
ATOM   1051  C   TYR   136      -5.204 -34.093  15.069  1.00  0.00           C
ATOM   1052  O   TYR   136      -5.554 -33.065  14.466  1.00  0.00           O
ATOM   1053  N   LEU   137      -5.097 -34.182  16.393  1.00  0.00           N
ATOM   1054  CA  LEU   137       7.982 -19.684   3.510 50.00  0.00           C
ATOM   1055  CB  LEU   137      -4.926 -33.462  18.723  1.00  0.00           C
ATOM   1056  CG  LEU   137      -6.061 -34.114  19.486  1.00  0.00           C
ATOM   1057  CD1 LEU   137      -5.575 -34.590  20.827  1.00  0.00           C
ATOM   1058  CD2 LEU   137      -7.169 -33.096  19.630  1.00  0.00           C
ATOM   1059  C   LEU   137      -4.430 -31.882  16.912  1.00  0.00           C
ATOM   1060  O   LEU   137      -4.899 -30.765  16.884  1.00  0.00           O
ATOM   1061  N   ARG   138      -3.159 -32.143  16.638  1.00  0.00           N
ATOM   1062  CA  ARG   138       7.845 -17.810   2.705 50.00  0.00           C
ATOM   1063  CB  ARG   138      -0.873 -31.672  15.968  1.00  0.00           C
ATOM   1064  CG  ARG   138      -0.167 -32.065  17.268  1.00  0.00           C
ATOM   1065  CD  ARG   138       1.177 -32.761  17.048  1.00  0.00           C
ATOM   1066  NE  ARG   138       2.090 -31.778  16.398  1.00  0.00           N
ATOM   1067  CZ  ARG   138       3.285 -32.192  15.884  1.00  0.00           C
ATOM   1068  NH1 ARG   138       3.387 -33.536  16.100  1.00  0.00           N
ATOM   1069  NH2 ARG   138       3.911 -31.101  15.356  1.00  0.00           N
ATOM   1070  C   ARG   138      -2.753 -30.437  14.896  1.00  0.00           C
ATOM   1071  O   ARG   138      -2.679 -29.230  14.740  1.00  0.00           O
ATOM   1072  N   ALA   139      -3.485 -31.184  14.073  1.00  0.00           N
ATOM   1073  CA  ALA   139       6.733 -17.657   2.918 50.00  0.00           C
ATOM   1074  CB  ALA   139      -4.793 -31.678  12.028  1.00  0.00           C
ATOM   1075  C   ALA   139      -4.929 -29.443  13.086  1.00  0.00           C
ATOM   1076  O   ALA   139      -4.861 -28.414  12.409  1.00  0.00           O
ATOM   1077  N   LEU   140      -5.806 -29.568  14.080  1.00  0.00           N
ATOM   1078  CA  LEU   140       6.882 -15.452   4.024 50.00  0.00           C
ATOM   1079  CB  LEU   140      -7.845 -28.692  15.462  1.00  0.00           C
ATOM   1080  CG  LEU   140      -8.937 -29.663  15.008  1.00  0.00           C
ATOM   1081  CD1 LEU   140      -9.981 -30.003  16.070  1.00  0.00           C
ATOM   1082  CD2 LEU   140      -9.776 -29.184  13.825  1.00  0.00           C
ATOM   1083  C   LEU   140      -5.969 -27.350  15.074  1.00  0.00           C
ATOM   1084  O   LEU   140      -6.329 -26.175  14.963  1.00  0.00           O
ATOM   1085  N   ALA   141      -4.888 -27.695  15.768  1.00  0.00           N
ATOM   1086  CA  ALA   141       7.591 -13.467   3.611 50.00  0.00           C
ATOM   1087  CB  ALA   141      -2.964 -27.315  17.293  1.00  0.00           C
ATOM   1088  C   ALA   141      -3.391 -25.834  15.350  1.00  0.00           C
ATOM   1089  O   ALA   141      -3.328 -24.605  15.448  1.00  0.00           O
ATOM   1090  N   ILE   142      -2.887 -26.483  14.308  1.00  0.00           N
ATOM   1091  CA  ILE   142       6.799 -12.747   2.843 50.00  0.00           C
ATOM   1092  CB  ILE   142      -1.832 -26.207  11.804  1.00  0.00           C
ATOM   1093  CG2 ILE   142      -1.279 -25.061  10.941  1.00  0.00           C
ATOM   1094  CG1 ILE   142      -0.732 -27.282  11.797  1.00  0.00           C
ATOM   1095  CD1 ILE   142       0.592 -26.803  12.390  1.00  0.00           C
ATOM   1096  C   ILE   142      -3.206 -24.698  12.687  1.00  0.00           C
ATOM   1097  O   ILE   142      -2.839 -23.536  12.490  1.00  0.00           O
ATOM   1098  N   ASN   143      -4.455 -25.087  12.442  1.00  0.00           N
ATOM   1099  CA  ASN   143       6.076 -11.326   3.746 50.00  0.00           C
ATOM   1100  CB  ASN   143      -6.744 -24.883  11.451  1.00  0.00           C
ATOM   1101  CG  ASN   143      -6.426 -25.625  10.161  1.00  0.00           C
ATOM   1102  OD1 ASN   143      -5.457 -25.310   9.471  1.00  0.00           O
ATOM   1103  ND2 ASN   143      -7.225 -26.651   9.764  1.00  0.00           N
ATOM   1104  C   ASN   143      -5.917 -23.108  12.916  1.00  0.00           C
ATOM   1105  O   ASN   143      -6.539 -22.105  12.555  1.00  0.00           O
ATOM   1106  N   ALA   144      -5.606 -23.312  14.192  1.00  0.00           N
ATOM   1107  CA  ALA   144       6.955  -9.259   4.405 50.00  0.00           C
ATOM   1108  CB  ALA   144      -5.452 -20.956  14.890  1.00  0.00           C
ATOM   1109  C   ALA   144      -7.499 -22.224  15.366  1.00  0.00           C
ATOM   1110  O   ALA   144      -8.031 -21.166  15.712  1.00  0.00           O
ATOM   1111  N   GLU   145      -8.213 -23.318  15.117  1.00  0.00           N
ATOM   1112  CA  GLU   145       6.705 -10.250   5.368 50.00  0.00           C
ATOM   1113  CB  GLU   145     -10.516 -22.576  14.235  1.00  0.00           C
ATOM   1114  CG  GLU   145     -10.427 -23.149  12.819  1.00  0.00           C
ATOM   1115  CD  GLU   145     -11.205 -22.229  11.889  1.00  0.00           C
ATOM   1116  OE1 GLU   145     -11.712 -21.185  12.379  1.00  0.00           O
ATOM   1117  OE2 GLU   145     -11.303 -22.559  10.677  1.00  0.00           O
ATOM   1118  C   GLU   145     -10.202 -24.755  15.130  1.00  0.00           C
ATOM   1119  O   GLU   145      -9.568 -25.646  14.558  1.00  0.00           O
ATOM   1120  N   SER   146     -11.482 -24.927  15.693  1.00  0.00           N
ATOM   1121  CA  SER   146       6.693 -13.272   5.761 50.00  0.00           C
ATOM   1122  CB  SER   146     -10.917 -27.139  14.809  1.00  0.00           C
ATOM   1123  OG  SER   146     -11.418 -27.413  13.513  1.00  0.00           O
ATOM   1124  C   SER   146     -12.109 -26.950  17.000  1.00  0.00           C
ATOM   1125  O   SER   146     -11.941 -28.160  17.173  1.00  0.00           O
ATOM   1126  N   ALA   147     -12.468 -26.128  17.980  1.00  0.00           N
ATOM   1127  CA  ALA   147       7.683 -13.233   6.305 50.00  0.00           C
ATOM   1128  CB  ALA   147     -12.909 -25.453  20.324  1.00  0.00           C
ATOM   1129  C   ALA   147     -13.662 -27.759  19.526  1.00  0.00           C
ATOM   1130  O   ALA   147     -13.334 -28.770  20.145  1.00  0.00           O
ATOM   1131  N   LEU   148     -14.893 -27.615  18.990  1.00  0.00           N
ATOM   1132  CA  LEU   148       7.390 -15.380   7.028 50.00  0.00           C
ATOM   1133  CB  LEU   148     -17.029 -28.238  18.265  1.00  0.00           C
ATOM   1134  CG  LEU   148     -17.777 -27.044  18.862  1.00  0.00           C
ATOM   1135  CD1 LEU   148     -18.840 -26.429  17.953  1.00  0.00           C
ATOM   1136  CD2 LEU   148     -18.542 -27.341  20.151  1.00  0.00           C
ATOM   1137  C   LEU   148     -15.342 -30.053  18.586  1.00  0.00           C
ATOM   1138  O   LEU   148     -15.593 -31.116  19.180  1.00  0.00           O
ATOM   1139  N   GLU   149     -14.619 -30.000  17.472  1.00  0.00           N
ATOM   1140  CA  GLU   149       7.015 -18.023   6.100 50.00  0.00           C
ATOM   1141  CB  GLU   149     -13.503 -30.938  15.498  1.00  0.00           C
ATOM   1142  CG  GLU   149     -14.589 -30.668  14.455  1.00  0.00           C
ATOM   1143  CD  GLU   149     -13.904 -30.291  13.149  1.00  0.00           C
ATOM   1144  OE1 GLU   149     -12.648 -30.191  13.147  1.00  0.00           O
ATOM   1145  OE2 GLU   149     -14.628 -30.099  12.136  1.00  0.00           O
ATOM   1146  C   GLU   149     -12.977 -31.821  17.788  1.00  0.00           C
ATOM   1147  O   GLU   149     -12.877 -33.047  17.889  1.00  0.00           O
ATOM   1148  N   VAL   150     -12.170 -30.977  18.426  1.00  0.00           N
ATOM   1149  CA  VAL   150       7.972 -17.995   5.502 50.00  0.00           C
ATOM   1150  CB  VAL   150     -10.334 -30.271  19.867  1.00  0.00           C
ATOM   1151  CG1 VAL   150      -9.373 -30.640  20.998  1.00  0.00           C
ATOM   1152  CG2 VAL   150      -9.457 -29.629  18.790  1.00  0.00           C
ATOM   1153  C   VAL   150     -11.782 -32.127  20.565  1.00  0.00           C
ATOM   1154  O   VAL   150     -11.392 -33.213  20.969  1.00  0.00           O
ATOM   1155  N   LEU   151     -12.770 -31.473  21.145  1.00  0.00           N
ATOM   1156  CA  LEU   151       8.613 -18.384   6.391 50.00  0.00           C
ATOM   1157  CB  LEU   151     -14.503 -31.018  22.816  1.00  0.00           C
ATOM   1158  CG  LEU   151     -13.853 -29.816  23.504  1.00  0.00           C
ATOM   1159  CD1 LEU   151     -14.817 -28.698  23.898  1.00  0.00           C
ATOM   1160  CD2 LEU   151     -13.125 -30.134  24.809  1.00  0.00           C
ATOM   1161  C   LEU   151     -14.154 -33.381  21.923  1.00  0.00           C
ATOM   1162  O   LEU   151     -14.126 -34.325  22.728  1.00  0.00           O
ATOM   1163  N   GLN   152     -14.735 -33.456  20.724  1.00  0.00           N
ATOM   1164  CA  GLN   152       7.967 -21.224   6.486 50.00  0.00           C
ATOM   1165  CB  GLN   152     -15.884 -34.465  18.843  1.00  0.00           C
ATOM   1166  CG  GLN   152     -16.587 -35.694  18.263  1.00  0.00           C
ATOM   1167  CD  GLN   152     -17.761 -36.035  19.169  1.00  0.00           C
ATOM   1168  OE1 GLN   152     -18.529 -35.161  19.567  1.00  0.00           O
ATOM   1169  NE2 GLN   152     -17.965 -37.327  19.544  1.00  0.00           N
ATOM   1170  C   GLN   152     -14.212 -35.775  20.100  1.00  0.00           C
ATOM   1171  O   GLN   152     -14.426 -36.934  20.469  1.00  0.00           O
ATOM   1172  N   LEU   153     -13.055 -35.370  19.599  1.00  0.00           N
ATOM   1173  CA  LEU   153       8.104 -22.783   5.332 50.00  0.00           C
ATOM   1174  CB  LEU   153     -10.782 -35.448  18.812  1.00  0.00           C
ATOM   1175  CG  LEU   153      -9.983 -35.993  17.648  1.00  0.00           C
ATOM   1176  CD1 LEU   153      -9.154 -34.840  17.102  1.00  0.00           C
ATOM   1177  CD2 LEU   153      -9.088 -37.087  18.109  1.00  0.00           C
ATOM   1178  C   LEU   153     -11.452 -36.833  20.750  1.00  0.00           C
ATOM   1179  O   LEU   153     -11.145 -38.013  20.827  1.00  0.00           O
ATOM   1180  N   TYR   154     -11.391 -36.000  21.788  1.00  0.00           N
ATOM   1181  CA  TYR   154       9.232 -22.634   5.426 50.00  0.00           C
ATOM   1182  CB  TYR   154     -10.785 -35.263  24.049  1.00  0.00           C
ATOM   1183  CG  TYR   154     -10.406 -35.800  25.386  1.00  0.00           C
ATOM   1184  CD1 TYR   154      -9.106 -36.274  25.604  1.00  0.00           C
ATOM   1185  CD2 TYR   154     -11.328 -35.840  26.455  1.00  0.00           C
ATOM   1186  CE1 TYR   154      -8.709 -36.786  26.855  1.00  0.00           C
ATOM   1187  CE2 TYR   154     -10.941 -36.358  27.734  1.00  0.00           C
ATOM   1188  CZ  TYR   154      -9.624 -36.827  27.913  1.00  0.00           C
ATOM   1189  OH  TYR   154      -9.208 -37.337  29.123  1.00  0.00           O
ATOM   1190  C   TYR   154     -12.088 -37.357  23.737  1.00  0.00           C
ATOM   1191  O   TYR   154     -11.787 -38.301  24.479  1.00  0.00           O
ATOM   1192  N   GLN   155     -13.345 -37.049  23.434  1.00  0.00           N
ATOM   1193  CA  GLN   155       9.191 -24.061   6.340 50.00  0.00           C
ATOM   1194  CB  GLN   155     -15.742 -37.307  23.328  1.00  0.00           C
ATOM   1195  CG  GLN   155     -16.985 -38.087  23.760  1.00  0.00           C
ATOM   1196  CD  GLN   155     -18.204 -37.395  23.166  1.00  0.00           C
ATOM   1197  OE1 GLN   155     -18.081 -36.438  22.404  1.00  0.00           O
ATOM   1198  NE2 GLN   155     -19.449 -37.842  23.484  1.00  0.00           N
ATOM   1199  C   GLN   155     -14.284 -39.295  23.331  1.00  0.00           C
ATOM   1200  O   GLN   155     -14.398 -40.294  24.053  1.00  0.00           O
ATOM   1201  N   MET   156     -13.998 -39.402  22.036  1.00  0.00           N
ATOM   1202  CA  MET   156       8.650 -27.053   5.728 50.00  0.00           C
ATOM   1203  CB  MET   156     -13.540 -40.940  19.926  1.00  0.00           C
ATOM   1204  CG  MET   156     -14.714 -40.575  19.016  1.00  0.00           C
ATOM   1205  SD  MET   156     -16.246 -41.493  19.356  1.00  0.00           S
ATOM   1206  CE  MET   156     -15.586 -43.101  18.830  1.00  0.00           C
ATOM   1207  C   MET   156     -12.586 -41.425  21.932  1.00  0.00           C
ATOM   1208  O   MET   156     -12.438 -42.641  21.786  1.00  0.00           O
ATOM   1209  N   MET   157     -11.671 -40.684  22.550  1.00  0.00           N
ATOM   1210  CA  MET   157       9.447 -27.443   4.881 50.00  0.00           C
ATOM   1211  CB  MET   157      -9.079 -40.625  23.108  1.00  0.00           C
ATOM   1212  CG  MET   157      -8.377 -40.625  21.748  1.00  0.00           C
ATOM   1213  SD  MET   157      -6.765 -39.785  21.732  1.00  0.00           S
ATOM   1214  CE  MET   157      -5.941 -41.005  22.796  1.00  0.00           C
ATOM   1215  C   MET   157     -10.532 -41.550  24.587  1.00  0.00           C
ATOM   1216  O   MET   157      -9.567 -41.345  25.329  1.00  0.00           O
ATOM   1217  N   GLU   158     -11.689 -42.005  25.057  1.00  0.00           N
ATOM   1218  CA  GLU   158      10.296 -28.156   5.507 50.00  0.00           C
ATOM   1219  CB  GLU   158     -13.256 -42.701  26.968  1.00  0.00           C
ATOM   1220  CG  GLU   158     -14.305 -41.591  26.873  1.00  0.00           C
ATOM   1221  CD  GLU   158     -15.625 -42.148  27.388  1.00  0.00           C
ATOM   1222  OE1 GLU   158     -15.638 -43.328  27.829  1.00  0.00           O
ATOM   1223  OE2 GLU   158     -16.638 -41.399  27.347  1.00  0.00           O
ATOM   1224  C   GLU   158     -10.997 -43.422  26.984  1.00  0.00           C
ATOM   1225  O   GLU   158     -10.202 -43.261  27.914  1.00  0.00           O
ATOM   1226  N   GLY   159     -11.127 -44.602  26.385  1.00  0.00           N
ATOM   1227  CA  GLY   159      10.654 -31.524   4.802 50.00  0.00           C
ATOM   1228  C   GLY   159      -8.831 -45.423  26.630  1.00  0.00           C
ATOM   1229  O   GLY   159      -7.980 -45.803  27.439  1.00  0.00           O
ATOM   1230  N   PRO   160      -8.466 -44.784  25.516  1.00  0.00           N
ATOM   1231  CA  PRO   160      10.448 -30.199   3.894 50.00  0.00           C
ATOM   1232  CD  PRO   160      -9.332 -44.764  23.455  1.00  0.00           C
ATOM   1233  CB  PRO   160      -6.926 -43.944  23.765  1.00  0.00           C
ATOM   1234  CG  PRO   160      -7.943 -44.607  22.834  1.00  0.00           C
ATOM   1235  C   PRO   160      -6.811 -43.114  26.116  1.00  0.00           C
ATOM   1236  O   PRO   160      -7.472 -42.083  26.068  1.00  0.00           O
ATOM   1237  N   VAL   161      -5.792 -43.297  26.940  1.00  0.00           N
ATOM   1238  CA  VAL   161      11.308 -28.022   3.839 50.00  0.00           C
ATOM   1239  CB  VAL   161      -4.922 -43.203  29.187  1.00  0.00           C
ATOM   1240  CG1 VAL   161      -4.501 -42.347  30.383  1.00  0.00           C
ATOM   1241  CG2 VAL   161      -5.916 -44.219  29.753  1.00  0.00           C
ATOM   1242  C   VAL   161      -4.447 -41.390  27.545  1.00  0.00           C
ATOM   1243  O   VAL   161      -3.353 -41.780  27.132  1.00  0.00           O
ATOM   1244  N   THR   162      -4.797 -40.093  27.588  1.00  0.00           N
ATOM   1245  CA  THR   162      11.097 -24.718   3.590 50.00  0.00           C
ATOM   1246  CB  THR   162      -4.650 -37.763  27.659  1.00  0.00           C
ATOM   1247  OG1 THR   162      -3.985 -36.589  27.217  1.00  0.00           O
ATOM   1248  CG2 THR   162      -4.693 -37.776  29.196  1.00  0.00           C
ATOM   1249  C   THR   162      -2.533 -39.079  27.802  1.00  0.00           C
ATOM   1250  O   THR   162      -2.401 -39.479  28.951  1.00  0.00           O
ATOM   1251  N   THR   163      -1.508 -38.718  27.046  1.00  0.00           N
ATOM   1252  CA  THR   163      11.517 -24.287   2.594 50.00  0.00           C
ATOM   1253  CB  THR   163       0.774 -39.580  26.770  1.00  0.00           C
ATOM   1254  OG1 THR   163       0.845 -39.130  25.425  1.00  0.00           O
ATOM   1255  CG2 THR   163       0.228 -41.018  26.798  1.00  0.00           C
ATOM   1256  C   THR   163       0.355 -37.218  27.609  1.00  0.00           C
ATOM   1257  O   THR   163      -0.459 -36.289  27.598  1.00  0.00           O
ATOM   1258  N   ARG   164       1.677 -37.027  27.659  1.00  0.00           N
ATOM   1259  CA  ARG   164      11.654 -21.285   2.169 50.00  0.00           C
ATOM   1260  CB  ARG   164       3.781 -35.795  27.827  1.00  0.00           C
ATOM   1261  CG  ARG   164       4.228 -36.266  29.212  1.00  0.00           C
ATOM   1262  CD  ARG   164       5.745 -36.416  29.348  1.00  0.00           C
ATOM   1263  NE  ARG   164       6.023 -36.926  30.720  1.00  0.00           N
ATOM   1264  CZ  ARG   164       7.304 -37.210  31.096  1.00  0.00           C
ATOM   1265  NH1 ARG   164       8.119 -36.939  30.035  1.00  0.00           N
ATOM   1266  NH2 ARG   164       7.271 -37.655  32.386  1.00  0.00           N
ATOM   1267  C   ARG   164       1.934 -34.852  26.448  1.00  0.00           C
ATOM   1268  O   ARG   164       1.474 -33.702  26.553  1.00  0.00           O
ATOM   1269  N   VAL   165       2.170 -35.450  25.283  1.00  0.00           N
ATOM   1270  CA  VAL   165      10.540 -20.741   1.981 50.00  0.00           C
ATOM   1271  CB  VAL   165       2.366 -35.731  22.857  1.00  0.00           C
ATOM   1272  CG1 VAL   165       1.942 -35.239  21.472  1.00  0.00           C
ATOM   1273  CG2 VAL   165       3.887 -35.880  22.781  1.00  0.00           C
ATOM   1274  C   VAL   165       0.426 -34.486  23.788  1.00  0.00           C
ATOM   1275  O   VAL   165       0.099 -33.438  23.243  1.00  0.00           O
ATOM   1276  N   ALA   166      -0.464 -35.378  24.203  1.00  0.00           N
ATOM   1277  CA  ALA   166      10.244 -21.084   3.043 50.00  0.00           C
ATOM   1278  CB  ALA   166      -2.703 -36.233  24.765  1.00  0.00           C
ATOM   1279  C   ALA   166      -2.252 -33.775  24.775  1.00  0.00           C
ATOM   1280  O   ALA   166      -2.912 -32.920  24.183  1.00  0.00           O
ATOM   1281  N   THR   167      -1.780 -33.610  26.004  1.00  0.00           N
ATOM   1282  CA  THR   167      10.929 -18.276   3.607 50.00  0.00           C
ATOM   1283  CB  THR   167      -1.694 -32.751  28.278  1.00  0.00           C
ATOM   1284  OG1 THR   167      -2.312 -33.919  28.798  1.00  0.00           O
ATOM   1285  CG2 THR   167      -2.067 -31.544  29.155  1.00  0.00           C
ATOM   1286  C   THR   167      -1.563 -31.227  26.278  1.00  0.00           C
ATOM   1287  O   THR   167      -2.265 -30.234  26.033  1.00  0.00           O
ATOM   1288  N   ALA   168      -0.260 -31.286  26.050  1.00  0.00           N
ATOM   1289  CA  ALA   168      10.682 -16.059   2.901 50.00  0.00           C
ATOM   1290  CB  ALA   168       1.869 -30.673  25.157  1.00  0.00           C
ATOM   1291  C   ALA   168      -0.193 -29.704  24.158  1.00  0.00           C
ATOM   1292  O   ALA   168      -0.404 -28.498  23.965  1.00  0.00           O
ATOM   1293  N   ALA   169      -0.533 -30.646  23.280  1.00  0.00           N
ATOM   1294  CA  ALA   169       9.563 -16.494   3.014 50.00  0.00           C
ATOM   1295  CB  ALA   169      -1.303 -31.532  21.081  1.00  0.00           C
ATOM   1296  C   ALA   169      -2.518 -29.659  22.151  1.00  0.00           C
ATOM   1297  O   ALA   169      -2.815 -28.685  21.450  1.00  0.00           O
ATOM   1298  N   LEU   170      -3.351 -30.193  23.045  1.00  0.00           N
ATOM   1299  CA  LEU   170       9.569 -15.775   4.195 50.00  0.00           C
ATOM   1300  CB  LEU   170      -5.538 -30.507  24.164  1.00  0.00           C
ATOM   1301  CG  LEU   170      -5.916 -31.861  23.560  1.00  0.00           C
ATOM   1302  CD1 LEU   170      -6.611 -32.825  24.521  1.00  0.00           C
ATOM   1303  CD2 LEU   170      -6.874 -31.793  22.372  1.00  0.00           C
ATOM   1304  C   LEU   170      -4.619 -28.221  23.781  1.00  0.00           C
ATOM   1305  O   LEU   170      -5.298 -27.293  23.306  1.00  0.00           O
ATOM   1306  N   HIS   171      -3.755 -28.052  24.771  1.00  0.00           N
ATOM   1307  CA  HIS   171      10.147 -12.694   4.307 50.00  0.00           C
ATOM   1308  ND1 HIS   171      -4.696 -26.892  28.348  1.00  0.00           N
ATOM   1309  CG  HIS   171      -3.564 -27.471  27.812  1.00  0.00           C
ATOM   1310  CB  HIS   171      -2.759 -26.865  26.700  1.00  0.00           C
ATOM   1311  NE2 HIS   171      -4.422 -28.704  29.412  1.00  0.00           N
ATOM   1312  CD2 HIS   171      -3.399 -28.625  28.499  1.00  0.00           C
ATOM   1313  CE1 HIS   171      -5.188 -27.637  29.291  1.00  0.00           C
ATOM   1314  C   HIS   171      -2.987 -25.709  24.405  1.00  0.00           C
ATOM   1315  O   HIS   171      -3.445 -24.574  24.396  1.00  0.00           O
ATOM   1316  N   ARG   172      -2.039 -26.107  23.561  1.00  0.00           N
ATOM   1317  CA  ARG   172       9.444 -11.445   3.498 50.00  0.00           C
ATOM   1318  CB  ARG   172      -0.218 -25.780  21.932  1.00  0.00           C
ATOM   1319  CG  ARG   172       0.468 -24.839  20.940  1.00  0.00           C
ATOM   1320  CD  ARG   172       1.835 -25.337  20.468  1.00  0.00           C
ATOM   1321  NE  ARG   172       2.372 -24.331  19.508  1.00  0.00           N
ATOM   1322  CZ  ARG   172       3.533 -24.581  18.835  1.00  0.00           C
ATOM   1323  NH1 ARG   172       3.985 -25.808  19.227  1.00  0.00           N
ATOM   1324  NH2 ARG   172       3.788 -23.507  18.033  1.00  0.00           N
ATOM   1325  C   ARG   172      -2.509 -24.883  21.437  1.00  0.00           C
ATOM   1326  O   ARG   172      -2.517 -23.790  20.859  1.00  0.00           O
ATOM   1327  N   PHE   173      -3.364 -25.857  21.155  1.00  0.00           N
ATOM   1328  CA  PHE   173       8.549 -12.028   4.132 50.00  0.00           C
ATOM   1329  CB  PHE   173      -5.216 -26.981  20.057  1.00  0.00           C
ATOM   1330  CG  PHE   173      -6.028 -27.162  21.293  1.00  0.00           C
ATOM   1331  CD1 PHE   173      -7.346 -26.664  21.403  1.00  0.00           C
ATOM   1332  CD2 PHE   173      -5.486 -27.850  22.394  1.00  0.00           C
ATOM   1333  CE1 PHE   173      -8.117 -26.845  22.588  1.00  0.00           C
ATOM   1334  CE2 PHE   173      -6.236 -28.046  23.593  1.00  0.00           C
ATOM   1335  CZ  PHE   173      -7.558 -27.539  23.689  1.00  0.00           C
ATOM   1336  C   PHE   173      -5.351 -24.608  20.734  1.00  0.00           C
ATOM   1337  O   PHE   173      -5.944 -23.846  19.975  1.00  0.00           O
ATOM   1338  N   LEU   174      -5.458 -24.559  22.055  1.00  0.00           N
ATOM   1339  CA  LEU   174       9.092  -9.711   5.043 50.00  0.00           C
ATOM   1340  CB  LEU   174      -6.533 -23.859  24.149  1.00  0.00           C
ATOM   1341  CG  LEU   174      -7.573 -24.976  24.260  1.00  0.00           C
ATOM   1342  CD1 LEU   174      -7.798 -25.511  25.674  1.00  0.00           C
ATOM   1343  CD2 LEU   174      -8.979 -24.598  23.796  1.00  0.00           C
ATOM   1344  C   LEU   174      -5.494 -22.106  22.683  1.00  0.00           C
ATOM   1345  O   LEU   174      -6.113 -21.091  22.353  1.00  0.00           O
ATOM   1346  N   THR   175      -4.220 -22.075  23.024  1.00  0.00           N
ATOM   1347  CA  THR   175       9.145  -7.171   4.486 50.00  0.00           C
ATOM   1348  CB  THR   175      -2.041 -21.051  23.478  1.00  0.00           C
ATOM   1349  OG1 THR   175      -2.009 -21.646  24.767  1.00  0.00           O
ATOM   1350  CG2 THR   175      -1.305 -19.701  23.522  1.00  0.00           C
ATOM   1351  C   THR   175      -3.517 -20.147  21.699  1.00  0.00           C
ATOM   1352  O   THR   175      -3.604 -18.944  21.623  1.00  0.00           O
ATOM   1353  N   ASN   176      -3.929 -18.878  21.639  1.00  0.00           N
ATOM   1354  CA  ASN   176       9.202  -4.134   5.037 50.00  0.00           C
ATOM   1355  CB  ASN   176      -4.091 -16.521  22.379  1.00  0.00           C
ATOM   1356  CG  ASN   176      -5.131 -16.150  21.331  1.00  0.00           C
ATOM   1357  OD1 ASN   176      -5.837 -17.011  20.807  1.00  0.00           O
ATOM   1358  ND2 ASN   176      -5.282 -14.848  20.969  1.00  0.00           N
ATOM   1359  C   ASN   176      -5.706 -18.112  23.263  1.00  0.00           C
ATOM   1360  O   ASN   176      -6.228 -17.195  23.903  1.00  0.00           O
ATOM   1361  N   ASP   177      -6.365 -19.245  23.007  1.00  0.00           N
ATOM   1362  CA  ASP   177       9.159  -5.611   5.951 50.00  0.00           C
ATOM   1363  CB  ASP   177      -8.476 -20.677  23.144  1.00  0.00           C
ATOM   1364  CG  ASP   177      -8.831 -20.552  21.669  1.00  0.00           C
ATOM   1365  OD1 ASP   177      -8.637 -19.442  21.106  1.00  0.00           O
ATOM   1366  OD2 ASP   177      -9.300 -21.566  21.086  1.00  0.00           O
ATOM   1367  C   ASP   177      -7.680 -19.485  25.044  1.00  0.00           C
ATOM   1368  O   ASP   177      -7.029 -20.342  25.647  1.00  0.00           O
ATOM   1369  N   ALA   178      -8.413 -18.544  25.693  1.00  0.00           N
ATOM   1370  CA  ALA   178       9.949  -4.381   6.497 50.00  0.00           C
ATOM   1371  CB  ALA   178      -8.843 -17.001  27.571  1.00  0.00           C
ATOM   1372  C   ALA   178      -9.529 -19.358  27.773  1.00  0.00           C
ATOM   1373  O   ALA   178      -9.205 -20.196  28.620  1.00  0.00           O
ATOM   1374  N   SER   179     -10.785 -19.233  27.374  1.00  0.00           N
ATOM   1375  CA  SER   179       9.916  -5.961   7.423 50.00  0.00           C
ATOM   1376  CB  SER   179     -13.369 -19.811  27.759  1.00  0.00           C
ATOM   1377  OG  SER   179     -14.094 -20.874  28.360  1.00  0.00           O
ATOM   1378  C   SER   179     -11.854 -21.420  27.278  1.00  0.00           C
ATOM   1379  O   SER   179     -12.201 -21.588  26.106  1.00  0.00           O
ATOM   1380  N   GLU   180     -11.534 -22.406  28.091  1.00  0.00           N
ATOM   1381  CA  GLU   180       9.971  -9.703   7.037 50.00  0.00           C
ATOM   1382  CB  GLU   180     -11.801 -23.690  26.061  1.00  0.00           C
ATOM   1383  CG  GLU   180     -12.035 -22.235  25.664  1.00  0.00           C
ATOM   1384  CD  GLU   180     -13.314 -21.972  24.881  1.00  0.00           C
ATOM   1385  OE1 GLU   180     -13.653 -20.773  24.713  1.00  0.00           O
ATOM   1386  OE2 GLU   180     -13.968 -22.935  24.422  1.00  0.00           O
ATOM   1387  C   GLU   180     -10.265 -24.533  27.895  1.00  0.00           C
ATOM   1388  O   GLU   180     -10.300 -25.692  28.298  1.00  0.00           O
ATOM   1389  N   VAL   181      -9.129 -23.861  27.757  1.00  0.00           N
ATOM   1390  CA  VAL   181      10.496 -10.245   5.976 50.00  0.00           C
ATOM   1391  CB  VAL   181      -6.714 -23.538  27.744  1.00  0.00           C
ATOM   1392  CG1 VAL   181      -5.356 -23.994  28.282  1.00  0.00           C
ATOM   1393  CG2 VAL   181      -6.523 -23.415  26.231  1.00  0.00           C
ATOM   1394  C   VAL   181      -7.814 -24.719  29.622  1.00  0.00           C
ATOM   1395  O   VAL   181      -7.386 -25.786  30.039  1.00  0.00           O
ATOM   1396  N   SER   182      -8.271 -23.753  30.418  1.00  0.00           N
ATOM   1397  CA  SER   182      11.492  -9.377   6.496 50.00  0.00           C
ATOM   1398  CB  SER   182      -8.717 -22.587  32.592  1.00  0.00           C
ATOM   1399  OG  SER   182      -7.784 -21.543  32.356  1.00  0.00           O
ATOM   1400  C   SER   182      -9.139 -25.056  32.306  1.00  0.00           C
ATOM   1401  O   SER   182      -8.776 -25.842  33.190  1.00  0.00           O
ATOM   1402  N   GLY   183     -10.291 -25.167  31.648  1.00  0.00           N
ATOM   1403  CA  GLY   183      11.313 -11.777   7.141 50.00  0.00           C
ATOM   1404  C   GLY   183     -10.609 -27.606  31.587  1.00  0.00           C
ATOM   1405  O   GLY   183     -10.716 -28.564  32.382  1.00  0.00           O
ATOM   1406  N   LEU   184      -9.960 -27.688  30.427  1.00  0.00           N
ATOM   1407  CA  LEU   184      11.038 -14.567   6.199 50.00  0.00           C
ATOM   1408  CB  LEU   184      -8.718 -28.804  28.616  1.00  0.00           C
ATOM   1409  CG  LEU   184      -8.021 -30.079  28.137  1.00  0.00           C
ATOM   1410  CD1 LEU   184      -8.928 -31.302  28.010  1.00  0.00           C
ATOM   1411  CD2 LEU   184      -7.366 -29.978  26.760  1.00  0.00           C
ATOM   1412  C   LEU   184      -8.274 -29.321  31.047  1.00  0.00           C
ATOM   1413  O   LEU   184      -8.258 -30.449  31.521  1.00  0.00           O
ATOM   1414  N   LEU   185      -7.416 -28.373  31.403  1.00  0.00           N
ATOM   1415  CA  LEU   185      11.965 -14.004   5.598 50.00  0.00           C
ATOM   1416  CB  LEU   185      -5.529 -27.337  32.619  1.00  0.00           C
ATOM   1417  CG  LEU   185      -4.392 -27.527  33.625  1.00  0.00           C
ATOM   1418  CD1 LEU   185      -3.307 -28.514  33.195  1.00  0.00           C
ATOM   1419  CD2 LEU   185      -3.606 -26.260  33.958  1.00  0.00           C
ATOM   1420  C   LEU   185      -6.914 -29.204  33.661  1.00  0.00           C
ATOM   1421  O   LEU   185      -6.412 -30.208  34.186  1.00  0.00           O
ATOM   1422  N   GLN   186      -7.983 -28.571  34.145  1.00  0.00           N
ATOM   1423  CA  GLN   186      12.632 -14.193   6.539 50.00  0.00           C
ATOM   1424  CB  GLN   186      -9.848 -28.101  35.704  1.00  0.00           C
ATOM   1425  CG  GLN   186      -9.414 -26.726  36.214  1.00  0.00           C
ATOM   1426  CD  GLN   186     -10.663 -25.875  36.396  1.00  0.00           C
ATOM   1427  OE1 GLN   186     -11.770 -26.299  36.069  1.00  0.00           O
ATOM   1428  NE2 GLN   186     -10.553 -24.628  36.928  1.00  0.00           N
ATOM   1429  C   GLN   186      -9.137 -30.466  35.217  1.00  0.00           C
ATOM   1430  O   GLN   186      -8.817 -31.289  36.074  1.00  0.00           O
ATOM   1431  N   PHE   187      -9.877 -30.771  34.151  1.00  0.00           N
ATOM   1432  CA  PHE   187      12.161 -17.445   6.607 50.00  0.00           C
ATOM   1433  CB  PHE   187     -10.953 -32.250  32.505  1.00  0.00           C
ATOM   1434  CG  PHE   187     -11.330 -33.675  32.288  1.00  0.00           C
ATOM   1435  CD1 PHE   187     -12.487 -34.241  32.869  1.00  0.00           C
ATOM   1436  CD2 PHE   187     -10.523 -34.499  31.482  1.00  0.00           C
ATOM   1437  CE1 PHE   187     -12.841 -35.604  32.652  1.00  0.00           C
ATOM   1438  CE2 PHE   187     -10.855 -35.868  31.248  1.00  0.00           C
ATOM   1439  CZ  PHE   187     -12.020 -36.422  31.839  1.00  0.00           C
ATOM   1440  C   PHE   187      -9.142 -33.123  33.978  1.00  0.00           C
ATOM   1441  O   PHE   187      -9.165 -34.083  34.740  1.00  0.00           O
ATOM   1442  N   LEU   188      -8.117 -32.834  33.186  1.00  0.00           N
ATOM   1443  CA  LEU   188      12.314 -18.862   5.429 50.00  0.00           C
ATOM   1444  CB  LEU   188      -5.853 -32.749  32.328  1.00  0.00           C
ATOM   1445  CG  LEU   188      -4.860 -33.403  31.361  1.00  0.00           C
ATOM   1446  CD1 LEU   188      -3.722 -34.080  32.109  1.00  0.00           C
ATOM   1447  CD2 LEU   188      -5.573 -34.391  30.444  1.00  0.00           C
ATOM   1448  C   LEU   188      -6.291 -33.843  34.540  1.00  0.00           C
ATOM   1449  O   LEU   188      -5.653 -34.875  34.783  1.00  0.00           O
ATOM   1450  N   LYS   189      -6.510 -32.896  35.452  1.00  0.00           N
ATOM   1451  CA  LYS   189      13.419 -17.980   5.595 50.00  0.00           C
ATOM   1452  CB  LYS   189      -5.859 -31.625  37.451  1.00  0.00           C
ATOM   1453  CG  LYS   189      -4.420 -31.185  37.677  1.00  0.00           C
ATOM   1454  CD  LYS   189      -4.335 -29.665  37.832  1.00  0.00           C
ATOM   1455  CE  LYS   189      -5.378 -29.113  38.819  1.00  0.00           C
ATOM   1456  NZ  LYS   189      -5.293 -27.615  38.994  1.00  0.00           N
ATOM   1457  C   LYS   189      -6.930 -33.876  37.692  1.00  0.00           C
ATOM   1458  O   LYS   189      -6.455 -34.745  38.421  1.00  0.00           O
ATOM   1459  N   LEU   190      -8.236 -33.618  37.612  1.00  0.00           N
ATOM   1460  CA  LEU   190      13.619 -19.217   6.578 50.00  0.00           C
ATOM   1461  CB  LEU   190     -10.641 -33.817  38.072  1.00  0.00           C
ATOM   1462  CG  LEU   190     -10.877 -32.431  38.675  1.00  0.00           C
ATOM   1463  CD1 LEU   190     -12.193 -31.767  38.270  1.00  0.00           C
ATOM   1464  CD2 LEU   190     -10.913 -32.388  40.202  1.00  0.00           C
ATOM   1465  C   LEU   190      -9.180 -35.858  38.002  1.00  0.00           C
ATOM   1466  O   LEU   190      -9.307 -36.716  38.875  1.00  0.00           O
ATOM   1467  N   ASN   191      -8.942 -36.168  36.732  1.00  0.00           N
ATOM   1468  CA  ASN   191      13.181 -22.551   5.950 50.00  0.00           C
ATOM   1469  CB  ASN   191      -8.946 -37.696  34.808  1.00  0.00           C
ATOM   1470  CG  ASN   191     -10.403 -37.428  34.460  1.00  0.00           C
ATOM   1471  OD1 ASN   191     -11.289 -37.547  35.306  1.00  0.00           O
ATOM   1472  ND2 ASN   191     -10.732 -37.051  33.196  1.00  0.00           N
ATOM   1473  C   ASN   191      -7.440 -38.105  36.763  1.00  0.00           C
ATOM   1474  O   ASN   191      -7.010 -39.157  36.298  1.00  0.00           O
ATOM   1475  N   ASN   192      -6.775 -37.387  37.662  1.00  0.00           N
ATOM   1476  CA  ASN   192      13.994 -22.702   5.213 50.00  0.00           C
ATOM   1477  CB  ASN   192      -5.893 -39.171  39.061  1.00  0.00           C
ATOM   1478  CG  ASN   192      -6.791 -38.817  40.238  1.00  0.00           C
ATOM   1479  OD1 ASN   192      -6.758 -37.696  40.744  1.00  0.00           O
ATOM   1480  ND2 ASN   192      -7.642 -39.753  40.737  1.00  0.00           N
ATOM   1481  C   ASN   192      -4.390 -38.122  37.384  1.00  0.00           C
ATOM   1482  O   ASN   192      -3.592 -39.046  37.604  1.00  0.00           O
ATOM   1483  N   VAL   193      -4.278 -37.301  36.345  1.00  0.00           N
ATOM   1484  CA  VAL   193      13.383 -22.378   4.280 50.00  0.00           C
ATOM   1485  CB  VAL   193      -3.462 -37.045  34.029  1.00  0.00           C
ATOM   1486  CG1 VAL   193      -2.243 -36.967  33.108  1.00  0.00           C
ATOM   1487  CG2 VAL   193      -4.379 -38.090  33.390  1.00  0.00           C
ATOM   1488  C   VAL   193      -2.108 -36.342  36.040  1.00  0.00           C
ATOM   1489  O   VAL   193      -2.513 -35.332  36.618  1.00  0.00           O
ATOM   1490  N   PRO   194      -0.812 -36.625  35.891  1.00  0.00           N
ATOM   1491  CA  PRO   194      13.922 -20.665   3.561 50.00  0.00           C
ATOM   1492  CD  PRO   194       0.050 -38.544  36.045  1.00  0.00           C
ATOM   1493  CB  PRO   194       1.259 -36.622  37.229  1.00  0.00           C
ATOM   1494  CG  PRO   194       1.323 -38.074  36.752  1.00  0.00           C
ATOM   1495  C   PRO   194       0.937 -34.890  35.432  1.00  0.00           C
ATOM   1496  O   PRO   194       1.832 -35.366  34.718  1.00  0.00           O
ATOM   1497  N   LEU   195       0.553 -33.592  35.371  1.00  0.00           N
ATOM   1498  CA  LEU   195      13.370 -17.663   3.355 50.00  0.00           C
ATOM   1499  CB  LEU   195       0.380 -31.314  34.832  1.00  0.00           C
ATOM   1500  CG  LEU   195      -1.135 -31.323  34.622  1.00  0.00           C
ATOM   1501  CD1 LEU   195      -1.869 -30.095  35.159  1.00  0.00           C
ATOM   1502  CD2 LEU   195      -1.585 -31.394  33.163  1.00  0.00           C
ATOM   1503  C   LEU   195       2.578 -32.451  34.491  1.00  0.00           C
ATOM   1504  O   LEU   195       3.113 -32.360  35.598  1.00  0.00           O
ATOM   1505  N   THR   196       3.260 -32.420  33.350  1.00  0.00           N
ATOM   1506  CA  THR   196      13.355 -17.320   2.285 50.00  0.00           C
ATOM   1507  CB  THR   196       5.386 -33.298  32.488  1.00  0.00           C
ATOM   1508  OG1 THR   196       5.013 -33.136  31.128  1.00  0.00           O
ATOM   1509  CG2 THR   196       4.953 -34.692  32.972  1.00  0.00           C
ATOM   1510  C   THR   196       5.106 -30.840  32.856  1.00  0.00           C
ATOM   1511  O   THR   196       4.257 -30.052  32.445  1.00  0.00           O
ATOM   1512  N   ARG   197       6.402 -30.555  32.898  1.00  0.00           N
ATOM   1513  CA  ARG   197      13.139 -14.478   1.804 50.00  0.00           C
ATOM   1514  CB  ARG   197       8.469 -29.404  32.211  1.00  0.00           C
ATOM   1515  CG  ARG   197       9.117 -28.099  31.745  1.00  0.00           C
ATOM   1516  CD  ARG   197      10.645 -28.159  31.693  1.00  0.00           C
ATOM   1517  NE  ARG   197      11.124 -26.874  31.110  1.00  0.00           N
ATOM   1518  CZ  ARG   197      12.459 -26.591  31.085  1.00  0.00           C
ATOM   1519  NH1 ARG   197      13.118 -27.644  31.651  1.00  0.00           N
ATOM   1520  NH2 ARG   197      12.615 -25.365  30.504  1.00  0.00           N
ATOM   1521  C   ARG   197       6.308 -28.849  31.073  1.00  0.00           C
ATOM   1522  O   ARG   197       6.088 -27.659  30.845  1.00  0.00           O
ATOM   1523  N   GLU   198       5.999 -29.814  30.215  1.00  0.00           N
ATOM   1524  CA  GLU   198      12.044 -14.997   1.853 50.00  0.00           C
ATOM   1525  CB  GLU   198       5.611 -30.811  27.983  1.00  0.00           C
ATOM   1526  CG  GLU   198       7.031 -31.295  27.683  1.00  0.00           C
ATOM   1527  CD  GLU   198       6.929 -32.619  26.939  1.00  0.00           C
ATOM   1528  OE1 GLU   198       5.783 -33.114  26.767  1.00  0.00           O
ATOM   1529  OE2 GLU   198       7.995 -33.153  26.533  1.00  0.00           O
ATOM   1530  C   GLU   198       4.186 -28.870  28.862  1.00  0.00           C
ATOM   1531  O   GLU   198       3.918 -27.891  28.144  1.00  0.00           O
ATOM   1532  N   PHE   199       3.329 -29.384  29.726  1.00  0.00           N
ATOM   1533  CA  PHE   199      11.985 -14.442   2.978 50.00  0.00           C
ATOM   1534  CB  PHE   199       1.092 -30.070  30.394  1.00  0.00           C
ATOM   1535  CG  PHE   199      -0.328 -29.624  30.328  1.00  0.00           C
ATOM   1536  CD1 PHE   199      -1.089 -29.721  29.141  1.00  0.00           C
ATOM   1537  CD2 PHE   199      -0.948 -29.087  31.471  1.00  0.00           C
ATOM   1538  CE1 PHE   199      -2.448 -29.295  29.088  1.00  0.00           C
ATOM   1539  CE2 PHE   199      -2.308 -28.653  31.445  1.00  0.00           C
ATOM   1540  CZ  PHE   199      -3.060 -28.756  30.246  1.00  0.00           C
ATOM   1541  C   PHE   199       1.861 -27.648  30.505  1.00  0.00           C
ATOM   1542  O   PHE   199       1.078 -26.769  30.146  1.00  0.00           O
ATOM   1543  N   HIS   200       2.684 -27.502  31.538  1.00  0.00           N
ATOM   1544  CA  HIS   200      12.653 -11.509   3.212 50.00  0.00           C
ATOM   1545  ND1 HIS   200       1.890 -26.849  35.284  1.00  0.00           N
ATOM   1546  CG  HIS   200       3.031 -27.218  34.601  1.00  0.00           C
ATOM   1547  CB  HIS   200       3.656 -26.401  33.510  1.00  0.00           C
ATOM   1548  NE2 HIS   200       2.471 -28.713  36.108  1.00  0.00           N
ATOM   1549  CD2 HIS   200       3.392 -28.409  35.136  1.00  0.00           C
ATOM   1550  CE1 HIS   200       1.575 -27.747  36.169  1.00  0.00           C
ATOM   1551  C   HIS   200       3.228 -25.140  31.346  1.00  0.00           C
ATOM   1552  O   HIS   200       2.574 -24.115  31.216  1.00  0.00           O
ATOM   1553  N   ASP   201       4.344 -25.378  30.666  1.00  0.00           N
ATOM   1554  CA  ASP   201      12.052  -9.911   2.496 50.00  0.00           C
ATOM   1555  CB  ASP   201       6.179 -25.015  29.087  1.00  0.00           C
ATOM   1556  CG  ASP   201       7.267 -24.883  30.143  1.00  0.00           C
ATOM   1557  OD1 ASP   201       7.026 -24.172  31.155  1.00  0.00           O
ATOM   1558  OD2 ASP   201       8.354 -25.492  29.952  1.00  0.00           O
ATOM   1559  C   ASP   201       3.866 -24.109  28.582  1.00  0.00           C
ATOM   1560  O   ASP   201       3.850 -22.985  28.066  1.00  0.00           O
ATOM   1561  N   GLN   202       2.998 -25.055  28.249  1.00  0.00           N
ATOM   1562  CA  GLN   202      11.125 -10.512   3.070 50.00  0.00           C
ATOM   1563  CB  GLN   202       1.358 -26.093  26.772  1.00  0.00           C
ATOM   1564  CG  GLN   202       2.331 -26.953  25.963  1.00  0.00           C
ATOM   1565  CD  GLN   202       2.757 -26.156  24.739  1.00  0.00           C
ATOM   1566  OE1 GLN   202       1.931 -25.549  24.059  1.00  0.00           O
ATOM   1567  NE2 GLN   202       4.071 -26.113  24.391  1.00  0.00           N
ATOM   1568  C   GLN   202       0.881 -23.867  27.808  1.00  0.00           C
ATOM   1569  O   GLN   202       0.422 -22.929  27.127  1.00  0.00           O
ATOM   1570  N   LEU   203       0.472 -24.150  29.042  1.00  0.00           N
ATOM   1571  CA  LEU   203      11.549  -8.935   4.115 50.00  0.00           C
ATOM   1572  CB  LEU   203      -0.754 -23.948  31.131  1.00  0.00           C
ATOM   1573  CG  LEU   203      -2.070 -24.668  31.397  1.00  0.00           C
ATOM   1574  CD1 LEU   203      -2.302 -24.706  32.907  1.00  0.00           C
ATOM   1575  CD2 LEU   203      -3.239 -23.987  30.663  1.00  0.00           C
ATOM   1576  C   LEU   203      -0.093 -21.907  29.887  1.00  0.00           C
ATOM   1577  O   LEU   203      -0.860 -20.945  29.642  1.00  0.00           O
ATOM   1578  N   GLU   204       1.167 -21.778  30.284  1.00  0.00           N
ATOM   1579  CA  GLU   204      11.812  -5.990   3.738 50.00  0.00           C
ATOM   1580  CB  GLU   204       3.326 -20.681  30.691  1.00  0.00           C
ATOM   1581  CG  GLU   204       4.090 -19.358  30.777  1.00  0.00           C
ATOM   1582  CD  GLU   204       5.509 -19.664  31.237  1.00  0.00           C
ATOM   1583  OE1 GLU   204       5.806 -20.864  31.482  1.00  0.00           O
ATOM   1584  OE2 GLU   204       6.314 -18.701  31.350  1.00  0.00           O
ATOM   1585  C   GLU   204       1.783 -19.829  28.963  1.00  0.00           C
ATOM   1586  O   GLU   204       1.532 -18.641  28.938  1.00  0.00           O
ATOM   1587  N   LYS   205       1.994 -20.546  27.857  1.00  0.00           N
ATOM   1588  CA  LYS   205      10.724  -5.744   3.449 50.00  0.00           C
ATOM   1589  CB  LYS   205       2.169 -20.831  25.375  1.00  0.00           C
ATOM   1590  CG  LYS   205       3.681 -21.032  25.250  1.00  0.00           C
ATOM   1591  CD  LYS   205       4.083 -21.944  24.090  1.00  0.00           C
ATOM   1592  CE  LYS   205       5.594 -22.165  23.980  1.00  0.00           C
ATOM   1593  NZ  LYS   205       5.891 -23.082  22.857  1.00  0.00           N
ATOM   1594  C   LYS   205       0.348 -19.379  26.298  1.00  0.00           C
ATOM   1595  O   LYS   205       0.183 -18.307  25.730  1.00  0.00           O
ATOM   1596  N   ASN   206      -0.670 -20.115  26.730  1.00  0.00           N
ATOM   1597  CA  ASN   206      10.386  -5.512   4.534 50.00  0.00           C
ATOM   1598  CB  ASN   206      -3.078 -20.666  27.052  1.00  0.00           C
ATOM   1599  CG  ASN   206      -4.467 -20.123  26.750  1.00  0.00           C
ATOM   1600  OD1 ASN   206      -4.836 -19.933  25.592  1.00  0.00           O
ATOM   1601  ND2 ASN   206      -5.315 -19.843  27.777  1.00  0.00           N
ATOM   1602  C   ASN   206      -2.305 -18.317  27.291  1.00  0.00           C
ATOM   1603  O   ASN   206      -2.850 -17.350  26.722  1.00  0.00           O
ATOM   1604  N   PHE   207      -1.927 -18.281  28.562  1.00  0.00           N
ATOM   1605  CA  PHE   207      11.088  -2.711   5.035 50.00  0.00           C
ATOM   1606  CB  PHE   207      -1.896 -17.313  30.796  1.00  0.00           C
ATOM   1607  CG  PHE   207      -2.980 -18.122  31.408  1.00  0.00           C
ATOM   1608  CD1 PHE   207      -4.291 -17.680  31.368  1.00  0.00           C
ATOM   1609  CD2 PHE   207      -2.699 -19.322  32.031  1.00  0.00           C
ATOM   1610  CE1 PHE   207      -5.314 -18.434  31.928  1.00  0.00           C
ATOM   1611  CE2 PHE   207      -3.716 -20.078  32.603  1.00  0.00           C
ATOM   1612  CZ  PHE   207      -5.024 -19.630  32.550  1.00  0.00           C
ATOM   1613  C   PHE   207      -1.163 -15.941  28.833  1.00  0.00           C
ATOM   1614  O   PHE   207      -1.470 -14.754  28.960  1.00  0.00           O
ATOM   1615  N   VAL   208      -0.022 -16.328  28.278  1.00  0.00           N
ATOM   1616  CA  VAL   208      10.862  -1.136   4.179 50.00  0.00           C
ATOM   1617  CB  VAL   208       2.269 -16.059  27.409  1.00  0.00           C
ATOM   1618  CG1 VAL   208       3.258 -15.137  26.693  1.00  0.00           C
ATOM   1619  CG2 VAL   208       3.028 -16.590  28.627  1.00  0.00           C
ATOM   1620  C   VAL   208       0.380 -14.653  26.554  1.00  0.00           C
ATOM   1621  O   VAL   208       0.459 -13.429  26.455  1.00  0.00           O
ATOM   1622  N   MET   209      -0.180 -15.417  25.627  1.00  0.00           N
ATOM   1623  CA  MET   209       9.697  -0.939   4.380 50.00  0.00           C
ATOM   1624  CB  MET   209      -1.041 -15.905  23.372  1.00  0.00           C
ATOM   1625  CG  MET   209       0.228 -16.520  22.778  1.00  0.00           C
ATOM   1626  SD  MET   209       1.337 -15.328  21.970  1.00  0.00           S
ATOM   1627  CE  MET   209       0.203 -14.959  20.600  1.00  0.00           C
ATOM   1628  C   MET   209      -2.049 -14.097  24.744  1.00  0.00           C
ATOM   1629  O   MET   209      -2.441 -13.196  24.002  1.00  0.00           O
ATOM   1630  N   ARG   210      -2.719 -14.475  25.829  1.00  0.00           N
ATOM   1631  CA  ARG   210       9.877   0.158   5.549 50.00  0.00           C
ATOM   1632  CB  ARG   210      -4.835 -14.722  27.025  1.00  0.00           C
ATOM   1633  CG  ARG   210      -6.130 -14.063  27.503  1.00  0.00           C
ATOM   1634  CD  ARG   210      -7.100 -13.723  26.369  1.00  0.00           C
ATOM   1635  NE  ARG   210      -8.265 -13.017  26.972  1.00  0.00           N
ATOM   1636  CZ  ARG   210      -8.204 -11.672  27.199  1.00  0.00           C
ATOM   1637  NH1 ARG   210      -6.979 -11.235  26.787  1.00  0.00           N
ATOM   1638  NH2 ARG   210      -9.393 -11.290  27.751  1.00  0.00           N
ATOM   1639  C   ARG   210      -3.740 -12.483  26.889  1.00  0.00           C
ATOM   1640  O   ARG   210      -4.584 -11.576  26.853  1.00  0.00           O
ATOM   1641  N   MET   211      -2.591 -12.366  27.536  1.00  0.00           N
ATOM   1642  CA  MET   211      10.629   3.033   5.531 50.00  0.00           C
ATOM   1643  CB  MET   211      -2.774  -9.916  27.605  1.00  0.00           C
ATOM   1644  CG  MET   211      -2.010  -9.630  26.311  1.00  0.00           C
ATOM   1645  SD  MET   211      -2.492  -8.088  25.476  1.00  0.00           S
ATOM   1646  CE  MET   211      -4.087  -8.723  24.882  1.00  0.00           C
ATOM   1647  C   MET   211      -2.990 -11.321  29.663  1.00  0.00           C
ATOM   1648  O   MET   211      -3.298 -10.336  30.336  1.00  0.00           O
ATOM   1649  N   LYS   212      -3.232 -12.571  30.042  1.00  0.00           N
ATOM   1650  CA  LYS   212      11.364   1.551   6.027 50.00  0.00           C
ATOM   1651  CB  LYS   212      -4.649 -14.150  31.173  1.00  0.00           C
ATOM   1652  CG  LYS   212      -5.916 -13.975  30.333  1.00  0.00           C
ATOM   1653  CD  LYS   212      -6.700 -15.272  30.130  1.00  0.00           C
ATOM   1654  CE  LYS   212      -7.380 -15.783  31.402  1.00  0.00           C
ATOM   1655  NZ  LYS   212      -8.537 -14.923  31.738  1.00  0.00           N
ATOM   1656  C   LYS   212      -2.804 -13.034  32.424  1.00  0.00           C
ATOM   1657  O   LYS   212      -2.212 -14.089  32.595  1.00  0.00           O
ATOM   1658  N   THR   213      -2.583 -11.957  33.167  1.00  0.00           N
ATOM   1659  CA  THR   213      12.290   2.842   5.700 50.00  0.00           C
ATOM   1660  CB  THR   213      -1.176 -10.392  34.426  1.00  0.00           C
ATOM   1661  OG1 THR   213      -0.841  -9.659  33.257  1.00  0.00           O
ATOM   1662  CG2 THR   213      -0.012 -10.312  35.428  1.00  0.00           C
ATOM   1663  C   THR   213      -1.643 -12.756  35.301  1.00  0.00           C
ATOM   1664  O   THR   213      -0.866 -13.682  35.570  1.00  0.00           O
ATOM   1665  N   GLU   214      -2.703 -12.464  36.037  1.00  0.00           N
ATOM   1666  CA  GLU   214      13.108   1.772   6.378 50.00  0.00           C
ATOM   1667  CB  GLU   214      -4.438 -12.809  37.725  1.00  0.00           C
ATOM   1668  CG  GLU   214      -4.860 -13.508  39.019  1.00  0.00           C
ATOM   1669  CD  GLU   214      -6.248 -13.009  39.392  1.00  0.00           C
ATOM   1670  OE1 GLU   214      -6.782 -12.137  38.655  1.00  0.00           O
ATOM   1671  OE2 GLU   214      -6.795 -13.494  40.419  1.00  0.00           O
ATOM   1672  C   GLU   214      -2.954 -14.710  37.097  1.00  0.00           C
ATOM   1673  O   GLU   214      -2.237 -15.402  37.842  1.00  0.00           O
ATOM   1674  N   ASP   215      -3.687 -15.219  36.112  1.00  0.00           N
ATOM   1675  CA  ASP   215      12.755  -1.797   6.176 50.00  0.00           C
ATOM   1676  CB  ASP   215      -4.757 -16.960  34.759  1.00  0.00           C
ATOM   1677  CG  ASP   215      -6.160 -16.735  35.304  1.00  0.00           C
ATOM   1678  OD1 ASP   215      -6.296 -16.597  36.549  1.00  0.00           O
ATOM   1679  OD2 ASP   215      -7.115 -16.698  34.483  1.00  0.00           O
ATOM   1680  C   ASP   215      -2.381 -17.279  35.533  1.00  0.00           C
ATOM   1681  O   ASP   215      -2.075 -18.410  35.919  1.00  0.00           O
ATOM   1682  N   ALA   216      -1.544 -16.515  34.833  1.00  0.00           N
ATOM   1683  CA  ALA   216      12.696  -2.155   5.021 50.00  0.00           C
ATOM   1684  CB  ALA   216       0.448 -16.021  33.511  1.00  0.00           C
ATOM   1685  C   ALA   216       0.637 -17.100  35.740  1.00  0.00           C
ATOM   1686  O   ALA   216       1.392 -18.060  35.897  1.00  0.00           O
ATOM   1687  N   GLU   217       0.501 -16.144  36.643  1.00  0.00           N
ATOM   1688  CA  GLU   217      13.767  -1.107   4.986 50.00  0.00           C
ATOM   1689  CB  GLU   217       1.080 -14.953  38.725  1.00  0.00           C
ATOM   1690  CG  GLU   217       1.803 -13.742  38.133  1.00  0.00           C
ATOM   1691  CD  GLU   217       1.477 -12.531  38.995  1.00  0.00           C
ATOM   1692  OE1 GLU   217       0.665 -12.684  39.946  1.00  0.00           O
ATOM   1693  OE2 GLU   217       2.037 -11.437  38.715  1.00  0.00           O
ATOM   1694  C   GLU   217       0.777 -17.445  38.692  1.00  0.00           C
ATOM   1695  O   GLU   217       1.580 -18.107  39.372  1.00  0.00           O
ATOM   1696  N   GLU   218      -0.491 -17.807  38.584  1.00  0.00           N
ATOM   1697  CA  GLU   218      14.060  -3.784   5.519 50.00  0.00           C
ATOM   1698  CB  GLU   218      -2.461 -19.142  39.300  1.00  0.00           C
ATOM   1699  CG  GLU   218      -3.156 -18.108  40.187  1.00  0.00           C
ATOM   1700  CD  GLU   218      -4.659 -18.259  39.998  1.00  0.00           C
ATOM   1701  OE1 GLU   218      -5.066 -19.095  39.148  1.00  0.00           O
ATOM   1702  OE2 GLU   218      -5.419 -17.540  40.700  1.00  0.00           O
ATOM   1703  C   GLU   218      -0.315 -20.232  38.680  1.00  0.00           C
ATOM   1704  O   GLU   218       0.198 -21.078  39.383  1.00  0.00           O
ATOM   1705  N   VAL   219      -0.344 -20.331  37.365  1.00  0.00           N
ATOM   1706  CA  VAL   219      13.521  -6.392   4.706 50.00  0.00           C
ATOM   1707  CB  VAL   219       0.328 -21.309  35.194  1.00  0.00           C
ATOM   1708  CG1 VAL   219       1.201 -22.359  34.569  1.00  0.00           C
ATOM   1709  CG2 VAL   219      -1.082 -21.447  34.702  1.00  0.00           C
ATOM   1710  C   VAL   219       1.816 -21.559  37.164  1.00  0.00           C
ATOM   1711  O   VAL   219       2.266 -22.660  37.469  1.00  0.00           O
ATOM   1712  N   PHE   220       2.552 -20.451  37.207  1.00  0.00           N
ATOM   1713  CA  PHE   220      14.102  -5.306   3.874 50.00  0.00           C
ATOM   1714  CB  PHE   220       4.560 -19.102  37.743  1.00  0.00           C
ATOM   1715  CG  PHE   220       5.937 -19.246  38.293  1.00  0.00           C
ATOM   1716  CD1 PHE   220       7.011 -19.725  37.510  1.00  0.00           C
ATOM   1717  CD2 PHE   220       6.197 -18.898  39.632  1.00  0.00           C
ATOM   1718  CE1 PHE   220       8.325 -19.856  38.044  1.00  0.00           C
ATOM   1719  CE2 PHE   220       7.505 -19.019  40.192  1.00  0.00           C
ATOM   1720  CZ  PHE   220       8.573 -19.503  39.392  1.00  0.00           C
ATOM   1721  C   PHE   220       3.973 -21.058  39.149  1.00  0.00           C
ATOM   1722  O   PHE   220       4.864 -21.847  39.488  1.00  0.00           O
ATOM   1723  N   LYS   221       3.023 -20.631  39.980  1.00  0.00           N
ATOM   1724  CA  LYS   221      15.058  -5.618   4.437 50.00  0.00           C
ATOM   1725  CB  LYS   221       1.834 -20.383  42.104  1.00  0.00           C
ATOM   1726  CG  LYS   221       2.133 -18.908  42.381  1.00  0.00           C
ATOM   1727  CD  LYS   221       1.015 -18.189  43.138  1.00  0.00           C
ATOM   1728  CE  LYS   221       1.303 -16.708  43.394  1.00  0.00           C
ATOM   1729  NZ  LYS   221       0.160 -16.084  44.098  1.00  0.00           N
ATOM   1730  C   LYS   221       2.741 -22.604  41.407  1.00  0.00           C
ATOM   1731  O   LYS   221       3.392 -23.310  42.191  1.00  0.00           O
ATOM   1732  N   ASP   222       1.827 -23.097  40.573  1.00  0.00           N
ATOM   1733  CA  ASP   222      14.805  -9.128   4.470 50.00  0.00           C
ATOM   1734  CB  ASP   222       0.389 -24.844  39.599  1.00  0.00           C
ATOM   1735  CG  ASP   222      -0.872 -24.446  40.353  1.00  0.00           C
ATOM   1736  OD1 ASP   222      -0.764 -24.156  41.574  1.00  0.00           O
ATOM   1737  OD2 ASP   222      -1.960 -24.425  39.717  1.00  0.00           O
ATOM   1738  C   ASP   222       2.822 -25.311  40.031  1.00  0.00           C
ATOM   1739  O   ASP   222       3.139 -26.340  40.599  1.00  0.00           O
ATOM   1740  N   MET   223       3.828 -24.622  39.505  1.00  0.00           N
ATOM   1741  CA  MET   223      14.750 -10.006   3.289 50.00  0.00           C
ATOM   1742  CB  MET   223       6.279 -24.721  38.475  1.00  0.00           C
ATOM   1743  CG  MET   223       6.040 -24.168  37.069  1.00  0.00           C
ATOM   1744  SD  MET   223       5.546 -25.415  35.842  1.00  0.00           S
ATOM   1745  CE  MET   223       3.767 -25.082  35.985  1.00  0.00           C
ATOM   1746  C   MET   223       5.759 -25.983  40.314  1.00  0.00           C
ATOM   1747  O   MET   223       6.281 -27.098  40.239  1.00  0.00           O
ATOM   1748  N   ARG   224       5.798 -25.274  41.443  1.00  0.00           N
ATOM   1749  CA  ARG   224      15.927 -10.147   3.243 50.00  0.00           C
ATOM   1750  CB  ARG   224       6.318 -25.100  44.019  1.00  0.00           C
ATOM   1751  CG  ARG   224       7.225 -23.869  44.076  1.00  0.00           C
ATOM   1752  CD  ARG   224       7.136 -23.103  45.397  1.00  0.00           C
ATOM   1753  NE  ARG   224       8.026 -21.913  45.286  1.00  0.00           N
ATOM   1754  CZ  ARG   224       8.093 -21.013  46.310  1.00  0.00           C
ATOM   1755  NH1 ARG   224       7.254 -21.442  47.297  1.00  0.00           N
ATOM   1756  NH2 ARG   224       8.974 -20.035  45.947  1.00  0.00           N
ATOM   1757  C   ARG   224       5.625 -27.137  43.068  1.00  0.00           C
ATOM   1758  O   ARG   224       6.227 -28.173  43.360  1.00  0.00           O
ATOM   1759  N   ASN   225       4.290 -27.075  43.071  1.00  0.00           N
ATOM   1760  CA  ASN   225      15.971 -12.510   3.915 50.00  0.00           C
ATOM   1761  CB  ASN   225       1.983 -28.003  43.421  1.00  0.00           C
ATOM   1762  CG  ASN   225       1.641 -27.157  44.639  1.00  0.00           C
ATOM   1763  OD1 ASN   225       2.405 -27.092  45.600  1.00  0.00           O
ATOM   1764  ND2 ASN   225       0.472 -26.462  44.665  1.00  0.00           N
ATOM   1765  C   ASN   225       3.750 -29.388  42.484  1.00  0.00           C
ATOM   1766  O   ASN   225       3.760 -30.557  42.881  1.00  0.00           O
ATOM   1767  N   ALA   226       3.963 -29.095  41.216  1.00  0.00           N
ATOM   1768  CA  ALA   226      15.186 -14.689   3.255 50.00  0.00           C
ATOM   1769  CB  ALA   226       3.811 -29.733  38.859  1.00  0.00           C
ATOM   1770  C   ALA   226       5.615 -30.717  40.181  1.00  0.00           C
ATOM   1771  O   ALA   226       5.875 -31.698  39.480  1.00  0.00           O
ATOM   1772  N   GLY   227       6.571 -30.101  40.890  1.00  0.00           N
ATOM   1773  CA  GLY   227      15.698 -14.994   2.229 50.00  0.00           C
ATOM   1774  C   GLY   227       8.758 -29.973  39.775  1.00  0.00           C
ATOM   1775  O   GLY   227       9.941 -30.272  39.591  1.00  0.00           O
ATOM   1776  N   VAL   228       8.099 -29.043  39.012  1.00  0.00           N
ATOM   1777  CA  VAL   228      14.821 -13.034   1.858 50.00  0.00           C
ATOM   1778  CB  VAL   228       7.836 -27.874  36.855  1.00  0.00           C
ATOM   1779  CG1 VAL   228       8.507 -27.078  35.733  1.00  0.00           C
ATOM   1780  CG2 VAL   228       7.089 -29.008  36.150  1.00  0.00           C
ATOM   1781  C   VAL   228       9.726 -27.215  38.413  1.00  0.00           C
ATOM   1782  O   VAL   228      10.957 -27.319  38.312  1.00  0.00           O
ATOM   1783  N   GLU   229       9.150 -26.104  38.857  1.00  0.00           N
ATOM   1784  CA  GLU   229      15.240  -9.541   1.966 50.00  0.00           C
ATOM   1785  CB  GLU   229      11.058 -25.471  40.316  1.00  0.00           C
ATOM   1786  CG  GLU   229      10.565 -26.160  41.590  1.00  0.00           C
ATOM   1787  CD  GLU   229      11.783 -26.646  42.363  1.00  0.00           C
ATOM   1788  OE1 GLU   229      12.920 -26.453  41.857  1.00  0.00           O
ATOM   1789  OE2 GLU   229      11.592 -27.215  43.471  1.00  0.00           O
ATOM   1790  C   GLU   229      10.507 -23.980  38.418  1.00  0.00           C
ATOM   1791  O   GLU   229      11.620 -24.126  37.934  1.00  0.00           O
ATOM   1792  N   PRO   230       9.756 -22.932  38.152  1.00  0.00           N
ATOM   1793  CA  PRO   230      14.588  -6.677   1.915 50.00  0.00           C
ATOM   1794  CD  PRO   230       8.485 -22.562  38.802  1.00  0.00           C
ATOM   1795  CB  PRO   230       9.289 -20.734  37.478  1.00  0.00           C
ATOM   1796  CG  PRO   230       8.057 -21.316  38.050  1.00  0.00           C
ATOM   1797  C   PRO   230      11.598 -21.638  37.034  1.00  0.00           C
ATOM   1798  O   PRO   230      12.293 -21.627  38.038  1.00  0.00           O
ATOM   1799  N   VAL   231      12.057 -21.376  35.820  1.00  0.00           N
ATOM   1800  CA  VAL   231      14.399  -5.989   0.801 50.00  0.00           C
ATOM   1801  CB  VAL   231      13.791 -22.020  34.291  1.00  0.00           C
ATOM   1802  CG1 VAL   231      13.571 -23.520  34.497  1.00  0.00           C
ATOM   1803  CG2 VAL   231      12.956 -21.659  33.061  1.00  0.00           C
ATOM   1804  C   VAL   231      13.734 -19.728  35.178  1.00  0.00           C
ATOM   1805  O   VAL   231      12.883 -18.865  35.344  1.00  0.00           O
ATOM   1806  N   SER   232      14.936 -19.463  34.690  1.00  0.00           N
ATOM   1807  CA  SER   232      13.954  -3.199   0.542 50.00  0.00           C
ATOM   1808  CB  SER   232      16.707 -17.953  33.961  1.00  0.00           C
ATOM   1809  OG  SER   232      17.332 -18.932  33.145  1.00  0.00           O
ATOM   1810  C   SER   232      14.638 -18.381  32.638  1.00  0.00           C
ATOM   1811  O   SER   232      14.225 -17.427  31.983  1.00  0.00           O
ATOM   1812  N   THR   233      14.605 -19.622  32.192  1.00  0.00           N
ATOM   1813  CA  THR   233      13.000  -5.186   0.447 50.00  0.00           C
ATOM   1814  CB  THR   233      14.411 -21.373  30.458  1.00  0.00           C
ATOM   1815  OG1 THR   233      15.817 -21.562  30.400  1.00  0.00           O
ATOM   1816  CG2 THR   233      13.803 -21.652  29.073  1.00  0.00           C
ATOM   1817  C   THR   233      12.585 -19.682  30.853  1.00  0.00           C
ATOM   1818  O   THR   233      12.055 -19.083  29.911  1.00  0.00           O
ATOM   1819  N   HIS   234      11.906 -20.130  31.903  1.00  0.00           N
ATOM   1820  CA  HIS   234      13.025  -5.198   1.501 50.00  0.00           C
ATOM   1821  ND1 HIS   234       7.481 -21.179  32.582  1.00  0.00           N
ATOM   1822  CG  HIS   234       8.400 -20.485  33.342  1.00  0.00           C
ATOM   1823  CB  HIS   234       9.886 -20.651  33.221  1.00  0.00           C
ATOM   1824  NE2 HIS   234       6.340 -19.930  33.857  1.00  0.00           N
ATOM   1825  CD2 HIS   234       7.660 -19.690  34.151  1.00  0.00           C
ATOM   1826  CE1 HIS   234       6.270 -20.838  32.903  1.00  0.00           C
ATOM   1827  C   HIS   234      10.028 -18.536  31.910  1.00  0.00           C
ATOM   1828  O   HIS   234       9.201 -18.160  31.068  1.00  0.00           O
ATOM   1829  N   PRO   235      10.607 -17.722  32.799  1.00  0.00           N
ATOM   1830  CA  PRO   235      13.155  -1.449   1.921 50.00  0.00           C
ATOM   1831  CD  PRO   235      11.654 -17.998  34.789  1.00  0.00           C
ATOM   1832  CB  PRO   235      11.029 -15.628  34.048  1.00  0.00           C
ATOM   1833  CG  PRO   235      11.273 -16.581  35.220  1.00  0.00           C
ATOM   1834  C   PRO   235      10.716 -15.565  31.605  1.00  0.00           C
ATOM   1835  O   PRO   235      10.011 -14.658  31.173  1.00  0.00           O
ATOM   1836  N   TYR   236      11.837 -15.947  31.015  1.00  0.00           N
ATOM   1837  CA  TYR   236      12.393  -0.729   1.163 50.00  0.00           C
ATOM   1838  CB  TYR   236      13.624 -15.734  29.335  1.00  0.00           C
ATOM   1839  CG  TYR   236      13.908 -15.057  28.038  1.00  0.00           C
ATOM   1840  CD1 TYR   236      14.275 -13.705  28.024  1.00  0.00           C
ATOM   1841  CD2 TYR   236      13.823 -15.748  26.809  1.00  0.00           C
ATOM   1842  CE1 TYR   236      14.555 -13.031  26.818  1.00  0.00           C
ATOM   1843  CE2 TYR   236      14.104 -15.078  25.575  1.00  0.00           C
ATOM   1844  CZ  TYR   236      14.469 -13.717  25.602  1.00  0.00           C
ATOM   1845  OH  TYR   236      14.744 -13.029  24.439  1.00  0.00           O
ATOM   1846  C   TYR   236      11.210 -15.574  28.676  1.00  0.00           C
ATOM   1847  O   TYR   236      10.851 -14.664  27.926  1.00  0.00           O
ATOM   1848  N   ASN   237      10.730 -16.807  28.584  1.00  0.00           N
ATOM   1849  CA  ASN   237      11.662  -2.727   1.529 50.00  0.00           C
ATOM   1850  CB  ASN   237       9.583 -18.628  27.398  1.00  0.00           C
ATOM   1851  CG  ASN   237      10.823 -19.154  26.689  1.00  0.00           C
ATOM   1852  OD1 ASN   237      11.539 -18.405  26.026  1.00  0.00           O
ATOM   1853  ND2 ASN   237      11.144 -20.471  26.790  1.00  0.00           N
ATOM   1854  C   ASN   237       8.391 -16.442  27.844  1.00  0.00           C
ATOM   1855  O   ASN   237       7.693 -16.060  26.908  1.00  0.00           O
ATOM   1856  N   PHE   238       8.049 -16.270  29.122  1.00  0.00           N
ATOM   1857  CA  PHE   238      11.891  -1.109   2.636 50.00  0.00           C
ATOM   1858  CB  PHE   238       6.604 -15.671  31.025  1.00  0.00           C
ATOM   1859  CG  PHE   238       5.364 -14.908  31.342  1.00  0.00           C
ATOM   1860  CD1 PHE   238       4.076 -15.443  31.114  1.00  0.00           C
ATOM   1861  CD2 PHE   238       5.457 -13.614  31.886  1.00  0.00           C
ATOM   1862  CE1 PHE   238       2.896 -14.707  31.424  1.00  0.00           C
ATOM   1863  CE2 PHE   238       4.291 -12.855  32.205  1.00  0.00           C
ATOM   1864  CZ  PHE   238       3.004 -13.405  31.970  1.00  0.00           C
ATOM   1865  C   PHE   238       6.818 -14.129  29.127  1.00  0.00           C
ATOM   1866  O   PHE   238       5.930 -13.644  28.401  1.00  0.00           O
ATOM   1867  N   ILE   239       7.793 -13.414  29.680  1.00  0.00           N
ATOM   1868  CA  ILE   239      11.637   2.370   2.565 50.00  0.00           C
ATOM   1869  CB  ILE   239       9.297 -11.535  29.927  1.00  0.00           C
ATOM   1870  CG2 ILE   239       9.666 -10.109  29.485  1.00  0.00           C
ATOM   1871  CG1 ILE   239       9.341 -11.504  31.464  1.00  0.00           C
ATOM   1872  CD1 ILE   239      10.744 -11.288  32.029  1.00  0.00           C
ATOM   1873  C   ILE   239       7.939 -11.878  27.857  1.00  0.00           C
ATOM   1874  O   ILE   239       7.183 -11.070  27.312  1.00  0.00           O
ATOM   1875  N   PHE   240       8.720 -12.712  27.184  1.00  0.00           N
ATOM   1876  CA  PHE   240      10.803   1.472   2.018 50.00  0.00           C
ATOM   1877  CB  PHE   240       9.799 -13.725  25.255  1.00  0.00           C
ATOM   1878  CG  PHE   240       9.842 -13.651  23.767  1.00  0.00           C
ATOM   1879  CD1 PHE   240      10.574 -12.651  23.088  1.00  0.00           C
ATOM   1880  CD2 PHE   240       9.139 -14.595  22.998  1.00  0.00           C
ATOM   1881  CE1 PHE   240      10.613 -12.590  21.665  1.00  0.00           C
ATOM   1882  CE2 PHE   240       9.161 -14.557  21.570  1.00  0.00           C
ATOM   1883  CZ  PHE   240       9.901 -13.548  20.903  1.00  0.00           C
ATOM   1884  C   PHE   240       7.485 -12.875  25.029  1.00  0.00           C
ATOM   1885  O   PHE   240       7.221 -12.275  24.001  1.00  0.00           O
ATOM   1886  N   MET   241       6.651 -13.739  25.578  1.00  0.00           N
ATOM   1887  CA  MET   241      10.416  -0.030   2.740 50.00  0.00           C
ATOM   1888  CB  MET   241       4.895 -15.443  25.431  1.00  0.00           C
ATOM   1889  CG  MET   241       5.778 -16.629  25.039  1.00  0.00           C
ATOM   1890  SD  MET   241       5.963 -16.872  23.247  1.00  0.00           S
ATOM   1891  CE  MET   241       4.240 -17.379  22.978  1.00  0.00           C
ATOM   1892  C   MET   241       4.383 -12.993  25.093  1.00  0.00           C
ATOM   1893  O   MET   241       3.652 -12.670  24.154  1.00  0.00           O
ATOM   1894  N   LYS   242       4.336 -12.413  26.292  1.00  0.00           N
ATOM   1895  CA  LYS   242      10.598   2.932   3.644 50.00  0.00           C
ATOM   1896  CB  LYS   242       3.793 -10.693  27.939  1.00  0.00           C
ATOM   1897  CG  LYS   242       3.358 -11.652  29.049  1.00  0.00           C
ATOM   1898  CD  LYS   242       1.844 -11.864  29.117  1.00  0.00           C
ATOM   1899  CE  LYS   242       1.407 -12.804  30.242  1.00  0.00           C
ATOM   1900  NZ  LYS   242      -0.056 -13.021  30.180  1.00  0.00           N
ATOM   1901  C   LYS   242       3.936 -10.141  25.544  1.00  0.00           C
ATOM   1902  O   LYS   242       3.121  -9.648  24.767  1.00  0.00           O
ATOM   1903  N   THR   243       5.213  -9.779  25.573  1.00  0.00           N
ATOM   1904  CA  THR   243      10.164   5.218   3.042 50.00  0.00           C
ATOM   1905  CB  THR   243       7.213  -8.582  24.914  1.00  0.00           C
ATOM   1906  OG1 THR   243       7.451  -8.169  26.252  1.00  0.00           O
ATOM   1907  CG2 THR   243       7.758  -7.517  23.948  1.00  0.00           C
ATOM   1908  C   THR   243       5.440  -8.987  23.200  1.00  0.00           C
ATOM   1909  O   THR   243       4.979  -8.086  22.521  1.00  0.00           O
ATOM   1910  N   PRO   244       5.674 -10.179  22.684  1.00  0.00           N
ATOM   1911  CA  PRO   244       9.231   3.243   2.706 50.00  0.00           C
ATOM   1912  CD  PRO   244       7.015 -11.849  23.119  1.00  0.00           C
ATOM   1913  CB  PRO   244       5.891 -11.785  20.819  1.00  0.00           C
ATOM   1914  CG  PRO   244       7.088 -12.257  21.646  1.00  0.00           C
ATOM   1915  C   PRO   244       3.791 -10.433  21.050  1.00  0.00           C
ATOM   1916  O   PRO   244       3.358 -10.279  19.911  1.00  0.00           O
ATOM   1917  N   PRO   245       2.987 -10.608  22.101  1.00  0.00           N
ATOM   1918  CA  PRO   245       9.078   3.102   3.822 50.00  0.00           C
ATOM   1919  CD  PRO   245       3.140 -11.430  24.181  1.00  0.00           C
ATOM   1920  CB  PRO   245       0.876 -11.463  22.986  1.00  0.00           C
ATOM   1921  CG  PRO   245       1.618 -11.472  24.324  1.00  0.00           C
ATOM   1922  C   PRO   245       0.987  -9.194  22.027  1.00  0.00           C
ATOM   1923  O   PRO   245      -0.177  -8.955  21.724  1.00  0.00           O
ATOM   1924  N   PHE   246       1.827  -8.263  22.456  1.00  0.00           N
ATOM   1925  CA  PHE   246       9.159   6.865   4.165 50.00  0.00           C
ATOM   1926  CB  PHE   246       0.110  -6.673  21.770  1.00  0.00           C
ATOM   1927  CG  PHE   246       0.366  -6.850  20.313  1.00  0.00           C
ATOM   1928  CD1 PHE   246       0.067  -8.057  19.641  1.00  0.00           C
ATOM   1929  CD2 PHE   246       0.920  -5.795  19.565  1.00  0.00           C
ATOM   1930  CE1 PHE   246       0.311  -8.216  18.246  1.00  0.00           C
ATOM   1931  CE2 PHE   246       1.175  -5.928  18.166  1.00  0.00           C
ATOM   1932  CZ  PHE   246       0.870  -7.146  17.506  1.00  0.00           C
ATOM   1933  C   PHE   246       1.333  -6.339  23.907  1.00  0.00           C
ATOM   1934  O   PHE   246       1.133  -5.146  24.091  1.00  0.00           O
ATOM   1935  N   LYS   247       1.459  -7.214  24.900  1.00  0.00           N
ATOM   1936  CA  LYS   247      10.264   7.290   4.612 50.00  0.00           C
ATOM   1937  CB  LYS   247       0.419  -7.863  27.031  1.00  0.00           C
ATOM   1938  CG  LYS   247       0.032  -7.442  28.450  1.00  0.00           C
ATOM   1939  CD  LYS   247      -0.869  -6.207  28.498  1.00  0.00           C
ATOM   1940  CE  LYS   247      -2.275  -6.452  27.947  1.00  0.00           C
ATOM   1941  NZ  LYS   247      -3.069  -5.205  28.011  1.00  0.00           N
ATOM   1942  C   LYS   247       2.564  -6.520  27.012  1.00  0.00           C
ATOM   1943  O   LYS   247       3.100  -7.333  27.760  1.00  0.00           O
ATOM   1944  N   TYR   248       3.053  -5.313  26.767  1.00  0.00           N
ATOM   1945  CA  TYR   248      10.659   9.377   4.010 50.00  0.00           C
ATOM   1946  CB  TYR   248       4.707  -3.582  26.536  1.00  0.00           C
ATOM   1947  CG  TYR   248       3.819  -2.496  27.039  1.00  0.00           C
ATOM   1948  CD1 TYR   248       4.192  -1.758  28.169  1.00  0.00           C
ATOM   1949  CD2 TYR   248       2.600  -2.182  26.398  1.00  0.00           C
ATOM   1950  CE1 TYR   248       3.380  -0.722  28.673  1.00  0.00           C
ATOM   1951  CE2 TYR   248       1.763  -1.132  26.897  1.00  0.00           C
ATOM   1952  CZ  TYR   248       2.172  -0.413  28.038  1.00  0.00           C
ATOM   1953  OH  TYR   248       1.398   0.602  28.556  1.00  0.00           O
ATOM   1954  C   TYR   248       4.300  -4.576  28.744  1.00  0.00           C
ATOM   1955  O   TYR   248       5.294  -4.752  29.421  1.00  0.00           O
ATOM   1956  N   MET   249       3.179  -4.109  29.257  1.00  0.00           N
ATOM   1957  CA  MET   249      11.472  10.798   4.701 50.00  0.00           C
ATOM   1958  CB  MET   249       1.842  -2.984  30.973  1.00  0.00           C
ATOM   1959  CG  MET   249       1.795  -2.414  32.392  1.00  0.00           C
ATOM   1960  SD  MET   249       3.024  -1.117  32.725  1.00  0.00           S
ATOM   1961  CE  MET   249       2.196   0.148  31.721  1.00  0.00           C
ATOM   1962  C   MET   249       3.350  -4.934  31.458  1.00  0.00           C
ATOM   1963  O   MET   249       4.090  -4.958  32.441  1.00  0.00           O
ATOM   1964  N   ASP   250       2.706  -5.995  30.990  1.00  0.00           N
ATOM   1965  CA  ASP   250      11.900   7.285   4.667 50.00  0.00           C
ATOM   1966  CB  ASP   250       1.896  -8.278  30.848  1.00  0.00           C
ATOM   1967  CG  ASP   250       0.467  -7.942  31.250  1.00  0.00           C
ATOM   1968  OD1 ASP   250       0.295  -7.137  32.204  1.00  0.00           O
ATOM   1969  OD2 ASP   250      -0.471  -8.485  30.609  1.00  0.00           O
ATOM   1970  C   ASP   250       4.265  -7.816  31.513  1.00  0.00           C
ATOM   1971  O   ASP   250       4.869  -8.129  32.537  1.00  0.00           O
ATOM   1972  N   LEU   251       4.799  -7.883  30.300  1.00  0.00           N
ATOM   1973  CA  LEU   251      11.802   6.229   3.457 50.00  0.00           C
ATOM   1974  CB  LEU   251       6.525  -7.978  28.606  1.00  0.00           C
ATOM   1975  CG  LEU   251       7.972  -8.319  28.244  1.00  0.00           C
ATOM   1976  CD1 LEU   251       8.344  -9.794  28.392  1.00  0.00           C
ATOM   1977  CD2 LEU   251       8.371  -7.996  26.805  1.00  0.00           C
ATOM   1978  C   LEU   251       7.180  -7.530  31.001  1.00  0.00           C
ATOM   1979  O   LEU   251       8.055  -8.157  31.601  1.00  0.00           O
ATOM   1980  N   LEU   252       7.032  -6.217  31.135  1.00  0.00           N
ATOM   1981  CA  LEU   252      12.330   9.235   3.429 50.00  0.00           C
ATOM   1982  CB  LEU   252       7.620  -3.953  31.772  1.00  0.00           C
ATOM   1983  CG  LEU   252       8.089  -3.469  30.399  1.00  0.00           C
ATOM   1984  CD1 LEU   252       7.712  -2.027  30.059  1.00  0.00           C
ATOM   1985  CD2 LEU   252       9.602  -3.499  30.185  1.00  0.00           C
ATOM   1986  C   LEU   252       7.649  -5.760  33.477  1.00  0.00           C
ATOM   1987  O   LEU   252       8.573  -5.702  34.281  1.00  0.00           O
ATOM   1988  N   ASP   253       6.416  -6.107  33.828  1.00  0.00           N
ATOM   1989  CA  ASP   253      13.094   8.314   4.103 50.00  0.00           C
ATOM   1990  CB  ASP   253       4.633  -6.796  35.389  1.00  0.00           C
ATOM   1991  CG  ASP   253       3.956  -5.435  35.461  1.00  0.00           C
ATOM   1992  OD1 ASP   253       4.687  -4.417  35.592  1.00  0.00           O
ATOM   1993  OD2 ASP   253       2.699  -5.395  35.385  1.00  0.00           O
ATOM   1994  C   ASP   253       6.929  -7.850  35.437  1.00  0.00           C
ATOM   1995  O   ASP   253       7.492  -8.060  36.519  1.00  0.00           O
ATOM   1996  N   TYR   254       6.994  -8.686  34.407  1.00  0.00           N
ATOM   1997  CA  TYR   254      13.172   4.931   3.288 50.00  0.00           C
ATOM   1998  CB  TYR   254       7.563 -10.729  33.190  1.00  0.00           C
ATOM   1999  CG  TYR   254       6.165 -11.243  33.175  1.00  0.00           C
ATOM   2000  CD1 TYR   254       5.186 -10.580  32.422  1.00  0.00           C
ATOM   2001  CD2 TYR   254       5.793 -12.396  33.901  1.00  0.00           C
ATOM   2002  CE1 TYR   254       3.854 -11.038  32.380  1.00  0.00           C
ATOM   2003  CE2 TYR   254       4.443 -12.875  33.869  1.00  0.00           C
ATOM   2004  CZ  TYR   254       3.487 -12.179  33.101  1.00  0.00           C
ATOM   2005  OH  TYR   254       2.176 -12.600  33.044  1.00  0.00           O
ATOM   2006  C   TYR   254       9.211  -9.677  34.710  1.00  0.00           C
ATOM   2007  O   TYR   254       9.771 -10.221  35.656  1.00  0.00           O
ATOM   2008  N   TYR   255       9.835  -8.875  33.849  1.00  0.00           N
ATOM   2009  CA  TYR   255      13.318   6.430   2.405 50.00  0.00           C
ATOM   2010  CB  TYR   255      11.660  -7.470  32.994  1.00  0.00           C
ATOM   2011  CG  TYR   255      13.096  -7.154  33.236  1.00  0.00           C
ATOM   2012  CD1 TYR   255      14.088  -8.043  32.801  1.00  0.00           C
ATOM   2013  CD2 TYR   255      13.492  -5.968  33.892  1.00  0.00           C
ATOM   2014  CE1 TYR   255      15.456  -7.778  33.009  1.00  0.00           C
ATOM   2015  CE2 TYR   255      14.878  -5.682  34.111  1.00  0.00           C
ATOM   2016  CZ  TYR   255      15.846  -6.603  33.661  1.00  0.00           C
ATOM   2017  OH  TYR   255      17.192  -6.373  33.853  1.00  0.00           O
ATOM   2018  C   TYR   255      11.490  -7.941  35.415  1.00  0.00           C
ATOM   2019  O   TYR   255      12.556  -8.148  35.993  1.00  0.00           O
ATOM   2020  N   GLU   256      10.492  -7.241  35.939  1.00  0.00           N
ATOM   2021  CA  GLU   256      14.095   8.475   3.014 50.00  0.00           C
ATOM   2022  CB  GLU   256       9.262  -5.926  37.601  1.00  0.00           C
ATOM   2023  CG  GLU   256       9.252  -5.294  38.994  1.00  0.00           C
ATOM   2024  CD  GLU   256       7.949  -4.522  39.151  1.00  0.00           C
ATOM   2025  OE1 GLU   256       7.148  -4.510  38.179  1.00  0.00           O
ATOM   2026  OE2 GLU   256       7.738  -3.933  40.245  1.00  0.00           O
ATOM   2027  C   GLU   256      10.712  -7.858  38.283  1.00  0.00           C
ATOM   2028  O   GLU   256      11.625  -7.839  39.115  1.00  0.00           O
ATOM   2029  N   ASP   257       9.828  -8.849  38.187  1.00  0.00           N
ATOM   2030  CA  ASP   257      14.655   5.329   3.150 50.00  0.00           C
ATOM   2031  CB  ASP   257       8.753 -11.010  38.981  1.00  0.00           C
ATOM   2032  CG  ASP   257       7.510 -10.383  39.595  1.00  0.00           C
ATOM   2033  OD1 ASP   257       7.652  -9.331  40.275  1.00  0.00           O
ATOM   2034  OD2 ASP   257       6.401 -10.946  39.392  1.00  0.00           O
ATOM   2035  C   ASP   257      11.144 -10.838  38.854  1.00  0.00           C
ATOM   2036  O   ASP   257      11.724 -11.392  39.792  1.00  0.00           O
ATOM   2037  N   MET   258      11.585 -10.952  37.605  1.00  0.00           N
ATOM   2038  CA  MET   258      14.482   3.535   2.019 50.00  0.00           C
ATOM   2039  CB  MET   258      13.452 -11.892  35.956  1.00  0.00           C
ATOM   2040  CG  MET   258      12.668 -12.832  35.038  1.00  0.00           C
ATOM   2041  SD  MET   258      13.533 -13.284  33.505  1.00  0.00           S
ATOM   2042  CE  MET   258      13.298 -11.669  32.708  1.00  0.00           C
ATOM   2043  C   MET   258      13.997 -11.010  38.001  1.00  0.00           C
ATOM   2044  O   MET   258      14.836 -11.643  38.647  1.00  0.00           O
ATOM   2045  N   LEU   259      14.082  -9.693  37.853  1.00  0.00           N
ATOM   2046  CA  LEU   259      14.923   6.405   1.670 50.00  0.00           C
ATOM   2047  CB  LEU   259      15.299  -7.428  38.316  1.00  0.00           C
ATOM   2048  CG  LEU   259      16.529  -6.842  39.012  1.00  0.00           C
ATOM   2049  CD1 LEU   259      17.871  -7.352  38.488  1.00  0.00           C
ATOM   2050  CD2 LEU   259      16.664  -5.323  38.915  1.00  0.00           C
ATOM   2051  C   LEU   259      15.112  -9.063  39.979  1.00  0.00           C
ATOM   2052  O   LEU   259      16.116  -9.275  40.664  1.00  0.00           O
ATOM   2053  N   ASP   260      13.911  -8.922  40.522  1.00  0.00           N
ATOM   2054  CA  ASP   260      15.792   6.636   2.413 50.00  0.00           C
ATOM   2055  CB  ASP   260      12.320  -8.736  42.498  1.00  0.00           C
ATOM   2056  CG  ASP   260      12.046  -7.244  42.377  1.00  0.00           C
ATOM   2057  OD1 ASP   260      13.018  -6.483  42.121  1.00  0.00           O
ATOM   2058  OD2 ASP   260      10.862  -6.844  42.539  1.00  0.00           O
ATOM   2059  C   ASP   260      14.062 -10.397  42.494  1.00  0.00           C
ATOM   2060  O   ASP   260      14.472 -10.577  43.644  1.00  0.00           O
ATOM   2061  N   ARG   261      13.885 -11.406  41.646  1.00  0.00           N
ATOM   2062  CA  ARG   261      15.952   2.886   1.998 50.00  0.00           C
ATOM   2063  CB  ARG   261      13.338 -13.946  41.411  1.00  0.00           C
ATOM   2064  CG  ARG   261      11.861 -13.933  41.809  1.00  0.00           C
ATOM   2065  CD  ARG   261      11.063 -15.109  41.241  1.00  0.00           C
ATOM   2066  NE  ARG   261       9.626 -14.868  41.553  1.00  0.00           N
ATOM   2067  CZ  ARG   261       9.121 -15.237  42.766  1.00  0.00           C
ATOM   2068  NH1 ARG   261      10.136 -15.795  43.488  1.00  0.00           N
ATOM   2069  NH2 ARG   261       7.797 -14.904  42.784  1.00  0.00           N
ATOM   2070  C   ARG   261      15.543 -13.280  41.645  1.00  0.00           C
ATOM   2071  O   ARG   261      15.903 -14.439  41.867  1.00  0.00           O
ATOM   2072  N   GLY   262      16.352 -12.399  41.072  1.00  0.00           N
ATOM   2073  CA  GLY   262      15.885   2.817   0.864 50.00  0.00           C
ATOM   2074  C   GLY   262      17.748 -13.698  39.465  1.00  0.00           C
ATOM   2075  O   GLY   262      18.714 -14.431  39.237  1.00  0.00           O
ATOM   2076  N   ILE   263      16.695 -13.651  38.651  1.00  0.00           N
ATOM   2077  CA  ILE   263      14.909   0.868   0.707 50.00  0.00           C
ATOM   2078  CB  ILE   263      15.411 -14.914  36.567  1.00  0.00           C
ATOM   2079  CG2 ILE   263      15.818 -15.769  35.355  1.00  0.00           C
ATOM   2080  CG1 ILE   263      14.414 -15.781  37.355  1.00  0.00           C
ATOM   2081  CD1 ILE   263      15.031 -17.060  37.918  1.00  0.00           C
ATOM   2082  C   ILE   263      17.164 -13.540  36.312  1.00  0.00           C
ATOM   2083  O   ILE   263      16.717 -12.406  36.122  1.00  0.00           O
ATOM   2084  N   VAL   264      18.074 -13.937  35.574  1.00  0.00           N
ATOM   2085  CA  VAL   264      14.267   1.863  -0.077 50.00  0.00           C
ATOM   2086  CB  VAL   264      20.120 -13.943  34.207  1.00  0.00           C
ATOM   2087  CG1 VAL   264      20.896 -13.317  33.046  1.00  0.00           C
ATOM   2088  CG2 VAL   264      21.090 -13.930  35.390  1.00  0.00           C
ATOM   2089  C   VAL   264      17.973 -13.164  33.220  1.00  0.00           C
ATOM   2090  O   VAL   264      17.666 -14.188  32.619  1.00  0.00           O
ATOM   2091  N   PRO   265      17.605 -11.949  32.798  1.00  0.00           N
ATOM   2092  CA  PRO   265      13.224   3.018   0.269 50.00  0.00           C
ATOM   2093  CD  PRO   265      17.489 -10.723  33.599  1.00  0.00           C
ATOM   2094  CB  PRO   265      16.437 -10.371  31.492  1.00  0.00           C
ATOM   2095  CG  PRO   265      17.193  -9.686  32.572  1.00  0.00           C
ATOM   2096  C   PRO   265      17.865 -12.090  30.332  1.00  0.00           C
ATOM   2097  O   PRO   265      19.075 -11.949  30.482  1.00  0.00           O
ATOM   2098  N   MET   266      17.303 -12.480  29.192  1.00  0.00           N
ATOM   2099  CA  MET   266      12.307   1.869  -0.520 50.00  0.00           C
ATOM   2100  CB  MET   266      17.209 -13.425  26.953  1.00  0.00           C
ATOM   2101  CG  MET   266      16.812 -14.843  27.369  1.00  0.00           C
ATOM   2102  SD  MET   266      18.212 -15.980  27.600  1.00  0.00           S
ATOM   2103  CE  MET   266      18.639 -16.066  25.837  1.00  0.00           C
ATOM   2104  C   MET   266      18.599 -11.433  27.352  1.00  0.00           C
ATOM   2105  O   MET   266      18.854 -10.444  28.027  1.00  0.00           O
ATOM   2106  N   ASN   267      18.792 -11.474  26.044  1.00  0.00           N
ATOM   2107  CA  ASN   267      11.572   4.078  -0.774 50.00  0.00           C
ATOM   2108  CB  ASN   267      20.296 -10.388  24.424  1.00  0.00           C
ATOM   2109  CG  ASN   267      21.485 -10.537  25.363  1.00  0.00           C
ATOM   2110  OD1 ASN   267      21.485 -10.007  26.473  1.00  0.00           O
ATOM   2111  ND2 ASN   267      22.563 -11.266  24.968  1.00  0.00           N
ATOM   2112  C   ASN   267      17.798 -10.151  24.408  1.00  0.00           C
ATOM   2113  O   ASN   267      17.180  -9.091  24.309  1.00  0.00           O
ATOM   2114  N   SER   268      17.452 -11.276  23.796  1.00  0.00           N
ATOM   2115  CA  SER   268      10.715   2.686  -0.327 50.00  0.00           C
ATOM   2116  CB  SER   268      16.188 -12.820  22.416  1.00  0.00           C
ATOM   2117  OG  SER   268      17.297 -13.144  21.591  1.00  0.00           O
ATOM   2118  C   SER   268      15.011 -10.973  23.596  1.00  0.00           C
ATOM   2119  O   SER   268      14.115 -10.403  22.956  1.00  0.00           O
ATOM   2120  N   THR   269      14.888 -11.262  24.887  1.00  0.00           N
ATOM   2121  CA  THR   269      11.215   3.385   0.679 50.00  0.00           C
ATOM   2122  CB  THR   269      13.419 -11.702  26.867  1.00  0.00           C
ATOM   2123  OG1 THR   269      14.168 -11.150  27.954  1.00  0.00           O
ATOM   2124  CG2 THR   269      13.778 -13.176  26.654  1.00  0.00           C
ATOM   2125  C   THR   269      13.657  -9.371  25.819  1.00  0.00           C
ATOM   2126  O   THR   269      12.646  -8.708  25.584  1.00  0.00           O
ATOM   2127  N   PHE   270      14.796  -8.837  26.262  1.00  0.00           N
ATOM   2128  CA  PHE   270      11.455   6.939   0.692 50.00  0.00           C
ATOM   2129  CB  PHE   270      16.463  -7.086  26.721  1.00  0.00           C
ATOM   2130  CG  PHE   270      16.718  -7.528  28.121  1.00  0.00           C
ATOM   2131  CD1 PHE   270      15.666  -7.847  29.009  1.00  0.00           C
ATOM   2132  CD2 PHE   270      18.039  -7.640  28.592  1.00  0.00           C
ATOM   2133  CE1 PHE   270      15.919  -8.273  30.345  1.00  0.00           C
ATOM   2134  CE2 PHE   270      18.321  -8.064  29.926  1.00  0.00           C
ATOM   2135  CZ  PHE   270      17.254  -8.380  30.806  1.00  0.00           C
ATOM   2136  C   PHE   270      14.589  -6.695  25.110  1.00  0.00           C
ATOM   2137  O   PHE   270      13.872  -5.696  25.124  1.00  0.00           O
ATOM   2138  N   ARG   271      15.033  -7.250  23.992  1.00  0.00           N
ATOM   2139  CA  ARG   271      10.349   7.355   0.460 50.00  0.00           C
ATOM   2140  CB  ARG   271      15.587  -7.336  21.604  1.00  0.00           C
ATOM   2141  CG  ARG   271      17.082  -7.018  21.679  1.00  0.00           C
ATOM   2142  CD  ARG   271      17.919  -7.760  20.635  1.00  0.00           C
ATOM   2143  NE  ARG   271      19.340  -7.360  20.835  1.00  0.00           N
ATOM   2144  CZ  ARG   271      20.329  -7.980  20.126  1.00  0.00           C
ATOM   2145  NH1 ARG   271      19.742  -8.912  19.320  1.00  0.00           N
ATOM   2146  NH2 ARG   271      21.519  -7.426  20.501  1.00  0.00           N
ATOM   2147  C   ARG   271      13.301  -6.745  22.347  1.00  0.00           C
ATOM   2148  O   ARG   271      12.762  -5.868  21.670  1.00  0.00           O
ATOM   2149  N   ILE   272      12.672  -7.822  22.804  1.00  0.00           N
ATOM   2150  CA  ILE   272      10.025   5.796   1.295 50.00  0.00           C
ATOM   2151  CB  ILE   272      10.933  -9.588  22.713  1.00  0.00           C
ATOM   2152  CG2 ILE   272       9.457  -9.925  22.440  1.00  0.00           C
ATOM   2153  CG1 ILE   272      11.737 -10.562  21.836  1.00  0.00           C
ATOM   2154  CD1 ILE   272      11.519 -10.354  20.337  1.00  0.00           C
ATOM   2155  C   ILE   272      10.353  -7.134  23.197  1.00  0.00           C
ATOM   2156  O   ILE   272       9.373  -6.627  22.617  1.00  0.00           O
ATOM   2157  N   LEU   273      10.688  -6.856  24.456  1.00  0.00           N
ATOM   2158  CA  LEU   273      10.594   8.234   2.132 50.00  0.00           C
ATOM   2159  CB  LEU   273      10.351  -5.912  26.717  1.00  0.00           C
ATOM   2160  CG  LEU   273       9.503  -4.996  27.602  1.00  0.00           C
ATOM   2161  CD1 LEU   273       8.007  -5.307  27.608  1.00  0.00           C
ATOM   2162  CD2 LEU   273       9.874  -5.007  29.085  1.00  0.00           C
ATOM   2163  C   LEU   273      10.189  -4.480  24.671  1.00  0.00           C
ATOM   2164  O   LEU   273       9.271  -3.645  24.639  1.00  0.00           O
ATOM   2165  N   SER   274      11.351  -4.242  24.057  1.00  0.00           N
ATOM   2166  CA  SER   274      10.127  11.008   1.746 50.00  0.00           C
ATOM   2167  CB  SER   274      12.972  -2.348  23.084  1.00  0.00           C
ATOM   2168  OG  SER   274      13.535  -3.087  22.010  1.00  0.00           O
ATOM   2169  C   SER   274      11.020  -2.879  22.061  1.00  0.00           C
ATOM   2170  O   SER   274      10.719  -1.809  21.527  1.00  0.00           O
ATOM   2171  N   ARG   275      10.847  -4.043  21.439  1.00  0.00           N
ATOM   2172  CA  ARG   275       9.130   9.556   1.712 50.00  0.00           C
ATOM   2173  CB  ARG   275      10.146  -5.414  19.311  1.00  0.00           C
ATOM   2174  CG  ARG   275      11.495  -5.915  18.790  1.00  0.00           C
ATOM   2175  CD  ARG   275      11.410  -7.259  18.063  1.00  0.00           C
ATOM   2176  NE  ARG   275      12.787  -7.617  17.622  1.00  0.00           N
ATOM   2177  CZ  ARG   275      13.018  -8.818  17.015  1.00  0.00           C
ATOM   2178  NH1 ARG   275      11.828  -9.483  16.942  1.00  0.00           N
ATOM   2179  NH2 ARG   275      14.349  -8.885  16.719  1.00  0.00           N
ATOM   2180  C   ARG   275       8.760  -3.753  20.213  1.00  0.00           C
ATOM   2181  O   ARG   275       8.205  -3.005  19.404  1.00  0.00           O
ATOM   2182  N   GLN   276       8.104  -4.300  21.235  1.00  0.00           N
ATOM   2183  CA  GLN   276       9.188   9.660   2.860 50.00  0.00           C
ATOM   2184  CB  GLN   276       5.893  -4.717  22.606  1.00  0.00           C
ATOM   2185  CG  GLN   276       4.398  -4.391  22.584  1.00  0.00           C
ATOM   2186  CD  GLN   276       3.818  -4.930  21.285  1.00  0.00           C
ATOM   2187  OE1 GLN   276       4.098  -6.060  20.887  1.00  0.00           O
ATOM   2188  NE2 GLN   276       2.978  -4.151  20.552  1.00  0.00           N
ATOM   2189  C   GLN   276       6.465  -2.584  21.876  1.00  0.00           C
ATOM   2190  O   GLN   276       5.465  -1.952  21.524  1.00  0.00           O
ATOM   2191  N   THR   277       7.418  -2.033  22.624  1.00  0.00           N
ATOM   2192  CA  THR   277       9.552  13.174   3.084 50.00  0.00           C
ATOM   2193  CB  THR   277       7.894   0.694  23.601  1.00  0.00           C
ATOM   2194  OG1 THR   277       8.351   0.515  24.933  1.00  0.00           O
ATOM   2195  CG2 THR   277       6.772   1.745  23.585  1.00  0.00           C
ATOM   2196  C   THR   277       7.383   0.224  21.773  1.00  0.00           C
ATOM   2197  O   THR   277       6.611   1.173  21.609  1.00  0.00           O
ATOM   2198  N   GLN   278       8.290  -0.114  20.862  1.00  0.00           N
ATOM   2199  CA  GLN   278       8.663  14.176   2.524 50.00  0.00           C
ATOM   2200  CB  GLN   278       9.475   0.208  18.544  1.00  0.00           C
ATOM   2201  CG  GLN   278       9.537   1.174  17.359  1.00  0.00           C
ATOM   2202  CD  GLN   278      10.001   2.527  17.880  1.00  0.00           C
ATOM   2203  OE1 GLN   278      10.956   2.616  18.649  1.00  0.00           O
ATOM   2204  NE2 GLN   278       9.349   3.655  17.488  1.00  0.00           N
ATOM   2205  C   GLN   278       7.195   0.519  18.737  1.00  0.00           C
ATOM   2206  O   GLN   278       6.744   1.507  18.152  1.00  0.00           O
ATOM   2207  N   ASN   279       6.614  -0.672  18.640  1.00  0.00           N
ATOM   2208  CA  ASN   279       7.953  12.549   3.102 50.00  0.00           C
ATOM   2209  CB  ASN   279       4.934  -2.294  17.728  1.00  0.00           C
ATOM   2210  CG  ASN   279       5.899  -3.053  16.829  1.00  0.00           C
ATOM   2211  OD1 ASN   279       6.642  -2.457  16.051  1.00  0.00           O
ATOM   2212  ND2 ASN   279       5.941  -4.411  16.887  1.00  0.00           N
ATOM   2213  C   ASN   279       4.256  -0.027  18.368  1.00  0.00           C
ATOM   2214  O   ASN   279       3.582   0.704  17.637  1.00  0.00           O
ATOM   2215  N   GLY   280       4.001  -0.138  19.667  1.00  0.00           N
ATOM   2216  CA  GLY   280       8.394  14.133   4.161 50.00  0.00           C
ATOM   2217  C   GLY   280       3.217   2.110  20.142  1.00  0.00           C
ATOM   2218  O   GLY   280       2.325   2.928  19.903  1.00  0.00           O
ATOM   2219  N   GLY   281       4.458   2.498  20.281  1.00  0.00           N
ATOM   2220  CA  GLY   281       8.454  17.403   3.859 50.00  0.00           C
ATOM   2221  C   GLY   281       5.805   4.281  21.327  1.00  0.00           C
ATOM   2222  O   GLY   281       6.674   5.138  21.167  1.00  0.00           O
ATOM   2223  N   LEU   282       5.608   3.657  22.484  1.00  0.00           N
ATOM   2224  CA  LEU   282       9.548  17.812   3.763 50.00  0.00           C
ATOM   2225  CB  LEU   282       6.046   3.015  24.813  1.00  0.00           C
ATOM   2226  CG  LEU   282       4.622   3.225  25.332  1.00  0.00           C
ATOM   2227  CD1 LEU   282       4.162   2.214  26.381  1.00  0.00           C
ATOM   2228  CD2 LEU   282       4.372   4.573  26.005  1.00  0.00           C
ATOM   2229  C   LEU   282       7.906   3.829  23.348  1.00  0.00           C
ATOM   2230  O   LEU   282       8.468   2.734  23.364  1.00  0.00           O
ATOM   2231  N   VAL   283       8.540   4.963  23.064  1.00  0.00           N
ATOM   2232  CA  VAL   283       9.566  18.811   2.759 50.00  0.00           C
ATOM   2233  CB  VAL   283      10.292   6.077  21.711  1.00  0.00           C
ATOM   2234  CG1 VAL   283      10.015   5.570  20.303  1.00  0.00           C
ATOM   2235  CG2 VAL   283       9.493   7.339  21.997  1.00  0.00           C
ATOM   2236  C   VAL   283      10.802   5.220  24.000  1.00  0.00           C
ATOM   2237  O   VAL   283      11.690   4.430  24.319  1.00  0.00           O
ATOM   2238  N   ASN   284      10.514   6.310  24.704  1.00  0.00           N
ATOM   2239  CA  ASN   284      10.518  20.758   2.838 50.00  0.00           C
ATOM   2240  CB  ASN   284      10.521   7.763  26.684  1.00  0.00           C
ATOM   2241  CG  ASN   284      10.725   9.063  25.920  1.00  0.00           C
ATOM   2242  OD1 ASN   284      11.603   9.165  25.065  1.00  0.00           O
ATOM   2243  ND2 ASN   284       9.923  10.129  26.186  1.00  0.00           N
ATOM   2244  C   ASN   284      11.417   5.428  26.818  1.00  0.00           C
ATOM   2245  O   ASN   284      12.526   5.121  27.255  1.00  0.00           O
ATOM   2246  N   GLN   285      10.315   4.736  27.089  1.00  0.00           N
ATOM   2247  CA  GLN   285      11.160  17.881   3.035 50.00  0.00           C
ATOM   2248  CB  GLN   285       8.934   3.108  28.304  1.00  0.00           C
ATOM   2249  CG  GLN   285       8.230   4.068  29.265  1.00  0.00           C
ATOM   2250  CD  GLN   285       6.817   3.550  29.495  1.00  0.00           C
ATOM   2251  OE1 GLN   285       6.408   2.546  28.915  1.00  0.00           O
ATOM   2252  NE2 GLN   285       5.994   4.209  30.354  1.00  0.00           N
ATOM   2253  C   GLN   285      11.130   2.389  27.311  1.00  0.00           C
ATOM   2254  O   GLN   285      11.704   1.582  28.022  1.00  0.00           O
ATOM   2255  N   HIS   286      11.152   2.327  25.982  1.00  0.00           N
ATOM   2256  CA  HIS   286      10.657  15.564   2.045 50.00  0.00           C
ATOM   2257  ND1 HIS   286      12.685   0.602  21.557  1.00  0.00           N
ATOM   2258  CG  HIS   286      12.619   0.605  22.935  1.00  0.00           C
ATOM   2259  CB  HIS   286      11.847   1.611  23.736  1.00  0.00           C
ATOM   2260  NE2 HIS   286      13.862  -1.043  22.188  1.00  0.00           N
ATOM   2261  CD2 HIS   286      13.374  -0.449  23.326  1.00  0.00           C
ATOM   2262  CE1 HIS   286      13.425  -0.378  21.136  1.00  0.00           C
ATOM   2263  C   HIS   286      13.494   1.837  25.597  1.00  0.00           C
ATOM   2264  O   HIS   286      14.298   0.977  25.950  1.00  0.00           O
ATOM   2265  N   PHE   287      13.811   3.124  25.516  1.00  0.00           N
ATOM   2266  CA  PHE   287      10.935  17.760   1.487 50.00  0.00           C
ATOM   2267  CB  PHE   287      15.407   4.975  25.249  1.00  0.00           C
ATOM   2268  CG  PHE   287      15.509   4.902  23.764  1.00  0.00           C
ATOM   2269  CD1 PHE   287      14.441   5.281  22.920  1.00  0.00           C
ATOM   2270  CD2 PHE   287      16.698   4.445  23.165  1.00  0.00           C
ATOM   2271  CE1 PHE   287      14.550   5.212  21.501  1.00  0.00           C
ATOM   2272  CE2 PHE   287      16.833   4.366  21.746  1.00  0.00           C
ATOM   2273  CZ  PHE   287      15.752   4.750  20.911  1.00  0.00           C
ATOM   2274  C   PHE   287      15.539   3.422  27.235  1.00  0.00           C
ATOM   2275  O   PHE   287      16.693   3.168  27.562  1.00  0.00           O
ATOM   2276  N   SER   288      14.568   3.586  28.123  1.00  0.00           N
ATOM   2277  CA  SER   288      11.952  17.893   1.905 50.00  0.00           C
ATOM   2278  CB  SER   288      13.598   3.712  30.363  1.00  0.00           C
ATOM   2279  OG  SER   288      13.283   5.093  30.258  1.00  0.00           O
ATOM   2280  C   SER   288      15.251   1.950  29.750  1.00  0.00           C
ATOM   2281  O   SER   288      16.194   1.635  30.491  1.00  0.00           O
ATOM   2282  N   ALA   289      14.540   1.074  29.050  1.00  0.00           N
ATOM   2283  CA  ALA   289      11.951  14.128   1.510 50.00  0.00           C
ATOM   2284  CB  ALA   289      13.716  -1.112  28.340  1.00  0.00           C
ATOM   2285  C   ALA   289      16.178  -0.661  28.477  1.00  0.00           C
ATOM   2286  O   ALA   289      16.841  -1.605  28.890  1.00  0.00           O
ATOM   2287  N   LEU   290      16.590   0.137  27.499  1.00  0.00           N
ATOM   2288  CA  LEU   290      11.583  14.294   0.509 50.00  0.00           C
ATOM   2289  CB  LEU   290      18.003   0.844  25.643  1.00  0.00           C
ATOM   2290  CG  LEU   290      17.086   0.439  24.487  1.00  0.00           C
ATOM   2291  CD1 LEU   290      17.046   1.424  23.319  1.00  0.00           C
ATOM   2292  CD2 LEU   290      17.438  -0.889  23.819  1.00  0.00           C
ATOM   2293  C   LEU   290      18.995   0.296  27.803  1.00  0.00           C
ATOM   2294  O   LEU   290      19.956  -0.445  27.932  1.00  0.00           O
ATOM   2295  N   ALA   291      18.873   1.442  28.457  1.00  0.00           N
ATOM   2296  CA  ALA   291      12.465  16.421   0.351 50.00  0.00           C
ATOM   2297  CB  ALA   291      19.421   3.123  30.192  1.00  0.00           C
ATOM   2298  C   ALA   291      19.905   0.719  30.536  1.00  0.00           C
ATOM   2299  O   ALA   291      20.992   0.383  30.979  1.00  0.00           O
ATOM   2300  N   ARG   292      18.766   0.158  30.927  1.00  0.00           N
ATOM   2301  CA  ARG   292      13.145  13.885   0.643 50.00  0.00           C
ATOM   2302  CB  ARG   292      17.330  -1.392  32.182  1.00  0.00           C
ATOM   2303  CG  ARG   292      16.496  -0.348  32.927  1.00  0.00           C
ATOM   2304  CD  ARG   292      15.056  -0.791  33.195  1.00  0.00           C
ATOM   2305  NE  ARG   292      14.365   0.329  33.893  1.00  0.00           N
ATOM   2306  CZ  ARG   292      13.038   0.230  34.195  1.00  0.00           C
ATOM   2307  NH1 ARG   292      12.599  -0.984  33.749  1.00  0.00           N
ATOM   2308  NH2 ARG   292      12.667   1.383  34.825  1.00  0.00           N
ATOM   2309  C   ARG   292      19.589  -2.102  31.357  1.00  0.00           C
ATOM   2310  O   ARG   292      20.468  -2.626  32.041  1.00  0.00           O
ATOM   2311  N   GLY   293      20.045  -3.373  31.306  1.00  0.00           N
ATOM   2312  CA  GLY   293       5.249  25.707 -30.170 99.00  0.00           C
ATOM   2313  C   GLY   293       5.531  24.717 -31.257  1.00  0.00           C
ATOM   2314  O   GLY   293       5.308  24.992 -32.433  1.00  0.00           O
ATOM   2315  N   THR   294      21.973  -2.098  29.215  1.00  0.00           N
ATOM   2316  CA  THR   294      12.780  12.954  -0.967 50.00  0.00           C
ATOM   2317  CB  THR   294      23.602  -0.406  28.473  1.00  0.00           C
ATOM   2318  OG1 THR   294      23.168  -0.506  27.125  1.00  0.00           O
ATOM   2319  CG2 THR   294      25.095  -0.037  28.502  1.00  0.00           C
ATOM   2320  C   THR   294      24.021  -1.674  30.590  1.00  0.00           C
ATOM   2321  O   THR   294      25.205  -1.978  30.763  1.00  0.00           O
ATOM   2322  N   THR   295      23.228  -1.253  31.571  1.00  0.00           N
ATOM   2323  CA  THR   295      13.842  13.843  -0.649 50.00  0.00           C
ATOM   2324  CB  THR   295      22.621  -0.547  33.844  1.00  0.00           C
ATOM   2325  OG1 THR   295      22.273   0.758  33.405  1.00  0.00           O
ATOM   2326  CG2 THR   295      23.142  -0.476  35.289  1.00  0.00           C
ATOM   2327  C   THR   295      24.150  -2.492  33.468  1.00  0.00           C
ATOM   2328  O   THR   295      25.221  -2.624  34.062  1.00  0.00           O
ATOM   2329  N   THR   296      23.316  -3.500  33.242  1.00  0.00           N
ATOM   2330  CA  THR   296      14.176  10.206  -0.869 50.00  0.00           C
ATOM   2331  CB  THR   296      22.400  -5.796  33.538  1.00  0.00           C
ATOM   2332  OG1 THR   296      22.315  -6.303  32.201  1.00  0.00           O
ATOM   2333  CG2 THR   296      21.107  -5.057  33.906  1.00  0.00           C
ATOM   2334  C   THR   296      24.730  -5.486  32.805  1.00  0.00           C
ATOM   2335  O   THR   296      24.721  -6.687  32.547  1.00  0.00           O
ATOM   2336  N   GLY   297      25.665  -4.659  32.347  1.00  0.00           N
ATOM   2337  CA  GLY   297      13.912   9.804  -2.027 50.00  0.00           C
ATOM   2338  C   GLY   297      26.662  -5.784  30.296  1.00  0.00           C
ATOM   2339  O   GLY   297      27.651  -5.989  29.584  1.00  0.00           O
ATOM   2340  N   SER   298      25.419  -6.059  29.923  1.00  0.00           N
ATOM   2341  CA  SER   298      12.957   8.146  -1.921 50.00  0.00           C
ATOM   2342  CB  SER   298      23.636  -6.644  28.341  1.00  0.00           C
ATOM   2343  OG  SER   298      23.381  -7.108  27.023  1.00  0.00           O
ATOM   2344  C   SER   298      25.766  -5.719  27.527  1.00  0.00           C
ATOM   2345  O   SER   298      25.627  -4.510  27.541  1.00  0.00           O
ATOM   2346  N   ASP   299      26.456  -6.333  26.590  1.00  0.00           N
ATOM   2347  CA  ASP   299      12.189   8.923  -2.659 50.00  0.00           C
ATOM   2348  CB  ASP   299      28.468  -5.886  25.211  1.00  0.00           C
ATOM   2349  CG  ASP   299      28.956  -4.994  24.079  1.00  0.00           C
ATOM   2350  OD1 ASP   299      28.119  -4.233  23.523  1.00  0.00           O
ATOM   2351  OD2 ASP   299      30.172  -5.061  23.755  1.00  0.00           O
ATOM   2352  C   ASP   299      26.179  -5.754  24.205  1.00  0.00           C
ATOM   2353  O   ASP   299      26.241  -6.822  23.597  1.00  0.00           O
ATOM   2354  N   ARG   300      25.410  -4.736  23.787  1.00  0.00           N
ATOM   2355  CA  ARG   300      11.146   9.245  -2.184 50.00  0.00           C
ATOM   2356  CB  ARG   300      23.585  -3.719  22.732  1.00  0.00           C
ATOM   2357  CG  ARG   300      22.660  -3.720  23.951  1.00  0.00           C
ATOM   2358  CD  ARG   300      21.829  -2.443  24.090  1.00  0.00           C
ATOM   2359  NE  ARG   300      20.891  -2.390  22.933  1.00  0.00           N
ATOM   2360  CZ  ARG   300      20.230  -1.232  22.643  1.00  0.00           C
ATOM   2361  NH1 ARG   300      20.609  -0.295  23.560  1.00  0.00           N
ATOM   2362  NH2 ARG   300      19.447  -1.463  21.549  1.00  0.00           N
ATOM   2363  C   ARG   300      25.218  -4.999  21.318  1.00  0.00           C
ATOM   2364  O   ARG   300      26.280  -4.386  21.151  1.00  0.00           O
ATOM   2365  N   ARG   301      24.648  -5.731  20.366  1.00  0.00           N
ATOM   2366  CA  ARG   301      10.225   8.099  -2.809 50.00  0.00           C
ATOM   2367  CB  ARG   301      25.542  -7.247  18.602  1.00  0.00           C
ATOM   2368  CG  ARG   301      26.651  -7.929  19.405  1.00  0.00           C
ATOM   2369  CD  ARG   301      26.974  -9.345  18.925  1.00  0.00           C
ATOM   2370  NE  ARG   301      25.832 -10.221  19.312  1.00  0.00           N
ATOM   2371  CZ  ARG   301      25.847 -11.546  18.985  1.00  0.00           C
ATOM   2372  NH1 ARG   301      27.010 -11.796  18.315  1.00  0.00           N
ATOM   2373  NH2 ARG   301      24.693 -12.103  19.456  1.00  0.00           N
ATOM   2374  C   ARG   301      24.575  -4.975  17.956  1.00  0.00           C
ATOM   2375  O   ARG   301      23.641  -4.217  18.196  1.00  0.00           O
ATOM   2376  N   HIS   302      -0.945  14.963 -19.995  1.00  0.00           N
ATOM   2377  CA  HIS   302      -1.956  16.630 -20.816 50.00  0.00           C
ATOM   2378  ND1 HIS   302       2.234  13.231 -18.537  1.00  0.00           N
ATOM   2379  CG  HIS   302       1.262  14.078 -18.045  1.00  0.00           C
ATOM   2380  CB  HIS   302      -0.194  13.726 -17.963  1.00  0.00           C
ATOM   2381  NE2 HIS   302       3.250  15.006 -17.982  1.00  0.00           N
ATOM   2382  CD2 HIS   302       1.922  15.207 -17.694  1.00  0.00           C
ATOM   2383  CE1 HIS   302       3.401  13.799 -18.492  1.00  0.00           C
ATOM   2384  C   HIS   302      -2.250  12.996 -19.197  1.00  0.00           C
ATOM   2385  O   HIS   302      -2.394  11.807 -19.485  1.00  0.00           O
ATOM   2386  N   GLU   303      -3.239  13.777 -18.757  1.00  0.00           N
ATOM   2387  CA  GLU   303      -5.371  15.823 -19.567 50.00  0.00           C
ATOM   2388  CB  GLU   303      -5.568  14.504 -18.556  1.00  0.00           C
ATOM   2389  CG  GLU   303      -5.884  14.897 -17.161  1.00  0.00           C
ATOM   2390  CD  GLU   303      -7.152  15.694 -17.103  1.00  0.00           C
ATOM   2391  OE1 GLU   303      -7.252  16.687 -17.857  1.00  0.00           O
ATOM   2392  OE2 GLU   303      -8.054  15.304 -16.326  1.00  0.00           O
ATOM   2393  C   GLU   303      -4.962  12.645 -19.983  1.00  0.00           C
ATOM   2394  O   GLU   303      -5.345  11.472 -20.000  1.00  0.00           O
ATOM   2395  N   THR   304      -4.793  13.391 -21.074  1.00  0.00           N
ATOM   2396  CA  THR   304      -6.394  15.075 -23.078 50.00  0.00           C
ATOM   2397  CB  THR   304      -4.871  13.883 -23.498  1.00  0.00           C
ATOM   2398  OG1 THR   304      -5.681  15.032 -23.296  1.00  0.00           O
ATOM   2399  CG2 THR   304      -5.210  13.254 -24.860  1.00  0.00           C
ATOM   2400  C   THR   304      -4.435  11.535 -22.669  1.00  0.00           C
ATOM   2401  O   THR   304      -5.071  10.596 -23.146  1.00  0.00           O
ATOM   2402  N   GLU   305      -3.148  11.449 -22.338  1.00  0.00           N
ATOM   2403  CA  GLU   305      -3.371  13.053 -23.760 50.00  0.00           C
ATOM   2404  CB  GLU   305      -0.959  10.365 -22.080  1.00  0.00           C
ATOM   2405  CG  GLU   305       0.039   9.424 -22.726  1.00  0.00           C
ATOM   2406  CD  GLU   305       1.472   9.771 -22.316  1.00  0.00           C
ATOM   2407  OE1 GLU   305       2.434   9.164 -22.830  1.00  0.00           O
ATOM   2408  OE2 GLU   305       1.633  10.670 -21.468  1.00  0.00           O
ATOM   2409  C   GLU   305      -3.064   9.062 -21.732  1.00  0.00           C
ATOM   2410  O   GLU   305      -3.139   7.936 -22.208  1.00  0.00           O
ATOM   2411  N   GLY   306      -3.573   9.395 -20.547  1.00  0.00           N
ATOM   2412  CA  GLY   306      -3.970  11.536 -20.537 50.00  0.00           C
ATOM   2413  C   GLY   306      -5.521   7.860 -20.067  1.00  0.00           C
ATOM   2414  O   GLY   306      -5.839   6.696 -19.797  1.00  0.00           O
ATOM   2415  N   HIS   307      -6.310   8.657 -20.776  1.00  0.00           N
ATOM   2416  CA  HIS   307      -7.039  10.111 -21.704 50.00  0.00           C
ATOM   2417  ND1 HIS   307      -9.542   9.282 -18.819  1.00  0.00           N
ATOM   2418  CG  HIS   307      -8.866   9.953 -19.817  1.00  0.00           C
ATOM   2419  CB  HIS   307      -8.624   9.395 -21.188  1.00  0.00           C
ATOM   2420  NE2 HIS   307      -8.980  11.143 -17.976  1.00  0.00           N
ATOM   2421  CD2 HIS   307      -8.513  11.139 -19.268  1.00  0.00           C
ATOM   2422  CE1 HIS   307      -9.597  10.002 -17.739  1.00  0.00           C
ATOM   2423  C   HIS   307      -7.655   7.524 -22.551  1.00  0.00           C
ATOM   2424  O   HIS   307      -8.719   7.163 -23.056  1.00  0.00           O
ATOM   2425  N   GLY   308      -6.470   7.331 -23.130  1.00  0.00           N
ATOM   2426  CA  GLY   308      -4.739   8.258 -23.839 50.00  0.00           C
ATOM   2427  C   GLY   308      -5.795   5.197 -24.241  1.00  0.00           C
ATOM   2428  O   GLY   308      -5.381   4.771 -23.156  1.00  0.00           O
ATOM   2429  N   GLY   309      -2.781   4.077 -16.666  1.00  0.00           N
ATOM   2430  CA  GLY   309      -3.295   7.208 -17.909 50.00  0.00           C
ATOM   2431  C   GLY   309      -4.599   3.771 -14.946  1.00  0.00           C
ATOM   2432  O   GLY   309      -5.765   3.440 -14.715  1.00  0.00           O
ATOM   2433  N   THR   310      -3.625   3.787 -14.030  1.00  0.00           N
ATOM   2434  CA  THR   310      -2.080   6.228 -14.556 50.00  0.00           C
ATOM   2435  CB  THR   310      -2.329   3.283 -12.000  1.00  0.00           C
ATOM   2436  OG1 THR   310      -1.963   4.647 -11.851  1.00  0.00           O
ATOM   2437  CG2 THR   310      -1.274   2.563 -12.857  1.00  0.00           C
ATOM   2438  C   THR   310      -4.798   3.730 -11.761  1.00  0.00           C
ATOM   2439  O   THR   310      -5.941   3.918 -12.168  1.00  0.00           O
ATOM   2440  N   GLN   311      -4.439   3.933 -10.497  1.00  0.00           N
ATOM   2441  CA  GLN   311      -3.061   7.110 -10.998 50.00  0.00           C
ATOM   2442  CB  GLN   311      -5.807   3.674  -8.488  1.00  0.00           C
ATOM   2443  CG  GLN   311      -6.827   2.674  -9.037  1.00  0.00           C
ATOM   2444  CD  GLN   311      -7.430   1.923  -7.858  1.00  0.00           C
ATOM   2445  OE1 GLN   311      -6.842   0.974  -7.344  1.00  0.00           O
ATOM   2446  NE2 GLN   311      -8.638   2.309  -7.366  1.00  0.00           N
ATOM   2447  C   GLN   311      -4.382   5.625  -8.857  1.00  0.00           C
ATOM   2448  O   GLN   311      -4.797   6.730  -8.530  1.00  0.00           O
ATOM   2449  N   ALA   312      -3.141   5.211  -8.627  1.00  0.00           N
ATOM   2450  CA  ALA   312      -0.080   9.139  -9.948 50.00  0.00           C
ATOM   2451  CB  ALA   312      -0.791   5.379  -7.957  1.00  0.00           C
ATOM   2452  C   ALA   312      -2.026   7.331  -9.006  1.00  0.00           C
ATOM   2453  O   ALA   312      -1.718   8.427  -8.557  1.00  0.00           O
ATOM   2454  N   PHE   313      -4.191   5.942  -9.728  1.00  0.00           N
ATOM   2455  CA  PHE   313      -2.877   9.333 -11.939 33.67  0.00           C
ATOM   2456  CB  PHE   313      -4.649   6.166 -12.022  1.00  0.00           C
ATOM   2457  CG  PHE   313      -4.190   6.241 -12.835  1.00  0.00           C
ATOM   2458  CD1 PHE   313      -3.590   5.969 -12.690  1.00  0.00           C
ATOM   2459  CD2 PHE   313      -4.352   6.592 -13.781  1.00  0.00           C
ATOM   2460  CE1 PHE   313      -3.155   6.041 -13.475  1.00  0.00           C
ATOM   2461  CE2 PHE   313      -3.924   6.673 -14.580  1.00  0.00           C
ATOM   2462  CZ  PHE   313      -3.323   6.396 -14.424  1.00  0.00           C
ATOM   2463  C   PHE   313      -5.301   7.901 -10.652  1.00  0.00           C
ATOM   2464  O   PHE   313      -5.055   9.037 -10.882  1.00  0.00           O
ATOM   2465  N   PHE   314      -6.415   7.519 -10.234  1.00  0.00           N
ATOM   2466  CA  PHE   314      -6.066  10.312 -10.779 33.67  0.00           C
ATOM   2467  CB  PHE   314      -8.543   7.715  -9.567  1.00  0.00           C
ATOM   2468  CG  PHE   314      -9.079   7.126 -10.641  1.00  0.00           C
ATOM   2469  CD1 PHE   314      -8.826   6.720 -11.248  1.00  0.00           C
ATOM   2470  CD2 PHE   314      -9.863   6.960 -11.078  1.00  0.00           C
ATOM   2471  CE1 PHE   314      -9.343   6.157 -12.272  1.00  0.00           C
ATOM   2472  CE2 PHE   314     -10.393   6.398 -12.103  1.00  0.00           C
ATOM   2473  CZ  PHE   314     -10.131   5.996 -12.702  1.00  0.00           C
ATOM   2474  C   PHE   314      -7.090   9.428  -9.091  1.00  0.00           C
ATOM   2475  O   PHE   314      -7.385  10.566  -9.325  1.00  0.00           O
ATOM   2476  N   ALA   315      -6.478   8.950  -8.036  1.00  0.00           N
ATOM   2477  CA  ALA   315      -4.551  11.629  -7.822 33.67  0.00           C
ATOM   2478  CB  ALA   315      -5.239   8.819  -6.160  1.00  0.00           C
ATOM   2479  C   ALA   315      -5.201  10.725  -7.663  1.00  0.00           C
ATOM   2480  O   ALA   315      -5.415  11.872  -7.455  1.00  0.00           O
ATOM   2481  N   THR   316      -4.277  10.280  -8.387  1.00  0.00           N
ATOM   2482  CA  THR   316      -2.295  13.495  -9.788 33.67  0.00           C
ATOM   2483  CB  THR   316      -2.490  10.340  -9.805  1.00  0.00           C
ATOM   2484  OG1 THR   316      -1.775   9.424  -9.021  1.00  0.00           O
ATOM   2485  CG2 THR   316      -1.605  11.235 -10.444  1.00  0.00           C
ATOM   2486  C   THR   316      -4.155  12.120  -9.921  1.00  0.00           C
ATOM   2487  O   THR   316      -3.800  13.245 -10.045  1.00  0.00           O
ATOM   2488  N   ARG   317      -5.172  11.679 -10.538  1.00  0.00           N
ATOM   2489  CA  ARG   317      -5.053  14.549 -12.033 33.67  0.00           C
ATOM   2490  CB  ARG   317      -6.806  11.656 -12.265  1.00  0.00           C
ATOM   2491  CG  ARG   317      -6.060  10.806 -13.207  1.00  0.00           C
ATOM   2492  CD  ARG   317      -6.918   9.929 -13.969  1.00  0.00           C
ATOM   2493  NE  ARG   317      -6.063   9.152 -14.826  1.00  0.00           N
ATOM   2494  CZ  ARG   317      -6.560   8.233 -15.559  1.00  0.00           C
ATOM   2495  NH1 ARG   317      -7.845   8.230 -15.322  1.00  0.00           N
ATOM   2496  NH2 ARG   317      -5.577   7.676 -16.243  1.00  0.00           N
ATOM   2497  C   ARG   317      -6.749  13.539 -10.818  1.00  0.00           C
ATOM   2498  O   ARG   317      -6.942  14.611 -11.272  1.00  0.00           O
ATOM   2499  N   LEU   318      -7.225  13.225  -9.768  1.00  0.00           N
ATOM   2500  CA  LEU   318      -7.060  15.875  -9.354 33.67  0.00           C
ATOM   2501  CB  LEU   318      -8.723  13.512  -7.932  1.00  0.00           C
ATOM   2502  CG  LEU   318      -9.942  12.864  -8.437  1.00  0.00           C
ATOM   2503  CD1 LEU   318     -10.597  12.102  -7.378  1.00  0.00           C
ATOM   2504  CD2 LEU   318     -10.937  13.805  -9.074  1.00  0.00           C
ATOM   2505  C   LEU   318      -6.963  15.142  -8.374  1.00  0.00           C
ATOM   2506  O   LEU   318      -7.219  16.333  -8.386  1.00  0.00           O
ATOM   2507  N   GLU   319      -5.858  14.640  -7.853  1.00  0.00           N
ATOM   2508  CA  GLU   319      -4.132  17.514  -7.814 33.67  0.00           C
ATOM   2509  CB  GLU   319      -3.765  14.650  -6.794  1.00  0.00           C
ATOM   2510  CG  GLU   319      -3.459  14.615  -6.189  1.00  0.00           C
ATOM   2511  CD  GLU   319      -2.365  13.713  -5.700  1.00  0.00           C
ATOM   2512  OE1 GLU   319      -1.922  13.149  -5.864  1.00  0.00           O
ATOM   2513  OE2 GLU   319      -1.959  13.577  -5.157  1.00  0.00           O
ATOM   2514  C   GLU   319      -4.425  16.471  -8.358  1.00  0.00           C
ATOM   2515  O   GLU   319      -4.320  17.645  -8.100  1.00  0.00           O
ATOM   2516  N   GLU   320      -4.209  15.997  -9.504  1.00  0.00           N
ATOM   2517  CA  GLU   320      -3.396  19.068 -11.006 33.67  0.00           C
ATOM   2518  CB  GLU   320      -3.465  16.103 -11.683  1.00  0.00           C
ATOM   2519  CG  GLU   320      -2.490  16.070 -12.223  1.00  0.00           C
ATOM   2520  CD  GLU   320      -2.301  15.279 -13.322  1.00  0.00           C
ATOM   2521  OE1 GLU   320      -2.945  14.783 -13.575  1.00  0.00           O
ATOM   2522  OE2 GLU   320      -1.511  15.159 -13.924  1.00  0.00           O
ATOM   2523  C   GLU   320      -4.735  17.909 -10.863  1.00  0.00           C
ATOM   2524  O   GLU   320      -4.483  19.076 -11.041  1.00  0.00           O
ATOM   2525  N   LEU   321      -5.867  17.516 -10.944  1.00  0.00           N
ATOM   2526  CA  LEU   321      -6.777  20.176 -11.487 33.67  0.00           C
ATOM   2527  CB  LEU   321      -8.160  17.846 -11.380  1.00  0.00           C
ATOM   2528  CG  LEU   321      -8.239  17.089 -12.542  1.00  0.00           C
ATOM   2529  CD1 LEU   321      -9.402  16.346 -12.681  1.00  0.00           C
ATOM   2530  CD2 LEU   321      -8.216  17.916 -13.759  1.00  0.00           C
ATOM   2531  C   LEU   321      -7.123  19.414 -10.135  1.00  0.00           C
ATOM   2532  O   LEU   321      -7.465  20.577 -10.335  1.00  0.00           O
ATOM   2533  N   ALA   322      -6.893  18.943  -8.994  1.00  0.00           N
ATOM   2534  CA  ALA   322      -6.800  21.495  -8.109 33.67  0.00           C
ATOM   2535  CB  ALA   322      -6.921  19.034  -6.678  1.00  0.00           C
ATOM   2536  C   ALA   322      -6.019  20.890  -7.893  1.00  0.00           C
ATOM   2537  O   ALA   322      -6.271  22.015  -7.499  1.00  0.00           O
ATOM   2538  N   ARG   323      -4.882  20.552  -8.320  1.00  0.00           N
ATOM   2539  CA  ARG   323      -3.887  23.578  -8.558 33.67  0.00           C
ATOM   2540  CB  ARG   323      -2.439  20.960  -8.285  1.00  0.00           C
ATOM   2541  CG  ARG   323      -2.093  20.292  -7.007  1.00  0.00           C
ATOM   2542  CD  ARG   323      -0.669  19.767  -6.980  1.00  0.00           C
ATOM   2543  NE  ARG   323      -0.493  19.111  -5.710  1.00  0.00           N
ATOM   2544  CZ  ARG   323       0.635  18.427  -5.500  1.00  0.00           C
ATOM   2545  NH1 ARG   323       1.422  18.503  -6.600  1.00  0.00           N
ATOM   2546  NH2 ARG   323       0.536  17.935  -4.272  1.00  0.00           N
ATOM   2547  C   ARG   323      -3.884  22.361  -9.522  1.00  0.00           C
ATOM   2548  O   ARG   323      -3.291  23.346  -9.550  1.00  0.00           O
ATOM   2549  N   LEU   324      -4.574  21.986 -10.532  1.00  0.00           N
ATOM   2550  CA  LEU   324      -5.088  24.785 -11.854 33.67  0.00           C
ATOM   2551  CB  LEU   324      -4.986  22.182 -12.887  1.00  0.00           C
ATOM   2552  CG  LEU   324      -4.109  21.212 -13.070  1.00  0.00           C
ATOM   2553  CD1 LEU   324      -4.397  20.567 -14.175  1.00  0.00           C
ATOM   2554  CD2 LEU   324      -2.933  21.640 -13.306  1.00  0.00           C
ATOM   2555  C   LEU   324      -5.656  23.773 -11.819  1.00  0.00           C
ATOM   2556  O   LEU   324      -6.170  24.388 -11.862  1.00  0.00           O
ATOM   2557  N   LYS   325      -6.767  25.312  -9.003  1.00  0.00           N
ATOM   2558  CA  LYS   325      -8.566  27.596  -9.561 50.00  0.00           C
ATOM   2559  CB  LYS   325      -8.840  26.242  -7.733  1.00  0.00           C
ATOM   2560  CG  LYS   325      -9.117  25.121  -6.729  1.00  0.00           C
ATOM   2561  CD  LYS   325      -9.813  25.600  -5.453  1.00  0.00           C
ATOM   2562  CE  LYS   325     -10.060  24.486  -4.435  1.00  0.00           C
ATOM   2563  NZ  LYS   325     -10.711  25.039  -3.227  1.00  0.00           N
ATOM   2564  C   LYS   325      -8.302  27.002  -9.882  1.00  0.00           C
ATOM   2565  O   LYS   325      -9.232  27.121 -10.685  1.00  0.00           O
ATOM   2566  N   SER   326      -7.375  27.959  -9.730  1.00  0.00           N
ATOM   2567  CA  SER   326      -8.620  31.026 -11.054 50.00  0.00           C
ATOM   2568  CB  SER   326      -6.602  30.456 -10.261  1.00  0.00           C
ATOM   2569  OG  SER   326      -5.250  30.182 -10.597  1.00  0.00           O
ATOM   2570  C   SER   326      -7.094  29.083 -11.943  1.00  0.00           C
ATOM   2571  O   SER   326      -7.447  29.931 -12.767  1.00  0.00           O
ATOM   2572  N   SER   327      -6.371  28.015 -12.308  1.00  0.00           N
ATOM   2573  CA  SER   327      -7.779  29.570 -14.433 50.00  0.00           C
ATOM   2574  CB  SER   327      -5.217  26.605 -14.254  1.00  0.00           C
ATOM   2575  OG  SER   327      -5.660  25.317 -13.852  1.00  0.00           O
ATOM   2576  C   SER   327      -7.403  27.095 -14.375  1.00  0.00           C
ATOM   2577  O   SER   327      -7.376  26.691 -15.541  1.00  0.00           O
ATOM   2578  N   THR   328      -8.525  27.038 -13.643  1.00  0.00           N
ATOM   2579  CA  THR   328     -11.089  27.838 -14.274 50.00  0.00           C
ATOM   2580  CB  THR   328     -10.945  26.231 -15.121  1.00  0.00           C
ATOM   2581  OG1 THR   328     -10.442  25.731 -16.351  1.00  0.00           O
ATOM   2582  CG2 THR   328     -11.661  27.569 -15.370  1.00  0.00           C
ATOM   2583  C   THR   328      -9.650  24.970 -14.529  1.00  0.00           C
ATOM   2584  O   THR   328     -10.109  24.523 -15.584  1.00  0.00           O
ATOM   2585  N   THR   329     -10.568  23.143 -14.067  1.00  0.00           N
ATOM   2586  CA  THR   329     -10.601  23.123 -13.767 33.67  0.00           C
ATOM   2587  CB  THR   329      -9.427  20.894 -13.174  1.00  0.00           C
ATOM   2588  OG1 THR   329      -9.520  21.013 -11.862  1.00  0.00           O
ATOM   2589  CG2 THR   329      -7.974  21.027 -13.651  1.00  0.00           C
ATOM   2590  C   THR   329     -11.250  21.229 -12.920  1.00  0.00           C
ATOM   2591  O   THR   329     -11.247  20.023 -12.719  1.00  0.00           O
ATOM   2592  N   LEU   330     -12.138  22.032 -12.359  1.00  0.00           N
ATOM   2593  CA  LEU   330     -13.279  22.209 -11.461 33.67  0.00           C
ATOM   2594  CB  LEU   330     -14.196  22.377 -11.121  1.00  0.00           C
ATOM   2595  CG  LEU   330     -15.248  21.719 -10.319  1.00  0.00           C
ATOM   2596  CD1 LEU   330     -14.800  20.998  -9.052  1.00  0.00           C
ATOM   2597  CD2 LEU   330     -16.284  22.628  -9.858  1.00  0.00           C
ATOM   2598  C   LEU   330     -13.971  20.433 -12.393  1.00  0.00           C
ATOM   2599  O   LEU   330     -14.285  19.339 -11.921  1.00  0.00           O
ATOM   2600  N   ASP   331     -14.279  20.808 -13.612  1.00  0.00           N
ATOM   2601  CA  ASP   331     -15.088  20.502 -14.249 33.67  0.00           C
ATOM   2602  CB  ASP   331     -15.225  20.502 -15.873  1.00  0.00           C
ATOM   2603  CG  ASP   331     -16.184  21.649 -15.794  1.00  0.00           C
ATOM   2604  OD1 ASP   331     -16.822  21.793 -14.734  1.00  0.00           O
ATOM   2605  OD2 ASP   331     -16.293  22.397 -16.794  1.00  0.00           O
ATOM   2606  C   ASP   331     -14.408  18.567 -14.627  1.00  0.00           C
ATOM   2607  O   ASP   331     -15.083  17.543 -14.627  1.00  0.00           O
ATOM   2608  N   ASP   332     -13.097  18.559 -14.745  1.00  0.00           N
ATOM   2609  CA  ASP   332     -12.181  18.348 -14.956 33.67  0.00           C
ATOM   2610  CB  ASP   332     -10.982  17.517 -15.435  1.00  0.00           C
ATOM   2611  CG  ASP   332     -11.116  17.979 -16.849  1.00  0.00           C
ATOM   2612  OD1 ASP   332     -12.015  17.621 -17.485  1.00  0.00           O
ATOM   2613  OD2 ASP   332     -10.320  18.696 -17.314  1.00  0.00           O
ATOM   2614  C   ASP   332     -12.228  16.577 -13.578  1.00  0.00           C
ATOM   2615  O   ASP   332     -12.182  15.343 -13.546  1.00  0.00           O
ATOM   2616  N   ALA   333     -12.159  17.297 -12.480  1.00  0.00           N
ATOM   2617  CA  ALA   333     -11.615  17.697 -11.309 33.67  0.00           C
ATOM   2618  CB  ALA   333     -11.872  17.612 -10.074  1.00  0.00           C
ATOM   2619  C   ALA   333     -13.443  15.913 -10.903  1.00  0.00           C
ATOM   2620  O   ALA   333     -13.443  14.812 -10.375  1.00  0.00           O
ATOM   2621  N   LEU   334     -14.558  16.543 -11.267  1.00  0.00           N
ATOM   2622  CA  LEU   334     -15.144  16.541 -11.077 33.67  0.00           C
ATOM   2623  CB  LEU   334     -16.983  16.921 -11.469  1.00  0.00           C
ATOM   2624  CG  LEU   334     -17.524  17.008 -10.949  1.00  0.00           C
ATOM   2625  CD1 LEU   334     -18.215  17.332 -10.424  1.00  0.00           C
ATOM   2626  CD2 LEU   334     -18.298  17.048 -10.537  1.00  0.00           C
ATOM   2627  C   LEU   334     -16.141  14.811 -12.009  1.00  0.00           C
ATOM   2628  O   LEU   334     -16.795  13.871 -11.599  1.00  0.00           O
ATOM   2629  N   ARG   335     -15.699  14.887 -13.260  1.00  0.00           N
ATOM   2630  CA  ARG   335     -14.845  14.469 -14.200 33.67  0.00           C
ATOM   2631  CB  ARG   335     -15.095  14.184 -15.493  1.00  0.00           C
ATOM   2632  CG  ARG   335     -15.923  14.199 -16.511  1.00  0.00           C
ATOM   2633  CD  ARG   335     -15.191  14.494 -17.796  1.00  0.00           C
ATOM   2634  NE  ARG   335     -14.990  14.633 -18.160  1.00  0.00           N
ATOM   2635  CZ  ARG   335     -14.603  14.346 -18.654  1.00  0.00           C
ATOM   2636  NH1 ARG   335     -14.390  13.777 -18.707  1.00  0.00           N
ATOM   2637  NH2 ARG   335     -14.505  14.482 -18.886  1.00  0.00           N
ATOM   2638  C   ARG   335     -14.970  12.600 -13.528  1.00  0.00           C
ATOM   2639  O   ARG   335     -15.479  11.507 -13.505  1.00  0.00           O
ATOM   2640  N   VAL   336     -13.795  12.824 -13.007  1.00  0.00           N
ATOM   2641  CA  VAL   336     -11.851  12.907 -12.684 33.67  0.00           C
ATOM   2642  CB  VAL   336     -11.614  12.245 -12.089  1.00  0.00           C
ATOM   2643  CG1 VAL   336     -10.788  11.246 -11.282  1.00  0.00           C
ATOM   2644  CG2 VAL   336     -10.788  12.536 -13.329  1.00  0.00           C
ATOM   2645  C   VAL   336     -13.630  11.181 -11.135  1.00  0.00           C
ATOM   2646  O   VAL   336     -13.400  10.011 -10.787  1.00  0.00           O
ATOM   2647  N   LEU   337     -14.385  11.997 -10.440  1.00  0.00           N
ATOM   2648  CA  LEU   337     -13.641  12.290  -9.488 33.67  0.00           C
ATOM   2649  CB  LEU   337     -15.725  12.645  -8.529  1.00  0.00           C
ATOM   2650  CG  LEU   337     -16.472  12.178  -7.318  1.00  0.00           C
ATOM   2651  CD1 LEU   337     -15.636  11.456  -6.280  1.00  0.00           C
ATOM   2652  CD2 LEU   337     -17.154  13.284  -6.553  1.00  0.00           C
ATOM   2653  C   LEU   337     -16.089  10.519  -9.759  1.00  0.00           C
ATOM   2654  O   LEU   337     -16.172   9.400  -9.253  1.00  0.00           O
ATOM   2655  N   ARG   338     -14.801  10.825 -12.413  1.00  0.00           N
ATOM   2656  CA  ARG   338     -14.292  10.775 -12.594 50.00  0.00           C
ATOM   2657  CB  ARG   338     -16.086  10.821 -14.469  1.00  0.00           C
ATOM   2658  CG  ARG   338     -17.048  10.076 -15.396  1.00  0.00           C
ATOM   2659  CD  ARG   338     -17.546  10.923 -16.570  1.00  0.00           C
ATOM   2660  NE  ARG   338     -18.431  10.059 -17.402  1.00  0.00           N
ATOM   2661  CZ  ARG   338     -18.975  10.549 -18.554  1.00  0.00           C
ATOM   2662  NH1 ARG   338     -18.549  11.837 -18.700  1.00  0.00           N
ATOM   2663  NH2 ARG   338     -19.733   9.559 -19.110  1.00  0.00           N
ATOM   2664  C   ARG   338     -14.949   8.735 -13.646  1.00  0.00           C
ATOM   2665  O   ARG   338     -15.536   7.647 -13.710  1.00  0.00           O
ATOM   2666  N   ARG   339     -13.653   8.878 -13.903  1.00  0.00           N
ATOM   2667  CA  ARG   339     -11.358   9.190 -14.240 50.00  0.00           C
ATOM   2668  CB  ARG   339     -11.534   8.348 -14.991  1.00  0.00           C
ATOM   2669  CG  ARG   339     -10.581   7.266 -15.503  1.00  0.00           C
ATOM   2670  CD  ARG   339     -11.172   6.412 -16.627  1.00  0.00           C
ATOM   2671  NE  ARG   339     -10.122   5.444 -17.052  1.00  0.00           N
ATOM   2672  CZ  ARG   339     -10.443   4.410 -17.884  1.00  0.00           C
ATOM   2673  NH1 ARG   339     -11.776   4.502 -18.166  1.00  0.00           N
ATOM   2674  NH2 ARG   339      -9.307   3.688 -18.109  1.00  0.00           N
ATOM   2675  C   ARG   339     -12.509   6.792 -13.237  1.00  0.00           C
ATOM   2676  O   ARG   339     -12.469   5.589 -13.468  1.00  0.00           O
ATOM   2677  N   MET   340     -14.649   7.202 -10.843  1.00  0.00           N
ATOM   2678  CA  MET   340     -12.277   7.353 -10.149 33.67  0.00           C
ATOM   2679  CB  MET   340     -13.645   7.073  -8.645  1.00  0.00           C
ATOM   2680  CG  MET   340     -12.814   6.824  -8.357  1.00  0.00           C
ATOM   2681  SD  MET   340     -11.949   7.602  -7.021  1.00  0.00           S
ATOM   2682  CE  MET   340     -11.566   8.682  -7.262  1.00  0.00           C
ATOM   2683  C   MET   340     -15.553   5.604  -9.207  1.00  0.00           C
ATOM   2684  O   MET   340     -15.505   4.410  -8.929  1.00  0.00           O
ATOM   2685  N   ASP   341     -16.642   6.347  -9.055  1.00  0.00           N
ATOM   2686  CA  ASP   341     -15.643   6.444  -8.784 33.67  0.00           C
ATOM   2687  CB  ASP   341     -18.785   6.939  -8.115  1.00  0.00           C
ATOM   2688  CG  ASP   341     -18.198   7.584  -6.917  1.00  0.00           C
ATOM   2689  OD1 ASP   341     -17.540   6.921  -6.087  1.00  0.00           O
ATOM   2690  OD2 ASP   341     -18.398   8.747  -6.816  1.00  0.00           O
ATOM   2691  C   ASP   341     -18.573   4.909  -9.533  1.00  0.00           C
ATOM   2692  O   ASP   341     -19.393   4.081  -9.169  1.00  0.00           O
ATOM   2693  N   VAL   342     -15.723   5.005 -12.486  1.00  0.00           N
ATOM   2694  CA  VAL   342     -14.010   5.212 -13.136 50.00  0.00           C
ATOM   2695  CB  VAL   342     -15.438   4.754 -14.903  1.00  0.00           C
ATOM   2696  CG1 VAL   342     -15.715   3.918 -16.154  1.00  0.00           C
ATOM   2697  CG2 VAL   342     -15.924   6.163 -15.247  1.00  0.00           C
ATOM   2698  C   VAL   342     -15.645   2.735 -13.439  1.00  0.00           C
ATOM   2699  O   VAL   342     -16.440   1.814 -13.621  1.00  0.00           O
ATOM   2700  N   GLU   343     -14.360   2.512 -13.781  1.00  0.00           N
ATOM   2701  CA  GLU   343     -11.469   2.918 -14.131 50.00  0.00           C
ATOM   2702  CB  GLU   343     -12.264   1.048 -13.855  1.00  0.00           C
ATOM   2703  CG  GLU   343     -11.777  -0.402 -13.849  1.00  0.00           C
ATOM   2704  CD  GLU   343     -10.260  -0.390 -13.978  1.00  0.00           C
ATOM   2705  OE1 GLU   343      -9.684   0.727 -14.075  1.00  0.00           O
ATOM   2706  OE2 GLU   343      -9.658  -1.496 -13.981  1.00  0.00           O
ATOM   2707  C   GLU   343     -14.081   0.402 -12.496  1.00  0.00           C
ATOM   2708  O   GLU   343     -14.051   0.857 -11.349  1.00  0.00           O
ATOM   2709  N   GLY   344     -16.319   0.778 -12.719  1.00  0.00           N
ATOM   2710  CA  GLY   344     -13.022   0.856 -12.647 33.67  0.00           C
ATOM   2711  C   GLY   344     -15.927  -0.149 -11.823  1.00  0.00           C
ATOM   2712  O   GLY   344     -15.530  -0.587 -11.567  1.00  0.00           O
ATOM   2713  N   THR   345     -16.494   0.064 -11.474  1.00  0.00           N
ATOM   2714  CA  THR   345     -14.746   0.406 -10.592 33.67  0.00           C
ATOM   2715  CB  THR   345     -17.108  -0.684  -9.764  1.00  0.00           C
ATOM   2716  OG1 THR   345     -17.298  -1.312  -8.791  1.00  0.00           O
ATOM   2717  CG2 THR   345     -17.233  -0.826  -9.447  1.00  0.00           C
ATOM   2718  C   THR   345     -16.873   0.543  -9.416  1.00  0.00           C
ATOM   2719  O   THR   345     -16.627   0.886  -8.564  1.00  0.00           O
ATOM   2720  N   PRO   346     -17.090   0.799  -9.325  1.00  0.00           N
ATOM   2721  CA  PRO   346     -14.502   2.489  -8.312 33.67  0.00           C
ATOM   2722  CD  PRO   346     -18.373  -0.120  -7.106  1.00  0.00           C
ATOM   2723  CB  PRO   346     -17.345   1.854  -6.954  1.00  0.00           C
ATOM   2724  CG  PRO   346     -18.100   0.864  -6.338  1.00  0.00           C
ATOM   2725  C   PRO   346     -15.902   1.661  -7.322  1.00  0.00           C
ATOM   2726  O   PRO   346     -15.496   1.624  -6.563  1.00  0.00           O
ATOM   2727  N   LEU   347     -15.392   1.804  -7.253  1.00  0.00           N
ATOM   2728  CA  LEU   347     -11.370   3.100  -7.033 33.67  0.00           C
ATOM   2729  CB  LEU   347     -13.224   2.859  -7.289  1.00  0.00           C
ATOM   2730  CG  LEU   347     -12.296   2.974  -7.730  1.00  0.00           C
ATOM   2731  CD1 LEU   347     -11.432   2.970  -7.794  1.00  0.00           C
ATOM   2732  CD2 LEU   347     -11.410   2.765  -8.330  1.00  0.00           C
ATOM   2733  C   LEU   347     -14.467   2.592  -5.239  1.00  0.00           C
ATOM   2734  O   LEU   347     -15.289   2.935  -5.031  1.00  0.00           O
ATOM   2735  N   ILE   348     -13.833   2.759  -4.364  1.00  0.00           N
ATOM   2736  CA  ILE   348     -11.165   4.443  -3.532 33.67  0.00           C
ATOM   2737  CB  ILE   348     -13.562   2.397  -1.959  1.00  0.00           C
ATOM   2738  CG2 ILE   348     -13.739   2.936  -0.538  1.00  0.00           C
ATOM   2739  CG1 ILE   348     -14.286   1.078  -1.980  1.00  0.00           C
ATOM   2740  CD1 ILE   348     -13.715   0.070  -1.006  1.00  0.00           C
ATOM   2741  C   ILE   348     -13.260   4.612  -2.926  1.00  0.00           C
ATOM   2742  O   ILE   348     -12.091   4.590  -2.701  1.00  0.00           O
ATOM   2743  N   SER   349     -13.914   5.719  -3.059  1.00  0.00           N
ATOM   2744  CA  SER   349     -10.972   8.103  -3.239 33.67  0.00           C
ATOM   2745  CB  SER   349     -14.349   7.999  -3.042  1.00  0.00           C
ATOM   2746  OG  SER   349     -14.513   8.718  -3.569  1.00  0.00           O
ATOM   2747  C   SER   349     -12.733   7.007  -1.457  1.00  0.00           C
ATOM   2748  O   SER   349     -13.293   6.496  -0.551  1.00  0.00           O
ATOM   2749  N   GLY   350     -11.626   7.566  -1.287  1.00  0.00           N
ATOM   2750  CA  GLY   350      -8.542   8.963  -0.555 33.67  0.00           C
ATOM   2751  C   GLY   350     -11.032   8.965   0.549  1.00  0.00           C
ATOM   2752  O   GLY   350     -11.344   9.934  -0.118  1.00  0.00           O
ATOM   2753  N   SER   351     -10.540   9.145   1.766  1.00  0.00           N
ATOM   2754  CA  SER   351      -8.339  11.731   1.989 33.67  0.00           C
ATOM   2755  CB  SER   351      -9.814  10.483   3.729  1.00  0.00           C
ATOM   2756  OG  SER   351      -9.944  10.664   4.472  1.00  0.00           O
ATOM   2757  C   SER   351      -9.921  11.519   1.615  1.00  0.00           C
ATOM   2758  O   SER   351     -10.470  12.614   1.503  1.00  0.00           O
ATOM   2759  N   VAL   352      -8.786  11.241   1.032  1.00  0.00           N
ATOM   2760  CA  VAL   352      -6.381  13.844  -0.315 33.67  0.00           C
ATOM   2761  CB  VAL   352      -6.908  12.063  -0.467  1.00  0.00           C
ATOM   2762  CG1 VAL   352      -6.630  13.103  -1.368  1.00  0.00           C
ATOM   2763  CG2 VAL   352      -5.728  11.985   0.482  1.00  0.00           C
ATOM   2764  C   VAL   352      -8.965  12.645  -1.020  1.00  0.00           C
ATOM   2765  O   VAL   352      -9.017  13.817  -1.394  1.00  0.00           O
ATOM   2766  N   VAL   353      -9.624  11.694  -1.645  1.00  0.00           N
ATOM   2767  CA  VAL   353      -8.786  13.349  -3.182 33.67  0.00           C
ATOM   2768  CB  VAL   353     -11.215  11.045  -3.627  1.00  0.00           C
ATOM   2769  CG1 VAL   353     -12.115  11.702  -4.610  1.00  0.00           C
ATOM   2770  CG2 VAL   353     -10.338  10.082  -4.389  1.00  0.00           C
ATOM   2771  C   VAL   353     -11.581  12.856  -2.511  1.00  0.00           C
ATOM   2772  O   VAL   353     -11.911  13.800  -3.223  1.00  0.00           O
ATOM   2773  N   LEU   354     -12.227  12.522  -1.431  1.00  0.00           N
ATOM   2774  CA  LEU   354     -11.686  14.167  -1.132 33.67  0.00           C
ATOM   2775  CB  LEU   354     -14.079  12.712   0.200  1.00  0.00           C
ATOM   2776  CG  LEU   354     -14.785  11.396  -0.031  1.00  0.00           C
ATOM   2777  CD1 LEU   354     -15.364  10.735   1.216  1.00  0.00           C
ATOM   2778  CD2 LEU   354     -15.957  11.476  -0.978  1.00  0.00           C
ATOM   2779  C   LEU   354     -12.983  14.677  -0.701  1.00  0.00           C
ATOM   2780  O   LEU   354     -13.679  15.612  -1.074  1.00  0.00           O
ATOM   2781  N   ASN   355     -11.867  14.834  -0.019  1.00  0.00           N
ATOM   2782  CA  ASN   355     -10.013  17.232   0.220 33.67  0.00           C
ATOM   2783  CB  ASN   355     -10.144  16.081   1.233  1.00  0.00           C
ATOM   2784  CG  ASN   355     -10.481  15.630   2.623  1.00  0.00           C
ATOM   2785  OD1 ASN   355     -11.610  15.720   3.025  1.00  0.00           O
ATOM   2786  ND2 ASN   355      -9.516  15.124   3.396  1.00  0.00           N
ATOM   2787  C   ASN   355     -11.089  17.022  -0.834  1.00  0.00           C
ATOM   2788  O   ASN   355     -11.437  18.185  -0.874  1.00  0.00           O
ATOM   2789  N   VAL   356     -10.450  16.463  -1.826  1.00  0.00           N
ATOM   2790  CA  VAL   356      -9.157  18.474  -3.171 33.67  0.00           C
ATOM   2791  CB  VAL   356      -9.450  16.688  -4.172  1.00  0.00           C
ATOM   2792  CG1 VAL   356      -9.456  17.623  -5.263  1.00  0.00           C
ATOM   2793  CG2 VAL   356      -8.049  16.353  -3.802  1.00  0.00           C
ATOM   2794  C   VAL   356     -11.373  17.634  -3.771  1.00  0.00           C
ATOM   2795  O   VAL   356     -11.450  18.706  -4.355  1.00  0.00           O
ATOM   2796  N   LEU   357     -12.366  16.795  -3.750  1.00  0.00           N
ATOM   2797  CA  LEU   357     -12.607  17.869  -4.355 33.67  0.00           C
ATOM   2798  CB  LEU   357     -14.563  15.916  -4.510  1.00  0.00           C
ATOM   2799  CG  LEU   357     -14.832  15.277  -5.495  1.00  0.00           C
ATOM   2800  CD1 LEU   357     -15.105  14.881  -6.301  1.00  0.00           C
ATOM   2801  CD2 LEU   357     -14.868  14.784  -6.196  1.00  0.00           C
ATOM   2802  C   LEU   357     -14.382  18.186  -3.752  1.00  0.00           C
ATOM   2803  O   LEU   357     -14.970  19.059  -4.395  1.00  0.00           O
ATOM   2804  N   LEU   358     -14.394  18.160  -2.425  1.00  0.00           N
ATOM   2805  CA  LEU   358     -14.037  19.758  -1.464 33.67  0.00           C
ATOM   2806  CB  LEU   358     -15.078  18.796  -0.233  1.00  0.00           C
ATOM   2807  CG  LEU   358     -15.659  18.696   0.766  1.00  0.00           C
ATOM   2808  CD1 LEU   358     -15.927  19.117   1.452  1.00  0.00           C
ATOM   2809  CD2 LEU   358     -16.562  18.475   1.309  1.00  0.00           C
ATOM   2810  C   LEU   358     -14.293  20.508  -1.850  1.00  0.00           C
ATOM   2811  O   LEU   358     -14.904  21.549  -2.061  1.00  0.00           O
ATOM   2812  N   SER   359     -12.992  20.437  -1.803  1.00  0.00           N
ATOM   2813  CA  SER   359     -11.623  22.507  -1.860 33.67  0.00           C
ATOM   2814  CB  SER   359     -10.732  21.243  -1.886  1.00  0.00           C
ATOM   2815  OG  SER   359     -10.317  21.354  -0.612  1.00  0.00           O
ATOM   2816  C   SER   359     -12.495  22.288  -3.318  1.00  0.00           C
ATOM   2817  O   SER   359     -12.751  23.483  -3.402  1.00  0.00           O
ATOM   2818  N   VAL   360     -12.482  21.516  -4.349  1.00  0.00           N
ATOM   2819  CA  VAL   360     -12.391  22.911  -5.482 33.67  0.00           C
ATOM   2820  CB  VAL   360     -12.378  21.010  -6.724  1.00  0.00           C
ATOM   2821  CG1 VAL   360     -12.763  21.493  -8.099  1.00  0.00           C
ATOM   2822  CG2 VAL   360     -10.920  20.616  -6.735  1.00  0.00           C
ATOM   2823  C   VAL   360     -14.028  22.669  -5.861  1.00  0.00           C
ATOM   2824  O   VAL   360     -14.179  23.740  -6.460  1.00  0.00           O
ATOM   2825  N   PHE   361     -15.023  22.010  -5.337  1.00  0.00           N
ATOM   2826  CA  PHE   361     -16.032  22.875  -5.011 33.67  0.00           C
ATOM   2827  CB  PHE   361     -17.326  21.562  -4.966  1.00  0.00           C
ATOM   2828  CG  PHE   361     -17.369  20.511  -5.935  1.00  0.00           C
ATOM   2829  CD1 PHE   361     -16.855  20.676  -7.198  1.00  0.00           C
ATOM   2830  CD2 PHE   361     -17.933  19.310  -5.602  1.00  0.00           C
ATOM   2831  CE1 PHE   361     -16.897  19.663  -8.117  1.00  0.00           C
ATOM   2832  CE2 PHE   361     -17.987  18.278  -6.504  1.00  0.00           C
ATOM   2833  CZ  PHE   361     -17.468  18.459  -7.768  1.00  0.00           C
ATOM   2834  C   PHE   361     -16.491  23.831  -4.687  1.00  0.00           C
ATOM   2835  O   PHE   361     -17.302  24.675  -5.062  1.00  0.00           O
ATOM   2836  N   CYS   362     -15.710  23.966  -3.645  1.00  0.00           N
ATOM   2837  CA  CYS   362     -15.546  25.563  -2.488 33.67  0.00           C
ATOM   2838  CB  CYS   362     -14.784  25.041  -1.738  1.00  0.00           C
ATOM   2839  SG  CYS   362     -15.339  24.226  -0.773  1.00  0.00           S
ATOM   2840  C   CYS   362     -15.136  26.357  -3.762  1.00  0.00           C
ATOM   2841  O   CYS   362     -15.503  27.472  -3.678  1.00  0.00           O
ATOM   2842  N   ARG   363     -14.013  26.145  -4.371  1.00  0.00           N
ATOM   2843  CA  ARG   363     -13.695  27.789  -4.773 33.67  0.00           C
ATOM   2844  CB  ARG   363     -12.104  26.787  -5.951  1.00  0.00           C
ATOM   2845  CG  ARG   363     -11.139  27.393  -5.851  1.00  0.00           C
ATOM   2846  CD  ARG   363      -9.876  27.024  -6.581  1.00  0.00           C
ATOM   2847  NE  ARG   363      -9.328  26.556  -6.754  1.00  0.00           N
ATOM   2848  CZ  ARG   363      -8.991  26.536  -7.588  1.00  0.00           C
ATOM   2849  NH1 ARG   363      -9.253  26.991  -8.125  1.00  0.00           N
ATOM   2850  NH2 ARG   363      -8.522  26.070  -7.558  1.00  0.00           N
ATOM   2851  C   ARG   363     -14.223  27.549  -6.388  1.00  0.00           C
ATOM   2852  O   ARG   363     -14.223  28.698  -6.812  1.00  0.00           O
ATOM   2853  N   ALA   364     -14.966  26.620  -6.940  1.00  0.00           N
ATOM   2854  CA  ALA   364     -16.238  27.282  -7.463 33.67  0.00           C
ATOM   2855  CB  ALA   364     -16.038  25.709  -8.890  1.00  0.00           C
ATOM   2856  C   ALA   364     -17.175  27.421  -7.634  1.00  0.00           C
ATOM   2857  O   ALA   364     -18.008  27.734  -8.466  1.00  0.00           O
ATOM   2858  N   GLY   365     -17.449  27.525  -6.361  1.00  0.00           N
ATOM   2859  CA  GLY   365     -19.070  27.912  -5.223 33.67  0.00           C
ATOM   2860  C   GLY   365     -19.804  27.074  -6.218  1.00  0.00           C
ATOM   2861  O   GLY   365     -20.968  27.429  -6.253  1.00  0.00           O
ATOM   2862  N   HIS   366     -19.491  25.810  -6.407  1.00  0.00           N
ATOM   2863  CA  HIS   366     -20.508  24.533  -5.947 33.67  0.00           C
ATOM   2864  ND1 HIS   366     -19.669  24.434  -9.681  1.00  0.00           N
ATOM   2865  CG  HIS   366     -19.882  23.903  -8.919  1.00  0.00           C
ATOM   2866  CB  HIS   366     -20.278  23.607  -7.529  1.00  0.00           C
ATOM   2867  NE2 HIS   366     -19.318  24.169 -10.885  1.00  0.00           N
ATOM   2868  CD2 HIS   366     -19.655  23.738  -9.698  1.00  0.00           C
ATOM   2869  CE1 HIS   366     -19.336  24.583 -10.838  1.00  0.00           C
ATOM   2870  C   HIS   366     -21.064  24.256  -5.428  1.00  0.00           C
ATOM   2871  O   HIS   366     -20.990  23.056  -5.203  1.00  0.00           O
ATOM   2872  N   ILE   367     -21.576  25.038  -4.596  1.00  0.00           N
ATOM   2873  CA  ILE   367     -21.508  24.215  -2.383 33.67  0.00           C
ATOM   2874  CB  ILE   367     -22.510  25.792  -2.310  1.00  0.00           C
ATOM   2875  CG2 ILE   367     -22.258  26.140  -1.476  1.00  0.00           C
ATOM   2876  CG1 ILE   367     -22.703  26.738  -2.393  1.00  0.00           C
ATOM   2877  CD1 ILE   367     -23.382  27.776  -1.560  1.00  0.00           C
ATOM   2878  C   ILE   367     -22.945  23.646  -3.138  1.00  0.00           C
ATOM   2879  O   ILE   367     -23.017  23.014  -2.146  1.00  0.00           O
ATOM   2880  N   GLU   368     -23.748  23.414  -4.164  1.00  0.00           N
ATOM   2881  CA  GLU   368     -24.038  21.603  -3.127 33.67  0.00           C
ATOM   2882  CB  GLU   368     -25.837  22.475  -5.188  1.00  0.00           C
ATOM   2883  CG  GLU   368     -26.999  22.529  -4.802  1.00  0.00           C
ATOM   2884  CD  GLU   368     -27.992  22.639  -5.932  1.00  0.00           C
ATOM   2885  OE1 GLU   368     -27.720  22.689  -6.940  1.00  0.00           O
ATOM   2886  OE2 GLU   368     -29.035  22.675  -5.803  1.00  0.00           O
ATOM   2887  C   GLU   368     -24.140  21.074  -4.309  1.00  0.00           C
ATOM   2888  O   GLU   368     -24.391  20.175  -3.633  1.00  0.00           O
ATOM   2889  N   ASP   369     -23.293  20.961  -5.214  1.00  0.00           N
ATOM   2890  CA  ASP   369     -21.741  19.294  -4.555 33.67  0.00           C
ATOM   2891  CB  ASP   369     -22.049  19.663  -6.557  1.00  0.00           C
ATOM   2892  CG  ASP   369     -23.103  19.635  -7.599  1.00  0.00           C
ATOM   2893  OD1 ASP   369     -24.210  19.445  -7.304  1.00  0.00           O
ATOM   2894  OD2 ASP   369     -22.816  19.803  -8.705  1.00  0.00           O
ATOM   2895  C   ASP   369     -21.706  19.386  -4.230  1.00  0.00           C
ATOM   2896  O   ASP   369     -21.579  18.252  -3.838  1.00  0.00           O
ATOM   2897  N   THR   370     -21.036  20.343  -3.724  1.00  0.00           N
ATOM   2898  CA  THR   370     -19.101  19.995  -2.052 33.67  0.00           C
ATOM   2899  CB  THR   370     -19.357  21.342  -2.312  1.00  0.00           C
ATOM   2900  OG1 THR   370     -18.603  21.749  -3.417  1.00  0.00           O
ATOM   2901  CG2 THR   370     -18.421  21.056  -1.149  1.00  0.00           C
ATOM   2902  C   THR   370     -20.798  19.582  -1.408  1.00  0.00           C
ATOM   2903  O   THR   370     -20.288  18.710  -0.701  1.00  0.00           O
ATOM   2904  N   GLU   371     -21.949  20.130  -1.151  1.00  0.00           N
ATOM   2905  CA  GLU   371     -21.428  19.150   0.639 33.67  0.00           C
ATOM   2906  CB  GLU   371     -23.975  20.455   0.028  1.00  0.00           C
ATOM   2907  CG  GLU   371     -24.813  19.979   1.136  1.00  0.00           C
ATOM   2908  CD  GLU   371     -25.964  20.920   1.194  1.00  0.00           C
ATOM   2909  OE1 GLU   371     -26.035  21.882   0.392  1.00  0.00           O
ATOM   2910  OE2 GLU   371     -26.787  20.690   2.040  1.00  0.00           O
ATOM   2911  C   GLU   371     -23.088  18.198  -0.378  1.00  0.00           C
ATOM   2912  O   GLU   371     -23.026  17.353   0.505  1.00  0.00           O
ATOM   2913  N   ARG   372     -23.436  17.944  -1.599  1.00  0.00           N
ATOM   2914  CA  ARG   372     -22.142  16.086  -1.257 33.67  0.00           C
ATOM   2915  CB  ARG   372     -24.141  16.567  -3.444  1.00  0.00           C
ATOM   2916  CG  ARG   372     -25.498  17.179  -3.739  1.00  0.00           C
ATOM   2917  CD  ARG   372     -25.883  17.208  -5.190  1.00  0.00           C
ATOM   2918  NE  ARG   372     -27.202  17.849  -5.328  1.00  0.00           N
ATOM   2919  CZ  ARG   372     -27.751  18.093  -6.535  1.00  0.00           C
ATOM   2920  NH1 ARG   372     -26.890  17.656  -7.427  1.00  0.00           N
ATOM   2921  NH2 ARG   372     -28.947  18.680  -6.377  1.00  0.00           N
ATOM   2922  C   ARG   372     -22.556  15.676  -1.751  1.00  0.00           C
ATOM   2923  O   ARG   372     -22.699  14.559  -1.398  1.00  0.00           O
ATOM   2924  N   VAL   373     -21.401  16.133  -1.940  1.00  0.00           N
ATOM   2925  CA  VAL   373     -18.550  15.333  -1.315 33.67  0.00           C
ATOM   2926  CB  VAL   373     -19.019  15.929  -2.387  1.00  0.00           C
ATOM   2927  CG1 VAL   373     -17.765  15.216  -1.968  1.00  0.00           C
ATOM   2928  CG2 VAL   373     -19.096  15.909  -3.876  1.00  0.00           C
ATOM   2929  C   VAL   373     -19.947  15.034  -0.315  1.00  0.00           C
ATOM   2930  O   VAL   373     -19.583  13.935   0.099  1.00  0.00           O
ATOM   2931  N   LEU   374     -20.122  16.040   0.443  1.00  0.00           N
ATOM   2932  CA  LEU   374     -18.255  15.764   2.310 33.67  0.00           C
ATOM   2933  CB  LEU   374     -20.238  17.137   2.493  1.00  0.00           C
ATOM   2934  CG  LEU   374     -20.120  17.009   3.953  1.00  0.00           C
ATOM   2935  CD1 LEU   374     -18.784  16.554   4.482  1.00  0.00           C
ATOM   2936  CD2 LEU   374     -20.410  18.267   4.656  1.00  0.00           C
ATOM   2937  C   LEU   374     -20.893  14.831   2.316  1.00  0.00           C
ATOM   2938  O   LEU   374     -20.487  13.882   2.983  1.00  0.00           O
ATOM   2939  N   GLN   375     -23.622  13.486   0.169  1.00  0.00           N
ATOM   2940  CA  GLN   375     -20.988  12.260   1.480 50.00  0.00           C
ATOM   2941  CB  GLN   375     -25.711  12.688  -0.837  1.00  0.00           C
ATOM   2942  CG  GLN   375     -26.731  11.554  -0.953  1.00  0.00           C
ATOM   2943  CD  GLN   375     -27.866  12.032  -1.848  1.00  0.00           C
ATOM   2944  OE1 GLN   375     -27.807  13.120  -2.420  1.00  0.00           O
ATOM   2945  NE2 GLN   375     -28.962  11.245  -2.018  1.00  0.00           N
ATOM   2946  C   GLN   375     -23.769  11.145  -0.498  1.00  0.00           C
ATOM   2947  O   GLN   375     -24.007  10.011  -0.083  1.00  0.00           O
ATOM   2948  N   LEU   376     -22.137  12.678   0.769  1.00  0.00           N
ATOM   2949  CA  LEU   376     -19.375  11.276   0.600 33.67  0.00           C
ATOM   2950  CB  LEU   376     -21.101  11.417  -0.973  1.00  0.00           C
ATOM   2951  CG  LEU   376     -20.645  10.099  -1.351  1.00  0.00           C
ATOM   2952  CD1 LEU   376     -21.700   9.146  -1.492  1.00  0.00           C
ATOM   2953  CD2 LEU   376     -20.009  10.061  -2.631  1.00  0.00           C
ATOM   2954  C   LEU   376     -20.659  10.769   1.230  1.00  0.00           C
ATOM   2955  O   LEU   376     -20.646   9.636   1.532  1.00  0.00           O
ATOM   2956  N   ALA   377     -19.785  11.483   1.658  1.00  0.00           N
ATOM   2957  CA  ALA   377     -16.376  11.178   2.692 33.67  0.00           C
ATOM   2958  CB  ALA   377     -17.820  11.887   2.824  1.00  0.00           C
ATOM   2959  C   ALA   377     -19.306  10.464   3.756  1.00  0.00           C
ATOM   2960  O   ALA   377     -18.993   9.377   4.173  1.00  0.00           O
ATOM   2961  N   LYS   378     -21.565   9.684   2.500  1.00  0.00           N
ATOM   2962  CA  LYS   378     -17.556   8.909   4.416 50.00  0.00           C
ATOM   2963  CB  LYS   378     -23.382   9.945   4.146  1.00  0.00           C
ATOM   2964  CG  LYS   378     -23.901   9.458   5.502  1.00  0.00           C
ATOM   2965  CD  LYS   378     -25.228  10.129   5.891  1.00  0.00           C
ATOM   2966  CE  LYS   378     -26.030   9.279   6.891  1.00  0.00           C
ATOM   2967  NZ  LYS   378     -27.457   9.716   6.980  1.00  0.00           N
ATOM   2968  C   LYS   378     -23.039   7.765   2.908  1.00  0.00           C
ATOM   2969  O   LYS   378     -23.240   6.756   3.591  1.00  0.00           O
ATOM   2970  N   GLU   379     -23.389   7.854   1.629  1.00  0.00           N
ATOM   2971  CA  GLU   379     -19.450   6.658   1.973 50.00  0.00           C
ATOM   2972  CB  GLU   379     -24.257   7.098  -0.539  1.00  0.00           C
ATOM   2973  CG  GLU   379     -24.998   6.011  -1.321  1.00  0.00           C
ATOM   2974  CD  GLU   379     -25.189   6.506  -2.747  1.00  0.00           C
ATOM   2975  OE1 GLU   379     -24.746   7.648  -3.042  1.00  0.00           O
ATOM   2976  OE2 GLU   379     -25.781   5.748  -3.562  1.00  0.00           O
ATOM   2977  C   GLU   379     -23.232   5.465   1.057  1.00  0.00           C
ATOM   2978  O   GLU   379     -23.463   4.658   1.958  1.00  0.00           O
ATOM   2979  N   ARG   380     -22.283   5.280   0.146  1.00  0.00           N
ATOM   2980  CA  ARG   380     -16.491   4.544   1.038 50.00  0.00           C
ATOM   2981  CB  ARG   380     -20.049   4.421  -0.403  1.00  0.00           C
ATOM   2982  CG  ARG   380     -19.122   3.206  -0.485  1.00  0.00           C
ATOM   2983  CD  ARG   380     -17.762   3.515  -1.115  1.00  0.00           C
ATOM   2984  NE  ARG   380     -16.997   2.238  -1.176  1.00  0.00           N
ATOM   2985  CZ  ARG   380     -16.278   1.820  -0.094  1.00  0.00           C
ATOM   2986  NH1 ARG   380     -16.435   2.750   0.893  1.00  0.00           N
ATOM   2987  NH2 ARG   380     -15.683   0.636  -0.421  1.00  0.00           N
ATOM   2988  C   ARG   380     -21.316   3.506   1.553  1.00  0.00           C
ATOM   2989  O   ARG   380     -22.049   2.587   1.917  1.00  0.00           O
ATOM   2990  N   GLY   381     -20.388   4.045   2.336  1.00  0.00           N
ATOM   2991  CA  GLY   381     -14.456   4.418   4.130 50.00  0.00           C
ATOM   2992  C   GLY   381     -19.175   4.438   4.448  1.00  0.00           C
ATOM   2993  O   GLY   381     -19.272   4.592   5.666  1.00  0.00           O
ATOM   2994  N   CYS   382     -16.881   6.712   5.610  1.00  0.00           N
ATOM   2995  CA  CYS   382     -12.970   7.914   5.364 33.67  0.00           C
ATOM   2996  CB  CYS   382     -15.274   8.194   4.470  1.00  0.00           C
ATOM   2997  SG  CYS   382     -14.107   8.765   4.620  1.00  0.00           S
ATOM   2998  C   CYS   382     -15.822   8.372   6.603  1.00  0.00           C
ATOM   2999  O   CYS   382     -16.689   9.092   6.622  1.00  0.00           O
ATOM   3000  N   THR   383     -17.304   7.287   6.297  1.00  0.00           N
ATOM   3001  CA  THR   383     -12.609   9.475   7.552 50.00  0.00           C
ATOM   3002  CB  THR   383     -18.118   7.782   8.583  1.00  0.00           C
ATOM   3003  OG1 THR   383     -19.058   6.719   8.522  1.00  0.00           O
ATOM   3004  CG2 THR   383     -18.678   8.906   9.470  1.00  0.00           C
ATOM   3005  C   THR   383     -16.944   9.568   7.249  1.00  0.00           C
ATOM   3006  O   THR   383     -15.739   9.457   7.513  1.00  0.00           O
ATOM   3007  N   PRO   384     -17.530  10.755   7.036  1.00  0.00           N
ATOM   3008  CA  PRO   384     -12.286  13.182   7.042 50.00  0.00           C
ATOM   3009  CD  PRO   384     -18.986  10.938   6.947  1.00  0.00           C
ATOM   3010  CB  PRO   384     -17.944  13.042   6.704  1.00  0.00           C
ATOM   3011  CG  PRO   384     -19.103  12.227   6.221  1.00  0.00           C
ATOM   3012  C   PRO   384     -16.012  12.400   8.165  1.00  0.00           C
ATOM   3013  O   PRO   384     -16.332  11.907   9.253  1.00  0.00           O
ATOM   3014  N   ASN   385     -14.993  13.246   8.014  1.00  0.00           N
ATOM   3015  CA  ASN   385      -9.918  15.369   9.009 50.00  0.00           C
ATOM   3016  CB  ASN   385     -12.728  13.351   9.038  1.00  0.00           C
ATOM   3017  CG  ASN   385     -12.185  14.060   7.805  1.00  0.00           C
ATOM   3018  OD1 ASN   385     -12.876  14.864   7.182  1.00  0.00           O
ATOM   3019  ND2 ASN   385     -10.917  13.800   7.387  1.00  0.00           N
ATOM   3020  C   ASN   385     -14.378  15.190   9.430  1.00  0.00           C
ATOM   3021  O   ASN   385     -15.342  15.804   9.016  1.00  0.00           O
ATOM   3022  N   GLY   386     -13.415  15.772  10.121  1.00  0.00           N
ATOM   3023  CA  GLY   386      -9.802  18.918  10.233 50.00  0.00           C
ATOM   3024  C   GLY   386     -13.260  17.994   9.120  1.00  0.00           C
ATOM   3025  O   GLY   386     -13.944  18.980   8.924  1.00  0.00           O
ATOM   3026  N   ALA   387     -12.325  17.579   8.263  1.00  0.00           N
ATOM   3027  CA  ALA   387      -8.940  20.291   6.822 50.00  0.00           C
ATOM   3028  CB  ALA   387     -10.903  17.679   6.235  1.00  0.00           C
ATOM   3029  C   ALA   387     -13.304  18.437   6.204  1.00  0.00           C
ATOM   3030  O   ALA   387     -13.545  19.450   5.539  1.00  0.00           O
ATOM   3031  N   THR   388     -14.150  17.415   6.303  1.00  0.00           N
ATOM   3032  CA  THR   388     -12.094  18.725   5.582 50.00  0.00           C
ATOM   3033  CB  THR   388     -16.047  16.002   5.608  1.00  0.00           C
ATOM   3034  OG1 THR   388     -15.016  15.012   5.676  1.00  0.00           O
ATOM   3035  CG2 THR   388     -16.871  15.786   4.374  1.00  0.00           C
ATOM   3036  C   THR   388     -16.439  18.371   6.126  1.00  0.00           C
ATOM   3037  O   THR   388     -16.928  19.249   5.407  1.00  0.00           O
ATOM   3038  N   LEU   389     -16.765  18.206   7.407  1.00  0.00           N
ATOM   3039  CA  LEU   389     -14.408  19.951   8.285 50.00  0.00           C
ATOM   3040  CB  LEU   389     -18.048  18.503   9.486  1.00  0.00           C
ATOM   3041  CG  LEU   389     -18.757  17.148   9.505  1.00  0.00           C
ATOM   3042  CD1 LEU   389     -18.906  16.516  10.889  1.00  0.00           C
ATOM   3043  CD2 LEU   389     -20.187  17.159   8.965  1.00  0.00           C
ATOM   3044  C   LEU   389     -17.369  20.514   8.093  1.00  0.00           C
ATOM   3045  O   LEU   389     -18.222  21.409   8.063  1.00  0.00           O
ATOM   3046  N   SER   390     -16.062  20.756   8.139  1.00  0.00           N
ATOM   3047  CA  SER   390     -12.913  23.352   8.227 50.00  0.00           C
ATOM   3048  CB  SER   390     -14.061  22.121   8.615  1.00  0.00           C
ATOM   3049  OG  SER   390     -14.013  21.734   9.980  1.00  0.00           O
ATOM   3050  C   SER   390     -15.595  22.675   6.714  1.00  0.00           C
ATOM   3051  O   SER   390     -15.846  23.873   6.540  1.00  0.00           O
ATOM   3052  N   ARG   391     -15.395  21.816   5.717  1.00  0.00           N
ATOM   3053  CA  ARG   391     -13.390  23.459   4.507 50.00  0.00           C
ATOM   3054  CB  ARG   391     -15.332  21.101   3.361  1.00  0.00           C
ATOM   3055  CG  ARG   391     -13.856  20.700   3.336  1.00  0.00           C
ATOM   3056  CD  ARG   391     -12.937  21.782   2.764  1.00  0.00           C
ATOM   3057  NE  ARG   391     -11.563  21.211   2.691  1.00  0.00           N
ATOM   3058  CZ  ARG   391     -10.511  22.005   2.337  1.00  0.00           C
ATOM   3059  NH1 ARG   391     -10.996  23.261   2.109  1.00  0.00           N
ATOM   3060  NH2 ARG   391      -9.385  21.235   2.350  1.00  0.00           N
ATOM   3061  C   ARG   391     -17.079  22.681   4.179  1.00  0.00           C
ATOM   3062  O   ARG   391     -17.352  23.725   3.560  1.00  0.00           O
ATOM   3063  N   VAL   392     -17.988  21.974   4.829  1.00  0.00           N
ATOM   3064  CA  VAL   392     -17.027  22.870   5.198 50.00  0.00           C
ATOM   3065  CB  VAL   392     -20.781  21.912   5.364  1.00  0.00           C
ATOM   3066  CG1 VAL   392     -21.910  22.864   4.965  1.00  0.00           C
ATOM   3067  CG2 VAL   392     -21.255  20.524   4.927  1.00  0.00           C
ATOM   3068  C   VAL   392     -19.639  23.663   5.477  1.00  0.00           C
ATOM   3069  O   VAL   392     -20.321  24.549   4.955  1.00  0.00           O
ATOM   3070  N   LEU   393     -19.069  23.832   6.668  1.00  0.00           N
ATOM   3071  CA  LEU   393     -17.313  25.522   7.867 50.00  0.00           C
ATOM   3072  CB  LEU   393     -18.615  25.279   8.788  1.00  0.00           C
ATOM   3073  CG  LEU   393     -19.223  24.392   9.876  1.00  0.00           C
ATOM   3074  CD1 LEU   393     -18.507  24.442  11.225  1.00  0.00           C
ATOM   3075  CD2 LEU   393     -20.674  24.711  10.233  1.00  0.00           C
ATOM   3076  C   LEU   393     -18.664  26.294   6.699  1.00  0.00           C
ATOM   3077  O   LEU   393     -19.184  27.408   6.800  1.00  0.00           O
ATOM   3078  N   LYS   394     -17.564  26.110   5.962  1.00  0.00           N
ATOM   3079  CA  LYS   394     -15.723  28.063   5.682 50.00  0.00           C
ATOM   3080  CB  LYS   394     -15.757  27.057   4.350  1.00  0.00           C
ATOM   3081  CG  LYS   394     -15.216  28.365   3.770  1.00  0.00           C
ATOM   3082  CD  LYS   394     -13.982  28.179   2.885  1.00  0.00           C
ATOM   3083  CE  LYS   394     -13.470  29.481   2.266  1.00  0.00           C
ATOM   3084  NZ  LYS   394     -12.295  29.207   1.409  1.00  0.00           N
ATOM   3085  C   LYS   394     -17.890  28.000   4.331  1.00  0.00           C
ATOM   3086  O   LYS   394     -17.962  29.232   4.339  1.00  0.00           O
ATOM   3087  N   LYS   395     -16.374  26.803   2.506  1.00  0.00           N
ATOM   3088  CA  LYS   395     -17.182  27.066   2.272 33.67  0.00           C
ATOM   3089  CB  LYS   395     -17.865  25.957   1.149  1.00  0.00           C
ATOM   3090  CG  LYS   395     -18.950  26.220   0.355  1.00  0.00           C
ATOM   3091  CD  LYS   395     -18.724  26.045   0.062  1.00  0.00           C
ATOM   3092  CE  LYS   395     -19.280  26.226   0.307  1.00  0.00           C
ATOM   3093  NZ  LYS   395     -19.031  26.002  -0.057  1.00  0.00           N
ATOM   3094  C   LYS   395     -18.695  27.611   2.344  1.00  0.00           C
ATOM   3095  O   LYS   395     -19.375  28.465   1.853  1.00  0.00           O
ATOM   3096  N   SER   396     -21.397  27.388   3.961  1.00  0.00           N
ATOM   3097  CA  SER   396     -21.063  27.257   5.408 50.00  0.00           C
ATOM   3098  CB  SER   396     -23.413  26.756   5.603  1.00  0.00           C
ATOM   3099  OG  SER   396     -23.719  25.484   5.051  1.00  0.00           O
ATOM   3100  C   SER   396     -22.660  28.898   5.436  1.00  0.00           C
ATOM   3101  O   SER   396     -23.675  29.562   5.661  1.00  0.00           O
ATOM   3102  N   ASN   397     -21.478  29.245   5.935  1.00  0.00           N
ATOM   3103  CA  ASN   397     -20.237  30.348   7.561 50.00  0.00           C
ATOM   3104  CB  ASN   397     -19.954  30.540   7.445  1.00  0.00           C
ATOM   3105  CG  ASN   397     -19.885  29.536   8.586  1.00  0.00           C
ATOM   3106  OD1 ASN   397     -20.627  29.634   9.562  1.00  0.00           O
ATOM   3107  ND2 ASN   397     -18.989  28.514   8.526  1.00  0.00           N
ATOM   3108  C   ASN   397     -21.568  31.708   6.018  1.00  0.00           C
ATOM   3109  O   ASN   397     -22.193  32.654   6.506  1.00  0.00           O
ATOM   3110  N   ALA   398     -19.705  31.376   2.971  1.00  0.00           N
ATOM   3111  CA  ALA   398     -20.398  32.062   3.130 33.67  0.00           C
ATOM   3112  CB  ALA   398     -19.975  32.249   0.888  1.00  0.00           C
ATOM   3113  C   ALA   398     -21.729  32.093   2.348  1.00  0.00           C
ATOM   3114  O   ALA   398     -22.527  32.642   2.092  1.00  0.00           O
ATOM   3115  N   ASP   399     -22.007  31.291   2.697  1.00  0.00           N
ATOM   3116  CA  ASP   399     -23.162  30.229   3.716 33.67  0.00           C
ATOM   3117  CB  ASP   399     -23.251  30.369   2.624  1.00  0.00           C
ATOM   3118  CG  ASP   399     -23.655  29.935   2.173  1.00  0.00           C
ATOM   3119  OD1 ASP   399     -24.006  30.174   1.995  1.00  0.00           O
ATOM   3120  OD2 ASP   399     -23.617  29.358   2.000  1.00  0.00           O
ATOM   3121  C   ASP   399     -24.227  31.038   3.499  1.00  0.00           C
ATOM   3122  O   ASP   399     -24.365  31.143   3.428  1.00  0.00           O
ATOM   3123  N   ARG   400     -24.933  30.959   4.216  1.00  0.00           N
ATOM   3124  CA  ARG   400     -25.491  29.784   6.103 33.67  0.00           C
ATOM   3125  CB  ARG   400     -26.736  30.134   4.361  1.00  0.00           C
ATOM   3126  CG  ARG   400     -27.460  30.419   3.077  1.00  0.00           C
ATOM   3127  CD  ARG   400     -28.362  29.739   2.529  1.00  0.00           C
ATOM   3128  NE  ARG   400     -28.983  30.098   1.280  1.00  0.00           N
ATOM   3129  CZ  ARG   400     -29.811  29.620   0.553  1.00  0.00           C
ATOM   3130  NH1 ARG   400     -29.893  28.850   1.189  1.00  0.00           N
ATOM   3131  NH2 ARG   400     -30.234  30.099  -0.527  1.00  0.00           N
ATOM   3132  C   ARG   400     -25.356  30.457   6.257  1.00  0.00           C
ATOM   3133  O   ARG   400     -24.868  29.407   6.525  1.00  0.00           O
ATOM   3134  N   PHE   401     -25.517  31.287   7.133  1.00  0.00           N
ATOM   3135  CA  PHE   401     -24.388  29.992   9.518 33.67  0.00           C
ATOM   3136  CB  PHE   401     -25.159  31.937   9.214  1.00  0.00           C
ATOM   3137  CG  PHE   401     -25.512  32.166  10.024  1.00  0.00           C
ATOM   3138  CD1 PHE   401     -25.321  32.363  10.968  1.00  0.00           C
ATOM   3139  CD2 PHE   401     -26.052  32.190   9.858  1.00  0.00           C
ATOM   3140  CE1 PHE   401     -25.662  32.583  11.739  1.00  0.00           C
ATOM   3141  CE2 PHE   401     -26.401  32.409  10.618  1.00  0.00           C
ATOM   3142  CZ  PHE   401     -26.205  32.605  11.562  1.00  0.00           C
ATOM   3143  C   PHE   401     -25.649  29.896   9.080  1.00  0.00           C
ATOM   3144  O   PHE   401     -25.053  29.382   9.967  1.00  0.00           O
ATOM   3145  N   LEU   402     -26.834  29.546   8.650  1.00  0.00           N
ATOM   3146  CA  LEU   402     -26.498  27.212  10.395 33.67  0.00           C
ATOM   3147  CB  LEU   402     -28.852  28.375   8.755  1.00  0.00           C
ATOM   3148  CG  LEU   402     -29.202  28.360   9.134  1.00  0.00           C
ATOM   3149  CD1 LEU   402     -30.104  28.423   9.759  1.00  0.00           C
ATOM   3150  CD2 LEU   402     -29.687  28.229   9.495  1.00  0.00           C
ATOM   3151  C   LEU   402     -26.882  27.187   8.844  1.00  0.00           C
ATOM   3152  O   LEU   402     -26.800  26.286   9.654  1.00  0.00           O
ATOM   3153  N   GLN   403     -26.450  27.104   7.629  1.00  0.00           N
ATOM   3154  CA  GLN   403     -24.731  24.957   8.185 33.67  0.00           C
ATOM   3155  CB  GLN   403     -25.577  25.935   5.789  1.00  0.00           C
ATOM   3156  CG  GLN   403     -26.741  25.965   4.952  1.00  0.00           C
ATOM   3157  CD  GLN   403     -26.334  26.013   3.596  1.00  0.00           C
ATOM   3158  OE1 GLN   403     -25.616  26.524   3.130  1.00  0.00           O
ATOM   3159  NE2 GLN   403     -26.770  25.483   2.888  1.00  0.00           N
ATOM   3160  C   GLN   403     -24.573  25.607   7.900  1.00  0.00           C
ATOM   3161  O   GLN   403     -24.238  24.485   8.246  1.00  0.00           O
ATOM   3162  N   ALA   404     -23.860  26.618   8.143  1.00  0.00           N
ATOM   3163  CA  ALA   404     -21.525  25.932   9.697 33.67  0.00           C
ATOM   3164  CB  ALA   404     -22.012  27.808   9.013  1.00  0.00           C
ATOM   3165  C   ALA   404     -22.907  25.973  10.278  1.00  0.00           C
ATOM   3166  O   ALA   404     -22.150  25.158  10.813  1.00  0.00           O
ATOM   3167  N   GLU   405     -23.985  26.437  10.857  1.00  0.00           N
ATOM   3168  CA  GLU   405     -22.892  25.247  13.042 33.67  0.00           C
ATOM   3169  CB  GLU   405     -25.435  26.767  12.724  1.00  0.00           C
ATOM   3170  CG  GLU   405     -25.524  26.479  14.121  1.00  0.00           C
ATOM   3171  CD  GLU   405     -26.865  26.829  14.716  1.00  0.00           C
ATOM   3172  OE1 GLU   405     -27.125  27.427  14.977  1.00  0.00           O
ATOM   3173  OE2 GLU   405     -27.650  26.507  14.921  1.00  0.00           O
ATOM   3174  C   GLU   405     -24.689  24.614  12.224  1.00  0.00           C
ATOM   3175  O   GLU   405     -24.372  23.892  13.164  1.00  0.00           O
ATOM   3176  N   LEU   406     -25.370  24.218  11.207  1.00  0.00           N
ATOM   3177  CA  LEU   406     -24.073  21.956  11.987 33.67  0.00           C
ATOM   3178  CB  LEU   406     -26.587  22.545   9.903  1.00  0.00           C
ATOM   3179  CG  LEU   406     -27.698  22.030   9.858  1.00  0.00           C
ATOM   3180  CD1 LEU   406     -28.375  21.776   9.804  1.00  0.00           C
ATOM   3181  CD2 LEU   406     -28.682  21.520   9.770  1.00  0.00           C
ATOM   3182  C   LEU   406     -24.582  21.968  11.064  1.00  0.00           C
ATOM   3183  O   LEU   406     -24.499  20.949  11.720  1.00  0.00           O
ATOM   3184  N   ILE   407     -23.639  22.345  10.274  1.00  0.00           N
ATOM   3185  CA  ILE   407     -20.710  21.117  10.783 33.67  0.00           C
ATOM   3186  CB  ILE   407     -21.348  21.967   9.273  1.00  0.00           C
ATOM   3187  CG2 ILE   407     -20.142  21.044   9.401  1.00  0.00           C
ATOM   3188  CG1 ILE   407     -21.789  21.950   7.899  1.00  0.00           C
ATOM   3189  CD1 ILE   407     -20.824  22.430   7.000  1.00  0.00           C
ATOM   3190  C   ILE   407     -21.663  21.494  11.499  1.00  0.00           C
ATOM   3191  O   ILE   407     -21.216  20.435  11.921  1.00  0.00           O
ATOM   3192  N   LEU   408     -21.520  22.606  12.150  1.00  0.00           N
ATOM   3193  CA  LEU   408     -19.084  22.306  13.946 33.67  0.00           C
ATOM   3194  CB  LEU   408     -20.818  24.006  14.095  1.00  0.00           C
ATOM   3195  CG  LEU   408     -19.736  24.837  14.246  1.00  0.00           C
ATOM   3196  CD1 LEU   408     -19.978  25.805  15.303  1.00  0.00           C
ATOM   3197  CD2 LEU   408     -18.456  24.121  14.376  1.00  0.00           C
ATOM   3198  C   LEU   408     -21.567  21.805  14.465  1.00  0.00           C
ATOM   3199  O   LEU   408     -20.956  21.166  15.317  1.00  0.00           O
ATOM   3200  N   PRO   409     -22.838  21.830  14.425  1.00  0.00           N
ATOM   3201  CA  PRO   409     -21.604  20.447  15.902 33.67  0.00           C
ATOM   3202  CD  PRO   409     -24.153  23.191  13.376  1.00  0.00           C
ATOM   3203  CB  PRO   409     -25.114  21.647  15.164  1.00  0.00           C
ATOM   3204  CG  PRO   409     -25.222  22.686  14.172  1.00  0.00           C
ATOM   3205  C   PRO   409     -23.665  19.604  15.024  1.00  0.00           C
ATOM   3206  O   PRO   409     -23.634  18.797  15.946  1.00  0.00           O
ATOM   3207  N   ASP   410     -24.457  19.170  14.048  1.00  0.00           N
ATOM   3208  CA  ASP   410     -22.170  17.098  14.305 33.67  0.00           C
ATOM   3209  CB  ASP   410     -25.656  17.378  12.841  1.00  0.00           C
ATOM   3210  CG  ASP   410     -26.889  17.455  13.620  1.00  0.00           C
ATOM   3211  OD1 ASP   410     -26.793  17.572  14.816  1.00  0.00           O
ATOM   3212  OD2 ASP   410     -27.943  17.399  13.030  1.00  0.00           O
ATOM   3213  C   ASP   410     -23.337  17.289  12.984  1.00  0.00           C
ATOM   3214  O   ASP   410     -22.835  17.090  12.665  1.00  0.00           O
ATOM   3215  N   ALA   411     -22.867  17.115  12.713  1.00  0.00           N
ATOM   3216  CA  ALA   411     -19.332  16.406  12.399 33.67  0.00           C
ATOM   3217  CB  ALA   411     -21.644  17.402  10.825  1.00  0.00           C
ATOM   3218  C   ALA   411     -20.511  16.791  12.739  1.00  0.00           C
ATOM   3219  O   ALA   411     -19.730  16.285  12.405  1.00  0.00           O
ATOM   3220  N   LEU   412     -20.353  17.476  13.726  1.00  0.00           N
ATOM   3221  CA  LEU   412     -16.814  17.770  14.666 33.67  0.00           C
ATOM   3222  CB  LEU   412     -18.983  19.043  15.065  1.00  0.00           C
ATOM   3223  CG  LEU   412     -18.627  19.676  15.157  1.00  0.00           C
ATOM   3224  CD1 LEU   412     -17.748  20.350  15.203  1.00  0.00           C
ATOM   3225  CD2 LEU   412     -18.292  20.310  14.910  1.00  0.00           C
ATOM   3226  C   LEU   412     -18.970  16.769  15.505  1.00  0.00           C
ATOM   3227  O   LEU   412     -17.882  16.548  15.953  1.00  0.00           O
ATOM   3228  N   GLU   413     -20.036  16.191  15.960  1.00  0.00           N
ATOM   3229  CA  GLU   413     -17.384  15.270  17.338 33.67  0.00           C
ATOM   3230  CB  GLU   413     -21.329  14.703  17.510  1.00  0.00           C
ATOM   3231  CG  GLU   413     -21.467  13.919  17.629  1.00  0.00           C
ATOM   3232  CD  GLU   413     -22.801  13.383  18.043  1.00  0.00           C
ATOM   3233  OE1 GLU   413     -23.530  13.650  18.192  1.00  0.00           O
ATOM   3234  OE2 GLU   413     -23.107  12.699  18.215  1.00  0.00           O
ATOM   3235  C   GLU   413     -19.131  14.145  16.997  1.00  0.00           C
ATOM   3236  O   GLU   413     -18.692  13.634  17.989  1.00  0.00           O
ATOM   3237  N   ALA   414     -18.836  13.684  15.794  1.00  0.00           N
ATOM   3238  CA  ALA   414     -15.098  12.883  15.580 33.67  0.00           C
ATOM   3239  CB  ALA   414     -18.129  11.979  14.206  1.00  0.00           C
ATOM   3240  C   ALA   414     -16.472  13.012  15.661  1.00  0.00           C
ATOM   3241  O   ALA   414     -15.605  12.190  15.846  1.00  0.00           O
ATOM   3242  N   GLY   415     -16.183  14.282  15.532  1.00  0.00           N
ATOM   3243  CA  GLY   415     -12.189  15.230  15.411 33.67  0.00           C
ATOM   3244  C   GLY   415     -14.581  16.034  16.193  1.00  0.00           C
ATOM   3245  O   GLY   415     -14.040  16.910  15.886  1.00  0.00           O
ATOM   3246  N   ASP   416     -14.887  16.195  17.161  1.00  0.00           N
ATOM   3247  CA  ASP   416     -12.382  17.957  17.770 33.67  0.00           C
ATOM   3248  CB  ASP   416     -13.868  17.523  18.866  1.00  0.00           C
ATOM   3249  CG  ASP   416     -14.128  17.755  19.762  1.00  0.00           C
ATOM   3250  OD1 ASP   416     -15.103  17.976  19.513  1.00  0.00           O
ATOM   3251  OD2 ASP   416     -13.353  17.717  20.706  1.00  0.00           O
ATOM   3252  C   ASP   416     -14.523  18.535  16.709  1.00  0.00           C
ATOM   3253  O   ASP   416     -13.658  18.328  15.867  1.00  0.00           O
ATOM   3254  N   VAL   417     -15.230  19.657  16.730  1.00  0.00           N
ATOM   3255  CA  VAL   417     -12.966  21.089  15.915 33.67  0.00           C
ATOM   3256  CB  VAL   417     -16.076  21.667  15.857  1.00  0.00           C
ATOM   3257  CG1 VAL   417     -15.983  22.760  16.126  1.00  0.00           C
ATOM   3258  CG2 VAL   417     -16.777  21.908  14.963  1.00  0.00           C
ATOM   3259  C   VAL   417     -13.618  21.442  16.269  1.00  0.00           C
ATOM   3260  O   VAL   417     -13.311  21.436  17.458  1.00  0.00           O
ATOM   3261  N   ARG   418     -12.882  22.024  15.337  1.00  0.00           N
ATOM   3262  CA  ARG   418     -10.149  23.501  15.705 33.67  0.00           C
ATOM   3263  CB  ARG   418     -10.566  22.471  14.870  1.00  0.00           C
ATOM   3264  CG  ARG   418     -10.238  21.037  14.908  1.00  0.00           C
ATOM   3265  CD  ARG   418      -9.461  20.346  15.245  1.00  0.00           C
ATOM   3266  NE  ARG   418      -8.922  19.123  15.384  1.00  0.00           N
ATOM   3267  CZ  ARG   418      -9.350  17.945  15.250  1.00  0.00           C
ATOM   3268  NH1 ARG   418     -10.268  17.957  15.035  1.00  0.00           N
ATOM   3269  NH2 ARG   418      -8.716  16.956  15.435  1.00  0.00           N
ATOM   3270  C   ARG   418     -12.050  24.289  15.587  1.00  0.00           C
ATOM   3271  O   ARG   418     -13.131  24.676  15.145  1.00  0.00           O
ATOM   3272  N   LEU   419     -11.074  25.096  15.960  1.00  0.00           N
ATOM   3273  CA  LEU   419      -9.958  27.230  16.024 33.67  0.00           C
ATOM   3274  CB  LEU   419      -9.912  27.208  16.360  1.00  0.00           C
ATOM   3275  CG  LEU   419      -9.971  28.726  16.361  1.00  0.00           C
ATOM   3276  CD1 LEU   419     -11.022  29.283  17.279  1.00  0.00           C
ATOM   3277  CD2 LEU   419      -8.650  29.354  16.767  1.00  0.00           C
ATOM   3278  C   LEU   419     -11.609  27.038  14.513  1.00  0.00           C
ATOM   3279  O   LEU   419     -12.459  27.905  14.382  1.00  0.00           O
ATOM   3280  N   SER   420     -11.024  26.491  13.484  1.00  0.00           N
ATOM   3281  CA  SER   420     -10.438  27.599  12.288 33.67  0.00           C
ATOM   3282  CB  SER   420     -10.471  26.300  11.101  1.00  0.00           C
ATOM   3283  OG  SER   420      -9.725  25.720  11.021  1.00  0.00           O
ATOM   3284  C   SER   420     -12.812  26.674  11.758  1.00  0.00           C
ATOM   3285  O   SER   420     -13.356  27.468  11.023  1.00  0.00           O
ATOM   3286  N   ASP   421     -13.424  25.608  12.268  1.00  0.00           N
ATOM   3287  CA  ASP   421     -13.612  25.643  12.282 33.67  0.00           C
ATOM   3288  CB  ASP   421     -15.246  23.975  12.520  1.00  0.00           C
ATOM   3289  CG  ASP   421     -14.635  22.947  11.618  1.00  0.00           C
ATOM   3290  OD1 ASP   421     -14.135  23.334  10.530  1.00  0.00           O
ATOM   3291  OD2 ASP   421     -14.660  21.762  12.005  1.00  0.00           O
ATOM   3292  C   ASP   421     -15.726  26.397  12.586  1.00  0.00           C
ATOM   3293  O   ASP   421     -16.697  26.796  11.978  1.00  0.00           O
ATOM   3294  N   ILE   422     -15.396  26.859  13.764  1.00  0.00           N
ATOM   3295  CA  ILE   422     -15.066  27.969  14.815 33.67  0.00           C
ATOM   3296  CB  ILE   422     -15.752  27.973  15.891  1.00  0.00           C
ATOM   3297  CG2 ILE   422     -16.410  29.129  16.620  1.00  0.00           C
ATOM   3298  CG1 ILE   422     -16.116  26.725  16.708  1.00  0.00           C
ATOM   3299  CD1 ILE   422     -16.648  26.713  17.542  1.00  0.00           C
ATOM   3300  C   ILE   422     -15.998  29.249  13.748  1.00  0.00           C
ATOM   3301  O   ILE   422     -16.976  29.953  13.600  1.00  0.00           O
ATOM   3302  N   THR   423     -14.795  29.607  13.328  1.00  0.00           N
ATOM   3303  CA  THR   423     -13.920  31.100  13.108 33.67  0.00           C
ATOM   3304  CB  THR   423     -13.111  30.989  12.224  1.00  0.00           C
ATOM   3305  OG1 THR   423     -12.260  30.942  13.334  1.00  0.00           O
ATOM   3306  CG2 THR   423     -12.884  32.310  11.498  1.00  0.00           C
ATOM   3307  C   THR   423     -15.452  31.159  11.488  1.00  0.00           C
ATOM   3308  O   THR   423     -15.909  32.213  11.331  1.00  0.00           O
ATOM   3309  N   ASP   424     -15.713  30.220  10.651  1.00  0.00           N
ATOM   3310  CA  ASP   424     -16.035  30.467  10.083 33.67  0.00           C
ATOM   3311  CB  ASP   424     -16.557  29.212   8.659  1.00  0.00           C
ATOM   3312  CG  ASP   424     -15.218  29.178   7.990  1.00  0.00           C
ATOM   3313  OD1 ASP   424     -14.510  30.197   8.034  1.00  0.00           O
ATOM   3314  OD2 ASP   424     -14.885  28.132   7.426  1.00  0.00           O
ATOM   3315  C   ASP   424     -18.020  30.752   9.947  1.00  0.00           C
ATOM   3316  O   ASP   424     -18.655  31.611   9.370  1.00  0.00           O
ATOM   3317  N   VAL   425     -18.509  30.082  10.963  1.00  0.00           N
ATOM   3318  CA  VAL   425     -19.067  29.987  12.193 33.67  0.00           C
ATOM   3319  CB  VAL   425     -20.303  29.293  12.403  1.00  0.00           C
ATOM   3320  CG1 VAL   425     -21.652  29.567  12.961  1.00  0.00           C
ATOM   3321  CG2 VAL   425     -20.403  27.975  11.712  1.00  0.00           C
ATOM   3322  C   VAL   425     -19.865  31.723  12.178  1.00  0.00           C
ATOM   3323  O   VAL   425     -20.719  32.557  11.892  1.00  0.00           O
ATOM   3324  N   MET   426     -18.931  31.940  13.086  1.00  0.00           N
ATOM   3325  CA  MET   426     -18.297  32.881  14.468 33.67  0.00           C
ATOM   3326  CB  MET   426     -17.737  33.189  14.835  1.00  0.00           C
ATOM   3327  CG  MET   426     -17.657  34.458  15.643  1.00  0.00           C
ATOM   3328  SD  MET   426     -16.506  35.011  15.726  1.00  0.00           S
ATOM   3329  CE  MET   426     -15.991  33.840  16.269  1.00  0.00           C
ATOM   3330  C   MET   426     -18.685  34.355  12.833  1.00  0.00           C
ATOM   3331  O   MET   426     -19.488  35.280  12.846  1.00  0.00           O
ATOM   3332  N   CYS   427     -17.667  34.301  11.993  1.00  0.00           N
ATOM   3333  CA  CYS   427     -17.294  35.211  11.754 33.67  0.00           C
ATOM   3334  CB  CYS   427     -16.125  35.151  10.299  1.00  0.00           C
ATOM   3335  SG  CYS   427     -14.979  35.302  11.115  1.00  0.00           S
ATOM   3336  C   CYS   427     -18.595  35.599  10.090  1.00  0.00           C
ATOM   3337  O   CYS   427     -18.903  36.737   9.743  1.00  0.00           O
ATOM   3338  N   HIS   428     -19.259  34.530   9.688  1.00  0.00           N
ATOM   3339  CA  HIS   428     -20.253  34.157   9.654 33.67  0.00           C
ATOM   3340  ND1 HIS   428     -21.050  32.447   6.830  1.00  0.00           N
ATOM   3341  CG  HIS   428     -21.030  32.897   7.556  1.00  0.00           C
ATOM   3342  CB  HIS   428     -20.884  33.227   8.411  1.00  0.00           C
ATOM   3343  NE2 HIS   428     -21.261  32.536   6.447  1.00  0.00           N
ATOM   3344  CD2 HIS   428     -21.164  32.950   7.304  1.00  0.00           C
ATOM   3345  CE1 HIS   428     -21.188  32.240   6.182  1.00  0.00           C
ATOM   3346  C   HIS   428     -21.581  35.366   9.449  1.00  0.00           C
ATOM   3347  O   HIS   428     -22.150  36.254   8.863  1.00  0.00           O
ATOM   3348  N   PHE   429     -21.922  35.015  10.660  1.00  0.00           N
ATOM   3349  CA  PHE   429     -22.771  34.883  12.328 33.67  0.00           C
ATOM   3350  CB  PHE   429     -23.330  34.994  12.680  1.00  0.00           C
ATOM   3351  CG  PHE   429     -24.147  33.768  12.534  1.00  0.00           C
ATOM   3352  CD1 PHE   429     -25.192  33.674  11.627  1.00  0.00           C
ATOM   3353  CD2 PHE   429     -23.889  32.705  13.339  1.00  0.00           C
ATOM   3354  CE1 PHE   429     -25.951  32.541  11.515  1.00  0.00           C
ATOM   3355  CE2 PHE   429     -24.641  31.574  13.228  1.00  0.00           C
ATOM   3356  CZ  PHE   429     -25.674  31.492  12.315  1.00  0.00           C
ATOM   3357  C   PHE   429     -22.673  37.132  11.647  1.00  0.00           C
ATOM   3358  O   PHE   429     -23.579  37.975  11.629  1.00  0.00           O
ATOM   3359  N   PHE   430     -21.407  37.436  11.917  1.00  0.00           N
ATOM   3360  CA  PHE   430     -21.144  38.187  13.017 33.67  0.00           C
ATOM   3361  CB  PHE   430     -19.622  38.945  12.630  1.00  0.00           C
ATOM   3362  CG  PHE   430     -19.576  39.256  13.135  1.00  0.00           C
ATOM   3363  CD1 PHE   430     -20.417  39.588  13.897  1.00  0.00           C
ATOM   3364  CD2 PHE   430     -18.676  39.225  12.856  1.00  0.00           C
ATOM   3365  CE1 PHE   430     -20.364  39.882  14.376  1.00  0.00           C
ATOM   3366  CE2 PHE   430     -18.603  39.516  13.324  1.00  0.00           C
ATOM   3367  CZ  PHE   430     -19.453  39.847  14.087  1.00  0.00           C
ATOM   3368  C   PHE   430     -21.242  39.574  10.887  1.00  0.00           C
ATOM   3369  O   PHE   430     -21.909  40.566  10.914  1.00  0.00           O
ATOM   3370  N   ASN   431     -20.670  39.096   9.801  1.00  0.00           N
ATOM   3371  CA  ASN   431     -21.177  39.100   9.400 33.67  0.00           C
ATOM   3372  CB  ASN   431     -20.197  38.870   7.447  1.00  0.00           C
ATOM   3373  CG  ASN   431     -18.686  38.939   7.539  1.00  0.00           C
ATOM   3374  OD1 ASN   431     -18.148  39.816   8.165  1.00  0.00           O
ATOM   3375  ND2 ASN   431     -17.918  38.017   6.919  1.00  0.00           N
ATOM   3376  C   ASN   431     -22.263  39.998   8.160  1.00  0.00           C
ATOM   3377  O   ASN   431     -22.576  41.034   7.653  1.00  0.00           O
ATOM   3378  N   ASN   432     -23.137  39.065   8.429  1.00  0.00           N
ATOM   3379  CA  ASN   432     -24.786  38.177   9.199 33.67  0.00           C
ATOM   3380  CB  ASN   432     -25.215  37.866   7.944  1.00  0.00           C
ATOM   3381  CG  ASN   432     -24.684  37.283   6.683  1.00  0.00           C
ATOM   3382  OD1 ASN   432     -24.271  37.153   6.440  1.00  0.00           O
ATOM   3383  ND2 ASN   432     -24.668  36.899   5.812  1.00  0.00           N
ATOM   3384  C   ASN   432     -25.308  39.996   9.172  1.00  0.00           C
ATOM   3385  O   ASN   432     -26.520  40.146   9.087  1.00  0.00           O
ATOM   3386  N   GLY   433     -24.595  40.497  10.150  1.00  0.00           N
ATOM   3387  CA  GLY   433     -25.528  40.124  12.335 33.67  0.00           C
ATOM   3388  C   GLY   433     -25.878  40.599  12.282  1.00  0.00           C
ATOM   3389  O   GLY   433     -26.231  41.200  13.237  1.00  0.00           O
ATOM   3390  N   MET   434     -26.050  39.311  12.177  1.00  0.00           N
ATOM   3391  CA  MET   434     -26.561  37.182  14.345 33.67  0.00           C
ATOM   3392  CB  MET   434     -27.125  37.249  12.576  1.00  0.00           C
ATOM   3393  CG  MET   434     -28.146  37.462  11.507  1.00  0.00           C
ATOM   3394  SD  MET   434     -29.687  38.228  12.053  1.00  0.00           S
ATOM   3395  CE  MET   434     -30.575  37.412  11.485  1.00  0.00           C
ATOM   3396  C   MET   434     -25.784  38.236  14.376  1.00  0.00           C
ATOM   3397  O   MET   434     -25.258  37.192  14.622  1.00  0.00           O
ATOM   3398  N   ARG   435     -25.642  39.159  15.128  1.00  0.00           N
ATOM   3399  CA  ARG   435     -24.671  37.973  17.382 33.67  0.00           C
ATOM   3400  CB  ARG   435     -24.560  40.359  16.820  1.00  0.00           C
ATOM   3401  CG  ARG   435     -25.763  40.876  17.448  1.00  0.00           C
ATOM   3402  CD  ARG   435     -25.621  42.187  17.953  1.00  0.00           C
ATOM   3403  NE  ARG   435     -26.857  42.544  18.533  1.00  0.00           N
ATOM   3404  CZ  ARG   435     -27.630  43.229  18.565  1.00  0.00           C
ATOM   3405  NH1 ARG   435     -27.046  43.454  18.005  1.00  0.00           N
ATOM   3406  NH2 ARG   435     -28.686  43.421  19.141  1.00  0.00           N
ATOM   3407  C   ARG   435     -25.160  38.136  17.253  1.00  0.00           C
ATOM   3408  O   ARG   435     -24.329  37.513  17.910  1.00  0.00           O
ATOM   3409  N   SER   436     -26.434  38.037  17.365  1.00  0.00           N
ATOM   3410  CA  SER   436     -26.399  35.875  19.659 33.67  0.00           C
ATOM   3411  CB  SER   436     -28.458  37.333  18.397  1.00  0.00           C
ATOM   3412  OG  SER   436     -28.859  38.586  18.770  1.00  0.00           O
ATOM   3413  C   SER   436     -26.519  35.834  18.161  1.00  0.00           C
ATOM   3414  O   SER   436     -26.042  35.133  19.027  1.00  0.00           O
ATOM   3415  N   GLN   437     -26.664  35.465  16.994  1.00  0.00           N
ATOM   3416  CA  GLN   437     -25.496  32.951  17.712 33.67  0.00           C
ATOM   3417  CB  GLN   437     -26.646  33.993  15.181  1.00  0.00           C
ATOM   3418  CG  GLN   437     -28.128  33.871  14.984  1.00  0.00           C
ATOM   3419  CD  GLN   437     -28.822  34.239  14.513  1.00  0.00           C
ATOM   3420  OE1 GLN   437     -28.558  34.219  14.353  1.00  0.00           O
ATOM   3421  NE2 GLN   437     -29.748  34.597  14.265  1.00  0.00           N
ATOM   3422  C   GLN   437     -24.725  33.963  16.759  1.00  0.00           C
ATOM   3423  O   GLN   437     -24.254  32.992  17.294  1.00  0.00           O
ATOM   3424  N   VAL   438     -23.999  34.869  16.320  1.00  0.00           N
ATOM   3425  CA  VAL   438     -22.006  34.031  17.486 33.67  0.00           C
ATOM   3426  CB  VAL   438     -22.022  36.080  16.083  1.00  0.00           C
ATOM   3427  CG1 VAL   438     -20.583  36.211  16.476  1.00  0.00           C
ATOM   3428  CG2 VAL   438     -22.096  36.233  14.637  1.00  0.00           C
ATOM   3429  C   VAL   438     -22.185  34.610  17.952  1.00  0.00           C
ATOM   3430  O   VAL   438     -21.238  33.876  18.225  1.00  0.00           O
ATOM   3431  N   VAL   439     -22.932  35.207  18.813  1.00  0.00           N
ATOM   3432  CA  VAL   439     -21.872  34.242  21.135 33.67  0.00           C
ATOM   3433  CB  VAL   439     -23.438  36.064  20.953  1.00  0.00           C
ATOM   3434  CG1 VAL   439     -23.330  35.842  22.400  1.00  0.00           C
ATOM   3435  CG2 VAL   439     -22.987  37.438  20.738  1.00  0.00           C
ATOM   3436  C   VAL   439     -22.976  33.706  20.643  1.00  0.00           C
ATOM   3437  O   VAL   439     -22.306  33.145  21.489  1.00  0.00           O
ATOM   3438  N   ARG   440     -23.993  33.181  20.090  1.00  0.00           N
ATOM   3439  CA  ARG   440     -23.113  30.860  21.430 33.67  0.00           C
ATOM   3440  CB  ARG   440     -25.557  31.523  19.739  1.00  0.00           C
ATOM   3441  CG  ARG   440     -26.498  30.983  20.026  1.00  0.00           C
ATOM   3442  CD  ARG   440     -27.325  30.536  18.842  1.00  0.00           C
ATOM   3443  NE  ARG   440     -28.525  31.208  18.836  1.00  0.00           N
ATOM   3444  CZ  ARG   440     -29.361  31.097  17.832  1.00  0.00           C
ATOM   3445  NH1 ARG   440     -28.961  30.337  16.820  1.00  0.00           N
ATOM   3446  NH2 ARG   440     -30.387  31.775  17.998  1.00  0.00           N
ATOM   3447  C   ARG   440     -23.248  30.898  20.046  1.00  0.00           C
ATOM   3448  O   ARG   440     -22.893  29.984  20.775  1.00  0.00           O
ATOM   3449  N   ILE   441     -22.746  31.115  18.890  1.00  0.00           N
ATOM   3450  CA  ILE   441     -20.485  29.655  19.228 33.67  0.00           C
ATOM   3451  CB  ILE   441     -21.418  30.641  17.037  1.00  0.00           C
ATOM   3452  CG2 ILE   441     -20.237  29.898  16.559  1.00  0.00           C
ATOM   3453  CG1 ILE   441     -22.534  30.360  16.118  1.00  0.00           C
ATOM   3454  CD1 ILE   441     -22.646  30.168  15.794  1.00  0.00           C
ATOM   3455  C   ILE   441     -20.459  30.446  19.260  1.00  0.00           C
ATOM   3456  O   ILE   441     -19.748  29.462  19.518  1.00  0.00           O
ATOM   3457  N   LEU   442     -20.234  31.653  19.690  1.00  0.00           N
ATOM   3458  CA  LEU   442     -18.082  31.532  21.239 33.67  0.00           C
ATOM   3459  CB  LEU   442     -19.046  33.310  20.818  1.00  0.00           C
ATOM   3460  CG  LEU   442     -17.921  33.561  21.742  1.00  0.00           C
ATOM   3461  CD1 LEU   442     -16.645  33.620  21.071  1.00  0.00           C
ATOM   3462  CD2 LEU   442     -18.034  34.839  22.548  1.00  0.00           C
ATOM   3463  C   LEU   442     -19.282  31.122  21.830  1.00  0.00           C
ATOM   3464  O   LEU   442     -18.326  30.492  22.318  1.00  0.00           O
ATOM   3465  N   ARG   443     -20.465  31.181  22.351  1.00  0.00           N
ATOM   3466  CA  ARG   443     -19.351  29.988  24.310 33.67  0.00           C
ATOM   3467  CB  ARG   443     -22.164  30.741  23.969  1.00  0.00           C
ATOM   3468  CG  ARG   443     -22.404  30.996  24.555  1.00  0.00           C
ATOM   3469  CD  ARG   443     -23.831  31.268  24.978  1.00  0.00           C
ATOM   3470  NE  ARG   443     -23.917  31.491  25.509  1.00  0.00           N
ATOM   3471  CZ  ARG   443     -25.105  31.747  25.913  1.00  0.00           C
ATOM   3472  NH1 ARG   443     -26.034  31.740  25.723  1.00  0.00           N
ATOM   3473  NH2 ARG   443     -24.902  31.909  26.348  1.00  0.00           N
ATOM   3474  C   ARG   443     -20.496  29.038  23.416  1.00  0.00           C
ATOM   3475  O   ARG   443     -19.972  28.387  24.294  1.00  0.00           O
ATOM   3476  N   ALA   444     -20.452  29.726  25.251  1.00  0.00           N
ATOM   3477  CA  ALA   444     -17.953  26.956  25.922 50.00  0.00           C
ATOM   3478  CB  ALA   444     -21.820  27.861  26.079  1.00  0.00           C
ATOM   3479  C   ALA   444     -20.221  27.420  24.275  1.00  0.00           C
ATOM   3480  O   ALA   444     -19.411  26.490  24.315  1.00  0.00           O
ATOM   3481  N   TYR   445     -20.623  27.306  21.161  1.00  0.00           N
ATOM   3482  CA  TYR   445     -18.378  26.413  20.895 33.67  0.00           C
ATOM   3483  CB  TYR   445     -20.066  27.633  19.061  1.00  0.00           C
ATOM   3484  CG  TYR   445     -21.181  27.361  18.525  1.00  0.00           C
ATOM   3485  CD1 TYR   445     -22.245  28.259  18.478  1.00  0.00           C
ATOM   3486  CD2 TYR   445     -21.196  26.213  18.054  1.00  0.00           C
ATOM   3487  CE1 TYR   445     -23.316  28.037  17.976  1.00  0.00           C
ATOM   3488  CE2 TYR   445     -22.278  25.972  17.540  1.00  0.00           C
ATOM   3489  CZ  TYR   445     -23.331  26.899  17.510  1.00  0.00           C
ATOM   3490  OH  TYR   445     -24.393  26.707  17.028  1.00  0.00           O
ATOM   3491  C   TYR   445     -18.567  27.022  20.342  1.00  0.00           C
ATOM   3492  O   TYR   445     -17.790  26.200  19.956  1.00  0.00           O
ATOM   3493  N   VAL   446     -18.295  28.033  20.913  1.00  0.00           N
ATOM   3494  CA  VAL   446     -15.614  28.155  21.554 33.67  0.00           C
ATOM   3495  CB  VAL   446     -16.184  29.211  21.603  1.00  0.00           C
ATOM   3496  CG1 VAL   446     -15.501  29.653  21.117  1.00  0.00           C
ATOM   3497  CG2 VAL   446     -16.755  29.805  21.796  1.00  0.00           C
ATOM   3498  C   VAL   446     -16.484  27.280  22.051  1.00  0.00           C
ATOM   3499  O   VAL   446     -15.561  26.778  22.101  1.00  0.00           O
ATOM   3500  N   VAL   447     -17.040  27.021  22.841  1.00  0.00           N
ATOM   3501  CA  VAL   447     -14.683  26.097  24.228 33.67  0.00           C
ATOM   3502  CB  VAL   447     -16.852  25.325  24.806  1.00  0.00           C
ATOM   3503  CG1 VAL   447     -15.845  24.322  25.323  1.00  0.00           C
ATOM   3504  CG2 VAL   447     -17.220  26.144  25.759  1.00  0.00           C
ATOM   3505  C   VAL   447     -16.140  24.792  23.302  1.00  0.00           C
ATOM   3506  O   VAL   447     -15.189  24.127  23.564  1.00  0.00           O
ATOM   3507  N   ARG   448     -16.875  24.421  22.618  1.00  0.00           N
ATOM   3508  CA  ARG   448     -14.648  23.107  22.523 33.67  0.00           C
ATOM   3509  CB  ARG   448     -17.540  22.600  21.637  1.00  0.00           C
ATOM   3510  CG  ARG   448     -18.433  22.317  22.417  1.00  0.00           C
ATOM   3511  CD  ARG   448     -19.113  21.314  22.939  1.00  0.00           C
ATOM   3512  NE  ARG   448     -19.964  21.180  23.652  1.00  0.00           N
ATOM   3513  CZ  ARG   448     -20.696  20.352  24.239  1.00  0.00           C
ATOM   3514  NH1 ARG   448     -20.465  19.778  24.022  1.00  0.00           N
ATOM   3515  NH2 ARG   448     -21.374  20.420  24.813  1.00  0.00           N
ATOM   3516  C   ARG   448     -15.811  23.144  21.229  1.00  0.00           C
ATOM   3517  O   ARG   448     -15.492  22.172  21.013  1.00  0.00           O
ATOM   3518  N   CYS   449     -15.487  24.223  20.664  1.00  0.00           N
ATOM   3519  CA  CYS   449     -13.017  24.460  19.909 33.67  0.00           C
ATOM   3520  CB  CYS   449     -14.832  24.911  18.376  1.00  0.00           C
ATOM   3521  SG  CYS   449     -14.849  25.040  18.196  1.00  0.00           S
ATOM   3522  C   CYS   449     -13.819  25.005  19.928  1.00  0.00           C
ATOM   3523  O   CYS   449     -13.623  25.530  19.065  1.00  0.00           O
ATOM   3524  N   TYR   450     -13.250  25.037  21.095  1.00  0.00           N
ATOM   3525  CA  TYR   450     -11.002  26.344  21.556 33.67  0.00           C
ATOM   3526  CB  TYR   450     -11.792  25.416  21.563  1.00  0.00           C
ATOM   3527  CG  TYR   450     -11.111  24.499  21.645  1.00  0.00           C
ATOM   3528  CD1 TYR   450     -10.815  24.727  22.211  1.00  0.00           C
ATOM   3529  CD2 TYR   450     -10.752  23.393  21.166  1.00  0.00           C
ATOM   3530  CE1 TYR   450     -10.173  23.882  22.308  1.00  0.00           C
ATOM   3531  CE2 TYR   450     -10.099  22.524  21.256  1.00  0.00           C
ATOM   3532  CZ  TYR   450      -9.819  22.787  21.832  1.00  0.00           C
ATOM   3533  OH  TYR   450      -9.190  21.978  21.939  1.00  0.00           O
ATOM   3534  C   TYR   450     -11.633  26.840  20.994  1.00  0.00           C
ATOM   3535  O   TYR   450     -11.153  27.572  20.680  1.00  0.00           O
ATOM   3536  N   ARG   451     -11.470  27.017  20.985  1.00  0.00           N
ATOM   3537  CA  ARG   451      -9.158  28.712  20.792 33.67  0.00           C
ATOM   3538  CB  ARG   451     -10.299  28.328  20.837  1.00  0.00           C
ATOM   3539  CG  ARG   451      -9.534  27.869  22.026  1.00  0.00           C
ATOM   3540  CD  ARG   451      -9.153  28.044  22.228  1.00  0.00           C
ATOM   3541  NE  ARG   451      -8.425  27.543  23.401  1.00  0.00           N
ATOM   3542  CZ  ARG   451      -8.055  27.475  23.800  1.00  0.00           C
ATOM   3543  NH1 ARG   451      -8.471  27.922  22.960  1.00  0.00           N
ATOM   3544  NH2 ARG   451      -7.411  26.986  24.884  1.00  0.00           N
ATOM   3545  C   ARG   451     -10.180  28.670  20.068  1.00  0.00           C
ATOM   3546  O   ARG   451      -9.471  28.834  20.267  1.00  0.00           O
ATOM   3547  N   ASN   452     -10.504  28.991  19.335  1.00  0.00           N
ATOM   3548  CA  ASN   452      -9.047  30.432  18.693 33.67  0.00           C
ATOM   3549  CB  ASN   452     -10.267  29.152  17.633  1.00  0.00           C
ATOM   3550  CG  ASN   452     -10.141  29.361  18.125  1.00  0.00           C
ATOM   3551  OD1 ASN   452      -9.967  29.660  18.699  1.00  0.00           O
ATOM   3552  ND2 ASN   452     -10.222  29.216  17.927  1.00  0.00           N
ATOM   3553  C   ASN   452     -10.073  31.023  19.101  1.00  0.00           C
ATOM   3554  O   ASN   452     -10.487  31.923  18.418  1.00  0.00           O
ATOM   3555  N   PRO   453      -9.674  31.269  20.214  1.00  0.00           N
ATOM   3556  CA  PRO   453      -9.073  33.285  20.924 33.67  0.00           C
ATOM   3557  CD  PRO   453      -8.086  31.601  22.691  1.00  0.00           C
ATOM   3558  CB  PRO   453      -9.396  33.567  21.729  1.00  0.00           C
ATOM   3559  CG  PRO   453      -8.234  33.112  22.548  1.00  0.00           C
ATOM   3560  C   PRO   453      -8.632  33.419  20.132  1.00  0.00           C
ATOM   3561  O   PRO   453      -8.150  33.844  20.280  1.00  0.00           O
ATOM   3562  N   THR   454      -8.205  33.662  19.469  1.00  0.00           N
ATOM   3563  CA  THR   454      -6.791  35.342  19.070 33.67  0.00           C
ATOM   3564  CB  THR   454      -6.441  34.375  17.715  1.00  0.00           C
ATOM   3565  OG1 THR   454      -6.337  35.157  17.379  1.00  0.00           O
ATOM   3566  CG2 THR   454      -6.636  33.806  16.645  1.00  0.00           C
ATOM   3567  C   THR   454      -7.552  35.876  18.829  1.00  0.00           C
ATOM   3568  O   THR   454      -7.813  36.325  18.445  1.00  0.00           O
ATOM   3569  N   GLU   455      -7.528  36.637  19.131  1.00  0.00           N
ATOM   3570  CA  GLU   455      -7.916  38.797  19.452 33.67  0.00           C
ATOM   3571  CB  GLU   455      -6.736  38.943  19.193  1.00  0.00           C
ATOM   3572  CG  GLU   455      -6.609  39.561  19.862  1.00  0.00           C
ATOM   3573  CD  GLU   455      -5.570  39.454  19.839  1.00  0.00           C
ATOM   3574  OE1 GLU   455      -4.956  38.939  19.276  1.00  0.00           O
ATOM   3575  OE2 GLU   455      -5.379  39.872  20.404  1.00  0.00           O
ATOM   3576  C   GLU   455      -8.857  38.424  17.967  1.00  0.00           C
ATOM   3577  O   GLU   455      -9.798  39.197  18.156  1.00  0.00           O
ATOM   3578  N   SER   456      -8.579  37.905  16.773  1.00  0.00           N
ATOM   3579  CA  SER   456      -9.533  38.902  16.016 33.67  0.00           C
ATOM   3580  CB  SER   456      -8.730  37.518  14.417  1.00  0.00           C
ATOM   3581  OG  SER   456      -7.437  38.036  14.170  1.00  0.00           O
ATOM   3582  C   SER   456     -10.830  37.828  15.772  1.00  0.00           C
ATOM   3583  O   SER   456     -11.718  38.550  15.340  1.00  0.00           O
ATOM   3584  N   ILE   457     -11.083  36.706  16.428  1.00  0.00           N
ATOM   3585  CA  ILE   457     -12.268  36.622  17.133 33.67  0.00           C
ATOM   3586  CB  ILE   457     -12.507  34.845  17.163  1.00  0.00           C
ATOM   3587  CG2 ILE   457     -13.918  34.378  17.523  1.00  0.00           C
ATOM   3588  CG1 ILE   457     -11.992  33.801  16.164  1.00  0.00           C
ATOM   3589  CD1 ILE   457     -12.417  33.098  15.889  1.00  0.00           C
ATOM   3590  C   ILE   457     -13.164  37.209  17.599  1.00  0.00           C
ATOM   3591  O   ILE   457     -14.330  37.585  17.395  1.00  0.00           O
ATOM   3592  N   PRO   458     -12.457  37.613  18.641  1.00  0.00           N
ATOM   3593  CA  PRO   458     -12.906  38.761  20.177 33.67  0.00           C
ATOM   3594  CD  PRO   458     -10.780  36.757  19.831  1.00  0.00           C
ATOM   3595  CB  PRO   458     -12.091  38.698  20.827  1.00  0.00           C
ATOM   3596  CG  PRO   458     -11.260  37.453  21.095  1.00  0.00           C
ATOM   3597  C   PRO   458     -13.247  39.928  18.962  1.00  0.00           C
ATOM   3598  O   PRO   458     -14.284  40.530  19.201  1.00  0.00           O
ATOM   3599  N   HIS   459     -12.319  40.384  18.135  1.00  0.00           N
ATOM   3600  CA  HIS   459     -12.815  41.843  18.066 33.67  0.00           C
ATOM   3601  ND1 HIS   459     -10.878  44.144  15.675  1.00  0.00           N
ATOM   3602  CG  HIS   459     -11.225  43.241  15.939  1.00  0.00           C
ATOM   3603  CB  HIS   459     -11.201  41.927  16.616  1.00  0.00           C
ATOM   3604  NE2 HIS   459     -11.420  45.031  14.946  1.00  0.00           N
ATOM   3605  CD2 HIS   459     -11.570  43.815  15.477  1.00  0.00           C
ATOM   3606  CE1 HIS   459     -10.999  45.199  15.077  1.00  0.00           C
ATOM   3607  C   HIS   459     -13.657  41.586  16.528  1.00  0.00           C
ATOM   3608  O   HIS   459     -14.316  42.595  16.348  1.00  0.00           O
ATOM   3609  N   THR   460     -13.918  40.427  15.943  1.00  0.00           N
ATOM   3610  CA  THR   460     -15.332  40.212  15.795 33.67  0.00           C
ATOM   3611  CB  THR   460     -14.993  38.881  14.349  1.00  0.00           C
ATOM   3612  OG1 THR   460     -13.895  38.873  13.440  1.00  0.00           O
ATOM   3613  CG2 THR   460     -16.279  38.644  13.579  1.00  0.00           C
ATOM   3614  C   THR   460     -16.351  40.358  15.832  1.00  0.00           C
ATOM   3615  O   THR   460     -17.218  41.102  15.434  1.00  0.00           O
ATOM   3616  N   VAL   461     -16.458  39.671  16.944  1.00  0.00           N
ATOM   3617  CA  VAL   461     -17.600  39.438  18.594 33.67  0.00           C
ATOM   3618  CB  VAL   461     -17.690  38.703  18.748  1.00  0.00           C
ATOM   3619  CG1 VAL   461     -17.777  38.752  20.019  1.00  0.00           C
ATOM   3620  CG2 VAL   461     -18.479  38.025  18.669  1.00  0.00           C
ATOM   3621  C   VAL   461     -17.800  41.178  18.328  1.00  0.00           C
ATOM   3622  O   VAL   461     -18.902  41.692  18.383  1.00  0.00           O
ATOM   3623  N   ALA   462     -16.713  41.814  18.730  1.00  0.00           N
ATOM   3624  CA  ALA   462     -17.139  42.845  20.101 33.67  0.00           C
ATOM   3625  CB  ALA   462     -15.397  43.686  19.674  1.00  0.00           C
ATOM   3626  C   ALA   462     -17.318  44.111  18.174  1.00  0.00           C
ATOM   3627  O   ALA   462     -18.210  44.875  18.446  1.00  0.00           O
ATOM   3628  N   LEU   463     -16.779  44.049  16.985  1.00  0.00           N
ATOM   3629  CA  LEU   463     -17.965  44.511  16.847 33.67  0.00           C
ATOM   3630  CB  LEU   463     -16.377  44.723  14.695  1.00  0.00           C
ATOM   3631  CG  LEU   463     -14.979  45.290  14.864  1.00  0.00           C
ATOM   3632  CD1 LEU   463     -14.033  44.991  13.737  1.00  0.00           C
ATOM   3633  CD2 LEU   463     -14.888  46.805  14.996  1.00  0.00           C
ATOM   3634  C   LEU   463     -18.676  44.590  15.543  1.00  0.00           C
ATOM   3635  O   LEU   463     -19.390  45.464  15.172  1.00  0.00           O
ATOM   3636  N   TRP   464     -19.085  43.330  15.691  1.00  0.00           N
ATOM   3637  CA  TRP   464     -20.920  42.233  16.529 33.67  0.00           C
ATOM   3638  CB  TRP   464     -20.799  41.468  15.649  1.00  0.00           C
ATOM   3639  CG  TRP   464     -22.243  41.121  15.429  1.00  0.00           C
ATOM   3640  CD2 TRP   464     -23.269  41.000  16.415  1.00  0.00           C
ATOM   3641  CD1 TRP   464     -22.887  40.855  14.278  1.00  0.00           C
ATOM   3642  NE1 TRP   464     -24.164  40.586  14.431  1.00  0.00           N
ATOM   3643  CE2 TRP   464     -24.462  40.661  15.752  1.00  0.00           C
ATOM   3644  CE3 TRP   464     -23.301  41.141  17.801  1.00  0.00           C
ATOM   3645  CZ2 TRP   464     -25.689  40.460  16.433  1.00  0.00           C
ATOM   3646  CZ3 TRP   464     -24.529  40.941  18.491  1.00  0.00           C
ATOM   3647  CH2 TRP   464     -25.702  40.606  17.796  1.00  0.00           C
ATOM   3648  C   TRP   464     -21.371  43.634  16.403  1.00  0.00           C
ATOM   3649  O   TRP   464     -22.332  44.223  16.003  1.00  0.00           O
ATOM   3650  N   ALA   465     -21.061  43.563  17.667  1.00  0.00           N
ATOM   3651  CA  ALA   465     -22.265  43.341  19.820 33.67  0.00           C
ATOM   3652  CB  ALA   465     -21.260  43.959  20.055  1.00  0.00           C
ATOM   3653  C   ALA   465     -21.878  45.685  18.461  1.00  0.00           C
ATOM   3654  O   ALA   465     -22.926  46.275  18.523  1.00  0.00           O
ATOM   3655  N   LEU   466     -20.733  46.289  18.202  1.00  0.00           N
ATOM   3656  CA  LEU   466     -21.457  46.914  19.034 33.67  0.00           C
ATOM   3657  CB  LEU   466     -19.279  48.219  17.635  1.00  0.00           C
ATOM   3658  CG  LEU   466     -19.032  49.005  18.325  1.00  0.00           C
ATOM   3659  CD1 LEU   466     -18.423  49.288  18.957  1.00  0.00           C
ATOM   3660  CD2 LEU   466     -18.604  50.060  18.507  1.00  0.00           C
ATOM   3661  C   LEU   466     -21.495  48.122  16.799  1.00  0.00           C
ATOM   3662  O   LEU   466     -22.169  49.121  16.875  1.00  0.00           O
ATOM   3663  N   ARG   467     -19.894  47.725  13.509  1.00  0.00           N
ATOM   3664  CA  ARG   467     -22.667  46.725  14.966 50.00  0.00           C
ATOM   3665  CB  ARG   467     -20.638  47.031  11.094  1.00  0.00           C
ATOM   3666  CG  ARG   467     -19.300  46.975  10.355  1.00  0.00           C
ATOM   3667  CD  ARG   467     -19.266  45.942   9.226  1.00  0.00           C
ATOM   3668  NE  ARG   467     -17.906  45.984   8.620  1.00  0.00           N
ATOM   3669  CZ  ARG   467     -17.558  45.079   7.660  1.00  0.00           C
ATOM   3670  NH1 ARG   467     -18.631  44.262   7.450  1.00  0.00           N
ATOM   3671  NH2 ARG   467     -16.272  45.343   7.285  1.00  0.00           N
ATOM   3672  C   ARG   467     -22.147  48.073  12.520  1.00  0.00           C
ATOM   3673  O   ARG   467     -22.824  48.872  11.866  1.00  0.00           O
ATOM   3674  N   LEU   468     -22.750  47.300  13.415  1.00  0.00           N
ATOM   3675  CA  LEU   468     -25.863  45.432  16.428 50.00  0.00           C
ATOM   3676  CB  LEU   468     -24.927  46.252  14.363  1.00  0.00           C
ATOM   3677  CG  LEU   468     -24.925  44.952  13.557  1.00  0.00           C
ATOM   3678  CD1 LEU   468     -25.572  43.757  14.256  1.00  0.00           C
ATOM   3679  CD2 LEU   468     -25.660  45.020  12.219  1.00  0.00           C
ATOM   3680  C   LEU   468     -24.560  48.530  14.635  1.00  0.00           C
ATOM   3681  O   LEU   468     -25.736  48.656  14.987  1.00  0.00           O
ATOM   3682  N   GLY   469     -23.644  49.379  15.093  1.00  0.00           N
ATOM   3683  CA  GLY   469     -25.841  48.198  19.087 50.00  0.00           C
ATOM   3684  C   GLY   469     -24.039  50.076  17.478  1.00  0.00           C
ATOM   3685  O   GLY   469     -24.376  50.904  18.328  1.00  0.00           O
ATOM   3686  N   PRO   470     -23.743  48.829  17.825  1.00  0.00           N
ATOM   3687  CA  PRO   470     -25.022  45.975  22.016 50.00  0.00           C
ATOM   3688  CD  PRO   470     -22.784  45.838  18.051  1.00  0.00           C
ATOM   3689  CB  PRO   470     -23.738  47.061  20.088  1.00  0.00           C
ATOM   3690  CG  PRO   470     -22.796  45.970  19.575  1.00  0.00           C
ATOM   3691  C   PRO   470     -22.489  48.456  19.935  1.00  0.00           C
ATOM   3692  O   PRO   470     -21.650  47.558  20.043  1.00  0.00           O
ATOM   3693  N   PHE   471     -22.308  49.682  20.488  1.00  0.00           N
ATOM   3694  CA  PHE   471     -22.400  47.934  23.951 50.00  0.00           C
ATOM   3695  CB  PHE   471     -20.793  51.511  21.111  1.00  0.00           C
ATOM   3696  CG  PHE   471     -20.710  52.089  19.730  1.00  0.00           C
ATOM   3697  CD1 PHE   471     -19.533  52.031  19.011  1.00  0.00           C
ATOM   3698  CD2 PHE   471     -21.807  52.705  19.159  1.00  0.00           C
ATOM   3699  CE1 PHE   471     -19.460  52.567  17.742  1.00  0.00           C
ATOM   3700  CE2 PHE   471     -21.741  53.247  17.893  1.00  0.00           C
ATOM   3701  CZ  PHE   471     -20.572  53.178  17.183  1.00  0.00           C
ATOM   3702  C   PHE   471     -20.910  49.498  22.569  1.00  0.00           C
ATOM   3703  O   PHE   471     -19.808  49.216  23.072  1.00  0.00           O
ATOM   3704  N   ASP   472     -22.057  49.386  23.219  1.00  0.00           N
ATOM   3705  CA  ASP   472     -22.927  46.214  27.215 50.00  0.00           C
ATOM   3706  CB  ASP   472     -23.593  48.822  25.008  1.00  0.00           C
ATOM   3707  CG  ASP   472     -23.966  50.261  25.335  1.00  0.00           C
ATOM   3708  OD1 ASP   472     -23.037  51.108  25.422  1.00  0.00           O
ATOM   3709  OD2 ASP   472     -25.185  50.532  25.503  1.00  0.00           O
ATOM   3710  C   ASP   472     -21.599  47.370  24.586  1.00  0.00           C
ATOM   3711  O   ASP   472     -20.880  46.989  25.513  1.00  0.00           O
ATOM   3712  N   TRP   473     -22.550  44.338  25.059  1.00  0.00           N
ATOM   3713  CA  TRP   473     -21.938  42.330  25.796 33.67  0.00           C
ATOM   3714  CB  TRP   473     -22.681  42.611  23.556  1.00  0.00           C
ATOM   3715  CG  TRP   473     -23.917  42.007  23.913  1.00  0.00           C
ATOM   3716  CD2 TRP   473     -24.024  40.896  24.626  1.00  0.00           C
ATOM   3717  CD1 TRP   473     -25.179  42.367  23.649  1.00  0.00           C
ATOM   3718  NE1 TRP   473     -26.037  41.624  24.104  1.00  0.00           N
ATOM   3719  CE2 TRP   473     -25.369  40.684  24.727  1.00  0.00           C
ATOM   3720  CE3 TRP   473     -23.112  40.053  25.192  1.00  0.00           C
ATOM   3721  CZ2 TRP   473     -25.834  39.653  25.378  1.00  0.00           C
ATOM   3722  CZ3 TRP   473     -23.571  39.016  25.848  1.00  0.00           C
ATOM   3723  CH2 TRP   473     -24.923  38.833  25.930  1.00  0.00           C
ATOM   3724  C   TRP   473     -20.529  43.544  24.283  1.00  0.00           C
ATOM   3725  O   TRP   473     -19.664  42.766  24.456  1.00  0.00           O
ATOM   3726  N   ALA   474     -20.338  44.651  23.785  1.00  0.00           N
ATOM   3727  CA  ALA   474     -19.340  44.401  24.482 33.67  0.00           C
ATOM   3728  CB  ALA   474     -19.056  46.385  22.939  1.00  0.00           C
ATOM   3729  C   ALA   474     -18.163  45.065  24.751  1.00  0.00           C
ATOM   3730  O   ALA   474     -17.060  44.547  24.806  1.00  0.00           O
ATOM   3731  N   LEU   475     -17.756  47.163  23.877  1.00  0.00           N
ATOM   3732  CA  LEU   475     -17.754  45.816  27.369 50.00  0.00           C
ATOM   3733  CB  LEU   475     -17.621  48.563  25.892  1.00  0.00           C
ATOM   3734  CG  LEU   475     -16.916  48.836  27.222  1.00  0.00           C
ATOM   3735  CD1 LEU   475     -15.437  49.202  27.106  1.00  0.00           C
ATOM   3736  CD2 LEU   475     -17.503  49.986  28.040  1.00  0.00           C
ATOM   3737  C   LEU   475     -17.067  46.121  25.976  1.00  0.00           C
ATOM   3738  O   LEU   475     -16.033  45.693  26.473  1.00  0.00           O
ATOM   3739  N   GLY   476     -18.607  43.481  27.383  1.00  0.00           N
ATOM   3740  CA  GLY   476     -18.238  41.641  28.717 33.67  0.00           C
ATOM   3741  C   GLY   476     -17.302  41.545  27.096  1.00  0.00           C
ATOM   3742  O   GLY   476     -16.443  40.941  27.665  1.00  0.00           O
ATOM   3743  N   PHE   477     -17.327  41.677  25.857  1.00  0.00           N
ATOM   3744  CA  PHE   477     -16.153  40.776  25.883 33.67  0.00           C
ATOM   3745  CB  PHE   477     -16.554  41.345  23.642  1.00  0.00           C
ATOM   3746  CG  PHE   477     -17.703  40.505  23.219  1.00  0.00           C
ATOM   3747  CD1 PHE   477     -18.170  39.482  23.972  1.00  0.00           C
ATOM   3748  CD2 PHE   477     -18.357  40.720  22.040  1.00  0.00           C
ATOM   3749  CE1 PHE   477     -19.273  38.685  23.563  1.00  0.00           C
ATOM   3750  CE2 PHE   477     -19.465  39.936  21.607  1.00  0.00           C
ATOM   3751  CZ  PHE   477     -19.922  38.913  22.372  1.00  0.00           C
ATOM   3752  C   PHE   477     -14.974  41.713  25.505  1.00  0.00           C
ATOM   3753  O   PHE   477     -14.064  41.046  25.681  1.00  0.00           O
ATOM   3754  N   LEU   478     -14.875  42.939  25.699  1.00  0.00           N
ATOM   3755  CA  LEU   478     -13.725  43.433  26.873 33.67  0.00           C
ATOM   3756  CB  LEU   478     -13.794  44.981  26.148  1.00  0.00           C
ATOM   3757  CG  LEU   478     -12.579  45.773  26.504  1.00  0.00           C
ATOM   3758  CD1 LEU   478     -11.488  45.478  25.604  1.00  0.00           C
ATOM   3759  CD2 LEU   478     -12.806  47.167  26.475  1.00  0.00           C
ATOM   3760  C   LEU   478     -13.081  43.101  27.396  1.00  0.00           C
ATOM   3761  O   LEU   478     -11.897  42.875  27.571  1.00  0.00           O
ATOM   3762  N   ALA   479     -13.928  42.924  28.256  1.00  0.00           N
ATOM   3763  CA  ALA   479     -13.394  42.236  30.271 33.67  0.00           C
ATOM   3764  CB  ALA   479     -14.651  42.451  30.323  1.00  0.00           C
ATOM   3765  C   ALA   479     -13.095  41.118  29.343  1.00  0.00           C
ATOM   3766  O   ALA   479     -12.183  40.758  30.036  1.00  0.00           O
ATOM   3767  N   ILE   480     -13.747  40.396  28.475  1.00  0.00           N
ATOM   3768  CA  ILE   480     -12.866  39.024  28.910 33.67  0.00           C
ATOM   3769  CB  ILE   480     -14.189  38.232  27.422  1.00  0.00           C
ATOM   3770  CG2 ILE   480     -13.554  37.059  27.171  1.00  0.00           C
ATOM   3771  CG1 ILE   480     -15.512  37.801  27.923  1.00  0.00           C
ATOM   3772  CD1 ILE   480     -16.444  37.081  27.024  1.00  0.00           C
ATOM   3773  C   ILE   480     -12.022  39.122  27.694  1.00  0.00           C
ATOM   3774  O   ILE   480     -11.248  38.319  28.011  1.00  0.00           O
ATOM   3775  N   LEU   481     -11.750  39.993  26.891  1.00  0.00           N
ATOM   3776  CA  LEU   481     -10.192  40.332  26.909 33.67  0.00           C
ATOM   3777  CB  LEU   481     -10.254  41.067  25.458  1.00  0.00           C
ATOM   3778  CG  LEU   481     -10.939  40.750  24.151  1.00  0.00           C
ATOM   3779  CD1 LEU   481     -10.996  41.741  23.291  1.00  0.00           C
ATOM   3780  CD2 LEU   481     -10.363  39.722  23.486  1.00  0.00           C
ATOM   3781  C   LEU   481      -9.411  40.390  27.465  1.00  0.00           C
ATOM   3782  O   LEU   481      -8.363  39.853  27.519  1.00  0.00           O
ATOM   3783  N   LEU   482      -9.676  41.255  28.292  1.00  0.00           N
ATOM   3784  CA  LEU   482      -8.625  41.826  29.933 33.67  0.00           C
ATOM   3785  CB  LEU   482      -9.343  42.427  30.424  1.00  0.00           C
ATOM   3786  CG  LEU   482      -8.421  42.628  31.625  1.00  0.00           C
ATOM   3787  CD1 LEU   482      -7.113  43.232  31.373  1.00  0.00           C
ATOM   3788  CD2 LEU   482      -8.935  43.473  32.674  1.00  0.00           C
ATOM   3789  C   LEU   482      -8.565  40.358  30.165  1.00  0.00           C
ATOM   3790  O   LEU   482      -7.450  40.087  30.514  1.00  0.00           O
ATOM   3791  N   ALA   483      -9.602  39.665  30.408  1.00  0.00           N
ATOM   3792  CA  ALA   483      -8.993  38.777  31.745 33.67  0.00           C
ATOM   3793  CB  ALA   483     -10.930  38.064  31.437  1.00  0.00           C
ATOM   3794  C   ALA   483      -8.841  37.466  30.563  1.00  0.00           C
ATOM   3795  O   ALA   483      -8.122  36.809  31.189  1.00  0.00           O
ATOM   3796  N   SER   484      -9.032  37.303  29.299  1.00  0.00           N
ATOM   3797  CA  SER   484      -7.814  36.635  29.005 33.67  0.00           C
ATOM   3798  CB  SER   484      -9.161  35.613  27.468  1.00  0.00           C
ATOM   3799  OG  SER   484     -10.299  35.043  27.793  1.00  0.00           O
ATOM   3800  C   SER   484      -7.227  36.545  28.054  1.00  0.00           C
ATOM   3801  O   SER   484      -6.518  35.689  27.691  1.00  0.00           O
ATOM   3802  N   ASN   485      -7.479  37.962  30.485  1.00  0.00           N
ATOM   3803  CA  ASN   485      -4.739  38.780  30.163 50.00  0.00           C
ATOM   3804  CB  ASN   485      -5.281  36.953  31.168  1.00  0.00           C
ATOM   3805  CG  ASN   485      -5.366  37.420  32.614  1.00  0.00           C
ATOM   3806  OD1 ASN   485      -5.522  38.609  32.888  1.00  0.00           O
ATOM   3807  ND2 ASN   485      -5.267  36.509  33.619  1.00  0.00           N
ATOM   3808  C   ASN   485      -5.644  37.674  28.858  1.00  0.00           C
ATOM   3809  O   ASN   485      -4.581  37.126  28.553  1.00  0.00           O
ATOM   3810  N   ASP   486      -5.851  39.021  27.090  1.00  0.00           N
ATOM   3811  CA  ASP   486      -5.199  40.152  27.002 33.67  0.00           C
ATOM   3812  CB  ASP   486      -5.300  38.855  26.598  1.00  0.00           C
ATOM   3813  CG  ASP   486      -5.595  37.500  27.193  1.00  0.00           C
ATOM   3814  OD1 ASP   486      -5.560  37.174  28.004  1.00  0.00           O
ATOM   3815  OD2 ASP   486      -5.859  36.772  26.842  1.00  0.00           O
ATOM   3816  C   ASP   486      -5.012  40.665  26.149  1.00  0.00           C
ATOM   3817  O   ASP   486      -5.338  41.736  26.637  1.00  0.00           O
ATOM   3818  N   GLU   487      -4.306  40.577  25.037  1.00  0.00           N
ATOM   3819  CA  GLU   487      -4.090  42.593  24.388 33.67  0.00           C
ATOM   3820  CB  GLU   487      -2.632  41.647  23.642  1.00  0.00           C
ATOM   3821  CG  GLU   487      -1.569  41.549  24.600  1.00  0.00           C
ATOM   3822  CD  GLU   487      -0.353  41.497  23.857  1.00  0.00           C
ATOM   3823  OE1 GLU   487      -0.408  41.512  22.661  1.00  0.00           O
ATOM   3824  OE2 GLU   487       0.648  41.442  24.475  1.00  0.00           O
ATOM   3825  C   GLU   487      -4.777  42.000  23.150  1.00  0.00           C
ATOM   3826  O   GLU   487      -5.246  41.154  22.661  1.00  0.00           O
ATOM   3827  N   PRO   488      -4.991  43.176  22.737  1.00  0.00           N
ATOM   3828  CA  PRO   488      -6.424  44.211  22.146 33.67  0.00           C
ATOM   3829  CD  PRO   488      -6.029  45.369  23.736  1.00  0.00           C
ATOM   3830  CB  PRO   488      -6.998  44.343  21.608  1.00  0.00           C
ATOM   3831  CG  PRO   488      -6.817  45.557  22.496  1.00  0.00           C
ATOM   3832  C   PRO   488      -5.077  44.208  20.635  1.00  0.00           C
ATOM   3833  O   PRO   488      -4.172  44.878  20.908  1.00  0.00           O
ATOM   3834  N   LEU   489      -5.479  44.142  19.435  1.00  0.00           N
ATOM   3835  CA  LEU   489      -5.665  45.817  18.927 33.67  0.00           C
ATOM   3836  CB  LEU   489      -4.901  44.477  16.958  1.00  0.00           C
ATOM   3837  CG  LEU   489      -4.434  43.090  16.626  1.00  0.00           C
ATOM   3838  CD1 LEU   489      -4.611  42.672  15.199  1.00  0.00           C
ATOM   3839  CD2 LEU   489      -2.968  42.820  16.850  1.00  0.00           C
ATOM   3840  C   LEU   489      -5.270  46.251  18.391  1.00  0.00           C
ATOM   3841  O   LEU   489      -6.341  46.640  18.794  1.00  0.00           O
ATOM   3842  N   SER   490      -4.521  47.035  17.947  1.00  0.00           N
ATOM   3843  CA  SER   490      -6.148  49.372  18.377 33.67  0.00           C
ATOM   3844  CB  SER   490      -3.945  49.083  16.890  1.00  0.00           C
ATOM   3845  OG  SER   490      -2.767  49.111  17.415  1.00  0.00           O
ATOM   3846  C   SER   490      -6.096  48.644  17.066  1.00  0.00           C
ATOM   3847  O   SER   490      -6.818  49.554  17.402  1.00  0.00           O
ATOM   3848  N   ASP   491      -6.378  47.802  16.121  1.00  0.00           N
ATOM   3849  CA  ASP   491      -8.934  48.525  16.149 33.67  0.00           C
ATOM   3850  CB  ASP   491      -7.704  46.757  14.432  1.00  0.00           C
ATOM   3851  CG  ASP   491      -6.812  47.110  13.349  1.00  0.00           C
ATOM   3852  OD1 ASP   491      -6.323  48.264  13.324  1.00  0.00           O
ATOM   3853  OD2 ASP   491      -6.609  46.229  12.533  1.00  0.00           O
ATOM   3854  C   ASP   491      -8.732  47.696  16.278  1.00  0.00           C
ATOM   3855  O   ASP   491      -9.697  48.281  16.059  1.00  0.00           O
ATOM   3856  N   ARG   492      -8.630  46.896  17.260  1.00  0.00           N
ATOM   3857  CA  ARG   492     -10.689  47.020  18.980 33.67  0.00           C
ATOM   3858  CB  ARG   492      -9.431  45.512  19.139  1.00  0.00           C
ATOM   3859  CG  ARG   492     -10.557  45.283  20.061  1.00  0.00           C
ATOM   3860  CD  ARG   492     -11.258  44.502  20.410  1.00  0.00           C
ATOM   3861  NE  ARG   492     -12.330  44.407  21.307  1.00  0.00           N
ATOM   3862  CZ  ARG   492     -12.732  44.498  22.000  1.00  0.00           C
ATOM   3863  NH1 ARG   492     -11.995  44.670  21.688  1.00  0.00           N
ATOM   3864  NH2 ARG   492     -13.713  44.380  22.734  1.00  0.00           N
ATOM   3865  C   ARG   492      -9.947  47.859  18.909  1.00  0.00           C
ATOM   3866  O   ARG   492     -11.009  48.202  19.093  1.00  0.00           O
ATOM   3867  N   ILE   493      -8.981  48.529  19.367  1.00  0.00           N
ATOM   3868  CA  ILE   493     -10.495  50.077  21.085 33.67  0.00           C
ATOM   3869  CB  ILE   493      -7.954  50.522  20.618  1.00  0.00           C
ATOM   3870  CG2 ILE   493      -8.253  51.837  21.304  1.00  0.00           C
ATOM   3871  CG1 ILE   493      -7.220  49.800  21.591  1.00  0.00           C
ATOM   3872  CD1 ILE   493      -5.954  50.436  21.941  1.00  0.00           C
ATOM   3873  C   ILE   493      -9.682  50.762  19.251  1.00  0.00           C
ATOM   3874  O   ILE   493     -10.541  51.433  19.603  1.00  0.00           O
ATOM   3875  N   GLN   494      -9.214  50.863  18.135  1.00  0.00           N
ATOM   3876  CA  GLN   494     -11.306  52.094  18.221 33.67  0.00           C
ATOM   3877  CB  GLN   494      -9.162  51.750  16.036  1.00  0.00           C
ATOM   3878  CG  GLN   494      -8.523  52.492  15.104  1.00  0.00           C
ATOM   3879  CD  GLN   494      -8.012  52.335  13.940  1.00  0.00           C
ATOM   3880  OE1 GLN   494      -8.156  51.623  13.839  1.00  0.00           O
ATOM   3881  NE2 GLN   494      -7.386  52.992  12.998  1.00  0.00           N
ATOM   3882  C   GLN   494     -11.094  51.542  17.024  1.00  0.00           C
ATOM   3883  O   GLN   494     -11.806  52.481  16.929  1.00  0.00           O
ATOM   3884  N   ARG   495     -11.496  50.278  16.950  1.00  0.00           N
ATOM   3885  CA  ARG   495     -14.166  49.897  17.597 33.67  0.00           C
ATOM   3886  CB  ARG   495     -13.067  48.395  16.558  1.00  0.00           C
ATOM   3887  CG  ARG   495     -13.039  47.883  15.129  1.00  0.00           C
ATOM   3888  CD  ARG   495     -13.966  46.939  14.757  1.00  0.00           C
ATOM   3889  NE  ARG   495     -14.363  47.358  13.888  1.00  0.00           N
ATOM   3890  CZ  ARG   495     -15.389  47.552  13.078  1.00  0.00           C
ATOM   3891  NH1 ARG   495     -16.025  47.317  13.147  1.00  0.00           N
ATOM   3892  NH2 ARG   495     -15.587  47.900  12.360  1.00  0.00           N
ATOM   3893  C   ARG   495     -13.583  50.224  17.742  1.00  0.00           C
ATOM   3894  O   ARG   495     -14.589  50.675  17.574  1.00  0.00           O
ATOM   3895  N   ARG   496     -13.130  49.966  18.847  1.00  0.00           N
ATOM   3896  CA  ARG   496     -15.179  50.228  20.963 33.67  0.00           C
ATOM   3897  CB  ARG   496     -13.214  49.908  21.101  1.00  0.00           C
ATOM   3898  CG  ARG   496     -13.467  49.252  22.073  1.00  0.00           C
ATOM   3899  CD  ARG   496     -13.049  48.788  23.254  1.00  0.00           C
ATOM   3900  NE  ARG   496     -12.568  49.101  24.289  1.00  0.00           N
ATOM   3901  CZ  ARG   496     -11.688  48.368  24.861  1.00  0.00           C
ATOM   3902  NH1 ARG   496     -11.419  47.426  24.306  1.00  0.00           N
ATOM   3903  NH2 ARG   496     -11.419  48.865  25.765  1.00  0.00           N
ATOM   3904  C   ARG   496     -13.925  51.844  19.950  1.00  0.00           C
ATOM   3905  O   ARG   496     -14.857  52.289  20.166  1.00  0.00           O
ATOM   3906  N   MET   497     -12.979  52.607  19.711  1.00  0.00           N
ATOM   3907  CA  MET   497     -14.663  53.934  20.757 33.67  0.00           C
ATOM   3908  CB  MET   497     -11.709  54.655  19.451  1.00  0.00           C
ATOM   3909  CG  MET   497     -10.942  55.260  19.573  1.00  0.00           C
ATOM   3910  SD  MET   497      -9.962  55.914  20.573  1.00  0.00           S
ATOM   3911  CE  MET   497      -9.595  56.416  19.608  1.00  0.00           C
ATOM   3912  C   MET   497     -13.864  54.491  18.769  1.00  0.00           C
ATOM   3913  O   MET   497     -14.527  55.376  19.042  1.00  0.00           O
ATOM   3914  N   HIS   498     -13.458  54.098  14.942  1.00  0.00           N
ATOM   3915  CA  HIS   498     -16.966  53.984  16.483 50.00  0.00           C
ATOM   3916  ND1 HIS   498     -13.176  54.608  10.376  1.00  0.00           N
ATOM   3917  CG  HIS   498     -13.632  53.576  11.171  1.00  0.00           C
ATOM   3918  CB  HIS   498     -13.211  53.366  12.595  1.00  0.00           C
ATOM   3919  NE2 HIS   498     -14.507  53.495   9.159  1.00  0.00           N
ATOM   3920  CD2 HIS   498     -14.480  52.873  10.384  1.00  0.00           C
ATOM   3921  CE1 HIS   498     -13.704  54.541   9.191  1.00  0.00           C
ATOM   3922  C   HIS   498     -15.342  54.308  13.390  1.00  0.00           C
ATOM   3923  O   HIS   498     -15.935  55.205  12.793  1.00  0.00           O
ATOM   3924  N   GLU   499     -15.964  53.252  13.905  1.00  0.00           N
ATOM   3925  CA  GLU   499     -20.188  52.078  16.838 50.00  0.00           C
ATOM   3926  CB  GLU   499     -17.926  51.803  14.299  1.00  0.00           C
ATOM   3927  CG  GLU   499     -19.442  51.634  14.181  1.00  0.00           C
ATOM   3928  CD  GLU   499     -19.799  51.616  12.701  1.00  0.00           C
ATOM   3929  OE1 GLU   499     -18.860  51.519  11.867  1.00  0.00           O
ATOM   3930  OE2 GLU   499     -21.016  51.698  12.385  1.00  0.00           O
ATOM   3931  C   GLU   499     -18.156  54.314  14.518  1.00  0.00           C
ATOM   3932  O   GLU   499     -19.141  54.843  13.996  1.00  0.00           O
ATOM   3933  N   VAL   500     -17.669  54.688  15.704  1.00  0.00           N
ATOM   3934  CA  VAL   500     -20.763  54.293  19.859 50.00  0.00           C
ATOM   3935  CB  VAL   500     -17.332  56.116  17.661  1.00  0.00           C
ATOM   3936  CG1 VAL   500     -17.636  57.454  18.337  1.00  0.00           C
ATOM   3937  CG2 VAL   500     -17.555  55.060  18.745  1.00  0.00           C
ATOM   3938  C   VAL   500     -18.062  57.124  15.490  1.00  0.00           C
ATOM   3939  O   VAL   500     -19.008  57.912  15.389  1.00  0.00           O
ATOM   3940  N   VAL   501     -16.918  57.297  14.835  1.00  0.00           N
ATOM   3941  CA  VAL   501     -20.588  57.090  17.657 50.00  0.00           C
ATOM   3942  CB  VAL   501     -15.151  58.632  13.718  1.00  0.00           C
ATOM   3943  CG1 VAL   501     -14.836  59.770  12.745  1.00  0.00           C
ATOM   3944  CG2 VAL   501     -14.281  58.904  14.947  1.00  0.00           C
ATOM   3945  C   VAL   501     -17.436  58.574  12.755  1.00  0.00           C
ATOM   3946  O   VAL   501     -17.687  59.668  12.249  1.00  0.00           O
ATOM   3947  N   LYS   502     -17.819  57.423  12.207  1.00  0.00           N
ATOM   3948  CA  LYS   502     -23.415  55.669  16.649 50.00  0.00           C
ATOM   3949  CB  LYS   502     -18.483  56.231  10.130  1.00  0.00           C
ATOM   3950  CG  LYS   502     -19.291  56.291   8.832  1.00  0.00           C
ATOM   3951  CD  LYS   502     -19.014  55.124   7.883  1.00  0.00           C
ATOM   3952  CE  LYS   502     -19.838  55.173   6.595  1.00  0.00           C
ATOM   3953  NZ  LYS   502     -19.494  54.023   5.730  1.00  0.00           N
ATOM   3954  C   LYS   502     -20.121  57.695  11.402  1.00  0.00           C
ATOM   3955  O   LYS   502     -20.874  58.326  10.656  1.00  0.00           O
ATOM   3956  N   GLU   503     -20.500  57.229  12.586  1.00  0.00           N
ATOM   3957  CA  GLU   503     -25.235  55.775  15.868 50.00  0.00           C
ATOM   3958  CB  GLU   503     -22.182  56.411  14.164  1.00  0.00           C
ATOM   3959  CG  GLU   503     -22.201  54.961  13.676  1.00  0.00           C
ATOM   3960  CD  GLU   503     -23.310  54.827  12.643  1.00  0.00           C
ATOM   3961  OE1 GLU   503     -24.472  55.185  12.972  1.00  0.00           O
ATOM   3962  OE2 GLU   503     -23.010  54.365  11.510  1.00  0.00           O
ATOM   3963  C   GLU   503     -22.107  58.847  13.561  1.00  0.00           C
ATOM   3964  O   GLU   503     -23.240  59.222  13.897  1.00  0.00           O
ATOM   3965  N   ASN   504     -21.029  59.632  13.583  1.00  0.00           N
ATOM   3966  CA  ASN   504     -24.372  59.503  16.827 50.00  0.00           C
ATOM   3967  CB  ASN   504     -22.050  61.879  13.304  1.00  0.00           C
ATOM   3968  CG  ASN   504     -21.552  62.064  11.878  1.00  0.00           C
ATOM   3969  OD1 ASN   504     -20.354  61.986  11.610  1.00  0.00           O
ATOM   3970  ND2 ASN   504     -22.445  62.320  10.884  1.00  0.00           N
ATOM   3971  C   ASN   504     -21.216  61.102  15.580  1.00  0.00           C
ATOM   3972  O   ASN   504     -21.722  62.097  16.095  1.00  0.00           O
ATOM   3973  N   ARG   505     -20.779  60.075  16.298  1.00  0.00           N
ATOM   3974  CA  ARG   505     -23.440  58.913  20.457 50.00  0.00           C
ATOM   3975  CB  ARG   505     -21.465  58.797  18.293  1.00  0.00           C
ATOM   3976  CG  ARG   505     -22.295  58.069  17.244  1.00  0.00           C
ATOM   3977  CD  ARG   505     -23.626  57.538  17.768  1.00  0.00           C
ATOM   3978  NE  ARG   505     -23.474  56.493  18.778  1.00  0.00           N
ATOM   3979  CZ  ARG   505     -24.482  56.001  19.493  1.00  0.00           C
ATOM   3980  NH1 ARG   505     -24.262  55.049  20.394  1.00  0.00           N
ATOM   3981  NH2 ARG   505     -25.714  56.472  19.315  1.00  0.00           N
ATOM   3982  C   ARG   505     -19.444  60.319  18.375  1.00  0.00           C
ATOM   3983  O   ARG   505     -18.883  59.442  19.036  1.00  0.00           O
ATOM   3984  N   SER   506     -18.911  61.521  18.153  1.00  0.00           N
ATOM   3985  CA  SER   506     -22.105  59.751  22.622 50.00  0.00           C
ATOM   3986  CB  SER   506     -17.143  63.174  17.831  1.00  0.00           C
ATOM   3987  OG  SER   506     -17.071  62.965  16.428  1.00  0.00           O
ATOM   3988  C   SER   506     -17.365  62.015  20.045  1.00  0.00           C
ATOM   3989  O   SER   506     -16.373  61.511  20.616  1.00  0.00           O
ATOM   3990  N   ARG   507     -18.326  62.690  20.682  1.00  0.00           N
ATOM   3991  CA  ARG   507     -21.910  60.484  26.195 50.00  0.00           C
ATOM   3992  CB  ARG   507     -19.631  63.557  22.588  1.00  0.00           C
ATOM   3993  CG  ARG   507     -19.691  63.807  24.096  1.00  0.00           C
ATOM   3994  CD  ARG   507     -21.003  64.442  24.560  1.00  0.00           C
ATOM   3995  NE  ARG   507     -22.088  63.446  24.335  1.00  0.00           N
ATOM   3996  CZ  ARG   507     -22.252  62.407  25.205  1.00  0.00           C
ATOM   3997  NH1 ARG   507     -21.300  62.523  26.175  1.00  0.00           N
ATOM   3998  NH2 ARG   507     -23.305  61.658  24.765  1.00  0.00           N
ATOM   3999  C   ARG   507     -18.332  61.415  22.776  1.00  0.00           C
ATOM   4000  O   ARG   507     -17.529  61.150  23.678  1.00  0.00           O
ATOM   4001  N   GLU   508     -19.183  60.529  22.266  1.00  0.00           N
ATOM   4002  CA  GLU   508     -21.881  56.765  26.448 50.00  0.00           C
ATOM   4003  CB  GLU   508     -20.196  58.347  21.839  1.00  0.00           C
ATOM   4004  CG  GLU   508     -21.615  58.475  22.336  1.00  0.00           C
ATOM   4005  CD  GLU   508     -22.204  57.134  22.760  1.00  0.00           C
ATOM   4006  OE1 GLU   508     -21.789  56.603  23.810  1.00  0.00           O
ATOM   4007  OE2 GLU   508     -23.080  56.613  22.036  1.00  0.00           O
ATOM   4008  C   GLU   508     -17.785  58.447  22.540  1.00  0.00           C
ATOM   4009  O   GLU   508     -17.312  57.860  23.506  1.00  0.00           O
ATOM   4010  N   ILE   509     -17.149  58.532  21.372  1.00  0.00           N
ATOM   4011  CA  ILE   509     -18.887  56.518  24.354 50.00  0.00           C
ATOM   4012  CB  ILE   509     -15.313  58.247  19.748  1.00  0.00           C
ATOM   4013  CG2 ILE   509     -15.025  59.735  19.487  1.00  0.00           C
ATOM   4014  CG1 ILE   509     -14.004  57.517  19.401  1.00  0.00           C
ATOM   4015  CD1 ILE   509     -14.139  55.995  19.393  1.00  0.00           C
ATOM   4016  C   ILE   509     -14.803  58.459  22.195  1.00  0.00           C
ATOM   4017  O   ILE   509     -14.026  57.673  22.748  1.00  0.00           O
ATOM   4018  N   LEU   510     -14.815  59.763  22.451  1.00  0.00           N
ATOM   4019  CA  LEU   510     -17.187  59.027  26.650 50.00  0.00           C
ATOM   4020  CB  LEU   510     -14.137  61.862  23.531  1.00  0.00           C
ATOM   4021  CG  LEU   510     -13.620  62.603  22.296  1.00  0.00           C
ATOM   4022  CD1 LEU   510     -13.956  64.093  22.249  1.00  0.00           C
ATOM   4023  CD2 LEU   510     -12.103  62.576  22.115  1.00  0.00           C
ATOM   4024  C   LEU   510     -14.235  59.713  24.813  1.00  0.00           C
ATOM   4025  O   LEU   510     -13.325  59.416  25.587  1.00  0.00           O
ATOM   4026  N   LYS   511     -15.519  59.505  25.086  1.00  0.00           N
ATOM   4027  CA  LYS   511     -18.427  57.038  29.500 50.00  0.00           C
ATOM   4028  CB  LYS   511     -17.486  58.832  26.380  1.00  0.00           C
ATOM   4029  CG  LYS   511     -18.095  60.213  26.631  1.00  0.00           C
ATOM   4030  CD  LYS   511     -19.623  60.204  26.706  1.00  0.00           C
ATOM   4031  CE  LYS   511     -20.233  61.589  26.933  1.00  0.00           C
ATOM   4032  NZ  LYS   511     -21.710  61.494  26.959  1.00  0.00           N
ATOM   4033  C   LYS   511     -15.427  57.420  26.415  1.00  0.00           C
ATOM   4034  O   LYS   511     -15.042  56.950  27.488  1.00  0.00           O
ATOM   4035  N   VAL   512     -15.412  56.735  25.279  1.00  0.00           N
ATOM   4036  CA  VAL   512     -17.016  53.972  27.920 50.00  0.00           C
ATOM   4037  CB  VAL   512     -15.214  54.821  23.781  1.00  0.00           C
ATOM   4038  CG1 VAL   512     -14.556  53.467  23.508  1.00  0.00           C
ATOM   4039  CG2 VAL   512     -16.705  54.601  23.520  1.00  0.00           C
ATOM   4040  C   VAL   512     -13.417  55.367  25.487  1.00  0.00           C
ATOM   4041  O   VAL   512     -12.902  54.459  26.148  1.00  0.00           O
ATOM   4042  N   LEU   513     -12.715  56.389  25.026  1.00  0.00           N
ATOM   4043  CA  LEU   513     -13.750  55.716  27.820 50.00  0.00           C
ATOM   4044  CB  LEU   513     -10.592  57.434  24.399  1.00  0.00           C
ATOM   4045  CG  LEU   513     -10.383  56.762  23.049  1.00  0.00           C
ATOM   4046  CD1 LEU   513      -9.662  57.707  22.147  1.00  0.00           C
ATOM   4047  CD2 LEU   513      -9.624  55.448  23.197  1.00  0.00           C
ATOM   4048  C   LEU   513     -11.044  56.757  26.765  1.00  0.00           C
ATOM   4049  O   LEU   513     -10.008  56.375  27.325  1.00  0.00           O
ATOM   4050  N   GLU   514     -11.991  57.453  27.387  1.00  0.00           N
ATOM   4051  CA  GLU   514     -14.140  56.668  31.441 50.00  0.00           C
ATOM   4052  CB  GLU   514     -12.965  58.817  29.181  1.00  0.00           C
ATOM   4053  CG  GLU   514     -12.774  60.202  28.559  1.00  0.00           C
ATOM   4054  CD  GLU   514     -13.985  61.050  28.918  1.00  0.00           C
ATOM   4055  OE1 GLU   514     -14.923  60.502  29.556  1.00  0.00           O
ATOM   4056  OE2 GLU   514     -13.988  62.258  28.559  1.00  0.00           O
ATOM   4057  C   GLU   514     -11.946  56.587  29.690  1.00  0.00           C
ATOM   4058  O   GLU   514     -11.191  56.462  30.654  1.00  0.00           O
ATOM   4059  N   GLY   515     -12.862  55.679  29.366  1.00  0.00           N
ATOM   4060  CA  GLY   515     -14.469  53.147  32.479 50.00  0.00           C
ATOM   4061  C   GLY   515     -11.919  53.457  29.849  1.00  0.00           C
ATOM   4062  O   GLY   515     -11.132  53.116  30.732  1.00  0.00           O
ATOM   4063  N   HIS   516     -11.858  52.992  28.606  1.00  0.00           N
ATOM   4064  CA  HIS   516     -12.144  51.514  30.119 50.00  0.00           C
ATOM   4065  ND1 HIS   516     -12.256  51.252  24.713  1.00  0.00           N
ATOM   4066  CG  HIS   516     -11.981  51.273  26.065  1.00  0.00           C
ATOM   4067  CB  HIS   516     -10.762  51.910  26.663  1.00  0.00           C
ATOM   4068  NE2 HIS   516     -13.865  50.231  25.638  1.00  0.00           N
ATOM   4069  CD2 HIS   516     -13.015  50.619  26.644  1.00  0.00           C
ATOM   4070  CE1 HIS   516     -13.373  50.631  24.482  1.00  0.00           C
ATOM   4071  C   HIS   516      -9.467  52.422  28.735  1.00  0.00           C
ATOM   4072  O   HIS   516      -8.603  51.569  28.936  1.00  0.00           O
ATOM   4073  N   LYS   517     -18.686  43.485  47.703  1.00  0.00           N
ATOM   4074  CA  LYS   517     -15.414  48.533  39.620 50.00  0.00           C
ATOM   4075  CB  LYS   517     -20.207  42.440  46.117  1.00  0.00           C
ATOM   4076  CG  LYS   517     -20.513  41.296  45.159  1.00  0.00           C
ATOM   4077  CD  LYS   517     -21.991  41.288  44.754  1.00  0.00           C
ATOM   4078  CE  LYS   517     -22.408  42.510  43.916  1.00  0.00           C
ATOM   4079  NZ  LYS   517     -21.965  42.475  42.488  1.00  0.00           N
ATOM   4080  C   LYS   517     -17.745  42.645  45.574  1.00  0.00           C
ATOM   4081  O   LYS   517     -17.190  41.667  45.074  1.00  0.00           O
ATOM   4082  N   TRP   518     -17.470  43.896  45.208  1.00  0.00           N
ATOM   4083  CA  TRP   518     -14.516  50.501  39.960 50.00  0.00           C
ATOM   4084  CB  TRP   518     -16.364  45.730  43.859  1.00  0.00           C
ATOM   4085  CG  TRP   518     -15.305  46.087  42.843  1.00  0.00           C
ATOM   4086  CD2 TRP   518     -13.975  46.554  43.110  1.00  0.00           C
ATOM   4087  CD1 TRP   518     -15.365  46.049  41.480  1.00  0.00           C
ATOM   4088  NE1 TRP   518     -14.253  46.431  40.883  1.00  0.00           N
ATOM   4089  CE2 TRP   518     -13.343  46.760  41.855  1.00  0.00           C
ATOM   4090  CE3 TRP   518     -13.250  46.822  44.291  1.00  0.00           C
ATOM   4091  CZ2 TRP   518     -12.005  47.225  41.743  1.00  0.00           C
ATOM   4092  CZ3 TRP   518     -11.905  47.290  44.185  1.00  0.00           C
ATOM   4093  CH2 TRP   518     -11.306  47.483  42.915  1.00  0.00           C
ATOM   4094  C   TRP   518     -15.054  43.834  44.765  1.00  0.00           C
ATOM   4095  O   TRP   518     -14.286  43.122  44.124  1.00  0.00           O
ATOM   4096  N   ASP   519     -14.782  44.326  45.969  1.00  0.00           N
ATOM   4097  CA  ASP   519     -12.456  51.793  40.194 50.00  0.00           C
ATOM   4098  CB  ASP   519     -13.869  44.413  48.206  1.00  0.00           C
ATOM   4099  CG  ASP   519     -12.595  44.166  49.000  1.00  0.00           C
ATOM   4100  OD1 ASP   519     -11.622  43.630  48.407  1.00  0.00           O
ATOM   4101  OD2 ASP   519     -12.577  44.513  50.211  1.00  0.00           O
ATOM   4102  C   ASP   519     -13.385  42.392  46.677  1.00  0.00           C
ATOM   4103  O   ASP   519     -12.329  41.939  46.206  1.00  0.00           O
ATOM   4104  N   ARG   520      -9.469  58.790  26.845  1.00  0.00           N
ATOM   4105  CA  ARG   520     -10.561  59.682  28.215 50.00  0.00           C
ATOM   4106  CB  ARG   520      -7.634  60.345  27.379  1.00  0.00           C
ATOM   4107  CG  ARG   520      -8.496  61.579  27.652  1.00  0.00           C
ATOM   4108  CD  ARG   520      -7.905  62.517  28.708  1.00  0.00           C
ATOM   4109  NE  ARG   520      -8.846  63.662  28.867  1.00  0.00           N
ATOM   4110  CZ  ARG   520      -8.594  64.627  29.798  1.00  0.00           C
ATOM   4111  NH1 ARG   520      -7.431  64.300  30.433  1.00  0.00           N
ATOM   4112  NH2 ARG   520      -9.604  65.543  29.730  1.00  0.00           N
ATOM   4113  C   ARG   520      -7.193  58.390  25.928  1.00  0.00           C
ATOM   4114  O   ARG   520      -6.986  57.398  26.633  1.00  0.00           O
ATOM   4115  N   ARG   521      -6.565  58.622  24.777  1.00  0.00           N
ATOM   4116  CA  ARG   521      -8.038  58.691  25.704 50.00  0.00           C
ATOM   4117  CB  ARG   521      -5.051  58.008  22.916  1.00  0.00           C
ATOM   4118  CG  ARG   521      -6.109  57.282  22.111  1.00  0.00           C
ATOM   4119  CD  ARG   521      -5.851  57.410  20.628  1.00  0.00           C
ATOM   4120  NE  ARG   521      -6.860  56.692  19.835  1.00  0.00           N
ATOM   4121  CZ  ARG   521      -6.587  55.728  18.970  1.00  0.00           C
ATOM   4122  NH1 ARG   521      -5.325  55.334  18.736  1.00  0.00           N
ATOM   4123  NH2 ARG   521      -7.586  55.148  18.314  1.00  0.00           N
ATOM   4124  C   ARG   521      -4.170  58.430  25.243  1.00  0.00           C
ATOM   4125  O   ARG   521      -3.968  59.626  25.344  1.00  0.00           O
ATOM   4126  N   SER   522      -3.425  57.529  25.870  1.00  0.00           N
ATOM   4127  CA  SER   522      -5.083  58.754  28.083 50.00  0.00           C
ATOM   4128  CB  SER   522      -3.184  58.087  28.217  1.00  0.00           C
ATOM   4129  OG  SER   522      -2.314  58.702  29.156  1.00  0.00           O
ATOM   4130  C   SER   522      -1.306  56.910  26.910  1.00  0.00           C
ATOM   4131  O   SER   522      -1.111  56.161  25.955  1.00  0.00           O
ATOM   4132  N   ILE   523      -0.542  56.954  28.000  1.00  0.00           N
ATOM   4133  CA  ILE   523      -2.928  57.096  30.412 50.00  0.00           C
ATOM   4134  CB  ILE   523       1.918  56.850  27.766  1.00  0.00           C
ATOM   4135  CG2 ILE   523       3.208  56.058  28.036  1.00  0.00           C
ATOM   4136  CG1 ILE   523       1.934  57.142  26.257  1.00  0.00           C
ATOM   4137  CD1 ILE   523       3.091  58.042  25.822  1.00  0.00           C
ATOM   4138  C   ILE   523       0.771  55.903  29.699  1.00  0.00           C
ATOM   4139  O   ILE   523       0.239  56.692  30.486  1.00  0.00           O
ATOM   4140  N   LEU   524       1.451  55.319  30.704  1.00  0.00           N
ATOM   4141  CA  LEU   524      -4.267  59.737  35.219 99.00  0.00           C
ATOM   4142  CB  LEU   524      -5.719  59.343  34.827  1.00  0.00           C
ATOM   4143  CG  LEU   524      -6.876  60.061  35.562  1.00  0.00           C
ATOM   4144  CD1 LEU   524      -8.233  59.552  35.052  1.00  0.00           C
ATOM   4145  CD2 LEU   524      -6.799  61.589  35.432  1.00  0.00           C
ATOM   4146  C   LEU   524      -3.969  59.442  36.679  1.00  0.00           C
ATOM   4147  O   LEU   524      -4.744  59.806  37.561  1.00  0.00           O
ATOM   4148  N   LEU   525       3.563  55.503  32.754  1.00  0.00           N
ATOM   4149  CA  LEU   525       1.604  56.608  34.972 50.00  0.00           C
ATOM   4150  CB  LEU   525       5.141  56.955  34.012  1.00  0.00           C
ATOM   4151  CG  LEU   525       4.600  58.293  33.504  1.00  0.00           C
ATOM   4152  CD1 LEU   525       4.605  59.424  34.531  1.00  0.00           C
ATOM   4153  CD2 LEU   525       5.359  58.888  32.317  1.00  0.00           C
ATOM   4154  C   LEU   525       5.636  54.514  33.466  1.00  0.00           C
ATOM   4155  O   LEU   525       4.964  53.475  33.580  1.00  0.00           O
ATOM   4156  N   ALA   526       6.924  54.562  33.770  1.00  0.00           N
ATOM   4157  CA  ALA   526       4.736  55.790  35.947 50.00  0.00           C
ATOM   4158  CB  ALA   526       9.158  53.521  34.082  1.00  0.00           C
ATOM   4159  C   ALA   526       7.158  52.596  35.243  1.00  0.00           C
ATOM   4160  O   ALA   526       7.457  51.422  35.412  1.00  0.00           O
ATOM   4161  N   PRO   527       6.417  53.277  36.111  1.00  0.00           N
ATOM   4162  CA  PRO   527       5.135  54.443  36.587 50.00  0.00           C
ATOM   4163  CD  PRO   527       6.267  55.435  36.752  1.00  0.00           C
ATOM   4164  CB  PRO   527       5.997  53.570  38.487  1.00  0.00           C
ATOM   4165  CG  PRO   527       5.735  55.036  38.131  1.00  0.00           C
ATOM   4166  C   PRO   527       4.455  52.127  37.144  1.00  0.00           C
ATOM   4167  O   PRO   527       3.864  51.601  38.092  1.00  0.00           O
ATOM   4168  N   GLN   528       3.937  52.256  35.934  1.00  0.00           N
ATOM   4169  CA  GLN   528       2.067  53.531  34.608 50.00  0.00           C
ATOM   4170  CB  GLN   528       2.292  50.478  36.279  1.00  0.00           C
ATOM   4171  CG  GLN   528       3.193  49.352  35.766  1.00  0.00           C
ATOM   4172  CD  GLN   528       2.750  48.056  36.431  1.00  0.00           C
ATOM   4173  OE1 GLN   528       1.963  48.066  37.377  1.00  0.00           O
ATOM   4174  NE2 GLN   528       3.233  46.870  35.975  1.00  0.00           N
ATOM   4175  C   GLN   528       1.526  52.829  36.057  1.00  0.00           C
ATOM   4176  O   GLN   528       0.381  52.439  36.343  1.00  0.00           O
ATOM   4177  N   LEU   529       1.910  54.110  36.117  1.00  0.00           N
ATOM   4178  CA  LEU   529       1.192  56.817  34.034 50.00  0.00           C
ATOM   4179  CB  LEU   529       1.595  56.276  37.389  1.00  0.00           C
ATOM   4180  CG  LEU   529       2.115  55.733  38.721  1.00  0.00           C
ATOM   4181  CD1 LEU   529       2.805  56.763  39.613  1.00  0.00           C
ATOM   4182  CD2 LEU   529       1.047  55.132  39.635  1.00  0.00           C
ATOM   4183  C   LEU   529       0.430  55.815  35.206  1.00  0.00           C
ATOM   4184  O   LEU   529       1.198  56.278  34.380  1.00  0.00           O
ATOM   4185  N   GLN   530      -0.893  55.754  35.022  1.00  0.00           N
ATOM   4186  CA  GLN   530      -1.880  57.208  32.228 50.00  0.00           C
ATOM   4187  CB  GLN   530      -3.113  56.126  33.923  1.00  0.00           C
ATOM   4188  CG  GLN   530      -3.513  54.655  33.792  1.00  0.00           C
ATOM   4189  CD  GLN   530      -3.059  54.164  32.425  1.00  0.00           C
ATOM   4190  OE1 GLN   530      -3.277  54.825  31.412  1.00  0.00           O
ATOM   4191  NE2 GLN   530      -2.403  52.978  32.324  1.00  0.00           N
ATOM   4192  C   GLN   530      -1.308  57.856  33.906  1.00  0.00           C
ATOM   4193  O   GLN   530      -1.568  58.525  34.902  1.00  0.00           O
ATOM   4194  N   GLU   531      -0.829  58.381  32.797  1.00  0.00           N
ATOM   4195  CA  GLU   531      -1.839  60.754  31.883 50.00  0.00           C
ATOM   4196  CB  GLU   531       0.469  60.051  31.621  1.00  0.00           C
ATOM   4197  CG  GLU   531       1.831  59.382  31.817  1.00  0.00           C
ATOM   4198  CD  GLU   531       2.680  59.677  30.589  1.00  0.00           C
ATOM   4199  OE1 GLU   531       2.843  60.880  30.257  1.00  0.00           O
ATOM   4200  OE2 GLU   531       3.178  58.700  29.965  1.00  0.00           O
ATOM   4201  C   GLU   531      -1.727  60.669  32.603  1.00  0.00           C
ATOM   4202  O   GLU   531      -1.653  61.857  32.838  1.00  0.00           O
ATOM   4203  N   PRO   532      -2.854  60.070  32.202  1.00  0.00           N
ATOM   4204  CA  PRO   532      -4.456  62.111  30.047 50.00  0.00           C
ATOM   4205  CD  PRO   532      -3.336  58.146  31.247  1.00  0.00           C
ATOM   4206  CB  PRO   532      -5.085  60.013  31.146  1.00  0.00           C
ATOM   4207  CG  PRO   532      -4.814  58.508  31.100  1.00  0.00           C
ATOM   4208  C   PRO   532      -4.736  60.924  33.434  1.00  0.00           C
ATOM   4209  O   PRO   532      -4.947  62.024  33.901  1.00  0.00           O
ATOM   4210  N   THR   533      -4.969  59.809  34.115  1.00  0.00           N
ATOM   4211  CA  THR   533      -3.854  60.917  30.621 50.00  0.00           C
ATOM   4212  CB  THR   533      -6.922  58.800  35.185  1.00  0.00           C
ATOM   4213  OG1 THR   533      -6.387  57.496  35.012  1.00  0.00           O
ATOM   4214  CG2 THR   533      -7.783  59.163  33.964  1.00  0.00           C
ATOM   4215  C   THR   533      -4.992  59.586  36.607  1.00  0.00           C
ATOM   4216  O   THR   533      -5.544  59.675  37.689  1.00  0.00           O
ATOM   4217  N   ASN   534      -3.716  59.232  36.474  1.00  0.00           N
ATOM   4218  CA  ASN   534      -2.409  58.556  31.918 50.00  0.00           C
ATOM   4219  CB  ASN   534      -2.749  60.052  38.622  1.00  0.00           C
ATOM   4220  CG  ASN   534      -1.984  61.193  37.966  1.00  0.00           C
ATOM   4221  OD1 ASN   534      -1.123  60.971  37.115  1.00  0.00           O
ATOM   4222  ND2 ASN   534      -2.257  62.476  38.324  1.00  0.00           N
ATOM   4223  C   ASN   534      -3.174  57.617  38.354  1.00  0.00           C
ATOM   4224  O   ASN   534      -2.657  57.412  39.417  1.00  0.00           O
ATOM   4225  N   SER   535      -4.013  56.746  37.823  1.00  0.00           N
ATOM   4226  CA  SER   535      -1.560  55.718  32.322 50.00  0.00           C
ATOM   4227  CB  SER   535      -5.433  54.768  37.902  1.00  0.00           C
ATOM   4228  OG  SER   535      -5.583  53.487  38.495  1.00  0.00           O
ATOM   4229  C   SER   535      -2.922  54.638  38.297  1.00  0.00           C
ATOM   4230  O   SER   535      -2.295  54.675  37.247  1.00  0.00           O
ATOM   4231  N   SER   536      -2.556  53.903  39.328  1.00  0.00           N
ATOM   4232  CA  SER   536       0.284  52.827  33.520 50.00  0.00           C
ATOM   4233  CB  SER   536      -0.709  53.031  40.681  1.00  0.00           C
ATOM   4234  OG  SER   536       0.408  52.155  40.676  1.00  0.00           O
ATOM   4235  C   SER   536      -1.841  51.646  38.940  1.00  0.00           C
ATOM   4236  O   SER   536      -2.564  51.036  39.700  1.00  0.00           O
ATOM   4237  N   ASP   537      -1.447  51.132  37.776  1.00  0.00           N
ATOM   4238  CA  ASP   537       0.643  50.916  34.331 50.00  0.00           C
ATOM   4239  CB  ASP   537      -1.491  49.440  35.992  1.00  0.00           C
ATOM   4240  CG  ASP   537      -2.426  50.210  35.071  1.00  0.00           C
ATOM   4241  OD1 ASP   537      -3.407  50.808  35.589  1.00  0.00           O
ATOM   4242  OD2 ASP   537      -2.173  50.212  33.836  1.00  0.00           O
ATOM   4243  C   ASP   537      -1.347  48.692  38.379  1.00  0.00           C
ATOM   4244  O   ASP   537      -0.375  48.881  39.071  1.00  0.00           O
ATOM   4245  N   GLY   538      -2.033  47.569  38.399  1.00  0.00           N
ATOM   4246  CA  GLY   538       0.557  48.096  36.746 50.00  0.00           C
ATOM   4247  C   GLY   538      -0.653  45.525  38.579  1.00  0.00           C
ATOM   4248  O   GLY   538      -0.599  45.399  37.377  1.00  0.00           O
ATOM   4249  N   ALA   539       0.184  44.921  39.414  1.00  0.00           N
ATOM   4250  CA  ALA   539       2.979  45.276  37.168 50.00  0.00           C
ATOM   4251  CB  ALA   539       2.427  43.947  39.708  1.00  0.00           C
ATOM   4252  C   ALA   539       0.340  42.522  39.461  1.00  0.00           C
ATOM   4253  O   ALA   539       0.124  42.282  40.623  1.00  0.00           O
ATOM   4254  N   GLY   540      -0.089  41.737  38.499  1.00  0.00           N
ATOM   4255  CA  GLY   540       0.739  42.424  37.741 50.00  0.00           C
ATOM   4256  C   GLY   540      -0.084  39.366  38.912  1.00  0.00           C
ATOM   4257  O   GLY   540       0.797  39.104  38.108  1.00  0.00           O
ATOM   4258  N   GLU   541      -0.291  38.668  40.026  1.00  0.00           N
ATOM   4259  CA  GLU   541       2.365  39.648  39.670 50.00  0.00           C
ATOM   4260  CB  GLU   541       0.024  36.952  41.742  1.00  0.00           C
ATOM   4261  CG  GLU   541       0.415  37.859  42.911  1.00  0.00           C
ATOM   4262  CD  GLU   541      -0.130  37.239  44.189  1.00  0.00           C
ATOM   4263  OE1 GLU   541      -0.819  36.188  44.091  1.00  0.00           O
ATOM   4264  OE2 GLU   541       0.134  37.809  45.282  1.00  0.00           O
ATOM   4265  C   GLU   541       0.282  36.351  39.398  1.00  0.00           C
ATOM   4266  O   GLU   541      -0.795  36.142  38.881  1.00  0.00           O
ATOM   4267  N   VAL   542       1.336  35.564  39.231  1.00  0.00           N
ATOM   4268  CA  VAL   542       2.199  36.225  38.621 50.00  0.00           C
ATOM   4269  CB  VAL   542       2.408  34.032  37.641  1.00  0.00           C
ATOM   4270  CG1 VAL   542       2.379  32.656  36.973  1.00  0.00           C
ATOM   4271  CG2 VAL   542       2.481  35.035  36.487  1.00  0.00           C
ATOM   4272  C   VAL   542       1.180  33.161  39.649  1.00  0.00           C
ATOM   4273  O   VAL   542       2.033  33.047  40.507  1.00  0.00           O
ATOM   4274  N   ASN   543       0.170  32.332  39.569  1.00  0.00           N
ATOM   4275  CA  ASN   543       1.832  33.694  41.160 50.00  0.00           C
ATOM   4276  CB  ASN   543      -1.316  30.547  40.244  1.00  0.00           C
ATOM   4277  CG  ASN   543      -1.588  29.403  41.213  1.00  0.00           C
ATOM   4278  OD1 ASN   543      -0.678  28.797  41.756  1.00  0.00           O
ATOM   4279  ND2 ASN   543      -2.855  29.110  41.425  1.00  0.00           N
ATOM   4280  C   ASN   543       1.118  30.190  40.256  1.00  0.00           C
ATOM   4281  O   ASN   543       1.156  29.584  39.171  1.00  0.00           O
ATOM   4282  N   GLY   544       0.800  28.908  40.412  1.00  0.00           N
ATOM   4283  CA  GLY   544       3.661  30.361  40.924 50.00  0.00           C
ATOM   4284  C   GLY   544       1.819  27.482  38.698  1.00  0.00           C
ATOM   4285  O   GLY   544       1.993  26.317  38.328  1.00  0.00           O
ATOM   4286  N   ARG   545       1.651  28.474  37.827  1.00  0.00           N
ATOM   4287  CA  ARG   545       2.300  29.964  37.751 50.00  0.00           C
ATOM   4288  CB  ARG   545       1.563  29.407  35.391  1.00  0.00           C
ATOM   4289  CG  ARG   545       0.202  30.106  35.373  1.00  0.00           C
ATOM   4290  CD  ARG   545       0.149  31.319  34.440  1.00  0.00           C
ATOM   4291  NE  ARG   545      -1.215  31.908  34.549  1.00  0.00           N
ATOM   4292  CZ  ARG   545      -1.518  33.057  33.876  1.00  0.00           C
ATOM   4293  NH1 ARG   545      -0.402  33.444  33.193  1.00  0.00           N
ATOM   4294  NH2 ARG   545      -2.818  33.368  34.151  1.00  0.00           N
ATOM   4295  C   ARG   545       2.880  27.562  35.810  1.00  0.00           C
ATOM   4296  O   ARG   545       2.799  26.747  34.887  1.00  0.00           O
ATOM   4297  N   SER   546       4.055  27.883  36.342  1.00  0.00           N
ATOM   4298  CA  SER   546       5.479  28.497  36.574 50.00  0.00           C
ATOM   4299  CB  SER   546       6.710  27.570  36.268  1.00  0.00           C
ATOM   4300  OG  SER   546       6.908  27.205  37.625  1.00  0.00           O
ATOM   4301  C   SER   546       5.329  25.786  36.246  1.00  0.00           C
ATOM   4302  O   SER   546       5.950  24.960  35.571  1.00  0.00           O
ATOM   4303  N   SER   547       4.680  25.424  37.353  1.00  0.00           N
ATOM   4304  CA  SER   547       5.839  25.881  38.997 50.00  0.00           C
ATOM   4305  CB  SER   547       3.749  23.362  38.801  1.00  0.00           C
ATOM   4306  OG  SER   547       4.032  21.974  38.895  1.00  0.00           O
ATOM   4307  C   SER   547       4.206  23.168  36.580  1.00  0.00           C
ATOM   4308  O   SER   547       4.908  22.285  36.082  1.00  0.00           O
ATOM   4309  N   GLU   548       2.958  23.380  36.172  1.00  0.00           N
ATOM   4310  CA  GLU   548       2.945  24.466  37.773 50.00  0.00           C
ATOM   4311  CB  GLU   548       0.891  23.223  34.874  1.00  0.00           C
ATOM   4312  CG  GLU   548      -0.030  22.653  35.955  1.00  0.00           C
ATOM   4313  CD  GLU   548      -1.401  23.292  35.790  1.00  0.00           C
ATOM   4314  OE1 GLU   548      -1.537  24.183  34.909  1.00  0.00           O
ATOM   4315  OE2 GLU   548      -2.332  22.897  36.542  1.00  0.00           O
ATOM   4316  C   GLU   548       3.035  23.100  33.711  1.00  0.00           C
ATOM   4317  O   GLU   548       3.266  22.244  32.832  1.00  0.00           O
ATOM   4318  N   GLY   549       3.398  24.369  33.589  1.00  0.00           N
ATOM   4319  CA  GLY   549       3.239  25.565  34.675 50.00  0.00           C
ATOM   4320  C   GLY   549       5.473  24.218  32.295  1.00  0.00           C
ATOM   4321  O   GLY   549       5.927  23.814  31.227  1.00  0.00           O
ATOM   4322  N   GLY   550       6.184  24.149  33.409  1.00  0.00           N
ATOM   4323  CA  GLY   550       6.636  24.215  34.626 50.00  0.00           C
ATOM   4324  C   GLY   550       7.437  22.026  32.929  1.00  0.00           C
ATOM   4325  O   GLY   550       8.257  21.574  32.111  1.00  0.00           O
ATOM   4326  N   ASN   551       6.446  21.305  33.426  1.00  0.00           N
ATOM   4327  CA  ASN   551       5.739  20.941  35.536 50.00  0.00           C
ATOM   4328  CB  ASN   551       5.027  19.350  33.776  1.00  0.00           C
ATOM   4329  CG  ASN   551       5.520  19.061  35.187  1.00  0.00           C
ATOM   4330  OD1 ASN   551       6.720  18.961  35.431  1.00  0.00           O
ATOM   4331  ND2 ASN   551       4.616  18.913  36.194  1.00  0.00           N
ATOM   4332  C   ASN   551       5.904  19.921  31.483  1.00  0.00           C
ATOM   4333  O   ASN   551       6.458  19.107  30.763  1.00  0.00           O
ATOM   4334  N   SER   552       5.070  20.836  30.982  1.00  0.00           N
ATOM   4335  CA  SER   552       3.244  21.200  33.193 50.00  0.00           C
ATOM   4336  CB  SER   552       3.742  21.931  29.150  1.00  0.00           C
ATOM   4337  OG  SER   552       2.580  21.711  29.903  1.00  0.00           O
ATOM   4338  C   SER   552       6.011  21.148  28.700  1.00  0.00           C
ATOM   4339  O   SER   552       6.259  20.528  27.661  1.00  0.00           O
ATOM   4340  N   GLY   553       6.810  22.079  29.187  1.00  0.00           N
ATOM   4341  CA  GLY   553       5.496  22.015  30.918 50.00  0.00           C
ATOM   4342  C   GLY   553       8.984  21.250  28.378  1.00  0.00           C
ATOM   4343  O   GLY   553       9.534  21.031  27.327  1.00  0.00           O
ATOM   4344  N   GLY   554       9.120  20.451  29.426  1.00  0.00           N
ATOM   4345  CA  GLY   554       7.770  19.083  31.648 50.00  0.00           C
ATOM   4346  C   GLY   554       9.517  18.261  28.389  1.00  0.00           C
ATOM   4347  O   GLY   554      10.272  17.758  27.554  1.00  0.00           O
ATOM   4348  N   GLY   555       8.215  17.992  28.437  1.00  0.00           N
ATOM   4349  CA  GLY   555       5.277  16.893  31.598 50.00  0.00           C
ATOM   4350  C   GLY   555       7.758  17.552  26.044  1.00  0.00           C
ATOM   4351  O   GLY   555       8.245  16.795  25.195  1.00  0.00           O
ATOM   4352  N   HIS   556       7.477  18.827  25.751  1.00  0.00           N
ATOM   4353  CA  HIS   556       4.005  18.172  28.698 50.00  0.00           C
ATOM   4354  ND1 HIS   556       6.602  20.986  21.804  1.00  0.00           N
ATOM   4355  CG  HIS   556       7.363  21.357  22.894  1.00  0.00           C
ATOM   4356  CB  HIS   556       7.183  20.793  24.272  1.00  0.00           C
ATOM   4357  NE2 HIS   556       7.965  22.438  21.081  1.00  0.00           N
ATOM   4358  CD2 HIS   556       8.229  22.284  22.420  1.00  0.00           C
ATOM   4359  CE1 HIS   556       6.973  21.638  20.743  1.00  0.00           C
ATOM   4360  C   HIS   556       9.110  19.270  23.907  1.00  0.00           C
ATOM   4361  O   HIS   556       9.414  18.914  22.709  1.00  0.00           O
ATOM   4362  N   HIS   557      10.023  19.635  24.797  1.00  0.00           N
ATOM   4363  CA  HIS   557       7.140  18.016  27.306 50.00  0.00           C
ATOM   4364  ND1 HIS   557      12.516  22.431  26.755  1.00  0.00           N
ATOM   4365  CG  HIS   557      12.084  21.699  25.667  1.00  0.00           C
ATOM   4366  CB  HIS   557      12.271  20.218  25.523  1.00  0.00           C
ATOM   4367  NE2 HIS   557      11.603  23.831  25.453  1.00  0.00           N
ATOM   4368  CD2 HIS   557      11.503  22.604  24.844  1.00  0.00           C
ATOM   4369  CE1 HIS   557      12.223  23.686  26.608  1.00  0.00           C
ATOM   4370  C   HIS   557      11.852  18.171  24.129  1.00  0.00           C
ATOM   4371  O   HIS   557      12.588  17.936  23.191  1.00  0.00           O
ATOM   4372  N   ARG   558      11.443  17.224  24.964  1.00  0.00           N
ATOM   4373  CA  ARG   558       8.014  14.557  28.354 50.00  0.00           C
ATOM   4374  CB  ARG   558      11.599  14.997  26.014  1.00  0.00           C
ATOM   4375  CG  ARG   558      12.482  15.370  27.207  1.00  0.00           C
ATOM   4376  CD  ARG   558      12.194  14.545  28.463  1.00  0.00           C
ATOM   4377  NE  ARG   558      10.830  14.912  28.937  1.00  0.00           N
ATOM   4378  CZ  ARG   558      10.320  14.333  30.064  1.00  0.00           C
ATOM   4379  NH1 ARG   558      11.260  13.470  30.550  1.00  0.00           N
ATOM   4380  NH2 ARG   558       9.071  14.843  30.266  1.00  0.00           N
ATOM   4381  C   ARG   558      11.287  15.147  23.559  1.00  0.00           C
ATOM   4382  O   ARG   558      11.842  14.143  23.090  1.00  0.00           O
ATOM   4383  N   ARG   559      10.219  15.729  23.015  1.00  0.00           N
ATOM   4384  CA  ARG   559       5.104  13.692  26.900 50.00  0.00           C
ATOM   4385  CB  ARG   559       8.228  15.857  21.557  1.00  0.00           C
ATOM   4386  CG  ARG   559       7.208  15.191  22.483  1.00  0.00           C
ATOM   4387  CD  ARG   559       5.784  15.724  22.309  1.00  0.00           C
ATOM   4388  NE  ARG   559       4.907  14.970  23.249  1.00  0.00           N
ATOM   4389  CZ  ARG   559       3.617  15.369  23.448  1.00  0.00           C
ATOM   4390  NH1 ARG   559       3.398  16.469  22.669  1.00  0.00           N
ATOM   4391  NH2 ARG   559       3.049  14.513  24.346  1.00  0.00           N
ATOM   4392  C   ARG   559      10.497  15.793  20.578  1.00  0.00           C
ATOM   4393  O   ARG   559      10.492  15.134  19.549  1.00  0.00           O
ATOM   4394  N   ASP   560      11.168  16.941  20.682  1.00  0.00           N
ATOM   4395  CA  ASP   560       6.043  14.832  23.904 50.00  0.00           C
ATOM   4396  CB  ASP   560      11.525  18.924  19.288  1.00  0.00           C
ATOM   4397  CG  ASP   560      10.036  18.966  18.976  1.00  0.00           C
ATOM   4398  OD1 ASP   560       9.616  18.281  18.005  1.00  0.00           O
ATOM   4399  OD2 ASP   560       9.299  19.683  19.704  1.00  0.00           O
ATOM   4400  C   ASP   560      13.426  17.343  19.677  1.00  0.00           C
ATOM   4401  O   ASP   560      14.121  17.141  18.702  1.00  0.00           O
ATOM   4402  N   MET   561      13.948  17.463  20.894  1.00  0.00           N
ATOM   4403  CA  MET   561       9.546  14.414  24.105 50.00  0.00           C
ATOM   4404  CB  MET   561      15.806  17.804  22.435  1.00  0.00           C
ATOM   4405  CG  MET   561      15.701  19.330  22.440  1.00  0.00           C
ATOM   4406  SD  MET   561      16.748  20.161  21.206  1.00  0.00           S
ATOM   4407  CE  MET   561      18.316  19.729  22.013  1.00  0.00           C
ATOM   4408  C   MET   561      15.568  15.670  21.185  1.00  0.00           C
ATOM   4409  O   MET   561      14.646  14.883  21.424  1.00  0.00           O
ATOM   4410  N   GLY   562      16.822  15.292  20.998  1.00  0.00           N
ATOM   4411  CA  GLY   562      11.674  11.317  24.576 50.00  0.00           C
ATOM   4412  C   GLY   562      17.135  13.903  22.923  1.00  0.00           C
ATOM   4413  O   GLY   562      17.689  14.741  23.679  1.00  0.00           O
ATOM   4414  N   PRO   563      16.415  12.885  23.372  1.00  0.00           N
ATOM   4415  CA  PRO   563      11.745  11.152  27.954 50.00  0.00           C
ATOM   4416  CD  PRO   563      15.149  11.275  22.439  1.00  0.00           C
ATOM   4417  CB  PRO   563      14.988  11.645  24.967  1.00  0.00           C
ATOM   4418  CG  PRO   563      14.277  11.185  23.693  1.00  0.00           C
ATOM   4419  C   PRO   563      17.059  12.848  25.751  1.00  0.00           C
ATOM   4420  O   PRO   563      17.084  13.606  26.731  1.00  0.00           O
ATOM   4421  N   SER   564      18.046  12.039  25.464  1.00  0.00           N
ATOM   4422  CA  SER   564      15.280  10.411  28.474 50.00  0.00           C
ATOM   4423  CB  SER   564      20.119  10.874  25.733  1.00  0.00           C
ATOM   4424  OG  SER   564      19.496   9.611  25.922  1.00  0.00           O
ATOM   4425  C   SER   564      20.020  13.308  26.336  1.00  0.00           C
ATOM   4426  O   SER   564      20.611  13.592  27.367  1.00  0.00           O
ATOM   4427  N   TRP   565      20.061  14.083  25.257  1.00  0.00           N
ATOM   4428  CA  TRP   565      15.850  12.951  26.332 50.00  0.00           C
ATOM   4429  CB  TRP   565      20.736  16.109  23.979  1.00  0.00           C
ATOM   4430  CG  TRP   565      21.774  15.541  23.040  1.00  0.00           C
ATOM   4431  CD2 TRP   565      23.198  15.683  23.139  1.00  0.00           C
ATOM   4432  CD1 TRP   565      21.604  14.785  21.916  1.00  0.00           C
ATOM   4433  NE1 TRP   565      22.731  14.450  21.319  1.00  0.00           N
ATOM   4434  CE2 TRP   565      23.766  14.986  22.041  1.00  0.00           C
ATOM   4435  CE3 TRP   565      24.056  16.335  24.049  1.00  0.00           C
ATOM   4436  CZ2 TRP   565      25.169  14.918  21.824  1.00  0.00           C
ATOM   4437  CZ3 TRP   565      25.466  16.272  23.839  1.00  0.00           C
ATOM   4438  CH2 TRP   565      25.999  15.564  22.731  1.00  0.00           C
ATOM   4439  C   TRP   565      19.958  16.350  26.306  1.00  0.00           C
ATOM   4440  O   TRP   565      20.586  17.013  27.115  1.00  0.00           O
ATOM   4441  N   MET   566      18.632  16.422  26.183  1.00  0.00           N
ATOM   4442  CA  MET   566      13.684  15.412  28.191 50.00  0.00           C
ATOM   4443  CB  MET   566      16.415  17.447  26.629  1.00  0.00           C
ATOM   4444  CG  MET   566      15.607  18.379  27.534  1.00  0.00           C
ATOM   4445  SD  MET   566      16.193  20.100  27.560  1.00  0.00           S
ATOM   4446  CE  MET   566      15.532  20.517  25.921  1.00  0.00           C
ATOM   4447  C   MET   566      17.912  16.798  28.511  1.00  0.00           C
ATOM   4448  O   MET   566      18.148  17.622  29.419  1.00  0.00           O
ATOM   4449  N   SER   567      17.760  15.493  28.747  1.00  0.00           N
ATOM   4450  CA  SER   567      14.005  15.134  28.152 50.00  0.00           C
ATOM   4451  CB  SER   567      17.723  13.403  30.108  1.00  0.00           C
ATOM   4452  OG  SER   567      16.363  13.110  29.825  1.00  0.00           O
ATOM   4453  C   SER   567      19.343  15.195  30.684  1.00  0.00           C
ATOM   4454  O   SER   567      19.467  15.470  31.867  1.00  0.00           O
ATOM   4455  N   ALA   568      20.387  15.083  29.886  1.00  0.00           N
ATOM   4456  CA  ALA   568      17.761  14.670  28.414 50.00  0.00           C
ATOM   4457  CB  ALA   568      22.741  15.321  29.291  1.00  0.00           C
ATOM   4458  C   ALA   568      21.740  16.854  30.917  1.00  0.00           C
ATOM   4459  O   ALA   568      22.311  17.095  31.948  1.00  0.00           O
ATOM   4460  N   MET   569      21.151  17.788  30.187  1.00  0.00           N
ATOM   4461  CA  MET   569      17.838  18.392  28.530 50.00  0.00           C
ATOM   4462  CB  MET   569      20.532  20.130  29.630  1.00  0.00           C
ATOM   4463  CG  MET   569      21.471  20.339  28.440  1.00  0.00           C
ATOM   4464  SD  MET   569      23.075  21.084  28.868  1.00  0.00           S
ATOM   4465  CE  MET   569      22.386  22.710  29.292  1.00  0.00           C
ATOM   4466  C   MET   569      20.277  19.308  31.972  1.00  0.00           C
ATOM   4467  O   MET   569      20.678  20.024  32.903  1.00  0.00           O
ATOM   4468  N   HIS   570      19.127  18.622  32.031  1.00  0.00           N
ATOM   4469  CA  HIS   570      16.063  18.442  31.638 50.00  0.00           C
ATOM   4470  ND1 HIS   570      15.371  19.563  32.423  1.00  0.00           N
ATOM   4471  CG  HIS   570      16.016  18.360  32.214  1.00  0.00           C
ATOM   4472  CB  HIS   570      17.052  17.793  33.138  1.00  0.00           C
ATOM   4473  NE2 HIS   570      14.584  18.802  30.610  1.00  0.00           N
ATOM   4474  CD2 HIS   570      15.505  17.884  31.054  1.00  0.00           C
ATOM   4475  CE1 HIS   570      14.531  19.805  31.464  1.00  0.00           C
ATOM   4476  C   HIS   570      19.106  18.204  34.436  1.00  0.00           C
ATOM   4477  O   HIS   570      19.020  18.799  35.527  1.00  0.00           O
ATOM   4478  N   LEU   571      19.876  17.142  34.249  1.00  0.00           N
ATOM   4479  CA  LEU   571      18.988  16.739  33.052 50.00  0.00           C
ATOM   4480  CB  LEU   571      21.199  15.178  34.963  1.00  0.00           C
ATOM   4481  CG  LEU   571      20.111  14.110  34.839  1.00  0.00           C
ATOM   4482  CD1 LEU   571      20.587  12.763  34.296  1.00  0.00           C
ATOM   4483  CD2 LEU   571      19.418  13.738  36.149  1.00  0.00           C
ATOM   4484  C   LEU   571      21.786  17.479  35.759  1.00  0.00           C
ATOM   4485  O   LEU   571      22.070  17.600  36.957  1.00  0.00           O
ATOM   4486  N   VAL   572      22.430  18.103  34.784  1.00  0.00           N
ATOM   4487  CA  VAL   572      21.041  18.776  31.311 50.00  0.00           C
ATOM   4488  CB  VAL   572      24.094  19.548  33.766  1.00  0.00           C
ATOM   4489  CG1 VAL   572      25.052  20.731  33.916  1.00  0.00           C
ATOM   4490  CG2 VAL   572      24.915  18.454  33.080  1.00  0.00           C
ATOM   4491  C   VAL   572      22.875  20.295  35.818  1.00  0.00           C
ATOM   4492  O   VAL   572      23.433  20.752  36.814  1.00  0.00           O
ATOM   4493  N   LEU   573      21.718  20.800  35.375  1.00  0.00           N
ATOM   4494  CA  LEU   573      19.066  21.892  32.144 50.00  0.00           C
ATOM   4495  CB  LEU   573      19.838  22.383  35.306  1.00  0.00           C
ATOM   4496  CG  LEU   573      19.262  23.730  35.691  1.00  0.00           C
ATOM   4497  CD1 LEU   573      20.281  24.814  35.305  1.00  0.00           C
ATOM   4498  CD2 LEU   573      17.883  23.991  35.009  1.00  0.00           C
ATOM   4499  C   LEU   573      20.797  21.653  37.492  1.00  0.00           C
ATOM   4500  O   LEU   573      20.991  22.474  38.378  1.00  0.00           O
ATOM   4501  N   LYS   574      20.311  20.435  37.764  1.00  0.00           N
ATOM   4502  CA  LYS   574      19.416  20.923  35.370 50.00  0.00           C
ATOM   4503  CB  LYS   574      19.414  18.597  39.214  1.00  0.00           C
ATOM   4504  CG  LYS   574      18.000  18.487  38.638  1.00  0.00           C
ATOM   4505  CD  LYS   574      17.423  17.071  38.697  1.00  0.00           C
ATOM   4506  CE  LYS   574      16.012  16.961  38.116  1.00  0.00           C
ATOM   4507  NZ  LYS   574      15.544  15.559  38.186  1.00  0.00           N
ATOM   4508  C   LYS   574      21.223  20.153  40.055  1.00  0.00           C
ATOM   4509  O   LYS   574      21.055  20.340  41.262  1.00  0.00           O
ATOM   4510  N   GLU   575      22.437  19.969  39.492  1.00  0.00           N
ATOM   4511  CA  GLU   575      22.884  20.887  34.955 50.00  0.00           C
ATOM   4512  CB  GLU   575      24.629  18.979  39.658  1.00  0.00           C
ATOM   4513  CG  GLU   575      24.165  17.536  39.867  1.00  0.00           C
ATOM   4514  CD  GLU   575      25.177  16.616  39.200  1.00  0.00           C
ATOM   4515  OE1 GLU   575      26.151  17.144  38.599  1.00  0.00           O
ATOM   4516  OE2 GLU   575      24.989  15.373  39.279  1.00  0.00           O
ATOM   4517  C   GLU   575      24.340  21.397  40.227  1.00  0.00           C
ATOM   4518  O   GLU   575      25.428  21.600  40.818  1.00  0.00           O
ATOM   4519  N   MET   576      23.728  22.349  39.522  1.00  0.00           N
ATOM   4520  CA  MET   576      22.770  23.828  33.081 50.00  0.00           C
ATOM   4521  CB  MET   576      24.836  23.898  38.107  1.00  0.00           C
ATOM   4522  CG  MET   576      23.765  23.828  37.017  1.00  0.00           C
ATOM   4523  SD  MET   576      24.382  24.124  35.333  1.00  0.00           S
ATOM   4524  CE  MET   576      24.652  25.904  35.575  1.00  0.00           C
ATOM   4525  C   MET   576      23.364  24.768  39.957  1.00  0.00           C
ATOM   4526  O   MET   576      22.258  24.530  40.461  1.00  0.00           O
ATOM   4527  N   GLY   577      23.869  25.993  39.877  1.00  0.00           N
ATOM   4528  CA  GLY   577      21.878  27.380  33.339 50.00  0.00           C
ATOM   4529  C   GLY   577      23.587  27.352  41.851  1.00  0.00           C
ATOM   4530  O   GLY   577      24.534  26.748  42.319  1.00  0.00           O
ATOM   4531  N   VAL   578      22.902  28.278  42.550  1.00  0.00           N
ATOM   4532  CA  VAL   578      20.884  29.954  35.081 50.00  0.00           C
ATOM   4533  CB  VAL   578      21.359  29.803  43.421  1.00  0.00           C
ATOM   4534  CG1 VAL   578      21.061  28.911  44.629  1.00  0.00           C
ATOM   4535  CG2 VAL   578      22.463  30.753  43.889  1.00  0.00           C
ATOM   4536  C   VAL   578      22.018  30.052  41.027  1.00  0.00           C
ATOM   4537  O   VAL   578      23.121  30.502  40.912  1.00  0.00           O
ATOM   4538  N   GLU   579      20.998  30.451  40.281  1.00  0.00           N
ATOM   4539  CA  GLU   579      19.137  31.529  32.397 50.00  0.00           C
ATOM   4540  CB  GLU   579      20.865  30.803  37.865  1.00  0.00           C
ATOM   4541  CG  GLU   579      21.940  29.777  37.498  1.00  0.00           C
ATOM   4542  CD  GLU   579      21.598  29.207  36.130  1.00  0.00           C
ATOM   4543  OE1 GLU   579      20.555  29.625  35.559  1.00  0.00           O
ATOM   4544  OE2 GLU   579      22.376  28.347  35.636  1.00  0.00           O
ATOM   4545  C   GLU   579      20.061  32.553  39.459  1.00  0.00           C
ATOM   4546  O   GLU   579      18.965  32.245  39.903  1.00  0.00           O
ATOM   4547  N   LYS   580      20.466  33.809  39.267  1.00  0.00           N
ATOM   4548  CA  LYS   580      17.473  34.932  32.175 50.00  0.00           C
ATOM   4549  CB  LYS   580      20.206  36.201  39.792  1.00  0.00           C
ATOM   4550  CG  LYS   580      20.409  36.068  41.303  1.00  0.00           C
ATOM   4551  CD  LYS   580      21.041  37.302  41.947  1.00  0.00           C
ATOM   4552  CE  LYS   580      21.204  37.186  43.463  1.00  0.00           C
ATOM   4553  NZ  LYS   580      21.801  38.428  44.005  1.00  0.00           N
ATOM   4554  C   LYS   580      19.140  35.297  37.699  1.00  0.00           C
ATOM   4555  O   LYS   580      18.118  35.943  37.462  1.00  0.00           O
ATOM   4556  N   GLU   581      19.896  34.818  36.742  1.00  0.00           N
ATOM   4557  CA  GLU   581      16.174  34.240  29.150 50.00  0.00           C
ATOM   4558  CB  GLU   581      20.289  36.392  34.987  1.00  0.00           C
ATOM   4559  CG  GLU   581      21.812  36.398  35.131  1.00  0.00           C
ATOM   4560  CD  GLU   581      22.308  37.805  34.834  1.00  0.00           C
ATOM   4561  OE1 GLU   581      21.456  38.679  34.521  1.00  0.00           O
ATOM   4562  OE2 GLU   581      23.546  38.026  34.916  1.00  0.00           O
ATOM   4563  C   GLU   581      20.222  33.976  34.511  1.00  0.00           C
ATOM   4564  O   GLU   581      21.263  33.366  34.863  1.00  0.00           O
ATOM   4565  N   VAL   582      19.547  33.780  33.384  1.00  0.00           N
ATOM   4566  CA  VAL   582      15.556  31.437  27.264 50.00  0.00           C
ATOM   4567  CB  VAL   582      18.808  31.768  32.103  1.00  0.00           C
ATOM   4568  CG1 VAL   582      19.044  30.911  30.873  1.00  0.00           C
ATOM   4569  CG2 VAL   582      18.698  30.875  33.359  1.00  0.00           C
ATOM   4570  C   VAL   582      20.044  33.733  31.027  1.00  0.00           C
ATOM   4571  O   VAL   582      19.150  34.499  30.707  1.00  0.00           O
ATOM   4572  N   LEU   583      21.161  33.599  30.338  1.00  0.00           N
ATOM   4573  CA  LEU   583      15.441  31.841  23.966 50.00  0.00           C
ATOM   4574  CB  LEU   583      22.802  34.978  29.122  1.00  0.00           C
ATOM   4575  CG  LEU   583      22.930  36.022  30.233  1.00  0.00           C
ATOM   4576  CD1 LEU   583      24.309  36.667  30.358  1.00  0.00           C
ATOM   4577  CD2 LEU   583      21.989  37.220  30.105  1.00  0.00           C
ATOM   4578  C   LEU   583      21.321  33.339  27.901  1.00  0.00           C
ATOM   4579  O   LEU   583      21.951  32.260  27.870  1.00  0.00           O
ATOM   4580  N   MET   584      20.558  33.786  26.886  1.00  0.00           N
ATOM   4581  CA  MET   584      13.477  30.199  21.865 50.00  0.00           C
ATOM   4582  CB  MET   584      19.310  33.679  24.801  1.00  0.00           C
ATOM   4583  CG  MET   584      19.124  33.094  23.399  1.00  0.00           C
ATOM   4584  SD  MET   584      17.446  33.272  22.724  1.00  0.00           S
ATOM   4585  CE  MET   584      17.870  32.641  21.075  1.00  0.00           C
ATOM   4586  C   MET   584      21.827  33.778  24.812  1.00  0.00           C
ATOM   4587  O   MET   584      21.850  34.918  24.298  1.00  0.00           O
ATOM   4588  N   GLY   585      22.888  33.004  24.891  1.00  0.00           N
ATOM   4589  CA  GLY   585      15.954  29.684  19.747 50.00  0.00           C
ATOM   4590  C   GLY   585      24.269  33.738  23.003  1.00  0.00           C
ATOM   4591  O   GLY   585      24.903  34.678  22.600  1.00  0.00           O
ATOM   4592  N   TYR   586      23.615  32.917  22.188  1.00  0.00           N
ATOM   4593  CA  TYR   586      15.086  28.124  16.721 50.00  0.00           C
ATOM   4594  CB  TYR   586      22.994  31.972  19.961  1.00  0.00           C
ATOM   4595  CG  TYR   586      23.970  30.845  19.981  1.00  0.00           C
ATOM   4596  CD1 TYR   586      23.748  29.747  20.821  1.00  0.00           C
ATOM   4597  CD2 TYR   586      25.124  30.852  19.165  1.00  0.00           C
ATOM   4598  CE1 TYR   586      24.646  28.662  20.865  1.00  0.00           C
ATOM   4599  CE2 TYR   586      26.046  29.757  19.199  1.00  0.00           C
ATOM   4600  CZ  TYR   586      25.789  28.670  20.057  1.00  0.00           C
ATOM   4601  OH  TYR   586      26.650  27.596  20.127  1.00  0.00           O
ATOM   4602  C   TYR   586      22.937  34.475  20.416  1.00  0.00           C
ATOM   4603  O   TYR   586      23.424  35.264  19.579  1.00  0.00           O
ATOM   4604  N   LYS   587      21.781  34.720  21.040  1.00  0.00           N
ATOM   4605  CA  LYS   587      15.618  28.848  16.845 50.00  0.00           C
ATOM   4606  CB  LYS   587      19.714  36.009  21.363  1.00  0.00           C
ATOM   4607  CG  LYS   587      18.744  35.137  20.563  1.00  0.00           C
ATOM   4608  CD  LYS   587      17.316  35.148  21.112  1.00  0.00           C
ATOM   4609  CE  LYS   587      16.348  34.272  20.314  1.00  0.00           C
ATOM   4610  NZ  LYS   587      15.002  34.316  20.930  1.00  0.00           N
ATOM   4611  C   LYS   587      21.947  37.208  21.338  1.00  0.00           C
ATOM   4612  O   LYS   587      22.025  38.240  20.660  1.00  0.00           O
ATOM   4613  N   ALA   588      22.687  37.021  22.426  1.00  0.00           N
ATOM   4614  CA  ALA   588      16.548  29.798  19.822 50.00  0.00           C
ATOM   4615  CB  ALA   588      24.059  37.751  24.357  1.00  0.00           C
ATOM   4616  C   ALA   588      24.662  38.422  22.056  1.00  0.00           C
ATOM   4617  O   ALA   588      24.959  39.582  21.762  1.00  0.00           O
ATOM   4618  N   LYS   589      25.340  37.381  21.582  1.00  0.00           N
ATOM   4619  CA  LYS   589      16.298  28.710  18.576 50.00  0.00           C
ATOM   4620  CB  LYS   589      27.230  36.322  20.131  1.00  0.00           C
ATOM   4621  CG  LYS   589      28.083  35.628  21.195  1.00  0.00           C
ATOM   4622  CD  LYS   589      28.852  34.414  20.668  1.00  0.00           C
ATOM   4623  CE  LYS   589      29.685  33.705  21.737  1.00  0.00           C
ATOM   4624  NZ  LYS   589      30.365  32.529  21.152  1.00  0.00           N
ATOM   4625  C   LYS   589      26.035  38.166  19.339  1.00  0.00           C
ATOM   4626  O   LYS   589      26.694  39.051  18.788  1.00  0.00           O
ATOM   4627  N   LEU   590      24.907  37.697  18.816  1.00  0.00           N
ATOM   4628  CA  LEU   590      15.967  27.832  15.698 50.00  0.00           C
ATOM   4629  CB  LEU   590      23.586  37.196  16.637  1.00  0.00           C
ATOM   4630  CG  LEU   590      24.366  35.961  16.182  1.00  0.00           C
ATOM   4631  CD1 LEU   590      23.554  34.942  15.385  1.00  0.00           C
ATOM   4632  CD2 LEU   590      25.562  36.250  15.276  1.00  0.00           C
ATOM   4633  C   LEU   590      23.420  39.310  17.539  1.00  0.00           C
ATOM   4634  O   LEU   590      23.122  39.928  16.513  1.00  0.00           O
ATOM   4635  N   SER   591      22.903  39.622  18.718  1.00  0.00           N
ATOM   4636  CA  SER   591      16.540  28.864  16.943 50.00  0.00           C
ATOM   4637  CB  SER   591      21.572  41.417  20.181  1.00  0.00           C
ATOM   4638  OG  SER   591      20.515  42.337  19.947  1.00  0.00           O
ATOM   4639  C   SER   591      22.465  41.982  18.188  1.00  0.00           C
ATOM   4640  O   SER   591      23.461  42.591  18.591  1.00  0.00           O
ATOM   4641  N   ALA   592      21.775  42.389  17.132  1.00  0.00           N
ATOM   4642  CA  ALA   592      17.086  28.603  16.414 50.00  0.00           C
ATOM   4643  CB  ALA   592      21.809  44.845  17.195  1.00  0.00           C
ATOM   4644  C   ALA   592      23.584  43.700  15.946  1.00  0.00           C
ATOM   4645  O   ALA   592      24.215  44.753  16.072  1.00  0.00           O
ATOM   4646  N   LEU   593      24.130  42.680  15.347  1.00  0.00           N
ATOM   4647  CA  LEU   593      16.988  27.572  15.434 50.00  0.00           C
ATOM   4648  CB  LEU   593      26.275  41.714  16.032  1.00  0.00           C
ATOM   4649  CG  LEU   593      27.798  41.822  15.934  1.00  0.00           C
ATOM   4650  CD1 LEU   593      28.364  43.218  16.188  1.00  0.00           C
ATOM   4651  CD2 LEU   593      28.573  40.942  16.913  1.00  0.00           C
ATOM   4652  C   LEU   593      25.640  41.987  13.622  1.00  0.00           C
ATOM   4653  O   LEU   593      24.799  41.228  13.207  1.00  0.00           O
ATOM   4654  N   ILE   594      26.628  42.019  12.707  1.00  0.00           N
ATOM   4655  CA  ILE   594       6.529  14.644  18.386 99.00  0.00           C
ATOM   4656  CB  ILE   594       5.002  14.971  18.243  1.00  0.00           C
ATOM   4657  CG2 ILE   594       4.640  15.715  16.938  1.00  0.00           C
ATOM   4658  CG1 ILE   594       4.079  13.752  18.495  1.00  0.00           C
ATOM   4659  CD1 ILE   594       2.579  14.062  18.358  1.00  0.00           C
ATOM   4660  C   ILE   594       6.995  13.564  17.424  1.00  0.00           C
ATOM   4661  O   ILE   594       7.926  13.776  16.649  1.00  0.00           O
ATOM   4662  N   ASN   595       6.358  12.378  17.449  1.00  0.00           N
ATOM   4663  CA  ASN   595       3.044  13.238  17.811 99.00  0.00           C
ATOM   4664  CB  ASN   595       2.582  14.177  16.651  1.00  0.00           C
ATOM   4665  CG  ASN   595       1.114  14.609  16.735  1.00  0.00           C
ATOM   4666  OD1 ASN   595       0.591  15.349  15.900  1.00  0.00           O
ATOM   4667  ND2 ASN   595       0.417  14.144  17.796  1.00  0.00           N
ATOM   4668  C   ASN   595       2.288  11.914  17.844  1.00  0.00           C
ATOM   4669  O   ASN   595       2.895  10.851  17.748  1.00  0.00           O
ATOM   4670  N   ASP   596       0.949  11.969  17.980  1.00  0.00           N
ATOM   4671  CA  ASP   596       1.751  13.356  21.385 99.00  0.00           C
ATOM   4672  CB  ASP   596       2.670  14.614  21.277  1.00  0.00           C
ATOM   4673  CG  ASP   596       3.016  15.204  22.637  1.00  0.00           C
ATOM   4674  OD1 ASP   596       3.753  16.221  22.668  1.00  0.00           O
ATOM   4675  OD2 ASP   596       2.706  14.541  23.655  1.00  0.00           O
ATOM   4676  C   ASP   596       0.397  13.626  22.049  1.00  0.00           C
ATOM   4677  O   ASP   596      -0.655  13.383  21.455  1.00  0.00           O
ATOM   4678  N   ALA   597       0.418  14.134  23.297  1.00  0.00           N
ATOM   4679  CA  ALA   597       2.457  17.091  21.372 99.00  0.00           C
ATOM   4680  CB  ALA   597       1.585  17.607  20.204  1.00  0.00           C
ATOM   4681  C   ALA   597       2.908  18.249  22.280  1.00  0.00           C
ATOM   4682  O   ALA   597       2.670  18.274  23.486  1.00  0.00           O
ATOM   4683  N   LYS   598       3.591  19.260  21.689  1.00  0.00           N
ATOM   4684  CA  LYS   598      -0.440  17.492  18.944 99.00  0.00           C
ATOM   4685  CB  LYS   598      -1.769  17.602  19.748  1.00  0.00           C
ATOM   4686  CG  LYS   598      -3.035  17.101  19.021  1.00  0.00           C
ATOM   4687  CD  LYS   598      -4.315  17.236  19.867  1.00  0.00           C
ATOM   4688  CE  LYS   598      -5.561  16.733  19.124  1.00  0.00           C
ATOM   4689  NZ  LYS   598      -6.748  16.879  19.961  1.00  0.00           N
ATOM   4690  C   LYS   598      -0.502  18.301  17.660  1.00  0.00           C
ATOM   4691  O   LYS   598       0.318  19.187  17.436  1.00  0.00           O
ATOM   4692  N   LYS   599      -1.486  18.004  16.789  1.00  0.00           N
ATOM   4693  CA  LYS   599      -3.068  16.234  21.383 99.00  0.00           C
ATOM   4694  CB  LYS   599      -3.121  17.372  22.445  1.00  0.00           C
ATOM   4695  CG  LYS   599      -3.582  16.951  23.856  1.00  0.00           C
ATOM   4696  CD  LYS   599      -3.613  18.119  24.860  1.00  0.00           C
ATOM   4697  CE  LYS   599      -4.074  17.677  26.255  1.00  0.00           C
ATOM   4698  NZ  LYS   599      -4.088  18.813  27.172  1.00  0.00           N
ATOM   4699  C   LYS   599      -4.435  15.605  21.178  1.00  0.00           C
ATOM   4700  O   LYS   599      -4.958  15.587  20.068  1.00  0.00           O
ATOM   4701  N   LYS   600      -5.039  15.074  22.260  1.00  0.00           N
ATOM   4702  CA  LYS   600      -2.019  18.210  24.459 99.00  0.00           C
ATOM   4703  CB  LYS   600      -2.260  19.669  23.973  1.00  0.00           C
ATOM   4704  CG  LYS   600      -1.354  20.746  24.608  1.00  0.00           C
ATOM   4705  CD  LYS   600      -1.641  22.167  24.088  1.00  0.00           C
ATOM   4706  CE  LYS   600      -0.729  23.219  24.733  1.00  0.00           C
ATOM   4707  NZ  LYS   600      -1.035  24.549  24.211  1.00  0.00           N
ATOM   4708  C   LYS   600      -2.245  18.081  25.955  1.00  0.00           C
ATOM   4709  O   LYS   600      -3.071  17.289  26.400  1.00  0.00           O
ATOM   4710  N   ARG   601      -1.504  18.866  26.762  1.00  0.00           N
ATOM   4711  CA  ARG   601      -0.769  21.361  22.749 99.00  0.00           C
ATOM   4712  CB  ARG   601      -2.122  21.782  22.107  1.00  0.00           C
ATOM   4713  CG  ARG   601      -2.025  22.599  20.797  1.00  0.00           C
ATOM   4714  CD  ARG   601      -3.388  22.991  20.201  1.00  0.00           C
ATOM   4715  NE  ARG   601      -3.114  23.771  18.945  1.00  0.00           N
ATOM   4716  CZ  ARG   601      -4.085  24.270  18.168  1.00  0.00           C
ATOM   4717  NH1 ARG   601      -3.729  24.945  17.078  1.00  0.00           N
ATOM   4718  NH2 ARG   601      -5.373  24.087  18.441  1.00  0.00           N
ATOM   4719  C   ARG   601       0.127  22.557  23.040  1.00  0.00           C
ATOM   4720  O   ARG   601       0.555  22.775  24.169  1.00  0.00           O
ATOM   4721  N   ASP   602       0.431  23.367  22.002  1.00  0.00           N
ATOM   4722  CA  ASP   602       1.061  22.952  25.676 99.00  0.00           C
ATOM   4723  CB  ASP   602       1.657  21.543  25.989  1.00  0.00           C
ATOM   4724  CG  ASP   602       1.926  21.338  27.473  1.00  0.00           C
ATOM   4725  OD1 ASP   602       2.408  20.236  27.838  1.00  0.00           O
ATOM   4726  OD2 ASP   602       1.484  22.199  28.270  1.00  0.00           O
ATOM   4727  C   ASP   602       1.971  24.118  26.073  1.00  0.00           C
ATOM   4728  O   ASP   602       2.337  24.945  25.235  1.00  0.00           O
ATOM   4729  N   ASN   603       2.345  24.191  27.366  1.00  0.00           N
ATOM   4730  CA  ASN   603      -1.743  22.527  28.204 99.00  0.00           C
ATOM   4731  CB  ASN   603      -1.951  21.163  27.472  1.00  0.00           C
ATOM   4732  CG  ASN   603      -3.323  20.525  27.705  1.00  0.00           C
ATOM   4733  OD1 ASN   603      -3.649  19.448  27.201  1.00  0.00           O
ATOM   4734  ND2 ASN   603      -4.178  21.219  28.490  1.00  0.00           N
ATOM   4735  C   ASN   603      -1.823  22.408  29.722  1.00  0.00           C
ATOM   4736  O   ASN   603      -0.821  22.765  30.397  1.00  0.00           O
ATOM   4737  OXT ASN   603      -2.887  21.959  30.225  1.00  0.00           O
TER    4738      ASN   603
END
```
