## Supplementary material for "The endosomal TbTpr86/TbUsp7/SkpZ (TUS) complex controls surface protein abundance in trypanosomes": DA1: ss_report 2.pdf

### Phyre2

| | |
|---|---|
| Email | |
| Description | Undefined |
| Date | Tue Dec 31 14:42:05 GMT 2019 |
| Unique Job ID | 55c641d1d7bceb76 |

#### Secondary structure and disorder prediction

| 1 – 60 | |
|---|---|
| Sequence | M G A E K K I N R P T N S G E V A E N V G A F T E V M V P V W S R L C D K V I A A Y K A G E D I V V L Q H F E E L R A T |
| Secondary structure | |
| SS confidence | |
| Disorder | ? ? ? ? ? ? ? ? ? ? ? ? ? — ? ? ? ? ? ? — ? ? ? — ? ? ? ? ? ? ? — ? ? ? ? |
| Disorder confidence | |

| 61 – 120 | |
|---|---|
| Sequence | P E L D V I A H D S L I S A A P D L T V G A V E K K E D D R A A K K E R L E Y C W G R A L L S A R H L A G D D D E H I N |
| Secondary structure | |
| SS confidence | |
| Disorder | ? ? ? — ? ? ? ? ? ? — ? — ? ? ? ? ? ? ? ? ? — ? ? — ? ? — |
| Disorder confidence | |

| 121 – 180 | |
|---|---|
| Sequence | S I V E S F E Q Q F G C V S G Y L R A L A I N A E S A L E V L Q L Y Q M M E G P V T T R V A T A A L H R F L T N D A S E |
| Secondary structure | |
| SS confidence | |
| Disorder | — ? ? — ? — |
| Disorder confidence | |

| 181 – 240 | |
|---|---|
| Sequence | V S G L L Q F L K L N N V P L T R E F H D Q L E K N F V M R M K T E D A E E V F K D M R N A G V E P V S T H P Y N F I F |
| Secondary structure | |
| SS confidence | |
| Disorder | — |
| Disorder confidence | |

| 241 – 300 | |
|---|---|
| Sequence | M K T P P F K Y M D L L D Y Y E D M L D R G I V P M N S T F R I L S R Q T Q N G G L V N Q H F S A L A R G T T T G S D R |
| Secondary structure | |
| SS confidence | |
| Disorder | — ? — |
| Disorder confidence | |

| 301 – 360 | |
|---|---|
| Sequence | R H E T E G H G G T Q A F F A T R L E E L A R L K S S T T L D D A L R V L R R M D V E G T P L I S G S V V L N V L L S V |
| Secondary structure | |
| SS confidence | |
| Disorder | — |
| Disorder confidence | |

| 361 – 420 | |
|---|---|
| Sequence | F C R A G H I E D T E R V L Q L A K E R G C T P N G A T L S R V L K K S N A D R F L Q A E L I L P D A L E A G D V R L S |
| Secondary structure | |
| SS confidence | |
| Disorder | — ? ? — |
| Disorder confidence | |

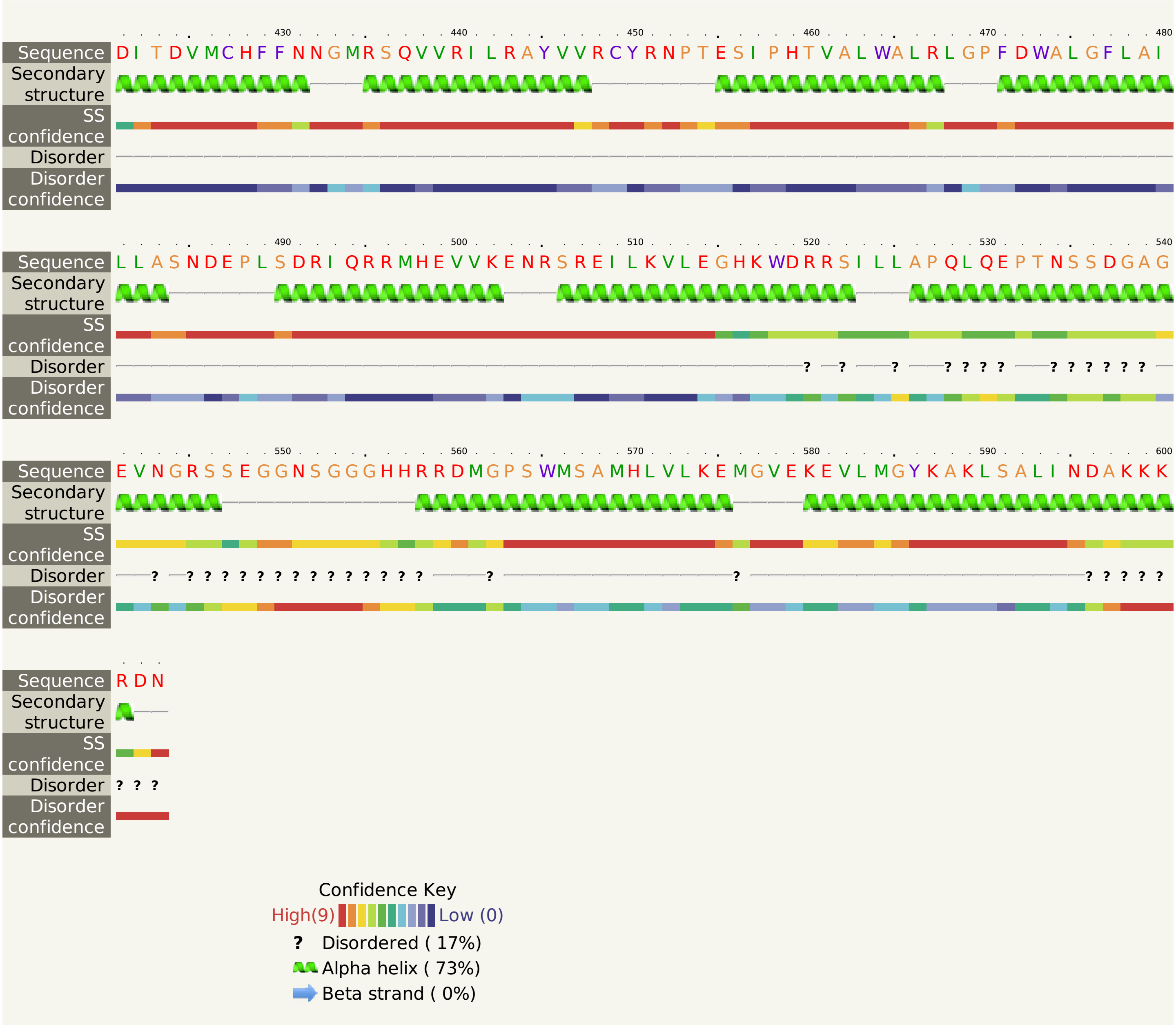

430 440 450 460 470 480
Sequence D I T D V M C H F F N N G M R S Q V V R I L R A Y V V R C Y R N P T E S I P H T V A L W A L R L G P F D W A L G F L A I
Secondary structure
SS confidence
Disorder
Disorder confidence
490 500 510 520 530 540
Sequence L L A S N D E P L S D R I Q R R M H E V V K E N R S R E I L K V L E G H K W D R R S I L L A P Q L Q E P T N S S D G A G
Secondary structure
SS confidence
Disorder ? ? ? ? ? ? ? ? ? ? ? ? ? ?
Disorder confidence
550 560 570 580 590 600
Sequence E V N G R S S E G G N S G G G H H R R D M G P S W M S A M H L V L K E M G V E K E V L M G Y K A K L S A L I N D A K K K
Secondary structure
SS confidence
Disorder ? ? ? ? ? ? ? ? ? ? ? ? ? ? ? ? ? ? ? ? ?
Disorder confidence
Sequence R D N
Secondary structure
SS confidence
Disorder ? ? ?
Disorder confidence
Confidence Key
High(9) Low (0)
? Disordered ( 17%)
Alpha helix ( 73%)
Beta strand ( 0%)
