## Supplementary figures and images for "The endosomal TbTpr86/TbUsp7/SkpZ (TUS) complex controls surface protein abundance in trypanosomes"

### c2xpiA_.19.big 3.png

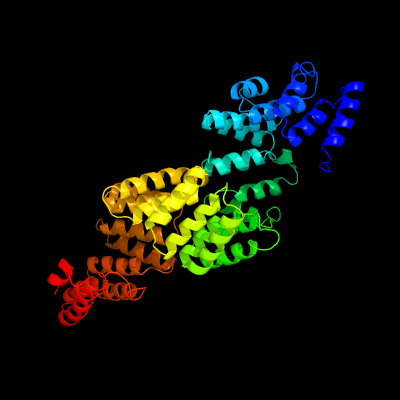

### c3eiqC_.14 2.png

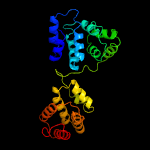

### c3eiqC_.14.big 2.png

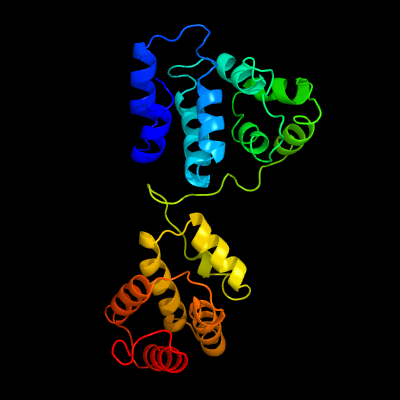

### c3jd5e_.2.big 3.png

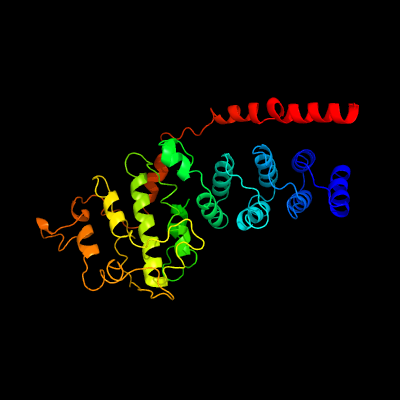

### c3jd5o_.9.big 2.png

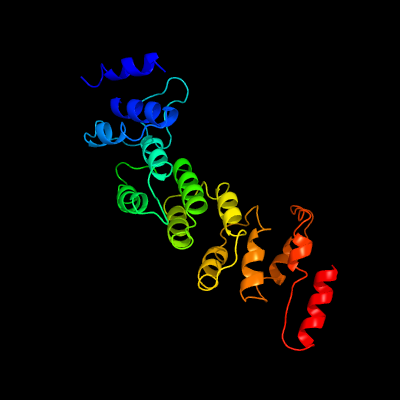

### c3jd5o_.9.png

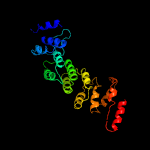

### c3spaA_.15.big 2.png

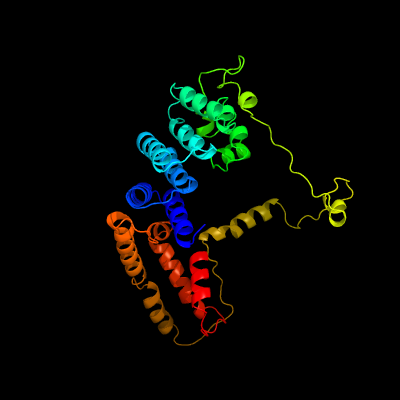

### c3spaA_.15.png

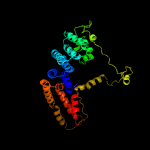

### c4g25A_.5.big 2.png

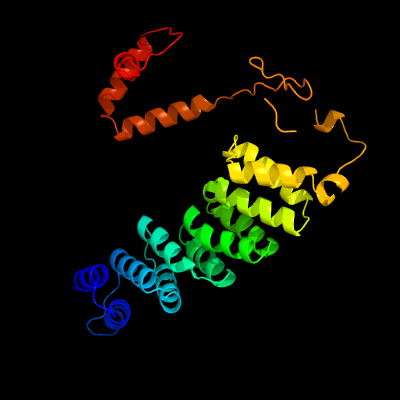

### c4g25A_.5.png

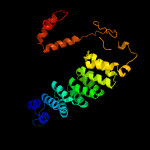

### c4leuA_.12.png

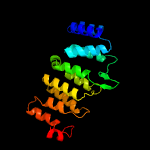

### c4m57A_.1 2.png

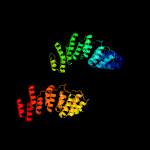

### c4n2sA_.11.big 2.png

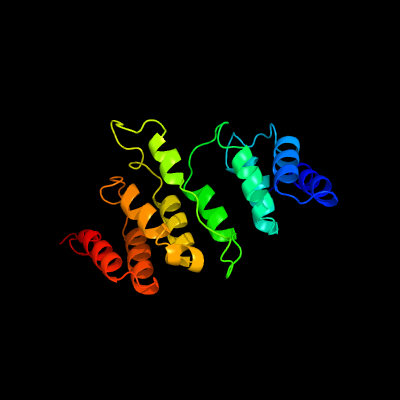

### c4n2sA_.11.png

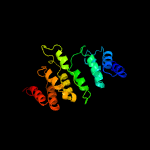

### c4ui9C_.20 3.png

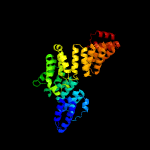

### c4ui9C_.20.big.png

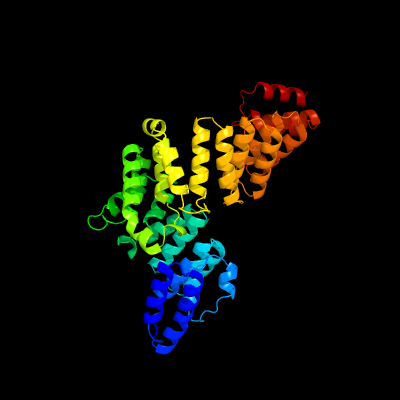

### c4uzyA_.16 2.png

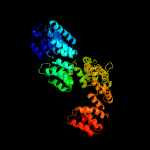

### c4uzyA_.16.big 2.png

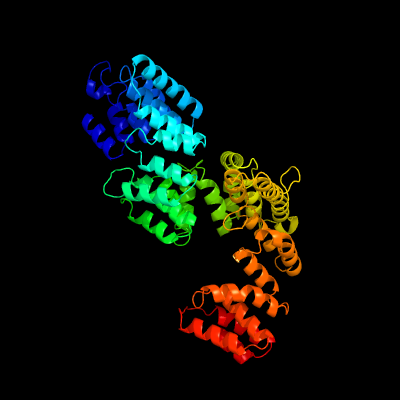

### c4wslA_.7.big.png

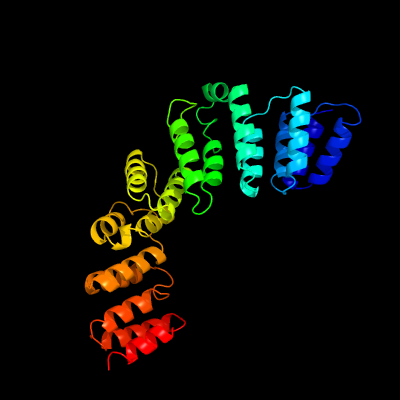

### c5dizB_.6 2.png

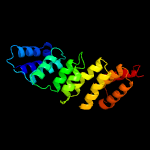

### c5i9fA_.3 2.png

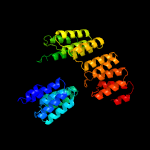

### c5iwwD_.4.big 3.png

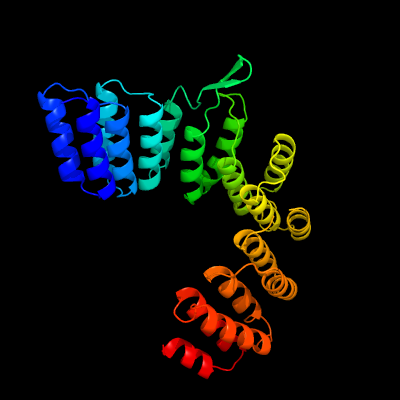

### c5izwA_.17.png

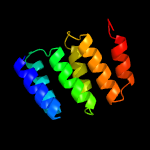

### c5mqfO_.18.big 2.png

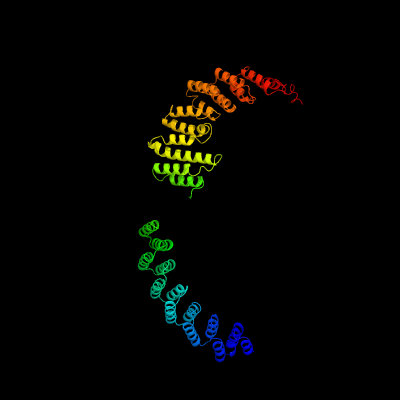

### c5mqfO_.18.png

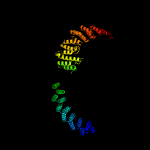

### c6neqo_.10.big.png

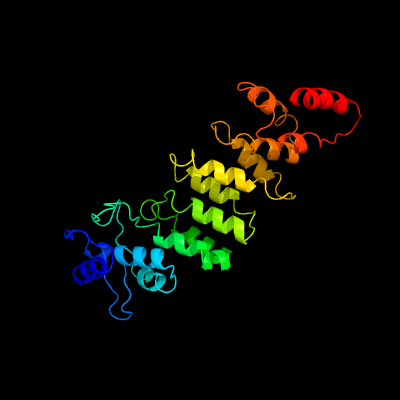

### c6nf8o_.8 2.png

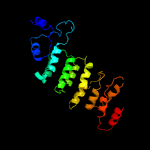

### final.casp.big 2.png

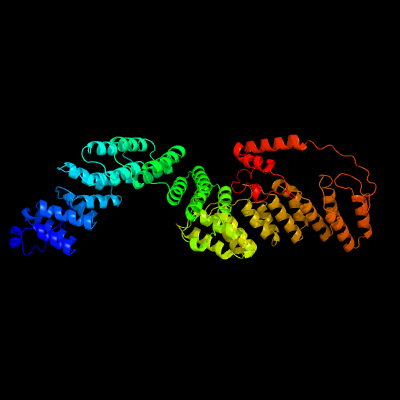

### final.pub.black 2.png

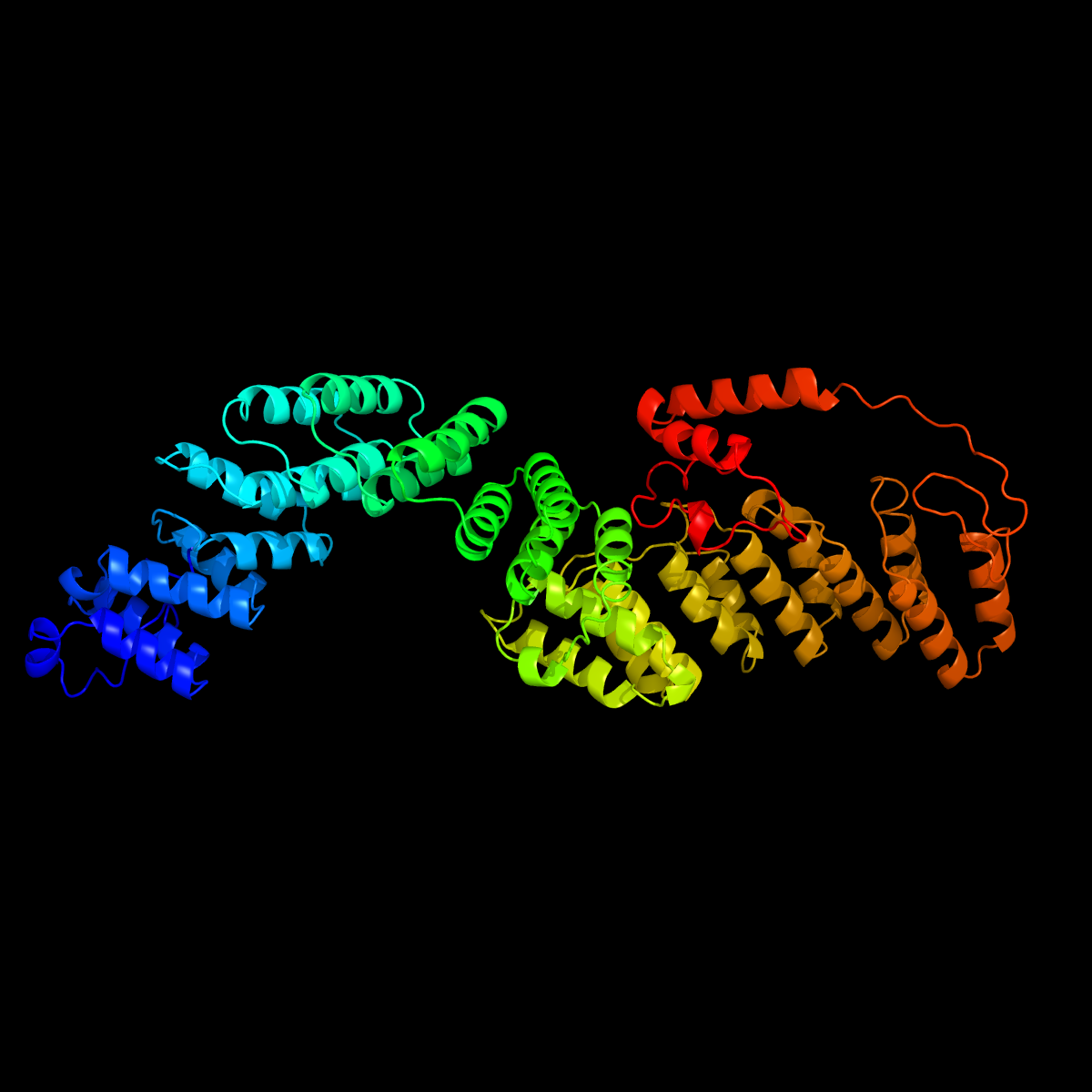

### final.pub.white.png

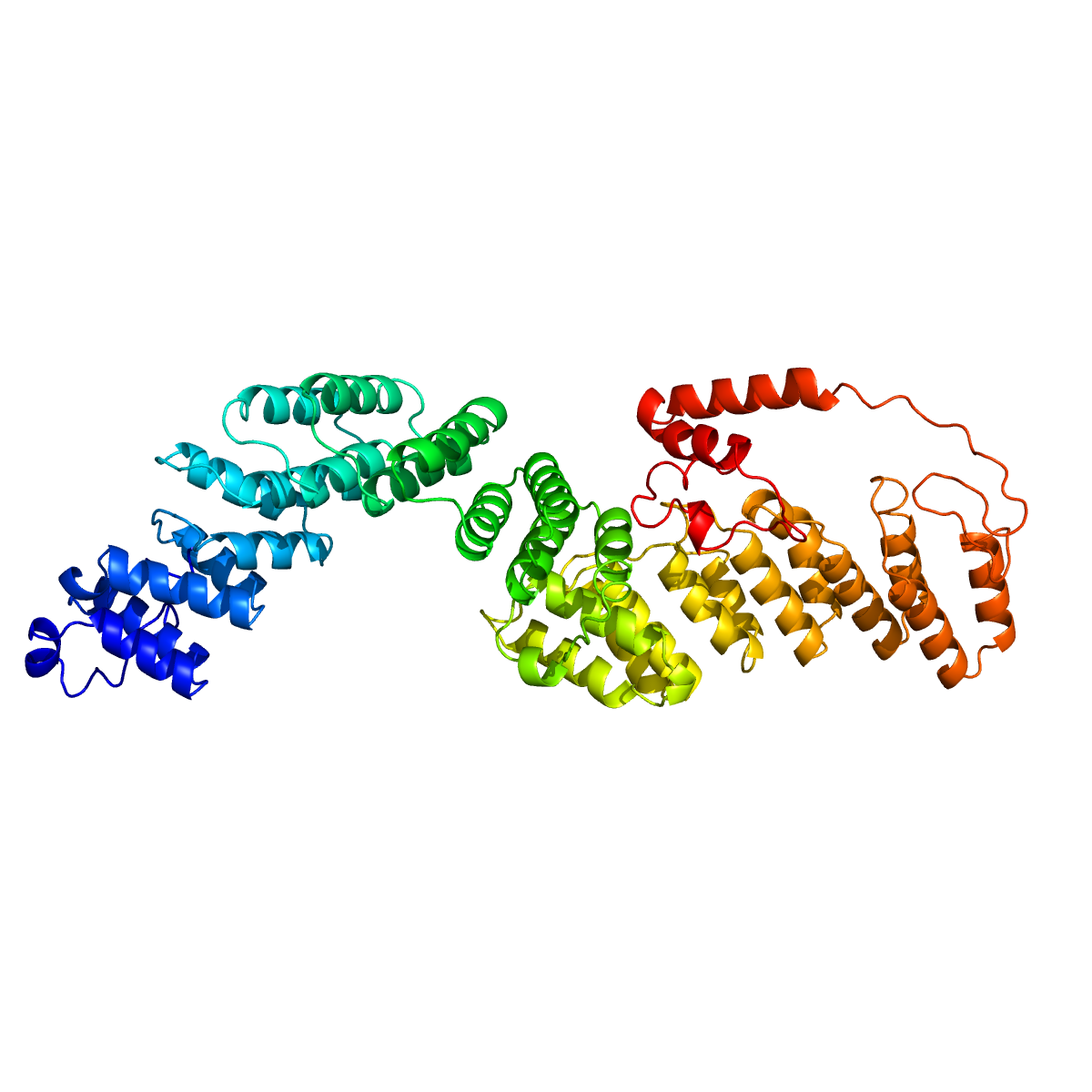
